# DISRUPTOR: Computational identification of oncogenic mutants disrupting protein interactions

**DOI:** 10.1101/2022.11.02.514903

**Authors:** V Kugler, A Lieb, N Guerin, BR Donald, E Stefan, T Kaserer

## Abstract

We report an Osprey-based computational protocol to prospectively identify oncogenic mutations that act via disruption of molecular interactions. It is applicable to analyze both protein-protein and protein-DNA interfaces and has been validated on a dataset of clinically relevant mutations. In addition, it was used to predict previously uncharacterized patient mutations in CDK6 and p16 genes, which were experimentally confirmed to impair complex formation.

## Main

Missense mutations play a central role in the onset and progression of cancer.^1^ Examples of relevant molecular mechanisms include oncogenic activation/inactivation of proteins,^1^ disruption of the contacts between proteins and their macromolecular interaction partners,^2-5^ or emergence of cancer drug resistance.^6^ The last has been previously addressed by a computational protocol^6,7^ predicting likely resistance mutations in the pharmacological target of targeted cancer drugs. However, to the best of our knowledge, no theoretical framework exists to systematically evaluate mutations within the interaction interfaces of critical signalling and regulatory components to identify disrupting mutations involved in the aetiology and progression of cancer (Fig. 1a).

**Fig. 1.**
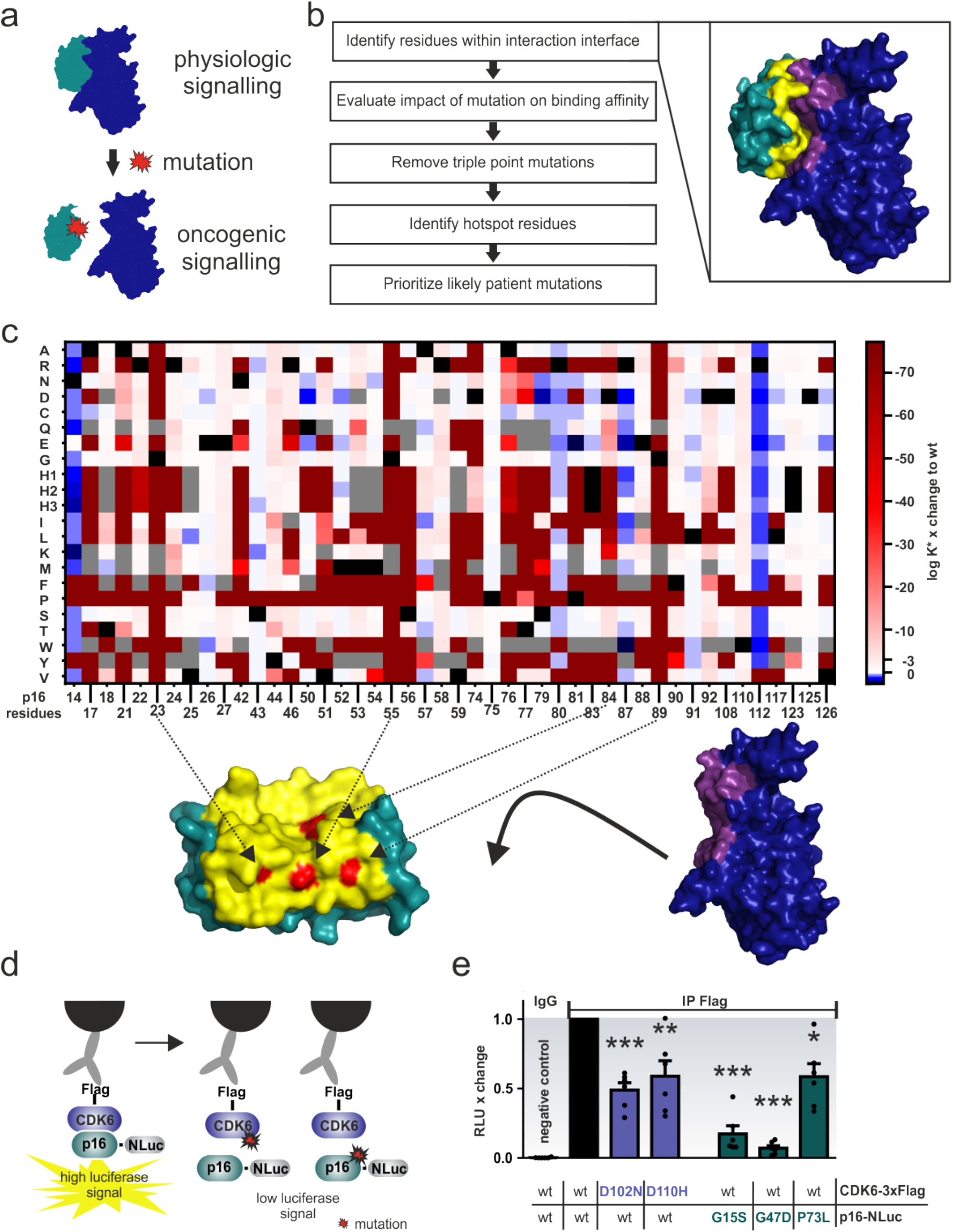
p16-CDK6 results. **a** Schematic representation of the molecular mechanism, where the two binding partners are presented in blue and green. Upon mutation (red), binding is disrupted. **b** Overview of the computational workflow. The inset shows the interaction between p16 (green) and CDK6 (blue), with the interaction interface coloured yellow (p16) and violet (CDK6). **c** Heatmap showing the changes in the Osprey^20^ log K* score for mutations (Y-axis, H1-3 correspond to different histidine protonation states) compared to wildtype (wt, marked black) p16 residues (X-axis). Triple point mutations are marked grey. Hotspot residues predicted to disrupt interaction with CDK6 are coloured red on the p16 surface below. Arrows indicate the p16 residue position. **d** Left: Schematic depiction of the LUMIER assay for the detection of protein:protein interactions. A p16 protein tagged with the NanoLuc Luciferase (NLuc) is transiently expressed in HEK293T cells together with Flag-tagged CDK6. The complex is immunoprecipitated with Flag antibodies and the emission of light is detected on-bead upon substrate (benzylcoelenterazine) addition if the bait protein is present. Expression profiles have been validated by Western Blot as shown in Supplementary Fig. 1. Right: introduction of dimerization interfering mutations to either CDK6 or p16 lower the detected luciferase signal. **e** LUMIER assay of Flag-tagged CDK6 variants in the presence of wildtype or mutated p16-NLuc. Please note, we introduced the p16 mutations into mouse, not human, p16 and thus G23S, G55D, and P81L correspond to mouse G15S, G47D and P73L. The bioluminescence signals were normalized on the corresponding input signals. Bars represent the luciferase intensity relative to wild-type CDK6 and p16 interactions. Error bars represent SEM of n=5 independ. experiments. Significance was determined by t-test *p< 0.05; **p<0.01; ***p<0.001.

We suggest that such mutations (1) have a high likelihood to be formed in a particular cancer type and (2) affect the molecular interactions formed by interaction partners, i.e., disrupt in the investigated cases. We report here the development and validation of a computational protocol, Disruptor, to address these aspects: It builds upon our previous work^6^ where we systematically evaluate the impact on binding affinity for all possible mutations within the binding interface using experimental structures of central protein complexes. In addition, we combine gene sequences and mutational signatures^8^ to calculate the probability with which a specific mutation is formed. Results of these analyses are used to predict and rank mutations that have a high probability to become clinically relevant for carcinogenesis (Fig. 1b). We have tested Disruptor on a dataset of known mutations involving p53:DNA (a consensus recognition sequence), p53:ASPP2 (also known as 53BP2), ERK2:DUSP6, p16 (also known as INK4a or CDKN2):CDK6 (Fig. 1b-c), and Smad4:Smad2 complexes. In all cases, Disruptor predicted clinically relevant mutations, which have been demonstrated to disrupt binding to their respective interaction partner (Table 1). For example, this includes highly prevalent p53 hotspot mutations, e.g., at residues R248, R249, and R273, which are known to interfere with binding of the transcription factor to its DNA response element and thus hamper transactivation.^2,5^ Furthermore, transactivation data deposited in the International Agency for Research on CANCER (IARC) TP53 database (version R20, July 2019)^9^ confirmed that 31% (67 out of 215) of our predicted mutations were indeed non-functional or only partially functional. In contrast, only 4% of mutations (10 out of the 215) showed transactivation activity despite their classification as disruptive. Unfortunately, for the vast majority of predicted p53 mutations (64%) within the DNA binding site, no functional or mechanistic data were available. This lack of data was not limited to p53, which is a thoroughly investigated target, but rather a general trend: For each interaction pair, we identified several mutations, that have not yet been investigated experimentally despite their detection in cancer patients, sometimes even multiple times (Supplementary Tables S1-S8). To investigate some of these understudied mutations in more detail, we selected three p16 (G23S, G55D, and P81L, Fig. 1c) and two CDK6 (D102N and D110N) patient mutations predicted by our method for experimental validation. Intriguingly, in a biochemical assay for quantifying binary complex formation of cellularly expressed proteins (termed LUMIER assay^10^; Fig. 1d) all five of our selected mutants showed a significant decrease in their binary interaction with the binding partner when compared to the non-mutated complex of p16:CDK6 (Fig. 1e).

**Table 1.** Exemplary, computationally predicted patient mutations confirmed to disrupt complex formation.

| Protein | Interaction partner | Mutation | Reference |
| --- | --- | --- | --- |
| p53 | DNA consensus sequence | R248Q | Merabet et al.<sup>2</sup> |
|  |  | R248W | Merabet et al. |
|  |  | R249S | Merabet et al. |
|  |  | R273C | Garg et al.<sup>5</sup> |
|  |  | R273H | Garg et al. |
|  |  | R273L | Garg et al. |
|  | ASPP2/53BP2 | R248W | Gorina et al.<sup>13</sup> |
|  |  | R249S | Gorina et al. |
|  |  | R273H | Gorina et al. |
| ERK2 | DUSP6 | D321N | Brenan et al.<sup>3</sup> Taylor et al.<sup>14</sup> |
| p16 | CDK6 | G23D | McKenzie et al.<sup>15</sup> |
|  |  | M53I | Harland et al.<sup>16</sup> |
|  |  | D84G | Yarbrough et al.<sup>17</sup> |
|  |  | D84H | Ruas et al.<sup>18</sup> |
|  |  | D84N | Ruas et al. |
|  |  | D84V | Yarbrough et al. |
|  |  | D84Y | Ruas et al. |
|  |  | R87P | Yarbrough et al. |
| smad4 | smad2 | R361C | Shi et al.<sup>4</sup> |
|  |  | D537E | Shi et al. |
|  |  | D537Y | Gori et al.<sup>19</sup> |

Besides providing validation of Disruptor, this indicates that there may be many more overlooked disease-relevant mutations in patients that occur only at low rates and thus only affect a small patient population or even individuals. We therefore suggest that our method could be an especially valuable tool in precision medicine.

However, some limitations of the current approach should be noted. There are many other mechanisms by which mutations can affect protein function that are not addressed in this computational framework. For example, many p53 mutations also exert a gain-of-function phenotype, e.g. via changes in protein stability or reprogramming of DNA or protein-protein interaction preferences.^11^ In addition, Disruptor requires structural data as input, which may not always be available. We are thus working on the extension of our computational toolbox towards additional molecular mechanisms and are investigating the suitability of computationally derived structural models (e.g. generated using AlphaFold^12^) as starting point for our analyses.

Taken together, we report a computational protocol to prospectively predict protein mutations affecting binding to macromolecular interaction partners. It can be applied to investigate data on novel patient mutations, guide selection of mutants for subsequent wet lab experiments, and even predict a potential mode of action on a molecular level. In addition, Disruptor can not only be used to systematically investigate all mutations within the interaction interface of a given target of interest, but also identify those that will most likely emerge in the clinic. Moreover, we highlight an adaptable computational workflow for anticipating and unveiling the functional relevance of less common and overlooked patient mutations.

## Methods

### Preparation of input structures

The following PDB entries were used for the analysis: 1tup (p53:DNA complex),^21^ 1ycs (p53:ASPP2),^13^ 2fys (ERK2:DUSP6),^22^ 1bi7 (p16:CDK6),^23^ and 1u7v (Smad4:Smad2).^24^ All structures were prepared using the default parameters of the Protein Preparation Wizard^25^ in Maestro release 2020-3^26^ and all water molecules and buffer components were deleted. In case of CDK6, ERK2, and p53:ASPP2, only residues within 12 Å of the interaction interface were included, and chains A and B were used for the calculation for both p53:ASPP2 and Smad4:Smad2. All three p53 copies were analysed in case of 1tup.

### Computational evaluation of mutations

Structures and definitions of mutable residues and allowed mutations were submitted in YAML file format. In case of histidine mutations, all three protonation states were considered. Mutable residues were either investigated alone or in pairs. Wildtype residues were set to continuously flexible, all other residues were kept rigid. ZAFF^27^ force field parameters were added for zinc ions and zinc coordinating residues and downloaded here: https://ambermd.org/tutorials/advanced/tutorial20/ZAFF.htm. Template coordinates and force field parameters for phosphoserines were calculated using Antechamber 19.0. An example input file for each of the interaction pairs is provided in the supplementary information. Osprey version 3^20^ was used for calculating K* scores, which predicts low-energy structural ensembles and provides an approximation to binding affinity. The stability threshold was disabled and an epsilon of 0.03 was used. OSPREY is free and open-source and available on GitHub at https://github.com/donaldlab/OSPREY3.

### Calculation of probabilities

A detailed description of the calculation of probabilities has been reported previously.^6,7,28^ Briefly, mutational signatures and their contribution to the mutational burden in a particular cancer type^8^ have been combined to calculate cancer-specific values for single base exchanges within a defined trinucleotide context. These have been used to calculate relative probabilities for generation of the DNA sequence mutations encoding for the investigated protein amino acid mutation. We only calculated probabilities for mutations that could be generated with single- or double base pair exchanges, because we considered mutation of the whole trinucleotide codon required for triple point mutations as extremely unlikely.^6^ Colorectal and cervical cancer associated probabilities have been calculated for ERK2, and melanoma and colorectal cancer associated probabilities have been calculated for p16 and smad4, respectively. No cancer-associated probabilities have been calculated for p53, given that p53 mutations have been observed in the majority of cancer types.

### Data analysis

Mutations with a change of Log_10_ K* scores greater than –3 in comparison to wildtype scores from the same run were considered to disrupt the interactions. Histidine mutations were included only if all three protonation states disrupted binding. Triple point mutations requiring mutations of all three bases of the codon were discarded. This led to a final set of mutations we considered clinically relevant. To prioritize mutations further, the number of individual mutations included for each residue position were analysed. Protein residues with the highest number of predicted individual mutations were considered as “mutational hotspots” and cancer-associated probabilities for all mutations at these positions were calculated to prioritize individual mutants further. An overview of the top-three mutational hotspots, and the individual mutations and their probabilities are reported in Supplementary Table S1-S8.

### Selection of mutants to be tested

Two of the three p16 mutations (human G23S and G55D (mouse G15S and G47D)) were chosen because they were prioritized by our protocol (Fig. 1c) and both have been associated with hereditary melanoma.^29,30^ P81L (corresponding to P73L in mouse) was included, because within the dataset of computationally predicted mutations it was among the most frequently reported in cancer patients (29 times). In contrast, CDK6 generally appears to be mutated at a very low rate, with only 97 unique missense mutations reported in COSMIC^31^ in total and the most common mutations observed only five times in patient samples. For comparison, the p16 H83Y mutation is reported 128 times and one of 387 unique missense mutations deposited. We therefore focused on two CDK6 mutations (i.e. D102N and D110N) that were also observed in the clinic.

### Cell culture and antibodies

HEK293 cells were grown in Dulbecco’s modified Eagle’s medium (DMEM) supplemented with 10% fetal bovine serum (FBS). Transient transfections were performed with TransFectin reagent (Bio-Rad, #1703352). Antibody used for LUMIER experiments was mouse anti-FLAG (Sigma-Aldrich, #F3165). The expression constructs were cloned using cDNA as PCR templates for amplifying the inserts (CDK6 (Gene ID: 12571, p16 (Gene ID: 12578)), digestion with restriction enzymes and ligation into a Flag or NLuc vector.

### Western Blotting

Expression of indicated Flag-tagged CDK6 constructs in HEK293 cells were determined via Western blotting with indicated Flag antibody [mouse anti-FLAG® M2-tag (Sigma-Aldrich, St. Louis, MO, USA, F3165-1MG)]. 5x SDS loading buffer was added to the lysate to reach a final concentration of 1x SDS LB.

### LUMIER experiments

HEK293 cells were transiently transfected with wild type or mutated p16-NLuc (NanoLuciferase) and 3xFlag-tagged wild type CDK6 constructs. Subsequent to homogenizing the cells with a syringe [lysis buffer: 150 mM NaCl, 10 mM sodium phosphate (pH 7.2), 0.05% Triton-X100 supplemented with standard protease inhibitors] the lysates were clarified by centrifugation at 13,000 rpm for 20 min. Cell extracts were incubated on an overhead shaker with anti-flag antibody (0.6 μg per sample) and protein G–Sepharose beads or IgG beads for 3 hours at 4°C. Isolated complexes were washed three times with lysis buffer and three times with PBS. Probes were transferred to 96-well white-walled plates and subjected to bioluminescence analysis using the PHERAstar FSX luminometer. As substrate benzylcoelenterazine is used. NLuc bioluminescence signals were integrated for 1.2 s following addition of luciferase substrate. Raw Data Bioluminescence signals are shown in Supplementary Fig. 1.

## Supporting information

exemplary input files

## Acknowledgements

This work has been funded by the Austrian Science Fund FWF (P34376 to T.K., P27606, P30441, P32960, and P35159 to E.S., and P35579 and P33222 to A.L.) and the NIH (R35-GM144042 to BRD). We would like to thank Veronika Sexl for providing the cDNAs for constructing the LUMIER-based reporter. The authors declare the following competing interests: B.R.D. is a founder of Ten63 Therapeutics, Inc. E.S. is a founder of KinCon Biolabs. KinCon reporters are subjects of pending patent applications.

