## Supplementary material for "DISRUPTOR: Computational identification of oncogenic mutants disrupting protein interactions": exemplary input files: CDK6_1bi7.14-15.yaml

```
---
osprey_version: 3.1
design_name: CDK6 and p16
protein:
  residue_configurations: []
  extra_templates: ""
  extra_templates_coordinates: ""
  coordinates: |
    ATOM   1823  N   GLU B 510     150.292  72.261 103.158  1.00 99.90           N1+
    ATOM   1824  CA  GLU B 510     150.131  71.514 101.893  1.00 99.90           C
    ATOM   1825  C   GLU B 510     150.441  70.012 102.169  1.00 99.90           C
    ATOM   1826  O   GLU B 510     151.418  69.716 102.863  1.00 99.90           O
    ATOM   1827  CB  GLU B 510     151.046  72.063 100.775  1.00  0.00           C
    ATOM   1828  CG  GLU B 510     150.742  73.525 100.379  1.00  0.00           C
    ATOM   1829  CD  GLU B 510     151.630  74.056  99.249  1.00  0.00           C
    ATOM   1830  OE1 GLU B 510     152.771  73.561  99.112  1.00  0.00           O
    ATOM   1831  OE2 GLU B 510     151.149  74.967  98.542  1.00  0.00           O1-
    ATOM   1832  H   GLU B 510     150.215  73.254 102.988  1.00  0.00           H
    ATOM   1833  HA  GLU B 510     149.103  71.658 101.568  1.00  0.00           H
    ATOM   1834  HB2 GLU B 510     152.090  71.983 101.089  1.00  0.00           H
    ATOM   1835  HB3 GLU B 510     150.953  71.436  99.886  1.00  0.00           H
    ATOM   1836  HG2 GLU B 510     149.699  73.610 100.071  1.00  0.00           H
    ATOM   1837  HG3 GLU B 510     150.871  74.184 101.237  1.00  0.00           H
    ATOM   1838  N   PRO B 511     149.616  69.079 101.628  1.00 99.90           N
    ATOM   1839  CA  PRO B 511     149.798  67.634 101.860  1.00 99.90           C
    ATOM   1840  C   PRO B 511     150.828  66.952 100.932  1.00 99.90           C
    ATOM   1841  O   PRO B 511     151.966  66.758 101.354  1.00 99.90           O
    ATOM   1842  CB  PRO B 511     148.373  67.081 101.705  1.00 99.90           C
    ATOM   1843  CG  PRO B 511     147.723  67.983 100.668  1.00 99.90           C
    ATOM   1844  CD  PRO B 511     148.337  69.348 100.964  1.00 99.90           C
    ATOM   1845  HA  PRO B 511     150.118  67.448 102.887  1.00  0.00           H
    ATOM   1846  HB2 PRO B 511     148.340  66.029 101.425  1.00  0.00           H
    ATOM   1847  HB3 PRO B 511     147.844  67.176 102.655  1.00  0.00           H
    ATOM   1848  HG2 PRO B 511     147.998  67.666  99.663  1.00  0.00           H
    ATOM   1849  HG3 PRO B 511     146.634  67.981 100.722  1.00  0.00           H
    ATOM   1850  HD2 PRO B 511     148.455  69.941 100.055  1.00  0.00           H
    ATOM   1851  HD3 PRO B 511     147.688  69.892 101.651  1.00  0.00           H
    ATOM   1852  N   SER B 512     150.400  66.520  99.731  1.00 99.90           N
    ATOM   1853  CA  SER B 512     151.108  65.632  98.795  1.00 99.90           C
    ATOM   1854  C   SER B 512     151.150  64.168  99.295  1.00 99.90           C
    ATOM   1855  O   SER B 512     150.872  63.239  98.540  1.00 99.90           O
    ATOM   1856  CB  SER B 512     152.504  66.157  98.408  1.00 99.90           C
    ATOM   1857  OG  SER B 512     152.995  65.474  97.271  1.00 99.90           O
    ATOM   1858  H   SER B 512     149.448  66.731  99.470  1.00  0.00           H
    ATOM   1859  HA  SER B 512     150.531  65.694  97.880  1.00  0.00           H
    ATOM   1860  HB2 SER B 512     152.459  67.221  98.170  1.00  0.00           H
    ATOM   1861  HB3 SER B 512     153.223  66.040  99.220  1.00  0.00           H
    ATOM   1862  HG  SER B 512     153.229  64.577  97.524  1.00  0.00           H
    ATOM   1863  N   ALA B 513     151.446  63.974 100.585  1.00 91.34           N
    ATOM   1864  CA  ALA B 513     151.292  62.710 101.284  1.00 91.34           C
    ATOM   1865  C   ALA B 513     149.844  62.604 101.788  1.00 91.34           C
    ATOM   1866  O   ALA B 513     149.554  62.759 102.972  1.00 91.34           O
    ATOM   1867  CB  ALA B 513     152.357  62.636 102.373  1.00 99.90           C
    ATOM   1868  H   ALA B 513     151.678  64.775 101.159  1.00  0.00           H
    ATOM   1869  HA  ALA B 513     151.474  61.878 100.602  1.00  0.00           H
    ATOM   1870  HB1 ALA B 513     152.241  61.736 102.976  1.00  0.00           H
    ATOM   1871  HB2 ALA B 513     153.353  62.599 101.933  1.00  0.00           H
    ATOM   1872  HB3 ALA B 513     152.322  63.517 103.014  1.00  0.00           H
    ATOM   1873  N   ASP B 514     148.971  62.380 100.808  1.00 77.49           N
    ATOM   1874  CA  ASP B 514     147.525  62.235 100.878  1.00 77.49           C
    ATOM   1875  C   ASP B 514     147.168  61.446  99.621  1.00 77.49           C
    ATOM   1876  O   ASP B 514     146.502  60.420  99.721  1.00 77.49           O
    ATOM   1877  CB  ASP B 514     146.797  63.595 100.947  1.00 99.90           C
    ATOM   1878  CG  ASP B 514     145.276  63.520 101.125  1.00 99.90           C
    ATOM   1879  OD1 ASP B 514     144.731  62.400 101.232  1.00 99.90           O
    ATOM   1880  OD2 ASP B 514     144.682  64.616 101.195  1.00 99.90           O1-
    ATOM   1881  H   ASP B 514     149.346  62.310  99.873  1.00  0.00           H
    ATOM   1882  HA  ASP B 514     147.278  61.635 101.756  1.00  0.00           H
    ATOM   1883  HB2 ASP B 514     147.183  64.133 101.812  1.00  0.00           H
    ATOM   1884  HB3 ASP B 514     147.018  64.181 100.053  1.00  0.00           H
    ATOM   1885  N   TRP B 515     147.762  61.860  98.486  1.00 61.24           N
    ATOM   1886  CA  TRP B 515     147.926  61.059  97.278  1.00 61.24           C
    ATOM   1887  C   TRP B 515     148.501  59.684  97.604  1.00 61.24           C
    ATOM   1888  O   TRP B 515     148.027  58.690  97.070  1.00 61.24           O
    ATOM   1889  CB  TRP B 515     148.909  61.758  96.330  1.00 99.90           C
    ATOM   1890  CG  TRP B 515     148.566  63.138  95.909  1.00 99.90           C
    ATOM   1891  CD1 TRP B 515     148.758  64.269  96.612  1.00 99.90           C
    ATOM   1892  CD2 TRP B 515     147.839  63.540  94.729  1.00 99.90           C
    ATOM   1893  NE1 TRP B 515     148.378  65.362  95.878  1.00 99.90           N
    ATOM   1894  CE2 TRP B 515     147.760  64.965  94.716  1.00 99.90           C
    ATOM   1895  CE3 TRP B 515     147.218  62.837  93.676  1.00 99.90           C
    ATOM   1896  CZ2 TRP B 515     147.110  65.666  93.694  1.00 99.90           C
    ATOM   1897  CZ3 TRP B 515     146.535  63.530  92.661  1.00 99.90           C
    ATOM   1898  CH2 TRP B 515     146.487  64.938  92.666  1.00 99.90           C
    ATOM   1899  H   TRP B 515     148.257  62.740  98.485  1.00  0.00           H
    ATOM   1900  HA  TRP B 515     146.953  60.947  96.796  1.00  0.00           H
    ATOM   1901  HB2 TRP B 515     149.901  61.813  96.771  1.00  0.00           H
    ATOM   1902  HB3 TRP B 515     149.017  61.165  95.421  1.00  0.00           H
    ATOM   1903  HD1 TRP B 515     149.064  64.298  97.633  1.00  0.00           H
    ATOM   1904  HE1 TRP B 515     148.448  66.311  96.215  1.00  0.00           H
    ATOM   1905  HE3 TRP B 515     147.250  61.758  93.660  1.00  0.00           H
    ATOM   1906  HZ2 TRP B 515     147.061  66.745  93.710  1.00  0.00           H
    ATOM   1907  HZ3 TRP B 515     146.052  62.980  91.868  1.00  0.00           H
    ATOM   1908  HH2 TRP B 515     145.964  65.461  91.880  1.00  0.00           H
    ATOM   1909  N   LEU B 516     149.512  59.686  98.489  1.00 47.40           N
    ATOM   1910  CA  LEU B 516     150.227  58.508  98.938  1.00 47.40           C
    ATOM   1911  C   LEU B 516     149.356  57.616  99.829  1.00 47.40           C
    ATOM   1912  O   LEU B 516     149.465  56.396  99.739  1.00 47.40           O
    ATOM   1913  CB  LEU B 516     151.539  58.982  99.598  1.00 64.05           C
    ATOM   1914  CG  LEU B 516     152.630  57.913  99.833  1.00 64.05           C
    ATOM   1915  CD1 LEU B 516     152.626  57.392 101.260  1.00 64.05           C
    ATOM   1916  CD2 LEU B 516     152.645  56.762  98.815  1.00 64.05           C
    ATOM   1917  H   LEU B 516     149.835  60.568  98.859  1.00  0.00           H
    ATOM   1918  HA  LEU B 516     150.455  57.940  98.040  1.00  0.00           H
    ATOM   1919  HB2 LEU B 516     151.978  59.732  98.940  1.00  0.00           H
    ATOM   1920  HB3 LEU B 516     151.321  59.531 100.514  1.00  0.00           H
    ATOM   1921  HG  LEU B 516     153.580  58.438  99.719  1.00  0.00           H
    ATOM   1922 HD11 LEU B 516     153.639  57.162 101.594  1.00  0.00           H
    ATOM   1923 HD12 LEU B 516     152.219  58.142 101.938  1.00  0.00           H
    ATOM   1924 HD13 LEU B 516     152.030  56.485 101.344  1.00  0.00           H
    ATOM   1925 HD21 LEU B 516     153.606  56.257  98.822  1.00  0.00           H
    ATOM   1926 HD22 LEU B 516     151.894  56.003  99.029  1.00  0.00           H
    ATOM   1927 HD23 LEU B 516     152.487  57.124  97.801  1.00  0.00           H
    ATOM   1928  N   ALA B 517     148.456  58.231 100.612  1.00 53.31           N
    ATOM   1929  CA  ALA B 517     147.450  57.533 101.401  1.00 53.31           C
    ATOM   1930  C   ALA B 517     146.459  56.787 100.508  1.00 53.31           C
    ATOM   1931  O   ALA B 517     146.363  55.574 100.658  1.00 53.31           O
    ATOM   1932  CB  ALA B 517     146.742  58.525 102.320  1.00 99.85           C
    ATOM   1933  H   ALA B 517     148.395  59.238 100.605  1.00  0.00           H
    ATOM   1934  HA  ALA B 517     147.968  56.808 102.031  1.00  0.00           H
    ATOM   1935  HB1 ALA B 517     146.276  58.029 103.167  1.00  0.00           H
    ATOM   1936  HB2 ALA B 517     147.465  59.233 102.727  1.00  0.00           H
    ATOM   1937  HB3 ALA B 517     145.967  59.098 101.813  1.00  0.00           H
    ATOM   1938  N   THR B 518     145.808  57.493  99.562  1.00 36.39           N
    ATOM   1939  CA  THR B 518     144.927  56.877  98.568  1.00 36.39           C
    ATOM   1940  C   THR B 518     145.662  55.847  97.704  1.00 36.39           C
    ATOM   1941  O   THR B 518     145.070  54.828  97.398  1.00 36.39           O
    ATOM   1942  CB  THR B 518     144.202  57.875  97.620  1.00 59.31           C
    ATOM   1943  OG1 THR B 518     144.918  58.212  96.441  1.00 59.31           O
    ATOM   1944  CG2 THR B 518     143.710  59.157  98.299  1.00 59.31           C
    ATOM   1945  H   THR B 518     145.949  58.493  99.491  1.00  0.00           H
    ATOM   1946  HA  THR B 518     144.149  56.359  99.128  1.00  0.00           H
    ATOM   1947  HB  THR B 518     143.311  57.356  97.264  1.00  0.00           H
    ATOM   1948 HG21 THR B 518     142.920  59.624  97.710  1.00  0.00           H
    ATOM   1949 HG22 THR B 518     143.312  58.965  99.296  1.00  0.00           H
    ATOM   1950 HG23 THR B 518     144.504  59.891  98.395  1.00  0.00           H
    ATOM   1951  HG1 THR B 518     145.710  58.696  96.689  1.00  0.00           H
    ATOM   1952  N   ALA B 519     146.942  56.089  97.372  1.00 20.56           N
    ATOM   1953  CA  ALA B 519     147.777  55.149  96.636  1.00 20.56           C
    ATOM   1954  C   ALA B 519     147.901  53.843  97.410  1.00 20.56           C
    ATOM   1955  O   ALA B 519     147.458  52.817  96.904  1.00 20.56           O
    ATOM   1956  CB  ALA B 519     149.155  55.753  96.331  1.00 53.07           C
    ATOM   1957  H   ALA B 519     147.375  56.965  97.634  1.00  0.00           H
    ATOM   1958  HA  ALA B 519     147.281  54.947  95.685  1.00  0.00           H
    ATOM   1959  HB1 ALA B 519     149.711  55.119  95.641  1.00  0.00           H
    ATOM   1960  HB2 ALA B 519     149.061  56.718  95.836  1.00  0.00           H
    ATOM   1961  HB3 ALA B 519     149.767  55.888  97.220  1.00  0.00           H
    ATOM   1962  N   ALA B 520     148.388  53.937  98.658  1.00 42.94           N
    ATOM   1963  CA  ALA B 520     148.470  52.821  99.582  1.00 42.94           C
    ATOM   1964  C   ALA B 520     147.115  52.128  99.747  1.00 42.94           C
    ATOM   1965  O   ALA B 520     147.062  50.911  99.611  1.00 42.94           O
    ATOM   1966  CB  ALA B 520     149.062  53.286 100.920  1.00 73.75           C
    ATOM   1967  H   ALA B 520     148.693  54.836  99.011  1.00  0.00           H
    ATOM   1968  HA  ALA B 520     149.160  52.116  99.134  1.00  0.00           H
    ATOM   1969  HB1 ALA B 520     149.241  52.442 101.581  1.00  0.00           H
    ATOM   1970  HB2 ALA B 520     150.019  53.785 100.770  1.00  0.00           H
    ATOM   1971  HB3 ALA B 520     148.404  53.983 101.439  1.00  0.00           H
    ATOM   1972  N   ALA B 521     146.055  52.917  99.989  1.00 57.12           N
    ATOM   1973  CA  ALA B 521     144.677  52.473 100.186  1.00 57.12           C
    ATOM   1974  C   ALA B 521     144.076  51.748  98.985  1.00 57.12           C
    ATOM   1975  O   ALA B 521     143.421  50.726  99.170  1.00 57.12           O
    ATOM   1976  CB  ALA B 521     143.796  53.666 100.581  1.00 58.93           C
    ATOM   1977  H   ALA B 521     146.205  53.916 100.082  1.00  0.00           H
    ATOM   1978  HA  ALA B 521     144.686  51.751 100.995  1.00  0.00           H
    ATOM   1979  HB1 ALA B 521     142.825  53.329 100.944  1.00  0.00           H
    ATOM   1980  HB2 ALA B 521     144.253  54.252 101.377  1.00  0.00           H
    ATOM   1981  HB3 ALA B 521     143.613  54.328  99.737  1.00  0.00           H
    ATOM   1982  N   ARG B 522     144.346  52.258  97.779  1.00 32.72           N
    ATOM   1983  CA  ARG B 522     143.987  51.632  96.513  1.00 32.72           C
    ATOM   1984  C   ARG B 522     144.938  50.504  96.122  1.00 32.72           C
    ATOM   1985  O   ARG B 522     144.691  49.862  95.103  1.00 32.72           O
    ATOM   1986  CB  ARG B 522     143.921  52.688  95.400  1.00 75.54           C
    ATOM   1987  CG  ARG B 522     142.805  53.711  95.640  1.00 75.54           C
    ATOM   1988  CD  ARG B 522     142.165  54.225  94.369  1.00 75.54           C
    ATOM   1989  NE  ARG B 522     143.069  54.824  93.382  1.00 75.54           N
    ATOM   1990  CZ  ARG B 522     143.394  54.314  92.183  1.00 75.54           C
    ATOM   1991  NH1 ARG B 522     143.010  53.090  91.805  1.00 75.54           N
    ATOM   1992  NH2 ARG B 522     144.119  55.059  91.345  1.00 75.54           N1+
    ATOM   1993  H   ARG B 522     144.865  53.126  97.711  1.00  0.00           H
    ATOM   1994  HA  ARG B 522     143.001  51.174  96.617  1.00  0.00           H
    ATOM   1995  HB2 ARG B 522     144.881  53.186  95.251  1.00  0.00           H
    ATOM   1996  HB3 ARG B 522     143.703  52.175  94.462  1.00  0.00           H
    ATOM   1997  HG2 ARG B 522     141.999  53.132  96.092  1.00  0.00           H
    ATOM   1998  HG3 ARG B 522     142.992  54.495  96.366  1.00  0.00           H
    ATOM   1999  HD2 ARG B 522     141.441  53.536  93.980  1.00  0.00           H
    ATOM   2000  HD3 ARG B 522     141.606  55.107  94.679  1.00  0.00           H
    ATOM   2001  HE  ARG B 522     143.443  55.729  93.633  1.00  0.00           H
    ATOM   2002 HH11 ARG B 522     142.472  52.516  92.437  1.00  0.00           H
    ATOM   2003 HH12 ARG B 522     143.226  52.738  90.884  1.00  0.00           H
    ATOM   2004 HH21 ARG B 522     144.383  56.002  91.592  1.00  0.00           H
    ATOM   2005 HH22 ARG B 522     144.402  54.698  90.447  1.00  0.00           H
    ATOM   2006  N   GLY B 523     145.980  50.268  96.939  1.00 47.56           N
    ATOM   2007  CA  GLY B 523     147.057  49.303  96.749  1.00 47.56           C
    ATOM   2008  C   GLY B 523     147.723  49.480  95.389  1.00 47.56           C
    ATOM   2009  O   GLY B 523     147.902  48.510  94.653  1.00 47.56           O
    ATOM   2010  H   GLY B 523     146.065  50.848  97.763  1.00  0.00           H
    ATOM   2011  HA2 GLY B 523     147.789  49.508  97.526  1.00  0.00           H
    ATOM   2012  HA3 GLY B 523     146.758  48.272  96.892  1.00  0.00           H
    ATOM   2013  N   ARG B 524     148.058  50.735  95.065  1.00 39.85           N
    ATOM   2014  CA  ARG B 524     148.767  51.127  93.859  1.00 39.85           C
    ATOM   2015  C   ARG B 524     150.245  51.052  94.127  1.00 39.85           C
    ATOM   2016  O   ARG B 524     150.873  52.085  94.277  1.00 39.85           O
    ATOM   2017  CB  ARG B 524     148.311  52.537  93.407  1.00 99.90           C
    ATOM   2018  CG  ARG B 524     147.142  52.607  92.416  1.00 99.90           C
    ATOM   2019  CD  ARG B 524     146.196  51.400  92.403  1.00 99.90           C
    ATOM   2020  NE  ARG B 524     146.781  50.271  91.659  1.00 99.90           N
    ATOM   2021  CZ  ARG B 524     146.315  49.018  91.609  1.00 99.90           C
    ATOM   2022  NH1 ARG B 524     145.165  48.685  92.209  1.00 99.90           N
    ATOM   2023  NH2 ARG B 524     147.024  48.093  90.955  1.00 99.90           N1+
    ATOM   2024  H   ARG B 524     147.854  51.480  95.722  1.00  0.00           H
    ATOM   2025  HA  ARG B 524     148.637  50.393  93.076  1.00  0.00           H
    ATOM   2026  HB2 ARG B 524     148.093  53.148  94.282  1.00  0.00           H
    ATOM   2027  HB3 ARG B 524     149.123  53.038  92.890  1.00  0.00           H
    ATOM   2028  HG2 ARG B 524     146.612  53.558  92.468  1.00  0.00           H
    ATOM   2029  HG3 ARG B 524     147.619  52.612  91.434  1.00  0.00           H
    ATOM   2030  HD2 ARG B 524     146.106  51.038  93.420  1.00  0.00           H
    ATOM   2031  HD3 ARG B 524     145.179  51.657  92.126  1.00  0.00           H
    ATOM   2032  HE  ARG B 524     147.637  50.475  91.167  1.00  0.00           H
    ATOM   2033 HH11 ARG B 524     144.660  49.373  92.749  1.00  0.00           H
    ATOM   2034 HH12 ARG B 524     144.809  47.736  92.197  1.00  0.00           H
    ATOM   2035 HH21 ARG B 524     147.911  48.323  90.531  1.00  0.00           H
    ATOM   2036 HH22 ARG B 524     146.692  47.140  90.916  1.00  0.00           H
    ATOM   2037  N   VAL B 525     150.769  49.820  94.125  1.00 68.55           N
    ATOM   2038  CA  VAL B 525     152.177  49.477  94.287  1.00 68.55           C
    ATOM   2039  C   VAL B 525     153.104  50.454  93.555  1.00 68.55           C
    ATOM   2040  O   VAL B 525     153.919  51.107  94.200  1.00 68.55           O
    ATOM   2041  CB  VAL B 525     152.501  48.051  93.760  1.00 78.99           C
    ATOM   2042  CG1 VAL B 525     153.642  47.452  94.584  1.00 78.99           C
    ATOM   2043  CG2 VAL B 525     151.293  47.093  93.721  1.00 78.99           C
    ATOM   2044  H   VAL B 525     150.148  49.033  94.001  1.00  0.00           H
    ATOM   2045  HA  VAL B 525     152.431  49.544  95.341  1.00  0.00           H
    ATOM   2046  HB  VAL B 525     152.861  48.086  92.730  1.00  0.00           H
    ATOM   2047 HG11 VAL B 525     153.900  46.471  94.197  1.00  0.00           H
    ATOM   2048 HG12 VAL B 525     154.541  48.067  94.534  1.00  0.00           H
    ATOM   2049 HG13 VAL B 525     153.364  47.350  95.633  1.00  0.00           H
    ATOM   2050 HG21 VAL B 525     151.600  46.091  93.419  1.00  0.00           H
    ATOM   2051 HG22 VAL B 525     150.815  47.004  94.697  1.00  0.00           H
    ATOM   2052 HG23 VAL B 525     150.539  47.420  93.005  1.00  0.00           H
    ATOM   2053  N   GLU B 526     152.858  50.616  92.247  1.00 49.24           N
    ATOM   2054  CA  GLU B 526     153.624  51.510  91.400  1.00 49.24           C
    ATOM   2055  C   GLU B 526     153.432  52.993  91.692  1.00 49.24           C
    ATOM   2056  O   GLU B 526     154.447  53.680  91.727  1.00 49.24           O
    ATOM   2057  CB  GLU B 526     153.405  51.177  89.915  1.00 99.90           C
    ATOM   2058  CG  GLU B 526     153.869  49.755  89.534  1.00 99.90           C
    ATOM   2059  CD  GLU B 526     155.309  49.445  89.969  1.00 99.90           C
    ATOM   2060  OE1 GLU B 526     156.190  50.301  89.731  1.00 99.90           O
    ATOM   2061  OE2 GLU B 526     155.504  48.375  90.585  1.00 99.90           O1-
    ATOM   2062  H   GLU B 526     152.139  50.071  91.796  1.00  0.00           H
    ATOM   2063  HA  GLU B 526     154.672  51.334  91.646  1.00  0.00           H
    ATOM   2064  HB2 GLU B 526     152.349  51.286  89.665  1.00  0.00           H
    ATOM   2065  HB3 GLU B 526     153.933  51.901  89.292  1.00  0.00           H
    ATOM   2066  HG2 GLU B 526     153.199  49.016  89.976  1.00  0.00           H
    ATOM   2067  HG3 GLU B 526     153.798  49.622  88.454  1.00  0.00           H
    ATOM   2068  N   GLU B 527     152.199  53.471  91.942  1.00 38.97           N
    ATOM   2069  CA  GLU B 527     152.027  54.867  92.345  1.00 38.97           C
    ATOM   2070  C   GLU B 527     152.599  55.120  93.734  1.00 38.97           C
    ATOM   2071  O   GLU B 527     153.315  56.088  93.856  1.00 38.97           O
    ATOM   2072  CB  GLU B 527     150.603  55.412  92.195  1.00 99.90           C
    ATOM   2073  CG  GLU B 527     150.092  55.296  90.750  1.00 99.90           C
    ATOM   2074  CD  GLU B 527     148.719  55.932  90.555  1.00 99.90           C
    ATOM   2075  OE1 GLU B 527     148.565  57.109  90.950  1.00 99.90           O
    ATOM   2076  OE2 GLU B 527     147.849  55.224  90.001  1.00 99.90           O1-
    ATOM   2077  H   GLU B 527     151.387  52.872  91.941  1.00  0.00           H
    ATOM   2078  HA  GLU B 527     152.620  55.466  91.651  1.00  0.00           H
    ATOM   2079  HB2 GLU B 527     149.924  54.949  92.905  1.00  0.00           H
    ATOM   2080  HB3 GLU B 527     150.616  56.467  92.476  1.00  0.00           H
    ATOM   2081  HG2 GLU B 527     150.786  55.785  90.066  1.00  0.00           H
    ATOM   2082  HG3 GLU B 527     150.044  54.248  90.450  1.00  0.00           H
    ATOM   2083  N   VAL B 528     152.397  54.226  94.711  1.00 42.59           N
    ATOM   2084  CA  VAL B 528     153.012  54.275  96.036  1.00 42.59           C
    ATOM   2085  C   VAL B 528     154.541  54.389  95.953  1.00 42.59           C
    ATOM   2086  O   VAL B 528     155.088  55.315  96.545  1.00 42.59           O
    ATOM   2087  CB  VAL B 528     152.652  53.034  96.898  1.00 37.40           C
    ATOM   2088  CG1 VAL B 528     153.425  52.937  98.219  1.00 37.40           C
    ATOM   2089  CG2 VAL B 528     151.170  53.019  97.254  1.00 37.40           C
    ATOM   2090  H   VAL B 528     151.819  53.414  94.524  1.00  0.00           H
    ATOM   2091  HA  VAL B 528     152.628  55.167  96.528  1.00  0.00           H
    ATOM   2092  HB  VAL B 528     152.856  52.133  96.322  1.00  0.00           H
    ATOM   2093 HG11 VAL B 528     153.069  52.112  98.833  1.00  0.00           H
    ATOM   2094 HG12 VAL B 528     154.477  52.765  98.061  1.00  0.00           H
    ATOM   2095 HG13 VAL B 528     153.311  53.855  98.791  1.00  0.00           H
    ATOM   2096 HG21 VAL B 528     150.885  52.059  97.680  1.00  0.00           H
    ATOM   2097 HG22 VAL B 528     150.951  53.796  97.985  1.00  0.00           H
    ATOM   2098 HG23 VAL B 528     150.526  53.186  96.399  1.00  0.00           H
    ATOM   2099  N   ARG B 529     155.175  53.497  95.167  1.00 78.77           N
    ATOM   2100  CA  ARG B 529     156.594  53.547  94.817  1.00 78.77           C
    ATOM   2101  C   ARG B 529     156.953  54.895  94.205  1.00 78.77           C
    ATOM   2102  O   ARG B 529     157.701  55.627  94.835  1.00 78.77           O
    ATOM   2103  CB  ARG B 529     156.985  52.384  93.880  1.00 99.90           C
    ATOM   2104  CG  ARG B 529     158.500  52.315  93.567  1.00 99.90           C
    ATOM   2105  CD  ARG B 529     158.836  51.699  92.201  1.00 99.90           C
    ATOM   2106  NE  ARG B 529     158.258  50.364  92.003  1.00 99.90           N
    ATOM   2107  CZ  ARG B 529     158.550  49.259  92.707  1.00 99.90           C
    ATOM   2108  NH1 ARG B 529     159.432  49.275  93.718  1.00 99.90           N
    ATOM   2109  NH2 ARG B 529     157.948  48.112  92.376  1.00 99.90           N1+
    ATOM   2110  H   ARG B 529     154.637  52.765  94.721  1.00  0.00           H
    ATOM   2111  HA  ARG B 529     157.192  53.456  95.719  1.00  0.00           H
    ATOM   2112  HB2 ARG B 529     156.671  51.432  94.307  1.00  0.00           H
    ATOM   2113  HB3 ARG B 529     156.433  52.481  92.946  1.00  0.00           H
    ATOM   2114  HG2 ARG B 529     158.873  53.331  93.460  1.00  0.00           H
    ATOM   2115  HG3 ARG B 529     159.089  51.912  94.390  1.00  0.00           H
    ATOM   2116  HD2 ARG B 529     158.584  52.380  91.387  1.00  0.00           H
    ATOM   2117  HD3 ARG B 529     159.914  51.548  92.147  1.00  0.00           H
    ATOM   2118  HE  ARG B 529     157.576  50.292  91.253  1.00  0.00           H
    ATOM   2119 HH11 ARG B 529     159.912  50.133  93.949  1.00  0.00           H
    ATOM   2120 HH12 ARG B 529     159.634  48.437  94.244  1.00  0.00           H
    ATOM   2121 HH21 ARG B 529     157.246  48.114  91.641  1.00  0.00           H
    ATOM   2122 HH22 ARG B 529     158.201  47.250  92.833  1.00  0.00           H
    ATOM   2123  N   ALA B 530     156.403  55.189  93.017  1.00 71.99           N
    ATOM   2124  CA  ALA B 530     156.637  56.404  92.244  1.00 71.99           C
    ATOM   2125  C   ALA B 530     156.499  57.662  93.086  1.00 71.99           C
    ATOM   2126  O   ALA B 530     157.464  58.393  93.192  1.00 71.99           O
    ATOM   2127  CB  ALA B 530     155.680  56.466  91.047  1.00 99.90           C
    ATOM   2128  H   ALA B 530     155.763  54.527  92.595  1.00  0.00           H
    ATOM   2129  HA  ALA B 530     157.659  56.355  91.865  1.00  0.00           H
    ATOM   2130  HB1 ALA B 530     155.861  57.357  90.444  1.00  0.00           H
    ATOM   2131  HB2 ALA B 530     155.815  55.604  90.394  1.00  0.00           H
    ATOM   2132  HB3 ALA B 530     154.637  56.488  91.360  1.00  0.00           H
    ATOM   2133  N   LEU B 531     155.351  57.819  93.745  1.00 54.26           N
    ATOM   2134  CA  LEU B 531     155.014  58.878  94.679  1.00 54.26           C
    ATOM   2135  C   LEU B 531     156.073  59.044  95.769  1.00 54.26           C
    ATOM   2136  O   LEU B 531     156.580  60.152  95.917  1.00 54.26           O
    ATOM   2137  CB  LEU B 531     153.613  58.559  95.209  1.00 72.01           C
    ATOM   2138  CG  LEU B 531     152.793  59.628  95.912  1.00 72.01           C
    ATOM   2139  CD1 LEU B 531     153.053  61.058  95.426  1.00 72.01           C
    ATOM   2140  CD2 LEU B 531     151.327  59.266  95.676  1.00 72.01           C
    ATOM   2141  H   LEU B 531     154.631  57.114  93.631  1.00  0.00           H
    ATOM   2142  HA  LEU B 531     154.983  59.792  94.084  1.00  0.00           H
    ATOM   2143  HB2 LEU B 531     153.038  58.360  94.305  1.00  0.00           H
    ATOM   2144  HB3 LEU B 531     153.616  57.645  95.804  1.00  0.00           H
    ATOM   2145  HG  LEU B 531     153.026  59.584  96.975  1.00  0.00           H
    ATOM   2146 HD11 LEU B 531     152.462  61.774  96.000  1.00  0.00           H
    ATOM   2147 HD12 LEU B 531     154.096  61.351  95.530  1.00  0.00           H
    ATOM   2148 HD13 LEU B 531     152.783  61.176  94.376  1.00  0.00           H
    ATOM   2149 HD21 LEU B 531     150.742  59.865  96.341  1.00  0.00           H
    ATOM   2150 HD22 LEU B 531     151.002  59.497  94.661  1.00  0.00           H
    ATOM   2151 HD23 LEU B 531     151.118  58.213  95.867  1.00  0.00           H
    ATOM   2152  N   LEU B 532     156.479  57.935  96.412  1.00 60.24           N
    ATOM   2153  CA  LEU B 532     157.613  57.905  97.335  1.00 60.24           C
    ATOM   2154  C   LEU B 532     158.942  58.324  96.691  1.00 60.24           C
    ATOM   2155  O   LEU B 532     159.516  59.317  97.128  1.00 60.24           O
    ATOM   2156  CB  LEU B 532     157.760  56.524  97.997  1.00 63.77           C
    ATOM   2157  CG  LEU B 532     156.755  56.234  99.111  1.00 63.77           C
    ATOM   2158  CD1 LEU B 532     156.890  54.787  99.598  1.00 63.77           C
    ATOM   2159  CD2 LEU B 532     156.887  57.215 100.275  1.00 63.77           C
    ATOM   2160  H   LEU B 532     156.038  57.046  96.213  1.00  0.00           H
    ATOM   2161  HA  LEU B 532     157.418  58.650  98.105  1.00  0.00           H
    ATOM   2162  HB2 LEU B 532     157.727  55.728  97.258  1.00  0.00           H
    ATOM   2163  HB3 LEU B 532     158.744  56.468  98.447  1.00  0.00           H
    ATOM   2164  HG  LEU B 532     155.755  56.360  98.709  1.00  0.00           H
    ATOM   2165 HD11 LEU B 532     156.938  54.707 100.685  1.00  0.00           H
    ATOM   2166 HD12 LEU B 532     156.033  54.210  99.262  1.00  0.00           H
    ATOM   2167 HD13 LEU B 532     157.791  54.317  99.209  1.00  0.00           H
    ATOM   2168 HD21 LEU B 532     156.312  56.856 101.126  1.00  0.00           H
    ATOM   2169 HD22 LEU B 532     157.923  57.384 100.570  1.00  0.00           H
    ATOM   2170 HD23 LEU B 532     156.437  58.175 100.025  1.00  0.00           H
    ATOM   2171  N   GLU B 533     159.389  57.597  95.655  1.00 99.90           N
    ATOM   2172  CA  GLU B 533     160.601  57.865  94.868  1.00 99.90           C
    ATOM   2173  C   GLU B 533     160.462  59.049  93.887  1.00 99.90           C
    ATOM   2174  O   GLU B 533     161.214  59.141  92.919  1.00 99.90           O
    ATOM   2175  CB  GLU B 533     161.080  56.571  94.179  1.00 99.90           C
    ATOM   2176  CG  GLU B 533     161.512  55.498  95.193  1.00 99.90           C
    ATOM   2177  CD  GLU B 533     162.025  54.210  94.551  1.00 99.90           C
    ATOM   2178  OE1 GLU B 533     162.653  54.292  93.473  1.00 99.90           O
    ATOM   2179  OE2 GLU B 533     161.784  53.153  95.175  1.00 99.90           O1-
    ATOM   2180  H   GLU B 533     158.825  56.818  95.332  1.00  0.00           H
    ATOM   2181  HA  GLU B 533     161.385  58.168  95.565  1.00  0.00           H
    ATOM   2182  HB2 GLU B 533     160.285  56.178  93.543  1.00  0.00           H
    ATOM   2183  HB3 GLU B 533     161.920  56.789  93.518  1.00  0.00           H
    ATOM   2184  HG2 GLU B 533     162.294  55.889  95.844  1.00  0.00           H
    ATOM   2185  HG3 GLU B 533     160.663  55.247  95.829  1.00  0.00           H
    ATOM   2186  N   ALA B 534     159.516  59.946  94.184  1.00 99.90           N
    ATOM   2187  CA  ALA B 534     159.286  61.249  93.580  1.00 99.90           C
    ATOM   2188  C   ALA B 534     159.216  62.309  94.690  1.00 99.90           C
    ATOM   2189  O   ALA B 534     158.737  63.413  94.437  1.00 99.90           O
    ATOM   2190  CB  ALA B 534     157.994  61.227  92.743  1.00 99.90           C
    ATOM   2191  H   ALA B 534     158.909  59.739  94.965  1.00  0.00           H
    ATOM   2192  HA  ALA B 534     160.123  61.514  92.932  1.00  0.00           H
    ATOM   2193  HB1 ALA B 534     157.807  62.196  92.280  1.00  0.00           H
    ATOM   2194  HB2 ALA B 534     158.067  60.507  91.928  1.00  0.00           H
    ATOM   2195  HB3 ALA B 534     157.121  60.982  93.348  1.00  0.00           H
    ATOM   2196  N   GLY B 535     159.700  61.963  95.895  1.00 89.08           N
    ATOM   2197  CA  GLY B 535     159.722  62.816  97.073  1.00 89.08           C
    ATOM   2198  C   GLY B 535     158.309  62.979  97.637  1.00 89.08           C
    ATOM   2199  O   GLY B 535     157.727  64.055  97.513  1.00 89.08           O
    ATOM   2200  H   GLY B 535     160.055  61.025  96.020  1.00  0.00           H
    ATOM   2201  HA2 GLY B 535     160.366  62.364  97.828  1.00  0.00           H
    ATOM   2202  HA3 GLY B 535     160.147  63.790  96.828  1.00  0.00           H
    ATOM   2203  N   ALA B 536     157.775  61.926  98.272  1.00 89.12           N
    ATOM   2204  CA  ALA B 536     156.552  62.000  99.075  1.00 89.12           C
    ATOM   2205  C   ALA B 536     156.849  61.417 100.445  1.00 89.12           C
    ATOM   2206  O   ALA B 536     157.596  60.443 100.532  1.00 89.12           O
    ATOM   2207  CB  ALA B 536     155.405  61.224  98.429  1.00 99.90           C
    ATOM   2208  H   ALA B 536     158.285  61.055  98.319  1.00  0.00           H
    ATOM   2209  HA  ALA B 536     156.235  63.036  99.207  1.00  0.00           H
    ATOM   2210  HB1 ALA B 536     154.490  61.320  99.015  1.00  0.00           H
    ATOM   2211  HB2 ALA B 536     155.187  61.626  97.443  1.00  0.00           H
    ATOM   2212  HB3 ALA B 536     155.619  60.159  98.348  1.00  0.00           H
    ATOM   2213  N   ASN B 537     156.265  62.029 101.486  1.00 83.48           N
    ATOM   2214  CA  ASN B 537     156.581  61.661 102.860  1.00 83.48           C
    ATOM   2215  C   ASN B 537     155.911  60.335 103.277  1.00 83.48           C
    ATOM   2216  O   ASN B 537     154.680  60.288 103.344  1.00 83.48           O
    ATOM   2217  CB  ASN B 537     156.219  62.801 103.838  1.00 99.90           C
    ATOM   2218  CG  ASN B 537     157.447  63.242 104.641  1.00 99.90           C
    ATOM   2219  OD1 ASN B 537     158.244  62.413 105.075  1.00 99.90           O
    ATOM   2220  ND2 ASN B 537     157.601  64.544 104.867  1.00 99.90           N
    ATOM   2221  H   ASN B 537     155.645  62.813 101.346  1.00  0.00           H
    ATOM   2222  HA  ASN B 537     157.662  61.566 102.847  1.00  0.00           H
    ATOM   2223  HB2 ASN B 537     155.800  63.657 103.306  1.00  0.00           H
    ATOM   2224  HB3 ASN B 537     155.461  62.494 104.558  1.00  0.00           H
    ATOM   2225 HD21 ASN B 537     156.934  65.212 104.509  1.00  0.00           H
    ATOM   2226 HD22 ASN B 537     158.397  64.873 105.394  1.00  0.00           H
    ATOM   2227  N   PRO B 538     156.733  59.299 103.579  1.00 68.86           N
    ATOM   2228  CA  PRO B 538     156.259  57.970 104.013  1.00 68.86           C
    ATOM   2229  C   PRO B 538     155.783  57.924 105.481  1.00 68.86           C
    ATOM   2230  O   PRO B 538     156.061  56.942 106.164  1.00 68.86           O
    ATOM   2231  CB  PRO B 538     157.511  57.090 103.805  1.00 52.60           C
    ATOM   2232  CG  PRO B 538     158.670  58.011 104.114  1.00 52.60           C
    ATOM   2233  CD  PRO B 538     158.204  59.330 103.542  1.00 52.60           C
    ATOM   2234  HA  PRO B 538     155.444  57.616 103.380  1.00  0.00           H
    ATOM   2235  HB2 PRO B 538     157.539  56.171 104.391  1.00  0.00           H
    ATOM   2236  HB3 PRO B 538     157.601  56.764 102.777  1.00  0.00           H
    ATOM   2237  HG2 PRO B 538     158.795  58.105 105.190  1.00  0.00           H
    ATOM   2238  HG3 PRO B 538     159.620  57.678 103.700  1.00  0.00           H
    ATOM   2239  HD2 PRO B 538     158.642  60.139 104.122  1.00  0.00           H
    ATOM   2240  HD3 PRO B 538     158.555  59.407 102.516  1.00  0.00           H
    ATOM   2241  N   ASN B 539     155.122  58.984 105.971  1.00 78.23           N
    ATOM   2242  CA  ASN B 539     154.812  59.136 107.395  1.00 78.23           C
    ATOM   2243  C   ASN B 539     153.740  60.180 107.700  1.00 78.23           C
    ATOM   2244  O   ASN B 539     153.326  60.269 108.856  1.00 78.23           O
    ATOM   2245  CB  ASN B 539     156.102  59.384 108.224  1.00 99.90           C
    ATOM   2246  CG  ASN B 539     157.169  60.290 107.614  1.00 99.90           C
    ATOM   2247  OD1 ASN B 539     158.007  59.829 106.844  1.00 99.90           O
    ATOM   2248  ND2 ASN B 539     157.168  61.569 107.976  1.00 99.90           N
    ATOM   2249  H   ASN B 539     154.918  59.765 105.366  1.00  0.00           H
    ATOM   2250  HA  ASN B 539     154.394  58.174 107.695  1.00  0.00           H
    ATOM   2251  HB2 ASN B 539     155.903  59.628 109.268  1.00  0.00           H
    ATOM   2252  HB3 ASN B 539     156.632  58.449 108.226  1.00  0.00           H
    ATOM   2253 HD21 ASN B 539     156.484  61.921 108.629  1.00  0.00           H
    ATOM   2254 HD22 ASN B 539     157.860  62.194 107.588  1.00  0.00           H
    ATOM   2255  N   ALA B 540     153.309  60.960 106.697  1.00 84.80           N
    ATOM   2256  CA  ALA B 540     152.414  62.091 106.921  1.00 84.80           C
    ATOM   2257  C   ALA B 540     150.960  61.662 107.211  1.00 84.80           C
    ATOM   2258  O   ALA B 540     150.500  60.681 106.630  1.00 84.80           O
    ATOM   2259  CB  ALA B 540     152.422  62.999 105.695  1.00 79.24           C
    ATOM   2260  H   ALA B 540     153.642  60.826 105.754  1.00  0.00           H
    ATOM   2261  HA  ALA B 540     152.868  62.651 107.734  1.00  0.00           H
    ATOM   2262  HB1 ALA B 540     152.074  64.000 105.951  1.00  0.00           H
    ATOM   2263  HB2 ALA B 540     153.413  63.101 105.259  1.00  0.00           H
    ATOM   2264  HB3 ALA B 540     151.740  62.614 104.942  1.00  0.00           H
    ATOM   2265  N   PRO B 541     150.238  62.452 108.037  1.00 74.06           N
    ATOM   2266  CA  PRO B 541     148.768  62.369 108.144  1.00 74.06           C
    ATOM   2267  C   PRO B 541     148.098  62.733 106.804  1.00 74.06           C
    ATOM   2268  O   PRO B 541     148.570  63.649 106.131  1.00 74.06           O
    ATOM   2269  CB  PRO B 541     148.426  63.395 109.244  1.00 99.90           C
    ATOM   2270  CG  PRO B 541     149.734  63.746 109.924  1.00 99.90           C
    ATOM   2271  CD  PRO B 541     150.762  63.562 108.832  1.00 99.90           C
    ATOM   2272  HA  PRO B 541     148.492  61.360 108.459  1.00  0.00           H
    ATOM   2273  HB2 PRO B 541     147.981  64.299 108.824  1.00  0.00           H
    ATOM   2274  HB3 PRO B 541     147.730  62.995 109.978  1.00  0.00           H
    ATOM   2275  HG2 PRO B 541     149.739  64.746 110.360  1.00  0.00           H
    ATOM   2276  HG3 PRO B 541     149.926  63.031 110.725  1.00  0.00           H
    ATOM   2277  HD2 PRO B 541     150.845  64.452 108.207  1.00  0.00           H
    ATOM   2278  HD3 PRO B 541     151.724  63.365 109.304  1.00  0.00           H
    ATOM   2279  N   ASN B 542     147.040  62.010 106.420  1.00 66.71           N
    ATOM   2280  CA  ASN B 542     146.453  62.094 105.077  1.00 66.71           C
    ATOM   2281  C   ASN B 542     145.328  63.131 104.883  1.00 66.71           C
    ATOM   2282  O   ASN B 542     144.452  62.870 104.065  1.00 66.71           O
    ATOM   2283  CB  ASN B 542     145.970  60.692 104.652  1.00 78.42           C
    ATOM   2284  CG  ASN B 542     144.707  60.197 105.372  1.00 78.42           C
    ATOM   2285  OD1 ASN B 542     144.515  60.446 106.560  1.00 78.42           O
    ATOM   2286  ND2 ASN B 542     143.840  59.503 104.644  1.00 78.42           N
    ATOM   2287  H   ASN B 542     146.692  61.259 106.999  1.00  0.00           H
    ATOM   2288  HA  ASN B 542     147.213  62.392 104.353  1.00  0.00           H
    ATOM   2289  HB2 ASN B 542     145.764  60.713 103.582  1.00  0.00           H
    ATOM   2290  HB3 ASN B 542     146.772  59.970 104.787  1.00  0.00           H
    ATOM   2291 HD21 ASN B 542     144.010  59.345 103.662  1.00  0.00           H
    ATOM   2292 HD22 ASN B 542     143.010  59.116 105.076  1.00  0.00           H
    ATOM   2293  N   SER B 543     145.283  64.237 105.645  1.00 70.96           N
    ATOM   2294  CA  SER B 543     144.159  65.200 105.662  1.00 70.96           C
    ATOM   2295  C   SER B 543     142.883  64.656 106.347  1.00 70.96           C
    ATOM   2296  O   SER B 543     142.036  65.440 106.768  1.00 70.96           O
    ATOM   2297  CB  SER B 543     143.868  65.800 104.254  1.00 50.74           C
    ATOM   2298  OG  SER B 543     142.904  65.093 103.490  1.00 50.74           O
    ATOM   2299  H   SER B 543     146.035  64.417 106.295  1.00  0.00           H
    ATOM   2300  HA  SER B 543     144.516  66.018 106.289  1.00  0.00           H
    ATOM   2301  HB2 SER B 543     143.485  66.814 104.375  1.00  0.00           H
    ATOM   2302  HB3 SER B 543     144.788  65.895 103.676  1.00  0.00           H
    ATOM   2303  HG  SER B 543     143.372  64.460 102.933  1.00  0.00           H
    ATOM   2304  N   TYR B 544     142.799  63.326 106.473  1.00 60.54           N
    ATOM   2305  CA  TYR B 544     141.858  62.569 107.284  1.00 60.54           C
    ATOM   2306  C   TYR B 544     142.558  62.064 108.555  1.00 60.54           C
    ATOM   2307  O   TYR B 544     141.944  61.304 109.305  1.00 60.54           O
    ATOM   2308  CB  TYR B 544     141.293  61.411 106.433  1.00 65.65           C
    ATOM   2309  CG  TYR B 544     140.306  61.865 105.381  1.00 65.65           C
    ATOM   2310  CD1 TYR B 544     140.734  62.528 104.211  1.00 65.65           C
    ATOM   2311  CD2 TYR B 544     138.933  61.685 105.613  1.00 65.65           C
    ATOM   2312  CE1 TYR B 544     139.786  63.062 103.319  1.00 65.65           C
    ATOM   2313  CE2 TYR B 544     137.986  62.195 104.711  1.00 65.65           C
    ATOM   2314  CZ  TYR B 544     138.412  62.913 103.585  1.00 65.65           C
    ATOM   2315  OH  TYR B 544     137.489  63.515 102.788  1.00 65.65           O
    ATOM   2316  H   TYR B 544     143.509  62.767 106.021  1.00  0.00           H
    ATOM   2317  HA  TYR B 544     141.032  63.207 107.602  1.00  0.00           H
    ATOM   2318  HB2 TYR B 544     142.078  60.849 105.947  1.00  0.00           H
    ATOM   2319  HB3 TYR B 544     140.780  60.690 107.072  1.00  0.00           H
    ATOM   2320  HD1 TYR B 544     141.786  62.662 104.014  1.00  0.00           H
    ATOM   2321  HD2 TYR B 544     138.591  61.223 106.522  1.00  0.00           H
    ATOM   2322  HE1 TYR B 544     140.121  63.607 102.448  1.00  0.00           H
    ATOM   2323  HE2 TYR B 544     136.931  62.107 104.919  1.00  0.00           H
    ATOM   2324  HH  TYR B 544     137.896  64.004 102.068  1.00  0.00           H
    ATOM   2325  N   GLY B 545     143.814  62.507 108.771  1.00 63.23           N
    ATOM   2326  CA  GLY B 545     144.689  62.188 109.889  1.00 63.23           C
    ATOM   2327  C   GLY B 545     144.792  60.682 110.074  1.00 63.23           C
    ATOM   2328  O   GLY B 545     144.050  60.137 110.874  1.00 63.23           O
    ATOM   2329  H   GLY B 545     144.201  63.130 108.079  1.00  0.00           H
    ATOM   2330  HA2 GLY B 545     145.665  62.614 109.687  1.00  0.00           H
    ATOM   2331  HA3 GLY B 545     144.310  62.655 110.799  1.00  0.00           H
    ATOM   2332  N   ARG B 546     145.671  60.038 109.307  1.00 49.99           N
    ATOM   2333  CA  ARG B 546     145.898  58.602 109.227  1.00 49.99           C
    ATOM   2334  C   ARG B 546     146.968  58.500 108.156  1.00 49.99           C
    ATOM   2335  O   ARG B 546     146.716  58.861 107.017  1.00 49.99           O
    ATOM   2336  CB  ARG B 546     144.632  57.813 108.778  1.00 49.60           C
    ATOM   2337  CG  ARG B 546     144.282  56.643 109.709  1.00 49.60           C
    ATOM   2338  CD  ARG B 546     143.731  57.116 111.051  1.00 49.60           C
    ATOM   2339  NE  ARG B 546     142.464  57.839 110.910  1.00 49.60           N
    ATOM   2340  CZ  ARG B 546     141.408  57.850 111.732  1.00 49.60           C
    ATOM   2341  NH1 ARG B 546     141.233  56.955 112.708  1.00 49.60           N
    ATOM   2342  NH2 ARG B 546     140.508  58.814 111.547  1.00 49.60           N1+
    ATOM   2343  H   ARG B 546     146.206  60.577 108.644  1.00  0.00           H
    ATOM   2344  HA  ARG B 546     146.287  58.247 110.182  1.00  0.00           H
    ATOM   2345  HB2 ARG B 546     143.771  58.473 108.664  1.00  0.00           H
    ATOM   2346  HB3 ARG B 546     144.763  57.390 107.780  1.00  0.00           H
    ATOM   2347  HG2 ARG B 546     143.502  56.080 109.219  1.00  0.00           H
    ATOM   2348  HG3 ARG B 546     145.096  55.931 109.832  1.00  0.00           H
    ATOM   2349  HD2 ARG B 546     143.755  56.369 111.824  1.00  0.00           H
    ATOM   2350  HD3 ARG B 546     144.425  57.852 111.435  1.00  0.00           H
    ATOM   2351  HE  ARG B 546     142.501  58.578 110.222  1.00  0.00           H
    ATOM   2352 HH11 ARG B 546     141.798  56.108 112.704  1.00  0.00           H
    ATOM   2353 HH12 ARG B 546     140.423  57.003 113.307  1.00  0.00           H
    ATOM   2354 HH21 ARG B 546     140.649  59.495 110.814  1.00  0.00           H
    ATOM   2355 HH22 ARG B 546     139.697  58.935 112.153  1.00  0.00           H
    ATOM   2356  N   ARG B 547     148.159  58.024 108.491  1.00 52.50           N
    ATOM   2357  CA  ARG B 547     149.191  57.784 107.493  1.00 52.50           C
    ATOM   2358  C   ARG B 547     148.803  56.626 106.553  1.00 52.50           C
    ATOM   2359  O   ARG B 547     147.922  55.841 106.896  1.00 52.50           O
    ATOM   2360  CB  ARG B 547     150.585  57.683 108.165  1.00 99.90           C
    ATOM   2361  CG  ARG B 547     150.688  57.214 109.631  1.00 99.90           C
    ATOM   2362  CD  ARG B 547     151.889  57.882 110.307  1.00 99.90           C
    ATOM   2363  NE  ARG B 547     152.235  57.324 111.616  1.00 99.90           N
    ATOM   2364  CZ  ARG B 547     151.564  57.431 112.774  1.00 99.90           C
    ATOM   2365  NH1 ARG B 547     150.387  58.068 112.857  1.00 99.90           N
    ATOM   2366  NH2 ARG B 547     152.104  56.902 113.880  1.00 99.90           N1+
    ATOM   2367  H   ARG B 547     148.318  57.655 109.419  1.00  0.00           H
    ATOM   2368  HA  ARG B 547     149.207  58.665 106.857  1.00  0.00           H
    ATOM   2369  HB2 ARG B 547     151.289  57.100 107.572  1.00  0.00           H
    ATOM   2370  HB3 ARG B 547     150.980  58.698 108.145  1.00  0.00           H
    ATOM   2371  HG2 ARG B 547     149.843  57.603 110.192  1.00  0.00           H
    ATOM   2372  HG3 ARG B 547     150.655  56.129 109.747  1.00  0.00           H
    ATOM   2373  HD2 ARG B 547     152.757  57.873 109.648  1.00  0.00           H
    ATOM   2374  HD3 ARG B 547     151.654  58.928 110.507  1.00  0.00           H
    ATOM   2375  HE  ARG B 547     153.101  56.802 111.619  1.00  0.00           H
    ATOM   2376 HH11 ARG B 547     149.985  58.481 112.029  1.00  0.00           H
    ATOM   2377 HH12 ARG B 547     149.910  58.161 113.742  1.00  0.00           H
    ATOM   2378 HH21 ARG B 547     153.004  56.444 113.845  1.00  0.00           H
    ATOM   2379 HH22 ARG B 547     151.639  56.995 114.771  1.00  0.00           H
    ATOM   2380  N   PRO B 548     149.498  56.468 105.403  1.00 67.17           N
    ATOM   2381  CA  PRO B 548     149.530  55.169 104.690  1.00 67.17           C
    ATOM   2382  C   PRO B 548     150.091  54.085 105.618  1.00 67.17           C
    ATOM   2383  O   PRO B 548     150.534  54.409 106.718  1.00 67.17           O
    ATOM   2384  CB  PRO B 548     150.514  55.428 103.550  1.00 37.03           C
    ATOM   2385  CG  PRO B 548     151.501  56.370 104.218  1.00 37.03           C
    ATOM   2386  CD  PRO B 548     150.551  57.343 104.884  1.00 37.03           C
    ATOM   2387  HA  PRO B 548     148.569  54.852 104.295  1.00  0.00           H
    ATOM   2388  HB2 PRO B 548     150.992  54.533 103.156  1.00  0.00           H
    ATOM   2389  HB3 PRO B 548     150.016  55.921 102.715  1.00  0.00           H
    ATOM   2390  HG2 PRO B 548     152.088  55.845 104.974  1.00  0.00           H
    ATOM   2391  HG3 PRO B 548     152.233  56.846 103.592  1.00  0.00           H
    ATOM   2392  HD2 PRO B 548     151.075  57.965 105.606  1.00  0.00           H
    ATOM   2393  HD3 PRO B 548     150.108  58.016 104.148  1.00  0.00           H
    ATOM   2394  N   ILE B 549     150.064  52.816 105.205  1.00 43.13           N
    ATOM   2395  CA  ILE B 549     150.536  51.720 106.066  1.00 43.13           C
    ATOM   2396  C   ILE B 549     149.658  51.565 107.336  1.00 43.13           C
    ATOM   2397  O   ILE B 549     150.042  50.972 108.335  1.00 43.13           O
    ATOM   2398  CB  ILE B 549     152.075  51.832 106.383  1.00 46.00           C
    ATOM   2399  CG1 ILE B 549     152.755  50.459 106.274  1.00 46.00           C
    ATOM   2400  CG2 ILE B 549     152.561  52.566 107.670  1.00 46.00           C
    ATOM   2401  CD1 ILE B 549     152.413  49.495 107.409  1.00 46.00           C
    ATOM   2402  H   ILE B 549     149.722  52.574 104.287  1.00  0.00           H
    ATOM   2403  HA  ILE B 549     150.365  50.825 105.485  1.00  0.00           H
    ATOM   2404  HB  ILE B 549     152.519  52.404 105.568  1.00  0.00           H
    ATOM   2405 HG12 ILE B 549     152.498  49.992 105.323  1.00  0.00           H
    ATOM   2406 HG13 ILE B 549     153.837  50.594 106.233  1.00  0.00           H
    ATOM   2407 HG21 ILE B 549     153.606  52.317 107.860  1.00  0.00           H
    ATOM   2408 HG22 ILE B 549     152.572  53.648 107.574  1.00  0.00           H
    ATOM   2409 HG23 ILE B 549     152.032  52.315 108.585  1.00  0.00           H
    ATOM   2410 HD11 ILE B 549     153.199  48.754 107.485  1.00  0.00           H
    ATOM   2411 HD12 ILE B 549     152.380  49.983 108.381  1.00  0.00           H
    ATOM   2412 HD13 ILE B 549     151.461  48.994 107.241  1.00  0.00           H
    ATOM   2413  N   GLN B 550     148.460  52.129 107.249  1.00 93.05           N
    ATOM   2414  CA  GLN B 550     147.472  52.283 108.292  1.00 93.05           C
    ATOM   2415  C   GLN B 550     146.153  52.301 107.528  1.00 93.05           C
    ATOM   2416  O   GLN B 550     145.214  51.609 107.910  1.00 93.05           O
    ATOM   2417  CB  GLN B 550     147.715  53.631 108.994  1.00 71.66           C
    ATOM   2418  CG  GLN B 550     147.304  53.682 110.457  1.00 71.66           C
    ATOM   2419  CD  GLN B 550     147.545  55.068 111.052  1.00 71.66           C
    ATOM   2420  OE1 GLN B 550     147.785  56.040 110.349  1.00 71.66           O
    ATOM   2421  NE2 GLN B 550     147.467  55.173 112.373  1.00 71.66           N
    ATOM   2422  H   GLN B 550     148.235  52.592 106.383  1.00  0.00           H
    ATOM   2423  HA  GLN B 550     147.503  51.444 108.991  1.00  0.00           H
    ATOM   2424  HB2 GLN B 550     148.764  53.925 108.912  1.00  0.00           H
    ATOM   2425  HB3 GLN B 550     147.126  54.399 108.504  1.00  0.00           H
    ATOM   2426  HG2 GLN B 550     146.254  53.424 110.551  1.00  0.00           H
    ATOM   2427  HG3 GLN B 550     147.865  52.939 111.026  1.00  0.00           H
    ATOM   2428 HE21 GLN B 550     147.250  54.361 112.934  1.00  0.00           H
    ATOM   2429 HE22 GLN B 550     147.598  56.066 112.825  1.00  0.00           H
    ATOM   2430  N   VAL B 551     146.154  53.066 106.423  1.00 51.04           N
    ATOM   2431  CA  VAL B 551     145.081  53.116 105.461  1.00 51.04           C
    ATOM   2432  C   VAL B 551     145.376  52.303 104.196  1.00 51.04           C
    ATOM   2433  O   VAL B 551     144.499  52.244 103.336  1.00 51.04           O
    ATOM   2434  CB  VAL B 551     144.782  54.582 105.081  1.00 39.27           C
    ATOM   2435  CG1 VAL B 551     145.796  55.282 104.155  1.00 39.27           C
    ATOM   2436  CG2 VAL B 551     143.373  54.699 104.512  1.00 39.27           C
    ATOM   2437  H   VAL B 551     146.941  53.670 106.234  1.00  0.00           H
    ATOM   2438  HA  VAL B 551     144.186  52.677 105.906  1.00  0.00           H
    ATOM   2439  HB  VAL B 551     144.762  55.148 106.015  1.00  0.00           H
    ATOM   2440 HG11 VAL B 551     145.341  56.182 103.746  1.00  0.00           H
    ATOM   2441 HG12 VAL B 551     146.660  55.610 104.718  1.00  0.00           H
    ATOM   2442 HG13 VAL B 551     146.126  54.686 103.305  1.00  0.00           H
    ATOM   2443 HG21 VAL B 551     142.949  55.652 104.803  1.00  0.00           H
    ATOM   2444 HG22 VAL B 551     143.347  54.600 103.429  1.00  0.00           H
    ATOM   2445 HG23 VAL B 551     142.705  53.951 104.928  1.00  0.00           H
    ATOM   2446  N   MET B 552     146.560  51.667 104.104  1.00 54.85           N
    ATOM   2447  CA  MET B 552     146.933  50.852 102.955  1.00 54.85           C
    ATOM   2448  C   MET B 552     145.932  49.726 102.682  1.00 54.85           C
    ATOM   2449  O   MET B 552     145.197  49.321 103.579  1.00 54.85           O
    ATOM   2450  CB  MET B 552     148.360  50.296 103.102  1.00 47.26           C
    ATOM   2451  CG  MET B 552     148.461  49.141 104.098  1.00 47.26           C
    ATOM   2452  SD  MET B 552     150.099  48.458 104.304  1.00 47.26           S
    ATOM   2453  CE  MET B 552     149.632  47.151 105.458  1.00 47.26           C
    ATOM   2454  H   MET B 552     147.220  51.715 104.865  1.00  0.00           H
    ATOM   2455  HA  MET B 552     146.920  51.556 102.146  1.00  0.00           H
    ATOM   2456  HB2 MET B 552     148.710  49.949 102.128  1.00  0.00           H
    ATOM   2457  HB3 MET B 552     149.045  51.090 103.393  1.00  0.00           H
    ATOM   2458  HG2 MET B 552     148.124  49.478 105.077  1.00  0.00           H
    ATOM   2459  HG3 MET B 552     147.825  48.307 103.805  1.00  0.00           H
    ATOM   2460  HE1 MET B 552     150.413  46.405 105.557  1.00  0.00           H
    ATOM   2461  HE2 MET B 552     149.396  47.568 106.436  1.00  0.00           H
    ATOM   2462  HE3 MET B 552     148.759  46.621 105.080  1.00  0.00           H
    ATOM   2463  N   MET B 553     145.969  49.186 101.465  1.00 46.06           N
    ATOM   2464  CA  MET B 553     145.322  47.929 101.185  1.00 46.06           C
    ATOM   2465  C   MET B 553     146.132  46.835 101.875  1.00 46.06           C
    ATOM   2466  O   MET B 553     147.164  46.437 101.343  1.00 46.06           O
    ATOM   2467  CB  MET B 553     145.189  47.726  99.670  1.00 49.33           C
    ATOM   2468  CG  MET B 553     144.651  46.336  99.299  1.00 49.33           C
    ATOM   2469  SD  MET B 553     143.508  46.294  97.904  1.00 49.33           S
    ATOM   2470  CE  MET B 553     144.456  47.090  96.625  1.00 49.33           C
    ATOM   2471  H   MET B 553     146.563  49.575 100.743  1.00  0.00           H
    ATOM   2472  HA  MET B 553     144.315  47.953 101.591  1.00  0.00           H
    ATOM   2473  HB2 MET B 553     144.534  48.493  99.266  1.00  0.00           H
    ATOM   2474  HB3 MET B 553     146.158  47.873  99.193  1.00  0.00           H
    ATOM   2475  HG2 MET B 553     145.448  45.613  99.144  1.00  0.00           H
    ATOM   2476  HG3 MET B 553     144.069  45.955 100.134  1.00  0.00           H
    ATOM   2477  HE1 MET B 553     143.882  47.174  95.705  1.00  0.00           H
    ATOM   2478  HE2 MET B 553     144.643  48.081  96.994  1.00  0.00           H
    ATOM   2479  HE3 MET B 553     145.384  46.562  96.432  1.00  0.00           H
    ATOM   2480  N   MET B 554     145.644  46.343 103.029  1.00 31.74           N
    ATOM   2481  CA  MET B 554     146.092  45.073 103.611  1.00 31.74           C
    ATOM   2482  C   MET B 554     145.976  43.996 102.522  1.00 31.74           C
    ATOM   2483  O   MET B 554     145.101  44.128 101.691  1.00 31.74           O
    ATOM   2484  CB  MET B 554     145.199  44.692 104.802  1.00 57.75           C
    ATOM   2485  CG  MET B 554     144.952  45.779 105.844  1.00 57.75           C
    ATOM   2486  SD  MET B 554     146.421  46.415 106.682  1.00 57.75           S
    ATOM   2487  CE  MET B 554     146.073  48.187 106.615  1.00 57.75           C
    ATOM   2488  H   MET B 554     144.773  46.717 103.389  1.00  0.00           H
    ATOM   2489  HA  MET B 554     147.115  45.189 103.955  1.00  0.00           H
    ATOM   2490  HB2 MET B 554     144.214  44.399 104.436  1.00  0.00           H
    ATOM   2491  HB3 MET B 554     145.625  43.821 105.298  1.00  0.00           H
    ATOM   2492  HG2 MET B 554     144.409  46.615 105.410  1.00  0.00           H
    ATOM   2493  HG3 MET B 554     144.289  45.358 106.593  1.00  0.00           H
    ATOM   2494  HE1 MET B 554     146.830  48.737 107.174  1.00  0.00           H
    ATOM   2495  HE2 MET B 554     146.096  48.527 105.592  1.00  0.00           H
    ATOM   2496  HE3 MET B 554     145.099  48.412 107.048  1.00  0.00           H
    ATOM   2497  N   GLY B 555     146.849  42.999 102.424  1.00 60.05           N
    ATOM   2498  CA  GLY B 555     146.783  42.069 101.287  1.00 60.05           C
    ATOM   2499  C   GLY B 555     147.296  42.670  99.970  1.00 60.05           C
    ATOM   2500  O   GLY B 555     147.334  41.954  98.972  1.00 60.05           O
    ATOM   2501  H   GLY B 555     147.574  42.869 103.120  1.00  0.00           H
    ATOM   2502  HA2 GLY B 555     147.412  41.208 101.505  1.00  0.00           H
    ATOM   2503  HA3 GLY B 555     145.771  41.692 101.133  1.00  0.00           H
    ATOM   2504  N   SER B 556     147.752  43.932  99.971  1.00 43.70           N
    ATOM   2505  CA  SER B 556     148.720  44.433  99.009  1.00 43.70           C
    ATOM   2506  C   SER B 556     150.050  44.419  99.759  1.00 43.70           C
    ATOM   2507  O   SER B 556     150.555  45.469 100.150  1.00 43.70           O
    ATOM   2508  CB  SER B 556     148.329  45.824  98.476  1.00 99.76           C
    ATOM   2509  OG  SER B 556     149.181  46.170  97.407  1.00 99.76           O
    ATOM   2510  H   SER B 556     147.623  44.505 100.795  1.00  0.00           H
    ATOM   2511  HA  SER B 556     148.805  43.762  98.152  1.00  0.00           H
    ATOM   2512  HB2 SER B 556     147.308  45.821  98.107  1.00  0.00           H
    ATOM   2513  HB3 SER B 556     148.394  46.603  99.233  1.00  0.00           H
    ATOM   2514  HG  SER B 556     148.917  47.029  97.070  1.00  0.00           H
    ATOM   2515  N   ALA B 557     150.555  43.196  99.994  1.00 51.18           N
    ATOM   2516  CA  ALA B 557     151.817  42.907 100.672  1.00 51.18           C
    ATOM   2517  C   ALA B 557     152.977  43.683 100.073  1.00 51.18           C
    ATOM   2518  O   ALA B 557     153.727  44.293 100.828  1.00 51.18           O
    ATOM   2519  CB  ALA B 557     152.089  41.409 100.608  1.00 99.90           C
    ATOM   2520  H   ALA B 557     150.046  42.391  99.659  1.00  0.00           H
    ATOM   2521  HA  ALA B 557     151.737  43.181 101.717  1.00  0.00           H
    ATOM   2522  HB1 ALA B 557     152.890  41.116 101.287  1.00  0.00           H
    ATOM   2523  HB2 ALA B 557     151.188  40.876 100.894  1.00  0.00           H
    ATOM   2524  HB3 ALA B 557     152.354  41.076  99.603  1.00  0.00           H
    ATOM   2525  N   ARG B 558     153.024  43.725  98.732  1.00 61.00           N
    ATOM   2526  CA  ARG B 558     153.946  44.560  97.981  1.00 61.00           C
    ATOM   2527  C   ARG B 558     153.897  46.036  98.400  1.00 61.00           C
    ATOM   2528  O   ARG B 558     154.966  46.600  98.561  1.00 61.00           O
    ATOM   2529  CB  ARG B 558     153.698  44.414  96.474  1.00 99.90           C
    ATOM   2530  CG  ARG B 558     154.244  43.114  95.859  1.00 99.90           C
    ATOM   2531  CD  ARG B 558     154.138  43.116  94.326  1.00 99.90           C
    ATOM   2532  NE  ARG B 558     154.987  44.135  93.693  1.00 99.90           N
    ATOM   2533  CZ  ARG B 558     154.838  44.614  92.444  1.00 99.90           C
    ATOM   2534  NH1 ARG B 558     153.959  44.091  91.581  1.00 99.90           N
    ATOM   2535  NH2 ARG B 558     155.583  45.651  92.049  1.00 99.90           N1+
    ATOM   2536  H   ARG B 558     152.347  43.206  98.192  1.00  0.00           H
    ATOM   2537  HA  ARG B 558     154.950  44.191  98.202  1.00  0.00           H
    ATOM   2538  HB2 ARG B 558     152.639  44.543  96.244  1.00  0.00           H
    ATOM   2539  HB3 ARG B 558     154.225  45.228  95.985  1.00  0.00           H
    ATOM   2540  HG2 ARG B 558     155.303  43.081  96.115  1.00  0.00           H
    ATOM   2541  HG3 ARG B 558     153.800  42.213  96.284  1.00  0.00           H
    ATOM   2542  HD2 ARG B 558     154.198  42.125  93.876  1.00  0.00           H
    ATOM   2543  HD3 ARG B 558     153.150  43.505  94.108  1.00  0.00           H
    ATOM   2544  HE  ARG B 558     155.722  44.514  94.271  1.00  0.00           H
    ATOM   2545 HH11 ARG B 558     153.353  43.316  91.860  1.00  0.00           H
    ATOM   2546 HH12 ARG B 558     153.868  44.464  90.648  1.00  0.00           H
    ATOM   2547 HH21 ARG B 558     156.240  46.069  92.688  1.00  0.00           H
    ATOM   2548 HH22 ARG B 558     155.446  46.079  91.142  1.00  0.00           H
    ATOM   2549  N   VAL B 559     152.708  46.616  98.635  1.00 27.72           N
    ATOM   2550  CA  VAL B 559     152.567  47.999  99.101  1.00 27.72           C
    ATOM   2551  C   VAL B 559     152.982  48.207 100.560  1.00 27.72           C
    ATOM   2552  O   VAL B 559     153.655  49.198 100.833  1.00 27.72           O
    ATOM   2553  CB  VAL B 559     151.128  48.530  98.921  1.00 29.57           C
    ATOM   2554  CG1 VAL B 559     150.793  49.837  99.664  1.00 29.57           C
    ATOM   2555  CG2 VAL B 559     150.845  48.726  97.434  1.00 29.57           C
    ATOM   2556  H   VAL B 559     151.854  46.085  98.535  1.00  0.00           H
    ATOM   2557  HA  VAL B 559     153.223  48.619  98.486  1.00  0.00           H
    ATOM   2558  HB  VAL B 559     150.436  47.779  99.289  1.00  0.00           H
    ATOM   2559 HG11 VAL B 559     149.778  50.130  99.408  1.00  0.00           H
    ATOM   2560 HG12 VAL B 559     150.799  49.721 100.748  1.00  0.00           H
    ATOM   2561 HG13 VAL B 559     151.470  50.647  99.403  1.00  0.00           H
    ATOM   2562 HG21 VAL B 559     149.782  48.803  97.244  1.00  0.00           H
    ATOM   2563 HG22 VAL B 559     151.293  49.658  97.098  1.00  0.00           H
    ATOM   2564 HG23 VAL B 559     151.228  47.911  96.823  1.00  0.00           H
    ATOM   2565  N   ALA B 560     152.594  47.292 101.464  1.00 56.40           N
    ATOM   2566  CA  ALA B 560     153.035  47.328 102.856  1.00 56.40           C
    ATOM   2567  C   ALA B 560     154.552  47.269 102.942  1.00 56.40           C
    ATOM   2568  O   ALA B 560     155.155  48.180 103.496  1.00 56.40           O
    ATOM   2569  CB  ALA B 560     152.434  46.159 103.626  1.00 50.06           C
    ATOM   2570  H   ALA B 560     152.025  46.506 101.178  1.00  0.00           H
    ATOM   2571  HA  ALA B 560     152.706  48.268 103.299  1.00  0.00           H
    ATOM   2572  HB1 ALA B 560     152.655  46.242 104.686  1.00  0.00           H
    ATOM   2573  HB2 ALA B 560     151.358  46.166 103.499  1.00  0.00           H
    ATOM   2574  HB3 ALA B 560     152.804  45.193 103.280  1.00  0.00           H
    ATOM   2575  N   GLU B 561     155.119  46.253 102.277  1.00 71.58           N
    ATOM   2576  CA  GLU B 561     156.546  46.034 102.107  1.00 71.58           C
    ATOM   2577  C   GLU B 561     157.243  47.249 101.509  1.00 71.58           C
    ATOM   2578  O   GLU B 561     158.295  47.635 102.015  1.00 71.58           O
    ATOM   2579  CB  GLU B 561     156.788  44.770 101.253  1.00 99.90           C
    ATOM   2580  CG  GLU B 561     158.278  44.414 101.112  1.00 99.90           C
    ATOM   2581  CD  GLU B 561     158.540  42.921 101.289  1.00 99.90           C
    ATOM   2582  OE1 GLU B 561     157.987  42.133 100.491  1.00 99.90           O
    ATOM   2583  OE2 GLU B 561     159.276  42.595 102.246  1.00 99.90           O1-
    ATOM   2584  H   GLU B 561     154.522  45.580 101.811  1.00  0.00           H
    ATOM   2585  HA  GLU B 561     156.970  45.875 103.099  1.00  0.00           H
    ATOM   2586  HB2 GLU B 561     156.237  43.918 101.642  1.00  0.00           H
    ATOM   2587  HB3 GLU B 561     156.378  44.922 100.254  1.00  0.00           H
    ATOM   2588  HG2 GLU B 561     158.663  44.741 100.145  1.00  0.00           H
    ATOM   2589  HG3 GLU B 561     158.861  44.937 101.869  1.00  0.00           H
    ATOM   2590  N   LEU B 562     156.610  47.850 100.488  1.00 67.61           N
    ATOM   2591  CA  LEU B 562     157.114  49.033  99.821  1.00 67.61           C
    ATOM   2592  C   LEU B 562     157.256  50.199 100.787  1.00 67.61           C
    ATOM   2593  O   LEU B 562     158.347  50.729 100.854  1.00 67.61           O
    ATOM   2594  CB  LEU B 562     156.285  49.406  98.571  1.00 67.75           C
    ATOM   2595  CG  LEU B 562     156.946  50.446  97.647  1.00 67.75           C
    ATOM   2596  CD1 LEU B 562     156.906  51.872  98.195  1.00 67.75           C
    ATOM   2597  CD2 LEU B 562     158.372  50.030  97.253  1.00 67.75           C
    ATOM   2598  H   LEU B 562     155.753  47.457 100.122  1.00  0.00           H
    ATOM   2599  HA  LEU B 562     158.118  48.757  99.500  1.00  0.00           H
    ATOM   2600  HB2 LEU B 562     156.175  48.523  97.951  1.00  0.00           H
    ATOM   2601  HB3 LEU B 562     155.279  49.725  98.840  1.00  0.00           H
    ATOM   2602  HG  LEU B 562     156.349  50.450  96.733  1.00  0.00           H
    ATOM   2603 HD11 LEU B 562     156.520  52.562  97.458  1.00  0.00           H
    ATOM   2604 HD12 LEU B 562     156.262  51.960  99.067  1.00  0.00           H
    ATOM   2605 HD13 LEU B 562     157.897  52.224  98.478  1.00  0.00           H
    ATOM   2606 HD21 LEU B 562     158.607  50.390  96.252  1.00  0.00           H
    ATOM   2607 HD22 LEU B 562     159.127  50.431  97.929  1.00  0.00           H
    ATOM   2608 HD23 LEU B 562     158.491  48.946  97.234  1.00  0.00           H
    ATOM   2609  N   LEU B 563     156.187  50.568 101.505  1.00 74.59           N
    ATOM   2610  CA  LEU B 563     156.190  51.668 102.467  1.00 74.59           C
    ATOM   2611  C   LEU B 563     157.170  51.411 103.620  1.00 74.59           C
    ATOM   2612  O   LEU B 563     158.027  52.253 103.870  1.00 74.59           O
    ATOM   2613  CB  LEU B 563     154.770  51.866 103.025  1.00 63.22           C
    ATOM   2614  CG  LEU B 563     153.713  52.274 101.981  1.00 63.22           C
    ATOM   2615  CD1 LEU B 563     152.314  51.900 102.487  1.00 63.22           C
    ATOM   2616  CD2 LEU B 563     153.794  53.761 101.608  1.00 63.22           C
    ATOM   2617  H   LEU B 563     155.315  50.067 101.395  1.00  0.00           H
    ATOM   2618  HA  LEU B 563     156.498  52.578 101.949  1.00  0.00           H
    ATOM   2619  HB2 LEU B 563     154.463  50.926 103.486  1.00  0.00           H
    ATOM   2620  HB3 LEU B 563     154.774  52.593 103.840  1.00  0.00           H
    ATOM   2621  HG  LEU B 563     153.871  51.705 101.067  1.00  0.00           H
    ATOM   2622 HD11 LEU B 563     151.585  51.962 101.680  1.00  0.00           H
    ATOM   2623 HD12 LEU B 563     152.276  50.881 102.874  1.00  0.00           H
    ATOM   2624 HD13 LEU B 563     151.997  52.567 103.283  1.00  0.00           H
    ATOM   2625 HD21 LEU B 563     152.809  54.187 101.416  1.00  0.00           H
    ATOM   2626 HD22 LEU B 563     154.258  54.355 102.396  1.00  0.00           H
    ATOM   2627 HD23 LEU B 563     154.394  53.903 100.715  1.00  0.00           H
    ATOM   2628  N   LEU B 564     157.024  50.259 104.296  1.00 73.50           N
    ATOM   2629  CA  LEU B 564     157.810  49.833 105.460  1.00 73.50           C
    ATOM   2630  C   LEU B 564     159.314  49.989 105.273  1.00 73.50           C
    ATOM   2631  O   LEU B 564     159.939  50.760 106.001  1.00 73.50           O
    ATOM   2632  CB  LEU B 564     157.517  48.365 105.747  1.00 29.70           C
    ATOM   2633  CG  LEU B 564     156.162  48.115 106.397  1.00 29.70           C
    ATOM   2634  CD1 LEU B 564     155.885  46.623 106.305  1.00 29.70           C
    ATOM   2635  CD2 LEU B 564     156.076  48.643 107.828  1.00 29.70           C
    ATOM   2636  H   LEU B 564     156.304  49.611 104.000  1.00  0.00           H
    ATOM   2637  HA  LEU B 564     157.530  50.420 106.330  1.00  0.00           H
    ATOM   2638  HB2 LEU B 564     157.580  47.825 104.801  1.00  0.00           H
    ATOM   2639  HB3 LEU B 564     158.285  47.933 106.392  1.00  0.00           H
    ATOM   2640  HG  LEU B 564     155.389  48.639 105.840  1.00  0.00           H
    ATOM   2641 HD11 LEU B 564     154.888  46.473 105.900  1.00  0.00           H
    ATOM   2642 HD12 LEU B 564     156.588  46.109 105.652  1.00  0.00           H
    ATOM   2643 HD13 LEU B 564     155.971  46.132 107.261  1.00  0.00           H
    ATOM   2644 HD21 LEU B 564     155.635  47.898 108.489  1.00  0.00           H
    ATOM   2645 HD22 LEU B 564     157.051  48.922 108.227  1.00  0.00           H
    ATOM   2646 HD23 LEU B 564     155.447  49.533 107.879  1.00  0.00           H
    ATOM   2647  N   LEU B 565     159.837  49.283 104.260  1.00 80.95           N
    ATOM   2648  CA  LEU B 565     161.240  49.271 103.863  1.00 80.95           C
    ATOM   2649  C   LEU B 565     161.538  50.509 102.989  1.00 80.95           C
    ATOM   2650  O   LEU B 565     162.106  50.387 101.904  1.00 80.95           O
    ATOM   2651  CB  LEU B 565     161.542  47.932 103.140  1.00 99.90           C
    ATOM   2652  CG  LEU B 565     161.640  46.659 104.020  1.00 99.90           C
    ATOM   2653  CD1 LEU B 565     160.462  46.417 104.980  1.00 99.90           C
    ATOM   2654  CD2 LEU B 565     161.831  45.429 103.117  1.00 99.90           C
    ATOM   2655  H   LEU B 565     159.224  48.711 103.694  1.00  0.00           H
    ATOM   2656  HA  LEU B 565     161.874  49.338 104.749  1.00  0.00           H
    ATOM   2657  HB2 LEU B 565     160.809  47.779 102.348  1.00  0.00           H
    ATOM   2658  HB3 LEU B 565     162.500  48.013 102.626  1.00  0.00           H
    ATOM   2659  HG  LEU B 565     162.535  46.752 104.637  1.00  0.00           H
    ATOM   2660 HD11 LEU B 565     160.426  45.382 105.322  1.00  0.00           H
    ATOM   2661 HD12 LEU B 565     160.539  47.044 105.868  1.00  0.00           H
    ATOM   2662 HD13 LEU B 565     159.510  46.617 104.490  1.00  0.00           H
    ATOM   2663 HD21 LEU B 565     161.876  44.506 103.696  1.00  0.00           H
    ATOM   2664 HD22 LEU B 565     161.019  45.330 102.398  1.00  0.00           H
    ATOM   2665 HD23 LEU B 565     162.761  45.501 102.552  1.00  0.00           H
    ATOM   2666  N   HIP B 566     161.108  51.687 103.469  1.00 61.99           N
    ATOM   2667  CA  HIP B 566     161.148  52.971 102.770  1.00 61.99           C
    ATOM   2668  C   HIP B 566     160.911  54.132 103.745  1.00 61.99           C
    ATOM   2669  O   HIP B 566     160.563  55.228 103.304  1.00 61.99           O
    ATOM   2670  CB  HIP B 566     160.129  52.937 101.623  1.00 86.03           C
    ATOM   2671  CG  HIP B 566     160.432  53.678 100.368  1.00 86.03           C
    ATOM   2672  ND1 HIP B 566     160.517  55.055 100.302  1.00 86.03           N
    ATOM   2673  CD2 HIP B 566     160.808  53.173  99.141  1.00 86.03           C
    ATOM   2674  CE1 HIP B 566     160.906  55.345  99.063  1.00 86.03           C
    ATOM   2675  NE2 HIP B 566     161.078  54.251  98.323  1.00 86.03           N
    ATOM   2676  H   HIP B 566     160.609  51.686 104.349  1.00  0.00           H
    ATOM   2677  HA  HIP B 566     162.153  53.081 102.358  1.00  0.00           H
    ATOM   2678  HB2 HIP B 566     160.169  51.932 101.242  1.00  0.00           H
    ATOM   2679  HB3 HIP B 566     159.109  53.159 101.925  1.00  0.00           H
    ATOM   2680  HD1 HIP B 566     160.342  55.703 101.060  1.00  0.00           H
    ATOM   2681  HD2 HIP B 566     160.910  52.160  98.780  1.00  0.00           H
    ATOM   2682  HE2 HIP B 566     161.376  54.206  97.357  1.00  0.00           H
    ATOM   2683  HE1 HIP B 566     161.072  56.350  98.700  1.00  0.00           H
    ATOM   2684  N   GLY B 567     161.120  53.875 105.046  1.00 99.90           N
    ATOM   2685  CA  GLY B 567     160.961  54.861 106.102  1.00 99.90           C
    ATOM   2686  C   GLY B 567     159.498  54.942 106.533  1.00 99.90           C
    ATOM   2687  O   GLY B 567     158.976  56.050 106.622  1.00 99.90           O
    ATOM   2688  H   GLY B 567     161.420  52.953 105.327  1.00  0.00           H
    ATOM   2689  HA2 GLY B 567     161.564  54.562 106.959  1.00  0.00           H
    ATOM   2690  HA3 GLY B 567     161.320  55.841 105.784  1.00  0.00           H
    ATOM   2691  N   ALA B 568     158.841  53.799 106.800  1.00 57.69           N
    ATOM   2692  CA  ALA B 568     157.483  53.796 107.336  1.00 57.69           C
    ATOM   2693  C   ALA B 568     157.419  53.025 108.651  1.00 57.69           C
    ATOM   2694  O   ALA B 568     157.360  51.796 108.665  1.00 57.69           O
    ATOM   2695  CB  ALA B 568     156.459  53.306 106.311  1.00 56.73           C
    ATOM   2696  H   ALA B 568     159.293  52.901 106.693  1.00  0.00           H
    ATOM   2697  HA  ALA B 568     157.166  54.815 107.557  1.00  0.00           H
    ATOM   2698  HB1 ALA B 568     155.450  53.544 106.651  1.00  0.00           H
    ATOM   2699  HB2 ALA B 568     156.588  53.810 105.354  1.00  0.00           H
    ATOM   2700  HB3 ALA B 568     156.483  52.235 106.163  1.00  0.00           H
    ATOM   2701  N   GLU B 569     157.387  53.826 109.724  1.00 65.26           N
    ATOM   2702  CA  GLU B 569     156.922  53.563 111.082  1.00 65.26           C
    ATOM   2703  C   GLU B 569     155.684  52.650 111.076  1.00 65.26           C
    ATOM   2704  O   GLU B 569     154.599  53.086 110.695  1.00 65.26           O
    ATOM   2705  CB  GLU B 569     156.628  54.915 111.800  1.00 99.90           C
    ATOM   2706  CG  GLU B 569     155.549  55.865 111.206  1.00 99.90           C
    ATOM   2707  CD  GLU B 569     155.833  56.247 109.758  1.00 99.90           C
    ATOM   2708  OE1 GLU B 569     156.993  56.653 109.535  1.00 99.90           O
    ATOM   2709  OE2 GLU B 569     154.965  55.976 108.900  1.00 99.90           O1-
    ATOM   2710  H   GLU B 569     157.487  54.818 109.536  1.00  0.00           H
    ATOM   2711  HA  GLU B 569     157.763  53.121 111.611  1.00  0.00           H
    ATOM   2712  HB2 GLU B 569     156.343  54.695 112.830  1.00  0.00           H
    ATOM   2713  HB3 GLU B 569     157.565  55.466 111.891  1.00  0.00           H
    ATOM   2714  HG2 GLU B 569     154.547  55.448 111.292  1.00  0.00           H
    ATOM   2715  HG3 GLU B 569     155.528  56.786 111.789  1.00  0.00           H
    ATOM   2716  N   PRO B 570     155.874  51.367 111.431  1.00 43.58           N
    ATOM   2717  CA  PRO B 570     154.770  50.406 111.384  1.00 43.58           C
    ATOM   2718  C   PRO B 570     153.818  50.498 112.594  1.00 43.58           C
    ATOM   2719  O   PRO B 570     152.750  49.894 112.541  1.00 43.58           O
    ATOM   2720  CB  PRO B 570     155.532  49.084 111.303  1.00 65.85           C
    ATOM   2721  CG  PRO B 570     156.762  49.292 112.160  1.00 65.85           C
    ATOM   2722  CD  PRO B 570     157.134  50.731 111.836  1.00 65.85           C
    ATOM   2723  HA  PRO B 570     154.175  50.529 110.476  1.00  0.00           H
    ATOM   2724  HB2 PRO B 570     154.962  48.194 111.535  1.00  0.00           H
    ATOM   2725  HB3 PRO B 570     155.879  48.959 110.291  1.00  0.00           H
    ATOM   2726  HG2 PRO B 570     156.498  49.200 113.214  1.00  0.00           H
    ATOM   2727  HG3 PRO B 570     157.564  48.585 111.947  1.00  0.00           H
    ATOM   2728  HD2 PRO B 570     157.589  51.210 112.703  1.00  0.00           H
    ATOM   2729  HD3 PRO B 570     157.845  50.759 111.010  1.00  0.00           H
    ATOM   2730  N   ASN B 571     154.200  51.238 113.651  1.00 99.90           N
    ATOM   2731  CA  ASN B 571     153.425  51.377 114.887  1.00 99.90           C
    ATOM   2732  C   ASN B 571     152.670  52.698 114.896  1.00 99.90           C
    ATOM   2733  O   ASN B 571     153.272  53.756 115.079  1.00 99.90           O
    ATOM   2734  CB  ASN B 571     154.367  51.255 116.100  1.00 99.90           C
    ATOM   2735  CG  ASN B 571     154.795  49.811 116.352  1.00 99.90           C
    ATOM   2736  OD1 ASN B 571     154.028  48.886 116.106  1.00 99.90           O
    ATOM   2737  ND2 ASN B 571     156.004  49.611 116.873  1.00 99.90           N
    ATOM   2738  H   ASN B 571     155.083  51.727 113.621  1.00  0.00           H
    ATOM   2739  HA  ASN B 571     152.695  50.570 114.929  1.00  0.00           H
    ATOM   2740  HB2 ASN B 571     155.235  51.909 115.990  1.00  0.00           H
    ATOM   2741  HB3 ASN B 571     153.849  51.577 117.004  1.00  0.00           H
    ATOM   2742 HD21 ASN B 571     156.613  50.393 117.067  1.00  0.00           H
    ATOM   2743 HD22 ASN B 571     156.336  48.672 117.041  1.00  0.00           H
    ATOM   2744  N   CYS B 572     151.351  52.597 114.674  1.00 76.40           N
    ATOM   2745  CA  CYS B 572     150.474  53.747 114.530  1.00 76.40           C
    ATOM   2746  C   CYS B 572     149.089  53.379 115.079  1.00 76.40           C
    ATOM   2747  O   CYS B 572     148.427  52.521 114.493  1.00 76.40           O
    ATOM   2748  CB  CYS B 572     150.326  54.184 113.054  1.00 68.89           C
    ATOM   2749  SG  CYS B 572     151.816  54.069 112.034  1.00 68.89           S
    ATOM   2750  H   CYS B 572     150.931  51.694 114.510  1.00  0.00           H
    ATOM   2751  HA  CYS B 572     150.864  54.585 115.111  1.00  0.00           H
    ATOM   2752  HB2 CYS B 572     149.606  53.555 112.533  1.00  0.00           H
    ATOM   2753  HB3 CYS B 572     149.965  55.210 113.007  1.00  0.00           H
    ATOM   2754  HG  CYS B 572     152.497  55.021 112.674  1.00  0.00           H
    ATOM   2755  N   ALA B 573     148.653  54.053 116.156  1.00 97.72           N
    ATOM   2756  CA  ALA B 573     147.272  53.980 116.637  1.00 97.72           C
    ATOM   2757  C   ALA B 573     146.388  54.870 115.757  1.00 97.72           C
    ATOM   2758  O   ALA B 573     146.788  55.983 115.418  1.00 97.72           O
    ATOM   2759  CB  ALA B 573     147.204  54.419 118.104  1.00 99.90           C
    ATOM   2760  H   ALA B 573     149.241  54.746 116.594  1.00  0.00           H
    ATOM   2761  HA  ALA B 573     146.921  52.947 116.577  1.00  0.00           H
    ATOM   2762  HB1 ALA B 573     146.182  54.363 118.482  1.00  0.00           H
    ATOM   2763  HB2 ALA B 573     147.817  53.774 118.734  1.00  0.00           H
    ATOM   2764  HB3 ALA B 573     147.551  55.445 118.237  1.00  0.00           H
    ATOM   2765  N   ASP B 574     145.226  54.341 115.372  1.00 61.97           N
    ATOM   2766  CA  ASP B 574     144.363  54.912 114.348  1.00 61.97           C
    ATOM   2767  C   ASP B 574     143.361  55.867 115.030  1.00 61.97           C
    ATOM   2768  O   ASP B 574     142.335  55.364 115.466  1.00 61.97           O
    ATOM   2769  CB  ASP B 574     143.724  53.711 113.628  1.00 92.76           C
    ATOM   2770  CG  ASP B 574     143.243  53.914 112.209  1.00 92.76           C
    ATOM   2771  OD1 ASP B 574     143.941  53.420 111.300  1.00 92.76           O
    ATOM   2772  OD2 ASP B 574     142.175  54.522 112.032  1.00 92.76           O1-
    ATOM   2773  H   ASP B 574     144.982  53.408 115.677  1.00  0.00           H
    ATOM   2774  HA  ASP B 574     144.975  55.425 113.613  1.00  0.00           H
    ATOM   2775  HB2 ASP B 574     144.553  53.044 113.436  1.00  0.00           H
    ATOM   2776  HB3 ASP B 574     143.042  53.127 114.240  1.00  0.00           H
    ATOM   2777  N   PRO B 575     143.662  57.191 115.147  1.00 85.92           N
    ATOM   2778  CA  PRO B 575     142.974  58.202 115.977  1.00 85.92           C
    ATOM   2779  C   PRO B 575     141.773  57.758 116.815  1.00 85.92           C
    ATOM   2780  O   PRO B 575     141.997  57.342 117.950  1.00 85.92           O
    ATOM   2781  CB  PRO B 575     142.712  59.328 114.977  1.00 74.50           C
    ATOM   2782  CG  PRO B 575     143.939  59.290 114.078  1.00 74.50           C
    ATOM   2783  CD  PRO B 575     144.633  57.954 114.368  1.00 74.50           C
    ATOM   2784  HA  PRO B 575     143.719  58.560 116.691  1.00  0.00           H
    ATOM   2785  HB2 PRO B 575     141.819  59.137 114.380  1.00  0.00           H
    ATOM   2786  HB3 PRO B 575     142.586  60.297 115.460  1.00  0.00           H
    ATOM   2787  HG2 PRO B 575     143.593  59.335 113.054  1.00  0.00           H
    ATOM   2788  HG3 PRO B 575     144.612  60.131 114.249  1.00  0.00           H
    ATOM   2789  HD2 PRO B 575     145.006  57.488 113.464  1.00  0.00           H
    ATOM   2790  HD3 PRO B 575     145.499  58.139 115.004  1.00  0.00           H
    ATOM   2791  N   ALA B 576     140.563  57.754 116.231  1.00 75.36           N
    ATOM   2792  CA  ALA B 576     139.362  57.195 116.851  1.00 75.36           C
    ATOM   2793  C   ALA B 576     139.521  55.717 117.256  1.00 75.36           C
    ATOM   2794  O   ALA B 576     139.732  55.449 118.437  1.00 75.36           O
    ATOM   2795  CB  ALA B 576     138.156  57.417 115.930  1.00 99.90           C
    ATOM   2796  H   ALA B 576     140.458  58.116 115.295  1.00  0.00           H
    ATOM   2797  HA  ALA B 576     139.179  57.760 117.767  1.00  0.00           H
    ATOM   2798  HB1 ALA B 576     137.253  56.960 116.336  1.00  0.00           H
    ATOM   2799  HB2 ALA B 576     137.957  58.484 115.824  1.00  0.00           H
    ATOM   2800  HB3 ALA B 576     138.319  57.009 114.932  1.00  0.00           H
    ATOM   2801  N   THR B 577     139.440  54.803 116.276  1.00 85.18           N
    ATOM   2802  CA  THR B 577     139.507  53.345 116.443  1.00 85.18           C
    ATOM   2803  C   THR B 577     140.633  52.795 117.332  1.00 85.18           C
    ATOM   2804  O   THR B 577     140.408  51.783 117.991  1.00 85.18           O
    ATOM   2805  CB  THR B 577     139.708  52.622 115.092  1.00 93.65           C
    ATOM   2806  OG1 THR B 577     140.601  53.346 114.278  1.00 93.65           O
    ATOM   2807  CG2 THR B 577     138.436  52.501 114.284  1.00 93.65           C
    ATOM   2808  H   THR B 577     139.321  55.115 115.323  1.00  0.00           H
    ATOM   2809  HA  THR B 577     138.561  53.038 116.889  1.00  0.00           H
    ATOM   2810  HB  THR B 577     140.084  51.611 115.238  1.00  0.00           H
    ATOM   2811 HG21 THR B 577     138.564  51.797 113.470  1.00  0.00           H
    ATOM   2812 HG22 THR B 577     137.616  52.146 114.891  1.00  0.00           H
    ATOM   2813 HG23 THR B 577     138.146  53.461 113.856  1.00  0.00           H
    ATOM   2814  HG1 THR B 577     140.781  52.844 113.478  1.00  0.00           H
    ATOM   2815  N   LEU B 578     141.822  53.416 117.313  1.00 55.83           N
    ATOM   2816  CA  LEU B 578     143.024  52.990 118.034  1.00 55.83           C
    ATOM   2817  C   LEU B 578     143.668  51.717 117.440  1.00 55.83           C
    ATOM   2818  O   LEU B 578     144.777  51.371 117.845  1.00 55.83           O
    ATOM   2819  CB  LEU B 578     142.677  52.784 119.528  1.00 74.72           C
    ATOM   2820  CG  LEU B 578     143.790  53.092 120.533  1.00 74.72           C
    ATOM   2821  CD1 LEU B 578     143.897  54.613 120.764  1.00 74.72           C
    ATOM   2822  CD2 LEU B 578     143.555  52.319 121.845  1.00 74.72           C
    ATOM   2823  H   LEU B 578     141.931  54.231 116.719  1.00  0.00           H
    ATOM   2824  HA  LEU B 578     143.739  53.806 117.934  1.00  0.00           H
    ATOM   2825  HB2 LEU B 578     141.804  53.373 119.816  1.00  0.00           H
    ATOM   2826  HB3 LEU B 578     142.380  51.748 119.655  1.00  0.00           H
    ATOM   2827  HG  LEU B 578     144.738  52.747 120.121  1.00  0.00           H
    ATOM   2828 HD11 LEU B 578     144.936  54.938 120.700  1.00  0.00           H
    ATOM   2829 HD12 LEU B 578     143.339  55.185 120.022  1.00  0.00           H
    ATOM   2830 HD13 LEU B 578     143.521  54.913 121.743  1.00  0.00           H
    ATOM   2831 HD21 LEU B 578     143.812  52.908 122.725  1.00  0.00           H
    ATOM   2832 HD22 LEU B 578     142.517  52.011 121.962  1.00  0.00           H
    ATOM   2833 HD23 LEU B 578     144.168  51.418 121.875  1.00  0.00           H
    ATOM   2834  N   THR B 579     142.985  51.050 116.489  1.00 54.21           N
    ATOM   2835  CA  THR B 579     143.454  49.879 115.749  1.00 54.21           C
    ATOM   2836  C   THR B 579     144.780  50.177 115.034  1.00 54.21           C
    ATOM   2837  O   THR B 579     145.116  51.324 114.774  1.00 54.21           O
    ATOM   2838  CB  THR B 579     142.418  49.442 114.678  1.00 72.56           C
    ATOM   2839  OG1 THR B 579     141.986  50.561 113.932  1.00 72.56           O
    ATOM   2840  CG2 THR B 579     141.178  48.753 115.269  1.00 72.56           C
    ATOM   2841  H   THR B 579     142.087  51.407 116.198  1.00  0.00           H
    ATOM   2842  HA  THR B 579     143.621  49.069 116.458  1.00  0.00           H
    ATOM   2843  HB  THR B 579     142.876  48.745 113.973  1.00  0.00           H
    ATOM   2844 HG21 THR B 579     140.437  48.536 114.500  1.00  0.00           H
    ATOM   2845 HG22 THR B 579     141.448  47.809 115.740  1.00  0.00           H
    ATOM   2846 HG23 THR B 579     140.693  49.378 116.019  1.00  0.00           H
    ATOM   2847  HG1 THR B 579     141.537  50.248 113.141  1.00  0.00           H
    ATOM   2848  N   ARG B 580     145.556  49.129 114.787  1.00 99.90           N
    ATOM   2849  CA  ARG B 580     146.856  49.226 114.145  1.00 99.90           C
    ATOM   2850  C   ARG B 580     146.828  48.426 112.841  1.00 99.90           C
    ATOM   2851  O   ARG B 580     145.921  47.620 112.656  1.00 99.90           O
    ATOM   2852  CB  ARG B 580     147.918  48.713 115.131  1.00 99.90           C
    ATOM   2853  CG  ARG B 580     148.145  49.639 116.341  1.00 99.90           C
    ATOM   2854  CD  ARG B 580     149.628  49.917 116.629  1.00 99.90           C
    ATOM   2855  NE  ARG B 580     150.409  48.695 116.849  1.00 99.90           N
    ATOM   2856  CZ  ARG B 580     151.121  48.003 115.946  1.00 99.90           C
    ATOM   2857  NH1 ARG B 580     151.240  48.397 114.674  1.00 99.90           N
    ATOM   2858  NH2 ARG B 580     151.741  46.884 116.322  1.00 99.90           N1+
    ATOM   2859  H   ARG B 580     145.264  48.200 115.062  1.00  0.00           H
    ATOM   2860  HA  ARG B 580     147.089  50.256 113.873  1.00  0.00           H
    ATOM   2861  HB2 ARG B 580     147.724  47.696 115.465  1.00  0.00           H
    ATOM   2862  HB3 ARG B 580     148.838  48.619 114.581  1.00  0.00           H
    ATOM   2863  HG2 ARG B 580     147.751  50.610 116.047  1.00  0.00           H
    ATOM   2864  HG3 ARG B 580     147.556  49.372 117.218  1.00  0.00           H
    ATOM   2865  HD2 ARG B 580     150.087  50.629 115.947  1.00  0.00           H
    ATOM   2866  HD3 ARG B 580     149.655  50.399 117.607  1.00  0.00           H
    ATOM   2867  HE  ARG B 580     150.346  48.316 117.783  1.00  0.00           H
    ATOM   2868 HH11 ARG B 580     150.771  49.231 114.356  1.00  0.00           H
    ATOM   2869 HH12 ARG B 580     151.841  47.902 114.031  1.00  0.00           H
    ATOM   2870 HH21 ARG B 580     151.628  46.512 117.255  1.00  0.00           H
    ATOM   2871 HH22 ARG B 580     152.291  46.359 115.660  1.00  0.00           H
    ATOM   2872  N   PRO B 581     147.818  48.627 111.944  1.00 39.24           N
    ATOM   2873  CA  PRO B 581     147.891  47.851 110.695  1.00 39.24           C
    ATOM   2874  C   PRO B 581     147.885  46.337 110.898  1.00 39.24           C
    ATOM   2875  O   PRO B 581     147.050  45.678 110.291  1.00 39.24           O
    ATOM   2876  CB  PRO B 581     149.184  48.328 110.035  1.00 59.49           C
    ATOM   2877  CG  PRO B 581     149.998  48.945 111.157  1.00 59.49           C
    ATOM   2878  CD  PRO B 581     148.916  49.593 112.002  1.00 59.49           C
    ATOM   2879  HA  PRO B 581     147.042  48.113 110.060  1.00  0.00           H
    ATOM   2880  HB2 PRO B 581     149.739  47.565 109.500  1.00  0.00           H
    ATOM   2881  HB3 PRO B 581     148.909  49.080 109.301  1.00  0.00           H
    ATOM   2882  HG2 PRO B 581     150.512  48.168 111.725  1.00  0.00           H
    ATOM   2883  HG3 PRO B 581     150.743  49.654 110.794  1.00  0.00           H
    ATOM   2884  HD2 PRO B 581     149.288  49.861 112.986  1.00  0.00           H
    ATOM   2885  HD3 PRO B 581     148.577  50.518 111.532  1.00  0.00           H
    ATOM   2886  N   VAL B 582     148.737  45.826 111.802  1.00 30.29           N
    ATOM   2887  CA  VAL B 582     148.779  44.408 112.151  1.00 30.29           C
    ATOM   2888  C   VAL B 582     147.416  43.829 112.574  1.00 30.29           C
    ATOM   2889  O   VAL B 582     147.155  42.678 112.234  1.00 30.29           O
    ATOM   2890  CB  VAL B 582     149.800  44.106 113.272  1.00 64.96           C
    ATOM   2891  CG1 VAL B 582     151.222  44.403 112.794  1.00 64.96           C
    ATOM   2892  CG2 VAL B 582     149.516  44.818 114.604  1.00 64.96           C
    ATOM   2893  H   VAL B 582     149.382  46.431 112.288  1.00  0.00           H
    ATOM   2894  HA  VAL B 582     149.088  43.890 111.244  1.00  0.00           H
    ATOM   2895  HB  VAL B 582     149.760  43.038 113.489  1.00  0.00           H
    ATOM   2896 HG11 VAL B 582     151.951  44.190 113.576  1.00  0.00           H
    ATOM   2897 HG12 VAL B 582     151.472  43.769 111.945  1.00  0.00           H
    ATOM   2898 HG13 VAL B 582     151.330  45.438 112.476  1.00  0.00           H
    ATOM   2899 HG21 VAL B 582     150.220  44.496 115.372  1.00  0.00           H
    ATOM   2900 HG22 VAL B 582     149.618  45.898 114.499  1.00  0.00           H
    ATOM   2901 HG23 VAL B 582     148.520  44.632 114.990  1.00  0.00           H
    ATOM   2902  N   HIE B 583     146.553  44.653 113.210  1.00 55.13           N
    ATOM   2903  CA  HIE B 583     145.150  44.329 113.472  1.00 55.13           C
    ATOM   2904  C   HIE B 583     144.366  44.117 112.188  1.00 55.13           C
    ATOM   2905  O   HIE B 583     143.820  43.038 112.014  1.00 55.13           O
    ATOM   2906  CB  HIE B 583     144.407  45.395 114.295  1.00 80.53           C
    ATOM   2907  CG  HIE B 583     144.840  45.499 115.712  1.00 80.53           C
    ATOM   2908  ND1 HIE B 583     145.544  46.591 116.209  1.00 80.53           N
    ATOM   2909  CD2 HIE B 583     144.606  44.612 116.730  1.00 80.53           C
    ATOM   2910  CE1 HIE B 583     145.723  46.328 117.489  1.00 80.53           C
    ATOM   2911  NE2 HIE B 583     145.212  45.157 117.838  1.00 80.53           N
    ATOM   2912  H   HIE B 583     146.831  45.603 113.410  1.00  0.00           H
    ATOM   2913  HA  HIE B 583     145.118  43.393 114.029  1.00  0.00           H
    ATOM   2914  HB2 HIE B 583     144.414  46.379 113.849  1.00  0.00           H
    ATOM   2915  HB3 HIE B 583     143.345  45.162 114.325  1.00  0.00           H
    ATOM   2916  HD2 HIE B 583     144.072  43.677 116.761  1.00  0.00           H
    ATOM   2917  HE2 HIE B 583     145.212  44.760 118.770  1.00  0.00           H
    ATOM   2918  HE1 HIE B 583     146.214  46.997 118.173  1.00  0.00           H
    ATOM   2919  N   ASP B 584     144.325  45.150 111.333  1.00 44.59           N
    ATOM   2920  CA  ASP B 584     143.672  45.134 110.024  1.00 44.59           C
    ATOM   2921  C   ASP B 584     144.093  43.994 109.114  1.00 44.59           C
    ATOM   2922  O   ASP B 584     143.226  43.291 108.601  1.00 44.59           O
    ATOM   2923  CB  ASP B 584     143.767  46.466 109.276  1.00 46.12           C
    ATOM   2924  CG  ASP B 584     142.922  47.538 109.928  1.00 46.12           C
    ATOM   2925  OD1 ASP B 584     141.719  47.253 110.125  1.00 46.12           O
    ATOM   2926  OD2 ASP B 584     143.446  48.653 110.130  1.00 46.12           O1-
    ATOM   2927  H   ASP B 584     144.813  46.003 111.572  1.00  0.00           H
    ATOM   2928  HA  ASP B 584     142.623  44.969 110.218  1.00  0.00           H
    ATOM   2929  HB2 ASP B 584     144.805  46.800 109.244  1.00  0.00           H
    ATOM   2930  HB3 ASP B 584     143.424  46.388 108.247  1.00  0.00           H
    ATOM   2931  N   ALA B 585     145.410  43.797 108.984  1.00 51.20           N
    ATOM   2932  CA  ALA B 585     145.992  42.659 108.291  1.00 51.20           C
    ATOM   2933  C   ALA B 585     145.457  41.339 108.855  1.00 51.20           C
    ATOM   2934  O   ALA B 585     145.037  40.485 108.079  1.00 51.20           O
    ATOM   2935  CB  ALA B 585     147.518  42.743 108.380  1.00 39.97           C
    ATOM   2936  H   ALA B 585     146.056  44.429 109.442  1.00  0.00           H
    ATOM   2937  HA  ALA B 585     145.701  42.719 107.242  1.00  0.00           H
    ATOM   2938  HB1 ALA B 585     147.979  41.938 107.811  1.00  0.00           H
    ATOM   2939  HB2 ALA B 585     147.884  43.683 107.966  1.00  0.00           H
    ATOM   2940  HB3 ALA B 585     147.867  42.676 109.410  1.00  0.00           H
    ATOM   2941  N   ALA B 586     145.375  41.256 110.193  1.00 43.77           N
    ATOM   2942  CA  ALA B 586     144.794  40.127 110.894  1.00 43.77           C
    ATOM   2943  C   ALA B 586     143.281  39.934 110.707  1.00 43.77           C
    ATOM   2944  O   ALA B 586     142.909  38.789 110.485  1.00 43.77           O
    ATOM   2945  CB  ALA B 586     145.216  40.132 112.367  1.00 47.32           C
    ATOM   2946  H   ALA B 586     145.711  42.017 110.768  1.00  0.00           H
    ATOM   2947  HA  ALA B 586     145.247  39.263 110.421  1.00  0.00           H
    ATOM   2948  HB1 ALA B 586     144.952  39.195 112.846  1.00  0.00           H
    ATOM   2949  HB2 ALA B 586     146.298  40.226 112.460  1.00  0.00           H
    ATOM   2950  HB3 ALA B 586     144.754  40.933 112.939  1.00  0.00           H
    ATOM   2951  N   ARG B 587     142.444  40.991 110.754  1.00 61.34           N
    ATOM   2952  CA  ARG B 587     140.985  40.866 110.570  1.00 61.34           C
    ATOM   2953  C   ARG B 587     140.551  40.613 109.121  1.00 61.34           C
    ATOM   2954  O   ARG B 587     139.481  40.038 108.929  1.00 61.34           O
    ATOM   2955  CB  ARG B 587     140.178  42.059 111.136  1.00 79.21           C
    ATOM   2956  CG  ARG B 587     140.484  43.457 110.579  1.00 79.21           C
    ATOM   2957  CD  ARG B 587     139.284  44.414 110.618  1.00 79.21           C
    ATOM   2958  NE  ARG B 587     138.820  44.634 111.992  1.00 79.21           N
    ATOM   2959  CZ  ARG B 587     139.252  45.579 112.838  1.00 79.21           C
    ATOM   2960  NH1 ARG B 587     140.120  46.527 112.464  1.00 79.21           N
    ATOM   2961  NH2 ARG B 587     138.801  45.557 114.094  1.00 79.21           N1+
    ATOM   2962  H   ARG B 587     142.810  41.913 110.956  1.00  0.00           H
    ATOM   2963  HA  ARG B 587     140.670  39.991 111.142  1.00  0.00           H
    ATOM   2964  HB2 ARG B 587     139.123  41.837 110.966  1.00  0.00           H
    ATOM   2965  HB3 ARG B 587     140.269  42.113 112.216  1.00  0.00           H
    ATOM   2966  HG2 ARG B 587     141.268  43.833 111.231  1.00  0.00           H
    ATOM   2967  HG3 ARG B 587     140.869  43.439 109.561  1.00  0.00           H
    ATOM   2968  HD2 ARG B 587     139.447  45.332 110.052  1.00  0.00           H
    ATOM   2969  HD3 ARG B 587     138.449  43.914 110.127  1.00  0.00           H
    ATOM   2970  HE  ARG B 587     138.183  43.936 112.345  1.00  0.00           H
    ATOM   2971 HH11 ARG B 587     140.510  46.536 111.525  1.00  0.00           H
    ATOM   2972 HH12 ARG B 587     140.441  47.211 113.132  1.00  0.00           H
    ATOM   2973 HH21 ARG B 587     138.160  44.837 114.392  1.00  0.00           H
    ATOM   2974 HH22 ARG B 587     139.071  46.271 114.757  1.00  0.00           H
    ATOM   2975  N   GLU B 588     141.359  41.036 108.142  1.00 61.58           N
    ATOM   2976  CA  GLU B 588     141.092  40.800 106.728  1.00 61.58           C
    ATOM   2977  C   GLU B 588     141.836  39.553 106.222  1.00 61.58           C
    ATOM   2978  O   GLU B 588     141.697  39.230 105.048  1.00 61.58           O
    ATOM   2979  CB  GLU B 588     141.457  42.073 105.940  1.00 87.07           C
    ATOM   2980  CG  GLU B 588     140.354  42.525 104.968  1.00 87.07           C
    ATOM   2981  CD  GLU B 588     139.189  43.170 105.714  1.00 87.07           C
    ATOM   2982  OE1 GLU B 588     139.325  44.369 106.042  1.00 87.07           O
    ATOM   2983  OE2 GLU B 588     138.198  42.451 105.959  1.00 87.07           O1-
    ATOM   2984  H   GLU B 588     142.206  41.543 108.365  1.00  0.00           H
    ATOM   2985  HA  GLU B 588     140.031  40.590 106.585  1.00  0.00           H
    ATOM   2986  HB2 GLU B 588     141.657  42.901 106.619  1.00  0.00           H
    ATOM   2987  HB3 GLU B 588     142.401  41.939 105.426  1.00  0.00           H
    ATOM   2988  HG2 GLU B 588     140.740  43.277 104.291  1.00  0.00           H
    ATOM   2989  HG3 GLU B 588     140.005  41.695 104.351  1.00  0.00           H
    ATOM   2990  N   GLY B 589     142.580  38.881 107.118  1.00 54.25           N
    ATOM   2991  CA  GLY B 589     143.204  37.575 106.944  1.00 54.25           C
    ATOM   2992  C   GLY B 589     144.428  37.567 106.023  1.00 54.25           C
    ATOM   2993  O   GLY B 589     144.864  36.491 105.616  1.00 54.25           O
    ATOM   2994  H   GLY B 589     142.641  39.266 108.050  1.00  0.00           H
    ATOM   2995  HA2 GLY B 589     143.523  37.235 107.928  1.00  0.00           H
    ATOM   2996  HA3 GLY B 589     142.458  36.861 106.618  1.00  0.00           H
    ATOM   2997  N   PHE B 590     144.993  38.738 105.700  1.00 34.43           N
    ATOM   2998  CA  PHE B 590     146.163  38.848 104.840  1.00 34.43           C
    ATOM   2999  C   PHE B 590     147.415  38.709 105.676  1.00 34.43           C
    ATOM   3000  O   PHE B 590     148.015  39.702 106.097  1.00 34.43           O
    ATOM   3001  CB  PHE B 590     146.127  40.163 104.082  1.00 51.97           C
    ATOM   3002  CG  PHE B 590     144.824  40.341 103.351  1.00 51.97           C
    ATOM   3003  CD1 PHE B 590     144.293  39.328 102.527  1.00 51.97           C
    ATOM   3004  CD2 PHE B 590     144.115  41.532 103.518  1.00 51.97           C
    ATOM   3005  CE1 PHE B 590     143.041  39.498 101.956  1.00 51.97           C
    ATOM   3006  CE2 PHE B 590     142.891  41.695 102.908  1.00 51.97           C
    ATOM   3007  CZ  PHE B 590     142.337  40.673 102.158  1.00 51.97           C
    ATOM   3008  H   PHE B 590     144.621  39.597 106.080  1.00  0.00           H
    ATOM   3009  HA  PHE B 590     146.167  38.042 104.104  1.00  0.00           H
    ATOM   3010  HB2 PHE B 590     146.252  41.001 104.768  1.00  0.00           H
    ATOM   3011  HB3 PHE B 590     146.957  40.194 103.379  1.00  0.00           H
    ATOM   3012  HD1 PHE B 590     144.828  38.405 102.363  1.00  0.00           H
    ATOM   3013  HD2 PHE B 590     144.498  42.294 104.165  1.00  0.00           H
    ATOM   3014  HE1 PHE B 590     142.616  38.715 101.346  1.00  0.00           H
    ATOM   3015  HE2 PHE B 590     142.344  42.606 103.059  1.00  0.00           H
    ATOM   3016  HZ  PHE B 590     141.356  40.796 101.725  1.00  0.00           H
    ATOM   3017  N   LEU B 591     147.784  37.438 105.870  1.00 50.46           N
    ATOM   3018  CA  LEU B 591     149.005  37.045 106.539  1.00 50.46           C
    ATOM   3019  C   LEU B 591     150.238  37.685 105.896  1.00 50.46           C
    ATOM   3020  O   LEU B 591     151.131  38.115 106.612  1.00 50.46           O
    ATOM   3021  CB  LEU B 591     149.095  35.511 106.553  1.00 38.97           C
    ATOM   3022  CG  LEU B 591     150.265  34.936 107.370  1.00 38.97           C
    ATOM   3023  CD1 LEU B 591     150.467  35.638 108.717  1.00 38.97           C
    ATOM   3024  CD2 LEU B 591     150.093  33.422 107.583  1.00 38.97           C
    ATOM   3025  H   LEU B 591     147.205  36.691 105.508  1.00  0.00           H
    ATOM   3026  HA  LEU B 591     148.898  37.425 107.554  1.00  0.00           H
    ATOM   3027  HB2 LEU B 591     148.164  35.118 106.965  1.00  0.00           H
    ATOM   3028  HB3 LEU B 591     149.144  35.128 105.532  1.00  0.00           H
    ATOM   3029  HG  LEU B 591     151.175  35.088 106.788  1.00  0.00           H
    ATOM   3030 HD11 LEU B 591     151.379  35.252 109.151  1.00  0.00           H
    ATOM   3031 HD12 LEU B 591     150.593  36.713 108.661  1.00  0.00           H
    ATOM   3032 HD13 LEU B 591     149.644  35.446 109.401  1.00  0.00           H
    ATOM   3033 HD21 LEU B 591     151.009  32.890 107.327  1.00  0.00           H
    ATOM   3034 HD22 LEU B 591     149.848  33.175 108.616  1.00  0.00           H
    ATOM   3035 HD23 LEU B 591     149.299  33.010 106.960  1.00  0.00           H
    ATOM   3036  N   ASP B 592     150.203  37.810 104.565  1.00 37.51           N
    ATOM   3037  CA  ASP B 592     151.154  38.505 103.709  1.00 37.51           C
    ATOM   3038  C   ASP B 592     151.570  39.872 104.267  1.00 37.51           C
    ATOM   3039  O   ASP B 592     152.703  40.001 104.733  1.00 37.51           O
    ATOM   3040  CB  ASP B 592     150.651  38.631 102.254  1.00 99.90           C
    ATOM   3041  CG  ASP B 592     149.954  37.390 101.696  1.00 99.90           C
    ATOM   3042  OD1 ASP B 592     148.835  37.087 102.167  1.00 99.90           O
    ATOM   3043  OD2 ASP B 592     150.516  36.812 100.741  1.00 99.90           O1-
    ATOM   3044  H   ASP B 592     149.409  37.423 104.068  1.00  0.00           H
    ATOM   3045  HA  ASP B 592     152.049  37.881 103.689  1.00  0.00           H
    ATOM   3046  HB2 ASP B 592     149.966  39.468 102.116  1.00  0.00           H
    ATOM   3047  HB3 ASP B 592     151.511  38.831 101.615  1.00  0.00           H
    ATOM   3048  N   THR B 593     150.628  40.832 104.292  1.00 28.43           N
    ATOM   3049  CA  THR B 593     150.846  42.148 104.890  1.00 28.43           C
    ATOM   3050  C   THR B 593     151.172  42.083 106.382  1.00 28.43           C
    ATOM   3051  O   THR B 593     152.043  42.842 106.790  1.00 28.43           O
    ATOM   3052  CB  THR B 593     149.649  43.119 104.728  1.00 71.41           C
    ATOM   3053  OG1 THR B 593     148.421  42.424 104.784  1.00 71.41           O
    ATOM   3054  CG2 THR B 593     149.712  43.824 103.380  1.00 71.41           C
    ATOM   3055  H   THR B 593     149.703  40.649 103.932  1.00  0.00           H
    ATOM   3056  HA  THR B 593     151.718  42.577 104.392  1.00  0.00           H
    ATOM   3057  HB  THR B 593     149.644  43.883 105.509  1.00  0.00           H
    ATOM   3058 HG21 THR B 593     149.030  44.663 103.298  1.00  0.00           H
    ATOM   3059 HG22 THR B 593     150.702  44.221 103.216  1.00  0.00           H
    ATOM   3060 HG23 THR B 593     149.510  43.131 102.565  1.00  0.00           H
    ATOM   3061  HG1 THR B 593     148.350  41.983 105.636  1.00  0.00           H
    ATOM   3062  N   LEU B 594     150.552  41.165 107.145  1.00 39.18           N
    ATOM   3063  CA  LEU B 594     150.855  40.997 108.563  1.00 39.18           C
    ATOM   3064  C   LEU B 594     152.321  40.649 108.822  1.00 39.18           C
    ATOM   3065  O   LEU B 594     152.947  41.325 109.624  1.00 39.18           O
    ATOM   3066  CB  LEU B 594     149.949  39.949 109.218  1.00 37.45           C
    ATOM   3067  CG  LEU B 594     150.220  39.763 110.727  1.00 37.45           C
    ATOM   3068  CD1 LEU B 594     149.983  41.054 111.522  1.00 37.45           C
    ATOM   3069  CD2 LEU B 594     149.384  38.609 111.253  1.00 37.45           C
    ATOM   3070  H   LEU B 594     149.873  40.532 106.742  1.00  0.00           H
    ATOM   3071  HA  LEU B 594     150.656  41.961 109.032  1.00  0.00           H
    ATOM   3072  HB2 LEU B 594     148.902  40.214 109.070  1.00  0.00           H
    ATOM   3073  HB3 LEU B 594     150.084  38.997 108.715  1.00  0.00           H
    ATOM   3074  HG  LEU B 594     151.252  39.453 110.886  1.00  0.00           H
    ATOM   3075 HD11 LEU B 594     149.427  40.875 112.443  1.00  0.00           H
    ATOM   3076 HD12 LEU B 594     150.932  41.509 111.801  1.00  0.00           H
    ATOM   3077 HD13 LEU B 594     149.416  41.775 110.941  1.00  0.00           H
    ATOM   3078 HD21 LEU B 594     149.925  38.046 112.007  1.00  0.00           H
    ATOM   3079 HD22 LEU B 594     148.445  38.966 111.669  1.00  0.00           H
    ATOM   3080 HD23 LEU B 594     149.134  37.910 110.455  1.00  0.00           H
    ATOM   3081  N   VAL B 595     152.837  39.606 108.164  1.00 37.86           N
    ATOM   3082  CA  VAL B 595     154.219  39.159 108.302  1.00 37.86           C
    ATOM   3083  C   VAL B 595     155.207  40.248 107.876  1.00 37.86           C
    ATOM   3084  O   VAL B 595     156.163  40.497 108.602  1.00 37.86           O
    ATOM   3085  CB  VAL B 595     154.507  37.890 107.463  1.00 37.46           C
    ATOM   3086  CG1 VAL B 595     156.000  37.507 107.384  1.00 37.46           C
    ATOM   3087  CG2 VAL B 595     153.720  36.688 107.994  1.00 37.46           C
    ATOM   3088  H   VAL B 595     152.256  39.081 107.521  1.00  0.00           H
    ATOM   3089  HA  VAL B 595     154.392  38.927 109.354  1.00  0.00           H
    ATOM   3090  HB  VAL B 595     154.152  38.075 106.447  1.00  0.00           H
    ATOM   3091 HG11 VAL B 595     156.127  36.523 106.932  1.00  0.00           H
    ATOM   3092 HG12 VAL B 595     156.574  38.200 106.768  1.00  0.00           H
    ATOM   3093 HG13 VAL B 595     156.456  37.475 108.374  1.00  0.00           H
    ATOM   3094 HG21 VAL B 595     153.579  35.941 107.213  1.00  0.00           H
    ATOM   3095 HG22 VAL B 595     154.245  36.198 108.808  1.00  0.00           H
    ATOM   3096 HG23 VAL B 595     152.743  36.982 108.357  1.00  0.00           H
    ATOM   3097  N   VAL B 596     154.940  40.901 106.740  1.00 36.76           N
    ATOM   3098  CA  VAL B 596     155.734  42.021 106.261  1.00 36.76           C
    ATOM   3099  C   VAL B 596     155.756  43.194 107.267  1.00 36.76           C
    ATOM   3100  O   VAL B 596     156.842  43.685 107.570  1.00 36.76           O
    ATOM   3101  CB  VAL B 596     155.213  42.465 104.879  1.00 43.67           C
    ATOM   3102  CG1 VAL B 596     155.619  43.861 104.460  1.00 43.67           C
    ATOM   3103  CG2 VAL B 596     155.615  41.474 103.777  1.00 43.67           C
    ATOM   3104  H   VAL B 596     154.132  40.647 106.186  1.00  0.00           H
    ATOM   3105  HA  VAL B 596     156.762  41.670 106.144  1.00  0.00           H
    ATOM   3106  HB  VAL B 596     154.123  42.460 104.915  1.00  0.00           H
    ATOM   3107 HG11 VAL B 596     155.253  44.021 103.457  1.00  0.00           H
    ATOM   3108 HG12 VAL B 596     155.078  44.579 105.057  1.00  0.00           H
    ATOM   3109 HG13 VAL B 596     156.695  44.031 104.504  1.00  0.00           H
    ATOM   3110 HG21 VAL B 596     155.160  41.738 102.821  1.00  0.00           H
    ATOM   3111 HG22 VAL B 596     156.696  41.460 103.634  1.00  0.00           H
    ATOM   3112 HG23 VAL B 596     155.300  40.458 104.015  1.00  0.00           H
    ATOM   3113  N   LEU B 597     154.583  43.541 107.832  1.00 47.92           N
    ATOM   3114  CA  LEU B 597     154.411  44.441 108.982  1.00 47.92           C
    ATOM   3115  C   LEU B 597     155.224  43.986 110.190  1.00 47.92           C
    ATOM   3116  O   LEU B 597     155.890  44.814 110.808  1.00 47.92           O
    ATOM   3117  CB  LEU B 597     152.920  44.546 109.382  1.00 45.81           C
    ATOM   3118  CG  LEU B 597     152.076  45.668 108.756  1.00 45.81           C
    ATOM   3119  CD1 LEU B 597     152.603  46.145 107.410  1.00 45.81           C
    ATOM   3120  CD2 LEU B 597     150.604  45.240 108.655  1.00 45.81           C
    ATOM   3121  H   LEU B 597     153.735  43.093 107.506  1.00  0.00           H
    ATOM   3122  HA  LEU B 597     154.785  45.433 108.763  1.00  0.00           H
    ATOM   3123  HB2 LEU B 597     152.433  43.599 109.198  1.00  0.00           H
    ATOM   3124  HB3 LEU B 597     152.838  44.686 110.459  1.00  0.00           H
    ATOM   3125  HG  LEU B 597     152.125  46.532 109.420  1.00  0.00           H
    ATOM   3126 HD11 LEU B 597     151.918  46.835 106.938  1.00  0.00           H
    ATOM   3127 HD12 LEU B 597     153.530  46.685 107.555  1.00  0.00           H
    ATOM   3128 HD13 LEU B 597     152.787  45.318 106.726  1.00  0.00           H
    ATOM   3129 HD21 LEU B 597     150.029  45.883 107.992  1.00  0.00           H
    ATOM   3130 HD22 LEU B 597     150.482  44.224 108.287  1.00  0.00           H
    ATOM   3131 HD23 LEU B 597     150.144  45.271 109.635  1.00  0.00           H
    ATOM   3132  N   HID B 598     155.150  42.684 110.496  1.00 38.94           N
    ATOM   3133  CA  HID B 598     155.796  42.082 111.644  1.00 38.94           C
    ATOM   3134  C   HID B 598     157.322  42.175 111.571  1.00 38.94           C
    ATOM   3135  O   HID B 598     157.896  42.797 112.462  1.00 38.94           O
    ATOM   3136  CB  HID B 598     155.307  40.644 111.895  1.00 75.47           C
    ATOM   3137  CG  HID B 598     155.853  40.046 113.171  1.00 75.47           C
    ATOM   3138  ND1 HID B 598     156.448  40.804 114.168  1.00 75.47           N
    ATOM   3139  CD2 HID B 598     155.912  38.753 113.630  1.00 75.47           C
    ATOM   3140  CE1 HID B 598     156.893  39.969 115.104  1.00 75.47           C
    ATOM   3141  NE2 HID B 598     156.607  38.704 114.837  1.00 75.47           N
    ATOM   3142  H   HID B 598     154.555  42.077 109.945  1.00  0.00           H
    ATOM   3143  HA  HID B 598     155.475  42.680 112.498  1.00  0.00           H
    ATOM   3144  HB2 HID B 598     154.218  40.622 111.953  1.00  0.00           H
    ATOM   3145  HB3 HID B 598     155.588  39.987 111.074  1.00  0.00           H
    ATOM   3146  HD1 HID B 598     156.569  41.808 114.165  1.00  0.00           H
    ATOM   3147  HD2 HID B 598     155.497  37.855 113.220  1.00  0.00           H
    ATOM   3148  HE1 HID B 598     157.421  40.285 115.992  1.00  0.00           H
    ATOM   3149  N   ARG B 599     157.938  41.596 110.525  1.00 69.55           N
    ATOM   3150  CA  ARG B 599     159.384  41.671 110.288  1.00 69.55           C
    ATOM   3151  C   ARG B 599     159.898  43.123 110.281  1.00 69.55           C
    ATOM   3152  O   ARG B 599     160.986  43.368 110.798  1.00 69.55           O
    ATOM   3153  CB  ARG B 599     159.760  40.994 108.955  1.00 99.90           C
    ATOM   3154  CG  ARG B 599     159.461  39.489 108.845  1.00 99.90           C
    ATOM   3155  CD  ARG B 599     159.681  39.020 107.401  1.00 99.90           C
    ATOM   3156  NE  ARG B 599     159.523  37.571 107.228  1.00 99.90           N
    ATOM   3157  CZ  ARG B 599     159.305  36.968 106.045  1.00 99.90           C
    ATOM   3158  NH1 ARG B 599     159.220  37.674 104.907  1.00 99.90           N
    ATOM   3159  NH2 ARG B 599     159.169  35.638 105.997  1.00 99.90           N1+
    ATOM   3160  H   ARG B 599     157.386  41.124 109.819  1.00  0.00           H
    ATOM   3161  HA  ARG B 599     159.880  41.140 111.103  1.00  0.00           H
    ATOM   3162  HB2 ARG B 599     159.245  41.516 108.146  1.00  0.00           H
    ATOM   3163  HB3 ARG B 599     160.826  41.137 108.772  1.00  0.00           H
    ATOM   3164  HG2 ARG B 599     160.013  38.885 109.567  1.00  0.00           H
    ATOM   3165  HG3 ARG B 599     158.407  39.349 109.078  1.00  0.00           H
    ATOM   3166  HD2 ARG B 599     158.855  39.443 106.829  1.00  0.00           H
    ATOM   3167  HD3 ARG B 599     160.594  39.411 106.951  1.00  0.00           H
    ATOM   3168  HE  ARG B 599     159.625  37.004 108.057  1.00  0.00           H
    ATOM   3169 HH11 ARG B 599     159.323  38.679 104.925  1.00  0.00           H
    ATOM   3170 HH12 ARG B 599     159.051  37.214 104.024  1.00  0.00           H
    ATOM   3171 HH21 ARG B 599     159.232  35.087 106.841  1.00  0.00           H
    ATOM   3172 HH22 ARG B 599     159.006  35.172 105.116  1.00  0.00           H
    ATOM   3173  N   ALA B 600     159.077  44.054 109.758  1.00 57.26           N
    ATOM   3174  CA  ALA B 600     159.350  45.489 109.736  1.00 57.26           C
    ATOM   3175  C   ALA B 600     159.403  46.158 111.118  1.00 57.26           C
    ATOM   3176  O   ALA B 600     160.190  47.090 111.276  1.00 57.26           O
    ATOM   3177  CB  ALA B 600     158.333  46.202 108.844  1.00 50.42           C
    ATOM   3178  H   ALA B 600     158.195  43.767 109.354  1.00  0.00           H
    ATOM   3179  HA  ALA B 600     160.332  45.619 109.278  1.00  0.00           H
    ATOM   3180  HB1 ALA B 600     158.556  47.268 108.791  1.00  0.00           H
    ATOM   3181  HB2 ALA B 600     158.375  45.820 107.824  1.00  0.00           H
    ATOM   3182  HB3 ALA B 600     157.315  46.097 109.208  1.00  0.00           H
    ATOM   3183  N   GLY B 601     158.619  45.670 112.091  1.00 69.41           N
    ATOM   3184  CA  GLY B 601     158.773  46.082 113.482  1.00 69.41           C
    ATOM   3185  C   GLY B 601     157.453  46.201 114.238  1.00 69.41           C
    ATOM   3186  O   GLY B 601     157.511  46.234 115.468  1.00 69.41           O
    ATOM   3187  H   GLY B 601     158.002  44.892 111.899  1.00  0.00           H
    ATOM   3188  HA2 GLY B 601     159.398  45.342 113.984  1.00  0.00           H
    ATOM   3189  HA3 GLY B 601     159.285  47.041 113.577  1.00  0.00           H
    ATOM   3190  N   ALA B 602     156.283  46.254 113.569  1.00 77.80           N
    ATOM   3191  CA  ALA B 602     155.001  46.332 114.271  1.00 77.80           C
    ATOM   3192  C   ALA B 602     154.772  45.127 115.183  1.00 77.80           C
    ATOM   3193  O   ALA B 602     154.838  43.995 114.706  1.00 77.80           O
    ATOM   3194  CB  ALA B 602     153.818  46.516 113.319  1.00 99.90           C
    ATOM   3195  H   ALA B 602     156.260  46.194 112.560  1.00  0.00           H
    ATOM   3196  HA  ALA B 602     155.076  47.213 114.893  1.00  0.00           H
    ATOM   3197  HB1 ALA B 602     152.890  46.256 113.819  1.00  0.00           H
    ATOM   3198  HB2 ALA B 602     153.701  47.556 113.042  1.00  0.00           H
    ATOM   3199  HB3 ALA B 602     153.909  45.909 112.419  1.00  0.00           H
    ATOM   3200  N   ARG B 603     154.561  45.393 116.481  1.00 57.39           N
    ATOM   3201  CA  ARG B 603     154.422  44.327 117.464  1.00 57.39           C
    ATOM   3202  C   ARG B 603     153.021  43.702 117.395  1.00 57.39           C
    ATOM   3203  O   ARG B 603     152.015  44.373 117.600  1.00 57.39           O
    ATOM   3204  CB  ARG B 603     154.872  44.779 118.868  1.00 99.90           C
    ATOM   3205  CG  ARG B 603     154.267  46.068 119.441  1.00 99.90           C
    ATOM   3206  CD  ARG B 603     154.788  46.332 120.861  1.00 99.90           C
    ATOM   3207  NE  ARG B 603     153.982  47.323 121.589  1.00 99.90           N
    ATOM   3208  CZ  ARG B 603     154.143  48.657 121.598  1.00 99.90           C
    ATOM   3209  NH1 ARG B 603     155.105  49.254 120.882  1.00 99.90           N
    ATOM   3210  NH2 ARG B 603     153.326  49.404 122.351  1.00 99.90           N1+
    ATOM   3211  H   ARG B 603     154.544  46.345 116.818  1.00  0.00           H
    ATOM   3212  HA  ARG B 603     155.138  43.546 117.194  1.00  0.00           H
    ATOM   3213  HB2 ARG B 603     154.682  43.957 119.561  1.00  0.00           H
    ATOM   3214  HB3 ARG B 603     155.956  44.900 118.854  1.00  0.00           H
    ATOM   3215  HG2 ARG B 603     154.562  46.896 118.797  1.00  0.00           H
    ATOM   3216  HG3 ARG B 603     153.192  46.041 119.433  1.00  0.00           H
    ATOM   3217  HD2 ARG B 603     154.853  45.402 121.427  1.00  0.00           H
    ATOM   3218  HD3 ARG B 603     155.804  46.723 120.817  1.00  0.00           H
    ATOM   3219  HE  ARG B 603     153.214  46.937 122.120  1.00  0.00           H
    ATOM   3220 HH11 ARG B 603     155.731  48.699 120.318  1.00  0.00           H
    ATOM   3221 HH12 ARG B 603     155.224  50.257 120.912  1.00  0.00           H
    ATOM   3222 HH21 ARG B 603     152.607  48.971 122.912  1.00  0.00           H
    ATOM   3223 HH22 ARG B 603     153.429  50.408 122.377  1.00  0.00           H
    ATOM   3224  N   LEU B 604     153.001  42.403 117.083  1.00 60.10           N
    ATOM   3225  CA  LEU B 604     151.821  41.552 116.932  1.00 60.10           C
    ATOM   3226  C   LEU B 604     150.995  41.260 118.212  1.00 60.10           C
    ATOM   3227  O   LEU B 604     150.156  40.362 118.187  1.00 60.10           O
    ATOM   3228  CB  LEU B 604     152.213  40.270 116.161  1.00 86.40           C
    ATOM   3229  CG  LEU B 604     153.593  39.621 116.427  1.00 86.40           C
    ATOM   3230  CD1 LEU B 604     154.131  39.706 117.870  1.00 86.40           C
    ATOM   3231  CD2 LEU B 604     153.567  38.159 115.947  1.00 86.40           C
    ATOM   3232  H   LEU B 604     153.880  41.933 116.918  1.00  0.00           H
    ATOM   3233  HA  LEU B 604     151.128  42.097 116.287  1.00  0.00           H
    ATOM   3234  HB2 LEU B 604     151.447  39.524 116.329  1.00  0.00           H
    ATOM   3235  HB3 LEU B 604     152.154  40.481 115.092  1.00  0.00           H
    ATOM   3236  HG  LEU B 604     154.303  40.160 115.804  1.00  0.00           H
    ATOM   3237 HD11 LEU B 604     154.676  38.806 118.150  1.00  0.00           H
    ATOM   3238 HD12 LEU B 604     154.834  40.533 117.979  1.00  0.00           H
    ATOM   3239 HD13 LEU B 604     153.341  39.836 118.607  1.00  0.00           H
    ATOM   3240 HD21 LEU B 604     154.533  37.673 116.072  1.00  0.00           H
    ATOM   3241 HD22 LEU B 604     152.846  37.569 116.514  1.00  0.00           H
    ATOM   3242 HD23 LEU B 604     153.299  38.091 114.893  1.00  0.00           H
    ATOM   3243  N   ASP B 605     151.211  42.031 119.287  1.00 99.90           N
    ATOM   3244  CA  ASP B 605     150.441  42.021 120.530  1.00 99.90           C
    ATOM   3245  C   ASP B 605     150.236  43.472 120.983  1.00 99.90           C
    ATOM   3246  O   ASP B 605     150.912  43.950 121.894  1.00 99.90           O
    ATOM   3247  CB  ASP B 605     151.052  41.131 121.643  1.00 99.90           C
    ATOM   3248  CG  ASP B 605     150.504  39.702 121.687  1.00 99.90           C
    ATOM   3249  OD1 ASP B 605     150.732  38.941 120.724  1.00 99.90           O
    ATOM   3250  OD2 ASP B 605     149.882  39.388 122.725  1.00 99.90           O1-
    ATOM   3251  H   ASP B 605     151.916  42.750 119.226  1.00  0.00           H
    ATOM   3252  HA  ASP B 605     149.442  41.640 120.306  1.00  0.00           H
    ATOM   3253  HB2 ASP B 605     152.137  41.078 121.556  1.00  0.00           H
    ATOM   3254  HB3 ASP B 605     150.878  41.560 122.631  1.00  0.00           H
    ATOM   3255  N   VAL B 606     149.301  44.148 120.305  1.00 61.43           N
    ATOM   3256  CA  VAL B 606     148.821  45.491 120.631  1.00 61.43           C
    ATOM   3257  C   VAL B 606     147.301  45.432 120.723  1.00 61.43           C
    ATOM   3258  O   VAL B 606     146.706  44.531 120.145  1.00 61.43           O
    ATOM   3259  CB  VAL B 606     149.215  46.521 119.536  1.00 44.05           C
    ATOM   3260  CG1 VAL B 606     148.604  47.928 119.704  1.00 44.05           C
    ATOM   3261  CG2 VAL B 606     150.730  46.656 119.566  1.00 44.05           C
    ATOM   3262  H   VAL B 606     148.798  43.681 119.564  1.00  0.00           H
    ATOM   3263  HA  VAL B 606     149.202  45.814 121.602  1.00  0.00           H
    ATOM   3264  HB  VAL B 606     148.928  46.140 118.554  1.00  0.00           H
    ATOM   3265 HG11 VAL B 606     149.091  48.665 119.075  1.00  0.00           H
    ATOM   3266 HG12 VAL B 606     147.546  47.970 119.462  1.00  0.00           H
    ATOM   3267 HG13 VAL B 606     148.720  48.280 120.729  1.00  0.00           H
    ATOM   3268 HG21 VAL B 606     151.130  47.356 118.844  1.00  0.00           H
    ATOM   3269 HG22 VAL B 606     151.071  46.988 120.547  1.00  0.00           H
    ATOM   3270 HG23 VAL B 606     151.171  45.686 119.390  1.00  0.00           H
    ATOM   3271  N   ARG B 607     146.693  46.382 121.439  1.00 72.41           N
    ATOM   3272  CA  ARG B 607     145.247  46.453 121.564  1.00 72.41           C
    ATOM   3273  C   ARG B 607     144.668  47.709 120.919  1.00 72.41           C
    ATOM   3274  O   ARG B 607     145.331  48.743 120.836  1.00 72.41           O
    ATOM   3275  CB  ARG B 607     144.855  46.265 123.034  1.00 91.17           C
    ATOM   3276  CG  ARG B 607     144.976  44.787 123.439  1.00 91.17           C
    ATOM   3277  CD  ARG B 607     144.280  44.435 124.757  1.00 91.17           C
    ATOM   3278  NE  ARG B 607     144.086  42.982 124.856  1.00 91.17           N
    ATOM   3279  CZ  ARG B 607     145.044  42.072 125.100  1.00 91.17           C
    ATOM   3280  NH1 ARG B 607     146.299  42.442 125.392  1.00 91.17           N
    ATOM   3281  NH2 ARG B 607     144.735  40.771 125.052  1.00 91.17           N1+
    ATOM   3282  H   ARG B 607     147.224  47.119 121.881  1.00  0.00           H
    ATOM   3283  HA  ARG B 607     144.793  45.637 121.022  1.00  0.00           H
    ATOM   3284  HB2 ARG B 607     145.427  46.915 123.698  1.00  0.00           H
    ATOM   3285  HB3 ARG B 607     143.820  46.569 123.137  1.00  0.00           H
    ATOM   3286  HG2 ARG B 607     144.596  44.128 122.661  1.00  0.00           H
    ATOM   3287  HG3 ARG B 607     146.040  44.567 123.525  1.00  0.00           H
    ATOM   3288  HD2 ARG B 607     144.917  44.713 125.597  1.00  0.00           H
    ATOM   3289  HD3 ARG B 607     143.351  44.985 124.890  1.00  0.00           H
    ATOM   3290  HE  ARG B 607     143.157  42.651 124.628  1.00  0.00           H
    ATOM   3291 HH11 ARG B 607     146.539  43.421 125.442  1.00  0.00           H
    ATOM   3292 HH12 ARG B 607     147.015  41.747 125.551  1.00  0.00           H
    ATOM   3293 HH21 ARG B 607     143.809  40.474 124.779  1.00  0.00           H
    ATOM   3294 HH22 ARG B 607     145.435  40.069 125.245  1.00  0.00           H
    ATOM   3295  N   ASP B 608     143.419  47.553 120.461  1.00 87.59           N
    ATOM   3296  CA  ASP B 608     142.581  48.609 119.901  1.00 87.59           C
    ATOM   3297  C   ASP B 608     141.668  49.171 121.010  1.00 87.59           C
    ATOM   3298  O   ASP B 608     141.793  48.772 122.170  1.00 87.59           O
    ATOM   3299  CB  ASP B 608     141.774  48.107 118.673  1.00 99.90           C
    ATOM   3300  CG  ASP B 608     140.833  46.911 118.887  1.00 99.90           C
    ATOM   3301  OD1 ASP B 608     140.605  46.529 120.052  1.00 99.90           O
    ATOM   3302  OD2 ASP B 608     140.291  46.431 117.869  1.00 99.90           O1-
    ATOM   3303  H   ASP B 608     142.965  46.656 120.555  1.00  0.00           H
    ATOM   3304  HA  ASP B 608     143.213  49.431 119.562  1.00  0.00           H
    ATOM   3305  HB2 ASP B 608     141.199  48.932 118.254  1.00  0.00           H
    ATOM   3306  HB3 ASP B 608     142.500  47.794 117.924  1.00  0.00           H
    ATOM   3307  N   ALA B 609     140.742  50.070 120.638  1.00 72.09           N
    ATOM   3308  CA  ALA B 609     139.769  50.675 121.552  1.00 72.09           C
    ATOM   3309  C   ALA B 609     138.841  49.665 122.249  1.00 72.09           C
    ATOM   3310  O   ALA B 609     138.447  49.906 123.387  1.00 72.09           O
    ATOM   3311  CB  ALA B 609     138.950  51.736 120.807  1.00 71.47           C
    ATOM   3312  H   ALA B 609     140.685  50.366 119.673  1.00  0.00           H
    ATOM   3313  HA  ALA B 609     140.331  51.182 122.338  1.00  0.00           H
    ATOM   3314  HB1 ALA B 609     138.210  52.195 121.463  1.00  0.00           H
    ATOM   3315  HB2 ALA B 609     139.588  52.538 120.439  1.00  0.00           H
    ATOM   3316  HB3 ALA B 609     138.415  51.317 119.958  1.00  0.00           H
    ATOM   3317  N   TRP B 610     138.557  48.534 121.584  1.00 48.83           N
    ATOM   3318  CA  TRP B 610     137.776  47.414 122.118  1.00 48.83           C
    ATOM   3319  C   TRP B 610     138.653  46.392 122.849  1.00 48.83           C
    ATOM   3320  O   TRP B 610     138.142  45.351 123.258  1.00 48.83           O
    ATOM   3321  CB  TRP B 610     136.992  46.739 120.982  1.00 76.74           C
    ATOM   3322  CG  TRP B 610     136.111  47.668 120.218  1.00 76.74           C
    ATOM   3323  CD1 TRP B 610     134.784  47.844 120.407  1.00 76.74           C
    ATOM   3324  CD2 TRP B 610     136.510  48.642 119.215  1.00 76.74           C
    ATOM   3325  NE1 TRP B 610     134.335  48.857 119.586  1.00 76.74           N
    ATOM   3326  CE2 TRP B 610     135.362  49.399 118.843  1.00 76.74           C
    ATOM   3327  CE3 TRP B 610     137.733  48.971 118.587  1.00 76.74           C
    ATOM   3328  CZ2 TRP B 610     135.429  50.448 117.917  1.00 76.74           C
    ATOM   3329  CZ3 TRP B 610     137.810  50.010 117.646  1.00 76.74           C
    ATOM   3330  CH2 TRP B 610     136.664  50.762 117.333  1.00 76.74           C
    ATOM   3331  H   TRP B 610     138.955  48.390 120.666  1.00  0.00           H
    ATOM   3332  HA  TRP B 610     137.054  47.795 122.842  1.00  0.00           H
    ATOM   3333  HB2 TRP B 610     137.691  46.293 120.272  1.00  0.00           H
    ATOM   3334  HB3 TRP B 610     136.382  45.920 121.365  1.00  0.00           H
    ATOM   3335  HD1 TRP B 610     134.187  47.283 121.110  1.00  0.00           H
    ATOM   3336  HE1 TRP B 610     133.373  49.165 119.569  1.00  0.00           H
    ATOM   3337  HE3 TRP B 610     138.619  48.410 118.823  1.00  0.00           H
    ATOM   3338  HZ2 TRP B 610     134.543  51.011 117.659  1.00  0.00           H
    ATOM   3339  HZ3 TRP B 610     138.744  50.220 117.153  1.00  0.00           H
    ATOM   3340  HH2 TRP B 610     136.714  51.577 116.637  1.00  0.00           H
    ATOM   3341  N   GLY B 611     139.948  46.701 123.003  1.00 82.21           N
    ATOM   3342  CA  GLY B 611     140.923  45.889 123.701  1.00 82.21           C
    ATOM   3343  C   GLY B 611     141.130  44.537 123.011  1.00 82.21           C
    ATOM   3344  O   GLY B 611     140.855  43.516 123.636  1.00 82.21           O
    ATOM   3345  H   GLY B 611     140.289  47.569 122.611  1.00  0.00           H
    ATOM   3346  HA2 GLY B 611     141.848  46.448 123.762  1.00  0.00           H
    ATOM   3347  HA3 GLY B 611     140.592  45.733 124.729  1.00  0.00           H
    ATOM   3348  N   ARG B 612     141.576  44.511 121.746  1.00 99.90           N
    ATOM   3349  CA  ARG B 612     141.797  43.263 121.014  1.00 99.90           C
    ATOM   3350  C   ARG B 612     143.185  43.201 120.408  1.00 99.90           C
    ATOM   3351  O   ARG B 612     143.572  44.123 119.705  1.00 99.90           O
    ATOM   3352  CB  ARG B 612     140.763  43.088 119.895  1.00 99.90           C
    ATOM   3353  CG  ARG B 612     139.362  42.767 120.413  1.00 99.90           C
    ATOM   3354  CD  ARG B 612     138.591  41.923 119.396  1.00 99.90           C
    ATOM   3355  NE  ARG B 612     137.181  41.803 119.753  1.00 99.90           N
    ATOM   3356  CZ  ARG B 612     136.267  42.760 119.546  1.00 99.90           C
    ATOM   3357  NH1 ARG B 612     136.598  43.924 118.967  1.00 99.90           N
    ATOM   3358  NH2 ARG B 612     135.002  42.547 119.923  1.00 99.90           N1+
    ATOM   3359  H   ARG B 612     141.663  45.370 121.218  1.00  0.00           H
    ATOM   3360  HA  ARG B 612     141.714  42.402 121.680  1.00  0.00           H
    ATOM   3361  HB2 ARG B 612     140.740  43.942 119.218  1.00  0.00           H
    ATOM   3362  HB3 ARG B 612     141.085  42.243 119.283  1.00  0.00           H
    ATOM   3363  HG2 ARG B 612     139.497  42.116 121.278  1.00  0.00           H
    ATOM   3364  HG3 ARG B 612     138.820  43.639 120.781  1.00  0.00           H
    ATOM   3365  HD2 ARG B 612     138.764  42.220 118.363  1.00  0.00           H
    ATOM   3366  HD3 ARG B 612     138.953  40.898 119.481  1.00  0.00           H
    ATOM   3367  HE  ARG B 612     136.894  40.922 120.158  1.00  0.00           H
    ATOM   3368 HH11 ARG B 612     137.556  44.093 118.693  1.00  0.00           H
    ATOM   3369 HH12 ARG B 612     135.906  44.643 118.818  1.00  0.00           H
    ATOM   3370 HH21 ARG B 612     134.741  41.664 120.338  1.00  0.00           H
    ATOM   3371 HH22 ARG B 612     134.292  43.246 119.763  1.00  0.00           H
    ATOM   3372  N   LEU B 613     143.853  42.061 120.627  1.00 76.56           N
    ATOM   3373  CA  LEU B 613     145.040  41.610 119.904  1.00 76.56           C
    ATOM   3374  C   LEU B 613     144.664  41.363 118.429  1.00 76.56           C
    ATOM   3375  O   LEU B 613     143.528  40.955 118.175  1.00 76.56           O
    ATOM   3376  CB  LEU B 613     145.524  40.309 120.590  1.00 71.12           C
    ATOM   3377  CG  LEU B 613     146.730  40.377 121.559  1.00 71.12           C
    ATOM   3378  CD1 LEU B 613     146.965  41.752 122.204  1.00 71.12           C
    ATOM   3379  CD2 LEU B 613     146.642  39.269 122.636  1.00 71.12           C
    ATOM   3380  H   LEU B 613     143.420  41.353 121.208  1.00  0.00           H
    ATOM   3381  HA  LEU B 613     145.795  42.374 119.965  1.00  0.00           H
    ATOM   3382  HB2 LEU B 613     144.692  39.876 121.122  1.00  0.00           H
    ATOM   3383  HB3 LEU B 613     145.730  39.555 119.841  1.00  0.00           H
    ATOM   3384  HG  LEU B 613     147.622  40.177 120.966  1.00  0.00           H
    ATOM   3385 HD11 LEU B 613     147.604  41.683 123.084  1.00  0.00           H
    ATOM   3386 HD12 LEU B 613     147.472  42.421 121.515  1.00  0.00           H
    ATOM   3387 HD13 LEU B 613     146.027  42.219 122.502  1.00  0.00           H
    ATOM   3388 HD21 LEU B 613     146.895  39.632 123.632  1.00  0.00           H
    ATOM   3389 HD22 LEU B 613     145.656  38.809 122.708  1.00  0.00           H
    ATOM   3390 HD23 LEU B 613     147.345  38.467 122.409  1.00  0.00           H
    ATOM   3391  N   PRO B 614     145.606  41.562 117.472  1.00 42.94           N
    ATOM   3392  CA  PRO B 614     145.388  41.129 116.075  1.00 42.94           C
    ATOM   3393  C   PRO B 614     145.014  39.649 115.995  1.00 42.94           C
    ATOM   3394  O   PRO B 614     144.134  39.272 115.233  1.00 42.94           O
    ATOM   3395  CB  PRO B 614     146.737  41.383 115.390  1.00 68.43           C
    ATOM   3396  CG  PRO B 614     147.421  42.417 116.252  1.00 68.43           C
    ATOM   3397  CD  PRO B 614     146.979  42.029 117.652  1.00 68.43           C
    ATOM   3398  HA  PRO B 614     144.604  41.741 115.627  1.00  0.00           H
    ATOM   3399  HB2 PRO B 614     147.364  40.494 115.374  1.00  0.00           H
    ATOM   3400  HB3 PRO B 614     146.619  41.701 114.365  1.00  0.00           H
    ATOM   3401  HG2 PRO B 614     148.504  42.441 116.130  1.00  0.00           H
    ATOM   3402  HG3 PRO B 614     147.023  43.401 116.014  1.00  0.00           H
    ATOM   3403  HD2 PRO B 614     147.584  41.212 118.047  1.00  0.00           H
    ATOM   3404  HD3 PRO B 614     147.077  42.882 118.315  1.00  0.00           H
    ATOM   3405  N   VAL B 615     145.638  38.885 116.901  1.00 41.98           N
    ATOM   3406  CA  VAL B 615     145.366  37.502 117.228  1.00 41.98           C
    ATOM   3407  C   VAL B 615     143.870  37.257 117.479  1.00 41.98           C
    ATOM   3408  O   VAL B 615     143.292  36.451 116.763  1.00 41.98           O
    ATOM   3409  CB  VAL B 615     146.144  37.057 118.498  1.00 41.79           C
    ATOM   3410  CG1 VAL B 615     146.266  35.526 118.540  1.00 41.79           C
    ATOM   3411  CG2 VAL B 615     147.531  37.729 118.648  1.00 41.79           C
    ATOM   3412  H   VAL B 615     146.347  39.317 117.476  1.00  0.00           H
    ATOM   3413  HA  VAL B 615     145.676  36.890 116.381  1.00  0.00           H
    ATOM   3414  HB  VAL B 615     145.590  37.324 119.399  1.00  0.00           H
    ATOM   3415 HG11 VAL B 615     146.749  35.191 119.459  1.00  0.00           H
    ATOM   3416 HG12 VAL B 615     145.290  35.049 118.494  1.00  0.00           H
    ATOM   3417 HG13 VAL B 615     146.853  35.148 117.703  1.00  0.00           H
    ATOM   3418 HG21 VAL B 615     148.164  37.191 119.354  1.00  0.00           H
    ATOM   3419 HG22 VAL B 615     148.066  37.803 117.708  1.00  0.00           H
    ATOM   3420 HG23 VAL B 615     147.469  38.751 119.009  1.00  0.00           H
    ATOM   3421  N   ASP B 616     143.266  38.007 118.420  1.00 74.90           N
    ATOM   3422  CA  ASP B 616     141.836  37.969 118.736  1.00 74.90           C
    ATOM   3423  C   ASP B 616     140.947  38.223 117.515  1.00 74.90           C
    ATOM   3424  O   ASP B 616     140.022  37.449 117.307  1.00 74.90           O
    ATOM   3425  CB  ASP B 616     141.431  38.893 119.907  1.00 99.90           C
    ATOM   3426  CG  ASP B 616     142.205  38.638 121.204  1.00 99.90           C
    ATOM   3427  OD1 ASP B 616     142.424  37.450 121.524  1.00 99.90           O
    ATOM   3428  OD2 ASP B 616     142.490  39.635 121.903  1.00 99.90           O1-
    ATOM   3429  H   ASP B 616     143.812  38.672 118.947  1.00  0.00           H
    ATOM   3430  HA  ASP B 616     141.633  36.943 119.052  1.00  0.00           H
    ATOM   3431  HB2 ASP B 616     141.596  39.928 119.612  1.00  0.00           H
    ATOM   3432  HB3 ASP B 616     140.366  38.798 120.124  1.00  0.00           H
    ATOM   3433  N   LEU B 617     141.283  39.235 116.698  1.00 51.95           N
    ATOM   3434  CA  LEU B 617     140.600  39.534 115.438  1.00 51.95           C
    ATOM   3435  C   LEU B 617     140.651  38.376 114.436  1.00 51.95           C
    ATOM   3436  O   LEU B 617     139.598  37.918 113.997  1.00 51.95           O
    ATOM   3437  CB  LEU B 617     141.206  40.794 114.802  1.00 28.20           C
    ATOM   3438  CG  LEU B 617     140.867  42.090 115.547  1.00 28.20           C
    ATOM   3439  CD1 LEU B 617     141.703  43.259 115.015  1.00 28.20           C
    ATOM   3440  CD2 LEU B 617     139.368  42.407 115.470  1.00 28.20           C
    ATOM   3441  H   LEU B 617     142.081  39.810 116.927  1.00  0.00           H
    ATOM   3442  HA  LEU B 617     139.547  39.706 115.664  1.00  0.00           H
    ATOM   3443  HB2 LEU B 617     142.284  40.697 114.728  1.00  0.00           H
    ATOM   3444  HB3 LEU B 617     140.864  40.871 113.775  1.00  0.00           H
    ATOM   3445  HG  LEU B 617     141.136  41.963 116.597  1.00  0.00           H
    ATOM   3446 HD11 LEU B 617     141.964  43.936 115.828  1.00  0.00           H
    ATOM   3447 HD12 LEU B 617     142.629  42.920 114.551  1.00  0.00           H
    ATOM   3448 HD13 LEU B 617     141.160  43.838 114.268  1.00  0.00           H
    ATOM   3449 HD21 LEU B 617     139.171  43.441 115.749  1.00  0.00           H
    ATOM   3450 HD22 LEU B 617     138.972  42.247 114.466  1.00  0.00           H
    ATOM   3451 HD23 LEU B 617     138.791  41.783 116.151  1.00  0.00           H
    ATOM   3452  N   ALA B 618     141.869  37.912 114.121  1.00 33.33           N
    ATOM   3453  CA  ALA B 618     142.113  36.773 113.242  1.00 33.33           C
    ATOM   3454  C   ALA B 618     141.457  35.480 113.729  1.00 33.33           C
    ATOM   3455  O   ALA B 618     140.815  34.810 112.925  1.00 33.33           O
    ATOM   3456  CB  ALA B 618     143.615  36.565 113.063  1.00 84.83           C
    ATOM   3457  H   ALA B 618     142.687  38.340 114.538  1.00  0.00           H
    ATOM   3458  HA  ALA B 618     141.684  37.009 112.267  1.00  0.00           H
    ATOM   3459  HB1 ALA B 618     143.834  35.589 112.628  1.00  0.00           H
    ATOM   3460  HB2 ALA B 618     144.013  37.309 112.380  1.00  0.00           H
    ATOM   3461  HB3 ALA B 618     144.157  36.635 114.007  1.00  0.00           H
    ATOM   3462  N   GLU B 619     141.578  35.180 115.033  1.00 46.36           N
    ATOM   3463  CA  GLU B 619     140.898  34.062 115.682  1.00 46.36           C
    ATOM   3464  C   GLU B 619     139.376  34.152 115.547  1.00 46.36           C
    ATOM   3465  O   GLU B 619     138.749  33.151 115.203  1.00 46.36           O
    ATOM   3466  CB  GLU B 619     141.263  33.962 117.176  1.00 99.90           C
    ATOM   3467  CG  GLU B 619     142.611  33.277 117.452  1.00 99.90           C
    ATOM   3468  CD  GLU B 619     142.771  32.935 118.934  1.00 99.90           C
    ATOM   3469  OE1 GLU B 619     141.839  32.303 119.478  1.00 99.90           O
    ATOM   3470  OE2 GLU B 619     143.834  33.273 119.496  1.00 99.90           O1-
    ATOM   3471  H   GLU B 619     142.136  35.773 115.637  1.00  0.00           H
    ATOM   3472  HA  GLU B 619     141.209  33.143 115.182  1.00  0.00           H
    ATOM   3473  HB2 GLU B 619     141.265  34.955 117.627  1.00  0.00           H
    ATOM   3474  HB3 GLU B 619     140.471  33.418 117.697  1.00  0.00           H
    ATOM   3475  HG2 GLU B 619     142.669  32.341 116.899  1.00  0.00           H
    ATOM   3476  HG3 GLU B 619     143.443  33.892 117.110  1.00  0.00           H
    ATOM   3477  N   GLU B 620     138.832  35.351 115.805  1.00 95.11           N
    ATOM   3478  CA  GLU B 620     137.408  35.657 115.771  1.00 95.11           C
    ATOM   3479  C   GLU B 620     136.785  35.441 114.386  1.00 95.11           C
    ATOM   3480  O   GLU B 620     135.849  34.650 114.282  1.00 95.11           O
    ATOM   3481  CB  GLU B 620     137.173  37.070 116.340  1.00 99.90           C
    ATOM   3482  CG  GLU B 620     135.704  37.521 116.400  1.00 99.90           C
    ATOM   3483  CD  GLU B 620     135.520  38.921 116.993  1.00 99.90           C
    ATOM   3484  OE1 GLU B 620     136.514  39.509 117.478  1.00 99.90           O
    ATOM   3485  OE2 GLU B 620     134.363  39.389 116.945  1.00 99.90           O1-
    ATOM   3486  H   GLU B 620     139.433  36.113 116.094  1.00  0.00           H
    ATOM   3487  HA  GLU B 620     136.923  34.955 116.452  1.00  0.00           H
    ATOM   3488  HB2 GLU B 620     137.557  37.086 117.360  1.00  0.00           H
    ATOM   3489  HB3 GLU B 620     137.753  37.802 115.779  1.00  0.00           H
    ATOM   3490  HG2 GLU B 620     135.268  37.529 115.401  1.00  0.00           H
    ATOM   3491  HG3 GLU B 620     135.122  36.815 116.993  1.00  0.00           H
    ATOM   3492  N   LEU B 621     137.339  36.090 113.346  1.00 68.07           N
    ATOM   3493  CA  LEU B 621     136.878  35.944 111.959  1.00 68.07           C
    ATOM   3494  C   LEU B 621     137.109  34.545 111.357  1.00 68.07           C
    ATOM   3495  O   LEU B 621     136.535  34.258 110.307  1.00 68.07           O
    ATOM   3496  CB  LEU B 621     137.498  37.047 111.064  1.00 95.77           C
    ATOM   3497  CG  LEU B 621     136.575  38.262 110.836  1.00 95.77           C
    ATOM   3498  CD1 LEU B 621     135.312  37.897 110.024  1.00 95.77           C
    ATOM   3499  CD2 LEU B 621     136.252  39.006 112.147  1.00 95.77           C
    ATOM   3500  H   LEU B 621     138.123  36.711 113.503  1.00  0.00           H
    ATOM   3501  HA  LEU B 621     135.795  36.072 111.974  1.00  0.00           H
    ATOM   3502  HB2 LEU B 621     138.456  37.372 111.473  1.00  0.00           H
    ATOM   3503  HB3 LEU B 621     137.744  36.646 110.080  1.00  0.00           H
    ATOM   3504  HG  LEU B 621     137.152  38.955 110.222  1.00  0.00           H
    ATOM   3505 HD11 LEU B 621     135.258  38.492 109.112  1.00  0.00           H
    ATOM   3506 HD12 LEU B 621     135.306  36.850 109.719  1.00  0.00           H
    ATOM   3507 HD13 LEU B 621     134.391  38.076 110.579  1.00  0.00           H
    ATOM   3508 HD21 LEU B 621     136.305  40.085 111.998  1.00  0.00           H
    ATOM   3509 HD22 LEU B 621     135.257  38.774 112.525  1.00  0.00           H
    ATOM   3510 HD23 LEU B 621     136.960  38.764 112.940  1.00  0.00           H
    ATOM   3511  N   GLY B 622     137.903  33.696 112.029  1.00 99.90           N
    ATOM   3512  CA  GLY B 622     138.150  32.320 111.616  1.00 99.90           C
    ATOM   3513  C   GLY B 622     139.302  32.232 110.608  1.00 99.90           C
    ATOM   3514  O   GLY B 622     139.341  31.281 109.828  1.00 99.90           O
    ATOM   3515  H   GLY B 622     138.328  33.999 112.895  1.00  0.00           H
    ATOM   3516  HA2 GLY B 622     138.416  31.739 112.499  1.00  0.00           H
    ATOM   3517  HA3 GLY B 622     137.250  31.867 111.198  1.00  0.00           H
    ATOM   3518  N   HIE B 623     140.231  33.202 110.616  1.00 58.81           N
    ATOM   3519  CA  HIE B 623     141.430  33.216 109.780  1.00 58.81           C
    ATOM   3520  C   HIE B 623     142.506  32.323 110.397  1.00 58.81           C
    ATOM   3521  O   HIE B 623     143.448  32.837 110.996  1.00 58.81           O
    ATOM   3522  CB  HIE B 623     141.921  34.662 109.587  1.00 61.27           C
    ATOM   3523  CG  HIE B 623     140.945  35.575 108.897  1.00 61.27           C
    ATOM   3524  ND1 HIE B 623     140.109  35.172 107.849  1.00 61.27           N
    ATOM   3525  CD2 HIE B 623     140.706  36.902 109.153  1.00 61.27           C
    ATOM   3526  CE1 HIE B 623     139.452  36.267 107.494  1.00 61.27           C
    ATOM   3527  NE2 HIE B 623     139.787  37.328 108.220  1.00 61.27           N
    ATOM   3528  H   HIE B 623     140.159  33.950 111.295  1.00  0.00           H
    ATOM   3529  HA  HIE B 623     141.194  32.810 108.793  1.00  0.00           H
    ATOM   3530  HB2 HIE B 623     142.179  35.110 110.547  1.00  0.00           H
    ATOM   3531  HB3 HIE B 623     142.836  34.666 108.993  1.00  0.00           H
    ATOM   3532  HD2 HIE B 623     141.078  37.559 109.913  1.00  0.00           H
    ATOM   3533  HE2 HIE B 623     139.408  38.264 108.143  1.00  0.00           H
    ATOM   3534  HE1 HIE B 623     138.726  36.295 106.695  1.00  0.00           H
    ATOM   3535  N   ARG B 624     142.319  31.000 110.246  1.00 50.15           N
    ATOM   3536  CA  ARG B 624     143.134  29.930 110.818  1.00 50.15           C
    ATOM   3537  C   ARG B 624     144.640  30.149 110.634  1.00 50.15           C
    ATOM   3538  O   ARG B 624     145.391  30.068 111.598  1.00 50.15           O
    ATOM   3539  CB  ARG B 624     142.695  28.573 110.220  1.00 99.90           C
    ATOM   3540  CG  ARG B 624     142.979  27.394 111.161  1.00 99.90           C
    ATOM   3541  CD  ARG B 624     144.434  26.903 111.226  1.00 99.90           C
    ATOM   3542  NE  ARG B 624     144.677  26.187 112.483  1.00 99.90           N
    ATOM   3543  CZ  ARG B 624     145.650  25.292 112.703  1.00 99.90           C
    ATOM   3544  NH1 ARG B 624     146.503  24.942 111.731  1.00 99.90           N
    ATOM   3545  NH2 ARG B 624     145.760  24.733 113.913  1.00 99.90           N1+
    ATOM   3546  H   ARG B 624     141.500  30.698 109.735  1.00  0.00           H
    ATOM   3547  HA  ARG B 624     142.926  29.926 111.889  1.00  0.00           H
    ATOM   3548  HB2 ARG B 624     141.613  28.601 110.078  1.00  0.00           H
    ATOM   3549  HB3 ARG B 624     143.116  28.404 109.227  1.00  0.00           H
    ATOM   3550  HG2 ARG B 624     142.571  27.583 112.155  1.00  0.00           H
    ATOM   3551  HG3 ARG B 624     142.395  26.562 110.764  1.00  0.00           H
    ATOM   3552  HD2 ARG B 624     144.514  26.131 110.460  1.00  0.00           H
    ATOM   3553  HD3 ARG B 624     145.214  27.618 110.987  1.00  0.00           H
    ATOM   3554  HE  ARG B 624     144.046  26.409 113.240  1.00  0.00           H
    ATOM   3555 HH11 ARG B 624     146.406  25.344 110.810  1.00  0.00           H
    ATOM   3556 HH12 ARG B 624     147.193  24.222 111.890  1.00  0.00           H
    ATOM   3557 HH21 ARG B 624     145.090  24.947 114.639  1.00  0.00           H
    ATOM   3558 HH22 ARG B 624     146.472  24.039 114.096  1.00  0.00           H
    ATOM   3559  N   ASP B 625     145.028  30.447 109.392  1.00 52.35           N
    ATOM   3560  CA  ASP B 625     146.360  30.824 108.926  1.00 52.35           C
    ATOM   3561  C   ASP B 625     147.015  31.906 109.798  1.00 52.35           C
    ATOM   3562  O   ASP B 625     148.007  31.615 110.466  1.00 52.35           O
    ATOM   3563  CB  ASP B 625     146.330  31.281 107.447  1.00 99.90           C
    ATOM   3564  CG  ASP B 625     145.088  32.106 107.076  1.00 99.90           C
    ATOM   3565  OD1 ASP B 625     144.004  31.496 106.950  1.00 99.90           O
    ATOM   3566  OD2 ASP B 625     145.243  33.339 106.967  1.00 99.90           O1-
    ATOM   3567  H   ASP B 625     144.314  30.516 108.675  1.00  0.00           H
    ATOM   3568  HA  ASP B 625     146.980  29.929 108.988  1.00  0.00           H
    ATOM   3569  HB2 ASP B 625     147.222  31.853 107.187  1.00  0.00           H
    ATOM   3570  HB3 ASP B 625     146.370  30.419 106.781  1.00  0.00           H
    ATOM   3571  N   VAL B 626     146.427  33.117 109.801  1.00 45.06           N
    ATOM   3572  CA  VAL B 626     146.887  34.236 110.615  1.00 45.06           C
    ATOM   3573  C   VAL B 626     146.818  33.923 112.108  1.00 45.06           C
    ATOM   3574  O   VAL B 626     147.759  34.265 112.822  1.00 45.06           O
    ATOM   3575  CB  VAL B 626     146.086  35.543 110.394  1.00 57.95           C
    ATOM   3576  CG1 VAL B 626     146.567  36.656 111.341  1.00 57.95           C
    ATOM   3577  CG2 VAL B 626     146.192  36.051 108.956  1.00 57.95           C
    ATOM   3578  H   VAL B 626     145.629  33.289 109.203  1.00  0.00           H
    ATOM   3579  HA  VAL B 626     147.931  34.413 110.359  1.00  0.00           H
    ATOM   3580  HB  VAL B 626     145.027  35.361 110.583  1.00  0.00           H
    ATOM   3581 HG11 VAL B 626     146.305  37.643 110.993  1.00  0.00           H
    ATOM   3582 HG12 VAL B 626     146.160  36.543 112.341  1.00  0.00           H
    ATOM   3583 HG13 VAL B 626     147.649  36.626 111.437  1.00  0.00           H
    ATOM   3584 HG21 VAL B 626     145.640  36.981 108.826  1.00  0.00           H
    ATOM   3585 HG22 VAL B 626     147.230  36.256 108.703  1.00  0.00           H
    ATOM   3586 HG23 VAL B 626     145.799  35.340 108.237  1.00  0.00           H
    ATOM   3587  N   ALA B 627     145.746  33.228 112.530  1.00 70.54           N
    ATOM   3588  CA  ALA B 627     145.613  32.703 113.877  1.00 70.54           C
    ATOM   3589  C   ALA B 627     146.842  31.874 114.252  1.00 70.54           C
    ATOM   3590  O   ALA B 627     147.370  32.119 115.317  1.00 70.54           O
    ATOM   3591  CB  ALA B 627     144.297  31.933 114.067  1.00 81.69           C
    ATOM   3592  H   ALA B 627     145.004  32.999 111.879  1.00  0.00           H
    ATOM   3593  HA  ALA B 627     145.577  33.568 114.543  1.00  0.00           H
    ATOM   3594  HB1 ALA B 627     144.154  31.679 115.117  1.00  0.00           H
    ATOM   3595  HB2 ALA B 627     143.443  32.543 113.770  1.00  0.00           H
    ATOM   3596  HB3 ALA B 627     144.252  31.006 113.507  1.00  0.00           H
    ATOM   3597  N   ARG B 628     147.364  31.038 113.338  1.00 52.52           N
    ATOM   3598  CA  ARG B 628     148.600  30.287 113.532  1.00 52.52           C
    ATOM   3599  C   ARG B 628     149.853  31.159 113.624  1.00 52.52           C
    ATOM   3600  O   ARG B 628     150.582  31.009 114.601  1.00 52.52           O
    ATOM   3601  CB  ARG B 628     148.778  29.197 112.457  1.00 72.29           C
    ATOM   3602  CG  ARG B 628     148.881  27.811 113.100  1.00 72.29           C
    ATOM   3603  CD  ARG B 628     149.358  26.706 112.164  1.00 72.29           C
    ATOM   3604  NE  ARG B 628     150.802  26.759 111.924  1.00 72.29           N
    ATOM   3605  CZ  ARG B 628     151.461  25.929 111.098  1.00 72.29           C
    ATOM   3606  NH1 ARG B 628     150.809  24.999 110.385  1.00 72.29           N
    ATOM   3607  NH2 ARG B 628     152.790  26.027 110.990  1.00 72.29           N1+
    ATOM   3608  H   ARG B 628     146.894  30.916 112.450  1.00  0.00           H
    ATOM   3609  HA  ARG B 628     148.492  29.801 114.504  1.00  0.00           H
    ATOM   3610  HB2 ARG B 628     147.940  29.191 111.759  1.00  0.00           H
    ATOM   3611  HB3 ARG B 628     149.660  29.386 111.842  1.00  0.00           H
    ATOM   3612  HG2 ARG B 628     149.617  27.885 113.904  1.00  0.00           H
    ATOM   3613  HG3 ARG B 628     147.941  27.522 113.557  1.00  0.00           H
    ATOM   3614  HD2 ARG B 628     149.028  25.724 112.501  1.00  0.00           H
    ATOM   3615  HD3 ARG B 628     148.909  26.874 111.184  1.00  0.00           H
    ATOM   3616  HE  ARG B 628     151.325  27.444 112.451  1.00  0.00           H
    ATOM   3617 HH11 ARG B 628     149.805  24.924 110.449  1.00  0.00           H
    ATOM   3618 HH12 ARG B 628     151.313  24.376 109.770  1.00  0.00           H
    ATOM   3619 HH21 ARG B 628     153.298  26.720 111.523  1.00  0.00           H
    ATOM   3620 HH22 ARG B 628     153.302  25.413 110.372  1.00  0.00           H
    ATOM   3621  N   TYR B 629     150.084  32.056 112.647  1.00 62.83           N
    ATOM   3622  CA  TYR B 629     151.267  32.926 112.655  1.00 62.83           C
    ATOM   3623  C   TYR B 629     151.348  33.867 113.871  1.00 62.83           C
    ATOM   3624  O   TYR B 629     152.433  34.355 114.183  1.00 62.83           O
    ATOM   3625  CB  TYR B 629     151.373  33.731 111.352  1.00 66.94           C
    ATOM   3626  CG  TYR B 629     152.652  34.552 111.291  1.00 66.94           C
    ATOM   3627  CD1 TYR B 629     153.889  33.881 111.243  1.00 66.94           C
    ATOM   3628  CD2 TYR B 629     152.622  35.961 111.389  1.00 66.94           C
    ATOM   3629  CE1 TYR B 629     155.090  34.605 111.289  1.00 66.94           C
    ATOM   3630  CE2 TYR B 629     153.827  36.689 111.419  1.00 66.94           C
    ATOM   3631  CZ  TYR B 629     155.061  36.008 111.356  1.00 66.94           C
    ATOM   3632  OH  TYR B 629     156.236  36.698 111.314  1.00 66.94           O
    ATOM   3633  H   TYR B 629     149.437  32.155 111.877  1.00  0.00           H
    ATOM   3634  HA  TYR B 629     152.134  32.265 112.718  1.00  0.00           H
    ATOM   3635  HB2 TYR B 629     151.359  33.057 110.496  1.00  0.00           H
    ATOM   3636  HB3 TYR B 629     150.503  34.382 111.261  1.00  0.00           H
    ATOM   3637  HD1 TYR B 629     153.918  32.803 111.180  1.00  0.00           H
    ATOM   3638  HD2 TYR B 629     151.679  36.483 111.452  1.00  0.00           H
    ATOM   3639  HE1 TYR B 629     156.034  34.082 111.237  1.00  0.00           H
    ATOM   3640  HE2 TYR B 629     153.797  37.767 111.474  1.00  0.00           H
    ATOM   3641  HH  TYR B 629     157.009  36.128 111.312  1.00  0.00           H
    ATOM   3642  N   LEU B 630     150.208  34.093 114.530  1.00 62.94           N
    ATOM   3643  CA  LEU B 630     150.098  34.931 115.708  1.00 62.94           C
    ATOM   3644  C   LEU B 630     149.990  34.130 117.012  1.00 62.94           C
    ATOM   3645  O   LEU B 630     150.446  34.624 118.039  1.00 62.94           O
    ATOM   3646  CB  LEU B 630     148.865  35.816 115.529  1.00 41.93           C
    ATOM   3647  CG  LEU B 630     148.993  36.898 114.441  1.00 41.93           C
    ATOM   3648  CD1 LEU B 630     147.697  37.708 114.405  1.00 41.93           C
    ATOM   3649  CD2 LEU B 630     150.203  37.816 114.669  1.00 41.93           C
    ATOM   3650  H   LEU B 630     149.349  33.684 114.188  1.00  0.00           H
    ATOM   3651  HA  LEU B 630     150.986  35.553 115.817  1.00  0.00           H
    ATOM   3652  HB2 LEU B 630     147.989  35.194 115.338  1.00  0.00           H
    ATOM   3653  HB3 LEU B 630     148.669  36.301 116.475  1.00  0.00           H
    ATOM   3654  HG  LEU B 630     149.111  36.409 113.474  1.00  0.00           H
    ATOM   3655 HD11 LEU B 630     147.614  38.336 113.524  1.00  0.00           H
    ATOM   3656 HD12 LEU B 630     146.820  37.064 114.433  1.00  0.00           H
    ATOM   3657 HD13 LEU B 630     147.654  38.354 115.276  1.00  0.00           H
    ATOM   3658 HD21 LEU B 630     149.964  38.864 114.489  1.00  0.00           H
    ATOM   3659 HD22 LEU B 630     150.553  37.728 115.694  1.00  0.00           H
    ATOM   3660 HD23 LEU B 630     151.030  37.546 114.012  1.00  0.00           H
    ATOM   3661  N   ARG B 631     149.452  32.902 116.966  1.00 69.32           N
    ATOM   3662  CA  ARG B 631     149.395  31.939 118.077  1.00 69.32           C
    ATOM   3663  C   ARG B 631     150.701  31.120 118.112  1.00 69.32           C
    ATOM   3664  O   ARG B 631     150.697  29.898 118.238  1.00 69.32           O
    ATOM   3665  CB  ARG B 631     148.160  31.055 117.857  1.00 99.90           C
    ATOM   3666  CG  ARG B 631     147.708  30.104 118.974  1.00 99.90           C
    ATOM   3667  CD  ARG B 631     147.109  28.814 118.394  1.00 99.90           C
    ATOM   3668  NE  ARG B 631     146.193  29.081 117.267  1.00 99.90           N
    ATOM   3669  CZ  ARG B 631     146.198  28.434 116.089  1.00 99.90           C
    ATOM   3670  NH1 ARG B 631     147.012  27.393 115.880  1.00 99.90           N
    ATOM   3671  NH2 ARG B 631     145.374  28.830 115.109  1.00 99.90           N1+
    ATOM   3672  H   ARG B 631     149.070  32.569 116.091  1.00  0.00           H
    ATOM   3673  HA  ARG B 631     149.293  32.465 119.028  1.00  0.00           H
    ATOM   3674  HB2 ARG B 631     147.296  31.689 117.662  1.00  0.00           H
    ATOM   3675  HB3 ARG B 631     148.357  30.478 116.952  1.00  0.00           H
    ATOM   3676  HG2 ARG B 631     148.591  29.789 119.526  1.00  0.00           H
    ATOM   3677  HG3 ARG B 631     147.072  30.587 119.716  1.00  0.00           H
    ATOM   3678  HD2 ARG B 631     147.885  28.075 118.196  1.00  0.00           H
    ATOM   3679  HD3 ARG B 631     146.459  28.373 119.151  1.00  0.00           H
    ATOM   3680  HE  ARG B 631     145.539  29.839 117.406  1.00  0.00           H
    ATOM   3681 HH11 ARG B 631     147.639  27.097 116.615  1.00  0.00           H
    ATOM   3682 HH12 ARG B 631     146.997  26.892 115.005  1.00  0.00           H
    ATOM   3683 HH21 ARG B 631     144.733  29.594 115.269  1.00  0.00           H
    ATOM   3684 HH22 ARG B 631     145.419  28.413 114.191  1.00  0.00           H
    ATOM   3685  N   ALA B 632     151.798  31.853 117.942  1.00 64.17           N
    ATOM   3686  CA  ALA B 632     153.171  31.468 117.675  1.00 64.17           C
    ATOM   3687  C   ALA B 632     153.853  32.787 117.306  1.00 64.17           C
    ATOM   3688  O   ALA B 632     153.174  33.744 116.928  1.00 64.17           O
    ATOM   3689  CB  ALA B 632     153.282  30.477 116.501  1.00 61.97           C
    ATOM   3690  H   ALA B 632     151.658  32.852 117.859  1.00  0.00           H
    ATOM   3691  HA  ALA B 632     153.612  31.051 118.582  1.00  0.00           H
    ATOM   3692  HB1 ALA B 632     154.319  30.181 116.345  1.00  0.00           H
    ATOM   3693  HB2 ALA B 632     152.717  29.562 116.674  1.00  0.00           H
    ATOM   3694  HB3 ALA B 632     152.932  30.919 115.568  1.00  0.00           H
    ATOM   3695  N   ALA B 633     155.182  32.859 117.475  1.00 99.90           N
    ATOM   3696  CA  ALA B 633     155.944  34.113 117.348  1.00 99.90           C
    ATOM   3697  C   ALA B 633     155.487  35.197 118.358  1.00 99.90           C
    ATOM   3698  O   ALA B 633     155.765  36.378 118.158  1.00 99.90           O
    ATOM   3699  CB  ALA B 633     155.901  34.620 115.886  1.00 99.90           C
    ATOM   3700  H   ALA B 633     155.699  32.047 117.780  1.00  0.00           H
    ATOM   3701  HA  ALA B 633     156.979  33.871 117.592  1.00  0.00           H
    ATOM   3702  HB1 ALA B 633     156.628  35.418 115.730  1.00  0.00           H
    ATOM   3703  HB2 ALA B 633     156.146  33.822 115.184  1.00  0.00           H
    ATOM   3704  HB3 ALA B 633     154.927  35.022 115.606  1.00  0.00           H
    ATOM   3705  N   ALA B 634     154.779  34.756 119.413  1.00 99.90           N
    ATOM   3706  CA  ALA B 634     154.069  35.542 120.416  1.00 99.90           C
    ATOM   3707  C   ALA B 634     153.517  34.468 121.334  1.00 99.90           C
    ATOM   3708  O   ALA B 634     152.400  34.516 121.849  1.00 99.90           O
    ATOM   3709  CB  ALA B 634     152.951  36.406 119.790  1.00 99.90           C
    ATOM   3710  H   ALA B 634     154.634  33.760 119.484  1.00  0.00           H
    ATOM   3711  HA  ALA B 634     154.775  36.158 120.975  1.00  0.00           H
    ATOM   3713  HB1 ALA B 634     152.350  36.896 120.555  1.00  0.00           H
    ATOM   3714  HB2 ALA B 634     153.360  37.191 119.158  1.00  0.00           H
    ATOM   3715  HB3 ALA B 634     152.281  35.815 119.171  1.00  0.00           H
  extra_rotamers: ""
ligand:
  residue_configurations:
  - mutability:
    - ALA
    - ARG
    - ASN
    - ASP
    - CYS
    - GLU
    - GLN
    - GLY
    - HIE
    - HID
    - HIP
    - ILE
    - LEU
    - LYS
    - MET
    - PHE
    - PRO
    - SER
    - THR
    - TRP
    - TYR
    - VAL
    flexibility:
      is_flexible: true
      include_structure_rotamer: true
      use_continuous: true
    identity:
      chain: A
      res_num: 14
      aa_type: GLU
  - mutability:
    - ALA
    - ARG
    - ASN
    - ASP
    - CYS
    - GLU
    - GLN
    - GLY
    - HIE
    - HID
    - HIP
    - ILE
    - LEU
    - LYS
    - MET
    - PHE
    - PRO
    - SER
    - THR
    - TRP
    - TYR
    - VAL
    flexibility:
      is_flexible: true
      include_structure_rotamer: true
      use_continuous: true
    identity:
      chain: A
      res_num: 15
      aa_type: CYS
  extra_templates: ""
  extra_templates_coordinates: ""
  coordinates: |
    ATOM      1  N   ASP A  10     123.857  61.443 110.351  1.00 51.43           N1+
    ATOM      2  CA  ASP A  10     125.302  61.561 110.638  1.00 51.43           C
    ATOM      3  C   ASP A  10     125.937  60.158 110.637  1.00 51.43           C
    ATOM      4  O   ASP A  10     125.311  59.176 111.036  1.00 51.43           O
    ATOM      5  CB  ASP A  10     125.579  62.240 112.013  1.00 43.01           C
    ATOM      6  CG  ASP A  10     126.198  63.638 111.954  1.00 43.01           C
    ATOM      7  OD1 ASP A  10     126.017  64.481 112.827  1.00 43.01           O
    ATOM      8  OD2 ASP A  10     127.019  63.840 110.885  1.00 43.01           O
    ATOM      9  H   ASP A  10     123.384  62.329 110.456  1.00  0.00           H
    ATOM     11  HA  ASP A  10     125.735  62.117 109.804  1.00  0.00           H
    ATOM     12  HB2 ASP A  10     124.660  62.309 112.598  1.00  0.00           H
    ATOM     13  HB3 ASP A  10     126.256  61.656 112.642  1.00  0.00           H
    ATOM     15  N   GLN A  11     127.220  60.111 110.251  1.00 35.86           N
    ATOM     16  CA  GLN A  11     127.978  58.876 110.116  1.00 35.86           C
    ATOM     17  C   GLN A  11     128.800  58.588 111.373  1.00 35.86           C
    ATOM     18  O   GLN A  11     129.624  59.394 111.806  1.00 35.86           O
    ATOM     19  CB  GLN A  11     128.913  58.941 108.901  1.00 86.13           C
    ATOM     20  CG  GLN A  11     128.217  59.389 107.609  1.00 86.13           C
    ATOM     21  CD  GLN A  11     128.394  60.886 107.349  1.00 86.13           C
    ATOM     22  OE1 GLN A  11     128.518  61.693 108.267  1.00 86.13           O
    ATOM     23  NE2 GLN A  11     128.407  61.280 106.083  1.00 86.13           N
    ATOM     24  H   GLN A  11     127.693  60.956 109.958  1.00  0.00           H
    ATOM     25  HA  GLN A  11     127.288  58.048 109.936  1.00  0.00           H
    ATOM     26  HB2 GLN A  11     129.779  59.575 109.106  1.00  0.00           H
    ATOM     27  HB3 GLN A  11     129.343  57.954 108.744  1.00  0.00           H
    ATOM     28  HG2 GLN A  11     128.692  58.854 106.794  1.00  0.00           H
    ATOM     29  HG3 GLN A  11     127.162  59.113 107.590  1.00  0.00           H
    ATOM     30 HE21 GLN A  11     128.222  60.619 105.336  1.00  0.00           H
    ATOM     31 HE22 GLN A  11     128.511  62.260 105.865  1.00  0.00           H
    ATOM     32  N   GLN A  12     128.581  57.374 111.870  1.00 80.65           N
    ATOM     33  CA  GLN A  12     129.406  56.664 112.836  1.00 80.65           C
    ATOM     34  C   GLN A  12     130.692  56.113 112.200  1.00 80.65           C
    ATOM     35  O   GLN A  12     131.712  56.008 112.875  1.00 80.65           O
    ATOM     36  CB  GLN A  12     128.556  55.521 113.412  1.00 99.90           C
    ATOM     37  CG  GLN A  12     127.716  54.797 112.335  1.00 99.90           C
    ATOM     38  CD  GLN A  12     127.513  53.348 112.690  1.00 99.90           C
    ATOM     39  OE1 GLN A  12     126.615  53.015 113.456  1.00 99.90           O
    ATOM     40  NE2 GLN A  12     128.378  52.500 112.141  1.00 99.90           N
    ATOM     41  H   GLN A  12     127.839  56.823 111.465  1.00  0.00           H
    ATOM     42  HA  GLN A  12     129.688  57.344 113.642  1.00  0.00           H
    ATOM     43  HB2 GLN A  12     129.211  54.823 113.938  1.00  0.00           H
    ATOM     44  HB3 GLN A  12     127.884  55.915 114.175  1.00  0.00           H
    ATOM     45  HG2 GLN A  12     126.745  55.282 112.214  1.00  0.00           H
    ATOM     46  HG3 GLN A  12     128.190  54.826 111.355  1.00  0.00           H
    ATOM     47 HE21 GLN A  12     129.083  52.830 111.499  1.00  0.00           H
    ATOM     48 HE22 GLN A  12     128.318  51.512 112.335  1.00  0.00           H
    ATOM     49  N   TYR A  13     130.613  55.788 110.901  1.00 55.68           N
    ATOM     50  CA  TYR A  13     131.722  55.358 110.065  1.00 55.68           C
    ATOM     51  C   TYR A  13     132.649  56.551 109.857  1.00 55.68           C
    ATOM     52  O   TYR A  13     132.188  57.571 109.342  1.00 55.68           O
    ATOM     53  CB  TYR A  13     131.207  54.869 108.697  1.00 42.90           C
    ATOM     54  CG  TYR A  13     129.866  54.171 108.737  1.00 42.90           C
    ATOM     55  CD1 TYR A  13     129.796  52.797 109.003  1.00 42.90           C
    ATOM     56  CD2 TYR A  13     128.679  54.896 108.503  1.00 42.90           C
    ATOM     57  CE1 TYR A  13     128.552  52.147 109.001  1.00 42.90           C
    ATOM     58  CE2 TYR A  13     127.430  54.251 108.561  1.00 42.90           C
    ATOM     59  CZ  TYR A  13     127.378  52.871 108.812  1.00 42.90           C
    ATOM     60  OH  TYR A  13     126.200  52.206 108.877  1.00 42.90           O
    ATOM     61  H   TYR A  13     129.732  55.908 110.423  1.00  0.00           H
    ATOM     62  HA  TYR A  13     132.239  54.552 110.575  1.00  0.00           H
    ATOM     63  HB2 TYR A  13     131.081  55.709 108.013  1.00  0.00           H
    ATOM     64  HB3 TYR A  13     131.951  54.225 108.225  1.00  0.00           H
    ATOM     65  HD1 TYR A  13     130.698  52.237 109.129  1.00  0.00           H
    ATOM     66  HD2 TYR A  13     128.716  55.949 108.281  1.00  0.00           H
    ATOM     67  HE1 TYR A  13     128.460  51.079 109.065  1.00  0.00           H
    ATOM     68  HE2 TYR A  13     126.522  54.812 108.397  1.00  0.00           H
    ATOM     69  HH  TYR A  13     125.443  52.765 108.662  1.00  0.00           H
    ATOM     70  N   GLU A  14     133.913  56.420 110.272  1.00 40.29           N
    ATOM     71  CA  GLU A  14     134.869  57.506 110.143  1.00 40.29           C
    ATOM     72  C   GLU A  14     135.697  57.302 108.887  1.00 40.29           C
    ATOM     73  O   GLU A  14     136.418  56.309 108.801  1.00 40.29           O
    ATOM     74  CB  GLU A  14     135.715  57.625 111.414  1.00 87.23           C
    ATOM     75  CG  GLU A  14     136.636  58.867 111.386  1.00 87.23           C
    ATOM     76  CD  GLU A  14     137.182  59.240 112.763  1.00 87.23           C
    ATOM     77  OE1 GLU A  14     136.459  59.021 113.757  1.00 87.23           O
    ATOM     78  OE2 GLU A  14     138.311  59.769 112.807  1.00 87.23           O1-
    ATOM     79  H   GLU A  14     134.237  55.557 110.690  1.00  0.00           H
    ATOM     80  HA  GLU A  14     134.340  58.458 110.056  1.00  0.00           H
    ATOM     81  HB2 GLU A  14     135.025  57.680 112.256  1.00  0.00           H
    ATOM     82  HB3 GLU A  14     136.311  56.724 111.568  1.00  0.00           H
    ATOM     83  HG2 GLU A  14     137.466  58.710 110.696  1.00  0.00           H
    ATOM     84  HG3 GLU A  14     136.085  59.734 111.021  1.00  0.00           H
    ATOM     85  N   CYS A  15     135.574  58.263 107.956  1.00 41.12           N
    ATOM     86  CA  CYS A  15     136.356  58.346 106.727  1.00 41.12           C
    ATOM     87  C   CYS A  15     137.845  58.264 107.044  1.00 41.12           C
    ATOM     88  O   CYS A  15     138.383  59.157 107.696  1.00 41.12           O
    ATOM     89  CB  CYS A  15     136.101  59.642 105.949  1.00 65.17           C
    ATOM     90  SG  CYS A  15     134.757  59.444 104.777  1.00 65.17           S
    ATOM     91  H   CYS A  15     134.944  59.033 108.128  1.00  0.00           H
    ATOM     92  HA  CYS A  15     136.091  57.496 106.099  1.00  0.00           H
    ATOM     93  HB2 CYS A  15     135.940  60.501 106.601  1.00  0.00           H
    ATOM     94  HB3 CYS A  15     136.959  59.855 105.315  1.00  0.00           H
    ATOM     95  HG  CYS A  15     135.402  58.613 103.953  1.00  0.00           H
    ATOM     96  N   VAL A  16     138.459  57.164 106.607  1.00 40.11           N
    ATOM     97  CA  VAL A  16     139.866  56.916 106.832  1.00 40.11           C
    ATOM     98  C   VAL A  16     140.714  57.388 105.637  1.00 40.11           C
    ATOM     99  O   VAL A  16     141.857  57.767 105.869  1.00 40.11           O
    ATOM    100  CB  VAL A  16     140.130  55.434 107.203  1.00 70.79           C
    ATOM    101  CG1 VAL A  16     140.140  54.412 106.065  1.00 70.79           C
    ATOM    102  CG2 VAL A  16     141.401  55.277 108.030  1.00 70.79           C
    ATOM    103  H   VAL A  16     137.957  56.478 106.059  1.00  0.00           H
    ATOM    104  HA  VAL A  16     140.198  57.509 107.687  1.00  0.00           H
    ATOM    105  HB  VAL A  16     139.317  55.137 107.860  1.00  0.00           H
    ATOM    106 HG11 VAL A  16     140.739  53.539 106.320  1.00  0.00           H
    ATOM    107 HG12 VAL A  16     139.142  54.031 105.910  1.00  0.00           H
    ATOM    108 HG13 VAL A  16     140.533  54.797 105.129  1.00  0.00           H
    ATOM    109 HG21 VAL A  16     141.545  54.249 108.363  1.00  0.00           H
    ATOM    110 HG22 VAL A  16     142.280  55.562 107.451  1.00  0.00           H
    ATOM    111 HG23 VAL A  16     141.346  55.897 108.925  1.00  0.00           H
    ATOM    112  N   ALA A  17     140.152  57.430 104.414  1.00 38.63           N
    ATOM    113  CA  ALA A  17     140.819  57.983 103.230  1.00 38.63           C
    ATOM    114  C   ALA A  17     139.910  57.949 102.018  1.00 38.63           C
    ATOM    115  O   ALA A  17     139.157  56.997 101.867  1.00 38.63           O
    ATOM    116  CB  ALA A  17     142.066  57.177 102.837  1.00 30.90           C
    ATOM    117  H   ALA A  17     139.205  57.102 104.281  1.00  0.00           H
    ATOM    118  HA  ALA A  17     141.094  59.020 103.435  1.00  0.00           H
    ATOM    119  HB1 ALA A  17     142.458  57.490 101.867  1.00  0.00           H
    ATOM    120  HB2 ALA A  17     142.887  57.323 103.532  1.00  0.00           H
    ATOM    121  HB3 ALA A  17     141.833  56.116 102.755  1.00  0.00           H
    ATOM    122  N   GLU A  18     140.101  58.915 101.110  1.00 44.54           N
    ATOM    123  CA  GLU A  18     139.543  58.880  99.759  1.00 44.54           C
    ATOM    124  C   GLU A  18     140.098  57.682  98.979  1.00 44.54           C
    ATOM    125  O   GLU A  18     141.287  57.388  99.083  1.00 44.54           O
    ATOM    126  CB  GLU A  18     139.879  60.193  99.031  1.00 99.90           C
    ATOM    127  CG  GLU A  18     139.210  61.414  99.676  1.00 99.90           C
    ATOM    128  CD  GLU A  18     139.438  62.709  98.904  1.00 99.90           C
    ATOM    129  OE1 GLU A  18     139.125  62.713  97.694  1.00 99.90           O
    ATOM    130  OE2 GLU A  18     139.859  63.694  99.548  1.00 99.90           O1-
    ATOM    131  H   GLU A  18     140.760  59.657 101.296  1.00  0.00           H
    ATOM    132  HA  GLU A  18     138.459  58.783  99.828  1.00  0.00           H
    ATOM    133  HB2 GLU A  18     140.959  60.344  99.011  1.00  0.00           H
    ATOM    134  HB3 GLU A  18     139.565  60.113  97.989  1.00  0.00           H
    ATOM    135  HG2 GLU A  18     138.140  61.243  99.666  1.00  0.00           H
    ATOM    136  HG3 GLU A  18     139.518  61.538 100.715  1.00  0.00           H
    ATOM    137  N   ILE A  19     139.226  57.001  98.232  1.00 29.62           N
    ATOM    138  CA  ILE A  19     139.599  55.839  97.427  1.00 29.62           C
    ATOM    139  C   ILE A  19     138.985  55.888  96.026  1.00 29.62           C
    ATOM    140  O   ILE A  19     139.106  54.905  95.300  1.00 29.62           O
    ATOM    141  CB  ILE A  19     139.219  54.492  98.101  1.00 26.84           C
    ATOM    142  CG1 ILE A  19     137.699  54.262  98.245  1.00 26.84           C
    ATOM    143  CG2 ILE A  19     139.972  54.329  99.428  1.00 26.84           C
    ATOM    144  CD1 ILE A  19     137.323  52.940  98.918  1.00 26.84           C
    ATOM    145  H   ILE A  19     138.249  57.267  98.211  1.00  0.00           H
    ATOM    146  HA  ILE A  19     140.676  55.837  97.249  1.00  0.00           H
    ATOM    147  HB  ILE A  19     139.597  53.691  97.464  1.00  0.00           H
    ATOM    148 HG12 ILE A  19     137.286  55.068  98.825  1.00  0.00           H
    ATOM    149 HG13 ILE A  19     137.208  54.285  97.276  1.00  0.00           H
    ATOM    150 HG21 ILE A  19     139.935  53.295  99.767  1.00  0.00           H
    ATOM    151 HG22 ILE A  19     141.029  54.571  99.309  1.00  0.00           H
    ATOM    152 HG23 ILE A  19     139.574  54.974 100.207  1.00  0.00           H
    ATOM    153 HD11 ILE A  19     136.252  52.754  98.916  1.00  0.00           H
    ATOM    154 HD12 ILE A  19     137.789  52.101  98.398  1.00  0.00           H
    ATOM    155 HD13 ILE A  19     137.649  52.897  99.955  1.00  0.00           H
    ATOM    156  N   GLY A  20     138.371  57.016  95.655  1.00 50.81           N
    ATOM    157  CA  GLY A  20     137.883  57.202  94.309  1.00 50.81           C
    ATOM    158  C   GLY A  20     136.731  58.185  94.280  1.00 50.81           C
    ATOM    159  O   GLY A  20     136.169  58.573  95.302  1.00 50.81           O
    ATOM    160  H   GLY A  20     138.270  57.787  96.300  1.00  0.00           H
    ATOM    161  HA2 GLY A  20     138.707  57.613  93.728  1.00  0.00           H
    ATOM    162  HA3 GLY A  20     137.547  56.267  93.860  1.00  0.00           H
    ATOM    163  N   GLU A  21     136.400  58.573  93.051  1.00 54.53           N
    ATOM    164  CA  GLU A  21     135.363  59.529  92.739  1.00 54.53           C
    ATOM    165  C   GLU A  21     134.538  58.859  91.649  1.00 54.53           C
    ATOM    166  O   GLU A  21     134.948  58.844  90.491  1.00 54.53           O
    ATOM    167  CB  GLU A  21     136.044  60.848  92.324  1.00 99.90           C
    ATOM    168  CG  GLU A  21     135.097  62.048  92.165  1.00 99.90           C
    ATOM    169  CD  GLU A  21     135.877  63.295  91.739  1.00 99.90           C
    ATOM    170  OE1 GLU A  21     136.343  63.299  90.578  1.00 99.90           O
    ATOM    171  OE2 GLU A  21     136.011  64.212  92.580  1.00 99.90           O1-
    ATOM    172  H   GLU A  21     136.895  58.197  92.255  1.00  0.00           H
    ATOM    173  HA  GLU A  21     134.725  59.718  93.595  1.00  0.00           H
    ATOM    174  HB2 GLU A  21     136.797  61.105  93.072  1.00  0.00           H
    ATOM    175  HB3 GLU A  21     136.600  60.697  91.397  1.00  0.00           H
    ATOM    176  HG2 GLU A  21     134.337  61.836  91.411  1.00  0.00           H
    ATOM    177  HG3 GLU A  21     134.568  62.244  93.098  1.00  0.00           H
    ATOM    178  N   GLY A  22     133.405  58.272  92.048  1.00 47.96           N
    ATOM    179  CA  GLY A  22     132.417  57.767  91.107  1.00 47.96           C
    ATOM    180  C   GLY A  22     131.586  58.938  90.571  1.00 47.96           C
    ATOM    181  O   GLY A  22     131.731  60.082  91.004  1.00 47.96           O
    ATOM    182  H   GLY A  22     133.126  58.336  93.017  1.00  0.00           H
    ATOM    183  HA2 GLY A  22     132.890  57.242  90.277  1.00  0.00           H
    ATOM    184  HA3 GLY A  22     131.782  57.050  91.611  1.00  0.00           H
    ATOM    185  N   TYR A  24     128.176  59.515  90.794  1.00 99.90           N1+
    ATOM    186  CA  TYR A  24     127.328  59.828  91.947  1.00 99.90           C
    ATOM    187  C   TYR A  24     127.985  60.644  93.081  1.00 99.90           C
    ATOM    188  O   TYR A  24     127.249  61.270  93.844  1.00 99.90           O
    ATOM    189  CB  TYR A  24     126.764  58.490  92.482  1.00 99.90           C
    ATOM    190  CG  TYR A  24     127.762  57.340  92.440  1.00 99.90           C
    ATOM    191  CD1 TYR A  24     128.871  57.334  93.310  1.00 99.90           C
    ATOM    192  CD2 TYR A  24     127.638  56.331  91.460  1.00 99.90           C
    ATOM    193  CE1 TYR A  24     129.863  56.347  93.180  1.00 99.90           C
    ATOM    194  CE2 TYR A  24     128.627  55.336  91.336  1.00 99.90           C
    ATOM    195  CZ  TYR A  24     129.743  55.349  92.195  1.00 99.90           C
    ATOM    196  OH  TYR A  24     130.720  54.404  92.094  1.00 99.90           O
    ATOM    197  H   TYR A  24     128.232  58.530  90.581  1.00  0.00           H
    ATOM    198  HA  TYR A  24     126.495  60.424  91.571  1.00  0.00           H
    ATOM    199  HB2 TYR A  24     126.358  58.597  93.487  1.00  0.00           H
    ATOM    200  HB3 TYR A  24     125.913  58.207  91.861  1.00  0.00           H
    ATOM    201  HD1 TYR A  24     128.981  58.096  94.066  1.00  0.00           H
    ATOM    202  HD2 TYR A  24     126.800  56.332  90.778  1.00  0.00           H
    ATOM    203  HE1 TYR A  24     130.717  56.357  93.842  1.00  0.00           H
    ATOM    204  HE2 TYR A  24     128.534  54.574  90.576  1.00  0.00           H
    ATOM    205  HH  TYR A  24     130.694  53.840  91.310  1.00  0.00           H
    ATOM    206  N   GLY A  25     129.324  60.639  93.184  1.00 99.90           N
    ATOM    207  CA  GLY A  25     130.035  61.383  94.218  1.00 99.90           C
    ATOM    208  C   GLY A  25     131.346  60.677  94.547  1.00 99.90           C
    ATOM    209  O   GLY A  25     131.955  60.038  93.700  1.00 99.90           O
    ATOM    210  H   GLY A  25     129.888  60.131  92.517  1.00  0.00           H
    ATOM    211  HA2 GLY A  25     130.250  62.389  93.856  1.00  0.00           H
    ATOM    212  HA3 GLY A  25     129.439  61.479  95.127  1.00  0.00           H
    ATOM    213  N   LYS A  26     131.812  60.834  95.788  1.00 84.58           N
    ATOM    214  CA  LYS A  26     133.071  60.250  96.238  1.00 84.58           C
    ATOM    215  C   LYS A  26     132.846  58.922  96.945  1.00 84.58           C
    ATOM    216  O   LYS A  26     131.767  58.680  97.485  1.00 84.58           O
    ATOM    217  CB  LYS A  26     133.793  61.239  97.157  1.00 99.90           C
    ATOM    218  CG  LYS A  26     134.305  62.476  96.410  1.00 99.90           C
    ATOM    219  CD  LYS A  26     135.179  63.339  97.323  1.00 99.90           C
    ATOM    220  CE  LYS A  26     135.702  64.603  96.635  1.00 99.90           C
    ATOM    221  NZ  LYS A  26     136.627  65.328  97.522  1.00 99.90           N1+
    ATOM    222  H   LYS A  26     131.283  61.357  96.470  1.00  0.00           H
    ATOM    223  HA  LYS A  26     133.729  60.064  95.396  1.00  0.00           H
    ATOM    224  HB2 LYS A  26     133.146  61.534  97.985  1.00  0.00           H
    ATOM    225  HB3 LYS A  26     134.654  60.742  97.608  1.00  0.00           H
    ATOM    226  HG2 LYS A  26     134.886  62.168  95.540  1.00  0.00           H
    ATOM    227  HG3 LYS A  26     133.466  63.064  96.034  1.00  0.00           H
    ATOM    228  HD2 LYS A  26     134.612  63.617  98.212  1.00  0.00           H
    ATOM    229  HD3 LYS A  26     136.024  62.745  97.675  1.00  0.00           H
    ATOM    230  HE2 LYS A  26     136.234  64.342  95.719  1.00  0.00           H
    ATOM    231  HE3 LYS A  26     134.877  65.259  96.356  1.00  0.00           H
    ATOM    232  HZ1 LYS A  26     137.411  64.728  97.745  1.00  0.00           H
    ATOM    233  HZ2 LYS A  26     136.968  66.154  97.052  1.00  0.00           H
    ATOM    234  HZ3 LYS A  26     136.153  65.593  98.373  1.00  0.00           H
    ATOM    235  N   VAL A  27     133.915  58.122  96.961  1.00 38.39           N
    ATOM    236  CA  VAL A  27     134.014  56.913  97.744  1.00 38.39           C
    ATOM    237  C   VAL A  27     135.310  56.943  98.549  1.00 38.39           C
    ATOM    238  O   VAL A  27     136.368  57.258  98.009  1.00 38.39           O
    ATOM    239  CB  VAL A  27     133.922  55.626  96.887  1.00 37.61           C
    ATOM    240  CG1 VAL A  27     134.947  55.548  95.756  1.00 37.61           C
    ATOM    241  CG2 VAL A  27     133.956  54.331  97.717  1.00 37.61           C
    ATOM    242  H   VAL A  27     134.761  58.397  96.475  1.00  0.00           H
    ATOM    243  HA  VAL A  27     133.185  56.934  98.428  1.00  0.00           H
    ATOM    244  HB  VAL A  27     132.961  55.653  96.386  1.00  0.00           H
    ATOM    245 HG11 VAL A  27     134.903  54.573  95.280  1.00  0.00           H
    ATOM    246 HG12 VAL A  27     134.731  56.292  94.991  1.00  0.00           H
    ATOM    247 HG13 VAL A  27     135.970  55.686  96.087  1.00  0.00           H
    ATOM    248 HG21 VAL A  27     133.851  53.455  97.084  1.00  0.00           H
    ATOM    249 HG22 VAL A  27     134.881  54.236  98.272  1.00  0.00           H
    ATOM    250 HG23 VAL A  27     133.144  54.279  98.432  1.00  0.00           H
    ATOM    251  N   PHE A  28     135.184  56.616  99.836  1.00 27.63           N
    ATOM    252  CA  PHE A  28     136.281  56.568 100.788  1.00 27.63           C
    ATOM    253  C   PHE A  28     136.298  55.170 101.366  1.00 27.63           C
    ATOM    254  O   PHE A  28     135.267  54.505 101.454  1.00 27.63           O
    ATOM    255  CB  PHE A  28     136.037  57.513 101.978  1.00 48.81           C
    ATOM    256  CG  PHE A  28     135.611  58.902 101.618  1.00 48.81           C
    ATOM    257  CD1 PHE A  28     134.283  59.102 101.201  1.00 48.81           C
    ATOM    258  CD2 PHE A  28     136.520  59.974 101.615  1.00 48.81           C
    ATOM    259  CE1 PHE A  28     133.862  60.358 100.815  1.00 48.81           C
    ATOM    260  CE2 PHE A  28     136.067  61.224 101.227  1.00 48.81           C
    ATOM    261  CZ  PHE A  28     134.751  61.419 100.838  1.00 48.81           C
    ATOM    262  H   PHE A  28     134.277  56.342 100.190  1.00  0.00           H
    ATOM    263  HA  PHE A  28     137.228  56.805 100.307  1.00  0.00           H
    ATOM    264  HB2 PHE A  28     135.251  57.110 102.621  1.00  0.00           H
    ATOM    265  HB3 PHE A  28     136.910  57.587 102.621  1.00  0.00           H
    ATOM    266  HD1 PHE A  28     133.580  58.282 101.184  1.00  0.00           H
    ATOM    267  HD2 PHE A  28     137.551  59.833 101.902  1.00  0.00           H
    ATOM    268  HE1 PHE A  28     132.844  60.509 100.490  1.00  0.00           H
    ATOM    269  HE2 PHE A  28     136.752  62.046 101.157  1.00  0.00           H
    ATOM    270  HZ  PHE A  28     134.427  62.398 100.516  1.00  0.00           H
    ATOM    271  N   LYS A  29     137.474  54.829 101.874  1.00 33.01           N
    ATOM    272  CA  LYS A  29     137.629  53.846 102.910  1.00 33.01           C
    ATOM    273  C   LYS A  29     137.252  54.550 104.214  1.00 33.01           C
    ATOM    274  O   LYS A  29     137.511  55.743 104.381  1.00 33.01           O
    ATOM    275  CB  LYS A  29     139.088  53.386 102.884  1.00 55.73           C
    ATOM    276  CG  LYS A  29     139.344  52.077 103.630  1.00 55.73           C
    ATOM    277  CD  LYS A  29     140.833  51.727 103.621  1.00 55.73           C
    ATOM    278  CE  LYS A  29     141.069  50.234 103.803  1.00 55.73           C
    ATOM    279  NZ  LYS A  29     142.475  49.952 104.113  1.00 55.73           N1+
    ATOM    280  H   LYS A  29     138.252  55.469 101.766  1.00  0.00           H
    ATOM    281  HA  LYS A  29     136.968  52.995 102.730  1.00  0.00           H
    ATOM    282  HB2 LYS A  29     139.359  53.181 101.853  1.00  0.00           H
    ATOM    283  HB3 LYS A  29     139.758  54.183 103.204  1.00  0.00           H
    ATOM    284  HG2 LYS A  29     138.977  52.121 104.655  1.00  0.00           H
    ATOM    285  HG3 LYS A  29     138.774  51.283 103.144  1.00  0.00           H
    ATOM    286  HD2 LYS A  29     141.308  52.062 102.697  1.00  0.00           H
    ATOM    287  HD3 LYS A  29     141.324  52.264 104.430  1.00  0.00           H
    ATOM    288  HE2 LYS A  29     140.442  49.841 104.595  1.00  0.00           H
    ATOM    289  HE3 LYS A  29     140.785  49.710 102.892  1.00  0.00           H
    ATOM    290  HZ1 LYS A  29     142.713  50.327 105.019  1.00  0.00           H
    ATOM    291  HZ2 LYS A  29     142.621  48.948 104.114  1.00  0.00           H
    ATOM    292  HZ3 LYS A  29     143.076  50.360 103.410  1.00  0.00           H
    ATOM    293  N   ALA A  30     136.652  53.789 105.118  1.00 31.06           N
    ATOM    294  CA  ALA A  30     136.239  54.268 106.417  1.00 31.06           C
    ATOM    295  C   ALA A  30     136.286  53.115 107.396  1.00 31.06           C
    ATOM    296  O   ALA A  30     136.346  51.958 106.994  1.00 31.06           O
    ATOM    297  CB  ALA A  30     134.829  54.857 106.318  1.00 30.73           C
    ATOM    298  H   ALA A  30     136.428  52.824 104.911  1.00  0.00           H
    ATOM    299  HA  ALA A  30     136.944  55.019 106.761  1.00  0.00           H
    ATOM    300  HB1 ALA A  30     134.475  55.247 107.271  1.00  0.00           H
    ATOM    301  HB2 ALA A  30     134.792  55.667 105.591  1.00  0.00           H
    ATOM    302  HB3 ALA A  30     134.119  54.094 106.003  1.00  0.00           H
    ATOM    303  N   ARG A  31     136.245  53.474 108.674  1.00 58.12           N
    ATOM    304  CA  ARG A  31     136.216  52.539 109.773  1.00 58.12           C
    ATOM    305  C   ARG A  31     134.834  52.642 110.372  1.00 58.12           C
    ATOM    306  O   ARG A  31     134.510  53.681 110.941  1.00 58.12           O
    ATOM    307  CB  ARG A  31     137.298  52.894 110.792  1.00 48.06           C
    ATOM    308  CG  ARG A  31     138.640  53.104 110.093  1.00 48.06           C
    ATOM    309  CD  ARG A  31     139.844  53.016 110.998  1.00 48.06           C
    ATOM    310  NE  ARG A  31     140.050  51.633 111.414  1.00 48.06           N
    ATOM    311  CZ  ARG A  31     141.107  50.871 111.132  1.00 48.06           C
    ATOM    312  NH1 ARG A  31     142.217  51.339 110.557  1.00 48.06           N
    ATOM    313  NH2 ARG A  31     141.020  49.581 111.431  1.00 48.06           N1+
    ATOM    314  H   ARG A  31     136.201  54.456 108.917  1.00  0.00           H
    ATOM    315  HA  ARG A  31     136.414  51.529 109.429  1.00  0.00           H
    ATOM    316  HB2 ARG A  31     137.058  53.808 111.339  1.00  0.00           H
    ATOM    317  HB3 ARG A  31     137.349  52.106 111.537  1.00  0.00           H
    ATOM    318  HG2 ARG A  31     138.742  52.296 109.368  1.00  0.00           H
    ATOM    319  HG3 ARG A  31     138.662  54.027 109.522  1.00  0.00           H
    ATOM    320  HD2 ARG A  31     140.698  53.462 110.498  1.00  0.00           H
    ATOM    321  HD3 ARG A  31     139.642  53.591 111.898  1.00  0.00           H
    ATOM    322  HE  ARG A  31     139.243  51.180 111.822  1.00  0.00           H
    ATOM    323 HH11 ARG A  31     142.381  52.334 110.455  1.00  0.00           H
    ATOM    324 HH12 ARG A  31     142.965  50.692 110.332  1.00  0.00           H
    ATOM    325 HH21 ARG A  31     140.143  49.199 111.757  1.00  0.00           H
    ATOM    326 HH22 ARG A  31     141.745  48.946 111.098  1.00  0.00           H
    ATOM    327  N   ASP A  32     134.044  51.578 110.221  1.00 43.96           N
    ATOM    328  CA  ASP A  32     132.753  51.442 110.867  1.00 43.96           C
    ATOM    329  C   ASP A  32     132.975  51.343 112.381  1.00 43.96           C
    ATOM    330  O   ASP A  32     133.427  50.298 112.849  1.00 43.96           O
    ATOM    331  CB  ASP A  32     131.996  50.240 110.282  1.00 58.66           C
    ATOM    332  CG  ASP A  32     130.783  49.808 111.085  1.00 58.66           C
    ATOM    333  OD1 ASP A  32     130.962  49.023 112.024  1.00 58.66           O
    ATOM    334  OD2 ASP A  32     129.666  50.195 110.725  1.00 58.66           O1-
    ATOM    335  H   ASP A  32     134.401  50.758 109.748  1.00  0.00           H
    ATOM    336  HA  ASP A  32     132.171  52.339 110.670  1.00  0.00           H
    ATOM    337  HB2 ASP A  32     131.692  50.390 109.251  1.00  0.00           H
    ATOM    338  HB3 ASP A  32     132.663  49.398 110.199  1.00  0.00           H
    ATOM    339  N   LEU A  33     132.682  52.434 113.101  1.00 66.69           N
    ATOM    340  CA  LEU A  33     132.901  52.485 114.542  1.00 66.69           C
    ATOM    341  C   LEU A  33     131.863  51.713 115.367  1.00 66.69           C
    ATOM    342  O   LEU A  33     132.158  51.469 116.536  1.00 66.69           O
    ATOM    343  CB  LEU A  33     133.015  53.942 115.033  1.00 76.19           C
    ATOM    344  CG  LEU A  33     134.259  54.692 114.518  1.00 76.19           C
    ATOM    345  CD1 LEU A  33     134.319  56.116 115.090  1.00 76.19           C
    ATOM    346  CD2 LEU A  33     135.556  53.935 114.823  1.00 76.19           C
    ATOM    347  H   LEU A  33     132.351  53.279 112.657  1.00  0.00           H
    ATOM    348  HA  LEU A  33     133.843  51.977 114.754  1.00  0.00           H
    ATOM    349  HB2 LEU A  33     132.102  54.481 114.789  1.00  0.00           H
    ATOM    350  HB3 LEU A  33     133.063  53.948 116.124  1.00  0.00           H
    ATOM    351  HG  LEU A  33     134.170  54.788 113.435  1.00  0.00           H
    ATOM    352 HD11 LEU A  33     134.801  56.797 114.390  1.00  0.00           H
    ATOM    353 HD12 LEU A  33     133.324  56.513 115.293  1.00  0.00           H
    ATOM    354 HD13 LEU A  33     134.880  56.154 116.024  1.00  0.00           H
    ATOM    355 HD21 LEU A  33     136.388  54.614 115.015  1.00  0.00           H
    ATOM    356 HD22 LEU A  33     135.460  53.288 115.694  1.00  0.00           H
    ATOM    357 HD23 LEU A  33     135.821  53.317 113.965  1.00  0.00           H
    ATOM    358  N   LYS A  34     130.707  51.315 114.794  1.00 99.90           N
    ATOM    359  CA  LYS A  34     129.739  50.485 115.528  1.00 99.90           C
    ATOM    360  C   LYS A  34     130.224  49.038 115.711  1.00 99.90           C
    ATOM    361  O   LYS A  34     130.009  48.479 116.783  1.00 99.90           O
    ATOM    362  CB  LYS A  34     128.317  50.540 114.932  1.00 99.90           C
    ATOM    363  CG  LYS A  34     128.103  49.756 113.628  1.00 99.90           C
    ATOM    364  CD  LYS A  34     126.658  49.718 113.104  1.00 99.90           C
    ATOM    365  CE  LYS A  34     126.512  49.671 111.571  1.00 99.90           C
    ATOM    366  NZ  LYS A  34     127.239  48.549 110.962  1.00 99.90           N1+
    ATOM    367  H   LYS A  34     130.506  51.529 113.828  1.00  0.00           H
    ATOM    368  HA  LYS A  34     129.650  50.910 116.530  1.00  0.00           H
    ATOM    369  HB2 LYS A  34     127.633  50.119 115.671  1.00  0.00           H
    ATOM    370  HB3 LYS A  34     128.024  51.579 114.838  1.00  0.00           H
    ATOM    371  HG2 LYS A  34     128.783  50.148 112.897  1.00  0.00           H
    ATOM    372  HG3 LYS A  34     128.390  48.728 113.789  1.00  0.00           H
    ATOM    373  HD2 LYS A  34     126.165  48.839 113.521  1.00  0.00           H
    ATOM    374  HD3 LYS A  34     126.090  50.560 113.494  1.00  0.00           H
    ATOM    375  HE2 LYS A  34     125.458  49.583 111.309  1.00  0.00           H
    ATOM    376  HE3 LYS A  34     126.865  50.594 111.117  1.00  0.00           H
    ATOM    377  HZ1 LYS A  34     126.943  47.680 111.382  1.00  0.00           H
    ATOM    378  HZ2 LYS A  34     127.041  48.539 109.973  1.00  0.00           H
    ATOM    379  HZ3 LYS A  34     128.231  48.694 111.105  1.00  0.00           H
    ATOM    380  N   ASN A  35     130.879  48.463 114.690  1.00 91.62           N
    ATOM    381  CA  ASN A  35     131.608  47.196 114.787  1.00 91.62           C
    ATOM    382  C   ASN A  35     133.062  47.524 115.150  1.00 91.62           C
    ATOM    383  O   ASN A  35     133.321  48.609 115.670  1.00 91.62           O
    ATOM    384  CB  ASN A  35     131.465  46.357 113.493  1.00 89.92           C
    ATOM    385  CG  ASN A  35     130.091  45.725 113.283  1.00 89.92           C
    ATOM    386  OD1 ASN A  35     129.956  44.509 113.368  1.00 89.92           O
    ATOM    387  ND2 ASN A  35     129.072  46.522 112.979  1.00 89.92           N
    ATOM    388  H   ASN A  35     130.996  48.965 113.815  1.00  0.00           H
    ATOM    389  HA  ASN A  35     131.182  46.611 115.604  1.00  0.00           H
    ATOM    390  HB2 ASN A  35     131.748  46.922 112.620  1.00  0.00           H
    ATOM    391  HB3 ASN A  35     132.155  45.513 113.521  1.00  0.00           H
    ATOM    392 HD21 ASN A  35     129.228  47.518 112.907  1.00  0.00           H
    ATOM    393 HD22 ASN A  35     128.147  46.136 112.865  1.00  0.00           H
    ATOM    394  N   GLY A  36     133.984  46.575 114.928  1.00 64.87           N
    ATOM    395  CA  GLY A  36     135.363  46.595 115.427  1.00 64.87           C
    ATOM    396  C   GLY A  36     136.242  47.788 115.007  1.00 64.87           C
    ATOM    397  O   GLY A  36     137.449  47.734 115.230  1.00 64.87           O
    ATOM    398  H   GLY A  36     133.685  45.712 114.498  1.00  0.00           H
    ATOM    399  HA2 GLY A  36     135.342  46.558 116.517  1.00  0.00           H
    ATOM    400  HA3 GLY A  36     135.851  45.677 115.098  1.00  0.00           H
    ATOM    401  N   GLY A  37     135.699  48.839 114.373  1.00 64.54           N
    ATOM    402  CA  GLY A  37     136.493  49.822 113.657  1.00 64.54           C
    ATOM    403  C   GLY A  37     136.924  49.208 112.324  1.00 64.54           C
    ATOM    404  O   GLY A  37     137.882  49.679 111.716  1.00 64.54           O
    ATOM    405  H   GLY A  37     134.692  48.904 114.286  1.00  0.00           H
    ATOM    406  HA2 GLY A  37     135.929  50.740 113.496  1.00  0.00           H
    ATOM    407  HA3 GLY A  37     137.377  50.062 114.243  1.00  0.00           H
    ATOM    408  N   ARG A  38     136.239  48.123 111.915  1.00 59.73           N
    ATOM    409  CA  ARG A  38     136.412  47.350 110.697  1.00 59.73           C
    ATOM    410  C   ARG A  38     136.320  48.264 109.474  1.00 59.73           C
    ATOM    411  O   ARG A  38     135.647  49.294 109.524  1.00 59.73           O
    ATOM    412  CB  ARG A  38     135.318  46.253 110.650  1.00 99.90           C
    ATOM    413  CG  ARG A  38     133.902  46.822 110.821  1.00 99.90           C
    ATOM    414  CD  ARG A  38     132.758  45.980 110.244  1.00 99.90           C
    ATOM    415  NE  ARG A  38     131.623  46.863 109.933  1.00 99.90           N
    ATOM    416  CZ  ARG A  38     130.355  46.521 109.675  1.00 99.90           C
    ATOM    417  NH1 ARG A  38     129.946  45.251 109.766  1.00 99.90           N
    ATOM    418  NH2 ARG A  38     129.487  47.469 109.309  1.00 99.90           N1+
    ATOM    419  H   ARG A  38     135.482  47.813 112.506  1.00  0.00           H
    ATOM    420  HA  ARG A  38     137.399  46.885 110.727  1.00  0.00           H
    ATOM    421  HB2 ARG A  38     135.394  45.739 109.691  1.00  0.00           H
    ATOM    422  HB3 ARG A  38     135.500  45.491 111.410  1.00  0.00           H
    ATOM    423  HG2 ARG A  38     133.760  47.004 111.885  1.00  0.00           H
    ATOM    424  HG3 ARG A  38     133.850  47.788 110.347  1.00  0.00           H
    ATOM    425  HD2 ARG A  38     133.059  45.350 109.406  1.00  0.00           H
    ATOM    426  HD3 ARG A  38     132.403  45.311 111.027  1.00  0.00           H
    ATOM    427  HE  ARG A  38     131.864  47.839 109.845  1.00  0.00           H
    ATOM    428 HH11 ARG A  38     130.597  44.533 110.051  1.00  0.00           H
    ATOM    429 HH12 ARG A  38     128.998  44.993 109.536  1.00  0.00           H
    ATOM    430 HH21 ARG A  38     129.771  48.441 109.293  1.00  0.00           H
    ATOM    431 HH22 ARG A  38     128.569  47.210 108.980  1.00  0.00           H
    ATOM    432  N   PHE A  39     136.982  47.862 108.389  1.00 65.10           N
    ATOM    433  CA  PHE A  39     136.957  48.647 107.170  1.00 65.10           C
    ATOM    434  C   PHE A  39     135.665  48.507 106.383  1.00 65.10           C
    ATOM    435  O   PHE A  39     135.105  47.416 106.293  1.00 65.10           O
    ATOM    436  CB  PHE A  39     138.188  48.357 106.317  1.00 33.62           C
    ATOM    437  CG  PHE A  39     139.449  48.991 106.860  1.00 33.62           C
    ATOM    438  CD1 PHE A  39     139.467  50.343 107.275  1.00 33.62           C
    ATOM    439  CD2 PHE A  39     140.665  48.287 106.779  1.00 33.62           C
    ATOM    440  CE1 PHE A  39     140.652  50.941 107.657  1.00 33.62           C
    ATOM    441  CE2 PHE A  39     141.846  48.918 107.133  1.00 33.62           C
    ATOM    442  CZ  PHE A  39     141.839  50.232 107.579  1.00 33.62           C
    ATOM    443  H   PHE A  39     137.509  47.001 108.378  1.00  0.00           H
    ATOM    444  HA  PHE A  39     136.985  49.684 107.477  1.00  0.00           H
    ATOM    445  HB2 PHE A  39     138.327  47.277 106.238  1.00  0.00           H
    ATOM    446  HB3 PHE A  39     138.052  48.720 105.297  1.00  0.00           H
    ATOM    447  HD1 PHE A  39     138.582  50.953 107.283  1.00  0.00           H
    ATOM    448  HD2 PHE A  39     140.690  47.267 106.420  1.00  0.00           H
    ATOM    449  HE1 PHE A  39     140.660  51.967 107.990  1.00  0.00           H
    ATOM    450  HE2 PHE A  39     142.780  48.382 107.060  1.00  0.00           H
    ATOM    451  HZ  PHE A  39     142.767  50.706 107.857  1.00  0.00           H
    ATOM    452  N   VAL A  40     135.230  49.642 105.834  1.00 41.75           N
    ATOM    453  CA  VAL A  40     133.967  49.767 105.149  1.00 41.75           C
    ATOM    454  C   VAL A  40     134.076  50.877 104.090  1.00 41.75           C
    ATOM    455  O   VAL A  40     134.665  51.924 104.348  1.00 41.75           O
    ATOM    456  CB  VAL A  40     132.841  50.048 106.180  1.00 43.90           C
    ATOM    457  CG1 VAL A  40     132.846  51.456 106.805  1.00 43.90           C
    ATOM    458  CG2 VAL A  40     131.477  49.734 105.591  1.00 43.90           C
    ATOM    459  H   VAL A  40     135.745  50.502 105.975  1.00  0.00           H
    ATOM    460  HA  VAL A  40     133.748  48.831 104.630  1.00  0.00           H
    ATOM    461  HB  VAL A  40     132.970  49.350 107.009  1.00  0.00           H
    ATOM    462 HG11 VAL A  40     132.017  51.583 107.493  1.00  0.00           H
    ATOM    463 HG12 VAL A  40     133.763  51.644 107.359  1.00  0.00           H
    ATOM    464 HG13 VAL A  40     132.724  52.239 106.059  1.00  0.00           H
    ATOM    465 HG21 VAL A  40     130.708  49.849 106.352  1.00  0.00           H
    ATOM    466 HG22 VAL A  40     131.256  50.376 104.743  1.00  0.00           H
    ATOM    467 HG23 VAL A  40     131.441  48.699 105.253  1.00  0.00           H
    ATOM    468  N   ALA A  41     133.524  50.617 102.902  1.00 35.03           N
    ATOM    469  CA  ALA A  41     133.501  51.557 101.790  1.00 35.03           C
    ATOM    470  C   ALA A  41     132.284  52.480 101.925  1.00 35.03           C
    ATOM    471  O   ALA A  41     131.165  51.985 101.890  1.00 35.03           O
    ATOM    472  CB  ALA A  41     133.466  50.762 100.475  1.00 62.28           C
    ATOM    473  H   ALA A  41     133.014  49.753 102.761  1.00  0.00           H
    ATOM    474  HA  ALA A  41     134.415  52.151 101.793  1.00  0.00           H
    ATOM    475  HB1 ALA A  41     133.349  51.423  99.615  1.00  0.00           H
    ATOM    476  HB2 ALA A  41     134.394  50.206 100.336  1.00  0.00           H
    ATOM    477  HB3 ALA A  41     132.652  50.039 100.461  1.00  0.00           H
    ATOM    478  N   LEU A  42     132.499  53.795 102.063  1.00 44.84           N
    ATOM    479  CA  LEU A  42     131.409  54.775 102.053  1.00 44.84           C
    ATOM    480  C   LEU A  42     131.292  55.364 100.654  1.00 44.84           C
    ATOM    481  O   LEU A  42     132.323  55.775 100.147  1.00 44.84           O
    ATOM    482  CB  LEU A  42     131.696  55.962 103.000  1.00 61.74           C
    ATOM    483  CG  LEU A  42     131.777  55.675 104.506  1.00 61.74           C
    ATOM    484  CD1 LEU A  42     131.930  57.003 105.282  1.00 61.74           C
    ATOM    485  CD2 LEU A  42     130.608  54.836 105.038  1.00 61.74           C
    ATOM    486  H   LEU A  42     133.443  54.159 102.066  1.00  0.00           H
    ATOM    487  HA  LEU A  42     130.474  54.302 102.326  1.00  0.00           H
    ATOM    488  HB2 LEU A  42     132.622  56.457 102.700  1.00  0.00           H
    ATOM    489  HB3 LEU A  42     130.908  56.704 102.854  1.00  0.00           H
    ATOM    490  HG  LEU A  42     132.682  55.091 104.651  1.00  0.00           H
    ATOM    491 HD11 LEU A  42     132.795  56.969 105.945  1.00  0.00           H
    ATOM    492 HD12 LEU A  42     132.074  57.854 104.616  1.00  0.00           H
    ATOM    493 HD13 LEU A  42     131.065  57.238 105.900  1.00  0.00           H
    ATOM    494 HD21 LEU A  42     130.185  55.245 105.952  1.00  0.00           H
    ATOM    495 HD22 LEU A  42     129.791  54.773 104.324  1.00  0.00           H
    ATOM    496 HD23 LEU A  42     130.941  53.825 105.265  1.00  0.00           H
    ATOM    497  N   LYS A  43     130.073  55.499 100.114  1.00 52.70           N
    ATOM    498  CA  LYS A  43     129.794  56.324  98.940  1.00 52.70           C
    ATOM    499  C   LYS A  43     128.802  57.407  99.361  1.00 52.70           C
    ATOM    500  O   LYS A  43     127.618  57.119  99.539  1.00 52.70           O
    ATOM    501  CB  LYS A  43     129.254  55.496  97.757  1.00 99.62           C
    ATOM    502  CG  LYS A  43     130.318  54.603  97.090  1.00 99.62           C
    ATOM    503  CD  LYS A  43     129.812  53.464  96.187  1.00 99.62           C
    ATOM    504  CE  LYS A  43     128.578  53.815  95.346  1.00 99.62           C
    ATOM    505  NZ  LYS A  43     128.269  52.763  94.364  1.00 99.62           N1+
    ATOM    506  H   LYS A  43     129.266  55.148 100.617  1.00  0.00           H
    ATOM    507  HA  LYS A  43     130.705  56.808  98.600  1.00  0.00           H
    ATOM    508  HB2 LYS A  43     128.372  54.956  98.062  1.00  0.00           H
    ATOM    509  HB3 LYS A  43     128.893  56.183  96.989  1.00  0.00           H
    ATOM    510  HG2 LYS A  43     130.923  55.260  96.473  1.00  0.00           H
    ATOM    511  HG3 LYS A  43     130.978  54.177  97.843  1.00  0.00           H
    ATOM    512  HD2 LYS A  43     130.625  53.181  95.516  1.00  0.00           H
    ATOM    513  HD3 LYS A  43     129.616  52.577  96.789  1.00  0.00           H
    ATOM    514  HE2 LYS A  43     127.706  53.945  95.988  1.00  0.00           H
    ATOM    515  HE3 LYS A  43     128.720  54.755  94.826  1.00  0.00           H
    ATOM    516  HZ1 LYS A  43     129.054  52.653  93.737  1.00  0.00           H
    ATOM    517  HZ2 LYS A  43     128.097  51.891  94.843  1.00  0.00           H
    ATOM    518  HZ3 LYS A  43     127.451  53.025  93.832  1.00  0.00           H
    ATOM    519  N   ARG A  44     129.315  58.636  99.521  1.00 68.25           N
    ATOM    520  CA  ARG A  44     128.519  59.824  99.820  1.00 68.25           C
    ATOM    521  C   ARG A  44     127.760  60.246  98.557  1.00 68.25           C
    ATOM    522  O   ARG A  44     128.409  60.653  97.592  1.00 68.25           O
    ATOM    523  CB  ARG A  44     129.445  60.966 100.301  1.00 73.70           C
    ATOM    524  CG  ARG A  44     129.051  61.538 101.667  1.00 73.70           C
    ATOM    525  CD  ARG A  44     129.930  61.037 102.813  1.00 73.70           C
    ATOM    526  NE  ARG A  44     131.252  61.671 102.768  1.00 73.70           N
    ATOM    527  CZ  ARG A  44     132.102  61.772 103.800  1.00 73.70           C
    ATOM    528  NH1 ARG A  44     131.801  61.255 104.999  1.00 73.70           N
    ATOM    529  NH2 ARG A  44     133.267  62.407 103.629  1.00 73.70           N1+
    ATOM    530  H   ARG A  44     130.295  58.790  99.332  1.00  0.00           H
    ATOM    531  HA  ARG A  44     127.804  59.585 100.603  1.00  0.00           H
    ATOM    532  HB2 ARG A  44     130.492  60.663 100.297  1.00  0.00           H
    ATOM    533  HB3 ARG A  44     129.415  61.798  99.595  1.00  0.00           H
    ATOM    534  HG2 ARG A  44     129.265  62.606 101.593  1.00  0.00           H
    ATOM    535  HG3 ARG A  44     127.985  61.508 101.881  1.00  0.00           H
    ATOM    536  HD2 ARG A  44     129.432  61.197 103.761  1.00  0.00           H
    ATOM    537  HD3 ARG A  44     130.114  59.966 102.714  1.00  0.00           H
    ATOM    538  HE  ARG A  44     131.540  62.029 101.869  1.00  0.00           H
    ATOM    539 HH11 ARG A  44     130.919  60.784 105.142  1.00  0.00           H
    ATOM    540 HH12 ARG A  44     132.455  61.321 105.766  1.00  0.00           H
    ATOM    541 HH21 ARG A  44     133.517  62.788 102.728  1.00  0.00           H
    ATOM    542 HH22 ARG A  44     133.923  62.487 104.392  1.00  0.00           H
    ATOM    543  N   VAL A  77     129.387  42.901  96.835  1.00 48.66           N1+
    ATOM    544  CA  VAL A  77     129.750  43.701  97.992  1.00 48.66           C
    ATOM    545  C   VAL A  77     128.454  44.272  98.619  1.00 48.66           C
    ATOM    546  O   VAL A  77     127.737  45.041  97.978  1.00 48.66           O
    ATOM    547  CB  VAL A  77     130.774  44.801  97.564  1.00 22.07           C
    ATOM    548  CG1 VAL A  77     130.190  46.067  96.903  1.00 22.07           C
    ATOM    549  CG2 VAL A  77     131.700  45.213  98.697  1.00 22.07           C
    ATOM    550  H   VAL A  77     129.717  43.249  95.944  1.00  0.00           H
    ATOM    551  HA  VAL A  77     130.232  43.033  98.709  1.00  0.00           H
    ATOM    552  HB  VAL A  77     131.440  44.338  96.839  1.00  0.00           H
    ATOM    553 HG11 VAL A  77     130.980  46.708  96.516  1.00  0.00           H
    ATOM    554 HG12 VAL A  77     129.538  45.816  96.066  1.00  0.00           H
    ATOM    555 HG13 VAL A  77     129.626  46.679  97.607  1.00  0.00           H
    ATOM    556 HG21 VAL A  77     132.456  45.922  98.359  1.00  0.00           H
    ATOM    557 HG22 VAL A  77     131.107  45.700  99.466  1.00  0.00           H
    ATOM    558 HG23 VAL A  77     132.220  44.357  99.126  1.00  0.00           H
    ATOM    559  N   ARG A  78     128.105  43.773  99.816  1.00 52.51           N
    ATOM    560  CA  ARG A  78     126.897  44.132 100.561  1.00 52.51           C
    ATOM    561  C   ARG A  78     126.815  45.631 100.865  1.00 52.51           C
    ATOM    562  O   ARG A  78     127.850  46.268 101.013  1.00 52.51           O
    ATOM    563  CB  ARG A  78     126.874  43.357 101.892  1.00 99.90           C
    ATOM    564  CG  ARG A  78     126.002  42.094 101.852  1.00 99.90           C
    ATOM    565  CD  ARG A  78     125.744  41.538 103.258  1.00 99.90           C
    ATOM    566  NE  ARG A  78     125.080  42.543 104.109  1.00 99.90           N
    ATOM    567  CZ  ARG A  78     123.765  42.782 104.226  1.00 99.90           C
    ATOM    568  NH1 ARG A  78     122.857  41.987 103.646  1.00 99.90           N
    ATOM    569  NH2 ARG A  78     123.360  43.840 104.941  1.00 99.90           N1+
    ATOM    570  H   ARG A  78     128.703  43.084 100.260  1.00  0.00           H
    ATOM    571  HA  ARG A  78     126.040  43.855  99.943  1.00  0.00           H
    ATOM    572  HB2 ARG A  78     127.874  43.138 102.255  1.00  0.00           H
    ATOM    573  HB3 ARG A  78     126.465  44.012 102.661  1.00  0.00           H
    ATOM    574  HG2 ARG A  78     125.035  42.403 101.454  1.00  0.00           H
    ATOM    575  HG3 ARG A  78     126.380  41.329 101.172  1.00  0.00           H
    ATOM    576  HD2 ARG A  78     125.261  40.561 103.256  1.00  0.00           H
    ATOM    577  HD3 ARG A  78     126.717  41.390 103.727  1.00  0.00           H
    ATOM    578  HE  ARG A  78     125.712  43.181 104.569  1.00  0.00           H
    ATOM    579 HH11 ARG A  78     123.162  41.192 103.104  1.00  0.00           H
    ATOM    580 HH12 ARG A  78     121.869  42.174 103.739  1.00  0.00           H
    ATOM    581 HH21 ARG A  78     124.031  44.465 105.360  1.00  0.00           H
    ATOM    582 HH22 ARG A  78     122.374  44.033 105.054  1.00  0.00           H
    ATOM    583  N   PHE A  80     125.896  47.640 103.798  1.00 82.07           N1+
    ATOM    584  CA  PHE A  80     125.710  47.753 105.245  1.00 82.07           C
    ATOM    585  C   PHE A  80     124.738  48.812 105.763  1.00 82.07           C
    ATOM    586  O   PHE A  80     124.157  48.556 106.817  1.00 82.07           O
    ATOM    587  CB  PHE A  80     127.055  47.889 105.995  1.00 99.90           C
    ATOM    588  CG  PHE A  80     127.892  46.636 106.101  1.00 99.90           C
    ATOM    589  CD1 PHE A  80     127.299  45.355 106.165  1.00 99.90           C
    ATOM    590  CD2 PHE A  80     129.275  46.754 106.339  1.00 99.90           C
    ATOM    591  CE1 PHE A  80     128.082  44.235 106.372  1.00 99.90           C
    ATOM    592  CE2 PHE A  80     130.042  45.621 106.567  1.00 99.90           C
    ATOM    593  CZ  PHE A  80     129.447  44.367 106.573  1.00 99.90           C
    ATOM    594  H   PHE A  80     126.844  47.754 103.468  1.00  0.00           H
    ATOM    595  HA  PHE A  80     125.205  46.837 105.537  1.00  0.00           H
    ATOM    596  HB2 PHE A  80     127.669  48.661 105.537  1.00  0.00           H
    ATOM    597  HB3 PHE A  80     126.887  48.226 107.019  1.00  0.00           H
    ATOM    598  HD1 PHE A  80     126.241  45.194 106.061  1.00  0.00           H
    ATOM    599  HD2 PHE A  80     129.738  47.726 106.357  1.00  0.00           H
    ATOM    600  HE1 PHE A  80     127.629  43.255 106.398  1.00  0.00           H
    ATOM    601  HE2 PHE A  80     131.104  45.716 106.743  1.00  0.00           H
    ATOM    602  HZ  PHE A  80     130.052  43.489 106.745  1.00  0.00           H
    ATOM    603  N   VAL A  85     122.239  62.026 105.455  1.00 33.54           N1+
    ATOM    604  CA  VAL A  85     123.000  63.167 105.952  1.00 33.54           C
    ATOM    605  C   VAL A  85     122.169  64.464 106.052  1.00 33.54           C
    ATOM    606  O   VAL A  85     122.385  65.195 107.017  1.00 33.54           O
    ATOM    607  CB  VAL A  85     124.273  63.413 105.096  1.00 99.63           C
    ATOM    608  CG1 VAL A  85     125.044  64.711 105.411  1.00 99.63           C
    ATOM    609  CG2 VAL A  85     125.220  62.208 105.118  1.00 99.63           C
    ATOM    610  H   VAL A  85     122.340  61.827 104.469  1.00  0.00           H
    ATOM    611  HA  VAL A  85     123.339  62.927 106.962  1.00  0.00           H
    ATOM    612  HB  VAL A  85     123.975  63.469 104.069  1.00  0.00           H
    ATOM    613 HG11 VAL A  85     125.975  64.760 104.847  1.00  0.00           H
    ATOM    614 HG12 VAL A  85     124.477  65.605 105.149  1.00  0.00           H
    ATOM    615 HG13 VAL A  85     125.298  64.771 106.470  1.00  0.00           H
    ATOM    616 HG21 VAL A  85     126.081  62.374 104.471  1.00  0.00           H
    ATOM    617 HG22 VAL A  85     125.589  62.017 106.126  1.00  0.00           H
    ATOM    618 HG23 VAL A  85     124.728  61.299 104.772  1.00  0.00           H
    ATOM    619  N   LEU A  94     123.226  61.692 101.355  1.00 56.41           N1+
    ATOM    620  CA  LEU A  94     123.212  60.372 101.963  1.00 56.41           C
    ATOM    621  C   LEU A  94     124.668  59.926 102.110  1.00 56.41           C
    ATOM    622  O   LEU A  94     125.592  60.673 101.791  1.00 56.41           O
    ATOM    623  CB  LEU A  94     122.448  59.351 101.093  1.00 26.80           C
    ATOM    624  CG  LEU A  94     121.020  59.714 100.659  1.00 26.80           C
    ATOM    625  CD1 LEU A  94     120.406  58.601  99.790  1.00 26.80           C
    ATOM    626  CD2 LEU A  94     120.123  59.992 101.862  1.00 26.80           C
    ATOM    627  H   LEU A  94     123.790  61.779 100.516  1.00  0.00           H
    ATOM    628  HA  LEU A  94     122.768  60.421 102.954  1.00  0.00           H
    ATOM    629  HB2 LEU A  94     123.041  59.168 100.216  1.00  0.00           H
    ATOM    630  HB3 LEU A  94     122.402  58.396 101.619  1.00  0.00           H
    ATOM    631  HG  LEU A  94     121.068  60.615 100.049  1.00  0.00           H
    ATOM    632 HD11 LEU A  94     119.829  59.028  98.969  1.00  0.00           H
    ATOM    633 HD12 LEU A  94     121.168  57.958  99.349  1.00  0.00           H
    ATOM    634 HD13 LEU A  94     119.736  57.959 100.364  1.00  0.00           H
    ATOM    635 HD21 LEU A  94     119.084  60.117 101.558  1.00  0.00           H
    ATOM    636 HD22 LEU A  94     120.158  59.164 102.570  1.00  0.00           H
    ATOM    637 HD23 LEU A  94     120.416  60.895 102.383  1.00  0.00           H
    ATOM    638  N   THR A  95     124.848  58.700 102.595  1.00 31.40           N
    ATOM    639  CA  THR A  95     126.129  58.056 102.763  1.00 31.40           C
    ATOM    640  C   THR A  95     125.826  56.573 102.814  1.00 31.40           C
    ATOM    641  O   THR A  95     125.529  56.017 103.870  1.00 31.40           O
    ATOM    642  CB  THR A  95     126.894  58.488 104.035  1.00 22.25           C
    ATOM    643  OG1 THR A  95     127.254  59.839 103.918  1.00 22.25           O
    ATOM    644  CG2 THR A  95     128.178  57.675 104.283  1.00 22.25           C
    ATOM    645  H   THR A  95     124.045  58.163 102.891  1.00  0.00           H
    ATOM    646  HA  THR A  95     126.751  58.244 101.890  1.00  0.00           H
    ATOM    647  HB  THR A  95     126.257  58.442 104.909  1.00  0.00           H
    ATOM    648 HG21 THR A  95     128.946  58.251 104.794  1.00  0.00           H
    ATOM    649 HG22 THR A  95     127.979  56.784 104.877  1.00  0.00           H
    ATOM    650 HG23 THR A  95     128.625  57.354 103.342  1.00  0.00           H
    ATOM    651  HG1 THR A  95     126.468  60.335 103.667  1.00  0.00           H
    ATOM    652  N   VAL A  97     126.938  52.778 102.651  1.00 43.71           N1+
    ATOM    653  CA  VAL A  97     128.010  52.203 103.428  1.00 43.71           C
    ATOM    654  C   VAL A  97     128.079  50.730 103.058  1.00 43.71           C
    ATOM    655  O   VAL A  97     127.258  49.959 103.524  1.00 43.71           O
    ATOM    656  CB  VAL A  97     127.728  52.331 104.941  1.00 30.88           C
    ATOM    657  CG1 VAL A  97     128.895  51.737 105.722  1.00 30.88           C
    ATOM    658  CG2 VAL A  97     127.427  53.776 105.378  1.00 30.88           C
    ATOM    659  H   VAL A  97     126.021  52.366 102.774  1.00  0.00           H
    ATOM    660  HA  VAL A  97     128.973  52.664 103.203  1.00  0.00           H
    ATOM    661  HB  VAL A  97     126.837  51.759 105.198  1.00  0.00           H
    ATOM    662 HG11 VAL A  97     129.073  52.265 106.637  1.00  0.00           H
    ATOM    663 HG12 VAL A  97     128.728  50.686 105.947  1.00  0.00           H
    ATOM    664 HG13 VAL A  97     129.818  51.850 105.164  1.00  0.00           H
    ATOM    665 HG21 VAL A  97     126.458  53.847 105.875  1.00  0.00           H
    ATOM    666 HG22 VAL A  97     128.177  54.160 106.056  1.00  0.00           H
    ATOM    667 HG23 VAL A  97     127.418  54.484 104.559  1.00  0.00           H
    ATOM    668  N   PHE A  98     129.018  50.379 102.186  1.00 38.75           N
    ATOM    669  CA  PHE A  98     129.165  49.050 101.620  1.00 38.75           C
    ATOM    670  C   PHE A  98     130.271  48.309 102.342  1.00 38.75           C
    ATOM    671  O   PHE A  98     131.264  48.930 102.708  1.00 38.75           O
    ATOM    672  CB  PHE A  98     129.533  49.149 100.133  1.00 36.51           C
    ATOM    673  CG  PHE A  98     128.567  49.972  99.326  1.00 36.51           C
    ATOM    674  CD1 PHE A  98     128.676  51.372  99.359  1.00 36.51           C
    ATOM    675  CD2 PHE A  98     127.466  49.375  98.680  1.00 36.51           C
    ATOM    676  CE1 PHE A  98     127.734  52.134  98.710  1.00 36.51           C
    ATOM    677  CE2 PHE A  98     126.536  50.172  98.025  1.00 36.51           C
    ATOM    678  CZ  PHE A  98     126.672  51.552  98.037  1.00 36.51           C
    ATOM    679  H   PHE A  98     129.700  51.066 101.884  1.00  0.00           H
    ATOM    680  HA  PHE A  98     128.234  48.513 101.737  1.00  0.00           H
    ATOM    681  HB2 PHE A  98     130.521  49.591 100.015  1.00  0.00           H
    ATOM    682  HB3 PHE A  98     129.596  48.151  99.695  1.00  0.00           H
    ATOM    683  HD1 PHE A  98     129.493  51.855  99.873  1.00  0.00           H
    ATOM    684  HD2 PHE A  98     127.343  48.302  98.693  1.00  0.00           H
    ATOM    685  HE1 PHE A  98     127.870  53.186  98.718  1.00  0.00           H
    ATOM    686  HE2 PHE A  98     125.699  49.716  97.518  1.00  0.00           H
    ATOM    687  HZ  PHE A  98     125.948  52.167  97.526  1.00  0.00           H
    ATOM    688  N   GLU A  99     130.126  46.979 102.450  1.00 51.62           N
    ATOM    689  CA  GLU A  99     131.149  46.096 103.004  1.00 51.62           C
    ATOM    690  C   GLU A  99     132.479  46.268 102.266  1.00 51.62           C
    ATOM    691  O   GLU A  99     132.504  46.738 101.126  1.00 51.62           O
    ATOM    692  CB  GLU A  99     130.671  44.625 103.004  1.00 83.55           C
    ATOM    693  CG  GLU A  99     130.907  43.839 101.698  1.00 83.55           C
    ATOM    694  CD  GLU A  99     130.658  42.348 101.851  1.00 83.55           C
    ATOM    695  OE1 GLU A  99     131.474  41.713 102.550  1.00 83.55           O
    ATOM    696  OE2 GLU A  99     129.695  41.849 101.231  1.00 83.55           O1-
    ATOM    697  H   GLU A  99     129.303  46.534 102.063  1.00  0.00           H
    ATOM    698  HA  GLU A  99     131.294  46.407 104.040  1.00  0.00           H
    ATOM    699  HB2 GLU A  99     131.171  44.111 103.827  1.00  0.00           H
    ATOM    700  HB3 GLU A  99     129.609  44.590 103.252  1.00  0.00           H
    ATOM    701  HG2 GLU A  99     130.295  44.258 100.912  1.00  0.00           H
    ATOM    702  HG3 GLU A  99     131.941  43.904 101.365  1.00  0.00           H
    ATOM    703  N   HIE A 100     133.565  45.882 102.930  1.00 65.42           N
    ATOM    704  CA  HIE A 100     134.853  46.032 102.301  1.00 65.42           C
    ATOM    705  C   HIE A 100     135.733  44.822 102.510  1.00 65.42           C
    ATOM    706  O   HIE A 100     135.949  44.366 103.630  1.00 65.42           O
    ATOM    707  CB  HIE A 100     135.522  47.333 102.729  1.00 65.13           C
    ATOM    708  CG  HIE A 100     136.685  47.630 101.834  1.00 65.13           C
    ATOM    709  ND1 HIE A 100     136.519  47.896 100.471  1.00 65.13           N
    ATOM    710  CD2 HIE A 100     138.028  47.591 102.130  1.00 65.13           C
    ATOM    711  CE1 HIE A 100     137.752  48.000  99.998  1.00 65.13           C
    ATOM    712  NE2 HIE A 100     138.683  47.836 100.939  1.00 65.13           N
    ATOM    713  H   HIE A 100     133.529  45.488 103.859  1.00  0.00           H
    ATOM    714  HA  HIE A 100     134.720  46.102 101.219  1.00  0.00           H
    ATOM    715  HB2 HIE A 100     134.826  48.166 102.651  1.00  0.00           H
    ATOM    716  HB3 HIE A 100     135.861  47.281 103.762  1.00  0.00           H
    ATOM    717  HD2 HIE A 100     138.554  47.391 103.051  1.00  0.00           H
    ATOM    718  HE2 HIE A 100     139.687  47.865 100.808  1.00  0.00           H
    ATOM    719  HE1 HIE A 100     137.981  48.198  98.962  1.00  0.00           H
    ATOM    720  N   VAL A 101     136.260  44.394 101.369  1.00 35.97           N
    ATOM    721  CA  VAL A 101     137.235  43.353 101.241  1.00 35.97           C
    ATOM    722  C   VAL A 101     138.496  44.058 100.753  1.00 35.97           C
    ATOM    723  O   VAL A 101     138.525  44.548  99.621  1.00 35.97           O
    ATOM    724  CB  VAL A 101     136.771  42.376 100.155  1.00 28.55           C
    ATOM    725  CG1 VAL A 101     137.816  41.301  99.875  1.00 28.55           C
    ATOM    726  CG2 VAL A 101     135.405  41.761 100.471  1.00 28.55           C
    ATOM    727  H   VAL A 101     136.026  44.872 100.511  1.00  0.00           H
    ATOM    728  HA  VAL A 101     137.416  42.829 102.183  1.00  0.00           H
    ATOM    729  HB  VAL A 101     136.620  42.938  99.244  1.00  0.00           H
    ATOM    730 HG11 VAL A 101     137.407  40.517  99.240  1.00  0.00           H
    ATOM    731 HG12 VAL A 101     138.700  41.694  99.370  1.00  0.00           H
    ATOM    732 HG13 VAL A 101     138.133  40.876 100.824  1.00  0.00           H
    ATOM    733 HG21 VAL A 101     135.175  41.020  99.713  1.00  0.00           H
    ATOM    734 HG22 VAL A 101     135.393  41.279 101.448  1.00  0.00           H
    ATOM    735 HG23 VAL A 101     134.605  42.503 100.453  1.00  0.00           H
    ATOM    736  N   ASP A 102     139.493  44.151 101.635  1.00 37.97           N
    ATOM    737  CA  ASP A 102     140.666  44.971 101.384  1.00 37.97           C
    ATOM    738  C   ASP A 102     141.682  44.217 100.513  1.00 37.97           C
    ATOM    739  O   ASP A 102     142.729  43.878 101.009  1.00 37.97           O
    ATOM    740  CB  ASP A 102     141.247  45.481 102.730  1.00 62.66           C
    ATOM    741  CG  ASP A 102     142.025  46.785 102.610  1.00 62.66           C
    ATOM    742  OD1 ASP A 102     141.690  47.595 101.719  1.00 62.66           O
    ATOM    743  OD2 ASP A 102     142.886  47.012 103.485  1.00 62.66           O1-
    ATOM    744  H   ASP A 102     139.411  43.738 102.554  1.00  0.00           H
    ATOM    745  HA  ASP A 102     140.353  45.823 100.783  1.00  0.00           H
    ATOM    746  HB2 ASP A 102     140.444  45.645 103.450  1.00  0.00           H
    ATOM    747  HB3 ASP A 102     141.919  44.748 103.172  1.00  0.00           H
    ATOM    748  N   GLN A 103     141.383  43.924  99.242  1.00 23.39           N
    ATOM    749  CA  GLN A 103     142.240  43.157  98.325  1.00 23.39           C
    ATOM    750  C   GLN A 103     141.465  43.016  97.021  1.00 23.39           C
    ATOM    751  O   GLN A 103     140.440  42.336  97.001  1.00 23.39           O
    ATOM    752  CB  GLN A 103     142.730  41.822  98.959  1.00 45.18           C
    ATOM    753  CG  GLN A 103     142.713  40.512  98.148  1.00 45.18           C
    ATOM    754  CD  GLN A 103     144.015  39.996  97.596  1.00 45.18           C
    ATOM    755  OE1 GLN A 103     144.282  38.802  97.694  1.00 45.18           O
    ATOM    756  NE2 GLN A 103     144.813  40.854  96.984  1.00 45.18           N
    ATOM    757  H   GLN A 103     140.483  44.204  98.875  1.00  0.00           H
    ATOM    758  HA  GLN A 103     143.149  43.713  98.120  1.00  0.00           H
    ATOM    759  HB2 GLN A 103     143.695  41.959  99.451  1.00  0.00           H
    ATOM    760  HB3 GLN A 103     142.031  41.610  99.762  1.00  0.00           H
    ATOM    761  HG2 GLN A 103     142.412  39.730  98.845  1.00  0.00           H
    ATOM    762  HG3 GLN A 103     142.014  40.495  97.322  1.00  0.00           H
    ATOM    763 HE21 GLN A 103     144.570  41.832  96.891  1.00  0.00           H
    ATOM    764 HE22 GLN A 103     145.691  40.538  96.599  1.00  0.00           H
    ATOM    765  N   ASP A 104     141.959  43.660  95.957  1.00 32.26           N
    ATOM    766  CA  ASP A 104     141.404  43.505  94.615  1.00 32.26           C
    ATOM    767  C   ASP A 104     141.974  42.244  93.946  1.00 32.26           C
    ATOM    768  O   ASP A 104     143.032  41.760  94.352  1.00 32.26           O
    ATOM    769  CB  ASP A 104     141.581  44.752  93.711  1.00 70.79           C
    ATOM    770  CG  ASP A 104     142.960  45.424  93.687  1.00 70.79           C
    ATOM    771  OD1 ASP A 104     143.915  44.860  94.257  1.00 70.79           O
    ATOM    772  OD2 ASP A 104     143.063  46.485  93.036  1.00 70.79           O1-
    ATOM    773  H   ASP A 104     142.801  44.215  96.032  1.00  0.00           H
    ATOM    774  HA  ASP A 104     140.329  43.360  94.708  1.00  0.00           H
    ATOM    775  HB2 ASP A 104     141.327  44.473  92.690  1.00  0.00           H
    ATOM    776  HB3 ASP A 104     140.856  45.501  94.033  1.00  0.00           H
    ATOM    777  N   LEU A 105     141.295  41.771  92.887  1.00 49.17           N
    ATOM    778  CA  LEU A 105     141.802  40.715  92.008  1.00 49.17           C
    ATOM    779  C   LEU A 105     143.140  41.092  91.350  1.00 49.17           C
    ATOM    780  O   LEU A 105     143.931  40.201  91.059  1.00 49.17           O
    ATOM    781  CB  LEU A 105     140.741  40.324  90.960  1.00 24.51           C
    ATOM    782  CG  LEU A 105     141.126  39.200  89.963  1.00 24.51           C
    ATOM    783  CD1 LEU A 105     141.385  37.832  90.622  1.00 24.51           C
    ATOM    784  CD2 LEU A 105     140.097  39.083  88.836  1.00 24.51           C
    ATOM    785  H   LEU A 105     140.420  42.204  92.618  1.00  0.00           H
    ATOM    786  HA  LEU A 105     141.961  39.860  92.653  1.00  0.00           H
    ATOM    787  HB2 LEU A 105     139.823  40.035  91.472  1.00  0.00           H
    ATOM    788  HB3 LEU A 105     140.495  41.221  90.390  1.00  0.00           H
    ATOM    789  HG  LEU A 105     142.046  39.503  89.469  1.00  0.00           H
    ATOM    790 HD11 LEU A 105     141.993  37.199  89.977  1.00  0.00           H
    ATOM    791 HD12 LEU A 105     141.924  37.925  91.560  1.00  0.00           H
    ATOM    792 HD13 LEU A 105     140.458  37.297  90.828  1.00  0.00           H
    ATOM    793 HD21 LEU A 105     140.617  38.908  87.896  1.00  0.00           H
    ATOM    794 HD22 LEU A 105     139.408  38.252  88.987  1.00  0.00           H
    ATOM    795 HD23 LEU A 105     139.496  39.986  88.720  1.00  0.00           H
    ATOM    796  N   THR A 106     143.396  42.398  91.170  1.00 41.60           N
    ATOM    797  CA  THR A 106     144.684  42.898  90.709  1.00 41.60           C
    ATOM    798  C   THR A 106     145.811  42.537  91.690  1.00 41.60           C
    ATOM    799  O   THR A 106     146.712  41.813  91.272  1.00 41.60           O
    ATOM    800  CB  THR A 106     144.677  44.423  90.464  1.00 56.29           C
    ATOM    801  OG1 THR A 106     143.863  44.703  89.351  1.00 56.29           O
    ATOM    802  CG2 THR A 106     146.052  45.044  90.186  1.00 56.29           C
    ATOM    803  H   THR A 106     142.705  43.089  91.422  1.00  0.00           H
    ATOM    804  HA  THR A 106     144.902  42.404  89.759  1.00  0.00           H
    ATOM    805  HB  THR A 106     144.234  44.935  91.313  1.00  0.00           H
    ATOM    806 HG21 THR A 106     145.939  46.041  89.762  1.00  0.00           H
    ATOM    807 HG22 THR A 106     146.653  45.131  91.092  1.00  0.00           H
    ATOM    808 HG23 THR A 106     146.616  44.452  89.463  1.00  0.00           H
    ATOM    809  HG1 THR A 106     144.266  44.315  88.570  1.00  0.00           H
    ATOM    810  N   THR A 107     145.726  42.967  92.966  1.00 23.19           N
    ATOM    811  CA  THR A 107     146.717  42.564  93.967  1.00 23.19           C
    ATOM    812  C   THR A 107     146.665  41.064  94.284  1.00 23.19           C
    ATOM    813  O   THR A 107     147.697  40.534  94.668  1.00 23.19           O
    ATOM    814  CB  THR A 107     146.694  43.357  95.291  1.00 30.41           C
    ATOM    815  OG1 THR A 107     145.416  43.383  95.889  1.00 30.41           O
    ATOM    816  CG2 THR A 107     147.186  44.788  95.067  1.00 30.41           C
    ATOM    817  H   THR A 107     144.960  43.551  93.282  1.00  0.00           H
    ATOM    818  HA  THR A 107     147.682  42.800  93.531  1.00  0.00           H
    ATOM    819  HB  THR A 107     147.367  42.894  96.015  1.00  0.00           H
    ATOM    820 HG21 THR A 107     147.146  45.364  95.982  1.00  0.00           H
    ATOM    821 HG22 THR A 107     148.222  44.798  94.724  1.00  0.00           H
    ATOM    822 HG23 THR A 107     146.587  45.314  94.322  1.00  0.00           H
    ATOM    823  HG1 THR A 107     144.842  43.937  95.341  1.00  0.00           H
    ATOM    824  N   TYR A 108     145.532  40.381  94.047  1.00 42.11           N
    ATOM    825  CA  TYR A 108     145.456  38.928  94.151  1.00 42.11           C
    ATOM    826  C   TYR A 108     146.398  38.256  93.147  1.00 42.11           C
    ATOM    827  O   TYR A 108     147.306  37.543  93.564  1.00 42.11           O
    ATOM    828  CB  TYR A 108     144.020  38.408  93.975  1.00 47.51           C
    ATOM    829  CG  TYR A 108     143.970  36.898  94.042  1.00 47.51           C
    ATOM    830  CD1 TYR A 108     144.090  36.270  95.294  1.00 47.51           C
    ATOM    831  CD2 TYR A 108     143.954  36.123  92.864  1.00 47.51           C
    ATOM    832  CE1 TYR A 108     144.202  34.874  95.374  1.00 47.51           C
    ATOM    833  CE2 TYR A 108     144.102  34.725  92.940  1.00 47.51           C
    ATOM    834  CZ  TYR A 108     144.224  34.103  94.198  1.00 47.51           C
    ATOM    835  OH  TYR A 108     144.357  32.750  94.288  1.00 47.51           O
    ATOM    836  H   TYR A 108     144.694  40.869  93.756  1.00  0.00           H
    ATOM    837  HA  TYR A 108     145.787  38.657  95.156  1.00  0.00           H
    ATOM    838  HB2 TYR A 108     143.352  38.837  94.722  1.00  0.00           H
    ATOM    839  HB3 TYR A 108     143.643  38.693  93.002  1.00  0.00           H
    ATOM    840  HD1 TYR A 108     144.115  36.854  96.199  1.00  0.00           H
    ATOM    841  HD2 TYR A 108     143.875  36.603  91.903  1.00  0.00           H
    ATOM    842  HE1 TYR A 108     144.293  34.397  96.338  1.00  0.00           H
    ATOM    843  HE2 TYR A 108     144.112  34.138  92.034  1.00  0.00           H
    ATOM    844  HH  TYR A 108     144.309  32.312  93.436  1.00  0.00           H
    ATOM    845  N   LEU A 109     146.156  38.509  91.853  1.00 43.50           N
    ATOM    846  CA  LEU A 109     146.921  37.956  90.738  1.00 43.50           C
    ATOM    847  C   LEU A 109     148.400  38.364  90.776  1.00 43.50           C
    ATOM    848  O   LEU A 109     149.256  37.537  90.471  1.00 43.50           O
    ATOM    849  CB  LEU A 109     146.258  38.381  89.414  1.00 35.96           C
    ATOM    850  CG  LEU A 109     144.860  37.767  89.203  1.00 35.96           C
    ATOM    851  CD1 LEU A 109     144.175  38.406  87.988  1.00 35.96           C
    ATOM    852  CD2 LEU A 109     144.901  36.235  89.084  1.00 35.96           C
    ATOM    853  H   LEU A 109     145.392  39.124  91.601  1.00  0.00           H
    ATOM    854  HA  LEU A 109     146.892  36.868  90.823  1.00  0.00           H
    ATOM    855  HB2 LEU A 109     146.187  39.470  89.388  1.00  0.00           H
    ATOM    856  HB3 LEU A 109     146.894  38.109  88.570  1.00  0.00           H
    ATOM    857  HG  LEU A 109     144.237  37.990  90.064  1.00  0.00           H
    ATOM    858 HD11 LEU A 109     143.173  38.004  87.836  1.00  0.00           H
    ATOM    859 HD12 LEU A 109     144.081  39.486  88.107  1.00  0.00           H
    ATOM    860 HD13 LEU A 109     144.743  38.223  87.076  1.00  0.00           H
    ATOM    861 HD21 LEU A 109     144.080  35.852  88.482  1.00  0.00           H
    ATOM    862 HD22 LEU A 109     145.830  35.891  88.628  1.00  0.00           H
    ATOM    863 HD23 LEU A 109     144.823  35.755  90.058  1.00  0.00           H
    ATOM    864  N   ASP A 110     148.668  39.601  91.218  1.00 38.23           N
    ATOM    865  CA  ASP A 110     149.992  40.137  91.527  1.00 38.23           C
    ATOM    866  C   ASP A 110     150.758  39.276  92.558  1.00 38.23           C
    ATOM    867  O   ASP A 110     151.948  39.026  92.381  1.00 38.23           O
    ATOM    868  CB  ASP A 110     149.822  41.591  92.014  1.00 98.08           C
    ATOM    869  CG  ASP A 110     151.030  42.243  92.675  1.00 98.08           C
    ATOM    870  OD1 ASP A 110     152.119  42.160  92.068  1.00 98.08           O
    ATOM    871  OD2 ASP A 110     150.830  42.851  93.749  1.00 98.08           O1-
    ATOM    872  H   ASP A 110     147.894  40.219  91.426  1.00  0.00           H
    ATOM    873  HA  ASP A 110     150.569  40.146  90.601  1.00  0.00           H
    ATOM    874  HB2 ASP A 110     149.449  42.240  91.222  1.00  0.00           H
    ATOM    875  HB3 ASP A 110     149.059  41.541  92.773  1.00  0.00           H
    ATOM    876  N   LYS A 111     150.052  38.873  93.623  1.00 57.21           N
    ATOM    877  CA  LYS A 111     150.615  38.299  94.839  1.00 57.21           C
    ATOM    878  C   LYS A 111     150.987  36.810  94.757  1.00 57.21           C
    ATOM    879  O   LYS A 111     151.964  36.420  95.394  1.00 57.21           O
    ATOM    880  CB  LYS A 111     149.592  38.575  95.946  1.00 99.90           C
    ATOM    881  CG  LYS A 111     149.992  38.282  97.388  1.00 99.90           C
    ATOM    882  CD  LYS A 111     148.771  38.423  98.314  1.00 99.90           C
    ATOM    883  CE  LYS A 111     147.762  37.269  98.188  1.00 99.90           C
    ATOM    884  NZ  LYS A 111     148.416  35.960  98.366  1.00 99.90           N1+
    ATOM    885  H   LYS A 111     149.070  39.110  93.672  1.00  0.00           H
    ATOM    886  HA  LYS A 111     151.526  38.852  95.079  1.00  0.00           H
    ATOM    887  HB2 LYS A 111     149.413  39.646  95.955  1.00  0.00           H
    ATOM    888  HB3 LYS A 111     148.653  38.087  95.696  1.00  0.00           H
    ATOM    889  HG2 LYS A 111     150.453  37.301  97.490  1.00  0.00           H
    ATOM    890  HG3 LYS A 111     150.758  38.996  97.696  1.00  0.00           H
    ATOM    891  HD2 LYS A 111     149.100  38.529  99.344  1.00  0.00           H
    ATOM    892  HD3 LYS A 111     148.255  39.358  98.091  1.00  0.00           H
    ATOM    893  HE2 LYS A 111     146.984  37.380  98.944  1.00  0.00           H
    ATOM    894  HE3 LYS A 111     147.257  37.283  97.222  1.00  0.00           H
    ATOM    895  HZ1 LYS A 111     148.877  35.948  99.268  1.00  0.00           H
    ATOM    896  HZ2 LYS A 111     147.727  35.223  98.327  1.00  0.00           H
    ATOM    897  HZ3 LYS A 111     149.106  35.825  97.642  1.00  0.00           H
    ATOM    898  N   VAL A 112     150.210  36.006  94.016  1.00 58.48           N
    ATOM    899  CA  VAL A 112     150.345  34.547  93.931  1.00 58.48           C
    ATOM    900  C   VAL A 112     151.780  34.041  93.590  1.00 58.48           C
    ATOM    901  O   VAL A 112     152.460  34.692  92.793  1.00 58.48           O
    ATOM    902  CB  VAL A 112     149.348  34.030  92.863  1.00 86.94           C
    ATOM    903  CG1 VAL A 112     149.639  32.652  92.282  1.00 86.94           C
    ATOM    904  CG2 VAL A 112     147.899  34.039  93.382  1.00 86.94           C
    ATOM    905  H   VAL A 112     149.435  36.410  93.508  1.00  0.00           H
    ATOM    906  HA  VAL A 112     150.043  34.197  94.917  1.00  0.00           H
    ATOM    907  HB  VAL A 112     149.381  34.726  92.021  1.00  0.00           H
    ATOM    908 HG11 VAL A 112     148.835  32.483  91.596  1.00  0.00           H
    ATOM    909 HG12 VAL A 112     150.553  32.619  91.694  1.00  0.00           H
    ATOM    910 HG13 VAL A 112     149.624  31.837  93.001  1.00  0.00           H
    ATOM    911 HG21 VAL A 112     147.188  33.788  92.595  1.00  0.00           H
    ATOM    912 HG22 VAL A 112     147.765  33.319  94.190  1.00  0.00           H
    ATOM    913 HG23 VAL A 112     147.615  35.015  93.769  1.00  0.00           H
    ATOM    914  N   PRO A 113     152.203  32.888  94.177  1.00 42.92           N
    ATOM    915  CA  PRO A 113     153.508  32.245  93.890  1.00 42.92           C
    ATOM    916  C   PRO A 113     153.724  31.915  92.399  1.00 42.92           C
    ATOM    917  O   PRO A 113     152.785  31.468  91.744  1.00 42.92           O
    ATOM    918  CB  PRO A 113     153.479  30.961  94.743  1.00 99.90           C
    ATOM    919  CG  PRO A 113     152.005  30.671  94.957  1.00 99.90           C
    ATOM    920  CD  PRO A 113     151.440  32.071  95.128  1.00 99.90           C
    ATOM    921  HA  PRO A 113     154.315  32.897  94.229  1.00  0.00           H
    ATOM    922  HB2 PRO A 113     154.002  30.117  94.291  1.00  0.00           H
    ATOM    923  HB3 PRO A 113     153.957  31.158  95.703  1.00  0.00           H
    ATOM    924  HG2 PRO A 113     151.581  30.204  94.066  1.00  0.00           H
    ATOM    925  HG3 PRO A 113     151.811  30.020  95.809  1.00  0.00           H
    ATOM    926  HD2 PRO A 113     150.361  32.061  94.996  1.00  0.00           H
    ATOM    927  HD3 PRO A 113     151.641  32.443  96.134  1.00  0.00           H
    ATOM    928  N   GLU A 114     154.962  32.127  91.913  1.00 54.75           N
    ATOM    929  CA  GLU A 114     155.359  32.205  90.498  1.00 54.75           C
    ATOM    930  C   GLU A 114     154.803  31.255  89.406  1.00 54.75           C
    ATOM    931  O   GLU A 114     154.667  31.752  88.289  1.00 54.75           O
    ATOM    932  CB  GLU A 114     156.893  32.378  90.362  1.00 99.90           C
    ATOM    933  CG  GLU A 114     157.433  32.650  88.929  1.00 99.90           C
    ATOM    934  CD  GLU A 114     156.785  33.874  88.260  1.00 99.90           C
    ATOM    935  OE1 GLU A 114     156.793  34.943  88.905  1.00 99.90           O
    ATOM    936  OE2 GLU A 114     156.242  33.722  87.143  1.00 99.90           O1-
    ATOM    937  H   GLU A 114     155.673  32.437  92.559  1.00  0.00           H
    ATOM    938  HA  GLU A 114     154.930  33.170  90.222  1.00  0.00           H
    ATOM    939  HB2 GLU A 114     157.212  33.196  91.009  1.00  0.00           H
    ATOM    940  HB3 GLU A 114     157.383  31.488  90.758  1.00  0.00           H
    ATOM    941  HG2 GLU A 114     158.507  32.829  88.982  1.00  0.00           H
    ATOM    942  HG3 GLU A 114     157.315  31.773  88.291  1.00  0.00           H
    ATOM    943  N   GLY A 116     151.729  30.582  88.944  1.00 90.84           N1+
    ATOM    944  CA  GLY A 116     150.546  31.363  88.664  1.00 90.84           C
    ATOM    945  C   GLY A 116     149.457  30.885  89.606  1.00 90.84           C
    ATOM    946  O   GLY A 116     149.681  30.071  90.507  1.00 90.84           O
    ATOM    947  H   GLY A 116     152.004  30.528  89.916  1.00  0.00           H
    ATOM    948  HA2 GLY A 116     150.761  32.409  88.875  1.00  0.00           H
    ATOM    949  HA3 GLY A 116     150.209  31.277  87.630  1.00  0.00           H
    ATOM    950  N   VAL A 117     148.262  31.457  89.421  1.00 41.73           N
    ATOM    951  CA  VAL A 117     147.064  31.092  90.170  1.00 41.73           C
    ATOM    952  C   VAL A 117     146.710  29.622  89.867  1.00 41.73           C
    ATOM    953  O   VAL A 117     146.729  29.245  88.694  1.00 41.73           O
    ATOM    954  CB  VAL A 117     145.894  32.039  89.808  1.00 45.10           C
    ATOM    955  CG1 VAL A 117     145.731  32.253  88.302  1.00 45.10           C
    ATOM    956  CG2 VAL A 117     144.553  31.629  90.441  1.00 45.10           C
    ATOM    957  H   VAL A 117     148.131  32.124  88.675  1.00  0.00           H
    ATOM    958  HA  VAL A 117     147.247  31.237  91.228  1.00  0.00           H
    ATOM    959  HB  VAL A 117     146.135  33.015  90.234  1.00  0.00           H
    ATOM    960 HG11 VAL A 117     144.812  32.799  88.092  1.00  0.00           H
    ATOM    961 HG12 VAL A 117     146.562  32.826  87.892  1.00  0.00           H
    ATOM    962 HG13 VAL A 117     145.689  31.313  87.763  1.00  0.00           H
    ATOM    963 HG21 VAL A 117     143.791  32.389  90.268  1.00  0.00           H
    ATOM    964 HG22 VAL A 117     144.167  30.698  90.032  1.00  0.00           H
    ATOM    965 HG23 VAL A 117     144.647  31.506  91.519  1.00  0.00           H
    ATOM    966  N   PRO A 118     146.463  28.795  90.910  1.00 39.29           N
    ATOM    967  CA  PRO A 118     146.006  27.408  90.701  1.00 39.29           C
    ATOM    968  C   PRO A 118     144.759  27.296  89.799  1.00 39.29           C
    ATOM    969  O   PRO A 118     143.892  28.166  89.841  1.00 39.29           O
    ATOM    970  CB  PRO A 118     145.753  26.897  92.127  1.00 99.90           C
    ATOM    971  CG  PRO A 118     146.692  27.715  92.996  1.00 99.90           C
    ATOM    972  CD  PRO A 118     146.661  29.083  92.334  1.00 99.90           C
    ATOM    973  HA  PRO A 118     146.836  26.856  90.257  1.00  0.00           H
    ATOM    974  HB2 PRO A 118     144.722  27.077  92.431  1.00  0.00           H
    ATOM    975  HB3 PRO A 118     145.935  25.826  92.224  1.00  0.00           H
    ATOM    976  HG2 PRO A 118     146.404  27.736  94.047  1.00  0.00           H
    ATOM    977  HG3 PRO A 118     147.699  27.301  92.929  1.00  0.00           H
    ATOM    978  HD2 PRO A 118     145.823  29.677  92.701  1.00  0.00           H
    ATOM    979  HD3 PRO A 118     147.590  29.611  92.550  1.00  0.00           H
    ATOM    980  N   THR A 121     141.086  27.026  91.507  1.00 67.49           N1+
    ATOM    981  CA  THR A 121     140.649  28.312  92.017  1.00 67.49           C
    ATOM    982  C   THR A 121     140.220  29.293  90.922  1.00 67.49           C
    ATOM    983  O   THR A 121     139.247  29.985  91.196  1.00 67.49           O
    ATOM    984  CB  THR A 121     141.702  28.996  92.920  1.00 99.90           C
    ATOM    985  OG1 THR A 121     142.974  29.038  92.313  1.00 99.90           O
    ATOM    986  CG2 THR A 121     141.856  28.303  94.276  1.00 99.90           C
    ATOM    987  H   THR A 121     142.033  26.761  91.736  1.00  0.00           H
    ATOM    988  HA  THR A 121     139.769  28.112  92.632  1.00  0.00           H
    ATOM    989  HB  THR A 121     141.404  30.029  93.112  1.00  0.00           H
    ATOM    990 HG21 THR A 121     142.570  28.829  94.911  1.00  0.00           H
    ATOM    991 HG22 THR A 121     140.906  28.267  94.810  1.00  0.00           H
    ATOM    992 HG23 THR A 121     142.208  27.277  94.161  1.00  0.00           H
    ATOM    993  HG1 THR A 121     143.616  29.313  92.971  1.00  0.00           H
    ATOM    994  N   MET A 125     136.777  31.166  91.216  1.00 74.23           N1+
    ATOM    995  CA  MET A 125     136.769  32.625  91.125  1.00 74.23           C
    ATOM    996  C   MET A 125     135.997  33.139  89.912  1.00 74.23           C
    ATOM    997  O   MET A 125     135.104  33.962  90.049  1.00 74.23           O
    ATOM    998  CB  MET A 125     138.202  33.133  91.004  1.00 59.10           C
    ATOM    999  CG  MET A 125     138.985  33.136  92.301  1.00 59.10           C
    ATOM   1000  SD  MET A 125     140.514  34.071  92.098  1.00 59.10           S
    ATOM   1001  CE  MET A 125     140.871  34.267  93.846  1.00 59.10           C
    ATOM   1002  H   MET A 125     137.690  30.728  91.257  1.00  0.00           H
    ATOM   1003  HA  MET A 125     136.305  33.031  92.025  1.00  0.00           H
    ATOM   1004  HB2 MET A 125     138.749  32.555  90.259  1.00  0.00           H
    ATOM   1005  HB3 MET A 125     138.181  34.152  90.639  1.00  0.00           H
    ATOM   1006  HG2 MET A 125     138.385  33.604  93.082  1.00  0.00           H
    ATOM   1007  HG3 MET A 125     139.214  32.126  92.640  1.00  0.00           H
    ATOM   1008  HE1 MET A 125     141.641  35.017  93.965  1.00  0.00           H
    ATOM   1009  HE2 MET A 125     139.983  34.597  94.382  1.00  0.00           H
    ATOM   1010  HE3 MET A 125     141.213  33.327  94.281  1.00  0.00           H
    ATOM   1011  N   GLN A 128     132.356  32.362  90.412  1.00 62.93           N1+
    ATOM   1012  CA  GLN A 128     131.879  33.344  91.362  1.00 62.93           C
    ATOM   1013  C   GLN A 128     131.588  34.690  90.707  1.00 62.93           C
    ATOM   1014  O   GLN A 128     130.471  35.183  90.842  1.00 62.93           O
    ATOM   1015  CB  GLN A 128     132.933  33.495  92.454  1.00 51.51           C
    ATOM   1016  CG  GLN A 128     132.956  32.344  93.458  1.00 51.51           C
    ATOM   1017  CD  GLN A 128     134.038  32.632  94.481  1.00 51.51           C
    ATOM   1018  OE1 GLN A 128     133.805  33.448  95.360  1.00 51.51           O
    ATOM   1019  NE2 GLN A 128     135.211  32.014  94.366  1.00 51.51           N
    ATOM   1020  H   GLN A 128     133.265  31.960  90.616  1.00  0.00           H
    ATOM   1021  HA  GLN A 128     130.951  32.973  91.803  1.00  0.00           H
    ATOM   1022  HB2 GLN A 128     133.922  33.558  92.019  1.00  0.00           H
    ATOM   1023  HB3 GLN A 128     132.825  34.450  92.953  1.00  0.00           H
    ATOM   1024  HG2 GLN A 128     131.998  32.282  93.974  1.00  0.00           H
    ATOM   1025  HG3 GLN A 128     133.106  31.372  92.995  1.00  0.00           H
    ATOM   1026 HE21 GLN A 128     135.347  31.301  93.663  1.00  0.00           H
    ATOM   1027 HE22 GLN A 128     135.954  32.196  95.027  1.00  0.00           H
    ATOM   1028  N   HIE A 143     126.995  46.955  85.234  1.00 52.87           N1+
    ATOM   1029  CA  HIE A 143     128.173  47.438  85.942  1.00 52.87           C
    ATOM   1030  C   HIE A 143     129.002  48.428  85.104  1.00 52.87           C
    ATOM   1031  O   HIE A 143     129.398  49.466  85.630  1.00 52.87           O
    ATOM   1032  CB  HIE A 143     128.975  46.213  86.419  1.00 69.54           C
    ATOM   1033  CG  HIE A 143     130.160  46.481  87.301  1.00 69.54           C
    ATOM   1034  ND1 HIE A 143     130.433  45.694  88.418  1.00 69.54           N
    ATOM   1035  CD2 HIE A 143     131.152  47.428  87.188  1.00 69.54           C
    ATOM   1036  CE1 HIE A 143     131.573  46.165  88.900  1.00 69.54           C
    ATOM   1037  NE2 HIE A 143     132.038  47.207  88.219  1.00 69.54           N
    ATOM   1038  H   HIE A 143     126.984  45.961  85.035  1.00  0.00           H
    ATOM   1039  HA  HIE A 143     127.816  47.970  86.827  1.00  0.00           H
    ATOM   1040  HB2 HIE A 143     128.307  45.543  86.963  1.00  0.00           H
    ATOM   1041  HB3 HIE A 143     129.357  45.632  85.591  1.00  0.00           H
    ATOM   1042  HD2 HIE A 143     131.314  48.226  86.492  1.00  0.00           H
    ATOM   1043  HE2 HIE A 143     132.885  47.727  88.406  1.00  0.00           H
    ATOM   1044  HE1 HIE A 143     132.070  45.727  89.751  1.00  0.00           H
    ATOM   1045  N   ASP A 145     131.889  49.283  84.213  1.00 80.80           N1+
    ATOM   1046  CA  ASP A 145     133.275  49.504  84.603  1.00 80.80           C
    ATOM   1047  C   ASP A 145     133.754  48.295  85.427  1.00 80.80           C
    ATOM   1048  O   ASP A 145     134.156  48.464  86.573  1.00 80.80           O
    ATOM   1049  CB  ASP A 145     133.376  50.842  85.382  1.00 99.90           C
    ATOM   1050  CG  ASP A 145     134.804  51.310  85.678  1.00 99.90           C
    ATOM   1051  OD1 ASP A 145     135.740  50.833  84.999  1.00 99.90           O
    ATOM   1052  OD2 ASP A 145     134.940  52.188  86.555  1.00 99.90           O1-
    ATOM   1053  H   ASP A 145     131.202  49.410  84.946  1.00  0.00           H
    ATOM   1054  HA  ASP A 145     133.907  49.561  83.715  1.00  0.00           H
    ATOM   1055  HB2 ASP A 145     132.882  51.629  84.811  1.00  0.00           H
    ATOM   1056  HB3 ASP A 145     132.842  50.747  86.330  1.00  0.00           H
    ATOM   1057  N   LEU A 146     133.648  47.076  84.877  1.00 23.34           N
    ATOM   1058  CA  LEU A 146     133.980  45.847  85.603  1.00 23.34           C
    ATOM   1059  C   LEU A 146     135.458  45.471  85.407  1.00 23.34           C
    ATOM   1060  O   LEU A 146     135.760  44.425  84.843  1.00 23.34           O
    ATOM   1061  CB  LEU A 146     132.995  44.737  85.191  1.00 34.89           C
    ATOM   1062  CG  LEU A 146     133.122  43.442  86.032  1.00 34.89           C
    ATOM   1063  CD1 LEU A 146     131.943  43.232  86.990  1.00 34.89           C
    ATOM   1064  CD2 LEU A 146     133.334  42.220  85.132  1.00 34.89           C
    ATOM   1065  H   LEU A 146     133.293  46.970  83.937  1.00  0.00           H
    ATOM   1066  HA  LEU A 146     133.845  45.996  86.675  1.00  0.00           H
    ATOM   1067  HB2 LEU A 146     131.977  45.125  85.248  1.00  0.00           H
    ATOM   1068  HB3 LEU A 146     133.133  44.523  84.131  1.00  0.00           H
    ATOM   1069  HG  LEU A 146     134.012  43.516  86.653  1.00  0.00           H
    ATOM   1070 HD11 LEU A 146     131.670  42.185  87.063  1.00  0.00           H
    ATOM   1071 HD12 LEU A 146     132.201  43.567  87.992  1.00  0.00           H
    ATOM   1072 HD13 LEU A 146     131.046  43.760  86.673  1.00  0.00           H
    ATOM   1073 HD21 LEU A 146     133.731  41.376  85.696  1.00  0.00           H
    ATOM   1074 HD22 LEU A 146     132.405  41.929  84.643  1.00  0.00           H
    ATOM   1075 HD23 LEU A 146     134.045  42.428  84.339  1.00  0.00           H
    ATOM   1076  N   LYS A 147     136.364  46.330  85.886  1.00 44.77           N
    ATOM   1077  CA  LYS A 147     137.805  46.071  85.916  1.00 44.77           C
    ATOM   1078  C   LYS A 147     138.178  45.103  87.069  1.00 44.77           C
    ATOM   1079  O   LYS A 147     137.376  44.944  87.985  1.00 44.77           O
    ATOM   1080  CB  LYS A 147     138.541  47.432  85.935  1.00 68.67           C
    ATOM   1081  CG  LYS A 147     138.077  48.461  86.992  1.00 68.67           C
    ATOM   1082  CD  LYS A 147     138.586  49.877  86.662  1.00 68.67           C
    ATOM   1083  CE  LYS A 147     138.425  50.904  87.790  1.00 68.67           C
    ATOM   1084  NZ  LYS A 147     137.025  51.240  88.104  1.00 68.67           N1+
    ATOM   1085  H   LYS A 147     136.049  47.183  86.329  1.00  0.00           H
    ATOM   1086  HA  LYS A 147     138.052  45.592  84.969  1.00  0.00           H
    ATOM   1087  HB2 LYS A 147     139.618  47.286  86.017  1.00  0.00           H
    ATOM   1088  HB3 LYS A 147     138.387  47.883  84.953  1.00  0.00           H
    ATOM   1089  HG2 LYS A 147     136.991  48.504  87.035  1.00  0.00           H
    ATOM   1090  HG3 LYS A 147     138.415  48.148  87.979  1.00  0.00           H
    ATOM   1091  HD2 LYS A 147     139.644  49.819  86.403  1.00  0.00           H
    ATOM   1092  HD3 LYS A 147     138.093  50.244  85.764  1.00  0.00           H
    ATOM   1093  HE2 LYS A 147     138.922  50.513  88.675  1.00  0.00           H
    ATOM   1094  HE3 LYS A 147     138.937  51.826  87.514  1.00  0.00           H
    ATOM   1095  HZ1 LYS A 147     136.551  51.615  87.286  1.00  0.00           H
    ATOM   1096  HZ2 LYS A 147     136.521  50.407  88.388  1.00  0.00           H
    ATOM   1097  HZ3 LYS A 147     136.982  51.925  88.842  1.00  0.00           H
    ATOM   1098  N   PRO A 148     139.373  44.463  87.043  1.00 67.16           N
    ATOM   1099  CA  PRO A 148     139.880  43.637  88.154  1.00 67.16           C
    ATOM   1100  C   PRO A 148     139.941  44.370  89.496  1.00 67.16           C
    ATOM   1101  O   PRO A 148     139.580  43.776  90.502  1.00 67.16           O
    ATOM   1102  CB  PRO A 148     141.272  43.192  87.712  1.00 61.95           C
    ATOM   1103  CG  PRO A 148     141.207  43.217  86.212  1.00 61.95           C
    ATOM   1104  CD  PRO A 148     140.286  44.399  85.923  1.00 61.95           C
    ATOM   1105  HA  PRO A 148     139.275  42.737  88.225  1.00  0.00           H
    ATOM   1106  HB2 PRO A 148     142.017  43.915  88.019  1.00  0.00           H
    ATOM   1107  HB3 PRO A 148     141.561  42.217  88.107  1.00  0.00           H
    ATOM   1108  HG2 PRO A 148     142.189  43.326  85.764  1.00  0.00           H
    ATOM   1109  HG3 PRO A 148     140.786  42.282  85.854  1.00  0.00           H
    ATOM   1110  HD2 PRO A 148     140.850  45.329  85.864  1.00  0.00           H
    ATOM   1111  HD3 PRO A 148     139.735  44.239  85.004  1.00  0.00           H
    ATOM   1112  N   GLN A 149     140.316  45.662  89.473  1.00 50.56           N
    ATOM   1113  CA  GLN A 149     140.356  46.572  90.627  1.00 50.56           C
    ATOM   1114  C   GLN A 149     139.017  46.833  91.332  1.00 50.56           C
    ATOM   1115  O   GLN A 149     138.984  47.528  92.347  1.00 50.56           O
    ATOM   1116  CB  GLN A 149     140.913  47.919  90.180  1.00 86.10           C
    ATOM   1117  CG  GLN A 149     142.357  47.838  89.702  1.00 86.10           C
    ATOM   1118  CD  GLN A 149     142.846  49.247  89.455  1.00 86.10           C
    ATOM   1119  OE1 GLN A 149     142.998  50.029  90.395  1.00 86.10           O
    ATOM   1120  NE2 GLN A 149     143.055  49.566  88.182  1.00 86.10           N
    ATOM   1121  H   GLN A 149     140.590  46.068  88.591  1.00  0.00           H
    ATOM   1122  HA  GLN A 149     141.037  46.150  91.359  1.00  0.00           H
    ATOM   1123  HB2 GLN A 149     140.288  48.335  89.387  1.00  0.00           H
    ATOM   1124  HB3 GLN A 149     140.857  48.627  91.009  1.00  0.00           H
    ATOM   1125  HG2 GLN A 149     142.990  47.377  90.461  1.00  0.00           H
    ATOM   1126  HG3 GLN A 149     142.445  47.229  88.802  1.00  0.00           H
    ATOM   1127 HE21 GLN A 149     142.909  48.881  87.455  1.00  0.00           H
    ATOM   1128 HE22 GLN A 149     143.360  50.493  87.930  1.00  0.00           H
    ATOM   1129  N   ASN A 150     137.940  46.316  90.745  1.00 30.81           N
    ATOM   1130  CA  ASN A 150     136.567  46.398  91.206  1.00 30.81           C
    ATOM   1131  C   ASN A 150     136.079  45.020  91.644  1.00 30.81           C
    ATOM   1132  O   ASN A 150     135.187  44.944  92.484  1.00 30.81           O
    ATOM   1133  CB  ASN A 150     135.791  46.985  90.009  1.00 40.82           C
    ATOM   1134  CG  ASN A 150     135.843  48.509  89.913  1.00 40.82           C
    ATOM   1135  OD1 ASN A 150     135.373  49.059  88.930  1.00 40.82           O
    ATOM   1136  ND2 ASN A 150     136.416  49.220  90.887  1.00 40.82           N
    ATOM   1137  H   ASN A 150     138.072  45.786  89.895  1.00  0.00           H
    ATOM   1138  HA  ASN A 150     136.529  47.048  92.081  1.00  0.00           H
    ATOM   1139  HB2 ASN A 150     136.182  46.622  89.069  1.00  0.00           H
    ATOM   1140  HB3 ASN A 150     134.767  46.646  89.953  1.00  0.00           H
    ATOM   1141 HD21 ASN A 150     136.824  48.752  91.685  1.00  0.00           H
    ATOM   1142 HD22 ASN A 150     136.458  50.224  90.831  1.00  0.00           H
    ATOM   1143  N   ILE A 151     136.719  43.956  91.146  1.00 43.16           N
    ATOM   1144  CA  ILE A 151     136.535  42.609  91.647  1.00 43.16           C
    ATOM   1145  C   ILE A 151     137.379  42.508  92.921  1.00 43.16           C
    ATOM   1146  O   ILE A 151     138.576  42.781  92.893  1.00 43.16           O
    ATOM   1147  CB  ILE A 151     137.043  41.538  90.642  1.00 36.55           C
    ATOM   1148  CG1 ILE A 151     136.686  41.863  89.179  1.00 36.55           C
    ATOM   1149  CG2 ILE A 151     136.546  40.132  91.015  1.00 36.55           C
    ATOM   1150  CD1 ILE A 151     135.250  42.322  88.935  1.00 36.55           C
    ATOM   1151  H   ILE A 151     137.476  44.081  90.489  1.00  0.00           H
    ATOM   1152  HA  ILE A 151     135.481  42.431  91.876  1.00  0.00           H
    ATOM   1153  HB  ILE A 151     138.131  41.500  90.676  1.00  0.00           H
    ATOM   1154 HG12 ILE A 151     137.319  42.673  88.856  1.00  0.00           H
    ATOM   1155 HG13 ILE A 151     136.935  41.024  88.529  1.00  0.00           H
    ATOM   1156 HG21 ILE A 151     136.950  39.383  90.335  1.00  0.00           H
    ATOM   1157 HG22 ILE A 151     136.854  39.846  92.022  1.00  0.00           H
    ATOM   1158 HG23 ILE A 151     135.459  40.070  90.970  1.00  0.00           H
    ATOM   1159 HD11 ILE A 151     134.885  41.939  87.984  1.00  0.00           H
    ATOM   1160 HD12 ILE A 151     134.591  41.979  89.726  1.00  0.00           H
    ATOM   1161 HD13 ILE A 151     135.178  43.409  88.893  1.00  0.00           H
    ATOM   1162  N   LEU A 152     136.733  42.164  94.028  1.00 47.34           N
    ATOM   1163  CA  LEU A 152     137.410  41.964  95.292  1.00 47.34           C
    ATOM   1164  C   LEU A 152     137.589  40.471  95.481  1.00 47.34           C
    ATOM   1165  O   LEU A 152     136.671  39.716  95.186  1.00 47.34           O
    ATOM   1166  CB  LEU A 152     136.571  42.541  96.434  1.00 32.31           C
    ATOM   1167  CG  LEU A 152     136.313  44.060  96.363  1.00 32.31           C
    ATOM   1168  CD1 LEU A 152     135.395  44.521  97.515  1.00 32.31           C
    ATOM   1169  CD2 LEU A 152     137.617  44.881  96.300  1.00 32.31           C
    ATOM   1170  H   LEU A 152     135.735  41.988  94.014  1.00  0.00           H
    ATOM   1171  HA  LEU A 152     138.390  42.438  95.305  1.00  0.00           H
    ATOM   1172  HB2 LEU A 152     135.618  42.016  96.490  1.00  0.00           H
    ATOM   1173  HB3 LEU A 152     137.106  42.309  97.348  1.00  0.00           H
    ATOM   1174  HG  LEU A 152     135.762  44.254  95.441  1.00  0.00           H
    ATOM   1175 HD11 LEU A 152     134.648  45.224  97.150  1.00  0.00           H
    ATOM   1176 HD12 LEU A 152     134.849  43.691  97.965  1.00  0.00           H
    ATOM   1177 HD13 LEU A 152     135.945  45.022  98.311  1.00  0.00           H
    ATOM   1178 HD21 LEU A 152     137.573  45.773  96.926  1.00  0.00           H
    ATOM   1179 HD22 LEU A 152     138.482  44.308  96.632  1.00  0.00           H
    ATOM   1180 HD23 LEU A 152     137.813  45.216  95.281  1.00  0.00           H
    ATOM   1181  N   VAL A 153     138.754  40.084  95.992  1.00 32.09           N
    ATOM   1182  CA  VAL A 153     139.036  38.713  96.362  1.00 32.09           C
    ATOM   1183  C   VAL A 153     139.248  38.740  97.875  1.00 32.09           C
    ATOM   1184  O   VAL A 153     140.108  39.471  98.339  1.00 32.09           O
    ATOM   1185  CB  VAL A 153     140.323  38.222  95.658  1.00 58.22           C
    ATOM   1186  CG1 VAL A 153     140.712  36.817  96.114  1.00 58.22           C
    ATOM   1187  CG2 VAL A 153     140.170  38.243  94.130  1.00 58.22           C
    ATOM   1188  H   VAL A 153     139.455  40.770  96.243  1.00  0.00           H
    ATOM   1189  HA  VAL A 153     138.214  38.033  96.127  1.00  0.00           H
    ATOM   1190  HB  VAL A 153     141.149  38.892  95.901  1.00  0.00           H
    ATOM   1191 HG11 VAL A 153     141.544  36.442  95.536  1.00  0.00           H
    ATOM   1192 HG12 VAL A 153     141.029  36.792  97.156  1.00  0.00           H
    ATOM   1193 HG13 VAL A 153     139.871  36.138  95.992  1.00  0.00           H
    ATOM   1194 HG21 VAL A 153     141.083  37.895  93.650  1.00  0.00           H
    ATOM   1195 HG22 VAL A 153     139.362  37.589  93.804  1.00  0.00           H
    ATOM   1196 HG23 VAL A 153     139.957  39.244  93.759  1.00  0.00           H
    ATOM   1197  N   THR A 154     138.452  38.003  98.654  1.00 21.90           N
    ATOM   1198  CA  THR A 154     138.618  37.980 100.110  1.00 21.90           C
    ATOM   1199  C   THR A 154     139.841  37.160 100.540  1.00 21.90           C
    ATOM   1200  O   THR A 154     140.508  36.548  99.703  1.00 21.90           O
    ATOM   1201  CB  THR A 154     137.363  37.434 100.834  1.00 53.67           C
    ATOM   1202  OG1 THR A 154     137.238  36.046 100.656  1.00 53.67           O
    ATOM   1203  CG2 THR A 154     136.055  38.176 100.530  1.00 53.67           C
    ATOM   1204  H   THR A 154     137.710  37.435  98.264  1.00  0.00           H
    ATOM   1205  HA  THR A 154     138.823  38.975 100.481  1.00  0.00           H
    ATOM   1206  HB  THR A 154     137.506  37.557 101.908  1.00  0.00           H
    ATOM   1207 HG21 THR A 154     135.188  37.616 100.877  1.00  0.00           H
    ATOM   1208 HG22 THR A 154     136.048  39.119 101.071  1.00  0.00           H
    ATOM   1209 HG23 THR A 154     135.913  38.426  99.484  1.00  0.00           H
    ATOM   1210  HG1 THR A 154     136.472  35.737 101.147  1.00  0.00           H
    ATOM   1211  N   SER A 155     140.081  37.141 101.863  1.00 58.51           N
    ATOM   1212  CA  SER A 155     141.018  36.240 102.531  1.00 58.51           C
    ATOM   1213  C   SER A 155     140.844  34.789 102.064  1.00 58.51           C
    ATOM   1214  O   SER A 155     141.812  34.150 101.660  1.00 58.51           O
    ATOM   1215  CB  SER A 155     140.757  36.282 104.043  1.00 99.90           C
    ATOM   1216  OG  SER A 155     141.737  35.537 104.732  1.00 99.90           O
    ATOM   1217  H   SER A 155     139.513  37.710 102.474  1.00  0.00           H
    ATOM   1218  HA  SER A 155     142.034  36.575 102.315  1.00  0.00           H
    ATOM   1219  HB2 SER A 155     140.712  37.290 104.413  1.00  0.00           H
    ATOM   1220  HB3 SER A 155     139.782  35.859 104.289  1.00  0.00           H
    ATOM   1221  HG  SER A 155     142.583  35.986 104.658  1.00  0.00           H
    ATOM   1222  N   SER A 156     139.579  34.349 102.088  1.00 71.13           N
    ATOM   1223  CA  SER A 156     139.127  33.028 101.691  1.00 71.13           C
    ATOM   1224  C   SER A 156     138.924  32.864 100.175  1.00 71.13           C
    ATOM   1225  O   SER A 156     138.266  31.903  99.780  1.00 71.13           O
    ATOM   1226  CB  SER A 156     137.867  32.694 102.517  1.00 86.74           C
    ATOM   1227  OG  SER A 156     136.769  33.534 102.210  1.00 86.74           O
    ATOM   1228  H   SER A 156     138.846  34.975 102.391  1.00  0.00           H
    ATOM   1229  HA  SER A 156     139.897  32.309 101.975  1.00  0.00           H
    ATOM   1230  HB2 SER A 156     137.565  31.659 102.350  1.00  0.00           H
    ATOM   1231  HB3 SER A 156     138.082  32.777 103.583  1.00  0.00           H
    ATOM   1232  HG  SER A 156     136.010  33.236 102.718  1.00  0.00           H
    ATOM   1233  N   GLY A 157     139.459  33.776  99.341  1.00 58.73           N
    ATOM   1234  CA  GLY A 157     139.341  33.713  97.882  1.00 58.73           C
    ATOM   1235  C   GLY A 157     137.889  33.843  97.385  1.00 58.73           C
    ATOM   1236  O   GLY A 157     137.604  33.499  96.238  1.00 58.73           O
    ATOM   1237  H   GLY A 157     139.981  34.553  99.725  1.00  0.00           H
    ATOM   1238  HA2 GLY A 157     139.949  34.507  97.458  1.00  0.00           H
    ATOM   1239  HA3 GLY A 157     139.760  32.771  97.523  1.00  0.00           H
    ATOM   1240  N   GLN A 158     136.977  34.301  98.258  1.00 48.94           N
    ATOM   1241  CA  GLN A 158     135.578  34.576  97.974  1.00 48.94           C
    ATOM   1242  C   GLN A 158     135.512  35.892  97.204  1.00 48.94           C
    ATOM   1243  O   GLN A 158     136.045  36.889  97.675  1.00 48.94           O
    ATOM   1244  CB  GLN A 158     134.759  34.514  99.294  1.00 99.90           C
    ATOM   1245  CG  GLN A 158     133.656  35.568  99.545  1.00 99.90           C
    ATOM   1246  CD  GLN A 158     132.291  35.167  98.994  1.00 99.90           C
    ATOM   1247  OE1 GLN A 158     131.293  35.281  99.699  1.00 99.90           O
    ATOM   1248  NE2 GLN A 158     132.228  34.695  97.749  1.00 99.90           N
    ATOM   1249  H   GLN A 158     137.288  34.567  99.183  1.00  0.00           H
    ATOM   1250  HA  GLN A 158     135.216  33.771  97.339  1.00  0.00           H
    ATOM   1251  HB2 GLN A 158     134.328  33.515  99.379  1.00  0.00           H
    ATOM   1252  HB3 GLN A 158     135.427  34.546 100.147  1.00  0.00           H
    ATOM   1253  HG2 GLN A 158     133.535  35.668 100.625  1.00  0.00           H
    ATOM   1254  HG3 GLN A 158     133.912  36.568  99.200  1.00  0.00           H
    ATOM   1255 HE21 GLN A 158     133.061  34.595  97.187  1.00  0.00           H
    ATOM   1256 HE22 GLN A 158     131.334  34.440  97.357  1.00  0.00           H
    ATOM   1257  N   ILE A 159     134.904  35.873  96.019  1.00 33.70           N
    ATOM   1258  CA  ILE A 159     134.849  37.041  95.160  1.00 33.70           C
    ATOM   1259  C   ILE A 159     133.611  37.890  95.405  1.00 33.70           C
    ATOM   1260  O   ILE A 159     132.549  37.341  95.664  1.00 33.70           O
    ATOM   1261  CB  ILE A 159     135.010  36.662  93.669  1.00 22.43           C
    ATOM   1262  CG1 ILE A 159     136.524  36.572  93.399  1.00 22.43           C
    ATOM   1263  CG2 ILE A 159     134.315  37.595  92.650  1.00 22.43           C
    ATOM   1264  CD1 ILE A 159     136.893  36.189  91.976  1.00 22.43           C
    ATOM   1265  H   ILE A 159     134.503  35.015  95.659  1.00  0.00           H
    ATOM   1266  HA  ILE A 159     135.700  37.652  95.426  1.00  0.00           H
    ATOM   1267  HB  ILE A 159     134.597  35.665  93.524  1.00  0.00           H
    ATOM   1268 HG12 ILE A 159     136.987  37.538  93.602  1.00  0.00           H
    ATOM   1269 HG13 ILE A 159     136.989  35.865  94.088  1.00  0.00           H
    ATOM   1270 HG21 ILE A 159     134.456  37.229  91.637  1.00  0.00           H
    ATOM   1271 HG22 ILE A 159     133.234  37.631  92.786  1.00  0.00           H
    ATOM   1272 HG23 ILE A 159     134.702  38.612  92.701  1.00  0.00           H
    ATOM   1273 HD11 ILE A 159     137.950  35.933  91.922  1.00  0.00           H
    ATOM   1274 HD12 ILE A 159     136.304  35.342  91.657  1.00  0.00           H
    ATOM   1275 HD13 ILE A 159     136.727  37.007  91.276  1.00  0.00           H
    ATOM   1276  N   LYS A 160     133.787  39.210  95.283  1.00 53.61           N
    ATOM   1277  CA  LYS A 160     132.730  40.198  95.388  1.00 53.61           C
    ATOM   1278  C   LYS A 160     132.920  41.264  94.318  1.00 53.61           C
    ATOM   1279  O   LYS A 160     134.009  41.816  94.196  1.00 53.61           O
    ATOM   1280  CB  LYS A 160     132.713  40.784  96.807  1.00 39.62           C
    ATOM   1281  CG  LYS A 160     132.030  39.836  97.799  1.00 39.62           C
    ATOM   1282  CD  LYS A 160     132.424  40.098  99.247  1.00 39.62           C
    ATOM   1283  CE  LYS A 160     131.692  39.176 100.224  1.00 39.62           C
    ATOM   1284  NZ  LYS A 160     130.232  39.351 100.154  1.00 39.62           N1+
    ATOM   1285  H   LYS A 160     134.710  39.570  95.075  1.00  0.00           H
    ATOM   1286  HA  LYS A 160     131.779  39.710  95.216  1.00  0.00           H
    ATOM   1287  HB2 LYS A 160     133.729  41.008  97.128  1.00  0.00           H
    ATOM   1288  HB3 LYS A 160     132.173  41.733  96.825  1.00  0.00           H
    ATOM   1289  HG2 LYS A 160     130.951  39.919  97.667  1.00  0.00           H
    ATOM   1290  HG3 LYS A 160     132.273  38.798  97.598  1.00  0.00           H
    ATOM   1291  HD2 LYS A 160     133.482  39.884  99.346  1.00  0.00           H
    ATOM   1292  HD3 LYS A 160     132.272  41.146  99.509  1.00  0.00           H
    ATOM   1293  HE2 LYS A 160     131.928  38.134 100.007  1.00  0.00           H
    ATOM   1294  HE3 LYS A 160     132.022  39.371 101.245  1.00  0.00           H
    ATOM   1295  HZ1 LYS A 160     129.993  40.303 100.421  1.00  0.00           H
    ATOM   1296  HZ2 LYS A 160     129.903  39.163  99.217  1.00  0.00           H
    ATOM   1297  HZ3 LYS A 160     129.784  38.713 100.796  1.00  0.00           H
    ATOM   1298  N   ALA A 162     132.238  45.065  92.989  1.00 48.68           N1+
    ATOM   1299  CA  ALA A 162     132.192  46.396  93.585  1.00 48.68           C
    ATOM   1300  C   ALA A 162     132.617  47.408  92.524  1.00 48.68           C
    ATOM   1301  O   ALA A 162     133.192  47.011  91.522  1.00 48.68           O
    ATOM   1302  CB  ALA A 162     133.133  46.452  94.803  1.00 12.50           C
    ATOM   1303  H   ALA A 162     132.810  44.990  92.158  1.00  0.00           H
    ATOM   1304  HA  ALA A 162     131.168  46.620  93.892  1.00  0.00           H
    ATOM   1305  HB1 ALA A 162     133.054  47.409  95.320  1.00  0.00           H
    ATOM   1306  HB2 ALA A 162     132.903  45.671  95.526  1.00  0.00           H
    ATOM   1307  HB3 ALA A 162     134.177  46.325  94.514  1.00  0.00           H
    ATOM   1308  N   ASP A 163     132.354  48.699  92.748  1.00 59.79           N
    ATOM   1309  CA  ASP A 163     132.847  49.768  91.881  1.00 59.79           C
    ATOM   1310  C   ASP A 163     133.359  50.906  92.759  1.00 59.79           C
    ATOM   1311  O   ASP A 163     132.688  51.268  93.727  1.00 59.79           O
    ATOM   1312  CB  ASP A 163     131.805  50.252  90.849  1.00 99.90           C
    ATOM   1313  CG  ASP A 163     132.260  51.495  90.069  1.00 99.90           C
    ATOM   1314  OD1 ASP A 163     133.286  51.391  89.361  1.00 99.90           O
    ATOM   1315  OD2 ASP A 163     131.603  52.544  90.244  1.00 99.90           O1-
    ATOM   1316  H   ASP A 163     131.878  48.989  93.591  1.00  0.00           H
    ATOM   1317  HA  ASP A 163     133.703  49.404  91.320  1.00  0.00           H
    ATOM   1318  HB2 ASP A 163     131.569  49.464  90.133  1.00  0.00           H
    ATOM   1319  HB3 ASP A 163     130.862  50.477  91.349  1.00  0.00           H
    ATOM   1320  N   PHE A 164     134.504  51.476  92.358  1.00 72.49           N
    ATOM   1321  CA  PHE A 164     135.124  52.586  93.066  1.00 72.49           C
    ATOM   1322  C   PHE A 164     135.288  53.858  92.215  1.00 72.49           C
    ATOM   1323  O   PHE A 164     135.499  54.909  92.814  1.00 72.49           O
    ATOM   1324  CB  PHE A 164     136.436  52.098  93.734  1.00 88.65           C
    ATOM   1325  CG  PHE A 164     136.270  51.002  94.785  1.00 88.65           C
    ATOM   1326  CD1 PHE A 164     135.277  51.102  95.783  1.00 88.65           C
    ATOM   1327  CD2 PHE A 164     137.178  49.921  94.846  1.00 88.65           C
    ATOM   1328  CE1 PHE A 164     135.152  50.122  96.757  1.00 88.65           C
    ATOM   1329  CE2 PHE A 164     137.041  48.950  95.831  1.00 88.65           C
    ATOM   1330  CZ  PHE A 164     136.027  49.045  96.777  1.00 88.65           C
    ATOM   1331  H   PHE A 164     134.956  51.148  91.518  1.00  0.00           H
    ATOM   1332  HA  PHE A 164     134.458  52.923  93.858  1.00  0.00           H
    ATOM   1333  HB2 PHE A 164     137.093  51.700  92.964  1.00  0.00           H
    ATOM   1334  HB3 PHE A 164     136.968  52.933  94.193  1.00  0.00           H
    ATOM   1335  HD1 PHE A 164     134.588  51.929  95.784  1.00  0.00           H
    ATOM   1336  HD2 PHE A 164     137.974  49.834  94.121  1.00  0.00           H
    ATOM   1337  HE1 PHE A 164     134.374  50.199  97.502  1.00  0.00           H
    ATOM   1338  HE2 PHE A 164     137.727  48.116  95.860  1.00  0.00           H
    ATOM   1339  HZ  PHE A 164     135.924  48.283  97.535  1.00  0.00           H
    ATOM   1340  N   GLY A 165     135.207  53.795  90.868  1.00 61.74           N
    ATOM   1341  CA  GLY A 165     135.368  54.948  89.949  1.00 61.74           C
    ATOM   1342  C   GLY A 165     136.730  55.680  90.033  1.00 61.74           C
    ATOM   1343  O   GLY A 165     136.989  56.621  89.287  1.00 61.74           O
    ATOM   1344  H   GLY A 165     134.951  52.918  90.435  1.00  0.00           H
    ATOM   1345  HA2 GLY A 165     135.226  54.596  88.927  1.00  0.00           H
    ATOM   1346  HA3 GLY A 165     134.575  55.669  90.144  1.00  0.00           H
    ATOM   1347  N   LEU A 166     137.582  55.213  90.948  1.00 99.90           N
    ATOM   1348  CA  LEU A 166     138.917  55.589  91.347  1.00 99.90           C
    ATOM   1349  C   LEU A 166     139.868  56.007  90.234  1.00 99.90           C
    ATOM   1350  O   LEU A 166     140.560  57.012  90.397  1.00 99.90           O
    ATOM   1351  CB  LEU A 166     139.441  54.433  92.227  1.00 99.90           C
    ATOM   1352  CG  LEU A 166     139.739  53.030  91.607  1.00 99.90           C
    ATOM   1353  CD1 LEU A 166     140.014  51.963  92.700  1.00 99.90           C
    ATOM   1354  CD2 LEU A 166     138.632  52.481  90.699  1.00 99.90           C
    ATOM   1355  H   LEU A 166     137.216  54.460  91.512  1.00  0.00           H
    ATOM   1356  HA  LEU A 166     138.798  56.474  91.957  1.00  0.00           H
    ATOM   1357  HB2 LEU A 166     140.329  54.819  92.710  1.00  0.00           H
    ATOM   1358  HB3 LEU A 166     138.700  54.280  93.007  1.00  0.00           H
    ATOM   1359  HG  LEU A 166     140.618  53.119  90.973  1.00  0.00           H
    ATOM   1360 HD11 LEU A 166     139.443  51.046  92.562  1.00  0.00           H
    ATOM   1361 HD12 LEU A 166     141.056  51.651  92.724  1.00  0.00           H
    ATOM   1362 HD13 LEU A 166     139.734  52.311  93.694  1.00  0.00           H
    ATOM   1363 HD21 LEU A 166     138.691  51.400  90.603  1.00  0.00           H
    ATOM   1364 HD22 LEU A 166     137.639  52.688  91.077  1.00  0.00           H
    ATOM   1365 HD23 LEU A 166     138.706  52.900  89.698  1.00  0.00           H
    ATOM   1366  N   ALA A 167     139.825  55.255  89.123  1.00 99.90           N
    ATOM   1367  CA  ALA A 167     140.579  55.468  87.897  1.00 99.90           C
    ATOM   1368  C   ALA A 167     142.086  55.337  88.145  1.00 99.90           C
    ATOM   1369  O   ALA A 167     142.737  56.317  88.500  1.00 99.90           O
    ATOM   1370  CB  ALA A 167     140.197  56.822  87.269  1.00 99.90           C
    ATOM   1371  H   ALA A 167     139.181  54.479  89.104  1.00  0.00           H
    ATOM   1372  HA  ALA A 167     140.286  54.684  87.197  1.00  0.00           H
    ATOM   1373  HB1 ALA A 167     140.841  57.059  86.429  1.00  0.00           H
    ATOM   1374  HB2 ALA A 167     139.168  56.809  86.907  1.00  0.00           H
    ATOM   1375  HB3 ALA A 167     140.290  57.660  87.960  1.00  0.00           H
    ATOM   1376  N   ARG A 168     142.618  54.121  87.967  1.00 99.90           N
    ATOM   1377  CA  ARG A 168     144.046  53.845  88.113  1.00 99.90           C
    ATOM   1378  C   ARG A 168     144.746  54.054  86.779  1.00 99.90           C
    ATOM   1379  O   ARG A 168     145.123  53.106  86.093  1.00 99.90           O
    ATOM   1380  CB  ARG A 168     144.201  52.433  88.629  1.00 99.90           C
    ATOM   1381  CG  ARG A 168     145.509  52.103  89.348  1.00 99.90           C
    ATOM   1382  CD  ARG A 168     146.616  51.447  88.528  1.00 99.90           C
    ATOM   1383  NE  ARG A 168     147.688  52.375  88.162  1.00 99.90           N
    ATOM   1384  CZ  ARG A 168     148.372  52.318  87.013  1.00 99.90           C
    ATOM   1385  NH1 ARG A 168     147.994  51.483  86.039  1.00 99.90           N
    ATOM   1386  NH2 ARG A 168     149.443  53.101  86.849  1.00 99.90           N1+
    ATOM   1387  H   ARG A 168     142.036  53.354  87.662  1.00  0.00           H
    ATOM   1388  HA  ARG A 168     144.508  54.530  88.823  1.00  0.00           H
    ATOM   1389  HB2 ARG A 168     143.400  52.280  89.346  1.00  0.00           H
    ATOM   1390  HB3 ARG A 168     144.003  51.739  87.821  1.00  0.00           H
    ATOM   1391  HG2 ARG A 168     145.856  52.954  89.934  1.00  0.00           H
    ATOM   1392  HG3 ARG A 168     145.210  51.326  90.036  1.00  0.00           H
    ATOM   1393  HD2 ARG A 168     146.983  50.508  88.944  1.00  0.00           H
    ATOM   1394  HD3 ARG A 168     146.139  51.165  87.593  1.00  0.00           H
    ATOM   1395  HE  ARG A 168     147.961  53.058  88.859  1.00  0.00           H
    ATOM   1396 HH11 ARG A 168     147.158  50.929  86.152  1.00  0.00           H
    ATOM   1397 HH12 ARG A 168     148.510  51.431  85.175  1.00  0.00           H
    ATOM   1398 HH21 ARG A 168     149.690  53.771  87.564  1.00  0.00           H
    ATOM   1399 HH22 ARG A 168     149.994  53.064  86.005  1.00  0.00           H
    ATOM   1400  N   ILE A 169     144.853  55.336  86.438  1.00 99.90           N
    ATOM   1401  CA  ILE A 169     145.339  55.842  85.173  1.00 99.90           C
    ATOM   1402  C   ILE A 169     146.836  55.450  84.994  1.00 99.90           C
    ATOM   1403  O   ILE A 169     147.642  55.717  85.882  1.00 99.90           O
    ATOM   1404  CB  ILE A 169     145.043  57.385  85.118  1.00 73.17           C
    ATOM   1405  CG1 ILE A 169     143.587  57.678  85.587  1.00 73.17           C
    ATOM   1406  CG2 ILE A 169     145.320  58.044  83.757  1.00 73.17           C
    ATOM   1407  CD1 ILE A 169     142.979  59.053  85.289  1.00 73.17           C
    ATOM   1408  H   ILE A 169     144.511  56.037  87.082  1.00  0.00           H
    ATOM   1409  HA  ILE A 169     144.768  55.313  84.422  1.00  0.00           H
    ATOM   1410  HB  ILE A 169     145.716  57.866  85.831  1.00  0.00           H
    ATOM   1411 HG12 ILE A 169     142.932  56.912  85.193  1.00  0.00           H
    ATOM   1412 HG13 ILE A 169     143.561  57.580  86.665  1.00  0.00           H
    ATOM   1413 HG21 ILE A 169     145.358  59.129  83.851  1.00  0.00           H
    ATOM   1414 HG22 ILE A 169     146.281  57.733  83.349  1.00  0.00           H
    ATOM   1415 HG23 ILE A 169     144.539  57.820  83.035  1.00  0.00           H
    ATOM   1416 HD11 ILE A 169     142.066  59.207  85.866  1.00  0.00           H
    ATOM   1417 HD12 ILE A 169     143.671  59.855  85.549  1.00  0.00           H
    ATOM   1418 HD13 ILE A 169     142.722  59.155  84.234  1.00  0.00           H
    ATOM   1419  N   TYR A 170     147.135  54.703  83.912  1.00 99.90           N
    ATOM   1420  CA  TYR A 170     148.392  54.004  83.591  1.00 99.90           C
    ATOM   1421  C   TYR A 170     149.330  54.785  82.656  1.00 99.90           C
    ATOM   1422  O   TYR A 170     148.918  55.777  82.068  1.00 99.90           O
    ATOM   1423  CB  TYR A 170     148.024  52.620  82.992  1.00 99.90           C
    ATOM   1424  CG  TYR A 170     149.119  51.720  82.421  1.00 99.90           C
    ATOM   1425  CD1 TYR A 170     149.918  50.919  83.265  1.00 99.90           C
    ATOM   1426  CD2 TYR A 170     149.305  51.630  81.023  1.00 99.90           C
    ATOM   1427  CE1 TYR A 170     150.816  49.982  82.717  1.00 99.90           C
    ATOM   1428  CE2 TYR A 170     150.165  50.662  80.474  1.00 99.90           C
    ATOM   1429  CZ  TYR A 170     150.921  49.833  81.320  1.00 99.90           C
    ATOM   1430  OH  TYR A 170     151.760  48.902  80.783  1.00 99.90           O
    ATOM   1431  H   TYR A 170     146.407  54.596  83.216  1.00  0.00           H
    ATOM   1432  HA  TYR A 170     148.943  53.848  84.512  1.00  0.00           H
    ATOM   1433  HB2 TYR A 170     147.438  52.039  83.696  1.00  0.00           H
    ATOM   1434  HB3 TYR A 170     147.327  52.776  82.187  1.00  0.00           H
    ATOM   1435  HD1 TYR A 170     149.839  50.990  84.335  1.00  0.00           H
    ATOM   1436  HD2 TYR A 170     148.737  52.250  80.351  1.00  0.00           H
    ATOM   1437  HE1 TYR A 170     151.407  49.362  83.375  1.00  0.00           H
    ATOM   1438  HE2 TYR A 170     150.252  50.563  79.403  1.00  0.00           H
    ATOM   1439  HH  TYR A 170     152.289  48.449  81.444  1.00  0.00           H
    ATOM   1440  N   SER A 171     150.569  54.270  82.553  1.00 99.90           N
    ATOM   1441  CA  SER A 171     151.683  54.678  81.701  1.00 99.90           C
    ATOM   1442  C   SER A 171     152.298  53.443  80.993  1.00 99.90           C
    ATOM   1443  O   SER A 171     152.603  52.468  81.676  1.00 99.90           O
    ATOM   1444  CB  SER A 171     152.724  55.394  82.574  1.00 99.90           C
    ATOM   1445  OG  SER A 171     153.182  54.599  83.645  1.00 99.90           O
    ATOM   1446  H   SER A 171     150.773  53.441  83.093  1.00  0.00           H
    ATOM   1447  HA  SER A 171     151.342  55.408  80.984  1.00  0.00           H
    ATOM   1448  HB2 SER A 171     153.608  55.584  81.978  1.00  0.00           H
    ATOM   1449  HB3 SER A 171     152.350  56.347  82.949  1.00  0.00           H
    ATOM   1450  HG  SER A 171     153.677  53.858  83.285  1.00  0.00           H
    ATOM   1451  N   PHE A 172     152.492  53.490  79.662  1.00 99.90           N
    ATOM   1452  CA  PHE A 172     152.970  52.365  78.825  1.00 99.90           C
    ATOM   1453  C   PHE A 172     154.460  51.956  78.908  1.00 99.90           C
    ATOM   1454  O   PHE A 172     155.278  52.682  79.469  1.00 99.90           O
    ATOM   1455  CB  PHE A 172     152.545  52.596  77.359  1.00 99.90           C
    ATOM   1456  CG  PHE A 172     151.154  52.117  77.035  1.00 99.90           C
    ATOM   1457  CD1 PHE A 172     150.039  52.793  77.549  1.00 99.90           C
    ATOM   1458  CD2 PHE A 172     150.960  50.908  76.346  1.00 99.90           C
    ATOM   1459  CE1 PHE A 172     148.765  52.328  77.337  1.00 99.90           C
    ATOM   1460  CE2 PHE A 172     149.675  50.435  76.159  1.00 99.90           C
    ATOM   1461  CZ  PHE A 172     148.593  51.139  76.659  1.00 99.90           C
    ATOM   1462  H   PHE A 172     152.204  54.321  79.165  1.00  0.00           H
    ATOM   1463  HA  PHE A 172     152.444  51.476  79.172  1.00  0.00           H
    ATOM   1464  HB2 PHE A 172     152.564  53.654  77.141  1.00  0.00           H
    ATOM   1465  HB3 PHE A 172     153.232  52.142  76.643  1.00  0.00           H
    ATOM   1466  HD1 PHE A 172     150.163  53.637  78.171  1.00  0.00           H
    ATOM   1467  HD2 PHE A 172     151.802  50.342  75.974  1.00  0.00           H
    ATOM   1468  HE1 PHE A 172     147.925  52.862  77.739  1.00  0.00           H
    ATOM   1469  HE2 PHE A 172     149.532  49.501  75.642  1.00  0.00           H
    ATOM   1470  HZ  PHE A 172     147.619  50.720  76.560  1.00  0.00           H
    ATOM   1471  N   GLN A 173     154.737  50.782  78.297  1.00 99.90           N
    ATOM   1472  CA  GLN A 173     155.990  50.035  78.058  1.00 99.90           C
    ATOM   1473  C   GLN A 173     157.337  50.680  78.451  1.00 99.90           C
    ATOM   1474  O   GLN A 173     158.139  50.955  77.561  1.00 99.90           O
    ATOM   1475  CB  GLN A 173     155.978  49.549  76.571  1.00 99.90           C
    ATOM   1476  CG  GLN A 173     155.492  48.104  76.357  1.00 99.90           C
    ATOM   1477  CD  GLN A 173     156.499  47.029  76.787  1.00 99.90           C
    ATOM   1478  OE1 GLN A 173     156.161  46.149  77.575  1.00 99.90           O
    ATOM   1479  NE2 GLN A 173     157.724  47.073  76.260  1.00 99.90           N
    ATOM   1480  H   GLN A 173     153.938  50.311  77.900  1.00  0.00           H
    ATOM   1481  HA  GLN A 173     155.913  49.155  78.697  1.00  0.00           H
    ATOM   1482  HB2 GLN A 173     155.340  50.213  75.986  1.00  0.00           H
    ATOM   1483  HB3 GLN A 173     156.942  49.628  76.070  1.00  0.00           H
    ATOM   1484  HG2 GLN A 173     154.545  47.948  76.876  1.00  0.00           H
    ATOM   1485  HG3 GLN A 173     155.286  47.949  75.297  1.00  0.00           H
    ATOM   1486 HE21 GLN A 173     157.978  47.803  75.610  1.00  0.00           H
    ATOM   1487 HE22 GLN A 173     158.408  46.375  76.512  1.00  0.00           H
    ATOM   1488  N   MET A 174     157.589  50.870  79.763  1.00 99.90           N
    ATOM   1489  CA  MET A 174     158.833  51.418  80.336  1.00 99.90           C
    ATOM   1490  C   MET A 174     159.239  52.772  79.708  1.00 99.90           C
    ATOM   1491  O   MET A 174     158.839  53.820  80.210  1.00 99.90           O
    ATOM   1492  CB  MET A 174     159.978  50.367  80.269  1.00  0.00           C
    ATOM   1493  CG  MET A 174     160.036  49.398  81.461  1.00  0.00           C
    ATOM   1494  SD  MET A 174     160.430  50.137  83.077  1.00  0.00           S
    ATOM   1495  CE  MET A 174     162.036  50.913  82.737  1.00  0.00           C
    ATOM   1496  H   MET A 174     156.872  50.632  80.433  1.00  0.00           H
    ATOM   1497  HA  MET A 174     158.619  51.636  81.383  1.00  0.00           H
    ATOM   1498  HB2 MET A 174     159.912  49.788  79.348  1.00  0.00           H
    ATOM   1499  HB3 MET A 174     160.948  50.859  80.208  1.00  0.00           H
    ATOM   1500  HG2 MET A 174     159.082  48.878  81.554  1.00  0.00           H
    ATOM   1501  HG3 MET A 174     160.782  48.629  81.260  1.00  0.00           H
    ATOM   1502  HE1 MET A 174     162.471  51.282  83.666  1.00  0.00           H
    ATOM   1503  HE2 MET A 174     162.725  50.194  82.296  1.00  0.00           H
    ATOM   1504  HE3 MET A 174     161.925  51.759  82.060  1.00  0.00           H
    ATOM   1505  N   THR A 177     156.032  55.438  77.880  1.00 99.90           N1+
    ATOM   1506  CA  THR A 177     155.022  55.916  78.839  1.00 99.90           C
    ATOM   1507  C   THR A 177     153.909  56.786  78.202  1.00 99.90           C
    ATOM   1508  O   THR A 177     153.951  58.013  78.255  1.00 99.90           O
    ATOM   1509  CB  THR A 177     155.730  56.652  80.022  1.00 99.90           C
    ATOM   1510  OG1 THR A 177     156.371  55.685  80.836  1.00 99.90           O
    ATOM   1511  CG2 THR A 177     154.877  57.564  80.936  1.00 99.90           C
    ATOM   1512  H   THR A 177     156.986  55.644  78.137  1.00  0.00           H
    ATOM   1513  HA  THR A 177     154.526  55.045  79.256  1.00  0.00           H
    ATOM   1514  HB  THR A 177     156.534  57.268  79.616  1.00  0.00           H
    ATOM   1515 HG21 THR A 177     155.190  57.510  81.979  1.00  0.00           H
    ATOM   1516 HG22 THR A 177     154.994  58.606  80.635  1.00  0.00           H
    ATOM   1517 HG23 THR A 177     153.811  57.357  80.887  1.00  0.00           H
    ATOM   1518  HG1 THR A 177     157.030  55.215  80.318  1.00  0.00           H
    ATOM   1519  N   SER A 178     152.912  56.113  77.610  1.00 99.90           N
    ATOM   1520  CA  SER A 178     151.688  56.712  77.085  1.00 99.90           C
    ATOM   1521  C   SER A 178     150.663  56.680  78.222  1.00 99.90           C
    ATOM   1522  O   SER A 178     150.528  55.650  78.862  1.00 99.90           O
    ATOM   1523  CB  SER A 178     151.184  55.895  75.874  1.00 99.90           C
    ATOM   1524  OG  SER A 178     152.134  55.894  74.827  1.00 99.90           O
    ATOM   1525  H   SER A 178     152.980  55.109  77.537  1.00  0.00           H
    ATOM   1526  HA  SER A 178     151.863  57.743  76.774  1.00  0.00           H
    ATOM   1527  HB2 SER A 178     150.952  54.867  76.138  1.00  0.00           H
    ATOM   1528  HB3 SER A 178     150.260  56.328  75.490  1.00  0.00           H
    ATOM   1529  HG  SER A 178     152.927  55.443  75.130  1.00  0.00           H
    ATOM   1530  N   VAL A 179     149.956  57.781  78.491  1.00 99.90           N
    ATOM   1531  CA  VAL A 179     148.927  57.790  79.535  1.00 99.90           C
    ATOM   1532  C   VAL A 179     147.629  57.077  79.072  1.00 99.90           C
    ATOM   1533  O   VAL A 179     147.435  56.905  77.868  1.00 99.90           O
    ATOM   1534  CB  VAL A 179     148.644  59.230  80.032  1.00 99.90           C
    ATOM   1535  CG1 VAL A 179     148.112  60.154  78.927  1.00 99.90           C
    ATOM   1536  CG2 VAL A 179     147.730  59.291  81.273  1.00 99.90           C
    ATOM   1537  H   VAL A 179     150.078  58.625  77.951  1.00  0.00           H
    ATOM   1538  HA  VAL A 179     149.353  57.270  80.381  1.00  0.00           H
    ATOM   1539  HB  VAL A 179     149.606  59.638  80.347  1.00  0.00           H
    ATOM   1540 HG11 VAL A 179     147.995  61.173  79.297  1.00  0.00           H
    ATOM   1541 HG12 VAL A 179     148.790  60.200  78.075  1.00  0.00           H
    ATOM   1542 HG13 VAL A 179     147.138  59.830  78.559  1.00  0.00           H
    ATOM   1543 HG21 VAL A 179     147.745  60.290  81.710  1.00  0.00           H
    ATOM   1544 HG22 VAL A 179     146.690  59.068  81.039  1.00  0.00           H
    ATOM   1545 HG23 VAL A 179     148.068  58.598  82.043  1.00  0.00           H
    ATOM   1546  N   VAL A 180     146.772  56.670  80.020  1.00 99.90           N
    ATOM   1547  CA  VAL A 180     145.474  56.032  79.772  1.00 99.90           C
    ATOM   1548  C   VAL A 180     144.742  55.822  81.098  1.00 99.90           C
    ATOM   1549  O   VAL A 180     145.417  55.602  82.085  1.00 99.90           O
    ATOM   1550  CB  VAL A 180     145.623  54.638  79.107  1.00 99.90           C
    ATOM   1551  CG1 VAL A 180     146.485  53.651  79.896  1.00 99.90           C
    ATOM   1552  CG2 VAL A 180     144.269  53.971  78.840  1.00 99.90           C
    ATOM   1553  H   VAL A 180     147.013  56.800  80.993  1.00  0.00           H
    ATOM   1554  HA  VAL A 180     144.880  56.692  79.137  1.00  0.00           H
    ATOM   1555  HB  VAL A 180     146.102  54.811  78.152  1.00  0.00           H
    ATOM   1556 HG11 VAL A 180     146.604  52.712  79.361  1.00  0.00           H
    ATOM   1557 HG12 VAL A 180     147.470  54.058  80.125  1.00  0.00           H
    ATOM   1558 HG13 VAL A 180     145.971  53.390  80.816  1.00  0.00           H
    ATOM   1559 HG21 VAL A 180     144.348  53.296  78.004  1.00  0.00           H
    ATOM   1560 HG22 VAL A 180     143.905  53.390  79.681  1.00  0.00           H
    ATOM   1561 HG23 VAL A 180     143.500  54.696  78.572  1.00  0.00           H
    ATOM   1562  N   VAL A 181     143.402  55.817  81.112  1.00 99.90           N
    ATOM   1563  CA  VAL A 181     142.605  55.552  82.315  1.00 99.90           C
    ATOM   1564  C   VAL A 181     142.148  54.080  82.443  1.00 99.90           C
    ATOM   1565  O   VAL A 181     141.043  53.831  82.922  1.00 99.90           O
    ATOM   1566  CB  VAL A 181     141.418  56.547  82.431  1.00 62.69           C
    ATOM   1567  CG1 VAL A 181     140.400  56.417  81.285  1.00 62.69           C
    ATOM   1568  CG2 VAL A 181     140.690  56.503  83.792  1.00 62.69           C
    ATOM   1569  H   VAL A 181     142.886  55.983  80.260  1.00  0.00           H
    ATOM   1570  HA  VAL A 181     143.205  55.749  83.192  1.00  0.00           H
    ATOM   1571  HB  VAL A 181     141.856  57.543  82.354  1.00  0.00           H
    ATOM   1572 HG11 VAL A 181     139.627  57.182  81.372  1.00  0.00           H
    ATOM   1573 HG12 VAL A 181     140.874  56.553  80.313  1.00  0.00           H
    ATOM   1574 HG13 VAL A 181     139.895  55.451  81.273  1.00  0.00           H
    ATOM   1575 HG21 VAL A 181     140.563  57.508  84.194  1.00  0.00           H
    ATOM   1576 HG22 VAL A 181     139.693  56.072  83.699  1.00  0.00           H
    ATOM   1577 HG23 VAL A 181     141.209  55.893  84.529  1.00  0.00           H
    ATOM   1578  N   THR A 182     143.006  53.121  82.037  1.00 90.28           N
    ATOM   1579  CA  THR A 182     142.778  51.665  82.076  1.00 90.28           C
    ATOM   1580  C   THR A 182     141.357  51.275  81.616  1.00 90.28           C
    ATOM   1581  O   THR A 182     140.458  51.028  82.419  1.00 90.28           O
    ATOM   1582  CB  THR A 182     143.057  51.051  83.474  1.00 99.90           C
    ATOM   1583  OG1 THR A 182     142.371  51.728  84.514  1.00 99.90           O
    ATOM   1584  CG2 THR A 182     144.552  51.057  83.801  1.00 99.90           C
    ATOM   1585  H   THR A 182     143.901  53.409  81.669  1.00  0.00           H
    ATOM   1586  HA  THR A 182     143.482  51.225  81.369  1.00  0.00           H
    ATOM   1587  HB  THR A 182     142.724  50.012  83.494  1.00  0.00           H
    ATOM   1588 HG21 THR A 182     144.745  50.667  84.801  1.00  0.00           H
    ATOM   1589 HG22 THR A 182     145.100  50.437  83.097  1.00  0.00           H
    ATOM   1590 HG23 THR A 182     144.968  52.063  83.741  1.00  0.00           H
    ATOM   1591  HG1 THR A 182     141.442  51.782  84.267  1.00  0.00           H
    ATOM   1592  N   LEU A 183     141.196  51.297  80.295  1.00 99.90           N
    ATOM   1593  CA  LEU A 183     139.963  51.013  79.572  1.00 99.90           C
    ATOM   1594  C   LEU A 183     140.067  49.650  78.869  1.00 99.90           C
    ATOM   1595  O   LEU A 183     139.378  49.392  77.891  1.00 99.90           O
    ATOM   1596  CB  LEU A 183     139.687  52.224  78.652  1.00 77.66           C
    ATOM   1597  CG  LEU A 183     140.457  52.348  77.315  1.00 77.66           C
    ATOM   1598  CD1 LEU A 183     140.240  53.757  76.752  1.00 77.66           C
    ATOM   1599  CD2 LEU A 183     141.963  52.032  77.349  1.00 77.66           C
    ATOM   1600  H   LEU A 183     141.986  51.537  79.713  1.00  0.00           H
    ATOM   1601  HA  LEU A 183     139.132  50.940  80.276  1.00  0.00           H
    ATOM   1602  HB2 LEU A 183     138.625  52.250  78.421  1.00  0.00           H
    ATOM   1603  HB3 LEU A 183     139.854  53.130  79.238  1.00  0.00           H
    ATOM   1604  HG  LEU A 183     140.002  51.654  76.606  1.00  0.00           H
    ATOM   1605 HD11 LEU A 183     140.530  53.800  75.708  1.00  0.00           H
    ATOM   1606 HD12 LEU A 183     139.195  54.063  76.817  1.00  0.00           H
    ATOM   1607 HD13 LEU A 183     140.838  54.497  77.286  1.00  0.00           H
    ATOM   1608 HD21 LEU A 183     142.521  52.702  76.694  1.00  0.00           H
    ATOM   1609 HD22 LEU A 183     142.407  52.113  78.332  1.00  0.00           H
    ATOM   1610 HD23 LEU A 183     142.146  51.025  76.978  1.00  0.00           H
    ATOM   1611  N   TRP A 184     140.943  48.794  79.402  1.00 69.76           N
    ATOM   1612  CA  TRP A 184     141.374  47.510  78.855  1.00 69.76           C
    ATOM   1613  C   TRP A 184     140.323  46.417  78.891  1.00 69.76           C
    ATOM   1614  O   TRP A 184     140.403  45.440  78.149  1.00 69.76           O
    ATOM   1615  CB  TRP A 184     142.584  47.075  79.661  1.00 72.21           C
    ATOM   1616  CG  TRP A 184     143.676  48.081  79.703  1.00 72.21           C
    ATOM   1617  CD1 TRP A 184     143.855  49.137  78.873  1.00 72.21           C
    ATOM   1618  CD2 TRP A 184     144.769  48.115  80.641  1.00 72.21           C
    ATOM   1619  NE1 TRP A 184     144.973  49.834  79.251  1.00 72.21           N
    ATOM   1620  CE2 TRP A 184     145.582  49.236  80.327  1.00 72.21           C
    ATOM   1621  CE3 TRP A 184     145.152  47.309  81.734  1.00 72.21           C
    ATOM   1622  CZ2 TRP A 184     146.732  49.536  81.050  1.00 72.21           C
    ATOM   1623  CZ3 TRP A 184     146.304  47.614  82.479  1.00 72.21           C
    ATOM   1624  CH2 TRP A 184     147.103  48.718  82.130  1.00 72.21           C
    ATOM   1625  H   TRP A 184     141.453  49.095  80.220  1.00  0.00           H
    ATOM   1626  HA  TRP A 184     141.681  47.659  77.834  1.00  0.00           H
    ATOM   1627  HB2 TRP A 184     142.282  46.912  80.694  1.00  0.00           H
    ATOM   1628  HB3 TRP A 184     142.976  46.115  79.335  1.00  0.00           H
    ATOM   1629  HD1 TRP A 184     143.220  49.384  78.037  1.00  0.00           H
    ATOM   1630  HE1 TRP A 184     145.298  50.650  78.744  1.00  0.00           H
    ATOM   1631  HE3 TRP A 184     144.554  46.450  82.002  1.00  0.00           H
    ATOM   1632  HZ2 TRP A 184     147.289  50.399  80.758  1.00  0.00           H
    ATOM   1633  HZ3 TRP A 184     146.580  46.994  83.320  1.00  0.00           H
    ATOM   1634  HH2 TRP A 184     147.992  48.940  82.700  1.00  0.00           H
    ATOM   1635  N   TYR A 185     139.367  46.620  79.788  1.00 55.74           N
    ATOM   1636  CA  TYR A 185     138.261  45.727  80.042  1.00 55.74           C
    ATOM   1637  C   TYR A 185     137.018  46.248  79.326  1.00 55.74           C
    ATOM   1638  O   TYR A 185     135.984  45.596  79.414  1.00 55.74           O
    ATOM   1639  CB  TYR A 185     138.085  45.628  81.568  1.00 39.86           C
    ATOM   1640  CG  TYR A 185     139.408  45.358  82.259  1.00 39.86           C
    ATOM   1641  CD1 TYR A 185     140.227  46.433  82.671  1.00 39.86           C
    ATOM   1642  CD2 TYR A 185     139.868  44.038  82.387  1.00 39.86           C
    ATOM   1643  CE1 TYR A 185     141.516  46.191  83.177  1.00 39.86           C
    ATOM   1644  CE2 TYR A 185     141.144  43.797  82.917  1.00 39.86           C
    ATOM   1645  CZ  TYR A 185     141.981  44.871  83.300  1.00 39.86           C
    ATOM   1646  OH  TYR A 185     143.221  44.652  83.826  1.00 39.86           O
    ATOM   1647  H   TYR A 185     139.404  47.451  80.360  1.00  0.00           H
    ATOM   1648  HA  TYR A 185     138.472  44.728  79.656  1.00  0.00           H
    ATOM   1649  HB2 TYR A 185     137.674  46.557  81.967  1.00  0.00           H
    ATOM   1650  HB3 TYR A 185     137.371  44.843  81.821  1.00  0.00           H
    ATOM   1651  HD1 TYR A 185     139.883  47.451  82.565  1.00  0.00           H
    ATOM   1652  HD2 TYR A 185     139.249  43.210  82.077  1.00  0.00           H
    ATOM   1653  HE1 TYR A 185     142.142  47.018  83.477  1.00  0.00           H
    ATOM   1654  HE2 TYR A 185     141.421  42.769  83.049  1.00  0.00           H
    ATOM   1655  HH  TYR A 185     143.456  43.714  83.840  1.00  0.00           H
    ATOM   1656  N   ARG A 186     137.128  47.386  78.614  1.00 83.81           N
    ATOM   1657  CA  ARG A 186     136.041  47.931  77.821  1.00 83.81           C
    ATOM   1658  C   ARG A 186     135.723  47.034  76.645  1.00 83.81           C
    ATOM   1659  O   ARG A 186     136.619  46.553  75.950  1.00 83.81           O
    ATOM   1660  CB  ARG A 186     136.335  49.358  77.334  1.00 90.66           C
    ATOM   1661  CG  ARG A 186     136.375  50.378  78.474  1.00 90.66           C
    ATOM   1662  CD  ARG A 186     135.014  50.533  79.171  1.00 90.66           C
    ATOM   1663  NE  ARG A 186     135.067  51.511  80.254  1.00 90.66           N
    ATOM   1664  CZ  ARG A 186     135.144  52.840  80.104  1.00 90.66           C
    ATOM   1665  NH1 ARG A 186     135.174  53.413  78.892  1.00 90.66           N
    ATOM   1666  NH2 ARG A 186     135.188  53.607  81.199  1.00 90.66           N1+
    ATOM   1667  H   ARG A 186     138.017  47.863  78.553  1.00  0.00           H
    ATOM   1668  HA  ARG A 186     135.160  47.939  78.462  1.00  0.00           H
    ATOM   1669  HB2 ARG A 186     137.252  49.378  76.749  1.00  0.00           H
    ATOM   1670  HB3 ARG A 186     135.563  49.676  76.639  1.00  0.00           H
    ATOM   1671  HG2 ARG A 186     137.128  50.090  79.206  1.00  0.00           H
    ATOM   1672  HG3 ARG A 186     136.680  51.340  78.069  1.00  0.00           H
    ATOM   1673  HD2 ARG A 186     134.229  50.805  78.467  1.00  0.00           H
    ATOM   1674  HD3 ARG A 186     134.711  49.604  79.650  1.00  0.00           H
    ATOM   1675  HE  ARG A 186     135.098  51.127  81.189  1.00  0.00           H
    ATOM   1676 HH11 ARG A 186     135.145  52.839  78.061  1.00  0.00           H
    ATOM   1677 HH12 ARG A 186     135.268  54.414  78.793  1.00  0.00           H
    ATOM   1678 HH21 ARG A 186     135.189  53.187  82.118  1.00  0.00           H
    ATOM   1679 HH22 ARG A 186     135.268  54.611  81.120  1.00  0.00           H
    ATOM   1680  N   ILE A 208     138.998  39.728  81.959  1.00 43.11           N1+
    ATOM   1681  CA  ILE A 208     139.389  39.308  83.292  1.00 43.11           C
    ATOM   1682  C   ILE A 208     139.709  37.810  83.344  1.00 43.11           C
    ATOM   1683  O   ILE A 208     140.717  37.465  83.941  1.00 43.11           O
    ATOM   1684  CB  ILE A 208     138.307  39.626  84.355  1.00 38.88           C
    ATOM   1685  CG1 ILE A 208     138.145  41.149  84.514  1.00 38.88           C
    ATOM   1686  CG2 ILE A 208     138.608  38.993  85.729  1.00 38.88           C
    ATOM   1687  CD1 ILE A 208     136.860  41.552  85.236  1.00 38.88           C
    ATOM   1688  H   ILE A 208     138.048  40.070  81.866  1.00  0.00           H
    ATOM   1689  HA  ILE A 208     140.286  39.870  83.554  1.00  0.00           H
    ATOM   1690  HB  ILE A 208     137.353  39.230  84.006  1.00  0.00           H
    ATOM   1691 HG12 ILE A 208     138.995  41.550  85.053  1.00  0.00           H
    ATOM   1692 HG13 ILE A 208     138.160  41.646  83.546  1.00  0.00           H
    ATOM   1693 HG21 ILE A 208     138.012  39.423  86.530  1.00  0.00           H
    ATOM   1694 HG22 ILE A 208     138.436  37.916  85.745  1.00  0.00           H
    ATOM   1695 HG23 ILE A 208     139.650  39.172  85.986  1.00  0.00           H
    ATOM   1696 HD11 ILE A 208     137.053  42.294  86.011  1.00  0.00           H
    ATOM   1697 HD12 ILE A 208     136.173  41.980  84.513  1.00  0.00           H
    ATOM   1698 HD13 ILE A 208     136.344  40.711  85.698  1.00  0.00           H
    ATOM   1699  N   GLU A 211     142.870  37.171  81.728  1.00 45.63           N1+
    ATOM   1700  CA  GLU A 211     143.747  37.761  82.724  1.00 45.63           C
    ATOM   1701  C   GLU A 211     144.033  36.908  83.976  1.00 45.63           C
    ATOM   1702  O   GLU A 211     145.110  37.062  84.544  1.00 45.63           O
    ATOM   1703  CB  GLU A 211     143.328  39.194  83.031  1.00 97.40           C
    ATOM   1704  CG  GLU A 211     144.340  39.965  83.896  1.00 97.40           C
    ATOM   1705  CD  GLU A 211     143.992  41.440  83.974  1.00 97.40           C
    ATOM   1706  OE1 GLU A 211     143.540  41.971  82.942  1.00 97.40           O
    ATOM   1707  OE2 GLU A 211     144.209  42.031  85.050  1.00 97.40           O1-
    ATOM   1708  H   GLU A 211     142.064  37.727  81.477  1.00  0.00           H
    ATOM   1709  HA  GLU A 211     144.654  37.928  82.180  1.00  0.00           H
    ATOM   1710  HB2 GLU A 211     143.191  39.705  82.077  1.00  0.00           H
    ATOM   1711  HB3 GLU A 211     142.368  39.218  83.532  1.00  0.00           H
    ATOM   1712  HG2 GLU A 211     144.371  39.564  84.909  1.00  0.00           H
    ATOM   1713  HG3 GLU A 211     145.345  39.877  83.482  1.00  0.00           H
    ATOM   1714  N   ARG A 214     146.623  34.690  82.857  1.00 36.83           N1+
    ATOM   1715  CA  ARG A 214     147.674  35.571  82.389  1.00 36.83           C
    ATOM   1716  C   ARG A 214     148.500  36.317  83.443  1.00 36.83           C
    ATOM   1717  O   ARG A 214     149.714  36.355  83.263  1.00 36.83           O
    ATOM   1718  CB  ARG A 214     147.119  36.601  81.427  1.00 61.66           C
    ATOM   1719  CG  ARG A 214     146.671  36.090  80.049  1.00 61.66           C
    ATOM   1720  CD  ARG A 214     146.553  37.240  79.048  1.00 61.66           C
    ATOM   1721  NE  ARG A 214     147.884  37.840  78.933  1.00 61.66           N
    ATOM   1722  CZ  ARG A 214     148.353  38.693  78.029  1.00 61.66           C
    ATOM   1723  NH1 ARG A 214     147.633  39.064  76.976  1.00 61.66           N
    ATOM   1724  NH2 ARG A 214     149.599  39.143  78.158  1.00 61.66           N1+
    ATOM   1725  H   ARG A 214     145.691  34.883  82.514  1.00  0.00           H
    ATOM   1726  HA  ARG A 214     148.372  34.942  81.832  1.00  0.00           H
    ATOM   1727  HB2 ARG A 214     146.387  37.192  81.927  1.00  0.00           H
    ATOM   1728  HB3 ARG A 214     147.946  37.288  81.269  1.00  0.00           H
    ATOM   1729  HG2 ARG A 214     147.489  35.457  79.706  1.00  0.00           H
    ATOM   1730  HG3 ARG A 214     145.815  35.429  80.021  1.00  0.00           H
    ATOM   1731  HD2 ARG A 214     146.138  36.899  78.100  1.00  0.00           H
    ATOM   1732  HD3 ARG A 214     145.896  38.021  79.434  1.00  0.00           H
    ATOM   1733  HE  ARG A 214     148.489  37.624  79.711  1.00  0.00           H
    ATOM   1734 HH11 ARG A 214     146.713  38.668  76.845  1.00  0.00           H
    ATOM   1735 HH12 ARG A 214     147.994  39.714  76.295  1.00  0.00           H
    ATOM   1736 HH21 ARG A 214     150.174  38.826  78.925  1.00  0.00           H
    ATOM   1737 HH22 ARG A 214     150.006  39.718  77.437  1.00  0.00           H
    ATOM   1738  N   ARG A 215     147.872  36.984  84.431  1.00 44.76           N
    ATOM   1739  CA  ARG A 215     148.543  37.905  85.370  1.00 44.76           C
    ATOM   1740  C   ARG A 215     149.249  39.062  84.619  1.00 44.76           C
    ATOM   1741  O   ARG A 215     150.337  39.502  84.990  1.00 44.76           O
    ATOM   1742  CB  ARG A 215     149.504  37.113  86.288  1.00 99.90           C
    ATOM   1743  CG  ARG A 215     148.828  36.009  87.130  1.00 99.90           C
    ATOM   1744  CD  ARG A 215     149.694  34.755  87.293  1.00 99.90           C
    ATOM   1745  NE  ARG A 215     151.060  35.078  87.736  1.00 99.90           N
    ATOM   1746  CZ  ARG A 215     152.171  34.388  87.418  1.00 99.90           C
    ATOM   1747  NH1 ARG A 215     152.122  33.296  86.645  1.00 99.90           N
    ATOM   1748  NH2 ARG A 215     153.360  34.798  87.875  1.00 99.90           N1+
    ATOM   1749  H   ARG A 215     146.863  36.922  84.513  1.00  0.00           H
    ATOM   1750  HA  ARG A 215     147.765  38.354  85.988  1.00  0.00           H
    ATOM   1751  HB2 ARG A 215     150.288  36.670  85.675  1.00  0.00           H
    ATOM   1752  HB3 ARG A 215     150.026  37.785  86.970  1.00  0.00           H
    ATOM   1753  HG2 ARG A 215     148.412  36.370  88.070  1.00  0.00           H
    ATOM   1754  HG3 ARG A 215     147.964  35.650  86.571  1.00  0.00           H
    ATOM   1755  HD2 ARG A 215     149.281  34.146  88.097  1.00  0.00           H
    ATOM   1756  HD3 ARG A 215     149.653  34.147  86.388  1.00  0.00           H
    ATOM   1757  HE  ARG A 215     151.143  35.858  88.374  1.00  0.00           H
    ATOM   1758 HH11 ARG A 215     151.248  32.983  86.250  1.00  0.00           H
    ATOM   1759 HH12 ARG A 215     152.964  32.764  86.466  1.00  0.00           H
    ATOM   1760 HH21 ARG A 215     153.438  35.619  88.457  1.00  0.00           H
    ATOM   1761 HH22 ARG A 215     154.210  34.297  87.631  1.00  0.00           H
    ATOM   1762  N   LYS A 216     148.608  39.477  83.523  1.00 42.64           N
    ATOM   1763  CA  LYS A 216     149.071  40.408  82.508  1.00 42.64           C
    ATOM   1764  C   LYS A 216     147.793  40.638  81.682  1.00 42.64           C
    ATOM   1765  O   LYS A 216     147.202  39.652  81.244  1.00 42.64           O
    ATOM   1766  CB  LYS A 216     150.197  39.755  81.661  1.00 84.82           C
    ATOM   1767  CG  LYS A 216     151.482  40.597  81.533  1.00 84.82           C
    ATOM   1768  CD  LYS A 216     151.400  41.896  80.712  1.00 84.82           C
    ATOM   1769  CE  LYS A 216     151.017  41.683  79.244  1.00 84.82           C
    ATOM   1770  NZ  LYS A 216     151.323  42.856  78.412  1.00 84.82           N1+
    ATOM   1771  H   LYS A 216     147.717  39.049  83.312  1.00  0.00           H
    ATOM   1772  HA  LYS A 216     149.447  41.300  83.004  1.00  0.00           H
    ATOM   1773  HB2 LYS A 216     150.502  38.814  82.121  1.00  0.00           H
    ATOM   1774  HB3 LYS A 216     149.835  39.457  80.682  1.00  0.00           H
    ATOM   1775  HG2 LYS A 216     151.833  40.842  82.537  1.00  0.00           H
    ATOM   1776  HG3 LYS A 216     152.264  39.968  81.106  1.00  0.00           H
    ATOM   1777  HD2 LYS A 216     150.711  42.590  81.187  1.00  0.00           H
    ATOM   1778  HD3 LYS A 216     152.375  42.383  80.759  1.00  0.00           H
    ATOM   1779  HE2 LYS A 216     151.567  40.837  78.833  1.00  0.00           H
    ATOM   1780  HE3 LYS A 216     149.949  41.478  79.172  1.00  0.00           H
    ATOM   1781  HZ1 LYS A 216     152.314  43.048  78.456  1.00  0.00           H
    ATOM   1782  HZ2 LYS A 216     151.059  42.676  77.455  1.00  0.00           H
    ATOM   1783  HZ3 LYS A 216     150.814  43.656  78.759  1.00  0.00           H
    ATOM   1784  N   PRO A 217     147.325  41.892  81.523  1.00 34.54           N
    ATOM   1785  CA  PRO A 217     146.087  42.170  80.769  1.00 34.54           C
    ATOM   1786  C   PRO A 217     146.088  41.653  79.329  1.00 34.54           C
    ATOM   1787  O   PRO A 217     147.142  41.664  78.689  1.00 34.54           O
    ATOM   1788  CB  PRO A 217     145.965  43.696  80.805  1.00 59.87           C
    ATOM   1789  CG  PRO A 217     146.771  44.127  82.015  1.00 59.87           C
    ATOM   1790  CD  PRO A 217     147.887  43.107  82.102  1.00 59.87           C
    ATOM   1791  HA  PRO A 217     145.269  41.703  81.308  1.00  0.00           H
    ATOM   1792  HB2 PRO A 217     146.388  44.132  79.903  1.00  0.00           H
    ATOM   1793  HB3 PRO A 217     144.928  44.029  80.868  1.00  0.00           H
    ATOM   1794  HG2 PRO A 217     147.151  45.143  81.936  1.00  0.00           H
    ATOM   1795  HG3 PRO A 217     146.143  44.062  82.905  1.00  0.00           H
    ATOM   1796  HD2 PRO A 217     148.746  43.426  81.521  1.00  0.00           H
    ATOM   1797  HD3 PRO A 217     148.187  42.999  83.145  1.00  0.00           H
    ATOM   1798  N   ARG A 220     146.037  45.926  76.524  1.00 64.52           N1+
    ATOM   1799  CA  ARG A 220     146.366  47.223  77.129  1.00 64.52           C
    ATOM   1800  C   ARG A 220     146.258  48.326  76.079  1.00 64.52           C
    ATOM   1801  O   ARG A 220     147.248  48.609  75.415  1.00 64.52           O
    ATOM   1802  CB  ARG A 220     147.762  47.292  77.785  1.00 99.90           C
    ATOM   1803  CG  ARG A 220     148.012  46.374  78.983  1.00 99.90           C
    ATOM   1804  CD  ARG A 220     148.890  45.148  78.706  1.00 99.90           C
    ATOM   1805  NE  ARG A 220     148.311  44.167  77.781  1.00 99.90           N
    ATOM   1806  CZ  ARG A 220     148.408  44.147  76.444  1.00 99.90           C
    ATOM   1807  NH1 ARG A 220     149.043  45.097  75.752  1.00 99.90           N
    ATOM   1808  NH2 ARG A 220     147.872  43.133  75.769  1.00 99.90           N1+
    ATOM   1809  H   ARG A 220     146.775  45.442  76.035  1.00  0.00           H
    ATOM   1810  HA  ARG A 220     145.664  47.442  77.927  1.00  0.00           H
    ATOM   1811  HB2 ARG A 220     148.578  47.220  77.070  1.00  0.00           H
    ATOM   1812  HB3 ARG A 220     147.843  48.308  78.176  1.00  0.00           H
    ATOM   1813  HG2 ARG A 220     148.649  46.981  79.628  1.00  0.00           H
    ATOM   1814  HG3 ARG A 220     147.132  46.187  79.597  1.00  0.00           H
    ATOM   1815  HD2 ARG A 220     149.934  45.396  78.515  1.00  0.00           H
    ATOM   1816  HD3 ARG A 220     148.896  44.587  79.639  1.00  0.00           H
    ATOM   1817  HE  ARG A 220     147.770  43.429  78.219  1.00  0.00           H
    ATOM   1818 HH11 ARG A 220     149.481  45.866  76.240  1.00  0.00           H
    ATOM   1819 HH12 ARG A 220     149.140  45.014  74.751  1.00  0.00           H
    ATOM   1820 HH21 ARG A 220     147.329  42.426  76.247  1.00  0.00           H
    ATOM   1821 HH22 ARG A 220     147.941  43.083  74.762  1.00  0.00           H
  extra_rotamers: ""
```
