## Supplementary material for "DISRUPTOR: Computational identification of oncogenic mutants disrupting protein interactions": exemplary input files: ERK2_2yfs.79-107.yaml

```
---
osprey_version: 3.1
design_name: ERK2 and DUSP6
protein:
  residue_configurations:
  - mutability:
    - ALA
    - ARG
    - ASN
    - ASP
    - CYS
    - GLU
    - GLN
    - GLY
    - HIE
    - HID
    - HIP
    - ILE
    - LEU
    - LYS
    - MET
    - PHE
    - PRO
    - SER
    - THR
    - TRP
    - TYR
    - VAL
    flexibility:
      is_flexible: true
      include_structure_rotamer: true
      use_continuous: true
    identity:
      chain: A
      res_num: 79
      aa_type: GLU
  - mutability:
    - ALA
    - ARG
    - ASN
    - ASP
    - CYS
    - GLU
    - GLN
    - GLY
    - HIE
    - HID
    - HIP
    - ILE
    - LEU
    - LYS
    - MET
    - PHE
    - PRO
    - SER
    - THR
    - TRP
    - TYR
    - VAL
    flexibility:
      is_flexible: true
      include_structure_rotamer: true
      use_continuous: true
    identity:
      chain: A
      res_num: 107
      aa_type: GLU
  extra_templates: ""
  extra_templates_coordinates: ""
  coordinates: |
    ATOM      1  N   TYR A  28      20.129 -17.515 -46.205  1.00 33.11           N1+
    ATOM      2  CA  TYR A  28      20.898 -17.308 -44.979  1.00 32.58           C
    ATOM      3  C   TYR A  28      21.923 -16.166 -45.141  1.00 31.73           C
    ATOM      4  O   TYR A  28      22.670 -16.156 -46.120  1.00 30.45           O
    ATOM      5  CB  TYR A  28      21.582 -18.657 -44.679  1.00 33.81           C
    ATOM      6  CG  TYR A  28      22.490 -18.696 -43.469  1.00 35.67           C
    ATOM      7  CD1 TYR A  28      21.986 -18.359 -42.198  1.00 36.31           C
    ATOM      8  CD2 TYR A  28      23.838 -19.083 -43.611  1.00 37.26           C
    ATOM      9  CE1 TYR A  28      22.826 -18.429 -41.074  1.00 36.02           C
    ATOM     10  CE2 TYR A  28      24.678 -19.145 -42.485  1.00 35.89           C
    ATOM     11  CZ  TYR A  28      24.169 -18.826 -41.213  1.00 35.76           C
    ATOM     12  OH  TYR A  28      24.978 -18.907 -40.120  1.00 37.87           O
    ATOM     13  H   TYR A  28      20.239 -18.414 -46.651  1.00  0.00           H
    ATOM     14  HA  TYR A  28      20.208 -17.062 -44.171  1.00  0.00           H
    ATOM     15  HB2 TYR A  28      20.819 -19.423 -44.532  1.00  0.00           H
    ATOM     16  HB3 TYR A  28      22.154 -18.977 -45.553  1.00  0.00           H
    ATOM     17  HD1 TYR A  28      20.957 -18.054 -42.078  1.00  0.00           H
    ATOM     18  HD2 TYR A  28      24.234 -19.339 -44.583  1.00  0.00           H
    ATOM     19  HE1 TYR A  28      22.428 -18.176 -40.109  1.00  0.00           H
    ATOM     20  HE2 TYR A  28      25.709 -19.445 -42.601  1.00  0.00           H
    ATOM     21  HH  TYR A  28      25.856 -19.190 -40.322  1.00  0.00           H
    ATOM     22  N   ILE A  29      21.953 -15.220 -44.181  1.00 31.54           N
    ATOM     23  CA  ILE A  29      22.813 -14.024 -44.179  1.00 32.23           C
    ATOM     24  C   ILE A  29      23.636 -13.882 -42.885  1.00 33.41           C
    ATOM     25  O   ILE A  29      24.055 -12.774 -42.546  1.00 34.22           O
    ATOM     26  CB  ILE A  29      22.051 -12.731 -44.603  1.00 30.73           C
    ATOM     27  CG1 ILE A  29      21.021 -12.245 -43.550  1.00 29.31           C
    ATOM     28  CG2 ILE A  29      21.447 -12.853 -46.012  1.00 30.63           C
    ATOM     29  CD1 ILE A  29      20.436 -10.849 -43.803  1.00 25.64           C
    ATOM     30  H   ILE A  29      21.325 -15.319 -43.395  1.00  0.00           H
    ATOM     31  HA  ILE A  29      23.588 -14.169 -44.934  1.00  0.00           H
    ATOM     32  HB  ILE A  29      22.805 -11.945 -44.679  1.00  0.00           H
    ATOM     33 HG12 ILE A  29      20.214 -12.960 -43.470  1.00  0.00           H
    ATOM     34 HG13 ILE A  29      21.478 -12.222 -42.562  1.00  0.00           H
    ATOM     35 HG21 ILE A  29      21.175 -11.876 -46.407  1.00  0.00           H
    ATOM     36 HG22 ILE A  29      22.159 -13.291 -46.712  1.00  0.00           H
    ATOM     37 HG23 ILE A  29      20.558 -13.477 -46.006  1.00  0.00           H
    ATOM     38 HD11 ILE A  29      19.785 -10.554 -42.980  1.00  0.00           H
    ATOM     39 HD12 ILE A  29      21.223 -10.100 -43.887  1.00  0.00           H
    ATOM     40 HD13 ILE A  29      19.839 -10.813 -44.709  1.00  0.00           H
    ATOM     41  N   GLY A  30      23.892 -14.996 -42.182  1.00 34.98           N
    ATOM     42  CA  GLY A  30      24.690 -15.056 -40.960  1.00 35.73           C
    ATOM     43  C   GLY A  30      23.833 -15.197 -39.700  1.00 35.77           C
    ATOM     44  O   GLY A  30      22.643 -15.493 -39.781  1.00 34.10           O
    ATOM     45  H   GLY A  30      23.500 -15.866 -42.512  1.00  0.00           H
    ATOM     46  HA2 GLY A  30      25.344 -14.188 -40.863  1.00  0.00           H
    ATOM     47  HA3 GLY A  30      25.341 -15.928 -41.026  1.00  0.00           H
    ATOM     48  N   GLU A  31      24.464 -15.018 -38.532  1.00 37.89           N
    ATOM     49  CA  GLU A  31      23.846 -15.161 -37.214  1.00 38.86           C
    ATOM     50  C   GLU A  31      23.858 -13.831 -36.459  1.00 37.93           C
    ATOM     51  O   GLU A  31      24.684 -12.952 -36.712  1.00 38.44           O
    ATOM     52  CB  GLU A  31      24.621 -16.213 -36.382  1.00 39.94           C
    ATOM     53  CG  GLU A  31      24.605 -17.649 -36.935  1.00 42.85           C
    ATOM     54  CD  GLU A  31      23.219 -18.296 -36.922  1.00 44.41           C
    ATOM     55  OE1 GLU A  31      23.005 -19.196 -36.081  1.00 44.59           O
    ATOM     56  OE2 GLU A  31      22.389 -17.883 -37.756  1.00 44.89           O1-
    ATOM     57  H   GLU A  31      25.442 -14.770 -38.548  1.00  0.00           H
    ATOM     58  HA  GLU A  31      22.801 -15.456 -37.304  1.00  0.00           H
    ATOM     59  HB2 GLU A  31      25.660 -15.896 -36.278  1.00  0.00           H
    ATOM     60  HB3 GLU A  31      24.225 -16.240 -35.365  1.00  0.00           H
    ATOM     61  HG2 GLU A  31      24.999 -17.657 -37.948  1.00  0.00           H
    ATOM     62  HG3 GLU A  31      25.280 -18.269 -36.344  1.00  0.00           H
    ATOM     63  N   SER A  39      17.877 -13.270 -45.056  1.00 25.56           N1+
    ATOM     64  CA  SER A  39      17.174 -12.328 -45.920  1.00 24.66           C
    ATOM     65  C   SER A  39      15.938 -13.000 -46.539  1.00 24.60           C
    ATOM     66  O   SER A  39      15.827 -14.225 -46.560  1.00 24.00           O
    ATOM     67  CB  SER A  39      18.115 -11.811 -47.028  1.00 24.85           C
    ATOM     68  OG  SER A  39      18.525 -12.807 -47.941  1.00 24.75           O
    ATOM     69  H   SER A  39      18.602 -13.836 -45.476  1.00  0.00           H
    ATOM     70  HA  SER A  39      16.856 -11.483 -45.313  1.00  0.00           H
    ATOM     71  HB2 SER A  39      18.993 -11.332 -46.608  1.00  0.00           H
    ATOM     72  HB3 SER A  39      17.604 -11.036 -47.600  1.00  0.00           H
    ATOM     73  HG  SER A  39      19.032 -13.462 -47.480  1.00  0.00           H
    ATOM     74  N   ARG A  48      14.039  -6.205 -50.607  1.00 24.76           N1+
    ATOM     75  CA  ARG A  48      14.735  -7.266 -49.896  1.00 23.98           C
    ATOM     76  C   ARG A  48      14.915  -6.808 -48.437  1.00 21.94           C
    ATOM     77  O   ARG A  48      15.507  -5.760 -48.178  1.00 21.61           O
    ATOM     78  CB  ARG A  48      16.103  -7.480 -50.580  1.00 25.60           C
    ATOM     79  CG  ARG A  48      16.763  -8.816 -50.209  1.00 27.65           C
    ATOM     80  CD  ARG A  48      16.197 -10.013 -50.982  1.00 27.89           C
    ATOM     81  NE  ARG A  48      16.708 -11.273 -50.430  1.00 28.42           N
    ATOM     82  CZ  ARG A  48      16.253 -12.500 -50.726  1.00 27.52           C
    ATOM     83  NH1 ARG A  48      15.262 -12.687 -51.606  1.00 28.69           N
    ATOM     84  NH2 ARG A  48      16.798 -13.564 -50.132  1.00 25.81           N1+
    ATOM     85  H   ARG A  48      14.569  -5.371 -50.815  1.00  0.00           H
    ATOM     86  HA  ARG A  48      14.159  -8.190 -49.936  1.00  0.00           H
    ATOM     87  HB2 ARG A  48      15.992  -7.439 -51.665  1.00  0.00           H
    ATOM     88  HB3 ARG A  48      16.781  -6.662 -50.329  1.00  0.00           H
    ATOM     89  HG2 ARG A  48      16.665  -8.997 -49.140  1.00  0.00           H
    ATOM     90  HG3 ARG A  48      17.830  -8.752 -50.421  1.00  0.00           H
    ATOM     91  HD2 ARG A  48      15.109 -10.026 -50.920  1.00  0.00           H
    ATOM     92  HD3 ARG A  48      16.457  -9.946 -52.039  1.00  0.00           H
    ATOM     93  HE  ARG A  48      17.482 -11.198 -49.785  1.00  0.00           H
    ATOM     94 HH11 ARG A  48      14.935 -13.624 -51.803  1.00  0.00           H
    ATOM     95 HH12 ARG A  48      14.846 -11.899 -52.079  1.00  0.00           H
    ATOM     96 HH21 ARG A  48      16.451 -14.493 -50.339  1.00  0.00           H
    ATOM     97 HH22 ARG A  48      17.538 -13.449 -49.445  1.00  0.00           H
    ATOM     98  N   ALA A  50      15.702  -8.103 -44.113  1.00 14.43           N1+
    ATOM     99  CA  ALA A  50      16.350  -9.020 -43.178  1.00 12.09           C
    ATOM    100  C   ALA A  50      15.297  -9.494 -42.166  1.00 11.74           C
    ATOM    101  O   ALA A  50      14.561  -8.671 -41.623  1.00 11.43           O
    ATOM    102  CB  ALA A  50      17.432  -8.238 -42.416  1.00 14.33           C
    ATOM    103  H   ALA A  50      15.375  -7.228 -43.720  1.00  0.00           H
    ATOM    104  HA  ALA A  50      16.802  -9.871 -43.688  1.00  0.00           H
    ATOM    105  HB1 ALA A  50      17.915  -8.861 -41.663  1.00  0.00           H
    ATOM    106  HB2 ALA A  50      18.211  -7.895 -43.098  1.00  0.00           H
    ATOM    107  HB3 ALA A  50      17.026  -7.358 -41.914  1.00  0.00           H
    ATOM    108  N   ARG A  77      12.678   6.250 -32.868  1.00 15.19           N1+
    ATOM    109  CA  ARG A  77      12.508   6.778 -34.209  1.00 13.33           C
    ATOM    110  C   ARG A  77      13.637   7.767 -34.486  1.00 12.06           C
    ATOM    111  O   ARG A  77      13.486   8.972 -34.285  1.00 10.98           O
    ATOM    112  CB  ARG A  77      11.072   7.316 -34.360  1.00 15.81           C
    ATOM    113  CG  ARG A  77      10.694   7.692 -35.799  1.00 21.37           C
    ATOM    114  CD  ARG A  77       9.194   7.997 -35.933  1.00 25.62           C
    ATOM    115  NE  ARG A  77       8.656   7.494 -37.203  1.00 28.05           N
    ATOM    116  CZ  ARG A  77       8.368   6.207 -37.474  1.00 30.51           C
    ATOM    117  NH1 ARG A  77       8.579   5.235 -36.574  1.00 27.65           N
    ATOM    118  NH2 ARG A  77       7.864   5.890 -38.673  1.00 31.22           N1+
    ATOM    119  H   ARG A  77      13.008   6.896 -32.162  1.00  0.00           H
    ATOM    120  HA  ARG A  77      12.628   5.959 -34.923  1.00  0.00           H
    ATOM    121  HB2 ARG A  77      10.379   6.546 -34.016  1.00  0.00           H
    ATOM    122  HB3 ARG A  77      10.916   8.171 -33.701  1.00  0.00           H
    ATOM    123  HG2 ARG A  77      11.274   8.558 -36.122  1.00  0.00           H
    ATOM    124  HG3 ARG A  77      10.974   6.876 -36.467  1.00  0.00           H
    ATOM    125  HD2 ARG A  77       8.617   7.551 -35.122  1.00  0.00           H
    ATOM    126  HD3 ARG A  77       9.025   9.073 -35.870  1.00  0.00           H
    ATOM    127  HE  ARG A  77       8.517   8.185 -37.927  1.00  0.00           H
    ATOM    128 HH11 ARG A  77       8.366   4.272 -36.792  1.00  0.00           H
    ATOM    129 HH12 ARG A  77       8.964   5.462 -35.668  1.00  0.00           H
    ATOM    130 HH21 ARG A  77       7.648   4.930 -38.899  1.00  0.00           H
    ATOM    131 HH22 ARG A  77       7.702   6.606 -39.365  1.00  0.00           H
    ATOM    132  N   HIE A  78      14.778   7.224 -34.928  1.00 11.61           N
    ATOM    133  CA  HIE A  78      16.005   7.945 -35.231  1.00 10.68           C
    ATOM    134  C   HIE A  78      16.638   7.347 -36.490  1.00 10.47           C
    ATOM    135  O   HIE A  78      16.495   6.153 -36.754  1.00 10.53           O
    ATOM    136  CB  HIE A  78      16.916   7.945 -33.990  1.00  9.44           C
    ATOM    137  CG  HIE A  78      18.088   8.885 -34.069  1.00  6.99           C
    ATOM    138  ND1 HIE A  78      19.310   8.515 -34.631  1.00  7.83           N
    ATOM    139  CD2 HIE A  78      18.168  10.196 -33.649  1.00  6.57           C
    ATOM    140  CE1 HIE A  78      20.065   9.597 -34.533  1.00  8.35           C
    ATOM    141  NE2 HIE A  78      19.445  10.625 -33.954  1.00  9.50           N
    ATOM    142  H   HIE A  78      14.803   6.225 -35.072  1.00  0.00           H
    ATOM    143  HA  HIE A  78      15.750   8.981 -35.464  1.00  0.00           H
    ATOM    144  HB2 HIE A  78      16.334   8.233 -33.115  1.00  0.00           H
    ATOM    145  HB3 HIE A  78      17.285   6.941 -33.793  1.00  0.00           H
    ATOM    146  HD2 HIE A  78      17.444  10.831 -33.163  1.00  0.00           H
    ATOM    147  HE2 HIE A  78      19.841  11.532 -33.755  1.00  0.00           H
    ATOM    148  HE1 HIE A  78      21.089   9.615 -34.863  1.00  0.00           H
    ATOM    149  N   GLU A  79      17.343   8.188 -37.263  1.00 11.70           N
    ATOM    150  CA  GLU A  79      18.010   7.831 -38.515  1.00 13.27           C
    ATOM    151  C   GLU A  79      19.157   6.827 -38.317  1.00 12.33           C
    ATOM    152  O   GLU A  79      19.344   5.951 -39.158  1.00 12.15           O
    ATOM    153  CB  GLU A  79      18.527   9.133 -39.163  1.00 15.25           C
    ATOM    154  CG  GLU A  79      19.294   8.958 -40.495  1.00 16.20           C
    ATOM    155  CD  GLU A  79      19.874  10.267 -41.032  1.00 18.95           C
    ATOM    156  OE1 GLU A  79      19.502  11.337 -40.501  1.00 17.16           O
    ATOM    157  OE2 GLU A  79      20.712  10.180 -41.955  1.00 19.53           O1-
    ATOM    158  H   GLU A  79      17.418   9.151 -36.967  1.00  0.00           H
    ATOM    159  HA  GLU A  79      17.271   7.378 -39.179  1.00  0.00           H
    ATOM    160  HB2 GLU A  79      17.678   9.798 -39.329  1.00  0.00           H
    ATOM    161  HB3 GLU A  79      19.174   9.644 -38.447  1.00  0.00           H
    ATOM    162  HG2 GLU A  79      20.133   8.271 -40.379  1.00  0.00           H
    ATOM    163  HG3 GLU A  79      18.638   8.529 -41.252  1.00  0.00           H
    ATOM    164  N   ASN A  80      19.902   6.951 -37.206  1.00 11.20           N
    ATOM    165  CA  ASN A  80      21.066   6.131 -36.865  1.00 10.30           C
    ATOM    166  C   ASN A  80      20.700   5.067 -35.814  1.00  7.61           C
    ATOM    167  O   ASN A  80      21.539   4.677 -35.001  1.00  5.18           O
    ATOM    168  CB  ASN A  80      22.226   7.024 -36.355  1.00 10.79           C
    ATOM    169  CG  ASN A  80      22.586   8.164 -37.313  1.00 10.21           C
    ATOM    170  OD1 ASN A  80      22.346   9.330 -37.014  1.00  9.93           O
    ATOM    171  ND2 ASN A  80      23.176   7.837 -38.462  1.00  7.53           N
    ATOM    172  H   ASN A  80      19.651   7.676 -36.546  1.00  0.00           H
    ATOM    173  HA  ASN A  80      21.402   5.599 -37.751  1.00  0.00           H
    ATOM    174  HB2 ASN A  80      23.121   6.420 -36.201  1.00  0.00           H
    ATOM    175  HB3 ASN A  80      21.977   7.452 -35.383  1.00  0.00           H
    ATOM    176 HD21 ASN A  80      23.428   8.553 -39.127  1.00  0.00           H
    ATOM    177 HD22 ASN A  80      23.352   6.863 -38.678  1.00  0.00           H
    ATOM    178  N   ILE A  81      19.455   4.570 -35.858  1.00  6.62           N
    ATOM    179  CA  ILE A  81      18.928   3.447 -35.087  1.00  6.53           C
    ATOM    180  C   ILE A  81      18.059   2.628 -36.050  1.00  7.61           C
    ATOM    181  O   ILE A  81      17.203   3.196 -36.731  1.00  9.12           O
    ATOM    182  CB  ILE A  81      18.181   3.928 -33.805  1.00  7.82           C
    ATOM    183  CG1 ILE A  81      19.080   4.792 -32.886  1.00  7.33           C
    ATOM    184  CG2 ILE A  81      17.566   2.747 -33.022  1.00 11.29           C
    ATOM    185  CD1 ILE A  81      18.386   5.314 -31.625  1.00  6.28           C
    ATOM    186  H   ILE A  81      18.827   4.957 -36.550  1.00  0.00           H
    ATOM    187  HA  ILE A  81      19.761   2.812 -34.780  1.00  0.00           H
    ATOM    188  HB  ILE A  81      17.351   4.557 -34.129  1.00  0.00           H
    ATOM    189 HG12 ILE A  81      19.977   4.241 -32.605  1.00  0.00           H
    ATOM    190 HG13 ILE A  81      19.421   5.670 -33.433  1.00  0.00           H
    ATOM    191 HG21 ILE A  81      17.013   3.078 -32.145  1.00  0.00           H
    ATOM    192 HG22 ILE A  81      16.853   2.186 -33.624  1.00  0.00           H
    ATOM    193 HG23 ILE A  81      18.337   2.052 -32.690  1.00  0.00           H
    ATOM    194 HD11 ILE A  81      18.965   6.124 -31.185  1.00  0.00           H
    ATOM    195 HD12 ILE A  81      17.392   5.701 -31.845  1.00  0.00           H
    ATOM    196 HD13 ILE A  81      18.282   4.531 -30.874  1.00  0.00           H
    ATOM    197  N   ILE A  82      18.263   1.300 -36.103  1.00  7.76           N
    ATOM    198  CA  ILE A  82      17.468   0.365 -36.910  1.00  8.08           C
    ATOM    199  C   ILE A  82      16.002   0.343 -36.426  1.00  9.92           C
    ATOM    200  O   ILE A  82      15.743   0.472 -35.231  1.00 10.49           O
    ATOM    201  CB  ILE A  82      18.140  -1.041 -36.882  1.00  6.33           C
    ATOM    202  CG1 ILE A  82      17.603  -2.008 -37.966  1.00  6.22           C
    ATOM    203  CG2 ILE A  82      18.074  -1.729 -35.501  1.00  4.74           C
    ATOM    204  CD1 ILE A  82      17.828  -1.549 -39.415  1.00  5.59           C
    ATOM    205  H   ILE A  82      18.983   0.901 -35.515  1.00  0.00           H
    ATOM    206  HA  ILE A  82      17.482   0.747 -37.930  1.00  0.00           H
    ATOM    207  HB  ILE A  82      19.199  -0.890 -37.101  1.00  0.00           H
    ATOM    208 HG12 ILE A  82      16.542  -2.202 -37.812  1.00  0.00           H
    ATOM    209 HG13 ILE A  82      18.096  -2.973 -37.846  1.00  0.00           H
    ATOM    210 HG21 ILE A  82      18.688  -2.630 -35.486  1.00  0.00           H
    ATOM    211 HG22 ILE A  82      18.429  -1.078 -34.703  1.00  0.00           H
    ATOM    212 HG23 ILE A  82      17.056  -2.027 -35.251  1.00  0.00           H
    ATOM    213 HD11 ILE A  82      17.554  -2.339 -40.115  1.00  0.00           H
    ATOM    214 HD12 ILE A  82      17.221  -0.678 -39.665  1.00  0.00           H
    ATOM    215 HD13 ILE A  82      18.872  -1.296 -39.598  1.00  0.00           H
    ATOM    216  N   ASP A 104      14.956  -3.651 -42.238  1.00 16.34           N1+
    ATOM    217  CA  ASP A 104      15.660  -3.283 -43.471  1.00 18.25           C
    ATOM    218  C   ASP A 104      16.946  -4.111 -43.606  1.00 18.28           C
    ATOM    219  O   ASP A 104      17.622  -4.363 -42.605  1.00 18.03           O
    ATOM    220  CB  ASP A 104      16.040  -1.782 -43.455  1.00 18.48           C
    ATOM    221  CG  ASP A 104      14.885  -0.840 -43.112  1.00 16.89           C
    ATOM    222  OD1 ASP A 104      13.833  -0.962 -43.777  1.00 18.07           O
    ATOM    223  OD2 ASP A 104      15.074  -0.024 -42.183  1.00 19.15           O1-
    ATOM    224  H   ASP A 104      14.961  -2.971 -41.491  1.00  0.00           H
    ATOM    225  HA  ASP A 104      15.003  -3.484 -44.320  1.00  0.00           H
    ATOM    226  HB2 ASP A 104      16.438  -1.488 -44.427  1.00  0.00           H
    ATOM    227  HB3 ASP A 104      16.843  -1.608 -42.735  1.00  0.00           H
    ATOM    228  N   LEU A 105      17.287  -4.523 -44.837  1.00 18.73           N
    ATOM    229  CA  LEU A 105      18.526  -5.243 -45.114  1.00 19.03           C
    ATOM    230  C   LEU A 105      19.628  -4.194 -45.306  1.00 18.63           C
    ATOM    231  O   LEU A 105      19.673  -3.491 -46.316  1.00 20.26           O
    ATOM    232  CB  LEU A 105      18.352  -6.102 -46.386  1.00 16.25           C
    ATOM    233  CG  LEU A 105      19.350  -7.279 -46.508  1.00 17.14           C
    ATOM    234  CD1 LEU A 105      18.873  -8.260 -47.578  1.00 18.93           C
    ATOM    235  CD2 LEU A 105      20.795  -6.852 -46.809  1.00 20.24           C
    ATOM    236  H   LEU A 105      16.701  -4.277 -45.621  1.00  0.00           H
    ATOM    237  HA  LEU A 105      18.767  -5.903 -44.278  1.00  0.00           H
    ATOM    238  HB2 LEU A 105      17.352  -6.534 -46.350  1.00  0.00           H
    ATOM    239  HB3 LEU A 105      18.361  -5.494 -47.293  1.00  0.00           H
    ATOM    240  HG  LEU A 105      19.353  -7.824 -45.562  1.00  0.00           H
    ATOM    241 HD11 LEU A 105      19.547  -9.112 -47.669  1.00  0.00           H
    ATOM    242 HD12 LEU A 105      17.887  -8.646 -47.325  1.00  0.00           H
    ATOM    243 HD13 LEU A 105      18.816  -7.770 -48.549  1.00  0.00           H
    ATOM    244 HD21 LEU A 105      21.426  -7.715 -47.024  1.00  0.00           H
    ATOM    245 HD22 LEU A 105      20.845  -6.181 -47.668  1.00  0.00           H
    ATOM    246 HD23 LEU A 105      21.245  -6.347 -45.959  1.00  0.00           H
    ATOM    247  N   MET A 106      20.513  -4.112 -44.306  1.00 17.65           N
    ATOM    248  CA  MET A 106      21.720  -3.300 -44.315  1.00 16.50           C
    ATOM    249  C   MET A 106      22.846  -4.207 -44.820  1.00 14.00           C
    ATOM    250  O   MET A 106      23.063  -5.295 -44.282  1.00 13.89           O
    ATOM    251  CB  MET A 106      22.026  -2.863 -42.870  1.00 17.73           C
    ATOM    252  CG  MET A 106      20.930  -1.965 -42.262  1.00 21.41           C
    ATOM    253  SD  MET A 106      20.735  -0.337 -43.033  1.00 23.89           S
    ATOM    254  CE  MET A 106      22.180   0.515 -42.350  1.00 22.79           C
    ATOM    255  H   MET A 106      20.383  -4.721 -43.511  1.00  0.00           H
    ATOM    256  HA  MET A 106      21.613  -2.426 -44.962  1.00  0.00           H
    ATOM    257  HB2 MET A 106      22.147  -3.742 -42.234  1.00  0.00           H
    ATOM    258  HB3 MET A 106      22.984  -2.341 -42.845  1.00  0.00           H
    ATOM    259  HG2 MET A 106      19.965  -2.472 -42.300  1.00  0.00           H
    ATOM    260  HG3 MET A 106      21.125  -1.797 -41.205  1.00  0.00           H
    ATOM    261  HE1 MET A 106      22.248   1.528 -42.746  1.00  0.00           H
    ATOM    262  HE2 MET A 106      22.117   0.569 -41.264  1.00  0.00           H
    ATOM    263  HE3 MET A 106      23.090  -0.015 -42.613  1.00  0.00           H
    ATOM    264  N   GLU A 107      23.519  -3.767 -45.892  1.00 12.58           N
    ATOM    265  CA  GLU A 107      24.430  -4.552 -46.725  1.00 11.77           C
    ATOM    266  C   GLU A 107      25.646  -5.111 -45.965  1.00  9.77           C
    ATOM    267  O   GLU A 107      26.123  -6.194 -46.301  1.00  8.12           O
    ATOM    268  CB  GLU A 107      24.905  -3.667 -47.895  1.00 11.70           C
    ATOM    269  CG  GLU A 107      23.894  -3.517 -49.051  1.00 15.76           C
    ATOM    270  CD  GLU A 107      22.619  -2.740 -48.714  1.00 14.81           C
    ATOM    271  OE1 GLU A 107      22.727  -1.730 -47.985  1.00  9.23           O
    ATOM    272  OE2 GLU A 107      21.553  -3.168 -49.203  1.00 16.49           O1-
    ATOM    273  H   GLU A 107      23.249  -2.867 -46.271  1.00  0.00           H
    ATOM    274  HA  GLU A 107      23.877  -5.405 -47.123  1.00  0.00           H
    ATOM    275  HB2 GLU A 107      25.208  -2.690 -47.519  1.00  0.00           H
    ATOM    276  HB3 GLU A 107      25.799  -4.111 -48.330  1.00  0.00           H
    ATOM    277  HG2 GLU A 107      23.619  -4.507 -49.420  1.00  0.00           H
    ATOM    278  HG3 GLU A 107      24.373  -3.014 -49.888  1.00  0.00           H
    ATOM    279  N   THR A 108      26.116  -4.401 -44.929  1.00  6.04           N
    ATOM    280  CA  THR A 108      27.211  -4.804 -44.057  1.00  6.17           C
    ATOM    281  C   THR A 108      26.996  -4.287 -42.626  1.00  4.90           C
    ATOM    282  O   THR A 108      25.968  -3.693 -42.301  1.00  6.38           O
    ATOM    283  CB  THR A 108      28.573  -4.465 -44.729  1.00  5.41           C
    ATOM    284  OG1 THR A 108      29.637  -5.154 -44.104  1.00  5.53           O
    ATOM    285  CG2 THR A 108      28.936  -2.971 -44.753  1.00  5.54           C
    ATOM    286  H   THR A 108      25.658  -3.526 -44.705  1.00  0.00           H
    ATOM    287  HA  THR A 108      27.159  -5.890 -43.962  1.00  0.00           H
    ATOM    288  HB  THR A 108      28.544  -4.807 -45.759  1.00  0.00           H
    ATOM    289 HG21 THR A 108      29.837  -2.797 -45.342  1.00  0.00           H
    ATOM    290 HG22 THR A 108      28.137  -2.377 -45.196  1.00  0.00           H
    ATOM    291 HG23 THR A 108      29.125  -2.586 -43.752  1.00  0.00           H
    ATOM    292  HG1 THR A 108      29.881  -4.684 -43.320  1.00  0.00           H
    ATOM    293  N   ASP A 109      28.007  -4.511 -41.784  1.00  4.58           N
    ATOM    294  CA  ASP A 109      28.223  -3.904 -40.479  1.00  2.99           C
    ATOM    295  C   ASP A 109      29.699  -3.484 -40.397  1.00  2.88           C
    ATOM    296  O   ASP A 109      30.509  -3.851 -41.253  1.00  1.85           O
    ATOM    297  CB  ASP A 109      27.733  -4.851 -39.352  1.00  3.25           C
    ATOM    298  CG  ASP A 109      28.372  -6.246 -39.357  1.00  4.21           C
    ATOM    299  OD1 ASP A 109      29.619  -6.311 -39.384  1.00  5.33           O
    ATOM    300  OD2 ASP A 109      27.604  -7.230 -39.355  1.00  6.14           O1-
    ATOM    301  H   ASP A 109      28.774  -5.072 -42.123  1.00  0.00           H
    ATOM    302  HA  ASP A 109      27.652  -2.978 -40.434  1.00  0.00           H
    ATOM    303  HB2 ASP A 109      26.652  -4.967 -39.432  1.00  0.00           H
    ATOM    304  HB3 ASP A 109      27.911  -4.401 -38.375  1.00  0.00           H
    ATOM    305  N   LEU A 110      30.049  -2.713 -39.357  1.00  2.53           N
    ATOM    306  CA  LEU A 110      31.400  -2.210 -39.121  1.00  3.78           C
    ATOM    307  C   LEU A 110      32.393  -3.322 -38.716  1.00  4.56           C
    ATOM    308  O   LEU A 110      33.586  -3.157 -38.958  1.00  3.14           O
    ATOM    309  CB  LEU A 110      31.332  -1.086 -38.060  1.00  1.85           C
    ATOM    310  CG  LEU A 110      32.648  -0.324 -37.786  1.00  1.85           C
    ATOM    311  CD1 LEU A 110      33.199   0.384 -39.035  1.00  1.85           C
    ATOM    312  CD2 LEU A 110      32.462   0.683 -36.646  1.00  1.85           C
    ATOM    313  H   LEU A 110      29.329  -2.444 -38.698  1.00  0.00           H
    ATOM    314  HA  LEU A 110      31.743  -1.777 -40.060  1.00  0.00           H
    ATOM    315  HB2 LEU A 110      30.572  -0.364 -38.363  1.00  0.00           H
    ATOM    316  HB3 LEU A 110      30.974  -1.502 -37.121  1.00  0.00           H
    ATOM    317  HG  LEU A 110      33.394  -1.037 -37.443  1.00  0.00           H
    ATOM    318 HD11 LEU A 110      34.110   0.930 -38.798  1.00  0.00           H
    ATOM    319 HD12 LEU A 110      33.446  -0.317 -39.831  1.00  0.00           H
    ATOM    320 HD13 LEU A 110      32.480   1.100 -39.434  1.00  0.00           H
    ATOM    321 HD21 LEU A 110      33.393   1.203 -36.422  1.00  0.00           H
    ATOM    322 HD22 LEU A 110      31.712   1.435 -36.894  1.00  0.00           H
    ATOM    323 HD23 LEU A 110      32.144   0.179 -35.735  1.00  0.00           H
    ATOM    324  N   TYR A 111      31.932  -4.469 -38.171  1.00  6.16           N
    ATOM    325  CA  TYR A 111      32.776  -5.638 -37.880  1.00  7.97           C
    ATOM    326  C   TYR A 111      33.333  -6.230 -39.177  1.00 10.08           C
    ATOM    327  O   TYR A 111      34.537  -6.430 -39.286  1.00 12.16           O
    ATOM    328  CB  TYR A 111      32.000  -6.727 -37.094  1.00  6.59           C
    ATOM    329  CG  TYR A 111      32.747  -8.038 -36.905  1.00  6.11           C
    ATOM    330  CD1 TYR A 111      33.609  -8.213 -35.804  1.00  7.28           C
    ATOM    331  CD2 TYR A 111      32.621  -9.066 -37.865  1.00  5.61           C
    ATOM    332  CE1 TYR A 111      34.375  -9.390 -35.691  1.00  8.57           C
    ATOM    333  CE2 TYR A 111      33.404 -10.229 -37.767  1.00  6.83           C
    ATOM    334  CZ  TYR A 111      34.292 -10.385 -36.685  1.00  8.85           C
    ATOM    335  OH  TYR A 111      35.079 -11.496 -36.604  1.00 12.02           O
    ATOM    336  H   TYR A 111      30.936  -4.563 -38.024  1.00  0.00           H
    ATOM    337  HA  TYR A 111      33.624  -5.313 -37.276  1.00  0.00           H
    ATOM    338  HB2 TYR A 111      31.747  -6.358 -36.107  1.00  0.00           H
    ATOM    339  HB3 TYR A 111      31.048  -6.956 -37.564  1.00  0.00           H
    ATOM    340  HD1 TYR A 111      33.704  -7.438 -35.058  1.00  0.00           H
    ATOM    341  HD2 TYR A 111      31.954  -8.945 -38.704  1.00  0.00           H
    ATOM    342  HE1 TYR A 111      35.048  -9.514 -34.856  1.00  0.00           H
    ATOM    343  HE2 TYR A 111      33.327 -10.993 -38.526  1.00  0.00           H
    ATOM    344  HH  TYR A 111      34.921 -12.118 -37.298  1.00  0.00           H
    ATOM    345  N   LYS A 112      32.454  -6.492 -40.153  1.00 10.76           N
    ATOM    346  CA  LYS A 112      32.790  -7.050 -41.456  1.00 11.02           C
    ATOM    347  C   LYS A 112      33.653  -6.075 -42.279  1.00 10.83           C
    ATOM    348  O   LYS A 112      34.600  -6.497 -42.943  1.00 10.49           O
    ATOM    349  CB  LYS A 112      31.467  -7.338 -42.190  1.00 12.46           C
    ATOM    350  CG  LYS A 112      30.686  -8.549 -41.653  1.00 14.67           C
    ATOM    351  CD  LYS A 112      29.226  -8.585 -42.135  1.00 17.40           C
    ATOM    352  CE  LYS A 112      29.056  -8.794 -43.647  1.00 18.38           C
    ATOM    353  NZ  LYS A 112      27.632  -8.810 -44.022  1.00 15.59           N1+
    ATOM    354  H   LYS A 112      31.473  -6.313 -39.966  1.00  0.00           H
    ATOM    355  HA  LYS A 112      33.348  -7.979 -41.321  1.00  0.00           H
    ATOM    356  HB2 LYS A 112      30.835  -6.451 -42.139  1.00  0.00           H
    ATOM    357  HB3 LYS A 112      31.659  -7.512 -43.243  1.00  0.00           H
    ATOM    358  HG2 LYS A 112      31.199  -9.473 -41.919  1.00  0.00           H
    ATOM    359  HG3 LYS A 112      30.671  -8.516 -40.564  1.00  0.00           H
    ATOM    360  HD2 LYS A 112      28.731  -7.657 -41.849  1.00  0.00           H
    ATOM    361  HD3 LYS A 112      28.703  -9.380 -41.602  1.00  0.00           H
    ATOM    362  HE2 LYS A 112      29.539  -7.991 -44.203  1.00  0.00           H
    ATOM    363  HE3 LYS A 112      29.519  -9.730 -43.959  1.00  0.00           H
    ATOM    364  HZ1 LYS A 112      27.201  -7.936 -43.755  1.00  0.00           H
    ATOM    365  HZ2 LYS A 112      27.543  -8.928 -45.022  1.00  0.00           H
    ATOM    366  HZ3 LYS A 112      27.162  -9.569 -43.549  1.00  0.00           H
    ATOM    367  N   LEU A 113      33.343  -4.771 -42.206  1.00  9.87           N
    ATOM    368  CA  LEU A 113      33.998  -3.716 -42.970  1.00  9.02           C
    ATOM    369  C   LEU A 113      35.447  -3.453 -42.519  1.00  9.91           C
    ATOM    370  O   LEU A 113      36.297  -3.224 -43.376  1.00  7.93           O
    ATOM    371  CB  LEU A 113      33.156  -2.434 -42.825  1.00  9.19           C
    ATOM    372  CG  LEU A 113      33.598  -1.258 -43.717  1.00  9.43           C
    ATOM    373  CD1 LEU A 113      33.469  -1.598 -45.210  1.00  8.61           C
    ATOM    374  CD2 LEU A 113      32.788  -0.008 -43.368  1.00  7.58           C
    ATOM    375  H   LEU A 113      32.560  -4.501 -41.625  1.00  0.00           H
    ATOM    376  HA  LEU A 113      34.008  -4.036 -44.012  1.00  0.00           H
    ATOM    377  HB2 LEU A 113      32.113  -2.661 -43.050  1.00  0.00           H
    ATOM    378  HB3 LEU A 113      33.176  -2.118 -41.782  1.00  0.00           H
    ATOM    379  HG  LEU A 113      34.641  -1.017 -43.511  1.00  0.00           H
    ATOM    380 HD11 LEU A 113      33.708  -0.735 -45.823  1.00  0.00           H
    ATOM    381 HD12 LEU A 113      34.152  -2.391 -45.511  1.00  0.00           H
    ATOM    382 HD13 LEU A 113      32.455  -1.910 -45.465  1.00  0.00           H
    ATOM    383 HD21 LEU A 113      33.184   0.870 -43.878  1.00  0.00           H
    ATOM    384 HD22 LEU A 113      31.748  -0.133 -43.663  1.00  0.00           H
    ATOM    385 HD23 LEU A 113      32.815   0.195 -42.297  1.00  0.00           H
    ATOM    386  N   LEU A 114      35.748  -3.522 -41.210  1.00 10.20           N
    ATOM    387  CA  LEU A 114      37.096  -3.305 -40.663  1.00 11.77           C
    ATOM    388  C   LEU A 114      38.042  -4.505 -40.847  1.00 12.89           C
    ATOM    389  O   LEU A 114      39.217  -4.406 -40.498  1.00 12.55           O
    ATOM    390  CB  LEU A 114      36.992  -2.975 -39.154  1.00  9.75           C
    ATOM    391  CG  LEU A 114      36.417  -1.573 -38.867  1.00 10.66           C
    ATOM    392  CD1 LEU A 114      36.201  -1.356 -37.366  1.00  8.00           C
    ATOM    393  CD2 LEU A 114      37.306  -0.453 -39.425  1.00 11.49           C
    ATOM    394  H   LEU A 114      35.004  -3.711 -40.552  1.00  0.00           H
    ATOM    395  HA  LEU A 114      37.570  -2.485 -41.200  1.00  0.00           H
    ATOM    396  HB2 LEU A 114      36.379  -3.732 -38.662  1.00  0.00           H
    ATOM    397  HB3 LEU A 114      37.971  -3.034 -38.676  1.00  0.00           H
    ATOM    398  HG  LEU A 114      35.448  -1.491 -39.355  1.00  0.00           H
    ATOM    399 HD11 LEU A 114      35.639  -0.442 -37.175  1.00  0.00           H
    ATOM    400 HD12 LEU A 114      35.664  -2.184 -36.903  1.00  0.00           H
    ATOM    401 HD13 LEU A 114      37.160  -1.238 -36.872  1.00  0.00           H
    ATOM    402 HD21 LEU A 114      36.904   0.520 -39.152  1.00  0.00           H
    ATOM    403 HD22 LEU A 114      38.324  -0.518 -39.040  1.00  0.00           H
    ATOM    404 HD23 LEU A 114      37.355  -0.465 -40.512  1.00  0.00           H
    ATOM    405  N   LYS A 115      37.546  -5.617 -41.408  1.00 13.19           N
    ATOM    406  CA  LYS A 115      38.293  -6.842 -41.687  1.00 13.64           C
    ATOM    407  C   LYS A 115      38.638  -6.955 -43.186  1.00 13.12           C
    ATOM    408  O   LYS A 115      39.277  -7.931 -43.579  1.00 12.80           O
    ATOM    409  CB  LYS A 115      37.424  -8.040 -41.226  1.00 14.44           C
    ATOM    410  CG  LYS A 115      37.609  -8.441 -39.745  1.00 16.07           C
    ATOM    411  CD  LYS A 115      37.551  -7.274 -38.737  1.00 17.82           C
    ATOM    412  CE  LYS A 115      37.004  -7.663 -37.362  1.00 18.06           C
    ATOM    413  NZ  LYS A 115      36.729  -6.468 -36.545  1.00 17.13           N1+
    ATOM    414  H   LYS A 115      36.568  -5.614 -41.661  1.00  0.00           H
    ATOM    415  HA  LYS A 115      39.242  -6.861 -41.147  1.00  0.00           H
    ATOM    416  HB2 LYS A 115      36.370  -7.827 -41.410  1.00  0.00           H
    ATOM    417  HB3 LYS A 115      37.639  -8.928 -41.824  1.00  0.00           H
    ATOM    418  HG2 LYS A 115      38.559  -8.962 -39.622  1.00  0.00           H
    ATOM    419  HG3 LYS A 115      36.838  -9.174 -39.504  1.00  0.00           H
    ATOM    420  HD2 LYS A 115      38.538  -6.821 -38.637  1.00  0.00           H
    ATOM    421  HD3 LYS A 115      36.913  -6.489 -39.123  1.00  0.00           H
    ATOM    422  HE2 LYS A 115      36.069  -8.205 -37.486  1.00  0.00           H
    ATOM    423  HE3 LYS A 115      37.698  -8.320 -36.837  1.00  0.00           H
    ATOM    424  HZ1 LYS A 115      36.069  -5.874 -37.026  1.00  0.00           H
    ATOM    425  HZ2 LYS A 115      36.342  -6.739 -35.653  1.00  0.00           H
    ATOM    426  HZ3 LYS A 115      37.580  -5.949 -36.376  1.00  0.00           H
    ATOM    427  N   THR A 116      38.243  -5.967 -44.014  1.00 14.99           N
    ATOM    428  CA  THR A 116      38.447  -5.928 -45.467  1.00 13.65           C
    ATOM    429  C   THR A 116      38.843  -4.544 -46.005  1.00 13.36           C
    ATOM    430  O   THR A 116      39.665  -4.483 -46.919  1.00 12.86           O
    ATOM    431  CB  THR A 116      37.257  -6.566 -46.225  1.00 13.89           C
    ATOM    432  OG1 THR A 116      36.049  -5.893 -45.916  1.00 14.04           O
    ATOM    433  CG2 THR A 116      37.109  -8.078 -45.996  1.00 14.48           C
    ATOM    434  H   THR A 116      37.754  -5.185 -43.602  1.00  0.00           H
    ATOM    435  HA  THR A 116      39.320  -6.545 -45.688  1.00  0.00           H
    ATOM    436  HB  THR A 116      37.422  -6.425 -47.294  1.00  0.00           H
    ATOM    437 HG21 THR A 116      36.356  -8.513 -46.651  1.00  0.00           H
    ATOM    438 HG22 THR A 116      38.047  -8.595 -46.203  1.00  0.00           H
    ATOM    439 HG23 THR A 116      36.832  -8.305 -44.968  1.00  0.00           H
    ATOM    440  HG1 THR A 116      35.890  -5.974 -44.987  1.00  0.00           H
    ATOM    441  N   GLN A 117      38.281  -3.449 -45.464  1.00 13.91           N
    ATOM    442  CA  GLN A 117      38.550  -2.078 -45.901  1.00 15.55           C
    ATOM    443  C   GLN A 117      39.321  -1.320 -44.811  1.00 16.15           C
    ATOM    444  O   GLN A 117      39.009  -1.433 -43.626  1.00 16.79           O
    ATOM    445  CB  GLN A 117      37.214  -1.348 -46.190  1.00 16.81           C
    ATOM    446  CG  GLN A 117      37.412  -0.052 -47.011  1.00 21.52           C
    ATOM    447  CD  GLN A 117      36.233   0.918 -46.943  1.00 24.25           C
    ATOM    448  OE1 GLN A 117      35.125   0.587 -47.351  1.00 26.77           O
    ATOM    449  NE2 GLN A 117      36.479   2.140 -46.465  1.00 25.84           N
    ATOM    450  H   GLN A 117      37.612  -3.558 -44.713  1.00  0.00           H
    ATOM    451  HA  GLN A 117      39.141  -2.081 -46.819  1.00  0.00           H
    ATOM    452  HB2 GLN A 117      36.547  -2.008 -46.746  1.00  0.00           H
    ATOM    453  HB3 GLN A 117      36.698  -1.129 -45.253  1.00  0.00           H
    ATOM    454  HG2 GLN A 117      38.304   0.479 -46.697  1.00  0.00           H
    ATOM    455  HG3 GLN A 117      37.583  -0.311 -48.056  1.00  0.00           H
    ATOM    456 HE21 GLN A 117      35.734   2.820 -46.429  1.00  0.00           H
    ATOM    457 HE22 GLN A 117      37.412   2.385 -46.164  1.00  0.00           H
    ATOM    458  N   HIE A 118      40.278  -0.488 -45.247  1.00 15.84           N
    ATOM    459  CA  HIE A 118      40.915   0.579 -44.481  1.00 15.67           C
    ATOM    460  C   HIE A 118      40.017   1.827 -44.571  1.00 15.40           C
    ATOM    461  O   HIE A 118      39.729   2.301 -45.672  1.00 16.25           O
    ATOM    462  CB  HIE A 118      42.273   0.880 -45.162  1.00 17.03           C
    ATOM    463  CG  HIE A 118      42.921   2.194 -44.793  1.00 21.38           C
    ATOM    464  ND1 HIE A 118      43.261   3.153 -45.753  1.00 23.98           N
    ATOM    465  CD2 HIE A 118      43.246   2.680 -43.543  1.00 22.54           C
    ATOM    466  CE1 HIE A 118      43.747   4.172 -45.059  1.00 24.13           C
    ATOM    467  NE2 HIE A 118      43.762   3.947 -43.745  1.00 26.25           N
    ATOM    468  H   HIE A 118      40.466  -0.484 -46.239  1.00  0.00           H
    ATOM    469  HA  HIE A 118      41.068   0.290 -43.439  1.00  0.00           H
    ATOM    470  HB2 HIE A 118      42.979   0.079 -44.938  1.00  0.00           H
    ATOM    471  HB3 HIE A 118      42.153   0.874 -46.247  1.00  0.00           H
    ATOM    472  HD2 HIE A 118      43.131   2.248 -42.559  1.00  0.00           H
    ATOM    473  HE2 HIE A 118      44.080   4.586 -43.030  1.00  0.00           H
    ATOM    474  HE1 HIE A 118      44.096   5.089 -45.511  1.00  0.00           H
    ATOM    475  N   LEU A 119      39.603   2.379 -43.420  1.00 14.72           N
    ATOM    476  CA  LEU A 119      38.819   3.612 -43.369  1.00 14.36           C
    ATOM    477  C   LEU A 119      39.759   4.826 -43.438  1.00 14.40           C
    ATOM    478  O   LEU A 119      40.757   4.896 -42.719  1.00 15.31           O
    ATOM    479  CB  LEU A 119      38.042   3.703 -42.038  1.00 11.94           C
    ATOM    480  CG  LEU A 119      37.037   2.572 -41.739  1.00 11.67           C
    ATOM    481  CD1 LEU A 119      36.251   2.905 -40.463  1.00 13.26           C
    ATOM    482  CD2 LEU A 119      36.068   2.292 -42.896  1.00 12.22           C
    ATOM    483  H   LEU A 119      39.886   1.966 -42.543  1.00  0.00           H
    ATOM    484  HA  LEU A 119      38.114   3.651 -44.201  1.00  0.00           H
    ATOM    485  HB2 LEU A 119      37.485   4.641 -42.045  1.00  0.00           H
    ATOM    486  HB3 LEU A 119      38.743   3.766 -41.203  1.00  0.00           H
    ATOM    487  HG  LEU A 119      37.601   1.657 -41.556  1.00  0.00           H
    ATOM    488 HD11 LEU A 119      35.527   2.125 -40.231  1.00  0.00           H
    ATOM    489 HD12 LEU A 119      36.921   3.001 -39.610  1.00  0.00           H
    ATOM    490 HD13 LEU A 119      35.702   3.842 -40.564  1.00  0.00           H
    ATOM    491 HD21 LEU A 119      35.326   1.546 -42.612  1.00  0.00           H
    ATOM    492 HD22 LEU A 119      35.535   3.194 -43.201  1.00  0.00           H
    ATOM    493 HD23 LEU A 119      36.595   1.896 -43.762  1.00  0.00           H
    ATOM    494  N   SER A 120      39.390   5.805 -44.279  1.00 13.72           N
    ATOM    495  CA  SER A 120      39.982   7.141 -44.340  1.00 14.14           C
    ATOM    496  C   SER A 120      39.534   7.968 -43.120  1.00 14.30           C
    ATOM    497  O   SER A 120      38.492   7.676 -42.536  1.00 12.50           O
    ATOM    498  CB  SER A 120      39.509   7.810 -45.652  1.00 14.23           C
    ATOM    499  OG  SER A 120      38.124   8.115 -45.658  1.00 14.78           O
    ATOM    500  H   SER A 120      38.560   5.662 -44.835  1.00  0.00           H
    ATOM    501  HA  SER A 120      41.070   7.052 -44.345  1.00  0.00           H
    ATOM    502  HB2 SER A 120      39.733   7.170 -46.507  1.00  0.00           H
    ATOM    503  HB3 SER A 120      40.058   8.740 -45.812  1.00  0.00           H
    ATOM    504  HG  SER A 120      37.897   8.480 -46.501  1.00  0.00           H
    ATOM    505  N   ASN A 121      40.297   9.017 -42.757  1.00 14.70           N
    ATOM    506  CA  ASN A 121      40.010   9.933 -41.640  1.00 17.71           C
    ATOM    507  C   ASN A 121      38.585  10.525 -41.684  1.00 19.33           C
    ATOM    508  O   ASN A 121      37.958  10.659 -40.635  1.00 19.84           O
    ATOM    509  CB  ASN A 121      41.079  11.050 -41.602  1.00 19.26           C
    ATOM    510  CG  ASN A 121      40.984  11.930 -40.347  1.00 22.57           C
    ATOM    511  OD1 ASN A 121      41.336  11.496 -39.254  1.00 23.97           O
    ATOM    512  ND2 ASN A 121      40.470  13.153 -40.495  1.00 25.48           N
    ATOM    513  H   ASN A 121      41.138   9.195 -43.287  1.00  0.00           H
    ATOM    514  HA  ASN A 121      40.092   9.341 -40.725  1.00  0.00           H
    ATOM    515  HB2 ASN A 121      42.075  10.605 -41.612  1.00  0.00           H
    ATOM    516  HB3 ASN A 121      41.015  11.672 -42.496  1.00  0.00           H
    ATOM    517 HD21 ASN A 121      40.385  13.757 -39.690  1.00  0.00           H
    ATOM    518 HD22 ASN A 121      40.175  13.476 -41.404  1.00  0.00           H
    ATOM    519  N   ASP A 122      38.074  10.831 -42.889  1.00 19.47           N
    ATOM    520  CA  ASP A 122      36.724  11.335 -43.133  1.00 19.03           C
    ATOM    521  C   ASP A 122      35.640  10.293 -42.795  1.00 17.49           C
    ATOM    522  O   ASP A 122      34.664  10.634 -42.131  1.00 18.12           O
    ATOM    523  CB  ASP A 122      36.630  11.800 -44.607  1.00 21.79           C
    ATOM    524  CG  ASP A 122      35.223  12.232 -45.033  1.00 25.49           C
    ATOM    525  OD1 ASP A 122      34.767  13.265 -44.497  1.00 25.45           O
    ATOM    526  OD2 ASP A 122      34.627  11.509 -45.863  1.00 25.48           O1-
    ATOM    527  H   ASP A 122      38.653  10.669 -43.700  1.00  0.00           H
    ATOM    528  HA  ASP A 122      36.565  12.202 -42.488  1.00  0.00           H
    ATOM    529  HB2 ASP A 122      37.307  12.638 -44.773  1.00  0.00           H
    ATOM    530  HB3 ASP A 122      36.964  11.003 -45.272  1.00  0.00           H
    ATOM    531  N   HID A 123      35.820   9.033 -43.231  1.00 15.15           N
    ATOM    532  CA  HID A 123      34.901   7.923 -42.967  1.00 13.83           C
    ATOM    533  C   HID A 123      34.883   7.519 -41.486  1.00 12.17           C
    ATOM    534  O   HID A 123      33.812   7.194 -40.986  1.00 11.34           O
    ATOM    535  CB  HID A 123      35.278   6.705 -43.837  1.00 16.42           C
    ATOM    536  CG  HID A 123      34.731   6.726 -45.246  1.00 19.33           C
    ATOM    537  ND1 HID A 123      34.457   7.889 -45.951  1.00 21.35           N
    ATOM    538  CD2 HID A 123      34.423   5.695 -46.108  1.00 19.50           C
    ATOM    539  CE1 HID A 123      34.006   7.524 -47.153  1.00 21.85           C
    ATOM    540  NE2 HID A 123      33.967   6.209 -47.324  1.00 20.78           N
    ATOM    541  H   HID A 123      36.656   8.824 -43.759  1.00  0.00           H
    ATOM    542  HA  HID A 123      33.886   8.239 -43.221  1.00  0.00           H
    ATOM    543  HB2 HID A 123      36.362   6.595 -43.887  1.00  0.00           H
    ATOM    544  HB3 HID A 123      34.910   5.783 -43.383  1.00  0.00           H
    ATOM    545  HD1 HID A 123      34.575   8.839 -45.623  1.00  0.00           H
    ATOM    546  HD2 HID A 123      34.496   4.630 -45.949  1.00  0.00           H
    ATOM    547  HE1 HID A 123      33.721   8.231 -47.917  1.00  0.00           H
    ATOM    548  N   ILE A 124      36.020   7.604 -40.771  1.00  9.97           N
    ATOM    549  CA  ILE A 124      36.110   7.390 -39.320  1.00  6.82           C
    ATOM    550  C   ILE A 124      35.357   8.504 -38.571  1.00  7.03           C
    ATOM    551  O   ILE A 124      34.572   8.201 -37.677  1.00  5.61           O
    ATOM    552  CB  ILE A 124      37.599   7.319 -38.869  1.00  3.80           C
    ATOM    553  CG1 ILE A 124      38.312   6.131 -39.550  1.00  4.90           C
    ATOM    554  CG2 ILE A 124      37.760   7.232 -37.332  1.00  1.85           C
    ATOM    555  CD1 ILE A 124      39.812   6.029 -39.273  1.00  1.85           C
    ATOM    556  H   ILE A 124      36.867   7.875 -41.253  1.00  0.00           H
    ATOM    557  HA  ILE A 124      35.628   6.439 -39.082  1.00  0.00           H
    ATOM    558  HB  ILE A 124      38.098   8.234 -39.196  1.00  0.00           H
    ATOM    559 HG12 ILE A 124      37.828   5.198 -39.269  1.00  0.00           H
    ATOM    560 HG13 ILE A 124      38.191   6.203 -40.623  1.00  0.00           H
    ATOM    561 HG21 ILE A 124      38.808   7.205 -37.034  1.00  0.00           H
    ATOM    562 HG22 ILE A 124      37.333   8.091 -36.815  1.00  0.00           H
    ATOM    563 HG23 ILE A 124      37.282   6.336 -36.941  1.00  0.00           H
    ATOM    564 HD11 ILE A 124      40.278   5.292 -39.927  1.00  0.00           H
    ATOM    565 HD12 ILE A 124      40.316   6.983 -39.431  1.00  0.00           H
    ATOM    566 HD13 ILE A 124      39.983   5.702 -38.252  1.00  0.00           H
    ATOM    567  N   CYS A 125      35.578   9.773 -38.961  1.00  8.11           N
    ATOM    568  CA  CYS A 125      34.969  10.969 -38.379  1.00  9.89           C
    ATOM    569  C   CYS A 125      33.443  10.995 -38.576  1.00  9.31           C
    ATOM    570  O   CYS A 125      32.716  11.290 -37.628  1.00 11.71           O
    ATOM    571  CB  CYS A 125      35.630  12.209 -39.020  1.00 11.48           C
    ATOM    572  SG  CYS A 125      35.165  13.744 -38.177  1.00 10.37           S
    ATOM    573  H   CYS A 125      36.246   9.924 -39.705  1.00  0.00           H
    ATOM    574  HA  CYS A 125      35.179  10.964 -37.309  1.00  0.00           H
    ATOM    575  HB2 CYS A 125      36.714  12.138 -38.960  1.00  0.00           H
    ATOM    576  HB3 CYS A 125      35.377  12.294 -40.078  1.00  0.00           H
    ATOM    577  HG  CYS A 125      33.850  13.623 -38.382  1.00  0.00           H
    ATOM    578  N   TYR A 126      32.962  10.654 -39.784  1.00 10.20           N
    ATOM    579  CA  TYR A 126      31.541  10.621 -40.118  1.00 11.35           C
    ATOM    580  C   TYR A 126      30.841   9.388 -39.517  1.00 10.94           C
    ATOM    581  O   TYR A 126      29.690   9.513 -39.110  1.00 12.26           O
    ATOM    582  CB  TYR A 126      31.361  10.629 -41.654  1.00 13.82           C
    ATOM    583  CG  TYR A 126      29.916  10.697 -42.141  1.00 17.01           C
    ATOM    584  CD1 TYR A 126      29.005  11.613 -41.569  1.00 16.17           C
    ATOM    585  CD2 TYR A 126      29.469   9.824 -43.157  1.00 16.91           C
    ATOM    586  CE1 TYR A 126      27.660  11.635 -41.983  1.00 16.37           C
    ATOM    587  CE2 TYR A 126      28.124   9.851 -43.578  1.00 17.74           C
    ATOM    588  CZ  TYR A 126      27.215  10.749 -42.983  1.00 17.89           C
    ATOM    589  OH  TYR A 126      25.905  10.756 -43.365  1.00 20.07           O
    ATOM    590  H   TYR A 126      33.619  10.432 -40.522  1.00  0.00           H
    ATOM    591  HA  TYR A 126      31.085  11.519 -39.699  1.00  0.00           H
    ATOM    592  HB2 TYR A 126      31.884  11.487 -42.074  1.00  0.00           H
    ATOM    593  HB3 TYR A 126      31.844   9.750 -42.084  1.00  0.00           H
    ATOM    594  HD1 TYR A 126      29.323  12.297 -40.797  1.00  0.00           H
    ATOM    595  HD2 TYR A 126      30.152   9.119 -43.609  1.00  0.00           H
    ATOM    596  HE1 TYR A 126      26.968  12.327 -41.526  1.00  0.00           H
    ATOM    597  HE2 TYR A 126      27.791   9.168 -44.347  1.00  0.00           H
    ATOM    598  HH  TYR A 126      25.680  10.048 -43.952  1.00  0.00           H
    ATOM    599  N   PHE A 127      31.511   8.226 -39.402  1.00 10.06           N
    ATOM    600  CA  PHE A 127      30.953   7.043 -38.735  1.00  9.53           C
    ATOM    601  C   PHE A 127      30.870   7.255 -37.217  1.00  7.27           C
    ATOM    602  O   PHE A 127      29.859   6.887 -36.627  1.00  8.11           O
    ATOM    603  CB  PHE A 127      31.807   5.788 -39.019  1.00  8.99           C
    ATOM    604  CG  PHE A 127      31.696   5.150 -40.400  1.00 11.45           C
    ATOM    605  CD1 PHE A 127      30.847   5.660 -41.411  1.00 10.73           C
    ATOM    606  CD2 PHE A 127      32.485   4.014 -40.676  1.00 10.97           C
    ATOM    607  CE1 PHE A 127      30.789   5.034 -42.670  1.00 10.91           C
    ATOM    608  CE2 PHE A 127      32.427   3.393 -41.937  1.00 10.38           C
    ATOM    609  CZ  PHE A 127      31.575   3.899 -42.935  1.00  9.67           C
    ATOM    610  H   PHE A 127      32.453   8.159 -39.766  1.00  0.00           H
    ATOM    611  HA  PHE A 127      29.933   6.869 -39.082  1.00  0.00           H
    ATOM    612  HB2 PHE A 127      32.858   6.002 -38.817  1.00  0.00           H
    ATOM    613  HB3 PHE A 127      31.533   5.001 -38.313  1.00  0.00           H
    ATOM    614  HD1 PHE A 127      30.237   6.534 -41.247  1.00  0.00           H
    ATOM    615  HD2 PHE A 127      33.145   3.617 -39.918  1.00  0.00           H
    ATOM    616  HE1 PHE A 127      30.142   5.427 -43.436  1.00  0.00           H
    ATOM    617  HE2 PHE A 127      33.044   2.530 -42.135  1.00  0.00           H
    ATOM    618  HZ  PHE A 127      31.527   3.422 -43.902  1.00  0.00           H
    ATOM    619  N   LEU A 128      31.875   7.903 -36.599  1.00  6.60           N
    ATOM    620  CA  LEU A 128      31.884   8.255 -35.177  1.00  6.01           C
    ATOM    621  C   LEU A 128      30.779   9.261 -34.825  1.00  5.08           C
    ATOM    622  O   LEU A 128      30.155   9.122 -33.774  1.00  7.33           O
    ATOM    623  CB  LEU A 128      33.265   8.818 -34.781  1.00  5.32           C
    ATOM    624  CG  LEU A 128      33.446   9.095 -33.271  1.00  4.13           C
    ATOM    625  CD1 LEU A 128      33.336   7.811 -32.433  1.00  4.14           C
    ATOM    626  CD2 LEU A 128      34.772   9.817 -33.007  1.00  7.87           C
    ATOM    627  H   LEU A 128      32.682   8.175 -37.146  1.00  0.00           H
    ATOM    628  HA  LEU A 128      31.701   7.341 -34.616  1.00  0.00           H
    ATOM    629  HB2 LEU A 128      34.038   8.114 -35.091  1.00  0.00           H
    ATOM    630  HB3 LEU A 128      33.458   9.732 -35.346  1.00  0.00           H
    ATOM    631  HG  LEU A 128      32.667   9.773 -32.925  1.00  0.00           H
    ATOM    632 HD11 LEU A 128      33.590   8.007 -31.392  1.00  0.00           H
    ATOM    633 HD12 LEU A 128      32.320   7.419 -32.444  1.00  0.00           H
    ATOM    634 HD13 LEU A 128      33.997   7.026 -32.798  1.00  0.00           H
    ATOM    635 HD21 LEU A 128      34.982   9.884 -31.940  1.00  0.00           H
    ATOM    636 HD22 LEU A 128      35.603   9.310 -33.492  1.00  0.00           H
    ATOM    637 HD23 LEU A 128      34.753  10.833 -33.394  1.00  0.00           H
    ATOM    638  N   TYR A 129      30.504  10.228 -35.719  1.00  3.37           N
    ATOM    639  CA  TYR A 129      29.407  11.183 -35.591  1.00  2.26           C
    ATOM    640  C   TYR A 129      28.042  10.483 -35.525  1.00  2.47           C
    ATOM    641  O   TYR A 129      27.240  10.819 -34.660  1.00  1.85           O
    ATOM    642  CB  TYR A 129      29.437  12.194 -36.757  1.00  1.85           C
    ATOM    643  CG  TYR A 129      28.267  13.160 -36.730  1.00  2.28           C
    ATOM    644  CD1 TYR A 129      28.183  14.111 -35.694  1.00  3.09           C
    ATOM    645  CD2 TYR A 129      27.224  13.057 -37.677  1.00  2.15           C
    ATOM    646  CE1 TYR A 129      27.059  14.945 -35.596  1.00  4.47           C
    ATOM    647  CE2 TYR A 129      26.102  13.904 -37.582  1.00  5.01           C
    ATOM    648  CZ  TYR A 129      26.018  14.848 -36.539  1.00  3.96           C
    ATOM    649  OH  TYR A 129      24.934  15.669 -36.437  1.00  3.92           O
    ATOM    650  H   TYR A 129      31.073  10.285 -36.552  1.00  0.00           H
    ATOM    651  HA  TYR A 129      29.551  11.727 -34.655  1.00  0.00           H
    ATOM    652  HB2 TYR A 129      30.361  12.773 -36.724  1.00  0.00           H
    ATOM    653  HB3 TYR A 129      29.446  11.675 -37.715  1.00  0.00           H
    ATOM    654  HD1 TYR A 129      28.971  14.187 -34.958  1.00  0.00           H
    ATOM    655  HD2 TYR A 129      27.273  12.319 -38.465  1.00  0.00           H
    ATOM    656  HE1 TYR A 129      27.006  15.658 -34.789  1.00  0.00           H
    ATOM    657  HE2 TYR A 129      25.303  13.823 -38.305  1.00  0.00           H
    ATOM    658  HH  TYR A 129      24.980  16.237 -35.678  1.00  0.00           H
    ATOM    659  N   GLN A 130      27.795   9.510 -36.414  1.00  1.85           N
    ATOM    660  CA  GLN A 130      26.537   8.777 -36.520  1.00  3.45           C
    ATOM    661  C   GLN A 130      26.321   7.802 -35.352  1.00  4.50           C
    ATOM    662  O   GLN A 130      25.186   7.648 -34.903  1.00  6.01           O
    ATOM    663  CB  GLN A 130      26.562   7.983 -37.828  1.00  3.71           C
    ATOM    664  CG  GLN A 130      26.485   8.889 -39.065  1.00  5.21           C
    ATOM    665  CD  GLN A 130      26.533   8.049 -40.329  1.00  5.72           C
    ATOM    666  OE1 GLN A 130      25.501   7.542 -40.747  1.00  7.12           O
    ATOM    667  NE2 GLN A 130      27.713   7.892 -40.930  1.00  6.54           N
    ATOM    668  H   GLN A 130      28.516   9.286 -37.086  1.00  0.00           H
    ATOM    669  HA  GLN A 130      25.705   9.484 -36.541  1.00  0.00           H
    ATOM    670  HB2 GLN A 130      25.722   7.289 -37.844  1.00  0.00           H
    ATOM    671  HB3 GLN A 130      27.463   7.367 -37.873  1.00  0.00           H
    ATOM    672  HG2 GLN A 130      25.547   9.444 -39.056  1.00  0.00           H
    ATOM    673  HG3 GLN A 130      27.267   9.644 -39.080  1.00  0.00           H
    ATOM    674 HE21 GLN A 130      27.773   7.339 -41.777  1.00  0.00           H
    ATOM    675 HE22 GLN A 130      28.533   8.348 -40.554  1.00  0.00           H
    ATOM    676  N   ILE A 131      27.408   7.203 -34.830  1.00  4.18           N
    ATOM    677  CA  ILE A 131      27.427   6.392 -33.610  1.00  5.29           C
    ATOM    678  C   ILE A 131      27.007   7.248 -32.402  1.00  5.18           C
    ATOM    679  O   ILE A 131      26.090   6.861 -31.684  1.00  3.35           O
    ATOM    680  CB  ILE A 131      28.832   5.743 -33.412  1.00  6.80           C
    ATOM    681  CG1 ILE A 131      29.124   4.669 -34.480  1.00  6.69           C
    ATOM    682  CG2 ILE A 131      29.017   5.120 -32.017  1.00  7.95           C
    ATOM    683  CD1 ILE A 131      30.613   4.324 -34.652  1.00  8.44           C
    ATOM    684  H   ILE A 131      28.299   7.374 -35.276  1.00  0.00           H
    ATOM    685  HA  ILE A 131      26.690   5.593 -33.721  1.00  0.00           H
    ATOM    686  HB  ILE A 131      29.580   6.530 -33.517  1.00  0.00           H
    ATOM    687 HG12 ILE A 131      28.572   3.762 -34.243  1.00  0.00           H
    ATOM    688 HG13 ILE A 131      28.739   4.998 -35.440  1.00  0.00           H
    ATOM    689 HG21 ILE A 131      29.976   4.610 -31.939  1.00  0.00           H
    ATOM    690 HG22 ILE A 131      29.006   5.875 -31.233  1.00  0.00           H
    ATOM    691 HG23 ILE A 131      28.236   4.391 -31.798  1.00  0.00           H
    ATOM    692 HD11 ILE A 131      30.760   3.645 -35.491  1.00  0.00           H
    ATOM    693 HD12 ILE A 131      31.211   5.209 -34.856  1.00  0.00           H
    ATOM    694 HD13 ILE A 131      31.025   3.841 -33.767  1.00  0.00           H
    ATOM    695  N   LEU A 132      27.643   8.415 -32.198  1.00  5.58           N
    ATOM    696  CA  LEU A 132      27.372   9.302 -31.067  1.00  6.03           C
    ATOM    697  C   LEU A 132      26.040  10.066 -31.196  1.00  6.47           C
    ATOM    698  O   LEU A 132      25.432  10.366 -30.172  1.00  7.31           O
    ATOM    699  CB  LEU A 132      28.509  10.330 -30.923  1.00  3.41           C
    ATOM    700  CG  LEU A 132      29.868   9.762 -30.461  1.00  2.95           C
    ATOM    701  CD1 LEU A 132      30.920  10.868 -30.532  1.00  1.85           C
    ATOM    702  CD2 LEU A 132      29.828   9.168 -29.045  1.00  1.85           C
    ATOM    703  H   LEU A 132      28.384   8.682 -32.834  1.00  0.00           H
    ATOM    704  HA  LEU A 132      27.323   8.703 -30.160  1.00  0.00           H
    ATOM    705  HB2 LEU A 132      28.635  10.854 -31.872  1.00  0.00           H
    ATOM    706  HB3 LEU A 132      28.213  11.087 -30.198  1.00  0.00           H
    ATOM    707  HG  LEU A 132      30.173   8.971 -31.145  1.00  0.00           H
    ATOM    708 HD11 LEU A 132      31.913  10.502 -30.266  1.00  0.00           H
    ATOM    709 HD12 LEU A 132      30.963  11.242 -31.550  1.00  0.00           H
    ATOM    710 HD13 LEU A 132      30.675  11.703 -29.876  1.00  0.00           H
    ATOM    711 HD21 LEU A 132      30.823   8.869 -28.716  1.00  0.00           H
    ATOM    712 HD22 LEU A 132      29.453   9.893 -28.321  1.00  0.00           H
    ATOM    713 HD23 LEU A 132      29.196   8.282 -28.995  1.00  0.00           H
    ATOM    714  N   ARG A 133      25.566  10.360 -32.420  1.00  6.14           N
    ATOM    715  CA  ARG A 133      24.284  11.021 -32.689  1.00  6.98           C
    ATOM    716  C   ARG A 133      23.110  10.071 -32.407  1.00  6.98           C
    ATOM    717  O   ARG A 133      22.110  10.495 -31.827  1.00  8.78           O
    ATOM    718  CB  ARG A 133      24.245  11.492 -34.162  1.00  6.18           C
    ATOM    719  CG  ARG A 133      22.916  12.138 -34.596  1.00  9.32           C
    ATOM    720  CD  ARG A 133      22.930  12.632 -36.044  1.00 10.76           C
    ATOM    721  NE  ARG A 133      21.599  13.106 -36.462  1.00 11.70           N
    ATOM    722  CZ  ARG A 133      20.917  12.734 -37.560  1.00 11.45           C
    ATOM    723  NH1 ARG A 133      21.387  11.817 -38.414  1.00 11.34           N
    ATOM    724  NH2 ARG A 133      19.731  13.296 -37.822  1.00 12.79           N1+
    ATOM    725  H   ARG A 133      26.139  10.124 -33.220  1.00  0.00           H
    ATOM    726  HA  ARG A 133      24.192  11.893 -32.038  1.00  0.00           H
    ATOM    727  HB2 ARG A 133      25.044  12.218 -34.317  1.00  0.00           H
    ATOM    728  HB3 ARG A 133      24.462  10.651 -34.823  1.00  0.00           H
    ATOM    729  HG2 ARG A 133      22.105  11.427 -34.500  1.00  0.00           H
    ATOM    730  HG3 ARG A 133      22.664  12.949 -33.916  1.00  0.00           H
    ATOM    731  HD2 ARG A 133      23.629  13.458 -36.141  1.00  0.00           H
    ATOM    732  HD3 ARG A 133      23.289  11.844 -36.703  1.00  0.00           H
    ATOM    733  HE  ARG A 133      21.222  13.871 -35.920  1.00  0.00           H
    ATOM    734 HH11 ARG A 133      20.840  11.568 -39.235  1.00  0.00           H
    ATOM    735 HH12 ARG A 133      22.242  11.319 -38.211  1.00  0.00           H
    ATOM    736 HH21 ARG A 133      19.219  13.019 -38.649  1.00  0.00           H
    ATOM    737 HH22 ARG A 133      19.380  14.047 -37.244  1.00  0.00           H
    ATOM    738  N   GLY A 134      23.256   8.787 -32.775  1.00  5.17           N
    ATOM    739  CA  GLY A 134      22.338   7.715 -32.417  1.00  6.98           C
    ATOM    740  C   GLY A 134      22.335   7.460 -30.909  1.00  7.70           C
    ATOM    741  O   GLY A 134      21.266   7.345 -30.320  1.00  7.46           O
    ATOM    742  H   GLY A 134      24.100   8.522 -33.266  1.00  0.00           H
    ATOM    743  HA2 GLY A 134      21.328   7.948 -32.751  1.00  0.00           H
    ATOM    744  HA3 GLY A 134      22.640   6.804 -32.934  1.00  0.00           H
    ATOM    745  N   LYS A 136      23.167   9.634 -28.483  1.00  4.84           N1+
    ATOM    746  CA  LYS A 136      22.581  10.810 -27.838  1.00  3.06           C
    ATOM    747  C   LYS A 136      21.058  10.677 -27.712  1.00  4.31           C
    ATOM    748  O   LYS A 136      20.515  11.010 -26.662  1.00  4.19           O
    ATOM    749  CB  LYS A 136      22.970  12.083 -28.626  1.00  2.65           C
    ATOM    750  CG  LYS A 136      22.217  13.370 -28.226  1.00  1.92           C
    ATOM    751  CD  LYS A 136      22.785  14.628 -28.895  1.00  3.16           C
    ATOM    752  CE  LYS A 136      21.828  15.825 -28.825  1.00  3.18           C
    ATOM    753  NZ  LYS A 136      22.442  17.032 -29.404  1.00  1.85           N1+
    ATOM    754  H   LYS A 136      23.616   9.780 -29.377  1.00  0.00           H
    ATOM    755  HA  LYS A 136      22.990  10.890 -26.830  1.00  0.00           H
    ATOM    756  HB2 LYS A 136      24.036  12.249 -28.494  1.00  0.00           H
    ATOM    757  HB3 LYS A 136      22.825  11.916 -29.694  1.00  0.00           H
    ATOM    758  HG2 LYS A 136      21.169  13.276 -28.511  1.00  0.00           H
    ATOM    759  HG3 LYS A 136      22.235  13.496 -27.143  1.00  0.00           H
    ATOM    760  HD2 LYS A 136      23.731  14.886 -28.419  1.00  0.00           H
    ATOM    761  HD3 LYS A 136      23.006  14.417 -29.943  1.00  0.00           H
    ATOM    762  HE2 LYS A 136      20.912  15.604 -29.375  1.00  0.00           H
    ATOM    763  HE3 LYS A 136      21.543  16.034 -27.793  1.00  0.00           H
    ATOM    764  HZ1 LYS A 136      21.758  17.773 -29.458  1.00  0.00           H
    ATOM    765  HZ2 LYS A 136      23.214  17.326 -28.824  1.00  0.00           H
    ATOM    766  HZ3 LYS A 136      22.799  16.833 -30.334  1.00  0.00           H
    ATOM    767  N   TYR A 137      20.386  10.159 -28.755  1.00  4.19           N
    ATOM    768  CA  TYR A 137      18.951   9.882 -28.759  1.00  4.10           C
    ATOM    769  C   TYR A 137      18.576   8.767 -27.758  1.00  4.78           C
    ATOM    770  O   TYR A 137      17.531   8.865 -27.116  1.00  6.74           O
    ATOM    771  CB  TYR A 137      18.510   9.516 -30.193  1.00  6.21           C
    ATOM    772  CG  TYR A 137      17.027   9.216 -30.344  1.00  4.27           C
    ATOM    773  CD1 TYR A 137      16.118  10.235 -30.695  1.00  2.44           C
    ATOM    774  CD2 TYR A 137      16.542   7.916 -30.093  1.00  4.76           C
    ATOM    775  CE1 TYR A 137      14.738   9.969 -30.768  1.00  1.85           C
    ATOM    776  CE2 TYR A 137      15.163   7.652 -30.157  1.00  5.99           C
    ATOM    777  CZ  TYR A 137      14.258   8.676 -30.487  1.00  4.02           C
    ATOM    778  OH  TYR A 137      12.920   8.414 -30.526  1.00  5.05           O
    ATOM    779  H   TYR A 137      20.910   9.904 -29.582  1.00  0.00           H
    ATOM    780  HA  TYR A 137      18.431  10.796 -28.468  1.00  0.00           H
    ATOM    781  HB2 TYR A 137      18.766  10.337 -30.863  1.00  0.00           H
    ATOM    782  HB3 TYR A 137      19.067   8.654 -30.559  1.00  0.00           H
    ATOM    783  HD1 TYR A 137      16.471  11.230 -30.897  1.00  0.00           H
    ATOM    784  HD2 TYR A 137      17.220   7.125 -29.811  1.00  0.00           H
    ATOM    785  HE1 TYR A 137      14.052  10.762 -31.026  1.00  0.00           H
    ATOM    786  HE2 TYR A 137      14.798   6.666 -29.936  1.00  0.00           H
    ATOM    787  HH  TYR A 137      12.388   9.187 -30.655  1.00  0.00           H
    ATOM    788  N   PRO A 150      31.425  -3.216 -33.093  1.00  2.61           N1+
    ATOM    789  CA  PRO A 150      31.402  -3.153 -34.567  1.00  3.37           C
    ATOM    790  C   PRO A 150      30.281  -3.971 -35.257  1.00  2.50           C
    ATOM    791  O   PRO A 150      29.720  -3.517 -36.253  1.00  1.85           O
    ATOM    792  CB  PRO A 150      32.793  -3.640 -34.997  1.00  3.95           C
    ATOM    793  CG  PRO A 150      33.698  -3.353 -33.819  1.00  3.99           C
    ATOM    794  CD  PRO A 150      32.782  -3.471 -32.607  1.00  2.20           C
    ATOM    795  HA  PRO A 150      31.290  -2.103 -34.841  1.00  0.00           H
    ATOM    796  HB2 PRO A 150      33.148  -3.156 -35.907  1.00  0.00           H
    ATOM    797  HB3 PRO A 150      32.801  -4.716 -35.156  1.00  0.00           H
    ATOM    798  HG2 PRO A 150      34.561  -4.018 -33.768  1.00  0.00           H
    ATOM    799  HG3 PRO A 150      34.072  -2.331 -33.892  1.00  0.00           H
    ATOM    800  HD2 PRO A 150      33.101  -2.808 -31.807  1.00  0.00           H
    ATOM    801  HD3 PRO A 150      32.831  -4.493 -32.234  1.00  0.00           H
    ATOM    802  N   SER A 151      29.913  -5.140 -34.700  1.00  1.87           N
    ATOM    803  CA  SER A 151      28.848  -6.022 -35.197  1.00  3.80           C
    ATOM    804  C   SER A 151      27.441  -5.436 -34.976  1.00  2.91           C
    ATOM    805  O   SER A 151      26.524  -5.739 -35.738  1.00  2.49           O
    ATOM    806  CB  SER A 151      28.925  -7.362 -34.437  1.00  5.24           C
    ATOM    807  OG  SER A 151      30.036  -8.120 -34.862  1.00  9.11           O
    ATOM    808  H   SER A 151      30.414  -5.453 -33.881  1.00  0.00           H
    ATOM    809  HA  SER A 151      28.982  -6.191 -36.268  1.00  0.00           H
    ATOM    810  HB2 SER A 151      28.033  -7.965 -34.616  1.00  0.00           H
    ATOM    811  HB3 SER A 151      28.987  -7.202 -33.359  1.00  0.00           H
    ATOM    812  HG  SER A 151      29.919  -8.342 -35.775  1.00  0.00           H
    ATOM    813  N   ASN A 152      27.286  -4.582 -33.951  1.00  3.37           N
    ATOM    814  CA  ASN A 152      26.053  -3.888 -33.576  1.00  4.65           C
    ATOM    815  C   ASN A 152      25.949  -2.496 -34.236  1.00  4.30           C
    ATOM    816  O   ASN A 152      25.146  -1.675 -33.800  1.00  3.46           O
    ATOM    817  CB  ASN A 152      25.990  -3.747 -32.031  1.00  5.15           C
    ATOM    818  CG  ASN A 152      25.637  -5.063 -31.332  1.00  8.55           C
    ATOM    819  OD1 ASN A 152      26.248  -6.095 -31.593  1.00 11.50           O
    ATOM    820  ND2 ASN A 152      24.624  -5.045 -30.466  1.00  8.48           N
    ATOM    821  H   ASN A 152      28.106  -4.357 -33.404  1.00  0.00           H
    ATOM    822  HA  ASN A 152      25.206  -4.483 -33.913  1.00  0.00           H
    ATOM    823  HB2 ASN A 152      26.938  -3.386 -31.632  1.00  0.00           H
    ATOM    824  HB3 ASN A 152      25.245  -3.010 -31.728  1.00  0.00           H
    ATOM    825 HD21 ASN A 152      24.367  -5.891 -29.978  1.00  0.00           H
    ATOM    826 HD22 ASN A 152      24.114  -4.189 -30.299  1.00  0.00           H
    ATOM    827  N   LEU A 153      26.732  -2.232 -35.295  1.00  4.34           N
    ATOM    828  CA  LEU A 153      26.732  -0.993 -36.068  1.00  3.56           C
    ATOM    829  C   LEU A 153      26.581  -1.361 -37.548  1.00  5.58           C
    ATOM    830  O   LEU A 153      27.574  -1.625 -38.221  1.00  5.43           O
    ATOM    831  CB  LEU A 153      28.020  -0.189 -35.780  1.00  4.62           C
    ATOM    832  CG  LEU A 153      28.235   0.233 -34.309  1.00  5.83           C
    ATOM    833  CD1 LEU A 153      29.630   0.841 -34.138  1.00  2.75           C
    ATOM    834  CD2 LEU A 153      27.153   1.183 -33.778  1.00  4.68           C
    ATOM    835  H   LEU A 153      27.368  -2.955 -35.600  1.00  0.00           H
    ATOM    836  HA  LEU A 153      25.878  -0.371 -35.803  1.00  0.00           H
    ATOM    837  HB2 LEU A 153      28.882  -0.782 -36.090  1.00  0.00           H
    ATOM    838  HB3 LEU A 153      28.033   0.702 -36.411  1.00  0.00           H
    ATOM    839  HG  LEU A 153      28.201  -0.652 -33.679  1.00  0.00           H
    ATOM    840 HD11 LEU A 153      29.747   1.298 -33.156  1.00  0.00           H
    ATOM    841 HD12 LEU A 153      30.398   0.076 -34.238  1.00  0.00           H
    ATOM    842 HD13 LEU A 153      29.819   1.607 -34.887  1.00  0.00           H
    ATOM    843 HD21 LEU A 153      27.379   1.513 -32.763  1.00  0.00           H
    ATOM    844 HD22 LEU A 153      27.056   2.066 -34.396  1.00  0.00           H
    ATOM    845 HD23 LEU A 153      26.179   0.703 -33.744  1.00  0.00           H
    ATOM    846  N   LEU A 154      25.334  -1.391 -38.046  1.00  6.25           N
    ATOM    847  CA  LEU A 154      24.977  -1.758 -39.419  1.00  6.33           C
    ATOM    848  C   LEU A 154      25.266  -0.595 -40.376  1.00  8.02           C
    ATOM    849  O   LEU A 154      25.101   0.559 -39.996  1.00  7.50           O
    ATOM    850  CB  LEU A 154      23.466  -2.049 -39.512  1.00  3.37           C
    ATOM    851  CG  LEU A 154      22.913  -3.090 -38.526  1.00  3.81           C
    ATOM    852  CD1 LEU A 154      21.413  -3.320 -38.769  1.00  6.06           C
    ATOM    853  CD2 LEU A 154      23.688  -4.418 -38.544  1.00  2.67           C
    ATOM    854  H   LEU A 154      24.576  -1.114 -37.436  1.00  0.00           H
    ATOM    855  HA  LEU A 154      25.540  -2.641 -39.721  1.00  0.00           H
    ATOM    856  HB2 LEU A 154      23.263  -2.398 -40.521  1.00  0.00           H
    ATOM    857  HB3 LEU A 154      22.900  -1.124 -39.388  1.00  0.00           H
    ATOM    858  HG  LEU A 154      23.018  -2.662 -37.537  1.00  0.00           H
    ATOM    859 HD11 LEU A 154      20.988  -4.009 -38.041  1.00  0.00           H
    ATOM    860 HD12 LEU A 154      20.857  -2.385 -38.693  1.00  0.00           H
    ATOM    861 HD13 LEU A 154      21.231  -3.741 -39.757  1.00  0.00           H
    ATOM    862 HD21 LEU A 154      23.225  -5.153 -37.887  1.00  0.00           H
    ATOM    863 HD22 LEU A 154      23.724  -4.843 -39.548  1.00  0.00           H
    ATOM    864 HD23 LEU A 154      24.712  -4.285 -38.196  1.00  0.00           H
    ATOM    865  N   LEU A 155      25.672  -0.911 -41.614  1.00  9.85           N
    ATOM    866  CA  LEU A 155      26.089   0.024 -42.659  1.00 12.21           C
    ATOM    867  C   LEU A 155      25.486  -0.382 -44.015  1.00 13.29           C
    ATOM    868  O   LEU A 155      25.514  -1.562 -44.362  1.00 13.26           O
    ATOM    869  CB  LEU A 155      27.631   0.089 -42.695  1.00 14.13           C
    ATOM    870  CG  LEU A 155      28.291   0.945 -41.592  1.00 13.93           C
    ATOM    871  CD1 LEU A 155      29.768   0.578 -41.414  1.00 14.64           C
    ATOM    872  CD2 LEU A 155      28.180   2.443 -41.904  1.00 16.81           C
    ATOM    873  H   LEU A 155      25.741  -1.894 -41.849  1.00  0.00           H
    ATOM    874  HA  LEU A 155      25.700   1.014 -42.430  1.00  0.00           H
    ATOM    875  HB2 LEU A 155      28.012  -0.931 -42.636  1.00  0.00           H
    ATOM    876  HB3 LEU A 155      27.958   0.463 -43.662  1.00  0.00           H
    ATOM    877  HG  LEU A 155      27.797   0.747 -40.641  1.00  0.00           H
    ATOM    878 HD11 LEU A 155      30.201   1.101 -40.561  1.00  0.00           H
    ATOM    879 HD12 LEU A 155      29.903  -0.491 -41.251  1.00  0.00           H
    ATOM    880 HD13 LEU A 155      30.337   0.864 -42.295  1.00  0.00           H
    ATOM    881 HD21 LEU A 155      28.651   3.043 -41.126  1.00  0.00           H
    ATOM    882 HD22 LEU A 155      28.674   2.690 -42.844  1.00  0.00           H
    ATOM    883 HD23 LEU A 155      27.146   2.761 -41.993  1.00  0.00           H
    ATOM    884  N   ASN A 156      24.961   0.591 -44.783  1.00 13.93           N
    ATOM    885  CA  ASN A 156      24.389   0.391 -46.124  1.00 13.73           C
    ATOM    886  C   ASN A 156      25.360   0.897 -47.217  1.00 14.25           C
    ATOM    887  O   ASN A 156      26.473   1.319 -46.914  1.00 12.41           O
    ATOM    888  CB  ASN A 156      23.008   1.105 -46.224  1.00 13.59           C
    ATOM    889  CG  ASN A 156      23.051   2.632 -46.391  1.00 14.77           C
    ATOM    890  OD1 ASN A 156      23.918   3.299 -45.836  1.00 18.05           O
    ATOM    891  ND2 ASN A 156      22.182   3.175 -47.246  1.00 15.40           N
    ATOM    892  H   ASN A 156      24.981   1.541 -44.436  1.00  0.00           H
    ATOM    893  HA  ASN A 156      24.234  -0.674 -46.292  1.00  0.00           H
    ATOM    894  HB2 ASN A 156      22.408   0.873 -45.345  1.00  0.00           H
    ATOM    895  HB3 ASN A 156      22.456   0.689 -47.067  1.00  0.00           H
    ATOM    896 HD21 ASN A 156      22.213   4.173 -47.414  1.00  0.00           H
    ATOM    897 HD22 ASN A 156      21.509   2.598 -47.728  1.00  0.00           H
    ATOM    898  N   THR A 157      24.917   0.891 -48.486  1.00 16.60           N
    ATOM    899  CA  THR A 157      25.633   1.352 -49.688  1.00 18.32           C
    ATOM    900  C   THR A 157      26.155   2.806 -49.702  1.00 17.79           C
    ATOM    901  O   THR A 157      27.030   3.100 -50.514  1.00 18.51           O
    ATOM    902  CB  THR A 157      24.761   1.085 -50.938  1.00 19.07           C
    ATOM    903  OG1 THR A 157      23.536   1.789 -50.864  1.00 20.42           O
    ATOM    904  CG2 THR A 157      24.482  -0.400 -51.190  1.00 19.66           C
    ATOM    905  H   THR A 157      23.986   0.529 -48.646  1.00  0.00           H
    ATOM    906  HA  THR A 157      26.524   0.738 -49.789  1.00  0.00           H
    ATOM    907  HB  THR A 157      25.287   1.462 -51.817  1.00  0.00           H
    ATOM    908 HG21 THR A 157      23.970  -0.550 -52.140  1.00  0.00           H
    ATOM    909 HG22 THR A 157      25.407  -0.976 -51.218  1.00  0.00           H
    ATOM    910 HG23 THR A 157      23.849  -0.814 -50.409  1.00  0.00           H
    ATOM    911  HG1 THR A 157      23.047   1.628 -51.657  1.00  0.00           H
    ATOM    912  N   THR A 158      25.655   3.708 -48.842  1.00 17.81           N
    ATOM    913  CA  THR A 158      25.936   5.150 -48.894  1.00 19.66           C
    ATOM    914  C   THR A 158      26.477   5.705 -47.571  1.00 19.04           C
    ATOM    915  O   THR A 158      26.238   6.867 -47.249  1.00 18.58           O
    ATOM    916  CB  THR A 158      24.746   5.928 -49.516  1.00 20.27           C
    ATOM    917  OG1 THR A 158      23.564   5.767 -48.756  1.00 21.28           O
    ATOM    918  CG2 THR A 158      24.456   5.551 -50.976  1.00 20.64           C
    ATOM    919  H   THR A 158      24.946   3.405 -48.188  1.00  0.00           H
    ATOM    920  HA  THR A 158      26.775   5.324 -49.567  1.00  0.00           H
    ATOM    921  HB  THR A 158      24.984   6.992 -49.512  1.00  0.00           H
    ATOM    922 HG21 THR A 158      23.684   6.195 -51.399  1.00  0.00           H
    ATOM    923 HG22 THR A 158      25.349   5.657 -51.593  1.00  0.00           H
    ATOM    924 HG23 THR A 158      24.109   4.521 -51.066  1.00  0.00           H
    ATOM    925  HG1 THR A 158      23.680   6.157 -47.897  1.00  0.00           H
    ATOM    926  N   CYS A 159      27.226   4.889 -46.807  1.00 19.61           N
    ATOM    927  CA  CYS A 159      27.976   5.285 -45.606  1.00 19.55           C
    ATOM    928  C   CYS A 159      27.093   5.581 -44.377  1.00 18.22           C
    ATOM    929  O   CYS A 159      27.601   6.083 -43.373  1.00 17.92           O
    ATOM    930  CB  CYS A 159      29.025   6.401 -45.897  1.00 19.74           C
    ATOM    931  SG  CYS A 159      30.559   5.689 -46.549  1.00 24.25           S
    ATOM    932  H   CYS A 159      27.344   3.937 -47.122  1.00  0.00           H
    ATOM    933  HA  CYS A 159      28.524   4.391 -45.302  1.00  0.00           H
    ATOM    934  HB2 CYS A 159      28.660   7.162 -46.583  1.00  0.00           H
    ATOM    935  HB3 CYS A 159      29.305   6.939 -44.992  1.00  0.00           H
    ATOM    936  HG  CYS A 159      31.229   6.842 -46.637  1.00  0.00           H
    ATOM    937  N   ASP A 160      25.780   5.287 -44.444  1.00 17.11           N
    ATOM    938  CA  ASP A 160      24.822   5.571 -43.376  1.00 14.69           C
    ATOM    939  C   ASP A 160      24.808   4.404 -42.390  1.00 12.23           C
    ATOM    940  O   ASP A 160      24.562   3.254 -42.758  1.00 10.88           O
    ATOM    941  CB  ASP A 160      23.388   5.795 -43.903  1.00 17.49           C
    ATOM    942  CG  ASP A 160      23.319   6.830 -45.021  1.00 19.81           C
    ATOM    943  OD1 ASP A 160      23.802   7.960 -44.782  1.00 19.20           O
    ATOM    944  OD2 ASP A 160      22.793   6.469 -46.096  1.00 20.99           O1-
    ATOM    945  H   ASP A 160      25.417   4.867 -45.289  1.00  0.00           H
    ATOM    946  HA  ASP A 160      25.129   6.492 -42.885  1.00  0.00           H
    ATOM    947  HB2 ASP A 160      22.924   4.869 -44.237  1.00  0.00           H
    ATOM    948  HB3 ASP A 160      22.757   6.153 -43.088  1.00  0.00           H
    ATOM    949  N   LEU A 161      25.111   4.740 -41.134  1.00  8.98           N
    ATOM    950  CA  LEU A 161      25.310   3.839 -40.015  1.00  7.41           C
    ATOM    951  C   LEU A 161      24.061   3.856 -39.127  1.00  7.74           C
    ATOM    952  O   LEU A 161      23.497   4.922 -38.882  1.00  9.41           O
    ATOM    953  CB  LEU A 161      26.537   4.374 -39.249  1.00  5.25           C
    ATOM    954  CG  LEU A 161      27.085   3.479 -38.119  1.00  1.85           C
    ATOM    955  CD1 LEU A 161      28.592   3.724 -37.944  1.00  1.85           C
    ATOM    956  CD2 LEU A 161      26.346   3.709 -36.789  1.00  1.85           C
    ATOM    957  H   LEU A 161      25.269   5.723 -40.949  1.00  0.00           H
    ATOM    958  HA  LEU A 161      25.522   2.844 -40.387  1.00  0.00           H
    ATOM    959  HB2 LEU A 161      27.331   4.520 -39.982  1.00  0.00           H
    ATOM    960  HB3 LEU A 161      26.340   5.373 -38.868  1.00  0.00           H
    ATOM    961  HG  LEU A 161      26.972   2.431 -38.402  1.00  0.00           H
    ATOM    962 HD11 LEU A 161      29.002   3.116 -37.138  1.00  0.00           H
    ATOM    963 HD12 LEU A 161      29.145   3.468 -38.848  1.00  0.00           H
    ATOM    964 HD13 LEU A 161      28.795   4.770 -37.723  1.00  0.00           H
    ATOM    965 HD21 LEU A 161      26.986   3.479 -35.947  1.00  0.00           H
    ATOM    966 HD22 LEU A 161      26.026   4.744 -36.667  1.00  0.00           H
    ATOM    967 HD23 LEU A 161      25.469   3.072 -36.699  1.00  0.00           H
    ATOM    968  N   LYS A 162      23.663   2.676 -38.624  1.00  5.79           N
    ATOM    969  CA  LYS A 162      22.574   2.475 -37.669  1.00  6.58           C
    ATOM    970  C   LYS A 162      23.027   1.584 -36.507  1.00  5.73           C
    ATOM    971  O   LYS A 162      23.604   0.524 -36.741  1.00  6.34           O
    ATOM    972  CB  LYS A 162      21.333   1.878 -38.363  1.00  7.14           C
    ATOM    973  CG  LYS A 162      20.725   2.758 -39.466  1.00  6.73           C
    ATOM    974  CD  LYS A 162      19.320   2.279 -39.866  1.00  3.71           C
    ATOM    975  CE  LYS A 162      18.705   3.039 -41.048  1.00  4.52           C
    ATOM    976  NZ  LYS A 162      19.342   2.673 -42.323  1.00  3.90           N1+
    ATOM    977  H   LYS A 162      24.187   1.852 -38.891  1.00  0.00           H
    ATOM    978  HA  LYS A 162      22.303   3.436 -37.245  1.00  0.00           H
    ATOM    979  HB2 LYS A 162      21.570   0.897 -38.777  1.00  0.00           H
    ATOM    980  HB3 LYS A 162      20.573   1.702 -37.602  1.00  0.00           H
    ATOM    981  HG2 LYS A 162      20.668   3.786 -39.115  1.00  0.00           H
    ATOM    982  HG3 LYS A 162      21.386   2.766 -40.332  1.00  0.00           H
    ATOM    983  HD2 LYS A 162      19.335   1.210 -40.083  1.00  0.00           H
    ATOM    984  HD3 LYS A 162      18.663   2.405 -39.007  1.00  0.00           H
    ATOM    985  HE2 LYS A 162      18.778   4.116 -40.904  1.00  0.00           H
    ATOM    986  HE3 LYS A 162      17.643   2.801 -41.122  1.00  0.00           H
    ATOM    987  HZ1 LYS A 162      20.321   2.917 -42.294  1.00  0.00           H
    ATOM    988  HZ2 LYS A 162      19.249   1.679 -42.479  1.00  0.00           H
    ATOM    989  HZ3 LYS A 162      18.895   3.172 -43.079  1.00  0.00           H
    ATOM    990  N   ILE A 163      22.714   1.980 -35.261  1.00  4.92           N
    ATOM    991  CA  ILE A 163      22.905   1.172 -34.049  1.00  5.78           C
    ATOM    992  C   ILE A 163      21.872   0.026 -34.043  1.00  7.18           C
    ATOM    993  O   ILE A 163      20.736   0.200 -34.485  1.00  4.76           O
    ATOM    994  CB  ILE A 163      22.778   2.066 -32.778  1.00  6.37           C
    ATOM    995  CG1 ILE A 163      23.855   3.178 -32.771  1.00  7.28           C
    ATOM    996  CG2 ILE A 163      22.860   1.248 -31.468  1.00  5.77           C
    ATOM    997  CD1 ILE A 163      23.680   4.216 -31.658  1.00  9.39           C
    ATOM    998  H   ILE A 163      22.246   2.870 -35.144  1.00  0.00           H
    ATOM    999  HA  ILE A 163      23.906   0.741 -34.076  1.00  0.00           H
    ATOM   1000  HB  ILE A 163      21.799   2.548 -32.803  1.00  0.00           H
    ATOM   1001 HG12 ILE A 163      24.849   2.735 -32.698  1.00  0.00           H
    ATOM   1002 HG13 ILE A 163      23.841   3.719 -33.717  1.00  0.00           H
    ATOM   1003 HG21 ILE A 163      22.778   1.878 -30.584  1.00  0.00           H
    ATOM   1004 HG22 ILE A 163      22.055   0.521 -31.382  1.00  0.00           H
    ATOM   1005 HG23 ILE A 163      23.802   0.703 -31.401  1.00  0.00           H
    ATOM   1006 HD11 ILE A 163      24.360   5.048 -31.813  1.00  0.00           H
    ATOM   1007 HD12 ILE A 163      22.666   4.616 -31.657  1.00  0.00           H
    ATOM   1008 HD13 ILE A 163      23.891   3.808 -30.670  1.00  0.00           H
    ATOM   1009  N   TRP A 190      35.405  -7.194 -25.342  1.00  7.07           N1+
    ATOM   1010  CA  TRP A 190      36.501  -6.286 -25.696  1.00  6.10           C
    ATOM   1011  C   TRP A 190      36.066  -4.807 -25.756  1.00  5.04           C
    ATOM   1012  O   TRP A 190      36.916  -3.928 -25.895  1.00  5.92           O
    ATOM   1013  CB  TRP A 190      37.035  -6.702 -27.086  1.00  6.67           C
    ATOM   1014  CG  TRP A 190      37.527  -8.113 -27.271  1.00  7.20           C
    ATOM   1015  CD1 TRP A 190      37.769  -9.022 -26.296  1.00  5.68           C
    ATOM   1016  CD2 TRP A 190      37.843  -8.794 -28.523  1.00  5.27           C
    ATOM   1017  NE1 TRP A 190      38.257 -10.187 -26.847  1.00  5.79           N
    ATOM   1018  CE2 TRP A 190      38.325 -10.106 -28.221  1.00  6.23           C
    ATOM   1019  CE3 TRP A 190      37.772  -8.439 -29.891  1.00  5.42           C
    ATOM   1020  CZ2 TRP A 190      38.740 -11.005 -29.218  1.00  7.04           C
    ATOM   1021  CZ3 TRP A 190      38.164  -9.340 -30.901  1.00  5.02           C
    ATOM   1022  CH2 TRP A 190      38.656 -10.617 -30.567  1.00  5.60           C
    ATOM   1023  H   TRP A 190      35.071  -7.792 -26.083  1.00  0.00           H
    ATOM   1024  HA  TRP A 190      37.298  -6.381 -24.959  1.00  0.00           H
    ATOM   1025  HB2 TRP A 190      37.858  -6.046 -27.374  1.00  0.00           H
    ATOM   1026  HB3 TRP A 190      36.255  -6.543 -27.831  1.00  0.00           H
    ATOM   1027  HD1 TRP A 190      37.620  -8.846 -25.244  1.00  0.00           H
    ATOM   1028  HE1 TRP A 190      38.541 -10.981 -26.286  1.00  0.00           H
    ATOM   1029  HE3 TRP A 190      37.406  -7.460 -30.166  1.00  0.00           H
    ATOM   1030  HZ2 TRP A 190      39.106 -11.986 -28.952  1.00  0.00           H
    ATOM   1031  HZ3 TRP A 190      38.093  -9.046 -31.938  1.00  0.00           H
    ATOM   1032  HH2 TRP A 190      38.962 -11.300 -31.346  1.00  0.00           H
    ATOM   1033  N   ILE A 215      35.637   1.100 -30.222  1.00  5.10           N1+
    ATOM   1034  CA  ILE A 215      35.255   1.055 -31.636  1.00  4.38           C
    ATOM   1035  C   ILE A 215      36.074   2.073 -32.454  1.00  5.59           C
    ATOM   1036  O   ILE A 215      36.496   1.729 -33.554  1.00  4.97           O
    ATOM   1037  CB  ILE A 215      33.725   1.318 -31.781  1.00  3.26           C
    ATOM   1038  CG1 ILE A 215      32.894   0.145 -31.218  1.00  2.00           C
    ATOM   1039  CG2 ILE A 215      33.273   1.585 -33.234  1.00  2.71           C
    ATOM   1040  CD1 ILE A 215      31.444   0.512 -30.858  1.00  1.85           C
    ATOM   1041  H   ILE A 215      34.972   1.514 -29.583  1.00  0.00           H
    ATOM   1042  HA  ILE A 215      35.475   0.059 -32.025  1.00  0.00           H
    ATOM   1043  HB  ILE A 215      33.494   2.211 -31.202  1.00  0.00           H
    ATOM   1044 HG12 ILE A 215      32.883  -0.654 -31.955  1.00  0.00           H
    ATOM   1045 HG13 ILE A 215      33.371  -0.286 -30.340  1.00  0.00           H
    ATOM   1046 HG21 ILE A 215      32.190   1.661 -33.306  1.00  0.00           H
    ATOM   1047 HG22 ILE A 215      33.669   2.521 -33.625  1.00  0.00           H
    ATOM   1048 HG23 ILE A 215      33.595   0.780 -33.895  1.00  0.00           H
    ATOM   1049 HD11 ILE A 215      31.223   0.246 -29.825  1.00  0.00           H
    ATOM   1050 HD12 ILE A 215      31.240   1.576 -30.974  1.00  0.00           H
    ATOM   1051 HD13 ILE A 215      30.740  -0.029 -31.489  1.00  0.00           H
    ATOM   1052  N   GLU A 218      39.631   0.834 -32.911  1.00  5.01           N1+
    ATOM   1053  CA  GLU A 218      39.599  -0.230 -33.904  1.00  4.19           C
    ATOM   1054  C   GLU A 218      39.284   0.264 -35.325  1.00  5.13           C
    ATOM   1055  O   GLU A 218      39.691  -0.392 -36.281  1.00  7.25           O
    ATOM   1056  CB  GLU A 218      38.583  -1.281 -33.413  1.00  3.90           C
    ATOM   1057  CG  GLU A 218      38.416  -2.522 -34.316  1.00  4.29           C
    ATOM   1058  CD  GLU A 218      37.728  -3.705 -33.637  1.00  3.19           C
    ATOM   1059  OE1 GLU A 218      37.513  -3.641 -32.410  1.00  8.14           O
    ATOM   1060  OE2 GLU A 218      37.422  -4.673 -34.366  1.00  5.41           O1-
    ATOM   1061  H   GLU A 218      38.806   0.952 -32.340  1.00  0.00           H
    ATOM   1062  HA  GLU A 218      40.587  -0.669 -33.964  1.00  0.00           H
    ATOM   1063  HB2 GLU A 218      38.910  -1.590 -32.419  1.00  0.00           H
    ATOM   1064  HB3 GLU A 218      37.606  -0.816 -33.271  1.00  0.00           H
    ATOM   1065  HG2 GLU A 218      37.832  -2.271 -35.192  1.00  0.00           H
    ATOM   1066  HG3 GLU A 218      39.383  -2.860 -34.679  1.00  0.00           H
    ATOM   1067  N   MET A 219      38.650   1.438 -35.481  1.00  4.80           N
    ATOM   1068  CA  MET A 219      38.462   2.104 -36.771  1.00  5.58           C
    ATOM   1069  C   MET A 219      39.778   2.715 -37.293  1.00  6.34           C
    ATOM   1070  O   MET A 219      39.987   2.746 -38.505  1.00  7.11           O
    ATOM   1071  CB  MET A 219      37.400   3.223 -36.650  1.00  2.83           C
    ATOM   1072  CG  MET A 219      35.951   2.727 -36.775  1.00  2.95           C
    ATOM   1073  SD  MET A 219      34.722   4.009 -37.160  1.00  6.97           S
    ATOM   1074  CE  MET A 219      34.707   4.976 -35.628  1.00  1.85           C
    ATOM   1075  H   MET A 219      38.324   1.921 -34.654  1.00  0.00           H
    ATOM   1076  HA  MET A 219      38.134   1.371 -37.504  1.00  0.00           H
    ATOM   1077  HB2 MET A 219      37.528   3.793 -35.731  1.00  0.00           H
    ATOM   1078  HB3 MET A 219      37.552   3.942 -37.453  1.00  0.00           H
    ATOM   1079  HG2 MET A 219      35.892   2.003 -37.586  1.00  0.00           H
    ATOM   1080  HG3 MET A 219      35.641   2.196 -35.878  1.00  0.00           H
    ATOM   1081  HE1 MET A 219      33.932   5.740 -35.682  1.00  0.00           H
    ATOM   1082  HE2 MET A 219      34.502   4.337 -34.771  1.00  0.00           H
    ATOM   1083  HE3 MET A 219      35.664   5.471 -35.474  1.00  0.00           H
    ATOM   1084  N   ASN A 222      42.281  -0.973 -37.605  1.00 12.03           N1+
    ATOM   1085  CA  ASN A 222      41.666  -2.201 -38.134  1.00 13.72           C
    ATOM   1086  C   ASN A 222      41.852  -3.404 -37.187  1.00 15.74           C
    ATOM   1087  O   ASN A 222      41.852  -4.553 -37.628  1.00 17.13           O
    ATOM   1088  CB  ASN A 222      42.078  -2.483 -39.602  1.00 14.55           C
    ATOM   1089  CG  ASN A 222      41.602  -1.382 -40.552  1.00 14.12           C
    ATOM   1090  OD1 ASN A 222      42.283  -0.380 -40.759  1.00 18.12           O
    ATOM   1091  ND2 ASN A 222      40.419  -1.580 -41.129  1.00 14.43           N
    ATOM   1092  H   ASN A 222      41.644  -0.242 -37.322  1.00  0.00           H
    ATOM   1093  HA  ASN A 222      40.591  -2.021 -38.133  1.00  0.00           H
    ATOM   1094  HB2 ASN A 222      43.156  -2.599 -39.705  1.00  0.00           H
    ATOM   1095  HB3 ASN A 222      41.654  -3.429 -39.941  1.00  0.00           H
    ATOM   1096 HD21 ASN A 222      40.055  -0.915 -41.797  1.00  0.00           H
    ATOM   1097 HD22 ASN A 222      39.910  -2.434 -40.939  1.00  0.00           H
    ATOM   1098  N   ARG A 223      42.050  -3.121 -35.889  1.00 16.81           N
    ATOM   1099  CA  ARG A 223      42.438  -4.050 -34.837  1.00 15.93           C
    ATOM   1100  C   ARG A 223      41.779  -3.654 -33.511  1.00 13.41           C
    ATOM   1101  O   ARG A 223      41.836  -2.476 -33.166  1.00 14.73           O
    ATOM   1102  CB  ARG A 223      43.976  -4.067 -34.699  1.00 20.37           C
    ATOM   1103  CG  ARG A 223      44.702  -4.923 -35.754  1.00 24.84           C
    ATOM   1104  CD  ARG A 223      45.649  -5.949 -35.120  1.00 25.96           C
    ATOM   1105  NE  ARG A 223      44.893  -6.890 -34.279  1.00 30.13           N
    ATOM   1106  CZ  ARG A 223      45.384  -7.962 -33.640  1.00 32.47           C
    ATOM   1107  NH1 ARG A 223      46.673  -8.311 -33.757  1.00 33.96           N
    ATOM   1108  NH2 ARG A 223      44.567  -8.692 -32.870  1.00 31.81           N1+
    ATOM   1109  H   ARG A 223      42.016  -2.146 -35.625  1.00  0.00           H
    ATOM   1110  HA  ARG A 223      42.091  -5.042 -35.122  1.00  0.00           H
    ATOM   1111  HB2 ARG A 223      44.363  -3.048 -34.738  1.00  0.00           H
    ATOM   1112  HB3 ARG A 223      44.240  -4.415 -33.699  1.00  0.00           H
    ATOM   1113  HG2 ARG A 223      43.989  -5.456 -36.383  1.00  0.00           H
    ATOM   1114  HG3 ARG A 223      45.258  -4.273 -36.429  1.00  0.00           H
    ATOM   1115  HD2 ARG A 223      46.409  -5.451 -34.516  1.00  0.00           H
    ATOM   1116  HD3 ARG A 223      46.167  -6.505 -35.901  1.00  0.00           H
    ATOM   1117  HE  ARG A 223      43.915  -6.671 -34.160  1.00  0.00           H
    ATOM   1118 HH11 ARG A 223      47.036  -9.120 -33.276  1.00  0.00           H
    ATOM   1119 HH12 ARG A 223      47.290  -7.762 -34.337  1.00  0.00           H
    ATOM   1120 HH21 ARG A 223      44.914  -9.506 -32.383  1.00  0.00           H
    ATOM   1121 HH22 ARG A 223      43.595  -8.439 -32.772  1.00  0.00           H
    ATOM   1122  N   PRO A 224      41.240  -4.602 -32.709  1.00 11.18           N
    ATOM   1123  CA  PRO A 224      40.685  -4.304 -31.383  1.00  9.93           C
    ATOM   1124  C   PRO A 224      41.804  -3.889 -30.420  1.00  8.56           C
    ATOM   1125  O   PRO A 224      42.893  -4.464 -30.450  1.00  8.39           O
    ATOM   1126  CB  PRO A 224      39.979  -5.595 -30.946  1.00 11.08           C
    ATOM   1127  CG  PRO A 224      40.710  -6.698 -31.695  1.00 12.28           C
    ATOM   1128  CD  PRO A 224      41.202  -6.039 -32.982  1.00 12.26           C
    ATOM   1129  HA  PRO A 224      39.959  -3.493 -31.452  1.00  0.00           H
    ATOM   1130  HB2 PRO A 224      39.981  -5.753 -29.866  1.00  0.00           H
    ATOM   1131  HB3 PRO A 224      38.937  -5.572 -31.265  1.00  0.00           H
    ATOM   1132  HG2 PRO A 224      41.574  -7.016 -31.112  1.00  0.00           H
    ATOM   1133  HG3 PRO A 224      40.094  -7.575 -31.880  1.00  0.00           H
    ATOM   1134  HD2 PRO A 224      42.178  -6.430 -33.268  1.00  0.00           H
    ATOM   1135  HD3 PRO A 224      40.499  -6.227 -33.795  1.00  0.00           H
    ATOM   1136  N   LEU A 306      30.256  14.761 -28.985  1.00 11.99           N1+
    ATOM   1137  CA  LEU A 306      29.681  15.365 -30.186  1.00 12.39           C
    ATOM   1138  C   LEU A 306      30.189  16.800 -30.401  1.00 14.63           C
    ATOM   1139  O   LEU A 306      30.537  17.155 -31.523  1.00 14.71           O
    ATOM   1140  CB  LEU A 306      28.141  15.368 -30.063  1.00 10.75           C
    ATOM   1141  CG  LEU A 306      27.454  14.025 -30.365  1.00 11.28           C
    ATOM   1142  CD1 LEU A 306      26.011  14.039 -29.852  1.00  8.75           C
    ATOM   1143  CD2 LEU A 306      27.472  13.698 -31.868  1.00 12.90           C
    ATOM   1144  H   LEU A 306      29.615  14.560 -28.229  1.00  0.00           H
    ATOM   1145  HA  LEU A 306      29.986  14.777 -31.053  1.00  0.00           H
    ATOM   1146  HB2 LEU A 306      27.865  15.713 -29.065  1.00  0.00           H
    ATOM   1147  HB3 LEU A 306      27.715  16.095 -30.745  1.00  0.00           H
    ATOM   1148  HG  LEU A 306      27.987  13.241 -29.832  1.00  0.00           H
    ATOM   1149 HD11 LEU A 306      25.502  13.107 -30.094  1.00  0.00           H
    ATOM   1150 HD12 LEU A 306      25.978  14.162 -28.769  1.00  0.00           H
    ATOM   1151 HD13 LEU A 306      25.444  14.854 -30.302  1.00  0.00           H
    ATOM   1152 HD21 LEU A 306      26.943  12.770 -32.078  1.00  0.00           H
    ATOM   1153 HD22 LEU A 306      26.988  14.484 -32.449  1.00  0.00           H
    ATOM   1154 HD23 LEU A 306      28.486  13.580 -32.248  1.00  0.00           H
    ATOM   1155  N   ALA A 307      30.284  17.595 -29.320  1.00 16.59           N
    ATOM   1156  CA  ALA A 307      30.760  18.979 -29.311  1.00 17.09           C
    ATOM   1157  C   ALA A 307      32.298  19.107 -29.227  1.00 17.84           C
    ATOM   1158  O   ALA A 307      32.799  20.190 -28.923  1.00 18.34           O
    ATOM   1159  CB  ALA A 307      30.132  19.673 -28.089  1.00 18.66           C
    ATOM   1160  H   ALA A 307      29.991  17.213 -28.432  1.00  0.00           H
    ATOM   1161  HA  ALA A 307      30.433  19.485 -30.219  1.00  0.00           H
    ATOM   1162  HB1 ALA A 307      30.371  20.737 -28.061  1.00  0.00           H
    ATOM   1163  HB2 ALA A 307      29.046  19.596 -28.112  1.00  0.00           H
    ATOM   1164  HB3 ALA A 307      30.475  19.230 -27.153  1.00  0.00           H
    ATOM   1165  N   TYR A 310      37.172  17.431 -33.119  1.00 17.39           N1+
    ATOM   1166  CA  TYR A 310      37.288  16.434 -34.190  1.00 15.99           C
    ATOM   1167  C   TYR A 310      35.990  16.354 -35.007  1.00 14.87           C
    ATOM   1168  O   TYR A 310      36.036  16.335 -36.235  1.00 15.81           O
    ATOM   1169  CB  TYR A 310      37.580  15.042 -33.581  1.00 13.94           C
    ATOM   1170  CG  TYR A 310      37.816  13.926 -34.591  1.00 12.43           C
    ATOM   1171  CD1 TYR A 310      38.799  14.066 -35.594  1.00 11.69           C
    ATOM   1172  CD2 TYR A 310      37.047  12.744 -34.539  1.00 11.36           C
    ATOM   1173  CE1 TYR A 310      39.006  13.042 -36.537  1.00 11.10           C
    ATOM   1174  CE2 TYR A 310      37.268  11.709 -35.469  1.00  9.99           C
    ATOM   1175  CZ  TYR A 310      38.251  11.856 -36.467  1.00 10.65           C
    ATOM   1176  OH  TYR A 310      38.476  10.848 -37.360  1.00  7.65           O
    ATOM   1177  H   TYR A 310      37.117  17.089 -32.167  1.00  0.00           H
    ATOM   1178  HA  TYR A 310      38.107  16.728 -34.848  1.00  0.00           H
    ATOM   1179  HB2 TYR A 310      38.458  15.092 -32.944  1.00  0.00           H
    ATOM   1180  HB3 TYR A 310      36.766  14.752 -32.914  1.00  0.00           H
    ATOM   1181  HD1 TYR A 310      39.398  14.962 -35.649  1.00  0.00           H
    ATOM   1182  HD2 TYR A 310      36.284  12.626 -33.785  1.00  0.00           H
    ATOM   1183  HE1 TYR A 310      39.764  13.162 -37.298  1.00  0.00           H
    ATOM   1184  HE2 TYR A 310      36.683  10.803 -35.414  1.00  0.00           H
    ATOM   1185  HH  TYR A 310      39.156  11.049 -37.986  1.00  0.00           H
    ATOM   1186  N   LEU A 311      34.846  16.332 -34.308  1.00 15.65           N
    ATOM   1187  CA  LEU A 311      33.508  16.188 -34.870  1.00 17.15           C
    ATOM   1188  C   LEU A 311      32.813  17.539 -35.104  1.00 18.50           C
    ATOM   1189  O   LEU A 311      31.639  17.534 -35.466  1.00 17.14           O
    ATOM   1190  CB  LEU A 311      32.663  15.347 -33.885  1.00 14.03           C
    ATOM   1191  CG  LEU A 311      33.203  13.922 -33.646  1.00 14.30           C
    ATOM   1192  CD1 LEU A 311      32.278  13.150 -32.713  1.00 12.33           C
    ATOM   1193  CD2 LEU A 311      33.384  13.135 -34.951  1.00 14.46           C
    ATOM   1194  H   LEU A 311      34.907  16.403 -33.302  1.00  0.00           H
    ATOM   1195  HA  LEU A 311      33.552  15.689 -35.839  1.00  0.00           H
    ATOM   1196  HB2 LEU A 311      31.639  15.263 -34.254  1.00  0.00           H
    ATOM   1197  HB3 LEU A 311      32.592  15.867 -32.927  1.00  0.00           H
    ATOM   1198  HG  LEU A 311      34.173  13.992 -33.152  1.00  0.00           H
    ATOM   1199 HD11 LEU A 311      32.744  12.209 -32.425  1.00  0.00           H
    ATOM   1200 HD12 LEU A 311      32.068  13.707 -31.807  1.00  0.00           H
    ATOM   1201 HD13 LEU A 311      31.324  12.929 -33.194  1.00  0.00           H
    ATOM   1202 HD21 LEU A 311      33.559  12.078 -34.769  1.00  0.00           H
    ATOM   1203 HD22 LEU A 311      32.503  13.213 -35.588  1.00  0.00           H
    ATOM   1204 HD23 LEU A 311      34.242  13.505 -35.508  1.00  0.00           H
    ATOM   1205  N   GLU A 312      33.521  18.676 -34.955  1.00 19.66           N
    ATOM   1206  CA  GLU A 312      33.081  20.046 -35.269  1.00 21.33           C
    ATOM   1207  C   GLU A 312      32.450  20.206 -36.665  1.00 20.18           C
    ATOM   1208  O   GLU A 312      31.457  20.916 -36.812  1.00 20.84           O
    ATOM   1209  CB  GLU A 312      34.240  21.038 -34.970  1.00 23.92           C
    ATOM   1210  CG  GLU A 312      34.309  22.377 -35.754  1.00 26.36           C
    ATOM   1211  CD  GLU A 312      34.904  22.299 -37.172  1.00 29.27           C
    ATOM   1212  OE1 GLU A 312      34.646  23.251 -37.940  1.00 30.18           O
    ATOM   1213  OE2 GLU A 312      35.596  21.302 -37.473  1.00 28.04           O1-
    ATOM   1214  H   GLU A 312      34.477  18.579 -34.645  1.00  0.00           H
    ATOM   1215  HA  GLU A 312      32.279  20.272 -34.563  1.00  0.00           H
    ATOM   1216  HB2 GLU A 312      34.216  21.267 -33.904  1.00  0.00           H
    ATOM   1217  HB3 GLU A 312      35.187  20.525 -35.113  1.00  0.00           H
    ATOM   1218  HG2 GLU A 312      33.324  22.845 -35.792  1.00  0.00           H
    ATOM   1219  HG3 GLU A 312      34.943  23.067 -35.198  1.00  0.00           H
    ATOM   1220  N   GLN A 313      33.025  19.512 -37.664  1.00 18.36           N
    ATOM   1221  CA  GLN A 313      32.615  19.472 -39.068  1.00 16.21           C
    ATOM   1222  C   GLN A 313      31.176  18.958 -39.271  1.00 13.20           C
    ATOM   1223  O   GLN A 313      30.538  19.314 -40.262  1.00 10.98           O
    ATOM   1224  CB  GLN A 313      33.639  18.588 -39.820  1.00 19.15           C
    ATOM   1225  CG  GLN A 313      33.385  18.412 -41.335  1.00 20.10           C
    ATOM   1226  CD  GLN A 313      34.417  17.524 -42.035  1.00 22.21           C
    ATOM   1227  OE1 GLN A 313      35.345  17.004 -41.418  1.00 23.41           O
    ATOM   1228  NE2 GLN A 313      34.246  17.334 -43.344  1.00 21.23           N
    ATOM   1229  H   GLN A 313      33.854  18.987 -37.429  1.00  0.00           H
    ATOM   1230  HA  GLN A 313      32.668  20.488 -39.461  1.00  0.00           H
    ATOM   1231  HB2 GLN A 313      34.633  19.015 -39.678  1.00  0.00           H
    ATOM   1232  HB3 GLN A 313      33.667  17.603 -39.349  1.00  0.00           H
    ATOM   1233  HG2 GLN A 313      32.412  17.953 -41.511  1.00  0.00           H
    ATOM   1234  HG3 GLN A 313      33.371  19.387 -41.822  1.00  0.00           H
    ATOM   1235 HE21 GLN A 313      34.885  16.734 -43.845  1.00  0.00           H
    ATOM   1236 HE22 GLN A 313      33.466  17.767 -43.814  1.00  0.00           H
    ATOM   1237  N   TYR A 314      30.679  18.131 -38.337  1.00 11.15           N
    ATOM   1238  CA  TYR A 314      29.381  17.471 -38.397  1.00 10.01           C
    ATOM   1239  C   TYR A 314      28.444  17.891 -37.259  1.00  8.24           C
    ATOM   1240  O   TYR A 314      27.233  17.734 -37.419  1.00  8.86           O
    ATOM   1241  CB  TYR A 314      29.606  15.949 -38.349  1.00  7.08           C
    ATOM   1242  CG  TYR A 314      30.354  15.404 -39.549  1.00  6.89           C
    ATOM   1243  CD1 TYR A 314      29.684  15.269 -40.781  1.00  7.74           C
    ATOM   1244  CD2 TYR A 314      31.720  15.066 -39.451  1.00  7.52           C
    ATOM   1245  CE1 TYR A 314      30.379  14.810 -41.915  1.00  8.70           C
    ATOM   1246  CE2 TYR A 314      32.415  14.606 -40.587  1.00  8.31           C
    ATOM   1247  CZ  TYR A 314      31.744  14.483 -41.820  1.00  9.10           C
    ATOM   1248  OH  TYR A 314      32.403  14.053 -42.933  1.00 10.42           O
    ATOM   1249  H   TYR A 314      31.267  17.914 -37.544  1.00  0.00           H
    ATOM   1250  HA  TYR A 314      28.867  17.713 -39.328  1.00  0.00           H
    ATOM   1251  HB2 TYR A 314      30.131  15.668 -37.434  1.00  0.00           H
    ATOM   1252  HB3 TYR A 314      28.642  15.444 -38.316  1.00  0.00           H
    ATOM   1253  HD1 TYR A 314      28.638  15.527 -40.862  1.00  0.00           H
    ATOM   1254  HD2 TYR A 314      32.243  15.173 -38.511  1.00  0.00           H
    ATOM   1255  HE1 TYR A 314      29.863  14.712 -42.858  1.00  0.00           H
    ATOM   1256  HE2 TYR A 314      33.464  14.360 -40.518  1.00  0.00           H
    ATOM   1257  HH  TYR A 314      33.322  13.855 -42.814  1.00  0.00           H
    ATOM   1258  N   TYR A 315      28.968  18.406 -36.128  1.00  7.90           N
    ATOM   1259  CA  TYR A 315      28.188  18.728 -34.933  1.00  7.22           C
    ATOM   1260  C   TYR A 315      27.185  19.847 -35.209  1.00  7.29           C
    ATOM   1261  O   TYR A 315      27.555  20.922 -35.682  1.00  8.25           O
    ATOM   1262  CB  TYR A 315      29.094  19.144 -33.756  1.00  7.46           C
    ATOM   1263  CG  TYR A 315      28.331  19.398 -32.455  1.00  8.68           C
    ATOM   1264  CD1 TYR A 315      27.452  18.417 -31.951  1.00 11.43           C
    ATOM   1265  CD2 TYR A 315      28.460  20.622 -31.763  1.00  9.14           C
    ATOM   1266  CE1 TYR A 315      26.719  18.643 -30.772  1.00 12.74           C
    ATOM   1267  CE2 TYR A 315      27.729  20.849 -30.579  1.00 10.71           C
    ATOM   1268  CZ  TYR A 315      26.859  19.858 -30.081  1.00 13.42           C
    ATOM   1269  OH  TYR A 315      26.157  20.074 -28.931  1.00 17.74           O
    ATOM   1270  H   TYR A 315      29.972  18.511 -36.066  1.00  0.00           H
    ATOM   1271  HA  TYR A 315      27.650  17.821 -34.662  1.00  0.00           H
    ATOM   1272  HB2 TYR A 315      29.825  18.359 -33.579  1.00  0.00           H
    ATOM   1273  HB3 TYR A 315      29.672  20.031 -34.020  1.00  0.00           H
    ATOM   1274  HD1 TYR A 315      27.336  17.477 -32.470  1.00  0.00           H
    ATOM   1275  HD2 TYR A 315      29.120  21.390 -32.138  1.00  0.00           H
    ATOM   1276  HE1 TYR A 315      26.056  17.878 -30.398  1.00  0.00           H
    ATOM   1277  HE2 TYR A 315      27.838  21.786 -30.053  1.00  0.00           H
    ATOM   1278  HH  TYR A 315      25.671  19.316 -28.647  1.00  0.00           H
    ATOM   1279  N   ASP A 316      25.917  19.561 -34.898  1.00  8.78           N
    ATOM   1280  CA  ASP A 316      24.805  20.469 -35.079  1.00  9.89           C
    ATOM   1281  C   ASP A 316      23.649  20.026 -34.167  1.00 10.15           C
    ATOM   1282  O   ASP A 316      22.883  19.154 -34.578  1.00  9.61           O
    ATOM   1283  CB  ASP A 316      24.454  20.567 -36.584  1.00 10.39           C
    ATOM   1284  CG  ASP A 316      23.217  21.410 -36.886  1.00 11.43           C
    ATOM   1285  OD1 ASP A 316      22.907  22.312 -36.076  1.00  9.92           O
    ATOM   1286  OD2 ASP A 316      22.594  21.135 -37.929  1.00 14.69           O1-
    ATOM   1287  H   ASP A 316      25.706  18.648 -34.509  1.00  0.00           H
    ATOM   1288  HA  ASP A 316      25.134  21.463 -34.783  1.00  0.00           H
    ATOM   1289  HB2 ASP A 316      25.286  20.995 -37.144  1.00  0.00           H
    ATOM   1290  HB3 ASP A 316      24.295  19.566 -36.989  1.00  0.00           H
    ATOM   1291  N   PRO A 317      23.469  20.617 -32.964  1.00  9.99           N
    ATOM   1292  CA  PRO A 317      22.373  20.264 -32.051  1.00  9.51           C
    ATOM   1293  C   PRO A 317      20.974  20.761 -32.494  1.00 10.51           C
    ATOM   1294  O   PRO A 317      20.014  20.552 -31.754  1.00  7.52           O
    ATOM   1295  CB  PRO A 317      22.807  20.831 -30.692  1.00  8.83           C
    ATOM   1296  CG  PRO A 317      23.683  22.022 -31.039  1.00  9.51           C
    ATOM   1297  CD  PRO A 317      24.328  21.645 -32.373  1.00 10.35           C
    ATOM   1298  HA  PRO A 317      22.314  19.181 -31.964  1.00  0.00           H
    ATOM   1299  HB2 PRO A 317      21.980  21.098 -30.032  1.00  0.00           H
    ATOM   1300  HB3 PRO A 317      23.410  20.086 -30.170  1.00  0.00           H
    ATOM   1301  HG2 PRO A 317      23.055  22.903 -31.174  1.00  0.00           H
    ATOM   1302  HG3 PRO A 317      24.416  22.251 -30.265  1.00  0.00           H
    ATOM   1303  HD2 PRO A 317      24.410  22.527 -33.010  1.00  0.00           H
    ATOM   1304  HD3 PRO A 317      25.327  21.236 -32.220  1.00  0.00           H
    ATOM   1305  N   SER A 318      20.838  21.353 -33.700  1.00 10.31           N
    ATOM   1306  CA  SER A 318      19.559  21.643 -34.359  1.00 10.77           C
    ATOM   1307  C   SER A 318      19.131  20.515 -35.331  1.00  9.84           C
    ATOM   1308  O   SER A 318      17.984  20.517 -35.775  1.00  9.15           O
    ATOM   1309  CB  SER A 318      19.667  22.998 -35.106  1.00 12.58           C
    ATOM   1310  OG  SER A 318      20.242  22.933 -36.398  1.00 13.89           O
    ATOM   1311  H   SER A 318      21.672  21.518 -34.248  1.00  0.00           H
    ATOM   1312  HA  SER A 318      18.780  21.745 -33.602  1.00  0.00           H
    ATOM   1313  HB2 SER A 318      20.216  23.727 -34.508  1.00  0.00           H
    ATOM   1314  HB3 SER A 318      18.665  23.408 -35.233  1.00  0.00           H
    ATOM   1315  HG  SER A 318      21.159  22.689 -36.319  1.00  0.00           H
    ATOM   1316  N   ASP A 319      20.020  19.545 -35.623  1.00  8.77           N
    ATOM   1317  CA  ASP A 319      19.809  18.386 -36.503  1.00  8.95           C
    ATOM   1318  C   ASP A 319      20.177  17.066 -35.789  1.00  9.15           C
    ATOM   1319  O   ASP A 319      20.514  16.059 -36.413  1.00  8.57           O
    ATOM   1320  CB  ASP A 319      20.472  18.639 -37.880  1.00  9.03           C
    ATOM   1321  CG  ASP A 319      20.258  17.551 -38.942  1.00 11.88           C
    ATOM   1322  OD1 ASP A 319      19.089  17.156 -39.150  1.00 14.46           O
    ATOM   1323  OD2 ASP A 319      21.272  17.111 -39.527  1.00 16.18           O1-
    ATOM   1324  H   ASP A 319      20.941  19.621 -35.211  1.00  0.00           H
    ATOM   1325  HA  ASP A 319      18.737  18.292 -36.686  1.00  0.00           H
    ATOM   1326  HB2 ASP A 319      20.107  19.581 -38.293  1.00  0.00           H
    ATOM   1327  HB3 ASP A 319      21.546  18.762 -37.733  1.00  0.00           H
    ATOM   1328  N   GLU A 320      20.095  17.069 -34.452  1.00  9.87           N
    ATOM   1329  CA  GLU A 320      20.334  15.925 -33.582  1.00  9.67           C
    ATOM   1330  C   GLU A 320      19.112  15.804 -32.665  1.00 10.30           C
    ATOM   1331  O   GLU A 320      19.142  16.346 -31.558  1.00 11.63           O
    ATOM   1332  CB  GLU A 320      21.641  16.139 -32.802  1.00  6.46           C
    ATOM   1333  CG  GLU A 320      22.910  16.067 -33.674  1.00  5.15           C
    ATOM   1334  CD  GLU A 320      24.151  16.629 -32.987  1.00  6.11           C
    ATOM   1335  OE1 GLU A 320      24.109  16.814 -31.750  1.00  7.21           O
    ATOM   1336  OE2 GLU A 320      25.126  16.877 -33.727  1.00  5.63           O1-
    ATOM   1337  H   GLU A 320      19.819  17.931 -34.005  1.00  0.00           H
    ATOM   1338  HA  GLU A 320      20.450  15.000 -34.147  1.00  0.00           H
    ATOM   1339  HB2 GLU A 320      21.723  15.384 -32.018  1.00  0.00           H
    ATOM   1340  HB3 GLU A 320      21.589  17.101 -32.293  1.00  0.00           H
    ATOM   1341  HG2 GLU A 320      23.117  15.043 -33.969  1.00  0.00           H
    ATOM   1342  HG3 GLU A 320      22.770  16.615 -34.604  1.00  0.00           H
    ATOM   1343  N   PRO A 321      18.030  15.119 -33.102  1.00 11.58           N
    ATOM   1344  CA  PRO A 321      16.817  14.965 -32.298  1.00 12.00           C
    ATOM   1345  C   PRO A 321      17.045  14.012 -31.117  1.00 12.76           C
    ATOM   1346  O   PRO A 321      17.958  13.183 -31.125  1.00 14.81           O
    ATOM   1347  CB  PRO A 321      15.749  14.448 -33.272  1.00 11.77           C
    ATOM   1348  CG  PRO A 321      16.531  13.743 -34.369  1.00 11.44           C
    ATOM   1349  CD  PRO A 321      17.888  14.450 -34.399  1.00 11.55           C
    ATOM   1350  HA  PRO A 321      16.507  15.938 -31.909  1.00  0.00           H
    ATOM   1351  HB2 PRO A 321      15.008  13.796 -32.807  1.00  0.00           H
    ATOM   1352  HB3 PRO A 321      15.212  15.295 -33.700  1.00  0.00           H
    ATOM   1353  HG2 PRO A 321      16.668  12.699 -34.096  1.00  0.00           H
    ATOM   1354  HG3 PRO A 321      16.023  13.767 -35.333  1.00  0.00           H
    ATOM   1355  HD2 PRO A 321      18.686  13.729 -34.581  1.00  0.00           H
    ATOM   1356  HD3 PRO A 321      17.906  15.196 -35.193  1.00  0.00           H
    ATOM   1357  N   ILE A 322      16.202  14.180 -30.095  1.00 13.17           N
    ATOM   1358  CA  ILE A 322      16.216  13.461 -28.829  1.00 14.51           C
    ATOM   1359  C   ILE A 322      14.837  12.840 -28.584  1.00 15.59           C
    ATOM   1360  O   ILE A 322      13.866  13.152 -29.277  1.00 16.73           O
    ATOM   1361  CB  ILE A 322      16.685  14.406 -27.688  1.00 12.38           C
    ATOM   1362  CG1 ILE A 322      15.840  15.693 -27.547  1.00 12.88           C
    ATOM   1363  CG2 ILE A 322      18.193  14.708 -27.803  1.00 12.33           C
    ATOM   1364  CD1 ILE A 322      16.013  16.387 -26.188  1.00 14.54           C
    ATOM   1365  H   ILE A 322      15.472  14.870 -30.200  1.00  0.00           H
    ATOM   1366  HA  ILE A 322      16.912  12.622 -28.887  1.00  0.00           H
    ATOM   1367  HB  ILE A 322      16.546  13.868 -26.755  1.00  0.00           H
    ATOM   1368 HG12 ILE A 322      16.081  16.390 -28.350  1.00  0.00           H
    ATOM   1369 HG13 ILE A 322      14.782  15.457 -27.660  1.00  0.00           H
    ATOM   1370 HG21 ILE A 322      18.567  15.238 -26.928  1.00  0.00           H
    ATOM   1371 HG22 ILE A 322      18.770  13.786 -27.888  1.00  0.00           H
    ATOM   1372 HG23 ILE A 322      18.413  15.318 -28.680  1.00  0.00           H
    ATOM   1373 HD11 ILE A 322      15.356  17.254 -26.114  1.00  0.00           H
    ATOM   1374 HD12 ILE A 322      15.763  15.714 -25.368  1.00  0.00           H
    ATOM   1375 HD13 ILE A 322      17.034  16.737 -26.040  1.00  0.00           H
  extra_rotamers: ""
ligand:
  residue_configurations: []
  extra_templates: ""
  extra_templates_coordinates: ""
  coordinates: |
    ATOM   1377  N   ARG B 564      22.551  19.212 -46.615  1.00 72.78           N1+
    ATOM   1378  CA  ARG B 564      21.736  18.646 -45.546  1.00 72.80           C
    ATOM   1379  C   ARG B 564      21.272  17.238 -45.913  1.00 72.62           C
    ATOM   1380  O   ARG B 564      20.497  17.072 -46.855  1.00 72.72           O
    ATOM   1381  CB  ARG B 564      20.529  19.561 -45.249  1.00 72.83           C
    ATOM   1382  CG  ARG B 564      20.900  20.906 -44.596  1.00 72.42           C
    ATOM   1383  CD  ARG B 564      21.125  20.823 -43.080  1.00 73.55           C
    ATOM   1384  NE  ARG B 564      22.395  20.167 -42.737  1.00 73.54           N
    ATOM   1385  CZ  ARG B 564      22.780  19.854 -41.491  1.00 73.18           C
    ATOM   1386  NH1 ARG B 564      22.009  20.184 -40.451  1.00 72.49           N
    ATOM   1387  NH2 ARG B 564      23.942  19.222 -41.282  1.00 73.25           N1+
    ATOM   1388  H   ARG B 564      22.014  19.331 -47.462  1.00  0.00           H
    ATOM   1389  HA  ARG B 564      22.367  18.554 -44.665  1.00  0.00           H
    ATOM   1390  HB2 ARG B 564      20.002  19.764 -46.183  1.00  0.00           H
    ATOM   1391  HB3 ARG B 564      19.806  19.044 -44.615  1.00  0.00           H
    ATOM   1392  HG2 ARG B 564      21.767  21.349 -45.088  1.00  0.00           H
    ATOM   1393  HG3 ARG B 564      20.082  21.605 -44.770  1.00  0.00           H
    ATOM   1394  HD2 ARG B 564      20.296  20.298 -42.604  1.00  0.00           H
    ATOM   1395  HD3 ARG B 564      21.145  21.830 -42.661  1.00  0.00           H
    ATOM   1396  HE  ARG B 564      23.012  19.958 -43.509  1.00  0.00           H
    ATOM   1397 HH11 ARG B 564      21.124  20.641 -40.604  1.00  0.00           H
    ATOM   1398 HH12 ARG B 564      22.335  20.087 -39.493  1.00  0.00           H
    ATOM   1399 HH21 ARG B 564      24.544  18.977 -42.053  1.00  0.00           H
    ATOM   1400 HH22 ARG B 564      24.218  18.982 -40.340  1.00  0.00           H
    ATOM   1401  N   ARG B 565      21.774  16.240 -45.165  1.00 71.88           N
    ATOM   1402  CA  ARG B 565      21.496  14.805 -45.259  1.00 70.59           C
    ATOM   1403  C   ARG B 565      22.347  14.167 -46.365  1.00 69.86           C
    ATOM   1404  O   ARG B 565      22.257  14.590 -47.517  1.00 69.50           O
    ATOM   1405  CB  ARG B 565      19.978  14.507 -45.379  1.00 70.13           C
    ATOM   1406  CG  ARG B 565      19.560  13.109 -44.887  1.00 69.03           C
    ATOM   1407  CD  ARG B 565      19.103  13.110 -43.418  1.00 68.13           C
    ATOM   1408  NE  ARG B 565      20.213  13.435 -42.509  1.00 67.46           N
    ATOM   1409  CZ  ARG B 565      20.214  14.340 -41.513  1.00 66.75           C
    ATOM   1410  NH1 ARG B 565      21.341  14.551 -40.825  1.00 66.58           N
    ATOM   1411  NH2 ARG B 565      19.122  15.049 -41.193  1.00 66.08           N1+
    ATOM   1412  H   ARG B 565      22.414  16.508 -44.432  1.00  0.00           H
    ATOM   1413  HA  ARG B 565      21.849  14.407 -44.309  1.00  0.00           H
    ATOM   1414  HB2 ARG B 565      19.398  15.259 -44.841  1.00  0.00           H
    ATOM   1415  HB3 ARG B 565      19.676  14.607 -46.423  1.00  0.00           H
    ATOM   1416  HG2 ARG B 565      20.359  12.382 -45.044  1.00  0.00           H
    ATOM   1417  HG3 ARG B 565      18.724  12.760 -45.495  1.00  0.00           H
    ATOM   1418  HD2 ARG B 565      18.726  12.122 -43.153  1.00  0.00           H
    ATOM   1419  HD3 ARG B 565      18.271  13.801 -43.289  1.00  0.00           H
    ATOM   1420  HE  ARG B 565      21.022  12.835 -42.598  1.00  0.00           H
    ATOM   1421 HH11 ARG B 565      21.374  15.307 -40.144  1.00  0.00           H
    ATOM   1422 HH12 ARG B 565      22.174  14.014 -41.012  1.00  0.00           H
    ATOM   1423 HH21 ARG B 565      19.161  15.753 -40.459  1.00  0.00           H
    ATOM   1424 HH22 ARG B 565      18.258  14.902 -41.691  1.00  0.00           H
    ATOM   1425  N   LEU B 566      23.168  13.156 -46.019  1.00 68.73           N
    ATOM   1426  CA  LEU B 566      24.015  12.379 -46.935  1.00 67.51           C
    ATOM   1427  C   LEU B 566      25.059  13.271 -47.637  1.00 66.78           C
    ATOM   1428  O   LEU B 566      24.960  13.537 -48.835  1.00 66.05           O
    ATOM   1429  CB  LEU B 566      23.150  11.538 -47.922  1.00 67.71           C
    ATOM   1430  CG  LEU B 566      22.960  10.076 -47.487  1.00 67.55           C
    ATOM   1431  CD1 LEU B 566      21.812   9.417 -48.265  1.00 67.96           C
    ATOM   1432  CD2 LEU B 566      24.258   9.280 -47.684  1.00 67.33           C
    ATOM   1433  H   LEU B 566      23.184  12.873 -45.050  1.00  0.00           H
    ATOM   1434  HA  LEU B 566      24.589  11.702 -46.303  1.00  0.00           H
    ATOM   1435  HB2 LEU B 566      22.172  11.999 -48.058  1.00  0.00           H
    ATOM   1436  HB3 LEU B 566      23.580  11.519 -48.923  1.00  0.00           H
    ATOM   1437  HG  LEU B 566      22.692  10.052 -46.429  1.00  0.00           H
    ATOM   1438 HD11 LEU B 566      21.664   8.385 -47.947  1.00  0.00           H
    ATOM   1439 HD12 LEU B 566      20.872   9.946 -48.105  1.00  0.00           H
    ATOM   1440 HD13 LEU B 566      22.013   9.409 -49.337  1.00  0.00           H
    ATOM   1441 HD21 LEU B 566      24.087   8.224 -47.496  1.00  0.00           H
    ATOM   1442 HD22 LEU B 566      24.631   9.367 -48.704  1.00  0.00           H
    ATOM   1443 HD23 LEU B 566      25.049   9.604 -47.009  1.00  0.00           H
    ATOM   1444  N   GLN B 567      26.037  13.772 -46.867  1.00 65.77           N
    ATOM   1445  CA  GLN B 567      27.055  14.709 -47.337  1.00 64.99           C
    ATOM   1446  C   GLN B 567      28.366  13.960 -47.583  1.00 64.11           C
    ATOM   1447  O   GLN B 567      29.248  13.931 -46.725  1.00 64.47           O
    ATOM   1448  CB  GLN B 567      27.226  15.824 -46.277  1.00 64.84           C
    ATOM   1449  CG  GLN B 567      26.056  16.832 -46.307  1.00 64.73           C
    ATOM   1450  CD  GLN B 567      25.988  17.694 -47.576  1.00 63.81           C
    ATOM   1451  OE1 GLN B 567      24.899  18.038 -48.023  1.00 63.54           O
    ATOM   1452  NE2 GLN B 567      27.128  18.019 -48.193  1.00 63.83           N
    ATOM   1453  H   GLN B 567      26.064  13.508 -45.894  1.00  0.00           H
    ATOM   1454  HA  GLN B 567      26.767  15.161 -48.287  1.00  0.00           H
    ATOM   1455  HB2 GLN B 567      27.295  15.382 -45.282  1.00  0.00           H
    ATOM   1456  HB3 GLN B 567      28.169  16.355 -46.420  1.00  0.00           H
    ATOM   1457  HG2 GLN B 567      25.106  16.312 -46.179  1.00  0.00           H
    ATOM   1458  HG3 GLN B 567      26.146  17.510 -45.457  1.00  0.00           H
    ATOM   1459 HE21 GLN B 567      27.093  18.572 -49.038  1.00  0.00           H
    ATOM   1460 HE22 GLN B 567      28.013  17.715 -47.817  1.00  0.00           H
    ATOM   1461  N   LYS B 568      28.472  13.375 -48.790  1.00 62.30           N
    ATOM   1462  CA  LYS B 568      29.649  12.719 -49.371  1.00 60.06           C
    ATOM   1463  C   LYS B 568      29.857  11.298 -48.824  1.00 59.70           C
    ATOM   1464  O   LYS B 568      29.254  10.902 -47.825  1.00 59.58           O
    ATOM   1465  CB  LYS B 568      30.929  13.586 -49.276  1.00 59.44           C
    ATOM   1466  CG  LYS B 568      30.792  15.020 -49.829  1.00 57.57           C
    ATOM   1467  CD  LYS B 568      31.840  15.980 -49.246  1.00 56.33           C
    ATOM   1468  CE  LYS B 568      31.561  16.353 -47.778  1.00 54.83           C
    ATOM   1469  NZ  LYS B 568      32.591  17.261 -47.251  1.00 52.73           N1+
    ATOM   1470  H   LYS B 568      27.659  13.428 -49.388  1.00  0.00           H
    ATOM   1471  HA  LYS B 568      29.413  12.604 -50.430  1.00  0.00           H
    ATOM   1472  HB2 LYS B 568      31.261  13.603 -48.240  1.00  0.00           H
    ATOM   1473  HB3 LYS B 568      31.745  13.097 -49.810  1.00  0.00           H
    ATOM   1474  HG2 LYS B 568      30.887  14.987 -50.915  1.00  0.00           H
    ATOM   1475  HG3 LYS B 568      29.804  15.437 -49.636  1.00  0.00           H
    ATOM   1476  HD2 LYS B 568      32.833  15.540 -49.343  1.00  0.00           H
    ATOM   1477  HD3 LYS B 568      31.851  16.891 -49.846  1.00  0.00           H
    ATOM   1478  HE2 LYS B 568      30.586  16.834 -47.687  1.00  0.00           H
    ATOM   1479  HE3 LYS B 568      31.542  15.472 -47.138  1.00  0.00           H
    ATOM   1480  HZ1 LYS B 568      32.621  18.103 -47.807  1.00  0.00           H
    ATOM   1481  HZ2 LYS B 568      32.372  17.495 -46.293  1.00  0.00           H
    ATOM   1482  HZ3 LYS B 568      33.490  16.801 -47.286  1.00  0.00           H
    ATOM   1483  N   GLY B 569      30.746  10.544 -49.485  1.00 58.90           N
    ATOM   1484  CA  GLY B 569      31.161   9.200 -49.107  1.00 57.56           C
    ATOM   1485  C   GLY B 569      30.152   8.141 -49.548  1.00 56.73           C
    ATOM   1486  O   GLY B 569      29.038   8.077 -49.029  1.00 55.32           O
    ATOM   1487  H   GLY B 569      31.192  10.945 -50.297  1.00  0.00           H
    ATOM   1488  HA2 GLY B 569      31.297   9.135 -48.026  1.00  0.00           H
    ATOM   1489  HA3 GLY B 569      32.135   9.005 -49.556  1.00  0.00           H
    ATOM   1490  N   ASN B 570      30.576   7.287 -50.488  1.00 55.98           N
    ATOM   1491  CA  ASN B 570      29.872   6.082 -50.913  1.00 56.14           C
    ATOM   1492  C   ASN B 570      30.620   4.905 -50.272  1.00 57.34           C
    ATOM   1493  O   ASN B 570      31.853   4.882 -50.299  1.00 57.18           O
    ATOM   1494  CB  ASN B 570      29.986   5.979 -52.453  1.00 54.61           C
    ATOM   1495  CG  ASN B 570      28.866   5.140 -53.071  1.00 52.39           C
    ATOM   1496  OD1 ASN B 570      27.915   5.687 -53.624  1.00 51.75           O
    ATOM   1497  ND2 ASN B 570      28.980   3.812 -53.005  1.00 50.51           N
    ATOM   1498  H   ASN B 570      31.511   7.410 -50.848  1.00  0.00           H
    ATOM   1499  HA  ASN B 570      28.827   6.104 -50.598  1.00  0.00           H
    ATOM   1500  HB2 ASN B 570      29.908   6.976 -52.891  1.00  0.00           H
    ATOM   1501  HB3 ASN B 570      30.957   5.595 -52.773  1.00  0.00           H
    ATOM   1502 HD21 ASN B 570      28.250   3.232 -53.393  1.00  0.00           H
    ATOM   1503 HD22 ASN B 570      29.773   3.390 -52.545  1.00  0.00           H
    ATOM   1504  N   LEU B 571      29.888   3.926 -49.716  1.00 58.85           N
    ATOM   1505  CA  LEU B 571      30.484   2.730 -49.128  1.00 60.46           C
    ATOM   1506  C   LEU B 571      30.476   1.604 -50.181  1.00 61.68           C
    ATOM   1507  O   LEU B 571      29.391   1.183 -50.589  1.00 62.45           O
    ATOM   1508  CB  LEU B 571      29.648   2.279 -47.903  1.00 60.16           C
    ATOM   1509  CG  LEU B 571      30.302   1.155 -47.060  1.00 60.08           C
    ATOM   1510  CD1 LEU B 571      31.700   1.538 -46.560  1.00 60.45           C
    ATOM   1511  CD2 LEU B 571      29.440   0.772 -45.853  1.00 61.61           C
    ATOM   1512  H   LEU B 571      28.879   3.977 -49.757  1.00  0.00           H
    ATOM   1513  HA  LEU B 571      31.484   2.975 -48.782  1.00  0.00           H
    ATOM   1514  HB2 LEU B 571      29.480   3.136 -47.254  1.00  0.00           H
    ATOM   1515  HB3 LEU B 571      28.654   1.964 -48.219  1.00  0.00           H
    ATOM   1516  HG  LEU B 571      30.407   0.263 -47.677  1.00  0.00           H
    ATOM   1517 HD11 LEU B 571      32.071   0.795 -45.864  1.00  0.00           H
    ATOM   1518 HD12 LEU B 571      32.432   1.587 -47.360  1.00  0.00           H
    ATOM   1519 HD13 LEU B 571      31.694   2.500 -46.047  1.00  0.00           H
    ATOM   1520 HD21 LEU B 571      29.952   0.053 -45.213  1.00  0.00           H
    ATOM   1521 HD22 LEU B 571      29.194   1.645 -45.248  1.00  0.00           H
    ATOM   1522 HD23 LEU B 571      28.517   0.292 -46.159  1.00  0.00           H
    ATOM   1523  N   PRO B 572      31.647   1.065 -50.596  1.00 62.14           N
    ATOM   1524  CA  PRO B 572      31.732  -0.172 -51.385  1.00 62.41           C
    ATOM   1525  C   PRO B 572      31.547  -1.383 -50.446  1.00 62.96           C
    ATOM   1526  O   PRO B 572      32.514  -2.038 -50.056  1.00 63.39           O
    ATOM   1527  CB  PRO B 572      33.131  -0.116 -52.022  1.00 62.36           C
    ATOM   1528  CG  PRO B 572      33.972   0.652 -51.015  1.00 63.06           C
    ATOM   1529  CD  PRO B 572      32.984   1.635 -50.388  1.00 62.73           C
    ATOM   1530  HA  PRO B 572      30.973  -0.205 -52.169  1.00  0.00           H
    ATOM   1531  HB2 PRO B 572      33.550  -1.092 -52.267  1.00  0.00           H
    ATOM   1532  HB3 PRO B 572      33.078   0.451 -52.952  1.00  0.00           H
    ATOM   1533  HG2 PRO B 572      34.350  -0.029 -50.251  1.00  0.00           H
    ATOM   1534  HG3 PRO B 572      34.830   1.148 -51.468  1.00  0.00           H
    ATOM   1535  HD2 PRO B 572      33.237   1.790 -49.343  1.00  0.00           H
    ATOM   1536  HD3 PRO B 572      33.042   2.596 -50.898  1.00  0.00           H
    ATOM   1537  N   VAL B 573      30.283  -1.617 -50.055  1.00 63.32           N
    ATOM   1538  CA  VAL B 573      29.784  -2.569 -49.062  1.00 63.65           C
    ATOM   1539  C   VAL B 573      30.409  -3.966 -49.106  1.00 63.63           C
    ATOM   1540  O   VAL B 573      30.604  -4.542 -50.175  1.00 63.92           O
    ATOM   1541  CB  VAL B 573      28.243  -2.641 -49.112  1.00 63.26           C
    ATOM   1542  CG1 VAL B 573      27.612  -1.400 -48.480  1.00 62.70           C
    ATOM   1543  CG2 VAL B 573      27.686  -2.897 -50.527  1.00 63.74           C
    ATOM   1544  H   VAL B 573      29.583  -1.000 -50.443  1.00  0.00           H
    ATOM   1545  HA  VAL B 573      30.061  -2.143 -48.096  1.00  0.00           H
    ATOM   1546  HB  VAL B 573      27.933  -3.484 -48.493  1.00  0.00           H
    ATOM   1547 HG11 VAL B 573      26.550  -1.348 -48.708  1.00  0.00           H
    ATOM   1548 HG12 VAL B 573      27.714  -1.425 -47.397  1.00  0.00           H
    ATOM   1549 HG13 VAL B 573      28.081  -0.484 -48.835  1.00  0.00           H
    ATOM   1550 HG21 VAL B 573      26.608  -3.043 -50.500  1.00  0.00           H
    ATOM   1551 HG22 VAL B 573      27.882  -2.059 -51.196  1.00  0.00           H
    ATOM   1552 HG23 VAL B 573      28.118  -3.791 -50.974  1.00  0.00           H
    ATOM   1553  N   ARG B 574      30.681  -4.504 -47.914  1.00 63.16           N
    ATOM   1554  CA  ARG B 574      31.195  -5.845 -47.705  1.00 62.89           C
    ATOM   1555  C   ARG B 574      30.037  -6.857 -47.580  1.00 63.21           C
    ATOM   1556  O   ARG B 574      28.849  -6.566 -47.456  1.00 63.58           O
    ATOM   1557  CB  ARG B 574      32.066  -5.793 -46.432  1.00 61.77           C
    ATOM   1558  CG  ARG B 574      32.439  -7.145 -45.815  1.00 60.98           C
    ATOM   1559  CD  ARG B 574      33.290  -8.042 -46.717  1.00 60.50           C
    ATOM   1560  NE  ARG B 574      33.283  -9.435 -46.252  1.00 60.42           N
    ATOM   1561  CZ  ARG B 574      33.808  -9.912 -45.109  1.00 59.54           C
    ATOM   1562  NH1 ARG B 574      34.398  -9.119 -44.206  1.00 58.75           N
    ATOM   1563  NH2 ARG B 574      33.736 -11.224 -44.859  1.00 60.44           N1+
    ATOM   1564  H   ARG B 574      30.475  -3.961 -47.088  1.00  0.00           H
    ATOM   1565  HA  ARG B 574      31.823  -6.141 -48.547  1.00  0.00           H
    ATOM   1567  HB2 ARG B 574      32.986  -5.252 -46.664  1.00  0.00           H
    ATOM   1568  HB3 ARG B 574      31.580  -5.191 -45.667  1.00  0.00           H
    ATOM   1569  HG2 ARG B 574      31.550  -7.677 -45.479  1.00  0.00           H
    ATOM   1570  HG3 ARG B 574      33.015  -6.928 -44.921  1.00  0.00           H
    ATOM   1571  HD2 ARG B 574      32.893  -8.055 -47.732  1.00  0.00           H
    ATOM   1572  HD3 ARG B 574      34.306  -7.666 -46.793  1.00  0.00           H
    ATOM   1573  HE  ARG B 574      32.838 -10.092 -46.877  1.00  0.00           H
    ATOM   1574 HH11 ARG B 574      34.459  -8.122 -44.361  1.00  0.00           H
    ATOM   1575 HH12 ARG B 574      34.793  -9.507 -43.362  1.00  0.00           H
    ATOM   1576 HH21 ARG B 574      33.304 -11.849 -45.524  1.00  0.00           H
    ATOM   1577 HH22 ARG B 574      34.128 -11.605 -44.010  1.00  0.00           H
  extra_rotamers: ""
```
