## Supplementary material for "DISRUPTOR: Computational identification of oncogenic mutants disrupting protein interactions": exemplary input files: p53_ASPP2_1ycs.137-165.yaml

```
---
osprey_version: 3.1
design_name: p53 and aspp2
protein:
  residue_configurations:
  - mutability:
    - ALA
    - ARG
    - ASN
    - ASP
    - CYS
    - GLU
    - GLN
    - GLY
    - HIE
    - HID
    - HIP
    - ILE
    - LEU
    - LYS
    - MET
    - PHE
    - PRO
    - SER
    - THR
    - TRP
    - TYR
    - VAL
    flexibility:
      is_flexible: true
      include_structure_rotamer: true
      use_continuous: true
    identity:
      chain: A
      res_num: 137
      aa_type: LEU
  - mutability:
    - ALA
    - ARG
    - ASN
    - ASP
    - CYS
    - GLU
    - GLN
    - GLY
    - HIE
    - HID
    - HIP
    - ILE
    - LEU
    - LYS
    - MET
    - PHE
    - PRO
    - SER
    - THR
    - TRP
    - TYR
    - VAL
    flexibility:
      is_flexible: true
      include_structure_rotamer: true
      use_continuous: true
    identity:
      chain: A
      res_num: 165
      aa_type: GLN
  extra_templates: |2
        0    0    2
    This is a remark line
    molecule.res

     CY1 INT 1
     CORR OMIT DU   BEG 
       0.00000
        1  DUMM  DU    M    0  -1  -2     0.000      .0        .0      .00000
        2  DUMM  DU    M    1   0  -1     1.449      .0        .0      .00000
        3  DUMM  DU    M    2   1   0     1.522   111.1        .0      .00000
        4  N     N     M    3   2   1     1.335   116.600   180.000  -0.46300
        5  HN    H     E    4   3   2     1.010   119.800     0.000   0.25200
        6  CA    CT    M    4   3   2     1.449   121.900   180.000   0.03500
        7  HA    H1    E    6   4   3     1.090   109.500   300.000   0.04800
        8  CB    CT    3    6   4   3     1.525   111.100    60.000  -0.54300
        9  HB3   H1    E    8   6   4     1.090   109.500    60.000   0.18377
       10  HB2   H1    E    8   6   4     1.090   109.500   300.000   0.18377
       11  SG    S1    E    8   6   4     1.810   116.000   180.000  -0.43963
       12  C     C     M    6   4   3     1.522   111.100   180.000   0.61600
       13  O     O     E   12   6   4     1.229   120.500     0.000  -0.50400
     
    IMPROPER
     -M   CA   N    HN 
     CA   +M   C    O  
     
    DONE
    Zinc(II) Ion
    
     ZN1 INT 1
     CORR OMIT DU   BEG 
       0.00000
        1  DUMM  DU    M    0  -1  -2     0.000      .0        .0      .00000
        2  DUMM  DU    M    1   0  -1     1.449      .0        .0      .00000
        3  DUMM  DU    M    2   1   0     1.522   111.1        .0      .00000
        4 ZN     ZN    M    3   2   1     1.000    90.000   180.000   0.52437
      
    DONE
    CYSTEINE with negative charge
    
     CY2 INT 1
     CORR OMIT DU   BEG 
       0.00000
        1  DUMM  DU    M    0  -1  -2     0.000      .0        .0      .00000
        2  DUMM  DU    M    1   0  -1     1.449      .0        .0      .00000
        3  DUMM  DU    M    2   1   0     1.522   111.1        .0      .00000
        4  N     N     M    3   2   1     1.335   116.600   180.000  -0.46300
        5  H     H     E    4   3   2     1.010   119.800     0.000   0.25200
        6  CA    CT    M    4   3   2     1.449   121.900   180.000   0.03500
        7  HA    H1    E    6   4   3     1.090   109.500   300.000   0.04800
        8  CB    CT    3    6   4   3     1.525   111.100    60.000  -0.42561
        9  HB3   H1    E    8   6   4     1.090   109.500    60.000   0.17337
       10  HB2   H1    E    8   6   4     1.090   109.500   300.000   0.17337
       11  SG    S2    E    8   6   4     1.810   116.000   180.000  -0.44155
       12  C     C     M    6   4   3     1.522   111.100   180.000   0.61600
       13  O     O     E   12   6   4     1.229   120.500     0.000  -0.50400
     
    IMPROPER
     -M   CA   N    HN 
     CA   +M   C    O  
     
    DONE
    HISTIDINE EPSILONH
    
     HE1 INT 1
     CORR OMIT DU   BEG 
       0.00000
        1  DUMM  DU    M    0  -1  -2     0.000      .0        .0      .00000
        2  DUMM  DU    M    1   0  -1     1.449      .0        .0      .00000
        3  DUMM  DU    M    2   1   0     1.522   111.1        .0      .00000
        4  N     N     M    3   2   1     1.335   116.600   180.000  -0.41570
        5  H     H     E    4   3   2     1.010   119.800     0.000   0.27190
        6  CA    CT    M    4   3   2     1.449   121.900   180.000  -0.05810
        7  HA    H1    E    6   4   3     1.090   109.500   300.000   0.13600
        8  CB    CT    3    6   4   3     1.525   111.100    60.000   0.04092
        9  HB2   HC    E    8   6   4     1.090   109.500   300.000   0.06506
       10  HB3   HC    E    8   6   4     1.090   109.500    60.000   0.06506
       11  CG    CC    S    8   6   4     1.510   115.000   180.000  -0.11300
       12  ND1   N1    S   11   8   6     1.390   122.000   180.000  -0.08206
       13  CE1   CR    B   12  11   8     1.320   108.000   180.000  -0.14656
       14  HE1   H5    E   13  12  11     1.090   120.000   180.000   0.19871
       15  NE2   NA    B   13  12  11     1.310   109.000     0.000  -0.01782
       16  HE2   H     E   15  13  12     1.010   125.000   180.000   0.27163
       17  CD2   CW    S   15  13  12     1.360   110.000     0.000  -0.19685
       18  HD2   H4    E   17  15  13     1.090   120.000   180.000   0.12407
       19  C     C     M    6   4   3     1.522   111.100   180.000   0.59730
       20  O     O     E   19   6   4     1.229   120.500     0.000  -0.56790
      
    LOOP
     CG   CD2
     
    IMPROPER
     -M   CA   N    H  
     CA   +M   C    O  
     CE1  CD2  NE2  HE2
     CG   NE2  CD2  HD2
     ND1  NE2  CE1  HE1
     ND1  CD2  CG   CB 
     
    DONE
    Zinc(II) Ion
    
     ZN2 INT 1
     CORR OMIT DU   BEG 
       0.00000
        1  DUMM  DU    M    0  -1  -2     0.000      .0        .0      .00000
        2  DUMM  DU    M    1   0  -1     1.449      .0        .0      .00000
        3  DUMM  DU    M    2   1   0     1.522   111.1        .0      .00000
        4  ZN    ZN    M    3   2   1     1.000    90.000   180.000   0.43655
      
    DONE
  extra_templates_coordinates: ""
  coordinates: |
    ATOM      1  N   LYS A 120      46.262  12.515  73.159  1.00 45.79           N
    ATOM      2  CA  LYS A 120      45.072  11.898  73.753  1.00 47.25           C
    ATOM      3  C   LYS A 120      43.960  12.910  74.113  1.00 48.76           C
    ATOM      4  O   LYS A 120      42.798  12.516  74.208  1.00 47.57           O
    ATOM      5  CB  LYS A 120      45.523  11.034  74.952  1.00 52.24           C
    ATOM      6  CG  LYS A 120      44.470  10.093  75.579  1.00 63.48           C
    ATOM      7  CD  LYS A 120      44.309   8.725  74.885  1.00 71.27           C
    ATOM      8  CE  LYS A 120      43.783   8.726  73.445  1.00 72.72           C
    ATOM      9  NZ  LYS A 120      42.509   9.457  73.330  1.00 71.11           N1+
    ATOM     10  H   LYS A 120      47.127  12.424  73.677  1.00  0.00           H
    ATOM     11  HA  LYS A 120      44.660  11.255  72.982  1.00  0.00           H
    ATOM     12  HB2 LYS A 120      46.381  10.427  74.662  1.00  0.00           H
    ATOM     13  HB3 LYS A 120      45.895  11.703  75.730  1.00  0.00           H
    ATOM     14  HG2 LYS A 120      44.782   9.897  76.605  1.00  0.00           H
    ATOM     15  HG3 LYS A 120      43.503  10.581  75.680  1.00  0.00           H
    ATOM     16  HD2 LYS A 120      45.275   8.225  74.900  1.00  0.00           H
    ATOM     17  HD3 LYS A 120      43.654   8.102  75.496  1.00  0.00           H
    ATOM     18  HE2 LYS A 120      44.517   9.140  72.756  1.00  0.00           H
    ATOM     19  HE3 LYS A 120      43.631   7.696  73.133  1.00  0.00           H
    ATOM     20  HZ1 LYS A 120      41.806   9.010  73.903  1.00  0.00           H
    ATOM     21  HZ2 LYS A 120      42.633  10.411  73.641  1.00  0.00           H
    ATOM     22  HZ3 LYS A 120      42.206   9.472  72.367  1.00  0.00           H
    ATOM     23  N   LEU A 130      45.379   8.801  55.233  1.00 38.56           N
    ATOM     24  CA  LEU A 130      44.206   7.928  55.131  1.00 33.77           C
    ATOM     25  C   LEU A 130      42.996   8.631  54.479  1.00 32.58           C
    ATOM     26  O   LEU A 130      42.052   7.931  54.114  1.00 37.11           O
    ATOM     27  CB  LEU A 130      43.836   7.427  56.550  1.00 30.41           C
    ATOM     28  CG  LEU A 130      44.593   6.187  57.085  1.00 31.10           C
    ATOM     29  CD1 LEU A 130      46.098   6.154  56.768  1.00 30.19           C
    ATOM     30  CD2 LEU A 130      44.346   6.010  58.598  1.00 30.50           C
    ATOM     31  H   LEU A 130      45.494   9.288  56.111  1.00  0.00           H
    ATOM     32  HA  LEU A 130      44.438   7.063  54.507  1.00  0.00           H
    ATOM     33  HB2 LEU A 130      43.931   8.257  57.246  1.00  0.00           H
    ATOM     34  HB3 LEU A 130      42.781   7.156  56.578  1.00  0.00           H
    ATOM     35  HG  LEU A 130      44.164   5.317  56.585  1.00  0.00           H
    ATOM     36 HD11 LEU A 130      46.603   5.360  57.319  1.00  0.00           H
    ATOM     37 HD12 LEU A 130      46.279   5.971  55.708  1.00  0.00           H
    ATOM     38 HD13 LEU A 130      46.582   7.093  57.037  1.00  0.00           H
    ATOM     39 HD21 LEU A 130      45.274   5.955  59.169  1.00  0.00           H
    ATOM     40 HD22 LEU A 130      43.762   6.828  59.021  1.00  0.00           H
    ATOM     41 HD23 LEU A 130      43.787   5.094  58.786  1.00  0.00           H
    ATOM     42  N   LYS A 132      40.984  10.577  56.311  1.00 29.59           N
    ATOM     43  CA  LYS A 132      40.115  10.011  57.330  1.00 29.53           C
    ATOM     44  C   LYS A 132      40.087  10.898  58.570  1.00 31.58           C
    ATOM     45  O   LYS A 132      41.138  11.173  59.146  1.00 36.45           O
    ATOM     46  CB  LYS A 132      40.572   8.575  57.647  1.00 29.87           C
    ATOM     47  CG  LYS A 132      39.679   7.836  58.654  1.00 31.93           C
    ATOM     48  CD  LYS A 132      40.040   6.350  58.755  1.00 31.50           C
    ATOM     49  CE  LYS A 132      39.122   5.608  59.729  1.00 32.49           C
    ATOM     50  NZ  LYS A 132      39.338   4.155  59.677  1.00 35.79           N1+
    ATOM     51  H   LYS A 132      41.901  10.879  56.613  1.00  0.00           H
    ATOM     52  HA  LYS A 132      39.105   9.952  56.936  1.00  0.00           H
    ATOM     53  HB2 LYS A 132      40.584   8.008  56.716  1.00  0.00           H
    ATOM     54  HB3 LYS A 132      41.597   8.585  58.017  1.00  0.00           H
    ATOM     55  HG2 LYS A 132      39.758   8.296  59.639  1.00  0.00           H
    ATOM     56  HG3 LYS A 132      38.635   7.938  58.357  1.00  0.00           H
    ATOM     57  HD2 LYS A 132      39.983   5.897  57.767  1.00  0.00           H
    ATOM     58  HD3 LYS A 132      41.078   6.243  59.069  1.00  0.00           H
    ATOM     59  HE2 LYS A 132      39.291   5.959  60.747  1.00  0.00           H
    ATOM     60  HE3 LYS A 132      38.080   5.810  59.483  1.00  0.00           H
    ATOM     61  HZ1 LYS A 132      40.298   3.949  59.944  1.00  0.00           H
    ATOM     62  HZ2 LYS A 132      38.709   3.692  60.316  1.00  0.00           H
    ATOM     63  HZ3 LYS A 132      39.162   3.825  58.738  1.00  0.00           H
    ATOM     64  N   PHE A 134      38.971  11.223  62.482  1.00 28.86           N
    ATOM     65  CA  PHE A 134      38.789  10.524  63.740  1.00 30.31           C
    ATOM     66  C   PHE A 134      38.174  11.547  64.692  1.00 33.32           C
    ATOM     67  O   PHE A 134      38.711  12.648  64.803  1.00 35.77           O
    ATOM     68  CB  PHE A 134      40.162  10.054  64.245  1.00 28.06           C
    ATOM     69  CG  PHE A 134      40.842   9.019  63.370  1.00 29.98           C
    ATOM     70  CD1 PHE A 134      41.509   9.397  62.185  1.00 30.82           C
    ATOM     71  CD2 PHE A 134      40.693   7.651  63.667  1.00 33.85           C
    ATOM     72  CE1 PHE A 134      42.059   8.427  61.359  1.00 33.84           C
    ATOM     73  CE2 PHE A 134      41.255   6.697  62.832  1.00 33.98           C
    ATOM     74  CZ  PHE A 134      41.938   7.084  61.687  1.00 36.09           C
    ATOM     75  H   PHE A 134      39.614  12.005  62.521  1.00  0.00           H
    ATOM     76  HA  PHE A 134      38.136   9.660  63.631  1.00  0.00           H
    ATOM     77  HB2 PHE A 134      40.835  10.900  64.368  1.00  0.00           H
    ATOM     78  HB3 PHE A 134      40.043   9.620  65.233  1.00  0.00           H
    ATOM     79  HD1 PHE A 134      41.603  10.440  61.919  1.00  0.00           H
    ATOM     80  HD2 PHE A 134      40.159   7.348  64.555  1.00  0.00           H
    ATOM     81  HE1 PHE A 134      42.583   8.717  60.461  1.00  0.00           H
    ATOM     82  HE2 PHE A 134      41.168   5.650  63.078  1.00  0.00           H
    ATOM     83  HZ  PHE A 134      42.381   6.334  61.049  1.00  0.00           H
    ATOM     84  N   CYS A 135      37.060  11.197  65.345  1.00 32.07           N
    ATOM     85  CA  CYS A 135      36.314  12.137  66.179  1.00 34.04           C
    ATOM     86  C   CYS A 135      35.552  11.410  67.288  1.00 33.67           C
    ATOM     87  O   CYS A 135      35.268  10.224  67.166  1.00 32.60           O
    ATOM     88  CB  CYS A 135      35.403  13.057  65.334  1.00 36.88           C
    ATOM     89  SG  CYS A 135      34.050  12.158  64.510  1.00 38.39           S
    ATOM     90  H   CYS A 135      36.648  10.282  65.211  1.00  0.00           H
    ATOM     91  HA  CYS A 135      37.039  12.766  66.694  1.00  0.00           H
    ATOM     92  HB2 CYS A 135      34.958  13.816  65.977  1.00  0.00           H
    ATOM     93  HB3 CYS A 135      35.982  13.595  64.582  1.00  0.00           H
    ATOM     94  HG  CYS A 135      33.369  11.871  65.623  1.00  0.00           H
    ATOM     95  N   GLN A 136      35.233  12.154  68.353  1.00 38.65           N
    ATOM     96  CA  GLN A 136      34.382  11.699  69.447  1.00 44.44           C
    ATOM     97  C   GLN A 136      32.905  11.969  69.144  1.00 40.80           C
    ATOM     98  O   GLN A 136      32.583  12.981  68.524  1.00 40.06           O
    ATOM     99  CB  GLN A 136      34.816  12.403  70.742  1.00 49.01           C
    ATOM    100  CG  GLN A 136      36.074  11.779  71.363  1.00 52.81           C
    ATOM    101  CD  GLN A 136      36.582  12.532  72.594  1.00 59.11           C
    ATOM    102  OE1 GLN A 136      36.366  13.734  72.742  1.00 58.79           O
    ATOM    103  NE2 GLN A 136      37.288  11.823  73.477  1.00 65.32           N
    ATOM    104  H   GLN A 136      35.534  13.119  68.394  1.00  0.00           H
    ATOM    105  HA  GLN A 136      34.508  10.625  69.581  1.00  0.00           H
    ATOM    106  HB2 GLN A 136      35.001  13.452  70.516  1.00  0.00           H
    ATOM    107  HB3 GLN A 136      34.010  12.377  71.480  1.00  0.00           H
    ATOM    108  HG2 GLN A 136      35.882  10.737  71.625  1.00  0.00           H
    ATOM    109  HG3 GLN A 136      36.872  11.773  70.629  1.00  0.00           H
    ATOM    110 HE21 GLN A 136      37.463  10.838  73.325  1.00  0.00           H
    ATOM    111 HE22 GLN A 136      37.668  12.271  74.298  1.00  0.00           H
    ATOM    112  N   LEU A 137      32.048  11.061  69.634  1.00 39.33           N
    ATOM    113  CA  LEU A 137      30.587  11.102  69.596  1.00 41.58           C
    ATOM    114  C   LEU A 137      30.027  12.425  70.157  1.00 46.12           C
    ATOM    115  O   LEU A 137      30.308  12.766  71.306  1.00 48.22           O
    ATOM    116  CB  LEU A 137      30.085   9.836  70.339  1.00 42.77           C
    ATOM    117  CG  LEU A 137      28.596   9.754  70.743  1.00 46.01           C
    ATOM    118  CD1 LEU A 137      27.638  10.010  69.578  1.00 50.18           C
    ATOM    119  CD2 LEU A 137      28.296   8.402  71.417  1.00 44.68           C
    ATOM    120  H   LEU A 137      32.430  10.260  70.120  1.00  0.00           H
    ATOM    121  HA  LEU A 137      30.294  11.019  68.549  1.00  0.00           H
    ATOM    122  HB2 LEU A 137      30.326   8.970  69.721  1.00  0.00           H
    ATOM    123  HB3 LEU A 137      30.668   9.718  71.255  1.00  0.00           H
    ATOM    124  HG  LEU A 137      28.413  10.524  71.494  1.00  0.00           H
    ATOM    125 HD11 LEU A 137      26.774   9.346  69.614  1.00  0.00           H
    ATOM    126 HD12 LEU A 137      27.256  11.029  69.616  1.00  0.00           H
    ATOM    127 HD13 LEU A 137      28.127   9.861  68.616  1.00  0.00           H
    ATOM    128 HD21 LEU A 137      27.641   8.535  72.279  1.00  0.00           H
    ATOM    129 HD22 LEU A 137      27.816   7.697  70.738  1.00  0.00           H
    ATOM    130 HD23 LEU A 137      29.204   7.919  71.778  1.00  0.00           H
    ATOM    131  N   ALA A 138      29.255  13.127  69.309  1.00 44.65           N
    ATOM    132  CA  ALA A 138      28.524  14.374  69.563  1.00 41.91           C
    ATOM    133  C   ALA A 138      29.380  15.646  69.745  1.00 40.60           C
    ATOM    134  O   ALA A 138      28.812  16.699  70.039  1.00 44.10           O
    ATOM    135  CB  ALA A 138      27.522  14.189  70.720  1.00 42.25           C
    ATOM    136  H   ALA A 138      29.095  12.735  68.388  1.00  0.00           H
    ATOM    137  HA  ALA A 138      27.943  14.541  68.657  1.00  0.00           H
    ATOM    138  HB1 ALA A 138      26.839  15.035  70.792  1.00  0.00           H
    ATOM    139  HB2 ALA A 138      26.913  13.297  70.573  1.00  0.00           H
    ATOM    140  HB3 ALA A 138      28.024  14.093  71.683  1.00  0.00           H
    ATOM    141  N   LYS A 139      30.705  15.553  69.554  1.00 38.50           N
    ATOM    142  CA  LYS A 139      31.625  16.683  69.690  1.00 38.90           C
    ATOM    143  C   LYS A 139      31.774  17.470  68.378  1.00 35.12           C
    ATOM    144  O   LYS A 139      31.440  16.961  67.307  1.00 35.40           O
    ATOM    145  CB  LYS A 139      32.997  16.164  70.139  1.00 45.91           C
    ATOM    146  CG  LYS A 139      33.029  15.408  71.478  1.00 49.49           C
    ATOM    147  CD  LYS A 139      32.625  16.253  72.693  1.00 55.66           C
    ATOM    148  CE  LYS A 139      32.661  15.463  74.010  1.00 63.43           C
    ATOM    149  NZ  LYS A 139      34.017  14.986  74.338  1.00 69.43           N1+
    ATOM    150  H   LYS A 139      31.115  14.663  69.310  1.00  0.00           H
    ATOM    151  HA  LYS A 139      31.243  17.374  70.443  1.00  0.00           H
    ATOM    152  HB2 LYS A 139      33.424  15.528  69.362  1.00  0.00           H
    ATOM    153  HB3 LYS A 139      33.645  17.030  70.223  1.00  0.00           H
    ATOM    154  HG2 LYS A 139      32.400  14.520  71.417  1.00  0.00           H
    ATOM    155  HG3 LYS A 139      34.039  15.046  71.644  1.00  0.00           H
    ATOM    156  HD2 LYS A 139      33.272  17.129  72.767  1.00  0.00           H
    ATOM    157  HD3 LYS A 139      31.614  16.630  72.549  1.00  0.00           H
    ATOM    158  HE2 LYS A 139      32.309  16.093  74.828  1.00  0.00           H
    ATOM    159  HE3 LYS A 139      31.987  14.608  73.954  1.00  0.00           H
    ATOM    160  HZ1 LYS A 139      34.361  14.387  73.599  1.00  0.00           H
    ATOM    161  HZ2 LYS A 139      34.638  15.777  74.437  1.00  0.00           H
    ATOM    162  HZ3 LYS A 139      33.992  14.469  75.205  1.00  0.00           H
    ATOM    163  N   TYR A 163      27.989   4.014  53.615  1.00 35.45           N
    ATOM    164  CA  TYR A 163      28.942   2.940  53.925  1.00 35.99           C
    ATOM    165  C   TYR A 163      30.042   2.848  52.857  1.00 35.37           C
    ATOM    166  O   TYR A 163      29.752   2.948  51.665  1.00 31.91           O
    ATOM    167  CB  TYR A 163      28.209   1.599  54.112  1.00 35.87           C
    ATOM    168  CG  TYR A 163      27.441   1.505  55.418  1.00 38.01           C
    ATOM    169  CD1 TYR A 163      28.129   1.495  56.650  1.00 36.96           C
    ATOM    170  CD2 TYR A 163      26.034   1.465  55.407  1.00 38.63           C
    ATOM    171  CE1 TYR A 163      27.409   1.513  57.859  1.00 37.69           C
    ATOM    172  CE2 TYR A 163      25.310   1.479  56.614  1.00 35.22           C
    ATOM    173  CZ  TYR A 163      26.000   1.532  57.839  1.00 37.16           C
    ATOM    174  OH  TYR A 163      25.306   1.624  59.005  1.00 37.76           O
    ATOM    175  H   TYR A 163      27.095   3.730  53.244  1.00  0.00           H
    ATOM    176  HA  TYR A 163      29.423   3.189  54.872  1.00  0.00           H
    ATOM    177  HB2 TYR A 163      27.548   1.419  53.266  1.00  0.00           H
    ATOM    178  HB3 TYR A 163      28.926   0.779  54.107  1.00  0.00           H
    ATOM    179  HD1 TYR A 163      29.209   1.509  56.674  1.00  0.00           H
    ATOM    180  HD2 TYR A 163      25.511   1.452  54.467  1.00  0.00           H
    ATOM    181  HE1 TYR A 163      27.943   1.536  58.798  1.00  0.00           H
    ATOM    182  HE2 TYR A 163      24.231   1.483  56.592  1.00  0.00           H
    ATOM    183  HH  TYR A 163      24.347   1.669  58.900  1.00  0.00           H
    ATOM    184  N   LYS A 164      31.288   2.683  53.328  1.00 34.93           N
    ATOM    185  CA  LYS A 164      32.500   2.627  52.513  1.00 35.83           C
    ATOM    186  C   LYS A 164      32.605   1.332  51.688  1.00 37.73           C
    ATOM    187  O   LYS A 164      33.047   1.390  50.542  1.00 41.25           O
    ATOM    188  CB  LYS A 164      33.714   2.797  53.452  1.00 33.84           C
    ATOM    189  CG  LYS A 164      35.093   2.804  52.766  1.00 32.51           C
    ATOM    190  CD  LYS A 164      36.235   2.666  53.781  1.00 38.37           C
    ATOM    191  CE  LYS A 164      37.628   2.656  53.137  1.00 43.22           C
    ATOM    192  NZ  LYS A 164      38.000   3.983  52.623  1.00 46.98           N1+
    ATOM    193  H   LYS A 164      31.428   2.606  54.327  1.00  0.00           H
    ATOM    194  HA  LYS A 164      32.480   3.470  51.820  1.00  0.00           H
    ATOM    195  HB2 LYS A 164      33.606   3.718  54.028  1.00  0.00           H
    ATOM    196  HB3 LYS A 164      33.693   1.989  54.183  1.00  0.00           H
    ATOM    197  HG2 LYS A 164      35.183   1.987  52.051  1.00  0.00           H
    ATOM    198  HG3 LYS A 164      35.202   3.725  52.193  1.00  0.00           H
    ATOM    199  HD2 LYS A 164      36.167   3.469  54.513  1.00  0.00           H
    ATOM    200  HD3 LYS A 164      36.106   1.739  54.340  1.00  0.00           H
    ATOM    201  HE2 LYS A 164      38.374   2.362  53.876  1.00  0.00           H
    ATOM    202  HE3 LYS A 164      37.673   1.925  52.330  1.00  0.00           H
    ATOM    203  HZ1 LYS A 164      38.059   4.627  53.404  1.00  0.00           H
    ATOM    204  HZ2 LYS A 164      38.903   3.934  52.174  1.00  0.00           H
    ATOM    205  HZ3 LYS A 164      37.309   4.309  51.961  1.00  0.00           H
    ATOM    206  N   GLN A 165      32.229   0.195  52.294  1.00 35.64           N
    ATOM    207  CA  GLN A 165      32.329  -1.130  51.689  1.00 39.23           C
    ATOM    208  C   GLN A 165      31.178  -1.366  50.703  1.00 43.97           C
    ATOM    209  O   GLN A 165      30.031  -1.048  51.021  1.00 46.94           O
    ATOM    210  CB  GLN A 165      32.345  -2.194  52.801  1.00 38.39           C
    ATOM    211  CG  GLN A 165      33.608  -2.125  53.689  1.00 37.36           C
    ATOM    212  CD  GLN A 165      33.464  -2.881  55.012  1.00 44.27           C
    ATOM    213  OE1 GLN A 165      32.452  -2.760  55.697  1.00 47.12           O
    ATOM    214  NE2 GLN A 165      34.499  -3.617  55.417  1.00 47.74           N
    ATOM    215  H   GLN A 165      31.854   0.227  53.233  1.00  0.00           H
    ATOM    216  HA  GLN A 165      33.276  -1.192  51.151  1.00  0.00           H
    ATOM    217  HB2 GLN A 165      31.435  -2.096  53.392  1.00  0.00           H
    ATOM    218  HB3 GLN A 165      32.296  -3.184  52.355  1.00  0.00           H
    ATOM    219  HG2 GLN A 165      34.470  -2.490  53.130  1.00  0.00           H
    ATOM    220  HG3 GLN A 165      33.831  -1.092  53.951  1.00  0.00           H
    ATOM    221 HE21 GLN A 165      35.333  -3.694  54.854  1.00  0.00           H
    ATOM    222 HE22 GLN A 165      34.440  -4.126  56.287  1.00  0.00           H
    ATOM    223  N   SER A 166      31.515  -1.930  49.532  1.00 46.20           N
    ATOM    224  CA  SER A 166      30.604  -2.199  48.416  1.00 49.12           C
    ATOM    225  C   SER A 166      29.427  -3.137  48.748  1.00 46.06           C
    ATOM    226  O   SER A 166      28.351  -2.959  48.176  1.00 43.80           O
    ATOM    227  CB  SER A 166      31.429  -2.693  47.210  1.00 56.88           C
    ATOM    228  OG  SER A 166      32.005  -3.965  47.445  1.00 62.68           O
    ATOM    229  H   SER A 166      32.484  -2.173  49.368  1.00  0.00           H
    ATOM    230  HA  SER A 166      30.167  -1.239  48.139  1.00  0.00           H
    ATOM    231  HB2 SER A 166      30.794  -2.763  46.326  1.00  0.00           H
    ATOM    232  HB3 SER A 166      32.220  -1.981  46.969  1.00  0.00           H
    ATOM    233  HG  SER A 166      32.421  -4.269  46.635  1.00  0.00           H
    ATOM    234  N   GLN A 167      29.636  -4.076  49.689  1.00 48.12           N
    ATOM    235  CA  GLN A 167      28.613  -4.989  50.201  1.00 51.11           C
    ATOM    236  C   GLN A 167      27.530  -4.314  51.072  1.00 54.33           C
    ATOM    237  O   GLN A 167      26.455  -4.895  51.216  1.00 59.32           O
    ATOM    238  CB  GLN A 167      29.287  -6.173  50.934  1.00 56.34           C
    ATOM    239  CG  GLN A 167      30.117  -5.813  52.190  1.00 62.29           C
    ATOM    240  CD  GLN A 167      31.631  -5.863  51.973  1.00 69.46           C
    ATOM    241  OE1 GLN A 167      32.137  -5.411  50.947  1.00 73.40           O
    ATOM    242  NE2 GLN A 167      32.368  -6.371  52.963  1.00 70.14           N
    ATOM    243  H   GLN A 167      30.559  -4.177  50.089  1.00  0.00           H
    ATOM    244  HA  GLN A 167      28.096  -5.405  49.334  1.00  0.00           H
    ATOM    245  HB2 GLN A 167      28.498  -6.858  51.250  1.00  0.00           H
    ATOM    246  HB3 GLN A 167      29.881  -6.755  50.227  1.00  0.00           H
    ATOM    247  HG2 GLN A 167      29.862  -4.824  52.563  1.00  0.00           H
    ATOM    248  HG3 GLN A 167      29.860  -6.511  52.989  1.00  0.00           H
    ATOM    249 HE21 GLN A 167      31.931  -6.711  53.808  1.00  0.00           H
    ATOM    250 HE22 GLN A 167      33.373  -6.418  52.874  1.00  0.00           H
    ATOM    251  N   HID A 168      27.814  -3.115  51.613  1.00 51.00           N
    ATOM    252  CA  HID A 168      26.925  -2.369  52.512  1.00 41.69           C
    ATOM    253  C   HID A 168      26.477  -1.011  51.948  1.00 40.62           C
    ATOM    254  O   HID A 168      25.669  -0.356  52.602  1.00 42.57           O
    ATOM    255  CB  HID A 168      27.625  -2.164  53.869  1.00 41.53           C
    ATOM    256  CG  HID A 168      28.063  -3.427  54.565  1.00 48.80           C
    ATOM    257  ND1 HID A 168      27.213  -4.483  54.850  1.00 54.96           N
    ATOM    258  CD2 HID A 168      29.296  -3.816  55.031  1.00 51.64           C
    ATOM    259  CE1 HID A 168      27.942  -5.424  55.455  1.00 56.67           C
    ATOM    260  NE2 HID A 168      29.218  -5.085  55.605  1.00 55.10           N
    ATOM    261  H   HID A 168      28.716  -2.693  51.435  1.00  0.00           H
    ATOM    262  HA  HID A 168      26.011  -2.935  52.692  1.00  0.00           H
    ATOM    263  HB2 HID A 168      28.509  -1.539  53.734  1.00  0.00           H
    ATOM    264  HB3 HID A 168      26.972  -1.625  54.557  1.00  0.00           H
    ATOM    265  HD1 HID A 168      26.225  -4.544  54.620  1.00  0.00           H
    ATOM    266  HD2 HID A 168      30.225  -3.277  54.992  1.00  0.00           H
    ATOM    267  HE1 HID A 168      27.538  -6.369  55.792  1.00  0.00           H
    ATOM    268  N   MET A 169      26.990  -0.597  50.775  1.00 36.95           N
    ATOM    269  CA  MET A 169      26.750   0.711  50.146  1.00 41.89           C
    ATOM    270  C   MET A 169      25.271   1.084  49.918  1.00 47.38           C
    ATOM    271  O   MET A 169      24.946   2.271  49.973  1.00 48.76           O
    ATOM    272  CB  MET A 169      27.523   0.779  48.812  1.00 45.67           C
    ATOM    273  CG  MET A 169      28.806   1.608  48.882  1.00 51.55           C
    ATOM    274  SD  MET A 169      29.640   1.734  47.280  1.00 56.59           S
    ATOM    275  CE  MET A 169      31.357   1.719  47.841  1.00 56.26           C
    ATOM    276  H   MET A 169      27.652  -1.193  50.300  1.00  0.00           H
    ATOM    277  HA  MET A 169      27.150   1.465  50.827  1.00  0.00           H
    ATOM    278  HB2 MET A 169      27.773  -0.227  48.476  1.00  0.00           H
    ATOM    279  HB3 MET A 169      26.902   1.195  48.020  1.00  0.00           H
    ATOM    280  HG2 MET A 169      28.587   2.619  49.228  1.00  0.00           H
    ATOM    281  HG3 MET A 169      29.486   1.165  49.609  1.00  0.00           H
    ATOM    282  HE1 MET A 169      32.032   1.878  47.001  1.00  0.00           H
    ATOM    283  HE2 MET A 169      31.515   2.516  48.568  1.00  0.00           H
    ATOM    284  HE3 MET A 169      31.602   0.765  48.305  1.00  0.00           H
    ATOM    285  N   THR A 170      24.411   0.081  49.672  1.00 48.56           N
    ATOM    286  CA  THR A 170      22.973   0.268  49.460  1.00 47.21           C
    ATOM    287  C   THR A 170      22.164   0.347  50.773  1.00 41.65           C
    ATOM    288  O   THR A 170      21.018   0.794  50.718  1.00 40.39           O
    ATOM    289  CB  THR A 170      22.373  -0.862  48.583  1.00 48.53           C
    ATOM    290  OG1 THR A 170      22.445  -2.129  49.211  1.00 49.40           O
    ATOM    291  CG2 THR A 170      23.040  -0.956  47.201  1.00 49.80           C
    ATOM    292  H   THR A 170      24.749  -0.871  49.636  1.00  0.00           H
    ATOM    293  HA  THR A 170      22.818   1.210  48.929  1.00  0.00           H
    ATOM    294  HB  THR A 170      21.315  -0.654  48.418  1.00  0.00           H
    ATOM    295 HG21 THR A 170      22.585  -1.743  46.599  1.00  0.00           H
    ATOM    296 HG22 THR A 170      22.927  -0.020  46.653  1.00  0.00           H
    ATOM    297 HG23 THR A 170      24.107  -1.167  47.270  1.00  0.00           H
    ATOM    298  HG1 THR A 170      23.296  -2.521  48.986  1.00  0.00           H
    ATOM    299  N   GLU A 171      22.754  -0.027  51.925  1.00 38.49           N
    ATOM    300  CA  GLU A 171      22.148   0.206  53.235  1.00 39.26           C
    ATOM    301  C   GLU A 171      22.307   1.685  53.615  1.00 38.87           C
    ATOM    302  O   GLU A 171      23.413   2.221  53.539  1.00 38.61           O
    ATOM    303  CB  GLU A 171      22.746  -0.748  54.290  1.00 41.53           C
    ATOM    304  CG  GLU A 171      22.058  -0.658  55.672  1.00 46.44           C
    ATOM    305  CD  GLU A 171      22.631  -1.627  56.707  1.00 52.19           C
    ATOM    306  OE1 GLU A 171      23.146  -2.690  56.295  1.00 53.76           O
    ATOM    307  OE2 GLU A 171      22.562  -1.289  57.908  1.00 56.25           O1-
    ATOM    308  H   GLU A 171      23.709  -0.360  51.925  1.00  0.00           H
    ATOM    309  HA  GLU A 171      21.088  -0.030  53.159  1.00  0.00           H
    ATOM    310  HB2 GLU A 171      22.674  -1.768  53.910  1.00  0.00           H
    ATOM    311  HB3 GLU A 171      23.811  -0.547  54.412  1.00  0.00           H
    ATOM    312  HG2 GLU A 171      22.139   0.352  56.075  1.00  0.00           H
    ATOM    313  HG3 GLU A 171      20.993  -0.868  55.569  1.00  0.00           H
    ATOM    314  N   VAL A 172      21.193   2.318  54.007  1.00 37.81           N
    ATOM    315  CA  VAL A 172      21.169   3.708  54.454  1.00 34.58           C
    ATOM    316  C   VAL A 172      21.836   3.824  55.835  1.00 38.75           C
    ATOM    317  O   VAL A 172      21.415   3.127  56.760  1.00 39.37           O
    ATOM    318  CB  VAL A 172      19.711   4.237  54.555  1.00 31.90           C
    ATOM    319  CG1 VAL A 172      19.598   5.654  55.150  1.00 21.74           C
    ATOM    320  CG2 VAL A 172      19.011   4.173  53.186  1.00 32.61           C
    ATOM    321  H   VAL A 172      20.313   1.820  54.032  1.00  0.00           H
    ATOM    322  HA  VAL A 172      21.715   4.316  53.731  1.00  0.00           H
    ATOM    323  HB  VAL A 172      19.145   3.582  55.218  1.00  0.00           H
    ATOM    324 HG11 VAL A 172      18.574   6.018  55.105  1.00  0.00           H
    ATOM    325 HG12 VAL A 172      19.891   5.691  56.199  1.00  0.00           H
    ATOM    326 HG13 VAL A 172      20.232   6.344  54.599  1.00  0.00           H
    ATOM    327 HG21 VAL A 172      18.032   4.654  53.206  1.00  0.00           H
    ATOM    328 HG22 VAL A 172      19.607   4.643  52.409  1.00  0.00           H
    ATOM    329 HG23 VAL A 172      18.858   3.140  52.877  1.00  0.00           H
    ATOM    330  N   VAL A 173      22.846   4.704  55.947  1.00 42.21           N
    ATOM    331  CA  VAL A 173      23.510   5.013  57.213  1.00 36.50           C
    ATOM    332  C   VAL A 173      22.558   5.817  58.110  1.00 36.63           C
    ATOM    333  O   VAL A 173      22.054   6.861  57.697  1.00 34.15           O
    ATOM    334  CB  VAL A 173      24.820   5.829  57.027  1.00 30.08           C
    ATOM    335  CG1 VAL A 173      25.467   6.273  58.357  1.00 27.84           C
    ATOM    336  CG2 VAL A 173      25.859   5.039  56.223  1.00 27.47           C
    ATOM    337  H   VAL A 173      23.134   5.244  55.142  1.00  0.00           H
    ATOM    338  HA  VAL A 173      23.767   4.071  57.702  1.00  0.00           H
    ATOM    339  HB  VAL A 173      24.590   6.729  56.454  1.00  0.00           H
    ATOM    340 HG11 VAL A 173      26.453   6.705  58.191  1.00  0.00           H
    ATOM    341 HG12 VAL A 173      24.874   7.028  58.874  1.00  0.00           H
    ATOM    342 HG13 VAL A 173      25.599   5.429  59.036  1.00  0.00           H
    ATOM    343 HG21 VAL A 173      26.745   5.646  56.046  1.00  0.00           H
    ATOM    344 HG22 VAL A 173      26.182   4.151  56.763  1.00  0.00           H
    ATOM    345 HG23 VAL A 173      25.470   4.721  55.255  1.00  0.00           H
    ATOM    346  N   ARG A 174      22.341   5.299  59.320  1.00 35.23           N
    ATOM    347  CA  ARG A 174      21.465   5.875  60.329  1.00 39.17           C
    ATOM    348  C   ARG A 174      21.916   5.370  61.701  1.00 38.92           C
    ATOM    349  O   ARG A 174      22.700   4.421  61.793  1.00 36.19           O
    ATOM    350  CB  ARG A 174      19.987   5.562  59.992  1.00 40.33           C
    ATOM    351  CG  ARG A 174      19.583   4.074  60.030  1.00 46.58           C
    ATOM    352  CD  ARG A 174      18.317   3.760  59.214  1.00 57.02           C
    ATOM    353  NE  ARG A 174      17.164   4.584  59.610  1.00 69.13           N
    ATOM    354  CZ  ARG A 174      16.462   4.487  60.754  1.00 77.29           C
    ATOM    355  NH1 ARG A 174      16.724   3.540  61.666  1.00 79.33           N
    ATOM    356  NH2 ARG A 174      15.471   5.352  60.995  1.00 78.66           N1+
    ATOM    357  H   ARG A 174      22.792   4.433  59.584  1.00  0.00           H
    ATOM    358  HA  ARG A 174      21.600   6.956  60.339  1.00  0.00           H
    ATOM    359  HB2 ARG A 174      19.338   6.119  60.668  1.00  0.00           H
    ATOM    360  HB3 ARG A 174      19.777   5.955  58.997  1.00  0.00           H
    ATOM    361  HG2 ARG A 174      20.376   3.525  59.524  1.00  0.00           H
    ATOM    362  HG3 ARG A 174      19.546   3.664  61.039  1.00  0.00           H
    ATOM    363  HD2 ARG A 174      18.526   3.833  58.146  1.00  0.00           H
    ATOM    364  HD3 ARG A 174      18.019   2.727  59.395  1.00  0.00           H
    ATOM    365  HE  ARG A 174      16.907   5.310  58.957  1.00  0.00           H
    ATOM    366 HH11 ARG A 174      17.465   2.874  61.507  1.00  0.00           H
    ATOM    367 HH12 ARG A 174      16.199   3.503  62.532  1.00  0.00           H
    ATOM    368 HH21 ARG A 174      15.219   6.067  60.320  1.00  0.00           H
    ATOM    369 HH22 ARG A 174      14.989   5.331  61.888  1.00  0.00           H
    ATOM    370  N   ARG A 175      21.422   6.037  62.752  1.00 38.57           N
    ATOM    371  CA  ARG A 175      21.738   5.715  64.143  1.00 37.77           C
    ATOM    372  C   ARG A 175      21.222   4.324  64.553  1.00 39.45           C
    ATOM    373  O   ARG A 175      20.258   3.824  63.969  1.00 40.08           O
    ATOM    374  CB  ARG A 175      21.177   6.818  65.060  1.00 31.75           C
    ATOM    375  CG  ARG A 175      21.744   8.216  64.743  1.00 29.45           C
    ATOM    376  CD  ARG A 175      21.174   9.304  65.658  1.00 33.63           C
    ATOM    377  NE  ARG A 175      21.797   9.313  66.978  1.00 37.66           N
    ATOM    378  CZ  ARG A 175      21.356   9.983  68.055  1.00 36.11           C
    ATOM    379  NH1 ARG A 175      20.229  10.706  68.036  1.00 28.06           N
    ATOM    380  NH2 ARG A 175      22.083   9.938  69.174  1.00 40.11           N1+
    ATOM    381  H   ARG A 175      20.774   6.800  62.605  1.00  0.00           H
    ATOM    382  HA  ARG A 175      22.823   5.718  64.231  1.00  0.00           H
    ATOM    383  HB2 ARG A 175      20.090   6.841  64.969  1.00  0.00           H
    ATOM    384  HB3 ARG A 175      21.386   6.573  66.102  1.00  0.00           H
    ATOM    385  HG2 ARG A 175      22.822   8.160  64.900  1.00  0.00           H
    ATOM    386  HG3 ARG A 175      21.613   8.505  63.702  1.00  0.00           H
    ATOM    387  HD2 ARG A 175      21.121  10.284  65.196  1.00  0.00           H
    ATOM    388  HD3 ARG A 175      20.152   9.012  65.877  1.00  0.00           H
    ATOM    389  HE  ARG A 175      22.685   8.832  67.055  1.00  0.00           H
    ATOM    390 HH11 ARG A 175      19.648  10.733  67.209  1.00  0.00           H
    ATOM    391 HH12 ARG A 175      19.925  11.182  68.877  1.00  0.00           H
    ATOM    392 HH21 ARG A 175      22.956   9.429  69.186  1.00  0.00           H
    ATOM    393 HH22 ARG A 175      21.838  10.519  69.975  1.00  0.00           H
    ATOM    394  N   CY2 A 176      21.868   3.733  65.571  1.00 40.65           N
    ATOM    395  CA  CY2 A 176      21.430   2.482  66.196  1.00 40.13           C
    ATOM    396  C   CY2 A 176      20.059   2.654  66.892  1.00 44.70           C
    ATOM    397  O   CY2 A 176      19.723   3.788  67.241  1.00 46.47           O
    ATOM    398  CB  CY2 A 176      22.501   1.901  67.153  1.00 34.41           C
    ATOM    399  SG  CY2 A 176      22.852   2.828  68.704  1.00 39.01           S
    ATOM    400  H   CY2 A 176      22.645   4.205  66.012  1.00  0.00           H
    ATOM    401  HA  CY2 A 176      21.316   1.780  65.371  1.00  0.00           H
    ATOM    402  HB2 CY2 A 176      22.220   0.889  67.435  1.00  0.00           H
    ATOM    403  HB3 CY2 A 176      23.426   1.773  66.598  1.00  0.00           H
    ATOM    404  N   PRO A 177      19.311   1.547  67.122  1.00 45.56           N
    ATOM    405  CA  PRO A 177      18.029   1.587  67.852  1.00 43.86           C
    ATOM    406  C   PRO A 177      18.034   2.331  69.197  1.00 45.35           C
    ATOM    407  O   PRO A 177      17.072   3.036  69.473  1.00 44.93           O
    ATOM    408  CB  PRO A 177      17.654   0.108  68.021  1.00 42.53           C
    ATOM    409  CG  PRO A 177      18.284  -0.574  66.821  1.00 40.35           C
    ATOM    410  CD  PRO A 177      19.583   0.196  66.623  1.00 41.90           C
    ATOM    411  HA  PRO A 177      17.305   2.064  67.187  1.00  0.00           H
    ATOM    412  HB2 PRO A 177      18.084  -0.300  68.939  1.00  0.00           H
    ATOM    413  HB3 PRO A 177      16.577  -0.051  68.062  1.00  0.00           H
    ATOM    414  HG2 PRO A 177      18.435  -1.645  66.956  1.00  0.00           H
    ATOM    415  HG3 PRO A 177      17.643  -0.429  65.950  1.00  0.00           H
    ATOM    416  HD2 PRO A 177      20.375  -0.261  67.212  1.00  0.00           H
    ATOM    417  HD3 PRO A 177      19.873   0.172  65.573  1.00  0.00           H
    ATOM    418  N   HIE A 178      19.122   2.212  69.974  1.00 47.74           N
    ATOM    419  CA  HIE A 178      19.307   2.909  71.245  1.00 49.40           C
    ATOM    420  C   HIE A 178      19.477   4.429  71.113  1.00 48.03           C
    ATOM    421  O   HIE A 178      18.741   5.164  71.773  1.00 52.80           O
    ATOM    422  CB  HIE A 178      20.472   2.267  72.016  1.00 51.74           C
    ATOM    423  CG  HIE A 178      19.977   1.264  73.013  1.00 58.07           C
    ATOM    424  ND1 HIE A 178      19.657  -0.051  72.665  1.00 58.90           N
    ATOM    425  CD2 HIE A 178      19.683   1.466  74.343  1.00 58.77           C
    ATOM    426  CE1 HIE A 178      19.184  -0.585  73.781  1.00 57.33           C
    ATOM    427  NE2 HIE A 178      19.172   0.269  74.802  1.00 58.07           N
    ATOM    428  H   HIE A 178      19.885   1.620  69.677  1.00  0.00           H
    ATOM    429  HA  HIE A 178      18.393   2.771  71.829  1.00  0.00           H
    ATOM    430  HB2 HIE A 178      21.178   1.783  71.339  1.00  0.00           H
    ATOM    431  HB3 HIE A 178      21.055   3.006  72.561  1.00  0.00           H
    ATOM    432  HD2 HIE A 178      19.780   2.339  74.969  1.00  0.00           H
    ATOM    433  HE2 HIE A 178      18.845   0.077  75.742  1.00  0.00           H
    ATOM    434  HE1 HIE A 178      18.831  -1.604  73.852  1.00  0.00           H
    ATOM    435  N   HE1 A 179      20.419   4.870  70.262  1.00 42.89           N
    ATOM    436  CA  HE1 A 179      20.674   6.288  69.994  1.00 38.37           C
    ATOM    437  C   HE1 A 179      19.500   7.010  69.309  1.00 37.61           C
    ATOM    438  O   HE1 A 179      19.304   8.197  69.564  1.00 37.56           O
    ATOM    439  CB  HE1 A 179      21.961   6.445  69.168  1.00 37.30           C
    ATOM    440  CG  HE1 A 179      23.233   6.575  69.969  1.00 34.39           C
    ATOM    441  ND1 HE1 A 179      24.301   5.694  69.829  1.00 34.18           N
    ATOM    442  CD2 HE1 A 179      23.597   7.555  70.870  1.00 32.41           C
    ATOM    443  CE1 HE1 A 179      25.235   6.126  70.662  1.00 35.16           C
    ATOM    444  NE2 HE1 A 179      24.881   7.247  71.286  1.00 32.74           N
    ATOM    445  H   HE1 A 179      20.980   4.202  69.748  1.00  0.00           H
    ATOM    446  HA  HE1 A 179      20.816   6.785  70.954  1.00  0.00           H
    ATOM    447  HB2 HE1 A 179      22.052   5.616  68.467  1.00  0.00           H
    ATOM    448  HB3 HE1 A 179      21.911   7.329  68.544  1.00  0.00           H
    ATOM    449  HD2 HE1 A 179      23.067   8.419  71.242  1.00  0.00           H
    ATOM    450  HE2 HE1 A 179      25.435   7.766  71.954  1.00  0.00           H
    ATOM    451  HE1 HE1 A 179      26.193   5.644  70.798  1.00  0.00           H
    ATOM    452  N   GLU A 180      18.720   6.280  68.494  1.00 33.62           N
    ATOM    453  CA  GLU A 180      17.516   6.776  67.831  1.00 34.07           C
    ATOM    454  C   GLU A 180      16.400   7.221  68.800  1.00 39.87           C
    ATOM    455  O   GLU A 180      15.650   8.125  68.443  1.00 39.75           O
    ATOM    456  CB  GLU A 180      17.041   5.736  66.791  1.00 34.43           C
    ATOM    457  CG  GLU A 180      15.760   6.128  66.017  1.00 42.05           C
    ATOM    458  CD  GLU A 180      15.396   5.164  64.886  1.00 50.46           C
    ATOM    459  OE1 GLU A 180      15.777   3.975  64.968  1.00 48.19           O
    ATOM    460  OE2 GLU A 180      14.751   5.642  63.926  1.00 54.83           O1-
    ATOM    461  H   GLU A 180      18.953   5.310  68.322  1.00  0.00           H
    ATOM    462  HA  GLU A 180      17.815   7.665  67.276  1.00  0.00           H
    ATOM    463  HB2 GLU A 180      17.849   5.567  66.077  1.00  0.00           H
    ATOM    464  HB3 GLU A 180      16.879   4.779  67.288  1.00  0.00           H
    ATOM    465  HG2 GLU A 180      14.903   6.173  66.689  1.00  0.00           H
    ATOM    466  HG3 GLU A 180      15.882   7.127  65.594  1.00  0.00           H
    ATOM    467  N   ARG A 181      16.321   6.630  70.005  1.00 44.80           N
    ATOM    468  CA  ARG A 181      15.325   6.983  71.027  1.00 41.51           C
    ATOM    469  C   ARG A 181      15.951   7.481  72.343  1.00 41.42           C
    ATOM    470  O   ARG A 181      15.211   7.640  73.312  1.00 43.27           O
    ATOM    471  CB  ARG A 181      14.271   5.856  71.211  1.00 41.63           C
    ATOM    472  CG  ARG A 181      14.791   4.417  71.101  1.00 48.77           C
    ATOM    473  CD  ARG A 181      13.736   3.369  71.481  1.00 59.39           C
    ATOM    474  NE  ARG A 181      14.208   1.995  71.257  1.00 69.92           N
    ATOM    475  CZ  ARG A 181      14.074   1.308  70.110  1.00 80.41           C
    ATOM    476  NH1 ARG A 181      13.487   1.854  69.035  1.00 80.65           N
    ATOM    477  NH2 ARG A 181      14.528   0.053  70.032  1.00 85.04           N1+
    ATOM    478  H   ARG A 181      16.983   5.906  70.251  1.00  0.00           H
    ATOM    479  HA  ARG A 181      14.760   7.852  70.687  1.00  0.00           H
    ATOM    480  HB2 ARG A 181      13.728   5.984  72.148  1.00  0.00           H
    ATOM    481  HB3 ARG A 181      13.516   5.977  70.433  1.00  0.00           H
    ATOM    482  HG2 ARG A 181      15.148   4.232  70.094  1.00  0.00           H
    ATOM    483  HG3 ARG A 181      15.639   4.321  71.779  1.00  0.00           H
    ATOM    484  HD2 ARG A 181      13.573   3.439  72.553  1.00  0.00           H
    ATOM    485  HD3 ARG A 181      12.758   3.564  71.039  1.00  0.00           H
    ATOM    486  HE  ARG A 181      14.696   1.559  72.028  1.00  0.00           H
    ATOM    487 HH11 ARG A 181      13.156   2.807  69.075  1.00  0.00           H
    ATOM    488 HH12 ARG A 181      13.397   1.336  68.174  1.00  0.00           H
    ATOM    489 HH21 ARG A 181      15.061  -0.344  70.799  1.00  0.00           H
    ATOM    490 HH22 ARG A 181      14.410  -0.485  69.186  1.00  0.00           H
    ATOM    491  N   CYS A 182      17.257   7.815  72.355  1.00 41.82           N
    ATOM    492  CA  CYS A 182      17.882   8.540  73.467  1.00 41.00           C
    ATOM    493  C   CYS A 182      17.514  10.042  73.416  1.00 41.32           C
    ATOM    494  O   CYS A 182      17.289  10.578  72.331  1.00 45.39           O
    ATOM    495  CB  CYS A 182      19.392   8.247  73.590  1.00 42.10           C
    ATOM    496  SG  CYS A 182      20.460   9.219  72.502  1.00 42.09           S
    ATOM    497  H   CYS A 182      17.822   7.687  71.527  1.00  0.00           H
    ATOM    498  HA  CYS A 182      17.458   8.120  74.372  1.00  0.00           H
    ATOM    499  HB2 CYS A 182      19.721   8.453  74.608  1.00  0.00           H
    ATOM    500  HB3 CYS A 182      19.580   7.189  73.419  1.00  0.00           H
    ATOM    501  HG  CYS A 182      20.249  10.390  73.110  1.00  0.00           H
    ATOM    502  N   SER A 183      17.437  10.686  74.590  1.00 41.57           N
    ATOM    503  CA  SER A 183      16.989  12.070  74.772  1.00 46.67           C
    ATOM    504  C   SER A 183      18.125  13.100  74.587  1.00 50.27           C
    ATOM    505  O   SER A 183      18.477  13.800  75.538  1.00 55.52           O
    ATOM    506  CB  SER A 183      16.254  12.169  76.132  1.00 53.12           C
    ATOM    507  OG  SER A 183      17.092  11.904  77.243  1.00 60.60           O
    ATOM    508  H   SER A 183      17.664  10.196  75.446  1.00  0.00           H
    ATOM    509  HA  SER A 183      16.244  12.293  74.005  1.00  0.00           H
    ATOM    510  HB2 SER A 183      15.833  13.168  76.256  1.00  0.00           H
    ATOM    511  HB3 SER A 183      15.413  11.479  76.165  1.00  0.00           H
    ATOM    512  HG  SER A 183      16.581  12.017  78.048  1.00  0.00           H
    ATOM    513  N   ASP A 184      18.665  13.196  73.359  1.00 45.91           N
    ATOM    514  CA  ASP A 184      19.760  14.116  73.020  1.00 36.72           C
    ATOM    515  C   ASP A 184      19.360  15.160  71.958  1.00 34.87           C
    ATOM    516  O   ASP A 184      20.258  15.770  71.381  1.00 38.21           O
    ATOM    517  CB  ASP A 184      21.073  13.384  72.633  1.00 31.82           C
    ATOM    518  CG  ASP A 184      21.027  12.455  71.414  1.00 33.23           C
    ATOM    519  OD1 ASP A 184      19.958  12.303  70.786  1.00 38.03           O
    ATOM    520  OD2 ASP A 184      22.102  11.890  71.126  1.00 35.00           O1-
    ATOM    521  H   ASP A 184      18.348  12.584  72.619  1.00  0.00           H
    ATOM    522  HA  ASP A 184      20.011  14.722  73.891  1.00  0.00           H
    ATOM    523  HB2 ASP A 184      21.869  14.114  72.475  1.00  0.00           H
    ATOM    524  HB3 ASP A 184      21.369  12.773  73.486  1.00  0.00           H
    ATOM    525  N   SER A 185      18.054  15.366  71.706  1.00 34.16           N
    ATOM    526  CA  SER A 185      17.542  16.319  70.713  1.00 32.81           C
    ATOM    527  C   SER A 185      17.993  17.769  70.983  1.00 36.09           C
    ATOM    528  O   SER A 185      17.964  18.211  72.132  1.00 43.23           O
    ATOM    529  CB  SER A 185      16.003  16.230  70.680  1.00 32.62           C
    ATOM    530  OG  SER A 185      15.451  17.024  69.648  1.00 32.79           O
    ATOM    531  H   SER A 185      17.360  14.852  72.231  1.00  0.00           H
    ATOM    532  HA  SER A 185      17.927  16.005  69.741  1.00  0.00           H
    ATOM    533  HB2 SER A 185      15.681  15.198  70.533  1.00  0.00           H
    ATOM    534  HB3 SER A 185      15.576  16.563  71.626  1.00  0.00           H
    ATOM    535  HG  SER A 185      15.486  16.518  68.830  1.00  0.00           H
    ATOM    536  N   ASP A 186      18.364  18.477  69.907  1.00 35.10           N
    ATOM    537  CA  ASP A 186      18.643  19.919  69.897  1.00 34.37           C
    ATOM    538  C   ASP A 186      17.386  20.745  69.532  1.00 34.08           C
    ATOM    539  O   ASP A 186      17.452  21.975  69.552  1.00 34.94           O
    ATOM    540  CB  ASP A 186      19.825  20.290  68.961  1.00 33.54           C
    ATOM    541  CG  ASP A 186      19.767  19.735  67.531  1.00 33.08           C
    ATOM    542  OD1 ASP A 186      18.737  19.136  67.152  1.00 33.74           O
    ATOM    543  OD2 ASP A 186      20.768  19.952  66.818  1.00 35.78           O1-
    ATOM    544  H   ASP A 186      18.363  18.042  68.994  1.00  0.00           H
    ATOM    545  HA  ASP A 186      18.931  20.231  70.903  1.00  0.00           H
    ATOM    546  HB2 ASP A 186      19.928  21.374  68.902  1.00  0.00           H
    ATOM    547  HB3 ASP A 186      20.736  19.905  69.421  1.00  0.00           H
    ATOM    548  N   PRO A 191      15.597  12.541  64.946  1.00 33.82           N
    ATOM    549  CA  PRO A 191      16.256  11.623  65.897  1.00 29.28           C
    ATOM    550  C   PRO A 191      17.084  10.466  65.298  1.00 29.55           C
    ATOM    551  O   PRO A 191      17.895   9.901  66.026  1.00 34.74           O
    ATOM    552  CB  PRO A 191      15.090  11.109  66.758  1.00 30.42           C
    ATOM    553  CG  PRO A 191      13.854  11.271  65.896  1.00 33.93           C
    ATOM    554  CD  PRO A 191      14.149  12.566  65.160  1.00 34.32           C
    ATOM    555  HA  PRO A 191      16.934  12.200  66.527  1.00  0.00           H
    ATOM    556  HB2 PRO A 191      15.209  10.082  67.086  1.00  0.00           H
    ATOM    557  HB3 PRO A 191      14.994  11.735  67.646  1.00  0.00           H
    ATOM    558  HG2 PRO A 191      13.793  10.446  65.185  1.00  0.00           H
    ATOM    559  HG3 PRO A 191      12.925  11.295  66.467  1.00  0.00           H
    ATOM    560  HD2 PRO A 191      13.565  12.635  64.244  1.00  0.00           H
    ATOM    561  HD3 PRO A 191      13.903  13.427  65.785  1.00  0.00           H
    ATOM    562  N   GLN A 192      16.884  10.154  64.006  1.00 31.04           N
    ATOM    563  CA  GLN A 192      17.543   9.066  63.276  1.00 32.45           C
    ATOM    564  C   GLN A 192      18.832   9.511  62.550  1.00 31.03           C
    ATOM    565  O   GLN A 192      19.582   8.640  62.106  1.00 32.02           O
    ATOM    566  CB  GLN A 192      16.569   8.424  62.243  1.00 39.67           C
    ATOM    567  CG  GLN A 192      15.066   8.757  62.341  1.00 52.47           C
    ATOM    568  CD  GLN A 192      14.702  10.103  61.710  1.00 55.07           C
    ATOM    569  OE1 GLN A 192      14.199  10.981  62.401  1.00 55.08           O
    ATOM    570  NE2 GLN A 192      14.939  10.271  60.407  1.00 45.82           N
    ATOM    571  H   GLN A 192      16.189  10.670  63.485  1.00  0.00           H
    ATOM    572  HA  GLN A 192      17.827   8.294  63.994  1.00  0.00           H
    ATOM    573  HB2 GLN A 192      16.899   8.613  61.223  1.00  0.00           H
    ATOM    574  HB3 GLN A 192      16.653   7.348  62.352  1.00  0.00           H
    ATOM    575  HG2 GLN A 192      14.492   7.991  61.821  1.00  0.00           H
    ATOM    576  HG3 GLN A 192      14.727   8.715  63.377  1.00  0.00           H
    ATOM    577 HE21 GLN A 192      15.335   9.521  59.858  1.00  0.00           H
    ATOM    578 HE22 GLN A 192      14.678  11.129  59.938  1.00  0.00           H
    ATOM    579  N   HIE A 193      19.051  10.830  62.396  1.00 27.27           N
    ATOM    580  CA  HIE A 193      20.153  11.409  61.623  1.00 30.32           C
    ATOM    581  C   HIE A 193      21.497  11.320  62.354  1.00 37.19           C
    ATOM    582  O   HIE A 193      21.643  11.912  63.421  1.00 35.13           O
    ATOM    583  CB  HIE A 193      19.843  12.882  61.288  1.00 23.63           C
    ATOM    584  CG  HIE A 193      18.931  13.082  60.109  1.00 28.03           C
    ATOM    585  ND1 HIE A 193      19.091  14.151  59.225  1.00 30.87           N
    ATOM    586  CD2 HIE A 193      17.870  12.314  59.684  1.00 26.46           C
    ATOM    587  CE1 HIE A 193      18.135  14.003  58.323  1.00 31.09           C
    ATOM    588  NE2 HIE A 193      17.385  12.929  58.547  1.00 28.49           N
    ATOM    589  H   HIE A 193      18.398  11.487  62.801  1.00  0.00           H
    ATOM    590  HA  HIE A 193      20.248  10.862  60.683  1.00  0.00           H
    ATOM    591  HB2 HIE A 193      19.416  13.396  62.148  1.00  0.00           H
    ATOM    592  HB3 HIE A 193      20.767  13.417  61.062  1.00  0.00           H
    ATOM    593  HD2 HIE A 193      17.436  11.408  60.080  1.00  0.00           H
    ATOM    594  HE2 HIE A 193      16.577  12.624  58.007  1.00  0.00           H
    ATOM    595  HE1 HIE A 193      17.979  14.687  57.503  1.00  0.00           H
    ATOM    596  N   LEU A 194      22.469  10.633  61.734  1.00 39.00           N
    ATOM    597  CA  LEU A 194      23.853  10.545  62.200  1.00 35.20           C
    ATOM    598  C   LEU A 194      24.519  11.926  62.317  1.00 30.97           C
    ATOM    599  O   LEU A 194      25.010  12.261  63.390  1.00 31.98           O
    ATOM    600  CB  LEU A 194      24.664   9.607  61.272  1.00 39.22           C
    ATOM    601  CG  LEU A 194      26.157   9.458  61.653  1.00 39.48           C
    ATOM    602  CD1 LEU A 194      26.316   8.866  63.056  1.00 38.37           C
    ATOM    603  CD2 LEU A 194      26.937   8.657  60.601  1.00 38.03           C
    ATOM    604  H   LEU A 194      22.263  10.165  60.860  1.00  0.00           H
    ATOM    605  HA  LEU A 194      23.822  10.104  63.198  1.00  0.00           H
    ATOM    606  HB2 LEU A 194      24.196   8.621  61.255  1.00  0.00           H
    ATOM    607  HB3 LEU A 194      24.603   9.979  60.249  1.00  0.00           H
    ATOM    608  HG  LEU A 194      26.624  10.443  61.659  1.00  0.00           H
    ATOM    609 HD11 LEU A 194      27.301   8.425  63.203  1.00  0.00           H
    ATOM    610 HD12 LEU A 194      26.199   9.635  63.817  1.00  0.00           H
    ATOM    611 HD13 LEU A 194      25.560   8.102  63.230  1.00  0.00           H
    ATOM    612 HD21 LEU A 194      27.914   9.105  60.426  1.00  0.00           H
    ATOM    613 HD22 LEU A 194      27.095   7.623  60.904  1.00  0.00           H
    ATOM    614 HD23 LEU A 194      26.422   8.643  59.642  1.00  0.00           H
    ATOM    615  N   ARG A 196      24.759  16.052  62.279  1.00 30.66           N
    ATOM    616  CA  ARG A 196      24.120  17.311  62.647  1.00 27.36           C
    ATOM    617  C   ARG A 196      25.116  18.450  62.464  1.00 30.39           C
    ATOM    618  O   ARG A 196      26.318  18.212  62.532  1.00 32.47           O
    ATOM    619  CB  ARG A 196      23.644  17.222  64.112  1.00 23.08           C
    ATOM    620  CG  ARG A 196      22.203  16.705  64.190  1.00 26.50           C
    ATOM    621  CD  ARG A 196      21.952  15.610  65.221  1.00 28.93           C
    ATOM    622  NE  ARG A 196      21.905  16.134  66.593  1.00 30.92           N
    ATOM    623  CZ  ARG A 196      21.215  15.612  67.622  1.00 33.57           C
    ATOM    624  NH1 ARG A 196      20.465  14.509  67.508  1.00 35.85           N
    ATOM    625  NH2 ARG A 196      21.276  16.224  68.805  1.00 31.30           N1+
    ATOM    626  H   ARG A 196      25.715  15.952  62.596  1.00  0.00           H
    ATOM    627  HA  ARG A 196      23.287  17.513  61.977  1.00  0.00           H
    ATOM    628  HB2 ARG A 196      24.328  16.601  64.692  1.00  0.00           H
    ATOM    629  HB3 ARG A 196      23.673  18.201  64.595  1.00  0.00           H
    ATOM    630  HG2 ARG A 196      21.618  17.580  64.453  1.00  0.00           H
    ATOM    631  HG3 ARG A 196      21.814  16.370  63.229  1.00  0.00           H
    ATOM    632  HD2 ARG A 196      21.082  15.018  64.949  1.00  0.00           H
    ATOM    633  HD3 ARG A 196      22.796  14.935  65.171  1.00  0.00           H
    ATOM    634  HE  ARG A 196      22.478  16.948  66.770  1.00  0.00           H
    ATOM    635 HH11 ARG A 196      20.395  14.011  66.626  1.00  0.00           H
    ATOM    636 HH12 ARG A 196      19.941  14.161  68.299  1.00  0.00           H
    ATOM    637 HH21 ARG A 196      21.882  17.029  68.930  1.00  0.00           H
    ATOM    638 HH22 ARG A 196      20.776  15.864  69.611  1.00  0.00           H
    ATOM    639  N   TYR A 236      28.629  13.995  65.033  1.00 27.61           N
    ATOM    640  CA  TYR A 236      28.274  12.627  64.652  1.00 31.18           C
    ATOM    641  C   TYR A 236      27.684  11.923  65.880  1.00 35.85           C
    ATOM    642  O   TYR A 236      28.374  11.802  66.889  1.00 31.05           O
    ATOM    643  CB  TYR A 236      29.521  11.890  64.129  1.00 33.37           C
    ATOM    644  CG  TYR A 236      30.056  12.417  62.808  1.00 31.39           C
    ATOM    645  CD1 TYR A 236      30.874  13.565  62.776  1.00 30.05           C
    ATOM    646  CD2 TYR A 236      29.721  11.767  61.603  1.00 29.84           C
    ATOM    647  CE1 TYR A 236      31.335  14.075  61.548  1.00 26.94           C
    ATOM    648  CE2 TYR A 236      30.189  12.269  60.375  1.00 31.53           C
    ATOM    649  CZ  TYR A 236      30.991  13.428  60.346  1.00 29.35           C
    ATOM    650  OH  TYR A 236      31.434  13.927  59.156  1.00 28.16           O
    ATOM    651  H   TYR A 236      29.406  14.083  65.678  1.00  0.00           H
    ATOM    652  HA  TYR A 236      27.533  12.629  63.852  1.00  0.00           H
    ATOM    653  HB2 TYR A 236      30.322  11.899  64.870  1.00  0.00           H
    ATOM    654  HB3 TYR A 236      29.255  10.844  63.980  1.00  0.00           H
    ATOM    655  HD1 TYR A 236      31.142  14.068  63.693  1.00  0.00           H
    ATOM    656  HD2 TYR A 236      29.098  10.884  61.615  1.00  0.00           H
    ATOM    657  HE1 TYR A 236      31.949  14.963  61.536  1.00  0.00           H
    ATOM    658  HE2 TYR A 236      29.918  11.763  59.462  1.00  0.00           H
    ATOM    659  HH  TYR A 236      31.959  14.725  59.259  1.00  0.00           H
    ATOM    660  N   MET A 237      26.412  11.506  65.775  1.00 37.23           N
    ATOM    661  CA  MET A 237      25.569  11.064  66.891  1.00 36.31           C
    ATOM    662  C   MET A 237      25.492   9.538  67.104  1.00 37.04           C
    ATOM    663  O   MET A 237      24.614   9.079  67.834  1.00 32.26           O
    ATOM    664  CB  MET A 237      24.169  11.687  66.754  1.00 31.83           C
    ATOM    665  CG  MET A 237      24.139  13.212  66.612  1.00 26.60           C
    ATOM    666  SD  MET A 237      25.055  14.209  67.822  1.00 32.33           S
    ATOM    667  CE  MET A 237      24.120  13.812  69.322  1.00 31.61           C
    ATOM    668  H   MET A 237      25.925  11.655  64.899  1.00  0.00           H
    ATOM    669  HA  MET A 237      25.989  11.463  67.812  1.00  0.00           H
    ATOM    670  HB2 MET A 237      23.661  11.251  65.892  1.00  0.00           H
    ATOM    671  HB3 MET A 237      23.564  11.433  67.621  1.00  0.00           H
    ATOM    672  HG2 MET A 237      24.487  13.500  65.621  1.00  0.00           H
    ATOM    673  HG3 MET A 237      23.094  13.504  66.689  1.00  0.00           H
    ATOM    674  HE1 MET A 237      24.446  14.442  70.149  1.00  0.00           H
    ATOM    675  HE2 MET A 237      23.054  13.972  69.167  1.00  0.00           H
    ATOM    676  HE3 MET A 237      24.277  12.771  69.602  1.00  0.00           H
    ATOM    677  N   CY2 A 238      26.447   8.784  66.548  1.00 37.79           N
    ATOM    678  CA  CY2 A 238      26.676   7.373  66.859  1.00 37.31           C
    ATOM    679  C   CY2 A 238      28.177   7.099  66.754  1.00 39.10           C
    ATOM    680  O   CY2 A 238      28.832   7.659  65.876  1.00 42.65           O
    ATOM    681  CB  CY2 A 238      25.890   6.406  65.936  1.00 40.28           C
    ATOM    682  SG  CY2 A 238      24.344   5.726  66.635  1.00 43.09           S
    ATOM    683  H   CY2 A 238      27.154   9.220  65.973  1.00  0.00           H
    ATOM    684  HA  CY2 A 238      26.402   7.185  67.900  1.00  0.00           H
    ATOM    685  HB2 CY2 A 238      25.678   6.887  64.986  1.00  0.00           H
    ATOM    686  HB3 CY2 A 238      26.500   5.537  65.677  1.00  0.00           H
    ATOM    687  N   ASN A 239      28.661   6.166  67.590  1.00 40.01           N
    ATOM    688  CA  ASN A 239      29.963   5.521  67.416  1.00 39.03           C
    ATOM    689  C   ASN A 239      29.926   4.587  66.204  1.00 38.04           C
    ATOM    690  O   ASN A 239      28.896   3.962  65.948  1.00 38.52           O
    ATOM    691  CB  ASN A 239      30.334   4.707  68.674  1.00 38.33           C
    ATOM    692  CG  ASN A 239      30.645   5.579  69.889  1.00 39.39           C
    ATOM    693  OD1 ASN A 239      31.408   6.536  69.788  1.00 38.76           O
    ATOM    694  ND2 ASN A 239      30.087   5.229  71.048  1.00 42.42           N
    ATOM    695  H   ASN A 239      28.042   5.724  68.255  1.00  0.00           H
    ATOM    696  HA  ASN A 239      30.703   6.305  67.270  1.00  0.00           H
    ATOM    697  HB2 ASN A 239      29.559   3.978  68.914  1.00  0.00           H
    ATOM    698  HB3 ASN A 239      31.237   4.126  68.481  1.00  0.00           H
    ATOM    699 HD21 ASN A 239      29.495   4.411  71.107  1.00  0.00           H
    ATOM    700 HD22 ASN A 239      30.296   5.753  71.890  1.00  0.00           H
    ATOM    701  N   SER A 240      31.080   4.452  65.534  1.00 35.83           N
    ATOM    702  CA  SER A 240      31.318   3.473  64.470  1.00 35.37           C
    ATOM    703  C   SER A 240      30.997   2.036  64.908  1.00 42.61           C
    ATOM    704  O   SER A 240      30.469   1.273  64.104  1.00 48.12           O
    ATOM    705  CB  SER A 240      32.785   3.567  64.014  1.00 31.77           C
    ATOM    706  OG  SER A 240      32.975   4.737  63.256  1.00 33.41           O
    ATOM    707  H   SER A 240      31.886   4.998  65.812  1.00  0.00           H
    ATOM    708  HA  SER A 240      30.663   3.711  63.630  1.00  0.00           H
    ATOM    709  HB2 SER A 240      33.475   3.560  64.857  1.00  0.00           H
    ATOM    710  HB3 SER A 240      33.048   2.720  63.379  1.00  0.00           H
    ATOM    711  HG  SER A 240      32.615   4.578  62.375  1.00  0.00           H
    ATOM    712  N   SER A 241      31.283   1.733  66.186  1.00 43.46           N
    ATOM    713  CA  SER A 241      31.071   0.447  66.833  1.00 44.92           C
    ATOM    714  C   SER A 241      29.614   0.074  67.155  1.00 45.91           C
    ATOM    715  O   SER A 241      29.373  -1.118  67.333  1.00 45.82           O
    ATOM    716  CB  SER A 241      31.965   0.372  68.089  1.00 44.87           C
    ATOM    717  OG  SER A 241      31.529   1.253  69.109  1.00 43.14           O
    ATOM    718  H   SER A 241      31.710   2.438  66.770  1.00  0.00           H
    ATOM    719  HA  SER A 241      31.439  -0.282  66.122  1.00  0.00           H
    ATOM    720  HB2 SER A 241      31.969  -0.642  68.492  1.00  0.00           H
    ATOM    721  HB3 SER A 241      32.998   0.605  67.835  1.00  0.00           H
    ATOM    722  HG  SER A 241      30.736   0.888  69.517  1.00  0.00           H
    ATOM    723  N   CY2 A 242      28.701   1.064  67.247  1.00 43.97           N
    ATOM    724  CA  CY2 A 242      27.307   0.932  67.703  1.00 40.64           C
    ATOM    725  C   CY2 A 242      26.564  -0.263  67.083  1.00 41.82           C
    ATOM    726  O   CY2 A 242      26.277  -0.235  65.886  1.00 38.01           O
    ATOM    727  CB  CY2 A 242      26.491   2.221  67.453  1.00 36.91           C
    ATOM    728  SG  CY2 A 242      26.719   3.572  68.669  1.00 39.70           S
    ATOM    729  H   CY2 A 242      28.994   2.013  67.060  1.00  0.00           H
    ATOM    730  HA  CY2 A 242      27.348   0.768  68.782  1.00  0.00           H
    ATOM    731  HB2 CY2 A 242      26.656   2.577  66.434  1.00  0.00           H
    ATOM    732  HB3 CY2 A 242      25.430   1.983  67.494  1.00  0.00           H
    ATOM    733  N   MET A 243      26.256  -1.267  67.923  1.00 44.12           N
    ATOM    734  CA  MET A 243      25.517  -2.466  67.538  1.00 46.13           C
    ATOM    735  C   MET A 243      24.090  -2.122  67.079  1.00 46.30           C
    ATOM    736  O   MET A 243      23.344  -1.488  67.826  1.00 47.08           O
    ATOM    737  CB  MET A 243      25.519  -3.479  68.699  1.00 50.04           C
    ATOM    738  CG  MET A 243      24.843  -4.826  68.366  1.00 52.36           C
    ATOM    739  SD  MET A 243      24.444  -5.875  69.791  1.00 52.73           S
    ATOM    740  CE  MET A 243      23.167  -4.874  70.600  1.00 51.68           C
    ATOM    741  H   MET A 243      26.528  -1.201  68.894  1.00  0.00           H
    ATOM    742  HA  MET A 243      26.063  -2.908  66.708  1.00  0.00           H
    ATOM    743  HB2 MET A 243      26.550  -3.678  68.988  1.00  0.00           H
    ATOM    744  HB3 MET A 243      25.053  -3.025  69.572  1.00  0.00           H
    ATOM    745  HG2 MET A 243      23.910  -4.682  67.825  1.00  0.00           H
    ATOM    746  HG3 MET A 243      25.490  -5.397  67.699  1.00  0.00           H
    ATOM    747  HE1 MET A 243      22.732  -5.431  71.429  1.00  0.00           H
    ATOM    748  HE2 MET A 243      23.582  -3.949  70.999  1.00  0.00           H
    ATOM    749  HE3 MET A 243      22.373  -4.627  69.896  1.00  0.00           H
    ATOM    750  N   GLY A 244      23.754  -2.556  65.855  1.00 43.40           N
    ATOM    751  CA  GLY A 244      22.477  -2.291  65.194  1.00 39.49           C
    ATOM    752  C   GLY A 244      22.498  -0.980  64.386  1.00 41.04           C
    ATOM    753  O   GLY A 244      21.570  -0.748  63.614  1.00 43.17           O
    ATOM    754  H   GLY A 244      24.432  -3.086  65.323  1.00  0.00           H
    ATOM    755  HA2 GLY A 244      22.264  -3.117  64.515  1.00  0.00           H
    ATOM    756  HA3 GLY A 244      21.661  -2.259  65.919  1.00  0.00           H
    ATOM    757  N   GLY A 245      23.541  -0.145  64.534  1.00 42.01           N
    ATOM    758  CA  GLY A 245      23.757   1.094  63.791  1.00 41.37           C
    ATOM    759  C   GLY A 245      24.800   0.802  62.714  1.00 42.95           C
    ATOM    760  O   GLY A 245      24.606  -0.091  61.885  1.00 42.75           O
    ATOM    761  H   GLY A 245      24.274  -0.404  65.181  1.00  0.00           H
    ATOM    762  HA2 GLY A 245      22.844   1.478  63.335  1.00  0.00           H
    ATOM    763  HA3 GLY A 245      24.124   1.858  64.474  1.00  0.00           H
    ATOM    764  N   MET A 246      25.911   1.558  62.740  1.00 42.35           N
    ATOM    765  CA  MET A 246      27.095   1.331  61.907  1.00 42.54           C
    ATOM    766  C   MET A 246      27.732  -0.050  62.110  1.00 42.19           C
    ATOM    767  O   MET A 246      28.180  -0.628  61.123  1.00 39.56           O
    ATOM    768  CB  MET A 246      28.108   2.480  62.067  1.00 40.89           C
    ATOM    769  CG  MET A 246      27.702   3.732  61.271  1.00 37.04           C
    ATOM    770  SD  MET A 246      28.827   5.140  61.443  1.00 34.36           S
    ATOM    771  CE  MET A 246      28.258   5.782  63.032  1.00 29.18           C
    ATOM    772  H   MET A 246      25.977   2.315  63.409  1.00  0.00           H
    ATOM    773  HA  MET A 246      26.770   1.334  60.874  1.00  0.00           H
    ATOM    774  HB2 MET A 246      28.235   2.723  63.123  1.00  0.00           H
    ATOM    775  HB3 MET A 246      29.090   2.163  61.712  1.00  0.00           H
    ATOM    776  HG2 MET A 246      27.659   3.485  60.210  1.00  0.00           H
    ATOM    777  HG3 MET A 246      26.697   4.053  61.544  1.00  0.00           H
    ATOM    778  HE1 MET A 246      28.888   6.611  63.353  1.00  0.00           H
    ATOM    779  HE2 MET A 246      27.227   6.123  62.953  1.00  0.00           H
    ATOM    780  HE3 MET A 246      28.294   5.003  63.785  1.00  0.00           H
    ATOM    781  N   ASN A 247      27.694  -0.579  63.346  1.00 41.67           N
    ATOM    782  CA  ASN A 247      28.046  -1.956  63.715  1.00 40.76           C
    ATOM    783  C   ASN A 247      29.460  -2.346  63.235  1.00 42.14           C
    ATOM    784  O   ASN A 247      29.645  -3.380  62.597  1.00 41.93           O
    ATOM    785  CB  ASN A 247      26.941  -2.929  63.227  1.00 40.53           C
    ATOM    786  CG  ASN A 247      26.833  -4.210  64.063  1.00 44.21           C
    ATOM    787  OD1 ASN A 247      25.843  -4.429  64.753  1.00 46.61           O
    ATOM    788  ND2 ASN A 247      27.831  -5.087  63.969  1.00 44.02           N
    ATOM    789  H   ASN A 247      27.294  -0.031  64.096  1.00  0.00           H
    ATOM    790  HA  ASN A 247      28.121  -2.024  64.801  1.00  0.00           H
    ATOM    791  HB2 ASN A 247      25.973  -2.432  63.323  1.00  0.00           H
    ATOM    792  HB3 ASN A 247      27.042  -3.162  62.166  1.00  0.00           H
    ATOM    793 HD21 ASN A 247      28.639  -4.879  63.396  1.00  0.00           H
    ATOM    794 HD22 ASN A 247      27.782  -5.978  64.448  1.00  0.00           H
    ATOM    795  N   ARG A 248      30.429  -1.473  63.535  1.00 44.63           N
    ATOM    796  CA  ARG A 248      31.853  -1.579  63.210  1.00 45.75           C
    ATOM    797  C   ARG A 248      32.194  -1.522  61.706  1.00 49.50           C
    ATOM    798  O   ARG A 248      33.296  -1.926  61.336  1.00 56.01           O
    ATOM    799  CB  ARG A 248      32.515  -2.798  63.892  1.00 43.00           C
    ATOM    800  CG  ARG A 248      31.985  -3.165  65.287  1.00 48.03           C
    ATOM    801  CD  ARG A 248      32.966  -4.005  66.083  1.00 48.02           C
    ATOM    802  NE  ARG A 248      33.414  -5.178  65.341  1.00 50.30           N
    ATOM    803  CZ  ARG A 248      34.679  -5.456  64.997  1.00 51.17           C
    ATOM    804  NH1 ARG A 248      35.728  -4.705  65.367  1.00 49.32           N
    ATOM    805  NH2 ARG A 248      34.878  -6.529  64.239  1.00 50.88           N1+
    ATOM    806  H   ARG A 248      30.173  -0.619  64.016  1.00  0.00           H
    ATOM    807  HA  ARG A 248      32.315  -0.687  63.635  1.00  0.00           H
    ATOM    808  HB2 ARG A 248      32.409  -3.680  63.263  1.00  0.00           H
    ATOM    809  HB3 ARG A 248      33.588  -2.608  63.954  1.00  0.00           H
    ATOM    810  HG2 ARG A 248      31.937  -2.238  65.842  1.00  0.00           H
    ATOM    811  HG3 ARG A 248      30.976  -3.576  65.301  1.00  0.00           H
    ATOM    812  HD2 ARG A 248      33.739  -3.402  66.532  1.00  0.00           H
    ATOM    813  HD3 ARG A 248      32.421  -4.458  66.898  1.00  0.00           H
    ATOM    814  HE  ARG A 248      32.680  -5.793  65.007  1.00  0.00           H
    ATOM    815 HH11 ARG A 248      35.605  -3.852  65.900  1.00  0.00           H
    ATOM    816 HH12 ARG A 248      36.668  -4.974  65.110  1.00  0.00           H
    ATOM    817 HH21 ARG A 248      34.097  -7.082  63.913  1.00  0.00           H
    ATOM    818 HH22 ARG A 248      35.807  -6.781  63.913  1.00  0.00           H
    ATOM    819  N   ARG A 249      31.271  -1.029  60.861  1.00 44.78           N
    ATOM    820  CA  ARG A 249      31.476  -0.915  59.416  1.00 38.54           C
    ATOM    821  C   ARG A 249      31.845   0.542  59.076  1.00 41.84           C
    ATOM    822  O   ARG A 249      31.093   1.438  59.468  1.00 43.54           O
    ATOM    823  CB  ARG A 249      30.192  -1.313  58.671  1.00 33.61           C
    ATOM    824  CG  ARG A 249      29.751  -2.762  58.936  1.00 33.85           C
    ATOM    825  CD  ARG A 249      28.406  -3.088  58.292  1.00 32.65           C
    ATOM    826  NE  ARG A 249      27.301  -2.391  58.955  1.00 34.35           N
    ATOM    827  CZ  ARG A 249      26.053  -2.307  58.475  1.00 33.48           C
    ATOM    828  NH1 ARG A 249      25.711  -2.926  57.338  1.00 32.81           N
    ATOM    829  NH2 ARG A 249      25.138  -1.591  59.136  1.00 33.25           N1+
    ATOM    830  H   ARG A 249      30.379  -0.711  61.215  1.00  0.00           H
    ATOM    831  HA  ARG A 249      32.241  -1.631  59.121  1.00  0.00           H
    ATOM    832  HB2 ARG A 249      29.387  -0.620  58.915  1.00  0.00           H
    ATOM    833  HB3 ARG A 249      30.358  -1.201  57.600  1.00  0.00           H
    ATOM    834  HG2 ARG A 249      30.505  -3.493  58.650  1.00  0.00           H
    ATOM    835  HG3 ARG A 249      29.619  -2.870  60.011  1.00  0.00           H
    ATOM    836  HD2 ARG A 249      28.410  -2.642  57.300  1.00  0.00           H
    ATOM    837  HD3 ARG A 249      28.244  -4.157  58.157  1.00  0.00           H
    ATOM    838  HE  ARG A 249      27.527  -1.908  59.818  1.00  0.00           H
    ATOM    839 HH11 ARG A 249      26.390  -3.475  56.830  1.00  0.00           H
    ATOM    840 HH12 ARG A 249      24.754  -2.863  56.991  1.00  0.00           H
    ATOM    841 HH21 ARG A 249      25.370  -1.103  59.990  1.00  0.00           H
    ATOM    842 HH22 ARG A 249      24.191  -1.518  58.768  1.00  0.00           H
    ATOM    843  N   PRO A 250      32.981   0.763  58.370  1.00 40.77           N
    ATOM    844  CA  PRO A 250      33.456   2.113  58.006  1.00 36.47           C
    ATOM    845  C   PRO A 250      32.482   2.883  57.099  1.00 31.50           C
    ATOM    846  O   PRO A 250      31.841   2.279  56.240  1.00 34.18           O
    ATOM    847  CB  PRO A 250      34.800   1.853  57.303  1.00 34.92           C
    ATOM    848  CG  PRO A 250      34.690   0.442  56.758  1.00 35.71           C
    ATOM    849  CD  PRO A 250      33.876  -0.262  57.830  1.00 38.17           C
    ATOM    850  HA  PRO A 250      33.630   2.692  58.913  1.00  0.00           H
    ATOM    851  HB2 PRO A 250      35.041   2.583  56.530  1.00  0.00           H
    ATOM    852  HB3 PRO A 250      35.604   1.890  58.038  1.00  0.00           H
    ATOM    853  HG2 PRO A 250      34.125   0.449  55.826  1.00  0.00           H
    ATOM    854  HG3 PRO A 250      35.653  -0.030  56.564  1.00  0.00           H
    ATOM    855  HD2 PRO A 250      33.352  -1.108  57.393  1.00  0.00           H
    ATOM    856  HD3 PRO A 250      34.519  -0.627  58.633  1.00  0.00           H
    ATOM    857  N   ILE A 251      32.395   4.201  57.322  1.00 29.17           N
    ATOM    858  CA  ILE A 251      31.568   5.111  56.535  1.00 29.44           C
    ATOM    859  C   ILE A 251      32.420   6.243  55.939  1.00 33.21           C
    ATOM    860  O   ILE A 251      33.483   6.567  56.464  1.00 36.27           O
    ATOM    861  CB  ILE A 251      30.434   5.753  57.388  1.00 29.03           C
    ATOM    862  CG1 ILE A 251      30.909   6.594  58.600  1.00 20.86           C
    ATOM    863  CG2 ILE A 251      29.377   4.714  57.803  1.00 31.06           C
    ATOM    864  CD1 ILE A 251      29.987   7.778  58.907  1.00 19.70           C
    ATOM    865  H   ILE A 251      32.970   4.632  58.031  1.00  0.00           H
    ATOM    866  HA  ILE A 251      31.121   4.579  55.696  1.00  0.00           H
    ATOM    867  HB  ILE A 251      29.906   6.443  56.727  1.00  0.00           H
    ATOM    868 HG12 ILE A 251      30.999   5.966  59.483  1.00  0.00           H
    ATOM    869 HG13 ILE A 251      31.905   7.000  58.445  1.00  0.00           H
    ATOM    870 HG21 ILE A 251      28.540   5.182  58.319  1.00  0.00           H
    ATOM    871 HG22 ILE A 251      28.970   4.205  56.932  1.00  0.00           H
    ATOM    872 HG23 ILE A 251      29.792   3.957  58.468  1.00  0.00           H
    ATOM    873 HD11 ILE A 251      30.211   8.200  59.888  1.00  0.00           H
    ATOM    874 HD12 ILE A 251      30.107   8.572  58.168  1.00  0.00           H
    ATOM    875 HD13 ILE A 251      28.940   7.475  58.905  1.00  0.00           H
    ATOM    876  N   GLU A 271      36.701   9.710  54.722  1.00 35.56           N
    ATOM    877  CA  GLU A 271      36.288   8.642  55.633  1.00 33.32           C
    ATOM    878  C   GLU A 271      36.087   9.196  57.052  1.00 34.79           C
    ATOM    879  O   GLU A 271      36.756  10.156  57.427  1.00 32.65           O
    ATOM    880  CB  GLU A 271      37.289   7.471  55.565  1.00 30.67           C
    ATOM    881  CG  GLU A 271      36.860   6.238  56.392  1.00 34.80           C
    ATOM    882  CD  GLU A 271      37.777   5.027  56.266  1.00 42.49           C
    ATOM    883  OE1 GLU A 271      38.580   4.979  55.309  1.00 47.43           O
    ATOM    884  OE2 GLU A 271      37.644   4.142  57.139  1.00 41.08           O1-
    ATOM    885  H   GLU A 271      37.686   9.747  54.490  1.00  0.00           H
    ATOM    886  HA  GLU A 271      35.332   8.258  55.293  1.00  0.00           H
    ATOM    887  HB2 GLU A 271      37.389   7.175  54.522  1.00  0.00           H
    ATOM    888  HB3 GLU A 271      38.278   7.806  55.879  1.00  0.00           H
    ATOM    889  HG2 GLU A 271      36.786   6.486  57.451  1.00  0.00           H
    ATOM    890  HG3 GLU A 271      35.874   5.911  56.072  1.00  0.00           H
    ATOM    891  N   VAL A 272      35.161   8.598  57.810  1.00 33.81           N
    ATOM    892  CA  VAL A 272      34.935   8.967  59.200  1.00 30.26           C
    ATOM    893  C   VAL A 272      34.945   7.705  60.065  1.00 32.09           C
    ATOM    894  O   VAL A 272      34.334   6.695  59.710  1.00 33.71           O
    ATOM    895  CB  VAL A 272      33.597   9.725  59.416  1.00 28.67           C
    ATOM    896  CG1 VAL A 272      33.369  10.152  60.884  1.00 27.39           C
    ATOM    897  CG2 VAL A 272      33.527  10.968  58.521  1.00 30.60           C
    ATOM    898  H   VAL A 272      34.621   7.821  57.449  1.00  0.00           H
    ATOM    899  HA  VAL A 272      35.732   9.618  59.553  1.00  0.00           H
    ATOM    900  HB  VAL A 272      32.767   9.077  59.132  1.00  0.00           H
    ATOM    901 HG11 VAL A 272      32.470  10.758  60.985  1.00  0.00           H
    ATOM    902 HG12 VAL A 272      33.238   9.297  61.546  1.00  0.00           H
    ATOM    903 HG13 VAL A 272      34.204  10.744  61.261  1.00  0.00           H
    ATOM    904 HG21 VAL A 272      32.646  11.562  58.747  1.00  0.00           H
    ATOM    905 HG22 VAL A 272      34.402  11.594  58.678  1.00  0.00           H
    ATOM    906 HG23 VAL A 272      33.483  10.708  57.463  1.00  0.00           H
    ATOM    907  N   ARG A 273      35.624   7.832  61.210  1.00 33.57           N
    ATOM    908  CA  ARG A 273      35.560   6.917  62.332  1.00 35.96           C
    ATOM    909  C   ARG A 273      35.152   7.741  63.554  1.00 36.21           C
    ATOM    910  O   ARG A 273      35.865   8.676  63.922  1.00 35.68           O
    ATOM    911  CB  ARG A 273      36.921   6.218  62.506  1.00 39.10           C
    ATOM    912  CG  ARG A 273      37.024   5.354  63.773  1.00 41.26           C
    ATOM    913  CD  ARG A 273      38.382   4.662  63.909  1.00 47.30           C
    ATOM    914  NE  ARG A 273      38.403   3.732  65.039  1.00 57.43           N
    ATOM    915  CZ  ARG A 273      37.875   2.500  65.066  1.00 64.82           C
    ATOM    916  NH1 ARG A 273      37.275   1.969  63.991  1.00 65.41           N
    ATOM    917  NH2 ARG A 273      37.957   1.792  66.197  1.00 67.60           N1+
    ATOM    918  H   ARG A 273      36.127   8.693  61.388  1.00  0.00           H
    ATOM    919  HA  ARG A 273      34.806   6.153  62.147  1.00  0.00           H
    ATOM    920  HB2 ARG A 273      37.086   5.582  61.639  1.00  0.00           H
    ATOM    921  HB3 ARG A 273      37.726   6.955  62.512  1.00  0.00           H
    ATOM    922  HG2 ARG A 273      36.934   6.033  64.619  1.00  0.00           H
    ATOM    923  HG3 ARG A 273      36.202   4.642  63.867  1.00  0.00           H
    ATOM    924  HD2 ARG A 273      38.770   4.251  62.976  1.00  0.00           H
    ATOM    925  HD3 ARG A 273      39.092   5.420  64.227  1.00  0.00           H
    ATOM    926  HE  ARG A 273      38.847   4.067  65.891  1.00  0.00           H
    ATOM    927 HH11 ARG A 273      37.218   2.493  63.131  1.00  0.00           H
    ATOM    928 HH12 ARG A 273      36.898   1.033  64.031  1.00  0.00           H
    ATOM    929 HH21 ARG A 273      38.419   2.193  67.004  1.00  0.00           H
    ATOM    930 HH22 ARG A 273      37.581   0.854  66.256  1.00  0.00           H
    ATOM    931  N   VAL A 274      34.029   7.357  64.172  1.00 34.93           N
    ATOM    932  CA  VAL A 274      33.573   7.942  65.423  1.00 34.73           C
    ATOM    933  C   VAL A 274      33.957   6.961  66.544  1.00 35.87           C
    ATOM    934  O   VAL A 274      33.472   5.830  66.562  1.00 37.53           O
    ATOM    935  CB  VAL A 274      32.048   8.198  65.454  1.00 35.50           C
    ATOM    936  CG1 VAL A 274      31.645   8.981  66.724  1.00 34.42           C
    ATOM    937  CG2 VAL A 274      31.557   8.944  64.203  1.00 35.43           C
    ATOM    938  H   VAL A 274      33.507   6.560  63.829  1.00  0.00           H
    ATOM    939  HA  VAL A 274      34.057   8.902  65.577  1.00  0.00           H
    ATOM    940  HB  VAL A 274      31.526   7.243  65.449  1.00  0.00           H
    ATOM    941 HG11 VAL A 274      30.596   9.271  66.709  1.00  0.00           H
    ATOM    942 HG12 VAL A 274      31.809   8.405  67.634  1.00  0.00           H
    ATOM    943 HG13 VAL A 274      32.223   9.902  66.811  1.00  0.00           H
    ATOM    944 HG21 VAL A 274      30.476   9.059  64.236  1.00  0.00           H
    ATOM    945 HG22 VAL A 274      32.004   9.934  64.118  1.00  0.00           H
    ATOM    946 HG23 VAL A 274      31.776   8.395  63.288  1.00  0.00           H
    ATOM    947  N   CYS A 275      34.875   7.398  67.413  1.00 35.59           N
    ATOM    948  CA  CYS A 275      35.577   6.556  68.377  1.00 34.89           C
    ATOM    949  C   CYS A 275      35.966   7.383  69.610  1.00 37.40           C
    ATOM    950  O   CYS A 275      36.163   8.595  69.511  1.00 38.09           O
    ATOM    951  CB  CYS A 275      36.785   5.851  67.715  1.00 34.40           C
    ATOM    952  SG  CYS A 275      38.067   7.022  67.161  1.00 34.66           S
    ATOM    953  H   CYS A 275      35.207   8.353  67.345  1.00  0.00           H
    ATOM    954  HA  CYS A 275      34.886   5.785  68.724  1.00  0.00           H
    ATOM    955  HB2 CYS A 275      37.242   5.153  68.415  1.00  0.00           H
    ATOM    956  HB3 CYS A 275      36.458   5.254  66.864  1.00  0.00           H
    ATOM    957  HG  CYS A 275      38.453   7.399  68.383  1.00  0.00           H
    ATOM    958  N   ALA A 276      36.054   6.705  70.765  1.00 39.31           N
    ATOM    959  CA  ALA A 276      36.360   7.312  72.064  1.00 42.07           C
    ATOM    960  C   ALA A 276      37.778   7.900  72.172  1.00 42.46           C
    ATOM    961  O   ALA A 276      37.979   8.810  72.978  1.00 41.94           O
    ATOM    962  CB  ALA A 276      36.127   6.271  73.166  1.00 43.64           C
    ATOM    963  H   ALA A 276      35.879   5.710  70.774  1.00  0.00           H
    ATOM    964  HA  ALA A 276      35.656   8.132  72.220  1.00  0.00           H
    ATOM    965  HB1 ALA A 276      36.320   6.689  74.155  1.00  0.00           H
    ATOM    966  HB2 ALA A 276      35.095   5.918  73.159  1.00  0.00           H
    ATOM    967  HB3 ALA A 276      36.778   5.404  73.039  1.00  0.00           H
    ATOM    968  N   CYS A 277      38.717   7.393  71.354  1.00 42.36           N
    ATOM    969  CA  CYS A 277      40.119   7.797  71.336  1.00 45.89           C
    ATOM    970  C   CYS A 277      40.542   8.179  69.900  1.00 42.25           C
    ATOM    971  O   CYS A 277      41.175   7.358  69.236  1.00 41.45           O
    ATOM    972  CB  CYS A 277      41.023   6.676  71.894  1.00 54.92           C
    ATOM    973  SG  CYS A 277      40.723   6.374  73.657  1.00 65.82           S
    ATOM    974  H   CYS A 277      38.472   6.629  70.737  1.00  0.00           H
    ATOM    975  HA  CYS A 277      40.268   8.678  71.960  1.00  0.00           H
    ATOM    976  HB2 CYS A 277      40.898   5.743  71.345  1.00  0.00           H
    ATOM    977  HB3 CYS A 277      42.061   6.969  71.788  1.00  0.00           H
    ATOM    978  HG  CYS A 277      39.481   5.905  73.504  1.00  0.00           H
    ATOM    979  N   PRO A 278      40.204   9.406  69.429  1.00 40.01           N
    ATOM    980  CA  PRO A 278      40.566   9.876  68.075  1.00 39.59           C
    ATOM    981  C   PRO A 278      42.061   9.827  67.720  1.00 37.97           C
    ATOM    982  O   PRO A 278      42.408   9.339  66.646  1.00 35.91           O
    ATOM    983  CB  PRO A 278      40.050  11.323  68.027  1.00 36.63           C
    ATOM    984  CG  PRO A 278      38.921  11.352  69.027  1.00 34.63           C
    ATOM    985  CD  PRO A 278      39.398  10.412  70.122  1.00 36.20           C
    ATOM    986  HA  PRO A 278      40.002   9.262  67.377  1.00  0.00           H
    ATOM    987  HB2 PRO A 278      40.814  12.031  68.348  1.00  0.00           H
    ATOM    988  HB3 PRO A 278      39.729  11.625  67.034  1.00  0.00           H
    ATOM    989  HG2 PRO A 278      38.695  12.355  69.386  1.00  0.00           H
    ATOM    990  HG3 PRO A 278      38.026  10.937  68.567  1.00  0.00           H
    ATOM    991  HD2 PRO A 278      40.016  10.941  70.846  1.00  0.00           H
    ATOM    992  HD3 PRO A 278      38.537  10.003  70.642  1.00  0.00           H
    ATOM    993  N   GLY A 279      42.900  10.332  68.638  1.00 33.24           N
    ATOM    994  CA  GLY A 279      44.343  10.453  68.475  1.00 33.33           C
    ATOM    995  C   GLY A 279      45.025   9.084  68.480  1.00 39.56           C
    ATOM    996  O   GLY A 279      45.862   8.846  67.613  1.00 39.85           O
    ATOM    997  H   GLY A 279      42.517  10.715  69.494  1.00  0.00           H
    ATOM    998  HA2 GLY A 279      44.570  10.972  67.544  1.00  0.00           H
    ATOM    999  HA3 GLY A 279      44.746  11.058  69.288  1.00  0.00           H
    ATOM   1000  N   ARG A 280      44.653   8.175  69.406  1.00 43.03           N
    ATOM   1001  CA  ARG A 280      45.192   6.808  69.457  1.00 43.69           C
    ATOM   1002  C   ARG A 280      44.878   6.019  68.182  1.00 41.71           C
    ATOM   1003  O   ARG A 280      45.804   5.469  67.594  1.00 41.00           O
    ATOM   1004  CB  ARG A 280      44.713   6.047  70.715  1.00 46.94           C
    ATOM   1005  CG  ARG A 280      45.209   4.585  70.795  1.00 52.50           C
    ATOM   1006  CD  ARG A 280      44.915   3.863  72.120  1.00 59.69           C
    ATOM   1007  NE  ARG A 280      43.485   3.789  72.460  1.00 65.21           N
    ATOM   1008  CZ  ARG A 280      42.562   3.004  71.874  1.00 70.53           C
    ATOM   1009  NH1 ARG A 280      42.861   2.230  70.821  1.00 73.39           N
    ATOM   1010  NH2 ARG A 280      41.318   2.977  72.367  1.00 72.00           N1+
    ATOM   1011  H   ARG A 280      43.938   8.405  70.078  1.00  0.00           H
    ATOM   1012  HA  ARG A 280      46.277   6.900  69.527  1.00  0.00           H
    ATOM   1013  HB2 ARG A 280      45.031   6.589  71.605  1.00  0.00           H
    ATOM   1014  HB3 ARG A 280      43.627   6.026  70.717  1.00  0.00           H
    ATOM   1015  HG2 ARG A 280      44.856   3.978  69.962  1.00  0.00           H
    ATOM   1016  HG3 ARG A 280      46.294   4.612  70.692  1.00  0.00           H
    ATOM   1017  HD2 ARG A 280      45.273   2.835  72.059  1.00  0.00           H
    ATOM   1018  HD3 ARG A 280      45.477   4.320  72.931  1.00  0.00           H
    ATOM   1019  HE  ARG A 280      43.192   4.363  73.237  1.00  0.00           H
    ATOM   1020 HH11 ARG A 280      43.790   2.246  70.429  1.00  0.00           H
    ATOM   1021 HH12 ARG A 280      42.163   1.631  70.401  1.00  0.00           H
    ATOM   1022 HH21 ARG A 280      41.069   3.539  73.168  1.00  0.00           H
    ATOM   1023 HH22 ARG A 280      40.624   2.368  71.957  1.00  0.00           H
    ATOM   1024  N   ASP A 281      43.596   5.998  67.781  1.00 40.42           N
    ATOM   1025  CA  ASP A 281      43.117   5.271  66.605  1.00 39.48           C
    ATOM   1026  C   ASP A 281      43.699   5.810  65.289  1.00 41.59           C
    ATOM   1027  O   ASP A 281      43.996   4.998  64.417  1.00 41.71           O
    ATOM   1028  CB  ASP A 281      41.578   5.120  66.518  1.00 42.44           C
    ATOM   1029  CG  ASP A 281      40.894   4.432  67.712  1.00 51.39           C
    ATOM   1030  OD1 ASP A 281      41.478   4.374  68.816  1.00 55.89           O
    ATOM   1031  OD2 ASP A 281      39.738   4.000  67.511  1.00 54.20           O1-
    ATOM   1032  H   ASP A 281      42.885   6.461  68.332  1.00  0.00           H
    ATOM   1033  HA  ASP A 281      43.511   4.257  66.706  1.00  0.00           H
    ATOM   1034  HB2 ASP A 281      41.128   6.106  66.400  1.00  0.00           H
    ATOM   1035  HB3 ASP A 281      41.342   4.540  65.623  1.00  0.00           H
    ATOM   1036  N   ARG A 282      43.924   7.135  65.183  1.00 37.97           N
    ATOM   1037  CA  ARG A 282      44.669   7.731  64.070  1.00 34.93           C
    ATOM   1038  C   ARG A 282      46.125   7.244  64.026  1.00 37.91           C
    ATOM   1039  O   ARG A 282      46.569   6.803  62.968  1.00 38.71           O
    ATOM   1040  CB  ARG A 282      44.587   9.277  64.087  1.00 34.87           C
    ATOM   1041  CG  ARG A 282      45.426   9.917  62.962  1.00 33.77           C
    ATOM   1042  CD  ARG A 282      45.238  11.422  62.759  1.00 30.17           C
    ATOM   1043  NE  ARG A 282      46.328  11.920  61.911  1.00 33.21           N
    ATOM   1044  CZ  ARG A 282      46.858  13.152  61.901  1.00 35.19           C
    ATOM   1045  NH1 ARG A 282      46.422  14.114  62.722  1.00 35.94           N
    ATOM   1046  NH2 ARG A 282      47.856  13.421  61.052  1.00 33.40           N1+
    ATOM   1047  H   ARG A 282      43.636   7.759  65.925  1.00  0.00           H
    ATOM   1048  HA  ARG A 282      44.195   7.391  63.150  1.00  0.00           H
    ATOM   1049  HB2 ARG A 282      43.549   9.591  64.003  1.00  0.00           H
    ATOM   1050  HB3 ARG A 282      44.937   9.655  65.049  1.00  0.00           H
    ATOM   1051  HG2 ARG A 282      46.489   9.690  63.049  1.00  0.00           H
    ATOM   1052  HG3 ARG A 282      45.093   9.433  62.045  1.00  0.00           H
    ATOM   1053  HD2 ARG A 282      44.326  11.613  62.193  1.00  0.00           H
    ATOM   1054  HD3 ARG A 282      45.131  11.946  63.706  1.00  0.00           H
    ATOM   1055  HE  ARG A 282      46.720  11.243  61.261  1.00  0.00           H
    ATOM   1056 HH11 ARG A 282      45.623  13.956  63.323  1.00  0.00           H
    ATOM   1057 HH12 ARG A 282      46.819  15.044  62.664  1.00  0.00           H
    ATOM   1058 HH21 ARG A 282      48.216  12.696  60.447  1.00  0.00           H
    ATOM   1059 HH22 ARG A 282      48.253  14.351  61.018  1.00  0.00           H
    ATOM   1060  N   ARG A 283      46.828   7.340  65.166  1.00 35.97           N
    ATOM   1061  CA  ARG A 283      48.242   6.991  65.296  1.00 33.74           C
    ATOM   1062  C   ARG A 283      48.511   5.496  65.010  1.00 34.91           C
    ATOM   1063  O   ARG A 283      49.469   5.189  64.305  1.00 39.25           O
    ATOM   1064  CB  ARG A 283      48.748   7.442  66.687  1.00 35.57           C
    ATOM   1065  CG  ARG A 283      50.276   7.593  66.776  1.00 37.03           C
    ATOM   1066  CD  ARG A 283      50.812   7.845  68.196  1.00 33.67           C
    ATOM   1067  NE  ARG A 283      50.447   9.157  68.758  1.00 36.77           N
    ATOM   1068  CZ  ARG A 283      50.882   9.623  69.944  1.00 39.25           C
    ATOM   1069  NH1 ARG A 283      51.720   8.907  70.708  1.00 38.87           N
    ATOM   1070  NH2 ARG A 283      50.480  10.823  70.377  1.00 42.63           N1+
    ATOM   1071  H   ARG A 283      46.381   7.706  65.996  1.00  0.00           H
    ATOM   1072  HA  ARG A 283      48.780   7.566  64.539  1.00  0.00           H
    ATOM   1073  HB2 ARG A 283      48.318   8.410  66.937  1.00  0.00           H
    ATOM   1074  HB3 ARG A 283      48.389   6.747  67.448  1.00  0.00           H
    ATOM   1075  HG2 ARG A 283      50.781   6.714  66.374  1.00  0.00           H
    ATOM   1076  HG3 ARG A 283      50.568   8.421  66.131  1.00  0.00           H
    ATOM   1077  HD2 ARG A 283      50.385   7.102  68.869  1.00  0.00           H
    ATOM   1078  HD3 ARG A 283      51.891   7.690  68.219  1.00  0.00           H
    ATOM   1079  HE  ARG A 283      49.837   9.741  68.199  1.00  0.00           H
    ATOM   1080 HH11 ARG A 283      52.030   7.998  70.397  1.00  0.00           H
    ATOM   1081 HH12 ARG A 283      52.040   9.261  71.598  1.00  0.00           H
    ATOM   1082 HH21 ARG A 283      49.822  11.360  69.828  1.00  0.00           H
    ATOM   1083 HH22 ARG A 283      50.807  11.189  71.260  1.00  0.00           H
    ATOM   1084  N   THR A 284      47.623   4.614  65.506  1.00 37.81           N
    ATOM   1085  CA  THR A 284      47.657   3.161  65.315  1.00 41.86           C
    ATOM   1086  C   THR A 284      47.250   2.705  63.895  1.00 41.88           C
    ATOM   1087  O   THR A 284      47.747   1.672  63.449  1.00 47.58           O
    ATOM   1088  CB  THR A 284      46.735   2.440  66.340  1.00 46.24           C
    ATOM   1089  OG1 THR A 284      47.144   2.771  67.653  1.00 49.84           O
    ATOM   1090  CG2 THR A 284      46.691   0.901  66.251  1.00 43.44           C
    ATOM   1091  H   THR A 284      46.870   4.953  66.092  1.00  0.00           H
    ATOM   1092  HA  THR A 284      48.682   2.827  65.487  1.00  0.00           H
    ATOM   1093  HB  THR A 284      45.716   2.816  66.227  1.00  0.00           H
    ATOM   1094 HG21 THR A 284      46.140   0.473  67.088  1.00  0.00           H
    ATOM   1095 HG22 THR A 284      46.192   0.560  65.343  1.00  0.00           H
    ATOM   1096 HG23 THR A 284      47.696   0.477  66.263  1.00  0.00           H
    ATOM   1097  HG1 THR A 284      46.738   2.150  68.262  1.00  0.00           H
    ATOM   1098  N   GLU A 285      46.389   3.461  63.190  1.00 40.55           N
    ATOM   1099  CA  GLU A 285      46.049   3.174  61.790  1.00 45.15           C
    ATOM   1100  C   GLU A 285      47.091   3.719  60.793  1.00 48.65           C
    ATOM   1101  O   GLU A 285      47.264   3.114  59.735  1.00 52.61           O
    ATOM   1102  CB  GLU A 285      44.629   3.663  61.453  1.00 48.75           C
    ATOM   1103  CG  GLU A 285      43.519   2.806  62.102  1.00 50.55           C
    ATOM   1104  CD  GLU A 285      42.107   3.253  61.716  1.00 51.85           C
    ATOM   1105  OE1 GLU A 285      41.950   3.787  60.595  1.00 50.41           O
    ATOM   1106  OE2 GLU A 285      41.188   3.036  62.535  1.00 51.92           O1-
    ATOM   1107  H   GLU A 285      45.975   4.287  63.602  1.00  0.00           H
    ATOM   1108  HA  GLU A 285      46.042   2.092  61.644  1.00  0.00           H
    ATOM   1109  HB2 GLU A 285      44.515   4.709  61.744  1.00  0.00           H
    ATOM   1110  HB3 GLU A 285      44.504   3.632  60.371  1.00  0.00           H
    ATOM   1111  HG2 GLU A 285      43.628   1.768  61.787  1.00  0.00           H
    ATOM   1112  HG3 GLU A 285      43.615   2.804  63.187  1.00  0.00           H
    ATOM   1113  N   GLU A 286      47.791   4.811  61.151  1.00 45.01           N
    ATOM   1114  CA  GLU A 286      48.932   5.352  60.403  1.00 44.88           C
    ATOM   1115  C   GLU A 286      50.229   4.533  60.578  1.00 52.79           C
    ATOM   1116  O   GLU A 286      51.114   4.643  59.730  1.00 54.86           O
    ATOM   1117  CB  GLU A 286      49.173   6.822  60.806  1.00 38.81           C
    ATOM   1118  CG  GLU A 286      48.133   7.812  60.237  1.00 39.41           C
    ATOM   1119  CD  GLU A 286      48.392   9.267  60.648  1.00 43.85           C
    ATOM   1120  OE1 GLU A 286      49.246   9.499  61.530  1.00 44.69           O
    ATOM   1121  OE2 GLU A 286      47.718  10.149  60.073  1.00 46.16           O1-
    ATOM   1122  H   GLU A 286      47.566   5.283  62.017  1.00  0.00           H
    ATOM   1123  HA  GLU A 286      48.688   5.327  59.340  1.00  0.00           H
    ATOM   1124  HB2 GLU A 286      49.187   6.891  61.895  1.00  0.00           H
    ATOM   1125  HB3 GLU A 286      50.166   7.138  60.478  1.00  0.00           H
    ATOM   1126  HG2 GLU A 286      48.135   7.761  59.148  1.00  0.00           H
    ATOM   1127  HG3 GLU A 286      47.126   7.538  60.552  1.00  0.00           H
    ATOM   1128  N   GLU A 287      50.309   3.721  61.651  1.00 53.92           N
    ATOM   1129  CA  GLU A 287      51.419   2.836  62.042  1.00 52.58           C
    ATOM   1130  C   GLU A 287      51.443   1.683  61.053  1.00 45.21           C
    ATOM   1131  O   GLU A 287      52.470   1.058  60.796  1.00 90.00           O
    ATOM   1132  CB  GLU A 287      51.127   2.351  63.486  1.00 58.94           C
    ATOM   1133  CG  GLU A 287      51.991   1.222  64.086  1.00 68.31           C
    ATOM   1134  CD  GLU A 287      51.296   0.641  65.319  1.00 76.27           C
    ATOM   1135  OE1 GLU A 287      51.355   1.301  66.379  1.00 76.78           O
    ATOM   1136  OE2 GLU A 287      50.679  -0.436  65.164  1.00 80.66           O1-
    ATOM   1137  H   GLU A 287      49.531   3.725  62.295  1.00  0.00           H
    ATOM   1138  HA  GLU A 287      52.363   3.383  62.017  1.00  0.00           H
    ATOM   1140  HB2 GLU A 287      51.185   3.207  64.159  1.00  0.00           H
    ATOM   1141  HB3 GLU A 287      50.096   2.021  63.531  1.00  0.00           H
    ATOM   1142  HG2 GLU A 287      52.141   0.408  63.377  1.00  0.00           H
    ATOM   1143  HG3 GLU A 287      52.980   1.594  64.354  1.00  0.00           H
    ATOM   2304  ZN  ZN2 A   1      24.679   4.442  68.470  1.00 42.36          ZN2+
  extra_rotamers: ""
ligand:
  residue_configurations: []
  extra_templates: ""
  extra_templates_coordinates: ""
  coordinates: |
    ATOM   1145  N   GLU B 359       2.236  -0.786  78.341  1.00 68.52           N1+
    ATOM   1146  CA  GLU B 359       3.386  -0.018  78.820  1.00 63.63           C
    ATOM   1147  C   GLU B 359       4.697  -0.824  78.744  1.00 50.89           C
    ATOM   1148  O   GLU B 359       5.759  -0.239  78.948  1.00 48.12           O
    ATOM   1149  CB  GLU B 359       3.101   0.375  80.282  1.00 70.28           C
    ATOM   1150  CG  GLU B 359       1.816   1.207  80.476  1.00 74.81           C
    ATOM   1151  CD  GLU B 359       1.198   0.949  81.848  1.00 80.12           C
    ATOM   1152  OE1 GLU B 359       1.810   1.384  82.846  1.00 83.58           O
    ATOM   1153  OE2 GLU B 359       0.143   0.279  81.877  1.00 79.47           O1-
    ATOM   1154  H   GLU B 359       1.818  -1.403  79.025  1.00  0.00           H
    ATOM   1155  HA  GLU B 359       3.506   0.890  78.227  1.00  0.00           H
    ATOM   1156  HB2 GLU B 359       3.056  -0.527  80.891  1.00  0.00           H
    ATOM   1157  HB3 GLU B 359       3.942   0.945  80.679  1.00  0.00           H
    ATOM   1158  HG2 GLU B 359       2.034   2.270  80.368  1.00  0.00           H
    ATOM   1159  HG3 GLU B 359       1.060   0.982  79.725  1.00  0.00           H
    ATOM   1160  N   GLY B 360       4.612  -2.135  78.465  1.00 42.66           N
    ATOM   1161  CA  GLY B 360       5.753  -3.038  78.363  1.00 47.23           C
    ATOM   1162  C   GLY B 360       6.177  -3.638  79.720  1.00 51.70           C
    ATOM   1163  O   GLY B 360       7.156  -4.383  79.749  1.00 51.89           O
    ATOM   1164  H   GLY B 360       3.699  -2.547  78.326  1.00  0.00           H
    ATOM   1165  HA2 GLY B 360       5.480  -3.854  77.693  1.00  0.00           H
    ATOM   1166  HA3 GLY B 360       6.584  -2.525  77.884  1.00  0.00           H
    ATOM   1167  N   HIE B 365       8.433  -7.838  85.951  1.00 51.36           N1+
    ATOM   1168  CA  HIE B 365       8.609  -6.436  86.351  1.00 50.31           C
    ATOM   1169  C   HIE B 365       7.847  -6.135  87.649  1.00 50.59           C
    ATOM   1170  O   HIE B 365       8.459  -5.640  88.590  1.00 50.97           O
    ATOM   1171  CB  HIE B 365       8.157  -5.446  85.262  1.00 50.41           C
    ATOM   1172  CG  HIE B 365       9.027  -5.355  84.041  1.00 47.08           C
    ATOM   1173  ND1 HIE B 365       8.856  -6.188  82.937  1.00 45.84           N
    ATOM   1174  CD2 HIE B 365      10.025  -4.447  83.761  1.00 42.66           C
    ATOM   1175  CE1 HIE B 365       9.706  -5.736  82.031  1.00 41.69           C
    ATOM   1176  NE2 HIE B 365      10.432  -4.712  82.469  1.00 39.99           N
    ATOM   1177  H   HIE B 365       7.853  -8.013  85.142  1.00  0.00           H
    ATOM   1178  HA  HIE B 365       9.671  -6.262  86.545  1.00  0.00           H
    ATOM   1179  HB2 HIE B 365       7.143  -5.668  84.937  1.00  0.00           H
    ATOM   1180  HB3 HIE B 365       8.110  -4.438  85.679  1.00  0.00           H
    ATOM   1181  HD2 HIE B 365      10.434  -3.632  84.338  1.00  0.00           H
    ATOM   1182  HE2 HIE B 365      11.127  -4.194  81.943  1.00  0.00           H
    ATOM   1183  HE1 HIE B 365       9.782  -6.145  81.036  1.00  0.00           H
    ATOM   1184  N   ASN B 385      10.841 -14.823  80.548  1.00 58.14           N1+
    ATOM   1185  CA  ASN B 385      11.946 -14.012  80.037  1.00 56.60           C
    ATOM   1186  C   ASN B 385      12.433 -13.004  81.089  1.00 52.12           C
    ATOM   1187  O   ASN B 385      11.731 -12.039  81.393  1.00 48.57           O
    ATOM   1188  CB  ASN B 385      11.511 -13.321  78.723  1.00 62.43           C
    ATOM   1189  CG  ASN B 385      12.611 -12.477  78.059  1.00 69.17           C
    ATOM   1190  OD1 ASN B 385      13.802 -12.747  78.204  1.00 68.21           O
    ATOM   1191  ND2 ASN B 385      12.208 -11.444  77.318  1.00 73.90           N
    ATOM   1192  H   ASN B 385       9.988 -14.791  80.008  1.00  0.00           H
    ATOM   1193  HA  ASN B 385      12.754 -14.706  79.801  1.00  0.00           H
    ATOM   1194  HB2 ASN B 385      11.200 -14.077  78.000  1.00  0.00           H
    ATOM   1195  HB3 ASN B 385      10.636 -12.692  78.902  1.00  0.00           H
    ATOM   1196 HD21 ASN B 385      11.226 -11.244  77.202  1.00  0.00           H
    ATOM   1197 HD22 ASN B 385      12.894 -10.878  76.833  1.00  0.00           H
    ATOM   1198  N   ASN B 387      14.823 -10.885  80.617  1.00 50.43           N1+
    ATOM   1199  CA  ASN B 387      15.304  -9.693  79.908  1.00 46.36           C
    ATOM   1200  C   ASN B 387      14.195  -8.933  79.165  1.00 48.03           C
    ATOM   1201  O   ASN B 387      14.517  -8.171  78.254  1.00 53.47           O
    ATOM   1202  CB  ASN B 387      16.425 -10.105  78.920  1.00 43.72           C
    ATOM   1203  CG  ASN B 387      17.632 -10.724  79.616  1.00 42.11           C
    ATOM   1204  OD1 ASN B 387      17.975 -10.318  80.721  1.00 40.11           O
    ATOM   1205  ND2 ASN B 387      18.276 -11.703  78.980  1.00 41.32           N
    ATOM   1206  H   ASN B 387      14.634 -11.690  80.034  1.00  0.00           H
    ATOM   1207  HA  ASN B 387      15.716  -8.974  80.619  1.00  0.00           H
    ATOM   1208  HB2 ASN B 387      16.035 -10.796  78.169  1.00  0.00           H
    ATOM   1209  HB3 ASN B 387      16.803  -9.235  78.382  1.00  0.00           H
    ATOM   1210 HD21 ASN B 387      17.952 -12.038  78.081  1.00  0.00           H
    ATOM   1211 HD22 ASN B 387      19.056 -12.167  79.427  1.00  0.00           H
    ATOM   1212  N   ALA B 388      12.920  -9.107  79.555  1.00 47.15           N
    ATOM   1213  CA  ALA B 388      11.800  -8.338  79.002  1.00 46.83           C
    ATOM   1214  C   ALA B 388      11.971  -6.840  79.298  1.00 47.91           C
    ATOM   1215  O   ALA B 388      12.219  -6.507  80.448  1.00 51.50           O
    ATOM   1216  CB  ALA B 388      10.493  -8.869  79.598  1.00 46.54           C
    ATOM   1217  H   ALA B 388      12.707  -9.743  80.310  1.00  0.00           H
    ATOM   1218  HA  ALA B 388      11.776  -8.490  77.921  1.00  0.00           H
    ATOM   1219  HB1 ALA B 388       9.638  -8.345  79.174  1.00  0.00           H
    ATOM   1220  HB2 ALA B 388      10.369  -9.928  79.372  1.00  0.00           H
    ATOM   1221  HB3 ALA B 388      10.458  -8.748  80.682  1.00  0.00           H
    ATOM   1222  N   ALA B 389      11.884  -5.983  78.272  1.00 46.44           N
    ATOM   1223  CA  ALA B 389      12.147  -4.547  78.386  1.00 43.96           C
    ATOM   1224  C   ALA B 389      10.850  -3.759  78.173  1.00 43.67           C
    ATOM   1225  O   ALA B 389      10.104  -4.088  77.248  1.00 47.33           O
    ATOM   1226  CB  ALA B 389      13.202  -4.175  77.335  1.00 44.67           C
    ATOM   1227  H   ALA B 389      11.666  -6.324  77.347  1.00  0.00           H
    ATOM   1228  HA  ALA B 389      12.557  -4.302  79.366  1.00  0.00           H
    ATOM   1229  HB1 ALA B 389      13.482  -3.128  77.413  1.00  0.00           H
    ATOM   1230  HB2 ALA B 389      14.111  -4.761  77.470  1.00  0.00           H
    ATOM   1231  HB3 ALA B 389      12.843  -4.349  76.319  1.00  0.00           H
    ATOM   1232  N   ASP B 390      10.604  -2.734  79.012  1.00 41.64           N
    ATOM   1233  CA  ASP B 390       9.385  -1.916  78.950  1.00 51.10           C
    ATOM   1234  C   ASP B 390       9.525  -0.751  77.940  1.00 62.33           C
    ATOM   1235  O   ASP B 390      10.438  -0.761  77.112  1.00 66.60           O
    ATOM   1236  CB  ASP B 390       8.864  -1.473  80.349  1.00 49.71           C
    ATOM   1237  CG  ASP B 390       9.691  -0.442  81.124  1.00 51.81           C
    ATOM   1238  OD1 ASP B 390      10.694   0.066  80.576  1.00 53.83           O
    ATOM   1239  OD2 ASP B 390       9.265  -0.141  82.259  1.00 53.26           O1-
    ATOM   1240  H   ASP B 390      11.260  -2.506  79.749  1.00  0.00           H
    ATOM   1241  HA  ASP B 390       8.608  -2.548  78.534  1.00  0.00           H
    ATOM   1242  HB2 ASP B 390       7.849  -1.088  80.256  1.00  0.00           H
    ATOM   1243  HB3 ASP B 390       8.803  -2.363  80.975  1.00  0.00           H
    ATOM   1244  N   SER B 391       8.618   0.237  78.043  1.00 61.37           N
    ATOM   1245  CA  SER B 391       8.567   1.474  77.255  1.00 58.74           C
    ATOM   1246  C   SER B 391       9.893   2.247  77.163  1.00 57.56           C
    ATOM   1247  O   SER B 391      10.256   2.659  76.061  1.00 63.14           O
    ATOM   1248  CB  SER B 391       7.470   2.403  77.821  1.00 60.23           C
    ATOM   1249  OG  SER B 391       6.210   2.077  77.279  1.00 63.16           O
    ATOM   1250  H   SER B 391       7.892   0.147  78.741  1.00  0.00           H
    ATOM   1251  HA  SER B 391       8.300   1.195  76.234  1.00  0.00           H
    ATOM   1252  HB2 SER B 391       7.430   2.357  78.911  1.00  0.00           H
    ATOM   1253  HB3 SER B 391       7.659   3.446  77.559  1.00  0.00           H
    ATOM   1254  HG  SER B 391       5.922   1.246  77.671  1.00  0.00           H
    ATOM   1255  N   ASP B 392      10.584   2.412  78.306  1.00 51.01           N
    ATOM   1256  CA  ASP B 392      11.858   3.130  78.398  1.00 48.96           C
    ATOM   1257  C   ASP B 392      13.047   2.154  78.520  1.00 43.50           C
    ATOM   1258  O   ASP B 392      14.149   2.592  78.835  1.00 43.59           O
    ATOM   1259  CB  ASP B 392      11.839   4.167  79.548  1.00 56.11           C
    ATOM   1260  CG  ASP B 392      12.818   5.324  79.324  1.00 62.98           C
    ATOM   1261  OD1 ASP B 392      12.568   6.101  78.378  1.00 67.67           O
    ATOM   1262  OD2 ASP B 392      13.784   5.429  80.111  1.00 63.38           O1-
    ATOM   1263  H   ASP B 392      10.241   2.003  79.165  1.00  0.00           H
    ATOM   1264  HA  ASP B 392      12.030   3.669  77.464  1.00  0.00           H
    ATOM   1265  HB2 ASP B 392      10.844   4.593  79.680  1.00  0.00           H
    ATOM   1266  HB3 ASP B 392      12.061   3.693  80.503  1.00  0.00           H
    ATOM   1267  N   GLY B 393      12.839   0.859  78.238  1.00 40.77           N
    ATOM   1268  CA  GLY B 393      13.907  -0.131  78.146  1.00 43.15           C
    ATOM   1269  C   GLY B 393      14.350  -0.688  79.502  1.00 47.64           C
    ATOM   1270  O   GLY B 393      15.376  -1.367  79.553  1.00 47.18           O
    ATOM   1271  H   GLY B 393      11.905   0.549  78.005  1.00  0.00           H
    ATOM   1272  HA2 GLY B 393      13.531  -0.957  77.550  1.00  0.00           H
    ATOM   1273  HA3 GLY B 393      14.768   0.277  77.616  1.00  0.00           H
    ATOM   1274  N   TRP B 394      13.597  -0.428  80.584  1.00 53.76           N
    ATOM   1275  CA  TRP B 394      13.815  -1.047  81.887  1.00 52.82           C
    ATOM   1276  C   TRP B 394      13.438  -2.524  81.840  1.00 51.65           C
    ATOM   1277  O   TRP B 394      12.314  -2.853  81.464  1.00 52.14           O
    ATOM   1278  CB  TRP B 394      12.991  -0.361  82.986  1.00 50.98           C
    ATOM   1279  CG  TRP B 394      13.452   0.989  83.421  1.00 53.52           C
    ATOM   1280  CD1 TRP B 394      13.534   2.106  82.666  1.00 55.77           C
    ATOM   1281  CD2 TRP B 394      13.892   1.368  84.752  1.00 54.93           C
    ATOM   1282  NE1 TRP B 394      14.022   3.144  83.434  1.00 56.34           N
    ATOM   1283  CE2 TRP B 394      14.272   2.743  84.731  1.00 57.40           C
    ATOM   1284  CE3 TRP B 394      14.007   0.680  85.979  1.00 56.57           C
    ATOM   1285  CZ2 TRP B 394      14.768   3.396  85.874  1.00 61.10           C
    ATOM   1286  CZ3 TRP B 394      14.492   1.326  87.128  1.00 60.06           C
    ATOM   1287  CH2 TRP B 394      14.882   2.678  87.079  1.00 60.84           C
    ATOM   1288  H   TRP B 394      12.739   0.103  80.485  1.00  0.00           H
    ATOM   1289  HA  TRP B 394      14.871  -0.958  82.151  1.00  0.00           H
    ATOM   1290  HB2 TRP B 394      11.941  -0.296  82.723  1.00  0.00           H
    ATOM   1291  HB3 TRP B 394      13.013  -0.988  83.879  1.00  0.00           H
    ATOM   1292  HD1 TRP B 394      13.273   2.156  81.620  1.00  0.00           H
    ATOM   1293  HE1 TRP B 394      14.154   4.074  83.059  1.00  0.00           H
    ATOM   1294  HE3 TRP B 394      13.720  -0.359  86.040  1.00  0.00           H
    ATOM   1295  HZ2 TRP B 394      15.049   4.438  85.830  1.00  0.00           H
    ATOM   1296  HZ3 TRP B 394      14.559   0.770  88.046  1.00  0.00           H
    ATOM   1297  HH2 TRP B 394      15.255   3.165  87.967  1.00  0.00           H
    ATOM   1298  N   THR B 395      14.385  -3.363  82.261  1.00 51.01           N
    ATOM   1299  CA  THR B 395      14.205  -4.792  82.462  1.00 49.23           C
    ATOM   1300  C   THR B 395      14.012  -5.085  83.968  1.00 45.95           C
    ATOM   1301  O   THR B 395      14.229  -4.185  84.783  1.00 44.87           O
    ATOM   1302  CB  THR B 395      15.433  -5.565  81.898  1.00 50.56           C
    ATOM   1303  OG1 THR B 395      16.559  -5.488  82.753  1.00 55.15           O
    ATOM   1304  CG2 THR B 395      15.811  -5.166  80.460  1.00 46.52           C
    ATOM   1305  H   THR B 395      15.292  -3.001  82.527  1.00  0.00           H
    ATOM   1306  HA  THR B 395      13.305  -5.102  81.952  1.00  0.00           H
    ATOM   1307  HB  THR B 395      15.187  -6.622  81.841  1.00  0.00           H
    ATOM   1308 HG21 THR B 395      16.579  -5.821  80.051  1.00  0.00           H
    ATOM   1309 HG22 THR B 395      14.945  -5.236  79.801  1.00  0.00           H
    ATOM   1310 HG23 THR B 395      16.186  -4.144  80.404  1.00  0.00           H
    ATOM   1311  HG1 THR B 395      17.321  -5.839  82.277  1.00  0.00           H
    ATOM   1312  N   PRO B 396      13.662  -6.341  84.336  1.00 49.87           N
    ATOM   1313  CA  PRO B 396      13.709  -6.806  85.732  1.00 49.58           C
    ATOM   1314  C   PRO B 396      15.029  -6.569  86.488  1.00 49.52           C
    ATOM   1315  O   PRO B 396      14.971  -6.361  87.694  1.00 48.17           O
    ATOM   1316  CB  PRO B 396      13.370  -8.298  85.631  1.00 48.79           C
    ATOM   1317  CG  PRO B 396      12.439  -8.375  84.436  1.00 47.57           C
    ATOM   1318  CD  PRO B 396      13.023  -7.344  83.482  1.00 50.79           C
    ATOM   1319  HA  PRO B 396      12.903  -6.297  86.263  1.00  0.00           H
    ATOM   1320  HB2 PRO B 396      14.268  -8.885  85.427  1.00  0.00           H
    ATOM   1321  HB3 PRO B 396      12.913  -8.691  86.537  1.00  0.00           H
    ATOM   1322  HG2 PRO B 396      12.363  -9.371  84.000  1.00  0.00           H
    ATOM   1323  HG3 PRO B 396      11.438  -8.064  84.740  1.00  0.00           H
    ATOM   1324  HD2 PRO B 396      13.749  -7.815  82.826  1.00  0.00           H
    ATOM   1325  HD3 PRO B 396      12.226  -6.920  82.878  1.00  0.00           H
    ATOM   1326  N   LEU B 397      16.184  -6.557  85.795  1.00 51.07           N
    ATOM   1327  CA  LEU B 397      17.470  -6.232  86.414  1.00 50.26           C
    ATOM   1328  C   LEU B 397      17.586  -4.739  86.772  1.00 49.06           C
    ATOM   1329  O   LEU B 397      18.053  -4.447  87.870  1.00 51.58           O
    ATOM   1330  CB  LEU B 397      18.640  -6.713  85.530  1.00 53.94           C
    ATOM   1331  CG  LEU B 397      20.001  -6.788  86.262  1.00 56.40           C
    ATOM   1332  CD1 LEU B 397      19.998  -7.862  87.369  1.00 55.54           C
    ATOM   1333  CD2 LEU B 397      21.166  -6.974  85.268  1.00 57.17           C
    ATOM   1334  H   LEU B 397      16.185  -6.709  84.795  1.00  0.00           H
    ATOM   1335  HA  LEU B 397      17.515  -6.789  87.351  1.00  0.00           H
    ATOM   1336  HB2 LEU B 397      18.416  -7.708  85.150  1.00  0.00           H
    ATOM   1337  HB3 LEU B 397      18.716  -6.064  84.655  1.00  0.00           H
    ATOM   1338  HG  LEU B 397      20.178  -5.834  86.754  1.00  0.00           H
    ATOM   1339 HD11 LEU B 397      20.909  -8.456  87.384  1.00  0.00           H
    ATOM   1340 HD12 LEU B 397      19.913  -7.394  88.349  1.00  0.00           H
    ATOM   1341 HD13 LEU B 397      19.164  -8.556  87.266  1.00  0.00           H
    ATOM   1342 HD21 LEU B 397      21.901  -6.178  85.384  1.00  0.00           H
    ATOM   1343 HD22 LEU B 397      21.693  -7.918  85.405  1.00  0.00           H
    ATOM   1344 HD23 LEU B 397      20.827  -6.954  84.232  1.00  0.00           H
    ATOM   1345  N   HIE B 398      17.117  -3.833  85.888  1.00 46.61           N
    ATOM   1346  CA  HIE B 398      17.015  -2.391  86.167  1.00 46.38           C
    ATOM   1347  C   HIE B 398      16.089  -2.084  87.356  1.00 50.26           C
    ATOM   1348  O   HIE B 398      16.434  -1.240  88.183  1.00 50.26           O
    ATOM   1349  CB  HIE B 398      16.531  -1.599  84.932  1.00 45.05           C
    ATOM   1350  CG  HIE B 398      17.423  -1.604  83.720  1.00 47.95           C
    ATOM   1351  ND1 HIE B 398      17.275  -2.524  82.681  1.00 49.47           N
    ATOM   1352  CD2 HIE B 398      18.428  -0.721  83.386  1.00 48.11           C
    ATOM   1353  CE1 HIE B 398      18.169  -2.168  81.772  1.00 47.51           C
    ATOM   1354  NE2 HIE B 398      18.884  -1.107  82.141  1.00 47.15           N
    ATOM   1355  H   HIE B 398      16.740  -4.144  85.003  1.00  0.00           H
    ATOM   1356  HA  HIE B 398      18.010  -2.033  86.433  1.00  0.00           H
    ATOM   1357  HB2 HIE B 398      15.545  -1.946  84.623  1.00  0.00           H
    ATOM   1358  HB3 HIE B 398      16.395  -0.553  85.204  1.00  0.00           H
    ATOM   1359  HD2 HIE B 398      18.833   0.134  83.903  1.00  0.00           H
    ATOM   1360  HE2 HIE B 398      19.618  -0.662  81.606  1.00  0.00           H
    ATOM   1361  HE1 HIE B 398      18.297  -2.685  80.833  1.00  0.00           H
    ATOM   1362  N   ALA B 418      20.736  -9.840  81.264  1.00 62.68           N1+
    ATOM   1363  CA  ALA B 418      21.564  -9.065  80.352  1.00 53.82           C
    ATOM   1364  C   ALA B 418      22.047  -7.805  81.068  1.00 52.54           C
    ATOM   1365  O   ALA B 418      21.342  -6.798  81.107  1.00 51.37           O
    ATOM   1366  CB  ALA B 418      20.747  -8.744  79.098  1.00 49.47           C
    ATOM   1367  H   ALA B 418      19.736  -9.736  81.146  1.00  0.00           H
    ATOM   1368  HA  ALA B 418      22.433  -9.653  80.053  1.00  0.00           H
    ATOM   1369  HB1 ALA B 418      21.323  -8.151  78.387  1.00  0.00           H
    ATOM   1370  HB2 ALA B 418      20.459  -9.668  78.602  1.00  0.00           H
    ATOM   1371  HB3 ALA B 418      19.828  -8.206  79.328  1.00  0.00           H
    ATOM   1372  N   VAL B 419      23.232  -7.915  81.676  1.00 53.77           N
    ATOM   1373  CA  VAL B 419      23.812  -6.898  82.551  1.00 55.99           C
    ATOM   1374  C   VAL B 419      24.149  -5.585  81.812  1.00 56.89           C
    ATOM   1375  O   VAL B 419      24.076  -4.520  82.421  1.00 57.41           O
    ATOM   1376  CB  VAL B 419      25.079  -7.461  83.245  1.00 57.29           C
    ATOM   1377  CG1 VAL B 419      25.726  -6.513  84.268  1.00 54.64           C
    ATOM   1378  CG2 VAL B 419      24.776  -8.809  83.928  1.00 58.00           C
    ATOM   1379  H   VAL B 419      23.735  -8.789  81.618  1.00  0.00           H
    ATOM   1380  HA  VAL B 419      23.077  -6.658  83.318  1.00  0.00           H
    ATOM   1381  HB  VAL B 419      25.824  -7.634  82.473  1.00  0.00           H
    ATOM   1382 HG11 VAL B 419      26.629  -6.948  84.697  1.00  0.00           H
    ATOM   1383 HG12 VAL B 419      26.015  -5.566  83.819  1.00  0.00           H
    ATOM   1384 HG13 VAL B 419      25.044  -6.299  85.092  1.00  0.00           H
    ATOM   1385 HG21 VAL B 419      25.607  -9.145  84.537  1.00  0.00           H
    ATOM   1386 HG22 VAL B 419      23.906  -8.729  84.580  1.00  0.00           H
    ATOM   1387 HG23 VAL B 419      24.583  -9.599  83.202  1.00  0.00           H
    ATOM   1388  N   PHE B 420      24.442  -5.678  80.505  1.00 54.94           N
    ATOM   1389  CA  PHE B 420      24.702  -4.539  79.626  1.00 54.64           C
    ATOM   1390  C   PHE B 420      23.476  -4.025  78.856  1.00 57.24           C
    ATOM   1391  O   PHE B 420      23.635  -3.081  78.079  1.00 61.57           O
    ATOM   1392  CB  PHE B 420      25.867  -4.884  78.684  1.00 53.64           C
    ATOM   1393  CG  PHE B 420      27.213  -4.901  79.373  1.00 53.21           C
    ATOM   1394  CD1 PHE B 420      27.780  -3.697  79.839  1.00 51.40           C
    ATOM   1395  CD2 PHE B 420      27.866  -6.123  79.633  1.00 55.46           C
    ATOM   1396  CE1 PHE B 420      28.994  -3.721  80.508  1.00 51.24           C
    ATOM   1397  CE2 PHE B 420      29.086  -6.123  80.294  1.00 55.02           C
    ATOM   1398  CZ  PHE B 420      29.647  -4.927  80.723  1.00 53.05           C
    ATOM   1399  H   PHE B 420      24.463  -6.590  80.070  1.00  0.00           H
    ATOM   1400  HA  PHE B 420      25.016  -3.687  80.231  1.00  0.00           H
    ATOM   1401  HB2 PHE B 420      25.696  -5.860  78.231  1.00  0.00           H
    ATOM   1402  HB3 PHE B 420      25.935  -4.170  77.863  1.00  0.00           H
    ATOM   1403  HD1 PHE B 420      27.274  -2.756  79.677  1.00  0.00           H
    ATOM   1404  HD2 PHE B 420      27.432  -7.058  79.308  1.00  0.00           H
    ATOM   1405  HE1 PHE B 420      29.435  -2.799  80.858  1.00  0.00           H
    ATOM   1406  HE2 PHE B 420      29.604  -7.054  80.471  1.00  0.00           H
    ATOM   1407  HZ  PHE B 420      30.599  -4.931  81.229  1.00  0.00           H
    ATOM   1408  N   ALA B 421      22.279  -4.592  79.103  1.00 56.16           N
    ATOM   1409  CA  ALA B 421      21.011  -4.028  78.632  1.00 51.69           C
    ATOM   1410  C   ALA B 421      20.803  -2.636  79.236  1.00 48.21           C
    ATOM   1411  O   ALA B 421      21.026  -2.472  80.435  1.00 45.34           O
    ATOM   1412  CB  ALA B 421      19.846  -4.950  79.010  1.00 50.67           C
    ATOM   1413  H   ALA B 421      22.211  -5.361  79.755  1.00  0.00           H
    ATOM   1414  HA  ALA B 421      21.060  -3.968  77.547  1.00  0.00           H
    ATOM   1415  HB1 ALA B 421      18.913  -4.611  78.563  1.00  0.00           H
    ATOM   1416  HB2 ALA B 421      20.019  -5.965  78.658  1.00  0.00           H
    ATOM   1417  HB3 ALA B 421      19.700  -4.980  80.089  1.00  0.00           H
    ATOM   1418  N   MET B 422      20.419  -1.666  78.399  1.00 49.72           N
    ATOM   1419  CA  MET B 422      20.276  -0.274  78.811  1.00 55.26           C
    ATOM   1420  C   MET B 422      18.913   0.283  78.400  1.00 56.23           C
    ATOM   1421  O   MET B 422      18.291  -0.206  77.457  1.00 56.38           O
    ATOM   1422  CB  MET B 422      21.458   0.590  78.304  1.00 60.10           C
    ATOM   1423  CG  MET B 422      22.063   0.193  76.947  1.00 66.30           C
    ATOM   1424  SD  MET B 422      23.163   1.430  76.210  1.00 70.23           S
    ATOM   1425  CE  MET B 422      24.493   1.441  77.442  1.00 69.14           C
    ATOM   1426  H   MET B 422      20.261  -1.873  77.422  1.00  0.00           H
    ATOM   1427  HA  MET B 422      20.282  -0.216  79.897  1.00  0.00           H
    ATOM   1428  HB2 MET B 422      21.179   1.643  78.284  1.00  0.00           H
    ATOM   1429  HB3 MET B 422      22.249   0.519  79.050  1.00  0.00           H
    ATOM   1430  HG2 MET B 422      22.624  -0.736  77.048  1.00  0.00           H
    ATOM   1431  HG3 MET B 422      21.272  -0.010  76.233  1.00  0.00           H
    ATOM   1432  HE1 MET B 422      25.293   2.108  77.122  1.00  0.00           H
    ATOM   1433  HE2 MET B 422      24.125   1.791  78.406  1.00  0.00           H
    ATOM   1434  HE3 MET B 422      24.903   0.438  77.567  1.00  0.00           H
    ATOM   1435  N   THR B 423      18.487   1.318  79.134  1.00 57.94           N
    ATOM   1436  CA  THR B 423      17.285   2.106  78.872  1.00 58.98           C
    ATOM   1437  C   THR B 423      17.410   2.937  77.574  1.00 62.80           C
    ATOM   1438  O   THR B 423      18.509   3.077  77.049  1.00 62.43           O
    ATOM   1439  CB  THR B 423      17.038   3.062  80.064  1.00 55.88           C
    ATOM   1440  OG1 THR B 423      18.172   3.872  80.323  1.00 57.26           O
    ATOM   1441  CG2 THR B 423      16.692   2.285  81.339  1.00 53.66           C
    ATOM   1442  H   THR B 423      19.058   1.651  79.899  1.00  0.00           H
    ATOM   1443  HA  THR B 423      16.448   1.418  78.768  1.00  0.00           H
    ATOM   1444  HB  THR B 423      16.210   3.737  79.842  1.00  0.00           H
    ATOM   1445 HG21 THR B 423      16.376   2.972  82.119  1.00  0.00           H
    ATOM   1446 HG22 THR B 423      15.870   1.591  81.162  1.00  0.00           H
    ATOM   1447 HG23 THR B 423      17.537   1.707  81.712  1.00  0.00           H
    ATOM   1448  HG1 THR B 423      17.866   4.750  80.591  1.00  0.00           H
    ATOM   1449  N   TYR B 424      16.285   3.445  77.050  1.00 65.14           N
    ATOM   1450  CA  TYR B 424      16.261   4.264  75.834  1.00 66.90           C
    ATOM   1451  C   TYR B 424      15.695   5.657  76.138  1.00 67.79           C
    ATOM   1452  O   TYR B 424      14.549   5.958  75.809  1.00 75.17           O
    ATOM   1453  CB  TYR B 424      15.597   3.555  74.627  1.00 68.22           C
    ATOM   1454  CG  TYR B 424      14.936   2.199  74.814  1.00 65.43           C
    ATOM   1455  CD1 TYR B 424      15.717   1.028  74.773  1.00 64.13           C
    ATOM   1456  CD2 TYR B 424      13.534   2.094  74.926  1.00 63.32           C
    ATOM   1457  CE1 TYR B 424      15.107  -0.240  74.809  1.00 64.81           C
    ATOM   1458  CE2 TYR B 424      12.918   0.827  74.938  1.00 64.61           C
    ATOM   1459  CZ  TYR B 424      13.704  -0.341  74.871  1.00 65.99           C
    ATOM   1460  OH  TYR B 424      13.113  -1.571  74.880  1.00 68.03           O
    ATOM   1461  H   TYR B 424      15.398   3.295  77.515  1.00  0.00           H
    ATOM   1462  HA  TYR B 424      17.283   4.463  75.505  1.00  0.00           H
    ATOM   1463  HB2 TYR B 424      14.898   4.219  74.122  1.00  0.00           H
    ATOM   1464  HB3 TYR B 424      16.376   3.400  73.878  1.00  0.00           H
    ATOM   1465  HD1 TYR B 424      16.787   1.109  74.683  1.00  0.00           H
    ATOM   1466  HD2 TYR B 424      12.925   2.986  74.963  1.00  0.00           H
    ATOM   1467  HE1 TYR B 424      15.716  -1.132  74.769  1.00  0.00           H
    ATOM   1468  HE2 TYR B 424      11.843   0.754  74.999  1.00  0.00           H
    ATOM   1469  HH  TYR B 424      12.152  -1.539  74.892  1.00  0.00           H
    ATOM   1470  N   SER B 425      16.560   6.474  76.750  1.00 58.40           N
    ATOM   1471  CA  SER B 425      16.413   7.906  76.998  1.00 52.46           C
    ATOM   1472  C   SER B 425      17.766   8.406  77.521  1.00 51.23           C
    ATOM   1473  O   SER B 425      18.437   9.191  76.853  1.00 55.40           O
    ATOM   1474  CB  SER B 425      15.197   8.259  77.892  1.00 51.34           C
    ATOM   1475  OG  SER B 425      15.295   7.781  79.218  1.00 51.18           O
    ATOM   1476  H   SER B 425      17.463   6.101  77.003  1.00  0.00           H
    ATOM   1477  HA  SER B 425      16.223   8.392  76.044  1.00  0.00           H
    ATOM   1478  HB2 SER B 425      15.098   9.344  77.947  1.00  0.00           H
    ATOM   1479  HB3 SER B 425      14.266   7.903  77.453  1.00  0.00           H
    ATOM   1480  HG  SER B 425      14.815   6.946  79.290  1.00  0.00           H
    ATOM   1481  N   ASP B 426      18.166   7.871  78.680  1.00 45.55           N
    ATOM   1482  CA  ASP B 426      19.413   8.154  79.381  1.00 47.69           C
    ATOM   1483  C   ASP B 426      20.527   7.145  79.039  1.00 49.66           C
    ATOM   1484  O   ASP B 426      21.696   7.480  79.230  1.00 51.03           O
    ATOM   1485  CB  ASP B 426      19.217   8.243  80.914  1.00 56.81           C
    ATOM   1486  CG  ASP B 426      18.418   7.123  81.601  1.00 64.25           C
    ATOM   1487  OD1 ASP B 426      17.548   6.480  80.973  1.00 71.26           O
    ATOM   1488  OD2 ASP B 426      18.662   6.962  82.812  1.00 62.98           O1-
    ATOM   1489  H   ASP B 426      17.529   7.262  79.182  1.00  0.00           H
    ATOM   1490  HA  ASP B 426      19.775   9.132  79.056  1.00  0.00           H
    ATOM   1491  HB2 ASP B 426      20.197   8.299  81.389  1.00  0.00           H
    ATOM   1492  HB3 ASP B 426      18.710   9.181  81.139  1.00  0.00           H
    ATOM   1493  N   MET B 427      20.157   5.947  78.542  1.00 53.34           N
    ATOM   1494  CA  MET B 427      21.051   4.858  78.123  1.00 56.63           C
    ATOM   1495  C   MET B 427      21.976   4.362  79.233  1.00 56.50           C
    ATOM   1496  O   MET B 427      23.199   4.452  79.130  1.00 54.65           O
    ATOM   1497  CB  MET B 427      21.784   5.190  76.817  1.00 61.75           C
    ATOM   1498  CG  MET B 427      20.833   5.438  75.641  1.00 67.13           C
    ATOM   1499  SD  MET B 427      21.642   5.274  74.036  1.00 71.70           S
    ATOM   1500  CE  MET B 427      22.926   6.530  74.216  1.00 71.67           C
    ATOM   1501  H   MET B 427      19.171   5.752  78.457  1.00  0.00           H
    ATOM   1502  HA  MET B 427      20.433   3.996  77.924  1.00  0.00           H
    ATOM   1503  HB2 MET B 427      22.405   6.071  76.974  1.00  0.00           H
    ATOM   1504  HB3 MET B 427      22.464   4.378  76.556  1.00  0.00           H
    ATOM   1505  HG2 MET B 427      20.008   4.727  75.658  1.00  0.00           H
    ATOM   1506  HG3 MET B 427      20.389   6.430  75.721  1.00  0.00           H
    ATOM   1507  HE1 MET B 427      23.549   6.526  73.332  1.00  0.00           H
    ATOM   1508  HE2 MET B 427      22.488   7.521  74.339  1.00  0.00           H
    ATOM   1509  HE3 MET B 427      23.576   6.320  75.063  1.00  0.00           H
    ATOM   1510  N   GLN B 428      21.326   3.854  80.283  1.00 58.25           N
    ATOM   1511  CA  GLN B 428      21.949   3.408  81.514  1.00 57.11           C
    ATOM   1512  C   GLN B 428      21.599   1.949  81.788  1.00 53.31           C
    ATOM   1513  O   GLN B 428      20.440   1.556  81.642  1.00 48.91           O
    ATOM   1514  CB  GLN B 428      21.463   4.336  82.637  1.00 57.27           C
    ATOM   1515  CG  GLN B 428      22.253   5.656  82.663  1.00 59.76           C
    ATOM   1516  CD  GLN B 428      21.734   6.642  83.707  1.00 62.63           C
    ATOM   1517  OE1 GLN B 428      21.004   6.274  84.626  1.00 64.04           O
    ATOM   1518  NE2 GLN B 428      22.124   7.910  83.571  1.00 66.26           N
    ATOM   1519  H   GLN B 428      20.315   3.818  80.267  1.00  0.00           H
    ATOM   1520  HA  GLN B 428      23.037   3.481  81.444  1.00  0.00           H
    ATOM   1521  HB2 GLN B 428      20.400   4.544  82.506  1.00  0.00           H
    ATOM   1522  HB3 GLN B 428      21.542   3.841  83.601  1.00  0.00           H
    ATOM   1523  HG2 GLN B 428      23.306   5.457  82.866  1.00  0.00           H
    ATOM   1524  HG3 GLN B 428      22.207   6.137  81.685  1.00  0.00           H
    ATOM   1525 HE21 GLN B 428      22.720   8.182  82.803  1.00  0.00           H
    ATOM   1526 HE22 GLN B 428      21.826   8.600  84.244  1.00  0.00           H
    ATOM   1527  N   THR B 429      22.617   1.192  82.220  1.00 53.83           N
    ATOM   1528  CA  THR B 429      22.502  -0.188  82.682  1.00 53.71           C
    ATOM   1529  C   THR B 429      21.900  -0.250  84.100  1.00 51.92           C
    ATOM   1530  O   THR B 429      21.756   0.785  84.752  1.00 51.11           O
    ATOM   1531  CB  THR B 429      23.895  -0.873  82.708  1.00 51.34           C
    ATOM   1532  OG1 THR B 429      24.726  -0.375  83.741  1.00 52.74           O
    ATOM   1533  CG2 THR B 429      24.640  -0.811  81.366  1.00 50.56           C
    ATOM   1534  H   THR B 429      23.523   1.622  82.365  1.00  0.00           H
    ATOM   1535  HA  THR B 429      21.846  -0.730  82.006  1.00  0.00           H
    ATOM   1536  HB  THR B 429      23.742  -1.929  82.927  1.00  0.00           H
    ATOM   1537 HG21 THR B 429      25.564  -1.389  81.405  1.00  0.00           H
    ATOM   1538 HG22 THR B 429      24.029  -1.221  80.561  1.00  0.00           H
    ATOM   1539 HG23 THR B 429      24.902   0.213  81.098  1.00  0.00           H
    ATOM   1540  HG1 THR B 429      25.558  -0.857  83.722  1.00  0.00           H
    ATOM   1541  N   ALA B 430      21.567  -1.469  84.556  1.00 51.63           N
    ATOM   1542  CA  ALA B 430      21.043  -1.735  85.894  1.00 51.39           C
    ATOM   1543  C   ALA B 430      21.955  -1.250  87.034  1.00 53.74           C
    ATOM   1544  O   ALA B 430      21.442  -0.665  87.983  1.00 56.21           O
    ATOM   1545  CB  ALA B 430      20.741  -3.228  86.031  1.00 50.47           C
    ATOM   1546  H   ALA B 430      21.691  -2.280  83.964  1.00  0.00           H
    ATOM   1547  HA  ALA B 430      20.102  -1.191  85.982  1.00  0.00           H
    ATOM   1548  HB1 ALA B 430      20.362  -3.459  87.023  1.00  0.00           H
    ATOM   1549  HB2 ALA B 430      19.985  -3.542  85.311  1.00  0.00           H
    ATOM   1550  HB3 ALA B 430      21.629  -3.837  85.869  1.00  0.00           H
    ATOM   1551  N   ALA B 431      23.281  -1.421  86.892  1.00 52.57           N
    ATOM   1552  CA  ALA B 431      24.306  -0.918  87.815  1.00 52.53           C
    ATOM   1553  C   ALA B 431      24.237   0.600  88.062  1.00 57.58           C
    ATOM   1554  O   ALA B 431      24.460   1.035  89.192  1.00 57.94           O
    ATOM   1555  CB  ALA B 431      25.693  -1.301  87.280  1.00 49.93           C
    ATOM   1556  H   ALA B 431      23.624  -1.923  86.083  1.00  0.00           H
    ATOM   1557  HA  ALA B 431      24.163  -1.412  88.775  1.00  0.00           H
    ATOM   1558  HB1 ALA B 431      26.448  -1.159  88.052  1.00  0.00           H
    ATOM   1559  HB2 ALA B 431      25.744  -2.343  86.973  1.00  0.00           H
    ATOM   1560  HB3 ALA B 431      25.972  -0.699  86.413  1.00  0.00           H
    ATOM   1561  N   ASP B 432      23.884   1.357  87.010  1.00 61.95           N
    ATOM   1562  CA  ASP B 432      23.670   2.806  87.029  1.00 61.12           C
    ATOM   1563  C   ASP B 432      22.339   3.220  87.688  1.00 61.10           C
    ATOM   1564  O   ASP B 432      22.242   4.360  88.141  1.00 61.61           O
    ATOM   1565  CB  ASP B 432      23.756   3.429  85.615  1.00 61.80           C
    ATOM   1566  CG  ASP B 432      24.969   3.010  84.777  1.00 65.61           C
    ATOM   1567  OD1 ASP B 432      26.050   2.803  85.374  1.00 67.43           O
    ATOM   1568  OD2 ASP B 432      24.810   2.983  83.536  1.00 66.12           O1-
    ATOM   1569  H   ASP B 432      23.717   0.906  86.121  1.00  0.00           H
    ATOM   1570  HA  ASP B 432      24.464   3.249  87.632  1.00  0.00           H
    ATOM   1571  HB2 ASP B 432      22.870   3.112  85.075  1.00  0.00           H
    ATOM   1572  HB3 ASP B 432      23.733   4.518  85.670  1.00  0.00           H
    ATOM   1573  N   LYS B 433      21.345   2.312  87.739  1.00 62.65           N
    ATOM   1574  CA  LYS B 433      20.046   2.552  88.379  1.00 63.32           C
    ATOM   1575  C   LYS B 433      20.076   2.441  89.904  1.00 61.56           C
    ATOM   1576  O   LYS B 433      19.128   2.917  90.529  1.00 59.09           O
    ATOM   1577  CB  LYS B 433      18.971   1.586  87.833  1.00 65.20           C
    ATOM   1578  CG  LYS B 433      18.741   1.667  86.323  1.00 64.49           C
    ATOM   1579  CD  LYS B 433      18.559   3.101  85.815  1.00 65.97           C
    ATOM   1580  CE  LYS B 433      18.176   3.152  84.342  1.00 69.46           C
    ATOM   1581  NZ  LYS B 433      17.897   4.534  83.934  1.00 70.10           N1+
    ATOM   1582  H   LYS B 433      21.491   1.390  87.351  1.00  0.00           H
    ATOM   1583  HA  LYS B 433      19.738   3.577  88.171  1.00  0.00           H
    ATOM   1584  HB2 LYS B 433      19.196   0.556  88.107  1.00  0.00           H
    ATOM   1585  HB3 LYS B 433      18.021   1.813  88.317  1.00  0.00           H
    ATOM   1586  HG2 LYS B 433      19.575   1.200  85.809  1.00  0.00           H
    ATOM   1587  HG3 LYS B 433      17.862   1.073  86.073  1.00  0.00           H
    ATOM   1588  HD2 LYS B 433      17.806   3.612  86.416  1.00  0.00           H
    ATOM   1589  HD3 LYS B 433      19.489   3.658  85.933  1.00  0.00           H
    ATOM   1590  HE2 LYS B 433      18.974   2.744  83.720  1.00  0.00           H
    ATOM   1591  HE3 LYS B 433      17.286   2.545  84.173  1.00  0.00           H
    ATOM   1592  HZ1 LYS B 433      17.641   4.566  82.957  1.00  0.00           H
    ATOM   1593  HZ2 LYS B 433      17.139   4.912  84.484  1.00  0.00           H
    ATOM   1594  HZ3 LYS B 433      18.714   5.119  84.062  1.00  0.00           H
    ATOM   1595  N   GLN B 452      29.946  -6.473  85.528  1.00 72.85           N1+
    ATOM   1596  CA  GLN B 452      30.354  -5.763  84.305  1.00 74.88           C
    ATOM   1597  C   GLN B 452      31.871  -5.799  84.025  1.00 76.30           C
    ATOM   1598  O   GLN B 452      32.277  -5.878  82.865  1.00 76.79           O
    ATOM   1599  CB  GLN B 452      29.875  -4.295  84.353  1.00 76.16           C
    ATOM   1600  CG  GLN B 452      28.355  -4.136  84.171  1.00 77.27           C
    ATOM   1601  CD  GLN B 452      27.840  -2.692  84.182  1.00 75.83           C
    ATOM   1602  OE1 GLN B 452      26.639  -2.481  84.057  1.00 75.79           O
    ATOM   1603  NE2 GLN B 452      28.712  -1.690  84.319  1.00 73.74           N
    ATOM   1604  H   GLN B 452      29.478  -5.919  86.234  1.00  0.00           H
    ATOM   1605  HA  GLN B 452      29.864  -6.260  83.466  1.00  0.00           H
    ATOM   1606  HB2 GLN B 452      30.186  -3.842  85.295  1.00  0.00           H
    ATOM   1607  HB3 GLN B 452      30.375  -3.729  83.565  1.00  0.00           H
    ATOM   1608  HG2 GLN B 452      28.056  -4.587  83.225  1.00  0.00           H
    ATOM   1609  HG3 GLN B 452      27.831  -4.680  84.953  1.00  0.00           H
    ATOM   1610 HE21 GLN B 452      29.699  -1.878  84.419  1.00  0.00           H
    ATOM   1611 HE22 GLN B 452      28.388  -0.734  84.324  1.00  0.00           H
    ATOM   1612  N   MET B 455      33.106  -9.958  83.989  1.00 75.68           N1+
    ATOM   1613  CA  MET B 455      32.501 -10.801  82.945  1.00 78.55           C
    ATOM   1614  C   MET B 455      33.498 -11.160  81.841  1.00 76.14           C
    ATOM   1615  O   MET B 455      34.260 -10.300  81.405  1.00 73.19           O
    ATOM   1616  CB  MET B 455      31.262 -10.137  82.292  1.00 80.11           C
    ATOM   1617  CG  MET B 455      29.980 -10.960  82.480  1.00 78.04           C
    ATOM   1618  SD  MET B 455      28.708 -10.650  81.229  1.00 75.57           S
    ATOM   1619  CE  MET B 455      27.957  -9.171  81.925  1.00 73.25           C
    ATOM   1620  H   MET B 455      32.694  -9.045  84.128  1.00  0.00           H
    ATOM   1621  HA  MET B 455      32.207 -11.726  83.440  1.00  0.00           H
    ATOM   1622  HB2 MET B 455      31.118  -9.124  82.659  1.00  0.00           H
    ATOM   1623  HB3 MET B 455      31.410  -9.992  81.220  1.00  0.00           H
    ATOM   1624  HG2 MET B 455      30.210 -12.021  82.418  1.00  0.00           H
    ATOM   1625  HG3 MET B 455      29.568 -10.789  83.473  1.00  0.00           H
    ATOM   1626  HE1 MET B 455      27.298  -8.723  81.184  1.00  0.00           H
    ATOM   1627  HE2 MET B 455      27.371  -9.439  82.802  1.00  0.00           H
    ATOM   1628  HE3 MET B 455      28.714  -8.443  82.215  1.00  0.00           H
    ATOM   1629  N   TYR B 464      27.935 -16.774  78.221  1.00 60.63           N1+
    ATOM   1630  CA  TYR B 464      26.801 -17.328  77.487  1.00 59.23           C
    ATOM   1631  C   TYR B 464      26.080 -16.237  76.693  1.00 53.91           C
    ATOM   1632  O   TYR B 464      25.840 -15.150  77.218  1.00 52.60           O
    ATOM   1633  CB  TYR B 464      25.836 -18.047  78.444  1.00 65.00           C
    ATOM   1634  CG  TYR B 464      26.340 -19.371  78.998  1.00 68.45           C
    ATOM   1635  CD1 TYR B 464      27.292 -19.387  80.035  1.00 68.94           C
    ATOM   1636  CD2 TYR B 464      25.832 -20.594  78.509  1.00 68.63           C
    ATOM   1637  CE1 TYR B 464      27.694 -20.596  80.626  1.00 69.36           C
    ATOM   1638  CE2 TYR B 464      26.235 -21.810  79.096  1.00 69.00           C
    ATOM   1639  CZ  TYR B 464      27.157 -21.811  80.162  1.00 72.02           C
    ATOM   1640  OH  TYR B 464      27.536 -22.988  80.737  1.00 78.78           O
    ATOM   1641  H   TYR B 464      27.738 -15.969  78.802  1.00  0.00           H
    ATOM   1642  HA  TYR B 464      27.164 -18.067  76.786  1.00  0.00           H
    ATOM   1643  HB2 TYR B 464      25.583 -17.385  79.271  1.00  0.00           H
    ATOM   1644  HB3 TYR B 464      24.894 -18.248  77.930  1.00  0.00           H
    ATOM   1645  HD1 TYR B 464      27.716 -18.462  80.381  1.00  0.00           H
    ATOM   1646  HD2 TYR B 464      25.112 -20.602  77.704  1.00  0.00           H
    ATOM   1647  HE1 TYR B 464      28.409 -20.585  81.435  1.00  0.00           H
    ATOM   1648  HE2 TYR B 464      25.830 -22.742  78.730  1.00  0.00           H
    ATOM   1649  HH  TYR B 464      28.165 -22.872  81.454  1.00  0.00           H
    ATOM   1650  N   ALA B 465      25.719 -16.581  75.448  1.00 55.55           N
    ATOM   1651  CA  ALA B 465      24.915 -15.760  74.550  1.00 59.09           C
    ATOM   1652  C   ALA B 465      23.433 -15.842  74.943  1.00 63.53           C
    ATOM   1653  O   ALA B 465      22.838 -16.911  74.835  1.00 65.26           O
    ATOM   1654  CB  ALA B 465      25.139 -16.251  73.111  1.00 58.51           C
    ATOM   1655  H   ALA B 465      25.970 -17.498  75.101  1.00  0.00           H
    ATOM   1656  HA  ALA B 465      25.250 -14.724  74.615  1.00  0.00           H
    ATOM   1657  HB1 ALA B 465      24.583 -15.649  72.396  1.00  0.00           H
    ATOM   1658  HB2 ALA B 465      26.190 -16.184  72.836  1.00  0.00           H
    ATOM   1659  HB3 ALA B 465      24.832 -17.289  72.980  1.00  0.00           H
    ATOM   1660  N   LEU B 466      22.869 -14.712  75.390  1.00 62.42           N
    ATOM   1661  CA  LEU B 466      21.457 -14.562  75.761  1.00 57.33           C
    ATOM   1662  C   LEU B 466      20.554 -14.271  74.544  1.00 60.24           C
    ATOM   1663  O   LEU B 466      19.340 -14.189  74.720  1.00 63.95           O
    ATOM   1664  CB  LEU B 466      21.309 -13.426  76.799  1.00 51.24           C
    ATOM   1665  CG  LEU B 466      22.266 -13.464  78.006  1.00 48.99           C
    ATOM   1666  CD1 LEU B 466      22.115 -12.170  78.819  1.00 50.17           C
    ATOM   1667  CD2 LEU B 466      22.080 -14.719  78.879  1.00 48.86           C
    ATOM   1668  H   LEU B 466      23.422 -13.865  75.444  1.00  0.00           H
    ATOM   1669  HA  LEU B 466      21.106 -15.492  76.211  1.00  0.00           H
    ATOM   1670  HB2 LEU B 466      21.476 -12.477  76.297  1.00  0.00           H
    ATOM   1671  HB3 LEU B 466      20.278 -13.393  77.157  1.00  0.00           H
    ATOM   1672  HG  LEU B 466      23.294 -13.474  77.642  1.00  0.00           H
    ATOM   1673 HD11 LEU B 466      22.515 -12.267  79.828  1.00  0.00           H
    ATOM   1674 HD12 LEU B 466      22.640 -11.346  78.336  1.00  0.00           H
    ATOM   1675 HD13 LEU B 466      21.069 -11.883  78.913  1.00  0.00           H
    ATOM   1676 HD21 LEU B 466      22.028 -14.482  79.941  1.00  0.00           H
    ATOM   1677 HD22 LEU B 466      21.168 -15.261  78.628  1.00  0.00           H
    ATOM   1678 HD23 LEU B 466      22.916 -15.406  78.747  1.00  0.00           H
    ATOM   1679  N   TRP B 467      21.157 -14.095  73.355  1.00 58.36           N
    ATOM   1680  CA  TRP B 467      20.504 -13.929  72.058  1.00 57.28           C
    ATOM   1681  C   TRP B 467      21.422 -14.455  70.950  1.00 62.95           C
    ATOM   1682  O   TRP B 467      22.634 -14.577  71.136  1.00 64.60           O
    ATOM   1683  CB  TRP B 467      20.183 -12.444  71.762  1.00 53.19           C
    ATOM   1684  CG  TRP B 467      19.142 -11.764  72.595  1.00 51.55           C
    ATOM   1685  CD1 TRP B 467      17.814 -11.739  72.337  1.00 51.24           C
    ATOM   1686  CD2 TRP B 467      19.317 -11.026  73.836  1.00 49.55           C
    ATOM   1687  NE1 TRP B 467      17.170 -10.992  73.303  1.00 49.97           N
    ATOM   1688  CE2 TRP B 467      18.052 -10.516  74.251  1.00 48.33           C
    ATOM   1689  CE3 TRP B 467      20.425 -10.727  74.650  1.00 48.70           C
    ATOM   1690  CZ2 TRP B 467      17.910  -9.716  75.396  1.00 47.80           C
    ATOM   1691  CZ3 TRP B 467      20.303  -9.918  75.795  1.00 46.95           C
    ATOM   1692  CH2 TRP B 467      19.046  -9.404  76.163  1.00 46.94           C
    ATOM   1693  H   TRP B 467      22.163 -14.168  73.311  1.00  0.00           H
    ATOM   1694  HA  TRP B 467      19.584 -14.518  72.035  1.00  0.00           H
    ATOM   1695  HB2 TRP B 467      21.098 -11.854  71.829  1.00  0.00           H
    ATOM   1696  HB3 TRP B 467      19.850 -12.337  70.728  1.00  0.00           H
    ATOM   1697  HD1 TRP B 467      17.348 -12.218  71.488  1.00  0.00           H
    ATOM   1698  HE1 TRP B 467      16.173 -10.829  73.292  1.00  0.00           H
    ATOM   1699  HE3 TRP B 467      21.379 -11.143  74.372  1.00  0.00           H
    ATOM   1700  HZ2 TRP B 467      16.945  -9.329  75.680  1.00  0.00           H
    ATOM   1701  HZ3 TRP B 467      21.180  -9.695  76.385  1.00  0.00           H
    ATOM   1702  HH2 TRP B 467      18.942  -8.774  77.033  1.00  0.00           H
    ATOM   1703  N   ASP B 468      20.809 -14.678  69.780  1.00 66.11           N
    ATOM   1704  CA  ASP B 468      21.473 -14.927  68.502  1.00 67.83           C
    ATOM   1705  C   ASP B 468      22.186 -13.663  67.989  1.00 69.98           C
    ATOM   1706  O   ASP B 468      21.794 -12.548  68.341  1.00 73.58           O
    ATOM   1707  CB  ASP B 468      20.519 -15.469  67.399  1.00 66.92           C
    ATOM   1708  CG  ASP B 468      19.470 -16.506  67.831  1.00 69.82           C
    ATOM   1709  OD1 ASP B 468      18.714 -16.231  68.788  1.00 75.84           O
    ATOM   1710  OD2 ASP B 468      19.382 -17.541  67.137  1.00 67.05           O1-
    ATOM   1711  H   ASP B 468      19.802 -14.611  69.727  1.00  0.00           H
    ATOM   1712  HA  ASP B 468      22.243 -15.674  68.688  1.00  0.00           H
    ATOM   1713  HB2 ASP B 468      19.987 -14.656  66.902  1.00  0.00           H
    ATOM   1714  HB3 ASP B 468      21.134 -15.928  66.623  1.00  0.00           H
    ATOM   1715  N   TYR B 469      23.201 -13.863  67.135  1.00 66.07           N
    ATOM   1716  CA  TYR B 469      23.898 -12.776  66.460  1.00 60.35           C
    ATOM   1717  C   TYR B 469      24.339 -13.204  65.058  1.00 62.42           C
    ATOM   1718  O   TYR B 469      25.108 -14.156  64.929  1.00 62.14           O
    ATOM   1719  CB  TYR B 469      25.073 -12.250  67.311  1.00 57.25           C
    ATOM   1720  CG  TYR B 469      25.787 -11.077  66.669  1.00 57.82           C
    ATOM   1721  CD1 TYR B 469      25.141  -9.826  66.598  1.00 60.87           C
    ATOM   1722  CD2 TYR B 469      27.050 -11.254  66.068  1.00 58.17           C
    ATOM   1723  CE1 TYR B 469      25.734  -8.765  65.890  1.00 60.83           C
    ATOM   1724  CE2 TYR B 469      27.641 -10.192  65.362  1.00 60.86           C
    ATOM   1725  CZ  TYR B 469      26.974  -8.958  65.253  1.00 60.64           C
    ATOM   1726  OH  TYR B 469      27.521  -7.951  64.520  1.00 58.06           O
    ATOM   1727  H   TYR B 469      23.464 -14.803  66.866  1.00  0.00           H
    ATOM   1728  HA  TYR B 469      23.195 -11.952  66.336  1.00  0.00           H
    ATOM   1729  HB2 TYR B 469      24.710 -11.933  68.287  1.00  0.00           H
    ATOM   1730  HB3 TYR B 469      25.796 -13.046  67.492  1.00  0.00           H
    ATOM   1731  HD1 TYR B 469      24.170  -9.688  67.051  1.00  0.00           H
    ATOM   1732  HD2 TYR B 469      27.547 -12.211  66.110  1.00  0.00           H
    ATOM   1733  HE1 TYR B 469      25.226  -7.815  65.814  1.00  0.00           H
    ATOM   1734  HE2 TYR B 469      28.593 -10.337  64.880  1.00  0.00           H
    ATOM   1735  HH  TYR B 469      28.343  -8.207  64.077  1.00  0.00           H
    ATOM   1736  N   GLU B 470      23.883 -12.435  64.058  1.00 65.05           N
    ATOM   1737  CA  GLU B 470      24.321 -12.500  62.667  1.00 69.34           C
    ATOM   1738  C   GLU B 470      25.472 -11.493  62.449  1.00 69.53           C
    ATOM   1739  O   GLU B 470      25.228 -10.293  62.593  1.00 70.41           O
    ATOM   1740  CB  GLU B 470      23.116 -12.144  61.761  1.00 76.17           C
    ATOM   1741  CG  GLU B 470      22.220 -13.349  61.414  1.00 83.19           C
    ATOM   1742  CD  GLU B 470      22.941 -14.368  60.526  1.00 89.29           C
    ATOM   1743  OE1 GLU B 470      23.225 -14.008  59.363  1.00 90.00           O
    ATOM   1744  OE2 GLU B 470      23.215 -15.477  61.033  1.00 90.00           O1-
    ATOM   1745  H   GLU B 470      23.241 -11.684  64.277  1.00  0.00           H
    ATOM   1746  HA  GLU B 470      24.631 -13.516  62.438  1.00  0.00           H
    ATOM   1747  HB2 GLU B 470      22.501 -11.380  62.241  1.00  0.00           H
    ATOM   1748  HB3 GLU B 470      23.459 -11.685  60.832  1.00  0.00           H
    ATOM   1749  HG2 GLU B 470      21.871 -13.835  62.326  1.00  0.00           H
    ATOM   1750  HG3 GLU B 470      21.331 -13.006  60.885  1.00  0.00           H
    ATOM   1751  N   PRO B 471      26.691 -11.975  62.103  1.00 67.74           N
    ATOM   1752  CA  PRO B 471      27.835 -11.122  61.716  1.00 66.12           C
    ATOM   1753  C   PRO B 471      27.544 -10.076  60.627  1.00 65.14           C
    ATOM   1754  O   PRO B 471      26.920 -10.399  59.616  1.00 64.80           O
    ATOM   1755  CB  PRO B 471      28.888 -12.132  61.235  1.00 63.94           C
    ATOM   1756  CG  PRO B 471      28.600 -13.377  62.045  1.00 64.03           C
    ATOM   1757  CD  PRO B 471      27.081 -13.387  62.073  1.00 66.14           C
    ATOM   1758  HA  PRO B 471      28.194 -10.632  62.620  1.00  0.00           H
    ATOM   1759  HB2 PRO B 471      28.757 -12.358  60.175  1.00  0.00           H
    ATOM   1760  HB3 PRO B 471      29.911 -11.785  61.363  1.00  0.00           H
    ATOM   1761  HG2 PRO B 471      29.037 -14.278  61.619  1.00  0.00           H
    ATOM   1762  HG3 PRO B 471      28.989 -13.253  63.057  1.00  0.00           H
    ATOM   1763  HD2 PRO B 471      26.680 -13.846  61.167  1.00  0.00           H
    ATOM   1764  HD3 PRO B 471      26.725 -13.950  62.932  1.00  0.00           H
    ATOM   1765  N   GLN B 472      28.023  -8.848  60.868  1.00 66.57           N
    ATOM   1766  CA  GLN B 472      28.052  -7.757  59.892  1.00 67.15           C
    ATOM   1767  C   GLN B 472      29.428  -7.692  59.216  1.00 72.01           C
    ATOM   1768  O   GLN B 472      29.495  -7.459  58.010  1.00 77.98           O
    ATOM   1769  CB  GLN B 472      27.739  -6.421  60.600  1.00 62.71           C
    ATOM   1770  CG  GLN B 472      26.373  -6.366  61.312  1.00 59.29           C
    ATOM   1771  CD  GLN B 472      25.190  -6.665  60.399  1.00 58.48           C
    ATOM   1772  OE1 GLN B 472      24.770  -5.805  59.629  1.00 63.53           O
    ATOM   1773  NE2 GLN B 472      24.651  -7.883  60.480  1.00 57.54           N
    ATOM   1774  H   GLN B 472      28.526  -8.679  61.733  1.00  0.00           H
    ATOM   1775  HA  GLN B 472      27.313  -7.929  59.107  1.00  0.00           H
    ATOM   1776  HB2 GLN B 472      28.518  -6.197  61.330  1.00  0.00           H
    ATOM   1777  HB3 GLN B 472      27.780  -5.612  59.871  1.00  0.00           H
    ATOM   1778  HG2 GLN B 472      26.357  -7.038  62.171  1.00  0.00           H
    ATOM   1779  HG3 GLN B 472      26.212  -5.364  61.699  1.00  0.00           H
    ATOM   1780 HE21 GLN B 472      25.045  -8.577  61.101  1.00  0.00           H
    ATOM   1781 HE22 GLN B 472      23.883  -8.135  59.877  1.00  0.00           H
    ATOM   1782  N   ASN B 473      30.487  -7.922  60.011  1.00 70.49           N
    ATOM   1783  CA  ASN B 473      31.878  -8.011  59.569  1.00 71.59           C
    ATOM   1784  C   ASN B 473      32.362  -9.456  59.750  1.00 72.01           C
    ATOM   1785  O   ASN B 473      31.864 -10.173  60.619  1.00 71.94           O
    ATOM   1786  CB  ASN B 473      32.769  -7.041  60.378  1.00 72.88           C
    ATOM   1787  CG  ASN B 473      32.318  -5.582  60.277  1.00 71.71           C
    ATOM   1788  OD1 ASN B 473      32.660  -4.879  59.328  1.00 71.14           O
    ATOM   1789  ND2 ASN B 473      31.538  -5.134  61.258  1.00 71.45           N
    ATOM   1790  H   ASN B 473      30.330  -8.125  60.989  1.00  0.00           H
    ATOM   1791  HA  ASN B 473      31.967  -7.772  58.508  1.00  0.00           H
    ATOM   1792  HB2 ASN B 473      32.801  -7.335  61.428  1.00  0.00           H
    ATOM   1793  HB3 ASN B 473      33.798  -7.098  60.019  1.00  0.00           H
    ATOM   1794 HD21 ASN B 473      31.300  -5.740  62.036  1.00  0.00           H
    ATOM   1795 HD22 ASN B 473      31.192  -4.185  61.254  1.00  0.00           H
    ATOM   1796  N   ASP B 474      33.335  -9.851  58.916  1.00 73.85           N
    ATOM   1797  CA  ASP B 474      33.883 -11.212  58.816  1.00 76.29           C
    ATOM   1798  C   ASP B 474      34.584 -11.727  60.085  1.00 74.41           C
    ATOM   1799  O   ASP B 474      34.586 -12.936  60.314  1.00 78.84           O
    ATOM   1800  CB  ASP B 474      34.827 -11.370  57.601  1.00 81.48           C
    ATOM   1801  CG  ASP B 474      34.214 -10.906  56.276  1.00 85.98           C
    ATOM   1802  OD1 ASP B 474      33.100 -11.379  55.967  1.00 86.09           O
    ATOM   1803  OD2 ASP B 474      34.892 -10.124  55.574  1.00 89.35           O1-
    ATOM   1804  H   ASP B 474      33.704  -9.185  58.250  1.00  0.00           H
    ATOM   1805  HA  ASP B 474      33.028 -11.873  58.656  1.00  0.00           H
    ATOM   1806  HB2 ASP B 474      35.732 -10.787  57.780  1.00  0.00           H
    ATOM   1807  HB3 ASP B 474      35.153 -12.404  57.482  1.00  0.00           H
    ATOM   1808  N   ASP B 475      35.147 -10.814  60.889  1.00 67.00           N
    ATOM   1809  CA  ASP B 475      35.875 -11.099  62.130  1.00 62.67           C
    ATOM   1810  C   ASP B 475      34.947 -11.301  63.351  1.00 62.23           C
    ATOM   1811  O   ASP B 475      35.453 -11.296  64.471  1.00 62.01           O
    ATOM   1812  CB  ASP B 475      36.976 -10.042  62.428  1.00 63.53           C
    ATOM   1813  CG  ASP B 475      36.515  -8.579  62.499  1.00 64.47           C
    ATOM   1814  OD1 ASP B 475      35.420  -8.262  61.985  1.00 61.52           O
    ATOM   1815  OD2 ASP B 475      37.285  -7.775  63.070  1.00 66.94           O1-
    ATOM   1816  H   ASP B 475      35.070  -9.831  60.658  1.00  0.00           H
    ATOM   1817  HA  ASP B 475      36.393 -12.050  61.996  1.00  0.00           H
    ATOM   1818  HB2 ASP B 475      37.485 -10.289  63.361  1.00  0.00           H
    ATOM   1819  HB3 ASP B 475      37.731 -10.102  61.644  1.00  0.00           H
    ATOM   1820  N   GLU B 476      33.629 -11.466  63.141  1.00 62.05           N
    ATOM   1821  CA  GLU B 476      32.629 -11.635  64.196  1.00 63.15           C
    ATOM   1822  C   GLU B 476      32.126 -13.089  64.231  1.00 64.76           C
    ATOM   1823  O   GLU B 476      31.842 -13.661  63.177  1.00 63.07           O
    ATOM   1824  CB  GLU B 476      31.478 -10.633  63.966  1.00 61.48           C
    ATOM   1825  CG  GLU B 476      31.919  -9.151  64.052  1.00 57.03           C
    ATOM   1826  CD  GLU B 476      30.875  -8.126  63.605  1.00 51.89           C
    ATOM   1827  OE1 GLU B 476      29.792  -8.537  63.134  1.00 44.91           O
    ATOM   1828  OE2 GLU B 476      31.192  -6.921  63.720  1.00 55.51           O1-
    ATOM   1829  H   GLU B 476      33.274 -11.457  62.195  1.00  0.00           H
    ATOM   1830  HA  GLU B 476      33.074 -11.406  65.159  1.00  0.00           H
    ATOM   1831  HB2 GLU B 476      31.014 -10.830  63.002  1.00  0.00           H
    ATOM   1832  HB3 GLU B 476      30.708 -10.813  64.714  1.00  0.00           H
    ATOM   1833  HG2 GLU B 476      32.180  -8.914  65.080  1.00  0.00           H
    ATOM   1834  HG3 GLU B 476      32.812  -8.980  63.450  1.00  0.00           H
    ATOM   1835  N   LEU B 477      32.009 -13.655  65.447  1.00 65.74           N
    ATOM   1836  CA  LEU B 477      31.472 -14.998  65.700  1.00 61.06           C
    ATOM   1837  C   LEU B 477      29.947 -15.020  65.489  1.00 59.38           C
    ATOM   1838  O   LEU B 477      29.260 -14.200  66.103  1.00 60.21           O
    ATOM   1839  CB  LEU B 477      31.755 -15.411  67.162  1.00 60.37           C
    ATOM   1840  CG  LEU B 477      33.100 -16.120  67.372  1.00 62.73           C
    ATOM   1841  CD1 LEU B 477      33.412 -16.259  68.870  1.00 63.39           C
    ATOM   1842  CD2 LEU B 477      33.191 -17.472  66.635  1.00 62.69           C
    ATOM   1843  H   LEU B 477      32.220 -13.100  66.266  1.00  0.00           H
    ATOM   1844  HA  LEU B 477      31.998 -15.680  65.038  1.00  0.00           H
    ATOM   1845  HB2 LEU B 477      31.710 -14.523  67.788  1.00  0.00           H
    ATOM   1846  HB3 LEU B 477      30.975 -16.077  67.538  1.00  0.00           H
    ATOM   1847  HG  LEU B 477      33.858 -15.477  66.941  1.00  0.00           H
    ATOM   1848 HD11 LEU B 477      33.955 -17.171  69.109  1.00  0.00           H
    ATOM   1849 HD12 LEU B 477      34.040 -15.430  69.184  1.00  0.00           H
    ATOM   1850 HD13 LEU B 477      32.510 -16.240  69.482  1.00  0.00           H
    ATOM   1851 HD21 LEU B 477      33.542 -18.280  67.274  1.00  0.00           H
    ATOM   1852 HD22 LEU B 477      32.227 -17.783  66.232  1.00  0.00           H
    ATOM   1853 HD23 LEU B 477      33.887 -17.407  65.799  1.00  0.00           H
    ATOM   1854  N   PRO B 478      29.432 -15.983  64.689  1.00 59.81           N
    ATOM   1855  CA  PRO B 478      27.996 -16.303  64.678  1.00 64.32           C
    ATOM   1856  C   PRO B 478      27.546 -16.902  66.020  1.00 71.36           C
    ATOM   1857  O   PRO B 478      28.032 -17.972  66.388  1.00 74.62           O
    ATOM   1858  CB  PRO B 478      27.833 -17.311  63.518  1.00 63.27           C
    ATOM   1859  CG  PRO B 478      29.152 -17.306  62.764  1.00 63.01           C
    ATOM   1860  CD  PRO B 478      30.171 -16.887  63.808  1.00 61.57           C
    ATOM   1861  HA  PRO B 478      27.438 -15.394  64.460  1.00  0.00           H
    ATOM   1862  HB2 PRO B 478      27.642 -18.323  63.881  1.00  0.00           H
    ATOM   1863  HB3 PRO B 478      26.998 -17.051  62.867  1.00  0.00           H
    ATOM   1864  HG2 PRO B 478      29.385 -18.259  62.289  1.00  0.00           H
    ATOM   1865  HG3 PRO B 478      29.111 -16.550  61.979  1.00  0.00           H
    ATOM   1866  HD2 PRO B 478      30.520 -17.744  64.385  1.00  0.00           H
    ATOM   1867  HD3 PRO B 478      31.024 -16.424  63.311  1.00  0.00           H
    ATOM   1868  N   MET B 479      26.642 -16.200  66.718  1.00 71.14           N
    ATOM   1869  CA  MET B 479      26.003 -16.703  67.933  1.00 67.33           C
    ATOM   1870  C   MET B 479      24.566 -17.145  67.637  1.00 69.22           C
    ATOM   1871  O   MET B 479      23.952 -16.715  66.659  1.00 69.70           O
    ATOM   1872  CB  MET B 479      26.006 -15.642  69.052  1.00 66.89           C
    ATOM   1873  CG  MET B 479      27.385 -15.041  69.362  1.00 68.52           C
    ATOM   1874  SD  MET B 479      27.489 -14.213  70.973  1.00 71.38           S
    ATOM   1875  CE  MET B 479      26.196 -12.958  70.804  1.00 71.63           C
    ATOM   1876  H   MET B 479      26.273 -15.339  66.337  1.00  0.00           H
    ATOM   1877  HA  MET B 479      26.546 -17.571  68.312  1.00  0.00           H
    ATOM   1878  HB2 MET B 479      25.317 -14.839  68.793  1.00  0.00           H
    ATOM   1879  HB3 MET B 479      25.604 -16.083  69.966  1.00  0.00           H
    ATOM   1880  HG2 MET B 479      28.140 -15.827  69.351  1.00  0.00           H
    ATOM   1881  HG3 MET B 479      27.671 -14.329  68.586  1.00  0.00           H
    ATOM   1882  HE1 MET B 479      26.214 -12.296  71.666  1.00  0.00           H
    ATOM   1883  HE2 MET B 479      26.365 -12.360  69.911  1.00  0.00           H
    ATOM   1884  HE3 MET B 479      25.209 -13.417  70.743  1.00  0.00           H
    ATOM   1885  N   MET B 485      30.780 -20.113  75.426  1.00 72.66           N1+
    ATOM   1886  CA  MET B 485      32.022 -19.697  74.771  1.00 73.13           C
    ATOM   1887  C   MET B 485      32.964 -19.058  75.797  1.00 71.45           C
    ATOM   1888  O   MET B 485      32.491 -18.474  76.769  1.00 66.98           O
    ATOM   1889  CB  MET B 485      31.694 -18.755  73.588  1.00 75.23           C
    ATOM   1890  CG  MET B 485      31.055 -17.407  73.952  1.00 77.01           C
    ATOM   1891  SD  MET B 485      30.312 -16.527  72.555  1.00 77.51           S
    ATOM   1892  CE  MET B 485      28.807 -17.509  72.331  1.00 74.86           C
    ATOM   1893  H   MET B 485      30.524 -19.595  76.258  1.00  0.00           H
    ATOM   1894  HA  MET B 485      32.523 -20.586  74.383  1.00  0.00           H
    ATOM   1895  HB2 MET B 485      32.611 -18.538  73.045  1.00  0.00           H
    ATOM   1896  HB3 MET B 485      31.063 -19.282  72.877  1.00  0.00           H
    ATOM   1897  HG2 MET B 485      30.290 -17.531  74.719  1.00  0.00           H
    ATOM   1898  HG3 MET B 485      31.823 -16.754  74.362  1.00  0.00           H
    ATOM   1899  HE1 MET B 485      28.220 -17.124  71.499  1.00  0.00           H
    ATOM   1900  HE2 MET B 485      29.048 -18.550  72.120  1.00  0.00           H
    ATOM   1901  HE3 MET B 485      28.197 -17.468  73.232  1.00  0.00           H
    ATOM   1902  N   ILE B 487      35.748 -16.189  76.454  1.00 71.96           N1+
    ATOM   1903  CA  ILE B 487      36.292 -14.903  76.036  1.00 69.18           C
    ATOM   1904  C   ILE B 487      37.834 -14.927  76.090  1.00 76.88           C
    ATOM   1905  O   ILE B 487      38.406 -14.979  77.179  1.00 81.82           O
    ATOM   1906  CB  ILE B 487      35.708 -13.748  76.893  1.00 59.41           C
    ATOM   1907  CG1 ILE B 487      34.170 -13.654  76.794  1.00 55.59           C
    ATOM   1908  CG2 ILE B 487      36.345 -12.368  76.639  1.00 54.60           C
    ATOM   1909  CD1 ILE B 487      33.632 -13.546  75.364  1.00 52.36           C
    ATOM   1910  H   ILE B 487      35.314 -16.210  77.368  1.00  0.00           H
    ATOM   1911  HA  ILE B 487      35.984 -14.720  75.018  1.00  0.00           H
    ATOM   1912  HB  ILE B 487      35.904 -14.021  77.920  1.00  0.00           H
    ATOM   1913 HG12 ILE B 487      33.733 -14.540  77.257  1.00  0.00           H
    ATOM   1914 HG13 ILE B 487      33.801 -12.815  77.382  1.00  0.00           H
    ATOM   1915 HG21 ILE B 487      35.838 -11.593  77.214  1.00  0.00           H
    ATOM   1916 HG22 ILE B 487      37.391 -12.347  76.942  1.00  0.00           H
    ATOM   1917 HG23 ILE B 487      36.292 -12.091  75.588  1.00  0.00           H
    ATOM   1918 HD11 ILE B 487      32.707 -12.977  75.317  1.00  0.00           H
    ATOM   1919 HD12 ILE B 487      34.343 -13.057  74.699  1.00  0.00           H
    ATOM   1920 HD13 ILE B 487      33.421 -14.540  74.970  1.00  0.00           H
    ATOM   1921  N   HID B 489      40.248 -12.845  76.034  1.00 86.84           N1+
    ATOM   1922  CA  HID B 489      40.360 -11.432  76.414  1.00 89.21           C
    ATOM   1923  C   HID B 489      39.388 -10.487  75.665  1.00 90.00           C
    ATOM   1924  O   HID B 489      38.563 -10.919  74.862  1.00 90.00           O
    ATOM   1925  CB  HID B 489      41.839 -10.968  76.432  1.00 90.00           C
    ATOM   1926  CG  HID B 489      42.498 -10.667  75.107  1.00 90.00           C
    ATOM   1927  ND1 HID B 489      42.242  -9.518  74.375  1.00 90.00           N
    ATOM   1928  CD2 HID B 489      43.437 -11.366  74.380  1.00 90.00           C
    ATOM   1929  CE1 HID B 489      42.981  -9.585  73.267  1.00 90.00           C
    ATOM   1930  NE2 HID B 489      43.732 -10.679  73.201  1.00 90.00           N
    ATOM   1931  H   HID B 489      39.945 -13.457  76.779  1.00  0.00           H
    ATOM   1932  HA  HID B 489      40.021 -11.418  77.451  1.00  0.00           H
    ATOM   1933  HB2 HID B 489      41.929 -10.073  77.047  1.00  0.00           H
    ATOM   1934  HB3 HID B 489      42.445 -11.718  76.940  1.00  0.00           H
    ATOM   1935  HD1 HID B 489      41.610  -8.766  74.622  1.00  0.00           H
    ATOM   1936  HD2 HID B 489      43.905 -12.312  74.604  1.00  0.00           H
    ATOM   1937  HE1 HID B 489      42.974  -8.820  72.504  1.00  0.00           H
    ATOM   1938  N   ARG B 490      39.449  -9.200  76.015  1.00 90.00           N
    ATOM   1939  CA  ARG B 490      38.469  -8.192  75.617  1.00 89.35           C
    ATOM   1940  C   ARG B 490      39.242  -6.872  75.485  1.00 90.00           C
    ATOM   1941  O   ARG B 490      40.163  -6.827  74.671  1.00 90.00           O
    ATOM   1942  CB  ARG B 490      37.233  -8.218  76.580  1.00 87.91           C
    ATOM   1943  CG  ARG B 490      37.420  -8.970  77.921  1.00 84.24           C
    ATOM   1944  CD  ARG B 490      36.350  -8.710  78.990  1.00 82.18           C
    ATOM   1945  NE  ARG B 490      36.552  -7.414  79.650  1.00 83.94           N
    ATOM   1946  CZ  ARG B 490      35.793  -6.897  80.633  1.00 87.18           C
    ATOM   1947  NH1 ARG B 490      34.729  -7.541  81.126  1.00 87.14           N
    ATOM   1948  NH2 ARG B 490      36.119  -5.708  81.150  1.00 89.19           N1+
    ATOM   1949  H   ARG B 490      40.157  -8.883  76.667  1.00  0.00           H
    ATOM   1950  HA  ARG B 490      38.113  -8.418  74.610  1.00  0.00           H
    ATOM   1951  HB2 ARG B 490      36.847  -7.217  76.763  1.00  0.00           H
    ATOM   1952  HB3 ARG B 490      36.424  -8.731  76.059  1.00  0.00           H
    ATOM   1953  HG2 ARG B 490      37.573 -10.039  77.791  1.00  0.00           H
    ATOM   1954  HG3 ARG B 490      38.352  -8.602  78.342  1.00  0.00           H
    ATOM   1955  HD2 ARG B 490      35.383  -8.627  78.500  1.00  0.00           H
    ATOM   1956  HD3 ARG B 490      36.275  -9.542  79.690  1.00  0.00           H
    ATOM   1957  HE  ARG B 490      37.399  -6.925  79.390  1.00  0.00           H
    ATOM   1958 HH11 ARG B 490      34.519  -8.485  80.824  1.00  0.00           H
    ATOM   1959 HH12 ARG B 490      34.170  -7.124  81.857  1.00  0.00           H
    ATOM   1960 HH21 ARG B 490      36.934  -5.214  80.813  1.00  0.00           H
    ATOM   1961 HH22 ARG B 490      35.580  -5.314  81.908  1.00  0.00           H
    ATOM   1962  N   GLU B 491      38.901  -5.864  76.305  1.00 90.00           N
    ATOM   1963  CA  GLU B 491      39.616  -4.598  76.506  1.00 90.00           C
    ATOM   1964  C   GLU B 491      39.454  -3.571  75.363  1.00 90.00           C
    ATOM   1965  O   GLU B 491      38.829  -3.845  74.341  1.00 90.00           O
    ATOM   1966  CB  GLU B 491      41.096  -4.809  76.951  1.00 90.00           C
    ATOM   1967  CG  GLU B 491      41.297  -5.343  78.387  1.00 90.00           C
    ATOM   1968  CD  GLU B 491      40.849  -6.789  78.602  1.00 90.00           C
    ATOM   1969  OE1 GLU B 491      41.425  -7.676  77.933  1.00 90.00           O
    ATOM   1970  OE2 GLU B 491      39.920  -6.982  79.416  1.00 90.00           O1-
    ATOM   1971  H   GLU B 491      38.122  -6.007  76.929  1.00  0.00           H
    ATOM   1972  HA  GLU B 491      39.104  -4.131  77.348  1.00  0.00           H
    ATOM   1973  HB2 GLU B 491      41.613  -5.476  76.262  1.00  0.00           H
    ATOM   1974  HB3 GLU B 491      41.653  -3.875  76.876  1.00  0.00           H
    ATOM   1975  HG2 GLU B 491      42.356  -5.293  78.640  1.00  0.00           H
    ATOM   1976  HG3 GLU B 491      40.787  -4.693  79.098  1.00  0.00           H
    ATOM   1977  N   ASP B 492      39.970  -2.364  75.639  1.00 89.87           N
    ATOM   1978  CA  ASP B 492      39.791  -1.098  74.922  1.00 90.00           C
    ATOM   1979  C   ASP B 492      40.415  -1.002  73.517  1.00 89.58           C
    ATOM   1980  O   ASP B 492      39.879  -0.246  72.703  1.00 88.24           O
    ATOM   1981  CB  ASP B 492      40.289   0.114  75.763  1.00 90.00           C
    ATOM   1982  CG  ASP B 492      41.664  -0.027  76.450  1.00 90.00           C
    ATOM   1983  OD1 ASP B 492      42.388  -1.011  76.182  1.00 90.00           O
    ATOM   1984  OD2 ASP B 492      41.986   0.897  77.228  1.00 90.00           O1-
    ATOM   1985  H   ASP B 492      40.528  -2.277  76.477  1.00  0.00           H
    ATOM   1986  HA  ASP B 492      38.714  -0.982  74.787  1.00  0.00           H
    ATOM   1987  HB2 ASP B 492      40.312   1.012  75.145  1.00  0.00           H
    ATOM   1988  HB3 ASP B 492      39.553   0.289  76.548  1.00  0.00           H
    ATOM   1989  N   GLU B 493      41.553  -1.688  73.293  1.00 90.00           N
    ATOM   1990  CA  GLU B 493      42.465  -1.553  72.143  1.00 90.00           C
    ATOM   1991  C   GLU B 493      41.770  -1.538  70.773  1.00 90.00           C
    ATOM   1992  O   GLU B 493      42.076  -0.683  69.941  1.00 90.00           O
    ATOM   1993  CB  GLU B 493      43.585  -2.626  72.218  1.00 90.00           C
    ATOM   1994  CG  GLU B 493      43.236  -4.085  71.816  1.00 89.94           C
    ATOM   1995  CD  GLU B 493      42.135  -4.722  72.667  1.00 90.00           C
    ATOM   1996  OE1 GLU B 493      42.266  -4.638  73.907  1.00 90.00           O
    ATOM   1997  OE2 GLU B 493      41.179  -5.253  72.063  1.00 90.00           O1-
    ATOM   1998  H   GLU B 493      41.887  -2.298  74.029  1.00  0.00           H
    ATOM   1999  HA  GLU B 493      42.939  -0.578  72.272  1.00  0.00           H
    ATOM   2000  HB2 GLU B 493      44.409  -2.296  71.583  1.00  0.00           H
    ATOM   2001  HB3 GLU B 493      44.001  -2.630  73.227  1.00  0.00           H
    ATOM   2002  HG2 GLU B 493      42.964  -4.142  70.762  1.00  0.00           H
    ATOM   2003  HG3 GLU B 493      44.126  -4.705  71.921  1.00  0.00           H
    ATOM   2004  N   ASP B 494      40.809  -2.453  70.617  1.00 90.00           N
    ATOM   2005  CA  ASP B 494      39.882  -2.517  69.513  1.00 90.00           C
    ATOM   2006  C   ASP B 494      38.565  -2.955  70.160  1.00 89.40           C
    ATOM   2007  O   ASP B 494      38.393  -4.133  70.473  1.00 87.47           O
    ATOM   2008  CB  ASP B 494      40.380  -3.474  68.404  1.00 90.00           C
    ATOM   2009  CG  ASP B 494      39.623  -3.325  67.080  1.00 90.00           C
    ATOM   2010  OD1 ASP B 494      39.632  -2.194  66.549  1.00 90.00           O
    ATOM   2011  OD2 ASP B 494      39.119  -4.355  66.579  1.00 90.00           O1-
    ATOM   2012  H   ASP B 494      40.646  -3.130  71.352  1.00  0.00           H
    ATOM   2013  HA  ASP B 494      39.750  -1.516  69.096  1.00  0.00           H
    ATOM   2014  HB2 ASP B 494      41.439  -3.314  68.202  1.00  0.00           H
    ATOM   2015  HB3 ASP B 494      40.321  -4.504  68.750  1.00  0.00           H
    ATOM   2016  N   GLU B 495      37.666  -1.972  70.333  1.00 89.62           N
    ATOM   2017  CA  GLU B 495      36.273  -2.101  70.764  1.00 90.00           C
    ATOM   2018  C   GLU B 495      36.093  -2.363  72.270  1.00 90.00           C
    ATOM   2019  O   GLU B 495      36.467  -3.428  72.755  1.00 90.00           O
    ATOM   2020  CB  GLU B 495      35.519  -3.185  69.955  1.00 90.00           C
    ATOM   2021  CG  GLU B 495      35.799  -3.286  68.441  1.00 88.93           C
    ATOM   2022  CD  GLU B 495      35.660  -1.983  67.646  1.00 90.00           C
    ATOM   2023  OE1 GLU B 495      34.964  -1.063  68.127  1.00 90.00           O
    ATOM   2024  OE2 GLU B 495      36.215  -1.954  66.527  1.00 90.00           O1-
    ATOM   2025  H   GLU B 495      37.934  -1.040  70.057  1.00  0.00           H
    ATOM   2026  HA  GLU B 495      35.820  -1.135  70.543  1.00  0.00           H
    ATOM   2027  HB2 GLU B 495      35.717  -4.168  70.384  1.00  0.00           H
    ATOM   2028  HB3 GLU B 495      34.456  -3.018  70.098  1.00  0.00           H
    ATOM   2029  HG2 GLU B 495      36.794  -3.688  68.259  1.00  0.00           H
    ATOM   2030  HG3 GLU B 495      35.130  -4.038  68.032  1.00  0.00           H
    ATOM   2031  N   ILE B 496      35.421  -1.425  72.957  1.00 88.03           N
    ATOM   2032  CA  ILE B 496      34.931  -1.603  74.328  1.00 82.53           C
    ATOM   2033  C   ILE B 496      33.614  -2.412  74.381  1.00 75.02           C
    ATOM   2034  O   ILE B 496      33.408  -3.160  75.336  1.00 71.58           O
    ATOM   2035  CB  ILE B 496      34.718  -0.230  75.040  1.00 84.40           C
    ATOM   2036  CG1 ILE B 496      34.361  -0.382  76.539  1.00 85.92           C
    ATOM   2037  CG2 ILE B 496      33.695   0.693  74.340  1.00 86.40           C
    ATOM   2038  CD1 ILE B 496      34.422   0.929  77.333  1.00 86.09           C
    ATOM   2039  H   ILE B 496      35.145  -0.572  72.493  1.00  0.00           H
    ATOM   2040  HA  ILE B 496      35.684  -2.158  74.891  1.00  0.00           H
    ATOM   2041  HB  ILE B 496      35.683   0.277  74.994  1.00  0.00           H
    ATOM   2042 HG12 ILE B 496      33.357  -0.791  76.644  1.00  0.00           H
    ATOM   2043 HG13 ILE B 496      35.031  -1.106  77.003  1.00  0.00           H
    ATOM   2044 HG21 ILE B 496      33.750   1.707  74.734  1.00  0.00           H
    ATOM   2045 HG22 ILE B 496      33.871   0.763  73.267  1.00  0.00           H
    ATOM   2046 HG23 ILE B 496      32.674   0.349  74.497  1.00  0.00           H
    ATOM   2047 HD11 ILE B 496      34.560   0.731  78.397  1.00  0.00           H
    ATOM   2048 HD12 ILE B 496      35.251   1.558  77.006  1.00  0.00           H
    ATOM   2049 HD13 ILE B 496      33.501   1.503  77.222  1.00  0.00           H
    ATOM   2050  N   GLU B 497      32.755  -2.251  73.360  1.00 71.76           N
    ATOM   2051  CA  GLU B 497      31.427  -2.863  73.285  1.00 67.99           C
    ATOM   2052  C   GLU B 497      31.453  -4.328  72.798  1.00 62.34           C
    ATOM   2053  O   GLU B 497      30.399  -4.965  72.809  1.00 63.33           O
    ATOM   2054  CB  GLU B 497      30.512  -2.010  72.374  1.00 72.65           C
    ATOM   2055  CG  GLU B 497      30.459  -0.503  72.700  1.00 76.70           C
    ATOM   2056  CD  GLU B 497      29.548   0.304  71.767  1.00 75.50           C
    ATOM   2057  OE1 GLU B 497      29.081  -0.260  70.752  1.00 71.81           O
    ATOM   2058  OE2 GLU B 497      29.360   1.501  72.075  1.00 74.35           O1-
    ATOM   2059  H   GLU B 497      32.999  -1.630  72.603  1.00  0.00           H
    ATOM   2060  HA  GLU B 497      30.992  -2.861  74.283  1.00  0.00           H
    ATOM   2061  HB2 GLU B 497      30.824  -2.119  71.339  1.00  0.00           H
    ATOM   2062  HB3 GLU B 497      29.497  -2.407  72.422  1.00  0.00           H
    ATOM   2063  HG2 GLU B 497      30.133  -0.353  73.730  1.00  0.00           H
    ATOM   2064  HG3 GLU B 497      31.452  -0.066  72.611  1.00  0.00           H
    ATOM   2065  N   TRP B 498      32.620  -4.823  72.346  1.00 57.69           N
    ATOM   2066  CA  TRP B 498      32.762  -6.112  71.673  1.00 53.91           C
    ATOM   2067  C   TRP B 498      33.924  -6.898  72.293  1.00 62.09           C
    ATOM   2068  O   TRP B 498      35.077  -6.488  72.157  1.00 64.86           O
    ATOM   2069  CB  TRP B 498      32.956  -5.898  70.153  1.00 45.66           C
    ATOM   2070  CG  TRP B 498      31.892  -5.104  69.450  1.00 43.11           C
    ATOM   2071  CD1 TRP B 498      31.737  -3.760  69.497  1.00 42.00           C
    ATOM   2072  CD2 TRP B 498      30.797  -5.594  68.622  1.00 40.71           C
    ATOM   2073  NE1 TRP B 498      30.607  -3.392  68.803  1.00 40.68           N
    ATOM   2074  CE2 TRP B 498      29.986  -4.481  68.238  1.00 40.85           C
    ATOM   2075  CE3 TRP B 498      30.411  -6.865  68.141  1.00 37.33           C
    ATOM   2076  CZ2 TRP B 498      28.833  -4.625  67.453  1.00 40.70           C
    ATOM   2077  CZ3 TRP B 498      29.265  -7.018  67.339  1.00 36.89           C
    ATOM   2078  CH2 TRP B 498      28.461  -5.908  67.020  1.00 39.09           C
    ATOM   2079  H   TRP B 498      33.455  -4.256  72.392  1.00  0.00           H
    ATOM   2080  HA  TRP B 498      31.851  -6.695  71.805  1.00  0.00           H
    ATOM   2081  HB2 TRP B 498      33.910  -5.417  69.951  1.00  0.00           H
    ATOM   2082  HB3 TRP B 498      33.015  -6.871  69.664  1.00  0.00           H
    ATOM   2083  HD1 TRP B 498      32.374  -3.084  70.041  1.00  0.00           H
    ATOM   2084  HE1 TRP B 498      30.282  -2.435  68.732  1.00  0.00           H
    ATOM   2085  HE3 TRP B 498      31.003  -7.733  68.390  1.00  0.00           H
    ATOM   2086  HZ2 TRP B 498      28.240  -3.763  67.188  1.00  0.00           H
    ATOM   2087  HZ3 TRP B 498      28.991  -7.997  66.978  1.00  0.00           H
    ATOM   2088  HH2 TRP B 498      27.563  -6.044  66.439  1.00  0.00           H
    ATOM   2089  N   TRP B 499      33.592  -8.026  72.944  1.00 64.63           N
    ATOM   2090  CA  TRP B 499      34.557  -9.010  73.438  1.00 62.60           C
    ATOM   2091  C   TRP B 499      35.076  -9.878  72.281  1.00 61.52           C
    ATOM   2092  O   TRP B 499      34.300 -10.194  71.384  1.00 57.51           O
    ATOM   2093  CB  TRP B 499      33.895  -9.938  74.481  1.00 65.52           C
    ATOM   2094  CG  TRP B 499      33.304  -9.387  75.750  1.00 68.87           C
    ATOM   2095  CD1 TRP B 499      32.548 -10.127  76.593  1.00 68.78           C
    ATOM   2096  CD2 TRP B 499      33.470  -8.092  76.415  1.00 68.95           C
    ATOM   2097  NE1 TRP B 499      32.221  -9.397  77.711  1.00 68.08           N
    ATOM   2098  CE2 TRP B 499      32.773  -8.137  77.665  1.00 69.93           C
    ATOM   2099  CE3 TRP B 499      34.149  -6.887  76.115  1.00 69.31           C
    ATOM   2100  CZ2 TRP B 499      32.759  -7.059  78.564  1.00 71.53           C
    ATOM   2101  CZ3 TRP B 499      34.170  -5.810  77.025  1.00 71.80           C
    ATOM   2102  CH2 TRP B 499      33.480  -5.894  78.248  1.00 72.66           C
    ATOM   2103  H   TRP B 499      32.620  -8.303  72.983  1.00  0.00           H
    ATOM   2104  HA  TRP B 499      35.404  -8.495  73.894  1.00  0.00           H
    ATOM   2105  HB2 TRP B 499      33.105 -10.510  73.995  1.00  0.00           H
    ATOM   2106  HB3 TRP B 499      34.635 -10.673  74.798  1.00  0.00           H
    ATOM   2107  HD1 TRP B 499      32.273 -11.152  76.418  1.00  0.00           H
    ATOM   2108  HE1 TRP B 499      31.667  -9.773  78.467  1.00  0.00           H
    ATOM   2109  HE3 TRP B 499      34.695  -6.786  75.191  1.00  0.00           H
    ATOM   2110  HZ2 TRP B 499      32.225  -7.133  79.500  1.00  0.00           H
    ATOM   2111  HZ3 TRP B 499      34.725  -4.916  76.781  1.00  0.00           H
    ATOM   2112  HH2 TRP B 499      33.506  -5.068  78.944  1.00  0.00           H
    ATOM   2113  N   TRP B 500      36.347 -10.304  72.358  1.00 63.77           N
    ATOM   2114  CA  TRP B 500      36.920 -11.302  71.455  1.00 65.18           C
    ATOM   2115  C   TRP B 500      36.772 -12.684  72.112  1.00 66.11           C
    ATOM   2116  O   TRP B 500      37.343 -12.895  73.178  1.00 66.36           O
    ATOM   2117  CB  TRP B 500      38.389 -10.937  71.162  1.00 68.93           C
    ATOM   2118  CG  TRP B 500      39.085 -11.832  70.179  1.00 73.67           C
    ATOM   2119  CD1 TRP B 500      39.910 -12.854  70.487  1.00 74.90           C
    ATOM   2120  CD2 TRP B 500      38.978 -11.844  68.725  1.00 73.22           C
    ATOM   2121  NE1 TRP B 500      40.284 -13.526  69.344  1.00 75.21           N
    ATOM   2122  CE2 TRP B 500      39.748 -12.937  68.220  1.00 74.69           C
    ATOM   2123  CE3 TRP B 500      38.311 -11.038  67.776  1.00 69.03           C
    ATOM   2124  CZ2 TRP B 500      39.846 -13.217  66.847  1.00 73.47           C
    ATOM   2125  CZ3 TRP B 500      38.433 -11.285  66.394  1.00 67.71           C
    ATOM   2126  CH2 TRP B 500      39.193 -12.376  65.928  1.00 70.59           C
    ATOM   2127  H   TRP B 500      36.930 -10.027  73.136  1.00  0.00           H
    ATOM   2128  HA  TRP B 500      36.386 -11.300  70.503  1.00  0.00           H
    ATOM   2129  HB2 TRP B 500      38.432  -9.924  70.759  1.00  0.00           H
    ATOM   2130  HB3 TRP B 500      38.971 -10.913  72.085  1.00  0.00           H
    ATOM   2131  HD1 TRP B 500      40.221 -13.093  71.484  1.00  0.00           H
    ATOM   2132  HE1 TRP B 500      40.890 -14.336  69.355  1.00  0.00           H
    ATOM   2133  HE3 TRP B 500      37.715 -10.210  68.124  1.00  0.00           H
    ATOM   2134  HZ2 TRP B 500      40.428 -14.059  66.502  1.00  0.00           H
    ATOM   2135  HZ3 TRP B 500      37.926 -10.644  65.689  1.00  0.00           H
    ATOM   2136  HH2 TRP B 500      39.267 -12.569  64.868  1.00  0.00           H
    ATOM   2137  N   ALA B 501      35.991 -13.584  71.497  1.00 66.34           N
    ATOM   2138  CA  ALA B 501      35.621 -14.877  72.074  1.00 69.80           C
    ATOM   2139  C   ALA B 501      36.096 -16.066  71.242  1.00 73.65           C
    ATOM   2140  O   ALA B 501      36.495 -15.887  70.094  1.00 72.93           O
    ATOM   2141  CB  ALA B 501      34.094 -14.915  72.255  1.00 68.89           C
    ATOM   2142  H   ALA B 501      35.582 -13.365  70.595  1.00  0.00           H
    ATOM   2143  HA  ALA B 501      36.101 -15.008  73.032  1.00  0.00           H
    ATOM   2144  HB1 ALA B 501      33.782 -15.766  72.862  1.00  0.00           H
    ATOM   2145  HB2 ALA B 501      33.737 -14.009  72.735  1.00  0.00           H
    ATOM   2146  HB3 ALA B 501      33.568 -14.974  71.306  1.00  0.00           H
    ATOM   2147  N   GLU B 507      36.996 -18.441  66.470  1.00 86.37           N1+
    ATOM   2148  CA  GLU B 507      37.199 -17.202  67.226  1.00 86.45           C
    ATOM   2149  C   GLU B 507      36.689 -15.984  66.441  1.00 84.62           C
    ATOM   2150  O   GLU B 507      36.552 -16.055  65.218  1.00 84.24           O
    ATOM   2151  CB  GLU B 507      38.686 -17.050  67.615  1.00 87.96           C
    ATOM   2152  CG  GLU B 507      39.170 -18.155  68.573  1.00 88.82           C
    ATOM   2153  CD  GLU B 507      40.613 -18.009  69.058  1.00 90.00           C
    ATOM   2154  OE1 GLU B 507      41.357 -17.190  68.475  1.00 90.00           O
    ATOM   2155  OE2 GLU B 507      40.950 -18.739  70.014  1.00 90.00           O1-
    ATOM   2156  H   GLU B 507      37.265 -18.409  65.497  1.00  0.00           H
    ATOM   2157  HA  GLU B 507      36.594 -17.248  68.127  1.00  0.00           H
    ATOM   2158  HB2 GLU B 507      39.300 -17.051  66.713  1.00  0.00           H
    ATOM   2159  HB3 GLU B 507      38.842 -16.077  68.087  1.00  0.00           H
    ATOM   2160  HG2 GLU B 507      38.525 -18.147  69.446  1.00  0.00           H
    ATOM   2161  HG3 GLU B 507      39.075 -19.137  68.112  1.00  0.00           H
    ATOM   2162  N   GLY B 508      36.401 -14.889  67.164  1.00 81.95           N
    ATOM   2163  CA  GLY B 508      35.835 -13.682  66.568  1.00 77.72           C
    ATOM   2164  C   GLY B 508      35.204 -12.781  67.638  1.00 72.67           C
    ATOM   2165  O   GLY B 508      35.109 -13.138  68.811  1.00 70.78           O
    ATOM   2166  H   GLY B 508      36.525 -14.897  68.169  1.00  0.00           H
    ATOM   2167  HA2 GLY B 508      36.612 -13.147  66.031  1.00  0.00           H
    ATOM   2168  HA3 GLY B 508      35.070 -13.931  65.831  1.00  0.00           H
    ATOM   2169  N   TYR B 509      34.764 -11.593  67.199  1.00 68.61           N
    ATOM   2170  CA  TYR B 509      34.064 -10.590  67.997  1.00 61.56           C
    ATOM   2171  C   TYR B 509      32.601 -10.963  68.266  1.00 57.73           C
    ATOM   2172  O   TYR B 509      31.893 -11.397  67.359  1.00 58.61           O
    ATOM   2173  CB  TYR B 509      34.152  -9.221  67.293  1.00 57.57           C
    ATOM   2174  CG  TYR B 509      35.464  -8.506  67.522  1.00 53.71           C
    ATOM   2175  CD1 TYR B 509      35.775  -8.023  68.810  1.00 48.92           C
    ATOM   2176  CD2 TYR B 509      36.384  -8.340  66.467  1.00 55.32           C
    ATOM   2177  CE1 TYR B 509      37.016  -7.410  69.053  1.00 50.27           C
    ATOM   2178  CE2 TYR B 509      37.618  -7.706  66.704  1.00 55.26           C
    ATOM   2179  CZ  TYR B 509      37.941  -7.264  68.002  1.00 51.88           C
    ATOM   2180  OH  TYR B 509      39.153  -6.701  68.250  1.00 51.17           O
    ATOM   2181  H   TYR B 509      34.903 -11.358  66.223  1.00  0.00           H
    ATOM   2182  HA  TYR B 509      34.589 -10.525  68.948  1.00  0.00           H
    ATOM   2183  HB2 TYR B 509      33.975  -9.330  66.225  1.00  0.00           H
    ATOM   2184  HB3 TYR B 509      33.375  -8.551  67.666  1.00  0.00           H
    ATOM   2185  HD1 TYR B 509      35.074  -8.145  69.622  1.00  0.00           H
    ATOM   2186  HD2 TYR B 509      36.155  -8.711  65.480  1.00  0.00           H
    ATOM   2187  HE1 TYR B 509      37.260  -7.058  70.046  1.00  0.00           H
    ATOM   2188  HE2 TYR B 509      38.325  -7.588  65.895  1.00  0.00           H
    ATOM   2189  HH  TYR B 509      39.517  -6.257  67.478  1.00  0.00           H
    ATOM   2190  N   VAL B 510      32.174 -10.732  69.512  1.00 49.65           N
    ATOM   2191  CA  VAL B 510      30.801 -10.895  69.973  1.00 45.92           C
    ATOM   2192  C   VAL B 510      30.402  -9.632  70.764  1.00 44.46           C
    ATOM   2193  O   VAL B 510      31.215  -9.138  71.547  1.00 45.58           O
    ATOM   2194  CB  VAL B 510      30.655 -12.136  70.903  1.00 40.68           C
    ATOM   2195  CG1 VAL B 510      30.877 -13.437  70.123  1.00 40.25           C
    ATOM   2196  CG2 VAL B 510      31.510 -12.138  72.185  1.00 36.18           C
    ATOM   2197  H   VAL B 510      32.827 -10.383  70.203  1.00  0.00           H
    ATOM   2198  HA  VAL B 510      30.156 -11.029  69.106  1.00  0.00           H
    ATOM   2199  HB  VAL B 510      29.615 -12.157  71.232  1.00  0.00           H
    ATOM   2200 HG11 VAL B 510      30.652 -14.308  70.735  1.00  0.00           H
    ATOM   2201 HG12 VAL B 510      30.235 -13.482  69.243  1.00  0.00           H
    ATOM   2202 HG13 VAL B 510      31.914 -13.527  69.800  1.00  0.00           H
    ATOM   2203 HG21 VAL B 510      31.366 -13.062  72.747  1.00  0.00           H
    ATOM   2204 HG22 VAL B 510      32.572 -12.060  71.956  1.00  0.00           H
    ATOM   2205 HG23 VAL B 510      31.245 -11.325  72.857  1.00  0.00           H
    ATOM   2206  N   PRO B 511      29.176  -9.107  70.545  1.00 42.00           N
    ATOM   2207  CA  PRO B 511      28.687  -7.947  71.305  1.00 40.56           C
    ATOM   2208  C   PRO B 511      28.391  -8.324  72.767  1.00 43.26           C
    ATOM   2209  O   PRO B 511      27.579  -9.218  73.002  1.00 46.45           O
    ATOM   2210  CB  PRO B 511      27.416  -7.542  70.548  1.00 36.88           C
    ATOM   2211  CG  PRO B 511      26.895  -8.827  69.924  1.00 37.95           C
    ATOM   2212  CD  PRO B 511      28.159  -9.619  69.621  1.00 39.52           C
    ATOM   2213  HA  PRO B 511      29.408  -7.128  71.271  1.00  0.00           H
    ATOM   2214  HB2 PRO B 511      26.675  -7.062  71.189  1.00  0.00           H
    ATOM   2215  HB3 PRO B 511      27.673  -6.830  69.764  1.00  0.00           H
    ATOM   2216  HG2 PRO B 511      26.302  -9.374  70.657  1.00  0.00           H
    ATOM   2217  HG3 PRO B 511      26.270  -8.657  69.048  1.00  0.00           H
    ATOM   2218  HD2 PRO B 511      27.984 -10.687  69.742  1.00  0.00           H
    ATOM   2219  HD3 PRO B 511      28.482  -9.445  68.596  1.00  0.00           H
    ATOM   2220  N   ARG B 512      29.043  -7.632  73.718  1.00 44.94           N
    ATOM   2221  CA  ARG B 512      28.859  -7.831  75.164  1.00 53.72           C
    ATOM   2222  C   ARG B 512      27.421  -7.562  75.658  1.00 59.55           C
    ATOM   2223  O   ARG B 512      27.050  -8.037  76.730  1.00 63.30           O
    ATOM   2224  CB  ARG B 512      29.881  -6.963  75.931  1.00 52.71           C
    ATOM   2225  CG  ARG B 512      29.487  -5.480  76.096  1.00 49.31           C
    ATOM   2226  CD  ARG B 512      30.544  -4.666  76.833  1.00 45.61           C
    ATOM   2227  NE  ARG B 512      30.196  -3.242  76.840  1.00 46.44           N
    ATOM   2228  CZ  ARG B 512      30.836  -2.289  77.535  1.00 51.15           C
    ATOM   2229  NH1 ARG B 512      31.857  -2.592  78.349  1.00 53.73           N
    ATOM   2230  NH2 ARG B 512      30.451  -1.014  77.405  1.00 52.20           N1+
    ATOM   2231  H   ARG B 512      29.693  -6.906  73.447  1.00  0.00           H
    ATOM   2232  HA  ARG B 512      29.084  -8.877  75.376  1.00  0.00           H
    ATOM   2233  HB2 ARG B 512      30.019  -7.386  76.927  1.00  0.00           H
    ATOM   2234  HB3 ARG B 512      30.853  -7.037  75.440  1.00  0.00           H
    ATOM   2235  HG2 ARG B 512      29.235  -5.012  75.144  1.00  0.00           H
    ATOM   2236  HG3 ARG B 512      28.584  -5.431  76.705  1.00  0.00           H
    ATOM   2237  HD2 ARG B 512      30.542  -4.975  77.878  1.00  0.00           H
    ATOM   2238  HD3 ARG B 512      31.543  -4.846  76.439  1.00  0.00           H
    ATOM   2239  HE  ARG B 512      29.452  -2.965  76.216  1.00  0.00           H
    ATOM   2240 HH11 ARG B 512      32.165  -3.550  78.429  1.00  0.00           H
    ATOM   2241 HH12 ARG B 512      32.337  -1.870  78.866  1.00  0.00           H
    ATOM   2242 HH21 ARG B 512      29.689  -0.769  76.790  1.00  0.00           H
    ATOM   2243 HH22 ARG B 512      30.922  -0.283  77.919  1.00  0.00           H
    ATOM   2244  N   ASN B 513      26.659  -6.783  74.871  1.00 59.86           N
    ATOM   2245  CA  ASN B 513      25.273  -6.383  75.118  1.00 64.15           C
    ATOM   2246  C   ASN B 513      24.302  -7.573  75.119  1.00 64.64           C
    ATOM   2247  O   ASN B 513      23.288  -7.516  75.816  1.00 66.02           O
    ATOM   2248  CB  ASN B 513      24.823  -5.343  74.064  1.00 71.36           C
    ATOM   2249  CG  ASN B 513      25.723  -4.106  73.923  1.00 79.80           C
    ATOM   2250  OD1 ASN B 513      25.814  -3.540  72.838  1.00 83.81           O
    ATOM   2251  ND2 ASN B 513      26.383  -3.666  74.998  1.00 81.34           N
    ATOM   2252  H   ASN B 513      27.061  -6.438  74.011  1.00  0.00           H
    ATOM   2253  HA  ASN B 513      25.200  -5.929  76.103  1.00  0.00           H
    ATOM   2254  HB2 ASN B 513      24.765  -5.821  73.084  1.00  0.00           H
    ATOM   2255  HB3 ASN B 513      23.815  -4.993  74.291  1.00  0.00           H
    ATOM   2256 HD21 ASN B 513      26.282  -4.131  75.888  1.00  0.00           H
    ATOM   2257 HD22 ASN B 513      26.953  -2.835  74.929  1.00  0.00           H
    ATOM   2258  N   LEU B 514      24.646  -8.621  74.354  1.00 61.74           N
    ATOM   2259  CA  LEU B 514      23.865  -9.846  74.233  1.00 61.89           C
    ATOM   2260  C   LEU B 514      24.341 -10.957  75.187  1.00 63.07           C
    ATOM   2261  O   LEU B 514      23.786 -12.050  75.131  1.00 65.53           O
    ATOM   2262  CB  LEU B 514      23.905 -10.340  72.768  1.00 63.85           C
    ATOM   2263  CG  LEU B 514      23.462  -9.319  71.694  1.00 62.20           C
    ATOM   2264  CD1 LEU B 514      23.457  -9.974  70.299  1.00 62.23           C
    ATOM   2265  CD2 LEU B 514      22.113  -8.644  72.005  1.00 59.30           C
    ATOM   2266  H   LEU B 514      25.504  -8.590  73.820  1.00  0.00           H
    ATOM   2267  HA  LEU B 514      22.840  -9.612  74.499  1.00  0.00           H
    ATOM   2268  HB2 LEU B 514      24.920 -10.661  72.534  1.00  0.00           H
    ATOM   2269  HB3 LEU B 514      23.280 -11.230  72.676  1.00  0.00           H
    ATOM   2270  HG  LEU B 514      24.212  -8.528  71.665  1.00  0.00           H
    ATOM   2271 HD11 LEU B 514      23.973  -9.348  69.572  1.00  0.00           H
    ATOM   2272 HD12 LEU B 514      23.954 -10.944  70.303  1.00  0.00           H
    ATOM   2273 HD13 LEU B 514      22.448 -10.141  69.924  1.00  0.00           H
    ATOM   2274 HD21 LEU B 514      21.563  -8.392  71.099  1.00  0.00           H
    ATOM   2275 HD22 LEU B 514      21.468  -9.284  72.603  1.00  0.00           H
    ATOM   2276 HD23 LEU B 514      22.262  -7.715  72.556  1.00  0.00           H
    ATOM   2277  N   LEU B 515      25.373 -10.691  76.004  1.00 64.40           N
    ATOM   2278  CA  LEU B 515      26.039 -11.684  76.843  1.00 62.00           C
    ATOM   2279  C   LEU B 515      25.619 -11.521  78.306  1.00 63.51           C
    ATOM   2280  O   LEU B 515      25.320 -10.417  78.765  1.00 62.61           O
    ATOM   2281  CB  LEU B 515      27.569 -11.514  76.709  1.00 55.38           C
    ATOM   2282  CG  LEU B 515      28.251 -12.317  75.577  1.00 49.67           C
    ATOM   2283  CD1 LEU B 515      27.550 -12.288  74.207  1.00 49.07           C
    ATOM   2284  CD2 LEU B 515      29.711 -11.862  75.439  1.00 46.01           C
    ATOM   2285  H   LEU B 515      25.765  -9.760  76.025  1.00  0.00           H
    ATOM   2286  HA  LEU B 515      25.770 -12.690  76.522  1.00  0.00           H
    ATOM   2287  HB2 LEU B 515      27.805 -10.458  76.599  1.00  0.00           H
    ATOM   2288  HB3 LEU B 515      28.058 -11.810  77.638  1.00  0.00           H
    ATOM   2289  HG  LEU B 515      28.268 -13.359  75.899  1.00  0.00           H
    ATOM   2290 HD11 LEU B 515      28.267 -12.268  73.385  1.00  0.00           H
    ATOM   2291 HD12 LEU B 515      26.933 -13.175  74.069  1.00  0.00           H
    ATOM   2292 HD13 LEU B 515      26.913 -11.419  74.079  1.00  0.00           H
    ATOM   2293 HD21 LEU B 515      30.356 -12.698  75.175  1.00  0.00           H
    ATOM   2294 HD22 LEU B 515      29.828 -11.100  74.668  1.00  0.00           H
    ATOM   2295 HD23 LEU B 515      30.078 -11.435  76.371  1.00  0.00           H
    ATOM   2296  N   GLY B 516      25.661 -12.654  79.010  1.00 64.17           N
    ATOM   2297  CA  GLY B 516      25.534 -12.741  80.452  1.00 65.16           C
    ATOM   2298  C   GLY B 516      26.508 -13.825  80.909  1.00 65.42           C
    ATOM   2299  O   GLY B 516      26.912 -14.691  80.129  1.00 68.81           O
    ATOM   2300  H   GLY B 516      25.902 -13.516  78.535  1.00  0.00           H
    ATOM   2301  HA2 GLY B 516      25.758 -11.795  80.947  1.00  0.00           H
    ATOM   2302  HA3 GLY B 516      24.519 -13.035  80.713  1.00  0.00           H
  extra_rotamers: ""
```
