## Supplementary material for "DISRUPTOR: Computational identification of oncogenic mutants disrupting protein interactions": exemplary input files: p53_DNA_1tup.A101-165.yaml

```
---
osprey_version: 3.1
design_name: p53 and 1tup
protein:
  residue_configurations:
  - mutability:
    - ALA
    - ARG
    - ASN
    - ASP
    - CYS
    - GLU
    - GLN
    - GLY
    - HIE
    - HID
    - HIP
    - ILE
    - LEU
    - LYS
    - MET
    - PHE
    - PRO
    - SER
    - THR
    - TRP
    - TYR
    - VAL
    flexibility:
      is_flexible: true
      include_structure_rotamer: true
      use_continuous: true
    identity:
      chain: A
      res_num: 101
      aa_type: LYS
  - mutability:
    - ALA
    - ARG
    - ASN
    - ASP
    - CYS
    - GLU
    - GLN
    - GLY
    - HIE
    - HID
    - HIP
    - ILE
    - LEU
    - LYS
    - MET
    - PHE
    - PRO
    - SER
    - THR
    - TRP
    - TYR
    - VAL
    flexibility:
      is_flexible: true
      include_structure_rotamer: true
      use_continuous: true
    identity:
      chain: A
      res_num: 165
      aa_type: GLN
  extra_templates: |2
        0    0    2
    This is a remark line
    molecule.res

     CY1 INT 1
     CORR OMIT DU   BEG 
       0.00000
        1  DUMM  DU    M    0  -1  -2     0.000      .0        .0      .00000
        2  DUMM  DU    M    1   0  -1     1.449      .0        .0      .00000
        3  DUMM  DU    M    2   1   0     1.522   111.1        .0      .00000
        4  N     N     M    3   2   1     1.335   116.600   180.000  -0.46300
        5  HN    H     E    4   3   2     1.010   119.800     0.000   0.25200
        6  CA    CT    M    4   3   2     1.449   121.900   180.000   0.03500
        7  HA    H1    E    6   4   3     1.090   109.500   300.000   0.04800
        8  CB    CT    3    6   4   3     1.525   111.100    60.000  -0.54300
        9  HB3   H1    E    8   6   4     1.090   109.500    60.000   0.18377
       10  HB2   H1    E    8   6   4     1.090   109.500   300.000   0.18377
       11  SG    S1    E    8   6   4     1.810   116.000   180.000  -0.43963
       12  C     C     M    6   4   3     1.522   111.100   180.000   0.61600
       13  O     O     E   12   6   4     1.229   120.500     0.000  -0.50400
     
    IMPROPER
     -M   CA   N    HN 
     CA   +M   C    O  
     
    DONE
    Zinc(II) Ion
    
     ZN1 INT 1
     CORR OMIT DU   BEG 
       0.00000
        1  DUMM  DU    M    0  -1  -2     0.000      .0        .0      .00000
        2  DUMM  DU    M    1   0  -1     1.449      .0        .0      .00000
        3  DUMM  DU    M    2   1   0     1.522   111.1        .0      .00000
        4 ZN     ZN    M    3   2   1     1.000    90.000   180.000   0.52437
      
    DONE
    CYSTEINE with negative charge
    
     CY2 INT 1
     CORR OMIT DU   BEG 
       0.00000
        1  DUMM  DU    M    0  -1  -2     0.000      .0        .0      .00000
        2  DUMM  DU    M    1   0  -1     1.449      .0        .0      .00000
        3  DUMM  DU    M    2   1   0     1.522   111.1        .0      .00000
        4  N     N     M    3   2   1     1.335   116.600   180.000  -0.46300
        5  H     H     E    4   3   2     1.010   119.800     0.000   0.25200
        6  CA    CT    M    4   3   2     1.449   121.900   180.000   0.03500
        7  HA    H1    E    6   4   3     1.090   109.500   300.000   0.04800
        8  CB    CT    3    6   4   3     1.525   111.100    60.000  -0.42561
        9  HB3   H1    E    8   6   4     1.090   109.500    60.000   0.17337
       10  HB2   H1    E    8   6   4     1.090   109.500   300.000   0.17337
       11  SG    S2    E    8   6   4     1.810   116.000   180.000  -0.44155
       12  C     C     M    6   4   3     1.522   111.100   180.000   0.61600
       13  O     O     E   12   6   4     1.229   120.500     0.000  -0.50400
     
    IMPROPER
     -M   CA   N    HN 
     CA   +M   C    O  
     
    DONE
    HISTIDINE EPSILONH
    
     HE1 INT 1
     CORR OMIT DU   BEG 
       0.00000
        1  DUMM  DU    M    0  -1  -2     0.000      .0        .0      .00000
        2  DUMM  DU    M    1   0  -1     1.449      .0        .0      .00000
        3  DUMM  DU    M    2   1   0     1.522   111.1        .0      .00000
        4  N     N     M    3   2   1     1.335   116.600   180.000  -0.41570
        5  H     H     E    4   3   2     1.010   119.800     0.000   0.27190
        6  CA    CT    M    4   3   2     1.449   121.900   180.000  -0.05810
        7  HA    H1    E    6   4   3     1.090   109.500   300.000   0.13600
        8  CB    CT    3    6   4   3     1.525   111.100    60.000   0.04092
        9  HB2   HC    E    8   6   4     1.090   109.500   300.000   0.06506
       10  HB3   HC    E    8   6   4     1.090   109.500    60.000   0.06506
       11  CG    CC    S    8   6   4     1.510   115.000   180.000  -0.11300
       12  ND1   N1    S   11   8   6     1.390   122.000   180.000  -0.08206
       13  CE1   CR    B   12  11   8     1.320   108.000   180.000  -0.14656
       14  HE1   H5    E   13  12  11     1.090   120.000   180.000   0.19871
       15  NE2   NA    B   13  12  11     1.310   109.000     0.000  -0.01782
       16  HE2   H     E   15  13  12     1.010   125.000   180.000   0.27163
       17  CD2   CW    S   15  13  12     1.360   110.000     0.000  -0.19685
       18  HD2   H4    E   17  15  13     1.090   120.000   180.000   0.12407
       19  C     C     M    6   4   3     1.522   111.100   180.000   0.59730
       20  O     O     E   19   6   4     1.229   120.500     0.000  -0.56790
      
    LOOP
     CG   CD2
     
    IMPROPER
     -M   CA   N    H  
     CA   +M   C    O  
     CE1  CD2  NE2  HE2
     CG   NE2  CD2  HD2
     ND1  NE2  CE1  HE1
     ND1  CD2  CG   CB 
     
    DONE
    Zinc(II) Ion
    
     ZN2 INT 1
     CORR OMIT DU   BEG 
       0.00000
        1  DUMM  DU    M    0  -1  -2     0.000      .0        .0      .00000
        2  DUMM  DU    M    1   0  -1     1.449      .0        .0      .00000
        3  DUMM  DU    M    2   1   0     1.522   111.1        .0      .00000
        4  ZN    ZN    M    3   2   1     1.000    90.000   180.000   0.43655
      
    DONE
  extra_templates_coordinates: ""
  coordinates: |
    ATOM      1  N   SER A  94      76.973  22.225  81.234  1.00 23.50           N1+
    ATOM      2  CA  SER A  94      75.613  21.900  80.696  1.00 24.59           C
    ATOM      3  C   SER A  94      75.814  20.923  79.525  1.00 26.11           C
    ATOM      4  O   SER A  94      76.940  20.595  79.143  1.00 23.93           O
    ATOM      5  CB  SER A  94      74.907  23.202  80.289  1.00 22.31           C
    ATOM      6  OG  SER A  94      74.764  23.994  81.446  1.00 20.52           O
    ATOM      7  H1  SER A  94      76.864  22.878  81.997  1.00  0.00           H
    ATOM      8  H2  SER A  94      77.416  21.377  81.585  1.00  0.00           H
    ATOM      9  H3  SER A  94      77.539  22.632  80.500  1.00  0.00           H
    ATOM     10  HA  SER A  94      75.087  21.395  81.509  1.00  0.00           H
    ATOM     11  HB2 SER A  94      75.485  23.755  79.546  1.00  0.00           H
    ATOM     12  HB3 SER A  94      73.924  23.026  79.856  1.00  0.00           H
    ATOM     13  HG  SER A  94      74.375  24.834  81.193  1.00  0.00           H
    ATOM     14  N   SER A  95      74.691  20.470  78.948  1.00 29.04           N
    ATOM     15  CA  SER A  95      74.686  19.624  77.762  1.00 23.53           C
    ATOM     16  C   SER A  95      75.353  20.387  76.607  1.00 23.17           C
    ATOM     17  O   SER A  95      74.928  21.503  76.299  1.00 23.61           O
    ATOM     18  CB  SER A  95      73.234  19.213  77.464  1.00 21.26           C
    ATOM     19  OG  SER A  95      73.159  17.908  76.946  1.00 31.17           O
    ATOM     20  H   SER A  95      73.786  20.742  79.313  1.00  0.00           H
    ATOM     21  HA  SER A  95      75.267  18.732  78.007  1.00  0.00           H
    ATOM     22  HB2 SER A  95      72.627  19.220  78.370  1.00  0.00           H
    ATOM     23  HB3 SER A  95      72.779  19.901  76.759  1.00  0.00           H
    ATOM     24  HG  SER A  95      73.532  17.295  77.588  1.00  0.00           H
    ATOM     25  N   SER A  96      76.441  19.820  76.070  1.00 24.75           N
    ATOM     26  CA  SER A  96      77.249  20.443  75.028  1.00 26.68           C
    ATOM     27  C   SER A  96      76.449  20.681  73.743  1.00 29.59           C
    ATOM     28  O   SER A  96      75.532  19.936  73.415  1.00 36.22           O
    ATOM     29  CB  SER A  96      78.535  19.627  74.815  1.00 31.31           C
    ATOM     30  OG  SER A  96      78.258  18.262  74.589  1.00 36.85           O
    ATOM     31  H   SER A  96      76.755  18.911  76.388  1.00  0.00           H
    ATOM     32  HA  SER A  96      77.553  21.420  75.407  1.00  0.00           H
    ATOM     33  HB2 SER A  96      79.113  20.021  73.977  1.00  0.00           H
    ATOM     34  HB3 SER A  96      79.174  19.704  75.696  1.00  0.00           H
    ATOM     35  HG  SER A  96      77.951  18.148  73.685  1.00  0.00           H
    ATOM     36  N   VAL A  97      76.797  21.771  73.073  1.00 22.57           N
    ATOM     37  CA  VAL A  97      76.098  22.306  71.918  1.00 23.40           C
    ATOM     38  C   VAL A  97      77.160  22.761  70.908  1.00 27.15           C
    ATOM     39  O   VAL A  97      78.288  23.038  71.328  1.00 29.57           O
    ATOM     40  CB  VAL A  97      75.250  23.530  72.351  1.00 29.18           C
    ATOM     41  CG1 VAL A  97      73.949  23.090  73.040  1.00 31.07           C
    ATOM     42  CG2 VAL A  97      76.026  24.531  73.235  1.00 34.23           C
    ATOM     43  H   VAL A  97      77.581  22.325  73.390  1.00  0.00           H
    ATOM     44  HA  VAL A  97      75.460  21.540  71.481  1.00  0.00           H
    ATOM     45  HB  VAL A  97      74.948  24.072  71.453  1.00  0.00           H
    ATOM     46 HG11 VAL A  97      73.330  23.946  73.296  1.00  0.00           H
    ATOM     47 HG12 VAL A  97      73.361  22.438  72.394  1.00  0.00           H
    ATOM     48 HG13 VAL A  97      74.146  22.547  73.963  1.00  0.00           H
    ATOM     49 HG21 VAL A  97      75.393  25.380  73.497  1.00  0.00           H
    ATOM     50 HG22 VAL A  97      76.366  24.085  74.167  1.00  0.00           H
    ATOM     51 HG23 VAL A  97      76.897  24.937  72.720  1.00  0.00           H
    ATOM     52  N   PRO A  98      76.810  22.848  69.603  1.00 26.71           N
    ATOM     53  CA  PRO A  98      77.723  23.407  68.598  1.00 26.98           C
    ATOM     54  C   PRO A  98      78.188  24.828  68.926  1.00 30.58           C
    ATOM     55  O   PRO A  98      77.384  25.682  69.305  1.00 32.88           O
    ATOM     56  CB  PRO A  98      76.934  23.380  67.284  1.00 24.50           C
    ATOM     57  CG  PRO A  98      75.883  22.312  67.492  1.00 23.33           C
    ATOM     58  CD  PRO A  98      75.568  22.388  68.981  1.00 22.20           C
    ATOM     59  HA  PRO A  98      78.574  22.727  68.523  1.00  0.00           H
    ATOM     60  HB2 PRO A  98      76.451  24.339  67.097  1.00  0.00           H
    ATOM     61  HB3 PRO A  98      77.568  23.166  66.422  1.00  0.00           H
    ATOM     62  HG2 PRO A  98      75.010  22.438  66.852  1.00  0.00           H
    ATOM     63  HG3 PRO A  98      76.329  21.342  67.265  1.00  0.00           H
    ATOM     64  HD2 PRO A  98      74.778  23.113  69.177  1.00  0.00           H
    ATOM     65  HD3 PRO A  98      75.246  21.411  69.337  1.00  0.00           H
    ATOM     66  N   SER A  99      79.497  25.025  68.767  1.00 30.53           N
    ATOM     67  CA  SER A  99      80.198  26.293  68.915  1.00 36.50           C
    ATOM     68  C   SER A  99      79.653  27.306  67.889  1.00 34.87           C
    ATOM     69  O   SER A  99      79.344  26.919  66.767  1.00 33.12           O
    ATOM     70  CB  SER A  99      81.686  26.009  68.638  1.00 45.47           C
    ATOM     71  OG  SER A  99      82.134  24.849  69.315  1.00 51.61           O
    ATOM     72  H   SER A  99      80.079  24.253  68.468  1.00  0.00           H
    ATOM     73  HA  SER A  99      80.063  26.663  69.933  1.00  0.00           H
    ATOM     74  HB2 SER A  99      81.845  25.836  67.573  1.00  0.00           H
    ATOM     75  HB3 SER A  99      82.306  26.862  68.918  1.00  0.00           H
    ATOM     76  HG  SER A  99      83.029  24.652  69.029  1.00  0.00           H
    ATOM     77  N   GLN A 100      79.528  28.582  68.256  1.00 32.67           N
    ATOM     78  CA  GLN A 100      79.027  29.599  67.321  1.00 28.20           C
    ATOM     79  C   GLN A 100      79.882  30.864  67.335  1.00 28.21           C
    ATOM     80  O   GLN A 100      79.496  31.823  66.674  1.00 31.37           O
    ATOM     81  CB  GLN A 100      77.562  29.930  67.650  1.00 24.44           C
    ATOM     82  CG  GLN A 100      77.435  30.730  68.953  1.00 26.66           C
    ATOM     83  CD  GLN A 100      76.003  30.872  69.407  1.00 33.43           C
    ATOM     84  OE1 GLN A 100      75.405  31.930  69.247  1.00 35.08           O
    ATOM     85  NE2 GLN A 100      75.470  29.808  70.004  1.00 38.80           N
    ATOM     86  H   GLN A 100      79.746  28.877  69.196  1.00  0.00           H
    ATOM     87  HA  GLN A 100      79.058  29.250  66.288  1.00  0.00           H
    ATOM     88  HB2 GLN A 100      77.127  30.523  66.846  1.00  0.00           H
    ATOM     89  HB3 GLN A 100      76.984  29.006  67.695  1.00  0.00           H
    ATOM     90  HG2 GLN A 100      77.977  30.221  69.752  1.00  0.00           H
    ATOM     91  HG3 GLN A 100      77.884  31.717  68.882  1.00  0.00           H
    ATOM     92 HE21 GLN A 100      75.998  28.955  70.120  1.00  0.00           H
    ATOM     93 HE22 GLN A 100      74.525  29.859  70.359  1.00  0.00           H
    ATOM     94  N   LYS A 101      80.973  30.883  68.114  1.00 31.06           N
    ATOM     95  CA  LYS A 101      81.824  32.056  68.266  1.00 26.06           C
    ATOM     96  C   LYS A 101      82.624  32.312  66.993  1.00 25.65           C
    ATOM     97  O   LYS A 101      83.176  31.376  66.410  1.00 21.93           O
    ATOM     98  CB  LYS A 101      82.768  31.868  69.462  1.00 31.79           C
    ATOM     99  CG  LYS A 101      82.049  31.848  70.822  1.00 37.92           C
    ATOM    100  CD  LYS A 101      81.715  33.247  71.372  1.00 43.32           C
    ATOM    101  CE  LYS A 101      80.250  33.393  71.811  1.00 50.61           C
    ATOM    102  NZ  LYS A 101      80.008  34.685  72.478  1.00 53.92           N1+
    ATOM    103  H   LYS A 101      81.236  30.063  68.640  1.00  0.00           H
    ATOM    104  HA  LYS A 101      81.181  32.917  68.464  1.00  0.00           H
    ATOM    105  HB2 LYS A 101      83.309  30.929  69.336  1.00  0.00           H
    ATOM    106  HB3 LYS A 101      83.533  32.646  69.478  1.00  0.00           H
    ATOM    107  HG2 LYS A 101      81.147  31.237  70.758  1.00  0.00           H
    ATOM    108  HG3 LYS A 101      82.686  31.336  71.543  1.00  0.00           H
    ATOM    109  HD2 LYS A 101      82.368  33.455  72.221  1.00  0.00           H
    ATOM    110  HD3 LYS A 101      81.942  34.013  70.630  1.00  0.00           H
    ATOM    111  HE2 LYS A 101      79.585  33.302  70.951  1.00  0.00           H
    ATOM    112  HE3 LYS A 101      79.987  32.597  72.507  1.00  0.00           H
    ATOM    113  HZ1 LYS A 101      80.187  35.443  71.836  1.00  0.00           H
    ATOM    114  HZ2 LYS A 101      79.047  34.734  72.791  1.00  0.00           H
    ATOM    115  HZ3 LYS A 101      80.615  34.775  73.281  1.00  0.00           H
    ATOM    116  N   THR A 102      82.665  33.592  66.612  1.00 29.20           N
    ATOM    117  CA  THR A 102      83.374  34.068  65.440  1.00 24.75           C
    ATOM    118  C   THR A 102      84.891  33.903  65.604  1.00 26.66           C
    ATOM    119  O   THR A 102      85.468  34.310  66.613  1.00 29.11           O
    ATOM    120  CB  THR A 102      83.048  35.552  65.160  1.00 20.65           C
    ATOM    121  OG1 THR A 102      81.676  35.642  64.837  1.00 23.23           O
    ATOM    122  CG2 THR A 102      83.853  36.201  64.021  1.00 14.51           C
    ATOM    123  H   THR A 102      82.181  34.296  67.150  1.00  0.00           H
    ATOM    124  HA  THR A 102      83.030  33.475  64.591  1.00  0.00           H
    ATOM    125  HB  THR A 102      83.204  36.138  66.068  1.00  0.00           H
    ATOM    126 HG21 THR A 102      83.468  37.192  63.780  1.00  0.00           H
    ATOM    127 HG22 THR A 102      84.904  36.329  64.283  1.00  0.00           H
    ATOM    128 HG23 THR A 102      83.810  35.598  63.113  1.00  0.00           H
    ATOM    129  HG1 THR A 102      81.493  36.536  64.533  1.00  0.00           H
    ATOM    130  N   TYR A 103      85.484  33.289  64.583  1.00 24.86           N
    ATOM    131  CA  TYR A 103      86.903  33.037  64.486  1.00 20.83           C
    ATOM    132  C   TYR A 103      87.271  33.286  63.036  1.00 22.44           C
    ATOM    133  O   TYR A 103      86.888  32.506  62.171  1.00 24.99           O
    ATOM    134  CB  TYR A 103      87.202  31.597  64.938  1.00 19.87           C
    ATOM    135  CG  TYR A 103      88.641  31.145  64.764  1.00 23.28           C
    ATOM    136  CD1 TYR A 103      89.709  31.898  65.298  1.00 28.35           C
    ATOM    137  CD2 TYR A 103      88.912  29.948  64.074  1.00 20.64           C
    ATOM    138  CE1 TYR A 103      91.035  31.449  65.146  1.00 25.59           C
    ATOM    139  CE2 TYR A 103      90.235  29.502  63.914  1.00 18.05           C
    ATOM    140  CZ  TYR A 103      91.298  30.249  64.453  1.00 20.10           C
    ATOM    141  OH  TYR A 103      92.580  29.813  64.291  1.00 24.17           O
    ATOM    142  H   TYR A 103      84.930  32.996  63.786  1.00  0.00           H
    ATOM    143  HA  TYR A 103      87.455  33.731  65.119  1.00  0.00           H
    ATOM    144  HB2 TYR A 103      86.931  31.465  65.983  1.00  0.00           H
    ATOM    145  HB3 TYR A 103      86.557  30.910  64.388  1.00  0.00           H
    ATOM    146  HD1 TYR A 103      89.517  32.817  65.832  1.00  0.00           H
    ATOM    147  HD2 TYR A 103      88.103  29.359  63.666  1.00  0.00           H
    ATOM    148  HE1 TYR A 103      91.848  32.029  65.557  1.00  0.00           H
    ATOM    149  HE2 TYR A 103      90.433  28.582  63.382  1.00  0.00           H
    ATOM    150  HH  TYR A 103      93.231  30.373  64.721  1.00  0.00           H
    ATOM    151  N   GLN A 104      87.994  34.378  62.782  1.00 23.72           N
    ATOM    152  CA  GLN A 104      88.406  34.702  61.423  1.00 19.26           C
    ATOM    153  C   GLN A 104      89.612  33.870  60.981  1.00 20.76           C
    ATOM    154  O   GLN A 104      89.764  33.649  59.787  1.00 23.88           O
    ATOM    155  CB  GLN A 104      88.561  36.221  61.271  1.00 19.77           C
    ATOM    156  CG  GLN A 104      87.213  36.905  61.609  1.00 21.47           C
    ATOM    157  CD  GLN A 104      87.097  38.390  61.290  1.00 33.72           C
    ATOM    158  OE1 GLN A 104      86.053  38.978  61.564  1.00 39.21           O
    ATOM    159  NE2 GLN A 104      88.133  39.013  60.726  1.00 36.55           N
    ATOM    160  H   GLN A 104      88.274  35.006  63.524  1.00  0.00           H
    ATOM    161  HA  GLN A 104      87.603  34.412  60.746  1.00  0.00           H
    ATOM    162  HB2 GLN A 104      89.352  36.597  61.921  1.00  0.00           H
    ATOM    163  HB3 GLN A 104      88.855  36.444  60.244  1.00  0.00           H
    ATOM    164  HG2 GLN A 104      86.394  36.406  61.092  1.00  0.00           H
    ATOM    165  HG3 GLN A 104      86.999  36.797  62.673  1.00  0.00           H
    ATOM    166 HE21 GLN A 104      88.983  38.510  60.518  1.00  0.00           H
    ATOM    167 HE22 GLN A 104      88.075  39.998  60.517  1.00  0.00           H
    ATOM    168  N   GLY A 105      90.394  33.350  61.939  1.00 21.45           N
    ATOM    169  CA  GLY A 105      91.495  32.429  61.697  1.00 21.55           C
    ATOM    170  C   GLY A 105      92.626  33.057  60.886  1.00 26.14           C
    ATOM    171  O   GLY A 105      92.660  34.262  60.637  1.00 35.50           O
    ATOM    172  H   GLY A 105      90.189  33.568  62.903  1.00  0.00           H
    ATOM    173  HA2 GLY A 105      91.912  32.131  62.655  1.00  0.00           H
    ATOM    174  HA3 GLY A 105      91.129  31.534  61.207  1.00  0.00           H
    ATOM    175  N   SER A 106      93.575  32.201  60.498  1.00 23.15           N
    ATOM    176  CA  SER A 106      94.814  32.589  59.828  1.00 28.41           C
    ATOM    177  C   SER A 106      94.595  33.153  58.416  1.00 33.57           C
    ATOM    178  O   SER A 106      95.475  33.850  57.918  1.00 35.06           O
    ATOM    179  CB  SER A 106      95.722  31.354  59.762  1.00 29.19           C
    ATOM    180  OG  SER A 106      95.837  30.765  61.043  1.00 32.27           O
    ATOM    181  H   SER A 106      93.508  31.227  60.756  1.00  0.00           H
    ATOM    182  HA  SER A 106      95.303  33.357  60.431  1.00  0.00           H
    ATOM    183  HB2 SER A 106      95.322  30.608  59.073  1.00  0.00           H
    ATOM    184  HB3 SER A 106      96.716  31.625  59.403  1.00  0.00           H
    ATOM    185  HG  SER A 106      96.414  30.000  60.983  1.00  0.00           H
    ATOM    186  N   TYR A 107      93.438  32.840  57.812  1.00 29.10           N
    ATOM    187  CA  TYR A 107      93.078  33.216  56.453  1.00 23.29           C
    ATOM    188  C   TYR A 107      92.119  34.409  56.396  1.00 25.77           C
    ATOM    189  O   TYR A 107      91.688  34.750  55.302  1.00 29.54           O
    ATOM    190  CB  TYR A 107      92.534  31.977  55.715  1.00 25.77           C
    ATOM    191  CG  TYR A 107      93.448  30.776  55.797  1.00 30.91           C
    ATOM    192  CD1 TYR A 107      94.569  30.693  54.948  1.00 28.68           C
    ATOM    193  CD2 TYR A 107      93.210  29.765  56.749  1.00 28.51           C
    ATOM    194  CE1 TYR A 107      95.475  29.626  55.078  1.00 28.09           C
    ATOM    195  CE2 TYR A 107      94.111  28.693  56.875  1.00 28.91           C
    ATOM    196  CZ  TYR A 107      95.251  28.633  56.051  1.00 33.57           C
    ATOM    197  OH  TYR A 107      96.134  27.604  56.193  1.00 37.69           O
    ATOM    198  H   TYR A 107      92.762  32.268  58.300  1.00  0.00           H
    ATOM    199  HA  TYR A 107      93.971  33.540  55.932  1.00  0.00           H
    ATOM    200  HB2 TYR A 107      91.542  31.702  56.073  1.00  0.00           H
    ATOM    201  HB3 TYR A 107      92.426  32.188  54.660  1.00  0.00           H
    ATOM    202  HD1 TYR A 107      94.742  31.444  54.196  1.00  0.00           H
    ATOM    203  HD2 TYR A 107      92.354  29.817  57.404  1.00  0.00           H
    ATOM    204  HE1 TYR A 107      96.336  29.573  54.429  1.00  0.00           H
    ATOM    205  HE2 TYR A 107      93.929  27.927  57.614  1.00  0.00           H
    ATOM    206  HH  TYR A 107      95.873  26.981  56.877  1.00  0.00           H
    ATOM    207  N   GLY A 108      91.812  35.050  57.539  1.00 25.41           N
    ATOM    208  CA  GLY A 108      90.941  36.225  57.612  1.00 27.15           C
    ATOM    209  C   GLY A 108      89.541  35.931  57.055  1.00 28.61           C
    ATOM    210  O   GLY A 108      89.032  36.685  56.235  1.00 23.49           O
    ATOM    211  H   GLY A 108      92.220  34.729  58.407  1.00  0.00           H
    ATOM    212  HA2 GLY A 108      90.855  36.540  58.653  1.00  0.00           H
    ATOM    213  HA3 GLY A 108      91.396  37.048  57.062  1.00  0.00           H
    ATOM    214  N   PHE A 109      88.942  34.800  57.442  1.00 28.87           N
    ATOM    215  CA  PHE A 109      87.617  34.393  57.007  1.00 24.11           C
    ATOM    216  C   PHE A 109      86.522  35.312  57.572  1.00 22.80           C
    ATOM    217  O   PHE A 109      86.336  35.362  58.785  1.00 25.48           O
    ATOM    218  CB  PHE A 109      87.403  32.892  57.278  1.00 25.27           C
    ATOM    219  CG  PHE A 109      86.000  32.352  57.045  1.00 24.55           C
    ATOM    220  CD1 PHE A 109      85.382  32.441  55.777  1.00 27.03           C
    ATOM    221  CD2 PHE A 109      85.297  31.751  58.108  1.00 20.54           C
    ATOM    222  CE1 PHE A 109      84.096  31.944  55.600  1.00 30.20           C
    ATOM    223  CE2 PHE A 109      84.017  31.255  57.909  1.00 22.08           C
    ATOM    224  CZ  PHE A 109      83.419  31.355  56.661  1.00 23.43           C
    ATOM    225  H   PHE A 109      89.388  34.212  58.135  1.00  0.00           H
    ATOM    226  HA  PHE A 109      87.619  34.477  55.927  1.00  0.00           H
    ATOM    227  HB2 PHE A 109      88.079  32.325  56.638  1.00  0.00           H
    ATOM    228  HB3 PHE A 109      87.702  32.659  58.301  1.00  0.00           H
    ATOM    229  HD1 PHE A 109      85.897  32.903  54.948  1.00  0.00           H
    ATOM    230  HD2 PHE A 109      85.745  31.689  59.087  1.00  0.00           H
    ATOM    231  HE1 PHE A 109      83.617  32.028  54.636  1.00  0.00           H
    ATOM    232  HE2 PHE A 109      83.480  30.807  58.729  1.00  0.00           H
    ATOM    233  HZ  PHE A 109      82.414  30.994  56.517  1.00  0.00           H
    ATOM    234  N   ARG A 110      85.831  36.014  56.669  1.00 25.26           N
    ATOM    235  CA  ARG A 110      84.619  36.782  56.941  1.00 25.44           C
    ATOM    236  C   ARG A 110      83.526  36.277  55.994  1.00 22.79           C
    ATOM    237  O   ARG A 110      83.825  35.701  54.947  1.00 22.93           O
    ATOM    238  CB  ARG A 110      84.825  38.293  56.694  1.00 33.06           C
    ATOM    239  CG  ARG A 110      86.006  38.969  57.412  1.00 40.33           C
    ATOM    240  CD  ARG A 110      86.473  40.224  56.658  1.00 48.29           C
    ATOM    241  NE  ARG A 110      87.060  39.848  55.362  1.00 56.43           N
    ATOM    242  CZ  ARG A 110      88.310  39.397  55.190  1.00 58.52           C
    ATOM    243  NH1 ARG A 110      89.183  39.421  56.205  1.00 60.26           N
    ATOM    244  NH2 ARG A 110      88.698  38.953  53.990  1.00 53.96           N1+
    ATOM    245  H   ARG A 110      86.071  35.914  55.690  1.00  0.00           H
    ATOM    246  HA  ARG A 110      84.283  36.629  57.969  1.00  0.00           H
    ATOM    247  HB2 ARG A 110      84.943  38.442  55.628  1.00  0.00           H
    ATOM    248  HB3 ARG A 110      83.912  38.835  56.948  1.00  0.00           H
    ATOM    249  HG2 ARG A 110      85.833  39.144  58.474  1.00  0.00           H
    ATOM    250  HG3 ARG A 110      86.846  38.276  57.365  1.00  0.00           H
    ATOM    251  HD2 ARG A 110      85.584  40.792  56.375  1.00  0.00           H
    ATOM    252  HD3 ARG A 110      87.081  40.900  57.260  1.00  0.00           H
    ATOM    253  HE  ARG A 110      86.433  39.857  54.571  1.00  0.00           H
    ATOM    254 HH11 ARG A 110      88.897  39.754  57.112  1.00  0.00           H
    ATOM    255 HH12 ARG A 110      90.157  39.177  56.052  1.00  0.00           H
    ATOM    256 HH21 ARG A 110      88.073  38.963  53.199  1.00  0.00           H
    ATOM    257 HH22 ARG A 110      89.672  38.712  53.849  1.00  0.00           H
    ATOM    258  N   LEU A 111      82.272  36.550  56.370  1.00 19.36           N
    ATOM    259  CA  LEU A 111      81.112  36.342  55.514  1.00 20.86           C
    ATOM    260  C   LEU A 111      80.667  37.680  54.924  1.00 26.13           C
    ATOM    261  O   LEU A 111      80.725  38.702  55.606  1.00 28.47           O
    ATOM    262  CB  LEU A 111      79.947  35.753  56.328  1.00 22.99           C
    ATOM    263  CG  LEU A 111      80.106  34.276  56.718  1.00 20.40           C
    ATOM    264  CD1 LEU A 111      78.860  33.811  57.494  1.00 17.68           C
    ATOM    265  CD2 LEU A 111      80.392  33.359  55.517  1.00 20.18           C
    ATOM    266  H   LEU A 111      82.101  37.033  57.240  1.00  0.00           H
    ATOM    267  HA  LEU A 111      81.368  35.669  54.703  1.00  0.00           H
    ATOM    268  HB2 LEU A 111      79.782  36.357  57.222  1.00  0.00           H
    ATOM    269  HB3 LEU A 111      79.028  35.836  55.744  1.00  0.00           H
    ATOM    270  HG  LEU A 111      80.969  34.219  57.373  1.00  0.00           H
    ATOM    271 HD11 LEU A 111      79.122  33.089  58.267  1.00  0.00           H
    ATOM    272 HD12 LEU A 111      78.354  34.642  57.988  1.00  0.00           H
    ATOM    273 HD13 LEU A 111      78.130  33.341  56.837  1.00  0.00           H
    ATOM    274 HD21 LEU A 111      79.897  32.395  55.610  1.00  0.00           H
    ATOM    275 HD22 LEU A 111      80.051  33.799  54.579  1.00  0.00           H
    ATOM    276 HD23 LEU A 111      81.460  33.167  55.422  1.00  0.00           H
    ATOM    277  N   GLY A 112      80.188  37.633  53.681  1.00 26.73           N
    ATOM    278  CA  GLY A 112      79.597  38.761  52.985  1.00 22.91           C
    ATOM    279  C   GLY A 112      78.221  38.343  52.483  1.00 26.14           C
    ATOM    280  O   GLY A 112      77.856  37.165  52.483  1.00 28.30           O
    ATOM    281  H   GLY A 112      80.182  36.746  53.192  1.00  0.00           H
    ATOM    282  HA2 GLY A 112      79.500  39.643  53.620  1.00  0.00           H
    ATOM    283  HA3 GLY A 112      80.211  39.040  52.134  1.00  0.00           H
    ATOM    284  N   PHE A 113      77.456  39.346  52.048  1.00 25.18           N
    ATOM    285  CA  PHE A 113      76.094  39.175  51.567  1.00 20.38           C
    ATOM    286  C   PHE A 113      75.831  40.202  50.461  1.00 21.49           C
    ATOM    287  O   PHE A 113      76.510  41.227  50.378  1.00 23.03           O
    ATOM    288  CB  PHE A 113      75.099  39.356  52.738  1.00 13.02           C
    ATOM    289  CG  PHE A 113      75.273  38.426  53.927  1.00 16.78           C
    ATOM    290  CD1 PHE A 113      74.736  37.122  53.899  1.00 18.91           C
    ATOM    291  CD2 PHE A 113      76.102  38.802  55.008  1.00 19.76           C
    ATOM    292  CE1 PHE A 113      74.968  36.255  54.960  1.00 18.59           C
    ATOM    293  CE2 PHE A 113      76.330  37.918  56.052  1.00 19.14           C
    ATOM    294  CZ  PHE A 113      75.757  36.655  56.031  1.00 17.71           C
    ATOM    295  H   PHE A 113      77.807  40.293  52.044  1.00  0.00           H
    ATOM    296  HA  PHE A 113      75.971  38.181  51.131  1.00  0.00           H
    ATOM    297  HB2 PHE A 113      75.144  40.376  53.107  1.00  0.00           H
    ATOM    298  HB3 PHE A 113      74.081  39.228  52.368  1.00  0.00           H
    ATOM    299  HD1 PHE A 113      74.140  36.796  53.059  1.00  0.00           H
    ATOM    300  HD2 PHE A 113      76.560  39.780  55.029  1.00  0.00           H
    ATOM    301  HE1 PHE A 113      74.547  35.260  54.949  1.00  0.00           H
    ATOM    302  HE2 PHE A 113      76.960  38.210  56.880  1.00  0.00           H
    ATOM    303  HZ  PHE A 113      75.950  35.978  56.847  1.00  0.00           H
    ATOM    304  N   LEU A 114      74.811  39.910  49.647  1.00 19.38           N
    ATOM    305  CA  LEU A 114      74.252  40.818  48.651  1.00 18.09           C
    ATOM    306  C   LEU A 114      73.431  41.908  49.345  1.00 27.87           C
    ATOM    307  O   LEU A 114      72.643  41.606  50.244  1.00 31.07           O
    ATOM    308  CB  LEU A 114      73.343  40.010  47.695  1.00  9.73           C
    ATOM    309  CG  LEU A 114      73.901  39.785  46.280  1.00 11.62           C
    ATOM    310  CD1 LEU A 114      75.368  39.321  46.271  1.00 11.31           C
    ATOM    311  CD2 LEU A 114      72.985  38.812  45.512  1.00  9.41           C
    ATOM    312  H   LEU A 114      74.300  39.046  49.787  1.00  0.00           H
    ATOM    313  HA  LEU A 114      75.063  41.293  48.095  1.00  0.00           H
    ATOM    314  HB2 LEU A 114      73.105  39.046  48.141  1.00  0.00           H
    ATOM    315  HB3 LEU A 114      72.374  40.499  47.577  1.00  0.00           H
    ATOM    316  HG  LEU A 114      73.870  40.744  45.760  1.00  0.00           H
    ATOM    317 HD11 LEU A 114      75.593  38.633  45.456  1.00  0.00           H
    ATOM    318 HD12 LEU A 114      76.047  40.168  46.167  1.00  0.00           H
    ATOM    319 HD13 LEU A 114      75.613  38.812  47.198  1.00  0.00           H
    ATOM    320 HD21 LEU A 114      72.497  39.326  44.685  1.00  0.00           H
    ATOM    321 HD22 LEU A 114      73.527  37.968  45.085  1.00  0.00           H
    ATOM    322 HD23 LEU A 114      72.189  38.404  46.135  1.00  0.00           H
    ATOM    323  N   HIS A 115      73.599  43.148  48.870  1.00 23.30           N
    ATOM    324  CA  HIS A 115      72.845  44.310  49.334  1.00 14.63           C
    ATOM    325  C   HIS A 115      71.588  44.442  48.473  1.00 19.59           C
    ATOM    326  O   HIS A 115      71.553  45.239  47.536  1.00 23.03           O
    ATOM    327  CB  HIS A 115      73.740  45.567  49.301  1.00 18.67           C
    ATOM    328  CG  HIS A 115      75.137  45.311  49.793  1.00 22.74           C
    ATOM    329  ND1 HIS A 115      76.279  45.506  49.012  1.00 24.35           N
    ATOM    330  CD2 HIS A 115      75.510  44.797  51.009  1.00 22.67           C
    ATOM    331  CE1 HIS A 115      77.281  45.079  49.773  1.00 23.11           C
    ATOM    332  NE2 HIS A 115      76.880  44.632  50.963  1.00 24.94           N
    ATOM    333  H   HIS A 115      74.255  43.313  48.120  1.00  0.00           H
    ATOM    334  HA  HIS A 115      72.546  44.154  50.373  1.00  0.00           H
    ATOM    335  HB2 HIS A 115      73.831  45.962  48.289  1.00  0.00           H
    ATOM    336  HB3 HIS A 115      73.293  46.360  49.902  1.00  0.00           H
    ATOM    337  HD2 HIS A 115      74.901  44.543  51.858  1.00  0.00           H
    ATOM    338  HE2 HIS A 115      77.463  44.265  51.701  1.00  0.00           H
    ATOM    339  HE1 HIS A 115      78.314  45.088  49.457  1.00  0.00           H
    ATOM    340  N   SER A 116      70.594  43.600  48.782  1.00 20.18           N
    ATOM    341  CA  SER A 116      69.379  43.449  47.989  1.00 19.43           C
    ATOM    342  C   SER A 116      68.345  44.559  48.215  1.00 22.14           C
    ATOM    343  O   SER A 116      67.502  44.753  47.345  1.00 29.27           O
    ATOM    344  CB  SER A 116      68.733  42.096  48.321  1.00 15.94           C
    ATOM    345  OG  SER A 116      69.621  41.031  48.063  1.00 17.86           O
    ATOM    346  H   SER A 116      70.720  42.935  49.532  1.00  0.00           H
    ATOM    347  HA  SER A 116      69.644  43.452  46.931  1.00  0.00           H
    ATOM    348  HB2 SER A 116      68.427  42.055  49.367  1.00  0.00           H
    ATOM    349  HB3 SER A 116      67.838  41.940  47.718  1.00  0.00           H
    ATOM    350  HG  SER A 116      69.177  40.209  48.294  1.00  0.00           H
    ATOM    351  N   GLY A 117      68.367  45.239  49.371  1.00 15.49           N
    ATOM    352  CA  GLY A 117      67.301  46.162  49.760  1.00 10.17           C
    ATOM    353  C   GLY A 117      66.054  45.375  50.194  1.00 14.73           C
    ATOM    354  O   GLY A 117      66.100  44.160  50.379  1.00 21.35           O
    ATOM    355  H   GLY A 117      69.097  45.058  50.046  1.00  0.00           H
    ATOM    356  HA2 GLY A 117      67.650  46.770  50.595  1.00  0.00           H
    ATOM    357  HA3 GLY A 117      67.057  46.843  48.942  1.00  0.00           H
    ATOM    358  N   THR A 118      64.939  46.087  50.385  1.00 22.37           N
    ATOM    359  CA  THR A 118      63.716  45.537  50.970  1.00 23.48           C
    ATOM    360  C   THR A 118      62.471  45.816  50.116  1.00 27.26           C
    ATOM    361  O   THR A 118      61.364  45.747  50.650  1.00 26.67           O
    ATOM    362  CB  THR A 118      63.495  46.122  52.387  1.00 20.02           C
    ATOM    363  OG1 THR A 118      63.496  47.536  52.342  1.00 21.60           O
    ATOM    364  CG2 THR A 118      64.533  45.653  53.403  1.00 13.02           C
    ATOM    365  H   THR A 118      64.943  47.085  50.232  1.00  0.00           H
    ATOM    366  HA  THR A 118      63.774  44.452  51.047  1.00  0.00           H
    ATOM    367  HB  THR A 118      62.527  45.824  52.780  1.00  0.00           H
    ATOM    368 HG21 THR A 118      64.299  46.035  54.396  1.00  0.00           H
    ATOM    369 HG22 THR A 118      64.534  44.566  53.453  1.00  0.00           H
    ATOM    370 HG23 THR A 118      65.540  45.981  53.146  1.00  0.00           H
    ATOM    371  HG1 THR A 118      63.681  47.865  53.227  1.00  0.00           H
    ATOM    372  N   ALA A 119      62.626  46.046  48.798  1.00 26.72           N
    ATOM    373  CA  ALA A 119      61.488  46.085  47.871  1.00 25.07           C
    ATOM    374  C   ALA A 119      60.708  44.760  47.915  1.00 31.36           C
    ATOM    375  O   ALA A 119      61.286  43.723  48.244  1.00 36.22           O
    ATOM    376  CB  ALA A 119      61.965  46.381  46.442  1.00 21.56           C
    ATOM    377  H   ALA A 119      63.551  46.062  48.385  1.00  0.00           H
    ATOM    378  HA  ALA A 119      60.830  46.894  48.193  1.00  0.00           H
    ATOM    379  HB1 ALA A 119      61.120  46.593  45.787  1.00  0.00           H
    ATOM    380  HB2 ALA A 119      62.620  47.252  46.414  1.00  0.00           H
    ATOM    381  HB3 ALA A 119      62.498  45.534  46.009  1.00  0.00           H
    ATOM    382  N   LYS A 120      59.400  44.818  47.631  1.00 36.53           N
    ATOM    383  CA  LYS A 120      58.493  43.679  47.809  1.00 39.17           C
    ATOM    384  C   LYS A 120      58.805  42.451  46.933  1.00 34.12           C
    ATOM    385  O   LYS A 120      58.353  41.357  47.265  1.00 32.16           O
    ATOM    386  CB  LYS A 120      57.022  44.127  47.676  1.00 43.97           C
    ATOM    387  CG  LYS A 120      56.454  44.878  48.897  1.00 54.99           C
    ATOM    388  CD  LYS A 120      56.919  46.340  49.031  1.00 56.97           C
    ATOM    389  CE  LYS A 120      56.371  47.042  50.275  1.00 53.65           C
    ATOM    390  NZ  LYS A 120      55.040  47.626  50.059  1.00 52.46           N1+
    ATOM    391  H   LYS A 120      58.980  45.699  47.364  1.00  0.00           H
    ATOM    392  HA  LYS A 120      58.632  43.335  48.836  1.00  0.00           H
    ATOM    393  HB2 LYS A 120      56.873  44.705  46.762  1.00  0.00           H
    ATOM    394  HB3 LYS A 120      56.401  43.237  47.560  1.00  0.00           H
    ATOM    395  HG2 LYS A 120      55.367  44.874  48.817  1.00  0.00           H
    ATOM    396  HG3 LYS A 120      56.670  44.313  49.803  1.00  0.00           H
    ATOM    397  HD2 LYS A 120      58.004  46.402  49.061  1.00  0.00           H
    ATOM    398  HD3 LYS A 120      56.599  46.889  48.145  1.00  0.00           H
    ATOM    399  HE2 LYS A 120      56.286  46.327  51.089  1.00  0.00           H
    ATOM    400  HE3 LYS A 120      57.044  47.839  50.585  1.00  0.00           H
    ATOM    401  HZ1 LYS A 120      54.727  48.079  50.905  1.00  0.00           H
    ATOM    402  HZ2 LYS A 120      55.082  48.302  49.310  1.00  0.00           H
    ATOM    403  HZ3 LYS A 120      54.389  46.894  49.812  1.00  0.00           H
    ATOM    404  N   SER A 121      59.605  42.628  45.874  1.00 35.36           N
    ATOM    405  CA  SER A 121      60.063  41.561  44.993  1.00 38.69           C
    ATOM    406  C   SER A 121      61.286  40.776  45.502  1.00 37.94           C
    ATOM    407  O   SER A 121      61.638  39.789  44.857  1.00 38.23           O
    ATOM    408  CB  SER A 121      60.258  42.145  43.580  1.00 39.52           C
    ATOM    409  OG  SER A 121      61.016  43.339  43.592  1.00 41.87           O
    ATOM    410  H   SER A 121      59.962  43.550  45.664  1.00  0.00           H
    ATOM    411  HA  SER A 121      59.266  40.819  44.908  1.00  0.00           H
    ATOM    412  HB2 SER A 121      60.736  41.425  42.915  1.00  0.00           H
    ATOM    413  HB3 SER A 121      59.286  42.369  43.138  1.00  0.00           H
    ATOM    414  HG  SER A 121      61.250  43.560  42.686  1.00  0.00           H
    ATOM    415  N   VAL A 122      61.908  41.174  46.626  1.00 36.46           N
    ATOM    416  CA  VAL A 122      63.104  40.492  47.116  1.00 32.64           C
    ATOM    417  C   VAL A 122      62.820  39.082  47.680  1.00 27.48           C
    ATOM    418  O   VAL A 122      61.858  38.857  48.412  1.00 26.55           O
    ATOM    419  CB  VAL A 122      63.925  41.359  48.097  1.00 32.79           C
    ATOM    420  CG1 VAL A 122      65.078  40.608  48.765  1.00 35.42           C
    ATOM    421  CG2 VAL A 122      64.508  42.575  47.358  1.00 30.32           C
    ATOM    422  H   VAL A 122      61.589  41.983  47.143  1.00  0.00           H
    ATOM    423  HA  VAL A 122      63.748  40.357  46.242  1.00  0.00           H
    ATOM    424  HB  VAL A 122      63.267  41.739  48.881  1.00  0.00           H
    ATOM    425 HG11 VAL A 122      65.913  41.280  48.923  1.00  0.00           H
    ATOM    426 HG12 VAL A 122      64.774  40.155  49.710  1.00  0.00           H
    ATOM    427 HG13 VAL A 122      65.475  39.830  48.129  1.00  0.00           H
    ATOM    428 HG21 VAL A 122      65.040  43.230  48.047  1.00  0.00           H
    ATOM    429 HG22 VAL A 122      65.209  42.275  46.579  1.00  0.00           H
    ATOM    430 HG23 VAL A 122      63.725  43.170  46.890  1.00  0.00           H
    ATOM    431  N   THR A 123      63.692  38.165  47.256  1.00 18.13           N
    ATOM    432  CA  THR A 123      63.650  36.729  47.481  1.00 16.46           C
    ATOM    433  C   THR A 123      64.603  36.333  48.632  1.00 21.21           C
    ATOM    434  O   THR A 123      64.444  35.258  49.210  1.00 21.44           O
    ATOM    435  CB  THR A 123      64.111  36.068  46.142  1.00 19.18           C
    ATOM    436  OG1 THR A 123      63.032  35.350  45.589  1.00 32.51           O
    ATOM    437  CG2 THR A 123      65.382  35.193  46.133  1.00 19.81           C
    ATOM    438  H   THR A 123      64.442  38.472  46.651  1.00  0.00           H
    ATOM    439  HA  THR A 123      62.638  36.417  47.747  1.00  0.00           H
    ATOM    440  HB  THR A 123      64.305  36.849  45.403  1.00  0.00           H
    ATOM    441 HG21 THR A 123      65.560  34.786  45.139  1.00  0.00           H
    ATOM    442 HG22 THR A 123      66.267  35.776  46.395  1.00  0.00           H
    ATOM    443 HG23 THR A 123      65.319  34.353  46.822  1.00  0.00           H
    ATOM    444  HG1 THR A 123      62.807  34.626  46.178  1.00  0.00           H
    ATOM    445  N   CYS A 124      65.576  37.207  48.923  1.00 22.12           N
    ATOM    446  CA  CYS A 124      66.633  37.027  49.898  1.00 21.19           C
    ATOM    447  C   CYS A 124      67.228  38.412  50.178  1.00 25.06           C
    ATOM    448  O   CYS A 124      67.722  39.042  49.244  1.00 26.63           O
    ATOM    449  CB  CYS A 124      67.695  36.012  49.421  1.00 19.11           C
    ATOM    450  SG  CYS A 124      69.076  35.883  50.593  1.00 29.29           S
    ATOM    451  H   CYS A 124      65.632  38.065  48.392  1.00  0.00           H
    ATOM    452  HA  CYS A 124      66.179  36.641  50.802  1.00  0.00           H
    ATOM    453  HB2 CYS A 124      67.254  35.022  49.302  1.00  0.00           H
    ATOM    454  HB3 CYS A 124      68.097  36.299  48.448  1.00  0.00           H
    ATOM    455  HG  CYS A 124      68.352  35.410  51.612  1.00  0.00           H
    ATOM    456  N   THR A 125      67.192  38.856  51.441  1.00 27.20           N
    ATOM    457  CA  THR A 125      67.773  40.126  51.862  1.00 27.83           C
    ATOM    458  C   THR A 125      68.380  40.012  53.258  1.00 24.83           C
    ATOM    459  O   THR A 125      67.795  39.384  54.132  1.00 31.54           O
    ATOM    460  CB  THR A 125      66.741  41.278  51.855  1.00 26.93           C
    ATOM    461  OG1 THR A 125      67.370  42.499  52.154  1.00 27.56           O
    ATOM    462  CG2 THR A 125      65.529  41.121  52.787  1.00 20.04           C
    ATOM    463  H   THR A 125      66.791  38.278  52.171  1.00  0.00           H
    ATOM    464  HA  THR A 125      68.587  40.388  51.182  1.00  0.00           H
    ATOM    465  HB  THR A 125      66.370  41.405  50.857  1.00  0.00           H
    ATOM    466 HG21 THR A 125      64.815  41.930  52.634  1.00  0.00           H
    ATOM    467 HG22 THR A 125      65.005  40.189  52.589  1.00  0.00           H
    ATOM    468 HG23 THR A 125      65.810  41.134  53.839  1.00  0.00           H
    ATOM    469  HG1 THR A 125      66.941  43.172  51.611  1.00  0.00           H
    ATOM    470  N   TYR A 126      69.537  40.649  53.443  1.00 15.43           N
    ATOM    471  CA  TYR A 126      70.272  40.618  54.694  1.00 15.65           C
    ATOM    472  C   TYR A 126      70.230  41.987  55.375  1.00 14.00           C
    ATOM    473  O   TYR A 126      70.500  42.996  54.728  1.00 19.87           O
    ATOM    474  CB  TYR A 126      71.697  40.142  54.381  1.00 13.08           C
    ATOM    475  CG  TYR A 126      72.673  40.270  55.528  1.00 16.66           C
    ATOM    476  CD1 TYR A 126      72.778  39.250  56.495  1.00 21.24           C
    ATOM    477  CD2 TYR A 126      73.478  41.420  55.623  1.00 16.50           C
    ATOM    478  CE1 TYR A 126      73.702  39.378  57.548  1.00 18.09           C
    ATOM    479  CE2 TYR A 126      74.413  41.536  56.662  1.00 19.32           C
    ATOM    480  CZ  TYR A 126      74.526  40.519  57.624  1.00 16.96           C
    ATOM    481  OH  TYR A 126      75.449  40.644  58.617  1.00 18.16           O
    ATOM    482  H   TYR A 126      69.944  41.188  52.694  1.00  0.00           H
    ATOM    483  HA  TYR A 126      69.839  39.898  55.381  1.00  0.00           H
    ATOM    484  HB2 TYR A 126      71.675  39.104  54.044  1.00  0.00           H
    ATOM    485  HB3 TYR A 126      72.098  40.712  53.541  1.00  0.00           H
    ATOM    486  HD1 TYR A 126      72.160  38.367  56.428  1.00  0.00           H
    ATOM    487  HD2 TYR A 126      73.379  42.209  54.894  1.00  0.00           H
    ATOM    488  HE1 TYR A 126      73.788  38.589  58.280  1.00  0.00           H
    ATOM    489  HE2 TYR A 126      75.050  42.404  56.717  1.00  0.00           H
    ATOM    490  HH  TYR A 126      75.455  39.894  59.223  1.00  0.00           H
    ATOM    491  N   SER A 127      69.956  41.981  56.684  1.00 21.45           N
    ATOM    492  CA  SER A 127      70.018  43.152  57.545  1.00 21.70           C
    ATOM    493  C   SER A 127      71.367  43.167  58.285  1.00 24.32           C
    ATOM    494  O   SER A 127      71.515  42.378  59.220  1.00 27.26           O
    ATOM    495  CB  SER A 127      68.864  43.066  58.557  1.00 16.99           C
    ATOM    496  OG  SER A 127      69.016  44.061  59.551  1.00 15.95           O
    ATOM    497  H   SER A 127      69.755  41.102  57.148  1.00  0.00           H
    ATOM    498  HA  SER A 127      69.858  44.062  56.974  1.00  0.00           H
    ATOM    499  HB2 SER A 127      67.903  43.198  58.063  1.00  0.00           H
    ATOM    500  HB3 SER A 127      68.833  42.086  59.032  1.00  0.00           H
    ATOM    501  HG  SER A 127      68.679  44.888  59.183  1.00  0.00           H
    ATOM    502  N   PRO A 128      72.309  44.067  57.917  1.00 20.46           N
    ATOM    503  CA  PRO A 128      73.562  44.213  58.678  1.00 21.33           C
    ATOM    504  C   PRO A 128      73.411  44.642  60.151  1.00 24.11           C
    ATOM    505  O   PRO A 128      74.247  44.234  60.956  1.00 30.45           O
    ATOM    506  CB  PRO A 128      74.421  45.176  57.840  1.00 16.19           C
    ATOM    507  CG  PRO A 128      73.449  45.903  56.928  1.00 16.17           C
    ATOM    508  CD  PRO A 128      72.314  44.911  56.718  1.00 17.69           C
    ATOM    509  HA  PRO A 128      74.063  43.246  58.711  1.00  0.00           H
    ATOM    510  HB2 PRO A 128      75.026  45.862  58.434  1.00  0.00           H
    ATOM    511  HB3 PRO A 128      75.112  44.597  57.227  1.00  0.00           H
    ATOM    512  HG2 PRO A 128      73.061  46.773  57.460  1.00  0.00           H
    ATOM    513  HG3 PRO A 128      73.898  46.258  56.000  1.00  0.00           H
    ATOM    514  HD2 PRO A 128      71.383  45.444  56.532  1.00  0.00           H
    ATOM    515  HD3 PRO A 128      72.529  44.281  55.855  1.00  0.00           H
    ATOM    516  N   ALA A 129      72.351  45.393  60.499  1.00 20.24           N
    ATOM    517  CA  ALA A 129      72.075  45.802  61.881  1.00 22.97           C
    ATOM    518  C   ALA A 129      71.541  44.669  62.778  1.00 23.67           C
    ATOM    519  O   ALA A 129      71.781  44.706  63.984  1.00 24.66           O
    ATOM    520  CB  ALA A 129      71.107  46.991  61.888  1.00 16.64           C
    ATOM    521  H   ALA A 129      71.696  45.711  59.795  1.00  0.00           H
    ATOM    522  HA  ALA A 129      73.012  46.147  62.322  1.00  0.00           H
    ATOM    523  HB1 ALA A 129      70.924  47.344  62.905  1.00  0.00           H
    ATOM    524  HB2 ALA A 129      71.509  47.831  61.321  1.00  0.00           H
    ATOM    525  HB3 ALA A 129      70.142  46.724  61.455  1.00  0.00           H
    ATOM    526  N   LEU A 130      70.838  43.687  62.193  1.00 17.42           N
    ATOM    527  CA  LEU A 130      70.335  42.513  62.913  1.00 16.46           C
    ATOM    528  C   LEU A 130      71.298  41.318  62.804  1.00 17.58           C
    ATOM    529  O   LEU A 130      71.092  40.339  63.521  1.00 22.80           O
    ATOM    530  CB  LEU A 130      68.959  42.107  62.331  1.00 19.55           C
    ATOM    531  CG  LEU A 130      67.671  42.730  62.919  1.00 23.20           C
    ATOM    532  CD1 LEU A 130      67.868  43.963  63.814  1.00 23.15           C
    ATOM    533  CD2 LEU A 130      66.627  42.991  61.823  1.00 23.09           C
    ATOM    534  H   LEU A 130      70.644  43.726  61.201  1.00  0.00           H
    ATOM    535  HA  LEU A 130      70.228  42.717  63.979  1.00  0.00           H
    ATOM    536  HB2 LEU A 130      68.984  42.199  61.251  1.00  0.00           H
    ATOM    537  HB3 LEU A 130      68.812  41.046  62.505  1.00  0.00           H
    ATOM    538  HG  LEU A 130      67.228  41.979  63.574  1.00  0.00           H
    ATOM    539 HD11 LEU A 130      66.947  44.539  63.913  1.00  0.00           H
    ATOM    540 HD12 LEU A 130      68.173  43.677  64.821  1.00  0.00           H
    ATOM    541 HD13 LEU A 130      68.617  44.646  63.415  1.00  0.00           H
    ATOM    542 HD21 LEU A 130      65.757  43.505  62.223  1.00  0.00           H
    ATOM    543 HD22 LEU A 130      67.020  43.622  61.031  1.00  0.00           H
    ATOM    544 HD23 LEU A 130      66.286  42.058  61.374  1.00  0.00           H
    ATOM    545  N   ASN A 131      72.269  41.380  61.871  1.00 21.06           N
    ATOM    546  CA  ASN A 131      73.054  40.257  61.344  1.00 20.14           C
    ATOM    547  C   ASN A 131      72.129  39.101  60.920  1.00 18.07           C
    ATOM    548  O   ASN A 131      72.339  37.957  61.320  1.00 20.66           O
    ATOM    549  CB  ASN A 131      74.165  39.812  62.325  1.00 23.52           C
    ATOM    550  CG  ASN A 131      75.156  38.806  61.715  1.00 23.06           C
    ATOM    551  OD1 ASN A 131      75.243  38.646  60.499  1.00 24.05           O
    ATOM    552  ND2 ASN A 131      75.887  38.096  62.566  1.00 18.79           N
    ATOM    553  H   ASN A 131      72.357  42.237  61.343  1.00  0.00           H
    ATOM    554  HA  ASN A 131      73.516  40.624  60.429  1.00  0.00           H
    ATOM    555  HB2 ASN A 131      74.742  40.684  62.638  1.00  0.00           H
    ATOM    556  HB3 ASN A 131      73.723  39.389  63.228  1.00  0.00           H
    ATOM    557 HD21 ASN A 131      75.804  38.250  63.562  1.00  0.00           H
    ATOM    558 HD22 ASN A 131      76.529  37.392  62.228  1.00  0.00           H
    ATOM    559  N   LYS A 132      71.074  39.452  60.177  1.00 20.17           N
    ATOM    560  CA  LYS A 132      69.963  38.557  59.934  1.00 16.46           C
    ATOM    561  C   LYS A 132      69.608  38.528  58.460  1.00 18.54           C
    ATOM    562  O   LYS A 132      69.262  39.560  57.893  1.00 17.47           O
    ATOM    563  CB  LYS A 132      68.784  38.986  60.802  1.00 12.94           C
    ATOM    564  CG  LYS A 132      67.570  38.064  60.714  1.00 12.83           C
    ATOM    565  CD  LYS A 132      66.567  38.414  61.811  1.00 13.14           C
    ATOM    566  CE  LYS A 132      65.389  37.450  61.803  1.00 18.24           C
    ATOM    567  NZ  LYS A 132      64.492  37.695  62.934  1.00 22.26           N1+
    ATOM    568  H   LYS A 132      70.969  40.409  59.867  1.00  0.00           H
    ATOM    569  HA  LYS A 132      70.225  37.554  60.245  1.00  0.00           H
    ATOM    570  HB2 LYS A 132      69.127  38.995  61.837  1.00  0.00           H
    ATOM    571  HB3 LYS A 132      68.478  40.000  60.546  1.00  0.00           H
    ATOM    572  HG2 LYS A 132      67.096  38.140  59.736  1.00  0.00           H
    ATOM    573  HG3 LYS A 132      67.888  37.027  60.824  1.00  0.00           H
    ATOM    574  HD2 LYS A 132      67.058  38.392  62.784  1.00  0.00           H
    ATOM    575  HD3 LYS A 132      66.216  39.437  61.679  1.00  0.00           H
    ATOM    576  HE2 LYS A 132      64.827  37.576  60.879  1.00  0.00           H
    ATOM    577  HE3 LYS A 132      65.742  36.421  61.844  1.00  0.00           H
    ATOM    578  HZ1 LYS A 132      63.704  37.060  62.907  1.00  0.00           H
    ATOM    579  HZ2 LYS A 132      65.007  37.544  63.793  1.00  0.00           H
    ATOM    580  HZ3 LYS A 132      64.154  38.654  62.907  1.00  0.00           H
    ATOM    581  N   MET A 133      69.647  37.319  57.896  1.00 22.59           N
    ATOM    582  CA  MET A 133      69.169  37.035  56.559  1.00 20.70           C
    ATOM    583  C   MET A 133      67.673  36.730  56.608  1.00 22.00           C
    ATOM    584  O   MET A 133      67.225  35.929  57.416  1.00 23.27           O
    ATOM    585  CB  MET A 133      69.959  35.862  55.970  1.00 21.84           C
    ATOM    586  CG  MET A 133      69.727  35.704  54.462  1.00 23.20           C
    ATOM    587  SD  MET A 133      70.465  34.209  53.775  1.00 31.82           S
    ATOM    588  CE  MET A 133      69.213  33.036  54.312  1.00 40.13           C
    ATOM    589  H   MET A 133      69.920  36.518  58.454  1.00  0.00           H
    ATOM    590  HA  MET A 133      69.344  37.898  55.921  1.00  0.00           H
    ATOM    591  HB2 MET A 133      71.025  36.030  56.131  1.00  0.00           H
    ATOM    592  HB3 MET A 133      69.721  34.936  56.495  1.00  0.00           H
    ATOM    593  HG2 MET A 133      68.669  35.724  54.203  1.00  0.00           H
    ATOM    594  HG3 MET A 133      70.174  36.553  53.944  1.00  0.00           H
    ATOM    595  HE1 MET A 133      68.935  32.367  53.507  1.00  0.00           H
    ATOM    596  HE2 MET A 133      69.599  32.458  55.149  1.00  0.00           H
    ATOM    597  HE3 MET A 133      68.317  33.556  54.642  1.00  0.00           H
    ATOM    598  N   PHE A 134      66.928  37.358  55.709  1.00 17.76           N
    ATOM    599  CA  PHE A 134      65.528  37.084  55.459  1.00 17.02           C
    ATOM    600  C   PHE A 134      65.476  36.449  54.081  1.00 21.15           C
    ATOM    601  O   PHE A 134      66.110  36.965  53.168  1.00 24.19           O
    ATOM    602  CB  PHE A 134      64.733  38.393  55.488  1.00 17.50           C
    ATOM    603  CG  PHE A 134      64.678  39.051  56.848  1.00 19.48           C
    ATOM    604  CD1 PHE A 134      65.770  39.808  57.325  1.00 20.82           C
    ATOM    605  CD2 PHE A 134      63.589  38.797  57.702  1.00 17.67           C
    ATOM    606  CE1 PHE A 134      65.729  40.351  58.600  1.00 20.55           C
    ATOM    607  CE2 PHE A 134      63.564  39.355  58.971  1.00 19.94           C
    ATOM    608  CZ  PHE A 134      64.625  40.133  59.414  1.00 18.95           C
    ATOM    609  H   PHE A 134      67.357  38.028  55.079  1.00  0.00           H
    ATOM    610  HA  PHE A 134      65.110  36.412  56.204  1.00  0.00           H
    ATOM    611  HB2 PHE A 134      65.137  39.105  54.771  1.00  0.00           H
    ATOM    612  HB3 PHE A 134      63.708  38.196  55.178  1.00  0.00           H
    ATOM    613  HD1 PHE A 134      66.634  39.976  56.699  1.00  0.00           H
    ATOM    614  HD2 PHE A 134      62.763  38.192  57.361  1.00  0.00           H
    ATOM    615  HE1 PHE A 134      66.556  40.944  58.961  1.00  0.00           H
    ATOM    616  HE2 PHE A 134      62.713  39.193  59.613  1.00  0.00           H
    ATOM    617  HZ  PHE A 134      64.596  40.572  60.399  1.00  0.00           H
    ATOM    618  N   CYS A 135      64.738  35.351  53.939  1.00 22.23           N
    ATOM    619  CA  CYS A 135      64.676  34.636  52.675  1.00 23.20           C
    ATOM    620  C   CYS A 135      63.326  33.961  52.510  1.00 25.02           C
    ATOM    621  O   CYS A 135      62.667  33.642  53.492  1.00 31.56           O
    ATOM    622  CB  CYS A 135      65.852  33.657  52.515  1.00 25.44           C
    ATOM    623  SG  CYS A 135      65.869  32.379  53.810  1.00 23.00           S
    ATOM    624  H   CYS A 135      64.234  34.952  54.721  1.00  0.00           H
    ATOM    625  HA  CYS A 135      64.727  35.361  51.862  1.00  0.00           H
    ATOM    626  HB2 CYS A 135      65.779  33.164  51.546  1.00  0.00           H
    ATOM    627  HB3 CYS A 135      66.802  34.192  52.528  1.00  0.00           H
    ATOM    628  HG  CYS A 135      64.812  31.701  53.353  1.00  0.00           H
    ATOM    629  N   GLN A 136      62.945  33.755  51.248  1.00 20.92           N
    ATOM    630  CA  GLN A 136      61.807  32.926  50.889  1.00 20.93           C
    ATOM    631  C   GLN A 136      62.167  31.454  50.969  1.00 22.57           C
    ATOM    632  O   GLN A 136      63.324  31.098  50.758  1.00 21.18           O
    ATOM    633  CB  GLN A 136      61.336  33.278  49.478  1.00 30.06           C
    ATOM    634  CG  GLN A 136      60.274  34.378  49.474  1.00 35.63           C
    ATOM    635  CD  GLN A 136      59.999  34.875  48.060  1.00 35.99           C
    ATOM    636  OE1 GLN A 136      60.162  34.142  47.087  1.00 36.49           O
    ATOM    637  NE2 GLN A 136      59.583  36.135  47.939  1.00 35.11           N
    ATOM    638  H   GLN A 136      63.521  34.086  50.485  1.00  0.00           H
    ATOM    639  HA  GLN A 136      60.989  33.092  51.582  1.00  0.00           H
    ATOM    640  HB2 GLN A 136      62.197  33.579  48.877  1.00  0.00           H
    ATOM    641  HB3 GLN A 136      60.909  32.402  48.984  1.00  0.00           H
    ATOM    642  HG2 GLN A 136      59.348  34.025  49.928  1.00  0.00           H
    ATOM    643  HG3 GLN A 136      60.629  35.214  50.074  1.00  0.00           H
    ATOM    644 HE21 GLN A 136      59.447  36.712  48.757  1.00  0.00           H
    ATOM    645 HE22 GLN A 136      59.395  36.518  47.024  1.00  0.00           H
    ATOM    646  N   LEU A 137      61.142  30.630  51.212  1.00 20.07           N
    ATOM    647  CA  LEU A 137      61.249  29.180  51.194  1.00 19.48           C
    ATOM    648  C   LEU A 137      61.714  28.706  49.809  1.00 24.24           C
    ATOM    649  O   LEU A 137      61.090  29.052  48.807  1.00 24.32           O
    ATOM    650  CB  LEU A 137      59.880  28.567  51.554  1.00 18.38           C
    ATOM    651  CG  LEU A 137      59.929  27.182  52.229  1.00 24.39           C
    ATOM    652  CD1 LEU A 137      58.512  26.622  52.425  1.00 32.39           C
    ATOM    653  CD2 LEU A 137      60.771  26.156  51.476  1.00 27.00           C
    ATOM    654  H   LEU A 137      60.218  31.006  51.378  1.00  0.00           H
    ATOM    655  HA  LEU A 137      61.981  28.900  51.954  1.00  0.00           H
    ATOM    656  HB2 LEU A 137      59.371  29.218  52.255  1.00  0.00           H
    ATOM    657  HB3 LEU A 137      59.236  28.537  50.674  1.00  0.00           H
    ATOM    658  HG  LEU A 137      60.378  27.310  53.215  1.00  0.00           H
    ATOM    659 HD11 LEU A 137      58.344  26.379  53.474  1.00  0.00           H
    ATOM    660 HD12 LEU A 137      57.739  27.333  52.130  1.00  0.00           H
    ATOM    661 HD13 LEU A 137      58.337  25.709  51.854  1.00  0.00           H
    ATOM    662 HD21 LEU A 137      60.583  25.143  51.832  1.00  0.00           H
    ATOM    663 HD22 LEU A 137      60.534  26.167  50.414  1.00  0.00           H
    ATOM    664 HD23 LEU A 137      61.826  26.351  51.609  1.00  0.00           H
    ATOM    665  N   ALA A 138      62.794  27.914  49.802  1.00 20.43           N
    ATOM    666  CA  ALA A 138      63.342  27.206  48.647  1.00 24.58           C
    ATOM    667  C   ALA A 138      64.004  28.088  47.576  1.00 29.90           C
    ATOM    668  O   ALA A 138      64.357  27.555  46.526  1.00 37.65           O
    ATOM    669  CB  ALA A 138      62.281  26.274  48.021  1.00 20.92           C
    ATOM    670  H   ALA A 138      63.248  27.709  50.683  1.00  0.00           H
    ATOM    671  HA  ALA A 138      64.139  26.579  49.044  1.00  0.00           H
    ATOM    672  HB1 ALA A 138      62.710  25.662  47.228  1.00  0.00           H
    ATOM    673  HB2 ALA A 138      61.890  25.577  48.759  1.00  0.00           H
    ATOM    674  HB3 ALA A 138      61.442  26.817  47.587  1.00  0.00           H
    ATOM    675  N   LYS A 139      64.172  29.396  47.825  1.00 21.62           N
    ATOM    676  CA  LYS A 139      64.790  30.309  46.863  1.00 18.52           C
    ATOM    677  C   LYS A 139      66.287  30.480  47.149  1.00 18.36           C
    ATOM    678  O   LYS A 139      66.751  30.173  48.249  1.00 20.58           O
    ATOM    679  CB  LYS A 139      64.055  31.659  46.886  1.00 20.28           C
    ATOM    680  CG  LYS A 139      62.546  31.606  46.577  1.00 31.42           C
    ATOM    681  CD  LYS A 139      62.139  31.387  45.108  1.00 40.22           C
    ATOM    682  CE  LYS A 139      62.247  29.945  44.584  1.00 52.90           C
    ATOM    683  NZ  LYS A 139      63.287  29.806  43.549  1.00 59.12           N1+
    ATOM    684  H   LYS A 139      63.867  29.789  48.704  1.00  0.00           H
    ATOM    685  HA  LYS A 139      64.717  29.907  45.854  1.00  0.00           H
    ATOM    686  HB2 LYS A 139      64.183  32.108  47.873  1.00  0.00           H
    ATOM    687  HB3 LYS A 139      64.526  32.343  46.180  1.00  0.00           H
    ATOM    688  HG2 LYS A 139      62.037  30.888  47.219  1.00  0.00           H
    ATOM    689  HG3 LYS A 139      62.144  32.574  46.859  1.00  0.00           H
    ATOM    690  HD2 LYS A 139      61.103  31.713  44.999  1.00  0.00           H
    ATOM    691  HD3 LYS A 139      62.712  32.068  44.476  1.00  0.00           H
    ATOM    692  HE2 LYS A 139      62.416  29.234  45.391  1.00  0.00           H
    ATOM    693  HE3 LYS A 139      61.302  29.663  44.121  1.00  0.00           H
    ATOM    694  HZ1 LYS A 139      64.187  30.050  43.937  1.00  0.00           H
    ATOM    695  HZ2 LYS A 139      63.313  28.850  43.225  1.00  0.00           H
    ATOM    696  HZ3 LYS A 139      63.079  30.420  42.774  1.00  0.00           H
    ATOM    697  N   THR A 140      67.022  30.964  46.138  1.00 20.22           N
    ATOM    698  CA  THR A 140      68.464  31.172  46.222  1.00 20.89           C
    ATOM    699  C   THR A 140      68.837  32.228  47.278  1.00 22.30           C
    ATOM    700  O   THR A 140      68.252  33.313  47.311  1.00 22.07           O
    ATOM    701  CB  THR A 140      69.059  31.613  44.863  1.00 25.60           C
    ATOM    702  OG1 THR A 140      68.766  30.639  43.883  1.00 28.60           O
    ATOM    703  CG2 THR A 140      70.576  31.848  44.856  1.00 24.79           C
    ATOM    704  H   THR A 140      66.594  31.199  45.254  1.00  0.00           H
    ATOM    705  HA  THR A 140      68.916  30.217  46.499  1.00  0.00           H
    ATOM    706  HB  THR A 140      68.578  32.537  44.542  1.00  0.00           H
    ATOM    707 HG21 THR A 140      70.946  32.003  43.842  1.00  0.00           H
    ATOM    708 HG22 THR A 140      70.866  32.720  45.431  1.00  0.00           H
    ATOM    709 HG23 THR A 140      71.097  30.990  45.275  1.00  0.00           H
    ATOM    710  HG1 THR A 140      69.113  29.778  44.154  1.00  0.00           H
    ATOM    711  N   CYS A 141      69.810  31.855  48.116  1.00 22.79           N
    ATOM    712  CA  CYS A 141      70.368  32.675  49.178  1.00 19.44           C
    ATOM    713  C   CYS A 141      71.879  32.672  48.964  1.00 19.36           C
    ATOM    714  O   CYS A 141      72.536  31.746  49.441  1.00 25.40           O
    ATOM    715  CB  CYS A 141      70.067  32.110  50.580  1.00 18.51           C
    ATOM    716  SG  CYS A 141      68.292  32.061  50.862  1.00 27.07           S
    ATOM    717  H   CYS A 141      70.191  30.921  48.048  1.00  0.00           H
    ATOM    718  HA  CYS A 141      69.979  33.688  49.130  1.00  0.00           H
    ATOM    719  HB2 CYS A 141      70.461  31.107  50.731  1.00  0.00           H
    ATOM    720  HB3 CYS A 141      70.520  32.742  51.342  1.00  0.00           H
    ATOM    721  HG  CYS A 141      68.034  31.130  49.937  1.00  0.00           H
    ATOM    722  N   PRO A 142      72.429  33.665  48.237  1.00 23.31           N
    ATOM    723  CA  PRO A 142      73.883  33.726  48.087  1.00 21.79           C
    ATOM    724  C   PRO A 142      74.555  34.154  49.399  1.00 26.02           C
    ATOM    725  O   PRO A 142      74.050  35.035  50.090  1.00 29.19           O
    ATOM    726  CB  PRO A 142      74.091  34.710  46.939  1.00 17.72           C
    ATOM    727  CG  PRO A 142      72.871  35.614  46.949  1.00 15.70           C
    ATOM    728  CD  PRO A 142      71.753  34.764  47.546  1.00 17.05           C
    ATOM    729  HA  PRO A 142      74.270  32.760  47.772  1.00  0.00           H
    ATOM    730  HB2 PRO A 142      75.027  35.270  46.997  1.00  0.00           H
    ATOM    731  HB3 PRO A 142      74.107  34.147  46.006  1.00  0.00           H
    ATOM    732  HG2 PRO A 142      73.065  36.463  47.606  1.00  0.00           H
    ATOM    733  HG3 PRO A 142      72.622  36.001  45.962  1.00  0.00           H
    ATOM    734  HD2 PRO A 142      71.141  35.364  48.221  1.00  0.00           H
    ATOM    735  HD3 PRO A 142      71.104  34.377  46.765  1.00  0.00           H
    ATOM    736  N   VAL A 143      75.659  33.487  49.739  1.00 20.65           N
    ATOM    737  CA  VAL A 143      76.480  33.808  50.895  1.00 18.54           C
    ATOM    738  C   VAL A 143      77.930  33.817  50.421  1.00 19.05           C
    ATOM    739  O   VAL A 143      78.392  32.836  49.836  1.00 22.91           O
    ATOM    740  CB  VAL A 143      76.298  32.786  52.048  1.00 22.24           C
    ATOM    741  CG1 VAL A 143      77.351  32.897  53.172  1.00 25.56           C
    ATOM    742  CG2 VAL A 143      74.887  32.894  52.656  1.00 17.35           C
    ATOM    743  H   VAL A 143      75.984  32.722  49.161  1.00  0.00           H
    ATOM    744  HA  VAL A 143      76.239  34.806  51.273  1.00  0.00           H
    ATOM    745  HB  VAL A 143      76.389  31.790  51.617  1.00  0.00           H
    ATOM    746 HG11 VAL A 143      77.113  32.228  53.999  1.00  0.00           H
    ATOM    747 HG12 VAL A 143      78.351  32.624  52.833  1.00  0.00           H
    ATOM    748 HG13 VAL A 143      77.398  33.911  53.574  1.00  0.00           H
    ATOM    749 HG21 VAL A 143      74.757  32.188  53.475  1.00  0.00           H
    ATOM    750 HG22 VAL A 143      74.699  33.894  53.044  1.00  0.00           H
    ATOM    751 HG23 VAL A 143      74.114  32.676  51.919  1.00  0.00           H
    ATOM    752  N   GLN A 144      78.603  34.946  50.670  1.00 21.46           N
    ATOM    753  CA  GLN A 144      79.954  35.178  50.192  1.00 23.53           C
    ATOM    754  C   GLN A 144      81.008  34.876  51.240  1.00 25.85           C
    ATOM    755  O   GLN A 144      80.897  35.346  52.363  1.00 29.75           O
    ATOM    756  CB  GLN A 144      80.088  36.597  49.640  1.00 25.26           C
    ATOM    757  CG  GLN A 144      79.412  36.689  48.261  1.00 30.76           C
    ATOM    758  CD  GLN A 144      78.244  37.644  48.250  1.00 36.33           C
    ATOM    759  OE1 GLN A 144      77.106  37.199  48.354  1.00 31.90           O
    ATOM    760  NE2 GLN A 144      78.521  38.942  48.124  1.00 38.52           N
    ATOM    761  H   GLN A 144      78.165  35.704  51.178  1.00  0.00           H
    ATOM    762  HA  GLN A 144      80.128  34.506  49.361  1.00  0.00           H
    ATOM    763  HB2 GLN A 144      79.699  37.331  50.343  1.00  0.00           H
    ATOM    764  HB3 GLN A 144      81.144  36.842  49.520  1.00  0.00           H
    ATOM    765  HG2 GLN A 144      80.086  36.974  47.468  1.00  0.00           H
    ATOM    766  HG3 GLN A 144      79.032  35.719  47.939  1.00  0.00           H
    ATOM    767 HE21 GLN A 144      79.475  39.253  48.016  1.00  0.00           H
    ATOM    768 HE22 GLN A 144      77.773  39.620  48.113  1.00  0.00           H
    ATOM    769  N   LEU A 145      82.032  34.121  50.837  1.00 24.45           N
    ATOM    770  CA  LEU A 145      83.203  33.865  51.656  1.00 17.74           C
    ATOM    771  C   LEU A 145      84.259  34.873  51.240  1.00 20.78           C
    ATOM    772  O   LEU A 145      84.658  34.908  50.077  1.00 23.02           O
    ATOM    773  CB  LEU A 145      83.756  32.448  51.440  1.00 17.56           C
    ATOM    774  CG  LEU A 145      82.854  31.247  51.809  1.00 17.95           C
    ATOM    775  CD1 LEU A 145      81.450  31.554  52.368  1.00 14.34           C
    ATOM    776  CD2 LEU A 145      82.775  30.267  50.638  1.00 17.92           C
    ATOM    777  H   LEU A 145      82.053  33.766  49.888  1.00  0.00           H
    ATOM    778  HA  LEU A 145      82.984  33.992  52.718  1.00  0.00           H
    ATOM    779  HB2 LEU A 145      84.088  32.354  50.410  1.00  0.00           H
    ATOM    780  HB3 LEU A 145      84.676  32.351  52.022  1.00  0.00           H
    ATOM    781  HG  LEU A 145      83.377  30.718  52.608  1.00  0.00           H
    ATOM    782 HD11 LEU A 145      81.098  30.724  52.982  1.00  0.00           H
    ATOM    783 HD12 LEU A 145      81.427  32.432  53.008  1.00  0.00           H
    ATOM    784 HD13 LEU A 145      80.719  31.694  51.570  1.00  0.00           H
    ATOM    785 HD21 LEU A 145      82.372  29.321  50.976  1.00  0.00           H
    ATOM    786 HD22 LEU A 145      82.121  30.642  49.849  1.00  0.00           H
    ATOM    787 HD23 LEU A 145      83.754  30.071  50.199  1.00  0.00           H
    ATOM    788  N   TRP A 146      84.701  35.645  52.226  1.00 30.23           N
    ATOM    789  CA  TRP A 146      85.808  36.569  52.113  1.00 30.30           C
    ATOM    790  C   TRP A 146      86.966  35.978  52.883  1.00 28.42           C
    ATOM    791  O   TRP A 146      86.776  35.457  53.977  1.00 28.31           O
    ATOM    792  CB  TRP A 146      85.393  37.917  52.687  1.00 37.69           C
    ATOM    793  CG  TRP A 146      84.384  38.598  51.839  1.00 52.88           C
    ATOM    794  CD1 TRP A 146      83.056  38.358  51.840  1.00 59.08           C
    ATOM    795  CD2 TRP A 146      84.632  39.522  50.751  1.00 56.67           C
    ATOM    796  NE1 TRP A 146      82.461  39.086  50.835  1.00 60.65           N
    ATOM    797  CE2 TRP A 146      83.381  39.841  50.146  1.00 58.53           C
    ATOM    798  CE3 TRP A 146      85.792  40.113  50.208  1.00 57.95           C
    ATOM    799  CZ2 TRP A 146      83.278  40.739  49.077  1.00 60.70           C
    ATOM    800  CZ3 TRP A 146      85.699  41.032  49.149  1.00 59.69           C
    ATOM    801  CH2 TRP A 146      84.444  41.358  48.601  1.00 61.66           C
    ATOM    802  H   TRP A 146      84.321  35.535  53.158  1.00  0.00           H
    ATOM    803  HA  TRP A 146      86.105  36.714  51.073  1.00  0.00           H
    ATOM    804  HB2 TRP A 146      84.977  37.789  53.678  1.00  0.00           H
    ATOM    805  HB3 TRP A 146      86.250  38.583  52.780  1.00  0.00           H
    ATOM    806  HD1 TRP A 146      82.565  37.668  52.509  1.00  0.00           H
    ATOM    807  HE1 TRP A 146      81.472  39.043  50.636  1.00  0.00           H
    ATOM    808  HE3 TRP A 146      86.756  39.874  50.626  1.00  0.00           H
    ATOM    809  HZ2 TRP A 146      82.317  40.965  48.638  1.00  0.00           H
    ATOM    810  HZ3 TRP A 146      86.595  41.497  48.765  1.00  0.00           H
    ATOM    811  HH2 TRP A 146      84.367  42.078  47.800  1.00  0.00           H
    ATOM    812  N   VAL A 147      88.143  36.068  52.276  1.00 23.15           N
    ATOM    813  CA  VAL A 147      89.353  35.499  52.820  1.00 25.21           C
    ATOM    814  C   VAL A 147      90.499  36.424  52.373  1.00 33.92           C
    ATOM    815  O   VAL A 147      90.470  36.954  51.261  1.00 40.96           O
    ATOM    816  CB  VAL A 147      89.470  34.013  52.353  1.00 24.79           C
    ATOM    817  CG1 VAL A 147      90.310  33.776  51.088  1.00 23.05           C
    ATOM    818  CG2 VAL A 147      89.867  33.074  53.491  1.00 27.52           C
    ATOM    819  H   VAL A 147      88.221  36.488  51.358  1.00  0.00           H
    ATOM    820  HA  VAL A 147      89.315  35.540  53.908  1.00  0.00           H
    ATOM    821  HB  VAL A 147      88.467  33.678  52.081  1.00  0.00           H
    ATOM    822 HG11 VAL A 147      90.311  32.727  50.805  1.00  0.00           H
    ATOM    823 HG12 VAL A 147      89.877  34.315  50.248  1.00  0.00           H
    ATOM    824 HG13 VAL A 147      91.339  34.106  51.188  1.00  0.00           H
    ATOM    825 HG21 VAL A 147      89.654  32.035  53.242  1.00  0.00           H
    ATOM    826 HG22 VAL A 147      90.929  33.161  53.652  1.00  0.00           H
    ATOM    827 HG23 VAL A 147      89.344  33.301  54.421  1.00  0.00           H
    ATOM    828  N   ASP A 148      91.450  36.662  53.281  1.00 34.78           N
    ATOM    829  CA  ASP A 148      92.624  37.522  53.081  1.00 32.86           C
    ATOM    830  C   ASP A 148      93.783  36.829  52.390  1.00 38.25           C
    ATOM    831  O   ASP A 148      94.610  37.491  51.765  1.00 45.00           O
    ATOM    832  CB  ASP A 148      93.137  38.100  54.413  1.00 34.48           C
    ATOM    833  CG  ASP A 148      92.219  39.176  54.967  1.00 37.05           C
    ATOM    834  OD1 ASP A 148      91.542  39.830  54.143  1.00 41.76           O
    ATOM    835  OD2 ASP A 148      92.246  39.380  56.198  1.00 38.18           O1-
    ATOM    836  H   ASP A 148      91.403  36.185  54.173  1.00  0.00           H
    ATOM    837  HA  ASP A 148      92.347  38.335  52.407  1.00  0.00           H
    ATOM    838  HB2 ASP A 148      93.196  37.292  55.142  1.00  0.00           H
    ATOM    839  HB3 ASP A 148      94.135  38.530  54.313  1.00  0.00           H
    ATOM    840  N   SER A 149      93.809  35.509  52.533  1.00 36.07           N
    ATOM    841  CA  SER A 149      94.768  34.616  51.933  1.00 39.91           C
    ATOM    842  C   SER A 149      94.006  33.337  51.622  1.00 40.28           C
    ATOM    843  O   SER A 149      93.162  32.944  52.426  1.00 38.21           O
    ATOM    844  CB  SER A 149      95.955  34.425  52.892  1.00 43.58           C
    ATOM    845  OG  SER A 149      95.543  33.903  54.135  1.00 48.69           O
    ATOM    846  H   SER A 149      93.090  35.059  53.082  1.00  0.00           H
    ATOM    847  HA  SER A 149      95.113  35.070  51.010  1.00  0.00           H
    ATOM    848  HB2 SER A 149      96.691  33.750  52.454  1.00  0.00           H
    ATOM    849  HB3 SER A 149      96.464  35.374  53.066  1.00  0.00           H
    ATOM    850  HG  SER A 149      95.270  32.992  54.006  1.00  0.00           H
    ATOM    851  N   THR A 150      94.268  32.714  50.466  1.00 41.46           N
    ATOM    852  CA  THR A 150      93.508  31.529  50.074  1.00 39.80           C
    ATOM    853  C   THR A 150      93.860  30.331  50.988  1.00 33.79           C
    ATOM    854  O   THR A 150      95.044  30.053  51.197  1.00 36.07           O
    ATOM    855  CB  THR A 150      93.709  31.132  48.593  1.00 42.10           C
    ATOM    856  OG1 THR A 150      95.073  31.040  48.230  1.00 46.49           O
    ATOM    857  CG2 THR A 150      93.009  32.124  47.652  1.00 39.65           C
    ATOM    858  H   THR A 150      94.968  33.069  49.827  1.00  0.00           H
    ATOM    859  HA  THR A 150      92.464  31.824  50.149  1.00  0.00           H
    ATOM    860  HB  THR A 150      93.269  30.148  48.422  1.00  0.00           H
    ATOM    861 HG21 THR A 150      93.130  31.827  46.609  1.00  0.00           H
    ATOM    862 HG22 THR A 150      91.937  32.171  47.853  1.00  0.00           H
    ATOM    863 HG23 THR A 150      93.411  33.132  47.758  1.00  0.00           H
    ATOM    864  HG1 THR A 150      95.451  31.923  48.242  1.00  0.00           H
    ATOM    865  N   PRO A 151      92.818  29.672  51.551  1.00 27.23           N
    ATOM    866  CA  PRO A 151      92.964  28.415  52.296  1.00 22.88           C
    ATOM    867  C   PRO A 151      93.729  27.324  51.526  1.00 32.12           C
    ATOM    868  O   PRO A 151      93.830  27.395  50.298  1.00 32.02           O
    ATOM    869  CB  PRO A 151      91.512  27.969  52.541  1.00 17.57           C
    ATOM    870  CG  PRO A 151      90.689  29.231  52.534  1.00 18.70           C
    ATOM    871  CD  PRO A 151      91.430  30.128  51.562  1.00 24.16           C
    ATOM    872  HA  PRO A 151      93.454  28.660  53.239  1.00  0.00           H
    ATOM    873  HB2 PRO A 151      91.165  27.321  51.733  1.00  0.00           H
    ATOM    874  HB3 PRO A 151      91.380  27.431  53.477  1.00  0.00           H
    ATOM    875  HG2 PRO A 151      89.640  29.077  52.281  1.00  0.00           H
    ATOM    876  HG3 PRO A 151      90.732  29.677  53.529  1.00  0.00           H
    ATOM    877  HD2 PRO A 151      91.029  30.061  50.557  1.00  0.00           H
    ATOM    878  HD3 PRO A 151      91.353  31.156  51.907  1.00  0.00           H
    ATOM    879  N   PRO A 152      94.208  26.299  52.260  1.00 34.57           N
    ATOM    880  CA  PRO A 152      94.902  25.166  51.640  1.00 39.45           C
    ATOM    881  C   PRO A 152      93.969  24.353  50.729  1.00 42.99           C
    ATOM    882  O   PRO A 152      92.748  24.514  50.798  1.00 42.47           O
    ATOM    883  CB  PRO A 152      95.417  24.355  52.843  1.00 41.28           C
    ATOM    884  CG  PRO A 152      94.498  24.707  54.000  1.00 44.15           C
    ATOM    885  CD  PRO A 152      94.087  26.140  53.707  1.00 38.18           C
    ATOM    886  HA  PRO A 152      95.738  25.527  51.044  1.00  0.00           H
    ATOM    887  HB2 PRO A 152      95.453  23.279  52.671  1.00  0.00           H
    ATOM    888  HB3 PRO A 152      96.432  24.678  53.076  1.00  0.00           H
    ATOM    889  HG2 PRO A 152      93.620  24.061  53.971  1.00  0.00           H
    ATOM    890  HG3 PRO A 152      94.965  24.585  54.978  1.00  0.00           H
    ATOM    891  HD2 PRO A 152      93.103  26.345  54.119  1.00  0.00           H
    ATOM    892  HD3 PRO A 152      94.798  26.826  54.156  1.00  0.00           H
    ATOM    893  N   PRO A 153      94.544  23.478  49.886  1.00 50.13           N
    ATOM    894  CA  PRO A 153      93.717  22.604  49.053  1.00 51.13           C
    ATOM    895  C   PRO A 153      93.030  21.527  49.903  1.00 48.47           C
    ATOM    896  O   PRO A 153      93.604  21.049  50.883  1.00 47.51           O
    ATOM    897  CB  PRO A 153      94.708  22.033  48.035  1.00 52.89           C
    ATOM    898  CG  PRO A 153      96.047  22.043  48.752  1.00 52.04           C
    ATOM    899  CD  PRO A 153      95.973  23.277  49.643  1.00 53.61           C
    ATOM    900  HA  PRO A 153      92.957  23.185  48.523  1.00  0.00           H
    ATOM    901  HB2 PRO A 153      94.436  21.043  47.664  1.00  0.00           H
    ATOM    902  HB3 PRO A 153      94.757  22.699  47.172  1.00  0.00           H
    ATOM    903  HG2 PRO A 153      96.107  21.157  49.387  1.00  0.00           H
    ATOM    904  HG3 PRO A 153      96.908  22.029  48.085  1.00  0.00           H
    ATOM    905  HD2 PRO A 153      96.562  23.153  50.552  1.00  0.00           H
    ATOM    906  HD3 PRO A 153      96.354  24.146  49.105  1.00  0.00           H
    ATOM    907  N   GLY A 154      91.791  21.204  49.517  1.00 43.87           N
    ATOM    908  CA  GLY A 154      90.946  20.259  50.234  1.00 35.44           C
    ATOM    909  C   GLY A 154      90.136  20.957  51.334  1.00 25.39           C
    ATOM    910  O   GLY A 154      89.369  20.282  52.020  1.00 26.72           O
    ATOM    911  H   GLY A 154      91.391  21.647  48.702  1.00  0.00           H
    ATOM    912  HA2 GLY A 154      90.256  19.804  49.525  1.00  0.00           H
    ATOM    913  HA3 GLY A 154      91.537  19.446  50.648  1.00  0.00           H
    ATOM    914  N   THR A 155      90.266  22.289  51.490  1.00 24.66           N
    ATOM    915  CA  THR A 155      89.439  23.067  52.406  1.00 21.99           C
    ATOM    916  C   THR A 155      87.964  22.981  51.982  1.00 17.78           C
    ATOM    917  O   THR A 155      87.659  22.984  50.793  1.00 16.68           O
    ATOM    918  CB  THR A 155      89.878  24.551  52.505  1.00 19.02           C
    ATOM    919  OG1 THR A 155      91.166  24.599  53.080  1.00 22.24           O
    ATOM    920  CG2 THR A 155      88.978  25.445  53.372  1.00 18.16           C
    ATOM    921  H   THR A 155      90.911  22.806  50.910  1.00  0.00           H
    ATOM    922  HA  THR A 155      89.549  22.611  53.390  1.00  0.00           H
    ATOM    923  HB  THR A 155      89.944  24.981  51.504  1.00  0.00           H
    ATOM    924 HG21 THR A 155      89.351  26.465  53.416  1.00  0.00           H
    ATOM    925 HG22 THR A 155      87.969  25.508  52.969  1.00  0.00           H
    ATOM    926 HG23 THR A 155      88.909  25.079  54.396  1.00  0.00           H
    ATOM    927  HG1 THR A 155      91.807  24.384  52.394  1.00  0.00           H
    ATOM    928  N   ARG A 156      87.104  22.837  52.984  1.00 16.52           N
    ATOM    929  CA  ARG A 156      85.676  22.622  52.871  1.00 16.61           C
    ATOM    930  C   ARG A 156      84.971  23.734  53.645  1.00 18.65           C
    ATOM    931  O   ARG A 156      85.520  24.261  54.612  1.00 18.72           O
    ATOM    932  CB  ARG A 156      85.390  21.220  53.447  1.00 17.92           C
    ATOM    933  CG  ARG A 156      85.492  20.122  52.373  1.00 20.09           C
    ATOM    934  CD  ARG A 156      86.006  18.766  52.879  1.00 19.55           C
    ATOM    935  NE  ARG A 156      85.107  18.091  53.823  1.00 24.41           N
    ATOM    936  CZ  ARG A 156      84.050  17.321  53.518  1.00 21.43           C
    ATOM    937  NH1 ARG A 156      83.644  17.153  52.255  1.00 22.99           N
    ATOM    938  NH2 ARG A 156      83.393  16.676  54.487  1.00 17.33           N1+
    ATOM    939  H   ARG A 156      87.459  22.849  53.934  1.00  0.00           H
    ATOM    940  HA  ARG A 156      85.367  22.676  51.827  1.00  0.00           H
    ATOM    941  HB2 ARG A 156      86.095  21.018  54.255  1.00  0.00           H
    ATOM    942  HB3 ARG A 156      84.413  21.171  53.919  1.00  0.00           H
    ATOM    943  HG2 ARG A 156      84.510  20.018  51.909  1.00  0.00           H
    ATOM    944  HG3 ARG A 156      86.163  20.425  51.568  1.00  0.00           H
    ATOM    945  HD2 ARG A 156      86.284  18.105  52.060  1.00  0.00           H
    ATOM    946  HD3 ARG A 156      86.925  18.951  53.436  1.00  0.00           H
    ATOM    947  HE  ARG A 156      85.370  18.188  54.799  1.00  0.00           H
    ATOM    948 HH11 ARG A 156      84.109  17.625  51.487  1.00  0.00           H
    ATOM    949 HH12 ARG A 156      82.905  16.483  52.051  1.00  0.00           H
    ATOM    950 HH21 ARG A 156      83.733  16.679  55.443  1.00  0.00           H
    ATOM    951 HH22 ARG A 156      82.617  16.066  54.252  1.00  0.00           H
    ATOM    952  N   VAL A 157      83.759  24.068  53.189  1.00 13.65           N
    ATOM    953  CA  VAL A 157      82.919  25.073  53.821  1.00 15.51           C
    ATOM    954  C   VAL A 157      81.622  24.383  54.215  1.00 18.38           C
    ATOM    955  O   VAL A 157      80.942  23.824  53.358  1.00 15.85           O
    ATOM    956  CB  VAL A 157      82.554  26.255  52.893  1.00 16.79           C
    ATOM    957  CG1 VAL A 157      81.901  27.398  53.697  1.00 15.66           C
    ATOM    958  CG2 VAL A 157      83.783  26.761  52.131  1.00 11.45           C
    ATOM    959  H   VAL A 157      83.370  23.591  52.385  1.00  0.00           H
    ATOM    960  HA  VAL A 157      83.402  25.479  54.710  1.00  0.00           H
    ATOM    961  HB  VAL A 157      81.837  25.934  52.136  1.00  0.00           H
    ATOM    962 HG11 VAL A 157      81.693  28.257  53.060  1.00  0.00           H
    ATOM    963 HG12 VAL A 157      80.950  27.091  54.133  1.00  0.00           H
    ATOM    964 HG13 VAL A 157      82.548  27.738  54.507  1.00  0.00           H
    ATOM    965 HG21 VAL A 157      83.557  27.633  51.528  1.00  0.00           H
    ATOM    966 HG22 VAL A 157      84.596  27.023  52.808  1.00  0.00           H
    ATOM    967 HG23 VAL A 157      84.142  25.998  51.442  1.00  0.00           H
    ATOM    968  N   ARG A 158      81.334  24.435  55.513  1.00 22.41           N
    ATOM    969  CA  ARG A 158      80.239  23.736  56.146  1.00 20.92           C
    ATOM    970  C   ARG A 158      79.230  24.740  56.692  1.00 22.83           C
    ATOM    971  O   ARG A 158      79.650  25.708  57.313  1.00 26.93           O
    ATOM    972  CB  ARG A 158      80.850  22.905  57.277  1.00 17.69           C
    ATOM    973  CG  ARG A 158      79.810  22.144  58.095  1.00 16.30           C
    ATOM    974  CD  ARG A 158      80.470  21.215  59.109  1.00 10.61           C
    ATOM    975  NE  ARG A 158      80.141  19.834  58.796  1.00 12.96           N
    ATOM    976  CZ  ARG A 158      80.833  18.968  58.055  1.00 21.55           C
    ATOM    977  NH1 ARG A 158      82.063  19.197  57.591  1.00 22.67           N
    ATOM    978  NH2 ARG A 158      80.229  17.817  57.799  1.00 29.26           N1+
    ATOM    979  H   ARG A 158      81.947  24.947  56.137  1.00  0.00           H
    ATOM    980  HA  ARG A 158      79.755  23.061  55.446  1.00  0.00           H
    ATOM    981  HB2 ARG A 158      81.569  22.200  56.864  1.00  0.00           H
    ATOM    982  HB3 ARG A 158      81.424  23.547  57.945  1.00  0.00           H
    ATOM    983  HG2 ARG A 158      79.122  22.811  58.613  1.00  0.00           H
    ATOM    984  HG3 ARG A 158      79.202  21.567  57.396  1.00  0.00           H
    ATOM    985  HD2 ARG A 158      81.540  21.347  59.178  1.00  0.00           H
    ATOM    986  HD3 ARG A 158      80.071  21.439  60.099  1.00  0.00           H
    ATOM    987  HE  ARG A 158      79.211  19.557  59.085  1.00  0.00           H
    ATOM    988 HH11 ARG A 158      82.571  20.030  57.866  1.00  0.00           H
    ATOM    989 HH12 ARG A 158      82.547  18.501  57.043  1.00  0.00           H
    ATOM    990 HH21 ARG A 158      79.313  17.632  58.178  1.00  0.00           H
    ATOM    991 HH22 ARG A 158      80.661  17.100  57.218  1.00  0.00           H
    ATOM    992  N   ALA A 159      77.935  24.453  56.521  1.00 22.98           N
    ATOM    993  CA  ALA A 159      76.852  25.223  57.120  1.00 14.03           C
    ATOM    994  C   ALA A 159      76.019  24.324  58.031  1.00 14.01           C
    ATOM    995  O   ALA A 159      75.728  23.187  57.662  1.00 16.65           O
    ATOM    996  CB  ALA A 159      75.986  25.836  56.018  1.00 13.29           C
    ATOM    997  H   ALA A 159      77.665  23.626  56.005  1.00  0.00           H
    ATOM    998  HA  ALA A 159      77.251  26.048  57.702  1.00  0.00           H
    ATOM    999  HB1 ALA A 159      75.193  26.456  56.437  1.00  0.00           H
    ATOM   1000  HB2 ALA A 159      76.586  26.470  55.369  1.00  0.00           H
    ATOM   1001  HB3 ALA A 159      75.519  25.073  55.399  1.00  0.00           H
    ATOM   1002  N   MET A 160      75.627  24.872  59.188  1.00 13.25           N
    ATOM   1003  CA  MET A 160      74.769  24.213  60.165  1.00 10.32           C
    ATOM   1004  C   MET A 160      73.836  25.249  60.790  1.00 12.97           C
    ATOM   1005  O   MET A 160      74.303  26.306  61.204  1.00 20.14           O
    ATOM   1006  CB  MET A 160      75.633  23.512  61.234  1.00 10.13           C
    ATOM   1007  CG  MET A 160      74.819  22.895  62.388  1.00 14.08           C
    ATOM   1008  SD  MET A 160      75.791  22.155  63.716  1.00 17.79           S
    ATOM   1009  CE  MET A 160      76.447  20.693  62.878  1.00 14.84           C
    ATOM   1010  H   MET A 160      75.923  25.811  59.423  1.00  0.00           H
    ATOM   1011  HA  MET A 160      74.153  23.461  59.666  1.00  0.00           H
    ATOM   1012  HB2 MET A 160      76.233  22.738  60.756  1.00  0.00           H
    ATOM   1013  HB3 MET A 160      76.346  24.226  61.650  1.00  0.00           H
    ATOM   1014  HG2 MET A 160      74.227  23.661  62.885  1.00  0.00           H
    ATOM   1015  HG3 MET A 160      74.106  22.163  62.018  1.00  0.00           H
    ATOM   1016  HE1 MET A 160      77.000  20.081  63.586  1.00  0.00           H
    ATOM   1017  HE2 MET A 160      75.636  20.093  62.467  1.00  0.00           H
    ATOM   1018  HE3 MET A 160      77.124  20.975  62.073  1.00  0.00           H
    ATOM   1019  N   ALA A 161      72.551  24.893  60.915  1.00 10.83           N
    ATOM   1020  CA  ALA A 161      71.566  25.676  61.647  1.00 10.75           C
    ATOM   1021  C   ALA A 161      71.504  25.208  63.104  1.00 14.32           C
    ATOM   1022  O   ALA A 161      71.438  24.004  63.335  1.00 17.69           O
    ATOM   1023  CB  ALA A 161      70.189  25.525  60.993  1.00  7.50           C
    ATOM   1024  H   ALA A 161      72.237  23.988  60.585  1.00  0.00           H
    ATOM   1025  HA  ALA A 161      71.835  26.721  61.581  1.00  0.00           H
    ATOM   1026  HB1 ALA A 161      69.434  26.085  61.544  1.00  0.00           H
    ATOM   1027  HB2 ALA A 161      70.184  25.907  59.975  1.00  0.00           H
    ATOM   1028  HB3 ALA A 161      69.868  24.483  60.954  1.00  0.00           H
    ATOM   1029  N   ILE A 162      71.475  26.149  64.051  1.00 14.46           N
    ATOM   1030  CA  ILE A 162      71.194  25.886  65.463  1.00 13.84           C
    ATOM   1031  C   ILE A 162      70.176  26.938  65.928  1.00 21.01           C
    ATOM   1032  O   ILE A 162      70.165  28.044  65.394  1.00 15.66           O
    ATOM   1033  CB  ILE A 162      72.474  25.972  66.350  1.00 12.75           C
    ATOM   1034  CG1 ILE A 162      73.086  27.390  66.486  1.00 21.65           C
    ATOM   1035  CG2 ILE A 162      73.538  24.953  65.898  1.00 10.85           C
    ATOM   1036  CD1 ILE A 162      74.221  27.484  67.510  1.00 22.34           C
    ATOM   1037  H   ILE A 162      71.543  27.125  63.788  1.00  0.00           H
    ATOM   1038  HA  ILE A 162      70.735  24.904  65.590  1.00  0.00           H
    ATOM   1039  HB  ILE A 162      72.167  25.663  67.352  1.00  0.00           H
    ATOM   1040 HG12 ILE A 162      73.450  27.724  65.514  1.00  0.00           H
    ATOM   1041 HG13 ILE A 162      72.345  28.124  66.797  1.00  0.00           H
    ATOM   1042 HG21 ILE A 162      74.321  24.854  66.646  1.00  0.00           H
    ATOM   1043 HG22 ILE A 162      73.105  23.962  65.757  1.00  0.00           H
    ATOM   1044 HG23 ILE A 162      74.008  25.248  64.960  1.00  0.00           H
    ATOM   1045 HD11 ILE A 162      74.404  28.525  67.776  1.00  0.00           H
    ATOM   1046 HD12 ILE A 162      73.984  26.944  68.427  1.00  0.00           H
    ATOM   1047 HD13 ILE A 162      75.152  27.081  67.111  1.00  0.00           H
    ATOM   1048  N   TYR A 163      69.346  26.606  66.923  1.00 24.50           N
    ATOM   1049  CA  TYR A 163      68.468  27.569  67.584  1.00 19.58           C
    ATOM   1050  C   TYR A 163      69.303  28.594  68.351  1.00 19.43           C
    ATOM   1051  O   TYR A 163      70.253  28.227  69.044  1.00 17.82           O
    ATOM   1052  CB  TYR A 163      67.464  26.865  68.513  1.00 17.06           C
    ATOM   1053  CG  TYR A 163      66.381  26.118  67.765  1.00 14.34           C
    ATOM   1054  CD1 TYR A 163      65.314  26.816  67.169  1.00  8.87           C
    ATOM   1055  CD2 TYR A 163      66.450  24.721  67.639  1.00 16.31           C
    ATOM   1056  CE1 TYR A 163      64.363  26.119  66.400  1.00  9.53           C
    ATOM   1057  CE2 TYR A 163      65.515  24.022  66.858  1.00 13.15           C
    ATOM   1058  CZ  TYR A 163      64.490  24.730  66.209  1.00 14.55           C
    ATOM   1059  OH  TYR A 163      63.626  24.068  65.391  1.00 20.05           O
    ATOM   1060  H   TYR A 163      69.412  25.690  67.354  1.00  0.00           H
    ATOM   1061  HA  TYR A 163      67.902  28.086  66.806  1.00  0.00           H
    ATOM   1062  HB2 TYR A 163      67.981  26.202  69.203  1.00  0.00           H
    ATOM   1063  HB3 TYR A 163      66.967  27.594  69.146  1.00  0.00           H
    ATOM   1064  HD1 TYR A 163      65.236  27.887  67.279  1.00  0.00           H
    ATOM   1065  HD2 TYR A 163      67.232  24.183  68.136  1.00  0.00           H
    ATOM   1066  HE1 TYR A 163      63.565  26.667  65.926  1.00  0.00           H
    ATOM   1067  HE2 TYR A 163      65.602  22.952  66.747  1.00  0.00           H
    ATOM   1068  HH  TYR A 163      63.768  23.115  65.362  1.00  0.00           H
    ATOM   1069  N   LYS A 164      68.935  29.866  68.173  1.00 21.07           N
    ATOM   1070  CA  LYS A 164      69.643  31.000  68.741  1.00 23.04           C
    ATOM   1071  C   LYS A 164      69.492  31.106  70.263  1.00 27.44           C
    ATOM   1072  O   LYS A 164      70.421  31.540  70.932  1.00 23.97           O
    ATOM   1073  CB  LYS A 164      69.165  32.294  68.064  1.00 19.81           C
    ATOM   1074  CG  LYS A 164      70.063  33.481  68.444  1.00 20.91           C
    ATOM   1075  CD  LYS A 164      69.746  34.788  67.718  1.00 24.22           C
    ATOM   1076  CE  LYS A 164      68.615  35.576  68.388  1.00 29.64           C
    ATOM   1077  NZ  LYS A 164      68.684  36.998  68.024  1.00 34.63           N1+
    ATOM   1078  H   LYS A 164      68.138  30.081  67.587  1.00  0.00           H
    ATOM   1079  HA  LYS A 164      70.704  30.866  68.520  1.00  0.00           H
    ATOM   1080  HB2 LYS A 164      69.192  32.160  66.983  1.00  0.00           H
    ATOM   1081  HB3 LYS A 164      68.127  32.500  68.321  1.00  0.00           H
    ATOM   1082  HG2 LYS A 164      70.019  33.673  69.515  1.00  0.00           H
    ATOM   1083  HG3 LYS A 164      71.101  33.211  68.256  1.00  0.00           H
    ATOM   1084  HD2 LYS A 164      70.651  35.396  67.678  1.00  0.00           H
    ATOM   1085  HD3 LYS A 164      69.477  34.583  66.682  1.00  0.00           H
    ATOM   1086  HE2 LYS A 164      67.642  35.169  68.122  1.00  0.00           H
    ATOM   1087  HE3 LYS A 164      68.698  35.525  69.474  1.00  0.00           H
    ATOM   1088  HZ1 LYS A 164      68.615  37.074  67.011  1.00  0.00           H
    ATOM   1089  HZ2 LYS A 164      67.916  37.500  68.445  1.00  0.00           H
    ATOM   1090  HZ3 LYS A 164      69.560  37.395  68.327  1.00  0.00           H
    ATOM   1091  N   GLN A 165      68.325  30.746  70.795  1.00 29.65           N
    ATOM   1092  CA  GLN A 165      68.039  30.858  72.216  1.00 26.03           C
    ATOM   1093  C   GLN A 165      68.656  29.684  72.977  1.00 26.58           C
    ATOM   1094  O   GLN A 165      68.601  28.555  72.487  1.00 29.91           O
    ATOM   1095  CB  GLN A 165      66.525  30.897  72.366  1.00 22.14           C
    ATOM   1096  CG  GLN A 165      65.906  32.173  71.774  1.00 20.93           C
    ATOM   1097  CD  GLN A 165      64.399  32.068  71.567  1.00 25.83           C
    ATOM   1098  OE1 GLN A 165      63.886  31.063  71.085  1.00 30.05           O
    ATOM   1099  NE2 GLN A 165      63.679  33.140  71.870  1.00 28.30           N
    ATOM   1100  H   GLN A 165      67.584  30.383  70.207  1.00  0.00           H
    ATOM   1101  HA  GLN A 165      68.450  31.792  72.606  1.00  0.00           H
    ATOM   1102  HB2 GLN A 165      66.118  29.999  71.909  1.00  0.00           H
    ATOM   1103  HB3 GLN A 165      66.274  30.857  73.413  1.00  0.00           H
    ATOM   1104  HG2 GLN A 165      66.144  33.025  72.411  1.00  0.00           H
    ATOM   1105  HG3 GLN A 165      66.336  32.396  70.798  1.00  0.00           H
    ATOM   1106 HE21 GLN A 165      64.101  33.962  72.278  1.00  0.00           H
    ATOM   1107 HE22 GLN A 165      62.678  33.138  71.721  1.00  0.00           H
    ATOM   1108  N   SER A 166      69.231  29.966  74.158  1.00 26.19           N
    ATOM   1109  CA  SER A 166      69.898  28.960  74.989  1.00 23.03           C
    ATOM   1110  C   SER A 166      68.980  27.802  75.406  1.00 17.17           C
    ATOM   1111  O   SER A 166      69.437  26.661  75.437  1.00 14.13           O
    ATOM   1112  CB  SER A 166      70.570  29.635  76.198  1.00 33.30           C
    ATOM   1113  OG  SER A 166      69.631  30.158  77.118  1.00 38.67           O
    ATOM   1114  H   SER A 166      69.228  30.912  74.513  1.00  0.00           H
    ATOM   1115  HA  SER A 166      70.697  28.536  74.379  1.00  0.00           H
    ATOM   1116  HB2 SER A 166      71.199  28.915  76.724  1.00  0.00           H
    ATOM   1117  HB3 SER A 166      71.229  30.439  75.868  1.00  0.00           H
    ATOM   1118  HG  SER A 166      70.105  30.641  77.800  1.00  0.00           H
    ATOM   1119  N   GLN A 167      67.701  28.111  75.664  1.00 17.30           N
    ATOM   1120  CA  GLN A 167      66.688  27.148  76.070  1.00 21.07           C
    ATOM   1121  C   GLN A 167      66.277  26.131  74.989  1.00 20.20           C
    ATOM   1122  O   GLN A 167      65.665  25.130  75.346  1.00 21.62           O
    ATOM   1123  CB  GLN A 167      65.488  27.876  76.698  1.00 19.39           C
    ATOM   1124  CG  GLN A 167      64.627  28.708  75.737  1.00 21.73           C
    ATOM   1125  CD  GLN A 167      65.028  30.168  75.577  1.00 23.31           C
    ATOM   1126  OE1 GLN A 167      66.163  30.556  75.848  1.00 22.48           O
    ATOM   1127  NE2 GLN A 167      64.090  30.983  75.089  1.00 23.11           N
    ATOM   1128  H   GLN A 167      67.401  29.078  75.622  1.00  0.00           H
    ATOM   1129  HA  GLN A 167      67.129  26.564  76.871  1.00  0.00           H
    ATOM   1130  HB2 GLN A 167      64.839  27.114  77.128  1.00  0.00           H
    ATOM   1131  HB3 GLN A 167      65.810  28.473  77.552  1.00  0.00           H
    ATOM   1132  HG2 GLN A 167      64.540  28.238  74.763  1.00  0.00           H
    ATOM   1133  HG3 GLN A 167      63.639  28.723  76.159  1.00  0.00           H
    ATOM   1134 HE21 GLN A 167      63.169  30.633  74.868  1.00  0.00           H
    ATOM   1135 HE22 GLN A 167      64.287  31.970  74.972  1.00  0.00           H
    ATOM   1136  N   HIS A 168      66.630  26.373  73.717  1.00 15.21           N
    ATOM   1137  CA  HIS A 168      66.335  25.478  72.595  1.00 12.91           C
    ATOM   1138  C   HIS A 168      67.600  24.955  71.908  1.00 11.63           C
    ATOM   1139  O   HIS A 168      67.479  24.244  70.913  1.00 14.78           O
    ATOM   1140  CB  HIS A 168      65.457  26.217  71.569  1.00 17.25           C
    ATOM   1141  CG  HIS A 168      64.151  26.747  72.092  1.00 15.22           C
    ATOM   1142  ND1 HIS A 168      63.089  25.931  72.488  1.00 17.84           N
    ATOM   1143  CD2 HIS A 168      63.773  28.058  72.244  1.00 16.91           C
    ATOM   1144  CE1 HIS A 168      62.134  26.769  72.868  1.00 17.90           C
    ATOM   1145  NE2 HIS A 168      62.485  28.046  72.738  1.00 19.50           N
    ATOM   1146  H   HIS A 168      67.142  27.216  73.493  1.00  0.00           H
    ATOM   1147  HA  HIS A 168      65.794  24.595  72.937  1.00  0.00           H
    ATOM   1148  HB2 HIS A 168      66.012  27.060  71.159  1.00  0.00           H
    ATOM   1149  HB3 HIS A 168      65.224  25.566  70.725  1.00  0.00           H
    ATOM   1150  HD2 HIS A 168      64.310  28.965  72.062  1.00  0.00           H
    ATOM   1151  HE2 HIS A 168      61.907  28.846  72.964  1.00  0.00           H
    ATOM   1152  HE1 HIS A 168      61.171  26.449  73.236  1.00  0.00           H
    ATOM   1153  N   MET A 169      68.790  25.322  72.404  1.00 15.07           N
    ATOM   1154  CA  MET A 169      70.058  25.053  71.733  1.00 17.32           C
    ATOM   1155  C   MET A 169      70.441  23.567  71.629  1.00 19.10           C
    ATOM   1156  O   MET A 169      71.189  23.213  70.720  1.00 21.44           O
    ATOM   1157  CB  MET A 169      71.160  25.890  72.386  1.00 24.71           C
    ATOM   1158  CG  MET A 169      72.282  26.269  71.409  1.00 30.00           C
    ATOM   1159  SD  MET A 169      73.263  27.662  72.009  1.00 40.41           S
    ATOM   1160  CE  MET A 169      72.165  29.029  71.564  1.00 34.43           C
    ATOM   1161  H   MET A 169      68.829  25.900  73.233  1.00  0.00           H
    ATOM   1162  HA  MET A 169      69.926  25.435  70.723  1.00  0.00           H
    ATOM   1163  HB2 MET A 169      70.725  26.805  72.781  1.00  0.00           H
    ATOM   1164  HB3 MET A 169      71.567  25.377  73.257  1.00  0.00           H
    ATOM   1165  HG2 MET A 169      72.942  25.418  71.251  1.00  0.00           H
    ATOM   1166  HG3 MET A 169      71.884  26.536  70.431  1.00  0.00           H
    ATOM   1167  HE1 MET A 169      72.431  29.910  72.146  1.00  0.00           H
    ATOM   1168  HE2 MET A 169      72.259  29.255  70.502  1.00  0.00           H
    ATOM   1169  HE3 MET A 169      71.127  28.788  71.772  1.00  0.00           H
    ATOM   1170  N   THR A 170      69.901  22.718  72.515  1.00 18.86           N
    ATOM   1171  CA  THR A 170      70.070  21.268  72.430  1.00 20.77           C
    ATOM   1172  C   THR A 170      69.073  20.613  71.461  1.00 15.74           C
    ATOM   1173  O   THR A 170      69.234  19.434  71.159  1.00 16.00           O
    ATOM   1174  CB  THR A 170      69.944  20.607  73.819  1.00 23.39           C
    ATOM   1175  OG1 THR A 170      68.707  20.899  74.445  1.00 29.02           O
    ATOM   1176  CG2 THR A 170      71.091  21.024  74.739  1.00 13.84           C
    ATOM   1177  H   THR A 170      69.293  23.057  73.248  1.00  0.00           H
    ATOM   1178  HA  THR A 170      71.068  21.042  72.051  1.00  0.00           H
    ATOM   1179  HB  THR A 170      70.001  19.523  73.710  1.00  0.00           H
    ATOM   1180 HG21 THR A 170      71.024  20.464  75.663  1.00  0.00           H
    ATOM   1181 HG22 THR A 170      72.061  20.800  74.292  1.00  0.00           H
    ATOM   1182 HG23 THR A 170      71.064  22.087  74.980  1.00  0.00           H
    ATOM   1183  HG1 THR A 170      68.760  20.637  75.368  1.00  0.00           H
    ATOM   1184  N   GLU A 171      68.097  21.364  70.937  1.00 15.17           N
    ATOM   1185  CA  GLU A 171      67.210  20.858  69.907  1.00 20.99           C
    ATOM   1186  C   GLU A 171      67.879  20.984  68.535  1.00 21.73           C
    ATOM   1187  O   GLU A 171      68.317  22.064  68.155  1.00 22.83           O
    ATOM   1188  CB  GLU A 171      65.870  21.592  69.974  1.00 22.48           C
    ATOM   1189  CG  GLU A 171      64.823  20.910  69.090  1.00 21.04           C
    ATOM   1190  CD  GLU A 171      63.509  21.658  69.077  1.00 25.50           C
    ATOM   1191  OE1 GLU A 171      63.086  22.098  70.165  1.00 24.00           O
    ATOM   1192  OE2 GLU A 171      62.943  21.794  67.973  1.00 25.87           O1-
    ATOM   1193  H   GLU A 171      68.000  22.341  71.184  1.00  0.00           H
    ATOM   1194  HA  GLU A 171      67.004  19.805  70.106  1.00  0.00           H
    ATOM   1195  HB2 GLU A 171      65.525  21.615  71.009  1.00  0.00           H
    ATOM   1196  HB3 GLU A 171      65.983  22.629  69.662  1.00  0.00           H
    ATOM   1197  HG2 GLU A 171      65.179  20.832  68.063  1.00  0.00           H
    ATOM   1198  HG3 GLU A 171      64.641  19.902  69.454  1.00  0.00           H
    ATOM   1199  N   VAL A 172      67.926  19.869  67.803  1.00 17.14           N
    ATOM   1200  CA  VAL A 172      68.428  19.827  66.439  1.00 16.82           C
    ATOM   1201  C   VAL A 172      67.417  20.514  65.518  1.00 19.96           C
    ATOM   1202  O   VAL A 172      66.253  20.109  65.487  1.00 21.13           O
    ATOM   1203  CB  VAL A 172      68.636  18.365  65.970  1.00 19.48           C
    ATOM   1204  CG1 VAL A 172      68.975  18.217  64.474  1.00 20.70           C
    ATOM   1205  CG2 VAL A 172      69.734  17.692  66.808  1.00 18.70           C
    ATOM   1206  H   VAL A 172      67.527  19.010  68.159  1.00  0.00           H
    ATOM   1207  HA  VAL A 172      69.385  20.350  66.400  1.00  0.00           H
    ATOM   1208  HB  VAL A 172      67.714  17.807  66.144  1.00  0.00           H
    ATOM   1209 HG11 VAL A 172      69.194  17.178  64.225  1.00  0.00           H
    ATOM   1210 HG12 VAL A 172      68.156  18.522  63.823  1.00  0.00           H
    ATOM   1211 HG13 VAL A 172      69.855  18.804  64.220  1.00  0.00           H
    ATOM   1212 HG21 VAL A 172      69.885  16.656  66.507  1.00  0.00           H
    ATOM   1213 HG22 VAL A 172      70.689  18.207  66.701  1.00  0.00           H
    ATOM   1214 HG23 VAL A 172      69.479  17.688  67.867  1.00  0.00           H
    ATOM   1215  N   VAL A 173      67.882  21.537  64.787  1.00 23.22           N
    ATOM   1216  CA  VAL A 173      67.068  22.196  63.775  1.00 21.13           C
    ATOM   1217  C   VAL A 173      66.875  21.216  62.613  1.00 17.08           C
    ATOM   1218  O   VAL A 173      67.845  20.687  62.077  1.00 10.45           O
    ATOM   1219  CB  VAL A 173      67.704  23.504  63.241  1.00 15.70           C
    ATOM   1220  CG1 VAL A 173      66.887  24.156  62.104  1.00 16.25           C
    ATOM   1221  CG2 VAL A 173      67.899  24.532  64.366  1.00 10.90           C
    ATOM   1222  H   VAL A 173      68.847  21.827  64.864  1.00  0.00           H
    ATOM   1223  HA  VAL A 173      66.098  22.445  64.214  1.00  0.00           H
    ATOM   1224  HB  VAL A 173      68.695  23.267  62.850  1.00  0.00           H
    ATOM   1225 HG11 VAL A 173      67.316  25.111  61.806  1.00  0.00           H
    ATOM   1226 HG12 VAL A 173      66.860  23.542  61.204  1.00  0.00           H
    ATOM   1227 HG13 VAL A 173      65.857  24.342  62.412  1.00  0.00           H
    ATOM   1228 HG21 VAL A 173      68.492  25.372  64.009  1.00  0.00           H
    ATOM   1229 HG22 VAL A 173      66.946  24.933  64.708  1.00  0.00           H
    ATOM   1230 HG23 VAL A 173      68.421  24.113  65.226  1.00  0.00           H
    ATOM   1231  N   ARG A 174      65.613  20.969  62.282  1.00 17.93           N
    ATOM   1232  CA  ARG A 174      65.222  20.101  61.190  1.00 18.26           C
    ATOM   1233  C   ARG A 174      63.882  20.605  60.685  1.00 20.64           C
    ATOM   1234  O   ARG A 174      63.206  21.391  61.357  1.00 19.76           O
    ATOM   1235  CB  ARG A 174      65.214  18.620  61.641  1.00 22.81           C
    ATOM   1236  CG  ARG A 174      64.206  18.276  62.753  1.00 16.28           C
    ATOM   1237  CD  ARG A 174      64.298  16.826  63.244  1.00 21.63           C
    ATOM   1238  NE  ARG A 174      63.722  15.876  62.286  1.00 23.48           N
    ATOM   1239  CZ  ARG A 174      63.758  14.538  62.395  1.00 27.25           C
    ATOM   1240  NH1 ARG A 174      64.398  13.938  63.406  1.00 26.14           N
    ATOM   1241  NH2 ARG A 174      63.142  13.784  61.477  1.00 25.51           N1+
    ATOM   1242  H   ARG A 174      64.857  21.433  62.767  1.00  0.00           H
    ATOM   1243  HA  ARG A 174      65.926  20.212  60.364  1.00  0.00           H
    ATOM   1244  HB2 ARG A 174      65.036  17.978  60.779  1.00  0.00           H
    ATOM   1245  HB3 ARG A 174      66.216  18.366  61.988  1.00  0.00           H
    ATOM   1246  HG2 ARG A 174      64.276  18.954  63.604  1.00  0.00           H
    ATOM   1247  HG3 ARG A 174      63.216  18.410  62.323  1.00  0.00           H
    ATOM   1248  HD2 ARG A 174      65.353  16.553  63.297  1.00  0.00           H
    ATOM   1249  HD3 ARG A 174      63.897  16.704  64.249  1.00  0.00           H
    ATOM   1250  HE  ARG A 174      63.289  16.277  61.460  1.00  0.00           H
    ATOM   1251 HH11 ARG A 174      64.835  14.486  64.134  1.00  0.00           H
    ATOM   1252 HH12 ARG A 174      64.441  12.931  63.459  1.00  0.00           H
    ATOM   1253 HH21 ARG A 174      62.622  14.205  60.716  1.00  0.00           H
    ATOM   1254 HH22 ARG A 174      63.191  12.777  61.531  1.00  0.00           H
    ATOM   1255  N   ARG A 175      63.517  20.131  59.492  1.00 22.46           N
    ATOM   1256  CA  ARG A 175      62.238  20.465  58.891  1.00 21.07           C
    ATOM   1257  C   ARG A 175      61.086  19.902  59.722  1.00 21.58           C
    ATOM   1258  O   ARG A 175      61.228  18.865  60.367  1.00 26.74           O
    ATOM   1259  CB  ARG A 175      62.176  19.925  57.462  1.00 17.97           C
    ATOM   1260  CG  ARG A 175      63.320  20.436  56.577  1.00 17.91           C
    ATOM   1261  CD  ARG A 175      63.309  19.762  55.215  1.00 24.08           C
    ATOM   1262  NE  ARG A 175      62.407  20.391  54.271  1.00 25.29           N
    ATOM   1263  CZ  ARG A 175      62.103  19.875  53.079  1.00 24.50           C
    ATOM   1264  NH1 ARG A 175      62.581  18.700  52.649  1.00 26.47           N
    ATOM   1265  NH2 ARG A 175      61.291  20.590  52.316  1.00 25.07           N1+
    ATOM   1266  H   ARG A 175      64.113  19.488  58.988  1.00  0.00           H
    ATOM   1267  HA  ARG A 175      62.167  21.552  58.863  1.00  0.00           H
    ATOM   1268  HB2 ARG A 175      62.217  18.834  57.508  1.00  0.00           H
    ATOM   1269  HB3 ARG A 175      61.219  20.176  57.003  1.00  0.00           H
    ATOM   1270  HG2 ARG A 175      63.219  21.509  56.451  1.00  0.00           H
    ATOM   1271  HG3 ARG A 175      64.296  20.295  57.031  1.00  0.00           H
    ATOM   1272  HD2 ARG A 175      64.287  19.535  54.808  1.00  0.00           H
    ATOM   1273  HD3 ARG A 175      62.818  18.806  55.367  1.00  0.00           H
    ATOM   1274  HE  ARG A 175      62.050  21.306  54.514  1.00  0.00           H
    ATOM   1275 HH11 ARG A 175      63.108  18.101  53.273  1.00  0.00           H
    ATOM   1276 HH12 ARG A 175      62.302  18.359  51.736  1.00  0.00           H
    ATOM   1277 HH21 ARG A 175      60.955  21.485  52.641  1.00  0.00           H
    ATOM   1278 HH22 ARG A 175      61.022  20.290  51.378  1.00  0.00           H
    ATOM   1279  N   CYS A 176      59.942  20.582  59.669  1.00 16.73           N
    ATOM   1280  CA  CYS A 176      58.691  20.083  60.222  1.00 13.13           C
    ATOM   1281  C   CYS A 176      58.281  18.759  59.526  1.00 17.61           C
    ATOM   1282  O   CYS A 176      58.800  18.472  58.443  1.00 19.56           O
    ATOM   1283  CB  CYS A 176      57.601  21.169  60.128  1.00 19.18           C
    ATOM   1284  SG  CYS A 176      56.949  21.516  58.443  1.00 30.55           S
    ATOM   1285  H   CYS A 176      59.890  21.437  59.130  1.00  0.00           H
    ATOM   1286  HA  CYS A 176      58.888  19.887  61.276  1.00  0.00           H
    ATOM   1287  HB2 CYS A 176      56.763  20.886  60.765  1.00  0.00           H
    ATOM   1288  HB3 CYS A 176      57.981  22.090  60.568  1.00  0.00           H
    ATOM   1289  N   PRO A 177      57.346  17.982  60.115  1.00 21.61           N
    ATOM   1290  CA  PRO A 177      56.828  16.768  59.463  1.00 23.04           C
    ATOM   1291  C   PRO A 177      56.253  16.973  58.055  1.00 29.08           C
    ATOM   1292  O   PRO A 177      56.547  16.164  57.176  1.00 33.20           O
    ATOM   1293  CB  PRO A 177      55.773  16.230  60.428  1.00 22.39           C
    ATOM   1294  CG  PRO A 177      56.203  16.753  61.784  1.00 22.72           C
    ATOM   1295  CD  PRO A 177      56.837  18.105  61.479  1.00 24.14           C
    ATOM   1296  HA  PRO A 177      57.641  16.048  59.441  1.00  0.00           H
    ATOM   1297  HB2 PRO A 177      54.787  16.635  60.193  1.00  0.00           H
    ATOM   1298  HB3 PRO A 177      55.698  15.142  60.405  1.00  0.00           H
    ATOM   1299  HG2 PRO A 177      55.393  16.804  62.511  1.00  0.00           H
    ATOM   1300  HG3 PRO A 177      56.969  16.088  62.188  1.00  0.00           H
    ATOM   1301  HD2 PRO A 177      56.086  18.891  61.521  1.00  0.00           H
    ATOM   1302  HD3 PRO A 177      57.615  18.329  62.209  1.00  0.00           H
    ATOM   1303  N   HIS A 178      55.492  18.064  57.849  1.00 28.73           N
    ATOM   1304  CA  HIS A 178      54.957  18.425  56.537  1.00 23.98           C
    ATOM   1305  C   HIS A 178      56.052  18.514  55.466  1.00 29.84           C
    ATOM   1306  O   HIS A 178      55.920  17.873  54.424  1.00 31.79           O
    ATOM   1307  CB  HIS A 178      54.165  19.743  56.597  1.00 27.38           C
    ATOM   1308  CG  HIS A 178      53.552  20.107  55.270  1.00 38.00           C
    ATOM   1309  ND1 HIS A 178      54.137  20.993  54.381  1.00 40.94           N
    ATOM   1310  CD2 HIS A 178      52.408  19.656  54.652  1.00 40.35           C
    ATOM   1311  CE1 HIS A 178      53.352  21.039  53.303  1.00 39.01           C
    ATOM   1312  NE2 HIS A 178      52.281  20.260  53.402  1.00 39.81           N
    ATOM   1313  H   HIS A 178      55.282  18.689  58.617  1.00  0.00           H
    ATOM   1314  HA  HIS A 178      54.269  17.630  56.245  1.00  0.00           H
    ATOM   1315  HB2 HIS A 178      53.365  19.672  57.328  1.00  0.00           H
    ATOM   1316  HB3 HIS A 178      54.799  20.566  56.921  1.00  0.00           H
    ATOM   1317  HD1 HIS A 178      54.999  21.503  54.512  1.00  0.00           H
    ATOM   1318  HD2 HIS A 178      51.675  18.946  55.006  1.00  0.00           H
    ATOM   1319  HE1 HIS A 178      53.569  21.647  52.437  1.00  0.00           H
    ATOM   1320  N   HIS A 179      57.106  19.294  55.759  1.00 31.52           N
    ATOM   1321  CA  HIS A 179      58.176  19.555  54.807  1.00 30.06           C
    ATOM   1322  C   HIS A 179      59.146  18.395  54.567  1.00 28.60           C
    ATOM   1323  O   HIS A 179      59.505  18.207  53.409  1.00 27.66           O
    ATOM   1324  CB  HIS A 179      58.872  20.894  55.088  1.00 28.00           C
    ATOM   1325  CG  HIS A 179      58.063  22.079  54.621  1.00 23.71           C
    ATOM   1326  ND1 HIS A 179      57.701  23.118  55.469  1.00 25.71           N
    ATOM   1327  CD2 HIS A 179      57.554  22.352  53.368  1.00 24.21           C
    ATOM   1328  CE1 HIS A 179      56.991  23.960  54.733  1.00 26.82           C
    ATOM   1329  NE2 HIS A 179      56.863  23.546  53.475  1.00 25.53           N
    ATOM   1330  H   HIS A 179      57.156  19.762  56.655  1.00  0.00           H
    ATOM   1331  HA  HIS A 179      57.696  19.679  53.837  1.00  0.00           H
    ATOM   1332  HB2 HIS A 179      59.091  20.990  56.152  1.00  0.00           H
    ATOM   1333  HB3 HIS A 179      59.830  20.947  54.577  1.00  0.00           H
    ATOM   1334  HD2 HIS A 179      57.627  21.813  52.436  1.00  0.00           H
    ATOM   1335  HE2 HIS A 179      56.386  24.030  52.727  1.00  0.00           H
    ATOM   1336  HE1 HIS A 179      56.569  24.885  55.098  1.00  0.00           H
    ATOM   1337  N   GLU A 180      59.515  17.588  55.580  1.00 23.62           N
    ATOM   1338  CA  GLU A 180      60.316  16.375  55.333  1.00 26.39           C
    ATOM   1339  C   GLU A 180      59.613  15.371  54.396  1.00 34.27           C
    ATOM   1340  O   GLU A 180      60.277  14.724  53.587  1.00 38.01           O
    ATOM   1341  CB  GLU A 180      60.759  15.701  56.650  1.00 23.38           C
    ATOM   1342  CG  GLU A 180      59.572  15.182  57.480  1.00 31.31           C
    ATOM   1343  CD  GLU A 180      59.914  14.414  58.750  1.00 42.80           C
    ATOM   1344  OE1 GLU A 180      61.072  14.515  59.216  1.00 43.65           O
    ATOM   1345  OE2 GLU A 180      58.981  13.762  59.268  1.00 45.78           O1-
    ATOM   1346  H   GLU A 180      59.196  17.760  56.525  1.00  0.00           H
    ATOM   1347  HA  GLU A 180      61.194  16.716  54.803  1.00  0.00           H
    ATOM   1348  HB2 GLU A 180      61.443  14.881  56.425  1.00  0.00           H
    ATOM   1349  HB3 GLU A 180      61.334  16.417  57.237  1.00  0.00           H
    ATOM   1350  HG2 GLU A 180      58.969  16.035  57.758  1.00  0.00           H
    ATOM   1351  HG3 GLU A 180      58.944  14.515  56.894  1.00  0.00           H
    ATOM   1352  N   ARG A 181      58.276  15.317  54.501  1.00 35.42           N
    ATOM   1353  CA  ARG A 181      57.396  14.511  53.665  1.00 36.07           C
    ATOM   1354  C   ARG A 181      57.116  15.126  52.288  1.00 37.05           C
    ATOM   1355  O   ARG A 181      56.511  14.434  51.469  1.00 37.63           O
    ATOM   1356  CB  ARG A 181      56.071  14.271  54.411  1.00 38.96           C
    ATOM   1357  CG  ARG A 181      56.201  13.247  55.544  1.00 42.62           C
    ATOM   1358  CD  ARG A 181      54.873  12.914  56.229  1.00 48.93           C
    ATOM   1359  NE  ARG A 181      54.236  14.100  56.826  1.00 54.92           N
    ATOM   1360  CZ  ARG A 181      53.559  14.132  57.987  1.00 54.22           C
    ATOM   1361  NH1 ARG A 181      53.440  13.041  58.757  1.00 54.44           N
    ATOM   1362  NH2 ARG A 181      52.991  15.275  58.389  1.00 50.02           N1+
    ATOM   1363  H   ARG A 181      57.816  15.890  55.197  1.00  0.00           H
    ATOM   1364  HA  ARG A 181      57.877  13.547  53.490  1.00  0.00           H
    ATOM   1365  HB2 ARG A 181      55.670  15.216  54.777  1.00  0.00           H
    ATOM   1366  HB3 ARG A 181      55.328  13.874  53.720  1.00  0.00           H
    ATOM   1367  HG2 ARG A 181      56.576  12.325  55.098  1.00  0.00           H
    ATOM   1368  HG3 ARG A 181      56.943  13.539  56.286  1.00  0.00           H
    ATOM   1369  HD2 ARG A 181      54.190  12.350  55.593  1.00  0.00           H
    ATOM   1370  HD3 ARG A 181      55.138  12.250  57.051  1.00  0.00           H
    ATOM   1371  HE  ARG A 181      54.285  14.943  56.271  1.00  0.00           H
    ATOM   1372 HH11 ARG A 181      53.872  12.175  58.469  1.00  0.00           H
    ATOM   1373 HH12 ARG A 181      52.933  13.075  59.629  1.00  0.00           H
    ATOM   1374 HH21 ARG A 181      53.068  16.110  57.825  1.00  0.00           H
    ATOM   1375 HH22 ARG A 181      52.478  15.316  59.258  1.00  0.00           H
    ATOM   1376  N   CYS A 182      57.537  16.379  52.032  1.00 37.39           N
    ATOM   1377  CA  CYS A 182      57.382  17.008  50.721  1.00 35.57           C
    ATOM   1378  C   CYS A 182      58.257  16.313  49.681  1.00 34.27           C
    ATOM   1379  O   CYS A 182      59.407  15.964  49.957  1.00 37.65           O
    ATOM   1380  CB  CYS A 182      57.716  18.514  50.695  1.00 33.56           C
    ATOM   1381  SG  CYS A 182      56.330  19.479  51.341  1.00 45.49           S
    ATOM   1382  H   CYS A 182      58.050  16.899  52.730  1.00  0.00           H
    ATOM   1383  HA  CYS A 182      56.340  16.886  50.417  1.00  0.00           H
    ATOM   1384  HB2 CYS A 182      58.635  18.752  51.222  1.00  0.00           H
    ATOM   1385  HB3 CYS A 182      57.865  18.858  49.670  1.00  0.00           H
    ATOM   1386  HG  CYS A 182      56.359  18.961  52.573  1.00  0.00           H
    ATOM   1387  N   SER A 183      57.694  16.197  48.475  1.00 32.86           N
    ATOM   1388  CA  SER A 183      58.431  15.811  47.287  1.00 36.20           C
    ATOM   1389  C   SER A 183      59.082  17.091  46.757  1.00 40.91           C
    ATOM   1390  O   SER A 183      58.514  17.758  45.895  1.00 43.67           O
    ATOM   1391  CB  SER A 183      57.459  15.218  46.251  1.00 39.33           C
    ATOM   1392  OG  SER A 183      56.738  14.135  46.792  1.00 43.24           O
    ATOM   1393  H   SER A 183      56.736  16.480  48.330  1.00  0.00           H
    ATOM   1394  HA  SER A 183      59.191  15.067  47.534  1.00  0.00           H
    ATOM   1395  HB2 SER A 183      56.727  15.955  45.916  1.00  0.00           H
    ATOM   1396  HB3 SER A 183      57.999  14.878  45.366  1.00  0.00           H
    ATOM   1397  HG  SER A 183      56.147  13.790  46.119  1.00  0.00           H
    ATOM   1398  N   ASP A 184      60.236  17.447  47.320  1.00 35.32           N
    ATOM   1399  CA  ASP A 184      60.985  18.622  46.884  1.00 24.44           C
    ATOM   1400  C   ASP A 184      62.470  18.302  46.813  1.00 20.08           C
    ATOM   1401  O   ASP A 184      63.278  19.225  46.752  1.00 27.73           O
    ATOM   1402  CB  ASP A 184      60.702  19.881  47.720  1.00 21.27           C
    ATOM   1403  CG  ASP A 184      60.859  19.720  49.227  1.00 18.13           C
    ATOM   1404  OD1 ASP A 184      61.520  18.767  49.705  1.00 17.67           O
    ATOM   1405  OD2 ASP A 184      60.345  20.643  49.884  1.00 23.49           O1-
    ATOM   1406  H   ASP A 184      60.624  16.911  48.083  1.00  0.00           H
    ATOM   1407  HA  ASP A 184      60.725  18.862  45.851  1.00  0.00           H
    ATOM   1408  HB2 ASP A 184      61.320  20.717  47.391  1.00  0.00           H
    ATOM   1409  HB3 ASP A 184      59.666  20.165  47.529  1.00  0.00           H
    ATOM   1410  N   SER A 185      62.800  17.004  46.812  1.00 19.17           N
    ATOM   1411  CA  SER A 185      64.158  16.519  46.736  1.00 26.41           C
    ATOM   1412  C   SER A 185      64.817  16.984  45.435  1.00 27.96           C
    ATOM   1413  O   SER A 185      64.271  16.779  44.349  1.00 30.17           O
    ATOM   1414  CB  SER A 185      64.156  14.989  46.870  1.00 31.42           C
    ATOM   1415  OG  SER A 185      65.481  14.539  47.028  1.00 35.90           O
    ATOM   1416  H   SER A 185      62.072  16.303  46.823  1.00  0.00           H
    ATOM   1417  HA  SER A 185      64.696  16.937  47.590  1.00  0.00           H
    ATOM   1418  HB2 SER A 185      63.578  14.675  47.740  1.00  0.00           H
    ATOM   1419  HB3 SER A 185      63.716  14.510  45.995  1.00  0.00           H
    ATOM   1420  HG  SER A 185      65.479  13.604  47.259  1.00  0.00           H
    ATOM   1421  N   ASP A 186      65.998  17.589  45.593  1.00 23.98           N
    ATOM   1422  CA  ASP A 186      66.911  17.891  44.494  1.00 20.72           C
    ATOM   1423  C   ASP A 186      67.720  16.636  44.094  1.00 20.65           C
    ATOM   1424  O   ASP A 186      68.590  16.725  43.229  1.00 24.29           O
    ATOM   1425  CB  ASP A 186      67.847  19.072  44.835  1.00 19.33           C
    ATOM   1426  CG  ASP A 186      68.705  18.898  46.094  1.00 21.96           C
    ATOM   1427  OD1 ASP A 186      68.644  17.827  46.738  1.00 21.72           O
    ATOM   1428  OD2 ASP A 186      69.437  19.863  46.386  1.00 18.62           O1-
    ATOM   1429  H   ASP A 186      66.380  17.716  46.520  1.00  0.00           H
    ATOM   1430  HA  ASP A 186      66.328  18.174  43.619  1.00  0.00           H
    ATOM   1431  HB2 ASP A 186      68.509  19.267  43.991  1.00  0.00           H
    ATOM   1432  HB3 ASP A 186      67.227  19.958  44.973  1.00  0.00           H
    ATOM   1433  N   GLY A 187      67.445  15.502  44.758  1.00 24.29           N
    ATOM   1434  CA  GLY A 187      68.104  14.224  44.553  1.00 26.44           C
    ATOM   1435  C   GLY A 187      69.402  14.099  45.359  1.00 24.32           C
    ATOM   1436  O   GLY A 187      70.011  13.033  45.305  1.00 26.92           O
    ATOM   1437  H   GLY A 187      66.733  15.523  45.476  1.00  0.00           H
    ATOM   1438  HA2 GLY A 187      67.421  13.433  44.865  1.00  0.00           H
    ATOM   1439  HA3 GLY A 187      68.315  14.064  43.495  1.00  0.00           H
    ATOM   1440  N   LEU A 188      69.838  15.139  46.091  1.00 26.50           N
    ATOM   1441  CA  LEU A 188      71.110  15.131  46.813  1.00 23.20           C
    ATOM   1442  C   LEU A 188      70.896  15.308  48.316  1.00 20.05           C
    ATOM   1443  O   LEU A 188      71.505  14.573  49.093  1.00 21.83           O
    ATOM   1444  CB  LEU A 188      72.034  16.230  46.252  1.00 21.20           C
    ATOM   1445  CG  LEU A 188      72.358  16.083  44.748  1.00 21.16           C
    ATOM   1446  CD1 LEU A 188      73.062  17.345  44.216  1.00 22.74           C
    ATOM   1447  CD2 LEU A 188      73.137  14.791  44.423  1.00 18.20           C
    ATOM   1448  H   LEU A 188      69.304  15.999  46.125  1.00  0.00           H
    ATOM   1449  HA  LEU A 188      71.622  14.175  46.700  1.00  0.00           H
    ATOM   1450  HB2 LEU A 188      71.550  17.194  46.413  1.00  0.00           H
    ATOM   1451  HB3 LEU A 188      72.966  16.268  46.818  1.00  0.00           H
    ATOM   1452  HG  LEU A 188      71.412  16.033  44.208  1.00  0.00           H
    ATOM   1453 HD11 LEU A 188      72.594  17.692  43.295  1.00  0.00           H
    ATOM   1454 HD12 LEU A 188      73.020  18.171  44.926  1.00  0.00           H
    ATOM   1455 HD13 LEU A 188      74.118  17.176  44.001  1.00  0.00           H
    ATOM   1456 HD21 LEU A 188      74.030  14.976  43.827  1.00  0.00           H
    ATOM   1457 HD22 LEU A 188      73.463  14.270  45.323  1.00  0.00           H
    ATOM   1458 HD23 LEU A 188      72.514  14.100  43.855  1.00  0.00           H
    ATOM   1459  N   ALA A 189      70.044  16.266  48.707  1.00 21.85           N
    ATOM   1460  CA  ALA A 189      69.793  16.580  50.106  1.00 22.79           C
    ATOM   1461  C   ALA A 189      68.889  15.530  50.761  1.00 21.88           C
    ATOM   1462  O   ALA A 189      67.873  15.156  50.167  1.00 30.35           O
    ATOM   1463  CB  ALA A 189      69.129  17.950  50.183  1.00 21.06           C
    ATOM   1464  H   ALA A 189      69.559  16.834  48.018  1.00  0.00           H
    ATOM   1465  HA  ALA A 189      70.749  16.662  50.608  1.00  0.00           H
    ATOM   1466  HB1 ALA A 189      69.016  18.274  51.218  1.00  0.00           H
    ATOM   1467  HB2 ALA A 189      69.706  18.704  49.652  1.00  0.00           H
    ATOM   1468  HB3 ALA A 189      68.141  17.923  49.723  1.00  0.00           H
    ATOM   1469  N   PRO A 190      69.247  15.087  51.986  1.00 15.40           N
    ATOM   1470  CA  PRO A 190      68.317  14.304  52.791  1.00 15.32           C
    ATOM   1471  C   PRO A 190      67.060  15.110  53.150  1.00 19.15           C
    ATOM   1472  O   PRO A 190      67.159  16.311  53.417  1.00 20.58           O
    ATOM   1473  CB  PRO A 190      69.106  13.933  54.040  1.00 16.31           C
    ATOM   1474  CG  PRO A 190      70.566  14.110  53.681  1.00 20.95           C
    ATOM   1475  CD  PRO A 190      70.541  15.220  52.645  1.00 16.89           C
    ATOM   1476  HA  PRO A 190      68.117  13.406  52.205  1.00  0.00           H
    ATOM   1477  HB2 PRO A 190      68.853  14.577  54.883  1.00  0.00           H
    ATOM   1478  HB3 PRO A 190      68.887  12.903  54.312  1.00  0.00           H
    ATOM   1479  HG2 PRO A 190      71.203  14.326  54.539  1.00  0.00           H
    ATOM   1480  HG3 PRO A 190      70.934  13.194  53.216  1.00  0.00           H
    ATOM   1481  HD2 PRO A 190      70.602  16.201  53.118  1.00  0.00           H
    ATOM   1482  HD3 PRO A 190      71.377  15.094  51.958  1.00  0.00           H
    ATOM   1483  N   PRO A 191      65.904  14.422  53.136  1.00 19.48           N
    ATOM   1484  CA  PRO A 191      64.600  15.072  53.269  1.00 21.80           C
    ATOM   1485  C   PRO A 191      64.365  15.879  54.554  1.00 25.65           C
    ATOM   1486  O   PRO A 191      63.488  16.722  54.519  1.00 26.77           O
    ATOM   1487  CB  PRO A 191      63.591  13.924  53.106  1.00 15.61           C
    ATOM   1488  CG  PRO A 191      64.365  12.652  53.398  1.00 19.97           C
    ATOM   1489  CD  PRO A 191      65.766  12.982  52.918  1.00 17.87           C
    ATOM   1490  HA  PRO A 191      64.481  15.767  52.435  1.00  0.00           H
    ATOM   1491  HB2 PRO A 191      62.715  14.012  53.748  1.00  0.00           H
    ATOM   1492  HB3 PRO A 191      63.238  13.896  52.075  1.00  0.00           H
    ATOM   1493  HG2 PRO A 191      64.368  12.479  54.474  1.00  0.00           H
    ATOM   1494  HG3 PRO A 191      63.941  11.768  52.920  1.00  0.00           H
    ATOM   1495  HD2 PRO A 191      66.505  12.383  53.451  1.00  0.00           H
    ATOM   1496  HD3 PRO A 191      65.861  12.776  51.850  1.00  0.00           H
    ATOM   1497  N   GLN A 192      65.131  15.668  55.634  1.00 26.50           N
    ATOM   1498  CA  GLN A 192      64.916  16.350  56.919  1.00 25.13           C
    ATOM   1499  C   GLN A 192      65.781  17.609  57.064  1.00 20.38           C
    ATOM   1500  O   GLN A 192      65.542  18.364  58.006  1.00 24.28           O
    ATOM   1501  CB  GLN A 192      65.224  15.410  58.105  1.00 25.76           C
    ATOM   1502  CG  GLN A 192      65.073  13.919  57.812  1.00 33.95           C
    ATOM   1503  CD  GLN A 192      66.426  13.384  57.370  1.00 39.63           C
    ATOM   1504  OE1 GLN A 192      67.417  13.577  58.071  1.00 40.99           O
    ATOM   1505  NE2 GLN A 192      66.479  12.751  56.200  1.00 28.78           N
    ATOM   1506  H   GLN A 192      65.864  14.974  55.608  1.00  0.00           H
    ATOM   1507  HA  GLN A 192      63.871  16.654  57.004  1.00  0.00           H
    ATOM   1508  HB2 GLN A 192      66.212  15.627  58.510  1.00  0.00           H
    ATOM   1509  HB3 GLN A 192      64.595  15.623  58.950  1.00  0.00           H
    ATOM   1510  HG2 GLN A 192      64.749  13.406  58.709  1.00  0.00           H
    ATOM   1511  HG3 GLN A 192      64.288  13.715  57.082  1.00  0.00           H
    ATOM   1512 HE21 GLN A 192      65.642  12.598  55.656  1.00  0.00           H
    ATOM   1513 HE22 GLN A 192      67.362  12.384  55.868  1.00  0.00           H
    ATOM   1514  N   HIS A 193      66.775  17.803  56.180  1.00 13.11           N
    ATOM   1515  CA  HIS A 193      67.697  18.931  56.249  1.00 15.23           C
    ATOM   1516  C   HIS A 193      67.029  20.231  55.823  1.00 16.36           C
    ATOM   1517  O   HIS A 193      66.543  20.337  54.698  1.00 18.62           O
    ATOM   1518  CB  HIS A 193      68.944  18.693  55.393  1.00 10.41           C
    ATOM   1519  CG  HIS A 193      69.911  17.701  55.970  1.00  6.77           C
    ATOM   1520  ND1 HIS A 193      71.288  17.852  55.812  1.00  9.76           N
    ATOM   1521  CD2 HIS A 193      69.669  16.550  56.689  1.00  5.25           C
    ATOM   1522  CE1 HIS A 193      71.820  16.790  56.392  1.00  9.84           C
    ATOM   1523  NE2 HIS A 193      70.907  15.991  56.934  1.00  5.47           N
    ATOM   1524  H   HIS A 193      66.906  17.157  55.413  1.00  0.00           H
    ATOM   1525  HA  HIS A 193      68.033  19.025  57.278  1.00  0.00           H
    ATOM   1526  HB2 HIS A 193      68.666  18.357  54.393  1.00  0.00           H
    ATOM   1527  HB3 HIS A 193      69.483  19.632  55.252  1.00  0.00           H
    ATOM   1528  HD2 HIS A 193      68.754  16.085  57.026  1.00  0.00           H
    ATOM   1529  HE2 HIS A 193      71.097  15.127  57.441  1.00  0.00           H
    ATOM   1530  HE1 HIS A 193      72.881  16.599  56.418  1.00  0.00           H
    ATOM   1531  N   LEU A 194      67.066  21.205  56.735  1.00 19.32           N
    ATOM   1532  CA  LEU A 194      66.576  22.555  56.521  1.00 20.34           C
    ATOM   1533  C   LEU A 194      67.429  23.317  55.489  1.00 18.35           C
    ATOM   1534  O   LEU A 194      66.851  23.980  54.636  1.00 22.16           O
    ATOM   1535  CB  LEU A 194      66.448  23.218  57.914  1.00 21.00           C
    ATOM   1536  CG  LEU A 194      65.982  24.690  58.014  1.00 18.76           C
    ATOM   1537  CD1 LEU A 194      67.155  25.663  58.152  1.00 17.22           C
    ATOM   1538  CD2 LEU A 194      64.975  25.137  56.941  1.00 17.07           C
    ATOM   1539  H   LEU A 194      67.483  21.020  57.639  1.00  0.00           H
    ATOM   1540  HA  LEU A 194      65.575  22.457  56.098  1.00  0.00           H
    ATOM   1541  HB2 LEU A 194      65.718  22.622  58.467  1.00  0.00           H
    ATOM   1542  HB3 LEU A 194      67.378  23.094  58.471  1.00  0.00           H
    ATOM   1543  HG  LEU A 194      65.454  24.754  58.967  1.00  0.00           H
    ATOM   1544 HD11 LEU A 194      66.824  26.624  58.546  1.00  0.00           H
    ATOM   1545 HD12 LEU A 194      67.918  25.283  58.832  1.00  0.00           H
    ATOM   1546 HD13 LEU A 194      67.634  25.820  57.188  1.00  0.00           H
    ATOM   1547 HD21 LEU A 194      64.216  25.784  57.374  1.00  0.00           H
    ATOM   1548 HD22 LEU A 194      65.459  25.715  56.156  1.00  0.00           H
    ATOM   1549 HD23 LEU A 194      64.456  24.304  56.473  1.00  0.00           H
    ATOM   1550  N   ILE A 195      68.763  23.174  55.536  1.00 14.66           N
    ATOM   1551  CA  ILE A 195      69.675  23.876  54.635  1.00 15.53           C
    ATOM   1552  C   ILE A 195      70.185  22.942  53.527  1.00 19.46           C
    ATOM   1553  O   ILE A 195      70.719  21.872  53.812  1.00 13.67           O
    ATOM   1554  CB  ILE A 195      70.913  24.467  55.368  1.00 16.85           C
    ATOM   1555  CG1 ILE A 195      70.471  25.371  56.539  1.00 20.14           C
    ATOM   1556  CG2 ILE A 195      71.826  25.267  54.406  1.00 17.31           C
    ATOM   1557  CD1 ILE A 195      71.625  26.077  57.263  1.00 16.53           C
    ATOM   1558  H   ILE A 195      69.189  22.605  56.257  1.00  0.00           H
    ATOM   1559  HA  ILE A 195      69.158  24.721  54.188  1.00  0.00           H
    ATOM   1560  HB  ILE A 195      71.501  23.648  55.785  1.00  0.00           H
    ATOM   1561 HG12 ILE A 195      69.783  26.125  56.160  1.00  0.00           H
    ATOM   1562 HG13 ILE A 195      69.912  24.791  57.274  1.00  0.00           H
    ATOM   1563 HG21 ILE A 195      72.735  25.610  54.898  1.00  0.00           H
    ATOM   1564 HG22 ILE A 195      72.165  24.687  53.552  1.00  0.00           H
    ATOM   1565 HG23 ILE A 195      71.309  26.143  54.014  1.00  0.00           H
    ATOM   1566 HD11 ILE A 195      71.290  26.526  58.195  1.00  0.00           H
    ATOM   1567 HD12 ILE A 195      72.435  25.386  57.499  1.00  0.00           H
    ATOM   1568 HD13 ILE A 195      72.032  26.890  56.662  1.00  0.00           H
    ATOM   1569  N   ARG A 196      70.073  23.416  52.284  1.00 21.40           N
    ATOM   1570  CA  ARG A 196      70.684  22.791  51.117  1.00 20.33           C
    ATOM   1571  C   ARG A 196      71.673  23.750  50.480  1.00 18.06           C
    ATOM   1572  O   ARG A 196      71.603  24.948  50.737  1.00 18.25           O
    ATOM   1573  CB  ARG A 196      69.604  22.411  50.101  1.00 14.85           C
    ATOM   1574  CG  ARG A 196      68.548  21.487  50.701  1.00 10.90           C
    ATOM   1575  CD  ARG A 196      67.612  20.985  49.618  1.00  9.17           C
    ATOM   1576  NE  ARG A 196      66.622  20.050  50.152  1.00 15.36           N
    ATOM   1577  CZ  ARG A 196      65.511  19.709  49.489  1.00 15.30           C
    ATOM   1578  NH1 ARG A 196      65.285  20.201  48.266  1.00 14.15           N
    ATOM   1579  NH2 ARG A 196      64.624  18.881  50.050  1.00 16.86           N1+
    ATOM   1580  H   ARG A 196      69.627  24.312  52.129  1.00  0.00           H
    ATOM   1581  HA  ARG A 196      71.238  21.899  51.403  1.00  0.00           H
    ATOM   1582  HB2 ARG A 196      69.126  23.299  49.690  1.00  0.00           H
    ATOM   1583  HB3 ARG A 196      70.076  21.907  49.256  1.00  0.00           H
    ATOM   1584  HG2 ARG A 196      68.997  20.652  51.240  1.00  0.00           H
    ATOM   1585  HG3 ARG A 196      67.975  22.054  51.434  1.00  0.00           H
    ATOM   1586  HD2 ARG A 196      67.049  21.852  49.274  1.00  0.00           H
    ATOM   1587  HD3 ARG A 196      68.138  20.580  48.754  1.00  0.00           H
    ATOM   1588  HE  ARG A 196      66.815  19.634  51.052  1.00  0.00           H
    ATOM   1589 HH11 ARG A 196      65.935  20.854  47.843  1.00  0.00           H
    ATOM   1590 HH12 ARG A 196      64.468  19.912  47.734  1.00  0.00           H
    ATOM   1591 HH21 ARG A 196      64.796  18.495  50.967  1.00  0.00           H
    ATOM   1592 HH22 ARG A 196      63.750  18.669  49.584  1.00  0.00           H
    ATOM   1593  N   VAL A 197      72.553  23.203  49.634  1.00 16.70           N
    ATOM   1594  CA  VAL A 197      73.358  24.010  48.733  1.00 16.77           C
    ATOM   1595  C   VAL A 197      72.917  23.717  47.302  1.00 24.08           C
    ATOM   1596  O   VAL A 197      72.743  22.564  46.916  1.00 25.01           O
    ATOM   1597  CB  VAL A 197      74.880  23.787  48.874  1.00 12.99           C
    ATOM   1598  CG1 VAL A 197      75.679  24.453  47.737  1.00 14.20           C
    ATOM   1599  CG2 VAL A 197      75.370  24.338  50.218  1.00 12.62           C
    ATOM   1600  H   VAL A 197      72.563  22.204  49.482  1.00  0.00           H
    ATOM   1601  HA  VAL A 197      73.186  25.069  48.921  1.00  0.00           H
    ATOM   1602  HB  VAL A 197      75.093  22.724  48.858  1.00  0.00           H
    ATOM   1603 HG11 VAL A 197      76.719  24.596  48.001  1.00  0.00           H
    ATOM   1604 HG12 VAL A 197      75.643  23.875  46.814  1.00  0.00           H
    ATOM   1605 HG13 VAL A 197      75.300  25.452  47.520  1.00  0.00           H
    ATOM   1606 HG21 VAL A 197      76.419  24.102  50.392  1.00  0.00           H
    ATOM   1607 HG22 VAL A 197      75.272  25.422  50.229  1.00  0.00           H
    ATOM   1608 HG23 VAL A 197      74.800  23.940  51.056  1.00  0.00           H
    ATOM   1609  N   GLU A 198      72.750  24.805  46.558  1.00 25.53           N
    ATOM   1610  CA  GLU A 198      72.249  24.852  45.207  1.00 26.37           C
    ATOM   1611  C   GLU A 198      73.423  24.890  44.215  1.00 28.44           C
    ATOM   1612  O   GLU A 198      74.422  25.565  44.467  1.00 23.37           O
    ATOM   1613  CB  GLU A 198      71.407  26.132  45.117  1.00 32.97           C
    ATOM   1614  CG  GLU A 198      70.367  26.087  44.000  1.00 36.80           C
    ATOM   1615  CD  GLU A 198      69.809  27.473  43.720  1.00 37.04           C
    ATOM   1616  OE1 GLU A 198      69.069  27.991  44.582  1.00 31.19           O
    ATOM   1617  OE2 GLU A 198      70.159  28.008  42.648  1.00 43.32           O1-
    ATOM   1618  H   GLU A 198      72.946  25.711  46.971  1.00  0.00           H
    ATOM   1619  HA  GLU A 198      71.615  23.981  45.022  1.00  0.00           H
    ATOM   1620  HB2 GLU A 198      70.898  26.320  46.063  1.00  0.00           H
    ATOM   1621  HB3 GLU A 198      72.060  26.985  44.962  1.00  0.00           H
    ATOM   1622  HG2 GLU A 198      70.806  25.697  43.081  1.00  0.00           H
    ATOM   1623  HG3 GLU A 198      69.550  25.416  44.268  1.00  0.00           H
    ATOM   1624  N   GLY A 199      73.245  24.198  43.079  1.00 25.55           N
    ATOM   1625  CA  GLY A 199      74.137  24.247  41.919  1.00 21.14           C
    ATOM   1626  C   GLY A 199      75.585  23.811  42.199  1.00 24.51           C
    ATOM   1627  O   GLY A 199      76.501  24.396  41.623  1.00 30.14           O
    ATOM   1628  H   GLY A 199      72.399  23.657  42.970  1.00  0.00           H
    ATOM   1629  HA2 GLY A 199      73.728  23.603  41.141  1.00  0.00           H
    ATOM   1630  HA3 GLY A 199      74.136  25.262  41.518  1.00  0.00           H
    ATOM   1631  N   ASN A 200      75.798  22.815  43.074  1.00 21.10           N
    ATOM   1632  CA  ASN A 200      77.118  22.283  43.392  1.00 18.72           C
    ATOM   1633  C   ASN A 200      76.999  20.761  43.538  1.00 28.10           C
    ATOM   1634  O   ASN A 200      76.560  20.288  44.582  1.00 32.82           O
    ATOM   1635  CB  ASN A 200      77.675  22.967  44.663  1.00 15.17           C
    ATOM   1636  CG  ASN A 200      79.174  22.733  44.875  1.00 22.63           C
    ATOM   1637  OD1 ASN A 200      79.698  21.650  44.617  1.00 29.17           O
    ATOM   1638  ND2 ASN A 200      79.871  23.755  45.363  1.00 22.65           N
    ATOM   1639  H   ASN A 200      75.014  22.367  43.535  1.00  0.00           H
    ATOM   1640  HA  ASN A 200      77.827  22.449  42.579  1.00  0.00           H
    ATOM   1641  HB2 ASN A 200      77.518  24.044  44.577  1.00  0.00           H
    ATOM   1642  HB3 ASN A 200      77.133  22.652  45.554  1.00  0.00           H
    ATOM   1643 HD21 ASN A 200      79.428  24.638  45.565  1.00  0.00           H
    ATOM   1644 HD22 ASN A 200      80.874  23.667  45.500  1.00  0.00           H
    ATOM   1645  N   LEU A 201      77.418  20.017  42.500  1.00 30.44           N
    ATOM   1646  CA  LEU A 201      77.425  18.548  42.498  1.00 33.85           C
    ATOM   1647  C   LEU A 201      78.443  17.892  43.439  1.00 33.23           C
    ATOM   1648  O   LEU A 201      78.367  16.677  43.620  1.00 24.91           O
    ATOM   1649  CB  LEU A 201      77.660  18.006  41.068  1.00 33.71           C
    ATOM   1650  CG  LEU A 201      76.397  17.721  40.236  1.00 31.25           C
    ATOM   1651  CD1 LEU A 201      76.780  16.860  39.017  1.00 30.08           C
    ATOM   1652  CD2 LEU A 201      75.251  17.052  41.025  1.00 30.82           C
    ATOM   1653  H   LEU A 201      77.761  20.471  41.664  1.00  0.00           H
    ATOM   1654  HA  LEU A 201      76.457  18.237  42.885  1.00  0.00           H
    ATOM   1655  HB2 LEU A 201      78.330  18.668  40.517  1.00  0.00           H
    ATOM   1656  HB3 LEU A 201      78.197  17.057  41.119  1.00  0.00           H
    ATOM   1657  HG  LEU A 201      76.024  18.679  39.871  1.00  0.00           H
    ATOM   1658 HD11 LEU A 201      76.236  17.189  38.131  1.00  0.00           H
    ATOM   1659 HD12 LEU A 201      77.843  16.930  38.779  1.00  0.00           H
    ATOM   1660 HD13 LEU A 201      76.553  15.804  39.170  1.00  0.00           H
    ATOM   1661 HD21 LEU A 201      74.660  16.386  40.396  1.00  0.00           H
    ATOM   1662 HD22 LEU A 201      75.618  16.449  41.857  1.00  0.00           H
    ATOM   1663 HD23 LEU A 201      74.563  17.798  41.424  1.00  0.00           H
    ATOM   1664  N   ARG A 202      79.363  18.674  44.017  1.00 32.12           N
    ATOM   1665  CA  ARG A 202      80.292  18.194  45.034  1.00 25.20           C
    ATOM   1666  C   ARG A 202      79.768  18.454  46.452  1.00 23.12           C
    ATOM   1667  O   ARG A 202      80.510  18.245  47.406  1.00 19.27           O
    ATOM   1668  CB  ARG A 202      81.694  18.775  44.797  1.00 23.67           C
    ATOM   1669  CG  ARG A 202      82.226  18.451  43.390  1.00 28.10           C
    ATOM   1670  CD  ARG A 202      83.747  18.573  43.275  1.00 29.11           C
    ATOM   1671  NE  ARG A 202      84.206  19.965  43.340  1.00 34.97           N
    ATOM   1672  CZ  ARG A 202      85.361  20.386  43.884  1.00 44.09           C
    ATOM   1673  NH1 ARG A 202      86.193  19.545  44.517  1.00 49.53           N
    ATOM   1674  NH2 ARG A 202      85.691  21.681  43.793  1.00 45.33           N1+
    ATOM   1675  H   ARG A 202      79.371  19.669  43.834  1.00  0.00           H
    ATOM   1676  HA  ARG A 202      80.395  17.109  44.964  1.00  0.00           H
    ATOM   1677  HB2 ARG A 202      81.715  19.852  44.968  1.00  0.00           H
    ATOM   1678  HB3 ARG A 202      82.365  18.334  45.536  1.00  0.00           H
    ATOM   1679  HG2 ARG A 202      81.945  17.444  43.076  1.00  0.00           H
    ATOM   1680  HG3 ARG A 202      81.744  19.129  42.684  1.00  0.00           H
    ATOM   1681  HD2 ARG A 202      84.153  18.084  44.159  1.00  0.00           H
    ATOM   1682  HD3 ARG A 202      84.160  18.028  42.427  1.00  0.00           H
    ATOM   1683  HE  ARG A 202      83.619  20.642  42.868  1.00  0.00           H
    ATOM   1684 HH11 ARG A 202      85.945  18.572  44.633  1.00  0.00           H
    ATOM   1685 HH12 ARG A 202      87.058  19.876  44.916  1.00  0.00           H
    ATOM   1686 HH21 ARG A 202      85.075  22.331  43.321  1.00  0.00           H
    ATOM   1687 HH22 ARG A 202      86.558  22.020  44.180  1.00  0.00           H
    ATOM   1688  N   VAL A 203      78.495  18.861  46.583  1.00 21.97           N
    ATOM   1689  CA  VAL A 203      77.819  19.012  47.865  1.00 22.96           C
    ATOM   1690  C   VAL A 203      77.712  17.672  48.621  1.00 27.72           C
    ATOM   1691  O   VAL A 203      77.471  16.631  48.011  1.00 29.56           O
    ATOM   1692  CB  VAL A 203      76.409  19.633  47.687  1.00 14.85           C
    ATOM   1693  CG1 VAL A 203      75.387  18.698  47.010  1.00  9.81           C
    ATOM   1694  CG2 VAL A 203      75.836  20.137  49.013  1.00 20.34           C
    ATOM   1695  H   VAL A 203      77.932  19.030  45.761  1.00  0.00           H
    ATOM   1696  HA  VAL A 203      78.431  19.689  48.459  1.00  0.00           H
    ATOM   1697  HB  VAL A 203      76.523  20.516  47.059  1.00  0.00           H
    ATOM   1698 HG11 VAL A 203      74.501  19.255  46.714  1.00  0.00           H
    ATOM   1699 HG12 VAL A 203      75.792  18.245  46.106  1.00  0.00           H
    ATOM   1700 HG13 VAL A 203      75.062  17.891  47.665  1.00  0.00           H
    ATOM   1701 HG21 VAL A 203      74.847  20.570  48.866  1.00  0.00           H
    ATOM   1702 HG22 VAL A 203      75.709  19.324  49.725  1.00  0.00           H
    ATOM   1703 HG23 VAL A 203      76.483  20.889  49.466  1.00  0.00           H
    ATOM   1704  N   GLU A 204      77.921  17.737  49.938  1.00 26.75           N
    ATOM   1705  CA  GLU A 204      77.906  16.603  50.845  1.00 22.86           C
    ATOM   1706  C   GLU A 204      77.065  16.953  52.068  1.00 25.66           C
    ATOM   1707  O   GLU A 204      77.211  18.038  52.616  1.00 24.82           O
    ATOM   1708  CB  GLU A 204      79.349  16.244  51.239  1.00 19.51           C
    ATOM   1709  CG  GLU A 204      79.449  15.179  52.352  1.00 15.34           C
    ATOM   1710  CD  GLU A 204      80.881  14.923  52.795  1.00 18.64           C
    ATOM   1711  OE1 GLU A 204      81.796  15.015  51.952  1.00 20.16           O
    ATOM   1712  OE2 GLU A 204      81.054  14.658  54.000  1.00 21.16           O1-
    ATOM   1713  H   GLU A 204      78.148  18.629  50.361  1.00  0.00           H
    ATOM   1714  HA  GLU A 204      77.455  15.737  50.359  1.00  0.00           H
    ATOM   1715  HB2 GLU A 204      79.871  15.888  50.350  1.00  0.00           H
    ATOM   1716  HB3 GLU A 204      79.875  17.148  51.554  1.00  0.00           H
    ATOM   1717  HG2 GLU A 204      78.901  15.478  53.245  1.00  0.00           H
    ATOM   1718  HG3 GLU A 204      79.012  14.238  52.016  1.00  0.00           H
    ATOM   1719  N   TYR A 205      76.247  15.992  52.499  1.00 23.55           N
    ATOM   1720  CA  TYR A 205      75.372  16.111  53.651  1.00 17.38           C
    ATOM   1721  C   TYR A 205      75.801  15.120  54.729  1.00 18.81           C
    ATOM   1722  O   TYR A 205      76.082  13.959  54.425  1.00 15.08           O
    ATOM   1723  CB  TYR A 205      73.935  15.853  53.188  1.00 12.58           C
    ATOM   1724  CG  TYR A 205      73.416  16.944  52.273  1.00 11.17           C
    ATOM   1725  CD1 TYR A 205      73.528  16.833  50.871  1.00  9.39           C
    ATOM   1726  CD2 TYR A 205      72.827  18.088  52.836  1.00 11.33           C
    ATOM   1727  CE1 TYR A 205      73.039  17.862  50.043  1.00 14.10           C
    ATOM   1728  CE2 TYR A 205      72.306  19.097  52.014  1.00 13.32           C
    ATOM   1729  CZ  TYR A 205      72.423  18.990  50.621  1.00 15.54           C
    ATOM   1730  OH  TYR A 205      71.936  19.991  49.844  1.00 13.04           O
    ATOM   1731  H   TYR A 205      76.197  15.113  51.997  1.00  0.00           H
    ATOM   1732  HA  TYR A 205      75.414  17.119  54.064  1.00  0.00           H
    ATOM   1733  HB2 TYR A 205      73.854  14.885  52.691  1.00  0.00           H
    ATOM   1734  HB3 TYR A 205      73.273  15.799  54.053  1.00  0.00           H
    ATOM   1735  HD1 TYR A 205      73.985  15.961  50.429  1.00  0.00           H
    ATOM   1736  HD2 TYR A 205      72.792  18.204  53.904  1.00  0.00           H
    ATOM   1737  HE1 TYR A 205      73.117  17.773  48.970  1.00  0.00           H
    ATOM   1738  HE2 TYR A 205      71.844  19.964  52.460  1.00  0.00           H
    ATOM   1739  HH  TYR A 205      72.000  19.802  48.898  1.00  0.00           H
    ATOM   1740  N   LEU A 206      75.822  15.614  55.972  1.00 19.17           N
    ATOM   1741  CA  LEU A 206      76.155  14.834  57.151  1.00 15.81           C
    ATOM   1742  C   LEU A 206      74.940  14.792  58.074  1.00 21.00           C
    ATOM   1743  O   LEU A 206      74.329  15.822  58.349  1.00 19.89           O
    ATOM   1744  CB  LEU A 206      77.374  15.471  57.859  1.00 19.46           C
    ATOM   1745  CG  LEU A 206      77.928  14.677  59.068  1.00 18.85           C
    ATOM   1746  CD1 LEU A 206      78.418  13.267  58.683  1.00 15.36           C
    ATOM   1747  CD2 LEU A 206      79.007  15.466  59.831  1.00 11.21           C
    ATOM   1748  H   LEU A 206      75.560  16.578  56.133  1.00  0.00           H
    ATOM   1749  HA  LEU A 206      76.409  13.812  56.866  1.00  0.00           H
    ATOM   1750  HB2 LEU A 206      78.175  15.604  57.131  1.00  0.00           H
    ATOM   1751  HB3 LEU A 206      77.102  16.475  58.183  1.00  0.00           H
    ATOM   1752  HG  LEU A 206      77.114  14.561  59.780  1.00  0.00           H
    ATOM   1753 HD11 LEU A 206      79.416  13.057  59.068  1.00  0.00           H
    ATOM   1754 HD12 LEU A 206      77.753  12.504  59.087  1.00  0.00           H
    ATOM   1755 HD13 LEU A 206      78.462  13.126  57.603  1.00  0.00           H
    ATOM   1756 HD21 LEU A 206      78.966  15.239  60.897  1.00  0.00           H
    ATOM   1757 HD22 LEU A 206      80.014  15.235  59.483  1.00  0.00           H
    ATOM   1758 HD23 LEU A 206      78.863  16.541  59.738  1.00  0.00           H
    ATOM   1759  N   ASP A 207      74.690  13.588  58.588  1.00 23.86           N
    ATOM   1760  CA  ASP A 207      73.946  13.350  59.810  1.00 20.68           C
    ATOM   1761  C   ASP A 207      74.991  12.889  60.789  1.00 22.11           C
    ATOM   1762  O   ASP A 207      75.489  11.770  60.646  1.00 17.87           O
    ATOM   1763  CB  ASP A 207      72.884  12.253  59.689  1.00 17.84           C
    ATOM   1764  CG  ASP A 207      71.703  12.630  58.825  1.00 28.30           C
    ATOM   1765  OD1 ASP A 207      71.621  13.789  58.374  1.00 34.38           O
    ATOM   1766  OD2 ASP A 207      70.852  11.732  58.682  1.00 27.19           O1-
    ATOM   1767  H   ASP A 207      75.224  12.797  58.260  1.00  0.00           H
    ATOM   1768  HA  ASP A 207      73.482  14.266  60.185  1.00  0.00           H
    ATOM   1769  HB2 ASP A 207      73.333  11.356  59.260  1.00  0.00           H
    ATOM   1770  HB3 ASP A 207      72.500  12.009  60.680  1.00  0.00           H
    ATOM   1771  N   ASP A 208      75.349  13.768  61.730  1.00 21.15           N
    ATOM   1772  CA  ASP A 208      76.401  13.445  62.672  1.00 18.86           C
    ATOM   1773  C   ASP A 208      75.954  12.254  63.528  1.00 18.90           C
    ATOM   1774  O   ASP A 208      74.882  12.302  64.131  1.00 20.07           O
    ATOM   1775  CB  ASP A 208      76.809  14.643  63.516  1.00 18.38           C
    ATOM   1776  CG  ASP A 208      78.152  14.431  64.218  1.00 21.05           C
    ATOM   1777  OD1 ASP A 208      78.199  13.579  65.132  1.00 26.44           O
    ATOM   1778  OD2 ASP A 208      79.103  15.151  63.846  1.00 20.64           O1-
    ATOM   1779  H   ASP A 208      74.910  14.677  61.802  1.00  0.00           H
    ATOM   1780  HA  ASP A 208      77.263  13.167  62.070  1.00  0.00           H
    ATOM   1781  HB2 ASP A 208      76.856  15.538  62.896  1.00  0.00           H
    ATOM   1782  HB3 ASP A 208      76.030  14.830  64.238  1.00  0.00           H
    ATOM   1783  N   ARG A 209      76.766  11.190  63.500  1.00 23.60           N
    ATOM   1784  CA  ARG A 209      76.469   9.937  64.183  1.00 27.27           C
    ATOM   1785  C   ARG A 209      76.376  10.099  65.708  1.00 22.56           C
    ATOM   1786  O   ARG A 209      75.640   9.343  66.340  1.00 23.95           O
    ATOM   1787  CB  ARG A 209      77.497   8.857  63.785  1.00 30.46           C
    ATOM   1788  CG  ARG A 209      78.936   9.085  64.298  1.00 35.52           C
    ATOM   1789  CD  ARG A 209      79.902   7.938  63.961  1.00 37.57           C
    ATOM   1790  NE  ARG A 209      79.446   6.603  64.393  1.00 45.66           N
    ATOM   1791  CZ  ARG A 209      78.837   6.255  65.542  1.00 51.63           C
    ATOM   1792  NH1 ARG A 209      78.570   7.128  66.514  1.00 55.05           N
    ATOM   1793  NH2 ARG A 209      78.479   4.982  65.738  1.00 52.95           N1+
    ATOM   1794  H   ARG A 209      77.622  11.223  62.959  1.00  0.00           H
    ATOM   1795  HA  ARG A 209      75.490   9.604  63.833  1.00  0.00           H
    ATOM   1796  HB2 ARG A 209      77.128   7.908  64.174  1.00  0.00           H
    ATOM   1797  HB3 ARG A 209      77.511   8.741  62.701  1.00  0.00           H
    ATOM   1798  HG2 ARG A 209      79.303   9.969  63.783  1.00  0.00           H
    ATOM   1799  HG3 ARG A 209      79.008   9.346  65.349  1.00  0.00           H
    ATOM   1800  HD2 ARG A 209      80.230   7.961  62.921  1.00  0.00           H
    ATOM   1801  HD3 ARG A 209      80.802   8.111  64.547  1.00  0.00           H
    ATOM   1802  HE  ARG A 209      79.646   5.863  63.735  1.00  0.00           H
    ATOM   1803 HH11 ARG A 209      78.867   8.091  66.454  1.00  0.00           H
    ATOM   1804 HH12 ARG A 209      78.074   6.817  67.342  1.00  0.00           H
    ATOM   1805 HH21 ARG A 209      78.659   4.278  65.039  1.00  0.00           H
    ATOM   1806 HH22 ARG A 209      78.012   4.727  66.598  1.00  0.00           H
    ATOM   1807  N   ASN A 210      77.111  11.078  66.260  1.00 23.73           N
    ATOM   1808  CA  ASN A 210      77.202  11.306  67.696  1.00 22.36           C
    ATOM   1809  C   ASN A 210      76.229  12.363  68.188  1.00 21.25           C
    ATOM   1810  O   ASN A 210      75.631  12.120  69.231  1.00 24.93           O
    ATOM   1811  CB  ASN A 210      78.656  11.635  68.085  1.00 23.40           C
    ATOM   1812  CG  ASN A 210      79.550  10.403  67.957  1.00 20.82           C
    ATOM   1813  OD1 ASN A 210      79.104   9.286  68.203  1.00 22.49           O
    ATOM   1814  ND2 ASN A 210      80.809  10.589  67.572  1.00 23.40           N
    ATOM   1815  H   ASN A 210      77.643  11.710  65.672  1.00  0.00           H
    ATOM   1816  HA  ASN A 210      76.911  10.378  68.190  1.00  0.00           H
    ATOM   1817  HB2 ASN A 210      79.043  12.466  67.493  1.00  0.00           H
    ATOM   1818  HB3 ASN A 210      78.702  11.949  69.129  1.00  0.00           H
    ATOM   1819 HD21 ASN A 210      81.156  11.518  67.383  1.00  0.00           H
    ATOM   1820 HD22 ASN A 210      81.424   9.793  67.458  1.00  0.00           H
    ATOM   1821  N   THR A 211      76.069  13.486  67.471  1.00 21.25           N
    ATOM   1822  CA  THR A 211      75.207  14.571  67.944  1.00 16.64           C
    ATOM   1823  C   THR A 211      73.824  14.642  67.280  1.00 20.66           C
    ATOM   1824  O   THR A 211      73.030  15.496  67.673  1.00 22.63           O
    ATOM   1825  CB  THR A 211      75.896  15.947  67.850  1.00 13.28           C
    ATOM   1826  OG1 THR A 211      77.128  15.901  68.533  1.00 22.49           O
    ATOM   1827  CG2 THR A 211      76.219  16.406  66.441  1.00 11.21           C
    ATOM   1828  H   THR A 211      76.625  13.654  66.641  1.00  0.00           H
    ATOM   1829  HA  THR A 211      74.972  14.417  68.985  1.00  0.00           H
    ATOM   1830  HB  THR A 211      75.273  16.700  68.337  1.00  0.00           H
    ATOM   1831 HG21 THR A 211      76.198  17.489  66.419  1.00  0.00           H
    ATOM   1832 HG22 THR A 211      77.208  16.082  66.126  1.00  0.00           H
    ATOM   1833 HG23 THR A 211      75.500  16.042  65.728  1.00  0.00           H
    ATOM   1834  HG1 THR A 211      76.969  15.543  69.410  1.00  0.00           H
    ATOM   1835  N   PHE A 212      73.558  13.769  66.293  1.00 16.71           N
    ATOM   1836  CA  PHE A 212      72.324  13.695  65.489  1.00 10.24           C
    ATOM   1837  C   PHE A 212      72.086  14.939  64.604  1.00 19.63           C
    ATOM   1838  O   PHE A 212      71.044  15.052  63.963  1.00 20.22           O
    ATOM   1839  CB  PHE A 212      71.091  13.376  66.371  1.00 12.11           C
    ATOM   1840  CG  PHE A 212      71.161  12.135  67.254  1.00 21.23           C
    ATOM   1841  CD1 PHE A 212      71.928  11.000  66.902  1.00 23.78           C
    ATOM   1842  CD2 PHE A 212      70.308  12.047  68.373  1.00 23.30           C
    ATOM   1843  CE1 PHE A 212      71.897   9.863  67.698  1.00 21.30           C
    ATOM   1844  CE2 PHE A 212      70.281  10.899  69.149  1.00 26.03           C
    ATOM   1845  CZ  PHE A 212      71.082   9.814  68.822  1.00 23.58           C
    ATOM   1846  H   PHE A 212      74.275  13.106  66.030  1.00  0.00           H
    ATOM   1847  HA  PHE A 212      72.455  12.872  64.790  1.00  0.00           H
    ATOM   1848  HB2 PHE A 212      70.871  14.227  67.014  1.00  0.00           H
    ATOM   1849  HB3 PHE A 212      70.210  13.262  65.738  1.00  0.00           H
    ATOM   1850  HD1 PHE A 212      72.558  11.003  66.025  1.00  0.00           H
    ATOM   1851  HD2 PHE A 212      69.670  12.873  68.637  1.00  0.00           H
    ATOM   1852  HE1 PHE A 212      72.509   9.011  67.439  1.00  0.00           H
    ATOM   1853  HE2 PHE A 212      69.648  10.860  70.021  1.00  0.00           H
    ATOM   1854  HZ  PHE A 212      71.069   8.927  69.438  1.00  0.00           H
    ATOM   1855  N   ARG A 213      73.059  15.852  64.591  1.00 22.02           N
    ATOM   1856  CA  ARG A 213      72.971  17.193  64.047  1.00 20.13           C
    ATOM   1857  C   ARG A 213      73.239  17.190  62.539  1.00 12.01           C
    ATOM   1858  O   ARG A 213      74.060  16.399  62.075  1.00 13.35           O
    ATOM   1859  CB  ARG A 213      73.992  18.023  64.836  1.00 18.87           C
    ATOM   1860  CG  ARG A 213      73.522  19.410  65.202  1.00 18.64           C
    ATOM   1861  CD  ARG A 213      73.531  19.658  66.717  1.00 19.36           C
    ATOM   1862  NE  ARG A 213      72.730  20.841  66.909  1.00 23.96           N
    ATOM   1863  CZ  ARG A 213      72.267  21.390  68.023  1.00 21.85           C
    ATOM   1864  NH1 ARG A 213      72.544  20.945  69.253  1.00 22.22           N
    ATOM   1865  NH2 ARG A 213      71.482  22.435  67.837  1.00 22.28           N1+
    ATOM   1866  H   ARG A 213      73.905  15.649  65.101  1.00  0.00           H
    ATOM   1867  HA  ARG A 213      71.959  17.564  64.221  1.00  0.00           H
    ATOM   1868  HB2 ARG A 213      74.077  17.542  65.803  1.00  0.00           H
    ATOM   1869  HB3 ARG A 213      74.988  18.018  64.388  1.00  0.00           H
    ATOM   1870  HG2 ARG A 213      74.205  20.105  64.722  1.00  0.00           H
    ATOM   1871  HG3 ARG A 213      72.545  19.629  64.771  1.00  0.00           H
    ATOM   1872  HD2 ARG A 213      73.168  18.812  67.302  1.00  0.00           H
    ATOM   1873  HD3 ARG A 213      74.536  19.896  67.060  1.00  0.00           H
    ATOM   1874  HE  ARG A 213      72.414  21.267  66.045  1.00  0.00           H
    ATOM   1875 HH11 ARG A 213      73.136  20.138  69.387  1.00  0.00           H
    ATOM   1876 HH12 ARG A 213      72.172  21.430  70.060  1.00  0.00           H
    ATOM   1877 HH21 ARG A 213      71.235  22.727  66.903  1.00  0.00           H
    ATOM   1878 HH22 ARG A 213      71.092  22.939  68.627  1.00  0.00           H
    ATOM   1879  N   HIS A 214      72.528  18.053  61.798  1.00 16.33           N
    ATOM   1880  CA  HIS A 214      72.604  18.108  60.335  1.00 16.31           C
    ATOM   1881  C   HIS A 214      73.537  19.223  59.868  1.00 17.93           C
    ATOM   1882  O   HIS A 214      73.601  20.275  60.502  1.00 19.66           O
    ATOM   1883  CB  HIS A 214      71.212  18.354  59.730  1.00 11.34           C
    ATOM   1884  CG  HIS A 214      70.154  17.343  60.081  1.00 11.14           C
    ATOM   1885  ND1 HIS A 214      68.793  17.648  60.001  1.00 13.43           N
    ATOM   1886  CD2 HIS A 214      70.290  16.034  60.490  1.00 11.15           C
    ATOM   1887  CE1 HIS A 214      68.169  16.543  60.373  1.00 13.48           C
    ATOM   1888  NE2 HIS A 214      69.008  15.554  60.675  1.00 14.68           N
    ATOM   1889  H   HIS A 214      71.880  18.694  62.236  1.00  0.00           H
    ATOM   1890  HA  HIS A 214      72.966  17.156  59.949  1.00  0.00           H
    ATOM   1891  HB2 HIS A 214      70.847  19.338  60.016  1.00  0.00           H
    ATOM   1892  HB3 HIS A 214      71.284  18.384  58.644  1.00  0.00           H
    ATOM   1893  HD2 HIS A 214      71.166  15.423  60.661  1.00  0.00           H
    ATOM   1894  HE2 HIS A 214      68.754  14.631  61.000  1.00  0.00           H
    ATOM   1895  HE1 HIS A 214      67.095  16.457  60.445  1.00  0.00           H
    ATOM   1896  N   SER A 215      74.197  19.000  58.728  1.00 12.51           N
    ATOM   1897  CA  SER A 215      75.019  20.012  58.075  1.00 15.33           C
    ATOM   1898  C   SER A 215      75.227  19.664  56.600  1.00 19.13           C
    ATOM   1899  O   SER A 215      75.046  18.512  56.203  1.00 18.86           O
    ATOM   1900  CB  SER A 215      76.352  20.212  58.837  1.00 17.87           C
    ATOM   1901  OG  SER A 215      77.222  19.112  58.673  1.00 19.02           O
    ATOM   1902  H   SER A 215      74.101  18.118  58.243  1.00  0.00           H
    ATOM   1903  HA  SER A 215      74.457  20.948  58.100  1.00  0.00           H
    ATOM   1904  HB2 SER A 215      76.876  21.089  58.463  1.00  0.00           H
    ATOM   1905  HB3 SER A 215      76.187  20.389  59.898  1.00  0.00           H
    ATOM   1906  HG  SER A 215      77.417  19.029  57.735  1.00  0.00           H
    ATOM   1907  N   VAL A 216      75.628  20.685  55.835  1.00 20.38           N
    ATOM   1908  CA  VAL A 216      75.944  20.583  54.419  1.00 15.94           C
    ATOM   1909  C   VAL A 216      77.321  21.201  54.157  1.00 20.90           C
    ATOM   1910  O   VAL A 216      77.582  22.317  54.598  1.00 21.29           O
    ATOM   1911  CB  VAL A 216      74.863  21.260  53.537  1.00 13.30           C
    ATOM   1912  CG1 VAL A 216      74.507  22.702  53.936  1.00 12.59           C
    ATOM   1913  CG2 VAL A 216      75.217  21.208  52.044  1.00 16.57           C
    ATOM   1914  H   VAL A 216      75.744  21.601  56.248  1.00  0.00           H
    ATOM   1915  HA  VAL A 216      75.980  19.532  54.140  1.00  0.00           H
    ATOM   1916  HB  VAL A 216      73.954  20.673  53.669  1.00  0.00           H
    ATOM   1917 HG11 VAL A 216      73.694  23.067  53.311  1.00  0.00           H
    ATOM   1918 HG12 VAL A 216      74.176  22.774  54.972  1.00  0.00           H
    ATOM   1919 HG13 VAL A 216      75.347  23.381  53.795  1.00  0.00           H
    ATOM   1920 HG21 VAL A 216      74.369  21.509  51.429  1.00  0.00           H
    ATOM   1921 HG22 VAL A 216      76.058  21.853  51.786  1.00  0.00           H
    ATOM   1922 HG23 VAL A 216      75.475  20.187  51.767  1.00  0.00           H
    ATOM   1923  N   VAL A 217      78.162  20.449  53.440  1.00 18.07           N
    ATOM   1924  CA  VAL A 217      79.535  20.793  53.117  1.00 17.85           C
    ATOM   1925  C   VAL A 217      79.697  20.846  51.599  1.00 23.22           C
    ATOM   1926  O   VAL A 217      79.120  20.032  50.887  1.00 24.57           O
    ATOM   1927  CB  VAL A 217      80.574  19.741  53.614  1.00 25.27           C
    ATOM   1928  CG1 VAL A 217      81.837  20.443  54.131  1.00 18.50           C
    ATOM   1929  CG2 VAL A 217      80.034  18.764  54.666  1.00 29.39           C
    ATOM   1930  H   VAL A 217      77.848  19.547  53.099  1.00  0.00           H
    ATOM   1931  HA  VAL A 217      79.767  21.763  53.539  1.00  0.00           H
    ATOM   1932  HB  VAL A 217      80.907  19.102  52.792  1.00  0.00           H
    ATOM   1933 HG11 VAL A 217      82.595  19.721  54.433  1.00  0.00           H
    ATOM   1934 HG12 VAL A 217      82.266  21.076  53.356  1.00  0.00           H
    ATOM   1935 HG13 VAL A 217      81.616  21.074  54.988  1.00  0.00           H
    ATOM   1936 HG21 VAL A 217      80.828  18.110  55.025  1.00  0.00           H
    ATOM   1937 HG22 VAL A 217      79.623  19.296  55.524  1.00  0.00           H
    ATOM   1938 HG23 VAL A 217      79.256  18.118  54.261  1.00  0.00           H
    ATOM   1939  N   VAL A 218      80.540  21.767  51.145  1.00 25.02           N
    ATOM   1940  CA  VAL A 218      81.015  21.831  49.773  1.00 20.32           C
    ATOM   1941  C   VAL A 218      82.535  22.030  49.840  1.00 19.75           C
    ATOM   1942  O   VAL A 218      83.025  22.547  50.845  1.00 14.09           O
    ATOM   1943  CB  VAL A 218      80.364  23.015  49.010  1.00 22.50           C
    ATOM   1944  CG1 VAL A 218      78.853  22.816  48.870  1.00 20.56           C
    ATOM   1945  CG2 VAL A 218      80.676  24.420  49.564  1.00 23.51           C
    ATOM   1946  H   VAL A 218      80.977  22.411  51.791  1.00  0.00           H
    ATOM   1947  HA  VAL A 218      80.790  20.897  49.257  1.00  0.00           H
    ATOM   1948  HB  VAL A 218      80.755  23.002  47.994  1.00  0.00           H
    ATOM   1949 HG11 VAL A 218      78.472  23.562  48.182  1.00  0.00           H
    ATOM   1950 HG12 VAL A 218      78.617  21.849  48.436  1.00  0.00           H
    ATOM   1951 HG13 VAL A 218      78.329  22.907  49.821  1.00  0.00           H
    ATOM   1952 HG21 VAL A 218      80.151  25.188  48.996  1.00  0.00           H
    ATOM   1953 HG22 VAL A 218      80.366  24.517  50.605  1.00  0.00           H
    ATOM   1954 HG23 VAL A 218      81.737  24.661  49.509  1.00  0.00           H
    ATOM   1955  N   PRO A 219      83.276  21.652  48.779  1.00 20.30           N
    ATOM   1956  CA  PRO A 219      84.678  22.063  48.671  1.00 19.66           C
    ATOM   1957  C   PRO A 219      84.773  23.592  48.564  1.00 22.44           C
    ATOM   1958  O   PRO A 219      83.989  24.194  47.837  1.00 25.40           O
    ATOM   1959  CB  PRO A 219      85.194  21.358  47.407  1.00 23.81           C
    ATOM   1960  CG  PRO A 219      84.035  20.597  46.783  1.00 20.08           C
    ATOM   1961  CD  PRO A 219      82.807  20.900  47.620  1.00 20.98           C
    ATOM   1962  HA  PRO A 219      85.235  21.710  49.541  1.00  0.00           H
    ATOM   1963  HB2 PRO A 219      85.621  22.059  46.688  1.00  0.00           H
    ATOM   1964  HB3 PRO A 219      85.984  20.656  47.653  1.00  0.00           H
    ATOM   1965  HG2 PRO A 219      83.866  20.923  45.759  1.00  0.00           H
    ATOM   1966  HG3 PRO A 219      84.240  19.526  46.763  1.00  0.00           H
    ATOM   1967  HD2 PRO A 219      82.074  21.465  47.046  1.00  0.00           H
    ATOM   1968  HD3 PRO A 219      82.344  19.967  47.936  1.00  0.00           H
    ATOM   1969  N   TYR A 220      85.699  24.207  49.306  1.00 26.69           N
    ATOM   1970  CA  TYR A 220      86.006  25.624  49.150  1.00 26.29           C
    ATOM   1971  C   TYR A 220      86.651  25.859  47.774  1.00 25.32           C
    ATOM   1972  O   TYR A 220      87.637  25.198  47.445  1.00 22.08           O
    ATOM   1973  CB  TYR A 220      86.901  26.123  50.306  1.00 18.07           C
    ATOM   1974  CG  TYR A 220      87.477  27.503  50.064  1.00 18.56           C
    ATOM   1975  CD1 TYR A 220      86.701  28.656  50.298  1.00 14.82           C
    ATOM   1976  CD2 TYR A 220      88.760  27.619  49.492  1.00 23.52           C
    ATOM   1977  CE1 TYR A 220      87.205  29.922  49.944  1.00 13.68           C
    ATOM   1978  CE2 TYR A 220      89.247  28.877  49.111  1.00 23.61           C
    ATOM   1979  CZ  TYR A 220      88.477  30.030  49.346  1.00 22.58           C
    ATOM   1980  OH  TYR A 220      88.972  31.248  48.987  1.00 25.76           O
    ATOM   1981  H   TYR A 220      86.321  23.675  49.903  1.00  0.00           H
    ATOM   1982  HA  TYR A 220      85.069  26.180  49.204  1.00  0.00           H
    ATOM   1983  HB2 TYR A 220      86.375  26.102  51.258  1.00  0.00           H
    ATOM   1984  HB3 TYR A 220      87.751  25.454  50.420  1.00  0.00           H
    ATOM   1985  HD1 TYR A 220      85.714  28.571  50.725  1.00  0.00           H
    ATOM   1986  HD2 TYR A 220      89.352  26.738  49.291  1.00  0.00           H
    ATOM   1987  HE1 TYR A 220      86.606  30.803  50.118  1.00  0.00           H
    ATOM   1988  HE2 TYR A 220      90.215  28.957  48.638  1.00  0.00           H
    ATOM   1989  HH  TYR A 220      88.343  31.968  49.123  1.00  0.00           H
    ATOM   1990  N   GLU A 221      86.091  26.806  47.020  1.00 29.77           N
    ATOM   1991  CA  GLU A 221      86.625  27.239  45.735  1.00 34.77           C
    ATOM   1992  C   GLU A 221      87.165  28.659  45.906  1.00 30.43           C
    ATOM   1993  O   GLU A 221      86.516  29.452  46.580  1.00 29.45           O
    ATOM   1994  CB  GLU A 221      85.484  27.217  44.706  1.00 40.74           C
    ATOM   1995  CG  GLU A 221      85.122  25.778  44.295  1.00 48.30           C
    ATOM   1996  CD  GLU A 221      83.849  25.660  43.457  1.00 56.08           C
    ATOM   1997  OE1 GLU A 221      83.144  26.679  43.277  1.00 59.48           O
    ATOM   1998  OE2 GLU A 221      83.567  24.507  43.060  1.00 59.93           O1-
    ATOM   1999  H   GLU A 221      85.257  27.282  47.341  1.00  0.00           H
    ATOM   2000  HA  GLU A 221      87.397  26.555  45.387  1.00  0.00           H
    ATOM   2001  HB2 GLU A 221      84.613  27.730  45.118  1.00  0.00           H
    ATOM   2002  HB3 GLU A 221      85.772  27.778  43.815  1.00  0.00           H
    ATOM   2003  HG2 GLU A 221      85.948  25.332  43.739  1.00  0.00           H
    ATOM   2004  HG3 GLU A 221      84.980  25.161  45.182  1.00  0.00           H
    ATOM   2005  N   PRO A 222      88.320  28.985  45.287  1.00 28.78           N
    ATOM   2006  CA  PRO A 222      88.757  30.384  45.160  1.00 29.61           C
    ATOM   2007  C   PRO A 222      87.721  31.281  44.452  1.00 29.64           C
    ATOM   2008  O   PRO A 222      86.884  30.763  43.706  1.00 29.06           O
    ATOM   2009  CB  PRO A 222      90.047  30.287  44.330  1.00 31.66           C
    ATOM   2010  CG  PRO A 222      90.578  28.894  44.603  1.00 32.27           C
    ATOM   2011  CD  PRO A 222      89.319  28.055  44.766  1.00 29.78           C
    ATOM   2012  HA  PRO A 222      88.984  30.715  46.172  1.00  0.00           H
    ATOM   2013  HB2 PRO A 222      89.829  30.372  43.263  1.00  0.00           H
    ATOM   2014  HB3 PRO A 222      90.772  31.063  44.578  1.00  0.00           H
    ATOM   2015  HG2 PRO A 222      91.264  28.521  43.842  1.00  0.00           H
    ATOM   2016  HG3 PRO A 222      91.114  28.910  45.554  1.00  0.00           H
    ATOM   2017  HD2 PRO A 222      88.984  27.670  43.802  1.00  0.00           H
    ATOM   2018  HD3 PRO A 222      89.507  27.212  45.432  1.00  0.00           H
    ATOM   2019  N   PRO A 223      87.814  32.611  44.661  1.00 33.60           N
    ATOM   2020  CA  PRO A 223      86.967  33.566  43.935  1.00 40.88           C
    ATOM   2021  C   PRO A 223      87.176  33.483  42.421  1.00 43.08           C
    ATOM   2022  O   PRO A 223      88.316  33.509  41.959  1.00 42.29           O
    ATOM   2023  CB  PRO A 223      87.397  34.934  44.476  1.00 38.03           C
    ATOM   2024  CG  PRO A 223      88.047  34.649  45.813  1.00 37.03           C
    ATOM   2025  CD  PRO A 223      88.707  33.296  45.596  1.00 34.62           C
    ATOM   2026  HA  PRO A 223      85.926  33.372  44.204  1.00  0.00           H
    ATOM   2027  HB2 PRO A 223      88.146  35.389  43.827  1.00  0.00           H
    ATOM   2028  HB3 PRO A 223      86.572  35.639  44.541  1.00  0.00           H
    ATOM   2029  HG2 PRO A 223      88.740  35.428  46.129  1.00  0.00           H
    ATOM   2030  HG3 PRO A 223      87.275  34.554  46.576  1.00  0.00           H
    ATOM   2031  HD2 PRO A 223      89.688  33.406  45.133  1.00  0.00           H
    ATOM   2032  HD3 PRO A 223      88.824  32.794  46.555  1.00  0.00           H
    ATOM   2033  N   GLU A 224      86.054  33.357  41.707  1.00 48.02           N
    ATOM   2034  CA  GLU A 224      85.922  33.280  40.258  1.00 52.40           C
    ATOM   2035  C   GLU A 224      86.661  34.454  39.592  1.00 52.99           C
    ATOM   2036  O   GLU A 224      86.105  35.540  39.618  1.00 50.52           O
    ATOM   2037  CB  GLU A 224      84.402  33.322  39.908  1.00 59.06           C
    ATOM   2038  CG  GLU A 224      83.439  32.471  40.778  1.00 67.38           C
    ATOM   2039  CD  GLU A 224      83.018  33.047  42.145  1.00 69.53           C
    ATOM   2040  OE1 GLU A 224      83.203  34.263  42.368  1.00 70.00           O
    ATOM   2041  OE2 GLU A 224      82.499  32.247  42.954  1.00 65.07           O1-
    ATOM   2042  H   GLU A 224      85.173  33.370  42.204  1.00  0.00           H
    ATOM   2043  HA  GLU A 224      86.349  32.333  39.927  1.00  0.00           H
    ATOM   2044  HB2 GLU A 224      84.016  34.342  39.900  1.00  0.00           H
    ATOM   2045  HB3 GLU A 224      84.297  32.983  38.877  1.00  0.00           H
    ATOM   2046  HG2 GLU A 224      82.517  32.315  40.218  1.00  0.00           H
    ATOM   2047  HG3 GLU A 224      83.867  31.484  40.922  1.00  0.00           H
    ATOM   2048  N   VAL A 225      87.886  34.255  39.069  1.00 57.24           N
    ATOM   2049  CA  VAL A 225      88.814  35.293  38.565  1.00 63.46           C
    ATOM   2050  C   VAL A 225      88.180  36.597  38.023  1.00 59.32           C
    ATOM   2051  O   VAL A 225      87.224  36.554  37.247  1.00 58.49           O
    ATOM   2052  CB  VAL A 225      89.777  34.738  37.480  1.00 69.02           C
    ATOM   2053  CG1 VAL A 225      90.969  34.003  38.106  1.00 70.00           C
    ATOM   2054  CG2 VAL A 225      89.073  33.845  36.440  1.00 70.00           C
    ATOM   2055  H   VAL A 225      88.268  33.321  39.095  1.00  0.00           H
    ATOM   2056  HA  VAL A 225      89.409  35.569  39.437  1.00  0.00           H
    ATOM   2057  HB  VAL A 225      90.208  35.577  36.929  1.00  0.00           H
    ATOM   2058 HG11 VAL A 225      91.657  33.644  37.340  1.00  0.00           H
    ATOM   2059 HG12 VAL A 225      91.536  34.647  38.775  1.00  0.00           H
    ATOM   2060 HG13 VAL A 225      90.640  33.139  38.685  1.00  0.00           H
    ATOM   2061 HG21 VAL A 225      89.767  33.545  35.655  1.00  0.00           H
    ATOM   2062 HG22 VAL A 225      88.676  32.933  36.886  1.00  0.00           H
    ATOM   2063 HG23 VAL A 225      88.247  34.368  35.958  1.00  0.00           H
    ATOM   2064  N   GLY A 226      88.676  37.735  38.530  1.00 52.46           N
    ATOM   2065  CA  GLY A 226      87.979  39.015  38.420  1.00 49.44           C
    ATOM   2066  C   GLY A 226      86.914  39.154  39.523  1.00 52.56           C
    ATOM   2067  O   GLY A 226      86.148  40.116  39.489  1.00 53.19           O
    ATOM   2068  H   GLY A 226      89.478  37.699  39.144  1.00  0.00           H
    ATOM   2069  HA2 GLY A 226      88.708  39.817  38.533  1.00  0.00           H
    ATOM   2070  HA3 GLY A 226      87.518  39.138  37.438  1.00  0.00           H
    ATOM   2071  N   SER A 227      86.871  38.217  40.491  1.00 50.81           N
    ATOM   2072  CA  SER A 227      85.992  38.270  41.651  1.00 47.67           C
    ATOM   2073  C   SER A 227      86.812  38.370  42.933  1.00 40.97           C
    ATOM   2074  O   SER A 227      87.950  37.911  43.049  1.00 35.97           O
    ATOM   2075  CB  SER A 227      85.008  37.084  41.694  1.00 53.09           C
    ATOM   2076  OG  SER A 227      83.967  37.280  42.623  1.00 57.19           O
    ATOM   2077  H   SER A 227      87.501  37.429  40.454  1.00  0.00           H
    ATOM   2078  HA  SER A 227      85.378  39.172  41.585  1.00  0.00           H
    ATOM   2079  HB2 SER A 227      84.509  36.972  40.732  1.00  0.00           H
    ATOM   2080  HB3 SER A 227      85.515  36.153  41.946  1.00  0.00           H
    ATOM   2081  HG  SER A 227      83.509  36.440  42.736  1.00  0.00           H
    ATOM   2082  N   ASP A 228      86.134  39.024  43.862  1.00 37.21           N
    ATOM   2083  CA  ASP A 228      86.557  39.472  45.166  1.00 40.14           C
    ATOM   2084  C   ASP A 228      86.347  38.394  46.244  1.00 34.99           C
    ATOM   2085  O   ASP A 228      87.074  38.372  47.237  1.00 33.57           O
    ATOM   2086  CB  ASP A 228      85.784  40.770  45.495  1.00 52.74           C
    ATOM   2087  CG  ASP A 228      84.323  40.817  44.992  1.00 60.60           C
    ATOM   2088  OD1 ASP A 228      83.448  40.267  45.689  1.00 63.12           O
    ATOM   2089  OD2 ASP A 228      84.113  41.377  43.895  1.00 60.06           O1-
    ATOM   2090  H   ASP A 228      85.197  39.317  43.617  1.00  0.00           H
    ATOM   2091  HA  ASP A 228      87.624  39.700  45.127  1.00  0.00           H
    ATOM   2092  HB2 ASP A 228      85.844  40.997  46.547  1.00  0.00           H
    ATOM   2093  HB3 ASP A 228      86.305  41.612  45.039  1.00  0.00           H
    ATOM   2094  N   CYS A 229      85.362  37.515  46.031  1.00 29.64           N
    ATOM   2095  CA  CYS A 229      84.899  36.547  47.015  1.00 32.57           C
    ATOM   2096  C   CYS A 229      84.283  35.332  46.318  1.00 32.33           C
    ATOM   2097  O   CYS A 229      83.992  35.381  45.124  1.00 34.22           O
    ATOM   2098  CB  CYS A 229      83.924  37.229  47.985  1.00 36.50           C
    ATOM   2099  SG  CYS A 229      82.477  37.853  47.103  1.00 44.56           S
    ATOM   2100  H   CYS A 229      84.818  37.565  45.179  1.00  0.00           H
    ATOM   2101  HA  CYS A 229      85.752  36.178  47.585  1.00  0.00           H
    ATOM   2102  HB2 CYS A 229      83.553  36.555  48.749  1.00  0.00           H
    ATOM   2103  HB3 CYS A 229      84.428  38.048  48.489  1.00  0.00           H
    ATOM   2104  HG  CYS A 229      83.095  38.871  46.480  1.00  0.00           H
    ATOM   2105  N   THR A 230      84.094  34.260  47.093  1.00 26.00           N
    ATOM   2106  CA  THR A 230      83.457  33.037  46.617  1.00 15.36           C
    ATOM   2107  C   THR A 230      81.998  33.045  47.042  1.00 15.00           C
    ATOM   2108  O   THR A 230      81.739  33.310  48.208  1.00 28.80           O
    ATOM   2109  CB  THR A 230      84.074  31.789  47.273  1.00 17.05           C
    ATOM   2110  OG1 THR A 230      85.461  31.841  47.042  1.00 15.87           O
    ATOM   2111  CG2 THR A 230      83.495  30.451  46.776  1.00 13.77           C
    ATOM   2112  H   THR A 230      84.345  34.288  48.072  1.00  0.00           H
    ATOM   2113  HA  THR A 230      83.532  32.942  45.531  1.00  0.00           H
    ATOM   2114  HB  THR A 230      83.946  31.827  48.351  1.00  0.00           H
    ATOM   2115 HG21 THR A 230      84.022  29.617  47.236  1.00  0.00           H
    ATOM   2116 HG22 THR A 230      82.443  30.329  47.031  1.00  0.00           H
    ATOM   2117 HG23 THR A 230      83.595  30.351  45.696  1.00  0.00           H
    ATOM   2118  HG1 THR A 230      85.798  30.938  46.973  1.00  0.00           H
    ATOM   2119  N   THR A 231      81.085  32.717  46.126  1.00 17.31           N
    ATOM   2120  CA  THR A 231      79.669  32.657  46.457  1.00 20.82           C
    ATOM   2121  C   THR A 231      79.206  31.195  46.557  1.00 21.52           C
    ATOM   2122  O   THR A 231      79.584  30.366  45.728  1.00 27.20           O
    ATOM   2123  CB  THR A 231      78.823  33.403  45.403  1.00 19.48           C
    ATOM   2124  OG1 THR A 231      79.224  34.760  45.376  1.00 23.99           O
    ATOM   2125  CG2 THR A 231      77.317  33.380  45.695  1.00 18.23           C
    ATOM   2126  H   THR A 231      81.355  32.484  45.179  1.00  0.00           H
    ATOM   2127  HA  THR A 231      79.466  33.141  47.414  1.00  0.00           H
    ATOM   2128  HB  THR A 231      79.004  32.985  44.412  1.00  0.00           H
    ATOM   2129 HG21 THR A 231      76.781  34.040  45.014  1.00  0.00           H
    ATOM   2130 HG22 THR A 231      76.908  32.379  45.576  1.00  0.00           H
    ATOM   2131 HG23 THR A 231      77.102  33.712  46.712  1.00  0.00           H
    ATOM   2132  HG1 THR A 231      80.145  34.799  45.106  1.00  0.00           H
    ATOM   2133  N   ILE A 232      78.379  30.927  47.575  1.00 16.03           N
    ATOM   2134  CA  ILE A 232      77.654  29.674  47.747  1.00 15.03           C
    ATOM   2135  C   ILE A 232      76.165  30.019  47.783  1.00 15.75           C
    ATOM   2136  O   ILE A 232      75.779  30.948  48.486  1.00 14.57           O
    ATOM   2137  CB  ILE A 232      78.020  28.945  49.072  1.00 17.65           C
    ATOM   2138  CG1 ILE A 232      79.499  28.499  49.053  1.00 14.99           C
    ATOM   2139  CG2 ILE A 232      77.102  27.727  49.340  1.00 17.90           C
    ATOM   2140  CD1 ILE A 232      80.039  28.035  50.413  1.00 15.44           C
    ATOM   2141  H   ILE A 232      78.160  31.653  48.246  1.00  0.00           H
    ATOM   2142  HA  ILE A 232      77.838  29.000  46.908  1.00  0.00           H
    ATOM   2143  HB  ILE A 232      77.898  29.646  49.901  1.00  0.00           H
    ATOM   2144 HG12 ILE A 232      79.635  27.705  48.317  1.00  0.00           H
    ATOM   2145 HG13 ILE A 232      80.134  29.319  48.720  1.00  0.00           H
    ATOM   2146 HG21 ILE A 232      77.424  27.138  50.195  1.00  0.00           H
    ATOM   2147 HG22 ILE A 232      76.072  28.012  49.552  1.00  0.00           H
    ATOM   2148 HG23 ILE A 232      77.090  27.055  48.481  1.00  0.00           H
    ATOM   2149 HD11 ILE A 232      81.069  27.699  50.312  1.00  0.00           H
    ATOM   2150 HD12 ILE A 232      80.005  28.838  51.149  1.00  0.00           H
    ATOM   2151 HD13 ILE A 232      79.488  27.188  50.817  1.00  0.00           H
    ATOM   2152  N   HIS A 233      75.356  29.254  47.043  1.00 14.42           N
    ATOM   2153  CA  HIS A 233      73.911  29.433  47.015  1.00 13.02           C
    ATOM   2154  C   HIS A 233      73.266  28.417  47.946  1.00 17.95           C
    ATOM   2155  O   HIS A 233      73.214  27.243  47.606  1.00 18.53           O
    ATOM   2156  CB  HIS A 233      73.404  29.247  45.583  1.00 19.17           C
    ATOM   2157  CG  HIS A 233      73.753  30.336  44.610  1.00 28.41           C
    ATOM   2158  ND1 HIS A 233      74.533  31.440  44.909  1.00 33.35           N
    ATOM   2159  CD2 HIS A 233      73.408  30.469  43.287  1.00 30.26           C
    ATOM   2160  CE1 HIS A 233      74.642  32.152  43.786  1.00 31.10           C
    ATOM   2161  NE2 HIS A 233      73.981  31.625  42.764  1.00 28.37           N
    ATOM   2162  H   HIS A 233      75.733  28.490  46.498  1.00  0.00           H
    ATOM   2163  HA  HIS A 233      73.617  30.432  47.342  1.00  0.00           H
    ATOM   2164  HB2 HIS A 233      73.808  28.323  45.169  1.00  0.00           H
    ATOM   2165  HB3 HIS A 233      72.318  29.143  45.583  1.00  0.00           H
    ATOM   2166  HD1 HIS A 233      74.964  31.650  45.798  1.00  0.00           H
    ATOM   2167  HD2 HIS A 233      72.803  29.819  42.671  1.00  0.00           H
    ATOM   2168  HE1 HIS A 233      75.224  33.055  43.701  1.00  0.00           H
    ATOM   2169  N   TYR A 234      72.789  28.891  49.099  1.00 14.43           N
    ATOM   2170  CA  TYR A 234      72.036  28.087  50.048  1.00 13.19           C
    ATOM   2171  C   TYR A 234      70.535  28.184  49.757  1.00 13.50           C
    ATOM   2172  O   TYR A 234      70.082  29.158  49.158  1.00 21.12           O
    ATOM   2173  CB  TYR A 234      72.353  28.551  51.480  1.00 15.55           C
    ATOM   2174  CG  TYR A 234      73.775  28.253  51.918  1.00 15.56           C
    ATOM   2175  CD1 TYR A 234      74.148  26.933  52.244  1.00 14.79           C
    ATOM   2176  CD2 TYR A 234      74.729  29.285  51.999  1.00 12.22           C
    ATOM   2177  CE1 TYR A 234      75.465  26.648  52.649  1.00 11.09           C
    ATOM   2178  CE2 TYR A 234      76.046  29.001  52.410  1.00  9.11           C
    ATOM   2179  CZ  TYR A 234      76.417  27.682  52.726  1.00 12.63           C
    ATOM   2180  OH  TYR A 234      77.704  27.413  53.084  1.00 14.54           O
    ATOM   2181  H   TYR A 234      72.864  29.880  49.307  1.00  0.00           H
    ATOM   2182  HA  TYR A 234      72.353  27.055  49.946  1.00  0.00           H
    ATOM   2183  HB2 TYR A 234      72.151  29.616  51.592  1.00  0.00           H
    ATOM   2184  HB3 TYR A 234      71.691  28.048  52.187  1.00  0.00           H
    ATOM   2185  HD1 TYR A 234      73.430  26.132  52.174  1.00  0.00           H
    ATOM   2186  HD2 TYR A 234      74.453  30.295  51.738  1.00  0.00           H
    ATOM   2187  HE1 TYR A 234      75.740  25.633  52.890  1.00  0.00           H
    ATOM   2188  HE2 TYR A 234      76.784  29.786  52.468  1.00  0.00           H
    ATOM   2189  HH  TYR A 234      77.886  26.467  53.151  1.00  0.00           H
    ATOM   2190  N   ASN A 235      69.788  27.170  50.201  1.00 12.77           N
    ATOM   2191  CA  ASN A 235      68.327  27.164  50.195  1.00 15.78           C
    ATOM   2192  C   ASN A 235      67.883  26.754  51.595  1.00 17.99           C
    ATOM   2193  O   ASN A 235      68.472  25.832  52.152  1.00 15.50           O
    ATOM   2194  CB  ASN A 235      67.719  26.135  49.207  1.00 21.08           C
    ATOM   2195  CG  ASN A 235      68.320  26.081  47.804  1.00 25.90           C
    ATOM   2196  OD1 ASN A 235      68.543  24.993  47.273  1.00 39.89           O
    ATOM   2197  ND2 ASN A 235      68.552  27.241  47.196  1.00 24.35           N
    ATOM   2198  H   ASN A 235      70.234  26.384  50.658  1.00  0.00           H
    ATOM   2199  HA  ASN A 235      67.898  28.152  50.015  1.00  0.00           H
    ATOM   2200  HB2 ASN A 235      67.812  25.129  49.615  1.00  0.00           H
    ATOM   2201  HB3 ASN A 235      66.647  26.311  49.112  1.00  0.00           H
    ATOM   2202 HD21 ASN A 235      68.354  28.112  47.668  1.00  0.00           H
    ATOM   2203 HD22 ASN A 235      68.899  27.276  46.241  1.00  0.00           H
    ATOM   2204  N   TYR A 236      66.827  27.390  52.110  1.00 17.76           N
    ATOM   2205  CA  TYR A 236      66.206  27.024  53.375  1.00 16.45           C
    ATOM   2206  C   TYR A 236      64.837  26.436  53.050  1.00 18.73           C
    ATOM   2207  O   TYR A 236      64.039  27.106  52.400  1.00 13.46           O
    ATOM   2208  CB  TYR A 236      66.107  28.248  54.289  1.00 11.33           C
    ATOM   2209  CG  TYR A 236      67.458  28.752  54.752  1.00  8.51           C
    ATOM   2210  CD1 TYR A 236      68.261  29.517  53.885  1.00 10.76           C
    ATOM   2211  CD2 TYR A 236      67.939  28.416  56.030  1.00  8.00           C
    ATOM   2212  CE1 TYR A 236      69.559  29.889  54.274  1.00 14.26           C
    ATOM   2213  CE2 TYR A 236      69.229  28.810  56.432  1.00 12.75           C
    ATOM   2214  CZ  TYR A 236      70.039  29.550  55.551  1.00 16.14           C
    ATOM   2215  OH  TYR A 236      71.281  29.957  55.932  1.00 18.48           O
    ATOM   2216  H   TYR A 236      66.378  28.139  51.596  1.00  0.00           H
    ATOM   2217  HA  TYR A 236      66.805  26.285  53.901  1.00  0.00           H
    ATOM   2218  HB2 TYR A 236      65.555  29.059  53.811  1.00  0.00           H
    ATOM   2219  HB3 TYR A 236      65.543  27.949  55.167  1.00  0.00           H
    ATOM   2220  HD1 TYR A 236      67.900  29.788  52.904  1.00  0.00           H
    ATOM   2221  HD2 TYR A 236      67.305  27.878  56.711  1.00  0.00           H
    ATOM   2222  HE1 TYR A 236      70.185  30.440  53.587  1.00  0.00           H
    ATOM   2223  HE2 TYR A 236      69.585  28.552  57.419  1.00  0.00           H
    ATOM   2224  HH  TYR A 236      71.727  30.463  55.249  1.00  0.00           H
    ATOM   2225  N   MET A 237      64.619  25.184  53.470  1.00 20.25           N
    ATOM   2226  CA  MET A 237      63.533  24.320  53.009  1.00 18.60           C
    ATOM   2227  C   MET A 237      62.344  24.200  53.967  1.00 19.09           C
    ATOM   2228  O   MET A 237      61.550  23.268  53.830  1.00 19.03           O
    ATOM   2229  CB  MET A 237      64.119  22.941  52.673  1.00 14.58           C
    ATOM   2230  CG  MET A 237      65.190  22.988  51.581  1.00 14.34           C
    ATOM   2231  SD  MET A 237      64.704  23.762  50.009  1.00 16.55           S
    ATOM   2232  CE  MET A 237      63.133  22.917  49.668  1.00  6.68           C
    ATOM   2233  H   MET A 237      65.344  24.723  54.008  1.00  0.00           H
    ATOM   2234  HA  MET A 237      63.123  24.723  52.088  1.00  0.00           H
    ATOM   2235  HB2 MET A 237      64.549  22.493  53.571  1.00  0.00           H
    ATOM   2236  HB3 MET A 237      63.342  22.255  52.347  1.00  0.00           H
    ATOM   2237  HG2 MET A 237      66.084  23.495  51.942  1.00  0.00           H
    ATOM   2238  HG3 MET A 237      65.487  21.966  51.373  1.00  0.00           H
    ATOM   2239  HE1 MET A 237      62.781  23.177  48.670  1.00  0.00           H
    ATOM   2240  HE2 MET A 237      63.263  21.836  49.712  1.00  0.00           H
    ATOM   2241  HE3 MET A 237      62.363  23.206  50.382  1.00  0.00           H
    ATOM   2242  N   CYS A 238      62.216  25.135  54.909  1.00 13.05           N
    ATOM   2243  CA  CYS A 238      61.184  25.085  55.934  1.00 15.10           C
    ATOM   2244  C   CYS A 238      61.018  26.510  56.449  1.00 13.76           C
    ATOM   2245  O   CYS A 238      62.024  27.200  56.604  1.00 16.12           O
    ATOM   2246  CB  CYS A 238      61.583  24.097  57.052  1.00 20.24           C
    ATOM   2247  SG  CYS A 238      60.226  23.326  57.985  1.00 24.56           S
    ATOM   2248  H   CYS A 238      62.873  25.901  54.970  1.00  0.00           H
    ATOM   2249  HA  CYS A 238      60.246  24.772  55.470  1.00  0.00           H
    ATOM   2250  HB2 CYS A 238      62.153  23.288  56.610  1.00  0.00           H
    ATOM   2251  HB3 CYS A 238      62.296  24.554  57.738  1.00  0.00           H
    ATOM   2252  N   ASN A 239      59.776  26.943  56.707  1.00 17.54           N
    ATOM   2253  CA  ASN A 239      59.535  28.302  57.196  1.00 20.67           C
    ATOM   2254  C   ASN A 239      59.765  28.384  58.700  1.00 21.15           C
    ATOM   2255  O   ASN A 239      59.524  27.411  59.417  1.00 20.54           O
    ATOM   2256  CB  ASN A 239      58.109  28.772  56.881  1.00 27.36           C
    ATOM   2257  CG  ASN A 239      57.813  28.835  55.388  1.00 30.54           C
    ATOM   2258  OD1 ASN A 239      58.543  29.462  54.624  1.00 32.96           O
    ATOM   2259  ND2 ASN A 239      56.718  28.197  54.978  1.00 33.84           N
    ATOM   2260  H   ASN A 239      58.975  26.337  56.605  1.00  0.00           H
    ATOM   2261  HA  ASN A 239      60.216  28.986  56.692  1.00  0.00           H
    ATOM   2262  HB2 ASN A 239      57.379  28.133  57.382  1.00  0.00           H
    ATOM   2263  HB3 ASN A 239      57.951  29.775  57.274  1.00  0.00           H
    ATOM   2264 HD21 ASN A 239      56.138  27.699  55.638  1.00  0.00           H
    ATOM   2265 HD22 ASN A 239      56.458  28.213  54.002  1.00  0.00           H
    ATOM   2266  N   SER A 240      60.164  29.580  59.156  1.00 17.90           N
    ATOM   2267  CA  SER A 240      60.275  29.928  60.570  1.00 20.76           C
    ATOM   2268  C   SER A 240      58.962  29.695  61.314  1.00 25.92           C
    ATOM   2269  O   SER A 240      59.002  29.193  62.423  1.00 27.38           O
    ATOM   2270  CB  SER A 240      60.689  31.396  60.717  1.00 17.73           C
    ATOM   2271  OG  SER A 240      62.023  31.547  60.300  1.00 22.57           O
    ATOM   2272  H   SER A 240      60.350  30.329  58.500  1.00  0.00           H
    ATOM   2273  HA  SER A 240      61.038  29.297  61.027  1.00  0.00           H
    ATOM   2274  HB2 SER A 240      60.048  32.059  60.134  1.00  0.00           H
    ATOM   2275  HB3 SER A 240      60.626  31.719  61.757  1.00  0.00           H
    ATOM   2276  HG  SER A 240      62.595  31.196  60.996  1.00  0.00           H
    ATOM   2277  N   SER A 241      57.836  30.005  60.658  1.00 24.22           N
    ATOM   2278  CA  SER A 241      56.490  29.845  61.186  1.00 22.93           C
    ATOM   2279  C   SER A 241      55.941  28.414  61.187  1.00 25.22           C
    ATOM   2280  O   SER A 241      54.857  28.239  61.744  1.00 23.88           O
    ATOM   2281  CB  SER A 241      55.550  30.796  60.431  1.00 27.33           C
    ATOM   2282  OG  SER A 241      55.346  30.372  59.097  1.00 29.72           O
    ATOM   2283  H   SER A 241      57.901  30.417  59.737  1.00  0.00           H
    ATOM   2284  HA  SER A 241      56.512  30.168  62.223  1.00  0.00           H
    ATOM   2285  HB2 SER A 241      54.579  30.843  60.926  1.00  0.00           H
    ATOM   2286  HB3 SER A 241      55.943  31.813  60.434  1.00  0.00           H
    ATOM   2287  HG  SER A 241      56.114  30.624  58.578  1.00  0.00           H
    ATOM   2288  N   CYS A 242      56.644  27.436  60.580  1.00 31.28           N
    ATOM   2289  CA  CYS A 242      56.216  26.037  60.571  1.00 28.87           C
    ATOM   2290  C   CYS A 242      56.117  25.523  62.009  1.00 27.97           C
    ATOM   2291  O   CYS A 242      57.144  25.276  62.641  1.00 29.15           O
    ATOM   2292  CB  CYS A 242      57.129  25.115  59.740  1.00 25.93           C
    ATOM   2293  SG  CYS A 242      56.916  25.203  57.926  1.00 28.16           S
    ATOM   2294  H   CYS A 242      57.528  27.638  60.133  1.00  0.00           H
    ATOM   2295  HA  CYS A 242      55.220  25.999  60.125  1.00  0.00           H
    ATOM   2296  HB2 CYS A 242      58.167  25.275  60.023  1.00  0.00           H
    ATOM   2297  HB3 CYS A 242      56.924  24.080  60.016  1.00  0.00           H
    ATOM   2298  N   MET A 243      54.873  25.376  62.491  1.00 27.94           N
    ATOM   2299  CA  MET A 243      54.586  24.725  63.763  1.00 39.01           C
    ATOM   2300  C   MET A 243      55.017  23.262  63.656  1.00 42.25           C
    ATOM   2301  O   MET A 243      54.791  22.643  62.620  1.00 49.18           O
    ATOM   2302  CB  MET A 243      53.086  24.821  64.084  1.00 51.13           C
    ATOM   2303  CG  MET A 243      52.632  26.227  64.491  1.00 60.72           C
    ATOM   2304  SD  MET A 243      50.828  26.385  64.572  1.00 67.06           S
    ATOM   2305  CE  MET A 243      50.619  27.046  66.245  1.00 64.40           C
    ATOM   2306  H   MET A 243      54.075  25.626  61.920  1.00  0.00           H
    ATOM   2307  HA  MET A 243      55.158  25.213  64.553  1.00  0.00           H
    ATOM   2308  HB2 MET A 243      52.498  24.475  63.233  1.00  0.00           H
    ATOM   2309  HB3 MET A 243      52.846  24.144  64.906  1.00  0.00           H
    ATOM   2310  HG2 MET A 243      53.061  26.486  65.458  1.00  0.00           H
    ATOM   2311  HG3 MET A 243      52.997  26.968  63.780  1.00  0.00           H
    ATOM   2312  HE1 MET A 243      49.561  27.202  66.460  1.00  0.00           H
    ATOM   2313  HE2 MET A 243      51.022  26.352  66.984  1.00  0.00           H
    ATOM   2314  HE3 MET A 243      51.135  28.001  66.345  1.00  0.00           H
    ATOM   2315  N   GLY A 244      55.679  22.741  64.690  1.00 40.79           N
    ATOM   2316  CA  GLY A 244      56.260  21.399  64.622  1.00 42.18           C
    ATOM   2317  C   GLY A 244      57.642  21.398  63.948  1.00 38.16           C
    ATOM   2318  O   GLY A 244      58.255  20.338  63.851  1.00 36.82           O
    ATOM   2319  H   GLY A 244      55.854  23.288  65.521  1.00  0.00           H
    ATOM   2320  HA2 GLY A 244      56.376  21.026  65.640  1.00  0.00           H
    ATOM   2321  HA3 GLY A 244      55.600  20.697  64.108  1.00  0.00           H
    ATOM   2322  N   GLY A 245      58.132  22.569  63.517  1.00 34.21           N
    ATOM   2323  CA  GLY A 245      59.479  22.799  63.030  1.00 33.46           C
    ATOM   2324  C   GLY A 245      60.088  23.844  63.957  1.00 36.68           C
    ATOM   2325  O   GLY A 245      60.010  23.731  65.182  1.00 37.30           O
    ATOM   2326  H   GLY A 245      57.550  23.393  63.594  1.00  0.00           H
    ATOM   2327  HA2 GLY A 245      60.092  21.902  63.026  1.00  0.00           H
    ATOM   2328  HA3 GLY A 245      59.429  23.165  62.006  1.00  0.00           H
    ATOM   2329  N   MET A 246      60.686  24.873  63.346  1.00 34.05           N
    ATOM   2330  CA  MET A 246      61.251  26.043  64.008  1.00 29.62           C
    ATOM   2331  C   MET A 246      60.244  26.811  64.871  1.00 30.76           C
    ATOM   2332  O   MET A 246      60.651  27.303  65.915  1.00 30.95           O
    ATOM   2333  CB  MET A 246      61.960  26.946  62.982  1.00 26.73           C
    ATOM   2334  CG  MET A 246      63.127  26.242  62.264  1.00 26.14           C
    ATOM   2335  SD  MET A 246      64.058  27.292  61.113  1.00 28.37           S
    ATOM   2336  CE  MET A 246      63.037  27.165  59.637  1.00 24.63           C
    ATOM   2337  H   MET A 246      60.714  24.890  62.331  1.00  0.00           H
    ATOM   2338  HA  MET A 246      62.001  25.683  64.699  1.00  0.00           H
    ATOM   2339  HB2 MET A 246      61.237  27.295  62.245  1.00  0.00           H
    ATOM   2340  HB3 MET A 246      62.341  27.839  63.481  1.00  0.00           H
    ATOM   2341  HG2 MET A 246      63.819  25.849  63.003  1.00  0.00           H
    ATOM   2342  HG3 MET A 246      62.799  25.360  61.722  1.00  0.00           H
    ATOM   2343  HE1 MET A 246      63.457  27.768  58.833  1.00  0.00           H
    ATOM   2344  HE2 MET A 246      62.967  26.129  59.310  1.00  0.00           H
    ATOM   2345  HE3 MET A 246      62.044  27.526  59.868  1.00  0.00           H
    ATOM   2346  N   ASN A 247      58.958  26.849  64.481  1.00 33.31           N
    ATOM   2347  CA  ASN A 247      57.837  27.371  65.278  1.00 32.35           C
    ATOM   2348  C   ASN A 247      58.080  28.796  65.833  1.00 34.17           C
    ATOM   2349  O   ASN A 247      57.887  29.050  67.019  1.00 34.21           O
    ATOM   2350  CB  ASN A 247      57.481  26.338  66.380  1.00 35.16           C
    ATOM   2351  CG  ASN A 247      56.173  26.615  67.134  1.00 41.59           C
    ATOM   2352  OD1 ASN A 247      55.291  27.325  66.650  1.00 48.40           O
    ATOM   2353  ND2 ASN A 247      56.052  26.054  68.338  1.00 40.39           N
    ATOM   2354  H   ASN A 247      58.697  26.404  63.611  1.00  0.00           H
    ATOM   2355  HA  ASN A 247      56.989  27.460  64.597  1.00  0.00           H
    ATOM   2356  HB2 ASN A 247      57.371  25.357  65.917  1.00  0.00           H
    ATOM   2357  HB3 ASN A 247      58.300  26.251  67.096  1.00  0.00           H
    ATOM   2358 HD21 ASN A 247      56.803  25.497  68.724  1.00  0.00           H
    ATOM   2359 HD22 ASN A 247      55.213  26.190  68.883  1.00  0.00           H
    ATOM   2360  N   ARG A 248      58.534  29.695  64.953  1.00 34.94           N
    ATOM   2361  CA  ARG A 248      58.805  31.119  65.163  1.00 35.37           C
    ATOM   2362  C   ARG A 248      60.049  31.408  66.011  1.00 36.12           C
    ATOM   2363  O   ARG A 248      60.284  32.568  66.350  1.00 33.38           O
    ATOM   2364  CB  ARG A 248      57.541  31.850  65.664  1.00 32.16           C
    ATOM   2365  CG  ARG A 248      56.396  31.774  64.638  1.00 34.71           C
    ATOM   2366  CD  ARG A 248      55.016  32.028  65.232  1.00 39.11           C
    ATOM   2367  NE  ARG A 248      54.545  30.855  65.985  1.00 45.00           N
    ATOM   2368  CZ  ARG A 248      53.967  30.864  67.195  1.00 47.22           C
    ATOM   2369  NH1 ARG A 248      53.851  31.991  67.906  1.00 49.44           N
    ATOM   2370  NH2 ARG A 248      53.504  29.717  67.707  1.00 47.16           N1+
    ATOM   2371  H   ARG A 248      58.700  29.378  64.003  1.00  0.00           H
    ATOM   2372  HA  ARG A 248      59.041  31.514  64.174  1.00  0.00           H
    ATOM   2373  HB2 ARG A 248      57.225  31.446  66.626  1.00  0.00           H
    ATOM   2374  HB3 ARG A 248      57.753  32.903  65.850  1.00  0.00           H
    ATOM   2375  HG2 ARG A 248      56.583  32.376  63.749  1.00  0.00           H
    ATOM   2376  HG3 ARG A 248      56.347  30.745  64.289  1.00  0.00           H
    ATOM   2377  HD2 ARG A 248      55.150  32.834  65.952  1.00  0.00           H
    ATOM   2378  HD3 ARG A 248      54.290  32.398  64.507  1.00  0.00           H
    ATOM   2379  HE  ARG A 248      54.607  29.970  65.500  1.00  0.00           H
    ATOM   2380 HH11 ARG A 248      54.286  32.849  67.594  1.00  0.00           H
    ATOM   2381 HH12 ARG A 248      53.307  31.995  68.762  1.00  0.00           H
    ATOM   2382 HH21 ARG A 248      53.652  28.843  67.221  1.00  0.00           H
    ATOM   2383 HH22 ARG A 248      53.085  29.706  68.627  1.00  0.00           H
    ATOM   2384  N   ARG A 249      60.846  30.371  66.308  1.00 31.02           N
    ATOM   2385  CA  ARG A 249      62.058  30.508  67.099  1.00 27.17           C
    ATOM   2386  C   ARG A 249      63.235  30.868  66.184  1.00 24.04           C
    ATOM   2387  O   ARG A 249      63.384  30.238  65.136  1.00 23.86           O
    ATOM   2388  CB  ARG A 249      62.361  29.203  67.834  1.00 28.74           C
    ATOM   2389  CG  ARG A 249      61.224  28.761  68.764  1.00 25.87           C
    ATOM   2390  CD  ARG A 249      61.491  27.388  69.367  1.00 20.38           C
    ATOM   2391  NE  ARG A 249      61.332  26.317  68.382  1.00 16.42           N
    ATOM   2392  CZ  ARG A 249      61.724  25.053  68.558  1.00 20.51           C
    ATOM   2393  NH1 ARG A 249      62.326  24.684  69.691  1.00 25.34           N
    ATOM   2394  NH2 ARG A 249      61.518  24.147  67.601  1.00 22.63           N1+
    ATOM   2395  H   ARG A 249      60.610  29.439  65.994  1.00  0.00           H
    ATOM   2396  HA  ARG A 249      61.842  31.255  67.859  1.00  0.00           H
    ATOM   2397  HB2 ARG A 249      62.582  28.419  67.112  1.00  0.00           H
    ATOM   2398  HB3 ARG A 249      63.268  29.329  68.426  1.00  0.00           H
    ATOM   2399  HG2 ARG A 249      61.010  29.494  69.542  1.00  0.00           H
    ATOM   2400  HG3 ARG A 249      60.311  28.678  68.177  1.00  0.00           H
    ATOM   2401  HD2 ARG A 249      62.551  27.353  69.624  1.00  0.00           H
    ATOM   2402  HD3 ARG A 249      60.951  27.221  70.295  1.00  0.00           H
    ATOM   2403  HE  ARG A 249      60.903  26.583  67.501  1.00  0.00           H
    ATOM   2404 HH11 ARG A 249      62.464  25.348  70.440  1.00  0.00           H
    ATOM   2405 HH12 ARG A 249      62.627  23.719  69.816  1.00  0.00           H
    ATOM   2406 HH21 ARG A 249      61.052  24.385  66.734  1.00  0.00           H
    ATOM   2407 HH22 ARG A 249      61.895  23.208  67.717  1.00  0.00           H
    ATOM   2408  N   PRO A 250      64.043  31.868  66.599  1.00 23.15           N
    ATOM   2409  CA  PRO A 250      65.215  32.316  65.840  1.00 21.46           C
    ATOM   2410  C   PRO A 250      66.296  31.248  65.658  1.00 20.96           C
    ATOM   2411  O   PRO A 250      66.635  30.556  66.615  1.00 24.02           O
    ATOM   2412  CB  PRO A 250      65.724  33.547  66.605  1.00 21.97           C
    ATOM   2413  CG  PRO A 250      65.111  33.463  67.990  1.00 19.41           C
    ATOM   2414  CD  PRO A 250      63.817  32.705  67.777  1.00 20.87           C
    ATOM   2415  HA  PRO A 250      64.853  32.608  64.862  1.00  0.00           H
    ATOM   2416  HB2 PRO A 250      66.811  33.627  66.629  1.00  0.00           H
    ATOM   2417  HB3 PRO A 250      65.356  34.443  66.106  1.00  0.00           H
    ATOM   2418  HG2 PRO A 250      65.768  32.866  68.622  1.00  0.00           H
    ATOM   2419  HG3 PRO A 250      64.977  34.433  68.471  1.00  0.00           H
    ATOM   2420  HD2 PRO A 250      63.586  32.136  68.676  1.00  0.00           H
    ATOM   2421  HD3 PRO A 250      62.994  33.393  67.573  1.00  0.00           H
    ATOM   2422  N   ILE A 251      66.790  31.139  64.418  1.00 20.28           N
    ATOM   2423  CA  ILE A 251      67.854  30.222  64.023  1.00 22.18           C
    ATOM   2424  C   ILE A 251      69.089  31.018  63.610  1.00 21.66           C
    ATOM   2425  O   ILE A 251      68.963  32.061  62.981  1.00 18.86           O
    ATOM   2426  CB  ILE A 251      67.397  29.290  62.848  1.00 21.95           C
    ATOM   2427  CG1 ILE A 251      66.923  27.941  63.408  1.00 24.47           C
    ATOM   2428  CG2 ILE A 251      68.426  28.990  61.733  1.00 26.20           C
    ATOM   2429  CD1 ILE A 251      65.554  28.011  64.057  1.00 25.71           C
    ATOM   2430  H   ILE A 251      66.457  31.759  63.694  1.00  0.00           H
    ATOM   2431  HA  ILE A 251      68.139  29.609  64.879  1.00  0.00           H
    ATOM   2432  HB  ILE A 251      66.554  29.764  62.343  1.00  0.00           H
    ATOM   2433 HG12 ILE A 251      66.853  27.211  62.603  1.00  0.00           H
    ATOM   2434 HG13 ILE A 251      67.653  27.539  64.109  1.00  0.00           H
    ATOM   2435 HG21 ILE A 251      68.000  28.333  60.975  1.00  0.00           H
    ATOM   2436 HG22 ILE A 251      68.744  29.887  61.201  1.00  0.00           H
    ATOM   2437 HG23 ILE A 251      69.312  28.491  62.127  1.00  0.00           H
    ATOM   2438 HD11 ILE A 251      65.160  27.009  64.158  1.00  0.00           H
    ATOM   2439 HD12 ILE A 251      65.615  28.461  65.045  1.00  0.00           H
    ATOM   2440 HD13 ILE A 251      64.850  28.599  63.475  1.00  0.00           H
    ATOM   2441  N   LEU A 252      70.255  30.459  63.930  1.00 20.08           N
    ATOM   2442  CA  LEU A 252      71.569  30.910  63.506  1.00 18.57           C
    ATOM   2443  C   LEU A 252      72.063  29.926  62.463  1.00 17.88           C
    ATOM   2444  O   LEU A 252      71.982  28.731  62.720  1.00 15.02           O
    ATOM   2445  CB  LEU A 252      72.506  30.842  64.732  1.00 20.22           C
    ATOM   2446  CG  LEU A 252      72.939  32.197  65.298  1.00 20.32           C
    ATOM   2447  CD1 LEU A 252      71.796  33.211  65.437  1.00 20.77           C
    ATOM   2448  CD2 LEU A 252      73.654  31.973  66.637  1.00 17.39           C
    ATOM   2449  H   LEU A 252      70.248  29.582  64.437  1.00  0.00           H
    ATOM   2450  HA  LEU A 252      71.548  31.908  63.076  1.00  0.00           H
    ATOM   2451  HB2 LEU A 252      72.028  30.269  65.529  1.00  0.00           H
    ATOM   2452  HB3 LEU A 252      73.414  30.280  64.508  1.00  0.00           H
    ATOM   2453  HG  LEU A 252      73.667  32.620  64.603  1.00  0.00           H
    ATOM   2454 HD11 LEU A 252      71.939  33.845  66.309  1.00  0.00           H
    ATOM   2455 HD12 LEU A 252      71.783  33.882  64.580  1.00  0.00           H
    ATOM   2456 HD13 LEU A 252      70.818  32.739  65.503  1.00  0.00           H
    ATOM   2457 HD21 LEU A 252      74.314  32.810  66.864  1.00  0.00           H
    ATOM   2458 HD22 LEU A 252      72.945  31.871  67.459  1.00  0.00           H
    ATOM   2459 HD23 LEU A 252      74.268  31.072  66.622  1.00  0.00           H
    ATOM   2460  N   THR A 253      72.639  30.419  61.365  1.00 18.92           N
    ATOM   2461  CA  THR A 253      73.409  29.570  60.474  1.00 11.54           C
    ATOM   2462  C   THR A 253      74.885  29.832  60.757  1.00 10.08           C
    ATOM   2463  O   THR A 253      75.341  30.965  60.635  1.00 14.15           O
    ATOM   2464  CB  THR A 253      73.110  29.810  58.985  1.00 14.89           C
    ATOM   2465  OG1 THR A 253      71.752  29.508  58.766  1.00 15.17           O
    ATOM   2466  CG2 THR A 253      73.928  28.897  58.058  1.00 11.74           C
    ATOM   2467  H   THR A 253      72.680  31.419  61.200  1.00  0.00           H
    ATOM   2468  HA  THR A 253      73.197  28.516  60.655  1.00  0.00           H
    ATOM   2469  HB  THR A 253      73.275  30.856  58.727  1.00  0.00           H
    ATOM   2470 HG21 THR A 253      73.652  29.039  57.013  1.00  0.00           H
    ATOM   2471 HG22 THR A 253      74.990  29.108  58.140  1.00  0.00           H
    ATOM   2472 HG23 THR A 253      73.782  27.844  58.303  1.00  0.00           H
    ATOM   2473  HG1 THR A 253      71.566  29.615  57.826  1.00  0.00           H
    ATOM   2474  N   ILE A 254      75.581  28.764  61.155  1.00 11.95           N
    ATOM   2475  CA  ILE A 254      76.994  28.766  61.476  1.00 17.30           C
    ATOM   2476  C   ILE A 254      77.736  28.272  60.229  1.00 20.62           C
    ATOM   2477  O   ILE A 254      77.528  27.129  59.827  1.00 16.31           O
    ATOM   2478  CB  ILE A 254      77.322  27.784  62.640  1.00 16.46           C
    ATOM   2479  CG1 ILE A 254      76.365  27.886  63.851  1.00 21.93           C
    ATOM   2480  CG2 ILE A 254      78.784  27.937  63.085  1.00 12.20           C
    ATOM   2481  CD1 ILE A 254      76.237  29.282  64.466  1.00 26.31           C
    ATOM   2482  H   ILE A 254      75.115  27.867  61.229  1.00  0.00           H
    ATOM   2483  HA  ILE A 254      77.330  29.770  61.740  1.00  0.00           H
    ATOM   2484  HB  ILE A 254      77.214  26.756  62.290  1.00  0.00           H
    ATOM   2485 HG12 ILE A 254      75.370  27.551  63.554  1.00  0.00           H
    ATOM   2486 HG13 ILE A 254      76.677  27.182  64.624  1.00  0.00           H
    ATOM   2487 HG21 ILE A 254      78.994  27.301  63.944  1.00  0.00           H
    ATOM   2488 HG22 ILE A 254      79.481  27.649  62.299  1.00  0.00           H
    ATOM   2489 HG23 ILE A 254      79.004  28.968  63.363  1.00  0.00           H
    ATOM   2490 HD11 ILE A 254      75.709  29.232  65.417  1.00  0.00           H
    ATOM   2491 HD12 ILE A 254      77.209  29.738  64.642  1.00  0.00           H
    ATOM   2492 HD13 ILE A 254      75.667  29.942  63.815  1.00  0.00           H
    ATOM   2493  N   ILE A 255      78.579  29.133  59.644  1.00 21.93           N
    ATOM   2494  CA  ILE A 255      79.417  28.780  58.505  1.00 18.14           C
    ATOM   2495  C   ILE A 255      80.842  28.565  59.020  1.00 17.80           C
    ATOM   2496  O   ILE A 255      81.430  29.491  59.571  1.00 13.33           O
    ATOM   2497  CB  ILE A 255      79.434  29.849  57.380  1.00 16.74           C
    ATOM   2498  CG1 ILE A 255      78.011  30.293  56.974  1.00  8.80           C
    ATOM   2499  CG2 ILE A 255      80.209  29.348  56.139  1.00 19.68           C
    ATOM   2500  CD1 ILE A 255      77.112  29.173  56.436  1.00  9.28           C
    ATOM   2501  H   ILE A 255      78.733  30.049  60.049  1.00  0.00           H
    ATOM   2502  HA  ILE A 255      79.049  27.865  58.055  1.00  0.00           H
    ATOM   2503  HB  ILE A 255      79.946  30.740  57.747  1.00  0.00           H
    ATOM   2504 HG12 ILE A 255      77.516  30.769  57.822  1.00  0.00           H
    ATOM   2505 HG13 ILE A 255      78.079  31.067  56.211  1.00  0.00           H
    ATOM   2506 HG21 ILE A 255      80.199  30.084  55.335  1.00  0.00           H
    ATOM   2507 HG22 ILE A 255      81.252  29.127  56.362  1.00  0.00           H
    ATOM   2508 HG23 ILE A 255      79.777  28.427  55.747  1.00  0.00           H
    ATOM   2509 HD11 ILE A 255      76.142  29.570  56.140  1.00  0.00           H
    ATOM   2510 HD12 ILE A 255      77.539  28.705  55.551  1.00  0.00           H
    ATOM   2511 HD13 ILE A 255      76.942  28.396  57.179  1.00  0.00           H
    ATOM   2512  N   THR A 256      81.344  27.339  58.846  1.00 15.94           N
    ATOM   2513  CA  THR A 256      82.661  26.921  59.295  1.00 20.91           C
    ATOM   2514  C   THR A 256      83.542  26.600  58.084  1.00 18.97           C
    ATOM   2515  O   THR A 256      83.134  25.854  57.199  1.00 22.05           O
    ATOM   2516  CB  THR A 256      82.589  25.663  60.195  1.00 18.18           C
    ATOM   2517  OG1 THR A 256      81.850  25.966  61.364  1.00 18.87           O
    ATOM   2518  CG2 THR A 256      83.962  25.122  60.631  1.00 14.69           C
    ATOM   2519  H   THR A 256      80.790  26.636  58.372  1.00  0.00           H
    ATOM   2520  HA  THR A 256      83.129  27.716  59.873  1.00  0.00           H
    ATOM   2521  HB  THR A 256      82.054  24.864  59.683  1.00  0.00           H
    ATOM   2522 HG21 THR A 256      83.903  24.537  61.547  1.00  0.00           H
    ATOM   2523 HG22 THR A 256      84.411  24.495  59.860  1.00  0.00           H
    ATOM   2524 HG23 THR A 256      84.652  25.937  60.835  1.00  0.00           H
    ATOM   2525  HG1 THR A 256      81.869  25.195  61.938  1.00  0.00           H
    ATOM   2526  N   LEU A 257      84.757  27.153  58.104  1.00 18.28           N
    ATOM   2527  CA  LEU A 257      85.837  26.835  57.191  1.00 22.65           C
    ATOM   2528  C   LEU A 257      86.679  25.750  57.869  1.00 27.08           C
    ATOM   2529  O   LEU A 257      87.116  25.964  58.997  1.00 24.85           O
    ATOM   2530  CB  LEU A 257      86.644  28.129  56.947  1.00 23.59           C
    ATOM   2531  CG  LEU A 257      87.646  28.047  55.782  1.00 24.96           C
    ATOM   2532  CD1 LEU A 257      86.911  27.825  54.449  1.00 25.15           C
    ATOM   2533  CD2 LEU A 257      88.534  29.306  55.735  1.00 24.54           C
    ATOM   2534  H   LEU A 257      85.014  27.735  58.893  1.00  0.00           H
    ATOM   2535  HA  LEU A 257      85.428  26.464  56.250  1.00  0.00           H
    ATOM   2536  HB2 LEU A 257      85.955  28.952  56.750  1.00  0.00           H
    ATOM   2537  HB3 LEU A 257      87.168  28.408  57.862  1.00  0.00           H
    ATOM   2538  HG  LEU A 257      88.305  27.194  55.945  1.00  0.00           H
    ATOM   2539 HD11 LEU A 257      87.496  28.162  53.592  1.00  0.00           H
    ATOM   2540 HD12 LEU A 257      86.687  26.771  54.298  1.00  0.00           H
    ATOM   2541 HD13 LEU A 257      85.961  28.360  54.423  1.00  0.00           H
    ATOM   2542 HD21 LEU A 257      89.589  29.036  55.795  1.00  0.00           H
    ATOM   2543 HD22 LEU A 257      88.397  29.881  54.819  1.00  0.00           H
    ATOM   2544 HD23 LEU A 257      88.332  29.980  56.567  1.00  0.00           H
    ATOM   2545  N   GLU A 258      86.875  24.610  57.200  1.00 25.30           N
    ATOM   2546  CA  GLU A 258      87.626  23.494  57.763  1.00 18.22           C
    ATOM   2547  C   GLU A 258      88.551  22.880  56.724  1.00 26.64           C
    ATOM   2548  O   GLU A 258      88.189  22.838  55.552  1.00 31.42           O
    ATOM   2549  CB  GLU A 258      86.672  22.458  58.386  1.00 15.34           C
    ATOM   2550  CG  GLU A 258      85.495  22.049  57.486  1.00 21.26           C
    ATOM   2551  CD  GLU A 258      84.651  20.994  58.173  1.00 24.29           C
    ATOM   2552  OE1 GLU A 258      84.770  19.815  57.789  1.00 21.79           O
    ATOM   2553  OE2 GLU A 258      83.892  21.377  59.081  1.00 22.95           O1-
    ATOM   2554  H   GLU A 258      86.486  24.476  56.275  1.00  0.00           H
    ATOM   2555  HA  GLU A 258      88.275  23.891  58.534  1.00  0.00           H
    ATOM   2556  HB2 GLU A 258      87.241  21.567  58.654  1.00  0.00           H
    ATOM   2557  HB3 GLU A 258      86.296  22.856  59.329  1.00  0.00           H
    ATOM   2558  HG2 GLU A 258      84.856  22.902  57.254  1.00  0.00           H
    ATOM   2559  HG3 GLU A 258      85.856  21.656  56.537  1.00  0.00           H
    ATOM   2560  N   ASP A 259      89.716  22.374  57.165  1.00 28.74           N
    ATOM   2561  CA  ASP A 259      90.654  21.677  56.280  1.00 25.67           C
    ATOM   2562  C   ASP A 259      90.045  20.334  55.822  1.00 28.50           C
    ATOM   2563  O   ASP A 259      88.981  19.935  56.302  1.00 25.80           O
    ATOM   2564  CB  ASP A 259      92.072  21.512  56.895  1.00 27.79           C
    ATOM   2565  CG  ASP A 259      92.194  20.668  58.164  1.00 34.90           C
    ATOM   2566  OD1 ASP A 259      91.228  19.958  58.509  1.00 36.66           O
    ATOM   2567  OD2 ASP A 259      93.306  20.682  58.734  1.00 37.87           O1-
    ATOM   2568  H   ASP A 259      89.940  22.395  58.151  1.00  0.00           H
    ATOM   2569  HA  ASP A 259      90.776  22.287  55.384  1.00  0.00           H
    ATOM   2570  HB2 ASP A 259      92.745  21.083  56.157  1.00  0.00           H
    ATOM   2571  HB3 ASP A 259      92.446  22.509  57.129  1.00  0.00           H
    ATOM   2572  N   SER A 260      90.747  19.626  54.932  1.00 30.76           N
    ATOM   2573  CA  SER A 260      90.344  18.298  54.465  1.00 26.84           C
    ATOM   2574  C   SER A 260      90.167  17.235  55.569  1.00 26.62           C
    ATOM   2575  O   SER A 260      89.453  16.266  55.324  1.00 29.16           O
    ATOM   2576  CB  SER A 260      91.339  17.826  53.396  1.00 26.43           C
    ATOM   2577  OG  SER A 260      92.593  17.503  53.958  1.00 33.14           O
    ATOM   2578  H   SER A 260      91.606  20.003  54.552  1.00  0.00           H
    ATOM   2579  HA  SER A 260      89.371  18.418  53.987  1.00  0.00           H
    ATOM   2580  HB2 SER A 260      90.957  16.941  52.884  1.00  0.00           H
    ATOM   2581  HB3 SER A 260      91.485  18.593  52.642  1.00  0.00           H
    ATOM   2582  HG  SER A 260      93.037  18.319  54.202  1.00  0.00           H
    ATOM   2583  N   SER A 261      90.792  17.432  56.744  1.00 27.66           N
    ATOM   2584  CA  SER A 261      90.692  16.543  57.903  1.00 30.83           C
    ATOM   2585  C   SER A 261      89.584  16.952  58.891  1.00 27.44           C
    ATOM   2586  O   SER A 261      89.396  16.255  59.887  1.00 28.42           O
    ATOM   2587  CB  SER A 261      92.056  16.485  58.613  1.00 35.69           C
    ATOM   2588  OG  SER A 261      92.990  15.791  57.812  1.00 41.36           O
    ATOM   2589  H   SER A 261      91.336  18.273  56.882  1.00  0.00           H
    ATOM   2590  HA  SER A 261      90.447  15.531  57.573  1.00  0.00           H
    ATOM   2591  HB2 SER A 261      92.430  17.484  58.838  1.00  0.00           H
    ATOM   2592  HB3 SER A 261      91.984  15.952  59.563  1.00  0.00           H
    ATOM   2593  HG  SER A 261      93.176  16.325  57.035  1.00  0.00           H
    ATOM   2594  N   GLY A 262      88.864  18.050  58.615  1.00 24.80           N
    ATOM   2595  CA  GLY A 262      87.730  18.507  59.412  1.00 24.30           C
    ATOM   2596  C   GLY A 262      88.131  19.438  60.561  1.00 26.96           C
    ATOM   2597  O   GLY A 262      87.262  19.797  61.353  1.00 28.80           O
    ATOM   2598  H   GLY A 262      89.108  18.609  57.807  1.00  0.00           H
    ATOM   2599  HA2 GLY A 262      87.059  19.062  58.762  1.00  0.00           H
    ATOM   2600  HA3 GLY A 262      87.159  17.665  59.803  1.00  0.00           H
    ATOM   2601  N   ASN A 263      89.405  19.849  60.656  1.00 23.82           N
    ATOM   2602  CA  ASN A 263      89.885  20.802  61.660  1.00 22.21           C
    ATOM   2603  C   ASN A 263      89.360  22.208  61.353  1.00 27.30           C
    ATOM   2604  O   ASN A 263      89.239  22.555  60.184  1.00 34.06           O
    ATOM   2605  CB  ASN A 263      91.427  20.842  61.707  1.00 26.41           C
    ATOM   2606  CG  ASN A 263      92.069  19.466  61.902  1.00 32.53           C
    ATOM   2607  OD1 ASN A 263      91.632  18.694  62.751  1.00 36.72           O
    ATOM   2608  ND2 ASN A 263      93.110  19.158  61.128  1.00 28.03           N
    ATOM   2609  H   ASN A 263      90.074  19.582  59.942  1.00  0.00           H
    ATOM   2610  HA  ASN A 263      89.508  20.519  62.646  1.00  0.00           H
    ATOM   2611  HB2 ASN A 263      91.827  21.319  60.815  1.00  0.00           H
    ATOM   2612  HB3 ASN A 263      91.739  21.466  62.543  1.00  0.00           H
    ATOM   2613 HD21 ASN A 263      93.387  19.786  60.379  1.00  0.00           H
    ATOM   2614 HD22 ASN A 263      93.584  18.272  61.230  1.00  0.00           H
    ATOM   2615  N   LEU A 264      89.083  22.992  62.402  1.00 24.61           N
    ATOM   2616  CA  LEU A 264      88.594  24.365  62.295  1.00 21.89           C
    ATOM   2617  C   LEU A 264      89.676  25.309  61.747  1.00 27.51           C
    ATOM   2618  O   LEU A 264      90.744  25.420  62.350  1.00 28.54           O
    ATOM   2619  CB  LEU A 264      88.145  24.823  63.697  1.00 21.02           C
    ATOM   2620  CG  LEU A 264      87.656  26.289  63.784  1.00 20.75           C
    ATOM   2621  CD1 LEU A 264      86.221  26.427  63.258  1.00 20.24           C
    ATOM   2622  CD2 LEU A 264      87.816  26.829  65.215  1.00 22.18           C
    ATOM   2623  H   LEU A 264      89.232  22.643  63.340  1.00  0.00           H
    ATOM   2624  HA  LEU A 264      87.729  24.374  61.628  1.00  0.00           H
    ATOM   2625  HB2 LEU A 264      87.384  24.149  64.092  1.00  0.00           H
    ATOM   2626  HB3 LEU A 264      89.001  24.709  64.366  1.00  0.00           H
    ATOM   2627  HG  LEU A 264      88.274  26.949  63.178  1.00  0.00           H
    ATOM   2628 HD11 LEU A 264      85.644  27.172  63.802  1.00  0.00           H
    ATOM   2629 HD12 LEU A 264      86.252  26.742  62.219  1.00  0.00           H
    ATOM   2630 HD13 LEU A 264      85.669  25.487  63.294  1.00  0.00           H
    ATOM   2631 HD21 LEU A 264      87.011  27.504  65.504  1.00  0.00           H
    ATOM   2632 HD22 LEU A 264      87.840  26.025  65.951  1.00  0.00           H
    ATOM   2633 HD23 LEU A 264      88.750  27.381  65.319  1.00  0.00           H
    ATOM   2634  N   LEU A 265      89.338  26.024  60.669  1.00 27.92           N
    ATOM   2635  CA  LEU A 265      90.144  27.112  60.122  1.00 20.28           C
    ATOM   2636  C   LEU A 265      89.481  28.468  60.341  1.00 18.19           C
    ATOM   2637  O   LEU A 265      90.199  29.461  60.377  1.00 25.26           O
    ATOM   2638  CB  LEU A 265      90.388  26.908  58.615  1.00 18.96           C
    ATOM   2639  CG  LEU A 265      91.058  25.576  58.237  1.00 20.57           C
    ATOM   2640  CD1 LEU A 265      91.452  25.589  56.749  1.00 20.12           C
    ATOM   2641  CD2 LEU A 265      92.250  25.197  59.137  1.00 22.54           C
    ATOM   2642  H   LEU A 265      88.449  25.859  60.211  1.00  0.00           H
    ATOM   2643  HA  LEU A 265      91.102  27.174  60.637  1.00  0.00           H
    ATOM   2644  HB2 LEU A 265      89.452  27.003  58.067  1.00  0.00           H
    ATOM   2645  HB3 LEU A 265      91.023  27.723  58.260  1.00  0.00           H
    ATOM   2646  HG  LEU A 265      90.312  24.795  58.364  1.00  0.00           H
    ATOM   2647 HD11 LEU A 265      91.278  24.618  56.287  1.00  0.00           H
    ATOM   2648 HD12 LEU A 265      90.872  26.317  56.179  1.00  0.00           H
    ATOM   2649 HD13 LEU A 265      92.504  25.837  56.607  1.00  0.00           H
    ATOM   2650 HD21 LEU A 265      93.052  24.730  58.564  1.00  0.00           H
    ATOM   2651 HD22 LEU A 265      92.691  26.056  59.640  1.00  0.00           H
    ATOM   2652 HD23 LEU A 265      91.951  24.474  59.896  1.00  0.00           H
    ATOM   2653  N   GLY A 266      88.153  28.508  60.483  1.00 16.22           N
    ATOM   2654  CA  GLY A 266      87.442  29.756  60.684  1.00 11.96           C
    ATOM   2655  C   GLY A 266      85.969  29.440  60.903  1.00 13.03           C
    ATOM   2656  O   GLY A 266      85.479  28.396  60.486  1.00 16.25           O
    ATOM   2657  H   GLY A 266      87.602  27.663  60.422  1.00  0.00           H
    ATOM   2658  HA2 GLY A 266      87.855  30.282  61.542  1.00  0.00           H
    ATOM   2659  HA3 GLY A 266      87.557  30.397  59.809  1.00  0.00           H
    ATOM   2660  N   ARG A 267      85.251  30.370  61.533  1.00 13.52           N
    ATOM   2661  CA  ARG A 267      83.818  30.286  61.746  1.00 16.05           C
    ATOM   2662  C   ARG A 267      83.217  31.684  61.802  1.00 15.69           C
    ATOM   2663  O   ARG A 267      83.834  32.606  62.329  1.00 10.45           O
    ATOM   2664  CB  ARG A 267      83.508  29.471  63.010  1.00 22.18           C
    ATOM   2665  CG  ARG A 267      82.004  29.353  63.333  1.00 23.78           C
    ATOM   2666  CD  ARG A 267      81.725  28.384  64.482  1.00 29.54           C
    ATOM   2667  NE  ARG A 267      82.541  28.709  65.650  1.00 32.24           N
    ATOM   2668  CZ  ARG A 267      83.508  27.955  66.184  1.00 24.84           C
    ATOM   2669  NH1 ARG A 267      83.752  26.711  65.779  1.00 19.66           N
    ATOM   2670  NH2 ARG A 267      84.259  28.467  67.154  1.00 22.57           N1+
    ATOM   2671  H   ARG A 267      85.698  31.227  61.841  1.00  0.00           H
    ATOM   2672  HA  ARG A 267      83.393  29.769  60.893  1.00  0.00           H
    ATOM   2673  HB2 ARG A 267      83.919  28.468  62.896  1.00  0.00           H
    ATOM   2674  HB3 ARG A 267      84.029  29.925  63.854  1.00  0.00           H
    ATOM   2675  HG2 ARG A 267      81.504  30.306  63.511  1.00  0.00           H
    ATOM   2676  HG3 ARG A 267      81.549  28.927  62.438  1.00  0.00           H
    ATOM   2677  HD2 ARG A 267      80.733  28.620  64.857  1.00  0.00           H
    ATOM   2678  HD3 ARG A 267      81.693  27.335  64.193  1.00  0.00           H
    ATOM   2679  HE  ARG A 267      82.452  29.660  65.991  1.00  0.00           H
    ATOM   2680 HH11 ARG A 267      83.169  26.265  65.083  1.00  0.00           H
    ATOM   2681 HH12 ARG A 267      84.501  26.173  66.191  1.00  0.00           H
    ATOM   2682 HH21 ARG A 267      84.121  29.422  67.460  1.00  0.00           H
    ATOM   2683 HH22 ARG A 267      85.011  27.937  67.571  1.00  0.00           H
    ATOM   2684  N   ASN A 268      81.997  31.789  61.275  1.00 18.66           N
    ATOM   2685  CA  ASN A 268      81.159  32.979  61.270  1.00 15.32           C
    ATOM   2686  C   ASN A 268      79.706  32.528  61.330  1.00 15.83           C
    ATOM   2687  O   ASN A 268      79.414  31.351  61.125  1.00 21.17           O
    ATOM   2688  CB  ASN A 268      81.439  33.763  59.980  1.00 22.83           C
    ATOM   2689  CG  ASN A 268      82.650  34.675  60.111  1.00 30.13           C
    ATOM   2690  OD1 ASN A 268      82.556  35.738  60.717  1.00 33.18           O
    ATOM   2691  ND2 ASN A 268      83.783  34.257  59.551  1.00 26.86           N
    ATOM   2692  H   ASN A 268      81.571  30.969  60.861  1.00  0.00           H
    ATOM   2693  HA  ASN A 268      81.413  33.565  62.157  1.00  0.00           H
    ATOM   2694  HB2 ASN A 268      81.543  33.079  59.136  1.00  0.00           H
    ATOM   2695  HB3 ASN A 268      80.604  34.426  59.751  1.00  0.00           H
    ATOM   2696 HD21 ASN A 268      83.819  33.375  59.062  1.00  0.00           H
    ATOM   2697 HD22 ASN A 268      84.621  34.823  59.602  1.00  0.00           H
    ATOM   2698  N   SER A 269      78.807  33.474  61.607  1.00 16.11           N
    ATOM   2699  CA  SER A 269      77.391  33.163  61.695  1.00 17.01           C
    ATOM   2700  C   SER A 269      76.511  34.372  61.405  1.00 17.22           C
    ATOM   2701  O   SER A 269      76.959  35.517  61.472  1.00 18.35           O
    ATOM   2702  CB  SER A 269      77.068  32.492  63.050  1.00 20.96           C
    ATOM   2703  OG  SER A 269      77.200  33.393  64.129  1.00 20.23           O
    ATOM   2704  H   SER A 269      79.079  34.434  61.766  1.00  0.00           H
    ATOM   2705  HA  SER A 269      77.185  32.462  60.888  1.00  0.00           H
    ATOM   2706  HB2 SER A 269      76.044  32.120  63.049  1.00  0.00           H
    ATOM   2707  HB3 SER A 269      77.711  31.631  63.230  1.00  0.00           H
    ATOM   2708  HG  SER A 269      78.144  33.519  64.289  1.00  0.00           H
    ATOM   2709  N   PHE A 270      75.253  34.061  61.093  1.00 13.73           N
    ATOM   2710  CA  PHE A 270      74.198  35.024  60.845  1.00 10.28           C
    ATOM   2711  C   PHE A 270      72.881  34.374  61.244  1.00 10.09           C
    ATOM   2712  O   PHE A 270      72.703  33.176  61.034  1.00 18.05           O
    ATOM   2713  CB  PHE A 270      74.229  35.490  59.371  1.00 13.04           C
    ATOM   2714  CG  PHE A 270      74.043  34.425  58.296  1.00 16.69           C
    ATOM   2715  CD1 PHE A 270      75.136  33.638  57.875  1.00 14.15           C
    ATOM   2716  CD2 PHE A 270      72.766  34.138  57.775  1.00 15.55           C
    ATOM   2717  CE1 PHE A 270      74.961  32.662  56.903  1.00 13.65           C
    ATOM   2718  CE2 PHE A 270      72.607  33.147  56.814  1.00 13.71           C
    ATOM   2719  CZ  PHE A 270      73.702  32.419  56.373  1.00 11.28           C
    ATOM   2720  H   PHE A 270      74.974  33.089  61.041  1.00  0.00           H
    ATOM   2721  HA  PHE A 270      74.350  35.890  61.492  1.00  0.00           H
    ATOM   2722  HB2 PHE A 270      73.462  36.253  59.226  1.00  0.00           H
    ATOM   2723  HB3 PHE A 270      75.174  36.001  59.180  1.00  0.00           H
    ATOM   2724  HD1 PHE A 270      76.116  33.808  58.293  1.00  0.00           H
    ATOM   2725  HD2 PHE A 270      71.914  34.716  58.089  1.00  0.00           H
    ATOM   2726  HE1 PHE A 270      75.806  32.082  56.564  1.00  0.00           H
    ATOM   2727  HE2 PHE A 270      71.631  32.951  56.397  1.00  0.00           H
    ATOM   2728  HZ  PHE A 270      73.575  31.656  55.619  1.00  0.00           H
    ATOM   2729  N   GLU A 271      71.964  35.183  61.787  1.00  6.66           N
    ATOM   2730  CA  GLU A 271      70.595  34.759  62.043  1.00  7.88           C
    ATOM   2731  C   GLU A 271      69.844  34.609  60.720  1.00 14.88           C
    ATOM   2732  O   GLU A 271      70.214  35.246  59.736  1.00 20.47           O
    ATOM   2733  CB  GLU A 271      69.914  35.739  63.010  1.00  9.49           C
    ATOM   2734  CG  GLU A 271      68.506  35.290  63.461  1.00 22.31           C
    ATOM   2735  CD  GLU A 271      67.885  36.210  64.501  1.00 29.53           C
    ATOM   2736  OE1 GLU A 271      68.661  36.830  65.258  1.00 30.89           O
    ATOM   2737  OE2 GLU A 271      66.637  36.260  64.536  1.00 29.88           O1-
    ATOM   2738  H   GLU A 271      72.156  36.172  61.886  1.00  0.00           H
    ATOM   2739  HA  GLU A 271      70.638  33.792  62.530  1.00  0.00           H
    ATOM   2740  HB2 GLU A 271      70.558  35.858  63.883  1.00  0.00           H
    ATOM   2741  HB3 GLU A 271      69.852  36.727  62.552  1.00  0.00           H
    ATOM   2742  HG2 GLU A 271      67.822  35.233  62.616  1.00  0.00           H
    ATOM   2743  HG3 GLU A 271      68.552  34.302  63.914  1.00  0.00           H
    ATOM   2744  N   VAL A 272      68.824  33.750  60.703  1.00 19.30           N
    ATOM   2745  CA  VAL A 272      68.037  33.529  59.507  1.00 14.73           C
    ATOM   2746  C   VAL A 272      66.564  33.448  59.860  1.00 17.17           C
    ATOM   2747  O   VAL A 272      66.204  32.817  60.853  1.00 21.48           O
    ATOM   2748  CB  VAL A 272      68.394  32.227  58.761  1.00 15.19           C
    ATOM   2749  CG1 VAL A 272      67.795  32.230  57.342  1.00 13.87           C
    ATOM   2750  CG2 VAL A 272      69.905  32.027  58.683  1.00 24.01           C
    ATOM   2751  H   VAL A 272      68.586  33.209  61.525  1.00  0.00           H
    ATOM   2752  HA  VAL A 272      68.187  34.356  58.826  1.00  0.00           H
    ATOM   2753  HB  VAL A 272      67.993  31.359  59.288  1.00  0.00           H
    ATOM   2754 HG11 VAL A 272      68.185  31.416  56.735  1.00  0.00           H
    ATOM   2755 HG12 VAL A 272      66.713  32.103  57.357  1.00  0.00           H
    ATOM   2756 HG13 VAL A 272      68.008  33.167  56.832  1.00  0.00           H
    ATOM   2757 HG21 VAL A 272      70.148  31.245  57.971  1.00  0.00           H
    ATOM   2758 HG22 VAL A 272      70.385  32.937  58.338  1.00  0.00           H
    ATOM   2759 HG23 VAL A 272      70.336  31.755  59.647  1.00  0.00           H
    ATOM   2760  N   ARG A 273      65.755  34.048  58.988  1.00 14.99           N
    ATOM   2761  CA  ARG A 273      64.320  33.949  59.017  1.00 14.64           C
    ATOM   2762  C   ARG A 273      63.833  33.546  57.631  1.00 17.45           C
    ATOM   2763  O   ARG A 273      64.049  34.284  56.673  1.00 17.86           O
    ATOM   2764  CB  ARG A 273      63.740  35.290  59.457  1.00 14.02           C
    ATOM   2765  CG  ARG A 273      62.228  35.192  59.541  1.00 15.34           C
    ATOM   2766  CD  ARG A 273      61.516  36.485  59.847  1.00 21.61           C
    ATOM   2767  NE  ARG A 273      60.111  36.130  59.982  1.00 33.75           N
    ATOM   2768  CZ  ARG A 273      59.095  36.973  60.145  1.00 39.26           C
    ATOM   2769  NH1 ARG A 273      59.323  38.278  60.277  1.00 40.31           N
    ATOM   2770  NH2 ARG A 273      57.853  36.484  60.200  1.00 36.90           N1+
    ATOM   2771  H   ARG A 273      66.150  34.560  58.208  1.00  0.00           H
    ATOM   2772  HA  ARG A 273      64.008  33.186  59.732  1.00  0.00           H
    ATOM   2773  HB2 ARG A 273      64.122  35.521  60.447  1.00  0.00           H
    ATOM   2774  HB3 ARG A 273      64.045  36.095  58.786  1.00  0.00           H
    ATOM   2775  HG2 ARG A 273      61.854  34.908  58.558  1.00  0.00           H
    ATOM   2776  HG3 ARG A 273      61.915  34.396  60.218  1.00  0.00           H
    ATOM   2777  HD2 ARG A 273      61.933  37.016  60.703  1.00  0.00           H
    ATOM   2778  HD3 ARG A 273      61.560  37.139  58.978  1.00  0.00           H
    ATOM   2779  HE  ARG A 273      59.904  35.144  59.916  1.00  0.00           H
    ATOM   2780 HH11 ARG A 273      60.276  38.616  60.279  1.00  0.00           H
    ATOM   2781 HH12 ARG A 273      58.578  38.933  60.466  1.00  0.00           H
    ATOM   2782 HH21 ARG A 273      57.690  35.494  60.099  1.00  0.00           H
    ATOM   2783 HH22 ARG A 273      57.070  37.108  60.327  1.00  0.00           H
    ATOM   2784  N   VAL A 274      63.133  32.412  57.571  1.00 19.63           N
    ATOM   2785  CA  VAL A 274      62.580  31.896  56.332  1.00 11.05           C
    ATOM   2786  C   VAL A 274      61.081  32.225  56.332  1.00 19.35           C
    ATOM   2787  O   VAL A 274      60.337  31.708  57.162  1.00 31.22           O
    ATOM   2788  CB  VAL A 274      62.788  30.375  56.208  1.00  9.65           C
    ATOM   2789  CG1 VAL A 274      62.404  29.871  54.800  1.00 15.63           C
    ATOM   2790  CG2 VAL A 274      64.231  29.970  56.548  1.00  9.66           C
    ATOM   2791  H   VAL A 274      62.940  31.884  58.412  1.00  0.00           H
    ATOM   2792  HA  VAL A 274      63.073  32.355  55.480  1.00  0.00           H
    ATOM   2793  HB  VAL A 274      62.168  29.871  56.946  1.00  0.00           H
    ATOM   2794 HG11 VAL A 274      62.636  28.814  54.673  1.00  0.00           H
    ATOM   2795 HG12 VAL A 274      61.341  29.994  54.596  1.00  0.00           H
    ATOM   2796 HG13 VAL A 274      62.951  30.410  54.025  1.00  0.00           H
    ATOM   2797 HG21 VAL A 274      64.307  28.887  56.573  1.00  0.00           H
    ATOM   2798 HG22 VAL A 274      64.946  30.367  55.829  1.00  0.00           H
    ATOM   2799 HG23 VAL A 274      64.539  30.301  57.539  1.00  0.00           H
    ATOM   2800  N   CYS A 275      60.682  33.141  55.447  1.00 16.57           N
    ATOM   2801  CA  CYS A 275      59.386  33.809  55.483  1.00 18.24           C
    ATOM   2802  C   CYS A 275      58.957  34.226  54.077  1.00 28.96           C
    ATOM   2803  O   CYS A 275      59.801  34.403  53.201  1.00 29.01           O
    ATOM   2804  CB  CYS A 275      59.440  35.024  56.421  1.00 18.03           C
    ATOM   2805  SG  CYS A 275      60.745  36.190  55.904  1.00 24.16           S
    ATOM   2806  H   CYS A 275      61.346  33.508  54.775  1.00  0.00           H
    ATOM   2807  HA  CYS A 275      58.632  33.111  55.849  1.00  0.00           H
    ATOM   2808  HB2 CYS A 275      58.482  35.540  56.432  1.00  0.00           H
    ATOM   2809  HB3 CYS A 275      59.636  34.709  57.446  1.00  0.00           H
    ATOM   2810  HG  CYS A 275      60.299  37.196  56.668  1.00  0.00           H
    ATOM   2811  N   ALA A 276      57.637  34.365  53.887  1.00 33.26           N
    ATOM   2812  CA  ALA A 276      57.028  34.675  52.594  1.00 32.70           C
    ATOM   2813  C   ALA A 276      57.432  36.033  52.002  1.00 29.50           C
    ATOM   2814  O   ALA A 276      57.635  36.096  50.791  1.00 30.26           O
    ATOM   2815  CB  ALA A 276      55.503  34.565  52.702  1.00 31.73           C
    ATOM   2816  H   ALA A 276      56.999  34.219  54.656  1.00  0.00           H
    ATOM   2817  HA  ALA A 276      57.361  33.912  51.890  1.00  0.00           H
    ATOM   2818  HB1 ALA A 276      55.023  34.780  51.746  1.00  0.00           H
    ATOM   2819  HB2 ALA A 276      55.204  33.558  52.996  1.00  0.00           H
    ATOM   2820  HB3 ALA A 276      55.104  35.262  53.440  1.00  0.00           H
    ATOM   2821  N   CYS A 277      57.581  37.074  52.837  1.00 26.45           N
    ATOM   2822  CA  CYS A 277      57.927  38.416  52.373  1.00 28.12           C
    ATOM   2823  C   CYS A 277      59.172  38.942  53.117  1.00 25.48           C
    ATOM   2824  O   CYS A 277      59.023  39.656  54.110  1.00 30.49           O
    ATOM   2825  CB  CYS A 277      56.713  39.366  52.418  1.00 35.97           C
    ATOM   2826  SG  CYS A 277      56.993  40.783  51.313  1.00 46.65           S
    ATOM   2827  H   CYS A 277      57.389  36.976  53.828  1.00  0.00           H
    ATOM   2828  HA  CYS A 277      58.192  38.380  51.316  1.00  0.00           H
    ATOM   2829  HB2 CYS A 277      55.821  38.850  52.060  1.00  0.00           H
    ATOM   2830  HB3 CYS A 277      56.490  39.702  53.431  1.00  0.00           H
    ATOM   2831  HG  CYS A 277      56.826  40.103  50.174  1.00  0.00           H
    ATOM   2832  N   PRO A 278      60.384  38.581  52.628  1.00 19.94           N
    ATOM   2833  CA  PRO A 278      61.665  39.038  53.199  1.00 17.78           C
    ATOM   2834  C   PRO A 278      61.861  40.554  53.316  1.00 24.14           C
    ATOM   2835  O   PRO A 278      62.437  40.994  54.308  1.00 32.91           O
    ATOM   2836  CB  PRO A 278      62.733  38.439  52.270  1.00 16.46           C
    ATOM   2837  CG  PRO A 278      62.075  37.228  51.657  1.00 17.57           C
    ATOM   2838  CD  PRO A 278      60.618  37.626  51.543  1.00 16.41           C
    ATOM   2839  HA  PRO A 278      61.738  38.587  54.189  1.00  0.00           H
    ATOM   2840  HB2 PRO A 278      62.997  39.134  51.471  1.00  0.00           H
    ATOM   2841  HB3 PRO A 278      63.653  38.184  52.791  1.00  0.00           H
    ATOM   2842  HG2 PRO A 278      62.522  36.905  50.716  1.00  0.00           H
    ATOM   2843  HG3 PRO A 278      62.145  36.408  52.370  1.00  0.00           H
    ATOM   2844  HD2 PRO A 278      60.415  38.097  50.583  1.00  0.00           H
    ATOM   2845  HD3 PRO A 278      60.010  36.733  51.631  1.00  0.00           H
    ATOM   2846  N   GLY A 279      61.391  41.316  52.315  1.00 17.64           N
    ATOM   2847  CA  GLY A 279      61.511  42.768  52.279  1.00 16.25           C
    ATOM   2848  C   GLY A 279      60.661  43.405  53.383  1.00 20.38           C
    ATOM   2849  O   GLY A 279      61.177  44.230  54.131  1.00 26.61           O
    ATOM   2850  H   GLY A 279      60.939  40.877  51.521  1.00  0.00           H
    ATOM   2851  HA2 GLY A 279      62.556  43.052  52.399  1.00  0.00           H
    ATOM   2852  HA3 GLY A 279      61.181  43.126  51.304  1.00  0.00           H
    ATOM   2853  N   ARG A 280      59.386  42.987  53.511  1.00 25.90           N
    ATOM   2854  CA  ARG A 280      58.445  43.451  54.536  1.00 27.30           C
    ATOM   2855  C   ARG A 280      58.968  43.187  55.952  1.00 22.89           C
    ATOM   2856  O   ARG A 280      59.039  44.114  56.757  1.00 23.38           O
    ATOM   2857  CB  ARG A 280      57.070  42.773  54.336  1.00 35.47           C
    ATOM   2858  CG  ARG A 280      55.940  43.260  55.274  1.00 45.40           C
    ATOM   2859  CD  ARG A 280      54.995  42.154  55.785  1.00 52.89           C
    ATOM   2860  NE  ARG A 280      55.580  41.353  56.873  1.00 58.02           N
    ATOM   2861  CZ  ARG A 280      55.707  41.712  58.165  1.00 58.91           C
    ATOM   2862  NH1 ARG A 280      55.320  42.918  58.595  1.00 60.17           N
    ATOM   2863  NH2 ARG A 280      56.241  40.852  59.040  1.00 58.03           N1+
    ATOM   2864  H   ARG A 280      59.040  42.273  52.885  1.00  0.00           H
    ATOM   2865  HA  ARG A 280      58.329  44.527  54.420  1.00  0.00           H
    ATOM   2866  HB2 ARG A 280      56.735  42.889  53.306  1.00  0.00           H
    ATOM   2867  HB3 ARG A 280      57.209  41.701  54.478  1.00  0.00           H
    ATOM   2868  HG2 ARG A 280      56.403  43.664  56.173  1.00  0.00           H
    ATOM   2869  HG3 ARG A 280      55.399  44.108  54.855  1.00  0.00           H
    ATOM   2870  HD2 ARG A 280      53.993  42.519  56.008  1.00  0.00           H
    ATOM   2871  HD3 ARG A 280      54.877  41.431  54.978  1.00  0.00           H
    ATOM   2872  HE  ARG A 280      56.073  40.508  56.596  1.00  0.00           H
    ATOM   2873 HH11 ARG A 280      54.942  43.588  57.942  1.00  0.00           H
    ATOM   2874 HH12 ARG A 280      55.431  43.181  59.564  1.00  0.00           H
    ATOM   2875 HH21 ARG A 280      56.620  39.982  58.686  1.00  0.00           H
    ATOM   2876 HH22 ARG A 280      56.366  41.097  60.011  1.00  0.00           H
    ATOM   2877  N   ASP A 281      59.329  41.923  56.202  1.00 20.10           N
    ATOM   2878  CA  ASP A 281      59.838  41.424  57.475  1.00 17.96           C
    ATOM   2879  C   ASP A 281      61.116  42.117  57.925  1.00 19.55           C
    ATOM   2880  O   ASP A 281      61.189  42.522  59.083  1.00 21.86           O
    ATOM   2881  CB  ASP A 281      60.053  39.901  57.472  1.00 17.11           C
    ATOM   2882  CG  ASP A 281      58.768  39.101  57.267  1.00 26.83           C
    ATOM   2883  OD1 ASP A 281      57.726  39.691  56.927  1.00 29.46           O
    ATOM   2884  OD2 ASP A 281      58.821  37.875  57.460  1.00 32.76           O1-
    ATOM   2885  H   ASP A 281      59.217  41.225  55.477  1.00  0.00           H
    ATOM   2886  HA  ASP A 281      59.077  41.654  58.225  1.00  0.00           H
    ATOM   2887  HB2 ASP A 281      60.743  39.636  56.669  1.00  0.00           H
    ATOM   2888  HB3 ASP A 281      60.514  39.605  58.413  1.00  0.00           H
    ATOM   2889  N   ARG A 282      62.078  42.275  57.003  1.00 23.83           N
    ATOM   2890  CA  ARG A 282      63.310  43.002  57.278  1.00 24.63           C
    ATOM   2891  C   ARG A 282      63.048  44.455  57.688  1.00 27.52           C
    ATOM   2892  O   ARG A 282      63.586  44.890  58.703  1.00 32.30           O
    ATOM   2893  CB  ARG A 282      64.279  42.921  56.083  1.00 20.79           C
    ATOM   2894  CG  ARG A 282      65.450  43.900  56.251  1.00 13.35           C
    ATOM   2895  CD  ARG A 282      66.648  43.671  55.355  1.00 11.08           C
    ATOM   2896  NE  ARG A 282      67.572  44.792  55.537  1.00 15.35           N
    ATOM   2897  CZ  ARG A 282      68.376  45.314  54.606  1.00 17.15           C
    ATOM   2898  NH1 ARG A 282      68.375  44.855  53.356  1.00 20.08           N
    ATOM   2899  NH2 ARG A 282      69.202  46.310  54.923  1.00 13.35           N1+
    ATOM   2900  H   ARG A 282      61.959  41.908  56.067  1.00  0.00           H
    ATOM   2901  HA  ARG A 282      63.788  42.507  58.125  1.00  0.00           H
    ATOM   2902  HB2 ARG A 282      64.642  41.900  55.975  1.00  0.00           H
    ATOM   2903  HB3 ARG A 282      63.755  43.160  55.158  1.00  0.00           H
    ATOM   2904  HG2 ARG A 282      65.157  44.950  56.221  1.00  0.00           H
    ATOM   2905  HG3 ARG A 282      65.817  43.730  57.264  1.00  0.00           H
    ATOM   2906  HD2 ARG A 282      67.197  42.783  55.670  1.00  0.00           H
    ATOM   2907  HD3 ARG A 282      66.333  43.508  54.328  1.00  0.00           H
    ATOM   2908  HE  ARG A 282      67.632  45.179  56.478  1.00  0.00           H
    ATOM   2909 HH11 ARG A 282      67.838  44.029  53.119  1.00  0.00           H
    ATOM   2910 HH12 ARG A 282      69.021  45.238  52.671  1.00  0.00           H
    ATOM   2911 HH21 ARG A 282      69.155  46.743  55.838  1.00  0.00           H
    ATOM   2912 HH22 ARG A 282      69.814  46.708  54.226  1.00  0.00           H
    ATOM   2913  N   ARG A 283      62.250  45.163  56.875  1.00 29.79           N
    ATOM   2914  CA  ARG A 283      61.928  46.578  57.041  1.00 28.89           C
    ATOM   2915  C   ARG A 283      61.308  46.866  58.411  1.00 28.05           C
    ATOM   2916  O   ARG A 283      61.747  47.785  59.094  1.00 24.98           O
    ATOM   2917  CB  ARG A 283      61.021  47.005  55.867  1.00 26.39           C
    ATOM   2918  CG  ARG A 283      61.029  48.513  55.553  1.00 26.52           C
    ATOM   2919  CD  ARG A 283      61.174  48.832  54.048  1.00 31.60           C
    ATOM   2920  NE  ARG A 283      59.934  49.188  53.352  1.00 27.52           N
    ATOM   2921  CZ  ARG A 283      59.812  49.260  52.012  1.00 24.25           C
    ATOM   2922  NH1 ARG A 283      60.782  48.841  51.184  1.00 24.13           N
    ATOM   2923  NH2 ARG A 283      58.697  49.782  51.489  1.00 27.49           N1+
    ATOM   2924  H   ARG A 283      61.852  44.712  56.061  1.00  0.00           H
    ATOM   2925  HA  ARG A 283      62.873  47.119  56.985  1.00  0.00           H
    ATOM   2926  HB2 ARG A 283      61.445  46.497  55.007  1.00  0.00           H
    ATOM   2927  HB3 ARG A 283      60.004  46.618  55.949  1.00  0.00           H
    ATOM   2928  HG2 ARG A 283      60.152  48.987  55.995  1.00  0.00           H
    ATOM   2929  HG3 ARG A 283      61.883  48.974  56.048  1.00  0.00           H
    ATOM   2930  HD2 ARG A 283      61.972  49.548  53.852  1.00  0.00           H
    ATOM   2931  HD3 ARG A 283      61.440  47.918  53.535  1.00  0.00           H
    ATOM   2932  HE  ARG A 283      59.171  49.506  53.933  1.00  0.00           H
    ATOM   2933 HH11 ARG A 283      61.630  48.439  51.562  1.00  0.00           H
    ATOM   2934 HH12 ARG A 283      60.676  48.914  50.183  1.00  0.00           H
    ATOM   2935 HH21 ARG A 283      57.961  50.116  52.094  1.00  0.00           H
    ATOM   2936 HH22 ARG A 283      58.597  49.874  50.489  1.00  0.00           H
    ATOM   2937  N   THR A 284      60.370  45.986  58.787  1.00 30.61           N
    ATOM   2938  CA  THR A 284      59.669  45.914  60.059  1.00 31.73           C
    ATOM   2939  C   THR A 284      60.602  45.652  61.257  1.00 31.01           C
    ATOM   2940  O   THR A 284      60.567  46.432  62.204  1.00 37.72           O
    ATOM   2941  CB  THR A 284      58.596  44.799  59.989  1.00 34.16           C
    ATOM   2942  OG1 THR A 284      57.607  45.190  59.055  1.00 36.95           O
    ATOM   2943  CG2 THR A 284      57.903  44.440  61.312  1.00 34.83           C
    ATOM   2944  H   THR A 284      60.135  45.250  58.135  1.00  0.00           H
    ATOM   2945  HA  THR A 284      59.159  46.864  60.214  1.00  0.00           H
    ATOM   2946  HB  THR A 284      59.044  43.888  59.595  1.00  0.00           H
    ATOM   2947 HG21 THR A 284      57.087  43.736  61.147  1.00  0.00           H
    ATOM   2948 HG22 THR A 284      58.586  43.967  62.019  1.00  0.00           H
    ATOM   2949 HG23 THR A 284      57.485  45.326  61.790  1.00  0.00           H
    ATOM   2950  HG1 THR A 284      58.003  45.180  58.178  1.00  0.00           H
    ATOM   2951  N   GLU A 285      61.430  44.592  61.217  1.00 19.03           N
    ATOM   2952  CA  GLU A 285      62.359  44.294  62.315  1.00 22.01           C
    ATOM   2953  C   GLU A 285      63.451  45.355  62.528  1.00 30.73           C
    ATOM   2954  O   GLU A 285      63.930  45.507  63.651  1.00 36.00           O
    ATOM   2955  CB  GLU A 285      62.983  42.900  62.181  1.00 20.96           C
    ATOM   2956  CG  GLU A 285      61.971  41.761  62.373  1.00 23.46           C
    ATOM   2957  CD  GLU A 285      62.640  40.393  62.478  1.00 26.89           C
    ATOM   2958  OE1 GLU A 285      63.784  40.336  62.982  1.00 23.11           O
    ATOM   2959  OE2 GLU A 285      61.988  39.411  62.060  1.00 28.91           O1-
    ATOM   2960  H   GLU A 285      61.440  43.966  60.422  1.00  0.00           H
    ATOM   2961  HA  GLU A 285      61.769  44.282  63.227  1.00  0.00           H
    ATOM   2962  HB2 GLU A 285      63.472  42.800  61.210  1.00  0.00           H
    ATOM   2963  HB3 GLU A 285      63.766  42.807  62.935  1.00  0.00           H
    ATOM   2964  HG2 GLU A 285      61.403  41.918  63.290  1.00  0.00           H
    ATOM   2965  HG3 GLU A 285      61.253  41.754  61.555  1.00  0.00           H
    ATOM   2966  N   GLU A 286      63.798  46.087  61.461  1.00 31.09           N
    ATOM   2967  CA  GLU A 286      64.685  47.246  61.516  1.00 29.92           C
    ATOM   2968  C   GLU A 286      63.955  48.522  61.924  1.00 37.30           C
    ATOM   2969  O   GLU A 286      64.594  49.441  62.429  1.00 38.02           O
    ATOM   2970  CB  GLU A 286      65.310  47.471  60.145  1.00 21.90           C
    ATOM   2971  CG  GLU A 286      66.303  46.370  59.756  1.00 22.96           C
    ATOM   2972  CD  GLU A 286      66.887  46.617  58.373  1.00 21.54           C
    ATOM   2973  OE1 GLU A 286      66.320  47.435  57.619  1.00 19.96           O
    ATOM   2974  OE2 GLU A 286      67.917  45.983  58.071  1.00 20.27           O1-
    ATOM   2975  H   GLU A 286      63.394  45.875  60.559  1.00  0.00           H
    ATOM   2976  HA  GLU A 286      65.485  47.064  62.232  1.00  0.00           H
    ATOM   2977  HB2 GLU A 286      64.513  47.536  59.401  1.00  0.00           H
    ATOM   2978  HB3 GLU A 286      65.825  48.434  60.123  1.00  0.00           H
    ATOM   2979  HG2 GLU A 286      67.117  46.334  60.481  1.00  0.00           H
    ATOM   2980  HG3 GLU A 286      65.835  45.387  59.763  1.00  0.00           H
    ATOM   2981  N   GLU A 287      62.634  48.562  61.706  1.00 40.40           N
    ATOM   2982  CA  GLU A 287      61.761  49.619  62.184  1.00 43.42           C
    ATOM   2983  C   GLU A 287      61.652  49.605  63.715  1.00 47.95           C
    ATOM   2984  O   GLU A 287      61.580  50.681  64.308  1.00 47.29           O
    ATOM   2985  CB  GLU A 287      60.438  49.606  61.383  1.00 44.55           C
    ATOM   2986  CG  GLU A 287      59.219  50.324  61.992  1.00 53.57           C
    ATOM   2987  CD  GLU A 287      58.292  49.386  62.770  1.00 66.08           C
    ATOM   2988  OE1 GLU A 287      58.764  48.759  63.742  1.00 69.99           O
    ATOM   2989  OE2 GLU A 287      57.112  49.303  62.370  1.00 70.00           O1-
    ATOM   2990  H   GLU A 287      62.177  47.782  61.252  1.00  0.00           H
    ATOM   2991  HA  GLU A 287      62.242  50.571  61.947  1.00  0.00           H
    ATOM   2992  HB2 GLU A 287      60.644  50.080  60.422  1.00  0.00           H
    ATOM   2993  HB3 GLU A 287      60.158  48.593  61.121  1.00  0.00           H
    ATOM   2994  HG2 GLU A 287      59.516  51.163  62.622  1.00  0.00           H
    ATOM   2995  HG3 GLU A 287      58.637  50.762  61.179  1.00  0.00           H
    ATOM   2996  N   ASN A 288      61.749  48.406  64.324  1.00 55.63           N
    ATOM   2997  CA  ASN A 288      61.909  48.244  65.773  1.00 58.55           C
    ATOM   2998  C   ASN A 288      63.302  48.711  66.254  1.00 60.68           C
    ATOM   2999  O   ASN A 288      63.458  48.972  67.444  1.00 61.83           O
    ATOM   3000  CB  ASN A 288      61.740  46.779  66.264  1.00 57.64           C
    ATOM   3001  CG  ASN A 288      60.731  45.848  65.586  1.00 62.87           C
    ATOM   3002  OD1 ASN A 288      60.995  44.650  65.515  1.00 64.88           O
    ATOM   3003  ND2 ASN A 288      59.589  46.341  65.107  1.00 62.47           N
    ATOM   3004  H   ASN A 288      61.766  47.563  63.766  1.00  0.00           H
    ATOM   3005  HA  ASN A 288      61.199  48.876  66.311  1.00  0.00           H
    ATOM   3006  HB2 ASN A 288      62.696  46.264  66.139  1.00  0.00           H
    ATOM   3007  HB3 ASN A 288      61.551  46.771  67.338  1.00  0.00           H
    ATOM   3008 HD21 ASN A 288      59.410  47.339  65.093  1.00  0.00           H
    ATOM   3009 HD22 ASN A 288      58.922  45.726  64.667  1.00  0.00           H
    ATOM   3010  N   LEU A 289      64.269  48.763  65.314  1.00 59.72           N
    ATOM   3011  CA  LEU A 289      65.705  49.058  65.405  1.00 52.31           C
    ATOM   3012  C   LEU A 289      66.397  47.730  65.211  1.00 50.96           C
    ATOM   3013  O   LEU A 289      67.103  47.183  66.052  1.00 49.07           O
    ATOM   3014  CB  LEU A 289      66.187  49.871  66.639  1.00 49.31           C
    ATOM   3015  CG  LEU A 289      65.760  51.357  66.658  1.00 50.21           C
    ATOM   3016  CD1 LEU A 289      66.712  52.169  67.556  1.00 53.81           C
    ATOM   3017  CD2 LEU A 289      65.660  52.001  65.262  1.00 46.63           C
    ATOM   3018  H   LEU A 289      63.975  48.537  64.374  1.00  0.00           H
    ATOM   3019  HA  LEU A 289      65.987  49.596  64.505  1.00  0.00           H
    ATOM   3021  HB2 LEU A 289      65.935  49.382  67.581  1.00  0.00           H
    ATOM   3022  HB3 LEU A 289      67.278  49.867  66.606  1.00  0.00           H
    ATOM   3023  HG  LEU A 289      64.764  51.406  67.101  1.00  0.00           H
    ATOM   3024 HD11 LEU A 289      66.149  52.860  68.183  1.00  0.00           H
    ATOM   3025 HD12 LEU A 289      67.290  51.532  68.227  1.00  0.00           H
    ATOM   3026 HD13 LEU A 289      67.423  52.757  66.975  1.00  0.00           H
    ATOM   3027 HD21 LEU A 289      65.627  53.088  65.335  1.00  0.00           H
    ATOM   3028 HD22 LEU A 289      66.516  51.751  64.634  1.00  0.00           H
    ATOM   3029 HD23 LEU A 289      64.750  51.691  64.747  1.00  0.00           H
    ATOM   3031  N   SER B  96      57.270  -4.072  85.505  1.00 60.24           N1+
    ATOM   3032  CA  SER B  96      57.257  -3.830  86.964  1.00 66.25           C
    ATOM   3033  C   SER B  96      57.175  -2.307  87.261  1.00 69.42           C
    ATOM   3034  O   SER B  96      56.847  -1.499  86.387  1.00 69.89           O
    ATOM   3035  CB  SER B  96      58.422  -4.555  87.696  1.00  0.00           C
    ATOM   3036  OG  SER B  96      59.642  -3.832  87.690  1.00  0.00           O
    ATOM   3038  H   SER B  96      56.400  -3.761  85.097  1.00  0.00           H
    ATOM   3039  HA  SER B  96      56.334  -4.260  87.355  1.00  0.00           H
    ATOM   3040  HB2 SER B  96      58.153  -4.731  88.739  1.00  0.00           H
    ATOM   3041  HB3 SER B  96      58.598  -5.542  87.265  1.00  0.00           H
    ATOM   3042  HG  SER B  96      59.965  -3.796  86.786  1.00  0.00           H
    ATOM   3043  N   VAL B  97      57.478  -1.961  88.529  1.00 70.00           N
    ATOM   3044  CA  VAL B  97      57.541  -0.608  89.072  1.00 70.00           C
    ATOM   3045  C   VAL B  97      58.613  -0.535  90.207  1.00 69.52           C
    ATOM   3046  O   VAL B  97      58.710  -1.475  90.997  1.00 66.98           O
    ATOM   3047  CB  VAL B  97      56.148  -0.158  89.604  1.00 68.79           C
    ATOM   3048  CG1 VAL B  97      56.165   1.061  90.551  1.00 67.96           C
    ATOM   3049  CG2 VAL B  97      55.179   0.141  88.443  1.00 65.53           C
    ATOM   3050  H   VAL B  97      57.752  -2.686  89.177  1.00  0.00           H
    ATOM   3051  HA  VAL B  97      57.784   0.041  88.235  1.00  0.00           H
    ATOM   3052  HB  VAL B  97      55.718  -0.982  90.175  1.00  0.00           H
    ATOM   3053 HG11 VAL B  97      55.152   1.357  90.826  1.00  0.00           H
    ATOM   3054 HG12 VAL B  97      56.683   0.856  91.488  1.00  0.00           H
    ATOM   3055 HG13 VAL B  97      56.638   1.926  90.084  1.00  0.00           H
    ATOM   3056 HG21 VAL B  97      54.258   0.581  88.821  1.00  0.00           H
    ATOM   3057 HG22 VAL B  97      55.605   0.841  87.725  1.00  0.00           H
    ATOM   3058 HG23 VAL B  97      54.887  -0.764  87.912  1.00  0.00           H
    ATOM   3059  N   PRO B  98      59.404   0.567  90.255  1.00 66.82           N
    ATOM   3060  CA  PRO B  98      60.449   0.809  91.282  1.00 62.56           C
    ATOM   3061  C   PRO B  98      60.046   1.786  92.417  1.00 56.09           C
    ATOM   3062  O   PRO B  98      59.256   2.703  92.190  1.00 57.22           O
    ATOM   3063  CB  PRO B  98      61.586   1.390  90.433  1.00 60.11           C
    ATOM   3064  CG  PRO B  98      60.889   2.178  89.337  1.00 58.67           C
    ATOM   3065  CD  PRO B  98      59.589   1.446  89.100  1.00 62.14           C
    ATOM   3066  HA  PRO B  98      60.782  -0.127  91.736  1.00  0.00           H
    ATOM   3067  HB2 PRO B  98      62.297   1.997  90.996  1.00  0.00           H
    ATOM   3068  HB3 PRO B  98      62.148   0.569  89.986  1.00  0.00           H
    ATOM   3069  HG2 PRO B  98      60.686   3.185  89.695  1.00  0.00           H
    ATOM   3070  HG3 PRO B  98      61.462   2.246  88.420  1.00  0.00           H
    ATOM   3071  HD2 PRO B  98      58.757   2.137  88.962  1.00  0.00           H
    ATOM   3072  HD3 PRO B  98      59.719   0.857  88.191  1.00  0.00           H
    ATOM   3073  N   SER B  99      60.621   1.583  93.618  1.00 48.69           N
    ATOM   3074  CA  SER B  99      60.281   2.359  94.815  1.00 48.23           C
    ATOM   3075  C   SER B  99      61.187   3.584  95.035  1.00 49.35           C
    ATOM   3076  O   SER B  99      62.349   3.594  94.630  1.00 59.42           O
    ATOM   3077  CB  SER B  99      60.298   1.422  96.031  1.00 52.92           C
    ATOM   3078  OG  SER B  99      59.601   2.001  97.113  1.00 58.16           O
    ATOM   3079  H   SER B  99      61.286   0.835  93.739  1.00  0.00           H
    ATOM   3080  HA  SER B  99      59.254   2.720  94.706  1.00  0.00           H
    ATOM   3081  HB2 SER B  99      59.809   0.483  95.777  1.00  0.00           H
    ATOM   3082  HB3 SER B  99      61.313   1.165  96.325  1.00  0.00           H
    ATOM   3083  HG  SER B  99      59.337   1.287  97.710  1.00  0.00           H
    ATOM   3084  N   GLN B 100      60.618   4.575  95.733  1.00 44.72           N
    ATOM   3085  CA  GLN B 100      61.272   5.800  96.184  1.00 46.53           C
    ATOM   3086  C   GLN B 100      61.592   5.730  97.687  1.00 44.78           C
    ATOM   3087  O   GLN B 100      61.578   6.764  98.355  1.00 49.25           O
    ATOM   3088  CB  GLN B 100      60.339   6.978  95.855  1.00 47.79           C
    ATOM   3089  CG  GLN B 100      58.998   6.985  96.633  1.00 49.38           C
    ATOM   3090  CD  GLN B 100      57.785   6.866  95.715  1.00 53.91           C
    ATOM   3091  OE1 GLN B 100      57.613   5.859  95.034  1.00 52.02           O
    ATOM   3092  NE2 GLN B 100      56.932   7.891  95.708  1.00 55.37           N
    ATOM   3093  H   GLN B 100      59.654   4.468  96.021  1.00  0.00           H
    ATOM   3094  HA  GLN B 100      62.214   5.946  95.653  1.00  0.00           H
    ATOM   3095  HB2 GLN B 100      60.864   7.914  96.046  1.00  0.00           H
    ATOM   3096  HB3 GLN B 100      60.170   6.969  94.779  1.00  0.00           H
    ATOM   3097  HG2 GLN B 100      58.934   6.167  97.351  1.00  0.00           H
    ATOM   3098  HG3 GLN B 100      58.929   7.901  97.222  1.00  0.00           H
    ATOM   3099 HE21 GLN B 100      57.091   8.700  96.289  1.00  0.00           H
    ATOM   3100 HE22 GLN B 100      56.110   7.855  95.121  1.00  0.00           H
    ATOM   3101  N   LYS B 101      61.821   4.516  98.212  1.00 33.59           N
    ATOM   3102  CA  LYS B 101      62.078   4.299  99.632  1.00 32.15           C
    ATOM   3103  C   LYS B 101      63.572   4.454  99.932  1.00 29.10           C
    ATOM   3104  O   LYS B 101      64.371   3.639  99.480  1.00 25.34           O
    ATOM   3105  CB  LYS B 101      61.493   2.940 100.085  1.00 40.03           C
    ATOM   3106  CG  LYS B 101      61.818   2.582 101.543  1.00 50.04           C
    ATOM   3107  CD  LYS B 101      61.413   1.156 101.925  1.00 54.58           C
    ATOM   3108  CE  LYS B 101      61.739   0.877 103.399  1.00 57.31           C
    ATOM   3109  NZ  LYS B 101      61.835  -0.564 103.674  1.00 54.87           N1+
    ATOM   3110  H   LYS B 101      61.730   3.700  97.623  1.00  0.00           H
    ATOM   3111  HA  LYS B 101      61.541   5.056 100.208  1.00  0.00           H
    ATOM   3112  HB2 LYS B 101      60.410   2.959  99.956  1.00  0.00           H
    ATOM   3113  HB3 LYS B 101      61.827   2.123  99.462  1.00  0.00           H
    ATOM   3114  HG2 LYS B 101      62.885   2.677 101.740  1.00  0.00           H
    ATOM   3115  HG3 LYS B 101      61.322   3.294 102.204  1.00  0.00           H
    ATOM   3116  HD2 LYS B 101      60.349   1.002 101.740  1.00  0.00           H
    ATOM   3117  HD3 LYS B 101      61.943   0.450 101.283  1.00  0.00           H
    ATOM   3118  HE2 LYS B 101      62.706   1.310 103.659  1.00  0.00           H
    ATOM   3119  HE3 LYS B 101      60.997   1.337 104.052  1.00  0.00           H
    ATOM   3120  HZ1 LYS B 101      62.556  -0.972 103.096  1.00  0.00           H
    ATOM   3121  HZ2 LYS B 101      62.064  -0.707 104.647  1.00  0.00           H
    ATOM   3122  HZ3 LYS B 101      60.951  -1.006 103.470  1.00  0.00           H
    ATOM   3123  N   THR B 102      63.924   5.466 100.736  1.00 37.07           N
    ATOM   3124  CA  THR B 102      65.298   5.658 101.193  1.00 46.23           C
    ATOM   3125  C   THR B 102      65.869   4.424 101.924  1.00 50.56           C
    ATOM   3126  O   THR B 102      65.236   3.872 102.824  1.00 56.09           O
    ATOM   3127  CB  THR B 102      65.551   6.995 101.961  1.00 49.38           C
    ATOM   3128  OG1 THR B 102      66.342   6.878 103.132  1.00 52.81           O
    ATOM   3129  CG2 THR B 102      64.292   7.817 102.280  1.00 49.91           C
    ATOM   3130  H   THR B 102      63.230   6.106 101.092  1.00  0.00           H
    ATOM   3131  HA  THR B 102      65.856   5.757 100.260  1.00  0.00           H
    ATOM   3132  HB  THR B 102      66.154   7.620 101.300  1.00  0.00           H
    ATOM   3133 HG21 THR B 102      64.549   8.736 102.806  1.00  0.00           H
    ATOM   3134 HG22 THR B 102      63.763   8.106 101.371  1.00  0.00           H
    ATOM   3135 HG23 THR B 102      63.600   7.259 102.912  1.00  0.00           H
    ATOM   3136  HG1 THR B 102      65.835   6.408 103.799  1.00  0.00           H
    ATOM   3137  N   TYR B 103      67.044   4.010 101.453  1.00 51.01           N
    ATOM   3138  CA  TYR B 103      67.851   2.948 102.014  1.00 47.70           C
    ATOM   3139  C   TYR B 103      69.232   3.563 102.104  1.00 50.02           C
    ATOM   3140  O   TYR B 103      70.116   3.212 101.338  1.00 48.05           O
    ATOM   3141  CB  TYR B 103      67.774   1.673 101.138  1.00 43.84           C
    ATOM   3142  CG  TYR B 103      68.574   0.451 101.573  1.00 50.08           C
    ATOM   3143  CD1 TYR B 103      68.174  -0.307 102.691  1.00 56.40           C
    ATOM   3144  CD2 TYR B 103      69.679   0.020 100.808  1.00 54.61           C
    ATOM   3145  CE1 TYR B 103      68.860  -1.492 103.028  1.00 59.92           C
    ATOM   3146  CE2 TYR B 103      70.363  -1.165 101.141  1.00 60.59           C
    ATOM   3147  CZ  TYR B 103      69.951  -1.926 102.248  1.00 61.86           C
    ATOM   3148  OH  TYR B 103      70.602  -3.083 102.560  1.00 57.96           O
    ATOM   3149  H   TYR B 103      67.483   4.540 100.711  1.00  0.00           H
    ATOM   3150  HA  TYR B 103      67.526   2.701 103.026  1.00  0.00           H
    ATOM   3151  HB2 TYR B 103      66.731   1.369 101.034  1.00  0.00           H
    ATOM   3152  HB3 TYR B 103      68.122   1.892 100.138  1.00  0.00           H
    ATOM   3153  HD1 TYR B 103      67.329   0.009 103.285  1.00  0.00           H
    ATOM   3154  HD2 TYR B 103      69.980   0.573  99.933  1.00  0.00           H
    ATOM   3155  HE1 TYR B 103      68.545  -2.070 103.884  1.00  0.00           H
    ATOM   3156  HE2 TYR B 103      71.189  -1.501 100.533  1.00  0.00           H
    ATOM   3157  HH  TYR B 103      71.311  -3.293 101.948  1.00  0.00           H
    ATOM   3158  N   GLN B 104      69.399   4.483 103.062  1.00 53.46           N
    ATOM   3159  CA  GLN B 104      70.701   4.850 103.598  1.00 55.32           C
    ATOM   3160  C   GLN B 104      71.275   3.545 104.169  1.00 55.30           C
    ATOM   3161  O   GLN B 104      70.786   3.083 105.198  1.00 59.81           O
    ATOM   3162  CB  GLN B 104      70.468   5.968 104.637  1.00 58.18           C
    ATOM   3163  CG  GLN B 104      71.744   6.455 105.343  1.00 63.49           C
    ATOM   3164  CD  GLN B 104      71.614   7.741 106.164  1.00 66.28           C
    ATOM   3165  OE1 GLN B 104      72.519   8.058 106.932  1.00 67.42           O
    ATOM   3166  NE2 GLN B 104      70.530   8.504 106.005  1.00 65.97           N
    ATOM   3167  H   GLN B 104      68.602   4.772 103.611  1.00  0.00           H
    ATOM   3168  HA  GLN B 104      71.339   5.223 102.794  1.00  0.00           H
    ATOM   3169  HB2 GLN B 104      70.001   6.799 104.105  1.00  0.00           H
    ATOM   3170  HB3 GLN B 104      69.746   5.643 105.388  1.00  0.00           H
    ATOM   3171  HG2 GLN B 104      72.126   5.668 105.992  1.00  0.00           H
    ATOM   3172  HG3 GLN B 104      72.495   6.655 104.588  1.00  0.00           H
    ATOM   3173 HE21 GLN B 104      69.804   8.231 105.359  1.00  0.00           H
    ATOM   3174 HE22 GLN B 104      70.425   9.351 106.543  1.00  0.00           H
    ATOM   3175  N   GLY B 105      72.106   2.870 103.362  1.00 52.97           N
    ATOM   3176  CA  GLY B 105      72.372   1.450 103.552  1.00 50.70           C
    ATOM   3177  C   GLY B 105      73.508   1.268 104.547  1.00 47.39           C
    ATOM   3178  O   GLY B 105      73.893   2.194 105.260  1.00 47.53           O
    ATOM   3179  H   GLY B 105      72.396   3.279 102.486  1.00  0.00           H
    ATOM   3180  HA2 GLY B 105      71.485   0.911 103.890  1.00  0.00           H
    ATOM   3181  HA3 GLY B 105      72.656   1.023 102.594  1.00  0.00           H
    ATOM   3182  N   SER B 106      74.093   0.066 104.535  1.00 43.59           N
    ATOM   3183  CA  SER B 106      75.324  -0.290 105.246  1.00 48.56           C
    ATOM   3184  C   SER B 106      76.544   0.632 104.984  1.00 48.12           C
    ATOM   3185  O   SER B 106      77.545   0.488 105.682  1.00 46.10           O
    ATOM   3186  CB  SER B 106      75.604  -1.786 104.985  1.00 55.95           C
    ATOM   3187  OG  SER B 106      75.560  -2.097 103.604  1.00 62.34           O
    ATOM   3188  H   SER B 106      73.743  -0.649 103.914  1.00  0.00           H
    ATOM   3189  HA  SER B 106      75.111  -0.184 106.311  1.00  0.00           H
    ATOM   3190  HB2 SER B 106      76.573  -2.082 105.388  1.00  0.00           H
    ATOM   3191  HB3 SER B 106      74.858  -2.396 105.494  1.00  0.00           H
    ATOM   3192  HG  SER B 106      75.769  -3.028 103.485  1.00  0.00           H
    ATOM   3193  N   TYR B 107      76.427   1.586 104.041  1.00 42.26           N
    ATOM   3194  CA  TYR B 107      77.418   2.619 103.737  1.00 31.97           C
    ATOM   3195  C   TYR B 107      77.003   4.027 104.154  1.00 34.42           C
    ATOM   3196  O   TYR B 107      77.861   4.904 104.125  1.00 35.32           O
    ATOM   3197  CB  TYR B 107      77.724   2.590 102.235  1.00 28.59           C
    ATOM   3198  CG  TYR B 107      78.249   1.233 101.852  1.00 29.17           C
    ATOM   3199  CD1 TYR B 107      79.561   0.867 102.208  1.00 34.87           C
    ATOM   3200  CD2 TYR B 107      77.381   0.289 101.271  1.00 29.17           C
    ATOM   3201  CE1 TYR B 107      80.009  -0.444 101.985  1.00 33.97           C
    ATOM   3202  CE2 TYR B 107      77.833  -1.020 101.043  1.00 30.44           C
    ATOM   3203  CZ  TYR B 107      79.145  -1.384 101.397  1.00 30.92           C
    ATOM   3204  OH  TYR B 107      79.579  -2.655 101.165  1.00 37.28           O
    ATOM   3205  H   TYR B 107      75.570   1.633 103.510  1.00  0.00           H
    ATOM   3206  HA  TYR B 107      78.341   2.407 104.273  1.00  0.00           H
    ATOM   3207  HB2 TYR B 107      76.833   2.822 101.649  1.00  0.00           H
    ATOM   3208  HB3 TYR B 107      78.468   3.341 101.966  1.00  0.00           H
    ATOM   3209  HD1 TYR B 107      80.218   1.567 102.692  1.00  0.00           H
    ATOM   3210  HD2 TYR B 107      76.362   0.554 101.029  1.00  0.00           H
    ATOM   3211  HE1 TYR B 107      81.015  -0.720 102.266  1.00  0.00           H
    ATOM   3212  HE2 TYR B 107      77.166  -1.748 100.609  1.00  0.00           H
    ATOM   3213  HH  TYR B 107      80.491  -2.794 101.428  1.00  0.00           H
    ATOM   3214  N   GLY B 108      75.736   4.236 104.538  1.00 31.82           N
    ATOM   3215  CA  GLY B 108      75.227   5.525 104.984  1.00 26.70           C
    ATOM   3216  C   GLY B 108      75.091   6.495 103.806  1.00 31.87           C
    ATOM   3217  O   GLY B 108      75.642   7.591 103.872  1.00 34.91           O
    ATOM   3218  H   GLY B 108      75.101   3.450 104.603  1.00  0.00           H
    ATOM   3219  HA2 GLY B 108      74.263   5.364 105.457  1.00  0.00           H
    ATOM   3220  HA3 GLY B 108      75.882   5.947 105.746  1.00  0.00           H
    ATOM   3221  N   PHE B 109      74.388   6.082 102.734  1.00 38.72           N
    ATOM   3222  CA  PHE B 109      74.170   6.876 101.522  1.00 39.32           C
    ATOM   3223  C   PHE B 109      73.215   8.051 101.759  1.00 36.46           C
    ATOM   3224  O   PHE B 109      72.093   7.851 102.223  1.00 41.46           O
    ATOM   3225  CB  PHE B 109      73.665   5.973 100.378  1.00 47.06           C
    ATOM   3226  CG  PHE B 109      73.473   6.661  99.032  1.00 48.27           C
    ATOM   3227  CD1 PHE B 109      74.557   7.312  98.405  1.00 48.79           C
    ATOM   3228  CD2 PHE B 109      72.224   6.631  98.379  1.00 46.86           C
    ATOM   3229  CE1 PHE B 109      74.379   7.915  97.168  1.00 49.44           C
    ATOM   3230  CE2 PHE B 109      72.063   7.247  97.145  1.00 45.04           C
    ATOM   3231  CZ  PHE B 109      73.140   7.883  96.543  1.00 48.94           C
    ATOM   3232  H   PHE B 109      73.969   5.165 102.747  1.00  0.00           H
    ATOM   3233  HA  PHE B 109      75.144   7.266 101.225  1.00  0.00           H
    ATOM   3234  HB2 PHE B 109      74.380   5.164 100.222  1.00  0.00           H
    ATOM   3235  HB3 PHE B 109      72.731   5.490 100.674  1.00  0.00           H
    ATOM   3236  HD1 PHE B 109      75.529   7.340  98.875  1.00  0.00           H
    ATOM   3237  HD2 PHE B 109      71.388   6.132  98.840  1.00  0.00           H
    ATOM   3238  HE1 PHE B 109      75.209   8.408  96.688  1.00  0.00           H
    ATOM   3239  HE2 PHE B 109      71.101   7.229  96.654  1.00  0.00           H
    ATOM   3240  HZ  PHE B 109      73.016   8.367  95.589  1.00  0.00           H
    ATOM   3241  N   ARG B 110      73.691   9.253 101.430  1.00 31.12           N
    ATOM   3242  CA  ARG B 110      72.968  10.511 101.548  1.00 34.13           C
    ATOM   3243  C   ARG B 110      73.344  11.381 100.350  1.00 31.57           C
    ATOM   3244  O   ARG B 110      74.449  11.256  99.818  1.00 30.51           O
    ATOM   3245  CB  ARG B 110      73.334  11.186 102.885  1.00 43.57           C
    ATOM   3246  CG  ARG B 110      72.742  10.480 104.128  1.00 53.06           C
    ATOM   3247  CD  ARG B 110      73.587  10.672 105.382  1.00 60.55           C
    ATOM   3248  NE  ARG B 110      74.884  10.008 105.207  1.00 68.40           N
    ATOM   3249  CZ  ARG B 110      76.082  10.555 105.443  1.00 69.02           C
    ATOM   3250  NH1 ARG B 110      76.216  11.743 106.045  1.00 70.00           N
    ATOM   3251  NH2 ARG B 110      77.164   9.901 105.030  1.00 63.10           N1+
    ATOM   3252  H   ARG B 110      74.638   9.333 101.079  1.00  0.00           H
    ATOM   3253  HA  ARG B 110      71.894  10.326 101.515  1.00  0.00           H
    ATOM   3254  HB2 ARG B 110      74.421  11.227 102.949  1.00  0.00           H
    ATOM   3255  HB3 ARG B 110      73.003  12.226 102.894  1.00  0.00           H
    ATOM   3256  HG2 ARG B 110      71.683  10.684 104.284  1.00  0.00           H
    ATOM   3257  HG3 ARG B 110      72.797   9.411 103.971  1.00  0.00           H
    ATOM   3258  HD2 ARG B 110      73.793  11.740 105.423  1.00  0.00           H
    ATOM   3259  HD3 ARG B 110      73.075  10.448 106.316  1.00  0.00           H
    ATOM   3260  HE  ARG B 110      74.863   9.073 104.818  1.00  0.00           H
    ATOM   3261 HH11 ARG B 110      75.401  12.245 106.364  1.00  0.00           H
    ATOM   3262 HH12 ARG B 110      77.137  12.124 106.210  1.00  0.00           H
    ATOM   3263 HH21 ARG B 110      77.065   8.995 104.594  1.00  0.00           H
    ATOM   3264 HH22 ARG B 110      78.090  10.288 105.183  1.00  0.00           H
    ATOM   3265  N   LEU B 111      72.410  12.253  99.949  1.00 27.19           N
    ATOM   3266  CA  LEU B 111      72.638  13.206  98.871  1.00 28.03           C
    ATOM   3267  C   LEU B 111      72.816  14.601  99.469  1.00 30.47           C
    ATOM   3268  O   LEU B 111      72.092  14.977 100.392  1.00 32.36           O
    ATOM   3269  CB  LEU B 111      71.479  13.217  97.856  1.00 27.56           C
    ATOM   3270  CG  LEU B 111      71.099  11.853  97.255  1.00 21.80           C
    ATOM   3271  CD1 LEU B 111      69.844  12.007  96.384  1.00 21.14           C
    ATOM   3272  CD2 LEU B 111      72.241  11.193  96.471  1.00 19.08           C
    ATOM   3273  H   LEU B 111      71.526  12.321 100.433  1.00  0.00           H
    ATOM   3274  HA  LEU B 111      73.532  12.923  98.330  1.00  0.00           H
    ATOM   3275  HB2 LEU B 111      70.595  13.626  98.347  1.00  0.00           H
    ATOM   3276  HB3 LEU B 111      71.716  13.912  97.048  1.00  0.00           H
    ATOM   3277  HG  LEU B 111      70.839  11.186  98.074  1.00  0.00           H
    ATOM   3278 HD11 LEU B 111      69.499  11.045  96.003  1.00  0.00           H
    ATOM   3279 HD12 LEU B 111      69.020  12.448  96.946  1.00  0.00           H
    ATOM   3280 HD13 LEU B 111      70.039  12.649  95.524  1.00  0.00           H
    ATOM   3281 HD21 LEU B 111      71.881  10.604  95.630  1.00  0.00           H
    ATOM   3282 HD22 LEU B 111      72.932  11.940  96.079  1.00  0.00           H
    ATOM   3283 HD23 LEU B 111      72.811  10.533  97.124  1.00  0.00           H
    ATOM   3284  N   GLY B 112      73.758  15.342  98.885  1.00 26.45           N
    ATOM   3285  CA  GLY B 112      73.988  16.746  99.167  1.00 22.81           C
    ATOM   3286  C   GLY B 112      73.612  17.523  97.910  1.00 22.74           C
    ATOM   3287  O   GLY B 112      73.626  17.000  96.793  1.00 25.47           O
    ATOM   3288  H   GLY B 112      74.293  14.940  98.125  1.00  0.00           H
    ATOM   3289  HA2 GLY B 112      73.417  17.104 100.026  1.00  0.00           H
    ATOM   3290  HA3 GLY B 112      75.043  16.906  99.388  1.00  0.00           H
    ATOM   3291  N   PHE B 113      73.290  18.800  98.116  1.00 18.52           N
    ATOM   3292  CA  PHE B 113      72.975  19.745  97.058  1.00 22.94           C
    ATOM   3293  C   PHE B 113      73.678  21.053  97.388  1.00 30.87           C
    ATOM   3294  O   PHE B 113      73.859  21.380  98.563  1.00 36.54           O
    ATOM   3295  CB  PHE B 113      71.451  19.934  96.951  1.00 21.85           C
    ATOM   3296  CG  PHE B 113      70.704  18.691  96.506  1.00 21.16           C
    ATOM   3297  CD1 PHE B 113      70.340  17.712  97.456  1.00 23.33           C
    ATOM   3298  CD2 PHE B 113      70.527  18.410  95.135  1.00 18.49           C
    ATOM   3299  CE1 PHE B 113      69.785  16.510  97.044  1.00 24.29           C
    ATOM   3300  CE2 PHE B 113      69.952  17.206  94.748  1.00 19.73           C
    ATOM   3301  CZ  PHE B 113      69.589  16.261  95.695  1.00 21.39           C
    ATOM   3302  H   PHE B 113      73.314  19.184  99.050  1.00  0.00           H
    ATOM   3303  HA  PHE B 113      73.361  19.391  96.100  1.00  0.00           H
    ATOM   3304  HB2 PHE B 113      71.052  20.260  97.912  1.00  0.00           H
    ATOM   3305  HB3 PHE B 113      71.232  20.734  96.245  1.00  0.00           H
    ATOM   3306  HD1 PHE B 113      70.504  17.888  98.509  1.00  0.00           H
    ATOM   3307  HD2 PHE B 113      70.835  19.119  94.382  1.00  0.00           H
    ATOM   3308  HE1 PHE B 113      69.509  15.765  97.776  1.00  0.00           H
    ATOM   3309  HE2 PHE B 113      69.804  16.971  93.709  1.00  0.00           H
    ATOM   3310  HZ  PHE B 113      69.167  15.319  95.382  1.00  0.00           H
    ATOM   3311  N   LEU B 114      74.055  21.786  96.334  1.00 33.52           N
    ATOM   3312  CA  LEU B 114      74.722  23.071  96.477  1.00 33.79           C
    ATOM   3313  C   LEU B 114      73.764  24.145  96.975  1.00 36.52           C
    ATOM   3314  O   LEU B 114      72.546  24.021  96.829  1.00 38.31           O
    ATOM   3315  CB  LEU B 114      75.326  23.534  95.140  1.00 34.75           C
    ATOM   3316  CG  LEU B 114      76.197  22.506  94.404  1.00 33.80           C
    ATOM   3317  CD1 LEU B 114      77.087  23.216  93.375  1.00 32.12           C
    ATOM   3318  CD2 LEU B 114      77.044  21.627  95.329  1.00 35.46           C
    ATOM   3319  H   LEU B 114      73.862  21.471  95.392  1.00  0.00           H
    ATOM   3320  HA  LEU B 114      75.515  22.968  97.220  1.00  0.00           H
    ATOM   3321  HB2 LEU B 114      74.544  23.880  94.464  1.00  0.00           H
    ATOM   3322  HB3 LEU B 114      75.938  24.414  95.346  1.00  0.00           H
    ATOM   3323  HG  LEU B 114      75.524  21.848  93.855  1.00  0.00           H
    ATOM   3324 HD11 LEU B 114      77.297  22.564  92.528  1.00  0.00           H
    ATOM   3325 HD12 LEU B 114      76.618  24.120  92.984  1.00  0.00           H
    ATOM   3326 HD13 LEU B 114      78.047  23.507  93.805  1.00  0.00           H
    ATOM   3327 HD21 LEU B 114      77.983  21.347  94.852  1.00  0.00           H
    ATOM   3328 HD22 LEU B 114      77.303  22.121  96.264  1.00  0.00           H
    ATOM   3329 HD23 LEU B 114      76.515  20.703  95.561  1.00  0.00           H
    ATOM   3330  N   HIS B 115      74.365  25.208  97.514  1.00 39.03           N
    ATOM   3331  CA  HIS B 115      73.637  26.390  97.920  1.00 40.79           C
    ATOM   3332  C   HIS B 115      73.605  27.383  96.754  1.00 41.41           C
    ATOM   3333  O   HIS B 115      74.504  28.213  96.635  1.00 47.89           O
    ATOM   3334  CB  HIS B 115      74.294  26.961  99.180  1.00 40.63           C
    ATOM   3335  CG  HIS B 115      73.342  27.904  99.843  1.00 41.41           C
    ATOM   3336  ND1 HIS B 115      73.358  29.270  99.635  1.00 42.65           N
    ATOM   3337  CD2 HIS B 115      72.267  27.649 100.659  1.00 41.09           C
    ATOM   3338  CE1 HIS B 115      72.317  29.770 100.304  1.00 42.79           C
    ATOM   3339  NE2 HIS B 115      71.606  28.844 100.938  1.00 41.09           N
    ATOM   3340  H   HIS B 115      75.371  25.247  97.589  1.00  0.00           H
    ATOM   3341  HA  HIS B 115      72.610  26.118  98.178  1.00  0.00           H
    ATOM   3342  HB2 HIS B 115      74.516  26.161  99.888  1.00  0.00           H
    ATOM   3343  HB3 HIS B 115      75.244  27.455  98.971  1.00  0.00           H
    ATOM   3344  HD1 HIS B 115      74.010  29.786  99.060  1.00  0.00           H
    ATOM   3345  HD2 HIS B 115      71.904  26.707 101.042  1.00  0.00           H
    ATOM   3346  HE1 HIS B 115      72.064  30.821 100.314  1.00  0.00           H
    ATOM   3347  N   SER B 116      72.603  27.247  95.877  1.00 40.64           N
    ATOM   3348  CA  SER B 116      72.595  27.966  94.605  1.00 38.62           C
    ATOM   3349  C   SER B 116      72.205  29.444  94.657  1.00 40.40           C
    ATOM   3350  O   SER B 116      72.660  30.194  93.795  1.00 43.46           O
    ATOM   3351  CB  SER B 116      71.692  27.235  93.622  1.00 38.97           C
    ATOM   3352  OG  SER B 116      72.410  26.146  93.099  1.00 43.62           O
    ATOM   3353  H   SER B 116      71.902  26.530  96.004  1.00  0.00           H
    ATOM   3354  HA  SER B 116      73.610  27.954  94.198  1.00  0.00           H
    ATOM   3355  HB2 SER B 116      70.783  26.903  94.119  1.00  0.00           H
    ATOM   3356  HB3 SER B 116      71.396  27.876  92.790  1.00  0.00           H
    ATOM   3357  HG  SER B 116      71.973  25.860  92.290  1.00  0.00           H
    ATOM   3358  N   GLY B 117      71.348  29.839  95.607  1.00 45.31           N
    ATOM   3359  CA  GLY B 117      70.725  31.158  95.581  1.00 47.96           C
    ATOM   3360  C   GLY B 117      69.446  31.074  94.744  1.00 52.54           C
    ATOM   3361  O   GLY B 117      69.128  30.034  94.160  1.00 56.77           O
    ATOM   3362  H   GLY B 117      71.012  29.174  96.290  1.00  0.00           H
    ATOM   3363  HA2 GLY B 117      70.475  31.443  96.604  1.00  0.00           H
    ATOM   3364  HA3 GLY B 117      71.391  31.926  95.184  1.00  0.00           H
    ATOM   3365  N   THR B 118      68.685  32.175  94.731  1.00 54.50           N
    ATOM   3366  CA  THR B 118      67.314  32.161  94.232  1.00 51.61           C
    ATOM   3367  C   THR B 118      67.049  33.226  93.160  1.00 42.01           C
    ATOM   3368  O   THR B 118      65.891  33.552  92.938  1.00 41.20           O
    ATOM   3369  CB  THR B 118      66.287  32.273  95.395  1.00 54.31           C
    ATOM   3370  OG1 THR B 118      66.659  31.435  96.470  1.00 57.97           O
    ATOM   3371  CG2 THR B 118      64.876  31.810  95.030  1.00 51.15           C
    ATOM   3372  H   THR B 118      68.989  33.015  95.202  1.00  0.00           H
    ATOM   3373  HA  THR B 118      67.101  31.226  93.711  1.00  0.00           H
    ATOM   3374  HB  THR B 118      66.250  33.296  95.773  1.00  0.00           H
    ATOM   3375 HG21 THR B 118      64.254  31.727  95.921  1.00  0.00           H
    ATOM   3376 HG22 THR B 118      64.394  32.498  94.352  1.00  0.00           H
    ATOM   3377 HG23 THR B 118      64.887  30.828  94.556  1.00  0.00           H
    ATOM   3378  HG1 THR B 118      67.500  31.739  96.819  1.00  0.00           H
    ATOM   3379  N   ALA B 119      68.085  33.716  92.449  1.00 35.43           N
    ATOM   3380  CA  ALA B 119      67.907  34.625  91.307  1.00 35.89           C
    ATOM   3381  C   ALA B 119      66.941  34.035  90.285  1.00 37.91           C
    ATOM   3382  O   ALA B 119      66.984  32.826  90.071  1.00 39.07           O
    ATOM   3383  CB  ALA B 119      69.251  34.894  90.619  1.00 34.91           C
    ATOM   3384  H   ALA B 119      69.026  33.401  92.634  1.00  0.00           H
    ATOM   3385  HA  ALA B 119      67.505  35.567  91.683  1.00  0.00           H
    ATOM   3386  HB1 ALA B 119      69.137  35.526  89.738  1.00  0.00           H
    ATOM   3387  HB2 ALA B 119      69.939  35.404  91.293  1.00  0.00           H
    ATOM   3388  HB3 ALA B 119      69.729  33.969  90.294  1.00  0.00           H
    ATOM   3389  N   LYS B 120      66.078  34.875  89.701  1.00 42.17           N
    ATOM   3390  CA  LYS B 120      65.118  34.417  88.709  1.00 40.58           C
    ATOM   3391  C   LYS B 120      65.869  33.898  87.481  1.00 48.00           C
    ATOM   3392  O   LYS B 120      66.644  34.647  86.887  1.00 49.75           O
    ATOM   3393  CB  LYS B 120      64.139  35.545  88.384  1.00 38.39           C
    ATOM   3394  CG  LYS B 120      63.106  35.166  87.317  1.00 41.12           C
    ATOM   3395  CD  LYS B 120      61.931  36.137  87.297  1.00 40.85           C
    ATOM   3396  CE  LYS B 120      61.057  36.008  86.051  1.00 40.97           C
    ATOM   3397  NZ  LYS B 120      61.676  36.734  84.975  1.00 44.78           N1+
    ATOM   3398  H   LYS B 120      66.122  35.867  89.886  1.00  0.00           H
    ATOM   3399  HA  LYS B 120      64.541  33.600  89.147  1.00  0.00           H
    ATOM   3400  HB2 LYS B 120      63.618  35.822  89.301  1.00  0.00           H
    ATOM   3401  HB3 LYS B 120      64.693  36.426  88.073  1.00  0.00           H
    ATOM   3402  HG2 LYS B 120      63.590  35.164  86.339  1.00  0.00           H
    ATOM   3403  HG3 LYS B 120      62.738  34.153  87.477  1.00  0.00           H
    ATOM   3404  HD2 LYS B 120      61.331  35.907  88.173  1.00  0.00           H
    ATOM   3405  HD3 LYS B 120      62.267  37.168  87.420  1.00  0.00           H
    ATOM   3406  HE2 LYS B 120      60.891  34.967  85.778  1.00  0.00           H
    ATOM   3407  HE3 LYS B 120      60.079  36.456  86.233  1.00  0.00           H
    ATOM   3408  HZ1 LYS B 120      61.690  37.721  85.199  1.00  0.00           H
    ATOM   3409  HZ2 LYS B 120      61.116  36.663  84.136  1.00  0.00           H
    ATOM   3410  HZ3 LYS B 120      62.602  36.352  84.842  1.00 44.78           H
    ATOM   3411  N   SER B 121      65.642  32.613  87.177  1.00 45.68           N
    ATOM   3412  CA  SER B 121      66.325  31.862  86.121  1.00 36.99           C
    ATOM   3413  C   SER B 121      67.684  31.322  86.536  1.00 36.69           C
    ATOM   3414  O   SER B 121      68.492  31.037  85.653  1.00 40.36           O
    ATOM   3415  CB  SER B 121      66.370  32.617  84.775  1.00 35.41           C
    ATOM   3416  OG  SER B 121      65.090  33.106  84.442  1.00 41.04           O
    ATOM   3417  H   SER B 121      64.990  32.087  87.738  1.00  0.00           H
    ATOM   3418  HA  SER B 121      65.790  30.940  85.947  1.00  0.00           H
    ATOM   3419  HB2 SER B 121      67.087  33.437  84.777  1.00  0.00           H
    ATOM   3420  HB3 SER B 121      66.683  31.946  83.975  1.00  0.00           H
    ATOM   3421  HG  SER B 121      64.940  33.911  84.946  1.00  0.00           H
    ATOM   3422  N   VAL B 122      67.900  31.096  87.844  1.00 34.50           N
    ATOM   3423  CA  VAL B 122      69.067  30.354  88.309  1.00 28.99           C
    ATOM   3424  C   VAL B 122      69.042  28.934  87.711  1.00 29.65           C
    ATOM   3425  O   VAL B 122      67.981  28.327  87.610  1.00 36.99           O
    ATOM   3426  CB  VAL B 122      69.162  30.298  89.855  1.00 23.76           C
    ATOM   3427  CG1 VAL B 122      68.061  29.455  90.517  1.00 22.80           C
    ATOM   3428  CG2 VAL B 122      70.551  29.841  90.330  1.00 23.49           C
    ATOM   3429  H   VAL B 122      67.231  31.385  88.544  1.00  0.00           H
    ATOM   3430  HA  VAL B 122      69.950  30.878  87.935  1.00  0.00           H
    ATOM   3431  HB  VAL B 122      69.045  31.319  90.217  1.00  0.00           H
    ATOM   3432 HG11 VAL B 122      68.072  29.585  91.599  1.00  0.00           H
    ATOM   3433 HG12 VAL B 122      67.080  29.767  90.169  1.00  0.00           H
    ATOM   3434 HG13 VAL B 122      68.165  28.394  90.294  1.00  0.00           H
    ATOM   3435 HG21 VAL B 122      70.628  29.901  91.416  1.00  0.00           H
    ATOM   3436 HG22 VAL B 122      70.765  28.811  90.046  1.00  0.00           H
    ATOM   3437 HG23 VAL B 122      71.339  30.472  89.916  1.00  0.00           H
    ATOM   3438  N   THR B 123      70.201  28.494  87.225  1.00 23.62           N
    ATOM   3439  CA  THR B 123      70.369  27.309  86.389  1.00 27.97           C
    ATOM   3440  C   THR B 123      69.899  25.999  87.057  1.00 30.25           C
    ATOM   3441  O   THR B 123      69.280  25.161  86.401  1.00 33.83           O
    ATOM   3442  CB  THR B 123      71.874  27.224  86.006  1.00 29.79           C
    ATOM   3443  OG1 THR B 123      71.998  27.588  84.650  1.00 35.94           O
    ATOM   3444  CG2 THR B 123      72.658  25.923  86.284  1.00 30.56           C
    ATOM   3445  H   THR B 123      71.022  29.073  87.329  1.00  0.00           H
    ATOM   3446  HA  THR B 123      69.755  27.450  85.500  1.00  0.00           H
    ATOM   3447  HB  THR B 123      72.424  28.000  86.541  1.00  0.00           H
    ATOM   3448 HG21 THR B 123      73.656  25.966  85.846  1.00  0.00           H
    ATOM   3449 HG22 THR B 123      72.798  25.769  87.355  1.00  0.00           H
    ATOM   3450 HG23 THR B 123      72.170  25.033  85.893  1.00  0.00           H
    ATOM   3451  HG1 THR B 123      72.913  27.463  84.385  1.00  0.00           H
    ATOM   3452  N   CYS B 124      70.205  25.880  88.350  1.00 32.16           N
    ATOM   3453  CA  CYS B 124      69.860  24.771  89.213  1.00 27.24           C
    ATOM   3454  C   CYS B 124      69.764  25.364  90.613  1.00 29.00           C
    ATOM   3455  O   CYS B 124      70.584  26.208  90.969  1.00 27.62           O
    ATOM   3456  CB  CYS B 124      70.876  23.615  89.118  1.00 22.07           C
    ATOM   3457  SG  CYS B 124      70.396  22.207  90.158  1.00 28.28           S
    ATOM   3458  H   CYS B 124      70.692  26.636  88.809  1.00  0.00           H
    ATOM   3459  HA  CYS B 124      68.880  24.397  88.921  1.00  0.00           H
    ATOM   3460  HB2 CYS B 124      70.926  23.263  88.089  1.00  0.00           H
    ATOM   3461  HB3 CYS B 124      71.877  23.943  89.399  1.00  0.00           H
    ATOM   3462  HG  CYS B 124      69.296  21.888  89.469  1.00  0.00           H
    ATOM   3463  N   THR B 125      68.770  24.923  91.381  1.00 31.69           N
    ATOM   3464  CA  THR B 125      68.597  25.353  92.751  1.00 32.99           C
    ATOM   3465  C   THR B 125      67.784  24.308  93.492  1.00 30.99           C
    ATOM   3466  O   THR B 125      66.848  23.736  92.939  1.00 29.86           O
    ATOM   3467  CB  THR B 125      67.932  26.750  92.869  1.00 36.82           C
    ATOM   3468  OG1 THR B 125      68.068  27.263  94.175  1.00 40.87           O
    ATOM   3469  CG2 THR B 125      66.436  26.807  92.513  1.00 35.49           C
    ATOM   3470  H   THR B 125      68.092  24.264  91.022  1.00  0.00           H
    ATOM   3471  HA  THR B 125      69.583  25.369  93.213  1.00  0.00           H
    ATOM   3472  HB  THR B 125      68.484  27.436  92.235  1.00  0.00           H
    ATOM   3473 HG21 THR B 125      66.061  27.829  92.579  1.00  0.00           H
    ATOM   3474 HG22 THR B 125      66.286  26.485  91.484  1.00  0.00           H
    ATOM   3475 HG23 THR B 125      65.814  26.197  93.166  1.00  0.00           H
    ATOM   3476  HG1 THR B 125      68.529  28.107  94.123  1.00  0.00           H
    ATOM   3477  N   TYR B 126      68.186  24.077  94.735  1.00 33.42           N
    ATOM   3478  CA  TYR B 126      67.605  23.057  95.567  1.00 30.38           C
    ATOM   3479  C   TYR B 126      66.904  23.703  96.755  1.00 40.22           C
    ATOM   3480  O   TYR B 126      67.443  24.635  97.349  1.00 42.59           O
    ATOM   3481  CB  TYR B 126      68.725  22.124  96.006  1.00 26.30           C
    ATOM   3482  CG  TYR B 126      68.196  21.039  96.899  1.00 26.01           C
    ATOM   3483  CD1 TYR B 126      67.451  19.980  96.340  1.00 25.23           C
    ATOM   3484  CD2 TYR B 126      68.389  21.132  98.288  1.00 22.72           C
    ATOM   3485  CE1 TYR B 126      66.921  18.986  97.179  1.00 21.65           C
    ATOM   3486  CE2 TYR B 126      67.865  20.136  99.122  1.00 21.76           C
    ATOM   3487  CZ  TYR B 126      67.147  19.066  98.568  1.00 25.79           C
    ATOM   3488  OH  TYR B 126      66.701  18.104  99.412  1.00 29.47           O
    ATOM   3489  H   TYR B 126      68.931  24.624  95.141  1.00  0.00           H
    ATOM   3490  HA  TYR B 126      66.877  22.463  95.014  1.00  0.00           H
    ATOM   3491  HB2 TYR B 126      69.195  21.664  95.136  1.00  0.00           H
    ATOM   3492  HB3 TYR B 126      69.513  22.675  96.525  1.00  0.00           H
    ATOM   3493  HD1 TYR B 126      67.289  19.926  95.273  1.00  0.00           H
    ATOM   3494  HD2 TYR B 126      68.941  21.956  98.716  1.00  0.00           H
    ATOM   3495  HE1 TYR B 126      66.363  18.162  96.758  1.00  0.00           H
    ATOM   3496  HE2 TYR B 126      68.036  20.185 100.187  1.00  0.00           H
    ATOM   3497  HH  TYR B 126      66.191  17.402  98.989  1.00  0.00           H
    ATOM   3498  N   SER B 127      65.734  23.150  97.082  1.00 48.58           N
    ATOM   3499  CA  SER B 127      64.854  23.639  98.120  1.00 43.81           C
    ATOM   3500  C   SER B 127      64.807  22.601  99.254  1.00 37.26           C
    ATOM   3501  O   SER B 127      64.158  21.577  99.066  1.00 34.58           O
    ATOM   3502  CB  SER B 127      63.504  23.880  97.440  1.00 47.11           C
    ATOM   3503  OG  SER B 127      62.769  24.833  98.162  1.00 51.86           O
    ATOM   3504  H   SER B 127      65.379  22.376  96.534  1.00  0.00           H
    ATOM   3505  HA  SER B 127      65.192  24.601  98.486  1.00  0.00           H
    ATOM   3506  HB2 SER B 127      63.648  24.278  96.439  1.00  0.00           H
    ATOM   3507  HB3 SER B 127      62.964  22.952  97.273  1.00  0.00           H
    ATOM   3508  HG  SER B 127      61.915  24.959  97.740  1.00  0.00           H
    ATOM   3509  N   PRO B 128      65.524  22.846 100.381  1.00 36.09           N
    ATOM   3510  CA  PRO B 128      65.635  21.892 101.508  1.00 34.25           C
    ATOM   3511  C   PRO B 128      64.298  21.456 102.122  1.00 35.81           C
    ATOM   3512  O   PRO B 128      64.118  20.270 102.396  1.00 37.43           O
    ATOM   3513  CB  PRO B 128      66.495  22.634 102.551  1.00 32.59           C
    ATOM   3514  CG  PRO B 128      67.243  23.691 101.766  1.00 33.32           C
    ATOM   3515  CD  PRO B 128      66.259  24.076 100.679  1.00 31.23           C
    ATOM   3516  HA  PRO B 128      66.168  21.011 101.153  1.00  0.00           H
    ATOM   3517  HB2 PRO B 128      65.871  23.136 103.294  1.00  0.00           H
    ATOM   3518  HB3 PRO B 128      67.167  21.967 103.094  1.00  0.00           H
    ATOM   3519  HG2 PRO B 128      67.586  24.532 102.369  1.00  0.00           H
    ATOM   3520  HG3 PRO B 128      68.118  23.233 101.303  1.00  0.00           H
    ATOM   3521  HD2 PRO B 128      65.553  24.822 101.049  1.00  0.00           H
    ATOM   3522  HD3 PRO B 128      66.811  24.495  99.842  1.00  0.00           H
    ATOM   3523  N   ALA B 129      63.396  22.434 102.307  1.00 39.32           N
    ATOM   3524  CA  ALA B 129      62.053  22.265 102.859  1.00 44.70           C
    ATOM   3525  C   ALA B 129      61.130  21.441 101.953  1.00 48.92           C
    ATOM   3526  O   ALA B 129      60.366  20.612 102.445  1.00 51.51           O
    ATOM   3527  CB  ALA B 129      61.449  23.657 103.089  1.00 44.94           C
    ATOM   3528  H   ALA B 129      63.643  23.379 102.049  1.00  0.00           H
    ATOM   3529  HA  ALA B 129      62.139  21.753 103.819  1.00  0.00           H
    ATOM   3530  HB1 ALA B 129      60.446  23.590 103.513  1.00  0.00           H
    ATOM   3531  HB2 ALA B 129      62.057  24.235 103.786  1.00  0.00           H
    ATOM   3532  HB3 ALA B 129      61.379  24.227 102.161  1.00  0.00           H
    ATOM   3533  N   LEU B 130      61.240  21.694 100.644  1.00 42.80           N
    ATOM   3534  CA  LEU B 130      60.488  21.009  99.604  1.00 37.44           C
    ATOM   3535  C   LEU B 130      61.096  19.647  99.231  1.00 33.27           C
    ATOM   3536  O   LEU B 130      60.383  18.821  98.666  1.00 33.59           O
    ATOM   3537  CB  LEU B 130      60.458  21.927  98.362  1.00 45.22           C
    ATOM   3538  CG  LEU B 130      59.163  22.703  98.103  1.00 52.22           C
    ATOM   3539  CD1 LEU B 130      57.938  21.774  98.136  1.00 56.06           C
    ATOM   3540  CD2 LEU B 130      59.022  23.942  99.003  1.00 55.27           C
    ATOM   3541  H   LEU B 130      61.899  22.391 100.329  1.00  0.00           H
    ATOM   3542  HA  LEU B 130      59.481  20.819  99.974  1.00  0.00           H
    ATOM   3543  HB2 LEU B 130      61.283  22.624  98.397  1.00  0.00           H
    ATOM   3544  HB3 LEU B 130      60.606  21.326  97.477  1.00  0.00           H
    ATOM   3545  HG  LEU B 130      59.240  23.077  97.080  1.00  0.00           H
    ATOM   3546 HD11 LEU B 130      57.190  22.094  97.414  1.00  0.00           H
    ATOM   3547 HD12 LEU B 130      58.197  20.742  97.892  1.00  0.00           H
    ATOM   3548 HD13 LEU B 130      57.454  21.770  99.112  1.00  0.00           H
    ATOM   3549 HD21 LEU B 130      58.126  23.904  99.623  1.00  0.00           H
    ATOM   3550 HD22 LEU B 130      59.873  24.068  99.672  1.00  0.00           H
    ATOM   3551 HD23 LEU B 130      58.953  24.848  98.399  1.00  0.00           H
    ATOM   3552  N   ASN B 131      62.404  19.468  99.488  1.00 31.93           N
    ATOM   3553  CA  ASN B 131      63.271  18.413  98.955  1.00 31.05           C
    ATOM   3554  C   ASN B 131      63.198  18.375  97.416  1.00 25.96           C
    ATOM   3555  O   ASN B 131      62.915  17.330  96.826  1.00 25.40           O
    ATOM   3556  CB  ASN B 131      62.975  17.051  99.632  1.00 33.58           C
    ATOM   3557  CG  ASN B 131      63.904  15.916  99.165  1.00 35.92           C
    ATOM   3558  OD1 ASN B 131      65.058  16.141  98.811  1.00 39.83           O
    ATOM   3559  ND2 ASN B 131      63.392  14.690  99.108  1.00 35.42           N
    ATOM   3560  H   ASN B 131      62.896  20.212  99.962  1.00  0.00           H
    ATOM   3561  HA  ASN B 131      64.274  18.730  99.228  1.00  0.00           H
    ATOM   3562  HB2 ASN B 131      63.092  17.150 100.711  1.00  0.00           H
    ATOM   3563  HB3 ASN B 131      61.939  16.757  99.458  1.00  0.00           H
    ATOM   3564 HD21 ASN B 131      62.428  14.522  99.359  1.00  0.00           H
    ATOM   3565 HD22 ASN B 131      63.965  13.919  98.792  1.00  0.00           H
    ATOM   3566  N   LYS B 132      63.377  19.545  96.797  1.00 25.72           N
    ATOM   3567  CA  LYS B 132      63.028  19.693  95.401  1.00 28.17           C
    ATOM   3568  C   LYS B 132      64.088  20.485  94.658  1.00 25.38           C
    ATOM   3569  O   LYS B 132      64.366  21.631  95.008  1.00 21.28           O
    ATOM   3570  CB  LYS B 132      61.637  20.314  95.319  1.00 32.17           C
    ATOM   3571  CG  LYS B 132      61.011  20.247  93.939  1.00 33.73           C
    ATOM   3572  CD  LYS B 132      59.594  20.813  93.938  1.00 37.93           C
    ATOM   3573  CE  LYS B 132      59.079  20.964  92.509  1.00 38.53           C
    ATOM   3574  NZ  LYS B 132      57.634  21.202  92.471  1.00 36.37           N1+
    ATOM   3575  H   LYS B 132      63.608  20.380  97.318  1.00  0.00           H
    ATOM   3576  HA  LYS B 132      62.960  18.722  94.920  1.00  0.00           H
    ATOM   3577  HB2 LYS B 132      60.971  19.768  95.989  1.00  0.00           H
    ATOM   3578  HB3 LYS B 132      61.678  21.346  95.659  1.00  0.00           H
    ATOM   3579  HG2 LYS B 132      61.621  20.814  93.236  1.00  0.00           H
    ATOM   3580  HG3 LYS B 132      61.016  19.221  93.584  1.00  0.00           H
    ATOM   3581  HD2 LYS B 132      58.942  20.159  94.515  1.00  0.00           H
    ATOM   3582  HD3 LYS B 132      59.563  21.772  94.452  1.00  0.00           H
    ATOM   3583  HE2 LYS B 132      59.593  21.792  92.022  1.00  0.00           H
    ATOM   3584  HE3 LYS B 132      59.288  20.061  91.936  1.00  0.00           H
    ATOM   3585  HZ1 LYS B 132      57.401  22.046  92.989  1.00  0.00           H
    ATOM   3586  HZ2 LYS B 132      57.317  21.295  91.518  1.00  0.00           H
    ATOM   3587  HZ3 LYS B 132      57.177  20.408  92.904  1.00  0.00           H
    ATOM   3588  N   MET B 133      64.622  19.848  93.612  1.00 24.43           N
    ATOM   3589  CA  MET B 133      65.460  20.494  92.625  1.00 25.30           C
    ATOM   3590  C   MET B 133      64.577  21.290  91.662  1.00 30.93           C
    ATOM   3591  O   MET B 133      63.563  20.784  91.198  1.00 35.18           O
    ATOM   3592  CB  MET B 133      66.281  19.425  91.895  1.00 24.14           C
    ATOM   3593  CG  MET B 133      67.259  20.023  90.875  1.00 28.94           C
    ATOM   3594  SD  MET B 133      68.322  18.785  90.115  1.00 30.05           S
    ATOM   3595  CE  MET B 133      69.547  18.588  91.429  1.00 31.51           C
    ATOM   3596  H   MET B 133      64.320  18.906  93.396  1.00  0.00           H
    ATOM   3597  HA  MET B 133      66.153  21.166  93.132  1.00  0.00           H
    ATOM   3598  HB2 MET B 133      66.832  18.841  92.632  1.00  0.00           H
    ATOM   3599  HB3 MET B 133      65.619  18.717  91.394  1.00  0.00           H
    ATOM   3600  HG2 MET B 133      66.715  20.511  90.066  1.00  0.00           H
    ATOM   3601  HG3 MET B 133      67.879  20.788  91.341  1.00  0.00           H
    ATOM   3602  HE1 MET B 133      70.199  17.747  91.205  1.00  0.00           H
    ATOM   3603  HE2 MET B 133      70.160  19.484  91.521  1.00  0.00           H
    ATOM   3604  HE3 MET B 133      69.044  18.419  92.375  1.00  0.00           H
    ATOM   3605  N   PHE B 134      64.988  22.517  91.362  1.00 31.89           N
    ATOM   3606  CA  PHE B 134      64.386  23.347  90.338  1.00 30.63           C
    ATOM   3607  C   PHE B 134      65.530  23.660  89.377  1.00 30.54           C
    ATOM   3608  O   PHE B 134      66.510  24.267  89.800  1.00 32.86           O
    ATOM   3609  CB  PHE B 134      63.757  24.621  90.941  1.00 38.73           C
    ATOM   3610  CG  PHE B 134      62.559  24.458  91.860  1.00 46.89           C
    ATOM   3611  CD1 PHE B 134      62.710  23.908  93.150  1.00 46.17           C
    ATOM   3612  CD2 PHE B 134      61.276  24.870  91.437  1.00 46.85           C
    ATOM   3613  CE1 PHE B 134      61.615  23.810  93.996  1.00 44.34           C
    ATOM   3614  CE2 PHE B 134      60.193  24.759  92.299  1.00 49.90           C
    ATOM   3615  CZ  PHE B 134      60.364  24.236  93.573  1.00 47.93           C
    ATOM   3616  H   PHE B 134      65.807  22.902  91.817  1.00  0.00           H
    ATOM   3617  HA  PHE B 134      63.599  22.816  89.817  1.00  0.00           H
    ATOM   3618  HB2 PHE B 134      64.504  25.188  91.486  1.00  0.00           H
    ATOM   3619  HB3 PHE B 134      63.424  25.257  90.123  1.00  0.00           H
    ATOM   3620  HD1 PHE B 134      63.685  23.606  93.499  1.00  0.00           H
    ATOM   3621  HD2 PHE B 134      61.132  25.288  90.452  1.00  0.00           H
    ATOM   3622  HE1 PHE B 134      61.740  23.428  94.996  1.00  0.00           H
    ATOM   3623  HE2 PHE B 134      59.213  25.085  91.980  1.00  0.00           H
    ATOM   3624  HZ  PHE B 134      59.520  24.168  94.243  1.00  0.00           H
    ATOM   3625  N   CYS B 135      65.411  23.244  88.114  1.00 31.58           N
    ATOM   3626  CA  CYS B 135      66.486  23.378  87.134  1.00 27.36           C
    ATOM   3627  C   CYS B 135      65.940  23.738  85.756  1.00 30.79           C
    ATOM   3628  O   CYS B 135      64.747  23.609  85.521  1.00 32.34           O
    ATOM   3629  CB  CYS B 135      67.348  22.109  87.100  1.00 24.65           C
    ATOM   3630  SG  CYS B 135      66.371  20.644  86.642  1.00 29.63           S
    ATOM   3631  H   CYS B 135      64.572  22.771  87.805  1.00  0.00           H
    ATOM   3632  HA  CYS B 135      67.123  24.215  87.415  1.00  0.00           H
    ATOM   3633  HB2 CYS B 135      68.156  22.235  86.379  1.00  0.00           H
    ATOM   3634  HB3 CYS B 135      67.810  21.937  88.072  1.00  0.00           H
    ATOM   3635  HG  CYS B 135      67.389  19.780  86.699  1.00  0.00           H
    ATOM   3636  N   GLN B 136      66.826  24.202  84.870  1.00 29.60           N
    ATOM   3637  CA  GLN B 136      66.485  24.481  83.479  1.00 24.36           C
    ATOM   3638  C   GLN B 136      66.933  23.314  82.617  1.00 22.52           C
    ATOM   3639  O   GLN B 136      67.886  22.617  82.966  1.00 27.61           O
    ATOM   3640  CB  GLN B 136      67.167  25.770  83.012  1.00 26.08           C
    ATOM   3641  CG  GLN B 136      66.822  26.985  83.874  1.00 25.99           C
    ATOM   3642  CD  GLN B 136      67.444  28.289  83.397  1.00 26.25           C
    ATOM   3643  OE1 GLN B 136      66.742  29.287  83.259  1.00 33.80           O
    ATOM   3644  NE2 GLN B 136      68.758  28.304  83.171  1.00 24.96           N
    ATOM   3645  H   GLN B 136      67.795  24.318  85.135  1.00  0.00           H
    ATOM   3646  HA  GLN B 136      65.411  24.590  83.346  1.00  0.00           H
    ATOM   3647  HB2 GLN B 136      68.232  25.607  83.042  1.00  0.00           H
    ATOM   3648  HB3 GLN B 136      66.901  25.974  81.974  1.00  0.00           H
    ATOM   3649  HG2 GLN B 136      65.744  27.106  83.879  1.00  0.00           H
    ATOM   3650  HG3 GLN B 136      67.157  26.833  84.897  1.00  0.00           H
    ATOM   3651 HE21 GLN B 136      69.319  27.483  83.345  1.00  0.00           H
    ATOM   3652 HE22 GLN B 136      69.207  29.167  82.901  1.00  0.00           H
    ATOM   3653  N   LEU B 137      66.242  23.158  81.483  1.00 20.14           N
    ATOM   3654  CA  LEU B 137      66.494  22.125  80.491  1.00 17.85           C
    ATOM   3655  C   LEU B 137      67.954  22.163  80.011  1.00 17.28           C
    ATOM   3656  O   LEU B 137      68.447  23.240  79.676  1.00 17.76           O
    ATOM   3657  CB  LEU B 137      65.476  22.313  79.348  1.00 15.32           C
    ATOM   3658  CG  LEU B 137      65.259  21.093  78.431  1.00 16.15           C
    ATOM   3659  CD1 LEU B 137      63.993  21.309  77.588  1.00 18.93           C
    ATOM   3660  CD2 LEU B 137      66.454  20.792  77.508  1.00 10.85           C
    ATOM   3661  H   LEU B 137      65.474  23.784  81.280  1.00  0.00           H
    ATOM   3662  HA  LEU B 137      66.282  21.192  81.001  1.00  0.00           H
    ATOM   3663  HB2 LEU B 137      64.511  22.535  79.806  1.00  0.00           H
    ATOM   3664  HB3 LEU B 137      65.722  23.195  78.754  1.00  0.00           H
    ATOM   3665  HG  LEU B 137      65.084  20.225  79.064  1.00  0.00           H
    ATOM   3666 HD11 LEU B 137      63.833  20.497  76.878  1.00  0.00           H
    ATOM   3667 HD12 LEU B 137      63.105  21.379  78.214  1.00  0.00           H
    ATOM   3668 HD13 LEU B 137      64.058  22.234  77.017  1.00  0.00           H
    ATOM   3669 HD21 LEU B 137      66.140  20.366  76.555  1.00  0.00           H
    ATOM   3670 HD22 LEU B 137      67.021  21.695  77.280  1.00  0.00           H
    ATOM   3671 HD23 LEU B 137      67.141  20.083  77.960  1.00  0.00           H
    ATOM   3672  N   ALA B 138      68.622  21.001  80.028  1.00 19.01           N
    ATOM   3673  CA  ALA B 138      70.006  20.807  79.597  1.00 17.03           C
    ATOM   3674  C   ALA B 138      71.085  21.428  80.506  1.00 20.43           C
    ATOM   3675  O   ALA B 138      72.260  21.165  80.260  1.00 22.72           O
    ATOM   3676  CB  ALA B 138      70.188  21.257  78.136  1.00 13.67           C
    ATOM   3677  H   ALA B 138      68.136  20.167  80.335  1.00  0.00           H
    ATOM   3678  HA  ALA B 138      70.163  19.732  79.609  1.00  0.00           H
    ATOM   3679  HB1 ALA B 138      71.205  21.074  77.805  1.00  0.00           H
    ATOM   3680  HB2 ALA B 138      69.518  20.712  77.473  1.00  0.00           H
    ATOM   3681  HB3 ALA B 138      70.017  22.321  77.988  1.00  0.00           H
    ATOM   3682  N   LYS B 139      70.710  22.216  81.530  1.00 20.40           N
    ATOM   3683  CA  LYS B 139      71.661  22.842  82.447  1.00 16.22           C
    ATOM   3684  C   LYS B 139      72.229  21.852  83.464  1.00 14.64           C
    ATOM   3685  O   LYS B 139      71.602  20.830  83.750  1.00 22.62           O
    ATOM   3686  CB  LYS B 139      71.041  24.055  83.162  1.00 13.88           C
    ATOM   3687  CG  LYS B 139      70.683  25.229  82.243  1.00 14.65           C
    ATOM   3688  CD  LYS B 139      71.860  25.922  81.540  1.00 23.98           C
    ATOM   3689  CE  LYS B 139      71.563  26.317  80.091  1.00 34.78           C
    ATOM   3690  NZ  LYS B 139      70.333  27.120  79.981  1.00 43.28           N1+
    ATOM   3691  H   LYS B 139      69.729  22.396  81.697  1.00  0.00           H
    ATOM   3692  HA  LYS B 139      72.491  23.185  81.843  1.00  0.00           H
    ATOM   3693  HB2 LYS B 139      70.160  23.747  83.726  1.00  0.00           H
    ATOM   3694  HB3 LYS B 139      71.749  24.433  83.894  1.00  0.00           H
    ATOM   3695  HG2 LYS B 139      69.927  24.904  81.527  1.00  0.00           H
    ATOM   3696  HG3 LYS B 139      70.232  25.997  82.865  1.00  0.00           H
    ATOM   3697  HD2 LYS B 139      72.120  26.826  82.090  1.00  0.00           H
    ATOM   3698  HD3 LYS B 139      72.751  25.315  81.572  1.00  0.00           H
    ATOM   3699  HE2 LYS B 139      72.398  26.879  79.674  1.00  0.00           H
    ATOM   3700  HE3 LYS B 139      71.442  25.423  79.478  1.00  0.00           H
    ATOM   3701  HZ1 LYS B 139      70.407  27.943  80.562  1.00  0.00           H
    ATOM   3702  HZ2 LYS B 139      70.198  27.400  79.020  1.00  0.00           H
    ATOM   3703  HZ3 LYS B 139      69.532  26.572  80.268  1.00  0.00           H
    ATOM   3704  N   THR B 140      73.416  22.178  83.993  1.00 17.73           N
    ATOM   3705  CA  THR B 140      74.093  21.360  84.992  1.00 21.98           C
    ATOM   3706  C   THR B 140      73.293  21.297  86.300  1.00 27.90           C
    ATOM   3707  O   THR B 140      72.957  22.340  86.861  1.00 29.54           O
    ATOM   3708  CB  THR B 140      75.504  21.897  85.316  1.00 19.30           C
    ATOM   3709  OG1 THR B 140      76.246  21.979  84.120  1.00 24.17           O
    ATOM   3710  CG2 THR B 140      76.294  20.998  86.272  1.00 16.53           C
    ATOM   3711  H   THR B 140      73.895  23.019  83.705  1.00  0.00           H
    ATOM   3712  HA  THR B 140      74.193  20.351  84.586  1.00  0.00           H
    ATOM   3713  HB  THR B 140      75.440  22.902  85.735  1.00  0.00           H
    ATOM   3714 HG21 THR B 140      77.305  21.379  86.420  1.00  0.00           H
    ATOM   3715 HG22 THR B 140      75.826  20.937  87.251  1.00  0.00           H
    ATOM   3716 HG23 THR B 140      76.373  19.989  85.874  1.00  0.00           H
    ATOM   3717  HG1 THR B 140      76.431  21.081  83.817  1.00  0.00           H
    ATOM   3718  N   CYS B 141      73.012  20.063  86.734  1.00 24.96           N
    ATOM   3719  CA  CYS B 141      72.296  19.740  87.957  1.00 20.48           C
    ATOM   3720  C   CYS B 141      73.249  18.915  88.823  1.00 23.06           C
    ATOM   3721  O   CYS B 141      73.256  17.688  88.694  1.00 23.34           O
    ATOM   3722  CB  CYS B 141      71.002  18.959  87.683  1.00 13.78           C
    ATOM   3723  SG  CYS B 141      69.868  20.018  86.770  1.00 23.87           S
    ATOM   3724  H   CYS B 141      73.288  19.266  86.174  1.00  0.00           H
    ATOM   3725  HA  CYS B 141      72.011  20.643  88.492  1.00  0.00           H
    ATOM   3726  HB2 CYS B 141      71.163  18.038  87.130  1.00  0.00           H
    ATOM   3727  HB3 CYS B 141      70.547  18.668  88.623  1.00  0.00           H
    ATOM   3728  HG  CYS B 141      70.596  20.060  85.650  1.00  0.00           H
    ATOM   3729  N   PRO B 142      74.070  19.587  89.664  1.00 19.56           N
    ATOM   3730  CA  PRO B 142      74.991  18.880  90.562  1.00 21.92           C
    ATOM   3731  C   PRO B 142      74.217  18.114  91.642  1.00 24.10           C
    ATOM   3732  O   PRO B 142      73.291  18.663  92.232  1.00 22.39           O
    ATOM   3733  CB  PRO B 142      75.862  19.994  91.176  1.00 16.94           C
    ATOM   3734  CG  PRO B 142      75.595  21.236  90.346  1.00 18.33           C
    ATOM   3735  CD  PRO B 142      74.169  21.032  89.870  1.00 20.82           C
    ATOM   3736  HA  PRO B 142      75.615  18.207  89.971  1.00  0.00           H
    ATOM   3737  HB2 PRO B 142      75.568  20.198  92.207  1.00  0.00           H
    ATOM   3738  HB3 PRO B 142      76.921  19.734  91.195  1.00  0.00           H
    ATOM   3739  HG2 PRO B 142      75.739  22.172  90.883  1.00  0.00           H
    ATOM   3740  HG3 PRO B 142      76.268  21.240  89.488  1.00  0.00           H
    ATOM   3741  HD2 PRO B 142      73.461  21.327  90.646  1.00  0.00           H
    ATOM   3742  HD3 PRO B 142      73.981  21.642  88.994  1.00  0.00           H
    ATOM   3743  N   VAL B 143      74.597  16.856  91.864  1.00 25.42           N
    ATOM   3744  CA  VAL B 143      74.055  16.018  92.926  1.00 30.90           C
    ATOM   3745  C   VAL B 143      75.265  15.366  93.605  1.00 34.09           C
    ATOM   3746  O   VAL B 143      76.004  14.641  92.940  1.00 34.00           O
    ATOM   3747  CB  VAL B 143      73.116  14.915  92.355  1.00 26.82           C
    ATOM   3748  CG1 VAL B 143      72.690  13.852  93.388  1.00 22.75           C
    ATOM   3749  CG2 VAL B 143      71.862  15.524  91.705  1.00 21.73           C
    ATOM   3750  H   VAL B 143      75.327  16.439  91.301  1.00  0.00           H
    ATOM   3751  HA  VAL B 143      73.514  16.608  93.669  1.00  0.00           H
    ATOM   3752  HB  VAL B 143      73.651  14.389  91.562  1.00  0.00           H
    ATOM   3753 HG11 VAL B 143      71.979  13.147  92.956  1.00  0.00           H
    ATOM   3754 HG12 VAL B 143      73.532  13.262  93.747  1.00  0.00           H
    ATOM   3755 HG13 VAL B 143      72.211  14.312  94.254  1.00  0.00           H
    ATOM   3756 HG21 VAL B 143      71.186  14.749  91.345  1.00  0.00           H
    ATOM   3757 HG22 VAL B 143      71.319  16.129  92.425  1.00  0.00           H
    ATOM   3758 HG23 VAL B 143      72.114  16.158  90.856  1.00  0.00           H
    ATOM   3759  N   GLN B 144      75.473  15.650  94.897  1.00 34.85           N
    ATOM   3760  CA  GLN B 144      76.627  15.128  95.631  1.00 30.66           C
    ATOM   3761  C   GLN B 144      76.276  13.822  96.335  1.00 27.97           C
    ATOM   3762  O   GLN B 144      75.212  13.734  96.936  1.00 35.10           O
    ATOM   3763  CB  GLN B 144      77.081  16.137  96.679  1.00 28.94           C
    ATOM   3764  CG  GLN B 144      77.529  17.482  96.094  1.00 28.70           C
    ATOM   3765  CD  GLN B 144      77.870  18.459  97.212  1.00 35.43           C
    ATOM   3766  OE1 GLN B 144      77.507  18.251  98.368  1.00 35.41           O
    ATOM   3767  NE2 GLN B 144      78.577  19.535  96.877  1.00 40.91           N
    ATOM   3768  H   GLN B 144      74.827  16.234  95.413  1.00  0.00           H
    ATOM   3769  HA  GLN B 144      77.467  14.958  94.956  1.00  0.00           H
    ATOM   3770  HB2 GLN B 144      76.265  16.291  97.385  1.00  0.00           H
    ATOM   3771  HB3 GLN B 144      77.903  15.711  97.257  1.00  0.00           H
    ATOM   3772  HG2 GLN B 144      78.396  17.342  95.450  1.00  0.00           H
    ATOM   3773  HG3 GLN B 144      76.741  17.927  95.486  1.00  0.00           H
    ATOM   3774 HE21 GLN B 144      78.873  19.675  95.922  1.00  0.00           H
    ATOM   3775 HE22 GLN B 144      78.834  20.211  97.581  1.00  0.00           H
    ATOM   3776  N   LEU B 145      77.198  12.858  96.293  1.00 24.92           N
    ATOM   3777  CA  LEU B 145      77.052  11.566  96.949  1.00 34.13           C
    ATOM   3778  C   LEU B 145      77.948  11.531  98.191  1.00 32.30           C
    ATOM   3779  O   LEU B 145      79.169  11.667  98.082  1.00 28.93           O
    ATOM   3780  CB  LEU B 145      77.416  10.419  95.986  1.00 38.07           C
    ATOM   3781  CG  LEU B 145      76.741  10.428  94.597  1.00 40.43           C
    ATOM   3782  CD1 LEU B 145      77.269   9.257  93.762  1.00 37.88           C
    ATOM   3783  CD2 LEU B 145      75.202  10.427  94.634  1.00 40.11           C
    ATOM   3784  H   LEU B 145      78.055  12.996  95.769  1.00  0.00           H
    ATOM   3785  HA  LEU B 145      76.019  11.417  97.269  1.00  0.00           H
    ATOM   3786  HB2 LEU B 145      78.488  10.436  95.830  1.00  0.00           H
    ATOM   3787  HB3 LEU B 145      77.213   9.468  96.479  1.00  0.00           H
    ATOM   3788  HG  LEU B 145      77.056  11.340  94.088  1.00  0.00           H
    ATOM   3789 HD11 LEU B 145      77.146   9.469  92.705  1.00  0.00           H
    ATOM   3790 HD12 LEU B 145      78.324   9.056  93.940  1.00  0.00           H
    ATOM   3791 HD13 LEU B 145      76.729   8.336  93.982  1.00  0.00           H
    ATOM   3792 HD21 LEU B 145      74.806  11.269  94.065  1.00  0.00           H
    ATOM   3793 HD22 LEU B 145      74.783   9.520  94.199  1.00  0.00           H
    ATOM   3794 HD23 LEU B 145      74.813  10.503  95.646  1.00  0.00           H
    ATOM   3795  N   TRP B 146      77.297  11.331  99.342  1.00 31.10           N
    ATOM   3796  CA  TRP B 146      77.908  11.239 100.660  1.00 27.05           C
    ATOM   3797  C   TRP B 146      77.678   9.850 101.230  1.00 29.19           C
    ATOM   3798  O   TRP B 146      76.532   9.460 101.435  1.00 34.84           O
    ATOM   3799  CB  TRP B 146      77.261  12.253 101.609  1.00 16.42           C
    ATOM   3800  CG  TRP B 146      77.555  13.672 101.284  1.00 16.24           C
    ATOM   3801  CD1 TRP B 146      76.947  14.410 100.332  1.00 23.25           C
    ATOM   3802  CD2 TRP B 146      78.586  14.515 101.861  1.00 17.20           C
    ATOM   3803  NE1 TRP B 146      77.526  15.659 100.287  1.00 19.23           N
    ATOM   3804  CE2 TRP B 146      78.529  15.786 101.222  1.00 18.39           C
    ATOM   3805  CE3 TRP B 146      79.564  14.337 102.863  1.00 22.03           C
    ATOM   3806  CZ2 TRP B 146      79.393  16.837 101.569  1.00 26.35           C
    ATOM   3807  CZ3 TRP B 146      80.417  15.388 103.239  1.00 24.97           C
    ATOM   3808  CH2 TRP B 146      80.335  16.634 102.592  1.00 28.22           C
    ATOM   3809  H   TRP B 146      76.290  11.224  99.332  1.00  0.00           H
    ATOM   3810  HA  TRP B 146      78.980  11.441 100.614  1.00  0.00           H
    ATOM   3811  HB2 TRP B 146      76.179  12.131 101.637  1.00  0.00           H
    ATOM   3812  HB3 TRP B 146      77.604  12.080 102.629  1.00  0.00           H
    ATOM   3813  HD1 TRP B 146      76.141  14.059  99.705  1.00  0.00           H
    ATOM   3814  HE1 TRP B 146      77.234  16.383  99.644  1.00  0.00           H
    ATOM   3815  HE3 TRP B 146      79.640  13.389 103.362  1.00  0.00           H
    ATOM   3816  HZ2 TRP B 146      79.328  17.789 101.063  1.00  0.00           H
    ATOM   3817  HZ3 TRP B 146      81.133  15.238 104.032  1.00  0.00           H
    ATOM   3818  HH2 TRP B 146      80.995  17.438 102.882  1.00  0.00           H
    ATOM   3819  N   VAL B 147      78.777   9.167 101.545  1.00 27.79           N
    ATOM   3820  CA  VAL B 147      78.763   7.913 102.282  1.00 28.72           C
    ATOM   3821  C   VAL B 147      79.536   8.098 103.597  1.00 33.21           C
    ATOM   3822  O   VAL B 147      80.439   8.930 103.681  1.00 29.51           O
    ATOM   3823  CB  VAL B 147      79.380   6.765 101.446  1.00 27.68           C
    ATOM   3824  CG1 VAL B 147      78.446   6.358 100.291  1.00 27.92           C
    ATOM   3825  CG2 VAL B 147      80.802   7.067 100.937  1.00 29.58           C
    ATOM   3826  H   VAL B 147      79.691   9.555 101.343  1.00  0.00           H
    ATOM   3827  HA  VAL B 147      77.747   7.622 102.538  1.00  0.00           H
    ATOM   3828  HB  VAL B 147      79.454   5.889 102.091  1.00  0.00           H
    ATOM   3829 HG11 VAL B 147      78.846   5.513  99.732  1.00  0.00           H
    ATOM   3830 HG12 VAL B 147      77.464   6.064 100.664  1.00  0.00           H
    ATOM   3831 HG13 VAL B 147      78.302   7.177  99.586  1.00  0.00           H
    ATOM   3832 HG21 VAL B 147      81.198   6.247 100.346  1.00  0.00           H
    ATOM   3833 HG22 VAL B 147      80.817   7.944 100.291  1.00  0.00           H
    ATOM   3834 HG23 VAL B 147      81.504   7.235 101.755  1.00  0.00           H
    ATOM   3835  N   ASP B 148      79.143   7.329 104.619  1.00 34.52           N
    ATOM   3836  CA  ASP B 148      79.857   7.206 105.895  1.00 34.49           C
    ATOM   3837  C   ASP B 148      81.090   6.310 105.792  1.00 34.53           C
    ATOM   3838  O   ASP B 148      82.023   6.464 106.579  1.00 31.88           O
    ATOM   3839  CB  ASP B 148      78.957   6.636 107.005  1.00 35.18           C
    ATOM   3840  CG  ASP B 148      77.874   7.576 107.512  1.00 36.89           C
    ATOM   3841  OD1 ASP B 148      78.124   8.800 107.510  1.00 38.09           O
    ATOM   3842  OD2 ASP B 148      76.856   7.041 108.000  1.00 43.77           O1-
    ATOM   3843  H   ASP B 148      78.389   6.668 104.480  1.00  0.00           H
    ATOM   3844  HA  ASP B 148      80.215   8.193 106.195  1.00  0.00           H
    ATOM   3845  HB2 ASP B 148      78.460   5.776 106.571  1.00  0.00           H
    ATOM   3846  HB3 ASP B 148      79.538   6.294 107.864  1.00  0.00           H
    ATOM   3847  N   SER B 149      81.056   5.380 104.837  1.00 35.07           N
    ATOM   3848  CA  SER B 149      82.085   4.387 104.608  1.00 37.28           C
    ATOM   3849  C   SER B 149      82.065   4.046 103.123  1.00 37.47           C
    ATOM   3850  O   SER B 149      80.989   3.841 102.565  1.00 37.19           O
    ATOM   3851  CB  SER B 149      81.864   3.195 105.565  1.00 41.88           C
    ATOM   3852  OG  SER B 149      80.516   2.769 105.612  1.00 48.42           O
    ATOM   3853  H   SER B 149      80.235   5.299 104.253  1.00  0.00           H
    ATOM   3854  HA  SER B 149      83.061   4.823 104.828  1.00  0.00           H
    ATOM   3855  HB2 SER B 149      82.498   2.353 105.283  1.00  0.00           H
    ATOM   3856  HB3 SER B 149      82.156   3.475 106.577  1.00  0.00           H
    ATOM   3857  HG  SER B 149      80.363   2.158 104.887  1.00  0.00           H
    ATOM   3858  N   THR B 150      83.252   4.040 102.495  1.00 39.89           N
    ATOM   3859  CA  THR B 150      83.387   3.821 101.057  1.00 48.22           C
    ATOM   3860  C   THR B 150      82.853   2.415 100.671  1.00 47.59           C
    ATOM   3861  O   THR B 150      83.292   1.425 101.263  1.00 47.71           O
    ATOM   3862  CB  THR B 150      84.863   3.996 100.605  1.00 50.46           C
    ATOM   3863  OG1 THR B 150      85.125   5.386 100.523  1.00 53.96           O
    ATOM   3864  CG2 THR B 150      85.241   3.389  99.241  1.00 45.65           C
    ATOM   3865  H   THR B 150      84.103   4.237 103.002  1.00  0.00           H
    ATOM   3866  HA  THR B 150      82.825   4.624 100.589  1.00  0.00           H
    ATOM   3867  HB  THR B 150      85.534   3.586 101.360  1.00  0.00           H
    ATOM   3868 HG21 THR B 150      86.204   3.769  98.894  1.00  0.00           H
    ATOM   3869 HG22 THR B 150      85.349   2.308  99.312  1.00  0.00           H
    ATOM   3870 HG23 THR B 150      84.502   3.616  98.471  1.00  0.00           H
    ATOM   3871  HG1 THR B 150      84.726   5.733  99.722  1.00  0.00           H
    ATOM   3872  N   PRO B 151      81.891   2.363  99.714  1.00 41.32           N
    ATOM   3873  CA  PRO B 151      81.427   1.112  99.089  1.00 36.37           C
    ATOM   3874  C   PRO B 151      82.563   0.294  98.458  1.00 35.72           C
    ATOM   3875  O   PRO B 151      83.625   0.858  98.190  1.00 32.58           O
    ATOM   3876  CB  PRO B 151      80.430   1.572  98.012  1.00 32.69           C
    ATOM   3877  CG  PRO B 151      79.972   2.950  98.430  1.00 31.14           C
    ATOM   3878  CD  PRO B 151      81.158   3.513  99.188  1.00 36.41           C
    ATOM   3879  HA  PRO B 151      80.918   0.539  99.851  1.00  0.00           H
    ATOM   3880  HB2 PRO B 151      80.926   1.657  97.044  1.00  0.00           H
    ATOM   3881  HB3 PRO B 151      79.593   0.883  97.891  1.00  0.00           H
    ATOM   3882  HG2 PRO B 151      79.649   3.577  97.598  1.00  0.00           H
    ATOM   3883  HG3 PRO B 151      79.132   2.848  99.119  1.00  0.00           H
    ATOM   3884  HD2 PRO B 151      81.813   4.085  98.537  1.00  0.00           H
    ATOM   3885  HD3 PRO B 151      80.786   4.156  99.981  1.00  0.00           H
    ATOM   3886  N   PRO B 152      82.323  -1.002  98.159  1.00 36.63           N
    ATOM   3887  CA  PRO B 152      83.344  -1.807  97.482  1.00 36.29           C
    ATOM   3888  C   PRO B 152      83.588  -1.305  96.040  1.00 45.08           C
    ATOM   3889  O   PRO B 152      82.831  -0.477  95.532  1.00 53.04           O
    ATOM   3890  CB  PRO B 152      82.780  -3.244  97.503  1.00 32.60           C
    ATOM   3891  CG  PRO B 152      81.460  -3.191  98.262  1.00 32.93           C
    ATOM   3892  CD  PRO B 152      81.059  -1.725  98.238  1.00 34.73           C
    ATOM   3893  HA  PRO B 152      84.255  -1.758  98.079  1.00  0.00           H
    ATOM   3894  HB2 PRO B 152      82.588  -3.609  96.492  1.00  0.00           H
    ATOM   3895  HB3 PRO B 152      83.473  -3.948  97.964  1.00  0.00           H
    ATOM   3896  HG2 PRO B 152      80.694  -3.850  97.850  1.00  0.00           H
    ATOM   3897  HG3 PRO B 152      81.633  -3.491  99.296  1.00  0.00           H
    ATOM   3898  HD2 PRO B 152      80.483  -1.520  97.339  1.00  0.00           H
    ATOM   3899  HD3 PRO B 152      80.441  -1.455  99.085  1.00  0.00           H
    ATOM   3900  N   PRO B 153      84.615  -1.847  95.365  1.00 47.30           N
    ATOM   3901  CA  PRO B 153      84.747  -1.693  93.908  1.00 45.53           C
    ATOM   3902  C   PRO B 153      83.593  -2.353  93.133  1.00 46.41           C
    ATOM   3903  O   PRO B 153      82.976  -3.292  93.633  1.00 48.23           O
    ATOM   3904  CB  PRO B 153      86.094  -2.359  93.597  1.00 48.28           C
    ATOM   3905  CG  PRO B 153      86.858  -2.352  94.911  1.00 53.07           C
    ATOM   3906  CD  PRO B 153      85.771  -2.499  95.964  1.00 51.02           C
    ATOM   3907  HA  PRO B 153      84.799  -0.627  93.679  1.00  0.00           H
    ATOM   3908  HB2 PRO B 153      85.952  -3.392  93.275  1.00  0.00           H
    ATOM   3909  HB3 PRO B 153      86.639  -1.844  92.806  1.00  0.00           H
    ATOM   3910  HG2 PRO B 153      87.629  -3.120  94.971  1.00  0.00           H
    ATOM   3911  HG3 PRO B 153      87.344  -1.383  95.034  1.00  0.00           H
    ATOM   3912  HD2 PRO B 153      85.535  -3.549  96.142  1.00  0.00           H
    ATOM   3913  HD3 PRO B 153      86.093  -2.048  96.903  1.00  0.00           H
    ATOM   3914  N   GLY B 154      83.320  -1.831  91.930  1.00 45.77           N
    ATOM   3915  CA  GLY B 154      82.226  -2.302  91.083  1.00 41.45           C
    ATOM   3916  C   GLY B 154      80.921  -1.564  91.399  1.00 38.83           C
    ATOM   3917  O   GLY B 154      79.979  -1.645  90.609  1.00 37.82           O
    ATOM   3918  H   GLY B 154      83.867  -1.055  91.586  1.00  0.00           H
    ATOM   3919  HA2 GLY B 154      82.491  -2.129  90.040  1.00  0.00           H
    ATOM   3920  HA3 GLY B 154      82.079  -3.375  91.203  1.00  0.00           H
    ATOM   3921  N   THR B 155      80.877  -0.816  92.516  1.00 37.38           N
    ATOM   3922  CA  THR B 155      79.772   0.053  92.872  1.00 32.22           C
    ATOM   3923  C   THR B 155      79.520   1.090  91.775  1.00 34.30           C
    ATOM   3924  O   THR B 155      80.448   1.749  91.318  1.00 32.85           O
    ATOM   3925  CB  THR B 155      79.950   0.754  94.232  1.00 31.98           C
    ATOM   3926  OG1 THR B 155      80.054  -0.231  95.228  1.00 39.47           O
    ATOM   3927  CG2 THR B 155      78.745   1.611  94.656  1.00 30.09           C
    ATOM   3928  H   THR B 155      81.677  -0.799  93.132  1.00  0.00           H
    ATOM   3929  HA  THR B 155      78.906  -0.570  92.973  1.00  0.00           H
    ATOM   3930  HB  THR B 155      80.863   1.351  94.243  1.00  0.00           H
    ATOM   3931 HG21 THR B 155      78.910   2.070  95.630  1.00  0.00           H
    ATOM   3932 HG22 THR B 155      78.534   2.414  93.958  1.00  0.00           H
    ATOM   3933 HG23 THR B 155      77.840   1.008  94.723  1.00  0.00           H
    ATOM   3934  HG1 THR B 155      80.787  -0.820  95.025  1.00  0.00           H
    ATOM   3935  N   ARG B 156      78.264   1.147  91.350  1.00 36.34           N
    ATOM   3936  CA  ARG B 156      77.724   2.059  90.371  1.00 35.51           C
    ATOM   3937  C   ARG B 156      76.608   2.870  91.026  1.00 32.14           C
    ATOM   3938  O   ARG B 156      76.047   2.480  92.047  1.00 34.26           O
    ATOM   3939  CB  ARG B 156      77.170   1.235  89.210  1.00 42.00           C
    ATOM   3940  CG  ARG B 156      78.227   0.803  88.176  1.00 48.36           C
    ATOM   3941  CD  ARG B 156      77.600   0.050  87.000  1.00 53.97           C
    ATOM   3942  NE  ARG B 156      76.743  -1.015  87.525  1.00 56.40           N
    ATOM   3943  CZ  ARG B 156      75.606  -1.478  86.997  1.00 55.36           C
    ATOM   3944  NH1 ARG B 156      75.144  -1.040  85.818  1.00 52.99           N
    ATOM   3945  NH2 ARG B 156      74.941  -2.417  87.675  1.00 54.38           N1+
    ATOM   3946  H   ARG B 156      77.581   0.539  91.791  1.00  0.00           H
    ATOM   3947  HA  ARG B 156      78.491   2.743  90.014  1.00  0.00           H
    ATOM   3948  HB2 ARG B 156      76.632   0.379  89.622  1.00  0.00           H
    ATOM   3949  HB3 ARG B 156      76.421   1.820  88.695  1.00  0.00           H
    ATOM   3950  HG2 ARG B 156      78.666   1.714  87.773  1.00  0.00           H
    ATOM   3951  HG3 ARG B 156      79.045   0.231  88.615  1.00  0.00           H
    ATOM   3952  HD2 ARG B 156      77.100   0.733  86.314  1.00  0.00           H
    ATOM   3953  HD3 ARG B 156      78.375  -0.464  86.431  1.00  0.00           H
    ATOM   3954  HE  ARG B 156      77.080  -1.456  88.369  1.00  0.00           H
    ATOM   3955 HH11 ARG B 156      75.655  -0.334  85.308  1.00  0.00           H
    ATOM   3956 HH12 ARG B 156      74.335  -1.466  85.392  1.00  0.00           H
    ATOM   3957 HH21 ARG B 156      75.240  -2.683  88.603  1.00  0.00           H
    ATOM   3958 HH22 ARG B 156      74.068  -2.783  87.325  1.00  0.00           H
    ATOM   3959  N   VAL B 157      76.314   4.002  90.391  1.00 27.18           N
    ATOM   3960  CA  VAL B 157      75.297   4.950  90.797  1.00 26.24           C
    ATOM   3961  C   VAL B 157      74.311   5.037  89.638  1.00 27.57           C
    ATOM   3962  O   VAL B 157      74.738   5.342  88.530  1.00 26.84           O
    ATOM   3963  CB  VAL B 157      75.881   6.376  90.970  1.00 22.73           C
    ATOM   3964  CG1 VAL B 157      74.833   7.347  91.546  1.00 22.38           C
    ATOM   3965  CG2 VAL B 157      77.153   6.377  91.827  1.00 22.98           C
    ATOM   3966  H   VAL B 157      76.830   4.247  89.554  1.00  0.00           H
    ATOM   3967  HA  VAL B 157      74.797   4.642  91.716  1.00  0.00           H
    ATOM   3968  HB  VAL B 157      76.193   6.772  90.001  1.00  0.00           H
    ATOM   3969 HG11 VAL B 157      75.261   8.337  91.702  1.00  0.00           H
    ATOM   3970 HG12 VAL B 157      73.982   7.478  90.877  1.00  0.00           H
    ATOM   3971 HG13 VAL B 157      74.457   6.994  92.507  1.00  0.00           H
    ATOM   3972 HG21 VAL B 157      77.596   7.370  91.866  1.00  0.00           H
    ATOM   3973 HG22 VAL B 157      76.950   6.051  92.847  1.00  0.00           H
    ATOM   3974 HG23 VAL B 157      77.921   5.728  91.412  1.00  0.00           H
    ATOM   3975  N   ARG B 158      73.028   4.790  89.905  1.00 31.72           N
    ATOM   3976  CA  ARG B 158      71.976   4.844  88.904  1.00 32.46           C
    ATOM   3977  C   ARG B 158      70.993   5.952  89.249  1.00 28.64           C
    ATOM   3978  O   ARG B 158      70.615   6.070  90.405  1.00 30.90           O
    ATOM   3979  CB  ARG B 158      71.275   3.483  88.831  1.00 33.95           C
    ATOM   3980  CG  ARG B 158      70.184   3.431  87.745  1.00 33.86           C
    ATOM   3981  CD  ARG B 158      69.437   2.097  87.721  1.00 33.11           C
    ATOM   3982  NE  ARG B 158      70.347   0.978  87.459  1.00 36.11           N
    ATOM   3983  CZ  ARG B 158      71.012   0.728  86.320  1.00 42.46           C
    ATOM   3984  NH1 ARG B 158      70.869   1.466  85.214  1.00 41.45           N
    ATOM   3985  NH2 ARG B 158      71.855  -0.305  86.287  1.00 43.40           N1+
    ATOM   3986  H   ARG B 158      72.748   4.528  90.843  1.00  0.00           H
    ATOM   3987  HA  ARG B 158      72.393   5.044  87.922  1.00  0.00           H
    ATOM   3988  HB2 ARG B 158      72.022   2.715  88.630  1.00  0.00           H
    ATOM   3989  HB3 ARG B 158      70.845   3.230  89.799  1.00  0.00           H
    ATOM   3990  HG2 ARG B 158      69.433   4.206  87.894  1.00  0.00           H
    ATOM   3991  HG3 ARG B 158      70.635   3.635  86.775  1.00  0.00           H
    ATOM   3992  HD2 ARG B 158      68.971   1.929  88.692  1.00  0.00           H
    ATOM   3993  HD3 ARG B 158      68.628   2.104  86.992  1.00  0.00           H
    ATOM   3994  HE  ARG B 158      70.559   0.400  88.266  1.00  0.00           H
    ATOM   3995 HH11 ARG B 158      70.149   2.180  85.162  1.00  0.00           H
    ATOM   3996 HH12 ARG B 158      71.401   1.253  84.382  1.00  0.00           H
    ATOM   3997 HH21 ARG B 158      72.052  -0.819  87.142  1.00  0.00           H
    ATOM   3998 HH22 ARG B 158      72.341  -0.537  85.436  1.00  0.00           H
    ATOM   3999  N   ALA B 159      70.551   6.690  88.228  1.00 26.83           N
    ATOM   4000  CA  ALA B 159      69.466   7.653  88.326  1.00 28.11           C
    ATOM   4001  C   ALA B 159      68.314   7.202  87.434  1.00 32.06           C
    ATOM   4002  O   ALA B 159      68.546   6.659  86.355  1.00 29.51           O
    ATOM   4003  CB  ALA B 159      69.970   9.030  87.892  1.00 26.32           C
    ATOM   4004  H   ALA B 159      70.905   6.517  87.296  1.00  0.00           H
    ATOM   4005  HA  ALA B 159      69.106   7.738  89.351  1.00  0.00           H
    ATOM   4006  HB1 ALA B 159      69.165   9.766  87.887  1.00  0.00           H
    ATOM   4007  HB2 ALA B 159      70.723   9.385  88.590  1.00  0.00           H
    ATOM   4008  HB3 ALA B 159      70.412   9.007  86.896  1.00  0.00           H
    ATOM   4009  N   MET B 160      67.091   7.472  87.895  1.00 32.47           N
    ATOM   4010  CA  MET B 160      65.860   7.147  87.194  1.00 28.23           C
    ATOM   4011  C   MET B 160      64.774   8.107  87.661  1.00 26.42           C
    ATOM   4012  O   MET B 160      64.688   8.370  88.858  1.00 26.69           O
    ATOM   4013  CB  MET B 160      65.487   5.678  87.469  1.00 29.40           C
    ATOM   4014  CG  MET B 160      64.004   5.320  87.273  1.00 30.44           C
    ATOM   4015  SD  MET B 160      63.585   3.626  87.690  1.00 28.36           S
    ATOM   4016  CE  MET B 160      64.843   2.709  86.776  1.00 21.89           C
    ATOM   4017  H   MET B 160      66.980   7.914  88.798  1.00  0.00           H
    ATOM   4018  HA  MET B 160      65.997   7.291  86.120  1.00  0.00           H
    ATOM   4019  HB2 MET B 160      66.115   5.056  86.836  1.00  0.00           H
    ATOM   4020  HB3 MET B 160      65.752   5.422  88.496  1.00  0.00           H
    ATOM   4021  HG2 MET B 160      63.351   5.928  87.898  1.00  0.00           H
    ATOM   4022  HG3 MET B 160      63.713   5.501  86.237  1.00  0.00           H
    ATOM   4023  HE1 MET B 160      64.610   1.644  86.801  1.00  0.00           H
    ATOM   4024  HE2 MET B 160      64.861   3.025  85.733  1.00  0.00           H
    ATOM   4025  HE3 MET B 160      65.831   2.845  87.215  1.00  0.00           H
    ATOM   4026  N   ALA B 161      63.921   8.538  86.725  1.00 26.58           N
    ATOM   4027  CA  ALA B 161      62.745   9.330  87.039  1.00 29.13           C
    ATOM   4028  C   ALA B 161      61.501   8.446  87.159  1.00 27.65           C
    ATOM   4029  O   ALA B 161      61.379   7.450  86.448  1.00 25.92           O
    ATOM   4030  CB  ALA B 161      62.528  10.407  85.976  1.00 31.93           C
    ATOM   4031  H   ALA B 161      64.021   8.243  85.762  1.00  0.00           H
    ATOM   4032  HA  ALA B 161      62.923   9.860  87.961  1.00  0.00           H
    ATOM   4033  HB1 ALA B 161      61.575  10.911  86.133  1.00  0.00           H
    ATOM   4034  HB2 ALA B 161      63.303  11.171  86.014  1.00  0.00           H
    ATOM   4035  HB3 ALA B 161      62.518   9.987  84.970  1.00  0.00           H
    ATOM   4036  N   ILE B 162      60.578   8.871  88.024  1.00 22.43           N
    ATOM   4037  CA  ILE B 162      59.227   8.332  88.129  1.00 23.26           C
    ATOM   4038  C   ILE B 162      58.289   9.502  88.420  1.00 31.31           C
    ATOM   4039  O   ILE B 162      58.722  10.449  89.060  1.00 29.53           O
    ATOM   4040  CB  ILE B 162      59.089   7.277  89.260  1.00 28.26           C
    ATOM   4041  CG1 ILE B 162      59.309   7.789  90.707  1.00 31.18           C
    ATOM   4042  CG2 ILE B 162      59.966   6.045  88.977  1.00 28.02           C
    ATOM   4043  CD1 ILE B 162      58.600   6.935  91.762  1.00 33.63           C
    ATOM   4044  H   ILE B 162      60.769   9.684  88.596  1.00  0.00           H
    ATOM   4045  HA  ILE B 162      58.932   7.876  87.185  1.00  0.00           H
    ATOM   4046  HB  ILE B 162      58.057   6.924  89.210  1.00  0.00           H
    ATOM   4047 HG12 ILE B 162      60.376   7.849  90.920  1.00  0.00           H
    ATOM   4048 HG13 ILE B 162      58.926   8.794  90.857  1.00  0.00           H
    ATOM   4049 HG21 ILE B 162      59.727   5.229  89.656  1.00  0.00           H
    ATOM   4050 HG22 ILE B 162      59.817   5.672  87.965  1.00  0.00           H
    ATOM   4051 HG23 ILE B 162      61.026   6.267  89.092  1.00  0.00           H
    ATOM   4052 HD11 ILE B 162      59.084   7.026  92.731  1.00  0.00           H
    ATOM   4053 HD12 ILE B 162      57.563   7.249  91.883  1.00  0.00           H
    ATOM   4054 HD13 ILE B 162      58.594   5.876  91.508  1.00  0.00           H
    ATOM   4055  N   TYR B 163      57.023   9.426  87.992  1.00 36.18           N
    ATOM   4056  CA  TYR B 163      56.004  10.390  88.388  1.00 36.49           C
    ATOM   4057  C   TYR B 163      55.713  10.245  89.872  1.00 30.49           C
    ATOM   4058  O   TYR B 163      55.622   9.135  90.390  1.00 31.35           O
    ATOM   4059  CB  TYR B 163      54.724  10.257  87.560  1.00 38.82           C
    ATOM   4060  CG  TYR B 163      54.938  10.641  86.115  1.00 31.18           C
    ATOM   4061  CD1 TYR B 163      55.178   9.646  85.152  1.00 26.95           C
    ATOM   4062  CD2 TYR B 163      54.965  11.998  85.741  1.00 29.29           C
    ATOM   4063  CE1 TYR B 163      55.490   9.996  83.829  1.00 25.31           C
    ATOM   4064  CE2 TYR B 163      55.299  12.353  84.421  1.00 28.75           C
    ATOM   4065  CZ  TYR B 163      55.580  11.351  83.474  1.00 27.25           C
    ATOM   4066  OH  TYR B 163      55.940  11.688  82.208  1.00 28.38           O
    ATOM   4067  H   TYR B 163      56.713   8.608  87.478  1.00  0.00           H
    ATOM   4068  HA  TYR B 163      56.409  11.381  88.202  1.00  0.00           H
    ATOM   4069  HB2 TYR B 163      54.348   9.241  87.637  1.00  0.00           H
    ATOM   4070  HB3 TYR B 163      53.949  10.910  87.966  1.00  0.00           H
    ATOM   4071  HD1 TYR B 163      55.130   8.608  85.417  1.00  0.00           H
    ATOM   4072  HD2 TYR B 163      54.797  12.770  86.478  1.00  0.00           H
    ATOM   4073  HE1 TYR B 163      55.674   9.226  83.093  1.00  0.00           H
    ATOM   4074  HE2 TYR B 163      55.371  13.396  84.154  1.00  0.00           H
    ATOM   4075  HH  TYR B 163      56.008  12.632  82.059  1.00  0.00           H
    ATOM   4076  N   LYS B 164      55.605  11.395  90.523  1.00 29.73           N
    ATOM   4077  CA  LYS B 164      55.388  11.498  91.948  1.00 42.52           C
    ATOM   4078  C   LYS B 164      53.988  11.007  92.358  1.00 58.43           C
    ATOM   4079  O   LYS B 164      53.845  10.402  93.420  1.00 59.13           O
    ATOM   4080  CB  LYS B 164      55.602  12.969  92.283  1.00 37.82           C
    ATOM   4081  CG  LYS B 164      55.612  13.263  93.771  1.00 34.37           C
    ATOM   4082  CD  LYS B 164      55.682  14.764  94.023  1.00 35.84           C
    ATOM   4083  CE  LYS B 164      55.438  15.102  95.495  1.00 36.93           C
    ATOM   4084  NZ  LYS B 164      56.362  14.365  96.371  1.00 44.88           N1+
    ATOM   4085  H   LYS B 164      55.700  12.267  90.016  1.00  0.00           H
    ATOM   4086  HA  LYS B 164      56.136  10.896  92.467  1.00  0.00           H
    ATOM   4087  HB2 LYS B 164      56.544  13.320  91.857  1.00  0.00           H
    ATOM   4088  HB3 LYS B 164      54.804  13.536  91.808  1.00  0.00           H
    ATOM   4089  HG2 LYS B 164      54.709  12.874  94.241  1.00  0.00           H
    ATOM   4090  HG3 LYS B 164      56.450  12.747  94.237  1.00  0.00           H
    ATOM   4091  HD2 LYS B 164      56.658  15.107  93.693  1.00  0.00           H
    ATOM   4092  HD3 LYS B 164      54.950  15.291  93.412  1.00  0.00           H
    ATOM   4093  HE2 LYS B 164      55.535  16.173  95.671  1.00  0.00           H
    ATOM   4094  HE3 LYS B 164      54.419  14.832  95.772  1.00  0.00           H
    ATOM   4095  HZ1 LYS B 164      57.308  14.648  96.151  1.00  0.00           H
    ATOM   4096  HZ2 LYS B 164      56.166  14.590  97.336  1.00  0.00           H
    ATOM   4097  HZ3 LYS B 164      56.260  13.371  96.223  1.00  0.00           H
    ATOM   4098  N   GLN B 165      52.998  11.276  91.492  1.00 62.22           N
    ATOM   4099  CA  GLN B 165      51.598  10.939  91.698  1.00 54.43           C
    ATOM   4100  C   GLN B 165      51.342   9.455  91.419  1.00 52.51           C
    ATOM   4101  O   GLN B 165      51.827   8.903  90.434  1.00 46.77           O
    ATOM   4102  CB  GLN B 165      50.722  11.839  90.815  1.00 52.92           C
    ATOM   4103  CG  GLN B 165      50.875  13.343  91.139  1.00 57.24           C
    ATOM   4104  CD  GLN B 165      50.914  14.224  89.891  1.00 65.58           C
    ATOM   4105  OE1 GLN B 165      51.549  13.884  88.894  1.00 68.37           O
    ATOM   4106  NE2 GLN B 165      50.267  15.388  89.955  1.00 66.18           N
    ATOM   4107  H   GLN B 165      53.216  11.750  90.627  1.00  0.00           H
    ATOM   4108  HA  GLN B 165      51.351  11.156  92.740  1.00  0.00           H
    ATOM   4109  HB2 GLN B 165      50.958  11.622  89.772  1.00  0.00           H
    ATOM   4110  HB3 GLN B 165      49.673  11.572  90.939  1.00  0.00           H
    ATOM   4111  HG2 GLN B 165      50.071  13.658  91.806  1.00  0.00           H
    ATOM   4112  HG3 GLN B 165      51.801  13.545  91.680  1.00  0.00           H
    ATOM   4113 HE21 GLN B 165      49.764  15.651  90.790  1.00  0.00           H
    ATOM   4114 HE22 GLN B 165      50.272  16.016  89.165  1.00  0.00           H
    ATOM   4115  N   SER B 166      50.560   8.882  92.330  1.00 53.46           N
    ATOM   4116  CA  SER B 166      50.136   7.499  92.491  1.00 49.95           C
    ATOM   4117  C   SER B 166      49.755   6.711  91.218  1.00 46.57           C
    ATOM   4118  O   SER B 166      50.432   5.740  90.891  1.00 45.66           O
    ATOM   4119  CB  SER B 166      49.033   7.570  93.553  1.00 48.45           C
    ATOM   4120  OG  SER B 166      48.029   8.501  93.172  1.00 48.91           O
    ATOM   4121  H   SER B 166      50.191   9.486  93.050  1.00  0.00           H
    ATOM   4122  HA  SER B 166      50.985   6.972  92.933  1.00  0.00           H
    ATOM   4123  HB2 SER B 166      48.637   6.580  93.757  1.00  0.00           H
    ATOM   4124  HB3 SER B 166      49.454   7.912  94.499  1.00  0.00           H
    ATOM   4125  HG  SER B 166      47.377   8.551  93.876  1.00  0.00           H
    ATOM   4126  N   GLN B 167      48.704   7.139  90.511  1.00 43.90           N
    ATOM   4127  CA  GLN B 167      48.260   6.634  89.204  1.00 46.13           C
    ATOM   4128  C   GLN B 167      49.365   6.642  88.128  1.00 52.11           C
    ATOM   4129  O   GLN B 167      49.562   5.640  87.446  1.00 51.85           O
    ATOM   4130  CB  GLN B 167      46.996   7.397  88.707  1.00 41.96           C
    ATOM   4131  CG  GLN B 167      46.478   8.581  89.563  1.00 43.62           C
    ATOM   4132  CD  GLN B 167      47.425   9.784  89.704  1.00 47.56           C
    ATOM   4133  OE1 GLN B 167      48.613   9.720  89.407  1.00 52.12           O
    ATOM   4134  NE2 GLN B 167      46.901  10.899  90.204  1.00 49.99           N
    ATOM   4135  H   GLN B 167      48.202   7.945  90.853  1.00  0.00           H
    ATOM   4136  HA  GLN B 167      47.978   5.592  89.363  1.00  0.00           H
    ATOM   4137  HB2 GLN B 167      47.124   7.743  87.679  1.00  0.00           H
    ATOM   4138  HB3 GLN B 167      46.182   6.673  88.644  1.00  0.00           H
    ATOM   4139  HG2 GLN B 167      45.547   8.941  89.123  1.00  0.00           H
    ATOM   4140  HG3 GLN B 167      46.216   8.231  90.563  1.00  0.00           H
    ATOM   4141 HE21 GLN B 167      45.927  10.931  90.471  1.00  0.00           H
    ATOM   4142 HE22 GLN B 167      47.474  11.722  90.322  1.00  0.00           H
    ATOM   4143  N   HIS B 168      50.068   7.777  88.019  1.00 51.90           N
    ATOM   4144  CA  HIS B 168      51.124   8.050  87.050  1.00 45.81           C
    ATOM   4145  C   HIS B 168      52.433   7.299  87.273  1.00 49.94           C
    ATOM   4146  O   HIS B 168      53.250   7.291  86.355  1.00 50.20           O
    ATOM   4147  CB  HIS B 168      51.406   9.557  87.040  1.00 37.57           C
    ATOM   4148  CG  HIS B 168      50.254  10.363  86.533  1.00 34.33           C
    ATOM   4149  ND1 HIS B 168      49.838  11.539  87.126  1.00 35.54           N
    ATOM   4150  CD2 HIS B 168      49.418  10.156  85.461  1.00 34.55           C
    ATOM   4151  CE1 HIS B 168      48.806  11.986  86.408  1.00 34.51           C
    ATOM   4152  NE2 HIS B 168      48.513  11.211  85.370  1.00 34.87           N
    ATOM   4153  H   HIS B 168      49.817   8.553  88.620  1.00  0.00           H
    ATOM   4154  HA  HIS B 168      50.766   7.748  86.064  1.00  0.00           H
    ATOM   4155  HB2 HIS B 168      51.691   9.912  88.031  1.00  0.00           H
    ATOM   4156  HB3 HIS B 168      52.237   9.779  86.373  1.00  0.00           H
    ATOM   4157  HD1 HIS B 168      50.233  11.978  87.946  1.00  0.00           H
    ATOM   4158  HD2 HIS B 168      49.409   9.348  84.745  1.00  0.00           H
    ATOM   4159  HE1 HIS B 168      48.277  12.899  86.636  1.00  0.00           H
    ATOM   4160  N   MET B 169      52.632   6.691  88.455  1.00 53.15           N
    ATOM   4161  CA  MET B 169      53.841   5.938  88.795  1.00 55.18           C
    ATOM   4162  C   MET B 169      54.156   4.767  87.855  1.00 51.12           C
    ATOM   4163  O   MET B 169      55.316   4.365  87.765  1.00 54.03           O
    ATOM   4164  CB  MET B 169      53.746   5.394  90.222  1.00 61.17           C
    ATOM   4165  CG  MET B 169      54.055   6.438  91.293  1.00 65.75           C
    ATOM   4166  SD  MET B 169      54.225   5.779  92.962  1.00 70.00           S
    ATOM   4167  CE  MET B 169      55.497   4.517  92.667  1.00 69.42           C
    ATOM   4168  H   MET B 169      51.919   6.743  89.169  1.00  0.00           H
    ATOM   4169  HA  MET B 169      54.685   6.626  88.751  1.00  0.00           H
    ATOM   4170  HB2 MET B 169      52.771   4.943  90.398  1.00  0.00           H
    ATOM   4171  HB3 MET B 169      54.454   4.574  90.323  1.00  0.00           H
    ATOM   4172  HG2 MET B 169      55.012   6.898  91.064  1.00  0.00           H
    ATOM   4173  HG3 MET B 169      53.308   7.222  91.311  1.00  0.00           H
    ATOM   4174  HE1 MET B 169      55.911   4.190  93.621  1.00  0.00           H
    ATOM   4175  HE2 MET B 169      55.079   3.637  92.180  1.00  0.00           H
    ATOM   4176  HE3 MET B 169      56.312   4.913  92.062  1.00  0.00           H
    ATOM   4177  N   THR B 170      53.135   4.239  87.177  1.00 44.59           N
    ATOM   4178  CA  THR B 170      53.297   3.171  86.205  1.00 42.43           C
    ATOM   4179  C   THR B 170      53.722   3.672  84.819  1.00 42.10           C
    ATOM   4180  O   THR B 170      54.279   2.878  84.062  1.00 49.60           O
    ATOM   4181  CB  THR B 170      52.005   2.339  86.019  1.00 47.29           C
    ATOM   4182  OG1 THR B 170      50.905   2.899  86.713  1.00 46.68           O
    ATOM   4183  CG2 THR B 170      52.187   0.919  86.571  1.00 49.66           C
    ATOM   4184  H   THR B 170      52.190   4.572  87.317  1.00  0.00           H
    ATOM   4185  HA  THR B 170      54.092   2.507  86.550  1.00  0.00           H
    ATOM   4186  HB  THR B 170      51.718   2.285  84.970  1.00  0.00           H
    ATOM   4187 HG21 THR B 170      51.290   0.315  86.438  1.00  0.00           H
    ATOM   4188 HG22 THR B 170      53.007   0.400  86.073  1.00  0.00           H
    ATOM   4189 HG23 THR B 170      52.397   0.942  87.639  1.00  0.00           H
    ATOM   4190  HG1 THR B 170      50.132   2.358  86.531  1.00  0.00           H
    ATOM   4191  N   GLU B 171      53.475   4.956  84.506  1.00 33.72           N
    ATOM   4192  CA  GLU B 171      53.869   5.531  83.231  1.00 32.74           C
    ATOM   4193  C   GLU B 171      55.377   5.777  83.245  1.00 27.07           C
    ATOM   4194  O   GLU B 171      55.853   6.600  84.029  1.00 27.34           O
    ATOM   4195  CB  GLU B 171      53.077   6.816  82.945  1.00 33.43           C
    ATOM   4196  CG  GLU B 171      53.199   7.262  81.473  1.00 30.34           C
    ATOM   4197  CD  GLU B 171      52.571   8.624  81.224  1.00 30.81           C
    ATOM   4198  OE1 GLU B 171      51.514   8.884  81.834  1.00 23.99           O
    ATOM   4199  OE2 GLU B 171      53.173   9.403  80.453  1.00 35.76           O1-
    ATOM   4200  H   GLU B 171      53.036   5.574  85.174  1.00  0.00           H
    ATOM   4201  HA  GLU B 171      53.616   4.812  82.450  1.00  0.00           H
    ATOM   4202  HB2 GLU B 171      52.025   6.646  83.182  1.00  0.00           H
    ATOM   4203  HB3 GLU B 171      53.404   7.619  83.608  1.00  0.00           H
    ATOM   4204  HG2 GLU B 171      54.247   7.315  81.178  1.00  0.00           H
    ATOM   4205  HG3 GLU B 171      52.726   6.534  80.815  1.00  0.00           H
    ATOM   4206  N   VAL B 172      56.089   5.059  82.367  1.00 28.52           N
    ATOM   4207  CA  VAL B 172      57.510   5.243  82.111  1.00 28.58           C
    ATOM   4208  C   VAL B 172      57.745   6.714  81.744  1.00 33.29           C
    ATOM   4209  O   VAL B 172      57.116   7.222  80.815  1.00 37.09           O
    ATOM   4210  CB  VAL B 172      57.998   4.335  80.950  1.00 31.31           C
    ATOM   4211  CG1 VAL B 172      59.468   4.565  80.546  1.00 32.85           C
    ATOM   4212  CG2 VAL B 172      57.794   2.850  81.297  1.00 34.23           C
    ATOM   4213  H   VAL B 172      55.610   4.417  81.747  1.00  0.00           H
    ATOM   4214  HA  VAL B 172      58.056   4.991  83.021  1.00  0.00           H
    ATOM   4215  HB  VAL B 172      57.390   4.542  80.068  1.00  0.00           H
    ATOM   4216 HG11 VAL B 172      59.785   3.849  79.786  1.00  0.00           H
    ATOM   4217 HG12 VAL B 172      59.638   5.558  80.129  1.00  0.00           H
    ATOM   4218 HG13 VAL B 172      60.127   4.442  81.403  1.00  0.00           H
    ATOM   4219 HG21 VAL B 172      58.127   2.205  80.484  1.00  0.00           H
    ATOM   4220 HG22 VAL B 172      58.359   2.572  82.186  1.00  0.00           H
    ATOM   4221 HG23 VAL B 172      56.747   2.613  81.484  1.00  0.00           H
    ATOM   4222  N   VAL B 173      58.562   7.389  82.558  1.00 32.75           N
    ATOM   4223  CA  VAL B 173      58.887   8.786  82.346  1.00 25.54           C
    ATOM   4224  C   VAL B 173      59.860   8.857  81.166  1.00 26.92           C
    ATOM   4225  O   VAL B 173      60.827   8.101  81.128  1.00 30.56           O
    ATOM   4226  CB  VAL B 173      59.568   9.413  83.584  1.00 28.58           C
    ATOM   4227  CG1 VAL B 173      59.975  10.860  83.313  1.00 37.16           C
    ATOM   4228  CG2 VAL B 173      58.666   9.396  84.821  1.00 25.01           C
    ATOM   4229  H   VAL B 173      59.029   6.921  83.323  1.00  0.00           H
    ATOM   4230  HA  VAL B 173      57.977   9.345  82.117  1.00  0.00           H
    ATOM   4231  HB  VAL B 173      60.464   8.839  83.828  1.00  0.00           H
    ATOM   4232 HG11 VAL B 173      60.283  11.379  84.218  1.00  0.00           H
    ATOM   4233 HG12 VAL B 173      60.798  10.928  82.604  1.00  0.00           H
    ATOM   4234 HG13 VAL B 173      59.122  11.374  82.880  1.00  0.00           H
    ATOM   4235 HG21 VAL B 173      59.246   9.682  85.694  1.00  0.00           H
    ATOM   4236 HG22 VAL B 173      57.872  10.130  84.740  1.00  0.00           H
    ATOM   4237 HG23 VAL B 173      58.226   8.418  85.009  1.00  0.00           H
    ATOM   4238  N   ARG B 174      59.586   9.742  80.213  1.00 28.08           N
    ATOM   4239  CA  ARG B 174      60.427   9.875  79.035  1.00 29.73           C
    ATOM   4240  C   ARG B 174      60.220  11.250  78.432  1.00 29.91           C
    ATOM   4241  O   ARG B 174      59.304  11.976  78.827  1.00 27.17           O
    ATOM   4242  CB  ARG B 174      60.172   8.724  78.036  1.00 32.67           C
    ATOM   4243  CG  ARG B 174      58.701   8.413  77.723  1.00 29.25           C
    ATOM   4244  CD  ARG B 174      58.480   7.745  76.357  1.00 28.40           C
    ATOM   4245  NE  ARG B 174      59.346   6.574  76.123  1.00 38.01           N
    ATOM   4246  CZ  ARG B 174      59.935   6.283  74.946  1.00 46.71           C
    ATOM   4247  NH1 ARG B 174      59.792   7.082  73.881  1.00 50.11           N
    ATOM   4248  NH2 ARG B 174      60.689   5.185  74.816  1.00 48.85           N1+
    ATOM   4249  H   ARG B 174      58.776  10.345  80.265  1.00  0.00           H
    ATOM   4250  HA  ARG B 174      61.474   9.846  79.330  1.00  0.00           H
    ATOM   4251  HB2 ARG B 174      60.719   8.918  77.112  1.00  0.00           H
    ATOM   4252  HB3 ARG B 174      60.597   7.812  78.445  1.00  0.00           H
    ATOM   4253  HG2 ARG B 174      58.198   7.905  78.541  1.00  0.00           H
    ATOM   4254  HG3 ARG B 174      58.206   9.381  77.633  1.00  0.00           H
    ATOM   4255  HD2 ARG B 174      57.473   7.337  76.311  1.00  0.00           H
    ATOM   4256  HD3 ARG B 174      58.522   8.499  75.571  1.00  0.00           H
    ATOM   4257  HE  ARG B 174      59.462   5.940  76.900  1.00  0.00           H
    ATOM   4258 HH11 ARG B 174      59.286   7.953  73.956  1.00  0.00           H
    ATOM   4259 HH12 ARG B 174      60.283   6.893  73.017  1.00  0.00           H
    ATOM   4260 HH21 ARG B 174      60.811   4.551  75.592  1.00  0.00           H
    ATOM   4261 HH22 ARG B 174      61.147   4.979  73.938  1.00  0.00           H
    ATOM   4262  N   ARG B 175      61.094  11.585  77.474  1.00 30.85           N
    ATOM   4263  CA  ARG B 175      60.959  12.825  76.727  1.00 28.87           C
    ATOM   4264  C   ARG B 175      59.672  12.860  75.925  1.00 28.57           C
    ATOM   4265  O   ARG B 175      59.208  11.831  75.424  1.00 32.29           O
    ATOM   4266  CB  ARG B 175      62.145  13.045  75.793  1.00 25.68           C
    ATOM   4267  CG  ARG B 175      63.396  13.406  76.583  1.00 24.79           C
    ATOM   4268  CD  ARG B 175      64.578  13.670  75.670  1.00 22.43           C
    ATOM   4269  NE  ARG B 175      64.756  15.064  75.283  1.00 19.49           N
    ATOM   4270  CZ  ARG B 175      65.766  15.419  74.480  1.00 18.71           C
    ATOM   4271  NH1 ARG B 175      66.567  14.494  73.944  1.00 13.67           N
    ATOM   4272  NH2 ARG B 175      65.988  16.705  74.224  1.00 22.59           N1+
    ATOM   4273  H   ARG B 175      61.825  10.947  77.190  1.00  0.00           H
    ATOM   4274  HA  ARG B 175      60.938  13.634  77.459  1.00  0.00           H
    ATOM   4275  HB2 ARG B 175      62.314  12.156  75.183  1.00  0.00           H
    ATOM   4276  HB3 ARG B 175      61.934  13.862  75.099  1.00  0.00           H
    ATOM   4277  HG2 ARG B 175      63.204  14.292  77.189  1.00  0.00           H
    ATOM   4278  HG3 ARG B 175      63.649  12.607  77.278  1.00  0.00           H
    ATOM   4279  HD2 ARG B 175      65.496  13.227  76.042  1.00  0.00           H
    ATOM   4280  HD3 ARG B 175      64.360  13.205  74.716  1.00  0.00           H
    ATOM   4281  HE  ARG B 175      64.167  15.763  75.715  1.00  0.00           H
    ATOM   4282 HH11 ARG B 175      66.385  13.514  74.115  1.00  0.00           H
    ATOM   4283 HH12 ARG B 175      67.275  14.745  73.268  1.00  0.00           H
    ATOM   4284 HH21 ARG B 175      65.415  17.424  74.645  1.00  0.00           H
    ATOM   4285 HH22 ARG B 175      66.776  16.983  73.643  1.00  0.00           H
    ATOM   4286  N   CYS B 176      59.153  14.086  75.804  1.00 26.09           N
    ATOM   4287  CA  CYS B 176      58.018  14.392  74.956  1.00 24.20           C
    ATOM   4288  C   CYS B 176      58.359  14.068  73.491  1.00 29.50           C
    ATOM   4289  O   CYS B 176      59.537  14.108  73.123  1.00 31.24           O
    ATOM   4290  CB  CYS B 176      57.540  15.845  75.147  1.00 22.59           C
    ATOM   4291  SG  CYS B 176      58.571  17.189  74.424  1.00 29.72           S
    ATOM   4292  H   CYS B 176      59.620  14.873  76.232  1.00  0.00           H
    ATOM   4293  HA  CYS B 176      57.223  13.730  75.302  1.00  0.00           H
    ATOM   4294  HB2 CYS B 176      56.539  15.956  74.737  1.00  0.00           H
    ATOM   4295  HB3 CYS B 176      57.413  15.991  76.214  1.00  0.00           H
    ATOM   4296  N   PRO B 177      57.332  13.736  72.688  1.00 29.46           N
    ATOM   4297  CA  PRO B 177      57.559  13.311  71.306  1.00 26.76           C
    ATOM   4298  C   PRO B 177      58.272  14.338  70.413  1.00 21.01           C
    ATOM   4299  O   PRO B 177      59.089  13.917  69.598  1.00 19.88           O
    ATOM   4300  CB  PRO B 177      56.181  12.859  70.807  1.00 29.41           C
    ATOM   4301  CG  PRO B 177      55.163  13.391  71.801  1.00 36.94           C
    ATOM   4302  CD  PRO B 177      55.941  13.550  73.096  1.00 34.09           C
    ATOM   4303  HA  PRO B 177      58.205  12.431  71.341  1.00  0.00           H
    ATOM   4304  HB2 PRO B 177      55.961  13.129  69.777  1.00  0.00           H
    ATOM   4305  HB3 PRO B 177      56.163  11.769  70.842  1.00  0.00           H
    ATOM   4306  HG2 PRO B 177      54.836  14.369  71.457  1.00  0.00           H
    ATOM   4307  HG3 PRO B 177      54.278  12.761  71.901  1.00  0.00           H
    ATOM   4308  HD2 PRO B 177      55.535  14.374  73.678  1.00  0.00           H
    ATOM   4309  HD3 PRO B 177      55.873  12.638  73.690  1.00  0.00           H
    ATOM   4310  N   HIS B 178      58.039  15.644  70.622  1.00 15.63           N
    ATOM   4311  CA  HIS B 178      58.763  16.702  69.922  1.00 15.90           C
    ATOM   4312  C   HIS B 178      60.270  16.678  70.217  1.00 18.62           C
    ATOM   4313  O   HIS B 178      61.052  16.558  69.277  1.00 20.61           O
    ATOM   4314  CB  HIS B 178      58.165  18.087  70.222  1.00 21.26           C
    ATOM   4315  CG  HIS B 178      58.972  19.215  69.642  1.00 32.57           C
    ATOM   4316  ND1 HIS B 178      59.112  19.414  68.280  1.00 31.29           N
    ATOM   4317  CD2 HIS B 178      59.763  20.160  70.246  1.00 30.47           C
    ATOM   4318  CE1 HIS B 178      59.942  20.444  68.117  1.00 28.85           C
    ATOM   4319  NE2 HIS B 178      60.356  20.954  69.268  1.00 30.03           N
    ATOM   4320  H   HIS B 178      57.347  15.939  71.300  1.00  0.00           H
    ATOM   4321  HA  HIS B 178      58.643  16.521  68.852  1.00  0.00           H
    ATOM   4322  HB2 HIS B 178      57.158  18.158  69.810  1.00  0.00           H
    ATOM   4323  HB3 HIS B 178      58.081  18.242  71.299  1.00  0.00           H
    ATOM   4324  HD1 HIS B 178      58.699  18.855  67.540  1.00  0.00           H
    ATOM   4325  HD2 HIS B 178      59.954  20.316  71.297  1.00  0.00           H
    ATOM   4326  HE1 HIS B 178      60.255  20.816  67.153  1.00  0.00           H
    ATOM   4327  N   HIS B 179      60.649  16.797  71.502  1.00 24.05           N
    ATOM   4328  CA  HIS B 179      62.058  16.809  71.898  1.00 22.52           C
    ATOM   4329  C   HIS B 179      62.785  15.496  71.650  1.00 21.32           C
    ATOM   4330  O   HIS B 179      63.982  15.545  71.386  1.00 22.05           O
    ATOM   4331  CB  HIS B 179      62.244  17.208  73.361  1.00 23.21           C
    ATOM   4332  CG  HIS B 179      62.043  18.669  73.617  1.00 18.34           C
    ATOM   4333  ND1 HIS B 179      61.359  19.119  74.734  1.00 23.57           N
    ATOM   4334  CD2 HIS B 179      62.456  19.760  72.884  1.00 19.24           C
    ATOM   4335  CE1 HIS B 179      61.376  20.438  74.663  1.00 20.40           C
    ATOM   4336  NE2 HIS B 179      62.011  20.872  73.575  1.00 21.37           N
    ATOM   4337  H   HIS B 179      59.959  16.878  72.237  1.00  0.00           H
    ATOM   4338  HA  HIS B 179      62.562  17.545  71.270  1.00  0.00           H
    ATOM   4339  HB2 HIS B 179      61.567  16.626  73.988  1.00  0.00           H
    ATOM   4340  HB3 HIS B 179      63.250  16.973  73.707  1.00  0.00           H
    ATOM   4341  HD2 HIS B 179      63.011  19.842  71.961  1.00  0.00           H
    ATOM   4342  HE2 HIS B 179      62.179  21.838  73.321  1.00  0.00           H
    ATOM   4343  HE1 HIS B 179      60.938  21.093  75.403  1.00  0.00           H
    ATOM   4344  N   GLU B 180      62.071  14.364  71.696  1.00 21.22           N
    ATOM   4345  CA  GLU B 180      62.635  13.095  71.266  1.00 16.89           C
    ATOM   4346  C   GLU B 180      62.933  13.086  69.764  1.00 17.13           C
    ATOM   4347  O   GLU B 180      64.011  12.644  69.368  1.00 17.82           O
    ATOM   4348  CB  GLU B 180      61.748  11.916  71.683  1.00 16.33           C
    ATOM   4349  CG  GLU B 180      62.207  10.595  71.027  1.00 18.82           C
    ATOM   4350  CD  GLU B 180      61.513   9.352  71.545  1.00 33.66           C
    ATOM   4351  OE1 GLU B 180      60.762   9.464  72.536  1.00 40.05           O
    ATOM   4352  OE2 GLU B 180      61.762   8.292  70.931  1.00 37.17           O1-
    ATOM   4353  H   GLU B 180      61.092  14.376  71.956  1.00  0.00           H
    ATOM   4354  HA  GLU B 180      63.589  12.963  71.781  1.00  0.00           H
    ATOM   4355  HB2 GLU B 180      61.781  11.827  72.771  1.00  0.00           H
    ATOM   4356  HB3 GLU B 180      60.708  12.115  71.422  1.00  0.00           H
    ATOM   4357  HG2 GLU B 180      62.039  10.610  69.949  1.00  0.00           H
    ATOM   4358  HG3 GLU B 180      63.277  10.455  71.183  1.00  0.00           H
    ATOM   4359  N   ARG B 181      61.973  13.572  68.964  1.00 20.13           N
    ATOM   4360  CA  ARG B 181      62.109  13.657  67.518  1.00 21.35           C
    ATOM   4361  C   ARG B 181      63.291  14.528  67.092  1.00 26.60           C
    ATOM   4362  O   ARG B 181      64.086  14.094  66.266  1.00 24.46           O
    ATOM   4363  CB  ARG B 181      60.789  14.137  66.895  1.00 22.85           C
    ATOM   4364  CG  ARG B 181      60.912  14.439  65.398  1.00 24.08           C
    ATOM   4365  CD  ARG B 181      59.568  14.726  64.748  1.00 21.66           C
    ATOM   4366  NE  ARG B 181      59.752  15.081  63.343  1.00 22.57           N
    ATOM   4367  CZ  ARG B 181      60.064  16.293  62.866  1.00 29.45           C
    ATOM   4368  NH1 ARG B 181      60.184  17.356  63.673  1.00 32.97           N
    ATOM   4369  NH2 ARG B 181      60.251  16.427  61.551  1.00 29.15           N1+
    ATOM   4370  H   ARG B 181      61.106  13.913  69.359  1.00  0.00           H
    ATOM   4371  HA  ARG B 181      62.313  12.652  67.154  1.00  0.00           H
    ATOM   4372  HB2 ARG B 181      60.021  13.380  67.055  1.00  0.00           H
    ATOM   4373  HB3 ARG B 181      60.434  15.038  67.390  1.00  0.00           H
    ATOM   4374  HG2 ARG B 181      61.622  15.230  65.152  1.00  0.00           H
    ATOM   4375  HG3 ARG B 181      61.307  13.529  64.948  1.00  0.00           H
    ATOM   4376  HD2 ARG B 181      59.035  13.776  64.695  1.00  0.00           H
    ATOM   4377  HD3 ARG B 181      58.926  15.394  65.316  1.00  0.00           H
    ATOM   4378  HE  ARG B 181      59.711  14.309  62.691  1.00  0.00           H
    ATOM   4379 HH11 ARG B 181      60.004  17.258  64.662  1.00  0.00           H
    ATOM   4380 HH12 ARG B 181      60.485  18.259  63.322  1.00  0.00           H
    ATOM   4381 HH21 ARG B 181      60.167  15.629  60.931  1.00  0.00           H
    ATOM   4382 HH22 ARG B 181      60.487  17.331  61.159  1.00  0.00           H
    ATOM   4383  N   CYS B 182      63.374  15.724  67.681  1.00 23.97           N
    ATOM   4384  CA  CYS B 182      64.385  16.729  67.383  1.00 21.13           C
    ATOM   4385  C   CYS B 182      65.573  16.625  68.350  1.00 20.63           C
    ATOM   4386  O   CYS B 182      66.290  17.607  68.529  1.00 21.86           O
    ATOM   4387  CB  CYS B 182      63.768  18.134  67.428  1.00 18.92           C
    ATOM   4388  SG  CYS B 182      62.376  18.257  66.282  1.00 23.20           S
    ATOM   4389  H   CYS B 182      62.668  15.990  68.357  1.00  0.00           H
    ATOM   4390  HA  CYS B 182      64.787  16.571  66.382  1.00  0.00           H
    ATOM   4391  HB2 CYS B 182      63.399  18.365  68.427  1.00  0.00           H
    ATOM   4392  HB3 CYS B 182      64.499  18.889  67.145  1.00  0.00           H
    ATOM   4393  HG  CYS B 182      61.549  17.484  66.992  1.00  0.00           H
    ATOM   4394  N   SER B 183      65.767  15.455  68.978  1.00 18.80           N
    ATOM   4395  CA  SER B 183      66.832  15.237  69.945  1.00 17.25           C
    ATOM   4396  C   SER B 183      68.220  15.378  69.316  1.00 17.55           C
    ATOM   4397  O   SER B 183      68.424  15.003  68.160  1.00 20.68           O
    ATOM   4398  CB  SER B 183      66.668  13.853  70.577  1.00 19.79           C
    ATOM   4399  OG  SER B 183      67.653  13.618  71.557  1.00 16.84           O
    ATOM   4400  H   SER B 183      65.140  14.679  68.812  1.00  0.00           H
    ATOM   4401  HA  SER B 183      66.724  15.986  70.732  1.00  0.00           H
    ATOM   4402  HB2 SER B 183      65.700  13.743  71.058  1.00  0.00           H
    ATOM   4403  HB3 SER B 183      66.727  13.097  69.796  1.00  0.00           H
    ATOM   4404  HG  SER B 183      68.519  13.567  71.134  1.00  0.00           H
    ATOM   4405  N   ASP B 184      69.144  15.877  70.142  1.00 19.17           N
    ATOM   4406  CA  ASP B 184      70.574  15.791  69.904  1.00 20.36           C
    ATOM   4407  C   ASP B 184      71.096  14.561  70.662  1.00 17.67           C
    ATOM   4408  O   ASP B 184      70.330  13.808  71.261  1.00 20.51           O
    ATOM   4409  CB  ASP B 184      71.331  17.081  70.321  1.00 28.39           C
    ATOM   4410  CG  ASP B 184      71.515  17.360  71.825  1.00 28.54           C
    ATOM   4411  OD1 ASP B 184      70.865  16.701  72.666  1.00 27.47           O
    ATOM   4412  OD2 ASP B 184      72.337  18.254  72.114  1.00 20.85           O1-
    ATOM   4413  H   ASP B 184      68.872  16.189  71.064  1.00  0.00           H
    ATOM   4414  HA  ASP B 184      70.761  15.628  68.845  1.00  0.00           H
    ATOM   4415  HB2 ASP B 184      72.327  17.040  69.877  1.00  0.00           H
    ATOM   4416  HB3 ASP B 184      70.834  17.939  69.871  1.00  0.00           H
    ATOM   4417  N   SER B 185      72.415  14.421  70.700  1.00 18.76           N
    ATOM   4418  CA  SER B 185      73.071  13.463  71.566  1.00 15.62           C
    ATOM   4419  C   SER B 185      74.510  13.920  71.792  1.00 16.75           C
    ATOM   4420  O   SER B 185      74.942  14.932  71.244  1.00 14.68           O
    ATOM   4421  CB  SER B 185      72.902  12.044  70.977  1.00 19.43           C
    ATOM   4422  OG  SER B 185      73.614  11.044  71.671  1.00 22.77           O
    ATOM   4423  H   SER B 185      73.016  15.075  70.218  1.00  0.00           H
    ATOM   4424  HA  SER B 185      72.596  13.490  72.550  1.00  0.00           H
    ATOM   4425  HB2 SER B 185      71.857  11.753  71.041  1.00  0.00           H
    ATOM   4426  HB3 SER B 185      73.175  12.016  69.922  1.00  0.00           H
    ATOM   4427  HG  SER B 185      73.030  10.673  72.344  1.00  0.00           H
    ATOM   4428  N   ASP B 186      75.216  13.169  72.631  1.00 23.12           N
    ATOM   4429  CA  ASP B 186      76.613  13.377  72.989  1.00 25.90           C
    ATOM   4430  C   ASP B 186      77.474  12.164  72.585  1.00 29.18           C
    ATOM   4431  O   ASP B 186      78.602  12.041  73.057  1.00 31.16           O
    ATOM   4432  CB  ASP B 186      76.756  13.727  74.492  1.00 23.68           C
    ATOM   4433  CG  ASP B 186      76.004  12.800  75.459  1.00 22.00           C
    ATOM   4434  OD1 ASP B 186      75.644  11.675  75.047  1.00 25.37           O
    ATOM   4435  OD2 ASP B 186      75.850  13.215  76.628  1.00 26.50           O1-
    ATOM   4436  H   ASP B 186      74.781  12.358  73.052  1.00  0.00           H
    ATOM   4437  HA  ASP B 186      77.032  14.202  72.417  1.00  0.00           H
    ATOM   4438  HB2 ASP B 186      77.806  13.763  74.784  1.00  0.00           H
    ATOM   4439  HB3 ASP B 186      76.355  14.732  74.629  1.00  0.00           H
    ATOM   4440  N   GLY B 187      76.925  11.284  71.734  1.00 27.02           N
    ATOM   4441  CA  GLY B 187      77.548  10.034  71.323  1.00 25.71           C
    ATOM   4442  C   GLY B 187      77.178   8.886  72.269  1.00 26.99           C
    ATOM   4443  O   GLY B 187      77.576   7.756  71.995  1.00 28.01           O
    ATOM   4444  H   GLY B 187      76.001  11.473  71.368  1.00  0.00           H
    ATOM   4445  HA2 GLY B 187      77.189   9.778  70.328  1.00  0.00           H
    ATOM   4446  HA3 GLY B 187      78.633  10.129  71.256  1.00  0.00           H
    ATOM   4447  N   LEU B 188      76.435   9.147  73.359  1.00 33.14           N
    ATOM   4448  CA  LEU B 188      76.067   8.144  74.359  1.00 32.22           C
    ATOM   4449  C   LEU B 188      74.561   8.171  74.609  1.00 24.84           C
    ATOM   4450  O   LEU B 188      73.919   7.129  74.490  1.00 28.90           O
    ATOM   4451  CB  LEU B 188      76.799   8.418  75.691  1.00 32.80           C
    ATOM   4452  CG  LEU B 188      78.339   8.412  75.631  1.00 32.68           C
    ATOM   4453  CD1 LEU B 188      78.955   9.295  76.732  1.00 31.55           C
    ATOM   4454  CD2 LEU B 188      78.908   6.984  75.645  1.00 30.96           C
    ATOM   4455  H   LEU B 188      76.149  10.098  73.556  1.00  0.00           H
    ATOM   4456  HA  LEU B 188      76.320   7.138  74.022  1.00  0.00           H
    ATOM   4457  HB2 LEU B 188      76.479   9.395  76.053  1.00  0.00           H
    ATOM   4458  HB3 LEU B 188      76.475   7.705  76.451  1.00  0.00           H
    ATOM   4459  HG  LEU B 188      78.649   8.868  74.692  1.00  0.00           H
    ATOM   4460 HD11 LEU B 188      79.606   8.728  77.396  1.00  0.00           H
    ATOM   4461 HD12 LEU B 188      79.579  10.073  76.289  1.00  0.00           H
    ATOM   4462 HD13 LEU B 188      78.212   9.804  77.344  1.00  0.00           H
    ATOM   4463 HD21 LEU B 188      79.859   6.942  75.112  1.00  0.00           H
    ATOM   4464 HD22 LEU B 188      79.077   6.621  76.659  1.00  0.00           H
    ATOM   4465 HD23 LEU B 188      78.235   6.278  75.158  1.00  0.00           H
    ATOM   4466  N   ALA B 189      74.044   9.351  74.986  1.00 18.44           N
    ATOM   4467  CA  ALA B 189      72.670   9.568  75.414  1.00 19.81           C
    ATOM   4468  C   ALA B 189      71.671   9.259  74.293  1.00 29.31           C
    ATOM   4469  O   ALA B 189      71.691   9.947  73.274  1.00 39.04           O
    ATOM   4470  CB  ALA B 189      72.530  11.023  75.854  1.00 16.62           C
    ATOM   4471  H   ALA B 189      74.653  10.161  75.051  1.00  0.00           H
    ATOM   4472  HA  ALA B 189      72.485   8.968  76.298  1.00  0.00           H
    ATOM   4473  HB1 ALA B 189      71.514  11.209  76.198  1.00  0.00           H
    ATOM   4474  HB2 ALA B 189      73.208  11.251  76.676  1.00  0.00           H
    ATOM   4475  HB3 ALA B 189      72.747  11.724  75.046  1.00  0.00           H
    ATOM   4476  N   PRO B 190      70.813   8.235  74.478  1.00 31.51           N
    ATOM   4477  CA  PRO B 190      69.728   7.982  73.525  1.00 29.03           C
    ATOM   4478  C   PRO B 190      68.642   9.076  73.610  1.00 27.56           C
    ATOM   4479  O   PRO B 190      68.492   9.699  74.662  1.00 29.13           O
    ATOM   4480  CB  PRO B 190      69.198   6.599  73.930  1.00 31.02           C
    ATOM   4481  CG  PRO B 190      70.232   6.001  74.866  1.00 28.43           C
    ATOM   4482  CD  PRO B 190      70.851   7.218  75.522  1.00 29.62           C
    ATOM   4483  HA  PRO B 190      70.181   7.920  72.535  1.00  0.00           H
    ATOM   4484  HB2 PRO B 190      68.256   6.700  74.454  1.00  0.00           H
    ATOM   4485  HB3 PRO B 190      69.011   5.957  73.069  1.00  0.00           H
    ATOM   4486  HG2 PRO B 190      69.817   5.286  75.576  1.00  0.00           H
    ATOM   4487  HG3 PRO B 190      70.993   5.485  74.279  1.00  0.00           H
    ATOM   4488  HD2 PRO B 190      70.253   7.557  76.369  1.00  0.00           H
    ATOM   4489  HD3 PRO B 190      71.849   6.970  75.869  1.00  0.00           H
    ATOM   4490  N   PRO B 191      67.908   9.294  72.501  1.00 22.30           N
    ATOM   4491  CA  PRO B 191      66.980  10.427  72.395  1.00 17.35           C
    ATOM   4492  C   PRO B 191      65.701  10.349  73.240  1.00 16.72           C
    ATOM   4493  O   PRO B 191      65.069  11.382  73.425  1.00 20.84           O
    ATOM   4494  CB  PRO B 191      66.696  10.517  70.890  1.00 22.73           C
    ATOM   4495  CG  PRO B 191      66.933   9.123  70.346  1.00 24.53           C
    ATOM   4496  CD  PRO B 191      68.041   8.576  71.233  1.00 20.27           C
    ATOM   4497  HA  PRO B 191      67.501  11.330  72.712  1.00  0.00           H
    ATOM   4498  HB2 PRO B 191      65.708  10.906  70.639  1.00  0.00           H
    ATOM   4499  HB3 PRO B 191      67.437  11.168  70.434  1.00  0.00           H
    ATOM   4500  HG2 PRO B 191      66.030   8.527  70.492  1.00  0.00           H
    ATOM   4501  HG3 PRO B 191      67.182   9.101  69.285  1.00  0.00           H
    ATOM   4502  HD2 PRO B 191      67.961   7.495  71.347  1.00  0.00           H
    ATOM   4503  HD3 PRO B 191      69.013   8.798  70.795  1.00  0.00           H
    ATOM   4504  N   GLN B 192      65.346   9.162  73.750  1.00 20.25           N
    ATOM   4505  CA  GLN B 192      64.178   8.952  74.598  1.00 20.75           C
    ATOM   4506  C   GLN B 192      64.430   9.281  76.085  1.00 23.62           C
    ATOM   4507  O   GLN B 192      63.466   9.589  76.785  1.00 24.88           O
    ATOM   4508  CB  GLN B 192      63.618   7.526  74.377  1.00 27.58           C
    ATOM   4509  CG  GLN B 192      64.470   6.326  74.859  1.00 33.55           C
    ATOM   4510  CD  GLN B 192      65.599   5.872  73.925  1.00 34.18           C
    ATOM   4511  OE1 GLN B 192      65.952   6.535  72.952  1.00 34.49           O
    ATOM   4512  NE2 GLN B 192      66.190   4.719  74.242  1.00 27.28           N
    ATOM   4513  H   GLN B 192      65.875   8.333  73.509  1.00  0.00           H
    ATOM   4514  HA  GLN B 192      63.394   9.637  74.265  1.00  0.00           H
    ATOM   4515  HB2 GLN B 192      62.666   7.474  74.907  1.00  0.00           H
    ATOM   4516  HB3 GLN B 192      63.352   7.396  73.327  1.00  0.00           H
    ATOM   4517  HG2 GLN B 192      64.861   6.503  75.860  1.00  0.00           H
    ATOM   4518  HG3 GLN B 192      63.800   5.471  74.965  1.00  0.00           H
    ATOM   4519 HE21 GLN B 192      65.878   4.197  75.054  1.00  0.00           H
    ATOM   4520 HE22 GLN B 192      66.935   4.352  73.669  1.00  0.00           H
    ATOM   4521  N   HIS B 193      65.695   9.229  76.544  1.00 20.87           N
    ATOM   4522  CA  HIS B 193      66.070   9.469  77.943  1.00 17.99           C
    ATOM   4523  C   HIS B 193      65.909  10.932  78.355  1.00 20.71           C
    ATOM   4524  O   HIS B 193      66.500  11.808  77.734  1.00 26.57           O
    ATOM   4525  CB  HIS B 193      67.515   9.013  78.201  1.00 15.22           C
    ATOM   4526  CG  HIS B 193      67.679   7.519  78.191  1.00 22.40           C
    ATOM   4527  ND1 HIS B 193      68.088   6.791  79.308  1.00 24.99           N
    ATOM   4528  CD2 HIS B 193      67.454   6.641  77.158  1.00 27.24           C
    ATOM   4529  CE1 HIS B 193      68.140   5.534  78.900  1.00 23.23           C
    ATOM   4530  NE2 HIS B 193      67.758   5.382  77.634  1.00 27.55           N
    ATOM   4531  H   HIS B 193      66.447   9.008  75.908  1.00  0.00           H
    ATOM   4532  HA  HIS B 193      65.424   8.869  78.584  1.00  0.00           H
    ATOM   4533  HB2 HIS B 193      68.186   9.432  77.451  1.00  0.00           H
    ATOM   4534  HB3 HIS B 193      67.871   9.391  79.160  1.00  0.00           H
    ATOM   4535  HD2 HIS B 193      67.090   6.813  76.162  1.00  0.00           H
    ATOM   4536  HE2 HIS B 193      67.670   4.508  77.133  1.00  0.00           H
    ATOM   4537  HE1 HIS B 193      68.457   4.723  79.534  1.00  0.00           H
    ATOM   4538  N   LEU B 194      65.152  11.166  79.433  1.00 16.90           N
    ATOM   4539  CA  LEU B 194      65.021  12.480  80.054  1.00 18.13           C
    ATOM   4540  C   LEU B 194      66.321  12.898  80.748  1.00 25.20           C
    ATOM   4541  O   LEU B 194      66.764  14.024  80.551  1.00 26.19           O
    ATOM   4542  CB  LEU B 194      63.840  12.453  81.047  1.00 18.89           C
    ATOM   4543  CG  LEU B 194      63.693  13.685  81.973  1.00 19.08           C
    ATOM   4544  CD1 LEU B 194      63.156  14.914  81.221  1.00 15.79           C
    ATOM   4545  CD2 LEU B 194      62.878  13.343  83.229  1.00 19.93           C
    ATOM   4546  H   LEU B 194      64.666  10.402  79.889  1.00  0.00           H
    ATOM   4547  HA  LEU B 194      64.804  13.216  79.277  1.00  0.00           H
    ATOM   4548  HB2 LEU B 194      62.904  12.270  80.517  1.00  0.00           H
    ATOM   4549  HB3 LEU B 194      63.988  11.589  81.695  1.00  0.00           H
    ATOM   4550  HG  LEU B 194      64.663  13.977  82.370  1.00  0.00           H
    ATOM   4551 HD11 LEU B 194      63.711  15.812  81.495  1.00  0.00           H
    ATOM   4552 HD12 LEU B 194      63.250  14.796  80.142  1.00  0.00           H
    ATOM   4553 HD13 LEU B 194      62.103  15.101  81.436  1.00  0.00           H
    ATOM   4554 HD21 LEU B 194      62.128  14.099  83.457  1.00  0.00           H
    ATOM   4555 HD22 LEU B 194      62.376  12.383  83.150  1.00  0.00           H
    ATOM   4556 HD23 LEU B 194      63.529  13.285  84.102  1.00  0.00           H
    ATOM   4557  N   ILE B 195      66.875  12.005  81.580  1.00 28.87           N
    ATOM   4558  CA  ILE B 195      68.039  12.315  82.396  1.00 28.12           C
    ATOM   4559  C   ILE B 195      69.297  11.856  81.648  1.00 28.37           C
    ATOM   4560  O   ILE B 195      69.367  10.707  81.217  1.00 27.96           O
    ATOM   4561  CB  ILE B 195      68.012  11.604  83.777  1.00 23.00           C
    ATOM   4562  CG1 ILE B 195      66.652  11.795  84.484  1.00 26.77           C
    ATOM   4563  CG2 ILE B 195      69.166  12.097  84.684  1.00 17.98           C
    ATOM   4564  CD1 ILE B 195      66.558  11.074  85.833  1.00 25.71           C
    ATOM   4565  H   ILE B 195      66.469  11.087  81.691  1.00  0.00           H
    ATOM   4566  HA  ILE B 195      68.083  13.387  82.585  1.00  0.00           H
    ATOM   4567  HB  ILE B 195      68.146  10.532  83.626  1.00  0.00           H
    ATOM   4568 HG12 ILE B 195      66.467  12.857  84.624  1.00  0.00           H
    ATOM   4569 HG13 ILE B 195      65.836  11.432  83.859  1.00  0.00           H
    ATOM   4570 HG21 ILE B 195      69.186  11.574  85.640  1.00  0.00           H
    ATOM   4571 HG22 ILE B 195      70.145  11.933  84.236  1.00  0.00           H
    ATOM   4572 HG23 ILE B 195      69.076  13.164  84.893  1.00  0.00           H
    ATOM   4573 HD11 ILE B 195      65.522  10.908  86.121  1.00  0.00           H
    ATOM   4574 HD12 ILE B 195      67.053  10.102  85.809  1.00  0.00           H
    ATOM   4575 HD13 ILE B 195      67.020  11.670  86.619  1.00  0.00           H
    ATOM   4576  N   ARG B 196      70.269  12.760  81.534  1.00 26.54           N
    ATOM   4577  CA  ARG B 196      71.611  12.463  81.050  1.00 16.72           C
    ATOM   4578  C   ARG B 196      72.610  12.761  82.157  1.00 15.60           C
    ATOM   4579  O   ARG B 196      72.318  13.582  83.020  1.00 14.47           O
    ATOM   4580  CB  ARG B 196      71.935  13.345  79.837  1.00 20.35           C
    ATOM   4581  CG  ARG B 196      71.086  13.045  78.601  1.00 25.12           C
    ATOM   4582  CD  ARG B 196      71.564  13.887  77.414  1.00 25.60           C
    ATOM   4583  NE  ARG B 196      70.658  13.761  76.274  1.00 25.34           N
    ATOM   4584  CZ  ARG B 196      70.731  14.457  75.134  1.00 23.18           C
    ATOM   4585  NH1 ARG B 196      71.708  15.346  74.925  1.00 13.29           N
    ATOM   4586  NH2 ARG B 196      69.803  14.262  74.192  1.00 19.29           N1+
    ATOM   4587  H   ARG B 196      70.124  13.696  81.893  1.00  0.00           H
    ATOM   4588  HA  ARG B 196      71.717  11.414  80.786  1.00  0.00           H
    ATOM   4589  HB2 ARG B 196      71.825  14.397  80.106  1.00  0.00           H
    ATOM   4590  HB3 ARG B 196      72.985  13.213  79.565  1.00  0.00           H
    ATOM   4591  HG2 ARG B 196      71.228  11.990  78.367  1.00  0.00           H
    ATOM   4592  HG3 ARG B 196      70.019  13.180  78.781  1.00  0.00           H
    ATOM   4593  HD2 ARG B 196      71.709  14.929  77.705  1.00  0.00           H
    ATOM   4594  HD3 ARG B 196      72.533  13.524  77.072  1.00  0.00           H
    ATOM   4595  HE  ARG B 196      69.903  13.087  76.361  1.00  0.00           H
    ATOM   4596 HH11 ARG B 196      72.423  15.515  75.619  1.00  0.00           H
    ATOM   4597 HH12 ARG B 196      71.686  15.912  74.079  1.00  0.00           H
    ATOM   4598 HH21 ARG B 196      69.058  13.596  74.354  1.00  0.00           H
    ATOM   4599 HH22 ARG B 196      69.883  14.735  73.301  1.00  0.00           H
    ATOM   4600  N   VAL B 197      73.785  12.126  82.075  1.00 18.19           N
    ATOM   4601  CA  VAL B 197      74.940  12.522  82.868  1.00 22.10           C
    ATOM   4602  C   VAL B 197      75.910  13.270  81.949  1.00 22.20           C
    ATOM   4603  O   VAL B 197      76.255  12.786  80.872  1.00 21.53           O
    ATOM   4604  CB  VAL B 197      75.673  11.330  83.533  1.00 21.86           C
    ATOM   4605  CG1 VAL B 197      77.022  11.723  84.168  1.00 16.81           C
    ATOM   4606  CG2 VAL B 197      74.794  10.684  84.613  1.00 26.51           C
    ATOM   4607  H   VAL B 197      73.954  11.465  81.329  1.00  0.00           H
    ATOM   4608  HA  VAL B 197      74.631  13.188  83.671  1.00  0.00           H
    ATOM   4609  HB  VAL B 197      75.874  10.579  82.774  1.00  0.00           H
    ATOM   4610 HG11 VAL B 197      77.444  10.899  84.736  1.00  0.00           H
    ATOM   4611 HG12 VAL B 197      77.765  12.010  83.425  1.00  0.00           H
    ATOM   4612 HG13 VAL B 197      76.909  12.555  84.861  1.00  0.00           H
    ATOM   4613 HG21 VAL B 197      75.251   9.771  84.998  1.00  0.00           H
    ATOM   4614 HG22 VAL B 197      74.670  11.366  85.452  1.00  0.00           H
    ATOM   4615 HG23 VAL B 197      73.806  10.422  84.238  1.00  0.00           H
    ATOM   4616  N   GLU B 198      76.326  14.440  82.430  1.00 22.59           N
    ATOM   4617  CA  GLU B 198      77.264  15.337  81.788  1.00 19.30           C
    ATOM   4618  C   GLU B 198      78.701  14.995  82.195  1.00 24.98           C
    ATOM   4619  O   GLU B 198      78.949  14.738  83.374  1.00 29.69           O
    ATOM   4620  CB  GLU B 198      76.876  16.741  82.259  1.00 13.85           C
    ATOM   4621  CG  GLU B 198      77.803  17.877  81.817  1.00 12.46           C
    ATOM   4622  CD  GLU B 198      77.389  19.212  82.415  1.00 17.38           C
    ATOM   4623  OE1 GLU B 198      76.254  19.333  82.930  1.00 18.03           O
    ATOM   4624  OE2 GLU B 198      78.229  20.129  82.321  1.00 23.24           O1-
    ATOM   4625  H   GLU B 198      76.005  14.731  83.345  1.00  0.00           H
    ATOM   4626  HA  GLU B 198      77.161  15.279  80.702  1.00  0.00           H
    ATOM   4627  HB2 GLU B 198      75.872  16.952  81.911  1.00  0.00           H
    ATOM   4628  HB3 GLU B 198      76.809  16.744  83.348  1.00  0.00           H
    ATOM   4629  HG2 GLU B 198      78.832  17.694  82.126  1.00  0.00           H
    ATOM   4630  HG3 GLU B 198      77.805  17.961  80.729  1.00  0.00           H
    ATOM   4631  N   GLY B 199      79.621  15.070  81.221  1.00 31.09           N
    ATOM   4632  CA  GLY B 199      81.067  15.003  81.434  1.00 30.25           C
    ATOM   4633  C   GLY B 199      81.553  13.693  82.076  1.00 30.96           C
    ATOM   4634  O   GLY B 199      82.472  13.740  82.893  1.00 36.26           O
    ATOM   4635  H   GLY B 199      79.317  15.282  80.281  1.00  0.00           H
    ATOM   4636  HA2 GLY B 199      81.566  15.121  80.472  1.00  0.00           H
    ATOM   4637  HA3 GLY B 199      81.374  15.848  82.052  1.00  0.00           H
    ATOM   4638  N   ASN B 200      80.947  12.543  81.741  1.00 27.51           N
    ATOM   4639  CA  ASN B 200      81.367  11.245  82.264  1.00 24.54           C
    ATOM   4640  C   ASN B 200      81.360  10.226  81.123  1.00 32.81           C
    ATOM   4641  O   ASN B 200      80.340   9.587  80.871  1.00 37.54           O
    ATOM   4642  CB  ASN B 200      80.482  10.823  83.460  1.00 23.24           C
    ATOM   4643  CG  ASN B 200      81.067   9.662  84.275  1.00 28.69           C
    ATOM   4644  OD1 ASN B 200      81.586   8.693  83.724  1.00 34.87           O
    ATOM   4645  ND2 ASN B 200      80.939   9.729  85.601  1.00 26.71           N
    ATOM   4646  H   ASN B 200      80.187  12.541  81.073  1.00  0.00           H
    ATOM   4647  HA  ASN B 200      82.404  11.267  82.611  1.00  0.00           H
    ATOM   4648  HB2 ASN B 200      80.358  11.677  84.130  1.00  0.00           H
    ATOM   4649  HB3 ASN B 200      79.481  10.548  83.128  1.00  0.00           H
    ATOM   4650 HD21 ASN B 200      80.500  10.526  86.036  1.00  0.00           H
    ATOM   4651 HD22 ASN B 200      81.291   8.983  86.182  1.00  0.00           H
    ATOM   4652  N   LEU B 201      82.529  10.059  80.493  1.00 37.28           N
    ATOM   4653  CA  LEU B 201      82.796   9.083  79.437  1.00 38.49           C
    ATOM   4654  C   LEU B 201      82.566   7.601  79.797  1.00 36.19           C
    ATOM   4655  O   LEU B 201      82.711   6.765  78.908  1.00 36.47           O
    ATOM   4656  CB  LEU B 201      84.248   9.282  78.941  1.00 42.71           C
    ATOM   4657  CG  LEU B 201      84.392  10.320  77.816  1.00 44.71           C
    ATOM   4658  CD1 LEU B 201      85.872  10.468  77.424  1.00 45.29           C
    ATOM   4659  CD2 LEU B 201      83.529   9.976  76.584  1.00 46.18           C
    ATOM   4660  H   LEU B 201      83.318  10.625  80.772  1.00  0.00           H
    ATOM   4661  HA  LEU B 201      82.085   9.294  78.639  1.00  0.00           H
    ATOM   4662  HB2 LEU B 201      84.893   9.541  79.783  1.00  0.00           H
    ATOM   4663  HB3 LEU B 201      84.659   8.349  78.553  1.00  0.00           H
    ATOM   4664  HG  LEU B 201      84.058  11.282  78.208  1.00  0.00           H
    ATOM   4665 HD11 LEU B 201      86.055  11.439  76.964  1.00  0.00           H
    ATOM   4666 HD12 LEU B 201      86.530  10.392  78.291  1.00  0.00           H
    ATOM   4667 HD13 LEU B 201      86.180   9.700  76.713  1.00  0.00           H
    ATOM   4668 HD21 LEU B 201      84.093  10.055  75.654  1.00  0.00           H
    ATOM   4669 HD22 LEU B 201      83.129   8.962  76.623  1.00  0.00           H
    ATOM   4670 HD23 LEU B 201      82.683  10.660  76.501  1.00  0.00           H
    ATOM   4671  N   ARG B 202      82.223   7.277  81.054  1.00 36.66           N
    ATOM   4672  CA  ARG B 202      81.906   5.914  81.482  1.00 35.74           C
    ATOM   4673  C   ARG B 202      80.422   5.712  81.793  1.00 31.70           C
    ATOM   4674  O   ARG B 202      80.067   4.648  82.298  1.00 39.27           O
    ATOM   4675  CB  ARG B 202      82.787   5.523  82.676  1.00 37.54           C
    ATOM   4676  CG  ARG B 202      84.243   5.282  82.256  1.00 39.07           C
    ATOM   4677  CD  ARG B 202      85.131   4.851  83.426  1.00 41.45           C
    ATOM   4678  NE  ARG B 202      84.958   3.442  83.810  1.00 47.69           N
    ATOM   4679  CZ  ARG B 202      85.015   2.933  85.056  1.00 48.15           C
    ATOM   4680  NH1 ARG B 202      85.120   3.701  86.149  1.00 50.03           N
    ATOM   4681  NH2 ARG B 202      84.969   1.606  85.215  1.00 44.14           N1+
    ATOM   4682  H   ARG B 202      82.118   8.001  81.752  1.00  0.00           H
    ATOM   4683  HA  ARG B 202      82.115   5.203  80.681  1.00  0.00           H
    ATOM   4684  HB2 ARG B 202      82.727   6.287  83.452  1.00  0.00           H
    ATOM   4685  HB3 ARG B 202      82.429   4.600  83.131  1.00  0.00           H
    ATOM   4686  HG2 ARG B 202      84.234   4.480  81.516  1.00  0.00           H
    ATOM   4687  HG3 ARG B 202      84.678   6.149  81.758  1.00  0.00           H
    ATOM   4688  HD2 ARG B 202      86.171   5.167  83.341  1.00  0.00           H
    ATOM   4689  HD3 ARG B 202      84.727   5.394  84.267  1.00  0.00           H
    ATOM   4690  HE  ARG B 202      84.886   2.796  83.037  1.00  0.00           H
    ATOM   4691 HH11 ARG B 202      85.120   4.715  86.070  1.00  0.00           H
    ATOM   4692 HH12 ARG B 202      85.175   3.287  87.068  1.00  0.00           H
    ATOM   4693 HH21 ARG B 202      84.888   0.999  84.412  1.00  0.00           H
    ATOM   4694 HH22 ARG B 202      85.019   1.198  86.137  1.00  0.00           H
    ATOM   4695  N   VAL B 203      79.576   6.691  81.450  1.00 30.12           N
    ATOM   4696  CA  VAL B 203      78.126   6.589  81.544  1.00 34.12           C
    ATOM   4697  C   VAL B 203      77.558   5.454  80.661  1.00 35.12           C
    ATOM   4698  O   VAL B 203      78.089   5.184  79.583  1.00 34.87           O
    ATOM   4699  CB  VAL B 203      77.490   7.947  81.139  1.00 28.42           C
    ATOM   4700  CG1 VAL B 203      77.597   8.271  79.642  1.00 27.37           C
    ATOM   4701  CG2 VAL B 203      76.037   8.069  81.572  1.00 23.30           C
    ATOM   4702  H   VAL B 203      79.937   7.542  81.040  1.00  0.00           H
    ATOM   4703  HA  VAL B 203      77.880   6.356  82.579  1.00  0.00           H
    ATOM   4704  HB  VAL B 203      78.027   8.728  81.677  1.00  0.00           H
    ATOM   4705 HG11 VAL B 203      77.254   9.286  79.442  1.00  0.00           H
    ATOM   4706 HG12 VAL B 203      78.631   8.196  79.314  1.00  0.00           H
    ATOM   4707 HG13 VAL B 203      77.004   7.600  79.023  1.00  0.00           H
    ATOM   4708 HG21 VAL B 203      75.611   9.018  81.250  1.00  0.00           H
    ATOM   4709 HG22 VAL B 203      75.420   7.286  81.132  1.00  0.00           H
    ATOM   4710 HG23 VAL B 203      75.961   8.011  82.656  1.00  0.00           H
    ATOM   4711  N   GLU B 204      76.485   4.820  81.146  1.00 34.40           N
    ATOM   4712  CA  GLU B 204      75.697   3.835  80.416  1.00 35.18           C
    ATOM   4713  C   GLU B 204      74.230   4.250  80.504  1.00 23.57           C
    ATOM   4714  O   GLU B 204      73.800   4.724  81.551  1.00 21.47           O
    ATOM   4715  CB  GLU B 204      75.880   2.432  81.032  1.00 49.22           C
    ATOM   4716  CG  GLU B 204      75.084   1.315  80.308  1.00 62.43           C
    ATOM   4717  CD  GLU B 204      74.371   0.352  81.254  1.00 70.00           C
    ATOM   4718  OE1 GLU B 204      75.087  -0.454  81.882  1.00 70.00           O
    ATOM   4719  OE2 GLU B 204      73.125   0.425  81.327  1.00 70.00           O1-
    ATOM   4720  H   GLU B 204      76.137   5.058  82.066  1.00  0.00           H
    ATOM   4721  HA  GLU B 204      75.998   3.810  79.369  1.00  0.00           H
    ATOM   4722  HB2 GLU B 204      76.939   2.173  81.011  1.00  0.00           H
    ATOM   4723  HB3 GLU B 204      75.623   2.472  82.087  1.00  0.00           H
    ATOM   4724  HG2 GLU B 204      74.349   1.696  79.609  1.00  0.00           H
    ATOM   4725  HG3 GLU B 204      75.771   0.736  79.691  1.00  0.00           H
    ATOM   4726  N   TYR B 205      73.479   4.013  79.425  1.00 27.39           N
    ATOM   4727  CA  TYR B 205      72.044   4.239  79.367  1.00 33.65           C
    ATOM   4728  C   TYR B 205      71.330   2.915  79.112  1.00 38.43           C
    ATOM   4729  O   TYR B 205      71.618   2.284  78.094  1.00 39.81           O
    ATOM   4730  CB  TYR B 205      71.753   5.229  78.233  1.00 28.01           C
    ATOM   4731  CG  TYR B 205      72.278   6.624  78.501  1.00 19.47           C
    ATOM   4732  CD1 TYR B 205      73.606   6.959  78.170  1.00 22.73           C
    ATOM   4733  CD2 TYR B 205      71.438   7.597  79.071  1.00  6.85           C
    ATOM   4734  CE1 TYR B 205      74.089   8.255  78.422  1.00 16.33           C
    ATOM   4735  CE2 TYR B 205      71.909   8.900  79.293  1.00 15.17           C
    ATOM   4736  CZ  TYR B 205      73.243   9.225  78.989  1.00 14.07           C
    ATOM   4737  OH  TYR B 205      73.719  10.478  79.237  1.00 20.54           O
    ATOM   4738  H   TYR B 205      73.907   3.623  78.593  1.00  0.00           H
    ATOM   4739  HA  TYR B 205      71.671   4.669  80.296  1.00  0.00           H
    ATOM   4740  HB2 TYR B 205      72.176   4.867  77.294  1.00  0.00           H
    ATOM   4741  HB3 TYR B 205      70.680   5.299  78.061  1.00  0.00           H
    ATOM   4742  HD1 TYR B 205      74.259   6.224  77.724  1.00  0.00           H
    ATOM   4743  HD2 TYR B 205      70.433   7.351  79.349  1.00  0.00           H
    ATOM   4744  HE1 TYR B 205      75.105   8.510  78.167  1.00  0.00           H
    ATOM   4745  HE2 TYR B 205      71.242   9.631  79.720  1.00  0.00           H
    ATOM   4746  HH  TYR B 205      74.604  10.627  78.893  1.00  0.00           H
    ATOM   4747  N   LEU B 206      70.381   2.546  79.993  1.00 34.94           N
```
