## Supplementary material for "DISRUPTOR: Computational identification of oncogenic mutants disrupting protein interactions": exemplary input files: smad4_1u7v.320-321.yaml

```
---
osprey_version: 3.1
design_name: smad4 and smad2
protein:
  residue_configurations:
  - mutability:
    - ALA
    - ARG
    - ASN
    - ASP
    - CYS
    - GLU
    - GLN
    - GLY
    - HIE
    - HID
    - HIP
    - ILE
    - LEU
    - LYS
    - MET
    - PHE
    - PRO
    - SER
    - THR
    - TRP
    - TYR
    - VAL
    flexibility:
      is_flexible: true
      include_structure_rotamer: true
      use_continuous: true
    identity:
      chain: B
      res_num: 320
      aa_type: PRO
  - mutability:
    - ALA
    - ARG
    - ASN
    - ASP
    - CYS
    - GLU
    - GLN
    - GLY
    - HIE
    - HID
    - HIP
    - ILE
    - LEU
    - LYS
    - MET
    - PHE
    - PRO
    - SER
    - THR
    - TRP
    - TYR
    - VAL
    flexibility:
      is_flexible: true
      include_structure_rotamer: true
      use_continuous: true
    identity:
      chain: B
      res_num: 321
      aa_type: GLU
  extra_templates: ""
  extra_templates_coordinates: ""
  coordinates: |
    ATOM   2891  N   ILE B 314     -11.978  75.870  55.586  1.00110.64           N1+
    ATOM   2892  CA  ILE B 314     -13.382  76.059  56.083  1.00110.65           C
    ATOM   2893  C   ILE B 314     -13.317  77.091  57.235  1.00110.47           C
    ATOM   2894  O   ILE B 314     -14.151  77.996  57.244  1.00109.66           O
    ATOM   2895  CB  ILE B 314     -14.112  74.756  56.540  1.00111.11           C
    ATOM   2896  CG1 ILE B 314     -14.150  73.667  55.441  1.00110.16           C
    ATOM   2897  CG2 ILE B 314     -15.533  74.994  57.093  1.00111.22           C
    ATOM   2898  CD1 ILE B 314     -15.030  74.008  54.232  1.00106.18           C
    ATOM   2899  H1  ILE B 314     -12.001  75.294  54.757  1.00  0.00           H
    ATOM   2900  H2  ILE B 314     -11.564  76.766  55.370  1.00  0.00           H
    ATOM   2901  H3  ILE B 314     -11.431  75.382  56.286  1.00  0.00           H
    ATOM   2902  HA  ILE B 314     -13.893  76.517  55.236  1.00  0.00           H
    ATOM   2903  HB  ILE B 314     -13.532  74.322  57.358  1.00  0.00           H
    ATOM   2904 HG12 ILE B 314     -13.145  73.445  55.087  1.00  0.00           H
    ATOM   2905 HG13 ILE B 314     -14.508  72.734  55.879  1.00  0.00           H
    ATOM   2906 HG21 ILE B 314     -16.026  74.048  57.312  1.00  0.00           H
    ATOM   2907 HG22 ILE B 314     -15.527  75.572  58.017  1.00  0.00           H
    ATOM   2908 HG23 ILE B 314     -16.154  75.530  56.375  1.00  0.00           H
    ATOM   2909 HD11 ILE B 314     -14.799  73.348  53.397  1.00  0.00           H
    ATOM   2910 HD12 ILE B 314     -16.086  73.887  54.471  1.00  0.00           H
    ATOM   2911 HD13 ILE B 314     -14.882  75.029  53.886  1.00  0.00           H
    ATOM   2912  N   SER B 315     -12.362  76.947  58.174  1.00111.06           N
    ATOM   2913  CA  SER B 315     -12.156  77.874  59.287  1.00112.45           C
    ATOM   2914  C   SER B 315     -10.695  78.351  59.334  1.00112.77           C
    ATOM   2915  O   SER B 315      -9.817  77.729  58.734  1.00112.89           O
    ATOM   2916  CB  SER B 315     -12.619  77.194  60.588  1.00113.00           C
    ATOM   2917  OG  SER B 315     -12.491  78.050  61.702  1.00116.58           O
    ATOM   2918  H   SER B 315     -11.708  76.178  58.126  1.00  0.00           H
    ATOM   2919  HA  SER B 315     -12.764  78.770  59.144  1.00  0.00           H
    ATOM   2920  HB2 SER B 315     -13.673  76.935  60.501  1.00  0.00           H
    ATOM   2921  HB3 SER B 315     -12.065  76.273  60.777  1.00  0.00           H
    ATOM   2922  HG  SER B 315     -12.911  77.635  62.463  1.00  0.00           H
    ATOM   2923  N   ASN B 316     -10.488  79.471  60.044  1.00112.44           N
    ATOM   2924  CA  ASN B 316      -9.211  80.177  60.168  1.00113.86           C
    ATOM   2925  C   ASN B 316      -8.525  79.933  61.522  1.00114.37           C
    ATOM   2926  O   ASN B 316      -7.333  80.225  61.620  1.00114.99           O
    ATOM   2927  CB  ASN B 316      -9.450  81.692  59.970  1.00115.51           C
    ATOM   2928  CG  ASN B 316      -9.995  82.040  58.581  1.00117.62           C
    ATOM   2929  ND2 ASN B 316     -11.131  82.738  58.527  1.00116.29           N
    ATOM   2930  OD1 ASN B 316      -9.392  81.690  57.569  1.00120.21           O
    ATOM   2931  H   ASN B 316     -11.274  79.892  60.520  1.00  0.00           H
    ATOM   2932  HA  ASN B 316      -8.522  79.831  59.394  1.00  0.00           H
    ATOM   2933  HB2 ASN B 316     -10.121  82.074  60.743  1.00  0.00           H
    ATOM   2934  HB3 ASN B 316      -8.510  82.234  60.086  1.00  0.00           H
    ATOM   2935 HD21 ASN B 316     -11.607  83.017  59.372  1.00  0.00           H
    ATOM   2936 HD22 ASN B 316     -11.521  82.994  57.631  1.00  0.00           H
    ATOM   2937  N   HIS B 317      -9.247  79.405  62.533  1.00114.49           N
    ATOM   2938  CA  HIS B 317      -8.690  79.076  63.854  1.00114.90           C
    ATOM   2939  C   HIS B 317      -7.625  77.954  63.765  1.00113.86           C
    ATOM   2940  O   HIS B 317      -7.693  77.155  62.825  1.00114.40           O
    ATOM   2941  CB  HIS B 317      -9.840  78.768  64.854  1.00115.23           C
    ATOM   2942  CG  HIS B 317     -10.743  77.582  64.595  1.00119.35           C
    ATOM   2943  CD2 HIS B 317     -12.114  77.462  64.653  1.00121.01           C
    ATOM   2944  ND1 HIS B 317     -10.283  76.318  64.270  1.00122.01           N
    ATOM   2945  CE1 HIS B 317     -11.349  75.534  64.103  1.00123.64           C
    ATOM   2946  NE2 HIS B 317     -12.497  76.161  64.317  1.00123.57           N
    ATOM   2947  H   HIS B 317     -10.217  79.165  62.383  1.00  0.00           H
    ATOM   2948  HA  HIS B 317      -8.192  79.984  64.199  1.00  0.00           H
    ATOM   2949  HB2 HIS B 317      -9.451  78.625  65.860  1.00  0.00           H
    ATOM   2950  HB3 HIS B 317     -10.479  79.649  64.924  1.00  0.00           H
    ATOM   2951  HD1 HIS B 317      -9.317  76.041  64.172  1.00  0.00           H
    ATOM   2952  HD2 HIS B 317     -12.854  78.212  64.892  1.00  0.00           H
    ATOM   2953  HE1 HIS B 317     -11.285  74.491  63.826  1.00  0.00           H
    ATOM   2954  N   PRO B 318      -6.669  77.899  64.727  1.00111.89           N
    ATOM   2955  CA  PRO B 318      -5.567  76.913  64.719  1.00109.25           C
    ATOM   2956  C   PRO B 318      -6.006  75.445  64.581  1.00106.41           C
    ATOM   2957  O   PRO B 318      -7.025  75.046  65.149  1.00105.93           O
    ATOM   2958  CB  PRO B 318      -4.821  77.162  66.045  1.00109.40           C
    ATOM   2959  CG  PRO B 318      -5.786  77.956  66.907  1.00109.49           C
    ATOM   2960  CD  PRO B 318      -6.527  78.794  65.878  1.00110.34           C
    ATOM   2961  HA  PRO B 318      -4.904  77.155  63.886  1.00  0.00           H
    ATOM   2962  HB2 PRO B 318      -4.482  76.252  66.544  1.00  0.00           H
    ATOM   2963  HB3 PRO B 318      -3.933  77.764  65.846  1.00  0.00           H
    ATOM   2964  HG2 PRO B 318      -6.485  77.276  67.398  1.00  0.00           H
    ATOM   2965  HG3 PRO B 318      -5.292  78.549  67.677  1.00  0.00           H
    ATOM   2966  HD2 PRO B 318      -7.454  79.174  66.297  1.00  0.00           H
    ATOM   2967  HD3 PRO B 318      -5.924  79.651  65.576  1.00  0.00           H
    ATOM   2968  N   ALA B 319      -5.216  74.690  63.798  1.00102.12           N
    ATOM   2969  CA  ALA B 319      -5.346  73.249  63.583  1.00 98.40           C
    ATOM   2970  C   ALA B 319      -5.269  72.468  64.913  1.00 93.89           C
    ATOM   2971  O   ALA B 319      -4.506  72.873  65.793  1.00 90.75           O
    ATOM   2972  CB  ALA B 319      -4.215  72.811  62.639  1.00 96.62           C
    ATOM   2973  H   ALA B 319      -4.409  75.118  63.367  1.00  0.00           H
    ATOM   2974  HA  ALA B 319      -6.301  73.087  63.088  1.00  0.00           H
    ATOM   2975  HB1 ALA B 319      -4.268  71.743  62.425  1.00  0.00           H
    ATOM   2976  HB2 ALA B 319      -4.275  73.337  61.686  1.00  0.00           H
    ATOM   2977  HB3 ALA B 319      -3.233  73.013  63.070  1.00  0.00           H
    ATOM   2978  N   PRO B 320      -6.067  71.385  65.054  1.00 88.74           N
    ATOM   2979  CA  PRO B 320      -6.107  70.607  66.301  1.00 84.09           C
    ATOM   2980  C   PRO B 320      -4.797  69.832  66.527  1.00 82.32           C
    ATOM   2981  O   PRO B 320      -4.117  69.495  65.557  1.00 82.83           O
    ATOM   2982  CB  PRO B 320      -7.311  69.680  66.087  1.00 85.00           C
    ATOM   2983  CG  PRO B 320      -7.343  69.427  64.587  1.00 85.72           C
    ATOM   2984  CD  PRO B 320      -6.885  70.764  64.009  1.00 87.23           C
    ATOM   2985  HA  PRO B 320      -6.290  71.260  67.157  1.00  0.00           H
    ATOM   2986  HB2 PRO B 320      -7.266  68.773  66.684  1.00  0.00           H
    ATOM   2987  HB3 PRO B 320      -8.223  70.191  66.387  1.00  0.00           H
    ATOM   2988  HG2 PRO B 320      -6.619  68.650  64.335  1.00  0.00           H
    ATOM   2989  HG3 PRO B 320      -8.319  69.107  64.222  1.00  0.00           H
    ATOM   2990  HD2 PRO B 320      -6.345  70.623  63.073  1.00  0.00           H
    ATOM   2991  HD3 PRO B 320      -7.737  71.410  63.819  1.00  0.00           H
    ATOM   2992  N   GLU B 321      -4.461  69.579  67.802  1.00 78.02           N
    ATOM   2993  CA  GLU B 321      -3.277  68.815  68.215  1.00 75.07           C
    ATOM   2994  C   GLU B 321      -3.252  67.388  67.632  1.00 73.26           C
    ATOM   2995  O   GLU B 321      -2.218  66.946  67.130  1.00 69.37           O
    ATOM   2996  CB  GLU B 321      -3.213  68.818  69.756  1.00 76.40           C
    ATOM   2997  CG  GLU B 321      -2.020  68.049  70.366  1.00 78.32           C
    ATOM   2998  CD  GLU B 321      -1.977  68.092  71.896  1.00 79.28           C
    ATOM   2999  OE1 GLU B 321      -0.986  67.559  72.439  1.00 77.35           O
    ATOM   3000  OE2 GLU B 321      -2.931  68.627  72.504  1.00 71.26           O1-
    ATOM   3001  H   GLU B 321      -5.056  69.912  68.547  1.00  0.00           H
    ATOM   3002  HA  GLU B 321      -2.399  69.341  67.836  1.00  0.00           H
    ATOM   3003  HB2 GLU B 321      -3.176  69.853  70.099  1.00  0.00           H
    ATOM   3004  HB3 GLU B 321      -4.144  68.407  70.152  1.00  0.00           H
    ATOM   3005  HG2 GLU B 321      -2.044  66.999  70.074  1.00  0.00           H
    ATOM   3006  HG3 GLU B 321      -1.084  68.458  69.983  1.00  0.00           H
    ATOM   3007  N   TYR B 322      -4.422  66.738  67.673  1.00 69.32           N
    ATOM   3008  CA  TYR B 322      -4.704  65.440  67.075  1.00 65.69           C
    ATOM   3009  C   TYR B 322      -5.796  65.651  66.025  1.00 63.62           C
    ATOM   3010  O   TYR B 322      -6.703  66.439  66.279  1.00 68.63           O
    ATOM   3011  CB  TYR B 322      -5.190  64.478  68.181  1.00 59.70           C
    ATOM   3012  CG  TYR B 322      -4.291  64.404  69.402  1.00 60.48           C
    ATOM   3013  CD1 TYR B 322      -3.113  63.634  69.365  1.00 60.15           C
    ATOM   3014  CD2 TYR B 322      -4.621  65.110  70.578  1.00 62.62           C
    ATOM   3015  CE1 TYR B 322      -2.268  63.562  70.485  1.00 61.27           C
    ATOM   3016  CE2 TYR B 322      -3.772  65.047  71.701  1.00 63.22           C
    ATOM   3017  CZ  TYR B 322      -2.595  64.274  71.654  1.00 65.53           C
    ATOM   3018  OH  TYR B 322      -1.764  64.220  72.733  1.00 68.09           O
    ATOM   3019  H   TYR B 322      -5.218  67.205  68.082  1.00  0.00           H
    ATOM   3020  HA  TYR B 322      -3.820  65.038  66.588  1.00  0.00           H
    ATOM   3021  HB2 TYR B 322      -6.175  64.783  68.525  1.00  0.00           H
    ATOM   3022  HB3 TYR B 322      -5.325  63.473  67.781  1.00  0.00           H
    ATOM   3023  HD1 TYR B 322      -2.857  63.081  68.484  1.00  0.00           H
    ATOM   3024  HD2 TYR B 322      -5.516  65.712  70.621  1.00  0.00           H
    ATOM   3025  HE1 TYR B 322      -1.369  62.965  70.445  1.00  0.00           H
    ATOM   3026  HE2 TYR B 322      -4.023  65.602  72.593  1.00  0.00           H
    ATOM   3027  HH  TYR B 322      -1.965  64.890  73.392  1.00  0.00           H
    ATOM   3028  N   TRP B 323      -5.726  64.953  64.881  1.00 59.86           N
    ATOM   3029  CA  TRP B 323      -6.787  65.023  63.869  1.00 59.39           C
    ATOM   3030  C   TRP B 323      -7.929  64.028  64.144  1.00 57.23           C
    ATOM   3031  O   TRP B 323      -9.014  64.225  63.598  1.00 61.61           O
    ATOM   3032  CB  TRP B 323      -6.204  64.875  62.453  1.00 56.28           C
    ATOM   3033  CG  TRP B 323      -5.704  63.529  62.027  1.00 54.21           C
    ATOM   3034  CD1 TRP B 323      -4.430  63.098  62.131  1.00 55.35           C
    ATOM   3035  CD2 TRP B 323      -6.449  62.420  61.442  1.00 47.15           C
    ATOM   3036  CE2 TRP B 323      -5.527  61.373  61.142  1.00 47.98           C
    ATOM   3037  CE3 TRP B 323      -7.804  62.201  61.095  1.00 47.85           C
    ATOM   3038  NE1 TRP B 323      -4.314  61.845  61.582  1.00 50.14           N
    ATOM   3039  CZ2 TRP B 323      -5.918  60.184  60.513  1.00 44.47           C
    ATOM   3040  CZ3 TRP B 323      -8.210  61.005  60.474  1.00 45.35           C
    ATOM   3041  CH2 TRP B 323      -7.268  60.009  60.167  1.00 40.95           C
    ATOM   3042  H   TRP B 323      -4.965  64.308  64.714  1.00  0.00           H
    ATOM   3043  HA  TRP B 323      -7.226  66.023  63.896  1.00  0.00           H
    ATOM   3044  HB2 TRP B 323      -6.977  65.150  61.733  1.00  0.00           H
    ATOM   3045  HB3 TRP B 323      -5.408  65.604  62.304  1.00  0.00           H
    ATOM   3046  HD1 TRP B 323      -3.627  63.672  62.546  1.00  0.00           H
    ATOM   3047  HE1 TRP B 323      -3.425  61.362  61.517  1.00  0.00           H
    ATOM   3048  HE3 TRP B 323      -8.537  62.968  61.295  1.00  0.00           H
    ATOM   3049  HZ2 TRP B 323      -5.182  59.430  60.289  1.00  0.00           H
    ATOM   3050  HZ3 TRP B 323      -9.247  60.850  60.224  1.00  0.00           H
    ATOM   3051  HH2 TRP B 323      -7.583  59.116  59.659  1.00  0.00           H
    ATOM   3052  N   CYS B 324      -7.688  63.008  64.991  1.00 57.38           N
    ATOM   3053  CA  CYS B 324      -8.708  62.048  65.403  1.00 54.36           C
    ATOM   3054  C   CYS B 324      -8.370  61.350  66.731  1.00 55.11           C
    ATOM   3055  O   CYS B 324      -7.222  61.369  67.176  1.00 54.38           O
    ATOM   3056  CB  CYS B 324      -9.087  61.046  64.282  1.00 48.66           C
    ATOM   3057  SG  CYS B 324      -7.831  59.761  64.000  1.00 48.38           S
    ATOM   3058  H   CYS B 324      -6.774  62.896  65.407  1.00  0.00           H
    ATOM   3059  HA  CYS B 324      -9.577  62.665  65.619  1.00  0.00           H
    ATOM   3060  HB2 CYS B 324     -10.011  60.535  64.549  1.00  0.00           H
    ATOM   3061  HB3 CYS B 324      -9.297  61.560  63.346  1.00  0.00           H
    ATOM   3062  HG  CYS B 324      -7.981  59.149  65.179  1.00  0.00           H
    ATOM   3063  N   SER B 325      -9.408  60.723  67.307  1.00 53.82           N
    ATOM   3064  CA  SER B 325      -9.372  59.893  68.508  1.00 51.14           C
    ATOM   3065  C   SER B 325     -10.096  58.575  68.211  1.00 52.73           C
    ATOM   3066  O   SER B 325     -11.249  58.631  67.793  1.00 53.60           O
    ATOM   3067  CB  SER B 325     -10.085  60.631  69.658  1.00 49.28           C
    ATOM   3068  OG  SER B 325      -9.176  61.451  70.354  1.00 53.09           O
    ATOM   3069  H   SER B 325     -10.316  60.790  66.865  1.00  0.00           H
    ATOM   3070  HA  SER B 325      -8.345  59.664  68.795  1.00  0.00           H
    ATOM   3071  HB2 SER B 325     -10.916  61.236  69.291  1.00  0.00           H
    ATOM   3072  HB3 SER B 325     -10.507  59.932  70.378  1.00  0.00           H
    ATOM   3073  HG  SER B 325      -9.107  62.288  69.882  1.00  0.00           H
    ATOM   3074  N   ILE B 326      -9.427  57.431  68.435  1.00 50.99           N
    ATOM   3075  CA  ILE B 326      -9.929  56.100  68.087  1.00 50.39           C
    ATOM   3076  C   ILE B 326     -10.095  55.271  69.377  1.00 46.37           C
    ATOM   3077  O   ILE B 326      -9.100  54.897  69.996  1.00 49.98           O
    ATOM   3078  CB  ILE B 326      -8.953  55.325  67.146  1.00 49.88           C
    ATOM   3079  CG1 ILE B 326      -8.454  56.180  65.958  1.00 51.18           C
    ATOM   3080  CG2 ILE B 326      -9.580  54.009  66.635  1.00 42.08           C
    ATOM   3081  CD1 ILE B 326      -7.312  55.523  65.175  1.00 56.14           C
    ATOM   3082  H   ILE B 326      -8.476  57.469  68.782  1.00  0.00           H
    ATOM   3083  HA  ILE B 326     -10.893  56.170  67.584  1.00  0.00           H
    ATOM   3084  HB  ILE B 326      -8.065  55.052  67.712  1.00  0.00           H
    ATOM   3085 HG12 ILE B 326      -9.282  56.399  65.286  1.00  0.00           H
    ATOM   3086 HG13 ILE B 326      -8.086  57.147  66.300  1.00  0.00           H
    ATOM   3087 HG21 ILE B 326      -8.873  53.426  66.046  1.00  0.00           H
    ATOM   3088 HG22 ILE B 326      -9.919  53.362  67.443  1.00  0.00           H
    ATOM   3089 HG23 ILE B 326     -10.444  54.217  66.003  1.00  0.00           H
    ATOM   3090 HD11 ILE B 326      -6.757  56.269  64.611  1.00  0.00           H
    ATOM   3091 HD12 ILE B 326      -6.607  55.013  65.832  1.00  0.00           H
    ATOM   3092 HD13 ILE B 326      -7.699  54.797  64.464  1.00  0.00           H
    ATOM   3093  N   ALA B 327     -11.350  54.981  69.744  1.00 43.23           N
    ATOM   3094  CA  ALA B 327     -11.701  54.124  70.875  1.00 42.19           C
    ATOM   3095  C   ALA B 327     -12.046  52.724  70.357  1.00 43.25           C
    ATOM   3096  O   ALA B 327     -12.978  52.605  69.566  1.00 41.22           O
    ATOM   3097  CB  ALA B 327     -12.899  54.751  71.605  1.00 42.76           C
    ATOM   3098  H   ALA B 327     -12.124  55.313  69.182  1.00  0.00           H
    ATOM   3099  HA  ALA B 327     -10.876  54.049  71.585  1.00  0.00           H
    ATOM   3100  HB1 ALA B 327     -13.289  54.085  72.374  1.00  0.00           H
    ATOM   3101  HB2 ALA B 327     -12.612  55.682  72.094  1.00  0.00           H
    ATOM   3102  HB3 ALA B 327     -13.717  54.978  70.919  1.00  0.00           H
    ATOM   3103  N   TYR B 328     -11.304  51.699  70.813  1.00 41.47           N
    ATOM   3104  CA  TYR B 328     -11.567  50.292  70.496  1.00 42.27           C
    ATOM   3105  C   TYR B 328     -12.645  49.700  71.416  1.00 45.11           C
    ATOM   3106  O   TYR B 328     -12.692  50.039  72.594  1.00 48.79           O
    ATOM   3107  CB  TYR B 328     -10.246  49.493  70.543  1.00 42.27           C
    ATOM   3108  CG  TYR B 328     -10.395  47.991  70.354  1.00 47.82           C
    ATOM   3109  CD1 TYR B 328     -10.846  47.479  69.122  1.00 48.10           C
    ATOM   3110  CD2 TYR B 328     -10.111  47.102  71.408  1.00 48.60           C
    ATOM   3111  CE1 TYR B 328     -11.019  46.093  68.949  1.00 52.96           C
    ATOM   3112  CE2 TYR B 328     -10.274  45.718  71.236  1.00 55.61           C
    ATOM   3113  CZ  TYR B 328     -10.736  45.210  70.008  1.00 56.04           C
    ATOM   3114  OH  TYR B 328     -10.916  43.867  69.855  1.00 69.33           O
    ATOM   3115  H   TYR B 328     -10.560  51.875  71.472  1.00  0.00           H
    ATOM   3116  HA  TYR B 328     -11.953  50.234  69.480  1.00  0.00           H
    ATOM   3117  HB2 TYR B 328      -9.571  49.862  69.771  1.00  0.00           H
    ATOM   3118  HB3 TYR B 328      -9.739  49.676  71.491  1.00  0.00           H
    ATOM   3119  HD1 TYR B 328     -11.066  48.149  68.306  1.00  0.00           H
    ATOM   3120  HD2 TYR B 328      -9.757  47.472  72.354  1.00  0.00           H
    ATOM   3121  HE1 TYR B 328     -11.374  45.725  68.000  1.00  0.00           H
    ATOM   3122  HE2 TYR B 328     -10.042  45.054  72.053  1.00  0.00           H
    ATOM   3123  HH  TYR B 328     -10.960  43.421  70.711  1.00  0.00           H
    ATOM   3124  N   PHE B 329     -13.476  48.816  70.848  1.00 45.61           N
    ATOM   3125  CA  PHE B 329     -14.592  48.146  71.503  1.00 46.83           C
    ATOM   3126  C   PHE B 329     -14.589  46.644  71.199  1.00 47.92           C
    ATOM   3127  O   PHE B 329     -14.131  46.228  70.135  1.00 51.17           O
    ATOM   3128  CB  PHE B 329     -15.918  48.748  70.997  1.00 43.82           C
    ATOM   3129  CG  PHE B 329     -16.250  50.157  71.448  1.00 43.13           C
    ATOM   3130  CD1 PHE B 329     -15.689  51.266  70.784  1.00 42.48           C
    ATOM   3131  CD2 PHE B 329     -17.027  50.368  72.605  1.00 43.72           C
    ATOM   3132  CE1 PHE B 329     -15.930  52.549  71.256  1.00 45.78           C
    ATOM   3133  CE2 PHE B 329     -17.260  51.660  73.057  1.00 44.10           C
    ATOM   3134  CZ  PHE B 329     -16.713  52.746  72.385  1.00 40.65           C
    ATOM   3135  H   PHE B 329     -13.360  48.591  69.867  1.00  0.00           H
    ATOM   3136  HA  PHE B 329     -14.517  48.265  72.582  1.00  0.00           H
    ATOM   3137  HB2 PHE B 329     -15.905  48.742  69.908  1.00  0.00           H
    ATOM   3138  HB3 PHE B 329     -16.752  48.110  71.296  1.00  0.00           H
    ATOM   3139  HD1 PHE B 329     -15.076  51.118  69.908  1.00  0.00           H
    ATOM   3140  HD2 PHE B 329     -17.452  49.530  73.136  1.00  0.00           H
    ATOM   3141  HE1 PHE B 329     -15.502  53.396  70.744  1.00  0.00           H
    ATOM   3142  HE2 PHE B 329     -17.864  51.819  73.938  1.00  0.00           H
    ATOM   3143  HZ  PHE B 329     -16.893  53.748  72.745  1.00  0.00           H
    ATOM   3144  N   GLU B 330     -15.186  45.882  72.125  1.00 45.87           N
    ATOM   3145  CA  GLU B 330     -15.583  44.489  71.957  1.00 43.79           C
    ATOM   3146  C   GLU B 330     -17.000  44.388  72.513  1.00 48.42           C
    ATOM   3147  O   GLU B 330     -17.173  44.457  73.731  1.00 49.07           O
    ATOM   3148  CB  GLU B 330     -14.622  43.543  72.705  1.00 45.18           C
    ATOM   3149  CG  GLU B 330     -13.200  43.578  72.132  1.00 44.93           C
    ATOM   3150  CD  GLU B 330     -12.207  42.644  72.816  1.00 44.15           C
    ATOM   3151  OE1 GLU B 330     -12.602  41.817  73.666  1.00 42.31           O
    ATOM   3152  OE2 GLU B 330     -11.020  42.783  72.460  1.00 43.57           O1-
    ATOM   3153  H   GLU B 330     -15.521  46.321  72.974  1.00  0.00           H
    ATOM   3154  HA  GLU B 330     -15.603  44.227  70.898  1.00  0.00           H
    ATOM   3155  HB2 GLU B 330     -14.595  43.795  73.766  1.00  0.00           H
    ATOM   3156  HB3 GLU B 330     -15.006  42.525  72.645  1.00  0.00           H
    ATOM   3157  HG2 GLU B 330     -13.227  43.330  71.073  1.00  0.00           H
    ATOM   3158  HG3 GLU B 330     -12.796  44.587  72.207  1.00  0.00           H
    ATOM   3159  N   MET B 331     -17.986  44.276  71.607  1.00 53.94           N
    ATOM   3160  CA  MET B 331     -19.418  44.360  71.908  1.00 53.52           C
    ATOM   3161  C   MET B 331     -19.778  45.787  72.385  1.00 52.85           C
    ATOM   3162  O   MET B 331     -19.393  46.753  71.728  1.00 43.89           O
    ATOM   3163  CB  MET B 331     -19.879  43.225  72.871  1.00 54.02           C
    ATOM   3164  CG  MET B 331     -19.462  41.800  72.460  1.00 60.96           C
    ATOM   3165  SD  MET B 331     -20.308  41.121  71.009  1.00 69.04           S
    ATOM   3166  CE  MET B 331     -21.899  40.669  71.753  1.00 62.06           C
    ATOM   3167  H   MET B 331     -17.753  44.212  70.625  1.00  0.00           H
    ATOM   3168  HA  MET B 331     -19.926  44.213  70.954  1.00  0.00           H
    ATOM   3169  HB2 MET B 331     -19.512  43.405  73.882  1.00  0.00           H
    ATOM   3170  HB3 MET B 331     -20.966  43.255  72.957  1.00  0.00           H
    ATOM   3171  HG2 MET B 331     -18.390  41.754  72.271  1.00  0.00           H
    ATOM   3172  HG3 MET B 331     -19.637  41.121  73.294  1.00  0.00           H
    ATOM   3173  HE1 MET B 331     -22.543  40.213  71.001  1.00  0.00           H
    ATOM   3174  HE2 MET B 331     -21.755  39.956  72.564  1.00  0.00           H
    ATOM   3175  HE3 MET B 331     -22.406  41.551  72.145  1.00  0.00           H
    ATOM   3176  N   ASP B 332     -20.509  45.900  73.504  1.00 56.51           N
    ATOM   3177  CA  ASP B 332     -20.947  47.153  74.127  1.00 57.45           C
    ATOM   3178  C   ASP B 332     -19.881  47.779  75.061  1.00 61.84           C
    ATOM   3179  O   ASP B 332     -20.169  48.807  75.673  1.00 68.00           O
    ATOM   3180  CB  ASP B 332     -22.279  46.976  74.903  1.00 57.09           C
    ATOM   3181  CG  ASP B 332     -22.287  45.896  75.999  1.00 60.79           C
    ATOM   3182  OD1 ASP B 332     -21.800  44.774  75.734  1.00 61.19           O
    ATOM   3183  OD2 ASP B 332     -22.861  46.194  77.067  1.00 57.77           O1-
    ATOM   3184  H   ASP B 332     -20.796  45.066  74.001  1.00  0.00           H
    ATOM   3185  HA  ASP B 332     -21.131  47.881  73.335  1.00  0.00           H
    ATOM   3186  HB2 ASP B 332     -22.550  47.926  75.361  1.00  0.00           H
    ATOM   3187  HB3 ASP B 332     -23.086  46.751  74.206  1.00  0.00           H
    ATOM   3188  N   VAL B 333     -18.698  47.155  75.191  1.00 56.86           N
    ATOM   3189  CA  VAL B 333     -17.668  47.542  76.156  1.00 49.71           C
    ATOM   3190  C   VAL B 333     -16.447  48.102  75.411  1.00 46.40           C
    ATOM   3191  O   VAL B 333     -15.942  47.432  74.510  1.00 41.18           O
    ATOM   3192  CB  VAL B 333     -17.215  46.306  76.989  1.00 48.63           C
    ATOM   3193  CG1 VAL B 333     -16.047  46.586  77.959  1.00 52.38           C
    ATOM   3194  CG2 VAL B 333     -18.395  45.695  77.767  1.00 51.39           C
    ATOM   3195  H   VAL B 333     -18.513  46.319  74.655  1.00  0.00           H
    ATOM   3196  HA  VAL B 333     -18.040  48.303  76.846  1.00  0.00           H
    ATOM   3197  HB  VAL B 333     -16.869  45.534  76.299  1.00  0.00           H
    ATOM   3198 HG11 VAL B 333     -15.845  45.722  78.592  1.00  0.00           H
    ATOM   3199 HG12 VAL B 333     -15.118  46.804  77.431  1.00  0.00           H
    ATOM   3200 HG13 VAL B 333     -16.268  47.430  78.613  1.00  0.00           H
    ATOM   3201 HG21 VAL B 333     -18.078  44.838  78.362  1.00  0.00           H
    ATOM   3202 HG22 VAL B 333     -18.840  46.424  78.446  1.00  0.00           H
    ATOM   3203 HG23 VAL B 333     -19.181  45.343  77.099  1.00  0.00           H
    ATOM   3204  N   GLN B 334     -15.975  49.295  75.824  1.00 48.32           N
    ATOM   3205  CA  GLN B 334     -14.717  49.871  75.344  1.00 49.60           C
    ATOM   3206  C   GLN B 334     -13.543  49.111  75.982  1.00 50.47           C
    ATOM   3207  O   GLN B 334     -13.451  49.051  77.208  1.00 52.95           O
    ATOM   3208  CB  GLN B 334     -14.638  51.385  75.646  1.00 46.74           C
    ATOM   3209  CG  GLN B 334     -13.442  52.067  74.938  1.00 42.15           C
    ATOM   3210  CD  GLN B 334     -13.248  53.540  75.292  1.00 50.22           C
    ATOM   3211  NE2 GLN B 334     -12.002  54.015  75.210  1.00 47.82           N
    ATOM   3212  OE1 GLN B 334     -14.204  54.253  75.588  1.00 56.41           O
    ATOM   3213  H   GLN B 334     -16.440  49.791  76.571  1.00  0.00           H
    ATOM   3214  HA  GLN B 334     -14.688  49.754  74.262  1.00  0.00           H
    ATOM   3215  HB2 GLN B 334     -15.561  51.868  75.325  1.00  0.00           H
    ATOM   3216  HB3 GLN B 334     -14.574  51.547  76.723  1.00  0.00           H
    ATOM   3217  HG2 GLN B 334     -12.513  51.537  75.152  1.00  0.00           H
    ATOM   3218  HG3 GLN B 334     -13.592  52.018  73.860  1.00  0.00           H
    ATOM   3219 HE21 GLN B 334     -11.239  53.402  74.964  1.00  0.00           H
    ATOM   3220 HE22 GLN B 334     -11.818  54.993  75.383  1.00  0.00           H
    ATOM   3221  N   VAL B 335     -12.680  48.552  75.125  1.00 49.95           N
    ATOM   3222  CA  VAL B 335     -11.500  47.791  75.512  1.00 47.21           C
    ATOM   3223  C   VAL B 335     -10.250  48.611  75.161  1.00 45.23           C
    ATOM   3224  O   VAL B 335     -10.165  49.169  74.067  1.00 47.81           O
    ATOM   3225  CB  VAL B 335     -11.483  46.422  74.783  1.00 45.90           C
    ATOM   3226  CG1 VAL B 335     -10.155  45.650  74.897  1.00 38.38           C
    ATOM   3227  CG2 VAL B 335     -12.647  45.553  75.283  1.00 33.57           C
    ATOM   3228  H   VAL B 335     -12.802  48.701  74.132  1.00  0.00           H
    ATOM   3229  HA  VAL B 335     -11.499  47.605  76.589  1.00  0.00           H
    ATOM   3230  HB  VAL B 335     -11.663  46.592  73.720  1.00  0.00           H
    ATOM   3231 HG11 VAL B 335     -10.251  44.637  74.507  1.00  0.00           H
    ATOM   3232 HG12 VAL B 335      -9.361  46.130  74.329  1.00  0.00           H
    ATOM   3233 HG13 VAL B 335      -9.834  45.581  75.934  1.00  0.00           H
    ATOM   3234 HG21 VAL B 335     -12.605  44.552  74.861  1.00  0.00           H
    ATOM   3235 HG22 VAL B 335     -12.630  45.454  76.368  1.00  0.00           H
    ATOM   3236 HG23 VAL B 335     -13.613  45.977  75.006  1.00  0.00           H
    ATOM   3237  N   GLY B 336      -9.318  48.704  76.124  1.00 44.41           N
    ATOM   3238  CA  GLY B 336      -8.167  49.604  76.058  1.00 49.25           C
    ATOM   3239  C   GLY B 336      -8.601  51.074  76.206  1.00 49.80           C
    ATOM   3240  O   GLY B 336      -9.776  51.383  76.418  1.00 49.52           O
    ATOM   3241  H   GLY B 336      -9.464  48.218  76.997  1.00  0.00           H
    ATOM   3242  HA2 GLY B 336      -7.471  49.350  76.858  1.00  0.00           H
    ATOM   3243  HA3 GLY B 336      -7.639  49.469  75.113  1.00  0.00           H
    ATOM   3244  N   GLU B 337      -7.624  51.982  76.093  1.00 52.14           N
    ATOM   3245  CA  GLU B 337      -7.835  53.429  76.148  1.00 54.96           C
    ATOM   3246  C   GLU B 337      -7.780  54.035  74.740  1.00 54.11           C
    ATOM   3247  O   GLU B 337      -7.182  53.449  73.834  1.00 56.36           O
    ATOM   3248  CB  GLU B 337      -6.790  54.055  77.091  1.00 57.07           C
    ATOM   3249  CG  GLU B 337      -7.024  53.635  78.558  1.00 71.53           C
    ATOM   3250  CD  GLU B 337      -6.028  54.222  79.556  1.00 75.03           C
    ATOM   3251  OE1 GLU B 337      -6.335  54.124  80.764  1.00 74.89           O
    ATOM   3252  OE2 GLU B 337      -4.962  54.711  79.122  1.00 77.63           O1-
    ATOM   3253  H   GLU B 337      -6.677  51.682  75.912  1.00  0.00           H
    ATOM   3254  HA  GLU B 337      -8.822  53.658  76.555  1.00  0.00           H
    ATOM   3255  HB2 GLU B 337      -5.787  53.760  76.774  1.00  0.00           H
    ATOM   3256  HB3 GLU B 337      -6.824  55.143  77.017  1.00  0.00           H
    ATOM   3257  HG2 GLU B 337      -8.028  53.930  78.866  1.00  0.00           H
    ATOM   3258  HG3 GLU B 337      -6.976  52.551  78.654  1.00  0.00           H
    ATOM   3259  N   THR B 338      -8.430  55.201  74.595  1.00 54.71           N
    ATOM   3260  CA  THR B 338      -8.560  55.953  73.346  1.00 53.57           C
    ATOM   3261  C   THR B 338      -7.196  56.379  72.770  1.00 51.95           C
    ATOM   3262  O   THR B 338      -6.376  56.956  73.487  1.00 52.52           O
    ATOM   3263  CB  THR B 338      -9.395  57.238  73.561  1.00 55.45           C
    ATOM   3264  CG2 THR B 338      -9.637  58.091  72.302  1.00 56.34           C
    ATOM   3265  OG1 THR B 338     -10.658  56.904  74.104  1.00 60.88           O
    ATOM   3266  H   THR B 338      -8.895  55.612  75.391  1.00  0.00           H
    ATOM   3267  HA  THR B 338      -9.074  55.311  72.630  1.00  0.00           H
    ATOM   3268  HB  THR B 338      -8.886  57.858  74.298  1.00  0.00           H
    ATOM   3269 HG21 THR B 338     -10.292  58.932  72.530  1.00  0.00           H
    ATOM   3270 HG22 THR B 338      -8.714  58.512  71.902  1.00  0.00           H
    ATOM   3271 HG23 THR B 338     -10.110  57.506  71.513  1.00  0.00           H
    ATOM   3272  HG1 THR B 338     -11.119  57.727  74.300  1.00  0.00           H
    ATOM   3273  N   PHE B 339      -7.002  56.079  71.480  1.00 48.39           N
    ATOM   3274  CA  PHE B 339      -5.773  56.322  70.743  1.00 50.77           C
    ATOM   3275  C   PHE B 339      -5.893  57.645  69.982  1.00 50.24           C
    ATOM   3276  O   PHE B 339      -6.631  57.721  69.001  1.00 51.20           O
    ATOM   3277  CB  PHE B 339      -5.545  55.135  69.794  1.00 51.41           C
    ATOM   3278  CG  PHE B 339      -4.171  55.109  69.164  1.00 51.94           C
    ATOM   3279  CD1 PHE B 339      -3.096  54.593  69.910  1.00 47.22           C
    ATOM   3280  CD2 PHE B 339      -3.916  55.732  67.923  1.00 51.98           C
    ATOM   3281  CE1 PHE B 339      -1.816  54.624  69.388  1.00 46.47           C
    ATOM   3282  CE2 PHE B 339      -2.622  55.749  67.416  1.00 50.12           C
    ATOM   3283  CZ  PHE B 339      -1.579  55.189  68.141  1.00 53.33           C
    ATOM   3284  H   PHE B 339      -7.738  55.612  70.964  1.00  0.00           H
    ATOM   3285  HA  PHE B 339      -4.931  56.370  71.438  1.00  0.00           H
    ATOM   3286  HB2 PHE B 339      -5.670  54.209  70.354  1.00  0.00           H
    ATOM   3287  HB3 PHE B 339      -6.297  55.104  69.008  1.00  0.00           H
    ATOM   3288  HD1 PHE B 339      -3.268  54.159  70.885  1.00  0.00           H
    ATOM   3289  HD2 PHE B 339      -4.724  56.178  67.362  1.00  0.00           H
    ATOM   3290  HE1 PHE B 339      -1.015  54.203  69.969  1.00  0.00           H
    ATOM   3291  HE2 PHE B 339      -2.419  56.202  66.457  1.00  0.00           H
    ATOM   3292  HZ  PHE B 339      -0.578  55.203  67.736  1.00  0.00           H
    ATOM   3293  N   LYS B 340      -5.154  58.657  70.450  1.00 52.65           N
    ATOM   3294  CA  LYS B 340      -5.114  59.985  69.847  1.00 58.61           C
    ATOM   3295  C   LYS B 340      -4.005  60.013  68.787  1.00 60.65           C
    ATOM   3296  O   LYS B 340      -2.866  59.669  69.103  1.00 59.68           O
    ATOM   3297  CB  LYS B 340      -4.848  61.034  70.940  1.00 60.31           C
    ATOM   3298  CG  LYS B 340      -5.812  60.970  72.133  1.00 64.13           C
    ATOM   3299  CD  LYS B 340      -5.531  62.088  73.148  1.00 66.66           C
    ATOM   3300  CE  LYS B 340      -6.323  61.923  74.450  1.00 70.22           C
    ATOM   3301  NZ  LYS B 340      -5.894  60.731  75.203  1.00 71.80           N1+
    ATOM   3302  H   LYS B 340      -4.558  58.514  71.253  1.00  0.00           H
    ATOM   3303  HA  LYS B 340      -6.074  60.216  69.381  1.00  0.00           H
    ATOM   3304  HB2 LYS B 340      -3.827  60.940  71.313  1.00  0.00           H
    ATOM   3305  HB3 LYS B 340      -4.928  62.022  70.494  1.00  0.00           H
    ATOM   3306  HG2 LYS B 340      -6.841  61.043  71.781  1.00  0.00           H
    ATOM   3307  HG3 LYS B 340      -5.724  60.000  72.621  1.00  0.00           H
    ATOM   3308  HD2 LYS B 340      -4.463  62.137  73.368  1.00  0.00           H
    ATOM   3309  HD3 LYS B 340      -5.789  63.047  72.699  1.00  0.00           H
    ATOM   3310  HE2 LYS B 340      -6.192  62.799  75.086  1.00  0.00           H
    ATOM   3311  HE3 LYS B 340      -7.388  61.846  74.228  1.00  0.00           H
    ATOM   3312  HZ1 LYS B 340      -4.942  60.836  75.521  1.00  0.00           H
    ATOM   3313  HZ2 LYS B 340      -6.517  60.612  75.993  1.00  0.00           H
    ATOM   3314  HZ3 LYS B 340      -5.973  59.915  74.613  1.00  0.00           H
    ATOM   3315  N   VAL B 341      -4.362  60.414  67.559  1.00 62.93           N
    ATOM   3316  CA  VAL B 341      -3.461  60.418  66.407  1.00 64.81           C
    ATOM   3317  C   VAL B 341      -3.073  61.880  66.104  1.00 67.01           C
    ATOM   3318  O   VAL B 341      -3.958  62.638  65.703  1.00 70.83           O
    ATOM   3319  CB  VAL B 341      -4.180  59.862  65.150  1.00 62.98           C
    ATOM   3320  CG1 VAL B 341      -3.255  59.817  63.917  1.00 63.16           C
    ATOM   3321  CG2 VAL B 341      -4.753  58.464  65.416  1.00 59.12           C
    ATOM   3322  H   VAL B 341      -5.316  60.700  67.380  1.00  0.00           H
    ATOM   3323  HA  VAL B 341      -2.597  59.779  66.584  1.00  0.00           H
    ATOM   3324  HB  VAL B 341      -5.029  60.501  64.901  1.00  0.00           H
    ATOM   3325 HG11 VAL B 341      -3.795  59.453  63.043  1.00  0.00           H
    ATOM   3326 HG12 VAL B 341      -2.859  60.796  63.653  1.00  0.00           H
    ATOM   3327 HG13 VAL B 341      -2.401  59.160  64.086  1.00  0.00           H
    ATOM   3328 HG21 VAL B 341      -5.251  58.082  64.532  1.00  0.00           H
    ATOM   3329 HG22 VAL B 341      -3.963  57.768  65.692  1.00  0.00           H
    ATOM   3330 HG23 VAL B 341      -5.495  58.465  66.215  1.00  0.00           H
    ATOM   3331  N   PRO B 342      -1.789  62.268  66.318  1.00 69.25           N
    ATOM   3332  CA  PRO B 342      -1.293  63.632  66.033  1.00 71.52           C
    ATOM   3333  C   PRO B 342      -1.572  64.113  64.602  1.00 73.82           C
    ATOM   3334  O   PRO B 342      -1.487  63.305  63.680  1.00 73.89           O
    ATOM   3335  CB  PRO B 342       0.220  63.532  66.286  1.00 71.13           C
    ATOM   3336  CG  PRO B 342       0.359  62.424  67.308  1.00 72.59           C
    ATOM   3337  CD  PRO B 342      -0.720  61.440  66.882  1.00 71.81           C
    ATOM   3338  HA  PRO B 342      -1.730  64.311  66.763  1.00  0.00           H
    ATOM   3339  HB2 PRO B 342       0.751  63.244  65.376  1.00  0.00           H
    ATOM   3340  HB3 PRO B 342       0.651  64.472  66.633  1.00  0.00           H
    ATOM   3341  HG2 PRO B 342       1.356  61.986  67.344  1.00  0.00           H
    ATOM   3342  HG3 PRO B 342       0.125  62.816  68.299  1.00  0.00           H
    ATOM   3343  HD2 PRO B 342      -0.349  60.767  66.107  1.00  0.00           H
    ATOM   3344  HD3 PRO B 342      -1.033  60.844  67.738  1.00  0.00           H
    ATOM   3345  N   SER B 343      -1.870  65.413  64.442  1.00 77.89           N
    ATOM   3346  CA  SER B 343      -2.027  66.079  63.139  1.00 81.56           C
    ATOM   3347  C   SER B 343      -0.783  65.994  62.228  1.00 80.75           C
    ATOM   3348  O   SER B 343      -0.931  66.014  61.006  1.00 83.14           O
    ATOM   3349  CB  SER B 343      -2.487  67.529  63.365  1.00 82.17           C
    ATOM   3350  OG  SER B 343      -3.884  67.549  63.568  1.00 78.35           O
    ATOM   3351  H   SER B 343      -1.945  66.010  65.255  1.00  0.00           H
    ATOM   3352  HA  SER B 343      -2.817  65.578  62.585  1.00  0.00           H
    ATOM   3353  HB2 SER B 343      -1.977  67.989  64.213  1.00  0.00           H
    ATOM   3354  HB3 SER B 343      -2.278  68.150  62.492  1.00  0.00           H
    ATOM   3355  HG  SER B 343      -4.099  68.326  64.096  1.00  0.00           H
    ATOM   3356  N   SER B 344       0.403  65.833  62.842  1.00 79.19           N
    ATOM   3357  CA  SER B 344       1.686  65.531  62.203  1.00 79.44           C
    ATOM   3358  C   SER B 344       1.821  64.085  61.673  1.00 81.15           C
    ATOM   3359  O   SER B 344       2.791  63.806  60.969  1.00 83.51           O
    ATOM   3360  CB  SER B 344       2.803  65.868  63.212  1.00 75.98           C
    ATOM   3361  OG  SER B 344       2.813  64.973  64.310  1.00 74.40           O
    ATOM   3362  H   SER B 344       0.422  65.817  63.852  1.00  0.00           H
    ATOM   3363  HA  SER B 344       1.793  66.198  61.346  1.00  0.00           H
    ATOM   3364  HB2 SER B 344       3.777  65.820  62.723  1.00  0.00           H
    ATOM   3365  HB3 SER B 344       2.692  66.888  63.583  1.00  0.00           H
    ATOM   3366  HG  SER B 344       3.546  65.203  64.887  1.00  0.00           H
    ATOM   3367  N   CYS B 345       0.863  63.207  62.009  1.00 81.38           N
    ATOM   3368  CA  CYS B 345       0.785  61.809  61.604  1.00 79.77           C
    ATOM   3369  C   CYS B 345      -0.469  61.645  60.719  1.00 80.59           C
    ATOM   3370  O   CYS B 345      -1.487  61.170  61.228  1.00 81.12           O
    ATOM   3371  CB  CYS B 345       0.785  60.888  62.849  1.00 78.33           C
    ATOM   3372  SG  CYS B 345       0.738  59.127  62.415  1.00 76.91           S
    ATOM   3373  H   CYS B 345       0.097  63.523  62.590  1.00  0.00           H
    ATOM   3374  HA  CYS B 345       1.664  61.527  61.026  1.00  0.00           H
    ATOM   3375  HB2 CYS B 345       1.684  61.065  63.441  1.00  0.00           H
    ATOM   3376  HB3 CYS B 345      -0.061  61.104  63.502  1.00  0.00           H
    ATOM   3377  HG  CYS B 345      -0.490  59.145  61.885  1.00  0.00           H
    ATOM   3378  N   PRO B 346      -0.413  62.082  59.434  1.00 82.82           N
    ATOM   3379  CA  PRO B 346      -1.592  62.077  58.550  1.00 81.77           C
    ATOM   3380  C   PRO B 346      -2.216  60.691  58.333  1.00 80.30           C
    ATOM   3381  O   PRO B 346      -3.439  60.615  58.365  1.00 78.25           O
    ATOM   3382  CB  PRO B 346      -1.108  62.725  57.241  1.00 84.51           C
    ATOM   3383  CG  PRO B 346       0.401  62.552  57.256  1.00 88.29           C
    ATOM   3384  CD  PRO B 346       0.745  62.647  58.736  1.00 86.38           C
    ATOM   3385  HA  PRO B 346      -2.359  62.716  58.993  1.00  0.00           H
    ATOM   3386  HB2 PRO B 346      -1.566  62.305  56.344  1.00  0.00           H
    ATOM   3387  HB3 PRO B 346      -1.354  63.788  57.259  1.00  0.00           H
    ATOM   3388  HG2 PRO B 346       0.664  61.561  56.894  1.00  0.00           H
    ATOM   3389  HG3 PRO B 346       0.928  63.283  56.642  1.00  0.00           H
    ATOM   3390  HD2 PRO B 346       1.683  62.133  58.941  1.00  0.00           H
    ATOM   3391  HD3 PRO B 346       0.848  63.694  59.022  1.00  0.00           H
    ATOM   3392  N   ILE B 347      -1.386  59.640  58.204  1.00 78.69           N
    ATOM   3393  CA  ILE B 347      -1.835  58.253  58.093  1.00 77.95           C
    ATOM   3394  C   ILE B 347      -1.514  57.509  59.405  1.00 78.02           C
    ATOM   3395  O   ILE B 347      -0.347  57.436  59.790  1.00 79.41           O
    ATOM   3396  CB  ILE B 347      -1.147  57.477  56.926  1.00 78.40           C
    ATOM   3397  CG1 ILE B 347      -1.200  58.244  55.584  1.00 82.69           C
    ATOM   3398  CG2 ILE B 347      -1.796  56.090  56.728  1.00 74.11           C
    ATOM   3399  CD1 ILE B 347      -0.072  59.268  55.383  1.00 83.33           C
    ATOM   3400  H   ILE B 347      -0.387  59.781  58.227  1.00  0.00           H
    ATOM   3401  HA  ILE B 347      -2.906  58.220  57.917  1.00  0.00           H
    ATOM   3402  HB  ILE B 347      -0.096  57.303  57.169  1.00  0.00           H
    ATOM   3403 HG12 ILE B 347      -1.147  57.540  54.752  1.00  0.00           H
    ATOM   3404 HG13 ILE B 347      -2.168  58.730  55.485  1.00  0.00           H
    ATOM   3405 HG21 ILE B 347      -1.312  55.549  55.916  1.00  0.00           H
    ATOM   3406 HG22 ILE B 347      -1.715  55.456  57.611  1.00  0.00           H
    ATOM   3407 HG23 ILE B 347      -2.854  56.184  56.481  1.00  0.00           H
    ATOM   3408 HD11 ILE B 347      -0.482  60.267  55.241  1.00  0.00           H
    ATOM   3409 HD12 ILE B 347       0.614  59.300  56.229  1.00  0.00           H
    ATOM   3410 HD13 ILE B 347       0.517  59.028  54.497  1.00  0.00           H
    ATOM   3411  N   VAL B 348      -2.554  56.931  60.024  1.00 76.06           N
    ATOM   3412  CA  VAL B 348      -2.460  55.974  61.130  1.00 71.60           C
    ATOM   3413  C   VAL B 348      -2.902  54.591  60.620  1.00 68.32           C
    ATOM   3414  O   VAL B 348      -3.766  54.523  59.751  1.00 68.65           O
    ATOM   3415  CB  VAL B 348      -3.377  56.398  62.319  1.00 71.55           C
    ATOM   3416  CG1 VAL B 348      -4.864  56.537  61.949  1.00 69.96           C
    ATOM   3417  CG2 VAL B 348      -3.244  55.519  63.580  1.00 71.37           C
    ATOM   3418  H   VAL B 348      -3.480  57.049  59.634  1.00  0.00           H
    ATOM   3419  HA  VAL B 348      -1.430  55.912  61.484  1.00  0.00           H
    ATOM   3420  HB  VAL B 348      -3.035  57.386  62.615  1.00  0.00           H
    ATOM   3421 HG11 VAL B 348      -5.468  56.847  62.795  1.00  0.00           H
    ATOM   3422 HG12 VAL B 348      -4.990  57.301  61.187  1.00  0.00           H
    ATOM   3423 HG13 VAL B 348      -5.296  55.605  61.586  1.00  0.00           H
    ATOM   3424 HG21 VAL B 348      -3.894  55.876  64.377  1.00  0.00           H
    ATOM   3425 HG22 VAL B 348      -3.523  54.482  63.400  1.00  0.00           H
    ATOM   3426 HG23 VAL B 348      -2.227  55.539  63.968  1.00  0.00           H
    ATOM   3427  N   THR B 349      -2.319  53.526  61.184  1.00 64.52           N
    ATOM   3428  CA  THR B 349      -2.681  52.133  60.947  1.00 63.32           C
    ATOM   3429  C   THR B 349      -3.342  51.573  62.218  1.00 62.34           C
    ATOM   3430  O   THR B 349      -2.798  51.747  63.307  1.00 58.60           O
    ATOM   3431  CB  THR B 349      -1.423  51.275  60.648  1.00 63.73           C
    ATOM   3432  CG2 THR B 349      -1.711  49.785  60.377  1.00 65.37           C
    ATOM   3433  OG1 THR B 349      -0.754  51.795  59.515  1.00 68.15           O
    ATOM   3434  H   THR B 349      -1.598  53.674  61.882  1.00  0.00           H
    ATOM   3435  HA  THR B 349      -3.375  52.049  60.112  1.00  0.00           H
    ATOM   3436  HB  THR B 349      -0.727  51.351  61.483  1.00  0.00           H
    ATOM   3437 HG21 THR B 349      -0.798  49.252  60.110  1.00  0.00           H
    ATOM   3438 HG22 THR B 349      -2.121  49.286  61.254  1.00  0.00           H
    ATOM   3439 HG23 THR B 349      -2.422  49.658  59.560  1.00  0.00           H
    ATOM   3440  HG1 THR B 349      -1.261  51.582  58.728  1.00  0.00           H
    ATOM   3441  N   VAL B 350      -4.491  50.904  62.057  1.00 58.33           N
    ATOM   3442  CA  VAL B 350      -5.211  50.251  63.146  1.00 59.59           C
    ATOM   3443  C   VAL B 350      -5.245  48.750  62.826  1.00 61.40           C
    ATOM   3444  O   VAL B 350      -6.105  48.320  62.058  1.00 64.23           O
    ATOM   3445  CB  VAL B 350      -6.661  50.793  63.298  1.00 59.12           C
    ATOM   3446  CG1 VAL B 350      -7.413  50.118  64.459  1.00 57.19           C
    ATOM   3447  CG2 VAL B 350      -6.685  52.315  63.510  1.00 60.25           C
    ATOM   3448  H   VAL B 350      -4.873  50.774  61.128  1.00  0.00           H
    ATOM   3449  HA  VAL B 350      -4.699  50.393  64.097  1.00  0.00           H
    ATOM   3450  HB  VAL B 350      -7.218  50.600  62.380  1.00  0.00           H
    ATOM   3451 HG11 VAL B 350      -8.402  50.548  64.592  1.00  0.00           H
    ATOM   3452 HG12 VAL B 350      -7.542  49.046  64.317  1.00  0.00           H
    ATOM   3453 HG13 VAL B 350      -6.864  50.273  65.385  1.00  0.00           H
    ATOM   3454 HG21 VAL B 350      -7.710  52.667  63.632  1.00  0.00           H
    ATOM   3455 HG22 VAL B 350      -6.122  52.607  64.396  1.00  0.00           H
    ATOM   3456 HG23 VAL B 350      -6.265  52.850  62.657  1.00  0.00           H
    ATOM   3457  N   ASP B 351      -4.268  48.008  63.373  1.00 61.94           N
    ATOM   3458  CA  ASP B 351      -4.010  46.601  63.055  1.00 59.97           C
    ATOM   3459  C   ASP B 351      -4.321  45.695  64.264  1.00 59.03           C
    ATOM   3460  O   ASP B 351      -4.556  46.183  65.368  1.00 56.62           O
    ATOM   3461  CB  ASP B 351      -2.566  46.393  62.522  1.00 52.75           C
    ATOM   3462  CG  ASP B 351      -1.422  46.874  63.419  1.00 55.77           C
    ATOM   3463  OD1 ASP B 351      -1.393  46.434  64.587  1.00 55.37           O
    ATOM   3464  OD2 ASP B 351      -0.507  47.542  62.890  1.00 59.68           O1-
    ATOM   3465  H   ASP B 351      -3.623  48.437  64.023  1.00  0.00           H
    ATOM   3466  HA  ASP B 351      -4.683  46.276  62.262  1.00  0.00           H
    ATOM   3467  HB2 ASP B 351      -2.374  45.355  62.267  1.00  0.00           H
    ATOM   3468  HB3 ASP B 351      -2.486  46.933  61.579  1.00  0.00           H
    ATOM   3469  N   GLY B 352      -4.296  44.379  64.015  1.00 58.14           N
    ATOM   3470  CA  GLY B 352      -4.534  43.312  64.982  1.00 59.74           C
    ATOM   3471  C   GLY B 352      -3.237  42.766  65.585  1.00 61.40           C
    ATOM   3472  O   GLY B 352      -3.296  41.748  66.274  1.00 61.63           O
    ATOM   3473  H   GLY B 352      -4.075  44.075  63.073  1.00  0.00           H
    ATOM   3474  HA2 GLY B 352      -5.218  43.624  65.772  1.00  0.00           H
    ATOM   3475  HA3 GLY B 352      -4.994  42.482  64.454  1.00  0.00           H
    ATOM   3476  N   TYR B 353      -2.082  43.405  65.326  1.00 60.93           N
    ATOM   3477  CA  TYR B 353      -0.781  43.037  65.892  1.00 63.40           C
    ATOM   3478  C   TYR B 353      -0.610  43.607  67.311  1.00 62.50           C
    ATOM   3479  O   TYR B 353      -1.618  43.877  67.962  1.00 62.81           O
    ATOM   3480  CB  TYR B 353       0.339  43.427  64.916  1.00 66.22           C
    ATOM   3481  CG  TYR B 353       0.204  42.806  63.540  1.00 67.44           C
    ATOM   3482  CD1 TYR B 353      -0.029  43.609  62.407  1.00 70.04           C
    ATOM   3483  CD2 TYR B 353       0.307  41.409  63.391  1.00 71.37           C
    ATOM   3484  CE1 TYR B 353      -0.161  43.017  61.139  1.00 69.53           C
    ATOM   3485  CE2 TYR B 353       0.185  40.819  62.121  1.00 71.57           C
    ATOM   3486  CZ  TYR B 353      -0.057  41.622  60.991  1.00 69.48           C
    ATOM   3487  OH  TYR B 353      -0.185  41.051  59.759  1.00 62.59           O
    ATOM   3488  H   TYR B 353      -2.094  44.246  64.764  1.00  0.00           H
    ATOM   3489  HA  TYR B 353      -0.746  41.952  66.007  1.00  0.00           H
    ATOM   3490  HB2 TYR B 353       0.385  44.510  64.830  1.00  0.00           H
    ATOM   3491  HB3 TYR B 353       1.301  43.097  65.306  1.00  0.00           H
    ATOM   3492  HD1 TYR B 353      -0.086  44.680  62.500  1.00  0.00           H
    ATOM   3493  HD2 TYR B 353       0.452  40.784  64.257  1.00  0.00           H
    ATOM   3494  HE1 TYR B 353      -0.327  43.642  60.278  1.00  0.00           H
    ATOM   3495  HE2 TYR B 353       0.262  39.749  62.020  1.00  0.00           H
    ATOM   3496  HH  TYR B 353      -0.113  40.093  59.778  1.00  0.00           H
    ATOM   3497  N   VAL B 354       0.633  43.676  67.823  1.00 61.21           N
    ATOM   3498  CA  VAL B 354       0.869  43.729  69.270  1.00 60.11           C
    ATOM   3499  C   VAL B 354       1.933  44.752  69.734  1.00 60.02           C
    ATOM   3500  O   VAL B 354       2.344  44.667  70.890  1.00 59.91           O
    ATOM   3501  CB  VAL B 354       1.203  42.309  69.830  1.00 56.35           C
    ATOM   3502  CG1 VAL B 354       0.084  41.285  69.573  1.00 51.38           C
    ATOM   3503  CG2 VAL B 354       2.550  41.746  69.359  1.00 53.43           C
    ATOM   3504  H   VAL B 354       1.440  43.489  67.243  1.00  0.00           H
    ATOM   3505  HA  VAL B 354      -0.038  44.067  69.769  1.00  0.00           H
    ATOM   3506  HB  VAL B 354       1.273  42.375  70.916  1.00  0.00           H
    ATOM   3507 HG11 VAL B 354       0.386  40.290  69.889  1.00  0.00           H
    ATOM   3508 HG12 VAL B 354      -0.821  41.549  70.113  1.00  0.00           H
    ATOM   3509 HG13 VAL B 354      -0.170  41.192  68.519  1.00  0.00           H
    ATOM   3510 HG21 VAL B 354       2.721  40.753  69.770  1.00  0.00           H
    ATOM   3511 HG22 VAL B 354       2.588  41.673  68.275  1.00  0.00           H
    ATOM   3512 HG23 VAL B 354       3.379  42.370  69.675  1.00  0.00           H
    ATOM   3513  N   ASP B 355       2.337  45.724  68.891  1.00 63.35           N
    ATOM   3514  CA  ASP B 355       3.282  46.786  69.278  1.00 63.16           C
    ATOM   3515  C   ASP B 355       2.594  47.789  70.239  1.00 64.69           C
    ATOM   3516  O   ASP B 355       1.639  48.447  69.831  1.00 64.57           O
    ATOM   3517  CB  ASP B 355       3.841  47.568  68.066  1.00 60.00           C
    ATOM   3518  CG  ASP B 355       5.222  48.161  68.367  1.00 58.50           C
    ATOM   3519  OD1 ASP B 355       5.287  49.040  69.252  1.00 57.33           O
    ATOM   3520  OD2 ASP B 355       6.192  47.732  67.706  1.00 60.17           O1-
    ATOM   3521  H   ASP B 355       1.971  45.767  67.950  1.00  0.00           H
    ATOM   3522  HA  ASP B 355       4.121  46.271  69.745  1.00  0.00           H
    ATOM   3523  HB2 ASP B 355       3.908  46.940  67.187  1.00  0.00           H
    ATOM   3524  HB3 ASP B 355       3.187  48.380  67.746  1.00  0.00           H
    ATOM   3525  N   PRO B 356       3.083  47.908  71.492  1.00 64.80           N
    ATOM   3526  CA  PRO B 356       2.469  48.800  72.486  1.00 66.06           C
    ATOM   3527  C   PRO B 356       2.780  50.291  72.274  1.00 65.61           C
    ATOM   3528  O   PRO B 356       1.988  51.121  72.721  1.00 66.20           O
    ATOM   3529  CB  PRO B 356       3.047  48.288  73.806  1.00 63.00           C
    ATOM   3530  CG  PRO B 356       4.442  47.823  73.427  1.00 64.23           C
    ATOM   3531  CD  PRO B 356       4.276  47.258  72.027  1.00 64.02           C
    ATOM   3532  HA  PRO B 356       1.386  48.683  72.491  1.00  0.00           H
    ATOM   3533  HB2 PRO B 356       3.055  49.033  74.603  1.00  0.00           H
    ATOM   3534  HB3 PRO B 356       2.455  47.442  74.157  1.00  0.00           H
    ATOM   3535  HG2 PRO B 356       5.127  48.672  73.408  1.00  0.00           H
    ATOM   3536  HG3 PRO B 356       4.838  47.084  74.107  1.00  0.00           H
    ATOM   3537  HD2 PRO B 356       5.165  47.446  71.426  1.00  0.00           H
    ATOM   3538  HD3 PRO B 356       4.114  46.187  72.064  1.00  0.00           H
    ATOM   3539  N   SER B 357       3.903  50.603  71.596  1.00 67.79           N
    ATOM   3540  CA  SER B 357       4.297  51.967  71.245  1.00 69.02           C
    ATOM   3541  C   SER B 357       3.290  52.541  70.243  1.00 72.51           C
    ATOM   3542  O   SER B 357       2.933  51.852  69.286  1.00 76.19           O
    ATOM   3543  CB  SER B 357       5.717  51.991  70.644  1.00 68.14           C
    ATOM   3544  OG  SER B 357       6.673  51.500  71.561  1.00 73.61           O
    ATOM   3545  H   SER B 357       4.476  49.866  71.206  1.00  0.00           H
    ATOM   3546  HA  SER B 357       4.291  52.571  72.155  1.00  0.00           H
    ATOM   3547  HB2 SER B 357       5.775  51.423  69.717  1.00  0.00           H
    ATOM   3548  HB3 SER B 357       5.996  53.015  70.390  1.00  0.00           H
    ATOM   3549  HG  SER B 357       6.778  50.557  71.407  1.00  0.00           H
    ATOM   3550  N   GLY B 358       2.836  53.775  70.509  1.00 70.84           N
    ATOM   3551  CA  GLY B 358       1.783  54.437  69.748  1.00 74.15           C
    ATOM   3552  C   GLY B 358       2.317  54.916  68.398  1.00 74.70           C
    ATOM   3553  O   GLY B 358       2.579  54.105  67.509  1.00 76.23           O
    ATOM   3554  H   GLY B 358       3.184  54.261  71.322  1.00  0.00           H
    ATOM   3555  HA2 GLY B 358       0.986  53.719  69.563  1.00  0.00           H
    ATOM   3556  HA3 GLY B 358       1.365  55.261  70.327  1.00  0.00           H
    ATOM   3557  N   GLY B 359       2.430  56.243  68.233  1.00 74.41           N
    ATOM   3558  CA  GLY B 359       2.890  56.865  66.994  1.00 73.80           C
    ATOM   3559  C   GLY B 359       1.838  56.659  65.897  1.00 71.81           C
    ATOM   3560  O   GLY B 359       0.705  57.121  66.031  1.00 65.45           O
    ATOM   3561  H   GLY B 359       2.174  56.863  68.988  1.00  0.00           H
    ATOM   3562  HA2 GLY B 359       3.029  57.934  67.160  1.00  0.00           H
    ATOM   3563  HA3 GLY B 359       3.857  56.455  66.698  1.00  0.00           H
    ATOM   3564  N   ASP B 360       2.243  55.970  64.820  1.00 71.96           N
    ATOM   3565  CA  ASP B 360       1.417  55.658  63.653  1.00 71.00           C
    ATOM   3566  C   ASP B 360       0.589  54.369  63.806  1.00 67.92           C
    ATOM   3567  O   ASP B 360      -0.282  54.155  62.970  1.00 61.94           O
    ATOM   3568  CB  ASP B 360       2.240  55.589  62.336  1.00 76.93           C
    ATOM   3569  CG  ASP B 360       3.447  54.634  62.314  1.00 83.34           C
    ATOM   3570  OD1 ASP B 360       4.020  54.502  61.211  1.00 88.14           O
    ATOM   3571  OD2 ASP B 360       3.793  54.060  63.371  1.00 89.36           O1-
    ATOM   3572  H   ASP B 360       3.181  55.592  64.787  1.00  0.00           H
    ATOM   3573  HA  ASP B 360       0.688  56.460  63.530  1.00  0.00           H
    ATOM   3574  HB2 ASP B 360       1.579  55.321  61.510  1.00  0.00           H
    ATOM   3575  HB3 ASP B 360       2.619  56.590  62.126  1.00  0.00           H
    ATOM   3576  N   ARG B 361       0.868  53.518  64.807  1.00 64.88           N
    ATOM   3577  CA  ARG B 361       0.244  52.200  64.928  1.00 61.96           C
    ATOM   3578  C   ARG B 361      -0.564  52.086  66.220  1.00 59.19           C
    ATOM   3579  O   ARG B 361      -0.020  52.280  67.306  1.00 59.88           O
    ATOM   3580  CB  ARG B 361       1.325  51.112  64.842  1.00 59.97           C
    ATOM   3581  CG  ARG B 361       1.878  50.958  63.413  1.00 62.90           C
    ATOM   3582  CD  ARG B 361       2.889  49.820  63.284  1.00 64.61           C
    ATOM   3583  NE  ARG B 361       2.245  48.512  63.458  1.00 62.65           N
    ATOM   3584  CZ  ARG B 361       2.842  47.413  63.934  1.00 63.88           C
    ATOM   3585  NH1 ARG B 361       4.138  47.416  64.269  1.00 61.38           N
    ATOM   3586  NH2 ARG B 361       2.116  46.305  64.088  1.00 67.34           N1+
    ATOM   3587  H   ARG B 361       1.590  53.742  65.479  1.00  0.00           H
    ATOM   3588  HA  ARG B 361      -0.436  52.022  64.096  1.00  0.00           H
    ATOM   3589  HB2 ARG B 361       2.132  51.330  65.544  1.00  0.00           H
    ATOM   3590  HB3 ARG B 361       0.899  50.157  65.155  1.00  0.00           H
    ATOM   3591  HG2 ARG B 361       1.026  50.708  62.781  1.00  0.00           H
    ATOM   3592  HG3 ARG B 361       2.288  51.881  63.007  1.00  0.00           H
    ATOM   3593  HD2 ARG B 361       3.522  49.892  62.400  1.00  0.00           H
    ATOM   3594  HD3 ARG B 361       3.554  49.924  64.142  1.00  0.00           H
    ATOM   3595  HE  ARG B 361       1.270  48.437  63.181  1.00  0.00           H
    ATOM   3596 HH11 ARG B 361       4.687  48.257  64.177  1.00  0.00           H
    ATOM   3597 HH12 ARG B 361       4.568  46.585  64.680  1.00  0.00           H
    ATOM   3598 HH21 ARG B 361       1.129  46.318  63.861  1.00  0.00           H
    ATOM   3599 HH22 ARG B 361       2.521  45.501  64.563  1.00  0.00           H
    ATOM   3600  N   PHE B 362      -1.844  51.719  66.054  1.00 58.94           N
    ATOM   3601  CA  PHE B 362      -2.749  51.331  67.125  1.00 59.65           C
    ATOM   3602  C   PHE B 362      -2.984  49.823  67.014  1.00 60.71           C
    ATOM   3603  O   PHE B 362      -3.761  49.372  66.175  1.00 64.84           O
    ATOM   3604  CB  PHE B 362      -4.063  52.131  67.031  1.00 64.31           C
    ATOM   3605  CG  PHE B 362      -5.103  51.842  68.109  1.00 63.14           C
    ATOM   3606  CD1 PHE B 362      -4.722  51.550  69.441  1.00 63.46           C
    ATOM   3607  CD2 PHE B 362      -6.472  52.037  67.829  1.00 62.50           C
    ATOM   3608  CE1 PHE B 362      -5.685  51.378  70.424  1.00 59.68           C
    ATOM   3609  CE2 PHE B 362      -7.421  51.864  68.828  1.00 57.13           C
    ATOM   3610  CZ  PHE B 362      -7.030  51.529  70.118  1.00 52.19           C
    ATOM   3611  H   PHE B 362      -2.201  51.597  65.114  1.00  0.00           H
    ATOM   3612  HA  PHE B 362      -2.280  51.549  68.085  1.00  0.00           H
    ATOM   3613  HB2 PHE B 362      -3.835  53.193  67.059  1.00  0.00           H
    ATOM   3614  HB3 PHE B 362      -4.521  51.960  66.056  1.00  0.00           H
    ATOM   3615  HD1 PHE B 362      -3.684  51.454  69.719  1.00  0.00           H
    ATOM   3616  HD2 PHE B 362      -6.788  52.317  66.835  1.00  0.00           H
    ATOM   3617  HE1 PHE B 362      -5.388  51.139  71.435  1.00  0.00           H
    ATOM   3618  HE2 PHE B 362      -8.469  51.995  68.606  1.00  0.00           H
    ATOM   3619  HZ  PHE B 362      -7.774  51.402  70.890  1.00  0.00           H
    ATOM   3620  N   CYS B 363      -2.265  49.078  67.857  1.00 55.97           N
    ATOM   3621  CA  CYS B 363      -2.264  47.624  67.878  1.00 54.65           C
    ATOM   3622  C   CYS B 363      -3.370  47.128  68.819  1.00 49.06           C
    ATOM   3623  O   CYS B 363      -3.277  47.328  70.030  1.00 51.73           O
    ATOM   3624  CB  CYS B 363      -0.881  47.137  68.314  1.00 55.89           C
    ATOM   3625  SG  CYS B 363       0.304  47.464  66.973  1.00 53.82           S
    ATOM   3626  H   CYS B 363      -1.632  49.528  68.501  1.00  0.00           H
    ATOM   3627  HA  CYS B 363      -2.464  47.234  66.878  1.00  0.00           H
    ATOM   3628  HB2 CYS B 363      -0.579  47.650  69.223  1.00  0.00           H
    ATOM   3629  HB3 CYS B 363      -0.874  46.083  68.556  1.00  0.00           H
    ATOM   3630  HG  CYS B 363      -0.370  46.803  66.021  1.00  0.00           H
    ATOM   3631  N   LEU B 364      -4.407  46.509  68.233  1.00 48.49           N
    ATOM   3632  CA  LEU B 364      -5.578  45.981  68.939  1.00 50.05           C
    ATOM   3633  C   LEU B 364      -5.297  44.625  69.606  1.00 49.82           C
    ATOM   3634  O   LEU B 364      -5.958  44.309  70.594  1.00 52.86           O
    ATOM   3635  CB  LEU B 364      -6.772  45.870  67.959  1.00 53.47           C
    ATOM   3636  CG  LEU B 364      -7.568  47.175  67.729  1.00 50.71           C
    ATOM   3637  CD1 LEU B 364      -6.717  48.423  67.472  1.00 58.11           C
    ATOM   3638  CD2 LEU B 364      -8.597  47.000  66.600  1.00 46.10           C
    ATOM   3639  H   LEU B 364      -4.419  46.414  67.224  1.00  0.00           H
    ATOM   3640  HA  LEU B 364      -5.849  46.673  69.736  1.00  0.00           H
    ATOM   3641  HB2 LEU B 364      -6.427  45.471  67.005  1.00  0.00           H
    ATOM   3642  HB3 LEU B 364      -7.485  45.131  68.330  1.00  0.00           H
    ATOM   3643  HG  LEU B 364      -8.115  47.368  68.651  1.00  0.00           H
    ATOM   3644 HD11 LEU B 364      -7.361  49.285  67.312  1.00  0.00           H
    ATOM   3645 HD12 LEU B 364      -6.079  48.670  68.320  1.00  0.00           H
    ATOM   3646 HD13 LEU B 364      -6.090  48.294  66.590  1.00  0.00           H
    ATOM   3647 HD21 LEU B 364      -9.317  47.818  66.594  1.00  0.00           H
    ATOM   3648 HD22 LEU B 364      -8.114  46.979  65.623  1.00  0.00           H
    ATOM   3649 HD23 LEU B 364      -9.157  46.074  66.714  1.00  0.00           H
    ATOM   3650  N   GLY B 365      -4.328  43.858  69.077  1.00 52.12           N
    ATOM   3651  CA  GLY B 365      -3.966  42.520  69.546  1.00 52.09           C
    ATOM   3652  C   GLY B 365      -3.309  42.513  70.937  1.00 50.38           C
    ATOM   3653  O   GLY B 365      -3.392  41.489  71.613  1.00 43.70           O
    ATOM   3654  H   GLY B 365      -3.803  44.209  68.287  1.00  0.00           H
    ATOM   3655  HA2 GLY B 365      -4.846  41.878  69.534  1.00  0.00           H
    ATOM   3656  HA3 GLY B 365      -3.262  42.074  68.846  1.00  0.00           H
    ATOM   3657  N   GLN B 366      -2.683  43.625  71.371  1.00 50.60           N
    ATOM   3658  CA  GLN B 366      -2.069  43.762  72.702  1.00 58.33           C
    ATOM   3659  C   GLN B 366      -3.010  44.348  73.768  1.00 60.97           C
    ATOM   3660  O   GLN B 366      -2.599  44.432  74.924  1.00 62.67           O
    ATOM   3661  CB  GLN B 366      -0.723  44.520  72.614  1.00 56.94           C
    ATOM   3662  CG  GLN B 366      -0.768  45.975  72.088  1.00 61.60           C
    ATOM   3663  CD  GLN B 366      -0.873  47.059  73.165  1.00 66.59           C
    ATOM   3664  NE2 GLN B 366      -1.364  48.238  72.780  1.00 70.02           N
    ATOM   3665  OE1 GLN B 366      -0.472  46.866  74.311  1.00 65.55           O
    ATOM   3666  H   GLN B 366      -2.638  44.436  70.771  1.00  0.00           H
    ATOM   3667  HA  GLN B 366      -1.814  42.765  73.068  1.00  0.00           H
    ATOM   3668  HB2 GLN B 366      -0.215  44.477  73.579  1.00  0.00           H
    ATOM   3669  HB3 GLN B 366      -0.072  43.937  71.970  1.00  0.00           H
    ATOM   3670  HG2 GLN B 366       0.164  46.182  71.564  1.00  0.00           H
    ATOM   3671  HG3 GLN B 366      -1.554  46.087  71.346  1.00  0.00           H
    ATOM   3672 HE21 GLN B 366      -1.677  48.378  71.830  1.00  0.00           H
    ATOM   3673 HE22 GLN B 366      -1.425  49.000  73.441  1.00  0.00           H
    ATOM   3674  N   LEU B 367      -4.248  44.717  73.396  1.00 62.08           N
    ATOM   3675  CA  LEU B 367      -5.276  45.158  74.340  1.00 60.68           C
    ATOM   3676  C   LEU B 367      -5.898  43.934  75.032  1.00 63.72           C
    ATOM   3677  O   LEU B 367      -6.168  42.939  74.359  1.00 65.57           O
    ATOM   3678  CB  LEU B 367      -6.343  45.994  73.598  1.00 54.35           C
    ATOM   3679  CG  LEU B 367      -5.786  47.259  72.902  1.00 57.82           C
    ATOM   3680  CD1 LEU B 367      -6.855  47.937  72.032  1.00 58.89           C
    ATOM   3681  CD2 LEU B 367      -5.154  48.258  73.889  1.00 59.89           C
    ATOM   3682  H   LEU B 367      -4.532  44.625  72.432  1.00  0.00           H
    ATOM   3683  HA  LEU B 367      -4.810  45.783  75.105  1.00  0.00           H
    ATOM   3684  HB2 LEU B 367      -6.850  45.365  72.866  1.00  0.00           H
    ATOM   3685  HB3 LEU B 367      -7.109  46.302  74.310  1.00  0.00           H
    ATOM   3686  HG  LEU B 367      -5.004  46.942  72.211  1.00  0.00           H
    ATOM   3687 HD11 LEU B 367      -6.394  48.566  71.270  1.00  0.00           H
    ATOM   3688 HD12 LEU B 367      -7.456  47.197  71.508  1.00  0.00           H
    ATOM   3689 HD13 LEU B 367      -7.524  48.564  72.623  1.00  0.00           H
    ATOM   3690 HD21 LEU B 367      -5.469  49.283  73.692  1.00  0.00           H
    ATOM   3691 HD22 LEU B 367      -5.417  48.036  74.923  1.00  0.00           H
    ATOM   3692 HD23 LEU B 367      -4.067  48.235  73.816  1.00  0.00           H
    ATOM   3693  N   SER B 368      -6.105  44.031  76.354  1.00 62.63           N
    ATOM   3694  CA  SER B 368      -6.666  42.964  77.188  1.00 62.36           C
    ATOM   3695  C   SER B 368      -8.103  43.297  77.609  1.00 60.10           C
    ATOM   3696  O   SER B 368      -8.444  44.471  77.753  1.00 55.78           O
    ATOM   3697  CB  SER B 368      -5.770  42.769  78.426  1.00 63.75           C
    ATOM   3698  OG  SER B 368      -4.637  42.008  78.065  1.00 72.16           O
    ATOM   3699  H   SER B 368      -5.864  44.884  76.837  1.00  0.00           H
    ATOM   3700  HA  SER B 368      -6.704  42.021  76.639  1.00  0.00           H
    ATOM   3701  HB2 SER B 368      -5.458  43.722  78.856  1.00  0.00           H
    ATOM   3702  HB3 SER B 368      -6.292  42.222  79.213  1.00  0.00           H
    ATOM   3703  HG  SER B 368      -4.019  42.584  77.604  1.00  0.00           H
    ATOM   3704  N   ASN B 369      -8.907  42.244  77.822  1.00 57.22           N
    ATOM   3705  CA  ASN B 369     -10.295  42.332  78.269  1.00 59.00           C
    ATOM   3706  C   ASN B 369     -10.637  41.041  79.025  1.00 61.85           C
    ATOM   3707  O   ASN B 369     -10.455  39.949  78.485  1.00 64.40           O
    ATOM   3708  CB  ASN B 369     -11.210  42.609  77.047  1.00 51.97           C
    ATOM   3709  CG  ASN B 369     -12.716  42.622  77.346  1.00 54.66           C
    ATOM   3710  ND2 ASN B 369     -13.534  42.260  76.356  1.00 55.11           N
    ATOM   3711  OD1 ASN B 369     -13.142  42.973  78.445  1.00 51.12           O
    ATOM   3712  H   ASN B 369      -8.552  41.306  77.693  1.00  0.00           H
    ATOM   3713  HA  ASN B 369     -10.364  43.178  78.957  1.00  0.00           H
    ATOM   3714  HB2 ASN B 369     -10.955  43.578  76.625  1.00  0.00           H
    ATOM   3715  HB3 ASN B 369     -11.007  41.883  76.261  1.00  0.00           H
    ATOM   3716 HD21 ASN B 369     -13.165  42.015  75.441  1.00  0.00           H
    ATOM   3717 HD22 ASN B 369     -14.534  42.264  76.495  1.00  0.00           H
    ATOM   3718  N   VAL B 370     -11.153  41.207  80.253  1.00 68.07           N
    ATOM   3719  CA  VAL B 370     -11.613  40.121  81.120  1.00 73.40           C
    ATOM   3720  C   VAL B 370     -12.940  39.486  80.652  1.00 77.03           C
    ATOM   3721  O   VAL B 370     -13.143  38.295  80.884  1.00 77.70           O
    ATOM   3722  CB  VAL B 370     -11.798  40.610  82.584  1.00 73.45           C
    ATOM   3723  CG1 VAL B 370     -10.468  41.082  83.192  1.00 72.64           C
    ATOM   3724  CG2 VAL B 370     -12.897  41.672  82.795  1.00 70.81           C
    ATOM   3725  H   VAL B 370     -11.271  42.140  80.619  1.00  0.00           H
    ATOM   3726  HA  VAL B 370     -10.854  39.336  81.113  1.00  0.00           H
    ATOM   3727  HB  VAL B 370     -12.097  39.742  83.171  1.00  0.00           H
    ATOM   3728 HG11 VAL B 370     -10.597  41.363  84.236  1.00  0.00           H
    ATOM   3729 HG12 VAL B 370      -9.715  40.295  83.153  1.00  0.00           H
    ATOM   3730 HG13 VAL B 370     -10.068  41.952  82.669  1.00  0.00           H
    ATOM   3731 HG21 VAL B 370     -12.911  42.002  83.833  1.00  0.00           H
    ATOM   3732 HG22 VAL B 370     -12.736  42.553  82.174  1.00  0.00           H
    ATOM   3733 HG23 VAL B 370     -13.893  41.286  82.580  1.00  0.00           H
    ATOM   3734  N   HIS B 371     -13.796  40.280  79.984  1.00 81.15           N
    ATOM   3735  CA  HIS B 371     -15.083  39.854  79.428  1.00 84.76           C
    ATOM   3736  C   HIS B 371     -14.962  39.130  78.078  1.00 83.23           C
    ATOM   3737  O   HIS B 371     -15.970  38.600  77.611  1.00 85.48           O
    ATOM   3738  CB  HIS B 371     -16.007  41.079  79.287  1.00 85.27           C
    ATOM   3739  CG  HIS B 371     -16.325  41.760  80.592  1.00 88.16           C
    ATOM   3740  CD2 HIS B 371     -16.916  41.287  81.742  1.00 88.19           C
    ATOM   3741  ND1 HIS B 371     -16.046  43.094  80.835  1.00 91.39           N
    ATOM   3742  CE1 HIS B 371     -16.455  43.359  82.079  1.00 90.08           C
    ATOM   3743  NE2 HIS B 371     -16.993  42.309  82.689  1.00 88.20           N
    ATOM   3744  H   HIS B 371     -13.548  41.247  79.824  1.00  0.00           H
    ATOM   3745  HA  HIS B 371     -15.548  39.152  80.123  1.00  0.00           H
    ATOM   3746  HB2 HIS B 371     -15.572  41.808  78.603  1.00  0.00           H
    ATOM   3747  HB3 HIS B 371     -16.959  40.784  78.844  1.00  0.00           H
    ATOM   3748  HD1 HIS B 371     -15.607  43.743  80.197  1.00  0.00           H
    ATOM   3749  HD2 HIS B 371     -17.285  40.296  81.961  1.00  0.00           H
    ATOM   3750  HE1 HIS B 371     -16.354  44.328  82.546  1.00  0.00           H
    ATOM   3751  N   ARG B 372     -13.758  39.131  77.475  1.00 81.93           N
    ATOM   3752  CA  ARG B 372     -13.462  38.540  76.172  1.00 77.54           C
    ATOM   3753  C   ARG B 372     -13.803  37.046  76.120  1.00 76.26           C
    ATOM   3754  O   ARG B 372     -13.393  36.292  77.003  1.00 76.39           O
    ATOM   3755  CB  ARG B 372     -11.983  38.779  75.833  1.00 72.23           C
    ATOM   3756  CG  ARG B 372     -11.580  38.336  74.414  1.00 67.08           C
    ATOM   3757  CD  ARG B 372     -10.136  38.711  74.070  1.00 64.70           C
    ATOM   3758  NE  ARG B 372      -9.973  40.165  73.953  1.00 59.00           N
    ATOM   3759  CZ  ARG B 372      -8.834  40.850  74.136  1.00 56.24           C
    ATOM   3760  NH1 ARG B 372      -7.686  40.246  74.473  1.00 58.07           N
    ATOM   3761  NH2 ARG B 372      -8.847  42.174  73.978  1.00 50.70           N1+
    ATOM   3762  H   ARG B 372     -12.979  39.568  77.945  1.00  0.00           H
    ATOM   3763  HA  ARG B 372     -14.070  39.074  75.439  1.00  0.00           H
    ATOM   3764  HB2 ARG B 372     -11.776  39.840  75.943  1.00  0.00           H
    ATOM   3765  HB3 ARG B 372     -11.353  38.267  76.563  1.00  0.00           H
    ATOM   3766  HG2 ARG B 372     -11.751  37.277  74.219  1.00  0.00           H
    ATOM   3767  HG3 ARG B 372     -12.240  38.877  73.737  1.00  0.00           H
    ATOM   3768  HD2 ARG B 372      -9.538  38.437  74.939  1.00  0.00           H
    ATOM   3769  HD3 ARG B 372      -9.728  38.148  73.230  1.00  0.00           H
    ATOM   3770  HE  ARG B 372     -10.800  40.688  73.680  1.00  0.00           H
    ATOM   3771 HH11 ARG B 372      -7.659  39.246  74.613  1.00  0.00           H
    ATOM   3772 HH12 ARG B 372      -6.845  40.794  74.598  1.00  0.00           H
    ATOM   3773 HH21 ARG B 372      -9.685  42.617  73.605  1.00  0.00           H
    ATOM   3774 HH22 ARG B 372      -7.998  42.712  74.104  1.00  0.00           H
    ATOM   3775  N   THR B 373     -14.549  36.672  75.076  1.00 76.33           N
    ATOM   3776  CA  THR B 373     -15.039  35.319  74.850  1.00 73.36           C
    ATOM   3777  C   THR B 373     -14.187  34.599  73.786  1.00 74.31           C
    ATOM   3778  O   THR B 373     -13.326  35.209  73.147  1.00 75.43           O
    ATOM   3779  CB  THR B 373     -16.515  35.371  74.362  1.00 70.17           C
    ATOM   3780  CG2 THR B 373     -17.448  36.142  75.310  1.00 66.03           C
    ATOM   3781  OG1 THR B 373     -16.641  35.897  73.052  1.00 65.24           O
    ATOM   3782  H   THR B 373     -14.837  37.356  74.392  1.00  0.00           H
    ATOM   3783  HA  THR B 373     -14.997  34.736  75.772  1.00  0.00           H
    ATOM   3784  HB  THR B 373     -16.894  34.349  74.315  1.00  0.00           H
    ATOM   3785 HG21 THR B 373     -18.487  36.064  74.988  1.00  0.00           H
    ATOM   3786 HG22 THR B 373     -17.388  35.747  76.325  1.00  0.00           H
    ATOM   3787 HG23 THR B 373     -17.198  37.203  75.351  1.00  0.00           H
    ATOM   3788  HG1 THR B 373     -17.576  35.958  72.839  1.00  0.00           H
    ATOM   3789  N   GLU B 374     -14.482  33.301  73.616  1.00 77.83           N
    ATOM   3790  CA  GLU B 374     -13.930  32.417  72.589  1.00 79.74           C
    ATOM   3791  C   GLU B 374     -14.134  32.912  71.141  1.00 80.79           C
    ATOM   3792  O   GLU B 374     -13.232  32.743  70.322  1.00 80.63           O
    ATOM   3793  CB  GLU B 374     -14.451  30.971  72.816  1.00 81.56           C
    ATOM   3794  CG  GLU B 374     -15.956  30.778  73.152  1.00 88.21           C
    ATOM   3795  CD  GLU B 374     -16.935  31.183  72.047  1.00 92.98           C
    ATOM   3796  OE1 GLU B 374     -17.838  31.992  72.354  1.00 94.22           O
    ATOM   3797  OE2 GLU B 374     -16.746  30.704  70.908  1.00 96.19           O1-
    ATOM   3798  H   GLU B 374     -15.195  32.883  74.195  1.00  0.00           H
    ATOM   3799  HA  GLU B 374     -12.849  32.396  72.743  1.00  0.00           H
    ATOM   3800  HB2 GLU B 374     -14.207  30.362  71.944  1.00  0.00           H
    ATOM   3801  HB3 GLU B 374     -13.870  30.531  73.627  1.00  0.00           H
    ATOM   3802  HG2 GLU B 374     -16.132  29.723  73.361  1.00  0.00           H
    ATOM   3803  HG3 GLU B 374     -16.211  31.303  74.073  1.00  0.00           H
    ATOM   3804  N   ALA B 375     -15.294  33.533  70.868  1.00 81.16           N
    ATOM   3805  CA  ALA B 375     -15.692  34.032  69.554  1.00 76.77           C
    ATOM   3806  C   ALA B 375     -15.010  35.357  69.194  1.00 76.73           C
    ATOM   3807  O   ALA B 375     -14.776  35.607  68.011  1.00 77.24           O
    ATOM   3808  CB  ALA B 375     -17.220  34.192  69.527  1.00 77.00           C
    ATOM   3809  H   ALA B 375     -15.990  33.612  71.597  1.00  0.00           H
    ATOM   3810  HA  ALA B 375     -15.415  33.290  68.802  1.00  0.00           H
    ATOM   3811  HB1 ALA B 375     -17.563  34.556  68.557  1.00  0.00           H
    ATOM   3812  HB2 ALA B 375     -17.714  33.236  69.706  1.00  0.00           H
    ATOM   3813  HB3 ALA B 375     -17.565  34.891  70.289  1.00  0.00           H
    ATOM   3814  N   ILE B 376     -14.696  36.170  70.216  1.00 78.53           N
    ATOM   3815  CA  ILE B 376     -14.007  37.444  70.057  1.00 78.38           C
    ATOM   3816  C   ILE B 376     -12.498  37.247  69.853  1.00 78.63           C
    ATOM   3817  O   ILE B 376     -11.942  37.860  68.947  1.00 75.97           O
    ATOM   3818  CB  ILE B 376     -14.256  38.404  71.253  1.00 76.83           C
    ATOM   3819  CG1 ILE B 376     -15.749  38.779  71.357  1.00 74.43           C
    ATOM   3820  CG2 ILE B 376     -13.402  39.686  71.190  1.00 78.57           C
    ATOM   3821  CD1 ILE B 376     -16.136  39.472  72.672  1.00 74.07           C
    ATOM   3822  H   ILE B 376     -14.924  35.894  71.162  1.00  0.00           H
    ATOM   3823  HA  ILE B 376     -14.392  37.920  69.158  1.00  0.00           H
    ATOM   3824  HB  ILE B 376     -13.995  37.874  72.171  1.00  0.00           H
    ATOM   3825 HG12 ILE B 376     -16.006  39.433  70.529  1.00  0.00           H
    ATOM   3826 HG13 ILE B 376     -16.378  37.897  71.237  1.00  0.00           H
    ATOM   3827 HG21 ILE B 376     -13.792  40.454  71.851  1.00  0.00           H
    ATOM   3828 HG22 ILE B 376     -12.378  39.495  71.501  1.00  0.00           H
    ATOM   3829 HG23 ILE B 376     -13.376  40.103  70.184  1.00  0.00           H
    ATOM   3830 HD11 ILE B 376     -17.078  39.080  73.056  1.00  0.00           H
    ATOM   3831 HD12 ILE B 376     -15.384  39.329  73.447  1.00  0.00           H
    ATOM   3832 HD13 ILE B 376     -16.262  40.544  72.524  1.00  0.00           H
    ATOM   3833  N   GLU B 377     -11.881  36.364  70.654  1.00 79.48           N
    ATOM   3834  CA  GLU B 377     -10.473  35.977  70.545  1.00 84.48           C
    ATOM   3835  C   GLU B 377     -10.145  35.296  69.197  1.00 86.62           C
    ATOM   3836  O   GLU B 377      -9.042  35.476  68.680  1.00 90.57           O
    ATOM   3837  CB  GLU B 377     -10.138  35.126  71.786  1.00 85.00           C
    ATOM   3838  CG  GLU B 377      -8.642  34.827  72.002  1.00 87.18           C
    ATOM   3839  CD  GLU B 377      -8.369  34.230  73.387  1.00 89.05           C
    ATOM   3840  OE1 GLU B 377      -7.317  34.589  73.961  1.00 87.30           O
    ATOM   3841  OE2 GLU B 377      -9.215  33.439  73.861  1.00 89.46           O1-
    ATOM   3842  H   GLU B 377     -12.413  35.893  71.375  1.00  0.00           H
    ATOM   3843  HA  GLU B 377      -9.881  36.891  70.599  1.00  0.00           H
    ATOM   3844  HB2 GLU B 377     -10.520  35.638  72.671  1.00  0.00           H
    ATOM   3845  HB3 GLU B 377     -10.689  34.185  71.732  1.00  0.00           H
    ATOM   3846  HG2 GLU B 377      -8.275  34.136  71.243  1.00  0.00           H
    ATOM   3847  HG3 GLU B 377      -8.064  35.747  71.898  1.00  0.00           H
    ATOM   3848  N   ARG B 378     -11.151  34.611  68.623  1.00 86.14           N
    ATOM   3849  CA  ARG B 378     -11.156  34.051  67.276  1.00 84.27           C
    ATOM   3850  C   ARG B 378     -11.240  35.129  66.176  1.00 82.23           C
    ATOM   3851  O   ARG B 378     -10.503  35.017  65.205  1.00 82.21           O
    ATOM   3852  CB  ARG B 378     -12.298  33.015  67.193  1.00 85.03           C
    ATOM   3853  CG  ARG B 378     -12.420  32.250  65.862  1.00 83.73           C
    ATOM   3854  CD  ARG B 378     -13.588  31.249  65.859  1.00 84.92           C
    ATOM   3855  NE  ARG B 378     -14.891  31.927  65.795  1.00 87.14           N
    ATOM   3856  CZ  ARG B 378     -16.088  31.370  66.049  1.00 89.46           C
    ATOM   3857  NH1 ARG B 378     -16.197  30.088  66.424  1.00 90.42           N
    ATOM   3858  NH2 ARG B 378     -17.196  32.110  65.924  1.00 91.67           N1+
    ATOM   3859  H   ARG B 378     -12.016  34.512  69.136  1.00  0.00           H
    ATOM   3860  HA  ARG B 378     -10.212  33.517  67.141  1.00  0.00           H
    ATOM   3861  HB2 ARG B 378     -12.144  32.281  67.983  1.00  0.00           H
    ATOM   3862  HB3 ARG B 378     -13.247  33.502  67.414  1.00  0.00           H
    ATOM   3863  HG2 ARG B 378     -12.619  32.984  65.080  1.00  0.00           H
    ATOM   3864  HG3 ARG B 378     -11.490  31.757  65.580  1.00  0.00           H
    ATOM   3865  HD2 ARG B 378     -13.471  30.464  65.111  1.00  0.00           H
    ATOM   3866  HD3 ARG B 378     -13.592  30.752  66.829  1.00  0.00           H
    ATOM   3867  HE  ARG B 378     -14.886  32.852  65.388  1.00  0.00           H
    ATOM   3868 HH11 ARG B 378     -15.371  29.514  66.503  1.00  0.00           H
    ATOM   3869 HH12 ARG B 378     -17.106  29.678  66.581  1.00  0.00           H
    ATOM   3870 HH21 ARG B 378     -17.155  33.052  65.562  1.00  0.00           H
    ATOM   3871 HH22 ARG B 378     -18.106  31.693  66.053  1.00  0.00           H
    ATOM   3872  N   ALA B 379     -12.103  36.150  66.335  1.00 76.94           N
    ATOM   3873  CA  ALA B 379     -12.264  37.243  65.365  1.00 75.30           C
    ATOM   3874  C   ALA B 379     -11.077  38.225  65.334  1.00 76.27           C
    ATOM   3875  O   ALA B 379     -10.694  38.664  64.251  1.00 78.21           O
    ATOM   3876  CB  ALA B 379     -13.572  37.991  65.650  1.00 74.47           C
    ATOM   3877  H   ALA B 379     -12.689  36.188  67.157  1.00  0.00           H
    ATOM   3878  HA  ALA B 379     -12.347  36.800  64.371  1.00  0.00           H
    ATOM   3879  HB1 ALA B 379     -13.741  38.785  64.921  1.00  0.00           H
    ATOM   3880  HB2 ALA B 379     -14.428  37.316  65.603  1.00  0.00           H
    ATOM   3881  HB3 ALA B 379     -13.561  38.445  66.641  1.00  0.00           H
    ATOM   3882  N   ARG B 380     -10.497  38.523  66.511  1.00 73.16           N
    ATOM   3883  CA  ARG B 380      -9.306  39.365  66.694  1.00 74.12           C
    ATOM   3884  C   ARG B 380      -8.052  38.836  65.983  1.00 76.60           C
    ATOM   3885  O   ARG B 380      -7.221  39.638  65.557  1.00 76.74           O
    ATOM   3886  CB  ARG B 380      -9.001  39.487  68.193  1.00 71.02           C
    ATOM   3887  CG  ARG B 380      -9.970  40.391  68.963  1.00 66.08           C
    ATOM   3888  CD  ARG B 380      -9.877  40.152  70.474  1.00 60.41           C
    ATOM   3889  NE  ARG B 380      -8.510  40.295  70.993  1.00 52.68           N
    ATOM   3890  CZ  ARG B 380      -7.775  41.417  71.019  1.00 47.28           C
    ATOM   3891  NH1 ARG B 380      -8.244  42.585  70.564  1.00 47.46           N
    ATOM   3892  NH2 ARG B 380      -6.538  41.365  71.520  1.00 45.88           N1+
    ATOM   3893  H   ARG B 380     -10.890  38.129  67.357  1.00  0.00           H
    ATOM   3894  HA  ARG B 380      -9.541  40.350  66.297  1.00  0.00           H
    ATOM   3895  HB2 ARG B 380      -8.986  38.488  68.634  1.00  0.00           H
    ATOM   3896  HB3 ARG B 380      -7.999  39.895  68.335  1.00  0.00           H
    ATOM   3897  HG2 ARG B 380      -9.863  41.442  68.693  1.00  0.00           H
    ATOM   3898  HG3 ARG B 380     -10.985  40.135  68.667  1.00  0.00           H
    ATOM   3899  HD2 ARG B 380     -10.481  40.895  70.982  1.00  0.00           H
    ATOM   3900  HD3 ARG B 380     -10.290  39.182  70.742  1.00  0.00           H
    ATOM   3901  HE  ARG B 380      -8.083  39.442  71.324  1.00  0.00           H
    ATOM   3902 HH11 ARG B 380      -9.180  42.654  70.190  1.00  0.00           H
    ATOM   3903 HH12 ARG B 380      -7.652  43.407  70.583  1.00  0.00           H
    ATOM   3904 HH21 ARG B 380      -6.145  40.492  71.840  1.00  0.00           H
    ATOM   3905 HH22 ARG B 380      -5.975  42.204  71.569  1.00  0.00           H
    ATOM   3906  N   LEU B 381      -7.966  37.502  65.854  1.00 77.75           N
    ATOM   3907  CA  LEU B 381      -6.941  36.751  65.128  1.00 78.27           C
    ATOM   3908  C   LEU B 381      -6.834  37.151  63.639  1.00 75.54           C
    ATOM   3909  O   LEU B 381      -5.749  37.037  63.070  1.00 71.62           O
    ATOM   3910  CB  LEU B 381      -7.284  35.243  65.275  1.00 84.95           C
    ATOM   3911  CG  LEU B 381      -6.156  34.345  65.811  1.00 86.04           C
    ATOM   3912  CD1 LEU B 381      -5.823  34.672  67.280  1.00 88.87           C
    ATOM   3913  CD2 LEU B 381      -6.498  32.852  65.622  1.00 86.37           C
    ATOM   3914  H   LEU B 381      -8.706  36.937  66.248  1.00  0.00           H
    ATOM   3915  HA  LEU B 381      -5.982  36.967  65.602  1.00  0.00           H
    ATOM   3916  HB2 LEU B 381      -8.125  35.109  65.954  1.00  0.00           H
    ATOM   3917  HB3 LEU B 381      -7.630  34.847  64.319  1.00  0.00           H
    ATOM   3918  HG  LEU B 381      -5.266  34.554  65.214  1.00  0.00           H
    ATOM   3919 HD11 LEU B 381      -4.744  34.712  67.411  1.00  0.00           H
    ATOM   3920 HD12 LEU B 381      -6.232  35.631  67.597  1.00  0.00           H
    ATOM   3921 HD13 LEU B 381      -6.208  33.925  67.975  1.00  0.00           H
    ATOM   3922 HD21 LEU B 381      -6.335  32.262  66.524  1.00  0.00           H
    ATOM   3923 HD22 LEU B 381      -7.537  32.703  65.325  1.00  0.00           H
    ATOM   3924 HD23 LEU B 381      -5.874  32.414  64.842  1.00  0.00           H
    ATOM   3925  N   HIS B 382      -7.957  37.605  63.052  1.00 72.47           N
    ATOM   3926  CA  HIS B 382      -8.128  37.888  61.625  1.00 71.90           C
    ATOM   3927  C   HIS B 382      -8.211  39.377  61.279  1.00 70.22           C
    ATOM   3928  O   HIS B 382      -8.400  39.684  60.100  1.00 64.78           O
    ATOM   3929  CB  HIS B 382      -9.383  37.162  61.117  1.00 74.92           C
    ATOM   3930  CG  HIS B 382      -9.298  35.677  61.283  1.00 81.51           C
    ATOM   3931  CD2 HIS B 382      -8.768  34.740  60.426  1.00 85.85           C
    ATOM   3932  ND1 HIS B 382      -9.754  35.038  62.434  1.00 82.51           N
    ATOM   3933  CE1 HIS B 382      -9.492  33.755  62.241  1.00 82.86           C
    ATOM   3934  NE2 HIS B 382      -8.919  33.526  61.062  1.00 84.84           N
    ATOM   3935  H   HIS B 382      -8.798  37.681  63.609  1.00  0.00           H
    ATOM   3936  HA  HIS B 382      -7.273  37.497  61.071  1.00  0.00           H
    ATOM   3937  HB2 HIS B 382     -10.268  37.507  61.651  1.00  0.00           H
    ATOM   3938  HB3 HIS B 382      -9.561  37.370  60.062  1.00  0.00           H
    ATOM   3939  HD2 HIS B 382      -8.314  34.834  59.451  1.00  0.00           H
    ATOM   3940  HE2 HIS B 382      -8.681  32.615  60.691  1.00  0.00           H
    ATOM   3941  HE1 HIS B 382      -9.738  32.981  62.953  1.00  0.00           H
    ATOM   3942  N   ILE B 383      -8.031  40.281  62.262  1.00 69.55           N
    ATOM   3943  CA  ILE B 383      -7.893  41.719  62.002  1.00 69.43           C
    ATOM   3944  C   ILE B 383      -6.673  41.998  61.092  1.00 70.64           C
    ATOM   3945  O   ILE B 383      -6.815  42.760  60.139  1.00 73.52           O
    ATOM   3946  CB  ILE B 383      -7.786  42.564  63.309  1.00 66.84           C
    ATOM   3947  CG1 ILE B 383      -9.055  42.437  64.181  1.00 65.06           C
    ATOM   3948  CG2 ILE B 383      -7.463  44.061  63.069  1.00 65.07           C
    ATOM   3949  CD1 ILE B 383      -8.892  43.004  65.601  1.00 63.17           C
    ATOM   3950  H   ILE B 383      -7.884  39.970  63.212  1.00  0.00           H
    ATOM   3951  HA  ILE B 383      -8.788  42.042  61.466  1.00  0.00           H
    ATOM   3952  HB  ILE B 383      -6.982  42.126  63.893  1.00  0.00           H
    ATOM   3953 HG12 ILE B 383      -9.884  42.945  63.692  1.00  0.00           H
    ATOM   3954 HG13 ILE B 383      -9.359  41.394  64.257  1.00  0.00           H
    ATOM   3955 HG21 ILE B 383      -7.362  44.610  64.004  1.00  0.00           H
    ATOM   3956 HG22 ILE B 383      -6.531  44.216  62.529  1.00  0.00           H
    ATOM   3957 HG23 ILE B 383      -8.251  44.539  62.491  1.00  0.00           H
    ATOM   3958 HD11 ILE B 383      -9.695  42.663  66.253  1.00  0.00           H
    ATOM   3959 HD12 ILE B 383      -7.946  42.700  66.050  1.00  0.00           H
    ATOM   3960 HD13 ILE B 383      -8.930  44.092  65.594  1.00  0.00           H
    ATOM   3961  N   GLY B 384      -5.544  41.307  61.346  1.00 71.98           N
    ATOM   3962  CA  GLY B 384      -4.348  41.313  60.498  1.00 74.15           C
    ATOM   3963  C   GLY B 384      -3.722  42.709  60.473  1.00 74.90           C
    ATOM   3964  O   GLY B 384      -3.587  43.337  61.518  1.00 76.27           O
    ATOM   3965  H   GLY B 384      -5.526  40.685  62.141  1.00  0.00           H
    ATOM   3966  HA2 GLY B 384      -3.622  40.603  60.896  1.00  0.00           H
    ATOM   3967  HA3 GLY B 384      -4.608  40.983  59.490  1.00  0.00           H
    ATOM   3968  N   LYS B 385      -3.366  43.195  59.274  1.00 77.87           N
    ATOM   3969  CA  LYS B 385      -2.911  44.571  59.029  1.00 78.94           C
    ATOM   3970  C   LYS B 385      -4.028  45.629  59.186  1.00 78.59           C
    ATOM   3971  O   LYS B 385      -3.716  46.820  59.210  1.00 79.01           O
    ATOM   3972  CB  LYS B 385      -2.285  44.646  57.625  1.00 82.13           C
    ATOM   3973  CG  LYS B 385      -1.095  43.701  57.401  1.00 84.50           C
    ATOM   3974  CD  LYS B 385      -0.555  43.769  55.964  1.00 86.94           C
    ATOM   3975  CE  LYS B 385       0.530  42.720  55.674  1.00 88.02           C
    ATOM   3976  NZ  LYS B 385       1.706  42.880  56.547  1.00 91.60           N1+
    ATOM   3977  H   LYS B 385      -3.485  42.620  58.453  1.00  0.00           H
    ATOM   3978  HA  LYS B 385      -2.145  44.820  59.762  1.00  0.00           H
    ATOM   3979  HB2 LYS B 385      -3.055  44.417  56.891  1.00  0.00           H
    ATOM   3980  HB3 LYS B 385      -1.952  45.667  57.435  1.00  0.00           H
    ATOM   3981  HG2 LYS B 385      -0.299  43.959  58.098  1.00  0.00           H
    ATOM   3982  HG3 LYS B 385      -1.383  42.673  57.621  1.00  0.00           H
    ATOM   3983  HD2 LYS B 385      -1.377  43.632  55.260  1.00  0.00           H
    ATOM   3984  HD3 LYS B 385      -0.159  44.767  55.771  1.00  0.00           H
    ATOM   3985  HE2 LYS B 385       0.127  41.715  55.808  1.00  0.00           H
    ATOM   3986  HE3 LYS B 385       0.853  42.797  54.635  1.00  0.00           H
    ATOM   3987  HZ1 LYS B 385       1.427  42.768  57.512  1.00  0.00           H
    ATOM   3988  HZ2 LYS B 385       2.397  42.178  56.319  1.00  0.00           H
    ATOM   3989  HZ3 LYS B 385       2.105  43.799  56.416  1.00  0.00           H
    ATOM   3990  N   GLY B 386      -5.288  45.174  59.321  1.00 78.16           N
    ATOM   3991  CA  GLY B 386      -6.468  45.965  59.650  1.00 75.03           C
    ATOM   3992  C   GLY B 386      -6.757  46.994  58.562  1.00 73.78           C
    ATOM   3993  O   GLY B 386      -6.920  46.639  57.398  1.00 76.36           O
    ATOM   3994  H   GLY B 386      -5.436  44.175  59.290  1.00  0.00           H
    ATOM   3995  HA2 GLY B 386      -7.332  45.310  59.749  1.00  0.00           H
    ATOM   3996  HA3 GLY B 386      -6.330  46.424  60.621  1.00  0.00           H
    ATOM   3997  N   VAL B 387      -6.849  48.266  58.960  1.00 69.23           N
    ATOM   3998  CA  VAL B 387      -7.036  49.384  58.045  1.00 66.33           C
    ATOM   3999  C   VAL B 387      -5.974  50.459  58.294  1.00 66.71           C
    ATOM   4000  O   VAL B 387      -5.209  50.380  59.255  1.00 64.47           O
    ATOM   4001  CB  VAL B 387      -8.435  50.045  58.211  1.00 64.88           C
    ATOM   4002  CG1 VAL B 387      -9.560  49.047  57.930  1.00 64.65           C
    ATOM   4003  CG2 VAL B 387      -8.658  50.769  59.558  1.00 65.13           C
    ATOM   4004  H   VAL B 387      -6.692  48.503  59.931  1.00  0.00           H
    ATOM   4005  HA  VAL B 387      -6.914  49.054  57.014  1.00  0.00           H
    ATOM   4006  HB  VAL B 387      -8.533  50.799  57.429  1.00  0.00           H
    ATOM   4007 HG11 VAL B 387     -10.498  49.567  57.755  1.00  0.00           H
    ATOM   4008 HG12 VAL B 387      -9.356  48.442  57.048  1.00  0.00           H
    ATOM   4009 HG13 VAL B 387      -9.703  48.376  58.773  1.00  0.00           H
    ATOM   4010 HG21 VAL B 387      -9.650  51.211  59.606  1.00  0.00           H
    ATOM   4011 HG22 VAL B 387      -8.563  50.079  60.396  1.00  0.00           H
    ATOM   4012 HG23 VAL B 387      -7.958  51.588  59.720  1.00  0.00           H
    ATOM   4013  N   GLN B 388      -6.011  51.478  57.431  1.00 70.31           N
    ATOM   4014  CA  GLN B 388      -5.400  52.773  57.656  1.00 72.21           C
    ATOM   4015  C   GLN B 388      -6.482  53.850  57.597  1.00 74.68           C
    ATOM   4016  O   GLN B 388      -7.475  53.672  56.896  1.00 76.10           O
    ATOM   4017  CB  GLN B 388      -4.323  53.035  56.594  1.00 69.48           C
    ATOM   4018  CG  GLN B 388      -3.160  52.030  56.670  1.00 76.97           C
    ATOM   4019  CD  GLN B 388      -1.975  52.370  55.765  1.00 81.15           C
    ATOM   4020  NE2 GLN B 388      -2.175  53.185  54.726  1.00 85.02           N
    ATOM   4021  OE1 GLN B 388      -0.872  51.884  56.001  1.00 82.13           O
    ATOM   4022  H   GLN B 388      -6.642  51.428  56.642  1.00  0.00           H
    ATOM   4023  HA  GLN B 388      -4.960  52.788  58.648  1.00  0.00           H
    ATOM   4024  HB2 GLN B 388      -4.773  53.017  55.601  1.00  0.00           H
    ATOM   4025  HB3 GLN B 388      -3.932  54.043  56.740  1.00  0.00           H
    ATOM   4026  HG2 GLN B 388      -2.798  51.978  57.696  1.00  0.00           H
    ATOM   4027  HG3 GLN B 388      -3.507  51.029  56.412  1.00  0.00           H
    ATOM   4028 HE21 GLN B 388      -3.086  53.584  54.555  1.00  0.00           H
    ATOM   4029 HE22 GLN B 388      -1.399  53.436  54.131  1.00  0.00           H
    ATOM   4030  N   LEU B 389      -6.244  54.952  58.318  1.00 76.58           N
    ATOM   4031  CA  LEU B 389      -7.065  56.156  58.302  1.00 76.97           C
    ATOM   4032  C   LEU B 389      -6.116  57.304  57.929  1.00 81.08           C
    ATOM   4033  O   LEU B 389      -5.153  57.526  58.659  1.00 79.52           O
    ATOM   4034  CB  LEU B 389      -7.724  56.390  59.686  1.00 71.77           C
    ATOM   4035  CG  LEU B 389      -8.406  55.170  60.341  1.00 67.89           C
    ATOM   4036  CD1 LEU B 389      -8.785  55.487  61.795  1.00 65.69           C
    ATOM   4037  CD2 LEU B 389      -9.626  54.700  59.537  1.00 64.30           C
    ATOM   4038  H   LEU B 389      -5.395  55.006  58.869  1.00  0.00           H
    ATOM   4039  HA  LEU B 389      -7.854  56.073  57.557  1.00  0.00           H
    ATOM   4040  HB2 LEU B 389      -6.986  56.779  60.380  1.00  0.00           H
    ATOM   4041  HB3 LEU B 389      -8.471  57.178  59.588  1.00  0.00           H
    ATOM   4042  HG  LEU B 389      -7.693  54.347  60.394  1.00  0.00           H
    ATOM   4043 HD11 LEU B 389      -9.077  54.585  62.333  1.00  0.00           H
    ATOM   4044 HD12 LEU B 389      -7.948  55.934  62.332  1.00  0.00           H
    ATOM   4045 HD13 LEU B 389      -9.612  56.194  61.843  1.00  0.00           H
    ATOM   4046 HD21 LEU B 389     -10.464  54.441  60.181  1.00  0.00           H
    ATOM   4047 HD22 LEU B 389      -9.971  55.478  58.858  1.00  0.00           H
    ATOM   4048 HD23 LEU B 389      -9.388  53.815  58.945  1.00  0.00           H
    ATOM   4049  N   GLU B 390      -6.378  57.989  56.806  1.00 83.90           N
    ATOM   4050  CA  GLU B 390      -5.566  59.100  56.303  1.00 86.49           C
    ATOM   4051  C   GLU B 390      -6.362  60.406  56.338  1.00 86.40           C
    ATOM   4052  O   GLU B 390      -7.479  60.411  55.837  1.00 87.21           O
    ATOM   4053  CB  GLU B 390      -5.065  58.783  54.877  1.00 87.98           C
    ATOM   4054  CG  GLU B 390      -4.383  59.977  54.159  1.00 88.40           C
    ATOM   4055  CD  GLU B 390      -3.862  59.647  52.764  1.00 87.15           C
    ATOM   4056  OE1 GLU B 390      -3.550  58.463  52.511  1.00 83.07           O
    ATOM   4057  OE2 GLU B 390      -3.808  60.593  51.948  1.00 87.03           O1-
    ATOM   4058  H   GLU B 390      -7.193  57.747  56.255  1.00  0.00           H
    ATOM   4059  HA  GLU B 390      -4.681  59.219  56.914  1.00  0.00           H
    ATOM   4060  HB2 GLU B 390      -4.366  57.949  54.942  1.00  0.00           H
    ATOM   4061  HB3 GLU B 390      -5.895  58.422  54.269  1.00  0.00           H
    ATOM   4062  HG2 GLU B 390      -5.086  60.802  54.036  1.00  0.00           H
    ATOM   4063  HG3 GLU B 390      -3.559  60.366  54.757  1.00  0.00           H
    ATOM   4064  N   CYS B 391      -5.745  61.493  56.832  1.00 86.35           N
    ATOM   4065  CA  CYS B 391      -6.283  62.852  56.777  1.00 85.20           C
    ATOM   4066  C   CYS B 391      -5.645  63.614  55.604  1.00 87.33           C
    ATOM   4067  O   CYS B 391      -4.435  63.846  55.631  1.00 87.29           O
    ATOM   4068  CB  CYS B 391      -6.080  63.588  58.113  1.00 81.37           C
    ATOM   4069  SG  CYS B 391      -6.893  65.215  58.143  1.00 74.64           S
    ATOM   4070  H   CYS B 391      -4.817  61.398  57.226  1.00  0.00           H
    ATOM   4071  HA  CYS B 391      -7.358  62.793  56.622  1.00  0.00           H
    ATOM   4072  HB2 CYS B 391      -6.523  62.989  58.899  1.00  0.00           H
    ATOM   4073  HB3 CYS B 391      -5.022  63.701  58.354  1.00  0.00           H
    ATOM   4074  HG  CYS B 391      -6.077  65.796  57.258  1.00  0.00           H
    ATOM   4075  N   LYS B 392      -6.470  63.998  54.614  1.00 89.89           N
    ATOM   4076  CA  LYS B 392      -6.062  64.785  53.449  1.00 90.38           C
    ATOM   4077  C   LYS B 392      -6.243  66.295  53.689  1.00 91.96           C
    ATOM   4078  O   LYS B 392      -5.290  66.942  54.122  1.00 92.24           O
    ATOM   4079  CB  LYS B 392      -6.790  64.298  52.174  1.00 85.26           C
    ATOM   4080  CG  LYS B 392      -6.225  62.997  51.579  1.00 83.19           C
    ATOM   4081  CD  LYS B 392      -6.596  62.853  50.092  1.00 85.34           C
    ATOM   4082  CE  LYS B 392      -6.191  61.513  49.464  1.00 85.16           C
    ATOM   4083  NZ  LYS B 392      -4.730  61.371  49.368  1.00 84.67           N1+
    ATOM   4084  H   LYS B 392      -7.457  63.784  54.686  1.00  0.00           H
    ATOM   4085  HA  LYS B 392      -4.994  64.635  53.280  1.00  0.00           H
    ATOM   4086  HB2 LYS B 392      -7.866  64.210  52.336  1.00  0.00           H
    ATOM   4087  HB3 LYS B 392      -6.665  65.065  51.408  1.00  0.00           H
    ATOM   4088  HG2 LYS B 392      -5.140  62.980  51.684  1.00  0.00           H
    ATOM   4089  HG3 LYS B 392      -6.600  62.144  52.147  1.00  0.00           H
    ATOM   4090  HD2 LYS B 392      -7.664  63.011  49.961  1.00  0.00           H
    ATOM   4091  HD3 LYS B 392      -6.120  63.658  49.531  1.00  0.00           H
    ATOM   4092  HE2 LYS B 392      -6.602  60.680  50.036  1.00  0.00           H
    ATOM   4093  HE3 LYS B 392      -6.606  61.441  48.458  1.00  0.00           H
    ATOM   4094  HZ1 LYS B 392      -4.339  61.339  50.305  1.00  0.00           H
    ATOM   4095  HZ2 LYS B 392      -4.511  60.498  48.909  1.00  0.00           H
    ATOM   4096  HZ3 LYS B 392      -4.331  62.142  48.854  1.00  0.00           H
    ATOM   4097  N   GLY B 393      -7.418  66.841  53.329  1.00 92.80           N
    ATOM   4098  CA  GLY B 393      -7.633  68.282  53.221  1.00 95.97           C
    ATOM   4099  C   GLY B 393      -8.382  68.791  54.449  1.00 97.57           C
    ATOM   4100  O   GLY B 393      -9.611  68.742  54.466  1.00 98.37           O
    ATOM   4101  H   GLY B 393      -8.161  66.245  52.996  1.00  0.00           H
    ATOM   4102  HA2 GLY B 393      -6.701  68.832  53.081  1.00  0.00           H
    ATOM   4103  HA3 GLY B 393      -8.240  68.472  52.335  1.00  0.00           H
    ATOM   4104  N   GLU B 394      -7.630  69.319  55.435  1.00 97.02           N
    ATOM   4105  CA  GLU B 394      -8.071  70.004  56.664  1.00 96.46           C
    ATOM   4106  C   GLU B 394      -8.777  69.090  57.691  1.00 94.54           C
    ATOM   4107  O   GLU B 394      -8.458  69.170  58.877  1.00 93.86           O
    ATOM   4108  CB  GLU B 394      -8.887  71.276  56.313  1.00 96.28           C
    ATOM   4109  CG  GLU B 394      -9.258  72.190  57.506  1.00 97.35           C
    ATOM   4110  CD  GLU B 394     -10.170  73.365  57.135  1.00 98.02           C
    ATOM   4111  OE1 GLU B 394     -10.803  73.301  56.061  1.00 95.75           O
    ATOM   4112  OE2 GLU B 394     -10.256  74.313  57.949  1.00 97.73           O1-
    ATOM   4113  H   GLU B 394      -6.628  69.287  55.318  1.00  0.00           H
    ATOM   4114  HA  GLU B 394      -7.149  70.339  57.141  1.00  0.00           H
    ATOM   4115  HB2 GLU B 394      -8.322  71.867  55.590  1.00  0.00           H
    ATOM   4116  HB3 GLU B 394      -9.798  70.977  55.795  1.00  0.00           H
    ATOM   4117  HG2 GLU B 394      -9.784  71.623  58.264  1.00  0.00           H
    ATOM   4118  HG3 GLU B 394      -8.352  72.576  57.974  1.00  0.00           H
    ATOM   4119  N   GLY B 395      -9.706  68.243  57.225  1.00 92.09           N
    ATOM   4120  CA  GLY B 395     -10.441  67.301  58.052  1.00 88.42           C
    ATOM   4121  C   GLY B 395     -11.030  66.144  57.232  1.00 87.19           C
    ATOM   4122  O   GLY B 395     -11.733  65.344  57.836  1.00 86.84           O
    ATOM   4123  H   GLY B 395      -9.925  68.266  56.238  1.00  0.00           H
    ATOM   4124  HA2 GLY B 395      -9.790  66.876  58.819  1.00  0.00           H
    ATOM   4125  HA3 GLY B 395     -11.249  67.821  58.566  1.00  0.00           H
    ATOM   4126  N   ASP B 396     -10.788  66.031  55.908  1.00 84.81           N
    ATOM   4127  CA  ASP B 396     -11.269  64.914  55.072  1.00 86.14           C
    ATOM   4128  C   ASP B 396     -10.507  63.616  55.361  1.00 85.32           C
    ATOM   4129  O   ASP B 396      -9.278  63.647  55.356  1.00 84.99           O
    ATOM   4130  CB  ASP B 396     -11.215  65.175  53.547  1.00 86.37           C
    ATOM   4131  CG  ASP B 396     -12.107  66.306  53.032  1.00 88.66           C
    ATOM   4132  OD1 ASP B 396     -12.428  67.229  53.811  1.00 90.45           O
    ATOM   4133  OD2 ASP B 396     -12.445  66.229  51.831  1.00 92.77           O1-
    ATOM   4134  H   ASP B 396     -10.236  66.729  55.431  1.00  0.00           H
    ATOM   4135  HA  ASP B 396     -12.317  64.754  55.333  1.00  0.00           H
    ATOM   4136  HB2 ASP B 396     -10.190  65.392  53.243  1.00  0.00           H
    ATOM   4137  HB3 ASP B 396     -11.518  64.263  53.030  1.00  0.00           H
    ATOM   4138  N   VAL B 397     -11.244  62.507  55.559  1.00 83.81           N
    ATOM   4139  CA  VAL B 397     -10.666  61.212  55.922  1.00 83.04           C
    ATOM   4140  C   VAL B 397     -10.892  60.166  54.818  1.00 81.00           C
    ATOM   4141  O   VAL B 397     -11.975  60.099  54.238  1.00 79.65           O
    ATOM   4142  CB  VAL B 397     -11.228  60.655  57.257  1.00 82.68           C
    ATOM   4143  CG1 VAL B 397     -10.564  59.340  57.725  1.00 88.40           C
    ATOM   4144  CG2 VAL B 397     -11.100  61.711  58.362  1.00 82.09           C
    ATOM   4145  H   VAL B 397     -12.250  62.549  55.481  1.00  0.00           H
    ATOM   4146  HA  VAL B 397      -9.602  61.341  56.072  1.00  0.00           H
    ATOM   4147  HB  VAL B 397     -12.286  60.429  57.138  1.00  0.00           H
    ATOM   4148 HG11 VAL B 397     -10.918  59.048  58.714  1.00  0.00           H
    ATOM   4149 HG12 VAL B 397     -10.787  58.505  57.061  1.00  0.00           H
    ATOM   4150 HG13 VAL B 397      -9.479  59.441  57.782  1.00  0.00           H
    ATOM   4151 HG21 VAL B 397     -11.401  61.291  59.312  1.00  0.00           H
    ATOM   4152 HG22 VAL B 397     -10.076  62.073  58.453  1.00  0.00           H
    ATOM   4153 HG23 VAL B 397     -11.739  62.573  58.180  1.00  0.00           H
    ATOM   4154  N   TRP B 398      -9.848  59.359  54.586  1.00 80.99           N
    ATOM   4155  CA  TRP B 398      -9.807  58.221  53.674  1.00 83.53           C
    ATOM   4156  C   TRP B 398      -9.454  56.950  54.450  1.00 83.70           C
    ATOM   4157  O   TRP B 398      -8.711  57.025  55.426  1.00 83.75           O
    ATOM   4158  CB  TRP B 398      -8.746  58.482  52.588  1.00 88.22           C
    ATOM   4159  CG  TRP B 398      -9.115  59.494  51.550  1.00 89.82           C
    ATOM   4160  CD1 TRP B 398      -9.262  60.824  51.746  1.00 89.52           C
    ATOM   4161  CD2 TRP B 398      -9.433  59.257  50.147  1.00 88.76           C
    ATOM   4162  CE2 TRP B 398      -9.761  60.508  49.541  1.00 90.39           C
    ATOM   4163  CE3 TRP B 398      -9.480  58.109  49.321  1.00 86.92           C
    ATOM   4164  NE1 TRP B 398      -9.661  61.421  50.569  1.00 90.18           N
    ATOM   4165  CZ2 TRP B 398     -10.107  60.615  48.182  1.00 90.75           C
    ATOM   4166  CZ3 TRP B 398      -9.849  58.200  47.964  1.00 91.49           C
    ATOM   4167  CH2 TRP B 398     -10.160  59.450  47.394  1.00 91.82           C
    ATOM   4168  H   TRP B 398      -8.997  59.513  55.115  1.00  0.00           H
    ATOM   4169  HA  TRP B 398     -10.779  58.062  53.209  1.00  0.00           H
    ATOM   4170  HB2 TRP B 398      -7.802  58.785  53.041  1.00  0.00           H
    ATOM   4171  HB3 TRP B 398      -8.537  57.553  52.056  1.00  0.00           H
    ATOM   4172  HD1 TRP B 398      -9.105  61.328  52.689  1.00  0.00           H
    ATOM   4173  HE1 TRP B 398      -9.843  62.412  50.499  1.00  0.00           H
    ATOM   4174  HE3 TRP B 398      -9.240  57.142  49.739  1.00  0.00           H
    ATOM   4175  HZ2 TRP B 398     -10.351  61.576  47.754  1.00  0.00           H
    ATOM   4176  HZ3 TRP B 398      -9.887  57.307  47.359  1.00  0.00           H
    ATOM   4177  HH2 TRP B 398     -10.439  59.514  46.353  1.00  0.00           H
    ATOM   4178  N   VAL B 399      -9.975  55.807  53.979  1.00 81.80           N
    ATOM   4179  CA  VAL B 399      -9.741  54.487  54.560  1.00 80.60           C
    ATOM   4180  C   VAL B 399      -9.141  53.560  53.499  1.00 81.26           C
    ATOM   4181  O   VAL B 399      -9.678  53.497  52.394  1.00 81.09           O
    ATOM   4182  CB  VAL B 399     -11.055  53.840  55.087  1.00 80.69           C
    ATOM   4183  CG1 VAL B 399     -10.914  52.377  55.567  1.00 81.25           C
    ATOM   4184  CG2 VAL B 399     -11.625  54.676  56.235  1.00 81.88           C
    ATOM   4185  H   VAL B 399     -10.563  55.827  53.155  1.00  0.00           H
    ATOM   4186  HA  VAL B 399      -9.046  54.575  55.387  1.00  0.00           H
    ATOM   4187  HB  VAL B 399     -11.794  53.852  54.283  1.00  0.00           H
    ATOM   4188 HG11 VAL B 399     -11.833  52.028  56.040  1.00  0.00           H
    ATOM   4189 HG12 VAL B 399     -10.706  51.689  54.748  1.00  0.00           H
    ATOM   4190 HG13 VAL B 399     -10.114  52.280  56.301  1.00  0.00           H
    ATOM   4191 HG21 VAL B 399     -12.589  54.297  56.574  1.00  0.00           H
    ATOM   4192 HG22 VAL B 399     -10.945  54.668  57.081  1.00  0.00           H
    ATOM   4193 HG23 VAL B 399     -11.754  55.710  55.937  1.00  0.00           H
    ATOM   4194  N   ARG B 400      -8.081  52.827  53.878  1.00 81.59           N
    ATOM   4195  CA  ARG B 400      -7.509  51.752  53.073  1.00 78.77           C
    ATOM   4196  C   ARG B 400      -7.552  50.450  53.876  1.00 76.17           C
    ATOM   4197  O   ARG B 400      -6.913  50.387  54.923  1.00 74.28           O
    ATOM   4198  CB  ARG B 400      -6.068  52.094  52.648  1.00 80.28           C
    ATOM   4199  CG  ARG B 400      -5.505  51.100  51.614  1.00 81.33           C
    ATOM   4200  CD  ARG B 400      -4.124  51.499  51.074  1.00 83.99           C
    ATOM   4201  NE  ARG B 400      -4.222  52.568  50.067  1.00 87.65           N
    ATOM   4202  CZ  ARG B 400      -3.334  53.548  49.830  1.00 89.52           C
    ATOM   4203  NH1 ARG B 400      -2.253  53.726  50.602  1.00 92.34           N
    ATOM   4204  NH2 ARG B 400      -3.535  54.368  48.792  1.00 90.94           N1+
    ATOM   4205  H   ARG B 400      -7.709  52.934  54.813  1.00  0.00           H
    ATOM   4206  HA  ARG B 400      -8.095  51.617  52.170  1.00  0.00           H
    ATOM   4207  HB2 ARG B 400      -6.026  53.099  52.237  1.00  0.00           H
    ATOM   4208  HB3 ARG B 400      -5.417  52.115  53.522  1.00  0.00           H
    ATOM   4209  HG2 ARG B 400      -5.505  50.070  51.967  1.00  0.00           H
    ATOM   4210  HG3 ARG B 400      -6.201  51.109  50.773  1.00  0.00           H
    ATOM   4211  HD2 ARG B 400      -3.593  51.948  51.913  1.00  0.00           H
    ATOM   4212  HD3 ARG B 400      -3.517  50.653  50.753  1.00  0.00           H
    ATOM   4213  HE  ARG B 400      -4.999  52.479  49.429  1.00  0.00           H
    ATOM   4214 HH11 ARG B 400      -2.084  53.106  51.380  1.00  0.00           H
    ATOM   4215 HH12 ARG B 400      -1.602  54.475  50.414  1.00  0.00           H
    ATOM   4216 HH21 ARG B 400      -4.356  54.269  48.213  1.00  0.00           H
    ATOM   4217 HH22 ARG B 400      -2.885  55.116  48.597  1.00  0.00           H
    ATOM   4218  N   CYS B 401      -8.263  49.433  53.366  1.00 72.85           N
    ATOM   4219  CA  CYS B 401      -8.335  48.105  53.977  1.00 73.28           C
    ATOM   4220  C   CYS B 401      -7.084  47.288  53.611  1.00 76.38           C
    ATOM   4221  O   CYS B 401      -7.009  46.757  52.507  1.00 79.53           O
    ATOM   4222  CB  CYS B 401      -9.652  47.395  53.600  1.00 68.20           C
    ATOM   4223  SG  CYS B 401      -9.813  45.761  54.378  1.00 70.50           S
    ATOM   4224  H   CYS B 401      -8.748  49.552  52.486  1.00  0.00           H
    ATOM   4225  HA  CYS B 401      -8.346  48.226  55.060  1.00  0.00           H
    ATOM   4226  HB2 CYS B 401     -10.500  48.001  53.920  1.00  0.00           H
    ATOM   4227  HB3 CYS B 401      -9.730  47.282  52.519  1.00  0.00           H
    ATOM   4228  HG  CYS B 401     -11.011  45.473  53.861  1.00  0.00           H
    ATOM   4229  N   LEU B 402      -6.124  47.209  54.544  1.00 77.48           N
    ATOM   4230  CA  LEU B 402      -4.892  46.424  54.413  1.00 78.96           C
    ATOM   4231  C   LEU B 402      -5.081  44.946  54.827  1.00 80.95           C
    ATOM   4232  O   LEU B 402      -4.195  44.139  54.549  1.00 83.44           O
    ATOM   4233  CB  LEU B 402      -3.776  47.106  55.245  1.00 76.05           C
    ATOM   4234  CG  LEU B 402      -2.845  48.046  54.451  1.00 77.27           C
    ATOM   4235  CD1 LEU B 402      -3.576  49.220  53.780  1.00 79.39           C
    ATOM   4236  CD2 LEU B 402      -1.694  48.545  55.341  1.00 78.97           C
    ATOM   4237  H   LEU B 402      -6.259  47.670  55.434  1.00  0.00           H
    ATOM   4238  HA  LEU B 402      -4.583  46.410  53.365  1.00  0.00           H
    ATOM   4239  HB2 LEU B 402      -4.201  47.620  56.110  1.00  0.00           H
    ATOM   4240  HB3 LEU B 402      -3.126  46.340  55.659  1.00  0.00           H
    ATOM   4241  HG  LEU B 402      -2.387  47.459  53.654  1.00  0.00           H
    ATOM   4242 HD11 LEU B 402      -3.083  50.176  53.955  1.00  0.00           H
    ATOM   4243 HD12 LEU B 402      -3.609  49.066  52.703  1.00  0.00           H
    ATOM   4244 HD13 LEU B 402      -4.597  49.328  54.137  1.00  0.00           H
    ATOM   4245 HD21 LEU B 402      -1.146  49.364  54.876  1.00  0.00           H
    ATOM   4246 HD22 LEU B 402      -2.056  48.892  56.309  1.00  0.00           H
    ATOM   4247 HD23 LEU B 402      -0.975  47.746  55.527  1.00  0.00           H
    ATOM   4248  N   SER B 403      -6.208  44.619  55.483  1.00 79.50           N
    ATOM   4249  CA  SER B 403      -6.566  43.288  55.969  1.00 77.42           C
    ATOM   4250  C   SER B 403      -6.969  42.344  54.826  1.00 76.43           C
    ATOM   4251  O   SER B 403      -7.522  42.791  53.821  1.00 75.05           O
    ATOM   4252  CB  SER B 403      -7.723  43.439  56.981  1.00 75.53           C
    ATOM   4253  OG  SER B 403      -8.100  42.212  57.576  1.00 81.13           O
    ATOM   4254  H   SER B 403      -6.875  45.350  55.688  1.00  0.00           H
    ATOM   4255  HA  SER B 403      -5.699  42.871  56.486  1.00  0.00           H
    ATOM   4256  HB2 SER B 403      -7.430  44.107  57.784  1.00  0.00           H
    ATOM   4257  HB3 SER B 403      -8.597  43.888  56.509  1.00  0.00           H
    ATOM   4258  HG  SER B 403      -7.620  42.118  58.406  1.00  0.00           H
    ATOM   4259  N   ASP B 404      -6.739  41.041  55.054  1.00 78.83           N
    ATOM   4260  CA  ASP B 404      -7.209  39.930  54.215  1.00 80.08           C
    ATOM   4261  C   ASP B 404      -8.737  39.734  54.295  1.00 80.34           C
    ATOM   4262  O   ASP B 404      -9.320  39.175  53.366  1.00 80.69           O
    ATOM   4263  CB  ASP B 404      -6.508  38.587  54.546  1.00 79.93           C
    ATOM   4264  CG  ASP B 404      -4.979  38.638  54.677  1.00 82.32           C
    ATOM   4265  OD1 ASP B 404      -4.444  37.675  55.267  1.00 80.01           O
    ATOM   4266  OD2 ASP B 404      -4.362  39.586  54.145  1.00 87.94           O1-
    ATOM   4267  H   ASP B 404      -6.255  40.771  55.898  1.00  0.00           H
    ATOM   4268  HA  ASP B 404      -6.973  40.182  53.179  1.00  0.00           H
    ATOM   4269  HB2 ASP B 404      -6.901  38.216  55.494  1.00  0.00           H
    ATOM   4270  HB3 ASP B 404      -6.751  37.851  53.778  1.00  0.00           H
    ATOM   4271  N   HIS B 405      -9.345  40.199  55.398  1.00 81.61           N
    ATOM   4272  CA  HIS B 405     -10.785  40.199  55.641  1.00 82.39           C
    ATOM   4273  C   HIS B 405     -11.362  41.594  55.372  1.00 81.52           C
    ATOM   4274  O   HIS B 405     -10.643  42.591  55.444  1.00 77.03           O
    ATOM   4275  CB  HIS B 405     -11.054  39.758  57.097  1.00 81.78           C
    ATOM   4276  CG  HIS B 405     -10.847  38.282  57.343  1.00 81.37           C
    ATOM   4277  CD2 HIS B 405     -11.710  37.300  57.776  1.00 79.22           C
    ATOM   4278  ND1 HIS B 405      -9.643  37.636  57.117  1.00 80.80           N
    ATOM   4279  CE1 HIS B 405      -9.826  36.343  57.391  1.00 76.08           C
    ATOM   4280  NE2 HIS B 405     -11.057  36.065  57.801  1.00 76.76           N
    ATOM   4281  H   HIS B 405      -8.791  40.684  56.092  1.00  0.00           H
    ATOM   4282  HA  HIS B 405     -11.285  39.499  54.970  1.00  0.00           H
    ATOM   4283  HB2 HIS B 405     -10.425  40.318  57.791  1.00  0.00           H
    ATOM   4284  HB3 HIS B 405     -12.084  39.988  57.375  1.00  0.00           H
    ATOM   4285  HD1 HIS B 405      -8.788  38.063  56.789  1.00  0.00           H
    ATOM   4286  HD2 HIS B 405     -12.750  37.387  58.053  1.00  0.00           H
    ATOM   4287  HE1 HIS B 405      -9.051  35.598  57.289  1.00  0.00           H
    ATOM   4288  N   ALA B 406     -12.671  41.632  55.080  1.00 83.05           N
    ATOM   4289  CA  ALA B 406     -13.434  42.862  54.879  1.00 84.34           C
    ATOM   4290  C   ALA B 406     -13.624  43.659  56.179  1.00 84.94           C
    ATOM   4291  O   ALA B 406     -13.483  43.114  57.277  1.00 84.87           O
    ATOM   4292  CB  ALA B 406     -14.793  42.497  54.269  1.00 80.97           C
    ATOM   4293  H   ALA B 406     -13.202  40.773  55.052  1.00  0.00           H
    ATOM   4294  HA  ALA B 406     -12.891  43.489  54.168  1.00  0.00           H
    ATOM   4295  HB1 ALA B 406     -15.376  43.390  54.045  1.00  0.00           H
    ATOM   4296  HB2 ALA B 406     -14.668  41.950  53.334  1.00  0.00           H
    ATOM   4297  HB3 ALA B 406     -15.381  41.875  54.946  1.00  0.00           H
    ATOM   4298  N   VAL B 407     -13.949  44.947  56.009  1.00 85.85           N
    ATOM   4299  CA  VAL B 407     -14.209  45.887  57.095  1.00 87.90           C
    ATOM   4300  C   VAL B 407     -15.459  46.711  56.749  1.00 87.25           C
    ATOM   4301  O   VAL B 407     -15.671  47.054  55.587  1.00 87.62           O
    ATOM   4302  CB  VAL B 407     -12.997  46.828  57.344  1.00 88.99           C
    ATOM   4303  CG1 VAL B 407     -11.757  46.026  57.780  1.00 86.46           C
    ATOM   4304  CG2 VAL B 407     -12.666  47.775  56.173  1.00 91.84           C
    ATOM   4305  H   VAL B 407     -14.056  45.319  55.075  1.00  0.00           H
    ATOM   4306  HA  VAL B 407     -14.420  45.337  58.014  1.00  0.00           H
    ATOM   4307  HB  VAL B 407     -13.249  47.463  58.196  1.00  0.00           H
    ATOM   4308 HG11 VAL B 407     -10.947  46.677  58.091  1.00  0.00           H
    ATOM   4309 HG12 VAL B 407     -11.987  45.375  58.623  1.00  0.00           H
    ATOM   4310 HG13 VAL B 407     -11.376  45.399  56.975  1.00  0.00           H
    ATOM   4311 HG21 VAL B 407     -11.718  48.286  56.326  1.00  0.00           H
    ATOM   4312 HG22 VAL B 407     -12.588  47.233  55.231  1.00  0.00           H
    ATOM   4313 HG23 VAL B 407     -13.423  48.551  56.064  1.00  0.00           H
    ATOM   4314  N   PHE B 408     -16.283  46.970  57.769  1.00 86.05           N
    ATOM   4315  CA  PHE B 408     -17.636  47.499  57.634  1.00 88.93           C
    ATOM   4316  C   PHE B 408     -17.679  48.901  58.237  1.00 87.66           C
    ATOM   4317  O   PHE B 408     -17.611  49.016  59.456  1.00 91.37           O
    ATOM   4318  CB  PHE B 408     -18.602  46.544  58.364  1.00 91.48           C
    ATOM   4319  CG  PHE B 408     -18.746  45.181  57.718  1.00 91.14           C
    ATOM   4320  CD1 PHE B 408     -17.766  44.184  57.920  1.00 89.49           C
    ATOM   4321  CD2 PHE B 408     -19.774  44.950  56.783  1.00 93.21           C
    ATOM   4322  CE1 PHE B 408     -17.857  42.977  57.243  1.00 91.22           C
    ATOM   4323  CE2 PHE B 408     -19.854  43.730  56.131  1.00 94.43           C
    ATOM   4324  CZ  PHE B 408     -18.906  42.743  56.364  1.00 93.10           C
    ATOM   4325  H   PHE B 408     -16.018  46.677  58.701  1.00  0.00           H
    ATOM   4326  HA  PHE B 408     -17.941  47.556  56.587  1.00  0.00           H
    ATOM   4327  HB2 PHE B 408     -18.259  46.383  59.385  1.00  0.00           H
    ATOM   4328  HB3 PHE B 408     -19.591  46.995  58.441  1.00  0.00           H
    ATOM   4329  HD1 PHE B 408     -16.943  44.354  58.599  1.00  0.00           H
    ATOM   4330  HD2 PHE B 408     -20.504  45.713  56.568  1.00  0.00           H
    ATOM   4331  HE1 PHE B 408     -17.106  42.217  57.393  1.00  0.00           H
    ATOM   4332  HE2 PHE B 408     -20.649  43.564  55.422  1.00  0.00           H
    ATOM   4333  HZ  PHE B 408     -18.973  41.798  55.844  1.00  0.00           H
    ATOM   4334  N   VAL B 409     -17.765  49.928  57.382  1.00 87.27           N
    ATOM   4335  CA  VAL B 409     -17.726  51.336  57.772  1.00 89.77           C
    ATOM   4336  C   VAL B 409     -19.145  51.933  57.798  1.00 90.82           C
    ATOM   4337  O   VAL B 409     -19.951  51.651  56.913  1.00 90.00           O
    ATOM   4338  CB  VAL B 409     -16.864  52.168  56.775  1.00 89.71           C
    ATOM   4339  CG1 VAL B 409     -16.796  53.678  57.097  1.00 86.61           C
    ATOM   4340  CG2 VAL B 409     -15.439  51.602  56.686  1.00 90.63           C
    ATOM   4341  H   VAL B 409     -17.850  49.748  56.388  1.00  0.00           H
    ATOM   4342  HA  VAL B 409     -17.280  51.432  58.764  1.00  0.00           H
    ATOM   4343  HB  VAL B 409     -17.299  52.080  55.777  1.00  0.00           H
    ATOM   4344 HG11 VAL B 409     -16.092  54.191  56.443  1.00  0.00           H
    ATOM   4345 HG12 VAL B 409     -17.756  54.173  56.954  1.00  0.00           H
    ATOM   4346 HG13 VAL B 409     -16.476  53.851  58.125  1.00  0.00           H
    ATOM   4347 HG21 VAL B 409     -14.771  52.272  56.148  1.00  0.00           H
    ATOM   4348 HG22 VAL B 409     -15.022  51.463  57.677  1.00  0.00           H
    ATOM   4349 HG23 VAL B 409     -15.419  50.635  56.182  1.00  0.00           H
    ATOM   4350  N   GLN B 410     -19.388  52.789  58.800  1.00 91.40           N
    ATOM   4351  CA  GLN B 410     -20.549  53.663  58.910  1.00 89.57           C
    ATOM   4352  C   GLN B 410     -20.075  55.103  58.695  1.00 89.26           C
    ATOM   4353  O   GLN B 410     -19.191  55.567  59.417  1.00 83.65           O
    ATOM   4354  CB  GLN B 410     -21.176  53.486  60.301  1.00 90.38           C
    ATOM   4355  CG  GLN B 410     -22.411  54.375  60.554  1.00 92.86           C
    ATOM   4356  CD  GLN B 410     -23.088  54.076  61.891  1.00 94.70           C
    ATOM   4357  NE2 GLN B 410     -24.421  54.117  61.913  1.00 96.83           N
    ATOM   4358  OE1 GLN B 410     -22.426  53.834  62.898  1.00 95.64           O
    ATOM   4359  H   GLN B 410     -18.671  52.936  59.501  1.00  0.00           H
    ATOM   4360  HA  GLN B 410     -21.297  53.407  58.156  1.00  0.00           H
    ATOM   4361  HB2 GLN B 410     -21.453  52.441  60.418  1.00  0.00           H
    ATOM   4362  HB3 GLN B 410     -20.421  53.686  61.059  1.00  0.00           H
    ATOM   4363  HG2 GLN B 410     -22.137  55.430  60.550  1.00  0.00           H
    ATOM   4364  HG3 GLN B 410     -23.130  54.235  59.745  1.00  0.00           H
    ATOM   4365 HE21 GLN B 410     -24.941  54.322  61.072  1.00  0.00           H
    ATOM   4366 HE22 GLN B 410     -24.919  53.937  62.772  1.00  0.00           H
    ATOM   4367  N   SER B 411     -20.693  55.776  57.718  1.00 92.11           N
    ATOM   4368  CA  SER B 411     -20.360  57.136  57.323  1.00 97.74           C
    ATOM   4369  C   SER B 411     -21.577  57.784  56.666  1.00102.51           C
    ATOM   4370  O   SER B 411     -22.131  57.196  55.739  1.00104.43           O
    ATOM   4371  CB  SER B 411     -19.164  57.098  56.352  1.00 94.87           C
    ATOM   4372  OG  SER B 411     -19.013  58.338  55.696  1.00 89.43           O
    ATOM   4373  H   SER B 411     -21.429  55.335  57.185  1.00  0.00           H
    ATOM   4374  HA  SER B 411     -20.090  57.723  58.204  1.00  0.00           H
    ATOM   4375  HB2 SER B 411     -18.241  56.838  56.865  1.00  0.00           H
    ATOM   4376  HB3 SER B 411     -19.324  56.331  55.599  1.00  0.00           H
    ATOM   4377  HG  SER B 411     -18.071  58.549  55.642  1.00  0.00           H
    ATOM   4378  N   TYR B 412     -21.893  59.020  57.085  1.00106.52           N
    ATOM   4379  CA  TYR B 412     -22.904  59.857  56.438  1.00109.92           C
    ATOM   4380  C   TYR B 412     -22.472  60.406  55.078  1.00110.04           C
    ATOM   4381  O   TYR B 412     -23.352  60.615  54.247  1.00111.08           O
    ATOM   4382  CB  TYR B 412     -23.297  61.033  57.338  1.00113.80           C
    ATOM   4383  CG  TYR B 412     -23.944  60.673  58.657  1.00117.33           C
    ATOM   4384  CD1 TYR B 412     -24.930  59.671  58.752  1.00118.93           C
    ATOM   4385  CD2 TYR B 412     -23.592  61.408  59.799  1.00118.34           C
    ATOM   4386  CE1 TYR B 412     -25.557  59.412  59.986  1.00119.28           C
    ATOM   4387  CE2 TYR B 412     -24.246  61.178  61.022  1.00118.97           C
    ATOM   4388  CZ  TYR B 412     -25.227  60.178  61.117  1.00118.90           C
    ATOM   4389  OH  TYR B 412     -25.874  59.963  62.296  1.00118.03           O
    ATOM   4390  H   TYR B 412     -21.385  59.439  57.851  1.00  0.00           H
    ATOM   4391  HA  TYR B 412     -23.783  59.241  56.256  1.00  0.00           H
    ATOM   4392  HB2 TYR B 412     -22.419  61.655  57.523  1.00  0.00           H
    ATOM   4393  HB3 TYR B 412     -24.010  61.673  56.815  1.00  0.00           H
    ATOM   4394  HD1 TYR B 412     -25.231  59.105  57.886  1.00  0.00           H
    ATOM   4395  HD2 TYR B 412     -22.850  62.177  59.702  1.00  0.00           H
    ATOM   4396  HE1 TYR B 412     -26.310  58.643  60.057  1.00  0.00           H
    ATOM   4397  HE2 TYR B 412     -24.021  61.786  61.879  1.00  0.00           H
    ATOM   4398  HH  TYR B 412     -26.542  59.276  62.235  1.00  0.00           H
    ATOM   4399  N   TYR B 413     -21.157  60.602  54.855  1.00109.56           N
    ATOM   4400  CA  TYR B 413     -20.609  60.927  53.538  1.00108.68           C
    ATOM   4401  C   TYR B 413     -20.895  59.799  52.533  1.00107.96           C
    ATOM   4402  O   TYR B 413     -21.365  60.089  51.438  1.00108.30           O
    ATOM   4403  CB  TYR B 413     -19.105  61.277  53.633  1.00107.38           C
    ATOM   4404  CG  TYR B 413     -18.432  61.546  52.295  1.00108.86           C
    ATOM   4405  CD1 TYR B 413     -18.682  62.755  51.617  1.00109.29           C
    ATOM   4406  CD2 TYR B 413     -17.603  60.573  51.696  1.00108.33           C
    ATOM   4407  CE1 TYR B 413     -18.139  62.976  50.337  1.00107.90           C
    ATOM   4408  CE2 TYR B 413     -17.056  60.798  50.417  1.00105.71           C
    ATOM   4409  CZ  TYR B 413     -17.335  61.995  49.731  1.00104.25           C
    ATOM   4410  OH  TYR B 413     -16.833  62.205  48.481  1.00 98.00           O
    ATOM   4411  H   TYR B 413     -20.488  60.388  55.582  1.00  0.00           H
    ATOM   4412  HA  TYR B 413     -21.129  61.820  53.184  1.00  0.00           H
    ATOM   4413  HB2 TYR B 413     -18.970  62.153  54.269  1.00  0.00           H
    ATOM   4414  HB3 TYR B 413     -18.570  60.466  54.125  1.00  0.00           H
    ATOM   4415  HD1 TYR B 413     -19.305  63.508  52.074  1.00  0.00           H
    ATOM   4416  HD2 TYR B 413     -17.406  59.639  52.202  1.00  0.00           H
    ATOM   4417  HE1 TYR B 413     -18.340  63.899  49.817  1.00  0.00           H
    ATOM   4418  HE2 TYR B 413     -16.440  60.041  49.955  1.00  0.00           H
    ATOM   4419  HH  TYR B 413     -17.237  62.957  48.039  1.00  0.00           H
    ATOM   4420  N   LEU B 414     -20.684  58.540  52.954  1.00106.76           N
    ATOM   4421  CA  LEU B 414     -20.980  57.352  52.156  1.00106.07           C
    ATOM   4422  C   LEU B 414     -22.490  57.083  51.995  1.00105.78           C
    ATOM   4423  O   LEU B 414     -22.882  56.650  50.912  1.00104.51           O
    ATOM   4424  CB  LEU B 414     -20.253  56.125  52.745  1.00105.76           C
    ATOM   4425  CG  LEU B 414     -18.710  56.227  52.771  1.00106.31           C
    ATOM   4426  CD1 LEU B 414     -18.091  55.017  53.505  1.00107.64           C
    ATOM   4427  CD2 LEU B 414     -18.118  56.415  51.361  1.00105.32           C
    ATOM   4428  H   LEU B 414     -20.318  58.378  53.883  1.00  0.00           H
    ATOM   4429  HA  LEU B 414     -20.588  57.528  51.154  1.00  0.00           H
    ATOM   4430  HB2 LEU B 414     -20.629  55.942  53.752  1.00  0.00           H
    ATOM   4431  HB3 LEU B 414     -20.519  55.243  52.162  1.00  0.00           H
    ATOM   4432  HG  LEU B 414     -18.439  57.110  53.348  1.00  0.00           H
    ATOM   4433 HD11 LEU B 414     -17.455  55.347  54.326  1.00  0.00           H
    ATOM   4434 HD12 LEU B 414     -18.852  54.367  53.936  1.00  0.00           H
    ATOM   4435 HD13 LEU B 414     -17.481  54.394  52.850  1.00  0.00           H
    ATOM   4436 HD21 LEU B 414     -17.172  55.891  51.234  1.00  0.00           H
    ATOM   4437 HD22 LEU B 414     -18.792  56.056  50.584  1.00  0.00           H
    ATOM   4438 HD23 LEU B 414     -17.928  57.470  51.162  1.00  0.00           H
    ATOM   4439  N   ASP B 415     -23.314  57.363  53.028  1.00113.58           N
    ATOM   4440  CA  ASP B 415     -24.787  57.297  52.960  1.00117.10           C
    ATOM   4441  C   ASP B 415     -25.369  58.284  51.935  1.00119.08           C
    ATOM   4442  O   ASP B 415     -26.194  57.873  51.122  1.00121.11           O
    ATOM   4443  CB  ASP B 415     -25.519  57.482  54.316  1.00114.31           C
    ATOM   4444  CG  ASP B 415     -25.124  56.512  55.432  1.00110.95           C
    ATOM   4445  OD1 ASP B 415     -24.828  55.343  55.110  1.00107.04           O
    ATOM   4446  OD2 ASP B 415     -25.209  56.929  56.606  1.00109.29           O1-
    ATOM   4447  H   ASP B 415     -22.921  57.672  53.907  1.00  0.00           H
    ATOM   4448  HA  ASP B 415     -25.030  56.295  52.600  1.00  0.00           H
    ATOM   4449  HB2 ASP B 415     -25.329  58.495  54.671  1.00  0.00           H
    ATOM   4450  HB3 ASP B 415     -26.598  57.398  54.173  1.00  0.00           H
    ATOM   4451  N   ARG B 416     -24.893  59.542  51.970  1.00122.19           N
    ATOM   4452  CA  ARG B 416     -25.209  60.617  51.021  1.00124.00           C
    ATOM   4453  C   ARG B 416     -24.874  60.246  49.569  1.00122.85           C
    ATOM   4454  O   ARG B 416     -25.733  60.407  48.703  1.00124.54           O
    ATOM   4455  CB  ARG B 416     -24.446  61.899  51.434  1.00126.37           C
    ATOM   4456  CG  ARG B 416     -25.280  62.869  52.283  1.00132.37           C
    ATOM   4457  CD  ARG B 416     -26.127  63.794  51.395  1.00136.83           C
    ATOM   4458  NE  ARG B 416     -27.069  64.626  52.156  1.00142.44           N
    ATOM   4459  CZ  ARG B 416     -27.581  65.798  51.738  1.00142.50           C
    ATOM   4460  NH1 ARG B 416     -27.204  66.359  50.579  1.00141.97           N
    ATOM   4461  NH2 ARG B 416     -28.492  66.422  52.495  1.00140.56           N1+
    ATOM   4462  H   ARG B 416     -24.222  59.784  52.688  1.00  0.00           H
    ATOM   4463  HA  ARG B 416     -26.285  60.794  51.084  1.00  0.00           H
    ATOM   4464  HB2 ARG B 416     -23.532  61.630  51.960  1.00  0.00           H
    ATOM   4465  HB3 ARG B 416     -24.092  62.447  50.557  1.00  0.00           H
    ATOM   4466  HG2 ARG B 416     -25.836  62.391  53.091  1.00  0.00           H
    ATOM   4467  HG3 ARG B 416     -24.546  63.513  52.769  1.00  0.00           H
    ATOM   4468  HD2 ARG B 416     -25.420  64.517  50.988  1.00  0.00           H
    ATOM   4469  HD3 ARG B 416     -26.575  63.295  50.535  1.00  0.00           H
    ATOM   4470  HE  ARG B 416     -27.414  64.227  53.017  1.00  0.00           H
    ATOM   4471 HH11 ARG B 416     -26.524  65.896  49.994  1.00  0.00           H
    ATOM   4472 HH12 ARG B 416     -27.598  67.238  50.278  1.00  0.00           H
    ATOM   4473 HH21 ARG B 416     -28.774  66.029  53.382  1.00  0.00           H
    ATOM   4474 HH22 ARG B 416     -28.887  67.303  52.201  1.00  0.00           H
    ATOM   4475  N   GLU B 417     -23.650  59.735  49.347  1.00120.38           N
    ATOM   4476  CA  GLU B 417     -23.149  59.262  48.052  1.00114.91           C
    ATOM   4477  C   GLU B 417     -23.943  58.072  47.476  1.00113.82           C
    ATOM   4478  O   GLU B 417     -24.036  57.957  46.255  1.00116.26           O
    ATOM   4479  CB  GLU B 417     -21.650  58.907  48.193  1.00115.53           C
    ATOM   4480  CG  GLU B 417     -20.706  60.117  48.365  1.00112.82           C
    ATOM   4481  CD  GLU B 417     -20.546  60.950  47.096  1.00111.43           C
    ATOM   4482  OE1 GLU B 417     -19.990  60.400  46.121  1.00111.73           O
    ATOM   4483  OE2 GLU B 417     -20.972  62.125  47.129  1.00112.49           O1-
    ATOM   4484  H   GLU B 417     -23.002  59.654  50.119  1.00  0.00           H
    ATOM   4485  HA  GLU B 417     -23.254  60.080  47.337  1.00  0.00           H
    ATOM   4486  HB2 GLU B 417     -21.520  58.246  49.050  1.00  0.00           H
    ATOM   4487  HB3 GLU B 417     -21.322  58.321  47.332  1.00  0.00           H
    ATOM   4488  HG2 GLU B 417     -21.059  60.782  49.149  1.00  0.00           H
    ATOM   4489  HG3 GLU B 417     -19.717  59.773  48.672  1.00  0.00           H
    ATOM   4490  N   ALA B 418     -24.520  57.234  48.355  1.00114.63           N
    ATOM   4491  CA  ALA B 418     -25.366  56.093  48.000  1.00117.21           C
    ATOM   4492  C   ALA B 418     -26.850  56.454  47.786  1.00118.80           C
    ATOM   4493  O   ALA B 418     -27.610  55.581  47.368  1.00122.97           O
    ATOM   4494  CB  ALA B 418     -25.224  55.029  49.101  1.00117.03           C
    ATOM   4495  H   ALA B 418     -24.380  57.388  49.344  1.00  0.00           H
    ATOM   4496  HA  ALA B 418     -25.001  55.658  47.068  1.00  0.00           H
    ATOM   4497  HB1 ALA B 418     -25.800  54.133  48.868  1.00  0.00           H
    ATOM   4498  HB2 ALA B 418     -24.184  54.723  49.218  1.00  0.00           H
    ATOM   4499  HB3 ALA B 418     -25.567  55.402  50.068  1.00  0.00           H
    ATOM   4500  N   GLY B 419     -27.254  57.702  48.091  1.00120.78           N
    ATOM   4501  CA  GLY B 419     -28.645  58.164  48.027  1.00122.60           C
    ATOM   4502  C   GLY B 419     -29.510  57.609  49.177  1.00123.67           C
    ATOM   4503  O   GLY B 419     -30.734  57.716  49.116  1.00123.80           O
    ATOM   4504  H   GLY B 419     -26.569  58.371  48.420  1.00  0.00           H
    ATOM   4505  HA2 GLY B 419     -28.651  59.252  48.079  1.00  0.00           H
    ATOM   4506  HA3 GLY B 419     -29.089  57.888  47.069  1.00  0.00           H
    ATOM   4507  N   ARG B 420     -28.882  57.012  50.203  1.00124.51           N
    ATOM   4508  CA  ARG B 420     -29.513  56.438  51.389  1.00125.27           C
    ATOM   4509  C   ARG B 420     -29.682  57.511  52.472  1.00126.93           C
    ATOM   4510  O   ARG B 420     -28.943  58.498  52.494  1.00127.22           O
    ATOM   4511  CB  ARG B 420     -28.644  55.259  51.896  1.00124.82           C
    ATOM   4512  CG  ARG B 420     -28.477  54.141  50.845  1.00124.56           C
    ATOM   4513  CD  ARG B 420     -29.683  53.190  50.717  1.00125.52           C
    ATOM   4514  NE  ARG B 420     -29.593  52.043  51.637  1.00129.12           N
    ATOM   4515  CZ  ARG B 420     -30.603  51.391  52.240  1.00130.18           C
    ATOM   4516  NH1 ARG B 420     -31.864  51.843  52.198  1.00132.41           N
    ATOM   4517  NH2 ARG B 420     -30.342  50.257  52.903  1.00130.45           N1+
    ATOM   4518  H   ARG B 420     -27.871  57.001  50.206  1.00  0.00           H
    ATOM   4519  HA  ARG B 420     -30.502  56.064  51.123  1.00  0.00           H
    ATOM   4520  HB2 ARG B 420     -27.655  55.642  52.157  1.00  0.00           H
    ATOM   4521  HB3 ARG B 420     -29.053  54.840  52.817  1.00  0.00           H
    ATOM   4522  HG2 ARG B 420     -28.176  54.524  49.871  1.00  0.00           H
    ATOM   4523  HG3 ARG B 420     -27.630  53.542  51.181  1.00  0.00           H
    ATOM   4524  HD2 ARG B 420     -30.563  53.748  51.032  1.00  0.00           H
    ATOM   4525  HD3 ARG B 420     -29.871  52.881  49.689  1.00  0.00           H
    ATOM   4526  HE  ARG B 420     -28.668  51.651  51.734  1.00  0.00           H
    ATOM   4527 HH11 ARG B 420     -32.072  52.715  51.734  1.00  0.00           H
    ATOM   4528 HH12 ARG B 420     -32.593  51.383  52.723  1.00  0.00           H
    ATOM   4529 HH21 ARG B 420     -29.397  49.911  52.970  1.00  0.00           H
    ATOM   4530 HH22 ARG B 420     -31.081  49.774  53.393  1.00  0.00           H
    ATOM   4531  N   ALA B 421     -30.649  57.277  53.373  1.00128.69           N
    ATOM   4532  CA  ALA B 421     -30.879  58.108  54.554  1.00129.36           C
    ATOM   4533  C   ALA B 421     -29.717  57.978  55.568  1.00130.22           C
    ATOM   4534  O   ALA B 421     -29.072  56.927  55.584  1.00131.62           O
    ATOM   4535  CB  ALA B 421     -32.217  57.694  55.186  1.00127.50           C
    ATOM   4536  H   ALA B 421     -31.213  56.442  53.300  1.00  0.00           H
    ATOM   4537  HA  ALA B 421     -30.974  59.129  54.189  1.00  0.00           H
    ATOM   4538  HB1 ALA B 421     -32.438  58.286  56.075  1.00  0.00           H
    ATOM   4539  HB2 ALA B 421     -33.040  57.840  54.486  1.00  0.00           H
    ATOM   4540  HB3 ALA B 421     -32.220  56.645  55.479  1.00  0.00           H
    ATOM   4541  N   PRO B 422     -29.464  59.024  56.394  1.00129.89           N
    ATOM   4542  CA  PRO B 422     -28.441  58.988  57.462  1.00129.35           C
    ATOM   4543  C   PRO B 422     -28.544  57.780  58.415  1.00130.39           C
    ATOM   4544  O   PRO B 422     -29.528  57.657  59.144  1.00130.46           O
    ATOM   4545  CB  PRO B 422     -28.637  60.326  58.201  1.00128.68           C
    ATOM   4546  CG  PRO B 422     -29.211  61.257  57.152  1.00126.85           C
    ATOM   4547  CD  PRO B 422     -30.112  60.339  56.340  1.00128.07           C
    ATOM   4548  HA  PRO B 422     -27.468  58.992  56.967  1.00  0.00           H
    ATOM   4549  HB2 PRO B 422     -29.354  60.229  59.018  1.00  0.00           H
    ATOM   4550  HB3 PRO B 422     -27.711  60.716  58.625  1.00  0.00           H
    ATOM   4551  HG2 PRO B 422     -29.733  62.118  57.571  1.00  0.00           H
    ATOM   4552  HG3 PRO B 422     -28.403  61.627  56.519  1.00  0.00           H
    ATOM   4553  HD2 PRO B 422     -31.100  60.266  56.795  1.00  0.00           H
    ATOM   4554  HD3 PRO B 422     -30.219  60.740  55.333  1.00  0.00           H
    ATOM   4555  N   GLY B 423     -27.526  56.905  58.364  1.00130.61           N
    ATOM   4556  CA  GLY B 423     -27.410  55.714  59.205  1.00130.68           C
    ATOM   4557  C   GLY B 423     -28.216  54.513  58.679  1.00130.01           C
    ATOM   4558  O   GLY B 423     -28.245  53.487  59.358  1.00132.22           O
    ATOM   4559  H   GLY B 423     -26.766  57.062  57.710  1.00  0.00           H
    ATOM   4560  HA2 GLY B 423     -26.358  55.432  59.249  1.00  0.00           H
    ATOM   4561  HA3 GLY B 423     -27.718  55.939  60.227  1.00  0.00           H
    ATOM   4562  N   ASP B 424     -28.849  54.618  57.495  1.00128.04           N
    ATOM   4563  CA  ASP B 424     -29.661  53.572  56.862  1.00124.89           C
    ATOM   4564  C   ASP B 424     -28.824  52.786  55.822  1.00123.26           C
    ATOM   4565  O   ASP B 424     -29.326  52.465  54.747  1.00122.60           O
    ATOM   4566  CB  ASP B 424     -30.954  54.185  56.257  1.00123.95           C
    ATOM   4567  CG  ASP B 424     -32.030  53.173  55.831  1.00126.31           C
    ATOM   4568  OD1 ASP B 424     -32.186  52.151  56.534  1.00127.46           O
    ATOM   4569  OD2 ASP B 424     -32.721  53.474  54.833  1.00128.35           O1-
    ATOM   4570  H   ASP B 424     -28.775  55.484  56.976  1.00  0.00           H
    ATOM   4571  HA  ASP B 424     -29.958  52.851  57.624  1.00  0.00           H
    ATOM   4572  HB2 ASP B 424     -31.406  54.879  56.966  1.00  0.00           H
    ATOM   4573  HB3 ASP B 424     -30.677  54.768  55.378  1.00  0.00           H
    ATOM   4574  N   ALA B 425     -27.559  52.478  56.153  1.00118.30           N
    ATOM   4575  CA  ALA B 425     -26.660  51.666  55.332  1.00110.68           C
    ATOM   4576  C   ALA B 425     -25.365  51.353  56.091  1.00109.56           C
    ATOM   4577  O   ALA B 425     -24.891  52.178  56.874  1.00104.15           O
    ATOM   4578  CB  ALA B 425     -26.347  52.352  53.985  1.00111.12           C
    ATOM   4579  H   ALA B 425     -27.187  52.785  57.041  1.00  0.00           H
    ATOM   4580  HA  ALA B 425     -27.162  50.717  55.131  1.00  0.00           H
    ATOM   4581  HB1 ALA B 425     -25.518  51.866  53.469  1.00  0.00           H
    ATOM   4582  HB2 ALA B 425     -27.200  52.317  53.309  1.00  0.00           H
    ATOM   4583  HB3 ALA B 425     -26.081  53.398  54.119  1.00  0.00           H
    ATOM   4584  N   VAL B 426     -24.811  50.165  55.802  1.00106.50           N
    ATOM   4585  CA  VAL B 426     -23.508  49.704  56.275  1.00103.98           C
    ATOM   4586  C   VAL B 426     -22.664  49.390  55.026  1.00103.25           C
    ATOM   4587  O   VAL B 426     -23.116  48.638  54.161  1.00104.57           O
    ATOM   4588  CB  VAL B 426     -23.629  48.419  57.147  1.00101.92           C
    ATOM   4589  CG1 VAL B 426     -22.266  47.879  57.623  1.00103.56           C
    ATOM   4590  CG2 VAL B 426     -24.538  48.649  58.369  1.00 98.95           C
    ATOM   4591  H   VAL B 426     -25.264  49.558  55.135  1.00  0.00           H
    ATOM   4592  HA  VAL B 426     -23.007  50.483  56.855  1.00  0.00           H
    ATOM   4593  HB  VAL B 426     -24.095  47.632  56.551  1.00  0.00           H
    ATOM   4594 HG11 VAL B 426     -22.386  46.993  58.247  1.00  0.00           H
    ATOM   4595 HG12 VAL B 426     -21.625  47.598  56.790  1.00  0.00           H
    ATOM   4596 HG13 VAL B 426     -21.731  48.625  58.211  1.00  0.00           H
    ATOM   4597 HG21 VAL B 426     -24.560  47.775  59.019  1.00  0.00           H
    ATOM   4598 HG22 VAL B 426     -24.187  49.492  58.966  1.00  0.00           H
    ATOM   4599 HG23 VAL B 426     -25.566  48.855  58.073  1.00  0.00           H
    ATOM   4600  N   HIS B 427     -21.471  50.001  54.947  1.00101.53           N
    ATOM   4601  CA  HIS B 427     -20.616  50.010  53.760  1.00 97.30           C
    ATOM   4602  C   HIS B 427     -19.439  49.047  53.948  1.00 95.76           C
    ATOM   4603  O   HIS B 427     -18.495  49.370  54.668  1.00 97.61           O
    ATOM   4604  CB  HIS B 427     -20.128  51.453  53.494  1.00 99.75           C
    ATOM   4605  CG  HIS B 427     -21.243  52.461  53.380  1.00103.85           C
    ATOM   4606  CD2 HIS B 427     -21.856  53.178  54.387  1.00104.75           C
    ATOM   4607  ND1 HIS B 427     -21.856  52.772  52.163  1.00107.52           N
    ATOM   4608  CE1 HIS B 427     -22.817  53.630  52.478  1.00110.01           C
    ATOM   4609  NE2 HIS B 427     -22.860  53.905  53.781  1.00108.12           N
    ATOM   4610  H   HIS B 427     -21.153  50.577  55.716  1.00  0.00           H
    ATOM   4611  HA  HIS B 427     -21.190  49.697  52.885  1.00  0.00           H
    ATOM   4612  HB2 HIS B 427     -19.459  51.790  54.286  1.00  0.00           H
    ATOM   4613  HB3 HIS B 427     -19.543  51.483  52.573  1.00  0.00           H
    ATOM   4614  HD2 HIS B 427     -21.684  53.207  55.453  1.00  0.00           H
    ATOM   4615  HE2 HIS B 427     -23.531  54.514  54.248  1.00  0.00           H
    ATOM   4616  HE1 HIS B 427     -23.495  54.059  51.756  1.00  0.00           H
    ATOM   4617  N   LYS B 428     -19.513  47.885  53.281  1.00 92.52           N
    ATOM   4618  CA  LYS B 428     -18.451  46.882  53.258  1.00 91.83           C
    ATOM   4619  C   LYS B 428     -17.332  47.309  52.290  1.00 92.42           C
    ATOM   4620  O   LYS B 428     -17.548  47.307  51.077  1.00 91.57           O
    ATOM   4621  CB  LYS B 428     -19.066  45.518  52.878  1.00 88.66           C
    ATOM   4622  CG  LYS B 428     -18.074  44.341  52.904  1.00 88.76           C
    ATOM   4623  CD  LYS B 428     -18.737  43.015  52.503  1.00 86.55           C
    ATOM   4624  CE  LYS B 428     -17.774  41.825  52.581  1.00 85.69           C
    ATOM   4625  NZ  LYS B 428     -18.440  40.570  52.204  1.00 81.64           N1+
    ATOM   4626  H   LYS B 428     -20.324  47.688  52.712  1.00  0.00           H
    ATOM   4627  HA  LYS B 428     -18.037  46.786  54.264  1.00  0.00           H
    ATOM   4628  HB2 LYS B 428     -19.881  45.297  53.568  1.00  0.00           H
    ATOM   4629  HB3 LYS B 428     -19.522  45.584  51.889  1.00  0.00           H
    ATOM   4630  HG2 LYS B 428     -17.235  44.532  52.233  1.00  0.00           H
    ATOM   4631  HG3 LYS B 428     -17.653  44.252  53.906  1.00  0.00           H
    ATOM   4632  HD2 LYS B 428     -19.597  42.829  53.148  1.00  0.00           H
    ATOM   4633  HD3 LYS B 428     -19.129  43.101  51.488  1.00  0.00           H
    ATOM   4634  HE2 LYS B 428     -16.914  41.982  51.930  1.00  0.00           H
    ATOM   4635  HE3 LYS B 428     -17.403  41.712  53.599  1.00  0.00           H
    ATOM   4636  HZ1 LYS B 428     -19.201  40.391  52.855  1.00  0.00           H
    ATOM   4637  HZ2 LYS B 428     -18.802  40.619  51.264  1.00  0.00           H
    ATOM   4638  HZ3 LYS B 428     -17.792  39.797  52.290  1.00  0.00           H
    ATOM   4639  N   ILE B 429     -16.159  47.647  52.847  1.00 91.77           N
    ATOM   4640  CA  ILE B 429     -14.934  47.922  52.098  1.00 89.99           C
    ATOM   4641  C   ILE B 429     -14.150  46.609  51.934  1.00 91.52           C
    ATOM   4642  O   ILE B 429     -13.931  45.891  52.913  1.00 93.88           O
    ATOM   4643  CB  ILE B 429     -14.039  48.984  52.811  1.00 88.60           C
    ATOM   4644  CG1 ILE B 429     -14.770  50.346  52.865  1.00 87.70           C
    ATOM   4645  CG2 ILE B 429     -12.638  49.161  52.175  1.00 86.11           C
    ATOM   4646  CD1 ILE B 429     -13.983  51.466  53.561  1.00 92.05           C
    ATOM   4647  H   ILE B 429     -16.059  47.609  53.854  1.00  0.00           H
    ATOM   4648  HA  ILE B 429     -15.191  48.308  51.109  1.00  0.00           H
    ATOM   4649  HB  ILE B 429     -13.893  48.655  53.837  1.00  0.00           H
    ATOM   4650 HG12 ILE B 429     -15.001  50.669  51.849  1.00  0.00           H
    ATOM   4651 HG13 ILE B 429     -15.731  50.237  53.368  1.00  0.00           H
    ATOM   4652 HG21 ILE B 429     -12.022  49.862  52.737  1.00  0.00           H
    ATOM   4653 HG22 ILE B 429     -12.067  48.236  52.138  1.00  0.00           H
    ATOM   4654 HG23 ILE B 429     -12.723  49.543  51.158  1.00  0.00           H
    ATOM   4655 HD11 ILE B 429     -14.647  52.282  53.844  1.00  0.00           H
    ATOM   4656 HD12 ILE B 429     -13.485  51.102  54.460  1.00  0.00           H
    ATOM   4657 HD13 ILE B 429     -13.224  51.888  52.902  1.00  0.00           H
    ATOM   4658  N   TYR B 430     -13.742  46.337  50.686  1.00 93.06           N
    ATOM   4659  CA  TYR B 430     -12.995  45.143  50.301  1.00 90.62           C
    ATOM   4660  C   TYR B 430     -11.478  45.335  50.506  1.00 86.82           C
    ATOM   4661  O   TYR B 430     -11.006  46.464  50.375  1.00 87.31           O
    ATOM   4662  CB  TYR B 430     -13.325  44.827  48.829  1.00 94.92           C
    ATOM   4663  CG  TYR B 430     -14.756  44.356  48.653  1.00 99.84           C
    ATOM   4664  CD1 TYR B 430     -15.101  43.038  49.016  1.00101.43           C
    ATOM   4665  CD2 TYR B 430     -15.754  45.240  48.193  1.00101.28           C
    ATOM   4666  CE1 TYR B 430     -16.438  42.607  48.932  1.00103.78           C
    ATOM   4667  CE2 TYR B 430     -17.093  44.810  48.112  1.00102.79           C
    ATOM   4668  CZ  TYR B 430     -17.436  43.496  48.486  1.00104.11           C
    ATOM   4669  OH  TYR B 430     -18.735  43.085  48.417  1.00106.55           O
    ATOM   4670  H   TYR B 430     -13.957  46.984  49.942  1.00  0.00           H
    ATOM   4671  HA  TYR B 430     -13.373  44.328  50.916  1.00  0.00           H
    ATOM   4672  HB2 TYR B 430     -13.133  45.693  48.193  1.00  0.00           H
    ATOM   4673  HB3 TYR B 430     -12.672  44.034  48.460  1.00  0.00           H
    ATOM   4674  HD1 TYR B 430     -14.343  42.358  49.376  1.00  0.00           H
    ATOM   4675  HD2 TYR B 430     -15.502  46.256  47.924  1.00  0.00           H
    ATOM   4676  HE1 TYR B 430     -16.694  41.598  49.219  1.00  0.00           H
    ATOM   4677  HE2 TYR B 430     -17.857  45.494  47.772  1.00  0.00           H
    ATOM   4678  HH  TYR B 430     -18.855  42.158  48.630  1.00  0.00           H
    ATOM   4679  N   PRO B 431     -10.737  44.232  50.784  1.00 81.57           N
    ATOM   4680  CA  PRO B 431      -9.257  44.203  50.793  1.00 76.27           C
    ATOM   4681  C   PRO B 431      -8.573  44.912  49.608  1.00 77.71           C
    ATOM   4682  O   PRO B 431      -8.949  44.675  48.460  1.00 78.11           O
    ATOM   4683  CB  PRO B 431      -8.936  42.700  50.805  1.00 72.65           C
    ATOM   4684  CG  PRO B 431     -10.090  42.087  51.573  1.00 71.44           C
    ATOM   4685  CD  PRO B 431     -11.280  42.906  51.099  1.00 75.41           C
    ATOM   4686  HA  PRO B 431      -8.930  44.647  51.734  1.00  0.00           H
    ATOM   4687  HB2 PRO B 431      -8.926  42.294  49.792  1.00  0.00           H
    ATOM   4688  HB3 PRO B 431      -7.970  42.476  51.255  1.00  0.00           H
    ATOM   4689  HG2 PRO B 431     -10.206  41.016  51.408  1.00  0.00           H
    ATOM   4690  HG3 PRO B 431      -9.939  42.248  52.641  1.00  0.00           H
    ATOM   4691  HD2 PRO B 431     -11.708  42.470  50.195  1.00  0.00           H
    ATOM   4692  HD3 PRO B 431     -12.045  42.926  51.876  1.00  0.00           H
    ATOM   4693  N   SER B 432      -7.607  45.783  49.944  1.00 81.20           N
    ATOM   4694  CA  SER B 432      -6.793  46.660  49.090  1.00 85.06           C
    ATOM   4695  C   SER B 432      -7.465  47.986  48.681  1.00 82.86           C
    ATOM   4696  O   SER B 432      -6.750  48.886  48.238  1.00 82.25           O
    ATOM   4697  CB  SER B 432      -6.166  45.901  47.892  1.00 90.40           C
    ATOM   4698  OG  SER B 432      -7.018  45.825  46.764  1.00 98.02           O
    ATOM   4699  H   SER B 432      -7.413  45.898  50.932  1.00  0.00           H
    ATOM   4700  HA  SER B 432      -5.964  46.946  49.739  1.00  0.00           H
    ATOM   4701  HB2 SER B 432      -5.260  46.417  47.573  1.00  0.00           H
    ATOM   4702  HB3 SER B 432      -5.857  44.896  48.183  1.00  0.00           H
    ATOM   4703  HG  SER B 432      -7.787  45.301  47.011  1.00  0.00           H
    ATOM   4704  N   ALA B 433      -8.797  48.098  48.824  1.00 79.70           N
    ATOM   4705  CA  ALA B 433      -9.578  49.248  48.369  1.00 80.77           C
    ATOM   4706  C   ALA B 433      -9.316  50.504  49.217  1.00 82.18           C
    ATOM   4707  O   ALA B 433      -9.207  50.393  50.438  1.00 83.67           O
    ATOM   4708  CB  ALA B 433     -11.063  48.860  48.367  1.00 78.61           C
    ATOM   4709  H   ALA B 433      -9.323  47.330  49.221  1.00  0.00           H
    ATOM   4710  HA  ALA B 433      -9.285  49.450  47.337  1.00  0.00           H
    ATOM   4711  HB1 ALA B 433     -11.637  49.535  47.736  1.00  0.00           H
    ATOM   4712  HB2 ALA B 433     -11.209  47.858  47.963  1.00  0.00           H
    ATOM   4713  HB3 ALA B 433     -11.490  48.879  49.369  1.00  0.00           H
    ATOM   4714  N   TYR B 434      -9.223  51.660  48.537  1.00 81.68           N
    ATOM   4715  CA  TYR B 434      -8.868  52.963  49.099  1.00 83.52           C
    ATOM   4716  C   TYR B 434      -9.984  53.975  48.789  1.00 86.43           C
    ATOM   4717  O   TYR B 434     -10.005  54.538  47.694  1.00 85.15           O
    ATOM   4718  CB  TYR B 434      -7.501  53.376  48.512  1.00 82.59           C
    ATOM   4719  CG  TYR B 434      -6.945  54.728  48.936  1.00 82.66           C
    ATOM   4720  CD1 TYR B 434      -6.817  55.057  50.302  1.00 82.70           C
    ATOM   4721  CD2 TYR B 434      -6.506  55.646  47.959  1.00 83.75           C
    ATOM   4722  CE1 TYR B 434      -6.192  56.257  50.692  1.00 84.78           C
    ATOM   4723  CE2 TYR B 434      -5.905  56.858  48.347  1.00 83.39           C
    ATOM   4724  CZ  TYR B 434      -5.725  57.153  49.711  1.00 85.55           C
    ATOM   4725  OH  TYR B 434      -5.086  58.302  50.069  1.00 86.08           O
    ATOM   4726  H   TYR B 434      -9.356  51.647  47.536  1.00  0.00           H
    ATOM   4727  HA  TYR B 434      -8.758  52.882  50.178  1.00  0.00           H
    ATOM   4728  HB2 TYR B 434      -6.762  52.622  48.780  1.00  0.00           H
    ATOM   4729  HB3 TYR B 434      -7.556  53.352  47.423  1.00  0.00           H
    ATOM   4730  HD1 TYR B 434      -7.193  54.385  51.056  1.00  0.00           H
    ATOM   4731  HD2 TYR B 434      -6.609  55.418  46.909  1.00  0.00           H
    ATOM   4732  HE1 TYR B 434      -6.074  56.487  51.742  1.00  0.00           H
    ATOM   4733  HE2 TYR B 434      -5.556  57.550  47.593  1.00  0.00           H
    ATOM   4734  HH  TYR B 434      -4.786  58.315  50.986  1.00  0.00           H
    ATOM   4735  N   ILE B 435     -10.909  54.145  49.746  1.00 90.16           N
    ATOM   4736  CA  ILE B 435     -12.144  54.923  49.604  1.00 90.16           C
    ATOM   4737  C   ILE B 435     -12.123  56.142  50.543  1.00 91.84           C
    ATOM   4738  O   ILE B 435     -11.675  56.025  51.683  1.00 95.96           O
    ATOM   4739  CB  ILE B 435     -13.395  54.035  49.937  1.00 88.56           C
    ATOM   4740  CG1 ILE B 435     -13.676  53.006  48.818  1.00 90.31           C
    ATOM   4741  CG2 ILE B 435     -14.704  54.808  50.210  1.00 86.35           C
    ATOM   4742  CD1 ILE B 435     -12.755  51.783  48.824  1.00 92.88           C
    ATOM   4743  H   ILE B 435     -10.789  53.679  50.636  1.00  0.00           H
    ATOM   4744  HA  ILE B 435     -12.241  55.291  48.581  1.00  0.00           H
    ATOM   4745  HB  ILE B 435     -13.182  53.478  50.852  1.00  0.00           H
    ATOM   4746 HG12 ILE B 435     -14.694  52.626  48.911  1.00  0.00           H
    ATOM   4747 HG13 ILE B 435     -13.641  53.496  47.844  1.00  0.00           H
    ATOM   4748 HG21 ILE B 435     -15.552  54.132  50.324  1.00  0.00           H
    ATOM   4749 HG22 ILE B 435     -14.652  55.362  51.146  1.00  0.00           H
    ATOM   4750 HG23 ILE B 435     -14.942  55.502  49.404  1.00  0.00           H
    ATOM   4751 HD11 ILE B 435     -13.333  50.870  48.681  1.00  0.00           H
    ATOM   4752 HD12 ILE B 435     -12.030  51.847  48.013  1.00  0.00           H
    ATOM   4753 HD13 ILE B 435     -12.208  51.675  49.761  1.00  0.00           H
    ATOM   4754  N   LYS B 436     -12.668  57.273  50.058  1.00 91.82           N
    ATOM   4755  CA  LYS B 436     -12.965  58.460  50.860  1.00 92.56           C
    ATOM   4756  C   LYS B 436     -14.219  58.210  51.705  1.00 91.55           C
    ATOM   4757  O   LYS B 436     -15.260  57.881  51.137  1.00 91.29           O
    ATOM   4758  CB  LYS B 436     -13.188  59.668  49.934  1.00 93.33           C
    ATOM   4759  CG  LYS B 436     -13.358  60.994  50.696  1.00 93.30           C
    ATOM   4760  CD  LYS B 436     -13.490  62.199  49.759  1.00 93.43           C
    ATOM   4761  CE  LYS B 436     -13.944  63.451  50.515  1.00 95.42           C
    ATOM   4762  NZ  LYS B 436     -14.140  64.592  49.607  1.00 97.61           N1+
    ATOM   4763  H   LYS B 436     -13.017  57.287  49.111  1.00  0.00           H
    ATOM   4764  HA  LYS B 436     -12.116  58.676  51.509  1.00  0.00           H
    ATOM   4765  HB2 LYS B 436     -12.347  59.755  49.255  1.00  0.00           H
    ATOM   4766  HB3 LYS B 436     -14.056  59.499  49.295  1.00  0.00           H
    ATOM   4767  HG2 LYS B 436     -14.243  60.946  51.331  1.00  0.00           H
    ATOM   4768  HG3 LYS B 436     -12.513  61.147  51.369  1.00  0.00           H
    ATOM   4769  HD2 LYS B 436     -12.529  62.389  49.279  1.00  0.00           H
    ATOM   4770  HD3 LYS B 436     -14.193  61.968  48.958  1.00  0.00           H
    ATOM   4771  HE2 LYS B 436     -14.884  63.244  51.023  1.00  0.00           H
    ATOM   4772  HE3 LYS B 436     -13.217  63.721  51.280  1.00  0.00           H
    ATOM   4773  HZ1 LYS B 436     -14.797  64.344  48.882  1.00  0.00           H
    ATOM   4774  HZ2 LYS B 436     -14.512  65.383  50.126  1.00  0.00           H
    ATOM   4775  HZ3 LYS B 436     -13.258  64.859  49.196  1.00  0.00           H
    ATOM   4776  N   VAL B 437     -14.103  58.394  53.025  1.00 88.05           N
    ATOM   4777  CA  VAL B 437     -15.187  58.129  53.969  1.00 81.23           C
    ATOM   4778  C   VAL B 437     -15.665  59.392  54.701  1.00 79.44           C
    ATOM   4779  O   VAL B 437     -16.552  59.267  55.538  1.00 75.82           O
    ATOM   4780  CB  VAL B 437     -14.766  57.079  55.032  1.00 81.35           C
    ATOM   4781  CG1 VAL B 437     -14.382  55.744  54.372  1.00 79.79           C
    ATOM   4782  CG2 VAL B 437     -13.674  57.563  56.008  1.00 79.86           C
    ATOM   4783  H   VAL B 437     -13.225  58.704  53.422  1.00  0.00           H
    ATOM   4784  HA  VAL B 437     -16.065  57.740  53.452  1.00  0.00           H
    ATOM   4785  HB  VAL B 437     -15.649  56.855  55.634  1.00  0.00           H
    ATOM   4786 HG11 VAL B 437     -14.269  54.954  55.112  1.00  0.00           H
    ATOM   4787 HG12 VAL B 437     -15.143  55.416  53.665  1.00  0.00           H
    ATOM   4788 HG13 VAL B 437     -13.441  55.823  53.827  1.00  0.00           H
    ATOM   4789 HG21 VAL B 437     -13.496  56.816  56.780  1.00  0.00           H
    ATOM   4790 HG22 VAL B 437     -12.734  57.749  55.490  1.00  0.00           H
    ATOM   4791 HG23 VAL B 437     -13.944  58.480  56.530  1.00  0.00           H
    ATOM   4792  N   PHE B 438     -15.099  60.577  54.425  1.00 78.90           N
    ATOM   4793  CA  PHE B 438     -15.512  61.792  55.118  1.00 79.23           C
    ATOM   4794  C   PHE B 438     -15.056  63.035  54.354  1.00 79.32           C
    ATOM   4795  O   PHE B 438     -13.913  63.089  53.900  1.00 74.79           O
    ATOM   4796  CB  PHE B 438     -15.004  61.762  56.577  1.00 74.95           C
    ATOM   4797  CG  PHE B 438     -15.446  62.918  57.441  1.00 73.20           C
    ATOM   4798  CD1 PHE B 438     -16.798  63.057  57.806  1.00 73.20           C
    ATOM   4799  CD2 PHE B 438     -14.521  63.907  57.814  1.00 72.09           C
    ATOM   4800  CE1 PHE B 438     -17.190  64.154  58.556  1.00 74.33           C
    ATOM   4801  CE2 PHE B 438     -14.929  64.988  58.579  1.00 72.12           C
    ATOM   4802  CZ  PHE B 438     -16.260  65.108  58.949  1.00 72.84           C
    ATOM   4803  H   PHE B 438     -14.353  60.658  53.749  1.00  0.00           H
    ATOM   4804  HA  PHE B 438     -16.604  61.810  55.139  1.00  0.00           H
    ATOM   4805  HB2 PHE B 438     -15.340  60.861  57.088  1.00  0.00           H
    ATOM   4806  HB3 PHE B 438     -13.918  61.717  56.576  1.00  0.00           H
    ATOM   4807  HD1 PHE B 438     -17.528  62.321  57.501  1.00  0.00           H
    ATOM   4808  HD2 PHE B 438     -13.499  63.808  57.498  1.00  0.00           H
    ATOM   4809  HE1 PHE B 438     -18.227  64.267  58.819  1.00  0.00           H
    ATOM   4810  HE2 PHE B 438     -14.212  65.736  58.878  1.00  0.00           H
    ATOM   4811  HZ  PHE B 438     -16.575  65.957  59.532  1.00  0.00           H
    ATOM   4812  N   ASP B 439     -15.968  64.013  54.272  1.00 79.78           N
    ATOM   4813  CA  ASP B 439     -15.752  65.327  53.686  1.00 81.95           C
    ATOM   4814  C   ASP B 439     -16.091  66.354  54.774  1.00 81.97           C
    ATOM   4815  O   ASP B 439     -17.228  66.379  55.251  1.00 78.54           O
    ATOM   4816  CB  ASP B 439     -16.599  65.500  52.406  1.00 86.34           C
    ATOM   4817  CG  ASP B 439     -16.352  66.802  51.646  1.00 88.54           C
    ATOM   4818  OD1 ASP B 439     -16.799  67.849  52.155  1.00 91.27           O
    ATOM   4819  OD2 ASP B 439     -15.778  66.722  50.539  1.00 86.45           O1-
    ATOM   4820  H   ASP B 439     -16.873  63.880  54.700  1.00  0.00           H
    ATOM   4821  HA  ASP B 439     -14.704  65.442  53.413  1.00  0.00           H
    ATOM   4822  HB2 ASP B 439     -16.401  64.676  51.726  1.00  0.00           H
    ATOM   4823  HB3 ASP B 439     -17.664  65.414  52.615  1.00  0.00           H
    ATOM   4824  N   LEU B 440     -15.095  67.177  55.139  1.00 83.25           N
    ATOM   4825  CA  LEU B 440     -15.205  68.213  56.166  1.00 83.59           C
    ATOM   4826  C   LEU B 440     -16.205  69.323  55.804  1.00 86.79           C
    ATOM   4827  O   LEU B 440     -16.918  69.789  56.692  1.00 87.79           O
    ATOM   4828  CB  LEU B 440     -13.808  68.812  56.430  1.00 81.43           C
    ATOM   4829  CG  LEU B 440     -13.736  69.782  57.632  1.00 82.83           C
    ATOM   4830  CD1 LEU B 440     -13.998  69.059  58.969  1.00 84.65           C
    ATOM   4831  CD2 LEU B 440     -12.412  70.560  57.629  1.00 78.87           C
    ATOM   4832  H   LEU B 440     -14.192  67.105  54.681  1.00  0.00           H
    ATOM   4833  HA  LEU B 440     -15.559  67.724  57.075  1.00  0.00           H
    ATOM   4834  HB2 LEU B 440     -13.090  68.007  56.582  1.00  0.00           H
    ATOM   4835  HB3 LEU B 440     -13.480  69.326  55.524  1.00  0.00           H
    ATOM   4836  HG  LEU B 440     -14.506  70.543  57.516  1.00  0.00           H
    ATOM   4837 HD11 LEU B 440     -13.304  69.360  59.752  1.00  0.00           H
    ATOM   4838 HD12 LEU B 440     -15.004  69.272  59.331  1.00  0.00           H
    ATOM   4839 HD13 LEU B 440     -13.914  67.978  58.872  1.00  0.00           H
    ATOM   4840 HD21 LEU B 440     -12.577  71.615  57.852  1.00  0.00           H
    ATOM   4841 HD22 LEU B 440     -11.711  70.170  58.361  1.00  0.00           H
    ATOM   4842 HD23 LEU B 440     -11.912  70.513  56.662  1.00  0.00           H
    ATOM   4843  N   ARG B 441     -16.236  69.722  54.520  1.00 90.48           N
    ATOM   4844  CA  ARG B 441     -17.128  70.756  53.999  1.00 94.17           C
    ATOM   4845  C   ARG B 441     -18.601  70.342  54.081  1.00 94.67           C
    ATOM   4846  O   ARG B 441     -19.390  71.122  54.604  1.00 92.95           O
    ATOM   4847  CB  ARG B 441     -16.730  71.135  52.561  1.00 96.87           C
    ATOM   4848  CG  ARG B 441     -17.534  72.308  51.967  1.00100.46           C
    ATOM   4849  CD  ARG B 441     -17.151  72.601  50.511  1.00102.28           C
    ATOM   4850  NE  ARG B 441     -15.747  73.015  50.391  1.00107.06           N
    ATOM   4851  CZ  ARG B 441     -15.228  74.198  50.757  1.00108.77           C
    ATOM   4852  NH1 ARG B 441     -16.001  75.185  51.231  1.00110.75           N
    ATOM   4853  NH2 ARG B 441     -13.909  74.393  50.642  1.00108.94           N1+
    ATOM   4854  H   ARG B 441     -15.649  69.256  53.840  1.00  0.00           H
    ATOM   4855  HA  ARG B 441     -16.997  71.638  54.628  1.00  0.00           H
    ATOM   4856  HB2 ARG B 441     -15.665  71.367  52.538  1.00  0.00           H
    ATOM   4857  HB3 ARG B 441     -16.857  70.277  51.905  1.00  0.00           H
    ATOM   4858  HG2 ARG B 441     -18.611  72.153  52.020  1.00  0.00           H
    ATOM   4859  HG3 ARG B 441     -17.334  73.187  52.578  1.00  0.00           H
    ATOM   4860  HD2 ARG B 441     -17.226  71.670  49.946  1.00  0.00           H
    ATOM   4861  HD3 ARG B 441     -17.838  73.301  50.034  1.00  0.00           H
    ATOM   4862  HE  ARG B 441     -15.115  72.299  50.060  1.00  0.00           H
    ATOM   4863 HH11 ARG B 441     -16.995  75.040  51.307  1.00  0.00           H
    ATOM   4864 HH12 ARG B 441     -15.602  76.076  51.486  1.00  0.00           H
    ATOM   4865 HH21 ARG B 441     -13.315  73.658  50.286  1.00  0.00           H
    ATOM   4866 HH22 ARG B 441     -13.495  75.278  50.896  1.00  0.00           H
    ATOM   4867  N   GLN B 442     -18.928  69.127  53.601  1.00 93.82           N
    ATOM   4868  CA  GLN B 442     -20.262  68.525  53.700  1.00 96.19           C
    ATOM   4869  C   GLN B 442     -20.742  68.353  55.145  1.00 96.98           C
    ATOM   4870  O   GLN B 442     -21.926  68.561  55.404  1.00 94.97           O
    ATOM   4871  CB  GLN B 442     -20.316  67.165  52.979  1.00 99.76           C
    ATOM   4872  CG  GLN B 442     -20.318  67.284  51.444  1.00106.09           C
    ATOM   4873  CD  GLN B 442     -20.531  65.956  50.710  1.00108.83           C
    ATOM   4874  NE2 GLN B 442     -21.344  65.044  51.252  1.00109.72           N
    ATOM   4875  OE1 GLN B 442     -19.978  65.762  49.632  1.00110.42           O
    ATOM   4876  H   GLN B 442     -18.208  68.560  53.163  1.00  0.00           H
    ATOM   4877  HA  GLN B 442     -20.961  69.205  53.210  1.00  0.00           H
    ATOM   4878  HB2 GLN B 442     -19.488  66.538  53.308  1.00  0.00           H
    ATOM   4879  HB3 GLN B 442     -21.225  66.648  53.293  1.00  0.00           H
    ATOM   4880  HG2 GLN B 442     -21.110  67.963  51.128  1.00  0.00           H
    ATOM   4881  HG3 GLN B 442     -19.381  67.727  51.107  1.00  0.00           H
    ATOM   4882 HE21 GLN B 442     -21.800  65.219  52.135  1.00  0.00           H
    ATOM   4883 HE22 GLN B 442     -21.503  64.170  50.772  1.00  0.00           H
    ATOM   4884  N   CYS B 443     -19.810  68.010  56.048  1.00 99.22           N
    ATOM   4885  CA  CYS B 443     -20.045  67.905  57.481  1.00100.77           C
    ATOM   4886  C   CYS B 443     -20.407  69.267  58.092  1.00100.55           C
    ATOM   4887  O   CYS B 443     -21.462  69.367  58.711  1.00100.42           O
    ATOM   4888  CB  CYS B 443     -18.856  67.239  58.190  1.00102.35           C
    ATOM   4889  SG  CYS B 443     -19.197  66.944  59.944  1.00 99.75           S
    ATOM   4890  H   CYS B 443     -18.861  67.844  55.739  1.00  0.00           H
    ATOM   4891  HA  CYS B 443     -20.908  67.249  57.611  1.00  0.00           H
    ATOM   4892  HB2 CYS B 443     -18.649  66.284  57.711  1.00  0.00           H
    ATOM   4893  HB3 CYS B 443     -17.951  67.838  58.104  1.00  0.00           H
    ATOM   4894  HG  CYS B 443     -20.038  65.922  59.757  1.00  0.00           H
    ATOM   4895  N   HIS B 444     -19.562  70.290  57.860  1.00 99.13           N
    ATOM   4896  CA  HIS B 444     -19.780  71.675  58.287  1.00 99.16           C
    ATOM   4897  C   HIS B 444     -21.079  72.284  57.737  1.00100.46           C
    ATOM   4898  O   HIS B 444     -21.780  72.963  58.486  1.00 99.81           O
    ATOM   4899  CB  HIS B 444     -18.558  72.538  57.917  1.00 98.72           C
    ATOM   4900  CG  HIS B 444     -18.716  74.009  58.228  1.00 99.10           C
    ATOM   4901  CD2 HIS B 444     -18.779  74.621  59.463  1.00101.58           C
    ATOM   4902  ND1 HIS B 444     -18.868  74.981  57.231  1.00 98.26           N
    ATOM   4903  CE1 HIS B 444     -18.984  76.125  57.894  1.00100.95           C
    ATOM   4904  NE2 HIS B 444     -18.947  75.971  59.218  1.00101.67           N
    ATOM   4905  H   HIS B 444     -18.712  70.119  57.337  1.00  0.00           H
    ATOM   4906  HA  HIS B 444     -19.868  71.673  59.375  1.00  0.00           H
    ATOM   4907  HB2 HIS B 444     -17.680  72.175  58.451  1.00  0.00           H
    ATOM   4908  HB3 HIS B 444     -18.330  72.433  56.855  1.00  0.00           H
    ATOM   4909  HD2 HIS B 444     -18.731  74.221  60.464  1.00  0.00           H
    ATOM   4910  HE2 HIS B 444     -19.032  76.704  59.909  1.00  0.00           H
    ATOM   4911  HE1 HIS B 444     -19.114  77.083  57.412  1.00  0.00           H
    ATOM   4912  N   ARG B 445     -21.388  71.995  56.459  1.00103.21           N
    ATOM   4913  CA  ARG B 445     -22.634  72.376  55.801  1.00105.21           C
    ATOM   4914  C   ARG B 445     -23.849  71.793  56.530  1.00105.08           C
    ATOM   4915  O   ARG B 445     -24.772  72.539  56.846  1.00105.47           O
    ATOM   4916  CB  ARG B 445     -22.649  71.921  54.321  1.00108.21           C
    ATOM   4917  CG  ARG B 445     -23.466  72.877  53.434  1.00110.12           C
    ATOM   4918  CD  ARG B 445     -22.588  73.882  52.668  1.00113.13           C
    ATOM   4919  NE  ARG B 445     -21.571  74.500  53.534  1.00115.12           N
    ATOM   4920  CZ  ARG B 445     -20.401  75.019  53.128  1.00116.01           C
    ATOM   4921  NH1 ARG B 445     -20.082  75.077  51.826  1.00116.79           N
    ATOM   4922  NH2 ARG B 445     -19.535  75.477  54.041  1.00114.90           N1+
    ATOM   4923  H   ARG B 445     -20.739  71.448  55.906  1.00  0.00           H
    ATOM   4924  HA  ARG B 445     -22.710  73.463  55.851  1.00  0.00           H
    ATOM   4925  HB2 ARG B 445     -21.646  71.838  53.910  1.00  0.00           H
    ATOM   4926  HB3 ARG B 445     -23.056  70.911  54.243  1.00  0.00           H
    ATOM   4927  HG2 ARG B 445     -23.887  72.238  52.657  1.00  0.00           H
    ATOM   4928  HG3 ARG B 445     -24.332  73.323  53.926  1.00  0.00           H
    ATOM   4929  HD2 ARG B 445     -22.172  73.418  51.773  1.00  0.00           H
    ATOM   4930  HD3 ARG B 445     -23.205  74.717  52.335  1.00  0.00           H
    ATOM   4931  HE  ARG B 445     -21.788  74.523  54.520  1.00  0.00           H
    ATOM   4932 HH11 ARG B 445     -20.735  74.733  51.137  1.00  0.00           H
    ATOM   4933 HH12 ARG B 445     -19.231  75.522  51.522  1.00  0.00           H
    ATOM   4934 HH21 ARG B 445     -19.733  75.391  55.030  1.00  0.00           H
    ATOM   4935 HH22 ARG B 445     -18.658  75.890  53.760  1.00  0.00           H
    ATOM   4936  N   GLN B 446     -23.777  70.479  56.808  1.00105.59           N
    ATOM   4937  CA  GLN B 446     -24.812  69.703  57.472  1.00106.46           C
    ATOM   4938  C   GLN B 446     -25.028  70.098  58.947  1.00108.48           C
    ATOM   4939  O   GLN B 446     -26.160  70.002  59.404  1.00108.16           O
    ATOM   4940  CB  GLN B 446     -24.547  68.193  57.277  1.00106.01           C
    ATOM   4941  CG  GLN B 446     -25.673  67.256  57.769  1.00108.02           C
    ATOM   4942  CD  GLN B 446     -27.037  67.549  57.130  1.00109.16           C
    ATOM   4943  NE2 GLN B 446     -28.068  67.732  57.956  1.00109.38           N
    ATOM   4944  OE1 GLN B 446     -27.160  67.602  55.908  1.00106.87           O
    ATOM   4945  H   GLN B 446     -22.962  69.957  56.513  1.00  0.00           H
    ATOM   4946  HA  GLN B 446     -25.728  69.927  56.932  1.00  0.00           H
    ATOM   4947  HB2 GLN B 446     -24.395  68.000  56.215  1.00  0.00           H
    ATOM   4948  HB3 GLN B 446     -23.612  67.916  57.764  1.00  0.00           H
    ATOM   4949  HG2 GLN B 446     -25.408  66.223  57.545  1.00  0.00           H
    ATOM   4950  HG3 GLN B 446     -25.759  67.321  58.853  1.00  0.00           H
    ATOM   4951 HE21 GLN B 446     -27.938  67.692  58.956  1.00  0.00           H
    ATOM   4952 HE22 GLN B 446     -28.987  67.925  57.585  1.00  0.00           H
    ATOM   4953  N   MET B 447     -23.993  70.581  59.658  1.00110.79           N
    ATOM   4954  CA  MET B 447     -24.132  71.128  61.014  1.00110.18           C
    ATOM   4955  C   MET B 447     -24.968  72.417  61.039  1.00110.66           C
    ATOM   4956  O   MET B 447     -25.883  72.520  61.854  1.00111.08           O
    ATOM   4957  CB  MET B 447     -22.760  71.436  61.627  1.00109.27           C
    ATOM   4958  CG  MET B 447     -21.895  70.234  61.990  1.00107.25           C
    ATOM   4959  SD  MET B 447     -20.310  70.855  62.594  1.00109.58           S
    ATOM   4960  CE  MET B 447     -19.147  69.786  61.737  1.00107.99           C
    ATOM   4961  H   MET B 447     -23.071  70.618  59.243  1.00  0.00           H
    ATOM   4962  HA  MET B 447     -24.633  70.386  61.639  1.00  0.00           H
    ATOM   4963  HB2 MET B 447     -22.198  72.097  60.966  1.00  0.00           H
    ATOM   4964  HB3 MET B 447     -22.900  71.995  62.551  1.00  0.00           H
    ATOM   4965  HG2 MET B 447     -22.366  69.640  62.773  1.00  0.00           H
    ATOM   4966  HG3 MET B 447     -21.745  69.574  61.141  1.00  0.00           H
    ATOM   4967  HE1 MET B 447     -18.134  70.166  61.844  1.00  0.00           H
    ATOM   4968  HE2 MET B 447     -19.193  68.780  62.149  1.00  0.00           H
    ATOM   4969  HE3 MET B 447     -19.374  69.752  60.676  1.00  0.00           H
    ATOM   4970  N   GLN B 448     -24.644  73.355  60.131  1.00110.14           N
    ATOM   4971  CA  GLN B 448     -25.366  74.613  59.925  1.00110.02           C
    ATOM   4972  C   GLN B 448     -26.813  74.400  59.446  1.00110.52           C
    ATOM   4973  O   GLN B 448     -27.674  75.211  59.774  1.00107.02           O
    ATOM   4974  CB  GLN B 448     -24.602  75.481  58.911  1.00108.73           C
    ATOM   4975  CG  GLN B 448     -23.213  75.938  59.401  1.00110.03           C
    ATOM   4976  CD  GLN B 448     -22.423  76.690  58.327  1.00112.36           C
    ATOM   4977  NE2 GLN B 448     -22.418  76.191  57.088  1.00112.62           N
    ATOM   4978  OE1 GLN B 448     -21.807  77.712  58.613  1.00115.32           O
    ATOM   4979  H   GLN B 448     -23.869  73.184  59.504  1.00  0.00           H
    ATOM   4980  HA  GLN B 448     -25.396  75.142  60.878  1.00  0.00           H
    ATOM   4981  HB2 GLN B 448     -24.509  74.917  57.982  1.00  0.00           H
    ATOM   4982  HB3 GLN B 448     -25.192  76.366  58.666  1.00  0.00           H
    ATOM   4983  HG2 GLN B 448     -23.327  76.581  60.275  1.00  0.00           H
    ATOM   4984  HG3 GLN B 448     -22.611  75.090  59.726  1.00  0.00           H
    ATOM   4985 HE21 GLN B 448     -22.931  75.346  56.881  1.00  0.00           H
    ATOM   4986 HE22 GLN B 448     -21.922  76.675  56.355  1.00  0.00           H
    ATOM   4987  N   GLN B 449     -27.045  73.295  58.714  1.00112.42           N
    ATOM   4988  CA  GLN B 449     -28.346  72.808  58.253  1.00115.22           C
    ATOM   4989  C   GLN B 449     -29.261  72.474  59.447  1.00115.42           C
    ATOM   4990  O   GLN B 449     -30.420  72.881  59.448  1.00116.31           O
    ATOM   4991  CB  GLN B 449     -28.110  71.541  57.396  1.00115.54           C
    ATOM   4992  CG  GLN B 449     -29.070  71.344  56.208  1.00113.60           C
    ATOM   4993  CD  GLN B 449     -28.481  71.743  54.848  1.00112.32           C
    ATOM   4994  NE2 GLN B 449     -29.219  71.450  53.776  1.00110.81           N
    ATOM   4995  OE1 GLN B 449     -27.389  72.301  54.745  1.00112.52           O
    ATOM   4996  H   GLN B 449     -26.257  72.711  58.470  1.00  0.00           H
    ATOM   4997  HA  GLN B 449     -28.806  73.591  57.647  1.00  0.00           H
    ATOM   4998  HB2 GLN B 449     -27.085  71.529  57.044  1.00  0.00           H
    ATOM   4999  HB3 GLN B 449     -28.185  70.651  58.019  1.00  0.00           H
    ATOM   5000  HG2 GLN B 449     -29.324  70.285  56.144  1.00  0.00           H
    ATOM   5001  HG3 GLN B 449     -30.010  71.871  56.377  1.00  0.00           H
    ATOM   5002 HE21 GLN B 449     -30.117  71.001  53.879  1.00  0.00           H
    ATOM   5003 HE22 GLN B 449     -28.886  71.693  52.855  1.00  0.00           H
    ATOM   5004  N   GLN B 450     -28.704  71.766  60.447  1.00115.10           N
    ATOM   5005  CA  GLN B 450     -29.386  71.393  61.687  1.00116.13           C
    ATOM   5006  C   GLN B 450     -29.564  72.582  62.646  1.00116.03           C
    ATOM   5007  O   GLN B 450     -30.609  72.668  63.287  1.00115.84           O
    ATOM   5008  CB  GLN B 450     -28.625  70.238  62.381  1.00117.79           C
    ATOM   5009  CG  GLN B 450     -28.474  68.911  61.603  1.00118.43           C
    ATOM   5010  CD  GLN B 450     -29.771  68.232  61.157  1.00118.95           C
    ATOM   5011  NE2 GLN B 450     -29.933  66.957  61.513  1.00117.48           N
    ATOM   5012  OE1 GLN B 450     -30.586  68.813  60.445  1.00121.61           O
    ATOM   5013  H   GLN B 450     -27.741  71.471  60.366  1.00  0.00           H
    ATOM   5014  HA  GLN B 450     -30.392  71.060  61.437  1.00  0.00           H
    ATOM   5015  HB2 GLN B 450     -27.626  70.583  62.648  1.00  0.00           H
    ATOM   5016  HB3 GLN B 450     -29.110  70.010  63.327  1.00  0.00           H
    ATOM   5017  HG2 GLN B 450     -27.900  69.075  60.701  1.00  0.00           H
    ATOM   5018  HG3 GLN B 450     -27.886  68.217  62.206  1.00  0.00           H
    ATOM   5019 HE21 GLN B 450     -29.243  66.489  62.083  1.00  0.00           H
    ATOM   5020 HE22 GLN B 450     -30.760  66.455  61.223  1.00  0.00           H
    ATOM   5021  N   ALA B 451     -28.570  73.486  62.705  1.00115.45           N
    ATOM   5022  CA  ALA B 451     -28.613  74.713  63.502  1.00116.51           C
    ATOM   5023  C   ALA B 451     -29.670  75.715  63.010  1.00117.61           C
    ATOM   5024  O   ALA B 451     -30.350  76.313  63.841  1.00119.95           O
    ATOM   5025  CB  ALA B 451     -27.219  75.354  63.531  1.00114.68           C
    ATOM   5026  H   ALA B 451     -27.728  73.333  62.166  1.00  0.00           H
    ATOM   5027  HA  ALA B 451     -28.874  74.433  64.523  1.00  0.00           H
    ATOM   5028  HB1 ALA B 451     -27.209  76.267  64.127  1.00  0.00           H
    ATOM   5029  HB2 ALA B 451     -26.490  74.670  63.964  1.00  0.00           H
    ATOM   5030  HB3 ALA B 451     -26.881  75.616  62.530  1.00  0.00           H
    ATOM   5031  N   ALA B 452     -29.813  75.843  61.678  1.00116.87           N
    ATOM   5032  CA  ALA B 452     -30.794  76.695  61.005  1.00116.37           C
    ATOM   5033  C   ALA B 452     -32.246  76.264  61.253  1.00117.00           C
    ATOM   5034  O   ALA B 452     -33.071  77.108  61.602  1.00119.93           O
    ATOM   5035  CB  ALA B 452     -30.489  76.736  59.500  1.00114.81           C
    ATOM   5036  H   ALA B 452     -29.194  75.325  61.068  1.00  0.00           H
    ATOM   5037  HA  ALA B 452     -30.682  77.706  61.391  1.00  0.00           H
    ATOM   5038  HB1 ALA B 452     -31.225  77.334  58.962  1.00  0.00           H
    ATOM   5039  HB2 ALA B 452     -29.512  77.183  59.314  1.00  0.00           H
    ATOM   5040  HB3 ALA B 452     -30.486  75.737  59.062  1.00  0.00           H
    ATOM   5041  N   THR B 453     -32.525  74.959  61.097  1.00115.62           N
    ATOM   5042  CA  THR B 453     -33.843  74.373  61.361  1.00115.65           C
    ATOM   5043  C   THR B 453     -34.197  74.339  62.862  1.00116.81           C
    ATOM   5044  O   THR B 453     -35.384  74.398  63.173  1.00117.72           O
    ATOM   5045  CB  THR B 453     -33.967  72.931  60.803  1.00113.36           C
    ATOM   5046  CG2 THR B 453     -33.800  72.861  59.278  1.00111.04           C
    ATOM   5047  OG1 THR B 453     -33.059  72.035  61.418  1.00115.46           O
    ATOM   5048  H   THR B 453     -31.797  74.322  60.805  1.00  0.00           H
    ATOM   5049  HA  THR B 453     -34.590  74.992  60.859  1.00  0.00           H
    ATOM   5050  HB  THR B 453     -34.964  72.554  61.033  1.00  0.00           H
    ATOM   5051 HG21 THR B 453     -33.849  71.831  58.923  1.00  0.00           H
    ATOM   5052 HG22 THR B 453     -34.595  73.416  58.779  1.00  0.00           H
    ATOM   5053 HG23 THR B 453     -32.856  73.285  58.943  1.00  0.00           H
    ATOM   5054  HG1 THR B 453     -32.185  72.178  61.046  1.00  0.00           H
    ATOM   5055  N   ALA B 454     -33.191  74.298  63.755  1.00116.96           N
    ATOM   5056  CA  ALA B 454     -33.371  74.385  65.204  1.00117.30           C
    ATOM   5057  C   ALA B 454     -33.727  75.805  65.670  1.00116.47           C
    ATOM   5058  O   ALA B 454     -34.616  75.938  66.506  1.00116.50           O
    ATOM   5059  CB  ALA B 454     -32.111  73.877  65.912  1.00119.29           C
    ATOM   5060  H   ALA B 454     -32.238  74.225  63.428  1.00  0.00           H
    ATOM   5061  HA  ALA B 454     -34.195  73.727  65.486  1.00  0.00           H
    ATOM   5062  HB1 ALA B 454     -32.194  73.973  66.995  1.00  0.00           H
    ATOM   5063  HB2 ALA B 454     -31.937  72.823  65.692  1.00  0.00           H
    ATOM   5064  HB3 ALA B 454     -31.231  74.433  65.593  1.00  0.00           H
    ATOM   5065  N   GLN B 455     -33.079  76.830  65.087  1.00116.66           N
    ATOM   5066  CA  GLN B 455     -33.414  78.250  65.256  1.00117.80           C
    ATOM   5067  C   GLN B 455     -34.858  78.578  64.836  1.00116.91           C
    ATOM   5068  O   GLN B 455     -35.533  79.340  65.528  1.00117.77           O
    ATOM   5069  CB  GLN B 455     -32.411  79.111  64.460  1.00120.17           C
    ATOM   5070  CG  GLN B 455     -31.036  79.240  65.138  1.00123.78           C
    ATOM   5071  CD  GLN B 455     -29.948  79.785  64.205  1.00127.20           C
    ATOM   5072  NE2 GLN B 455     -29.044  80.605  64.740  1.00128.37           N
    ATOM   5073  OE1 GLN B 455     -29.899  79.458  63.022  1.00128.24           O
    ATOM   5074  H   GLN B 455     -32.340  76.631  64.425  1.00  0.00           H
    ATOM   5075  HA  GLN B 455     -33.321  78.490  66.317  1.00  0.00           H
    ATOM   5076  HB2 GLN B 455     -32.302  78.688  63.462  1.00  0.00           H
    ATOM   5077  HB3 GLN B 455     -32.808  80.116  64.311  1.00  0.00           H
    ATOM   5078  HG2 GLN B 455     -31.150  79.898  65.998  1.00  0.00           H
    ATOM   5079  HG3 GLN B 455     -30.698  78.278  65.523  1.00  0.00           H
    ATOM   5080 HE21 GLN B 455     -29.084  80.846  65.719  1.00  0.00           H
    ATOM   5081 HE22 GLN B 455     -28.294  80.966  64.170  1.00  0.00           H
    ATOM   5082  N   ALA B 456     -35.300  77.957  63.728  1.00125.03           N
    ATOM   5083  CA  ALA B 456     -36.658  78.041  63.205  1.00127.60           C
    ATOM   5084  C   ALA B 456     -37.677  77.295  64.079  1.00127.72           C
    ATOM   5085  O   ALA B 456     -38.688  77.893  64.433  1.00129.19           O
    ATOM   5086  CB  ALA B 456     -36.679  77.524  61.760  1.00126.08           C
    ATOM   5087  H   ALA B 456     -34.666  77.359  63.216  1.00  0.00           H
    ATOM   5088  HA  ALA B 456     -36.943  79.095  63.186  1.00  0.00           H
    ATOM   5089  HB1 ALA B 456     -37.674  77.615  61.322  1.00  0.00           H
    ATOM   5090  HB2 ALA B 456     -35.992  78.094  61.133  1.00  0.00           H
    ATOM   5091  HB3 ALA B 456     -36.386  76.476  61.700  1.00  0.00           H
    ATOM   5092  N   ALA B 457     -37.389  76.029  64.435  1.00127.86           N
    ATOM   5093  CA  ALA B 457     -38.237  75.175  65.276  1.00126.97           C
    ATOM   5094  C   ALA B 457     -38.399  75.675  66.724  1.00127.21           C
    ATOM   5095  O   ALA B 457     -39.433  75.395  67.329  1.00127.95           O
    ATOM   5096  CB  ALA B 457     -37.699  73.737  65.256  1.00126.68           C
    ATOM   5097  H   ALA B 457     -36.538  75.600  64.093  1.00  0.00           H
    ATOM   5098  HA  ALA B 457     -39.229  75.158  64.823  1.00  0.00           H
    ATOM   5099  HB1 ALA B 457     -38.317  73.074  65.864  1.00  0.00           H
    ATOM   5100  HB2 ALA B 457     -37.692  73.334  64.243  1.00  0.00           H
    ATOM   5101  HB3 ALA B 457     -36.681  73.688  65.643  1.00  0.00           H
    ATOM   5102  N   ALA B 458     -37.401  76.424  67.229  1.00128.34           N
    ATOM   5103  CA  ALA B 458     -37.409  77.095  68.530  1.00128.44           C
    ATOM   5104  C   ALA B 458     -38.294  78.353  68.554  1.00127.57           C
    ATOM   5105  O   ALA B 458     -38.918  78.621  69.580  1.00126.77           O
    ATOM   5106  CB  ALA B 458     -35.970  77.448  68.932  1.00127.72           C
    ATOM   5107  H   ALA B 458     -36.568  76.564  66.673  1.00  0.00           H
    ATOM   5108  HA  ALA B 458     -37.791  76.397  69.273  1.00  0.00           H
    ATOM   5109  HB1 ALA B 458     -35.942  77.969  69.890  1.00  0.00           H
    ATOM   5110  HB2 ALA B 458     -35.363  76.549  69.039  1.00  0.00           H
    ATOM   5111  HB3 ALA B 458     -35.490  78.089  68.191  1.00  0.00           H
    ATOM   5112  N   ALA B 459     -38.351  79.085  67.429  1.00125.77           N
    ATOM   5113  CA  ALA B 459     -39.229  80.242  67.246  1.00124.12           C
    ATOM   5114  C   ALA B 459     -40.667  79.838  66.872  1.00124.30           C
    ATOM   5115  O   ALA B 459     -41.601  80.550  67.239  1.00123.90           O
    ATOM   5116  CB  ALA B 459     -38.623  81.159  66.174  1.00121.43           C
    ATOM   5117  H   ALA B 459     -37.796  78.814  66.629  1.00  0.00           H
    ATOM   5118  HA  ALA B 459     -39.272  80.805  68.181  1.00  0.00           H
    ATOM   5119  HB1 ALA B 459     -39.238  82.046  66.018  1.00  0.00           H
    ATOM   5120  HB2 ALA B 459     -37.629  81.498  66.467  1.00  0.00           H
    ATOM   5121  HB3 ALA B 459     -38.527  80.647  65.216  1.00  0.00           H
    ATOM   5122  N   ALA B 460     -40.816  78.704  66.165  1.00124.91           N
    ATOM   5123  CA  ALA B 460     -42.091  78.141  65.721  1.00124.98           C
    ATOM   5124  C   ALA B 460     -42.909  77.491  66.850  1.00126.50           C
    ATOM   5125  O   ALA B 460     -44.129  77.412  66.715  1.00128.37           O
    ATOM   5126  CB  ALA B 460     -41.828  77.123  64.602  1.00121.66           C
    ATOM   5127  H   ALA B 460     -39.989  78.200  65.872  1.00  0.00           H
    ATOM   5128  HA  ALA B 460     -42.691  78.952  65.305  1.00  0.00           H
    ATOM   5129  HB1 ALA B 460     -42.760  76.709  64.217  1.00  0.00           H
    ATOM   5130  HB2 ALA B 460     -41.308  77.585  63.763  1.00  0.00           H
    ATOM   5131  HB3 ALA B 460     -41.220  76.291  64.958  1.00  0.00           H
    ATOM   5132  N   GLN B 461     -42.244  77.049  67.937  1.00127.69           N
    ATOM   5133  CA  GLN B 461     -42.873  76.486  69.140  1.00127.69           C
    ATOM   5134  C   GLN B 461     -43.164  77.658  70.063  1.00128.04           C
    ATOM   5135  O   GLN B 461     -43.477  77.510  71.244  1.00130.41           O
    ATOM   5136  CB  GLN B 461     -41.969  75.392  69.774  1.00128.31           C
    ATOM   5137  CG  GLN B 461     -40.654  75.876  70.422  1.00130.31           C
    ATOM   5138  CD  GLN B 461     -39.827  74.729  70.999  1.00132.80           C
    ATOM   5139  NE2 GLN B 461     -40.190  74.268  72.197  1.00131.87           N
    ATOM   5140  OE1 GLN B 461     -38.871  74.267  70.381  1.00134.54           O
    ATOM   5141  H   GLN B 461     -41.237  77.131  67.963  1.00  0.00           H
    ATOM   5142  HA  GLN B 461     -43.818  76.007  68.872  1.00  0.00           H
    ATOM   5144  HB2 GLN B 461     -42.551  74.865  70.532  1.00  0.00           H
    ATOM   5145  HB3 GLN B 461     -41.739  74.641  69.016  1.00  0.00           H
    ATOM   5146  HG2 GLN B 461     -40.061  76.383  69.669  1.00  0.00           H
    ATOM   5147  HG3 GLN B 461     -40.841  76.602  71.213  1.00  0.00           H
    ATOM   5148 HE21 GLN B 461     -40.977  74.671  72.684  1.00  0.00           H
    ATOM   5149 HE22 GLN B 461     -39.675  73.511  72.622  1.00  0.00           H
    ATOM   5151  N   ILE B 490     -33.054  67.692  67.926  1.00 90.27           N1+
    ATOM   5152  CA  ILE B 490     -31.697  68.038  67.462  1.00 90.63           C
    ATOM   5153  C   ILE B 490     -30.948  68.629  68.684  1.00 91.16           C
    ATOM   5154  O   ILE B 490     -31.101  69.810  69.005  1.00 89.54           O
    ATOM   5155  CB  ILE B 490     -31.690  69.035  66.253  1.00 92.14           C
    ATOM   5156  CG1 ILE B 490     -32.368  68.440  64.993  1.00 95.49           C
    ATOM   5157  CG2 ILE B 490     -30.309  69.630  65.887  1.00 90.90           C
    ATOM   5158  CD1 ILE B 490     -31.736  67.144  64.456  1.00 92.69           C
    ATOM   5160  H   ILE B 490     -33.007  66.920  68.577  1.00  0.00           H
    ATOM   5161  HA  ILE B 490     -31.177  67.131  67.148  1.00  0.00           H
    ATOM   5162  HB  ILE B 490     -32.305  69.890  66.540  1.00  0.00           H
    ATOM   5163 HG12 ILE B 490     -33.416  68.240  65.216  1.00  0.00           H
    ATOM   5164 HG13 ILE B 490     -32.384  69.188  64.199  1.00  0.00           H
    ATOM   5165 HG21 ILE B 490     -30.415  70.345  65.073  1.00  0.00           H
    ATOM   5166 HG22 ILE B 490     -29.856  70.179  66.711  1.00  0.00           H
    ATOM   5167 HG23 ILE B 490     -29.599  68.862  65.581  1.00  0.00           H
    ATOM   5168 HD11 ILE B 490     -32.100  66.933  63.451  1.00  0.00           H
    ATOM   5169 HD12 ILE B 490     -30.649  67.206  64.405  1.00  0.00           H
    ATOM   5170 HD13 ILE B 490     -31.994  66.289  65.081  1.00  0.00           H
    ATOM   5171  N   GLY B 491     -30.156  67.769  69.343  1.00 92.74           N
    ATOM   5172  CA  GLY B 491     -29.256  68.114  70.438  1.00 93.38           C
    ATOM   5173  C   GLY B 491     -27.806  67.876  70.003  1.00 93.91           C
    ATOM   5174  O   GLY B 491     -27.536  67.475  68.867  1.00 94.48           O
    ATOM   5175  H   GLY B 491     -30.146  66.799  69.046  1.00  0.00           H
    ATOM   5176  HA2 GLY B 491     -29.372  69.150  70.760  1.00  0.00           H
    ATOM   5177  HA3 GLY B 491     -29.484  67.476  71.292  1.00  0.00           H
    ATOM   5178  N   VAL B 492     -26.871  68.114  70.943  1.00 93.75           N
    ATOM   5179  CA  VAL B 492     -25.425  67.921  70.772  1.00 92.46           C
    ATOM   5180  C   VAL B 492     -25.020  66.482  70.387  1.00 92.60           C
    ATOM   5181  O   VAL B 492     -24.099  66.311  69.592  1.00 92.49           O
    ATOM   5182  CB  VAL B 492     -24.614  68.380  72.020  1.00 91.82           C
    ATOM   5183  CG1 VAL B 492     -24.972  67.630  73.320  1.00 90.04           C
    ATOM   5184  CG2 VAL B 492     -23.089  68.329  71.788  1.00 92.14           C
    ATOM   5185  H   VAL B 492     -27.162  68.441  71.853  1.00  0.00           H
    ATOM   5186  HA  VAL B 492     -25.132  68.578  69.957  1.00  0.00           H
    ATOM   5187  HB  VAL B 492     -24.866  69.430  72.180  1.00  0.00           H
    ATOM   5188 HG11 VAL B 492     -24.440  68.057  74.172  1.00  0.00           H
    ATOM   5189 HG12 VAL B 492     -26.037  67.692  73.543  1.00  0.00           H
    ATOM   5190 HG13 VAL B 492     -24.702  66.575  73.270  1.00  0.00           H
    ATOM   5191 HG21 VAL B 492     -22.553  68.858  72.571  1.00  0.00           H
    ATOM   5192 HG22 VAL B 492     -22.707  67.307  71.774  1.00  0.00           H
    ATOM   5193 HG23 VAL B 492     -22.818  68.795  70.840  1.00  0.00           H
    ATOM   5194  N   ASP B 493     -25.743  65.486  70.916  1.00 93.04           N
    ATOM   5195  CA  ASP B 493     -25.534  64.068  70.633  1.00 91.97           C
    ATOM   5196  C   ASP B 493     -25.990  63.622  69.244  1.00 90.99           C
    ATOM   5197  O   ASP B 493     -25.353  62.738  68.674  1.00 88.62           O
    ATOM   5198  CB  ASP B 493     -26.131  63.173  71.718  1.00 92.56           C
    ATOM   5199  CG  ASP B 493     -25.466  63.432  73.061  1.00 92.39           C
    ATOM   5200  OD1 ASP B 493     -24.301  62.998  73.173  1.00 90.94           O
    ATOM   5201  OD2 ASP B 493     -26.106  64.059  73.932  1.00 94.19           O1-
    ATOM   5202  H   ASP B 493     -26.488  65.703  71.564  1.00  0.00           H
    ATOM   5203  HA  ASP B 493     -24.462  63.903  70.664  1.00  0.00           H
    ATOM   5204  HB2 ASP B 493     -27.198  63.378  71.794  1.00  0.00           H
    ATOM   5205  HB3 ASP B 493     -26.045  62.112  71.479  1.00  0.00           H
    ATOM   5206  N   ASP B 494     -27.047  64.247  68.709  1.00 89.19           N
    ATOM   5207  CA  ASP B 494     -27.563  63.976  67.367  1.00 86.80           C
    ATOM   5208  C   ASP B 494     -26.603  64.435  66.260  1.00 84.65           C
    ATOM   5209  O   ASP B 494     -26.413  63.690  65.299  1.00 83.26           O
    ATOM   5210  CB  ASP B 494     -28.977  64.553  67.143  1.00 87.83           C
    ATOM   5211  CG  ASP B 494     -29.978  64.108  68.212  1.00 89.93           C
    ATOM   5212  OD1 ASP B 494     -30.167  62.879  68.335  1.00 87.64           O
    ATOM   5213  OD2 ASP B 494     -30.571  65.003  68.850  1.00 95.03           O1-
    ATOM   5214  H   ASP B 494     -27.525  64.966  69.234  1.00  0.00           H
    ATOM   5215  HA  ASP B 494     -27.647  62.890  67.274  1.00  0.00           H
    ATOM   5216  HB2 ASP B 494     -28.929  65.644  67.148  1.00  0.00           H
    ATOM   5217  HB3 ASP B 494     -29.372  64.283  66.163  1.00  0.00           H
    ATOM   5218  N   LEU B 495     -26.008  65.630  66.421  1.00 84.24           N
    ATOM   5219  CA  LEU B 495     -25.163  66.256  65.403  1.00 87.37           C
    ATOM   5220  C   LEU B 495     -23.648  66.015  65.603  1.00 88.10           C
    ATOM   5221  O   LEU B 495     -22.907  66.245  64.650  1.00 88.28           O
    ATOM   5222  CB  LEU B 495     -25.597  67.735  65.205  1.00 90.63           C
    ATOM   5223  CG  LEU B 495     -24.721  68.835  65.833  1.00 93.38           C
    ATOM   5224  CD1 LEU B 495     -25.198  70.243  65.407  1.00 95.55           C
    ATOM   5225  CD2 LEU B 495     -24.607  68.682  67.360  1.00 96.47           C
    ATOM   5226  H   LEU B 495     -26.211  66.182  67.243  1.00  0.00           H
    ATOM   5227  HA  LEU B 495     -25.389  65.775  64.450  1.00  0.00           H
    ATOM   5228  HB2 LEU B 495     -25.617  67.921  64.130  1.00  0.00           H
    ATOM   5229  HB3 LEU B 495     -26.633  67.866  65.524  1.00  0.00           H
    ATOM   5230  HG  LEU B 495     -23.727  68.713  65.412  1.00  0.00           H
    ATOM   5231 HD11 LEU B 495     -25.414  70.898  66.250  1.00  0.00           H
    ATOM   5232 HD12 LEU B 495     -24.442  70.744  64.800  1.00  0.00           H
    ATOM   5233 HD13 LEU B 495     -26.106  70.201  64.804  1.00  0.00           H
    ATOM   5234 HD21 LEU B 495     -24.873  69.591  67.897  1.00  0.00           H
    ATOM   5235 HD22 LEU B 495     -25.263  67.898  67.726  1.00  0.00           H
    ATOM   5236 HD23 LEU B 495     -23.594  68.413  67.659  1.00  0.00           H
    ATOM   5237  N   ARG B 496     -23.191  65.520  66.774  1.00 89.34           N
    ATOM   5238  CA  ARG B 496     -21.800  65.053  66.950  1.00 88.39           C
    ATOM   5239  C   ARG B 496     -21.519  63.741  66.198  1.00 87.17           C
    ATOM   5240  O   ARG B 496     -20.354  63.446  65.941  1.00 86.52           O
    ATOM   5241  CB  ARG B 496     -21.410  64.925  68.440  1.00 87.07           C
    ATOM   5242  CG  ARG B 496     -21.987  63.680  69.134  1.00 85.60           C
    ATOM   5243  CD  ARG B 496     -21.668  63.590  70.625  1.00 80.50           C
    ATOM   5244  NE  ARG B 496     -22.103  62.291  71.150  1.00 68.91           N
    ATOM   5245  CZ  ARG B 496     -21.390  61.158  71.050  1.00 67.82           C
    ATOM   5246  NH1 ARG B 496     -20.101  61.184  70.684  1.00 70.36           N
    ATOM   5247  NH2 ARG B 496     -21.982  59.985  71.307  1.00 68.82           N1+
    ATOM   5248  H   ARG B 496     -23.821  65.385  67.552  1.00  0.00           H
    ATOM   5249  HA  ARG B 496     -21.131  65.790  66.509  1.00  0.00           H
    ATOM   5250  HB2 ARG B 496     -20.323  64.869  68.506  1.00  0.00           H
    ATOM   5251  HB3 ARG B 496     -21.683  65.832  68.978  1.00  0.00           H
    ATOM   5252  HG2 ARG B 496     -23.061  63.772  69.020  1.00  0.00           H
    ATOM   5253  HG3 ARG B 496     -21.718  62.742  68.650  1.00  0.00           H
    ATOM   5254  HD2 ARG B 496     -20.633  63.844  70.851  1.00  0.00           H
    ATOM   5255  HD3 ARG B 496     -22.297  64.314  71.141  1.00  0.00           H
    ATOM   5256  HE  ARG B 496     -23.049  62.260  71.510  1.00  0.00           H
    ATOM   5257 HH11 ARG B 496     -19.647  62.069  70.509  1.00  0.00           H
    ATOM   5258 HH12 ARG B 496     -19.563  60.337  70.577  1.00  0.00           H
    ATOM   5259 HH21 ARG B 496     -22.959  59.955  71.561  1.00  0.00           H
    ATOM   5260 HH22 ARG B 496     -21.473  59.121  71.190  1.00  0.00           H
    ATOM   5261  N   ARG B 497     -22.584  62.996  65.846  1.00 85.29           N
    ATOM   5262  CA  ARG B 497     -22.541  61.841  64.951  1.00 85.52           C
    ATOM   5263  C   ARG B 497     -22.008  62.198  63.554  1.00 84.64           C
    ATOM   5264  O   ARG B 497     -21.394  61.338  62.925  1.00 85.36           O
    ATOM   5265  CB  ARG B 497     -23.944  61.224  64.825  1.00 85.55           C
    ATOM   5266  CG  ARG B 497     -24.443  60.505  66.086  1.00 89.97           C
    ATOM   5267  CD  ARG B 497     -25.914  60.080  65.957  1.00 91.59           C
    ATOM   5268  NE  ARG B 497     -26.246  58.958  66.847  1.00100.49           N
    ATOM   5269  CZ  ARG B 497     -26.521  58.995  68.162  1.00101.74           C
    ATOM   5270  NH1 ARG B 497     -26.538  60.138  68.858  1.00 99.04           N
    ATOM   5271  NH2 ARG B 497     -26.804  57.851  68.795  1.00103.18           N1+
    ATOM   5272  H   ARG B 497     -23.507  63.304  66.119  1.00  0.00           H
    ATOM   5273  HA  ARG B 497     -21.872  61.102  65.394  1.00  0.00           H
    ATOM   5274  HB2 ARG B 497     -24.655  61.988  64.512  1.00  0.00           H
    ATOM   5275  HB3 ARG B 497     -23.929  60.482  64.026  1.00  0.00           H
    ATOM   5276  HG2 ARG B 497     -23.799  59.669  66.364  1.00  0.00           H
    ATOM   5277  HG3 ARG B 497     -24.379  61.218  66.905  1.00  0.00           H
    ATOM   5278  HD2 ARG B 497     -26.562  60.903  66.261  1.00  0.00           H
    ATOM   5279  HD3 ARG B 497     -26.175  59.856  64.924  1.00  0.00           H
    ATOM   5280  HE  ARG B 497     -26.229  58.053  66.401  1.00  0.00           H
    ATOM   5281 HH11 ARG B 497     -26.292  61.015  68.415  1.00  0.00           H
    ATOM   5282 HH12 ARG B 497     -26.875  60.160  69.813  1.00  0.00           H
    ATOM   5283 HH21 ARG B 497     -26.823  56.976  68.295  1.00  0.00           H
    ATOM   5284 HH22 ARG B 497     -27.051  57.867  69.776  1.00  0.00           H
    ATOM   5285  N   LEU B 498     -22.224  63.454  63.110  1.00 83.98           N
    ATOM   5286  CA  LEU B 498     -21.660  64.010  61.875  1.00 82.79           C
    ATOM   5287  C   LEU B 498     -20.124  64.008  61.887  1.00 82.19           C
    ATOM   5288  O   LEU B 498     -19.515  63.821  60.839  1.00 82.83           O
    ATOM   5289  CB  LEU B 498     -22.200  65.435  61.600  1.00 85.18           C
    ATOM   5290  CG  LEU B 498     -23.739  65.587  61.567  1.00 86.48           C
    ATOM   5291  CD1 LEU B 498     -24.129  67.073  61.467  1.00 84.96           C
    ATOM   5292  CD2 LEU B 498     -24.399  64.762  60.447  1.00 85.00           C
    ATOM   5293  H   LEU B 498     -22.737  64.104  63.689  1.00  0.00           H
    ATOM   5294  HA  LEU B 498     -21.958  63.367  61.060  1.00  0.00           H
    ATOM   5295  HB2 LEU B 498     -21.789  66.119  62.344  1.00  0.00           H
    ATOM   5296  HB3 LEU B 498     -21.806  65.783  60.645  1.00  0.00           H
    ATOM   5297  HG  LEU B 498     -24.146  65.223  62.510  1.00  0.00           H
    ATOM   5298 HD11 LEU B 498     -25.195  67.203  61.283  1.00  0.00           H
    ATOM   5299 HD12 LEU B 498     -23.897  67.599  62.394  1.00  0.00           H
    ATOM   5300 HD13 LEU B 498     -23.588  67.574  60.663  1.00  0.00           H
    ATOM   5301 HD21 LEU B 498     -25.121  65.343  59.877  1.00  0.00           H
    ATOM   5302 HD22 LEU B 498     -23.669  64.381  59.734  1.00  0.00           H
    ATOM   5303 HD23 LEU B 498     -24.940  63.913  60.862  1.00  0.00           H
    ATOM   5304  N   CYS B 499     -19.548  64.178  63.085  1.00 81.17           N
    ATOM   5305  CA  CYS B 499     -18.119  64.248  63.363  1.00 81.57           C
    ATOM   5306  C   CYS B 499     -17.530  62.869  63.731  1.00 81.37           C
    ATOM   5307  O   CYS B 499     -16.385  62.830  64.180  1.00 82.54           O
    ATOM   5308  CB  CYS B 499     -17.853  65.240  64.516  1.00 82.83           C
    ATOM   5309  SG  CYS B 499     -18.743  66.804  64.271  1.00 81.85           S
    ATOM   5310  H   CYS B 499     -20.144  64.292  63.893  1.00  0.00           H
    ATOM   5311  HA  CYS B 499     -17.598  64.605  62.473  1.00  0.00           H
    ATOM   5312  HB2 CYS B 499     -18.170  64.824  65.471  1.00  0.00           H
    ATOM   5313  HB3 CYS B 499     -16.789  65.456  64.600  1.00  0.00           H
    ATOM   5314  HG  CYS B 499     -18.015  67.225  63.234  1.00  0.00           H
    ATOM   5315  N   ILE B 500     -18.306  61.778  63.581  1.00 81.15           N
    ATOM   5316  CA  ILE B 500     -17.928  60.418  63.974  1.00 80.70           C
    ATOM   5317  C   ILE B 500     -18.017  59.447  62.785  1.00 80.07           C
    ATOM   5318  O   ILE B 500     -18.940  59.525  61.975  1.00 81.22           O
    ATOM   5319  CB  ILE B 500     -18.817  59.894  65.148  1.00 80.95           C
    ATOM   5320  CG1 ILE B 500     -18.452  60.626  66.456  1.00 82.98           C
    ATOM   5321  CG2 ILE B 500     -18.788  58.365  65.406  1.00 79.78           C
    ATOM   5322  CD1 ILE B 500     -19.508  60.487  67.558  1.00 86.29           C
    ATOM   5323  H   ILE B 500     -19.235  61.877  63.193  1.00  0.00           H
    ATOM   5324  HA  ILE B 500     -16.895  60.423  64.317  1.00  0.00           H
    ATOM   5325  HB  ILE B 500     -19.847  60.150  64.903  1.00  0.00           H
    ATOM   5326 HG12 ILE B 500     -17.494  60.253  66.818  1.00  0.00           H
    ATOM   5327 HG13 ILE B 500     -18.298  61.689  66.272  1.00  0.00           H
    ATOM   5328 HG21 ILE B 500     -19.384  58.092  66.277  1.00  0.00           H
    ATOM   5329 HG22 ILE B 500     -19.198  57.795  64.572  1.00  0.00           H
    ATOM   5330 HG23 ILE B 500     -17.771  58.018  65.588  1.00  0.00           H
    ATOM   5331 HD11 ILE B 500     -19.617  61.432  68.087  1.00  0.00           H
    ATOM   5332 HD12 ILE B 500     -20.489  60.222  67.162  1.00  0.00           H
    ATOM   5333 HD13 ILE B 500     -19.225  59.720  68.279  1.00  0.00           H
    ATOM   5334  N   LEU B 501     -17.059  58.512  62.771  1.00 78.77           N
    ATOM   5335  CA  LEU B 501     -17.017  57.313  61.944  1.00 74.03           C
    ATOM   5336  C   LEU B 501     -16.992  56.106  62.890  1.00 69.79           C
    ATOM   5337  O   LEU B 501     -16.338  56.162  63.929  1.00 67.43           O
    ATOM   5338  CB  LEU B 501     -15.720  57.337  61.110  1.00 76.69           C
    ATOM   5339  CG  LEU B 501     -15.724  58.339  59.942  1.00 79.88           C
    ATOM   5340  CD1 LEU B 501     -14.289  58.618  59.466  1.00 80.97           C
    ATOM   5341  CD2 LEU B 501     -16.622  57.869  58.787  1.00 76.56           C
    ATOM   5342  H   LEU B 501     -16.341  58.553  63.485  1.00  0.00           H
    ATOM   5343  HA  LEU B 501     -17.892  57.246  61.294  1.00  0.00           H
    ATOM   5344  HB2 LEU B 501     -14.895  57.582  61.772  1.00  0.00           H
    ATOM   5345  HB3 LEU B 501     -15.495  56.342  60.720  1.00  0.00           H
    ATOM   5346  HG  LEU B 501     -16.118  59.287  60.309  1.00  0.00           H
    ATOM   5347 HD11 LEU B 501     -14.255  59.483  58.803  1.00  0.00           H
    ATOM   5348 HD12 LEU B 501     -13.623  58.830  60.302  1.00  0.00           H
    ATOM   5349 HD13 LEU B 501     -13.876  57.764  58.931  1.00  0.00           H
    ATOM   5350 HD21 LEU B 501     -16.098  57.892  57.832  1.00  0.00           H
    ATOM   5351 HD22 LEU B 501     -16.971  56.847  58.932  1.00  0.00           H
    ATOM   5352 HD23 LEU B 501     -17.500  58.508  58.693  1.00  0.00           H
    ATOM   5353  N   ARG B 502     -17.664  55.019  62.496  1.00 66.62           N
    ATOM   5354  CA  ARG B 502     -17.599  53.731  63.184  1.00 67.07           C
    ATOM   5355  C   ARG B 502     -17.195  52.661  62.171  1.00 69.38           C
    ATOM   5356  O   ARG B 502     -17.582  52.755  61.007  1.00 72.12           O
    ATOM   5357  CB  ARG B 502     -18.959  53.413  63.824  1.00 64.73           C
    ATOM   5358  CG  ARG B 502     -19.214  54.153  65.142  1.00 65.84           C
    ATOM   5359  CD  ARG B 502     -20.450  53.601  65.860  1.00 69.64           C
    ATOM   5360  NE  ARG B 502     -20.756  54.336  67.089  1.00 74.73           N
    ATOM   5361  CZ  ARG B 502     -21.723  54.023  67.966  1.00 78.53           C
    ATOM   5362  NH1 ARG B 502     -22.565  53.001  67.751  1.00 80.00           N
    ATOM   5363  NH2 ARG B 502     -21.836  54.754  69.079  1.00 81.28           N1+
    ATOM   5364  H   ARG B 502     -18.169  55.029  61.620  1.00  0.00           H
    ATOM   5365  HA  ARG B 502     -16.836  53.749  63.962  1.00  0.00           H
    ATOM   5366  HB2 ARG B 502     -19.770  53.605  63.127  1.00  0.00           H
    ATOM   5367  HB3 ARG B 502     -18.996  52.351  64.038  1.00  0.00           H
    ATOM   5368  HG2 ARG B 502     -18.352  53.943  65.775  1.00  0.00           H
    ATOM   5369  HG3 ARG B 502     -19.255  55.236  65.026  1.00  0.00           H
    ATOM   5370  HD2 ARG B 502     -21.309  53.500  65.200  1.00  0.00           H
    ATOM   5371  HD3 ARG B 502     -20.202  52.601  66.207  1.00  0.00           H
    ATOM   5372  HE  ARG B 502     -20.150  55.117  67.323  1.00  0.00           H
    ATOM   5373 HH11 ARG B 502     -22.478  52.452  66.908  1.00  0.00           H
    ATOM   5374 HH12 ARG B 502     -23.284  52.777  68.422  1.00  0.00           H
    ATOM   5375 HH21 ARG B 502     -21.161  55.496  69.240  1.00  0.00           H
    ATOM   5376 HH22 ARG B 502     -22.548  54.562  69.768  1.00  0.00           H
    ATOM   5377  N   MET B 503     -16.415  51.671  62.630  1.00 72.70           N
    ATOM   5378  CA  MET B 503     -15.842  50.653  61.760  1.00 71.47           C
    ATOM   5379  C   MET B 503     -15.638  49.332  62.503  1.00 73.11           C
    ATOM   5380  O   MET B 503     -14.883  49.306  63.470  1.00 74.83           O
    ATOM   5381  CB  MET B 503     -14.551  51.197  61.122  1.00 70.07           C
    ATOM   5382  CG  MET B 503     -13.922  50.262  60.081  1.00 68.65           C
    ATOM   5383  SD  MET B 503     -12.396  50.872  59.319  1.00 74.91           S
    ATOM   5384  CE  MET B 503     -12.675  52.654  59.126  1.00 71.97           C
    ATOM   5385  H   MET B 503     -16.120  51.657  63.599  1.00  0.00           H
    ATOM   5386  HA  MET B 503     -16.550  50.478  60.958  1.00  0.00           H
    ATOM   5387  HB2 MET B 503     -14.788  52.145  60.642  1.00  0.00           H
    ATOM   5388  HB3 MET B 503     -13.815  51.423  61.893  1.00  0.00           H
    ATOM   5389  HG2 MET B 503     -13.679  49.300  60.531  1.00  0.00           H
    ATOM   5390  HG3 MET B 503     -14.645  50.043  59.296  1.00  0.00           H
    ATOM   5391  HE1 MET B 503     -11.835  53.083  58.584  1.00  0.00           H
    ATOM   5392  HE2 MET B 503     -13.585  52.870  58.572  1.00  0.00           H
    ATOM   5393  HE3 MET B 503     -12.730  53.145  60.097  1.00  0.00           H
    ATOM   5394  N   SER B 504     -16.274  48.259  62.005  1.00 74.20           N
    ATOM   5395  CA  SER B 504     -16.099  46.900  62.511  1.00 75.57           C
    ATOM   5396  C   SER B 504     -15.125  46.120  61.627  1.00 74.06           C
    ATOM   5397  O   SER B 504     -15.249  46.151  60.402  1.00 73.64           O
    ATOM   5398  CB  SER B 504     -17.462  46.193  62.598  1.00 76.42           C
    ATOM   5399  OG  SER B 504     -17.332  44.904  63.155  1.00 77.57           O
    ATOM   5400  H   SER B 504     -16.855  48.362  61.183  1.00  0.00           H
    ATOM   5401  HA  SER B 504     -15.682  46.933  63.514  1.00  0.00           H
    ATOM   5402  HB2 SER B 504     -18.155  46.770  63.211  1.00  0.00           H
    ATOM   5403  HB3 SER B 504     -17.909  46.080  61.613  1.00  0.00           H
    ATOM   5404  HG  SER B 504     -17.072  44.291  62.459  1.00  0.00           H
    ATOM   5405  N   PHE B 505     -14.212  45.394  62.286  1.00 75.55           N
    ATOM   5406  CA  PHE B 505     -13.314  44.436  61.655  1.00 76.89           C
    ATOM   5407  C   PHE B 505     -13.984  43.058  61.590  1.00 78.45           C
    ATOM   5408  O   PHE B 505     -14.635  42.656  62.556  1.00 82.06           O
    ATOM   5409  CB  PHE B 505     -12.010  44.346  62.468  1.00 74.73           C
    ATOM   5410  CG  PHE B 505     -11.163  45.604  62.476  1.00 74.65           C
    ATOM   5411  CD1 PHE B 505     -10.323  45.895  61.381  1.00 73.38           C
    ATOM   5412  CD2 PHE B 505     -11.306  46.559  63.506  1.00 76.63           C
    ATOM   5413  CE1 PHE B 505      -9.583  47.069  61.372  1.00 73.19           C
    ATOM   5414  CE2 PHE B 505     -10.563  47.732  63.470  1.00 75.86           C
    ATOM   5415  CZ  PHE B 505      -9.698  47.980  62.412  1.00 75.50           C
    ATOM   5416  H   PHE B 505     -14.210  45.396  63.297  1.00  0.00           H
    ATOM   5417  HA  PHE B 505     -13.073  44.763  60.642  1.00  0.00           H
    ATOM   5418  HB2 PHE B 505     -12.226  44.066  63.499  1.00  0.00           H
    ATOM   5419  HB3 PHE B 505     -11.395  43.543  62.059  1.00  0.00           H
    ATOM   5420  HD1 PHE B 505     -10.233  45.197  60.561  1.00  0.00           H
    ATOM   5421  HD2 PHE B 505     -11.981  46.377  64.329  1.00  0.00           H
    ATOM   5422  HE1 PHE B 505      -8.913  47.275  60.554  1.00  0.00           H
    ATOM   5423  HE2 PHE B 505     -10.656  48.454  64.266  1.00  0.00           H
    ATOM   5424  HZ  PHE B 505      -9.120  48.891  62.393  1.00  0.00           H
    ATOM   5425  N   VAL B 506     -13.752  42.352  60.469  1.00 78.79           N
    ATOM   5426  CA  VAL B 506     -14.121  40.957  60.199  1.00 80.12           C
    ATOM   5427  C   VAL B 506     -15.640  40.741  60.033  1.00 81.80           C
    ATOM   5428  O   VAL B 506     -16.085  40.519  58.908  1.00 85.33           O
    ATOM   5429  CB  VAL B 506     -13.522  39.954  61.236  1.00 80.92           C
    ATOM   5430  CG1 VAL B 506     -13.969  38.490  61.051  1.00 75.60           C
    ATOM   5431  CG2 VAL B 506     -11.985  40.022  61.231  1.00 82.61           C
    ATOM   5432  H   VAL B 506     -13.228  42.793  59.726  1.00  0.00           H
    ATOM   5433  HA  VAL B 506     -13.672  40.729  59.231  1.00  0.00           H
    ATOM   5434  HB  VAL B 506     -13.841  40.238  62.239  1.00  0.00           H
    ATOM   5435 HG11 VAL B 506     -13.421  37.848  61.737  1.00  0.00           H
    ATOM   5436 HG12 VAL B 506     -15.028  38.342  61.257  1.00  0.00           H
    ATOM   5437 HG13 VAL B 506     -13.772  38.137  60.039  1.00  0.00           H
    ATOM   5438 HG21 VAL B 506     -11.565  39.341  61.965  1.00  0.00           H
    ATOM   5439 HG22 VAL B 506     -11.576  39.749  60.257  1.00  0.00           H
    ATOM   5440 HG23 VAL B 506     -11.618  41.017  61.479  1.00  0.00           H
    ATOM   5441  N   LYS B 507     -16.397  40.795  61.141  1.00 82.02           N
    ATOM   5442  CA  LYS B 507     -17.850  40.640  61.163  1.00 81.61           C
    ATOM   5443  C   LYS B 507     -18.568  41.914  60.703  1.00 81.09           C
    ATOM   5444  O   LYS B 507     -18.092  43.019  60.972  1.00 83.85           O
    ATOM   5445  CB  LYS B 507     -18.304  40.288  62.588  1.00 81.66           C
    ATOM   5446  CG  LYS B 507     -17.826  38.927  63.133  1.00 80.66           C
    ATOM   5447  CD  LYS B 507     -18.372  37.727  62.350  1.00 83.79           C
    ATOM   5448  CE  LYS B 507     -18.039  36.357  62.961  1.00 80.69           C
    ATOM   5449  NZ  LYS B 507     -16.608  36.054  62.929  1.00 83.39           N1+
    ATOM   5450  H   LYS B 507     -15.958  41.025  62.021  1.00  0.00           H
    ATOM   5451  HA  LYS B 507     -18.130  39.837  60.486  1.00  0.00           H
    ATOM   5452  HB2 LYS B 507     -17.968  41.089  63.241  1.00  0.00           H
    ATOM   5453  HB3 LYS B 507     -19.393  40.307  62.630  1.00  0.00           H
    ATOM   5454  HG2 LYS B 507     -16.736  38.899  63.157  1.00  0.00           H
    ATOM   5455  HG3 LYS B 507     -18.156  38.832  64.166  1.00  0.00           H
    ATOM   5456  HD2 LYS B 507     -19.454  37.817  62.251  1.00  0.00           H
    ATOM   5457  HD3 LYS B 507     -17.977  37.769  61.341  1.00  0.00           H
    ATOM   5458  HE2 LYS B 507     -18.380  36.286  63.991  1.00  0.00           H
    ATOM   5459  HE3 LYS B 507     -18.536  35.576  62.393  1.00  0.00           H
    ATOM   5460  HZ1 LYS B 507     -16.090  36.714  63.488  1.00  0.00           H
    ATOM   5461  HZ2 LYS B 507     -16.456  35.114  63.274  1.00  0.00           H
    ATOM   5462  HZ3 LYS B 507     -16.288  36.091  61.969  1.00  0.00           H
    ATOM   5463  N   GLY B 508     -19.725  41.712  60.058  1.00 80.26           N
    ATOM   5464  CA  GLY B 508     -20.626  42.777  59.637  1.00 78.11           C
    ATOM   5465  C   GLY B 508     -21.872  42.769  60.522  1.00 77.13           C
    ATOM   5466  O   GLY B 508     -22.255  41.736  61.076  1.00 73.46           O
    ATOM   5467  H   GLY B 508     -20.041  40.766  59.889  1.00  0.00           H
    ATOM   5468  HA2 GLY B 508     -20.151  43.758  59.668  1.00  0.00           H
    ATOM   5469  HA3 GLY B 508     -20.929  42.593  58.607  1.00  0.00           H
    ATOM   5470  N   TRP B 509     -22.525  43.937  60.592  1.00 82.04           N
    ATOM   5471  CA  TRP B 509     -23.844  44.131  61.186  1.00 86.84           C
    ATOM   5472  C   TRP B 509     -24.772  44.772  60.142  1.00 92.15           C
    ATOM   5473  O   TRP B 509     -24.357  45.039  59.013  1.00 93.65           O
    ATOM   5474  CB  TRP B 509     -23.714  44.990  62.470  1.00 83.61           C
    ATOM   5475  CG  TRP B 509     -23.596  46.477  62.274  1.00 80.08           C
    ATOM   5476  CD1 TRP B 509     -24.622  47.354  62.362  1.00 82.87           C
    ATOM   5477  CD2 TRP B 509     -22.441  47.261  61.850  1.00 74.54           C
    ATOM   5478  CE2 TRP B 509     -22.856  48.620  61.693  1.00 74.35           C
    ATOM   5479  CE3 TRP B 509     -21.088  46.970  61.564  1.00 71.89           C
    ATOM   5480  NE1 TRP B 509     -24.191  48.617  62.029  1.00 79.12           N
    ATOM   5481  CZ2 TRP B 509     -21.981  49.632  61.272  1.00 70.25           C
    ATOM   5482  CZ3 TRP B 509     -20.202  47.984  61.151  1.00 69.15           C
    ATOM   5483  CH2 TRP B 509     -20.642  49.311  60.994  1.00 66.76           C
    ATOM   5484  H   TRP B 509     -22.166  44.731  60.082  1.00  0.00           H
    ATOM   5485  HA  TRP B 509     -24.269  43.163  61.437  1.00  0.00           H
    ATOM   5486  HB2 TRP B 509     -24.596  44.822  63.090  1.00  0.00           H
    ATOM   5487  HB3 TRP B 509     -22.875  44.643  63.073  1.00  0.00           H
    ATOM   5488  HD1 TRP B 509     -25.633  47.083  62.624  1.00  0.00           H
    ATOM   5489  HE1 TRP B 509     -24.806  49.418  62.010  1.00  0.00           H
    ATOM   5490  HE3 TRP B 509     -20.726  45.957  61.668  1.00  0.00           H
    ATOM   5491  HZ2 TRP B 509     -22.346  50.641  61.168  1.00  0.00           H
    ATOM   5492  HZ3 TRP B 509     -19.172  47.748  60.955  1.00  0.00           H
    ATOM   5493  HH2 TRP B 509     -19.953  50.080  60.674  1.00  0.00           H
    ATOM   5494  N   GLY B 510     -26.012  45.033  60.571  1.00 98.27           N
    ATOM   5495  CA  GLY B 510     -27.019  45.731  59.787  1.00102.82           C
    ATOM   5496  C   GLY B 510     -28.058  44.726  59.266  1.00107.31           C
    ATOM   5497  O   GLY B 510     -28.141  43.605  59.774  1.00106.93           O
    ATOM   5498  H   GLY B 510     -26.288  44.728  61.495  1.00  0.00           H
    ATOM   5499  HA2 GLY B 510     -27.518  46.455  60.432  1.00  0.00           H
    ATOM   5500  HA3 GLY B 510     -26.558  46.293  58.974  1.00  0.00           H
    ATOM   5501  N   PRO B 511     -28.888  45.134  58.281  1.00110.38           N
    ATOM   5502  CA  PRO B 511     -30.093  44.387  57.865  1.00110.94           C
    ATOM   5503  C   PRO B 511     -29.857  43.152  56.963  1.00110.20           C
    ATOM   5504  O   PRO B 511     -30.813  42.697  56.335  1.00112.69           O
    ATOM   5505  CB  PRO B 511     -30.939  45.476  57.178  1.00112.91           C
    ATOM   5506  CG  PRO B 511     -29.918  46.423  56.573  1.00112.77           C
    ATOM   5507  CD  PRO B 511     -28.815  46.441  57.621  1.00112.30           C
    ATOM   5508  HA  PRO B 511     -30.630  44.028  58.744  1.00  0.00           H
    ATOM   5509  HB2 PRO B 511     -31.646  45.104  56.436  1.00  0.00           H
    ATOM   5510  HB3 PRO B 511     -31.523  46.004  57.934  1.00  0.00           H
    ATOM   5511  HG2 PRO B 511     -29.535  45.996  55.644  1.00  0.00           H
    ATOM   5512  HG3 PRO B 511     -30.319  47.412  56.349  1.00  0.00           H
    ATOM   5513  HD2 PRO B 511     -27.845  46.627  57.157  1.00  0.00           H
    ATOM   5514  HD3 PRO B 511     -29.004  47.224  58.357  1.00  0.00           H
    ATOM   5515  N   ASP B 512     -28.627  42.614  56.923  1.00109.65           N
    ATOM   5516  CA  ASP B 512     -28.264  41.425  56.143  1.00107.08           C
    ATOM   5517  C   ASP B 512     -27.266  40.548  56.934  1.00104.92           C
    ATOM   5518  O   ASP B 512     -26.512  39.782  56.334  1.00103.44           O
    ATOM   5519  CB  ASP B 512     -27.749  41.829  54.734  1.00107.10           C
    ATOM   5520  CG  ASP B 512     -27.954  40.784  53.627  1.00107.00           C
    ATOM   5521  OD1 ASP B 512     -27.854  41.202  52.453  1.00106.94           O
    ATOM   5522  OD2 ASP B 512     -28.173  39.594  53.945  1.00107.21           O1-
    ATOM   5523  H   ASP B 512     -27.886  43.025  57.472  1.00  0.00           H
    ATOM   5524  HA  ASP B 512     -29.154  40.803  56.033  1.00  0.00           H
    ATOM   5525  HB2 ASP B 512     -28.277  42.731  54.420  1.00  0.00           H
    ATOM   5526  HB3 ASP B 512     -26.688  42.077  54.782  1.00  0.00           H
    ATOM   5527  N   TYR B 513     -27.265  40.673  58.272  1.00102.43           N
    ATOM   5528  CA  TYR B 513     -26.394  39.927  59.181  1.00 98.61           C
    ATOM   5529  C   TYR B 513     -27.185  39.565  60.451  1.00 96.71           C
    ATOM   5530  O   TYR B 513     -28.117  40.297  60.793  1.00 93.54           O
    ATOM   5531  CB  TYR B 513     -25.146  40.774  59.520  1.00 97.47           C
    ATOM   5532  CG  TYR B 513     -24.225  41.009  58.336  1.00 95.30           C
    ATOM   5533  CD1 TYR B 513     -23.255  40.046  57.992  1.00 93.45           C
    ATOM   5534  CD2 TYR B 513     -24.379  42.160  57.536  1.00 94.32           C
    ATOM   5535  CE1 TYR B 513     -22.477  40.208  56.830  1.00 91.30           C
    ATOM   5536  CE2 TYR B 513     -23.599  42.328  56.378  1.00 91.72           C
    ATOM   5537  CZ  TYR B 513     -22.663  41.342  56.013  1.00 91.87           C
    ATOM   5538  OH  TYR B 513     -21.952  41.488  54.859  1.00 94.14           O
    ATOM   5539  H   TYR B 513     -27.920  41.302  58.716  1.00  0.00           H
    ATOM   5540  HA  TYR B 513     -26.080  39.018  58.672  1.00  0.00           H
    ATOM   5541  HB2 TYR B 513     -25.449  41.734  59.939  1.00  0.00           H
    ATOM   5542  HB3 TYR B 513     -24.561  40.281  60.298  1.00  0.00           H
    ATOM   5543  HD1 TYR B 513     -23.124  39.169  58.608  1.00  0.00           H
    ATOM   5544  HD2 TYR B 513     -25.131  42.894  57.783  1.00  0.00           H
    ATOM   5545  HE1 TYR B 513     -21.753  39.453  56.560  1.00  0.00           H
    ATOM   5546  HE2 TYR B 513     -23.741  43.199  55.756  1.00  0.00           H
    ATOM   5547  HH  TYR B 513     -21.316  40.769  54.704  1.00  0.00           H
    ATOM   5548  N   PRO B 514     -26.783  38.483  61.166  1.00 96.58           N
    ATOM   5549  CA  PRO B 514     -27.344  38.149  62.497  1.00 96.62           C
    ATOM   5550  C   PRO B 514     -27.225  39.258  63.563  1.00 95.27           C
    ATOM   5551  O   PRO B 514     -28.033  39.295  64.489  1.00 94.47           O
    ATOM   5552  CB  PRO B 514     -26.567  36.891  62.926  1.00 96.96           C
    ATOM   5553  CG  PRO B 514     -26.083  36.276  61.626  1.00 96.13           C
    ATOM   5554  CD  PRO B 514     -25.777  37.494  60.766  1.00 95.84           C
    ATOM   5555  HA  PRO B 514     -28.396  37.897  62.356  1.00  0.00           H
    ATOM   5556  HB2 PRO B 514     -25.700  37.156  63.536  1.00  0.00           H
    ATOM   5557  HB3 PRO B 514     -27.176  36.198  63.507  1.00  0.00           H
    ATOM   5558  HG2 PRO B 514     -25.235  35.603  61.748  1.00  0.00           H
    ATOM   5559  HG3 PRO B 514     -26.896  35.708  61.171  1.00  0.00           H
    ATOM   5560  HD2 PRO B 514     -24.781  37.884  60.985  1.00  0.00           H
    ATOM   5561  HD3 PRO B 514     -25.815  37.209  59.715  1.00  0.00           H
    ATOM   5562  N   ARG B 515     -26.227  40.138  63.387  1.00 93.58           N
    ATOM   5563  CA  ARG B 515     -25.945  41.293  64.226  1.00 91.06           C
    ATOM   5564  C   ARG B 515     -26.747  42.495  63.720  1.00 89.69           C
    ATOM   5565  O   ARG B 515     -26.644  42.841  62.544  1.00 90.87           O
    ATOM   5566  CB  ARG B 515     -24.434  41.582  64.187  1.00 87.62           C
    ATOM   5567  CG  ARG B 515     -23.571  40.339  64.427  1.00 87.46           C
    ATOM   5568  CD  ARG B 515     -22.080  40.658  64.390  1.00 90.66           C
    ATOM   5569  NE  ARG B 515     -21.285  39.430  64.327  1.00 93.52           N
    ATOM   5570  CZ  ARG B 515     -21.210  38.480  65.271  1.00 94.58           C
    ATOM   5571  NH1 ARG B 515     -21.758  38.624  66.485  1.00 94.26           N
    ATOM   5572  NH2 ARG B 515     -20.571  37.348  64.982  1.00 97.38           N1+
    ATOM   5573  H   ARG B 515     -25.624  40.030  62.585  1.00  0.00           H
    ATOM   5574  HA  ARG B 515     -26.229  41.063  65.253  1.00  0.00           H
    ATOM   5575  HB2 ARG B 515     -24.164  41.971  63.210  1.00  0.00           H
    ATOM   5576  HB3 ARG B 515     -24.184  42.363  64.906  1.00  0.00           H
    ATOM   5577  HG2 ARG B 515     -23.831  39.984  65.425  1.00  0.00           H
    ATOM   5578  HG3 ARG B 515     -23.786  39.517  63.743  1.00  0.00           H
    ATOM   5579  HD2 ARG B 515     -21.826  41.362  63.600  1.00  0.00           H
    ATOM   5580  HD3 ARG B 515     -21.820  41.133  65.327  1.00  0.00           H
    ATOM   5581  HE  ARG B 515     -20.866  39.237  63.428  1.00  0.00           H
    ATOM   5582 HH11 ARG B 515     -22.187  39.502  66.750  1.00  0.00           H
    ATOM   5583 HH12 ARG B 515     -21.704  37.879  67.164  1.00  0.00           H
    ATOM   5584 HH21 ARG B 515     -20.591  36.561  65.619  1.00  0.00           H
    ATOM   5585 HH22 ARG B 515     -20.208  37.193  64.054  1.00  0.00           H
    ATOM   5586  N   GLN B 516     -27.535  43.100  64.617  1.00 88.46           N
    ATOM   5587  CA  GLN B 516     -28.438  44.206  64.299  1.00 88.18           C
    ATOM   5588  C   GLN B 516     -27.747  45.572  64.405  1.00 86.75           C
    ATOM   5589  O   GLN B 516     -28.000  46.436  63.565  1.00 81.96           O
    ATOM   5590  CB  GLN B 516     -29.656  44.139  65.237  1.00 92.89           C
    ATOM   5591  CG  GLN B 516     -30.505  42.870  65.027  1.00 98.52           C
    ATOM   5592  CD  GLN B 516     -31.708  42.813  65.970  1.00100.17           C
    ATOM   5593  NE2 GLN B 516     -32.857  42.370  65.458  1.00 97.82           N
    ATOM   5594  OE1 GLN B 516     -31.602  43.139  67.150  1.00100.13           O
    ATOM   5595  H   GLN B 516     -27.544  42.770  65.577  1.00  0.00           H
    ATOM   5596  HA  GLN B 516     -28.797  44.096  63.273  1.00  0.00           H
    ATOM   5597  HB2 GLN B 516     -29.316  44.189  66.274  1.00  0.00           H
    ATOM   5598  HB3 GLN B 516     -30.285  45.017  65.080  1.00  0.00           H
    ATOM   5599  HG2 GLN B 516     -30.844  42.823  63.992  1.00  0.00           H
    ATOM   5600  HG3 GLN B 516     -29.906  41.975  65.201  1.00  0.00           H
    ATOM   5601 HE21 GLN B 516     -32.915  42.093  64.488  1.00  0.00           H
    ATOM   5602 HE22 GLN B 516     -33.676  42.303  66.044  1.00  0.00           H
    ATOM   5603  N   SER B 517     -26.888  45.727  65.424  1.00 86.44           N
    ATOM   5604  CA  SER B 517     -26.120  46.937  65.706  1.00 84.30           C
    ATOM   5605  C   SER B 517     -24.625  46.600  65.791  1.00 81.74           C
    ATOM   5606  O   SER B 517     -24.257  45.434  65.938  1.00 82.45           O
    ATOM   5607  CB  SER B 517     -26.671  47.600  66.991  1.00 85.64           C
    ATOM   5608  OG  SER B 517     -26.286  46.938  68.183  1.00 85.44           O
    ATOM   5609  H   SER B 517     -26.714  44.944  66.040  1.00  0.00           H
    ATOM   5610  HA  SER B 517     -26.243  47.649  64.888  1.00  0.00           H
    ATOM   5611  HB2 SER B 517     -26.306  48.626  67.056  1.00  0.00           H
    ATOM   5612  HB3 SER B 517     -27.759  47.665  66.954  1.00  0.00           H
    ATOM   5613  HG  SER B 517     -26.697  46.069  68.197  1.00  0.00           H
    ATOM   5614  N   ILE B 518     -23.784  47.641  65.698  1.00 78.54           N
    ATOM   5615  CA  ILE B 518     -22.329  47.514  65.785  1.00 76.63           C
    ATOM   5616  C   ILE B 518     -21.824  47.139  67.200  1.00 74.87           C
    ATOM   5617  O   ILE B 518     -20.737  46.579  67.310  1.00 74.35           O
    ATOM   5618  CB  ILE B 518     -21.626  48.805  65.291  1.00 75.51           C
    ATOM   5619  CG1 ILE B 518     -20.132  48.560  64.982  1.00 75.06           C
    ATOM   5620  CG2 ILE B 518     -21.832  50.014  66.223  1.00 77.31           C
    ATOM   5621  CD1 ILE B 518     -19.443  49.703  64.234  1.00 74.21           C
    ATOM   5622  H   ILE B 518     -24.145  48.573  65.555  1.00  0.00           H
    ATOM   5623  HA  ILE B 518     -22.044  46.705  65.111  1.00  0.00           H
    ATOM   5624  HB  ILE B 518     -22.101  49.063  64.345  1.00  0.00           H
    ATOM   5625 HG12 ILE B 518     -19.585  48.380  65.904  1.00  0.00           H
    ATOM   5626 HG13 ILE B 518     -20.027  47.648  64.394  1.00  0.00           H
    ATOM   5627 HG21 ILE B 518     -21.535  50.928  65.716  1.00  0.00           H
    ATOM   5628 HG22 ILE B 518     -22.878  50.132  66.503  1.00  0.00           H
    ATOM   5629 HG23 ILE B 518     -21.245  49.932  67.139  1.00  0.00           H
    ATOM   5630 HD11 ILE B 518     -18.680  49.323  63.554  1.00  0.00           H
    ATOM   5631 HD12 ILE B 518     -20.149  50.291  63.648  1.00  0.00           H
    ATOM   5632 HD13 ILE B 518     -18.951  50.369  64.937  1.00  0.00           H
    ATOM   5633  N   LYS B 519     -22.651  47.355  68.242  1.00 73.68           N
    ATOM   5634  CA  LYS B 519     -22.415  46.919  69.627  1.00 71.13           C
    ATOM   5635  C   LYS B 519     -22.528  45.392  69.834  1.00 66.93           C
    ATOM   5636  O   LYS B 519     -22.516  44.938  70.978  1.00 65.81           O
    ATOM   5637  CB  LYS B 519     -23.413  47.622  70.567  1.00 70.86           C
    ATOM   5638  CG  LYS B 519     -23.424  49.152  70.461  1.00 74.53           C
    ATOM   5639  CD  LYS B 519     -24.213  49.830  71.593  1.00 81.35           C
    ATOM   5640  CE  LYS B 519     -25.639  49.308  71.838  1.00 83.94           C
    ATOM   5641  NZ  LYS B 519     -26.489  49.432  70.642  1.00 88.29           N1+
    ATOM   5642  H   LYS B 519     -23.549  47.780  68.066  1.00  0.00           H
    ATOM   5643  HA  LYS B 519     -21.403  47.213  69.909  1.00  0.00           H
    ATOM   5644  HB2 LYS B 519     -24.418  47.244  70.373  1.00  0.00           H
    ATOM   5645  HB3 LYS B 519     -23.179  47.355  71.600  1.00  0.00           H
    ATOM   5646  HG2 LYS B 519     -22.399  49.525  70.474  1.00  0.00           H
    ATOM   5647  HG3 LYS B 519     -23.840  49.454  69.499  1.00  0.00           H
    ATOM   5648  HD2 LYS B 519     -23.644  49.753  72.521  1.00  0.00           H
    ATOM   5649  HD3 LYS B 519     -24.270  50.892  71.358  1.00  0.00           H
    ATOM   5650  HE2 LYS B 519     -25.623  48.266  72.159  1.00  0.00           H
    ATOM   5651  HE3 LYS B 519     -26.099  49.874  72.648  1.00  0.00           H
    ATOM   5652  HZ1 LYS B 519     -26.106  48.870  69.895  1.00  0.00           H
    ATOM   5653  HZ2 LYS B 519     -27.420  49.105  70.858  1.00  0.00           H
    ATOM   5654  HZ3 LYS B 519     -26.531  50.399  70.354  1.00  0.00           H
    ATOM   5655  N   GLU B 520     -22.674  44.634  68.741  1.00 64.49           N
    ATOM   5656  CA  GLU B 520     -22.879  43.193  68.722  1.00 65.70           C
    ATOM   5657  C   GLU B 520     -21.787  42.487  67.897  1.00 64.97           C
    ATOM   5658  O   GLU B 520     -21.732  41.258  67.917  1.00 57.63           O
    ATOM   5659  CB  GLU B 520     -24.308  42.977  68.194  1.00 71.26           C
    ATOM   5660  CG  GLU B 520     -24.892  41.571  68.396  1.00 72.89           C
    ATOM   5661  CD  GLU B 520     -26.392  41.515  68.090  1.00 72.97           C
    ATOM   5662  OE1 GLU B 520     -27.011  40.524  68.530  1.00 74.54           O
    ATOM   5663  OE2 GLU B 520     -26.902  42.451  67.430  1.00 69.58           O1-
    ATOM   5664  H   GLU B 520     -22.704  45.087  67.839  1.00  0.00           H
    ATOM   5665  HA  GLU B 520     -22.819  42.792  69.735  1.00  0.00           H
    ATOM   5666  HB2 GLU B 520     -24.970  43.704  68.670  1.00  0.00           H
    ATOM   5667  HB3 GLU B 520     -24.333  43.204  67.129  1.00  0.00           H
    ATOM   5668  HG2 GLU B 520     -24.382  40.846  67.761  1.00  0.00           H
    ATOM   5669  HG3 GLU B 520     -24.740  41.254  69.429  1.00  0.00           H
    ATOM   5670  N   THR B 521     -20.914  43.260  67.224  1.00 64.47           N
    ATOM   5671  CA  THR B 521     -19.710  42.767  66.560  1.00 68.78           C
    ATOM   5672  C   THR B 521     -18.586  42.541  67.603  1.00 70.23           C
    ATOM   5673  O   THR B 521     -18.504  43.293  68.578  1.00 68.82           O
    ATOM   5674  CB  THR B 521     -19.226  43.759  65.466  1.00 70.05           C
    ATOM   5675  CG2 THR B 521     -20.316  44.171  64.463  1.00 73.16           C
    ATOM   5676  OG1 THR B 521     -18.590  44.911  65.972  1.00 72.32           O
    ATOM   5677  H   THR B 521     -20.992  44.265  67.279  1.00  0.00           H
    ATOM   5678  HA  THR B 521     -19.959  41.828  66.075  1.00  0.00           H
    ATOM   5679  HB  THR B 521     -18.456  43.249  64.889  1.00  0.00           H
    ATOM   5680 HG21 THR B 521     -19.949  44.903  63.746  1.00  0.00           H
    ATOM   5681 HG22 THR B 521     -20.665  43.311  63.893  1.00  0.00           H
    ATOM   5682 HG23 THR B 521     -21.179  44.610  64.960  1.00  0.00           H
    ATOM   5683  HG1 THR B 521     -18.274  45.428  65.226  1.00  0.00           H
    ATOM   5684  N   PRO B 522     -17.761  41.487  67.402  1.00 69.49           N
    ATOM   5685  CA  PRO B 522     -16.726  41.085  68.366  1.00 66.34           C
    ATOM   5686  C   PRO B 522     -15.570  42.081  68.551  1.00 61.49           C
    ATOM   5687  O   PRO B 522     -14.933  42.043  69.600  1.00 63.69           O
    ATOM   5688  CB  PRO B 522     -16.201  39.753  67.804  1.00 69.32           C
    ATOM   5689  CG  PRO B 522     -16.458  39.833  66.313  1.00 70.34           C
    ATOM   5690  CD  PRO B 522     -17.779  40.582  66.257  1.00 68.03           C
    ATOM   5691  HA  PRO B 522     -17.192  40.927  69.340  1.00  0.00           H
    ATOM   5692  HB2 PRO B 522     -15.146  39.586  68.014  1.00  0.00           H
    ATOM   5693  HB3 PRO B 522     -16.763  38.923  68.235  1.00  0.00           H
    ATOM   5694  HG2 PRO B 522     -15.677  40.426  65.835  1.00  0.00           H
    ATOM   5695  HG3 PRO B 522     -16.486  38.856  65.831  1.00  0.00           H
    ATOM   5696  HD2 PRO B 522     -17.876  41.117  65.321  1.00  0.00           H
    ATOM   5697  HD3 PRO B 522     -18.619  39.893  66.362  1.00  0.00           H
    ATOM   5698  N   CYS B 523     -15.309  42.924  67.541  1.00 57.58           N
    ATOM   5699  CA  CYS B 523     -14.174  43.838  67.507  1.00 57.03           C
    ATOM   5700  C   CYS B 523     -14.474  44.972  66.514  1.00 56.76           C
    ATOM   5701  O   CYS B 523     -14.749  44.709  65.342  1.00 56.02           O
    ATOM   5702  CB  CYS B 523     -12.833  43.101  67.257  1.00 58.52           C
    ATOM   5703  SG  CYS B 523     -12.787  42.150  65.707  1.00 65.86           S
    ATOM   5704  H   CYS B 523     -15.886  42.914  66.712  1.00  0.00           H
    ATOM   5705  HA  CYS B 523     -14.098  44.289  68.497  1.00  0.00           H
    ATOM   5706  HB2 CYS B 523     -12.011  43.811  67.219  1.00  0.00           H
    ATOM   5707  HB3 CYS B 523     -12.613  42.421  68.080  1.00  0.00           H
    ATOM   5708  HG  CYS B 523     -13.699  41.246  66.075  1.00  0.00           H
    ATOM   5709  N   TRP B 524     -14.437  46.216  67.016  1.00 55.86           N
    ATOM   5710  CA  TRP B 524     -14.750  47.428  66.258  1.00 55.58           C
    ATOM   5711  C   TRP B 524     -14.129  48.663  66.924  1.00 55.30           C
    ATOM   5712  O   TRP B 524     -13.640  48.574  68.046  1.00 58.78           O
    ATOM   5713  CB  TRP B 524     -16.281  47.567  66.077  1.00 51.48           C
    ATOM   5714  CG  TRP B 524     -17.131  47.804  67.292  1.00 45.72           C
    ATOM   5715  CD1 TRP B 524     -17.507  46.868  68.191  1.00 45.13           C
    ATOM   5716  CD2 TRP B 524     -17.713  49.060  67.757  1.00 41.84           C
    ATOM   5717  CE2 TRP B 524     -18.509  48.790  68.911  1.00 41.09           C
    ATOM   5718  CE3 TRP B 524     -17.670  50.399  67.309  1.00 40.16           C
    ATOM   5719  NE1 TRP B 524     -18.337  47.441  69.130  1.00 43.64           N
    ATOM   5720  CZ2 TRP B 524     -19.239  49.792  69.574  1.00 42.32           C
    ATOM   5721  CZ3 TRP B 524     -18.393  51.412  67.966  1.00 42.95           C
    ATOM   5722  CH2 TRP B 524     -19.182  51.112  69.091  1.00 39.81           C
    ATOM   5723  H   TRP B 524     -14.208  46.359  67.992  1.00  0.00           H
    ATOM   5724  HA  TRP B 524     -14.278  47.334  65.279  1.00  0.00           H
    ATOM   5725  HB2 TRP B 524     -16.486  48.391  65.394  1.00  0.00           H
    ATOM   5726  HB3 TRP B 524     -16.670  46.685  65.573  1.00  0.00           H
    ATOM   5727  HD1 TRP B 524     -17.223  45.826  68.149  1.00  0.00           H
    ATOM   5728  HE1 TRP B 524     -18.748  46.919  69.895  1.00  0.00           H
    ATOM   5729  HE3 TRP B 524     -17.086  50.642  66.437  1.00  0.00           H
    ATOM   5730  HZ2 TRP B 524     -19.829  49.551  70.446  1.00  0.00           H
    ATOM   5731  HZ3 TRP B 524     -18.353  52.423  67.596  1.00  0.00           H
    ATOM   5732  HH2 TRP B 524     -19.741  51.898  69.574  1.00  0.00           H
    ATOM   5733  N   ILE B 525     -14.156  49.794  66.206  1.00 57.28           N
    ATOM   5734  CA  ILE B 525     -13.612  51.081  66.637  1.00 58.16           C
    ATOM   5735  C   ILE B 525     -14.594  52.226  66.326  1.00 60.49           C
    ATOM   5736  O   ILE B 525     -15.322  52.155  65.336  1.00 58.05           O
    ATOM   5737  CB  ILE B 525     -12.251  51.395  65.940  1.00 54.84           C
    ATOM   5738  CG1 ILE B 525     -12.305  51.330  64.391  1.00 59.14           C
    ATOM   5739  CG2 ILE B 525     -11.119  50.514  66.498  1.00 48.47           C
    ATOM   5740  CD1 ILE B 525     -11.019  51.767  63.676  1.00 53.64           C
    ATOM   5741  H   ILE B 525     -14.563  49.782  65.279  1.00  0.00           H
    ATOM   5742  HA  ILE B 525     -13.468  51.056  67.715  1.00  0.00           H
    ATOM   5743  HB  ILE B 525     -11.987  52.421  66.200  1.00  0.00           H
    ATOM   5744 HG12 ILE B 525     -12.531  50.310  64.081  1.00  0.00           H
    ATOM   5745 HG13 ILE B 525     -13.117  51.948  64.013  1.00  0.00           H
    ATOM   5746 HG21 ILE B 525     -10.154  50.776  66.067  1.00  0.00           H
    ATOM   5747 HG22 ILE B 525     -11.014  50.645  67.572  1.00  0.00           H
    ATOM   5748 HG23 ILE B 525     -11.296  49.458  66.301  1.00  0.00           H
    ATOM   5749 HD11 ILE B 525     -11.246  52.167  62.688  1.00  0.00           H
    ATOM   5750 HD12 ILE B 525     -10.480  52.535  64.230  1.00  0.00           H
    ATOM   5751 HD13 ILE B 525     -10.353  50.919  63.527  1.00  0.00           H
    ATOM   5752  N   GLU B 526     -14.565  53.270  67.170  1.00 61.42           N
    ATOM   5753  CA  GLU B 526     -15.274  54.532  66.968  1.00 57.67           C
    ATOM   5754  C   GLU B 526     -14.228  55.648  66.887  1.00 54.44           C
    ATOM   5755  O   GLU B 526     -13.458  55.828  67.830  1.00 48.51           O
    ATOM   5756  CB  GLU B 526     -16.268  54.771  68.120  1.00 62.33           C
    ATOM   5757  CG  GLU B 526     -17.183  55.995  67.902  1.00 67.03           C
    ATOM   5758  CD  GLU B 526     -18.210  56.158  69.021  1.00 69.65           C
    ATOM   5759  OE1 GLU B 526     -17.791  56.097  70.195  1.00 71.42           O
    ATOM   5760  OE2 GLU B 526     -19.401  56.346  68.689  1.00 66.66           O1-
    ATOM   5761  H   GLU B 526     -13.952  53.237  67.975  1.00  0.00           H
    ATOM   5762  HA  GLU B 526     -15.838  54.503  66.034  1.00  0.00           H
    ATOM   5763  HB2 GLU B 526     -16.875  53.880  68.269  1.00  0.00           H
    ATOM   5764  HB3 GLU B 526     -15.712  54.902  69.049  1.00  0.00           H
    ATOM   5765  HG2 GLU B 526     -16.594  56.911  67.849  1.00  0.00           H
    ATOM   5766  HG3 GLU B 526     -17.702  55.908  66.948  1.00  0.00           H
    ATOM   5767  N   ILE B 527     -14.226  56.363  65.756  1.00 53.70           N
    ATOM   5768  CA  ILE B 527     -13.250  57.389  65.417  1.00 48.33           C
    ATOM   5769  C   ILE B 527     -13.972  58.751  65.426  1.00 46.83           C
    ATOM   5770  O   ILE B 527     -14.937  58.924  64.686  1.00 52.33           O
    ATOM   5771  CB  ILE B 527     -12.657  57.161  63.995  1.00 55.36           C
    ATOM   5772  CG1 ILE B 527     -12.122  55.728  63.756  1.00 60.31           C
    ATOM   5773  CG2 ILE B 527     -11.572  58.194  63.625  1.00 54.58           C
    ATOM   5774  CD1 ILE B 527     -12.315  55.266  62.307  1.00 58.70           C
    ATOM   5775  H   ILE B 527     -14.929  56.180  65.050  1.00  0.00           H
    ATOM   5776  HA  ILE B 527     -12.428  57.380  66.125  1.00  0.00           H
    ATOM   5777  HB  ILE B 527     -13.470  57.301  63.290  1.00  0.00           H
    ATOM   5778 HG12 ILE B 527     -11.071  55.660  64.033  1.00  0.00           H
    ATOM   5779 HG13 ILE B 527     -12.626  54.997  64.387  1.00  0.00           H
    ATOM   5780 HG21 ILE B 527     -11.198  58.038  62.614  1.00  0.00           H
    ATOM   5781 HG22 ILE B 527     -11.952  59.211  63.669  1.00  0.00           H
    ATOM   5782 HG23 ILE B 527     -10.722  58.131  64.305  1.00  0.00           H
    ATOM   5783 HD11 ILE B 527     -11.739  54.363  62.114  1.00  0.00           H
    ATOM   5784 HD12 ILE B 527     -13.362  55.036  62.108  1.00  0.00           H
    ATOM   5785 HD13 ILE B 527     -12.000  56.024  61.590  1.00  0.00           H
    ATOM   5786  N   HIS B 528     -13.486  59.692  66.248  1.00 45.05           N
    ATOM   5787  CA  HIS B 528     -13.984  61.067  66.327  1.00 48.82           C
    ATOM   5788  C   HIS B 528     -13.005  61.982  65.589  1.00 51.94           C
    ATOM   5789  O   HIS B 528     -11.837  61.998  65.966  1.00 50.88           O
    ATOM   5790  CB  HIS B 528     -14.065  61.507  67.802  1.00 56.08           C
    ATOM   5791  CG  HIS B 528     -15.168  60.884  68.615  1.00 63.54           C
    ATOM   5792  CD2 HIS B 528     -16.265  61.445  69.232  1.00 66.80           C
    ATOM   5793  ND1 HIS B 528     -15.222  59.533  68.911  1.00 65.93           N
    ATOM   5794  CE1 HIS B 528     -16.302  59.343  69.671  1.00 67.70           C
    ATOM   5795  NE2 HIS B 528     -16.987  60.457  69.902  1.00 70.49           N
    ATOM   5796  H   HIS B 528     -12.687  59.473  66.830  1.00  0.00           H
    ATOM   5797  HA  HIS B 528     -14.974  61.163  65.878  1.00  0.00           H
    ATOM   5798  HB2 HIS B 528     -13.120  61.312  68.311  1.00  0.00           H
    ATOM   5799  HB3 HIS B 528     -14.214  62.584  67.853  1.00  0.00           H
    ATOM   5800  HD1 HIS B 528     -14.560  58.825  68.624  1.00  0.00           H
    ATOM   5801  HD2 HIS B 528     -16.584  62.476  69.251  1.00  0.00           H
    ATOM   5802  HE1 HIS B 528     -16.583  58.382  70.071  1.00  0.00           H
    ATOM   5803  N   LEU B 529     -13.480  62.731  64.580  1.00 54.62           N
    ATOM   5804  CA  LEU B 529     -12.656  63.658  63.800  1.00 59.04           C
    ATOM   5805  C   LEU B 529     -12.642  65.032  64.476  1.00 62.44           C
    ATOM   5806  O   LEU B 529     -13.679  65.688  64.538  1.00 67.14           O
    ATOM   5807  CB  LEU B 529     -13.163  63.745  62.340  1.00 58.60           C
    ATOM   5808  CG  LEU B 529     -13.144  62.419  61.536  1.00 61.91           C
    ATOM   5809  CD1 LEU B 529     -11.929  61.524  61.837  1.00 67.00           C
    ATOM   5810  CD2 LEU B 529     -14.461  61.628  61.630  1.00 61.41           C
    ATOM   5811  H   LEU B 529     -14.462  62.687  64.335  1.00  0.00           H
    ATOM   5812  HA  LEU B 529     -11.624  63.305  63.777  1.00  0.00           H
    ATOM   5813  HB2 LEU B 529     -14.156  64.195  62.298  1.00  0.00           H
    ATOM   5814  HB3 LEU B 529     -12.512  64.448  61.818  1.00  0.00           H
    ATOM   5815  HG  LEU B 529     -13.065  62.717  60.490  1.00  0.00           H
    ATOM   5816 HD11 LEU B 529     -11.998  60.572  61.310  1.00  0.00           H
    ATOM   5817 HD12 LEU B 529     -11.001  62.012  61.540  1.00  0.00           H
    ATOM   5818 HD13 LEU B 529     -11.848  61.277  62.892  1.00  0.00           H
    ATOM   5819 HD21 LEU B 529     -14.755  61.264  60.647  1.00  0.00           H
    ATOM   5820 HD22 LEU B 529     -14.377  60.766  62.294  1.00  0.00           H
    ATOM   5821 HD23 LEU B 529     -15.282  62.244  61.985  1.00  0.00           H
    ATOM   5822  N   HIS B 530     -11.463  65.424  64.984  1.00 65.67           N
    ATOM   5823  CA  HIS B 530     -11.265  66.618  65.807  1.00 69.37           C
    ATOM   5824  C   HIS B 530     -11.550  67.949  65.112  1.00 72.24           C
    ATOM   5825  O   HIS B 530     -12.070  68.841  65.775  1.00 76.26           O
    ATOM   5826  CB  HIS B 530      -9.844  66.641  66.386  1.00 67.13           C
    ATOM   5827  CG  HIS B 530      -9.628  65.711  67.549  1.00 65.71           C
    ATOM   5828  CD2 HIS B 530     -10.285  64.572  67.960  1.00 60.40           C
    ATOM   5829  ND1 HIS B 530      -8.637  65.928  68.489  1.00 69.07           N
    ATOM   5830  CE1 HIS B 530      -8.725  64.949  69.390  1.00 71.19           C
    ATOM   5831  NE2 HIS B 530      -9.702  64.090  69.131  1.00 65.78           N
    ATOM   5832  H   HIS B 530     -10.651  64.835  64.857  1.00  0.00           H
    ATOM   5833  HA  HIS B 530     -11.967  66.553  66.639  1.00  0.00           H
    ATOM   5834  HB2 HIS B 530      -9.132  66.398  65.600  1.00  0.00           H
    ATOM   5835  HB3 HIS B 530      -9.584  67.644  66.729  1.00  0.00           H
    ATOM   5836  HD1 HIS B 530      -7.967  66.684  68.483  1.00  0.00           H
    ATOM   5837  HD2 HIS B 530     -11.129  64.069  67.513  1.00  0.00           H
    ATOM   5838  HE1 HIS B 530      -8.070  64.865  70.245  1.00  0.00           H
    ATOM   5839  N   ARG B 531     -11.229  68.074  63.813  1.00 74.20           N
    ATOM   5840  CA  ARG B 531     -11.504  69.301  63.065  1.00 71.22           C
    ATOM   5841  C   ARG B 531     -13.008  69.521  62.851  1.00 71.71           C
    ATOM   5842  O   ARG B 531     -13.467  70.652  62.985  1.00 69.37           O
    ATOM   5843  CB  ARG B 531     -10.722  69.314  61.741  1.00 75.98           C
    ATOM   5844  CG  ARG B 531     -10.776  70.654  60.982  1.00 80.25           C
    ATOM   5845  CD  ARG B 531     -10.228  71.835  61.790  1.00 84.85           C
    ATOM   5846  NE  ARG B 531     -10.012  73.026  60.960  1.00 88.18           N
    ATOM   5847  CZ  ARG B 531      -9.301  74.103  61.334  1.00 90.64           C
    ATOM   5848  NH1 ARG B 531      -8.700  74.160  62.530  1.00 93.18           N
    ATOM   5849  NH2 ARG B 531      -9.183  75.135  60.494  1.00 89.39           N1+
    ATOM   5850  H   ARG B 531     -10.799  67.309  63.313  1.00  0.00           H
    ATOM   5851  HA  ARG B 531     -11.138  70.125  63.679  1.00  0.00           H
    ATOM   5852  HB2 ARG B 531      -9.675  69.075  61.938  1.00  0.00           H
    ATOM   5853  HB3 ARG B 531     -11.090  68.520  61.089  1.00  0.00           H
    ATOM   5854  HG2 ARG B 531     -10.247  70.569  60.041  1.00  0.00           H
    ATOM   5855  HG3 ARG B 531     -11.814  70.867  60.728  1.00  0.00           H
    ATOM   5856  HD2 ARG B 531     -10.963  72.149  62.532  1.00  0.00           H
    ATOM   5857  HD3 ARG B 531      -9.331  71.536  62.328  1.00  0.00           H
    ATOM   5858  HE  ARG B 531     -10.463  73.042  60.054  1.00  0.00           H
    ATOM   5859 HH11 ARG B 531      -8.810  73.405  63.193  1.00  0.00           H
    ATOM   5860 HH12 ARG B 531      -8.135  74.962  62.777  1.00  0.00           H
    ATOM   5861 HH21 ARG B 531      -9.587  75.073  59.562  1.00  0.00           H
    ATOM   5862 HH22 ARG B 531      -8.671  75.965  60.758  1.00  0.00           H
    ATOM   5863  N   ALA B 532     -13.751  68.432  62.590  1.00 69.20           N
    ATOM   5864  CA  ALA B 532     -15.209  68.432  62.542  1.00 63.93           C
    ATOM   5865  C   ALA B 532     -15.839  68.781  63.902  1.00 64.19           C
    ATOM   5866  O   ALA B 532     -16.836  69.496  63.919  1.00 64.61           O
    ATOM   5867  CB  ALA B 532     -15.693  67.072  62.025  1.00 62.54           C
    ATOM   5868  H   ALA B 532     -13.302  67.531  62.514  1.00  0.00           H
    ATOM   5869  HA  ALA B 532     -15.517  69.194  61.823  1.00  0.00           H
    ATOM   5870  HB1 ALA B 532     -16.765  67.084  61.831  1.00  0.00           H
    ATOM   5871  HB2 ALA B 532     -15.189  66.818  61.093  1.00  0.00           H
    ATOM   5872  HB3 ALA B 532     -15.493  66.270  62.734  1.00  0.00           H
    ATOM   5873  N   LEU B 533     -15.228  68.325  65.011  1.00 62.89           N
    ATOM   5874  CA  LEU B 533     -15.626  68.678  66.376  1.00 62.80           C
    ATOM   5875  C   LEU B 533     -15.336  70.154  66.728  1.00 62.12           C
    ATOM   5876  O   LEU B 533     -16.147  70.750  67.431  1.00 65.99           O
    ATOM   5877  CB  LEU B 533     -14.953  67.733  67.394  1.00 63.30           C
    ATOM   5878  CG  LEU B 533     -15.399  66.258  67.369  1.00 56.65           C
    ATOM   5879  CD1 LEU B 533     -14.426  65.392  68.191  1.00 50.84           C
    ATOM   5880  CD2 LEU B 533     -16.849  66.059  67.845  1.00 55.01           C
    ATOM   5881  H   LEU B 533     -14.419  67.725  64.922  1.00  0.00           H
    ATOM   5882  HA  LEU B 533     -16.705  68.544  66.447  1.00  0.00           H
    ATOM   5883  HB2 LEU B 533     -13.875  67.785  67.252  1.00  0.00           H
    ATOM   5884  HB3 LEU B 533     -15.123  68.093  68.404  1.00  0.00           H
    ATOM   5885  HG  LEU B 533     -15.360  65.903  66.343  1.00  0.00           H
    ATOM   5886 HD11 LEU B 533     -14.948  64.709  68.862  1.00  0.00           H
    ATOM   5887 HD12 LEU B 533     -13.798  64.795  67.530  1.00  0.00           H
    ATOM   5888 HD13 LEU B 533     -13.757  65.994  68.806  1.00  0.00           H
    ATOM   5889 HD21 LEU B 533     -17.157  65.020  67.725  1.00  0.00           H
    ATOM   5890 HD22 LEU B 533     -16.956  66.310  68.899  1.00  0.00           H
    ATOM   5891 HD23 LEU B 533     -17.556  66.669  67.285  1.00  0.00           H
    ATOM   5892  N   GLN B 534     -14.231  70.737  66.225  1.00 62.78           N
    ATOM   5893  CA  GLN B 534     -13.903  72.161  66.382  1.00 67.08           C
    ATOM   5894  C   GLN B 534     -14.886  73.090  65.653  1.00 73.69           C
    ATOM   5895  O   GLN B 534     -15.315  74.089  66.229  1.00 79.44           O
    ATOM   5896  CB  GLN B 534     -12.478  72.453  65.885  1.00 66.14           C
    ATOM   5897  CG  GLN B 534     -11.358  72.000  66.833  1.00 66.11           C
    ATOM   5898  CD  GLN B 534      -9.983  72.367  66.275  1.00 66.40           C
    ATOM   5899  NE2 GLN B 534      -9.068  72.790  67.148  1.00 67.82           N
    ATOM   5900  OE1 GLN B 534      -9.739  72.258  65.074  1.00 71.85           O
    ATOM   5901  H   GLN B 534     -13.580  70.184  65.683  1.00  0.00           H
    ATOM   5902  HA  GLN B 534     -13.954  72.402  67.446  1.00  0.00           H
    ATOM   5903  HB2 GLN B 534     -12.338  72.014  64.897  1.00  0.00           H
    ATOM   5904  HB3 GLN B 534     -12.368  73.530  65.752  1.00  0.00           H
    ATOM   5905  HG2 GLN B 534     -11.494  72.471  67.808  1.00  0.00           H
    ATOM   5906  HG3 GLN B 534     -11.393  70.925  67.000  1.00  0.00           H
    ATOM   5907 HE21 GLN B 534      -9.294  72.859  68.131  1.00  0.00           H
    ATOM   5908 HE22 GLN B 534      -8.147  73.057  66.831  1.00  0.00           H
    ATOM   5909  N   LEU B 535     -15.245  72.716  64.412  1.00 75.75           N
    ATOM   5910  CA  LEU B 535     -16.287  73.365  63.617  1.00 77.93           C
    ATOM   5911  C   LEU B 535     -17.669  73.231  64.270  1.00 79.99           C
    ATOM   5912  O   LEU B 535     -18.456  74.172  64.197  1.00 82.56           O
    ATOM   5913  CB  LEU B 535     -16.312  72.750  62.202  1.00 74.37           C
    ATOM   5914  CG  LEU B 535     -15.073  73.063  61.341  1.00 75.68           C
    ATOM   5915  CD1 LEU B 535     -15.098  72.260  60.028  1.00 76.96           C
    ATOM   5916  CD2 LEU B 535     -14.881  74.568  61.100  1.00 76.19           C
    ATOM   5917  H   LEU B 535     -14.822  71.893  64.004  1.00  0.00           H
    ATOM   5918  HA  LEU B 535     -16.057  74.430  63.555  1.00  0.00           H
    ATOM   5919  HB2 LEU B 535     -16.415  71.669  62.296  1.00  0.00           H
    ATOM   5920  HB3 LEU B 535     -17.200  73.086  61.667  1.00  0.00           H
    ATOM   5921  HG  LEU B 535     -14.191  72.735  61.887  1.00  0.00           H
    ATOM   5922 HD11 LEU B 535     -14.152  71.739  59.890  1.00  0.00           H
    ATOM   5923 HD12 LEU B 535     -15.886  71.506  60.020  1.00  0.00           H
    ATOM   5924 HD13 LEU B 535     -15.251  72.888  59.152  1.00  0.00           H
    ATOM   5925 HD21 LEU B 535     -14.530  74.782  60.091  1.00  0.00           H
    ATOM   5926 HD22 LEU B 535     -15.805  75.127  61.249  1.00  0.00           H
    ATOM   5927 HD23 LEU B 535     -14.141  74.970  61.793  1.00  0.00           H
    ATOM   5928  N   LEU B 536     -17.921  72.085  64.927  1.00 79.36           N
    ATOM   5929  CA  LEU B 536     -19.138  71.840  65.689  1.00 75.91           C
    ATOM   5930  C   LEU B 536     -19.226  72.688  66.958  1.00 73.57           C
    ATOM   5931  O   LEU B 536     -20.307  73.186  67.251  1.00 72.51           O
    ATOM   5932  CB  LEU B 536     -19.299  70.337  65.988  1.00 72.72           C
    ATOM   5933  CG  LEU B 536     -20.568  69.964  66.779  1.00 70.70           C
    ATOM   5934  CD1 LEU B 536     -21.847  70.549  66.164  1.00 67.93           C
    ATOM   5935  CD2 LEU B 536     -20.687  68.447  66.965  1.00 66.22           C
    ATOM   5936  H   LEU B 536     -17.236  71.342  64.924  1.00  0.00           H
    ATOM   5937  HA  LEU B 536     -19.957  72.155  65.045  1.00  0.00           H
    ATOM   5938  HB2 LEU B 536     -19.295  69.782  65.051  1.00  0.00           H
    ATOM   5939  HB3 LEU B 536     -18.434  69.982  66.542  1.00  0.00           H
    ATOM   5940  HG  LEU B 536     -20.465  70.387  67.775  1.00  0.00           H
    ATOM   5941 HD11 LEU B 536     -22.687  70.387  66.833  1.00  0.00           H
    ATOM   5942 HD12 LEU B 536     -21.819  71.623  66.010  1.00  0.00           H
    ATOM   5943 HD13 LEU B 536     -22.056  70.098  65.196  1.00  0.00           H
    ATOM   5944 HD21 LEU B 536     -21.120  68.205  67.935  1.00  0.00           H
    ATOM   5945 HD22 LEU B 536     -21.332  68.015  66.202  1.00  0.00           H
    ATOM   5946 HD23 LEU B 536     -19.719  67.949  66.903  1.00  0.00           H
    ATOM   5947  N   ASP B 537     -18.100  72.875  67.662  1.00 75.13           N
    ATOM   5948  CA  ASP B 537     -18.008  73.724  68.846  1.00 77.29           C
    ATOM   5949  C   ASP B 537     -18.456  75.163  68.527  1.00 78.37           C
    ATOM   5950  O   ASP B 537     -19.294  75.706  69.242  1.00 77.47           O
    ATOM   5951  CB  ASP B 537     -16.587  73.701  69.439  1.00 73.86           C
    ATOM   5952  CG  ASP B 537     -16.502  74.422  70.780  1.00 73.04           C
    ATOM   5953  OD1 ASP B 537     -15.711  75.385  70.870  1.00 68.30           O
    ATOM   5954  OD2 ASP B 537     -17.187  73.962  71.715  1.00 74.01           O1-
    ATOM   5955  H   ASP B 537     -17.245  72.419  67.369  1.00  0.00           H
    ATOM   5956  HA  ASP B 537     -18.700  73.310  69.583  1.00  0.00           H
    ATOM   5957  HB2 ASP B 537     -16.221  72.683  69.564  1.00  0.00           H
    ATOM   5958  HB3 ASP B 537     -15.884  74.173  68.752  1.00  0.00           H
    ATOM   5959  N   GLU B 538     -17.955  75.688  67.398  1.00 80.23           N
    ATOM   5960  CA  GLU B 538     -18.321  76.972  66.816  1.00 82.71           C
    ATOM   5961  C   GLU B 538     -19.817  77.067  66.443  1.00 79.96           C
    ATOM   5962  O   GLU B 538     -20.445  78.067  66.790  1.00 75.54           O
    ATOM   5963  CB  GLU B 538     -17.360  77.238  65.639  1.00 85.37           C
    ATOM   5964  CG  GLU B 538     -17.518  78.600  64.940  1.00 90.01           C
    ATOM   5965  CD  GLU B 538     -16.517  78.788  63.794  1.00 92.58           C
    ATOM   5966  OE1 GLU B 538     -15.347  78.378  63.961  1.00 91.12           O
    ATOM   5967  OE2 GLU B 538     -16.941  79.349  62.761  1.00 95.11           O1-
    ATOM   5968  H   GLU B 538     -17.278  75.149  66.874  1.00  0.00           H
    ATOM   5969  HA  GLU B 538     -18.130  77.731  67.578  1.00  0.00           H
    ATOM   5970  HB2 GLU B 538     -16.338  77.151  66.014  1.00  0.00           H
    ATOM   5971  HB3 GLU B 538     -17.463  76.448  64.897  1.00  0.00           H
    ATOM   5972  HG2 GLU B 538     -18.528  78.708  64.541  1.00  0.00           H
    ATOM   5973  HG3 GLU B 538     -17.374  79.408  65.658  1.00  0.00           H
    ATOM   5974  N   VAL B 539     -20.375  76.021  65.802  1.00 79.52           N
    ATOM   5975  CA  VAL B 539     -21.797  75.941  65.440  1.00 79.98           C
    ATOM   5976  C   VAL B 539     -22.744  75.842  66.658  1.00 84.51           C
    ATOM   5977  O   VAL B 539     -23.801  76.465  66.619  1.00 87.78           O
    ATOM   5978  CB  VAL B 539     -22.102  74.784  64.436  1.00 76.25           C
    ATOM   5979  CG1 VAL B 539     -23.603  74.461  64.251  1.00 78.20           C
    ATOM   5980  CG2 VAL B 539     -21.493  75.079  63.052  1.00 71.73           C
    ATOM   5981  H   VAL B 539     -19.801  75.228  65.547  1.00  0.00           H
    ATOM   5982  HA  VAL B 539     -22.036  76.877  64.932  1.00  0.00           H
    ATOM   5983  HB  VAL B 539     -21.619  73.880  64.808  1.00  0.00           H
    ATOM   5984 HG11 VAL B 539     -23.772  73.790  63.414  1.00  0.00           H
    ATOM   5985 HG12 VAL B 539     -24.034  73.978  65.129  1.00  0.00           H
    ATOM   5986 HG13 VAL B 539     -24.174  75.367  64.046  1.00  0.00           H
    ATOM   5987 HG21 VAL B 539     -21.585  74.220  62.387  1.00  0.00           H
    ATOM   5988 HG22 VAL B 539     -21.995  75.921  62.574  1.00  0.00           H
    ATOM   5989 HG23 VAL B 539     -20.436  75.330  63.114  1.00  0.00           H
    ATOM   5990  N   LEU B 540     -22.357  75.107  67.717  1.00 86.94           N
    ATOM   5991  CA  LEU B 540     -23.132  74.946  68.956  1.00 88.18           C
    ATOM   5992  C   LEU B 540     -23.370  76.264  69.717  1.00 90.91           C
    ATOM   5993  O   LEU B 540     -24.422  76.405  70.341  1.00 90.82           O
    ATOM   5994  CB  LEU B 540     -22.440  73.923  69.885  1.00 88.07           C
    ATOM   5995  CG  LEU B 540     -22.621  72.445  69.481  1.00 88.02           C
    ATOM   5996  CD1 LEU B 540     -21.711  71.537  70.328  1.00 88.20           C
    ATOM   5997  CD2 LEU B 540     -24.090  71.983  69.534  1.00 85.88           C
    ATOM   5998  H   LEU B 540     -21.481  74.601  67.672  1.00  0.00           H
    ATOM   5999  HA  LEU B 540     -24.117  74.569  68.676  1.00  0.00           H
    ATOM   6000  HB2 LEU B 540     -21.379  74.166  69.947  1.00  0.00           H
    ATOM   6001  HB3 LEU B 540     -22.822  74.032  70.901  1.00  0.00           H
    ATOM   6002  HG  LEU B 540     -22.299  72.339  68.448  1.00  0.00           H
    ATOM   6003 HD11 LEU B 540     -21.425  70.648  69.769  1.00  0.00           H
    ATOM   6004 HD12 LEU B 540     -20.793  72.040  70.630  1.00  0.00           H
    ATOM   6005 HD13 LEU B 540     -22.212  71.204  71.237  1.00  0.00           H
    ATOM   6006 HD21 LEU B 540     -24.203  71.018  70.028  1.00  0.00           H
    ATOM   6007 HD22 LEU B 540     -24.724  72.694  70.067  1.00  0.00           H
    ATOM   6008 HD23 LEU B 540     -24.496  71.880  68.528  1.00  0.00           H
    ATOM   6009  N   HIS B 541     -22.422  77.212  69.618  1.00 94.99           N
    ATOM   6010  CA  HIS B 541     -22.550  78.572  70.152  1.00 98.55           C
    ATOM   6011  C   HIS B 541     -23.544  79.442  69.355  1.00101.39           C
    ATOM   6012  O   HIS B 541     -24.087  80.390  69.922  1.00102.67           O
    ATOM   6013  CB  HIS B 541     -21.161  79.244  70.217  1.00 98.67           C
    ATOM   6014  CG  HIS B 541     -20.228  78.720  71.286  1.00100.37           C
    ATOM   6015  CD2 HIS B 541     -19.551  79.363  72.299  1.00100.10           C
    ATOM   6016  ND1 HIS B 541     -19.871  77.389  71.404  1.00100.46           N
    ATOM   6017  CE1 HIS B 541     -19.013  77.287  72.420  1.00101.27           C
    ATOM   6018  NE2 HIS B 541     -18.779  78.448  73.018  1.00102.29           N
    ATOM   6019  H   HIS B 541     -21.589  77.022  69.078  1.00  0.00           H
    ATOM   6020  HA  HIS B 541     -22.939  78.507  71.171  1.00  0.00           H
    ATOM   6021  HB2 HIS B 541     -20.656  79.159  69.255  1.00  0.00           H
    ATOM   6022  HB3 HIS B 541     -21.276  80.314  70.397  1.00  0.00           H
    ATOM   6023  HD1 HIS B 541     -20.132  76.636  70.780  1.00  0.00           H
    ATOM   6024  HD2 HIS B 541     -19.551  80.412  72.558  1.00  0.00           H
    ATOM   6025  HE1 HIS B 541     -18.548  76.357  72.715  1.00  0.00           H
    ATOM   6026  N   THR B 542     -23.780  79.090  68.078  1.00102.16           N
    ATOM   6027  CA  THR B 542     -24.728  79.735  67.162  1.00102.15           C
    ATOM   6028  C   THR B 542     -26.051  78.946  67.039  1.00103.94           C
    ATOM   6029  O   THR B 542     -26.753  79.069  66.035  1.00106.29           O
    ATOM   6030  CB  THR B 542     -24.109  79.964  65.752  1.00102.52           C
    ATOM   6031  CG2 THR B 542     -22.733  80.642  65.784  1.00101.60           C
    ATOM   6032  OG1 THR B 542     -24.058  78.802  64.944  1.00102.34           O
    ATOM   6033  H   THR B 542     -23.305  78.279  67.704  1.00  0.00           H
    ATOM   6034  HA  THR B 542     -24.996  80.716  67.558  1.00  0.00           H
    ATOM   6035  HB  THR B 542     -24.768  80.650  65.218  1.00  0.00           H
    ATOM   6036 HG21 THR B 542     -22.372  80.844  64.775  1.00  0.00           H
    ATOM   6037 HG22 THR B 542     -22.781  81.593  66.314  1.00  0.00           H
    ATOM   6038 HG23 THR B 542     -21.987  80.026  66.282  1.00  0.00           H
    ATOM   6039  HG1 THR B 542     -23.615  78.104  65.435  1.00  0.00           H
    ATOM   6040  N   MET B 543     -26.418  78.221  68.104  1.00106.47           N
    ATOM   6041  CA  MET B 543     -27.757  77.687  68.320  1.00109.02           C
    ATOM   6042  C   MET B 543     -28.289  78.382  69.586  1.00112.07           C
    ATOM   6043  O   MET B 543     -28.173  77.767  70.647  1.00111.97           O
    ATOM   6044  CB  MET B 543     -27.686  76.144  68.453  1.00106.25           C
    ATOM   6045  CG  MET B 543     -27.197  75.452  67.171  1.00106.32           C
    ATOM   6046  SD  MET B 543     -27.100  73.642  67.230  1.00106.80           S
    ATOM   6047  CE  MET B 543     -28.861  73.217  67.161  1.00104.26           C
    ATOM   6048  H   MET B 543     -25.785  78.144  68.889  1.00  0.00           H
    ATOM   6049  HA  MET B 543     -28.427  77.908  67.488  1.00  0.00           H
    ATOM   6050  HB2 MET B 543     -27.022  75.863  69.273  1.00  0.00           H
    ATOM   6051  HB3 MET B 543     -28.672  75.756  68.715  1.00  0.00           H
    ATOM   6052  HG2 MET B 543     -27.842  75.737  66.341  1.00  0.00           H
    ATOM   6053  HG3 MET B 543     -26.204  75.809  66.916  1.00  0.00           H
    ATOM   6054  HE1 MET B 543     -29.051  72.533  66.334  1.00  0.00           H
    ATOM   6055  HE2 MET B 543     -29.169  72.735  68.089  1.00  0.00           H
    ATOM   6056  HE3 MET B 543     -29.470  74.108  67.021  1.00  0.00           H
    ATOM   6057  N   PRO B 544     -28.783  79.654  69.472  1.00116.95           N
    ATOM   6058  CA  PRO B 544     -29.277  80.491  70.593  1.00120.51           C
    ATOM   6059  C   PRO B 544     -30.095  79.699  71.601  1.00124.14           C
    ATOM   6060  O   PRO B 544     -31.156  79.211  71.210  1.00123.04           O
    ATOM   6061  CB  PRO B 544     -30.131  81.576  69.913  1.00120.00           C
    ATOM   6062  CG  PRO B 544     -29.448  81.783  68.585  1.00118.85           C
    ATOM   6063  CD  PRO B 544     -28.988  80.380  68.215  1.00118.29           C
    ATOM   6064  HA  PRO B 544     -28.409  80.949  71.070  1.00  0.00           H
    ATOM   6065  HB2 PRO B 544     -31.154  81.241  69.731  1.00  0.00           H
    ATOM   6066  HB3 PRO B 544     -30.187  82.496  70.496  1.00  0.00           H
    ATOM   6067  HG2 PRO B 544     -30.085  82.244  67.829  1.00  0.00           H
    ATOM   6068  HG3 PRO B 544     -28.577  82.424  68.727  1.00  0.00           H
    ATOM   6069  HD2 PRO B 544     -29.757  79.859  67.651  1.00  0.00           H
    ATOM   6070  HD3 PRO B 544     -28.098  80.467  67.605  1.00  0.00           H
    ATOM   6071  N   ILE B 545     -29.486  79.481  72.789  1.00127.48           N
    ATOM   6072  CA  ILE B 545     -29.742  78.351  73.694  1.00130.68           C
    ATOM   6073  C   ILE B 545     -31.234  78.050  73.792  1.00133.13           C
    ATOM   6074  O   ILE B 545     -31.626  77.057  73.185  1.00133.75           O
    ATOM   6075  CB  ILE B 545     -29.071  78.461  75.102  1.00131.02           C
    ATOM   6076  CG1 ILE B 545     -27.535  78.271  75.038  1.00130.68           C
    ATOM   6077  CG2 ILE B 545     -29.624  77.434  76.124  1.00128.39           C
    ATOM   6078  CD1 ILE B 545     -26.777  79.168  74.050  1.00129.26           C
    ATOM   6079  H   ILE B 545     -28.621  79.966  72.973  1.00  0.00           H
    ATOM   6080  HA  ILE B 545     -29.288  77.488  73.204  1.00  0.00           H
    ATOM   6081  HB  ILE B 545     -29.256  79.460  75.501  1.00  0.00           H
    ATOM   6082 HG12 ILE B 545     -27.113  78.425  76.032  1.00  0.00           H
    ATOM   6083 HG13 ILE B 545     -27.320  77.232  74.784  1.00  0.00           H
    ATOM   6084 HG21 ILE B 545     -28.987  77.351  77.002  1.00  0.00           H
    ATOM   6085 HG22 ILE B 545     -30.621  77.694  76.480  1.00  0.00           H
    ATOM   6086 HG23 ILE B 545     -29.674  76.435  75.688  1.00  0.00           H
    ATOM   6087 HD11 ILE B 545     -25.714  79.170  74.281  1.00  0.00           H
    ATOM   6088 HD12 ILE B 545     -26.873  78.802  73.027  1.00  0.00           H
    ATOM   6089 HD13 ILE B 545     -27.133  80.197  74.083  1.00  0.00           H
    ATOM   6090  N   ALA B 546     -31.992  79.007  74.370  1.00134.59           N
    ATOM   6091  CA  ALA B 546     -33.424  79.312  74.205  1.00135.95           C
    ATOM   6092  C   ALA B 546     -34.382  78.138  73.909  1.00137.44           C
    ATOM   6093  O   ALA B 546     -35.418  78.335  73.275  1.00138.24           O
    ATOM   6094  CB  ALA B 546     -33.579  80.460  73.188  1.00132.78           C
    ATOM   6095  H   ALA B 546     -31.485  79.748  74.831  1.00  0.00           H
    ATOM   6096  HA  ALA B 546     -33.741  79.698  75.174  1.00  0.00           H
    ATOM   6097  HB1 ALA B 546     -34.585  80.879  73.212  1.00  0.00           H
    ATOM   6098  HB2 ALA B 546     -32.885  81.274  73.403  1.00  0.00           H
    ATOM   6099  HB3 ALA B 546     -33.391  80.122  72.168  1.00  0.00           H
    ATOM   6100  N   ASP B 547     -34.015  76.946  74.399  1.00136.30           N
    ATOM   6101  CA  ASP B 547     -34.760  75.705  74.326  1.00135.73           C
    ATOM   6102  C   ASP B 547     -35.431  75.641  75.704  1.00136.22           C
    ATOM   6103  O   ASP B 547     -34.715  75.696  76.706  1.00137.29           O
    ATOM   6104  CB  ASP B 547     -33.879  74.460  74.000  1.00133.15           C
    ATOM   6105  CG  ASP B 547     -32.556  74.236  74.757  1.00131.48           C
    ATOM   6106  OD1 ASP B 547     -32.623  73.972  75.976  1.00130.02           O
    ATOM   6107  OD2 ASP B 547     -31.504  74.185  74.083  1.00129.61           O1-
    ATOM   6108  H   ASP B 547     -33.132  76.879  74.887  1.00  0.00           H
    ATOM   6109  HA  ASP B 547     -35.478  75.782  73.512  1.00  0.00           H
    ATOM   6110  HB2 ASP B 547     -34.470  73.548  74.082  1.00  0.00           H
    ATOM   6111  HB3 ASP B 547     -33.627  74.514  72.939  1.00  0.00           H
    ATOM   6112  N   PRO B 548     -36.780  75.690  75.755  1.00135.75           N
    ATOM   6113  CA  PRO B 548     -37.492  75.891  77.025  1.00136.70           C
    ATOM   6114  C   PRO B 548     -37.567  74.619  77.891  1.00137.18           C
    ATOM   6115  O   PRO B 548     -38.648  74.079  78.118  1.00136.21           O
    ATOM   6116  CB  PRO B 548     -38.849  76.458  76.580  1.00135.42           C
    ATOM   6117  CG  PRO B 548     -39.094  75.808  75.230  1.00135.24           C
    ATOM   6118  CD  PRO B 548     -37.701  75.754  74.617  1.00134.35           C
    ATOM   6119  HA  PRO B 548     -36.995  76.657  77.626  1.00  0.00           H
    ATOM   6120  HB2 PRO B 548     -39.664  76.288  77.284  1.00  0.00           H
    ATOM   6121  HB3 PRO B 548     -38.761  77.538  76.449  1.00  0.00           H
    ATOM   6122  HG2 PRO B 548     -39.469  74.794  75.378  1.00  0.00           H
    ATOM   6123  HG3 PRO B 548     -39.816  76.345  74.613  1.00  0.00           H
    ATOM   6124  HD2 PRO B 548     -37.594  74.904  73.941  1.00  0.00           H
    ATOM   6125  HD3 PRO B 548     -37.514  76.672  74.057  1.00  0.00           H
    ATOM   6126  N   GLN B 549     -36.391  74.233  78.412  1.00138.42           N
    ATOM   6127  CA  GLN B 549     -36.125  73.372  79.569  1.00139.60           C
    ATOM   6128  C   GLN B 549     -36.788  71.950  79.475  1.00145.00           C
    ATOM   6129  O   GLN B 549     -36.646  71.389  78.363  1.00146.16           O
    ATOM   6130  CB  GLN B 549     -36.431  74.139  80.896  1.00138.77           C
    ATOM   6131  CG  GLN B 549     -36.074  75.655  80.971  1.00137.64           C
    ATOM   6132  CD  GLN B 549     -34.688  76.104  80.471  1.00139.49           C
    ATOM   6133  NE2 GLN B 549     -34.407  77.403  80.594  1.00139.93           N
    ATOM   6134  OE1 GLN B 549     -33.881  75.321  79.978  1.00141.90           O
    ATOM   6135  OXT GLN B 549     -37.327  71.520  80.521  1.00 67.70           O1-
    ATOM   6136  H   GLN B 549     -35.570  74.726  78.081  1.00  0.00           H
    ATOM   6137  HA  GLN B 549     -35.059  73.162  79.522  1.00  0.00           H
    ATOM   6138  HB2 GLN B 549     -37.495  74.062  81.124  1.00  0.00           H
    ATOM   6139  HB3 GLN B 549     -35.922  73.625  81.713  1.00  0.00           H
    ATOM   6140  HG2 GLN B 549     -36.813  76.213  80.395  1.00  0.00           H
    ATOM   6141  HG3 GLN B 549     -36.191  75.992  82.001  1.00  0.00           H
    ATOM   6142 HE21 GLN B 549     -35.079  78.039  80.996  1.00  0.00           H
    ATOM   6143 HE22 GLN B 549     -33.509  77.752  80.290  1.00  0.00           H
  extra_rotamers: ""
ligand:
  residue_configurations: []
  extra_templates: |2
        0    0    2

    This is a remark line
    molecule.res
    SEP   INT  0
    CORRECT     OMIT DU   BEG
      0.0000
       1  DUMM  DU    M    0  -1  -2     0.000      .0        .0      .00000
       2  DUMM  DU    M    1   0  -1     1.449      .0        .0      .00000
       3  DUMM  DU    M    2   1   0     1.523   111.21       .0      .00000
       4  O     OS    M    3   2   1     1.540   111.208  -180.000 -0.305000
       5  C     CM    M    4   3   2     1.231    98.269   -87.835  0.242000
       6  CA    CT    M    5   4   3     1.541   119.173  -143.749 -0.101000
       7  N     DU    S    6   5   4     1.466   114.198   159.797 -0.316000
       8  H     H     E    7   6   5     1.011   115.100    98.370  0.224000
       9  HA    H1    E    6   5   4     1.084   105.732    42.190  0.049000
      10  CB    CT    M    6   5   4     1.543   109.616   -75.404  0.029000
      11  HB2   H1    E   10   6   5     1.091   112.138    56.381  0.053000
      12  HB3   H1    E   10   6   5     1.091   110.388   175.504  0.053000
      13  OG    OS    M   10   6   5     1.419   107.431   -64.478 -0.714000
      14  P     P     M   13  10   6     1.750   122.426  -165.589  1.548500
      15  O2P   O2    E   14  13  10     1.537   105.532   179.220 -0.920833
      16  O3P   O2    E   14  13  10     1.537   106.769   -60.104 -0.920833
      17  O1P   O2    M   14  13  10     1.538   105.805    59.470 -0.920833


    LOOP

    IMPROPER
       CA    O    C    O

    DONE
  extra_templates_coordinates: |
    SEP 14
    N    -17.631  36.875  57.242
    CA   -18.205  38.188  57.553
    C    -19.536  38.069  58.335
    O    -20.143  39.101  58.626
    CB   -18.393  39.013  56.266
    OG   -19.183  38.258  55.366
    P    -19.233  38.645  53.658
    O1P  -17.789  38.519  53.137
    O2P  -20.195  37.624  53.030
    O3P  -19.750  40.091  53.568
    H    -18.089  36.382  56.489
    HA   -17.514  38.750  58.180
    HB2  -18.860  39.983  56.452
    HB3  -17.420  39.220  55.817
  coordinates: |
    ATOM      1  N   GLU A 270      14.574  50.541  72.439  1.00 83.90           N1+
    ATOM      2  CA  GLU A 270      14.001  50.158  73.771  1.00 82.88           C
    ATOM      3  C   GLU A 270      12.853  51.179  74.014  1.00 79.44           C
    ATOM      4  O   GLU A 270      13.071  52.375  73.802  1.00 78.81           O
    ATOM      5  CB  GLU A 270      15.118  50.213  74.836  1.00 87.46           C
    ATOM      6  CG  GLU A 270      14.689  49.822  76.263  1.00 94.07           C
    ATOM      7  CD  GLU A 270      15.876  49.831  77.228  1.00100.14           C
    ATOM      8  OE1 GLU A 270      16.551  50.882  77.298  1.00104.05           O
    ATOM      9  OE2 GLU A 270      16.094  48.782  77.872  1.00102.43           O1-
    ATOM     10  H1  GLU A 270      15.352  49.931  72.230  1.00  0.00           H
    ATOM     11  H2  GLU A 270      13.865  50.456  71.725  1.00  0.00           H
    ATOM     12  H3  GLU A 270      14.899  51.497  72.496  1.00  0.00           H
    ATOM     13  HA  GLU A 270      13.675  49.131  73.620  1.00  0.00           H
    ATOM     14  HB2 GLU A 270      15.934  49.556  74.529  1.00  0.00           H
    ATOM     15  HB3 GLU A 270      15.545  51.219  74.861  1.00  0.00           H
    ATOM     16  HG2 GLU A 270      13.934  50.512  76.641  1.00  0.00           H
    ATOM     17  HG3 GLU A 270      14.233  48.830  76.257  1.00  0.00           H
    ATOM     18  N   PRO A 271      11.652  50.712  74.442  1.00 77.90           N
    ATOM     19  CA  PRO A 271      10.515  51.600  74.782  1.00 77.15           C
    ATOM     20  C   PRO A 271      10.797  52.614  75.905  1.00 77.31           C
    ATOM     21  O   PRO A 271      11.540  52.299  76.836  1.00 77.70           O
    ATOM     22  CB  PRO A 271       9.406  50.625  75.209  1.00 76.34           C
    ATOM     23  CG  PRO A 271       9.729  49.337  74.489  1.00 74.01           C
    ATOM     24  CD  PRO A 271      11.249  49.305  74.537  1.00 75.86           C
    ATOM     25  HA  PRO A 271      10.217  52.119  73.869  1.00  0.00           H
    ATOM     26  HB2 PRO A 271       9.455  50.436  76.282  1.00  0.00           H
    ATOM     27  HB3 PRO A 271       8.404  50.991  74.981  1.00  0.00           H
    ATOM     28  HG2 PRO A 271       9.261  48.466  74.942  1.00  0.00           H
    ATOM     29  HG3 PRO A 271       9.388  49.404  73.456  1.00  0.00           H
    ATOM     30  HD2 PRO A 271      11.601  48.901  75.488  1.00  0.00           H
    ATOM     31  HD3 PRO A 271      11.620  48.672  73.733  1.00  0.00           H
    ATOM     32  N   ALA A 272      10.162  53.796  75.802  1.00 76.19           N
    ATOM     33  CA  ALA A 272      10.195  54.861  76.813  1.00 76.66           C
    ATOM     34  C   ALA A 272       9.648  54.420  78.182  1.00 72.03           C
    ATOM     35  O   ALA A 272      10.236  54.767  79.205  1.00 73.43           O
    ATOM     36  CB  ALA A 272       9.408  56.075  76.296  1.00 75.35           C
    ATOM     37  H   ALA A 272       9.578  53.972  74.997  1.00  0.00           H
    ATOM     38  HA  ALA A 272      11.237  55.156  76.947  1.00  0.00           H
    ATOM     39  HB1 ALA A 272       9.425  56.893  77.017  1.00  0.00           H
    ATOM     40  HB2 ALA A 272       9.836  56.450  75.366  1.00  0.00           H
    ATOM     41  HB3 ALA A 272       8.363  55.826  76.103  1.00  0.00           H
    ATOM     42  N   PHE A 273       8.562  53.632  78.152  1.00 63.70           N
    ATOM     43  CA  PHE A 273       7.975  52.980  79.315  1.00 55.53           C
    ATOM     44  C   PHE A 273       8.206  51.479  79.130  1.00 53.20           C
    ATOM     45  O   PHE A 273       7.773  50.938  78.113  1.00 51.11           O
    ATOM     46  CB  PHE A 273       6.462  53.285  79.380  1.00 53.92           C
    ATOM     47  CG  PHE A 273       6.084  54.753  79.285  1.00 56.21           C
    ATOM     48  CD1 PHE A 273       5.740  55.318  78.039  1.00 51.65           C
    ATOM     49  CD2 PHE A 273       6.192  55.597  80.408  1.00 59.76           C
    ATOM     50  CE1 PHE A 273       5.445  56.672  77.948  1.00 52.41           C
    ATOM     51  CE2 PHE A 273       5.883  56.945  80.302  1.00 61.56           C
    ATOM     52  CZ  PHE A 273       5.508  57.479  79.077  1.00 56.35           C
    ATOM     53  H   PHE A 273       8.160  53.382  77.261  1.00  0.00           H
    ATOM     54  HA  PHE A 273       8.459  53.326  80.226  1.00  0.00           H
    ATOM     55  HB2 PHE A 273       5.923  52.751  78.595  1.00  0.00           H
    ATOM     56  HB3 PHE A 273       6.066  52.900  80.321  1.00  0.00           H
    ATOM     57  HD1 PHE A 273       5.696  54.700  77.154  1.00  0.00           H
    ATOM     58  HD2 PHE A 273       6.485  55.192  81.359  1.00  0.00           H
    ATOM     59  HE1 PHE A 273       5.168  57.098  76.996  1.00  0.00           H
    ATOM     60  HE2 PHE A 273       5.936  57.580  81.173  1.00  0.00           H
    ATOM     61  HZ  PHE A 273       5.272  58.531  78.999  1.00  0.00           H
    ATOM     62  N   TRP A 274       8.868  50.826  80.099  1.00 50.27           N
    ATOM     63  CA  TRP A 274       9.135  49.383  80.052  1.00 47.07           C
    ATOM     64  C   TRP A 274       7.893  48.525  80.360  1.00 48.97           C
    ATOM     65  O   TRP A 274       7.878  47.354  79.979  1.00 52.69           O
    ATOM     66  CB  TRP A 274      10.315  49.031  80.977  1.00 43.91           C
    ATOM     67  CG  TRP A 274      10.109  49.209  82.452  1.00 47.28           C
    ATOM     68  CD1 TRP A 274      10.304  50.359  83.133  1.00 48.48           C
    ATOM     69  CD2 TRP A 274       9.622  48.240  83.432  1.00 42.19           C
    ATOM     70  CE2 TRP A 274       9.584  48.873  84.711  1.00 41.93           C
    ATOM     71  CE3 TRP A 274       9.219  46.884  83.377  1.00 38.73           C
    ATOM     72  NE1 TRP A 274      10.003  50.162  84.463  1.00 47.63           N
    ATOM     73  CZ2 TRP A 274       9.191  48.194  85.877  1.00 40.64           C
    ATOM     74  CZ3 TRP A 274       8.801  46.198  84.535  1.00 38.95           C
    ATOM     75  CH2 TRP A 274       8.796  46.847  85.785  1.00 35.68           C
    ATOM     76  H   TRP A 274       9.191  51.324  80.918  1.00  0.00           H
    ATOM     77  HA  TRP A 274       9.447  49.137  79.034  1.00  0.00           H
    ATOM     78  HB2 TRP A 274      10.587  47.986  80.819  1.00  0.00           H
    ATOM     79  HB3 TRP A 274      11.194  49.605  80.681  1.00  0.00           H
    ATOM     80  HD1 TRP A 274      10.650  51.284  82.694  1.00  0.00           H
    ATOM     81  HE1 TRP A 274      10.101  50.902  85.149  1.00  0.00           H
    ATOM     82  HE3 TRP A 274       9.231  46.366  82.429  1.00  0.00           H
    ATOM     83  HZ2 TRP A 274       9.201  48.697  86.832  1.00  0.00           H
    ATOM     84  HZ3 TRP A 274       8.492  45.166  84.466  1.00  0.00           H
    ATOM     85  HH2 TRP A 274       8.494  46.305  86.667  1.00  0.00           H
    ATOM     86  N   CYS A 275       6.874  49.114  81.011  1.00 44.77           N
    ATOM     87  CA  CYS A 275       5.592  48.468  81.265  1.00 43.23           C
    ATOM     88  C   CYS A 275       4.466  49.499  81.424  1.00 41.27           C
    ATOM     89  O   CYS A 275       4.724  50.672  81.701  1.00 47.11           O
    ATOM     90  CB  CYS A 275       5.649  47.446  82.432  1.00 45.13           C
    ATOM     91  SG  CYS A 275       5.595  48.202  84.088  1.00 40.99           S
    ATOM     92  H   CYS A 275       6.952  50.078  81.304  1.00  0.00           H
    ATOM     93  HA  CYS A 275       5.368  47.928  80.347  1.00  0.00           H
    ATOM     94  HB2 CYS A 275       4.792  46.775  82.369  1.00  0.00           H
    ATOM     95  HB3 CYS A 275       6.528  46.808  82.353  1.00  0.00           H
    ATOM     96  HG  CYS A 275       4.336  48.642  84.003  1.00  0.00           H
    ATOM     97  N   SER A 276       3.229  49.002  81.274  1.00 37.75           N
    ATOM     98  CA  SER A 276       1.991  49.707  81.588  1.00 34.75           C
    ATOM     99  C   SER A 276       1.151  48.835  82.535  1.00 33.18           C
    ATOM    100  O   SER A 276       1.098  47.622  82.343  1.00 21.99           O
    ATOM    101  CB  SER A 276       1.268  50.090  80.280  1.00 28.07           C
    ATOM    102  OG  SER A 276       0.695  48.983  79.612  1.00 33.33           O
    ATOM    103  H   SER A 276       3.116  48.033  81.005  1.00  0.00           H
    ATOM    104  HA  SER A 276       2.249  50.623  82.114  1.00  0.00           H
    ATOM    105  HB2 SER A 276       0.475  50.805  80.495  1.00  0.00           H
    ATOM    106  HB3 SER A 276       1.956  50.593  79.600  1.00  0.00           H
    ATOM    107  HG  SER A 276      -0.039  48.652  80.136  1.00  0.00           H
    ATOM    108  N   ILE A 277       0.534  49.463  83.547  1.00 38.44           N
    ATOM    109  CA  ILE A 277      -0.162  48.788  84.641  1.00 40.58           C
    ATOM    110  C   ILE A 277      -1.598  49.332  84.730  1.00 41.37           C
    ATOM    111  O   ILE A 277      -1.785  50.539  84.884  1.00 35.53           O
    ATOM    112  CB  ILE A 277       0.530  49.029  86.019  1.00 45.71           C
    ATOM    113  CG1 ILE A 277       2.062  48.839  85.920  1.00 47.01           C
    ATOM    114  CG2 ILE A 277      -0.084  48.121  87.109  1.00 53.13           C
    ATOM    115  CD1 ILE A 277       2.800  48.867  87.261  1.00 51.35           C
    ATOM    116  H   ILE A 277       0.597  50.473  83.618  1.00  0.00           H
    ATOM    117  HA  ILE A 277      -0.201  47.714  84.462  1.00  0.00           H
    ATOM    118  HB  ILE A 277       0.364  50.061  86.330  1.00  0.00           H
    ATOM    119 HG12 ILE A 277       2.281  47.903  85.411  1.00  0.00           H
    ATOM    120 HG13 ILE A 277       2.491  49.620  85.290  1.00  0.00           H
    ATOM    121 HG21 ILE A 277       0.334  48.320  88.095  1.00  0.00           H
    ATOM    122 HG22 ILE A 277      -1.159  48.266  87.205  1.00  0.00           H
    ATOM    123 HG23 ILE A 277       0.087  47.069  86.880  1.00  0.00           H
    ATOM    124 HD11 ILE A 277       3.868  49.019  87.111  1.00  0.00           H
    ATOM    125 HD12 ILE A 277       2.424  49.669  87.895  1.00  0.00           H
    ATOM    126 HD13 ILE A 277       2.677  47.925  87.795  1.00  0.00           H
    ATOM    127  N   ALA A 278      -2.579  48.420  84.653  1.00 39.03           N
    ATOM    128  CA  ALA A 278      -4.004  48.718  84.758  1.00 31.93           C
    ATOM    129  C   ALA A 278      -4.555  48.110  86.051  1.00 31.59           C
    ATOM    130  O   ALA A 278      -4.432  46.902  86.236  1.00 35.89           O
    ATOM    131  CB  ALA A 278      -4.718  48.121  83.535  1.00 27.78           C
    ATOM    132  H   ALA A 278      -2.341  47.446  84.518  1.00  0.00           H
    ATOM    133  HA  ALA A 278      -4.184  49.795  84.765  1.00  0.00           H
    ATOM    134  HB1 ALA A 278      -5.798  48.239  83.608  1.00  0.00           H
    ATOM    135  HB2 ALA A 278      -4.397  48.619  82.619  1.00  0.00           H
    ATOM    136  HB3 ALA A 278      -4.507  47.056  83.423  1.00  0.00           H
    ATOM    137  N   TYR A 279      -5.163  48.950  86.905  1.00 30.41           N
    ATOM    138  CA  TYR A 279      -5.824  48.541  88.145  1.00 29.83           C
    ATOM    139  C   TYR A 279      -7.286  48.133  87.908  1.00 30.06           C
    ATOM    140  O   TYR A 279      -7.970  48.722  87.069  1.00 22.37           O
    ATOM    141  CB  TYR A 279      -5.674  49.670  89.187  1.00 29.88           C
    ATOM    142  CG  TYR A 279      -6.364  49.452  90.527  1.00 35.10           C
    ATOM    143  CD1 TYR A 279      -5.960  48.406  91.384  1.00 33.06           C
    ATOM    144  CD2 TYR A 279      -7.406  50.311  90.929  1.00 32.91           C
    ATOM    145  CE1 TYR A 279      -6.591  48.225  92.631  1.00 35.48           C
    ATOM    146  CE2 TYR A 279      -8.027  50.140  92.177  1.00 35.82           C
    ATOM    147  CZ  TYR A 279      -7.623  49.095  93.030  1.00 40.64           C
    ATOM    148  OH  TYR A 279      -8.238  48.924  94.235  1.00 35.20           O
    ATOM    149  H   TYR A 279      -5.213  49.933  86.680  1.00  0.00           H
    ATOM    150  HA  TYR A 279      -5.318  47.658  88.531  1.00  0.00           H
    ATOM    151  HB2 TYR A 279      -4.615  49.832  89.383  1.00  0.00           H
    ATOM    152  HB3 TYR A 279      -6.035  50.606  88.761  1.00  0.00           H
    ATOM    153  HD1 TYR A 279      -5.165  47.738  91.091  1.00  0.00           H
    ATOM    154  HD2 TYR A 279      -7.724  51.118  90.289  1.00  0.00           H
    ATOM    155  HE1 TYR A 279      -6.280  47.419  93.279  1.00  0.00           H
    ATOM    156  HE2 TYR A 279      -8.811  50.820  92.473  1.00  0.00           H
    ATOM    157  HH  TYR A 279      -9.080  49.396  94.270  1.00  0.00           H
    ATOM    158  N   TYR A 280      -7.712  47.116  88.669  1.00 27.90           N
    ATOM    159  CA  TYR A 280      -9.037  46.515  88.643  1.00 25.77           C
    ATOM    160  C   TYR A 280      -9.536  46.286  90.076  1.00 26.59           C
    ATOM    161  O   TYR A 280      -8.741  46.016  90.980  1.00 24.47           O
    ATOM    162  CB  TYR A 280      -8.975  45.152  87.924  1.00 29.80           C
    ATOM    163  CG  TYR A 280      -8.764  45.154  86.420  1.00 43.02           C
    ATOM    164  CD1 TYR A 280      -7.476  45.357  85.882  1.00 46.14           C
    ATOM    165  CD2 TYR A 280      -9.842  44.872  85.555  1.00 45.84           C
    ATOM    166  CE1 TYR A 280      -7.270  45.301  84.491  1.00 48.10           C
    ATOM    167  CE2 TYR A 280      -9.632  44.789  84.166  1.00 45.70           C
    ATOM    168  CZ  TYR A 280      -8.348  45.014  83.631  1.00 46.80           C
    ATOM    169  OH  TYR A 280      -8.147  44.943  82.283  1.00 47.94           O
    ATOM    170  H   TYR A 280      -7.070  46.689  89.327  1.00  0.00           H
    ATOM    171  HA  TYR A 280      -9.724  47.184  88.127  1.00  0.00           H
    ATOM    172  HB2 TYR A 280      -8.186  44.550  88.375  1.00  0.00           H
    ATOM    173  HB3 TYR A 280      -9.903  44.607  88.108  1.00  0.00           H
    ATOM    174  HD1 TYR A 280      -6.640  45.540  86.539  1.00  0.00           H
    ATOM    175  HD2 TYR A 280     -10.830  44.692  85.956  1.00  0.00           H
    ATOM    176  HE1 TYR A 280      -6.279  45.458  84.090  1.00  0.00           H
    ATOM    177  HE2 TYR A 280     -10.460  44.559  83.511  1.00  0.00           H
    ATOM    178  HH  TYR A 280      -7.216  44.919  82.043  1.00  0.00           H
    ATOM    179  N   GLU A 281     -10.868  46.321  90.219  1.00 31.49           N
    ATOM    180  CA  GLU A 281     -11.602  45.904  91.407  1.00 32.47           C
    ATOM    181  C   GLU A 281     -12.698  44.944  90.939  1.00 37.65           C
    ATOM    182  O   GLU A 281     -13.628  45.376  90.258  1.00 39.80           O
    ATOM    183  CB  GLU A 281     -12.182  47.129  92.145  1.00 23.93           C
    ATOM    184  CG  GLU A 281     -11.091  48.068  92.696  1.00 23.91           C
    ATOM    185  CD  GLU A 281     -11.616  49.308  93.422  1.00 29.18           C
    ATOM    186  OE1 GLU A 281     -10.758  50.082  93.893  1.00 26.90           O
    ATOM    187  OE2 GLU A 281     -12.850  49.487  93.515  1.00 38.89           O1-
    ATOM    188  H   GLU A 281     -11.440  46.543  89.413  1.00  0.00           H
    ATOM    189  HA  GLU A 281     -10.936  45.362  92.074  1.00  0.00           H
    ATOM    190  HB2 GLU A 281     -12.841  47.684  91.476  1.00  0.00           H
    ATOM    191  HB3 GLU A 281     -12.807  46.783  92.970  1.00  0.00           H
    ATOM    192  HG2 GLU A 281     -10.453  47.513  93.385  1.00  0.00           H
    ATOM    193  HG3 GLU A 281     -10.450  48.417  91.887  1.00  0.00           H
    ATOM    194  N   LEU A 282     -12.533  43.650  91.269  1.00 41.76           N
    ATOM    195  CA  LEU A 282     -13.302  42.519  90.733  1.00 41.52           C
    ATOM    196  C   LEU A 282     -13.114  42.434  89.201  1.00 41.48           C
    ATOM    197  O   LEU A 282     -11.977  42.400  88.743  1.00 48.16           O
    ATOM    198  CB  LEU A 282     -14.799  42.526  91.177  1.00 36.50           C
    ATOM    199  CG  LEU A 282     -15.091  42.409  92.689  1.00 33.86           C
    ATOM    200  CD1 LEU A 282     -14.476  41.161  93.352  1.00 28.24           C
    ATOM    201  CD2 LEU A 282     -14.797  43.701  93.457  1.00 42.12           C
    ATOM    202  H   LEU A 282     -11.746  43.392  91.848  1.00  0.00           H
    ATOM    203  HA  LEU A 282     -12.824  41.627  91.132  1.00  0.00           H
    ATOM    204  HB2 LEU A 282     -15.311  43.404  90.780  1.00  0.00           H
    ATOM    205  HB3 LEU A 282     -15.294  41.673  90.710  1.00  0.00           H
    ATOM    206  HG  LEU A 282     -16.172  42.278  92.750  1.00  0.00           H
    ATOM    207 HD11 LEU A 282     -15.229  40.614  93.919  1.00  0.00           H
    ATOM    208 HD12 LEU A 282     -14.070  40.465  92.619  1.00  0.00           H
    ATOM    209 HD13 LEU A 282     -13.674  41.410  94.044  1.00  0.00           H
    ATOM    210 HD21 LEU A 282     -15.472  43.803  94.306  1.00  0.00           H
    ATOM    211 HD22 LEU A 282     -13.775  43.714  93.828  1.00  0.00           H
    ATOM    212 HD23 LEU A 282     -14.929  44.586  92.833  1.00  0.00           H
    ATOM    213  N   ASN A 283     -14.209  42.406  88.431  1.00 45.33           N
    ATOM    214  CA  ASN A 283     -14.204  42.344  86.967  1.00 47.56           C
    ATOM    215  C   ASN A 283     -14.206  43.732  86.284  1.00 45.42           C
    ATOM    216  O   ASN A 283     -14.425  43.780  85.073  1.00 37.52           O
    ATOM    217  CB  ASN A 283     -15.380  41.451  86.492  1.00 48.25           C
    ATOM    218  CG  ASN A 283     -16.783  42.022  86.752  1.00 57.68           C
    ATOM    219  ND2 ASN A 283     -17.649  41.993  85.738  1.00 63.47           N
    ATOM    220  OD1 ASN A 283     -17.090  42.470  87.855  1.00 62.21           O
    ATOM    221  H   ASN A 283     -15.123  42.443  88.861  1.00  0.00           H
    ATOM    222  HA  ASN A 283     -13.287  41.863  86.622  1.00  0.00           H
    ATOM    223  HB2 ASN A 283     -15.270  41.247  85.425  1.00  0.00           H
    ATOM    224  HB3 ASN A 283     -15.327  40.480  86.983  1.00  0.00           H
    ATOM    225 HD21 ASN A 283     -17.369  41.655  84.827  1.00  0.00           H
    ATOM    226 HD22 ASN A 283     -18.584  42.350  85.865  1.00  0.00           H
    ATOM    227  N   GLN A 284     -13.994  44.825  87.040  1.00 44.42           N
    ATOM    228  CA  GLN A 284     -14.117  46.193  86.537  1.00 38.97           C
    ATOM    229  C   GLN A 284     -12.771  46.933  86.610  1.00 39.85           C
    ATOM    230  O   GLN A 284     -12.132  46.921  87.659  1.00 39.23           O
    ATOM    231  CB  GLN A 284     -15.207  46.917  87.358  1.00 41.06           C
    ATOM    232  CG  GLN A 284     -15.550  48.320  86.821  1.00 48.03           C
    ATOM    233  CD  GLN A 284     -16.584  49.038  87.683  1.00 52.30           C
    ATOM    234  NE2 GLN A 284     -16.142  50.067  88.402  1.00 44.96           N
    ATOM    235  OE1 GLN A 284     -17.761  48.684  87.688  1.00 53.92           O
    ATOM    236  H   GLN A 284     -13.805  44.727  88.028  1.00  0.00           H
    ATOM    237  HA  GLN A 284     -14.443  46.191  85.495  1.00  0.00           H
    ATOM    238  HB2 GLN A 284     -16.114  46.310  87.358  1.00  0.00           H
    ATOM    239  HB3 GLN A 284     -14.894  46.990  88.401  1.00  0.00           H
    ATOM    240  HG2 GLN A 284     -14.658  48.939  86.769  1.00  0.00           H
    ATOM    241  HG3 GLN A 284     -15.934  48.247  85.803  1.00  0.00           H
    ATOM    242 HE21 GLN A 284     -15.169  50.333  88.362  1.00  0.00           H
    ATOM    243 HE22 GLN A 284     -16.763  50.575  89.017  1.00  0.00           H
    ATOM    244  N   ARG A 285     -12.421  47.619  85.508  1.00 40.34           N
    ATOM    245  CA  ARG A 285     -11.345  48.613  85.397  1.00 40.97           C
    ATOM    246  C   ARG A 285     -11.628  49.828  86.293  1.00 39.59           C
    ATOM    247  O   ARG A 285     -12.694  50.431  86.170  1.00 38.00           O
    ATOM    248  CB  ARG A 285     -11.257  49.067  83.917  1.00 42.29           C
    ATOM    249  CG  ARG A 285     -10.277  48.229  83.079  1.00 45.70           C
    ATOM    250  CD  ARG A 285      -8.930  48.912  82.781  1.00 49.28           C
    ATOM    251  NE  ARG A 285      -8.292  49.479  83.981  1.00 48.07           N
    ATOM    252  CZ  ARG A 285      -7.429  50.511  83.985  1.00 50.55           C
    ATOM    253  NH1 ARG A 285      -7.049  51.115  82.850  1.00 53.40           N
    ATOM    254  NH2 ARG A 285      -6.943  50.956  85.149  1.00 53.78           N1+
    ATOM    255  H   ARG A 285     -13.017  47.551  84.696  1.00  0.00           H
    ATOM    256  HA  ARG A 285     -10.410  48.138  85.701  1.00  0.00           H
    ATOM    257  HB2 ARG A 285     -12.253  49.016  83.474  1.00  0.00           H
    ATOM    258  HB3 ARG A 285     -10.975  50.118  83.830  1.00  0.00           H
    ATOM    259  HG2 ARG A 285     -10.194  47.197  83.401  1.00  0.00           H
    ATOM    260  HG3 ARG A 285     -10.755  48.166  82.100  1.00  0.00           H
    ATOM    261  HD2 ARG A 285      -8.221  48.156  82.441  1.00  0.00           H
    ATOM    262  HD3 ARG A 285      -9.039  49.626  81.965  1.00  0.00           H
    ATOM    263  HE  ARG A 285      -8.477  49.008  84.856  1.00  0.00           H
    ATOM    264 HH11 ARG A 285      -7.405  50.795  81.961  1.00  0.00           H
    ATOM    265 HH12 ARG A 285      -6.401  51.891  82.867  1.00  0.00           H
    ATOM    266 HH21 ARG A 285      -7.270  50.552  86.018  1.00  0.00           H
    ATOM    267 HH22 ARG A 285      -6.335  51.763  85.177  1.00  0.00           H
    ATOM    268  N   VAL A 286     -10.654  50.178  87.145  1.00 39.12           N
    ATOM    269  CA  VAL A 286     -10.727  51.337  88.032  1.00 32.63           C
    ATOM    270  C   VAL A 286      -9.462  52.187  87.845  1.00 40.16           C
    ATOM    271  O   VAL A 286      -8.350  51.658  87.885  1.00 44.44           O
    ATOM    272  CB  VAL A 286     -10.885  50.907  89.519  1.00 34.91           C
    ATOM    273  CG1 VAL A 286     -10.750  52.060  90.535  1.00 25.57           C
    ATOM    274  CG2 VAL A 286     -12.234  50.194  89.731  1.00 33.94           C
    ATOM    275  H   VAL A 286      -9.795  49.645  87.185  1.00  0.00           H
    ATOM    276  HA  VAL A 286     -11.584  51.962  87.770  1.00  0.00           H
    ATOM    277  HB  VAL A 286     -10.105  50.180  89.754  1.00  0.00           H
    ATOM    278 HG11 VAL A 286     -10.985  51.728  91.546  1.00  0.00           H
    ATOM    279 HG12 VAL A 286      -9.737  52.458  90.565  1.00  0.00           H
    ATOM    280 HG13 VAL A 286     -11.427  52.878  90.292  1.00  0.00           H
    ATOM    281 HG21 VAL A 286     -12.397  49.953  90.778  1.00  0.00           H
    ATOM    282 HG22 VAL A 286     -13.066  50.820  89.408  1.00  0.00           H
    ATOM    283 HG23 VAL A 286     -12.286  49.257  89.177  1.00  0.00           H
    ATOM    284  N   GLY A 287      -9.673  53.496  87.631  1.00 42.73           N
    ATOM    285  CA  GLY A 287      -8.620  54.475  87.375  1.00 42.29           C
    ATOM    286  C   GLY A 287      -8.083  54.386  85.939  1.00 42.73           C
    ATOM    287  O   GLY A 287      -8.551  53.598  85.116  1.00 40.54           O
    ATOM    288  H   GLY A 287     -10.622  53.840  87.609  1.00  0.00           H
    ATOM    289  HA2 GLY A 287      -9.014  55.475  87.555  1.00  0.00           H
    ATOM    290  HA3 GLY A 287      -7.797  54.318  88.069  1.00  0.00           H
    ATOM    291  N   GLU A 288      -7.076  55.224  85.660  1.00 42.85           N
    ATOM    292  CA  GLU A 288      -6.333  55.265  84.399  1.00 43.53           C
    ATOM    293  C   GLU A 288      -5.231  54.193  84.375  1.00 46.25           C
    ATOM    294  O   GLU A 288      -4.828  53.695  85.430  1.00 46.76           O
    ATOM    295  CB  GLU A 288      -5.714  56.670  84.251  1.00 44.57           C
    ATOM    296  CG  GLU A 288      -6.761  57.794  84.152  1.00 52.07           C
    ATOM    297  CD  GLU A 288      -6.107  59.172  84.154  1.00 54.84           C
    ATOM    298  OE1 GLU A 288      -5.733  59.640  83.058  1.00 48.82           O
    ATOM    299  OE2 GLU A 288      -6.024  59.754  85.257  1.00 54.68           O1-
    ATOM    300  H   GLU A 288      -6.742  55.851  86.377  1.00  0.00           H
    ATOM    301  HA  GLU A 288      -7.013  55.085  83.563  1.00  0.00           H
    ATOM    302  HB2 GLU A 288      -5.047  56.865  85.093  1.00  0.00           H
    ATOM    303  HB3 GLU A 288      -5.084  56.701  83.359  1.00  0.00           H
    ATOM    304  HG2 GLU A 288      -7.352  57.679  83.242  1.00  0.00           H
    ATOM    305  HG3 GLU A 288      -7.464  57.749  84.984  1.00  0.00           H
    ATOM    306  N   THR A 289      -4.739  53.878  83.166  1.00 43.15           N
    ATOM    307  CA  THR A 289      -3.536  53.067  82.976  1.00 40.52           C
    ATOM    308  C   THR A 289      -2.285  53.901  83.307  1.00 38.29           C
    ATOM    309  O   THR A 289      -2.091  54.974  82.732  1.00 31.26           O
    ATOM    310  CB  THR A 289      -3.386  52.549  81.521  1.00 32.77           C
    ATOM    311  CG2 THR A 289      -2.164  51.645  81.281  1.00 31.69           C
    ATOM    312  OG1 THR A 289      -4.537  51.811  81.167  1.00 37.15           O
    ATOM    313  H   THR A 289      -5.137  54.293  82.331  1.00  0.00           H
    ATOM    314  HA  THR A 289      -3.588  52.204  83.642  1.00  0.00           H
    ATOM    315  HB  THR A 289      -3.328  53.395  80.835  1.00  0.00           H
    ATOM    316 HG21 THR A 289      -2.158  51.258  80.261  1.00  0.00           H
    ATOM    317 HG22 THR A 289      -1.227  52.183  81.420  1.00  0.00           H
    ATOM    318 HG23 THR A 289      -2.161  50.792  81.960  1.00  0.00           H
    ATOM    319  HG1 THR A 289      -5.242  52.445  80.992  1.00  0.00           H
    ATOM    320  N   PHE A 290      -1.464  53.377  84.224  1.00 41.61           N
    ATOM    321  CA  PHE A 290      -0.176  53.946  84.604  1.00 40.10           C
    ATOM    322  C   PHE A 290       0.901  53.423  83.646  1.00 40.06           C
    ATOM    323  O   PHE A 290       0.874  52.240  83.321  1.00 41.17           O
    ATOM    324  CB  PHE A 290       0.115  53.520  86.054  1.00 39.88           C
    ATOM    325  CG  PHE A 290       1.354  54.136  86.678  1.00 41.59           C
    ATOM    326  CD1 PHE A 290       2.629  53.567  86.466  1.00 43.64           C
    ATOM    327  CD2 PHE A 290       1.258  55.371  87.353  1.00 36.13           C
    ATOM    328  CE1 PHE A 290       3.759  54.200  86.964  1.00 43.62           C
    ATOM    329  CE2 PHE A 290       2.399  55.972  87.867  1.00 38.49           C
    ATOM    330  CZ  PHE A 290       3.643  55.386  87.677  1.00 39.66           C
    ATOM    331  H   PHE A 290      -1.687  52.479  84.634  1.00  0.00           H
    ATOM    332  HA  PHE A 290      -0.222  55.036  84.555  1.00  0.00           H
    ATOM    333  HB2 PHE A 290      -0.741  53.784  86.674  1.00  0.00           H
    ATOM    334  HB3 PHE A 290       0.202  52.435  86.117  1.00  0.00           H
    ATOM    335  HD1 PHE A 290       2.731  52.645  85.913  1.00  0.00           H
    ATOM    336  HD2 PHE A 290       0.296  55.844  87.490  1.00  0.00           H
    ATOM    337  HE1 PHE A 290       4.734  53.764  86.807  1.00  0.00           H
    ATOM    338  HE2 PHE A 290       2.322  56.903  88.410  1.00  0.00           H
    ATOM    339  HZ  PHE A 290       4.524  55.861  88.078  1.00  0.00           H
    ATOM    340  N   HIS A 291       1.840  54.290  83.246  1.00 42.30           N
    ATOM    341  CA  HIS A 291       2.963  53.932  82.380  1.00 44.03           C
    ATOM    342  C   HIS A 291       4.269  54.246  83.123  1.00 43.20           C
    ATOM    343  O   HIS A 291       4.502  55.400  83.483  1.00 36.03           O
    ATOM    344  CB  HIS A 291       2.852  54.681  81.038  1.00 42.32           C
    ATOM    345  CG  HIS A 291       1.630  54.347  80.216  1.00 41.17           C
    ATOM    346  CD2 HIS A 291       1.471  53.540  79.111  1.00 41.19           C
    ATOM    347  ND1 HIS A 291       0.373  54.873  80.472  1.00 41.82           N
    ATOM    348  CE1 HIS A 291      -0.460  54.373  79.557  1.00 46.88           C
    ATOM    349  NE2 HIS A 291       0.136  53.551  78.701  1.00 45.13           N
    ATOM    350  H   HIS A 291       1.823  55.244  83.579  1.00  0.00           H
    ATOM    351  HA  HIS A 291       2.958  52.865  82.154  1.00  0.00           H
    ATOM    352  HB2 HIS A 291       2.885  55.761  81.193  1.00  0.00           H
    ATOM    353  HB3 HIS A 291       3.711  54.430  80.422  1.00  0.00           H
    ATOM    354  HD1 HIS A 291       0.121  55.504  81.220  1.00  0.00           H
    ATOM    355  HD2 HIS A 291       2.209  52.952  78.586  1.00  0.00           H
    ATOM    356  HE1 HIS A 291      -1.515  54.605  79.521  1.00  0.00           H
    ATOM    357  N   ALA A 292       5.077  53.200  83.360  1.00 43.51           N
    ATOM    358  CA  ALA A 292       6.293  53.246  84.168  1.00 44.84           C
    ATOM    359  C   ALA A 292       7.523  53.591  83.322  1.00 43.45           C
    ATOM    360  O   ALA A 292       7.998  52.736  82.576  1.00 41.15           O
    ATOM    361  CB  ALA A 292       6.475  51.877  84.832  1.00 39.49           C
    ATOM    362  H   ALA A 292       4.828  52.289  82.996  1.00  0.00           H
    ATOM    363  HA  ALA A 292       6.185  53.992  84.957  1.00  0.00           H
    ATOM    364  HB1 ALA A 292       7.380  51.860  85.438  1.00  0.00           H
    ATOM    365  HB2 ALA A 292       5.629  51.646  85.476  1.00  0.00           H
    ATOM    366  HB3 ALA A 292       6.557  51.075  84.097  1.00  0.00           H
    ATOM    367  N   SER A 293       8.007  54.837  83.458  1.00 46.43           N
    ATOM    368  CA  SER A 293       9.188  55.351  82.760  1.00 49.74           C
    ATOM    369  C   SER A 293      10.504  54.973  83.461  1.00 54.99           C
    ATOM    370  O   SER A 293      11.472  54.629  82.784  1.00 54.91           O
    ATOM    371  CB  SER A 293       9.045  56.877  82.573  1.00 44.79           C
    ATOM    372  OG  SER A 293       9.067  57.588  83.798  1.00 54.58           O
    ATOM    373  H   SER A 293       7.551  55.486  84.084  1.00  0.00           H
    ATOM    374  HA  SER A 293       9.215  54.912  81.762  1.00  0.00           H
    ATOM    375  HB2 SER A 293       9.856  57.251  81.947  1.00  0.00           H
    ATOM    376  HB3 SER A 293       8.120  57.112  82.050  1.00  0.00           H
    ATOM    377  HG  SER A 293       9.019  58.527  83.604  1.00  0.00           H
    ATOM    378  N   PHE A 311       2.393  54.353  91.548  1.00 34.71           N
    ATOM    379  CA  PHE A 311       1.012  54.189  91.106  1.00 34.87           C
    ATOM    380  C   PHE A 311       0.123  54.034  92.346  1.00 30.93           C
    ATOM    381  O   PHE A 311      -0.139  52.920  92.800  1.00 29.36           O
    ATOM    382  CB  PHE A 311       0.896  53.008  90.122  1.00 38.26           C
    ATOM    383  CG  PHE A 311      -0.490  52.729  89.544  1.00 42.38           C
    ATOM    384  CD1 PHE A 311      -0.764  51.459  88.992  1.00 42.25           C
    ATOM    385  CD2 PHE A 311      -1.452  53.756  89.381  1.00 44.92           C
    ATOM    386  CE1 PHE A 311      -1.978  51.218  88.360  1.00 40.45           C
    ATOM    387  CE2 PHE A 311      -2.662  53.492  88.755  1.00 44.10           C
    ATOM    388  CZ  PHE A 311      -2.924  52.227  88.247  1.00 39.15           C
    ATOM    389  H   PHE A 311       2.852  53.517  91.886  1.00  0.00           H
    ATOM    390  HA  PHE A 311       0.707  55.090  90.571  1.00  0.00           H
    ATOM    391  HB2 PHE A 311       1.574  53.175  89.286  1.00  0.00           H
    ATOM    392  HB3 PHE A 311       1.252  52.103  90.609  1.00  0.00           H
    ATOM    393  HD1 PHE A 311      -0.030  50.669  89.054  1.00  0.00           H
    ATOM    394  HD2 PHE A 311      -1.270  54.758  89.737  1.00  0.00           H
    ATOM    395  HE1 PHE A 311      -2.187  50.242  87.947  1.00  0.00           H
    ATOM    396  HE2 PHE A 311      -3.396  54.277  88.647  1.00  0.00           H
    ATOM    397  HZ  PHE A 311      -3.864  52.032  87.751  1.00  0.00           H
    ATOM    398  N   LEU A 313      -2.965  53.777  94.299  1.00 28.89           N
    ATOM    399  CA  LEU A 313      -4.310  53.222  94.126  1.00 30.04           C
    ATOM    400  C   LEU A 313      -5.336  53.811  95.108  1.00 38.80           C
    ATOM    401  O   LEU A 313      -6.527  53.742  94.816  1.00 40.80           O
    ATOM    402  CB  LEU A 313      -4.289  51.684  94.304  1.00 28.52           C
    ATOM    403  CG  LEU A 313      -3.293  50.901  93.422  1.00 30.15           C
    ATOM    404  CD1 LEU A 313      -3.351  49.399  93.772  1.00 26.17           C
    ATOM    405  CD2 LEU A 313      -3.502  51.159  91.917  1.00 32.67           C
    ATOM    406  H   LEU A 313      -2.310  53.196  94.809  1.00  0.00           H
    ATOM    407  HA  LEU A 313      -4.650  53.453  93.117  1.00  0.00           H
    ATOM    408  HB2 LEU A 313      -4.072  51.451  95.348  1.00  0.00           H
    ATOM    409  HB3 LEU A 313      -5.293  51.297  94.119  1.00  0.00           H
    ATOM    410  HG  LEU A 313      -2.280  51.231  93.651  1.00  0.00           H
    ATOM    411 HD11 LEU A 313      -3.278  48.756  92.895  1.00  0.00           H
    ATOM    412 HD12 LEU A 313      -2.535  49.127  94.441  1.00  0.00           H
    ATOM    413 HD13 LEU A 313      -4.279  49.138  94.280  1.00  0.00           H
    ATOM    414 HD21 LEU A 313      -3.306  50.276  91.310  1.00  0.00           H
    ATOM    415 HD22 LEU A 313      -4.518  51.483  91.700  1.00  0.00           H
    ATOM    416 HD23 LEU A 313      -2.826  51.939  91.566  1.00  0.00           H
    ATOM    417  N   LEU A 316      -7.776  56.524  93.867  1.00 44.26           N
    ATOM    418  CA  LEU A 316      -8.719  56.062  92.845  1.00 37.97           C
    ATOM    419  C   LEU A 316     -10.088  55.772  93.470  1.00 38.78           C
    ATOM    420  O   LEU A 316     -10.162  55.088  94.491  1.00 36.54           O
    ATOM    421  CB  LEU A 316      -8.188  54.800  92.123  1.00 26.44           C
    ATOM    422  CG  LEU A 316      -6.872  55.000  91.338  1.00 33.40           C
    ATOM    423  CD1 LEU A 316      -6.312  53.660  90.835  1.00 34.68           C
    ATOM    424  CD2 LEU A 316      -7.018  56.015  90.187  1.00 34.22           C
    ATOM    425  H   LEU A 316      -7.011  55.902  94.084  1.00  0.00           H
    ATOM    426  HA  LEU A 316      -8.834  56.863  92.113  1.00  0.00           H
    ATOM    427  HB2 LEU A 316      -8.072  53.993  92.845  1.00  0.00           H
    ATOM    428  HB3 LEU A 316      -8.947  54.451  91.423  1.00  0.00           H
    ATOM    429  HG  LEU A 316      -6.126  55.372  92.038  1.00  0.00           H
    ATOM    430 HD11 LEU A 316      -5.225  53.702  90.749  1.00  0.00           H
    ATOM    431 HD12 LEU A 316      -6.547  52.851  91.526  1.00  0.00           H
    ATOM    432 HD13 LEU A 316      -6.708  53.385  89.859  1.00  0.00           H
    ATOM    433 HD21 LEU A 316      -6.456  55.712  89.303  1.00  0.00           H
    ATOM    434 HD22 LEU A 316      -8.059  56.143  89.888  1.00  0.00           H
    ATOM    435 HD23 LEU A 316      -6.644  56.996  90.469  1.00  0.00           H
    ATOM    436  N   ASN A 318     -14.201  54.394  92.955  1.00 33.98           N
    ATOM    437  CA  ASN A 318     -15.123  53.496  92.274  1.00 36.49           C
    ATOM    438  C   ASN A 318     -16.469  53.644  92.989  1.00 37.21           C
    ATOM    439  O   ASN A 318     -16.566  53.380  94.187  1.00 42.67           O
    ATOM    440  CB  ASN A 318     -14.548  52.058  92.256  1.00 35.79           C
    ATOM    441  CG  ASN A 318     -15.497  50.983  91.702  1.00 40.95           C
    ATOM    442  ND2 ASN A 318     -15.203  49.713  91.982  1.00 37.06           N
    ATOM    443  OD1 ASN A 318     -16.477  51.283  91.024  1.00 37.84           O
    ATOM    444  H   ASN A 318     -14.409  54.567  93.928  1.00  0.00           H
    ATOM    445  HA  ASN A 318     -15.223  53.840  91.242  1.00  0.00           H
    ATOM    446  HB2 ASN A 318     -13.635  52.040  91.661  1.00  0.00           H
    ATOM    447  HB3 ASN A 318     -14.250  51.774  93.264  1.00  0.00           H
    ATOM    448 HD21 ASN A 318     -14.380  49.497  92.540  1.00  0.00           H
    ATOM    449 HD22 ASN A 318     -15.785  48.962  91.642  1.00  0.00           H
    ATOM    450  N   ASN A 320     -19.163  51.838  92.342  1.00 41.62           N
    ATOM    451  CA  ASN A 320     -19.713  50.487  92.497  1.00 40.96           C
    ATOM    452  C   ASN A 320     -18.894  49.639  93.493  1.00 41.12           C
    ATOM    453  O   ASN A 320     -19.099  48.426  93.548  1.00 37.65           O
    ATOM    454  CB  ASN A 320     -19.776  49.795  91.110  1.00 39.93           C
    ATOM    455  CG  ASN A 320     -20.657  50.523  90.087  1.00 43.08           C
    ATOM    456  ND2 ASN A 320     -20.252  50.504  88.815  1.00 50.47           N
    ATOM    457  OD1 ASN A 320     -21.697  51.081  90.432  1.00 51.33           O
    ATOM    458  H   ASN A 320     -18.361  51.912  91.729  1.00  0.00           H
    ATOM    459  HA  ASN A 320     -20.725  50.541  92.902  1.00  0.00           H
    ATOM    460  HB2 ASN A 320     -18.769  49.677  90.705  1.00  0.00           H
    ATOM    461  HB3 ASN A 320     -20.187  48.790  91.210  1.00  0.00           H
    ATOM    462 HD21 ASN A 320     -19.409  50.010  88.550  1.00  0.00           H
    ATOM    463 HD22 ASN A 320     -20.788  50.978  88.103  1.00  0.00           H
    ATOM    464  N   TYR A 340      13.233  46.639  87.411  1.00 50.33           N
    ATOM    465  CA  TYR A 340      13.641  46.214  86.079  1.00 54.60           C
    ATOM    466  C   TYR A 340      15.170  46.303  85.999  1.00 60.93           C
    ATOM    467  O   TYR A 340      15.708  47.268  85.456  1.00 63.46           O
    ATOM    468  CB  TYR A 340      12.879  47.014  84.999  1.00 52.04           C
    ATOM    469  CG  TYR A 340      13.070  46.537  83.565  1.00 55.02           C
    ATOM    470  CD1 TYR A 340      12.926  45.170  83.240  1.00 56.70           C
    ATOM    471  CD2 TYR A 340      13.354  47.464  82.540  1.00 53.73           C
    ATOM    472  CE1 TYR A 340      13.055  44.733  81.909  1.00 54.63           C
    ATOM    473  CE2 TYR A 340      13.484  47.028  81.206  1.00 44.52           C
    ATOM    474  CZ  TYR A 340      13.327  45.664  80.889  1.00 51.68           C
    ATOM    475  OH  TYR A 340      13.435  45.247  79.595  1.00 51.70           O
    ATOM    476  H   TYR A 340      13.772  47.389  87.820  1.00  0.00           H
    ATOM    477  HA  TYR A 340      13.381  45.165  85.968  1.00  0.00           H
    ATOM    478  HB2 TYR A 340      11.811  46.943  85.202  1.00  0.00           H
    ATOM    479  HB3 TYR A 340      13.123  48.075  85.074  1.00  0.00           H
    ATOM    480  HD1 TYR A 340      12.710  44.443  84.006  1.00  0.00           H
    ATOM    481  HD2 TYR A 340      13.462  48.514  82.770  1.00  0.00           H
    ATOM    482  HE1 TYR A 340      12.937  43.685  81.675  1.00  0.00           H
    ATOM    483  HE2 TYR A 340      13.697  47.747  80.429  1.00  0.00           H
    ATOM    484  HH  TYR A 340      13.664  45.962  78.995  1.00  0.00           H
    ATOM    485  N   VAL A 345      12.589  41.315  85.616  1.00 43.99           N
    ATOM    486  CA  VAL A 345      11.633  42.030  86.462  1.00 38.50           C
    ATOM    487  C   VAL A 345      11.479  41.258  87.780  1.00 39.49           C
    ATOM    488  O   VAL A 345      11.391  40.031  87.750  1.00 41.42           O
    ATOM    489  CB  VAL A 345      10.228  42.130  85.797  1.00 39.78           C
    ATOM    490  CG1 VAL A 345       9.146  42.820  86.658  1.00 32.65           C
    ATOM    491  CG2 VAL A 345      10.304  42.828  84.428  1.00 34.70           C
    ATOM    492  H   VAL A 345      12.322  40.381  85.335  1.00  0.00           H
    ATOM    493  HA  VAL A 345      12.000  43.030  86.672  1.00  0.00           H
    ATOM    494  HB  VAL A 345       9.870  41.116  85.631  1.00  0.00           H
    ATOM    495 HG11 VAL A 345       8.207  42.899  86.109  1.00  0.00           H
    ATOM    496 HG12 VAL A 345       8.924  42.267  87.570  1.00  0.00           H
    ATOM    497 HG13 VAL A 345       9.440  43.829  86.940  1.00  0.00           H
    ATOM    498 HG21 VAL A 345       9.335  42.821  83.932  1.00  0.00           H
    ATOM    499 HG22 VAL A 345      10.610  43.868  84.538  1.00  0.00           H
    ATOM    500 HG23 VAL A 345      11.006  42.339  83.752  1.00  0.00           H
    ATOM    501  N   GLN A 358       0.599  35.742  89.047  1.00 42.52           N
    ATOM    502  CA  GLN A 358       0.468  34.789  87.954  1.00 48.41           C
    ATOM    503  C   GLN A 358       1.242  35.351  86.754  1.00 50.95           C
    ATOM    504  O   GLN A 358       0.699  36.159  85.999  1.00 51.96           O
    ATOM    505  CB  GLN A 358      -1.035  34.602  87.669  1.00 51.80           C
    ATOM    506  CG  GLN A 358      -1.371  33.546  86.600  1.00 51.37           C
    ATOM    507  CD  GLN A 358      -2.866  33.557  86.278  1.00 51.82           C
    ATOM    508  NE2 GLN A 358      -3.600  32.534  86.718  1.00 46.19           N
    ATOM    509  OE1 GLN A 358      -3.359  34.485  85.641  1.00 51.26           O
    ATOM    510  H   GLN A 358       0.227  36.666  88.864  1.00  0.00           H
    ATOM    511  HA  GLN A 358       0.895  33.822  88.223  1.00  0.00           H
    ATOM    512  HB2 GLN A 358      -1.524  34.314  88.599  1.00  0.00           H
    ATOM    513  HB3 GLN A 358      -1.470  35.564  87.390  1.00  0.00           H
    ATOM    514  HG2 GLN A 358      -0.831  33.745  85.673  1.00  0.00           H
    ATOM    515  HG3 GLN A 358      -1.058  32.556  86.932  1.00  0.00           H
    ATOM    516 HE21 GLN A 358      -3.175  31.780  87.243  1.00  0.00           H
    ATOM    517 HE22 GLN A 358      -4.586  32.496  86.512  1.00  0.00           H
    ATOM    518  N   SER A 359       2.502  34.913  86.620  1.00 56.31           N
    ATOM    519  CA  SER A 359       3.363  35.208  85.479  1.00 52.76           C
    ATOM    520  C   SER A 359       3.670  33.876  84.770  1.00 55.48           C
    ATOM    521  O   SER A 359       4.408  33.076  85.343  1.00 60.38           O
    ATOM    522  CB  SER A 359       4.663  35.863  85.978  1.00 51.82           C
    ATOM    523  OG  SER A 359       5.525  36.073  84.877  1.00 52.25           O
    ATOM    524  H   SER A 359       2.873  34.256  87.291  1.00  0.00           H
    ATOM    525  HA  SER A 359       2.895  35.926  84.806  1.00  0.00           H
    ATOM    526  HB2 SER A 359       4.465  36.821  86.455  1.00  0.00           H
    ATOM    527  HB3 SER A 359       5.167  35.250  86.723  1.00  0.00           H
    ATOM    528  HG  SER A 359       6.366  36.414  85.207  1.00  0.00           H
    ATOM    529  N   PRO A 360       3.143  33.661  83.542  1.00 54.62           N
    ATOM    530  CA  PRO A 360       3.475  32.476  82.724  1.00 55.81           C
    ATOM    531  C   PRO A 360       4.974  32.226  82.452  1.00 53.92           C
    ATOM    532  O   PRO A 360       5.362  31.062  82.363  1.00 56.48           O
    ATOM    533  CB  PRO A 360       2.687  32.697  81.424  1.00 54.64           C
    ATOM    534  CG  PRO A 360       1.516  33.571  81.833  1.00 54.40           C
    ATOM    535  CD  PRO A 360       2.130  34.487  82.881  1.00 55.41           C
    ATOM    536  HA  PRO A 360       3.061  31.610  83.244  1.00  0.00           H
    ATOM    537  HB2 PRO A 360       3.293  33.239  80.699  1.00  0.00           H
    ATOM    538  HB3 PRO A 360       2.368  31.764  80.959  1.00  0.00           H
    ATOM    539  HG2 PRO A 360       1.060  34.105  81.000  1.00  0.00           H
    ATOM    540  HG3 PRO A 360       0.747  32.951  82.296  1.00  0.00           H
    ATOM    541  HD2 PRO A 360       2.620  35.341  82.409  1.00  0.00           H
    ATOM    542  HD3 PRO A 360       1.360  34.857  83.559  1.00  0.00           H
    ATOM    543  N   PRO A 370       0.787  27.086  83.549  1.00 77.63           N
    ATOM    544  CA  PRO A 370       0.852  28.537  83.247  1.00 77.69           C
    ATOM    545  C   PRO A 370      -0.117  29.492  83.982  1.00 77.01           C
    ATOM    546  O   PRO A 370       0.163  30.689  84.017  1.00 80.14           O
    ATOM    547  CB  PRO A 370       0.649  28.598  81.727  1.00 78.20           C
    ATOM    548  CG  PRO A 370       1.214  27.284  81.222  1.00 80.25           C
    ATOM    549  CD  PRO A 370       0.829  26.296  82.316  1.00 78.32           C
    ATOM    550  HA  PRO A 370       1.871  28.853  83.479  1.00  0.00           H
    ATOM    551  HB2 PRO A 370      -0.414  28.654  81.484  1.00  0.00           H
    ATOM    552  HB3 PRO A 370       1.139  29.460  81.272  1.00  0.00           H
    ATOM    553  HG2 PRO A 370       0.849  27.002  80.234  1.00  0.00           H
    ATOM    554  HG3 PRO A 370       2.301  27.360  81.169  1.00  0.00           H
    ATOM    555  HD2 PRO A 370      -0.162  25.877  82.132  1.00  0.00           H
    ATOM    556  HD3 PRO A 370       1.555  25.484  82.342  1.00  0.00           H
    ATOM    557  N   ALA A 371      -1.202  28.956  84.569  1.00 71.41           N
    ATOM    558  CA  ALA A 371      -2.159  29.674  85.411  1.00 64.66           C
    ATOM    559  C   ALA A 371      -1.889  29.527  86.927  1.00 62.06           C
    ATOM    560  O   ALA A 371      -2.796  29.803  87.714  1.00 58.47           O
    ATOM    561  CB  ALA A 371      -3.577  29.202  85.045  1.00 64.04           C
    ATOM    562  H   ALA A 371      -1.362  27.960  84.489  1.00  0.00           H
    ATOM    563  HA  ALA A 371      -2.100  30.738  85.180  1.00  0.00           H
    ATOM    564  HB1 ALA A 371      -4.337  29.826  85.516  1.00  0.00           H
    ATOM    565  HB2 ALA A 371      -3.743  29.251  83.968  1.00  0.00           H
    ATOM    566  HB3 ALA A 371      -3.754  28.174  85.363  1.00  0.00           H
    ATOM    567  N   PHE A 385       9.857  38.012  85.821  1.00 52.22           N
    ATOM    568  CA  PHE A 385      10.077  37.893  84.377  1.00 51.86           C
    ATOM    569  C   PHE A 385      11.575  37.861  84.057  1.00 51.30           C
    ATOM    570  O   PHE A 385      12.391  38.348  84.840  1.00 48.27           O
    ATOM    571  CB  PHE A 385       9.360  39.052  83.648  1.00 49.88           C
    ATOM    572  CG  PHE A 385       9.192  38.888  82.149  1.00 47.64           C
    ATOM    573  CD1 PHE A 385       8.161  38.071  81.642  1.00 43.08           C
    ATOM    574  CD2 PHE A 385      10.122  39.458  81.253  1.00 46.20           C
    ATOM    575  CE1 PHE A 385       8.031  37.887  80.272  1.00 38.93           C
    ATOM    576  CE2 PHE A 385       9.969  39.256  79.889  1.00 45.22           C
    ATOM    577  CZ  PHE A 385       8.924  38.490  79.399  1.00 39.76           C
    ATOM    578  H   PHE A 385      10.563  38.497  86.359  1.00  0.00           H
    ATOM    579  HA  PHE A 385       9.656  36.946  84.035  1.00  0.00           H
    ATOM    580  HB2 PHE A 385       8.368  39.203  84.071  1.00  0.00           H
    ATOM    581  HB3 PHE A 385       9.898  39.983  83.812  1.00  0.00           H
    ATOM    582  HD1 PHE A 385       7.458  37.603  82.316  1.00  0.00           H
    ATOM    583  HD2 PHE A 385      10.945  40.055  81.618  1.00  0.00           H
    ATOM    584  HE1 PHE A 385       7.237  37.273  79.880  1.00  0.00           H
    ATOM    585  HE2 PHE A 385      10.658  39.693  79.190  1.00  0.00           H
    ATOM    586  HZ  PHE A 385       8.807  38.358  78.335  1.00  0.00           H
    ATOM    587  N   ASN A 387      13.685  38.394  80.170  1.00 59.34           N
    ATOM    588  CA  ASN A 387      13.665  38.731  78.744  1.00 58.75           C
    ATOM    589  C   ASN A 387      14.294  37.653  77.849  1.00 62.64           C
    ATOM    590  O   ASN A 387      13.721  37.383  76.797  1.00 65.84           O
    ATOM    591  CB  ASN A 387      14.379  40.086  78.517  1.00 54.27           C
    ATOM    592  CG  ASN A 387      13.591  41.326  78.962  1.00 53.83           C
    ATOM    593  ND2 ASN A 387      14.301  42.417  79.239  1.00 58.83           N
    ATOM    594  OD1 ASN A 387      12.366  41.340  78.993  1.00 55.68           O
    ATOM    595  H   ASN A 387      14.236  39.012  80.754  1.00  0.00           H
    ATOM    596  HA  ASN A 387      12.611  38.826  78.480  1.00  0.00           H
    ATOM    597  HB2 ASN A 387      15.364  40.077  78.986  1.00  0.00           H
    ATOM    598  HB3 ASN A 387      14.562  40.221  77.451  1.00  0.00           H
    ATOM    599 HD21 ASN A 387      15.308  42.405  79.173  1.00  0.00           H
    ATOM    600 HD22 ASN A 387      13.834  43.279  79.495  1.00  0.00           H
    ATOM    601  N   PHE A 390      11.962  34.595  77.447  1.00 52.70           N
    ATOM    602  CA  PHE A 390      10.906  34.981  76.515  1.00 54.11           C
    ATOM    603  C   PHE A 390      11.371  35.046  75.050  1.00 54.52           C
    ATOM    604  O   PHE A 390      10.657  34.553  74.180  1.00 45.21           O
    ATOM    605  CB  PHE A 390      10.216  36.280  76.984  1.00 54.06           C
    ATOM    606  CG  PHE A 390       9.187  36.823  76.006  1.00 51.41           C
    ATOM    607  CD1 PHE A 390       7.932  36.191  75.893  1.00 48.40           C
    ATOM    608  CD2 PHE A 390       9.550  37.803  75.057  1.00 49.93           C
    ATOM    609  CE1 PHE A 390       7.058  36.561  74.882  1.00 43.19           C
    ATOM    610  CE2 PHE A 390       8.663  38.153  74.048  1.00 47.51           C
    ATOM    611  CZ  PHE A 390       7.422  37.536  73.962  1.00 44.76           C
    ATOM    612  H   PHE A 390      12.292  35.312  78.078  1.00  0.00           H
    ATOM    613  HA  PHE A 390      10.147  34.196  76.558  1.00  0.00           H
    ATOM    614  HB2 PHE A 390       9.703  36.080  77.924  1.00  0.00           H
    ATOM    615  HB3 PHE A 390      10.955  37.053  77.201  1.00  0.00           H
    ATOM    616  HD1 PHE A 390       7.656  35.404  76.579  1.00  0.00           H
    ATOM    617  HD2 PHE A 390      10.528  38.262  75.095  1.00  0.00           H
    ATOM    618  HE1 PHE A 390       6.096  36.080  74.804  1.00  0.00           H
    ATOM    619  HE2 PHE A 390       8.946  38.897  73.318  1.00  0.00           H
    ATOM    620  HZ  PHE A 390       6.739  37.814  73.173  1.00  0.00           H
    ATOM    621  N   ALA A 391      12.553  35.639  74.810  1.00 52.31           N
    ATOM    622  CA  ALA A 391      13.175  35.767  73.491  1.00 55.30           C
    ATOM    623  C   ALA A 391      13.523  34.411  72.854  1.00 59.86           C
    ATOM    624  O   ALA A 391      13.362  34.260  71.644  1.00 61.16           O
    ATOM    625  CB  ALA A 391      14.424  36.654  73.605  1.00 49.16           C
    ATOM    626  H   ALA A 391      13.068  36.044  75.582  1.00  0.00           H
    ATOM    627  HA  ALA A 391      12.460  36.271  72.838  1.00  0.00           H
    ATOM    628  HB1 ALA A 391      14.896  36.797  72.632  1.00  0.00           H
    ATOM    629  HB2 ALA A 391      14.169  37.641  73.991  1.00  0.00           H
    ATOM    630  HB3 ALA A 391      15.167  36.217  74.273  1.00  0.00           H
    ATOM    631  N   LEU A 393      11.814  31.532  73.727  1.00 72.67           N
    ATOM    632  CA  LEU A 393      10.480  30.980  73.479  1.00 71.67           C
    ATOM    633  C   LEU A 393       9.842  31.531  72.188  1.00 72.93           C
    ATOM    634  O   LEU A 393       9.241  30.759  71.444  1.00 73.52           O
    ATOM    635  CB  LEU A 393       9.595  31.172  74.734  1.00 65.60           C
    ATOM    636  CG  LEU A 393       8.123  30.708  74.606  1.00 64.99           C
    ATOM    637  CD1 LEU A 393       7.989  29.226  74.185  1.00 62.48           C
    ATOM    638  CD2 LEU A 393       7.342  31.015  75.896  1.00 67.25           C
    ATOM    639  H   LEU A 393      11.900  32.147  74.525  1.00  0.00           H
    ATOM    640  HA  LEU A 393      10.614  29.908  73.345  1.00  0.00           H
    ATOM    641  HB2 LEU A 393      10.062  30.648  75.569  1.00  0.00           H
    ATOM    642  HB3 LEU A 393       9.597  32.226  75.010  1.00  0.00           H
    ATOM    643  HG  LEU A 393       7.657  31.304  73.824  1.00  0.00           H
    ATOM    644 HD11 LEU A 393       7.290  28.667  74.807  1.00  0.00           H
    ATOM    645 HD12 LEU A 393       7.628  29.148  73.159  1.00  0.00           H
    ATOM    646 HD13 LEU A 393       8.938  28.694  74.237  1.00  0.00           H
    ATOM    647 HD21 LEU A 393       6.735  30.181  76.247  1.00  0.00           H
    ATOM    648 HD22 LEU A 393       8.013  31.279  76.713  1.00  0.00           H
    ATOM    649 HD23 LEU A 393       6.667  31.853  75.737  1.00  0.00           H
    ATOM    650  N   LEU A 394      10.031  32.836  71.927  1.00 75.07           N
    ATOM    651  CA  LEU A 394       9.634  33.535  70.704  1.00 73.94           C
    ATOM    652  C   LEU A 394      10.330  32.985  69.445  1.00 77.30           C
    ATOM    653  O   LEU A 394       9.683  32.887  68.402  1.00 79.25           O
    ATOM    654  CB  LEU A 394       9.876  35.050  70.916  1.00 69.33           C
    ATOM    655  CG  LEU A 394       9.534  35.984  69.732  1.00 67.14           C
    ATOM    656  CD1 LEU A 394       8.047  35.920  69.346  1.00 65.87           C
    ATOM    657  CD2 LEU A 394       9.973  37.431  70.033  1.00 64.52           C
    ATOM    658  H   LEU A 394      10.515  33.401  72.614  1.00  0.00           H
    ATOM    659  HA  LEU A 394       8.567  33.369  70.565  1.00  0.00           H
    ATOM    660  HB2 LEU A 394       9.314  35.378  71.792  1.00  0.00           H
    ATOM    661  HB3 LEU A 394      10.926  35.193  71.172  1.00  0.00           H
    ATOM    662  HG  LEU A 394      10.116  35.667  68.867  1.00  0.00           H
    ATOM    663 HD11 LEU A 394       7.762  36.761  68.716  1.00  0.00           H
    ATOM    664 HD12 LEU A 394       7.825  35.014  68.782  1.00  0.00           H
    ATOM    665 HD13 LEU A 394       7.409  35.935  70.229  1.00  0.00           H
    ATOM    666 HD21 LEU A 394      10.483  37.873  69.177  1.00  0.00           H
    ATOM    667 HD22 LEU A 394       9.124  38.068  70.276  1.00  0.00           H
    ATOM    668 HD23 LEU A 394      10.656  37.487  70.881  1.00  0.00           H
    ATOM    669  N   SER A 397       9.070  29.375  68.465  1.00 93.53           N
    ATOM    670  CA  SER A 397       7.648  29.452  68.123  1.00 96.01           C
    ATOM    671  C   SER A 397       7.386  29.818  66.653  1.00 96.12           C
    ATOM    672  O   SER A 397       6.261  29.609  66.197  1.00 95.60           O
    ATOM    673  CB  SER A 397       6.949  30.456  69.055  1.00 97.26           C
    ATOM    674  OG  SER A 397       6.759  29.881  70.330  1.00 98.20           O
    ATOM    675  H   SER A 397       9.511  30.245  68.726  1.00  0.00           H
    ATOM    676  HA  SER A 397       7.183  28.475  68.272  1.00  0.00           H
    ATOM    677  HB2 SER A 397       7.517  31.383  69.142  1.00  0.00           H
    ATOM    678  HB3 SER A 397       5.959  30.722  68.679  1.00  0.00           H
    ATOM    679  HG  SER A 397       7.596  29.928  70.805  1.00  0.00           H
    ATOM    680  N   VAL A 398       8.407  30.302  65.916  1.00 97.44           N
    ATOM    681  CA  VAL A 398       8.323  30.572  64.471  1.00100.96           C
    ATOM    682  C   VAL A 398       8.100  29.291  63.630  1.00102.30           C
    ATOM    683  O   VAL A 398       7.665  29.384  62.483  1.00103.19           O
    ATOM    684  CB  VAL A 398       9.596  31.293  63.932  1.00100.24           C
    ATOM    685  CG1 VAL A 398       9.918  32.567  64.733  1.00100.80           C
    ATOM    686  CG2 VAL A 398      10.845  30.402  63.755  1.00101.54           C
    ATOM    687  H   VAL A 398       9.310  30.450  66.346  1.00  0.00           H
    ATOM    688  HA  VAL A 398       7.464  31.226  64.306  1.00  0.00           H
    ATOM    689  HB  VAL A 398       9.346  31.655  62.933  1.00  0.00           H
    ATOM    690 HG11 VAL A 398      10.719  33.138  64.262  1.00  0.00           H
    ATOM    691 HG12 VAL A 398       9.046  33.217  64.795  1.00  0.00           H
    ATOM    692 HG13 VAL A 398      10.234  32.337  65.749  1.00  0.00           H
    ATOM    693 HG21 VAL A 398      11.745  31.001  63.616  1.00  0.00           H
    ATOM    694 HG22 VAL A 398      11.003  29.751  64.613  1.00  0.00           H
    ATOM    695 HG23 VAL A 398      10.754  29.763  62.877  1.00  0.00           H
    ATOM    696  N   GLN A 400       5.941  26.918  64.676  1.00103.20           N
    ATOM    697  CA  GLN A 400       4.597  26.466  65.054  1.00100.30           C
    ATOM    698  C   GLN A 400       3.456  27.254  64.377  1.00 97.12           C
    ATOM    699  O   GLN A 400       2.305  26.839  64.505  1.00 91.44           O
    ATOM    700  CB  GLN A 400       4.464  26.510  66.590  1.00102.76           C
    ATOM    701  CG  GLN A 400       5.428  25.555  67.320  1.00106.14           C
    ATOM    702  CD  GLN A 400       5.076  25.410  68.801  1.00109.38           C
    ATOM    703  NE2 GLN A 400       5.284  26.469  69.587  1.00110.25           N
    ATOM    704  OE1 GLN A 400       4.618  24.354  69.231  1.00111.88           O
    ATOM    705  H   GLN A 400       6.253  27.779  65.105  1.00  0.00           H
    ATOM    706  HA  GLN A 400       4.456  25.427  64.749  1.00  0.00           H
    ATOM    707  HB2 GLN A 400       4.634  27.527  66.943  1.00  0.00           H
    ATOM    708  HB3 GLN A 400       3.440  26.253  66.867  1.00  0.00           H
    ATOM    709  HG2 GLN A 400       5.388  24.567  66.860  1.00  0.00           H
    ATOM    710  HG3 GLN A 400       6.458  25.901  67.227  1.00  0.00           H
    ATOM    711 HE21 GLN A 400       5.663  27.326  69.212  1.00  0.00           H
    ATOM    712 HE22 GLN A 400       5.066  26.414  70.571  1.00  0.00           H
    ATOM    713  N   GLY A 401       3.773  28.342  63.653  1.00 96.01           N
    ATOM    714  CA  GLY A 401       2.801  29.118  62.876  1.00 97.69           C
    ATOM    715  C   GLY A 401       2.229  30.300  63.675  1.00100.14           C
    ATOM    716  O   GLY A 401       2.582  30.520  64.836  1.00103.86           O
    ATOM    717  H   GLY A 401       4.739  28.631  63.598  1.00  0.00           H
    ATOM    718  HA2 GLY A 401       3.308  29.504  61.992  1.00  0.00           H
    ATOM    719  HA3 GLY A 401       1.985  28.487  62.519  1.00  0.00           H
    ATOM    720  N   PHE A 402       1.362  31.083  63.002  1.00 98.26           N
    ATOM    721  CA  PHE A 402       0.790  32.358  63.459  1.00 96.62           C
    ATOM    722  C   PHE A 402       0.067  32.278  64.814  1.00 96.74           C
    ATOM    723  O   PHE A 402       0.270  33.162  65.646  1.00 98.07           O
    ATOM    724  CB  PHE A 402      -0.142  32.926  62.358  1.00 93.36           C
    ATOM    725  CG  PHE A 402      -0.676  34.335  62.593  1.00 92.02           C
    ATOM    726  CD1 PHE A 402      -1.808  34.567  63.405  1.00 88.30           C
    ATOM    727  CD2 PHE A 402       0.054  35.443  62.120  1.00 93.05           C
    ATOM    728  CE1 PHE A 402      -2.206  35.868  63.692  1.00 87.77           C
    ATOM    729  CE2 PHE A 402      -0.356  36.733  62.416  1.00 93.23           C
    ATOM    730  CZ  PHE A 402      -1.482  36.947  63.200  1.00 90.65           C
    ATOM    731  H   PHE A 402       1.126  30.827  62.054  1.00  0.00           H
    ATOM    732  HA  PHE A 402       1.627  33.047  63.583  1.00  0.00           H
    ATOM    733  HB2 PHE A 402       0.393  32.937  61.407  1.00  0.00           H
    ATOM    734  HB3 PHE A 402      -0.990  32.255  62.206  1.00  0.00           H
    ATOM    735  HD1 PHE A 402      -2.365  33.736  63.812  1.00  0.00           H
    ATOM    736  HD2 PHE A 402       0.923  35.292  61.504  1.00  0.00           H
    ATOM    737  HE1 PHE A 402      -3.077  36.040  64.307  1.00  0.00           H
    ATOM    738  HE2 PHE A 402       0.210  37.567  62.033  1.00  0.00           H
    ATOM    739  HZ  PHE A 402      -1.795  37.955  63.430  1.00  0.00           H
    ATOM    740  N   GLU A 403      -0.740  31.215  65.001  1.00 97.27           N
    ATOM    741  CA  GLU A 403      -1.495  30.893  66.216  1.00 97.28           C
    ATOM    742  C   GLU A 403      -0.626  30.903  67.483  1.00 95.65           C
    ATOM    743  O   GLU A 403      -0.935  31.632  68.426  1.00 95.69           O
    ATOM    744  CB  GLU A 403      -2.189  29.523  66.025  1.00100.62           C
    ATOM    745  CG  GLU A 403      -2.908  28.965  67.278  1.00109.96           C
    ATOM    746  CD  GLU A 403      -3.453  27.553  67.069  1.00115.15           C
    ATOM    747  OE1 GLU A 403      -2.662  26.691  66.627  1.00120.27           O
    ATOM    748  OE2 GLU A 403      -4.645  27.354  67.388  1.00116.80           O1-
    ATOM    749  H   GLU A 403      -0.826  30.542  64.254  1.00  0.00           H
    ATOM    750  HA  GLU A 403      -2.267  31.656  66.338  1.00  0.00           H
    ATOM    751  HB2 GLU A 403      -2.918  29.609  65.218  1.00  0.00           H
    ATOM    752  HB3 GLU A 403      -1.450  28.800  65.675  1.00  0.00           H
    ATOM    753  HG2 GLU A 403      -2.239  28.900  68.135  1.00  0.00           H
    ATOM    754  HG3 GLU A 403      -3.719  29.634  67.567  1.00  0.00           H
    ATOM    755  N   ALA A 404       0.442  30.087  67.454  1.00 93.29           N
    ATOM    756  CA  ALA A 404       1.392  29.901  68.546  1.00 91.62           C
    ATOM    757  C   ALA A 404       2.100  31.197  68.961  1.00 87.85           C
    ATOM    758  O   ALA A 404       2.250  31.428  70.160  1.00 89.13           O
    ATOM    759  CB  ALA A 404       2.416  28.836  68.137  1.00 94.35           C
    ATOM    760  H   ALA A 404       0.613  29.535  66.626  1.00  0.00           H
    ATOM    761  HA  ALA A 404       0.835  29.530  69.409  1.00  0.00           H
    ATOM    762  HB1 ALA A 404       3.143  28.655  68.930  1.00  0.00           H
    ATOM    763  HB2 ALA A 404       1.926  27.887  67.920  1.00  0.00           H
    ATOM    764  HB3 ALA A 404       2.966  29.139  67.244  1.00  0.00           H
    ATOM    765  N   VAL A 405       2.482  32.020  67.969  1.00 83.64           N
    ATOM    766  CA  VAL A 405       3.156  33.298  68.182  1.00 81.07           C
    ATOM    767  C   VAL A 405       2.206  34.388  68.725  1.00 81.10           C
    ATOM    768  O   VAL A 405       2.621  35.107  69.630  1.00 82.21           O
    ATOM    769  CB  VAL A 405       3.877  33.795  66.898  1.00 78.93           C
    ATOM    770  CG1 VAL A 405       4.539  35.178  67.037  1.00 74.82           C
    ATOM    771  CG2 VAL A 405       4.957  32.794  66.453  1.00 81.14           C
    ATOM    772  H   VAL A 405       2.310  31.757  67.009  1.00  0.00           H
    ATOM    773  HA  VAL A 405       3.924  33.135  68.940  1.00  0.00           H
    ATOM    774  HB  VAL A 405       3.139  33.864  66.097  1.00  0.00           H
    ATOM    775 HG11 VAL A 405       5.000  35.490  66.102  1.00  0.00           H
    ATOM    776 HG12 VAL A 405       3.819  35.945  67.296  1.00  0.00           H
    ATOM    777 HG13 VAL A 405       5.312  35.173  67.807  1.00  0.00           H
    ATOM    778 HG21 VAL A 405       5.487  33.146  65.569  1.00  0.00           H
    ATOM    779 HG22 VAL A 405       5.702  32.645  67.235  1.00  0.00           H
    ATOM    780 HG23 VAL A 405       4.532  31.820  66.211  1.00  0.00           H
    ATOM    781  N   TYR A 406       0.953  34.469  68.231  1.00 78.20           N
    ATOM    782  CA  TYR A 406      -0.078  35.395  68.732  1.00 77.28           C
    ATOM    783  C   TYR A 406      -0.373  35.198  70.234  1.00 75.07           C
    ATOM    784  O   TYR A 406      -0.558  36.179  70.953  1.00 75.66           O
    ATOM    785  CB  TYR A 406      -1.369  35.270  67.884  1.00 77.36           C
    ATOM    786  CG  TYR A 406      -2.501  36.234  68.237  1.00 75.70           C
    ATOM    787  CD1 TYR A 406      -3.352  35.981  69.338  1.00 74.72           C
    ATOM    788  CD2 TYR A 406      -2.719  37.388  67.453  1.00 75.90           C
    ATOM    789  CE1 TYR A 406      -4.387  36.878  69.665  1.00 79.09           C
    ATOM    790  CE2 TYR A 406      -3.764  38.278  67.770  1.00 80.00           C
    ATOM    791  CZ  TYR A 406      -4.596  38.028  68.880  1.00 80.99           C
    ATOM    792  OH  TYR A 406      -5.603  38.894  69.191  1.00 85.69           O
    ATOM    793  H   TYR A 406       0.672  33.849  67.482  1.00  0.00           H
    ATOM    794  HA  TYR A 406       0.311  36.408  68.603  1.00  0.00           H
    ATOM    795  HB2 TYR A 406      -1.121  35.411  66.832  1.00  0.00           H
    ATOM    796  HB3 TYR A 406      -1.759  34.253  67.953  1.00  0.00           H
    ATOM    797  HD1 TYR A 406      -3.213  35.095  69.940  1.00  0.00           H
    ATOM    798  HD2 TYR A 406      -2.084  37.595  66.604  1.00  0.00           H
    ATOM    799  HE1 TYR A 406      -5.026  36.677  70.513  1.00  0.00           H
    ATOM    800  HE2 TYR A 406      -3.920  39.157  67.160  1.00  0.00           H
    ATOM    801  HH  TYR A 406      -5.667  39.621  68.567  1.00  0.00           H
    ATOM    802  N   GLN A 407      -0.386  33.930  70.675  1.00 71.26           N
    ATOM    803  CA  GLN A 407      -0.621  33.527  72.060  1.00 72.54           C
    ATOM    804  C   GLN A 407       0.540  33.808  73.038  1.00 69.21           C
    ATOM    805  O   GLN A 407       0.340  33.654  74.244  1.00 64.99           O
    ATOM    806  CB  GLN A 407      -1.073  32.056  72.079  1.00 72.65           C
    ATOM    807  CG  GLN A 407      -2.520  31.878  71.575  1.00 82.61           C
    ATOM    808  CD  GLN A 407      -3.074  30.466  71.788  1.00 88.89           C
    ATOM    809  NE2 GLN A 407      -4.074  30.088  70.991  1.00 90.14           N
    ATOM    810  OE1 GLN A 407      -2.638  29.735  72.676  1.00 93.76           O
    ATOM    811  H   GLN A 407      -0.225  33.179  70.018  1.00  0.00           H
    ATOM    812  HA  GLN A 407      -1.452  34.129  72.435  1.00  0.00           H
    ATOM    813  HB2 GLN A 407      -0.392  31.442  71.487  1.00  0.00           H
    ATOM    814  HB3 GLN A 407      -1.004  31.682  73.097  1.00  0.00           H
    ATOM    815  HG2 GLN A 407      -3.181  32.565  72.105  1.00  0.00           H
    ATOM    816  HG3 GLN A 407      -2.583  32.140  70.518  1.00  0.00           H
    ATOM    817 HE21 GLN A 407      -4.424  30.708  70.275  1.00  0.00           H
    ATOM    818 HE22 GLN A 407      -4.479  29.168  71.088  1.00  0.00           H
    ATOM    819  N   LEU A 408       1.701  34.267  72.534  1.00 68.75           N
    ATOM    820  CA  LEU A 408       2.822  34.758  73.347  1.00 69.59           C
    ATOM    821  C   LEU A 408       2.571  36.147  73.964  1.00 70.95           C
    ATOM    822  O   LEU A 408       3.396  36.592  74.759  1.00 69.85           O
    ATOM    823  CB  LEU A 408       4.123  34.775  72.516  1.00 72.28           C
    ATOM    824  CG  LEU A 408       4.575  33.392  72.016  1.00 73.20           C
    ATOM    825  CD1 LEU A 408       5.793  33.532  71.099  1.00 73.44           C
    ATOM    826  CD2 LEU A 408       4.860  32.405  73.160  1.00 75.60           C
    ATOM    827  H   LEU A 408       1.803  34.370  71.533  1.00  0.00           H
    ATOM    828  HA  LEU A 408       2.958  34.076  74.187  1.00  0.00           H
    ATOM    829  HB2 LEU A 408       4.006  35.460  71.678  1.00  0.00           H
    ATOM    830  HB3 LEU A 408       4.934  35.194  73.110  1.00  0.00           H
    ATOM    831  HG  LEU A 408       3.776  32.978  71.407  1.00  0.00           H
    ATOM    832 HD11 LEU A 408       5.926  32.640  70.489  1.00  0.00           H
    ATOM    833 HD12 LEU A 408       5.684  34.376  70.418  1.00  0.00           H
    ATOM    834 HD13 LEU A 408       6.696  33.685  71.688  1.00  0.00           H
    ATOM    835 HD21 LEU A 408       5.660  31.713  72.897  1.00  0.00           H
    ATOM    836 HD22 LEU A 408       5.153  32.919  74.074  1.00  0.00           H
    ATOM    837 HD23 LEU A 408       3.979  31.803  73.380  1.00  0.00           H
    ATOM    838  N   THR A 409       1.454  36.814  73.623  1.00 73.57           N
    ATOM    839  CA  THR A 409       1.014  38.052  74.276  1.00 73.66           C
    ATOM    840  C   THR A 409       0.693  37.841  75.767  1.00 73.25           C
    ATOM    841  O   THR A 409       0.988  38.733  76.557  1.00 75.17           O
    ATOM    842  CB  THR A 409      -0.259  38.638  73.617  1.00 69.68           C
    ATOM    843  CG2 THR A 409       0.017  39.187  72.217  1.00 59.44           C
    ATOM    844  OG1 THR A 409      -1.300  37.679  73.527  1.00 73.08           O
    ATOM    845  H   THR A 409       0.824  36.420  72.939  1.00  0.00           H
    ATOM    846  HA  THR A 409       1.822  38.783  74.201  1.00  0.00           H
    ATOM    847  HB  THR A 409      -0.640  39.464  74.222  1.00  0.00           H
    ATOM    848 HG21 THR A 409      -0.903  39.549  71.756  1.00  0.00           H
    ATOM    849 HG22 THR A 409       0.713  40.025  72.269  1.00  0.00           H
    ATOM    850 HG23 THR A 409       0.456  38.441  71.557  1.00  0.00           H
    ATOM    851  HG1 THR A 409      -1.187  37.182  72.711  1.00  0.00           H
    ATOM    852  N   ARG A 410       0.171  36.655  76.125  1.00 71.88           N
    ATOM    853  CA  ARG A 410      -0.143  36.241  77.493  1.00 70.00           C
    ATOM    854  C   ARG A 410       1.104  36.093  78.383  1.00 63.25           C
    ATOM    855  O   ARG A 410       1.001  36.369  79.576  1.00 63.28           O
    ATOM    856  CB  ARG A 410      -0.924  34.917  77.459  1.00 72.03           C
    ATOM    857  CG  ARG A 410      -2.185  34.943  76.567  1.00 78.64           C
    ATOM    858  CD  ARG A 410      -2.869  33.574  76.458  1.00 84.97           C
    ATOM    859  NE  ARG A 410      -1.930  32.564  75.952  1.00 94.42           N
    ATOM    860  CZ  ARG A 410      -2.193  31.270  75.721  1.00 99.96           C
    ATOM    861  NH1 ARG A 410      -3.428  30.773  75.864  1.00102.63           N
    ATOM    862  NH2 ARG A 410      -1.194  30.462  75.345  1.00102.58           N1+
    ATOM    863  H   ARG A 410      -0.022  35.975  75.404  1.00  0.00           H
    ATOM    864  HA  ARG A 410      -0.781  37.009  77.934  1.00  0.00           H
    ATOM    865  HB2 ARG A 410      -0.243  34.142  77.106  1.00  0.00           H
    ATOM    866  HB3 ARG A 410      -1.209  34.628  78.473  1.00  0.00           H
    ATOM    867  HG2 ARG A 410      -2.886  35.652  77.009  1.00  0.00           H
    ATOM    868  HG3 ARG A 410      -1.982  35.312  75.561  1.00  0.00           H
    ATOM    869  HD2 ARG A 410      -3.320  33.276  77.405  1.00  0.00           H
    ATOM    870  HD3 ARG A 410      -3.675  33.641  75.726  1.00  0.00           H
    ATOM    871  HE  ARG A 410      -0.996  32.900  75.754  1.00  0.00           H
    ATOM    872 HH11 ARG A 410      -4.187  31.379  76.139  1.00  0.00           H
    ATOM    873 HH12 ARG A 410      -3.617  29.801  75.667  1.00  0.00           H
    ATOM    874 HH21 ARG A 410      -0.258  30.823  75.235  1.00  0.00           H
    ATOM    875 HH22 ARG A 410      -1.378  29.494  75.122  1.00  0.00           H
    ATOM    876  N   MET A 411       2.253  35.713  77.790  1.00 60.91           N
    ATOM    877  CA  MET A 411       3.568  35.707  78.443  1.00 57.05           C
    ATOM    878  C   MET A 411       4.041  37.116  78.827  1.00 56.17           C
    ATOM    879  O   MET A 411       4.707  37.265  79.850  1.00 60.09           O
    ATOM    880  CB  MET A 411       4.623  35.076  77.515  1.00 53.88           C
    ATOM    881  CG  MET A 411       4.466  33.571  77.272  1.00 56.44           C
    ATOM    882  SD  MET A 411       5.110  32.498  78.585  1.00 64.23           S
    ATOM    883  CE  MET A 411       6.844  33.028  78.637  1.00 63.26           C
    ATOM    884  H   MET A 411       2.251  35.514  76.800  1.00  0.00           H
    ATOM    885  HA  MET A 411       3.499  35.119  79.359  1.00  0.00           H
    ATOM    886  HB2 MET A 411       4.639  35.588  76.560  1.00  0.00           H
    ATOM    887  HB3 MET A 411       5.613  35.265  77.928  1.00  0.00           H
    ATOM    888  HG2 MET A 411       3.420  33.318  77.098  1.00  0.00           H
    ATOM    889  HG3 MET A 411       4.991  33.311  76.355  1.00  0.00           H
    ATOM    890  HE1 MET A 411       7.462  32.249  79.084  1.00  0.00           H
    ATOM    891  HE2 MET A 411       7.221  33.240  77.638  1.00  0.00           H
    ATOM    892  HE3 MET A 411       6.942  33.933  79.232  1.00  0.00           H
    ATOM    893  N   CYS A 412       3.677  38.113  78.004  1.00 55.51           N
    ATOM    894  CA  CYS A 412       4.001  39.520  78.217  1.00 52.91           C
    ATOM    895  C   CYS A 412       3.077  40.185  79.255  1.00 53.00           C
    ATOM    896  O   CYS A 412       3.415  41.279  79.705  1.00 48.95           O
    ATOM    897  CB  CYS A 412       3.947  40.329  76.911  1.00 57.19           C
    ATOM    898  SG  CYS A 412       4.965  39.584  75.606  1.00 64.63           S
    ATOM    899  H   CYS A 412       3.117  37.905  77.189  1.00  0.00           H
    ATOM    900  HA  CYS A 412       5.022  39.582  78.602  1.00  0.00           H
    ATOM    901  HB2 CYS A 412       2.925  40.431  76.550  1.00  0.00           H
    ATOM    902  HB3 CYS A 412       4.326  41.332  77.105  1.00  0.00           H
    ATOM    903  HG  CYS A 412       4.221  38.485  75.445  1.00  0.00           H
    ATOM    904  N   THR A 413       1.951  39.538  79.618  1.00 47.74           N
    ATOM    905  CA  THR A 413       1.049  40.002  80.673  1.00 49.12           C
    ATOM    906  C   THR A 413       1.297  39.234  81.982  1.00 47.64           C
    ATOM    907  O   THR A 413       1.536  38.026  81.959  1.00 49.84           O
    ATOM    908  CB  THR A 413      -0.460  39.904  80.300  1.00 55.21           C
    ATOM    909  CG2 THR A 413      -0.810  40.434  78.906  1.00 52.24           C
    ATOM    910  OG1 THR A 413      -1.038  38.623  80.485  1.00 58.09           O
    ATOM    911  H   THR A 413       1.741  38.635  79.217  1.00  0.00           H
    ATOM    912  HA  THR A 413       1.262  41.050  80.849  1.00  0.00           H
    ATOM    913  HB  THR A 413      -0.985  40.551  81.005  1.00  0.00           H
    ATOM    914 HG21 THR A 413      -1.881  40.626  78.823  1.00  0.00           H
    ATOM    915 HG22 THR A 413      -0.285  41.363  78.687  1.00  0.00           H
    ATOM    916 HG23 THR A 413      -0.558  39.715  78.135  1.00  0.00           H
    ATOM    917  HG1 THR A 413      -0.497  37.972  80.031  1.00  0.00           H
    ATOM    918  N   ILE A 414       1.247  39.970  83.101  1.00 43.32           N
    ATOM    919  CA  ILE A 414       1.442  39.437  84.446  1.00 38.25           C
    ATOM    920  C   ILE A 414       0.398  40.094  85.352  1.00 35.50           C
    ATOM    921  O   ILE A 414       0.288  41.318  85.355  1.00 38.65           O
    ATOM    922  CB  ILE A 414       2.854  39.763  85.013  1.00 31.37           C
    ATOM    923  CG1 ILE A 414       3.974  39.165  84.132  1.00 32.84           C
    ATOM    924  CG2 ILE A 414       3.024  39.325  86.487  1.00 26.30           C
    ATOM    925  CD1 ILE A 414       5.330  39.849  84.309  1.00 22.07           C
    ATOM    926  H   ILE A 414       1.042  40.958  83.042  1.00  0.00           H
    ATOM    927  HA  ILE A 414       1.289  38.355  84.457  1.00  0.00           H
    ATOM    928  HB  ILE A 414       2.984  40.845  84.987  1.00  0.00           H
    ATOM    929 HG12 ILE A 414       4.049  38.095  84.303  1.00  0.00           H
    ATOM    930 HG13 ILE A 414       3.741  39.264  83.076  1.00  0.00           H
    ATOM    931 HG21 ILE A 414       4.047  39.444  86.834  1.00  0.00           H
    ATOM    932 HG22 ILE A 414       2.404  39.912  87.165  1.00  0.00           H
    ATOM    933 HG23 ILE A 414       2.747  38.279  86.617  1.00  0.00           H
    ATOM    934 HD11 ILE A 414       5.543  40.494  83.461  1.00  0.00           H
    ATOM    935 HD12 ILE A 414       5.360  40.489  85.189  1.00  0.00           H
    ATOM    936 HD13 ILE A 414       6.127  39.111  84.377  1.00  0.00           H
    ATOM    937  N   ARG A 415      -0.337  39.270  86.106  1.00 34.82           N
    ATOM    938  CA  ARG A 415      -1.419  39.722  86.974  1.00 35.47           C
    ATOM    939  C   ARG A 415      -1.026  39.480  88.427  1.00 37.26           C
    ATOM    940  O   ARG A 415      -0.522  38.401  88.732  1.00 38.74           O
    ATOM    941  CB  ARG A 415      -2.703  38.964  86.600  1.00 33.20           C
    ATOM    942  CG  ARG A 415      -3.202  39.336  85.195  1.00 31.51           C
    ATOM    943  CD  ARG A 415      -4.231  38.352  84.644  1.00 44.37           C
    ATOM    944  NE  ARG A 415      -4.828  38.862  83.407  1.00 53.95           N
    ATOM    945  CZ  ARG A 415      -5.670  38.198  82.604  1.00 51.65           C
    ATOM    946  NH1 ARG A 415      -6.076  36.953  82.891  1.00 51.74           N
    ATOM    947  NH2 ARG A 415      -6.102  38.805  81.493  1.00 48.86           N1+
    ATOM    948  H   ARG A 415      -0.170  38.273  86.077  1.00  0.00           H
    ATOM    949  HA  ARG A 415      -1.601  40.785  86.834  1.00  0.00           H
    ATOM    950  HB2 ARG A 415      -2.523  37.889  86.660  1.00  0.00           H
    ATOM    951  HB3 ARG A 415      -3.491  39.184  87.320  1.00  0.00           H
    ATOM    952  HG2 ARG A 415      -3.533  40.373  85.127  1.00  0.00           H
    ATOM    953  HG3 ARG A 415      -2.351  39.257  84.518  1.00  0.00           H
    ATOM    954  HD2 ARG A 415      -3.693  37.458  84.326  1.00  0.00           H
    ATOM    955  HD3 ARG A 415      -4.951  38.025  85.390  1.00  0.00           H
    ATOM    956  HE  ARG A 415      -4.540  39.788  83.107  1.00  0.00           H
    ATOM    957 HH11 ARG A 415      -5.735  36.494  83.723  1.00  0.00           H
    ATOM    958 HH12 ARG A 415      -6.709  36.463  82.276  1.00  0.00           H
    ATOM    959 HH21 ARG A 415      -5.742  39.731  81.275  1.00  0.00           H
    ATOM    960 HH22 ARG A 415      -6.738  38.351  80.855  1.00  0.00           H
    ATOM    961  N   MET A 416      -1.275  40.483  89.285  1.00 38.76           N
    ATOM    962  CA  MET A 416      -1.024  40.400  90.723  1.00 35.40           C
    ATOM    963  C   MET A 416      -2.260  40.845  91.500  1.00 32.84           C
    ATOM    964  O   MET A 416      -2.843  41.858  91.132  1.00 30.72           O
    ATOM    965  CB  MET A 416       0.191  41.241  91.166  1.00 37.51           C
    ATOM    966  CG  MET A 416       1.377  41.243  90.200  1.00 47.67           C
    ATOM    967  SD  MET A 416       2.879  41.969  90.896  1.00 56.66           S
    ATOM    968  CE  MET A 416       3.917  41.828  89.426  1.00 52.07           C
    ATOM    969  H   MET A 416      -1.713  41.333  88.955  1.00  0.00           H
    ATOM    970  HA  MET A 416      -0.814  39.367  90.951  1.00  0.00           H
    ATOM    971  HB2 MET A 416      -0.113  42.273  91.320  1.00  0.00           H
    ATOM    972  HB3 MET A 416       0.521  40.891  92.146  1.00  0.00           H
    ATOM    973  HG2 MET A 416       1.603  40.226  89.886  1.00  0.00           H
    ATOM    974  HG3 MET A 416       1.119  41.800  89.299  1.00  0.00           H
    ATOM    975  HE1 MET A 416       4.944  42.096  89.660  1.00  0.00           H
    ATOM    976  HE2 MET A 416       3.917  40.803  89.057  1.00  0.00           H
    ATOM    977  HE3 MET A 416       3.547  42.477  88.636  1.00  0.00           H
    ATOM    978  N   SER A 417      -2.585  40.125  92.584  1.00 26.75           N
    ATOM    979  CA  SER A 417      -3.647  40.475  93.524  1.00 29.39           C
    ATOM    980  C   SER A 417      -3.039  40.841  94.881  1.00 35.02           C
    ATOM    981  O   SER A 417      -2.153  40.136  95.364  1.00 31.19           O
    ATOM    982  CB  SER A 417      -4.637  39.303  93.645  1.00 31.58           C
    ATOM    983  OG  SER A 417      -5.662  39.585  94.571  1.00 34.45           O
    ATOM    984  H   SER A 417      -2.026  39.316  92.825  1.00  0.00           H
    ATOM    985  HA  SER A 417      -4.200  41.337  93.158  1.00  0.00           H
    ATOM    986  HB2 SER A 417      -5.093  39.083  92.678  1.00  0.00           H
    ATOM    987  HB3 SER A 417      -4.131  38.401  93.985  1.00  0.00           H
    ATOM    988  HG  SER A 417      -5.437  39.169  95.410  1.00  0.00           H
    ATOM    989  N   TRP A 422      -5.855  33.660  94.789  1.00 45.13           N
    ATOM    990  CA  TRP A 422      -6.763  32.988  93.862  1.00 46.13           C
    ATOM    991  C   TRP A 422      -6.081  31.776  93.204  1.00 47.60           C
    ATOM    992  O   TRP A 422      -4.921  31.473  93.489  1.00 42.44           O
    ATOM    993  CB  TRP A 422      -7.356  33.991  92.837  1.00 47.20           C
    ATOM    994  CG  TRP A 422      -6.580  34.244  91.574  1.00 51.15           C
    ATOM    995  CD1 TRP A 422      -6.781  33.610  90.395  1.00 52.63           C
    ATOM    996  CD2 TRP A 422      -5.426  35.107  91.361  1.00 48.55           C
    ATOM    997  CE2 TRP A 422      -4.998  34.966  90.004  1.00 46.17           C
    ATOM    998  CE3 TRP A 422      -4.692  35.994  92.176  1.00 46.59           C
    ATOM    999  NE1 TRP A 422      -5.861  34.039  89.466  1.00 46.92           N
    ATOM   1000  CZ2 TRP A 422      -3.911  35.682  89.479  1.00 45.72           C
    ATOM   1001  CZ3 TRP A 422      -3.595  36.707  91.659  1.00 43.79           C
    ATOM   1002  CH2 TRP A 422      -3.200  36.556  90.317  1.00 48.30           C
    ATOM   1003  H   TRP A 422      -4.870  33.569  94.582  1.00  0.00           H
    ATOM   1004  HA  TRP A 422      -7.577  32.596  94.469  1.00  0.00           H
    ATOM   1005  HB2 TRP A 422      -8.334  33.624  92.523  1.00  0.00           H
    ATOM   1006  HB3 TRP A 422      -7.561  34.943  93.322  1.00  0.00           H
    ATOM   1007  HD1 TRP A 422      -7.550  32.874  90.224  1.00  0.00           H
    ATOM   1008  HE1 TRP A 422      -5.834  33.687  88.519  1.00  0.00           H
    ATOM   1009  HE3 TRP A 422      -4.977  36.128  93.209  1.00  0.00           H
    ATOM   1010  HZ2 TRP A 422      -3.628  35.562  88.444  1.00  0.00           H
    ATOM   1011  HZ3 TRP A 422      -3.061  37.382  92.301  1.00  0.00           H
    ATOM   1012  HH2 TRP A 422      -2.358  37.111  89.937  1.00  0.00           H
    ATOM   1013  N   ARG A 428     -11.439  31.429  95.523  1.00 59.46           N
    ATOM   1014  CA  ARG A 428     -11.400  32.281  94.340  1.00 54.38           C
    ATOM   1015  C   ARG A 428     -10.588  31.560  93.260  1.00 55.88           C
    ATOM   1016  O   ARG A 428      -9.424  31.270  93.511  1.00 62.28           O
    ATOM   1017  CB  ARG A 428     -10.737  33.629  94.707  1.00 50.30           C
    ATOM   1018  CG  ARG A 428     -11.549  34.505  95.678  1.00 47.40           C
    ATOM   1019  CD  ARG A 428     -10.886  35.875  95.884  1.00 45.77           C
    ATOM   1020  NE  ARG A 428     -11.699  36.777  96.713  1.00 43.01           N
    ATOM   1021  CZ  ARG A 428     -11.724  36.818  98.057  1.00 41.97           C
    ATOM   1022  NH1 ARG A 428     -10.971  35.997  98.800  1.00 42.06           N
    ATOM   1023  NH2 ARG A 428     -12.516  37.701  98.678  1.00 33.40           N1+
    ATOM   1024  H   ARG A 428     -10.842  31.716  96.285  1.00  0.00           H
    ATOM   1025  HA  ARG A 428     -12.417  32.458  93.986  1.00  0.00           H
    ATOM   1026  HB2 ARG A 428      -9.747  33.453  95.130  1.00  0.00           H
    ATOM   1027  HB3 ARG A 428     -10.570  34.195  93.789  1.00  0.00           H
    ATOM   1028  HG2 ARG A 428     -12.552  34.651  95.276  1.00  0.00           H
    ATOM   1029  HG3 ARG A 428     -11.674  34.003  96.637  1.00  0.00           H
    ATOM   1030  HD2 ARG A 428      -9.881  35.783  96.296  1.00  0.00           H
    ATOM   1031  HD3 ARG A 428     -10.767  36.337  94.906  1.00  0.00           H
    ATOM   1032  HE  ARG A 428     -12.317  37.393  96.201  1.00  0.00           H
    ATOM   1033 HH11 ARG A 428     -10.344  35.332  98.371  1.00  0.00           H
    ATOM   1034 HH12 ARG A 428     -11.011  36.070  99.814  1.00  0.00           H
    ATOM   1035 HH21 ARG A 428     -13.086  38.354  98.161  1.00  0.00           H
    ATOM   1036 HH22 ARG A 428     -12.508  37.734  99.694  1.00  0.00           H
    ATOM   1037  N   GLN A 429     -11.209  31.246  92.115  1.00 51.94           N
    ATOM   1038  CA  GLN A 429     -10.550  30.516  91.024  1.00 52.70           C
    ATOM   1039  C   GLN A 429     -10.055  31.439  89.904  1.00 50.93           C
    ATOM   1040  O   GLN A 429      -9.172  31.030  89.151  1.00 57.22           O
    ATOM   1041  CB  GLN A 429     -11.506  29.441  90.460  1.00 54.24           C
    ATOM   1042  CG  GLN A 429     -11.911  28.335  91.457  1.00 55.61           C
    ATOM   1043  CD  GLN A 429     -10.724  27.654  92.146  1.00 61.33           C
    ATOM   1044  NE2 GLN A 429     -10.806  27.509  93.466  1.00 57.80           N
    ATOM   1045  OE1 GLN A 429      -9.756  27.261  91.499  1.00 66.30           O
    ATOM   1046  H   GLN A 429     -12.183  31.478  91.983  1.00  0.00           H
    ATOM   1047  HA  GLN A 429      -9.653  30.018  91.394  1.00  0.00           H
    ATOM   1048  HB2 GLN A 429     -12.410  29.917  90.078  1.00  0.00           H
    ATOM   1049  HB3 GLN A 429     -11.039  28.965  89.596  1.00  0.00           H
    ATOM   1050  HG2 GLN A 429     -12.574  28.758  92.213  1.00  0.00           H
    ATOM   1051  HG3 GLN A 429     -12.492  27.571  90.940  1.00  0.00           H
    ATOM   1052 HE21 GLN A 429     -11.611  27.852  93.969  1.00  0.00           H
    ATOM   1053 HE22 GLN A 429     -10.037  27.108  93.986  1.00  0.00           H
    ATOM   1054  N   THR A 430     -10.609  32.656  89.823  1.00 44.70           N
    ATOM   1055  CA  THR A 430     -10.258  33.671  88.835  1.00 41.86           C
    ATOM   1056  C   THR A 430      -9.799  34.946  89.560  1.00 45.78           C
    ATOM   1057  O   THR A 430     -10.238  35.221  90.677  1.00 37.74           O
    ATOM   1058  CB  THR A 430     -11.485  34.018  87.949  1.00 39.57           C
    ATOM   1059  CG2 THR A 430     -11.970  32.826  87.111  1.00 41.76           C
    ATOM   1060  OG1 THR A 430     -12.584  34.492  88.706  1.00 46.58           O
    ATOM   1061  H   THR A 430     -11.325  32.924  90.483  1.00  0.00           H
    ATOM   1062  HA  THR A 430      -9.441  33.329  88.195  1.00  0.00           H
    ATOM   1063  HB  THR A 430     -11.207  34.817  87.259  1.00  0.00           H
    ATOM   1064 HG21 THR A 430     -12.791  33.118  86.455  1.00  0.00           H
    ATOM   1065 HG22 THR A 430     -11.169  32.439  86.482  1.00  0.00           H
    ATOM   1066 HG23 THR A 430     -12.326  32.008  87.738  1.00  0.00           H
    ATOM   1067  HG1 THR A 430     -12.320  35.277  89.192  1.00  0.00           H
    ATOM   1068  N   VAL A 431      -8.917  35.709  88.894  1.00 42.06           N
    ATOM   1069  CA  VAL A 431      -8.382  36.975  89.398  1.00 40.13           C
    ATOM   1070  C   VAL A 431      -9.462  38.077  89.502  1.00 37.36           C
    ATOM   1071  O   VAL A 431      -9.353  38.949  90.359  1.00 33.61           O
    ATOM   1072  CB  VAL A 431      -7.203  37.471  88.515  1.00 31.64           C
    ATOM   1073  CG1 VAL A 431      -7.624  38.049  87.147  1.00 37.98           C
    ATOM   1074  CG2 VAL A 431      -6.294  38.454  89.272  1.00 32.89           C
    ATOM   1075  H   VAL A 431      -8.584  35.417  87.986  1.00  0.00           H
    ATOM   1076  HA  VAL A 431      -8.001  36.789  90.404  1.00  0.00           H
    ATOM   1077  HB  VAL A 431      -6.584  36.603  88.289  1.00  0.00           H
    ATOM   1078 HG11 VAL A 431      -6.754  38.321  86.556  1.00  0.00           H
    ATOM   1079 HG12 VAL A 431      -8.200  37.326  86.570  1.00  0.00           H
    ATOM   1080 HG13 VAL A 431      -8.223  38.952  87.246  1.00  0.00           H
    ATOM   1081 HG21 VAL A 431      -5.458  38.781  88.655  1.00  0.00           H
    ATOM   1082 HG22 VAL A 431      -6.837  39.337  89.593  1.00  0.00           H
    ATOM   1083 HG23 VAL A 431      -5.879  37.993  90.168  1.00  0.00           H
    ATOM   1084  N   THR A 432     -10.515  37.975  88.673  1.00 39.32           N
    ATOM   1085  CA  THR A 432     -11.705  38.824  88.696  1.00 40.31           C
    ATOM   1086  C   THR A 432     -12.634  38.568  89.909  1.00 41.30           C
    ATOM   1087  O   THR A 432     -13.578  39.335  90.090  1.00 45.27           O
    ATOM   1088  CB  THR A 432     -12.531  38.625  87.399  1.00 36.81           C
    ATOM   1089  CG2 THR A 432     -11.759  38.981  86.122  1.00 23.41           C
    ATOM   1090  OG1 THR A 432     -13.015  37.300  87.279  1.00 38.18           O
    ATOM   1091  H   THR A 432     -10.536  37.226  87.996  1.00  0.00           H
    ATOM   1092  HA  THR A 432     -11.373  39.861  88.745  1.00  0.00           H
    ATOM   1093  HB  THR A 432     -13.404  39.277  87.436  1.00  0.00           H
    ATOM   1094 HG21 THR A 432     -12.388  38.838  85.244  1.00  0.00           H
    ATOM   1095 HG22 THR A 432     -11.447  40.024  86.136  1.00  0.00           H
    ATOM   1096 HG23 THR A 432     -10.868  38.365  85.995  1.00  0.00           H
    ATOM   1097  HG1 THR A 432     -13.679  37.149  87.957  1.00  0.00           H
    ATOM   1098  N   SER A 433     -12.354  37.537  90.728  1.00 42.09           N
    ATOM   1099  CA  SER A 433     -13.018  37.315  92.016  1.00 40.87           C
    ATOM   1100  C   SER A 433     -12.321  38.071  93.161  1.00 44.77           C
    ATOM   1101  O   SER A 433     -12.944  38.253  94.208  1.00 45.62           O
    ATOM   1102  CB  SER A 433     -13.043  35.811  92.349  1.00 45.91           C
    ATOM   1103  OG  SER A 433     -13.825  35.094  91.421  1.00 58.23           O
    ATOM   1104  H   SER A 433     -11.570  36.932  90.524  1.00  0.00           H
    ATOM   1105  HA  SER A 433     -14.049  37.672  91.968  1.00  0.00           H
    ATOM   1106  HB2 SER A 433     -12.041  35.390  92.390  1.00  0.00           H
    ATOM   1107  HB3 SER A 433     -13.486  35.651  93.333  1.00  0.00           H
    ATOM   1108  HG  SER A 433     -14.746  35.344  91.533  1.00  0.00           H
    ATOM   1109  N   THR A 434     -11.049  38.473  92.970  1.00 40.15           N
    ATOM   1110  CA  THR A 434     -10.257  39.165  93.987  1.00 33.49           C
    ATOM   1111  C   THR A 434     -10.633  40.666  94.054  1.00 34.32           C
    ATOM   1112  O   THR A 434     -10.933  41.249  93.009  1.00 27.73           O
    ATOM   1113  CB  THR A 434      -8.731  39.029  93.743  1.00 30.30           C
    ATOM   1114  CG2 THR A 434      -8.257  37.586  93.521  1.00 28.46           C
    ATOM   1115  OG1 THR A 434      -8.224  39.868  92.725  1.00 31.94           O
    ATOM   1116  H   THR A 434     -10.600  38.325  92.077  1.00  0.00           H
    ATOM   1117  HA  THR A 434     -10.477  38.649  94.914  1.00  0.00           H
    ATOM   1118  HB  THR A 434      -8.249  39.373  94.659  1.00  0.00           H
    ATOM   1119 HG21 THR A 434      -7.180  37.545  93.357  1.00  0.00           H
    ATOM   1120 HG22 THR A 434      -8.474  36.968  94.391  1.00  0.00           H
    ATOM   1121 HG23 THR A 434      -8.736  37.127  92.656  1.00  0.00           H
    ATOM   1122  HG1 THR A 434      -8.598  39.594  91.880  1.00  0.00           H
    ATOM   1123  N   PRO A 435     -10.640  41.265  95.269  1.00 30.23           N
    ATOM   1124  CA  PRO A 435     -11.086  42.658  95.448  1.00 28.26           C
    ATOM   1125  C   PRO A 435     -10.165  43.733  94.846  1.00 31.83           C
    ATOM   1126  O   PRO A 435     -10.672  44.772  94.433  1.00 32.76           O
    ATOM   1127  CB  PRO A 435     -11.231  42.825  96.972  1.00 31.08           C
    ATOM   1128  CG  PRO A 435     -11.196  41.424  97.545  1.00 33.02           C
    ATOM   1129  CD  PRO A 435     -10.310  40.676  96.567  1.00 29.78           C
    ATOM   1130  HA  PRO A 435     -12.072  42.748  94.995  1.00  0.00           H
    ATOM   1131  HB2 PRO A 435     -10.410  43.411  97.386  1.00  0.00           H
    ATOM   1132  HB3 PRO A 435     -12.164  43.315  97.241  1.00  0.00           H
    ATOM   1133  HG2 PRO A 435     -10.846  41.385  98.576  1.00  0.00           H
    ATOM   1134  HG3 PRO A 435     -12.201  40.999  97.519  1.00  0.00           H
    ATOM   1135  HD2 PRO A 435      -9.256  40.867  96.773  1.00  0.00           H
    ATOM   1136  HD3 PRO A 435     -10.493  39.612  96.682  1.00  0.00           H
    ATOM   1137  N   CYS A 436      -8.845  43.485  94.830  1.00 34.77           N
    ATOM   1138  CA  CYS A 436      -7.835  44.444  94.386  1.00 30.14           C
    ATOM   1139  C   CYS A 436      -6.768  43.675  93.615  1.00 32.19           C
    ATOM   1140  O   CYS A 436      -6.115  42.813  94.203  1.00 38.02           O
    ATOM   1141  CB  CYS A 436      -7.163  45.169  95.569  1.00 27.42           C
    ATOM   1142  SG  CYS A 436      -8.334  46.262  96.399  1.00 33.15           S
    ATOM   1143  H   CYS A 436      -8.493  42.602  95.172  1.00  0.00           H
    ATOM   1144  HA  CYS A 436      -8.275  45.186  93.716  1.00  0.00           H
    ATOM   1145  HB2 CYS A 436      -6.775  44.454  96.296  1.00  0.00           H
    ATOM   1146  HB3 CYS A 436      -6.322  45.776  95.230  1.00  0.00           H
    ATOM   1147  HG  CYS A 436      -8.430  47.148  95.401  1.00  0.00           H
    ATOM   1148  N   TRP A 437      -6.584  44.022  92.334  1.00 30.62           N
    ATOM   1149  CA  TRP A 437      -5.562  43.421  91.484  1.00 30.30           C
    ATOM   1150  C   TRP A 437      -5.172  44.370  90.349  1.00 31.06           C
    ATOM   1151  O   TRP A 437      -5.961  45.233  89.970  1.00 30.37           O
    ATOM   1152  CB  TRP A 437      -6.032  42.039  90.966  1.00 27.52           C
    ATOM   1153  CG  TRP A 437      -7.216  42.003  90.048  1.00 20.91           C
    ATOM   1154  CD1 TRP A 437      -8.506  42.151  90.424  1.00 26.93           C
    ATOM   1155  CD2 TRP A 437      -7.236  41.826  88.596  1.00 21.71           C
    ATOM   1156  CE2 TRP A 437      -8.594  41.849  88.161  1.00 20.34           C
    ATOM   1157  CE3 TRP A 437      -6.250  41.636  87.600  1.00 24.32           C
    ATOM   1158  NE1 TRP A 437      -9.314  42.044  89.317  1.00 17.08           N
    ATOM   1159  CZ2 TRP A 437      -8.957  41.711  86.811  1.00 21.53           C
    ATOM   1160  CZ3 TRP A 437      -6.600  41.489  86.242  1.00 31.27           C
    ATOM   1161  CH2 TRP A 437      -7.949  41.536  85.845  1.00 28.09           C
    ATOM   1162  H   TRP A 437      -7.156  44.739  91.906  1.00  0.00           H
    ATOM   1163  HA  TRP A 437      -4.670  43.297  92.098  1.00  0.00           H
    ATOM   1164  HB2 TRP A 437      -5.221  41.558  90.423  1.00  0.00           H
    ATOM   1165  HB3 TRP A 437      -6.235  41.379  91.808  1.00  0.00           H
    ATOM   1166  HD1 TRP A 437      -8.835  42.314  91.440  1.00  0.00           H
    ATOM   1167  HE1 TRP A 437     -10.325  42.129  89.360  1.00  0.00           H
    ATOM   1168  HE3 TRP A 437      -5.210  41.595  87.887  1.00  0.00           H
    ATOM   1169  HZ2 TRP A 437      -9.995  41.760  86.521  1.00  0.00           H
    ATOM   1170  HZ3 TRP A 437      -5.831  41.336  85.501  1.00  0.00           H
    ATOM   1171  HH2 TRP A 437      -8.204  41.437  84.801  1.00  0.00           H
    ATOM   1172  N   ILE A 438      -3.957  44.169  89.819  1.00 25.24           N
    ATOM   1173  CA  ILE A 438      -3.442  44.864  88.644  1.00 23.29           C
    ATOM   1174  C   ILE A 438      -3.028  43.865  87.551  1.00 24.76           C
    ATOM   1175  O   ILE A 438      -2.840  42.682  87.835  1.00 27.00           O
    ATOM   1176  CB  ILE A 438      -2.222  45.782  88.964  1.00 21.22           C
    ATOM   1177  CG1 ILE A 438      -0.891  45.103  89.394  1.00 15.06           C
    ATOM   1178  CG2 ILE A 438      -2.596  46.973  89.859  1.00 15.20           C
    ATOM   1179  CD1 ILE A 438      -0.690  44.928  90.905  1.00 22.20           C
    ATOM   1180  H   ILE A 438      -3.375  43.425  90.184  1.00  0.00           H
    ATOM   1181  HA  ILE A 438      -4.236  45.477  88.222  1.00  0.00           H
    ATOM   1182  HB  ILE A 438      -1.980  46.251  88.011  1.00  0.00           H
    ATOM   1183 HG12 ILE A 438      -0.738  44.144  88.901  1.00  0.00           H
    ATOM   1184 HG13 ILE A 438      -0.070  45.731  89.045  1.00  0.00           H
    ATOM   1185 HG21 ILE A 438      -1.752  47.651  89.962  1.00  0.00           H
    ATOM   1186 HG22 ILE A 438      -3.396  47.560  89.422  1.00  0.00           H
    ATOM   1187 HG23 ILE A 438      -2.916  46.661  90.852  1.00  0.00           H
    ATOM   1188 HD11 ILE A 438       0.266  44.447  91.108  1.00  0.00           H
    ATOM   1189 HD12 ILE A 438      -0.665  45.887  91.420  1.00  0.00           H
    ATOM   1190 HD13 ILE A 438      -1.482  44.324  91.348  1.00  0.00           H
    ATOM   1191  N   GLU A 439      -2.877  44.389  86.327  1.00 31.80           N
    ATOM   1192  CA  GLU A 439      -2.383  43.680  85.155  1.00 32.54           C
    ATOM   1193  C   GLU A 439      -1.285  44.526  84.499  1.00 35.92           C
    ATOM   1194  O   GLU A 439      -1.576  45.603  83.979  1.00 31.77           O
    ATOM   1195  CB  GLU A 439      -3.559  43.394  84.202  1.00 39.20           C
    ATOM   1196  CG  GLU A 439      -3.158  42.675  82.897  1.00 47.12           C
    ATOM   1197  CD  GLU A 439      -4.368  42.336  82.028  1.00 50.45           C
    ATOM   1198  OE1 GLU A 439      -5.210  43.237  81.821  1.00 46.72           O
    ATOM   1199  OE2 GLU A 439      -4.429  41.171  81.577  1.00 55.44           O1-
    ATOM   1200  H   GLU A 439      -3.092  45.368  86.183  1.00  0.00           H
    ATOM   1201  HA  GLU A 439      -1.952  42.723  85.451  1.00  0.00           H
    ATOM   1202  HB2 GLU A 439      -4.299  42.793  84.729  1.00  0.00           H
    ATOM   1203  HB3 GLU A 439      -4.062  44.332  83.960  1.00  0.00           H
    ATOM   1204  HG2 GLU A 439      -2.487  43.295  82.303  1.00  0.00           H
    ATOM   1205  HG3 GLU A 439      -2.611  41.760  83.130  1.00  0.00           H
    ATOM   1206  N   LEU A 440      -0.052  44.002  84.526  1.00 40.14           N
    ATOM   1207  CA  LEU A 440       1.111  44.547  83.832  1.00 42.22           C
    ATOM   1208  C   LEU A 440       1.114  44.072  82.373  1.00 41.26           C
    ATOM   1209  O   LEU A 440       0.670  42.959  82.100  1.00 39.30           O
    ATOM   1210  CB  LEU A 440       2.411  44.069  84.516  1.00 45.50           C
    ATOM   1211  CG  LEU A 440       2.721  44.723  85.879  1.00 49.80           C
    ATOM   1212  CD1 LEU A 440       1.797  44.237  87.012  1.00 54.01           C
    ATOM   1213  CD2 LEU A 440       4.211  44.549  86.237  1.00 48.08           C
    ATOM   1214  H   LEU A 440       0.088  43.099  84.964  1.00  0.00           H
    ATOM   1215  HA  LEU A 440       1.074  45.635  83.866  1.00  0.00           H
    ATOM   1216  HB2 LEU A 440       2.422  42.984  84.605  1.00  0.00           H
    ATOM   1217  HB3 LEU A 440       3.236  44.315  83.844  1.00  0.00           H
    ATOM   1218  HG  LEU A 440       2.558  45.792  85.772  1.00  0.00           H
    ATOM   1219 HD11 LEU A 440       2.240  44.397  87.996  1.00  0.00           H
    ATOM   1220 HD12 LEU A 440       0.856  44.785  87.001  1.00  0.00           H
    ATOM   1221 HD13 LEU A 440       1.565  43.177  86.921  1.00  0.00           H
    ATOM   1222 HD21 LEU A 440       4.353  43.997  87.163  1.00  0.00           H
    ATOM   1223 HD22 LEU A 440       4.766  44.009  85.469  1.00  0.00           H
    ATOM   1224 HD23 LEU A 440       4.696  45.518  86.354  1.00  0.00           H
    ATOM   1225  N   HIS A 441       1.661  44.909  81.482  1.00 42.86           N
    ATOM   1226  CA  HIS A 441       1.991  44.595  80.092  1.00 42.11           C
    ATOM   1227  C   HIS A 441       3.435  45.036  79.878  1.00 46.76           C
    ATOM   1228  O   HIS A 441       3.707  46.230  79.971  1.00 43.77           O
    ATOM   1229  CB  HIS A 441       1.074  45.371  79.120  1.00 48.57           C
    ATOM   1230  CG  HIS A 441      -0.317  44.833  78.914  1.00 59.31           C
    ATOM   1231  CD2 HIS A 441      -1.256  45.111  77.945  1.00 61.01           C
    ATOM   1232  ND1 HIS A 441      -0.902  43.883  79.732  1.00 58.44           N
    ATOM   1233  CE1 HIS A 441      -2.124  43.648  79.255  1.00 60.21           C
    ATOM   1234  NE2 HIS A 441      -2.410  44.357  78.170  1.00 58.73           N
    ATOM   1235  H   HIS A 441       1.949  45.827  81.793  1.00  0.00           H
    ATOM   1236  HA  HIS A 441       1.916  43.524  79.892  1.00  0.00           H
    ATOM   1237  HB2 HIS A 441       0.982  46.405  79.448  1.00  0.00           H
    ATOM   1238  HB3 HIS A 441       1.534  45.414  78.132  1.00  0.00           H
    ATOM   1239  HD1 HIS A 441      -0.484  43.459  80.551  1.00  0.00           H
    ATOM   1240  HD2 HIS A 441      -1.192  45.791  77.107  1.00  0.00           H
    ATOM   1241  HE1 HIS A 441      -2.808  42.950  79.711  1.00  0.00           H
    ATOM   1242  N   LEU A 442       4.329  44.077  79.602  1.00 44.34           N
    ATOM   1243  CA  LEU A 442       5.749  44.326  79.378  1.00 39.79           C
    ATOM   1244  C   LEU A 442       5.960  44.760  77.926  1.00 41.74           C
    ATOM   1245  O   LEU A 442       5.803  43.948  77.013  1.00 39.12           O
    ATOM   1246  CB  LEU A 442       6.550  43.058  79.715  1.00 42.64           C
    ATOM   1247  CG  LEU A 442       6.364  42.569  81.163  1.00 47.39           C
    ATOM   1248  CD1 LEU A 442       7.068  41.219  81.343  1.00 50.13           C
    ATOM   1249  CD2 LEU A 442       6.822  43.617  82.199  1.00 40.71           C
    ATOM   1250  H   LEU A 442       4.029  43.112  79.546  1.00  0.00           H
    ATOM   1251  HA  LEU A 442       6.083  45.131  80.036  1.00  0.00           H
    ATOM   1252  HB2 LEU A 442       6.266  42.255  79.031  1.00  0.00           H
    ATOM   1253  HB3 LEU A 442       7.609  43.249  79.545  1.00  0.00           H
    ATOM   1254  HG  LEU A 442       5.305  42.383  81.347  1.00  0.00           H
    ATOM   1255 HD11 LEU A 442       7.625  41.161  82.277  1.00  0.00           H
    ATOM   1256 HD12 LEU A 442       6.346  40.402  81.334  1.00  0.00           H
    ATOM   1257 HD13 LEU A 442       7.772  41.022  80.537  1.00  0.00           H
    ATOM   1258 HD21 LEU A 442       7.237  43.164  83.098  1.00  0.00           H
    ATOM   1259 HD22 LEU A 442       7.575  44.293  81.792  1.00  0.00           H
    ATOM   1260 HD23 LEU A 442       5.976  44.226  82.522  1.00  0.00           H
    ATOM   1261  N   ASN A 443       6.277  46.051  77.751  1.00 45.59           N
    ATOM   1262  CA  ASN A 443       6.295  46.715  76.450  1.00 43.67           C
    ATOM   1263  C   ASN A 443       7.468  46.290  75.553  1.00 45.68           C
    ATOM   1264  O   ASN A 443       7.286  46.247  74.338  1.00 50.44           O
    ATOM   1265  CB  ASN A 443       6.271  48.248  76.646  1.00 46.88           C
    ATOM   1266  CG  ASN A 443       4.975  48.780  77.272  1.00 53.19           C
    ATOM   1267  ND2 ASN A 443       5.064  49.905  77.980  1.00 52.30           N
    ATOM   1268  OD1 ASN A 443       3.902  48.203  77.109  1.00 52.31           O
    ATOM   1269  H   ASN A 443       6.415  46.644  78.558  1.00  0.00           H
    ATOM   1270  HA  ASN A 443       5.363  46.395  75.989  1.00  0.00           H
    ATOM   1271  HB2 ASN A 443       7.125  48.551  77.252  1.00  0.00           H
    ATOM   1272  HB3 ASN A 443       6.385  48.748  75.683  1.00  0.00           H
    ATOM   1273 HD21 ASN A 443       5.958  50.369  78.085  1.00  0.00           H
    ATOM   1274 HD22 ASN A 443       4.239  50.303  78.404  1.00  0.00           H
    ATOM   1275  N   GLY A 444       8.632  45.969  76.146  1.00 48.07           N
    ATOM   1276  CA  GLY A 444       9.812  45.485  75.421  1.00 44.17           C
    ATOM   1277  C   GLY A 444       9.545  44.103  74.796  1.00 44.39           C
    ATOM   1278  O   GLY A 444       9.650  43.989  73.575  1.00 47.42           O
    ATOM   1279  H   GLY A 444       8.700  46.026  77.152  1.00  0.00           H
    ATOM   1280  HA2 GLY A 444      10.087  46.197  74.642  1.00  0.00           H
    ATOM   1281  HA3 GLY A 444      10.659  45.425  76.104  1.00  0.00           H
    ATOM   1282  N   PRO A 445       9.126  43.087  75.588  1.00 42.41           N
    ATOM   1283  CA  PRO A 445       8.588  41.802  75.104  1.00 45.35           C
    ATOM   1284  C   PRO A 445       7.495  41.870  74.024  1.00 51.62           C
    ATOM   1285  O   PRO A 445       7.648  41.183  73.017  1.00 53.79           O
    ATOM   1286  CB  PRO A 445       8.124  41.102  76.387  1.00 49.03           C
    ATOM   1287  CG  PRO A 445       9.117  41.590  77.423  1.00 45.85           C
    ATOM   1288  CD  PRO A 445       9.314  43.050  77.039  1.00 42.40           C
    ATOM   1289  HA  PRO A 445       9.429  41.243  74.695  1.00  0.00           H
    ATOM   1290  HB2 PRO A 445       7.120  41.427  76.667  1.00  0.00           H
    ATOM   1291  HB3 PRO A 445       8.111  40.015  76.304  1.00  0.00           H
    ATOM   1292  HG2 PRO A 445       8.782  41.453  78.449  1.00  0.00           H
    ATOM   1293  HG3 PRO A 445      10.057  41.051  77.300  1.00  0.00           H
    ATOM   1294  HD2 PRO A 445       8.559  43.671  77.513  1.00  0.00           H
    ATOM   1295  HD3 PRO A 445      10.297  43.406  77.352  1.00  0.00           H
    ATOM   1296  N   LEU A 446       6.458  42.711  74.210  1.00 54.88           N
    ATOM   1297  CA  LEU A 446       5.395  42.939  73.218  1.00 51.50           C
    ATOM   1298  C   LEU A 446       5.916  43.516  71.888  1.00 51.46           C
    ATOM   1299  O   LEU A 446       5.438  43.106  70.834  1.00 47.30           O
    ATOM   1300  CB  LEU A 446       4.310  43.880  73.790  1.00 49.51           C
    ATOM   1301  CG  LEU A 446       3.313  43.235  74.765  1.00 43.76           C
    ATOM   1302  CD1 LEU A 446       2.479  44.317  75.477  1.00 45.81           C
    ATOM   1303  CD2 LEU A 446       2.413  42.193  74.061  1.00 44.20           C
    ATOM   1304  H   LEU A 446       6.387  43.238  75.070  1.00  0.00           H
    ATOM   1305  HA  LEU A 446       4.946  41.970  72.990  1.00  0.00           H
    ATOM   1306  HB2 LEU A 446       4.796  44.730  74.268  1.00  0.00           H
    ATOM   1307  HB3 LEU A 446       3.716  44.291  72.980  1.00  0.00           H
    ATOM   1308  HG  LEU A 446       3.889  42.724  75.530  1.00  0.00           H
    ATOM   1309 HD11 LEU A 446       1.531  43.930  75.853  1.00  0.00           H
    ATOM   1310 HD12 LEU A 446       3.024  44.717  76.333  1.00  0.00           H
    ATOM   1311 HD13 LEU A 446       2.253  45.151  74.813  1.00  0.00           H
    ATOM   1312 HD21 LEU A 446       1.352  42.361  74.247  1.00  0.00           H
    ATOM   1313 HD22 LEU A 446       2.546  42.206  72.979  1.00  0.00           H
    ATOM   1314 HD23 LEU A 446       2.642  41.182  74.393  1.00  0.00           H
    ATOM   1315  N   GLN A 447       6.894  44.433  71.953  1.00 52.93           N
    ATOM   1316  CA  GLN A 447       7.524  45.045  70.784  1.00 54.87           C
    ATOM   1317  C   GLN A 447       8.376  44.055  69.963  1.00 56.39           C
    ATOM   1318  O   GLN A 447       8.332  44.118  68.736  1.00 48.65           O
    ATOM   1319  CB  GLN A 447       8.293  46.303  71.234  1.00 54.23           C
    ATOM   1320  CG  GLN A 447       9.056  47.034  70.117  1.00 61.62           C
    ATOM   1321  CD  GLN A 447       9.624  48.361  70.611  1.00 69.35           C
    ATOM   1322  NE2 GLN A 447      10.882  48.354  71.052  1.00 71.95           N
    ATOM   1323  OE1 GLN A 447       8.938  49.380  70.603  1.00 70.13           O
    ATOM   1324  H   GLN A 447       7.234  44.739  72.855  1.00  0.00           H
    ATOM   1325  HA  GLN A 447       6.719  45.385  70.127  1.00  0.00           H
    ATOM   1326  HB2 GLN A 447       7.576  46.996  71.679  1.00  0.00           H
    ATOM   1327  HB3 GLN A 447       8.991  46.044  72.031  1.00  0.00           H
    ATOM   1328  HG2 GLN A 447       9.868  46.417  69.734  1.00  0.00           H
    ATOM   1329  HG3 GLN A 447       8.388  47.235  69.278  1.00  0.00           H
    ATOM   1330 HE21 GLN A 447      11.415  47.497  71.068  1.00  0.00           H
    ATOM   1331 HE22 GLN A 447      11.293  49.211  71.389  1.00  0.00           H
    ATOM   1332  N   TRP A 448       9.096  43.136  70.634  1.00 54.46           N
    ATOM   1333  CA  TRP A 448       9.845  42.050  69.985  1.00 55.19           C
    ATOM   1334  C   TRP A 448       8.931  40.997  69.348  1.00 56.07           C
    ATOM   1335  O   TRP A 448       9.260  40.485  68.279  1.00 57.80           O
    ATOM   1336  CB  TRP A 448      10.783  41.361  70.991  1.00 57.31           C
    ATOM   1337  CG  TRP A 448      11.765  42.233  71.706  1.00 65.68           C
    ATOM   1338  CD1 TRP A 448      12.397  43.314  71.191  1.00 68.37           C
    ATOM   1339  CD2 TRP A 448      12.292  42.064  73.053  1.00 72.40           C
    ATOM   1340  CE2 TRP A 448      13.210  43.126  73.313  1.00 77.05           C
    ATOM   1341  CE3 TRP A 448      12.081  41.126  74.091  1.00 72.28           C
    ATOM   1342  NE1 TRP A 448      13.238  43.852  72.141  1.00 74.33           N
    ATOM   1343  CZ2 TRP A 448      13.867  43.264  74.545  1.00 80.04           C
    ATOM   1344  CZ3 TRP A 448      12.726  41.258  75.335  1.00 75.71           C
    ATOM   1345  CH2 TRP A 448      13.617  42.326  75.560  1.00 77.22           C
    ATOM   1346  H   TRP A 448       9.097  43.145  71.645  1.00  0.00           H
    ATOM   1347  HA  TRP A 448      10.451  42.486  69.189  1.00  0.00           H
    ATOM   1348  HB2 TRP A 448      10.188  40.858  71.755  1.00  0.00           H
    ATOM   1349  HB3 TRP A 448      11.351  40.576  70.488  1.00  0.00           H
    ATOM   1350  HD1 TRP A 448      12.244  43.696  70.193  1.00  0.00           H
    ATOM   1351  HE1 TRP A 448      13.803  44.672  71.975  1.00  0.00           H
    ATOM   1352  HE3 TRP A 448      11.404  40.300  73.929  1.00  0.00           H
    ATOM   1353  HZ2 TRP A 448      14.554  44.080  74.713  1.00  0.00           H
    ATOM   1354  HZ3 TRP A 448      12.536  40.535  76.115  1.00  0.00           H
    ATOM   1355  HH2 TRP A 448      14.122  42.432  76.505  1.00  0.00           H
    ATOM   1356  N   LEU A 449       7.796  40.720  70.013  1.00 57.37           N
    ATOM   1357  CA  LEU A 449       6.730  39.839  69.543  1.00 60.09           C
    ATOM   1358  C   LEU A 449       6.135  40.350  68.223  1.00 63.10           C
    ATOM   1359  O   LEU A 449       5.981  39.560  67.294  1.00 64.59           O
    ATOM   1360  CB  LEU A 449       5.655  39.731  70.647  1.00 56.52           C
    ATOM   1361  CG  LEU A 449       4.846  38.423  70.692  1.00 58.24           C
    ATOM   1362  CD1 LEU A 449       3.838  38.502  71.849  1.00 62.15           C
    ATOM   1363  CD2 LEU A 449       4.150  38.071  69.371  1.00 56.15           C
    ATOM   1364  H   LEU A 449       7.626  41.177  70.899  1.00  0.00           H
    ATOM   1365  HA  LEU A 449       7.164  38.855  69.373  1.00  0.00           H
    ATOM   1366  HB2 LEU A 449       6.138  39.809  71.619  1.00  0.00           H
    ATOM   1367  HB3 LEU A 449       4.979  40.584  70.595  1.00  0.00           H
    ATOM   1368  HG  LEU A 449       5.534  37.606  70.915  1.00  0.00           H
    ATOM   1369 HD11 LEU A 449       2.883  38.038  71.610  1.00  0.00           H
    ATOM   1370 HD12 LEU A 449       4.247  38.000  72.723  1.00  0.00           H
    ATOM   1371 HD13 LEU A 449       3.625  39.529  72.149  1.00  0.00           H
    ATOM   1372 HD21 LEU A 449       3.222  37.525  69.517  1.00  0.00           H
    ATOM   1373 HD22 LEU A 449       3.887  38.962  68.806  1.00  0.00           H
    ATOM   1374 HD23 LEU A 449       4.801  37.456  68.752  1.00  0.00           H
    ATOM   1375  N   ASP A 450       5.846  41.661  68.173  1.00 67.93           N
    ATOM   1376  CA  ASP A 450       5.279  42.367  67.029  1.00 71.35           C
    ATOM   1377  C   ASP A 450       6.097  42.194  65.738  1.00 72.43           C
    ATOM   1378  O   ASP A 450       5.511  41.880  64.703  1.00 72.00           O
    ATOM   1379  CB  ASP A 450       5.038  43.851  67.353  1.00 70.48           C
    ATOM   1380  CG  ASP A 450       4.460  44.615  66.168  1.00 66.33           C
    ATOM   1381  OD1 ASP A 450       3.228  44.532  65.981  1.00 60.35           O
    ATOM   1382  OD2 ASP A 450       5.265  45.257  65.461  1.00 64.59           O1-
    ATOM   1383  H   ASP A 450       6.003  42.226  68.997  1.00  0.00           H
    ATOM   1384  HA  ASP A 450       4.304  41.918  66.843  1.00  0.00           H
    ATOM   1385  HB2 ASP A 450       4.372  43.965  68.206  1.00  0.00           H
    ATOM   1386  HB3 ASP A 450       5.958  44.344  67.665  1.00  0.00           H
    ATOM   1387  N   LYS A 451       7.423  42.387  65.835  1.00 74.16           N
    ATOM   1388  CA  LYS A 451       8.372  42.246  64.728  1.00 75.51           C
    ATOM   1389  C   LYS A 451       8.289  40.876  64.037  1.00 76.01           C
    ATOM   1390  O   LYS A 451       8.360  40.819  62.811  1.00 77.80           O
    ATOM   1391  CB  LYS A 451       9.802  42.490  65.242  1.00 73.06           C
    ATOM   1392  CG  LYS A 451      10.049  43.928  65.720  1.00 77.65           C
    ATOM   1393  CD  LYS A 451      11.419  44.079  66.392  1.00 82.36           C
    ATOM   1394  CE  LYS A 451      11.621  45.471  66.999  1.00 84.20           C
    ATOM   1395  NZ  LYS A 451      12.960  45.597  67.596  1.00 91.18           N1+
    ATOM   1396  H   LYS A 451       7.815  42.653  66.728  1.00  0.00           H
    ATOM   1397  HA  LYS A 451       8.128  43.010  63.988  1.00  0.00           H
    ATOM   1398  HB2 LYS A 451      10.019  41.791  66.050  1.00  0.00           H
    ATOM   1399  HB3 LYS A 451      10.521  42.267  64.451  1.00  0.00           H
    ATOM   1400  HG2 LYS A 451       9.980  44.608  64.870  1.00  0.00           H
    ATOM   1401  HG3 LYS A 451       9.267  44.237  66.410  1.00  0.00           H
    ATOM   1402  HD2 LYS A 451      11.530  43.323  67.170  1.00  0.00           H
    ATOM   1403  HD3 LYS A 451      12.201  43.882  65.658  1.00  0.00           H
    ATOM   1404  HE2 LYS A 451      11.503  46.240  66.234  1.00  0.00           H
    ATOM   1405  HE3 LYS A 451      10.870  45.659  67.765  1.00  0.00           H
    ATOM   1406  HZ1 LYS A 451      13.067  46.522  67.989  1.00  0.00           H
    ATOM   1407  HZ2 LYS A 451      13.073  44.905  68.322  1.00  0.00           H
    ATOM   1408  HZ3 LYS A 451      13.660  45.453  66.883  1.00  0.00           H
    ATOM   1409  N   VAL A 452       8.086  39.818  64.839  1.00 72.11           N
    ATOM   1410  CA  VAL A 452       7.860  38.459  64.365  1.00 66.49           C
    ATOM   1411  C   VAL A 452       6.429  38.290  63.805  1.00 69.63           C
    ATOM   1412  O   VAL A 452       6.307  37.776  62.696  1.00 66.05           O
    ATOM   1413  CB  VAL A 452       8.139  37.413  65.484  1.00 63.19           C
    ATOM   1414  CG1 VAL A 452       7.884  35.952  65.065  1.00 57.82           C
    ATOM   1415  CG2 VAL A 452       9.583  37.537  66.009  1.00 54.88           C
    ATOM   1416  H   VAL A 452       8.023  39.957  65.838  1.00  0.00           H
    ATOM   1417  HA  VAL A 452       8.559  38.266  63.548  1.00  0.00           H
    ATOM   1418  HB  VAL A 452       7.475  37.620  66.323  1.00  0.00           H
    ATOM   1419 HG11 VAL A 452       8.126  35.265  65.877  1.00  0.00           H
    ATOM   1420 HG12 VAL A 452       6.844  35.771  64.805  1.00  0.00           H
    ATOM   1421 HG13 VAL A 452       8.493  35.676  64.203  1.00  0.00           H
    ATOM   1422 HG21 VAL A 452       9.791  36.784  66.769  1.00  0.00           H
    ATOM   1423 HG22 VAL A 452      10.309  37.403  65.206  1.00  0.00           H
    ATOM   1424 HG23 VAL A 452       9.767  38.509  66.465  1.00  0.00           H
    ATOM   1425  N   LEU A 453       5.393  38.757  64.533  1.00 72.12           N
    ATOM   1426  CA  LEU A 453       3.967  38.648  64.176  1.00 73.92           C
    ATOM   1427  C   LEU A 453       3.587  39.247  62.810  1.00 75.44           C
    ATOM   1428  O   LEU A 453       2.840  38.607  62.070  1.00 76.17           O
    ATOM   1429  CB  LEU A 453       3.074  39.293  65.270  1.00 67.00           C
    ATOM   1430  CG  LEU A 453       2.333  38.322  66.211  1.00 61.81           C
    ATOM   1431  CD1 LEU A 453       1.375  39.095  67.142  1.00 64.97           C
    ATOM   1432  CD2 LEU A 453       1.601  37.176  65.480  1.00 56.40           C
    ATOM   1433  H   LEU A 453       5.585  39.188  65.429  1.00  0.00           H
    ATOM   1434  HA  LEU A 453       3.768  37.578  64.113  1.00  0.00           H
    ATOM   1435  HB2 LEU A 453       3.664  39.987  65.866  1.00  0.00           H
    ATOM   1436  HB3 LEU A 453       2.317  39.921  64.807  1.00  0.00           H
    ATOM   1437  HG  LEU A 453       3.097  37.878  66.840  1.00  0.00           H
    ATOM   1438 HD11 LEU A 453       1.577  38.872  68.189  1.00  0.00           H
    ATOM   1439 HD12 LEU A 453       1.466  40.172  67.011  1.00  0.00           H
    ATOM   1440 HD13 LEU A 453       0.327  38.856  66.962  1.00  0.00           H
    ATOM   1441 HD21 LEU A 453       0.619  36.960  65.900  1.00  0.00           H
    ATOM   1442 HD22 LEU A 453       1.450  37.406  64.426  1.00  0.00           H
    ATOM   1443 HD23 LEU A 453       2.172  36.250  65.532  1.00  0.00           H
    ATOM   1444  N   THR A 454       4.116  40.443  62.497  1.00 77.79           N
    ATOM   1445  CA  THR A 454       3.929  41.153  61.222  1.00 80.23           C
    ATOM   1446  C   THR A 454       4.537  40.422  59.995  1.00 82.85           C
    ATOM   1447  O   THR A 454       4.331  40.882  58.871  1.00 84.31           O
    ATOM   1448  CB  THR A 454       4.528  42.588  61.290  1.00 80.08           C
    ATOM   1449  CG2 THR A 454       3.834  43.498  62.318  1.00 79.86           C
    ATOM   1450  OG1 THR A 454       5.922  42.564  61.548  1.00 76.52           O
    ATOM   1451  H   THR A 454       4.708  40.905  63.175  1.00  0.00           H
    ATOM   1452  HA  THR A 454       2.857  41.240  61.044  1.00  0.00           H
    ATOM   1453  HB  THR A 454       4.411  43.072  60.319  1.00  0.00           H
    ATOM   1454 HG21 THR A 454       4.338  44.461  62.402  1.00  0.00           H
    ATOM   1455 HG22 THR A 454       2.806  43.697  62.017  1.00  0.00           H
    ATOM   1456 HG23 THR A 454       3.793  43.058  63.313  1.00  0.00           H
    ATOM   1457  HG1 THR A 454       6.059  42.353  62.476  1.00  0.00           H
    ATOM   1458  N   GLN A 455       5.236  39.297  60.228  1.00 85.75           N
    ATOM   1459  CA  GLN A 455       5.878  38.445  59.229  1.00 87.89           C
    ATOM   1460  C   GLN A 455       5.345  36.995  59.247  1.00 88.40           C
    ATOM   1461  O   GLN A 455       5.621  36.272  58.290  1.00 87.63           O
    ATOM   1462  CB  GLN A 455       7.397  38.451  59.492  1.00 86.56           C
    ATOM   1463  CG  GLN A 455       8.041  39.850  59.403  1.00 89.46           C
    ATOM   1464  CD  GLN A 455       9.501  39.874  59.861  1.00 93.39           C
    ATOM   1465  NE2 GLN A 455      10.109  41.061  59.840  1.00 93.63           N
    ATOM   1466  OE1 GLN A 455      10.074  38.854  60.237  1.00 98.47           O
    ATOM   1467  H   GLN A 455       5.369  39.004  61.186  1.00  0.00           H
    ATOM   1468  HA  GLN A 455       5.698  38.838  58.226  1.00  0.00           H
    ATOM   1469  HB2 GLN A 455       7.589  38.024  60.477  1.00  0.00           H
    ATOM   1470  HB3 GLN A 455       7.895  37.790  58.781  1.00  0.00           H
    ATOM   1471  HG2 GLN A 455       7.977  40.225  58.381  1.00  0.00           H
    ATOM   1472  HG3 GLN A 455       7.506  40.561  60.033  1.00  0.00           H
    ATOM   1473 HE21 GLN A 455       9.611  41.886  59.541  1.00  0.00           H
    ATOM   1474 HE22 GLN A 455      11.070  41.138  60.139  1.00  0.00           H
    ATOM   1475  N   MET A 456       4.593  36.584  60.290  1.00 91.04           N
    ATOM   1476  CA  MET A 456       4.021  35.232  60.437  1.00 89.21           C
    ATOM   1477  C   MET A 456       2.858  34.903  59.471  1.00 87.86           C
    ATOM   1478  O   MET A 456       2.332  33.791  59.543  1.00 85.62           O
    ATOM   1479  CB  MET A 456       3.587  34.977  61.906  1.00 90.67           C
    ATOM   1480  CG  MET A 456       4.710  34.774  62.933  1.00 91.18           C
    ATOM   1481  SD  MET A 456       5.985  33.539  62.554  1.00 94.68           S
    ATOM   1482  CE  MET A 456       5.003  32.027  62.680  1.00 94.64           C
    ATOM   1483  H   MET A 456       4.391  37.231  61.039  1.00  0.00           H
    ATOM   1484  HA  MET A 456       4.803  34.515  60.181  1.00  0.00           H
    ATOM   1485  HB2 MET A 456       2.974  35.812  62.244  1.00  0.00           H
    ATOM   1486  HB3 MET A 456       2.943  34.101  61.967  1.00  0.00           H
    ATOM   1487  HG2 MET A 456       5.210  35.715  63.107  1.00  0.00           H
    ATOM   1488  HG3 MET A 456       4.262  34.507  63.890  1.00  0.00           H
    ATOM   1489  HE1 MET A 456       5.620  31.154  62.470  1.00  0.00           H
    ATOM   1490  HE2 MET A 456       4.599  31.929  63.685  1.00  0.00           H
    ATOM   1491  HE3 MET A 456       4.179  32.045  61.967  1.00  0.00           H
    ATOM   1492  N   GLY A 457       2.468  35.842  58.592  1.00 88.38           N
    ATOM   1493  CA  GLY A 457       1.373  35.664  57.642  1.00 90.03           C
    ATOM   1494  C   GLY A 457       0.042  35.914  58.359  1.00 92.14           C
    ATOM   1495  O   GLY A 457      -0.105  36.920  59.055  1.00 94.74           O
    ATOM   1496  H   GLY A 457       2.941  36.734  58.586  1.00  0.00           H
    ATOM   1497  HA2 GLY A 457       1.483  36.387  56.834  1.00  0.00           H
    ATOM   1498  HA3 GLY A 457       1.403  34.670  57.191  1.00  0.00           H
    ATOM   1499  N   SER A 458      -0.922  35.003  58.158  1.00 91.37           N
    ATOM   1500  CA  SER A 458      -2.264  35.049  58.744  1.00 92.66           C
    ATOM   1501  C   SER A 458      -2.710  33.637  59.196  1.00 92.16           C
    ATOM   1502  O   SER A 458      -2.084  32.655  58.789  1.00 91.21           O
    ATOM   1503  CB  SER A 458      -3.226  35.708  57.724  1.00 93.37           C
    ATOM   1504  OG  SER A 458      -3.585  34.828  56.673  1.00 94.51           O
    ATOM   1505  H   SER A 458      -0.736  34.198  57.577  1.00  0.00           H
    ATOM   1506  HA  SER A 458      -2.210  35.679  59.632  1.00  0.00           H
    ATOM   1507  HB2 SER A 458      -4.142  36.035  58.213  1.00  0.00           H
    ATOM   1508  HB3 SER A 458      -2.780  36.607  57.297  1.00  0.00           H
    ATOM   1509  HG  SER A 458      -4.214  34.183  57.007  1.00  0.00           H
    ATOM   1510  N   PRO A 459      -3.812  33.539  59.982  1.00 92.48           N
    ATOM   1511  CA  PRO A 459      -4.439  32.240  60.311  1.00 93.69           C
    ATOM   1512  C   PRO A 459      -5.033  31.519  59.084  1.00 95.04           C
    ATOM   1513  O   PRO A 459      -5.575  32.182  58.197  1.00 94.23           O
    ATOM   1514  CB  PRO A 459      -5.545  32.600  61.318  1.00 92.86           C
    ATOM   1515  CG  PRO A 459      -5.211  33.990  61.825  1.00 93.86           C
    ATOM   1516  CD  PRO A 459      -4.525  34.640  60.634  1.00 93.55           C
    ATOM   1517  HA  PRO A 459      -3.687  31.615  60.797  1.00  0.00           H
    ATOM   1518  HB2 PRO A 459      -6.513  32.625  60.821  1.00  0.00           H
    ATOM   1519  HB3 PRO A 459      -5.622  31.877  62.131  1.00  0.00           H
    ATOM   1520  HG2 PRO A 459      -6.086  34.535  62.176  1.00  0.00           H
    ATOM   1521  HG3 PRO A 459      -4.510  33.912  62.656  1.00  0.00           H
    ATOM   1522  HD2 PRO A 459      -5.272  35.038  59.946  1.00  0.00           H
    ATOM   1523  HD3 PRO A 459      -3.890  35.459  60.969  1.00  0.00           H
    ATOM   1524  N   SER A 460      -4.923  30.179  59.069  1.00 97.29           N
    ATOM   1525  CA  SER A 460      -5.430  29.324  57.990  1.00 99.88           C
    ATOM   1526  C   SER A 460      -6.964  29.195  57.972  1.00101.95           C
    ATOM   1527  O   SER A 460      -7.551  29.270  56.892  1.00102.69           O
    ATOM   1528  CB  SER A 460      -4.723  27.949  58.031  1.00 96.79           C
    ATOM   1529  OG  SER A 460      -5.214  27.089  59.043  1.00 96.54           O
    ATOM   1530  H   SER A 460      -4.465  29.702  59.832  1.00  0.00           H
```
