## supplementary information for "DISRUPTOR: Computational identification of oncogenic mutants disrupting protein interactions"

**Table S1.** Prioritized hotspot mutations for p53 chain A predicted to disrupt interactions DNA consensus sequence.

| wt residue[1] | mutant | K*(log10) | diff to wt[2] | COSMIC[3] |
|---|---|---|---|---|
| K101 | A | 2,34299 | -3,852917 | - |
| K101 | N | 2,383493 | -3,812414 | 1 |
| K101 | D | -0,10607 | -6,301977 | - |
| K101 | Q | 2,541033 | -3,654874 | - |
| K101 | E | -0,281937 | -6,477844 | - |
| K101 | G | 2,329185 | -3,866722 | - |
| K101 | H | 2,408996 | -3,786911 | - |
| K101 | I | 2,385238 | -3,810669 | - |
| K101 | L | 2,422414 | -3,773493 | - |
| K101 | M | 2,432355 | -3,763552 | - |
| K101 | P | 2,347536 | -3,848371 | - |
| K101 | S | 2,365646 | -3,830261 | - |
| K101 | T | 2,40944 | -3,786467 | - |
| K101 | Y | 2,590051 | -3,605856 | - |
| K101 | V | 2,376722 | -3,819185 | - |
| R249 | A | 0,517047 | -3,879753 | - |
| R249 | N | 0,321083 | -4,075717 | - |
| R249 | C | 0,476555 | -3,920245 | - |
| R249 | Q | 1,173259 | -3,223541 | - |
| R249 | E | -3,828045 | -8,224845 | - |
| R249 | G | 0,477414 | -3,919386 | 68 |
| R249 | I | 0,674289 | -3,722511 | 1 |
| R249 | L | 0,816013 | -3,580787 | - |
| R249 | M | 0,707806 | -3,688994 | 78 |
| R249 | P | none | X | - |
| R249 | S | 0,573587 | -3,823213 | 609 |
| R249 | T | 0,448275 | -3,948525 | 5046 |
| R249 | W | 1,102095 | -3,294705 | 61 |
| R249 | V | 0,629113 | -3,767687 | 1 |
| R248 | A | 0,203102 | -4,698122 | - |
| R248 | C | 0,321923 | -4,579301 | 1 |
| R248 | Q | 0,431464 | -4,46976 | 1520 |
| R248 | E | -3,03584 | -7,937064 | - |
| R248 | G | 0,181438 | -4,719786 | 36 |
| R248 | H | 0,37264 | -4,528584 | 1 |
| R248 | I | 0,288149 | -4,613075 | - |
| R248 | M | 0,307344 | -4,59388 | - |
| R248 | P | none | X | 40 |

| | | | | |
|---|---|---|---|---|
| R248 | S | 0,298687 | -4,602537 | - |
| R248 | T | 0,077249 | -4,823975 | - |
| R248 | W | 0,351411 | -4,549813 | 1193 |
| R248 | V | 0,265969 | -4,635255 | - |

[1] wildtype residue, [2]difference of K* score of mutant in relation to wt residue, [3] number of patient samples harboring the mutant reported in COSMIC, none… no binding at all was predicted for the mutant, X… thus no difference of the K* score to the wt could be calculated.

**Table S2**. Prioritized hotspot mutations for p53 chain B predicted to disrupt interactions DNA consensus sequence.

| wt residue[1] | mutant | K*(log10) | diff to wt[2] | COSMIC[3] |
|---|---|---|---|---|
| K120 | A | 2,691146 | -11,188792 | - |
| K120 | N | 4,108135 | -9,771803 | 3 |
| K120 | D | -4,373352 | -18,25329 | - |
| K120 | Q | 5,186083 | -8,693855 | 1 |
| K120 | E | -4,313321 | -18,193259 | 21 |
| K120 | G | 1,991899 | -11,888039 | - |
| K120 | H | -10,93979 | -24,819728 | - |
| K120 | I | -7,360472 | -21,24041 | - |
| K120 | L | 4,388371 | -9,491567 | - |
| K120 | M | 4,931815 | -8,948123 | 10 |
| K120 | P | none | X | - |
| K120 | S | 2,930995 | -10,948943 | - |
| K120 | T | 3,46453 | -10,415408 | 3 |
| K120 | W | none | X | - |
| K120 | Y | none | X | - |
| K120 | V | 4,020949 | -9,858989 | - |
| R273 | A | 10,016722 | -7,825299 | - |
| R273 | N | 10,008983 | -7,833038 | - |
| R273 | D | 5,055741 | -12,78628 | - |
| R273 | C | 9,978956 | -7,863065 | 1379 |
| R273 | Q | 10,768056 | -7,073965 | 2 |
| R273 | G | 9,989578 | -7,852443 | 27 |
| R273 | H | 10,095936 | -7,746085 | 1406 |
| R273 | I | 10,197308 | -7,644713 | - |
| R273 | L | 10,283987 | -7,558034 | 278 |
| R273 | F | 11,116865 | -6,725156 | - |
| R273 | P | none | X | 56 |
| R273 | S | 9,967608 | -7,874413 | 43 |
| R273 | T | 10,105834 | -7,736187 | - |
| R273 | W | 11,407796 | -6,434225 | - |
| R273 | Y | 11,329325 | -6,512696 | 1 |
| R273 | V | 10,108209 | -7,733812 | - |
| R283 | A | 0,606024 | -4,514618 | - |
| R283 | N | 0,89169 | -4,228952 | - |
| R283 | D | -2,922849 | -8,043491 | - |
| R283 | C | 0,706364 | -4,414278 | 37 |
| R283 | Q | 0,555647 | -4,564995 | - |
| R283 | G | 0,590585 | -4,530057 | 3 |

| | | | | |
|---|---|---|---|---|
| R283 | H | 0,742676 | -4,377966 | 23 |
| R283 | I | 0,695116 | -4,425526 | - |
| R283 | L | 0,669899 | -4,450743 | 4 |
| R283 | F | 0,658977 | -4,461665 | - |
| R283 | P | none | X | 56 |
| R283 | S | 0,630511 | -4,490131 | 1 |
| R283 | T | 0,638687 | -4,481955 | - |
| R283 | W | 0,647427 | -4,473215 | - |
| R283 | Y | 0,670915 | -4,449727 | - |
| R283 | V | 0,65277 | -4,467872 | - |

[1] wildtype residue, [2]difference of K* score of mutant in relation to wt residue, [3] number of patient samples harboring the mutant reported in COSMIC, none… no binding at all was predicted for the mutant, X… thus no difference of the K* score to the wt could be calculated.

**Table S3.** Prioritized hotspot mutations for p53 chain C predicted to disrupt interactions DNA consensus sequence.

| wt residue[1] | mutant | K*(log10) | diff to wt[2] | COSMIC[3] |
|---|---|---|---|---|
| K120 | A | 1,815678 | -6,933418 | - |
| K120 | N | 2,775805 | -5,973291 | 3 |
| K120 | D | -3,462315 | -12,211411 | - |
| K120 | Q | 2,493117 | -6,255979 | 1 |
| K120 | E | -3,730755 | -12,479851 | 21 |
| K120 | G | 1,873719 | -6,875377 | - |
| K120 | H | 3,06706 | -5,682036 | - |
| K120 | I | -39,45776 | -48,206856 | - |
| K120 | L | 2,954455 | -5,794641 | - |
| K120 | M | 3,706782 | -5,042314 | 10 |
| K120 | P | 2,544973 | -6,204123 | - |
| K120 | S | 2,16749 | -6,581606 | - |
| K120 | T | -19,04392 | -27,793016 | 3 |
| K120 | W | 4,1289 | -4,620196 | - |
| K120 | Y | 3,522574 | -5,226522 | - |
| K120 | V | -56,05519 | -64,804286 | - |
| R280 | A | -2,73018 | -4,932897 | - |
| R280 | N | -2,896371 | -5,099088 | - |
| R280 | C | -2,602158 | -4,804875 | - |
| R280 | Q | -2,396227 | -4,598944 | - |
| R280 | E | -7,085311 | -9,288028 | - |
| R280 | G | -2,799111 | -5,001828 | 78 |
| R280 | I | -2,419549 | -4,622266 | 50 |
| R280 | L | -2,313941 | -4,516658 | - |
| R280 | M | -2,239468 | -4,442185 | - |
| R280 | P | -2,619168 | -4,821885 | - |
| R280 | S | -2,751042 | -4,953759 | 39 |
| R280 | T | -2,635841 | -4,838558 | 174 |
| R280 | W | -1,659759 | -3,862476 | - |
| R280 | V | -2,540043 | -4,74276 | - |
| R248 | A | 1,724592 | -3,233005 | - |
| R248 | C | 1,856652 | -3,100945 | 1 |
| R248 | Q | 1,727043 | -3,230554 | 1520 |
| R248 | E | -1,084808 | -6,042405 | - |
| R248 | G | 1,556304 | -3,401293 | 36 |
| R248 | H | 1,888048 | -3,069549 | 1 |
| R248 | L | -2,518818 | -7,476415 | 159 |
| R248 | P | none | X | 40 |

| | | | | |
|---|---|---|---|---|
| R248 | S | 1,737028 | -3,220569 | - |
| R248 | W | -0,814039 | -5,771636 | 1193 |

[1] wildtype residue, [2]difference of K* score of mutant in relation to wt residue, [3] number of patient samples harboring the mutant reported in COSMIC, none… no binding at all was predicted for the mutant, X… thus no difference of the K* score to the wt could be calculated.

**Table S4**. Prioritized hotspot mutations for p53 predicted to disrupt interactions with DP2. Please note that four residue positions are reported, because an equal number of individual mutations was predicted for residues 248,249, 280.

| wt residue[1] | mutant | K*(log10) | diff to wt[2] | COSMIC[3] |
|---|---|---|---|---|
| R273 | A | 0,323495 | -4,932792 | - |
| R273 | N | 0,269833 | -4,986454 | - |
| R273 | D | -2,454937 | -7,711224 | - |
| R273 | C | 0,277161 | -4,979126 | 1379 |
| R273 | Q | 0,661741 | -4,594546 | 2 |
| R273 | G | 0,309248 | -4,947039 | 27 |
| R273 | H | 0,239307 | -5,01698 | 1406 |
| R273 | I | 0,346589 | -4,909698 | - |
| R273 | L | 0,363471 | -4,892816 | 278 |
| R273 | F | 0,560962 | -4,695325 | - |
| R273 | P | none | X | 56 |
| R273 | S | 0,295117 | -4,96117 | 43 |
| R273 | T | 0,351097 | -4,90519 | - |
| R273 | W | 0,891786 | -4,364501 | - |
| R273 | Y | 0,535113 | -4,721174 | 1 |
| R273 | V | 0,343263 | -4,913024 | - |
| R248 | A | 5,193638 | -4,511186 | - |
| R248 | C | 6,215555 | -13,489269 | 1 |
| R248 | Q | 3,323453 | -16,381371 | 1520 |
| R248 | E | -3,524651 | -23,229475 | - |
| R248 | G | 5,141793 | -14,563031 | 36 |
| R248 | H | 3,29916 | -16,405664 | 1 |
| R248 | L | 3,921772 | -15,783052 | 159 |
| R248 | K | 14,533442 | -5,171382 | - |
| R248 | M | 7,055204 | -12,64962 | - |
| R248 | P | none | X | 40 |
| R248 | S | 5,490434 | -14,21439 | - |
| R248 | T | -15,45115 | -35,155974 | - |
| R248 | W | -33,78764 | -53,492464 | 1193 |
| R248 | V | 6,466922 | -13,237902 | - |
| R249 | A | 15,437851 | -4,266973 | - |
| R249 | N | 14,964978 | -4,739846 | - |
| R249 | C | 15,689048 | -4,015776 | - |
| R249 | Q | 13,903387 | -5,801437 | - |
| R249 | E | 10,255063 | -9,449761 | - |
| R249 | G | 15,266362 | -4,438462 | 68 |
| R249 | I | 16,043865 | -3,660959 | 1 |

| | | | | |
|---|---|---|---|---|
| R249 | L | 6,003125 | -13,701699 | - |
| R249 | M | 16,406264 | -3,29856 | 78 |
| R249 | P | none | X | - |
| R249 | S | 15,557841 | -4,146983 | 609 |
| R249 | T | 15,519187 | -4,185637 | 50 |
| R249 | W | none | X | 61 |
| R249 | V | 16,015213 | -3,689611 | 1 |
| R280 | A | 0,475741 | -5,638805 | - |
| R280 | N | 0,825332 | -5,289214 | - |
| R280 | C | 0,533749 | -5,580797 | - |
| R280 | Q | 0,634461 | -5,480085 | - |
| R280 | E | -1,778061 | -7,892607 | - |
| R280 | G | 0,453977 | -5,660569 | 78 |
| R280 | I | 0,56953 | -5,545016 | 50 |
| R280 | L | 0,645035 | -5,469511 | - |
| R280 | M | 0,837309 | -5,277237 | - |
| R280 | P | none | X | - |
| R280 | S | 0,539367 | -5,575179 | 39 |
| R280 | T | 0,58243 | -5,532116 | 174 |
| R280 | W | 1,116796 | -4,99775 | - |
| R280 | V | 0,546488 | -5,568058 | - |

[1] wildtype residue, [2]difference of K* score of mutant in relation to wt residue, [3] number of patient samples harboring the mutant reported in COSMIC, none… no binding at all was predicted for the mutant, X… thus no difference of the K* score to the wt could be calculated.

**Table S5.** Prioritized hotspot mutations for ERK2 predicted to disrupt interactions with DUSP6.

| wt residue[1] | mutant | K*(log10) | diff to wt[2] | Pb[3] colorectal cancer | Pb[3] cervical cancer | COSMIC[4] |
|---|---|---|---|---|---|---|
| D321 | N | -1,182303 | -8,764402 | 0,00402124 | 0,00646836 | 8 |
| D321 | H | -14,64852 | -22,230619 | 0,00096405 | 0,00477121 | - |
| D321 | Y | None[5] | X[6] | 0,00128344 | 0,00352652 | 1 |
| D321 | G | 0,123075 | -7,459024 | 0,00604358 | 0,00298779 | 2 |
| D321 | V | 0,346748 | -7,235351 | 0,00064382 | 0,00246853 | 1 |
| D321 | A | 0,272173 | -7,309926 | 0,00091117 | 0,00146627 | 1 |
| D321 | E | -4,191007 | -11,773106 | 0,00042221 | 0,00142358 | 1 |
| D321 | S | -0,208939 | -7,791038 | 2,4673E-05 | 2,3709E-05 | - |
| D321 | I | -3,383509 | -10,965608 | 2,5077E-06 | 1,5454E-05 | - |
| D321 | R | -12,0264 | -19,608499 | 5,6435E-06 | 1,3797E-05 | - |
| D321 | L | -16,51716 | -24,099259 | 6,012E-07 | 1,1399E-05 | - |
| D321 | C | 0,639544 | -6,942555 | 7,5132E-06 | 1,0198E-05 | - |
| D321 | T | -3,179452 | -10,761551 | 3,5491E-06 | 9,1792E-06 | - |
| D321 | K | -9,397628 | -16,979727 | 1,6713E-06 | 9,0652E-06 | - |
| D321 | F | none | X | 8,0038E-07 | 8,4253E-06 | - |
| D321 | P | none | X | 8,5085E-07 | 6,7708E-06 | - |
| D321 | Q | -4,940341 | -12,52244 | 4,0068E-07 | 6,6867E-06 | - |
| D318 | N | -0,212286 | -4,309833 | 0,00999477 | 0,02316461 | 1 |
| D318 | H | 0,930958 | -3,166589 | 0,00011049 | 0,00341755 | - |
| D318 | G | 0,648646 | -3,448901 | 0,00457925 | 0,00240737 | - |
| D318 | Y | 0,909629 | -3,187918 | 0,00067654 | 0,00231833 | - |
| D318 | V | 0,788937 | -3,30861 | 0,00065276 | 0,00130214 | - |
| D318 | A | 0,693979 | -3,403568 | 0,00025986 | 0,00047436 | - |
| D318 | K | -1,872717 | -5,970264 | 2,3788E-05 | 0,00018536 | - |
| D318 | S | 0,647953 | -3,449594 | 4,5487E-05 | 5,5821E-05 | - |
| D318 | I | -0,010491 | -4,108038 | 6,4592E-06 | 0,00002961 | - |
| D318 | Q | 0,52897 | -3,568577 | 2,6297E-07 | 2,7347E-05 | - |
| D318 | T | 0,960081 | -3,137466 | 2,5714E-06 | 1,0787E-05 | - |
| D318 | R | -2,143087 | -6,240634 | 5,0092E-07 | 8,0762E-06 | - |
| D318 | C | 0,418232 | -3,679315 | 3,0672E-06 | 5,4786E-06 | - |
| D318 | L | -2,06998 | -6,167527 | 7,1404E-08 | 4,3684E-06 | - |
| D318 | F | 1,005684 | -3,091863 | 4,3722E-07 | 2,9633E-06 | - |
| D318 | P | none | X | 2,8426E-08 | 1,5914E-06 | - |
| E81 | K | -0,356733 | -5,721186 | 0,00397595 | 0,00679256 | 5 |
| E81 | Q | 2,323264 | -3,041189 | 0,00095319 | 0,00501034 | - |
| E81 | G | 1,556504 | -3,807949 | 0,00561014 | 0,0035114 | - |

| | | | | | | |
|---|---|---|---|---|---|---|
| E81 | V | 1,735473 | -3,62898 | 0,00093775 | 0,00209152 | - |
| E81 | A | 1,59246 | -3,771993 | 0,00070536 | 0,001344 | - |
| E81 | N | 1,803066 | -3,561387 | 2,7476E-05 | 0,000647 | - |
| E81 | H | 2,091465 | -3,272988 | 6,5871E-06 | 0,00047724 | - |
| E81 | Y | 2,151015 | -3,213438 | 8,7695E-06 | 0,00035274 | - |
| E81 | R | 0,386516 | -4,977937 | 2,7223E-05 | 3,3257E-05 | - |
| E81 | L | 2,071127 | -3,293326 | 2,0514E-06 | 1,4624E-05 | - |
| E81 | W | 2,223596 | -3,140857 | 7,0083E-06 | 1,0435E-05 | - |
| E81 | T | 1,724047 | -3,640406 | 2,7608E-06 | 7,3257E-06 | - |
| E81 | P | 1,603678 | -3,760775 | 6,6187E-07 | 5,4036E-06 | - |
| E81 | S | 1,64994 | -3,714513 | 8,8115E-07 | 3,9939E-06 | - |

[1] wildtype residue, [2]difference of K* score of mutant in relation to wt residue, [3] Pb relative Probability for a mutation to be formed in the indicated cancer type, [4] number of patient samples harboring the mutant reported in COSMIC, none… no binding at all was predicted for the mutant, X… thus no difference of the K* score to the wt could be calculated.

**Table S6.** Prioritized hotspot mutations for p16 predicted to disrupt interactions with CDK6. Please note that four residue positions are reported, because an equal number of individual mutations was predicted for residues G23, G55, G89.

| wt residue[1] | mutant | K*(log10) | diff to wt[2] | Pb[3] melanoma | COSMIC[4] |
|---|---|---|---|---|---|
| D84 | A | 0,705794 | -4,822505 | 5,1003E-05 | 1 |
| D84 | R | none | X | 4,9293E-05 | - |
| D84 | N | 0,267755 | -5,260544 | 0,07688685 | 37 |
| D84 | C | 1,413534 | -4,114765 | 2,2295E-07 | - |
| D84 | Q | -12,0721 | -17,600399 | 4,3045E-08 | - |
| D84 | E | -9,249456 | -14,777755 | 0,00015851 | - |
| D84 | G | 0,330964 | -5,197335 | 0,00064395 | 15 |
| D84 | H | none | X | 0,00027263 | 2 |
| D84 | I | none | X | 1,3085E-05 | - |
| D84 | L | -57,34838 | -62,876679 | 4,6397E-08 | - |
| D84 | K | -39,35921 | -44,887509 | 1,214E-05 | - |
| D84 | F | none | X | 5,9392E-08 | - |
| D84 | P | none | X | 1,3794E-08 | - |
| D84 | S | 1,151568 | -4,376731 | 4,9137E-05 | - |
| D84 | T | 1,457733 | -4,070566 | 3,8903E-06 | - |
| D84 | Y | none | X | 0,00034898 | 33 |
| D84 | V | none | X | 0,00017154 | 6 |
| G23 | A | none | X | 0,00024447 | - |
| G23 | R | none | X | 0,00049361 | 2 |
| G23 | N | none | X | 0,00125316 | - |
| G23 | D | none | X | 0,01622997 | 2 |
| G23 | C | none | X | 0,00073695 | 1 |
| G23 | E | none | X | 3,595E-06 | - |
| G23 | H | none | X | 7,7217E-06 | - |
| G23 | I | none | X | 6,9919E-05 | - |
| G23 | L | none | X | 4,3083E-07 | - |
| G23 | F | none | X | 6,6648E-07 | - |
| G23 | P | none | X | 1,1628E-07 | - |
| G23 | S | none | X | 0,07731202 | 6 |
| G23 | T | none | X | 1,8872E-05 | - |
| G23 | W | none | X | 3,741E-08 | - |
| G23 | Y | none | X | 1,1945E-05 | - |
| G23 | V | none | X | 0,00090574 | 5 |
| G55 | A | none | X | 6,096E-07 | - |
| G55 | R | none | X | 0,00057867 | 3 |
| G55 | N | none | X | 0,00136281 | - |

| | | | | | |
|---|---|---|---|---|---|
| G55 | D | none | X | 0,01766393 | 2 |
| G55 | C | none | X | 0,00073753 | 1 |
| G55 | E | none | X | 2,3125E-05 | - |
| G55 | H | none | X | 8,3974E-06 | - |
| G55 | I | none | X | 3,5426E-05 | - |
| G55 | L | none | X | 2,1829E-07 | - |
| G55 | F | none | X | 3,3769E-07 | - |
| G55 | P | none | X | 2,2125E-07 | - |
| G55 | S | none | X | 0,07737281 | - |
| G55 | T | none | X | 3,5907E-05 | - |
| G55 | W | none | X | 9,5684E-08 | - |
| G55 | Y | none | X | 1,2991E-05 | - |
| G55 | V | none | X | 0,00045977 | 3 |
| G89 | A | none | X | 0,00047142 | - |
| G89 | R | none | X | 0,00053999 | - |
| G89 | N | none | X | 0,00136281 | - |
| G89 | D | none | X | 0,01787906 | 1 |
| G89 | C | none | X | 0,00074651 | 2 |
| G89 | E | none | X | 1,3031E-05 | - |
| G89 | H | none | X | 8,3974E-06 | - |
| G89 | I | none | X | 3,5426E-05 | - |
| G89 | L | none | X | 2,1829E-07 | - |
| G89 | F | none | X | 3,3769E-07 | - |
| G89 | P | none | X | 2,2125E-07 | - |
| G89 | S | none | X | 0,07831512 | 6 |
| G89 | T | none | X | 3,5907E-05 | - |
| G89 | W | none | X | 1,4967E-07 | - |
| G89 | Y | none | X | 1,2991E-05 | - |
| G89 | V | none | X | 0,0004651 | 2 |

[1] wildtype residue, [2]difference of K* score of mutant in relation to wt residue, [3] Pb relative Probability for a mutation to be formed in the indicated cancer type, [4] number of patient samples harboring the mutant reported in COSMIC, none… no binding at all was predicted for the mutant, X… thus no difference of the K* score to the wt could be calculated.

**Table S7.** Prioritized hotspot mutations for CDK6 predicted to disrupt interactions with p16. Please note that four residue positions are reported, because an equal number of individual mutations was predicted for residues G37 and R31, G20 and K111.

| wt residue[1] | mutant | K*(log10) | diff to wt[2] | COSMIC[3] |
|---|---|---|---|---|
| G37 | A | none | X | |
| G37 | R | none | X | |
| G37 | N | none | X | |
| G37 | D | none | X | |
| G37 | C | none | X | |
| G37 | E | none | X | |
| G37 | H | none | X | |
| G37 | I | none | X | |
| G37 | L | none | X | |
| G37 | F | none | X | |
| G37 | P | none | X | |
| G37 | S | none | X | |
| G37 | T | none | X | |
| G37 | W | none | X | |
| G37 | Y | none | X | |
| G37 | V | none | X | |
| R31 | A | 0.798177 | -5.643494 | |
| R31 | N | 1.248857 | -5.192814 | |
| R31 | D | -1.09348 | -7.535152 | |
| R31 | C | 1.339103 | -5.102568 | |
| R31 | Q | 0.164235 | -6.277436 | |
| R31 | G | 0.394287 | -6.047384 | |
| R31 | H | 2.442278 | -3.999393 | |
| R31 | I | none | X | |
| R31 | L | -3.61789 | -10.059558 | |
| R31 | F | none | X | |
| R31 | P | none | X | |
| R31 | S | 0.799319 | -5.642352 | |
| R31 | T | 1.403068 | -5.038603 | |
| R31 | W | none | X | |
| R31 | Y | none | X | |
| R31 | V | 1.606236 | -4.835435 | |
| G20 | A | none | X | |
| G20 | R | none | X | |
| G20 | D | none | X | |
| G20 | C | none | X | |
| G20 | Q | none | X | |

| | | | |
|---|---|---|---|
| G20 | E | none | X |
| G20 | L | none | X |
| G20 | K | none | X |
| G20 | M | none | X |
| G20 | P | none | X |
| G20 | S | none | X |
| G20 | T | none | X |
| G20 | W | none | X |
| G20 | V | none | X |
| K111 | A | 1.493153 | -4.234931 |
| K111 | D | -0.173548 | -5.901632 |
| K111 | Q | 2.712797 | -3.015287 |
| K111 | E | 0.421897 | -5.306187 |
| K111 | G | 1.310223 | -4.417861 |
| K111 | H | -66.34503 | -72.073114 |
| K111 | I | -10.72942 | -16.457504 |
| K111 | L | 1.606365 | -4.121719 |
| K111 | M | 1.894562 | -3.833522 |
| K111 | P | none | X |
| K111 | S | 1.558725 | -4.169359 |
| K111 | T | 1.949978 | -3.778106 |
| K111 | Y | none | X |
| K111 | V | 2.363409 | -3.364675 |

[1] wildtype residue, [2]difference of K* score of mutant in relation to wt residue, [3] number of patient samples harboring the mutant reported in COSMIC, none… no binding at all was predicted for the mutant, X… thus no difference of the K* score to the wt could be calculated.

**Table S8.** Prioritized hotspot mutations for smad4 predicted to disrupt interactions with smad2. Please note that five residue positions are reported, because an equal number of individual mutations was predicted for residues 361, 365, 428, 507.

| wt residue[1] | mutant | K*(log10) | diff to wt[2] | Pb[3] colorectal | COSMIC[4] |
|---|---|---|---|---|---|
| D537 | A | 4.582058 | -4.821045 | 0.00026246 | 4 |
| D537 | R | none | X | 5.2273E-06 | |
| D537 | N | 4.470593 | -4.93251 | 0.00448332 | |
| D537 | C | 4.884818 | -4.518285 | 2.6362E-05 | |
| D537 | Q | -24.48158 | -33.884683 | 4.8495E-07 | |
| D537 | E | 2.371261 | -7.031842 | 0.00042221 | 7 |
| D537 | G | 3.761375 | -5.641728 | 0.00462508 | 20 |
| D537 | H | none | X | 0.00116682 | 6 |
| D537 | I | none | X | 2.8631E-06 | |
| D537 | L | none | X | 7.4514E-07 | |
| D537 | K | none | X | 1.8634E-06 | |
| D537 | F | none | X | 3.7578E-06 | |
| D537 | P | none | X | 2.9663E-07 | |
| D537 | S | 4.739755 | -4.663348 | 2.1581E-05 | |
| D537 | T | 2.886855 | -6.516248 | 1.1398E-06 | |
| D537 | Y | none | X | 0.00588437 | 29 |
| D537 | V | -19.50471 | -28.907813 | 0.00065929 | 14 |
| R361 | A | 0.974539 | -4.772984 | 2.8985E-09 | |
| R361 | N | 1.028212 | -4.719311 | 1.118E-05 | |
| R361 | D | 0.266894 | -5.480629 | 1.8259E-06 | |
| R361 | C | 1.065402 | -4.682121 | 0.01002855 | 167 |
| R361 | Q | 0.886778 | -4.860745 | 4.1914E-05 | |
| R361 | G | 0.959733 | -4.78779 | 0.00011114 | 15 |
| R361 | H | 1.062984 | -4.684539 | 0.01670801 | 210 |
| R361 | I | 1.070326 | -4.677197 | 2.5094E-07 | |
| R361 | L | 1.089429 | -4.658094 | 0.00037596 | 4 |
| R361 | F | 1.023549 | -4.723974 | 2.659E-05 | |
| R361 | P | none | X | 0.00067909 | 3 |
| R361 | S | 0.976897 | -4.770626 | 0.00067909 | 14 |
| R361 | T | 0.988378 | -4.759145 | 1.7748E-08 | |
| R361 | W | 1.545709 | -4.201814 | 6.2797E-06 | |
| R361 | Y | 1.015874 | -4.731649 | 0.00016517 | |
| R361 | V | 1.025202 | -4.722321 | 4.0982E-08 | |
| G365 | A | none | X | 0.00073185 | 1 |
| G365 | R | none | X | 0.00042811 | 1 |
| G365 | N | none | X | 1.4458E-05 | |

| | | | | | |
|---|---|---|---|---|---|
| G365 | D | none | X | 0.00383563 | 5 |
| G365 | C | none | X | 0.00274886 | |
| G365 | E | none | X | 3.4631E-06 | |
| G365 | H | none | X | 1.6127E-06 | |
| G365 | I | none | X | 6.2906E-06 | |
| G365 | L | none | X | 7.0167E-07 | |
| G365 | F | none | X | 4.5458E-06 | |
| G365 | P | none | X | 3.0744E-07 | |
| G365 | S | none | X | 0.0038059 | 4 |
| G365 | T | none | X | 2.7562E-06 | |
| G365 | W | none | X | 7.0669E-07 | |
| G365 | Y | none | X | 1.0448E-05 | |
| G365 | V | none | X | 0.00167031 | 2 |
| K428 | A | 0.126469 | -6.547002 | 3.6755E-06 | |
| K428 | N | 0.448996 | -6.224475 | 0.00694114 | |
| K428 | D | -1.760524 | -8.433995 | 1.4012E-05 | |
| K428 | Q | 0.903522 | -5.769949 | 0.00026279 | |
| K428 | E | -2.886629 | -9.5601 | 0.00202761 | |
| K428 | G | 0.102477 | -6.570994 | 1.3902E-05 | |
| K428 | H | 0.183194 | -6.490277 | 1.8161E-06 | |
| K428 | I | 0.333433 | -6.340038 | 1.1911E-05 | |
| K428 | L | 0.411409 | -6.262062 | 2.7446E-07 | |
| K428 | M | 0.175197 | -6.498274 | 0.00104902 | |
| K428 | P | 0.158751 | -6.51472 | 4.7637E-07 | |
| K428 | S | 0.14608 | -6.527391 | 4.927E-05 | |
| K428 | T | 0.184828 | -6.488643 | 0.00184139 | |
| K428 | W | 0.119359 | -6.554112 | 5.5573E-06 | |
| K428 | Y | 0.147132 | -6.526339 | 5.6013E-06 | |
| K428 | V | 0.196069 | -6.477402 | 2.1177E-06 | |
| K507 | A | 2.539634 | -8.242713 | 9.6383E-06 | |
| K507 | R | 5.220245 | -5.562102 | 0.00391035 | |
| K507 | N | 2.963475 | -7.818872 | 0.00286974 | 4 |
| K507 | D | -1.696103 | -12.47845 | 1.2285E-05 | |
| K507 | Q | 0.219739 | -10.562608 | 0.00073351 | 3 |
| K507 | E | -6.18591 | -16.968257 | 0.00431049 | 5 |
| K507 | G | 2.343533 | -8.438814 | 1.6626E-05 | |
| K507 | H | 3.447321 | -7.335026 | 2.0906E-06 | |
| K507 | I | none | X | 0.00096382 | |
| K507 | L | none | X | 1.6549E-06 | |
| K507 | M | 3.926115 | -6.856232 | 6.6185E-06 | |
| K507 | P | none | X | 1.6401E-06 | |
| K507 | S | 2.548982 | -8.233365 | 1.3382E-05 | |

| | | | | |
|---|---|---|---|---|
| K507 | T | 2.970434 | -7.811913 | 0.00227337 |
| K507 | Y | 4.805533 | -5.976814 | 2.8508E-06 |
| K507 | V | none | X | 4.1143E-06 |

[1] wildtype residue, [2]difference of K* score of mutant in relation to wt residue, [3] Pb relative Probability for a mutation to be formed in the indicated cancer type, [4] number of patient samples harboring the mutant reported in COSMIC, none… no binding at all was predicted for the mutant, X… thus no difference of the K* score to the wt could be calculated.

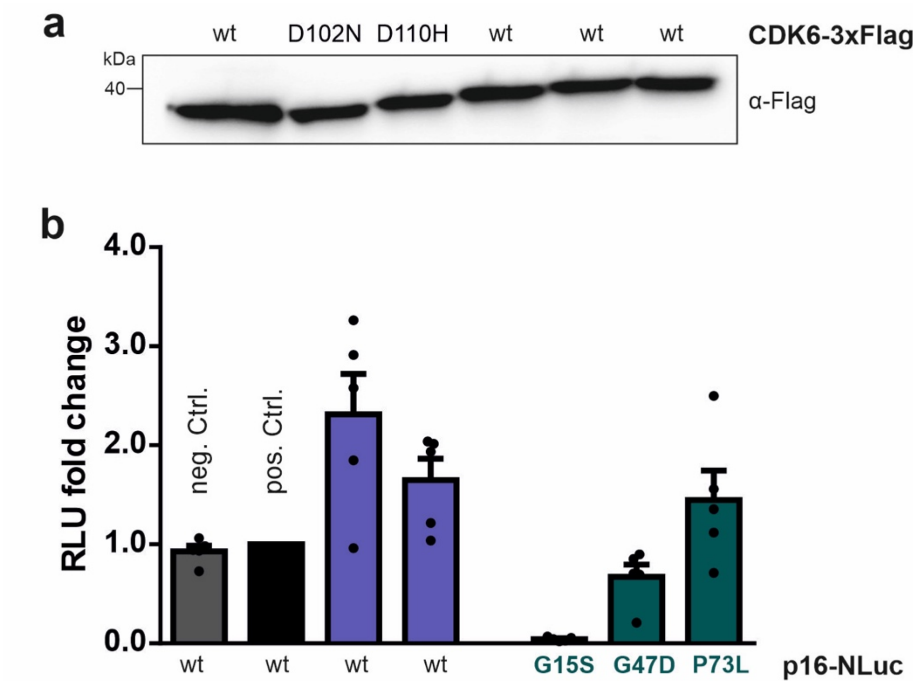

**Supplementary Fig. 1.** **a** Immunoblotting shows expression of Flag tagged CDK6 variants. **b** Bioluminescence signals of 1% input samples of LUMIER experiments. Bars represent the luciferase intensities of p16-NLuc variants relative to the wild-type. Input signals were used for normalization of signals after immunoprecipitation. Error bars represent SEM with n = 5 experiments.
